# Diverse CRISPR-Cas complexes require independent translation of small and large subunits from a single gene

**DOI:** 10.1101/2020.04.18.045682

**Authors:** Tess M. McBride, Evan A. Schwartz, Abhishek Kumar, David W. Taylor, Peter C. Fineran, Robert D. Fagerlund

## Abstract

CRISPR-Cas adaptive immune systems provide prokaryotes with defense against viruses by degradation of specific invading nucleic acids. We investigated the previously uncharacterized type I-D interference complex from *Synechocystis* and revealed it is a genetic and structural hybrid with similarity to both type I and III systems. Surprisingly, formation of the functional complex required internal in-frame translation of small subunits from within the large subunit gene. We further show that internal translation to generate small subunits is widespread across diverse type I-D, I-B and I-C systems, which account for roughly one quarter of CRISPR-Cas systems. Our work reveals the unexpected expansion of protein coding potential from within single *cas* genes, which has important implications for understanding CRISPR-Cas function and evolution.

**One Sentence Summary:** Internal translation of large subunit transcripts drives small subunit synthesis in diverse type I CRISPR-Cas interference complexes

## Main Text

Prokaryotes have evolved several strategies to provide defense against mobile genetic elements, such as viruses and plasmids (*1*). One mechanism is CRISPR-Cas immunity (*2*), which consists of CRISPR array(s), which store foreign sequences as spacers (*3*), and *cas* genes that encode proteins required for adaptive immunity (*4, 5*). During infection, CRISPR arrays are transcribed and processed into CRISPR RNAs (crRNAs), which assemble with the Cas protein(s) to form ribonucleoprotein interference complexes (*2*). The crRNAs guide these complexes to complementary invading nucleic acids (protospacers) and elicit interference by the degradation of the foreign element (*6*).

CRISPR-Cas systems are divided into two classes (*7*). Class 1 systems utilize characteristic multi-protein complexes, whereas class 2 systems utilize a single multi-domain protein for interference (e.g. Cas9). Class 1 encompass ~80% of all CRISPR-Cas systems found in sequenced microbes, demonstrating their evolutionary success, and can be further divided into types I, III, and IV (*4, 8*). Despite having distinct architecture and targeting mechanisms, type I and III interference complexes have a common evolutionary history and some structural similarities, consisting of multiple Cas7 proteins, which form a backbone capped at one end by Cas5 and the large subunit (Cas8 and Cas10 in type I and III systems, respectively) (*2, 6, 8*). In most type I complexes the opposite end is capped by Cas6, an RNA endonuclease that processes the precursor transcript^6^. In contrast, Cas6 dissociates after crRNA maturation in type III complexes (*9, 10*). Type III and two type I systems (type I-A and I-E) also include multiple Cas11 proteins (or small subunits), which form the minor filament along Cas7 (*9–13*). Type I interference complexes are termed Cascade (CRISPR associated complex for antiviral defence) and the genes encoding the type I-D Cascade have features that resemble both type I and III systems (*14*). Surprisingly, the type I-D system contains *cas10d*, a variant of the signature type III *cas10* gene (*14*). However, the palm domain that is associated with secondary messenger production in the type III counterpart is inactivated within *cas10d*. Additionally, the signature type I gene *cas3* is split, with the nuclease domain (*cas3’’*) fused to *cas10d*, and the helicase domain (*cas3’*) encoded separately (*15*). However, the lack of biochemical and structural information on the type I-D complex remains a bottleneck for understanding its chimeric nature. Here, we show that type I-D Cascade from *Synechocystis* sp. PCC 6803 (hereafter *Synechocystis*) has stoichiometry similar to type I complexes and an overall architecture similar to type III complexes. Importantly, we uncover an alternative internal translational initiation site within *cas10d* that leads to the expression of the small subunit, Cas11d. We further demonstrate that internal translation of small subunits from within *cas10d* or *cas8* large subunit genes is conserved in type I CRISPR-Cas systems that lack a separate *cas11* gene.

### Type I-D Cascade includes an unexpected small subunit

To characterize type I-D Cascade, we co-expressed *cas3’, cas10d, cas7d, cas5d* and *cas6d* from *Synechocystis* with the first repeat-spacer-repeat of the associated CRISPR array in *Escherichia coli* (fig. 1A). After affinity pulldown of the His6-tagged Cascade, size-exclusion chromatography indicated the assembled type I-D Cascade eluted at a size corresponding to ~470 kDa (fig. 1B). SDS-PAGE revealed Cas10d (108 kDa), Cas7d (37 kDa), Cas5d (29 kDa) and/or Cas6d (28 kDa), plus an additional unexpected band at >14 kDa (fig. 1C). Cas3’ (82 kDa) did not co-purify with the complex, suggesting that recruitment of this helicase may first require target recognition by Cascade – similar to recruitment of full length Cas3 proteins during interference in other type I systems (*16, 17*). Type I-D Cascade co-purified with a crRNA of ~72 nucleotides (fig. 1D), consistent with the full spacer-repeat crRNA unit observed previously by RNA-seq and Northern blot in *Synechocystis (18, 19*). Mass spectrometry (*20*) confirmed the presence of full length Cas10d, Cas7d, Cas5d and Cas6d in the complex (Table S1 and data S1). Interestingly, MS also identified the low molecular weight protein to be the C-terminus of Cas10d (fig. 1E,F). Hereafter, we refer to this polypeptide as Cas11d due to its similarity to other Cas11 small subunits (*11*). The stoichiometry of type I-D Cascade was estimated via peptide abundances to be Cas10d1:Cas7d6:Cas5d1:Cas6d1:Cas11d2 (Table S2 and data S1). Taken together, these results demonstrate that type I-D Cascade is similar to other class 1 interference complexes, but uniquely includes an additional small subunit polypeptide derived from within the *cas10d* gene.

**Fig. 1.**
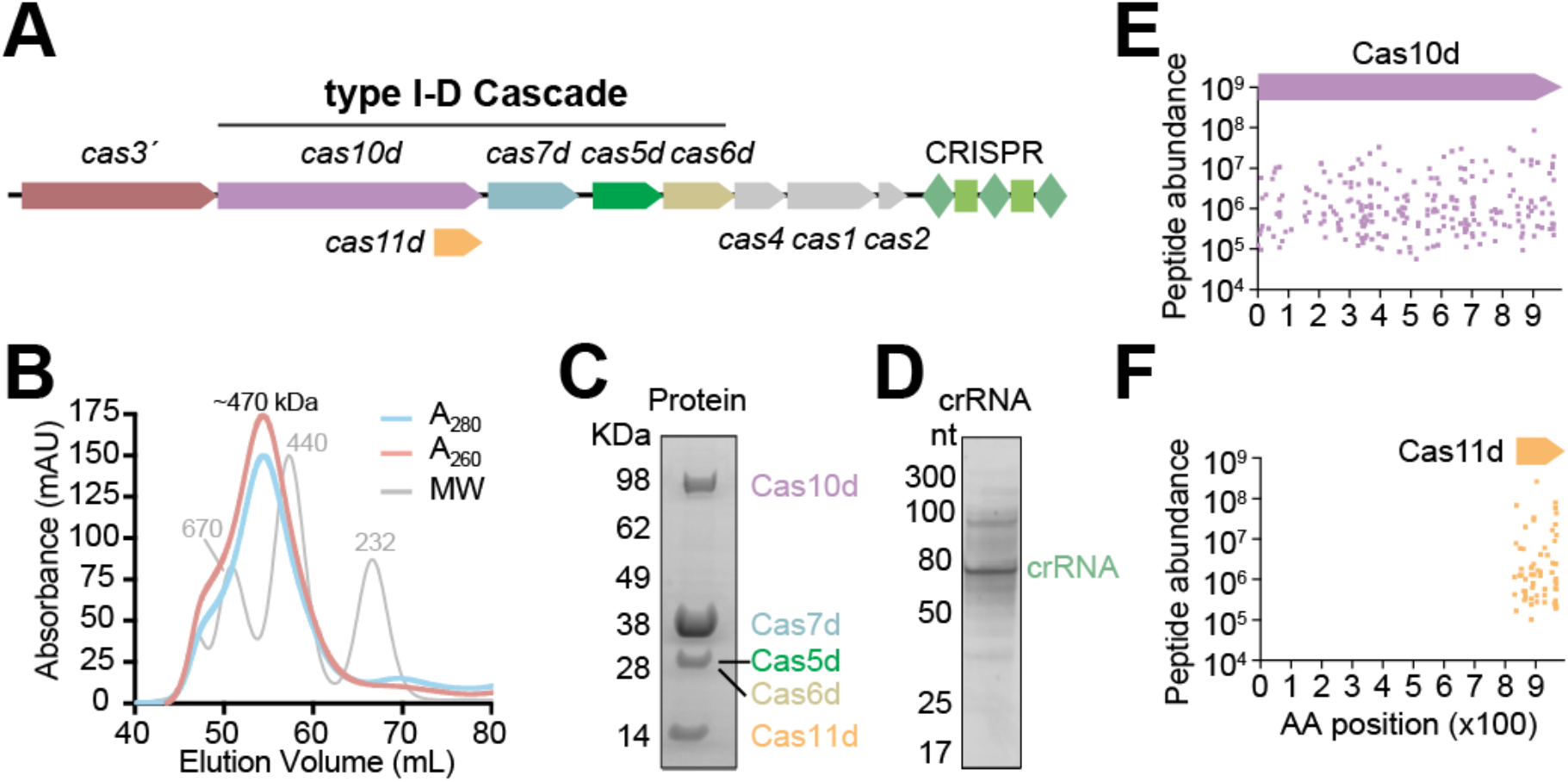
Type I-D Cascade contains a small subunit (Cas11d) derived from *cas10d*. (**A**) The *Synechocystis* sp. PCC 6803 type I-D locus. (**B**) Chromatogram of type I-D Cascade purified by SEC. (**C**) SDS-PAGE of type I-D Cascade. (**D**) crRNA co-purified with type I-D Cascade. MS analysis of peptides from (**E**) Cas10d and (**F**) Cas11d. Peptide abundance is plotted corresponding to position along Cas10d. The data presented in (**B**)-(**F**) are representatives of at least two replicates.

### Cas11d results from internal translation

We next investigated the origin of the Cas11d subunit. Multiple lines of evidence support internal translation as the source of Cas11d rather than proteolytic cleavage. Full length Cas10d was detected by MS in the complex, and the coverage of peptide abundance was consistent along the protein length (fig. 1E). Additionally, a defined Cas11d product was detected in the complex by MS (fig. 1F). The Cas11d N-terminus mapped to residue M830 of Cas10d by identification of peptides with either a non-tryptic or non-chymotryptic N-terminus by MS (fig. 2A and data S1). Translation from M830 is in-frame with *cas10d* and would result in a protein with an expected mass of 17.0 kDa – consistent with SDS-PAGE and MS (fig. 1C,F). This protein was not an artefact of the *E. coli* expression system, as Cas11d was also identified in samples from the native *Synechocystis* host, along with the other type I-D *cas* proteins (fig. S1). Eight bp upstream of the ATG that encodes M830 is a GA-rich region that resembles a ribosome binding site (RBS) (fig. 2A). *In silico* analysis predicted that this region is translationally-active and strongly expressed (ΔG_Total_ = −5.0 kcal/mol) (*21*). To investigate whether the predicted RBS and ATG start codon supported translation, a translation-reporter fusion system was established with the *cas11d* translation initiation site fused to *eyfp* (fig. 2B). As we predicted, eYFP was translated from the internal *cas11d* initiation region. Furthermore, mutations within the RBS and/or start codon, or a scrambled control, abolished expression (fig. 2B).

**Fig. 2.**
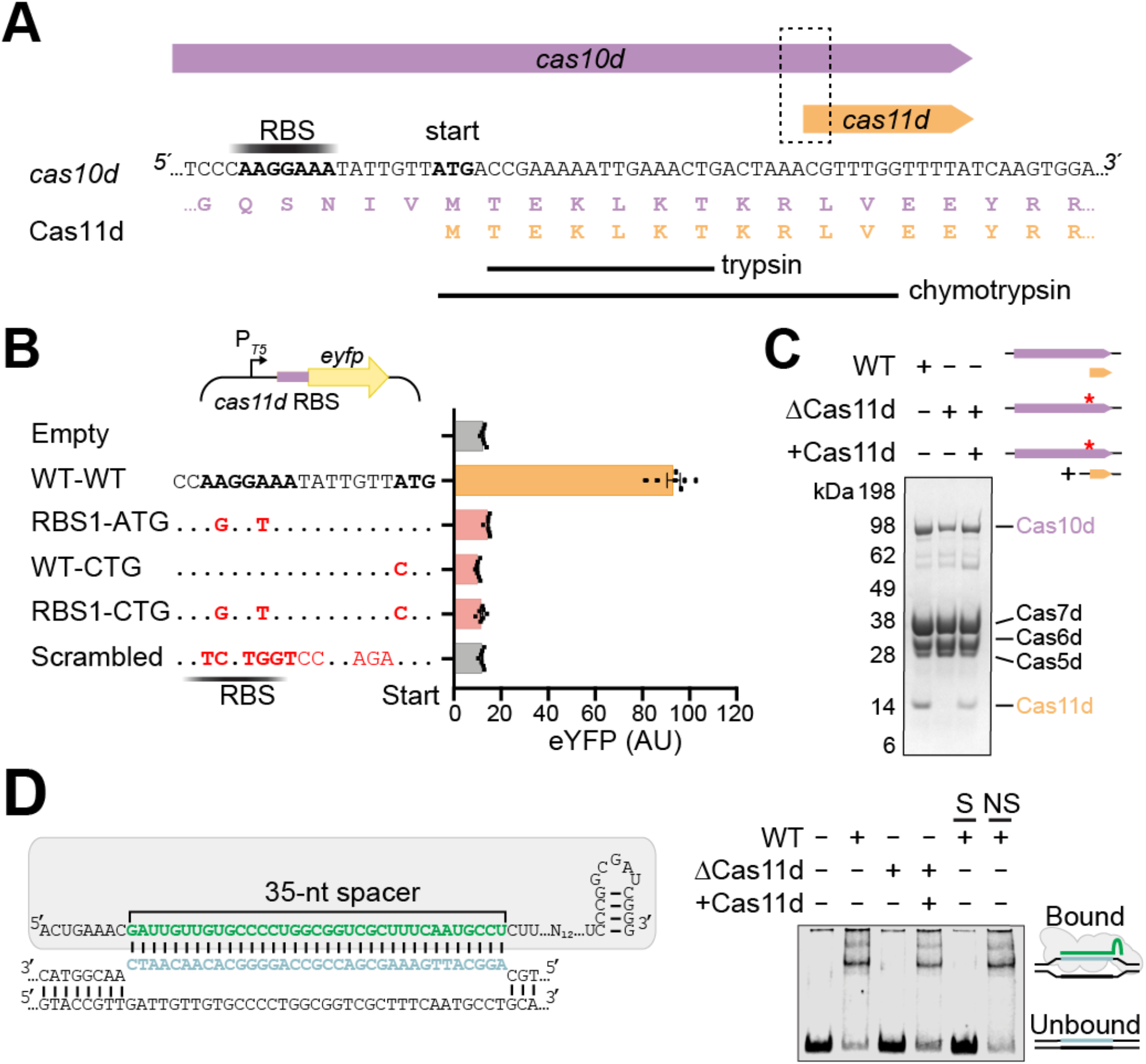
Cas11d is expressed from an internal translational initiation site in *cas10d*. (**A**) The *cas10d* and *cas11d* genes, including the translational start site of Cas11d. The N-terminal peptides from MS analysis after trypsin and chymotrypsin digests are shown. This was repeated with similar results. (**B**) Translation-reporter assay of *cas11d* RBS and start codon, with respective mutants. Error bars represent the standard error of the mean (n = 7). (**C**) Purified wild-type type I-D Cascade (WT), complex without Cas11d from attenuation of *cas11d* internal translation initiation (ΔCas11d), and ΔCas11d co-expressed with a complementary *cas11d-*containing plasmid (+Cas11d). The asterisks indicate the mutated *cas11d* RBS and ATG within *cas10d*. This experiment was repeated with similar results. (**D**) Schematic representation of type I-D Cascade (grey protein and green crRNA) binding to complementary target sequence (protospacer; blue) (left). Electrophoretic mobility shift assay involving type I-D Cascade variants binding to a specific fluorescently-labeled dsDNA protospacer (right). Non-fluorescent specific (S) and non-specific (NS) competitors are indicated. The data presented in (**A**)-(**D**) are representatives of at least two replicates.

Next, we investigated the role of internal translation of Cas11d in the formation of type I-D Cascade. Purification of the complex with a mutated *cas11d* RBS and ATG resulted in type I-D Cascade lacking Cas11d (ΔCas11d) (fig. 2C). The ΔCas11d complex displayed a similar Cas protein stoichiometry compared with the wild type complex by SDS-PAGE. We reasoned that since Cas11d is produced as a separate polypeptide, if produced *in trans*, it should associate with the complex lacking Cas11d. Indeed, complementation with *cas11d* expressed from a separate plasmid restored the protein constituents of the wild type complex (fig. 2C).

To test if the internal translation of Cas11d was important for the function of Cascade, we assessed DNA binding. Wild-type type I-D Cascade specifically bound dsDNA carrying a complementary protospacer (fig. 2D). In contrast, there was no detectable dsDNA binding by ΔCas11d. When ΔCas11d was complemented with *trans-*Cas11d, dsDNA binding was restored to levels similar to the WT complex. In agreement, the Cas11 small subunit (Cse2) from the type I-E system is important for the interaction of Cascade with DNA through stabilization of the R-loop formed during complex with target DNA (*9, 13, 16*). Thus, Cas11d is internally translated from *cas10d* and is required for specific binding of the type I-D Cascade complex to target dsDNA.

### Cas11d is analogous to the small subunit

To understand the architecture of type I-D Cascade, we employed electron microscopy on negatively stained complexes. Raw particles had an elongated shape which resembled a type III ‘sea worm’, rather than the typical type I ‘seahorse’ shape (fig. S2) (*10, 22*). The reference-free class averages of type I-D Cascade revealed a length of 250 Å and using *ab initio* reconstruction methods, we obtained a 3D structure at a resolution of ~19 Å (fig. 3A and fig. S2). The globular density at the foot of the complex is made up of the large subunit which can easily fit the structure of type III-B Cas10 (PDB: 3X1L) (*23*). The backbone of type I-D Cascade easily accommodated the crystal structure of type I-D Cas7d from *Thermophilum pedens* (PDB: 4TXD) (*24*) within the density for each of the backbone subunits. Cas6d and Cas5d densities both accommodated their type I-E counterparts (PDB: 4TVX and 6C66, respectively) (*25, 26*). Notably, the elongated shape of type I-D matches well with the type III-B complex, while the stoichiometry appears more consistent with the type I-E complex. Both type I-D and I-E have six Cas7 subunits, and two small subunits, while type III-B has three small subunits and four Cas7 subunits (fig. S2). Overall, the type I-D Cascade could be described as a hybrid of type I and type III CRISPR-Cas complexes (*10, 22, 27*).

**Fig. 3.**
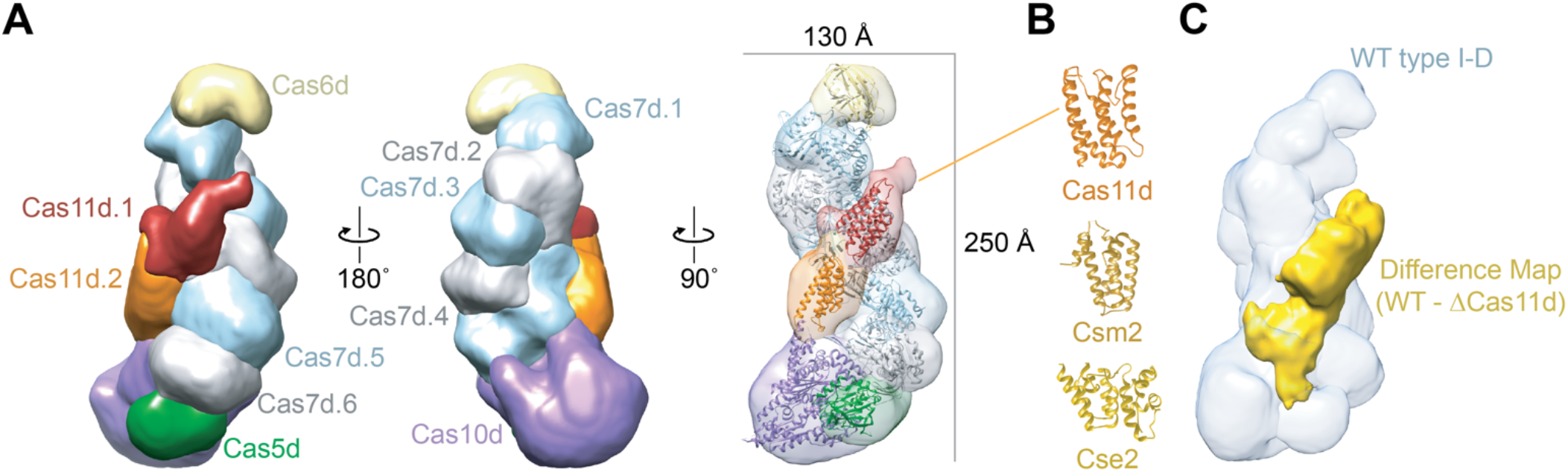
Cas11d assembles into the belly of type I-D Cascade analogous to a small subunit. (**A**) 3D reconstruction of type I-D Cascade at ~19 Å. The structure was segmented by subunit and colored as follows: Cas5d (*33*), Cas10d (purple), Cas7d (grey and light blue), Cas11d (orange and red), Cas6d (yellow). Type I-E Cas6e from *Escherichia coli* (PDB: 4TVX) (*25*), type III-B Cas10 from *Archaeoglobus fulgidus* (PDB: 3X1L) (*23*), type I-E Cas5e from *Thermobifida fusca* (PDB: 6C66) (*26*), and type I-D Cas7d from *Thermophilum pedens* (PDB: 4TXD) (*24*) were fit into their respective type I-D subunits. (**B**) A model for Cas11d was fit into the small subunit densities and is similar to *E. coli* Cse2 (PDB: 4TVX) (*25*) and *Staphylococcus epidermidis* Csm2 (PDB: 6AE1) (*32*) from type I-E and III-B complexes, respectively. (**C**) A difference map between ΔCas11d and WT complexes shows a density that overlays with the small subunits (yellow) in the original WT structure.

Interestingly, two smaller densities (orange and red) that run along the belly of the complex, parallel to the larger, helical backbone, remained unaccounted for after all other subunits were fit into the structure (fig. 3A). These smaller densities are ordered in a similar fashion to those of the Cas11 small subunits (Cse2, Csm2, and Cmr5) in the type I-E and III complexes (fig. S2) (*10, 22, 27, 28, 29*). Since we had identified an additional protein that was functionally similar to the small subunit, we hypothesized Cas11d would bind the type I-D Cascade in a manner analogous to the small subunits of other systems. To test this, we generated a homology model of Cas11d, which showed a six alpha-helical bundle, similar to other Cas11 proteins (fig. 3B) (*11, 30–32*). The homology model of Cas11d could easily fit into each of the two densities along the belly of the type I-D structure.

To test the predicted location of Cas11d in type I-D Cascade, we determined the structure of the ΔCas11d complex. Consistent with our prediction, the ΔCas11d complex lacked density for the two small subunits within the minor filament of the complex when compared with the WT structure. However, a protrusion from Cas10d is observed in the ΔCas11d complex that is likely the C-terminus of Cas10d, which we expect to structurally resemble Cas11d (fig. 3C and fig. S2). As predicted, when ΔCas11d was complemented with Cas11d, the density reappeared (fig. S2). Taken together, our results demonstrate that two Cas11d subunits are assembled into the complex and mimic the structural and functional role of the small subunits present in type I (-A and -E) and III systems.

### Internal translation of small subunits is widespread

We predicted the internal translation of the small subunit to be widespread in type I-D systems. To test this hypothesis, we created a phylogenetic tree of Cas10d proteins, and three groups emerged: two cyanobacterial and a third with mostly bacterial and archaeal species (fig. 4A). Analysis of Cyanobacteria lineage 1 revealed near complete conservation of a methionine start codon at the same position as M830 in *Synechocystis* (fig. 4B and data S2). Cyanobacteria lineage 2 and lineage 3, which encompasses bacterial and archaeal species, also displayed conservation of a translation start site (ATG, GTG or TTG) in similar positions to M830 in *Synechocystis* (fig. 4B and data S2). Furthermore, translationally active RBSs were predicted upstream of the proposed start codons, and translational-reporter assays using representatives from all three major groups showed that these regions were capable of initiating translation (fig. 4C). These results support that internal translation of Cas11d from within *cas10d* is conserved across type I-D CRISPR-Cas systems, including cyanobacteria and bacteria.

**Fig. 4.**
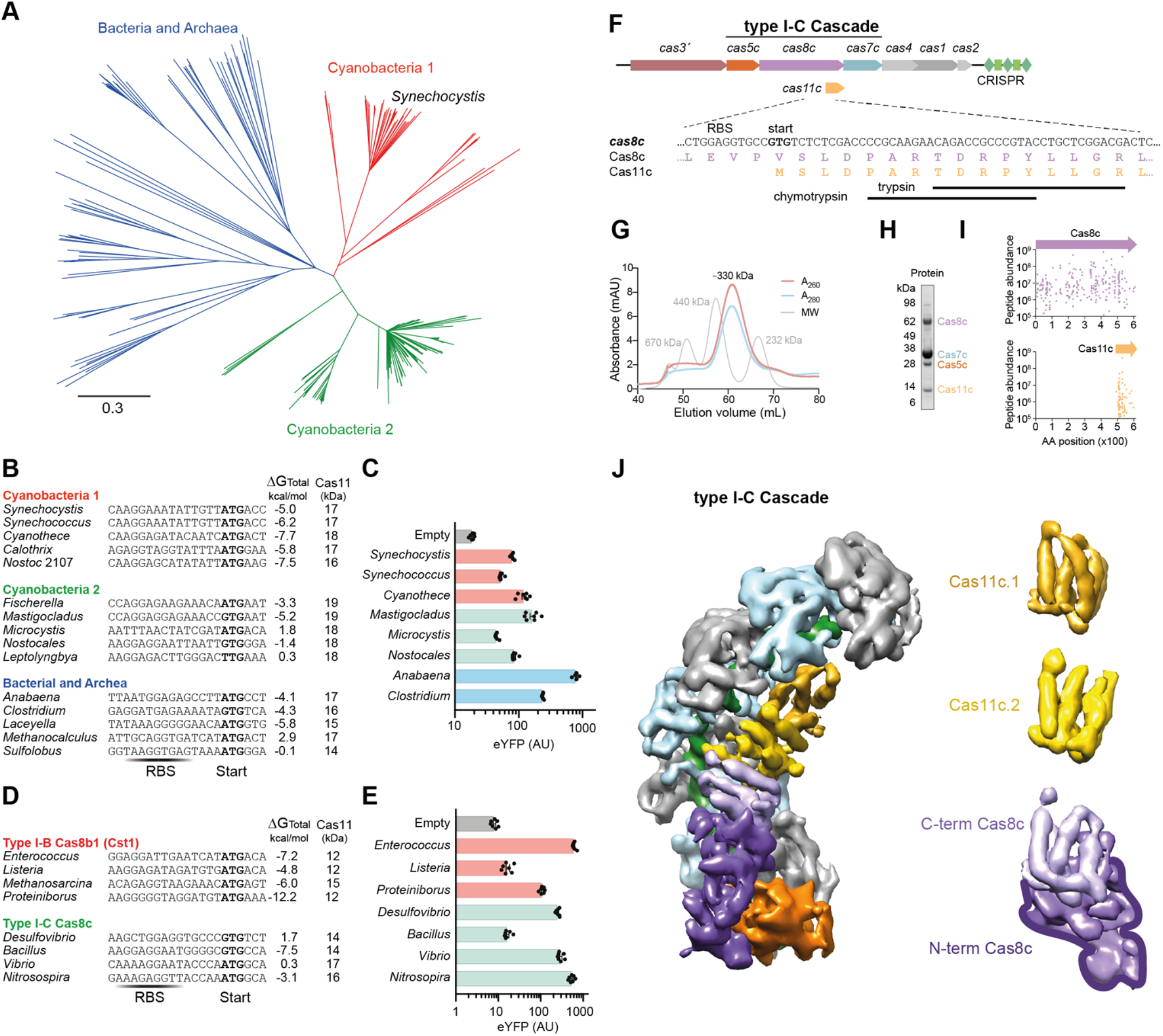
Internal translation of small subunits is conserved in multiple distinct type I CRISPR-Cas systems. (**A**) Maximum likelihood tree of Cas10d proteins grouped into three main lineages. (**B**) Alignment of selected *cas10d* sequences from the three lineages, with the potential start codon of Cas11d in bold and the RBS locations indicated. For each, the *in silico* predicted translation rates and mass of predicted Cas11d polypeptides is displayed. (**C**) Translation-reporter assay of Cas11d translation initiation regions. Error bars represent the standard error of the mean (n = 7). (**D**),(**E**) Similar to b,c, respectively, except analysis of selected *cas8b* and *cas8c* sequences from type I-B and I-C CRISPR-Cas systems (n = 7). (**F**) The *D. vulgaris* type I-C locus showing the *cas8c* and *cas11c* genes, including the translational start site of *cas11c*. The most N-terminal peptides identified by MS analysis after trypsin and chymotrypsin digests are displayed. (**G**) Chromatogram of type I-C Cascade purified by SEC. (**H**) SDS-PAGE of type I-C Cascade. (**I**) MS analysis of peptides from Cas8c and Cas11c. Peptide abundance is plotted corresponding to position along Cas8c. (**J**) Structure of a type I-C interference complex (*37*) segmented based on our results. Cas11c (yellow and gold) binds in the belly of the complex. The C-terminal domain of Cas8c (light purple) caps this filament at the foot.

Other type I systems (type I-B, I-C, I-F1 and I-G) have no *cas11* gene; instead, the C-terminus of the large subunit was predicted to be extended and functionally substitute for the small subunit (*7, 15, 34*). Therefore, we looked for evidence of internal translation of small subunits in other type I systems that lack a separate *cas11* gene. In contrast to the type I-D system, Cas8 is the large subunit in all other type I systems. Analysis of large subunit proteins revealed conservation of a potential translation start site near the C-terminus in both type I-B and type I-C systems (fig. 4D and data S2). Using both *in silico* predictions and translation reporter assays, we demonstrated that these regions within *cas8* supported translation initiation in representatives of type I-B and I-C systems (fig. 4E). In agreement with our findings for type I-B systems, a C-terminal portion of Cas8b containing an N-terminal methionine was previously detected; however, the authors proposed it resulted from proteolytic cleavage (*35*). Type I-F1 systems lack a *cas11* gene and alignment of Cas8f proteins did not detect a conserved internal translation start site, indicating internal translation initiation regions are not conserved in *cas8f* genes (data S2). This is corroborated by high-resolution type I-F1 Cascade structures that lack small subunits, but show a short minor filament due to the small subunit-like C-terminal extension of Cas8f (*36*). Type I-G (formally type I-U (*7*)) lack a *cas11* gene and alignment of Cas8u2 proteins did not reveal a conserved internal translation initiation regions (data S2).

To further investigate the association of internal translation of small subunits in type I complexes, we purified type I-C Cascade from *D. vulgaris* and discovered an additional 14 kDa protein, which we termed Cas11c (fig. 4F-J). Similar to the type I-D system (fig. 1E-F), MS revealed Cas8c was full length and Cas11c mapped to the C-terminus of the large subunit (fig. 4I and data S1), with the likely N-terminus being the highly-conserved M489 residue (fig. 4D,F and data S2). The predicted homology model of Cas11c revealed a six alpha helical bundle similar to other small subunits (fig. 4J). Indeed, the cryo-EM structure of type I-C Cascade possesses density where the minor filament formed by small subunits is expected to reside (*37*) (fig. 4J). The authors hypothesized that Cas8c (68 kDa) had an extended C-terminus that formed the minor filament as well as the foot of the complex. However, our data demonstrates that Cas11c subunits are translated as multiple copies from *cas8c* and contribute to the minor filament along the belly of type I-C Cascade. In summary, internal translation of small subunits from large subunit genes is widespread throughout type I-B, I-C, and I-D systems, which together comprise 23% of all CRISPR-Cas systems (*7*).

## DISCUSSION

CRISPR-Cas systems are divided into different types depending on their *cas* genes and their mode of CRISPR interference, which in turn has given rise to an expanding and broad range of novel genetic engineering tools (*7, 38, 39*). The type I-D CRISPR-Cas system appears to be a hybrid of type I and type III systems, yet the type I-D Cascade, involved in binding target nucleic acids, remained uncharacterized. Here, we present the first analysis of type I-D Cascade, which revealed similarities to both type I and III complexes and the inclusion of an unexpected small subunit protein expressed from an alternative internal translational initiation site within the large subunit gene. We demonstrate that the alternative expression of the small subunit is a widespread phenomenon among type I systems, which raises an unexpected aspect of CRISPR-Cas evolution with expanded coding potential within *cas* operons.

Previously, Makarova and colleagues proposed that type I systems that lack a *cas11* gene have an extended large subunit C-terminus, which might function as a small subunit by forming a shortened minor filament (*15*). This proposal was supported by high resolution structures of the type I-F1 system (*36*). Surprisingly, we discovered that three major type I systems that lack a separate *cas11* gene (I-B, I-C, and I-D), which account for ~23% of all CRISPR-Cas systems in microbes (*7*), have an internal translation initiation site within the large subunit gene that generates multiple copies of the Cas11 small subunit. Our findings are consistent with the prediction that the C-termini of large subunits exhibit structural similarity to small subunits (*15*), but greatly extends this by demonstrating that multiple Cas11 copies are derived via internal translation to form a full minor filament. We further reveal that small subunits produced from internal translation are functionally similar to the small subunits from other CRISPR-Cas systems in stabilizing Cascade-target DNA interactions. We speculate that having Cas11 fused to the C-terminus of the large subunit would aid in Cascade assembly by providing a ‘nucleation’ point for the separate Cas11 subunits to bind and complete extension of the minor filament.

Type I-D *cas* genes have similarities to both type I and type III systems, which might indicate that this CRISPR-Cas system is an evolutionary intermediate. The ancestor of CRISPR-Cas systems is proposed to be a relative of type III and, through replacement of the large subunit and gain of the Cas3 nuclease-helicase, resulted in type I systems (*7*). The absence of large-small subunit fusion genes in extant type III systems, yet their broad occurrence in type I systems, suggests that this fusion occurred after type I systems diverged. However, the absence of this fusion in some type I systems indicates it either reverted to two separate genes, or that fusion occurred independently multiple times. The internal translation of the small subunit from these fusions is likely to have facilitated the ease with which operon rearrangements could occur and still yield functional Cas complexes.

There has been significant recent interest in the biotechnological utilization of type I systems in both bacteria and eukaryotes. Internal translation in these type I systems needs to be considered when using CRISPR-Cas for applications in organisms with differing translational machinery, such as eukaryotes (*40, 41*). In these cases, *cas11* might need to be encoded from a separate gene. Importantly, we have shown that the coding capacity of CRISPR-Cas systems is richer than anticipated, with internal translation giving rise to alternative protein isoforms that are critical for bacterial adaptive immunity.

## ACKNOWLEDGMENTS

We thank Julian Eaton-Rye, University of Otago, for *Synechocystis* DNA and assistance growing *Synechocystis*. We are grateful to Torsten Kleffmann of the Centre for Protein Research, University of Otago, for MS assistance and Aguang Dai and the Sauer Structural Biology Lab, University of Texas at Austin for electron microscopy assistance and resources. We thank Sebastian Kieper, Cristobal Almendros, and Stan J.J. Brouns (Delft University of Technology, Netherlands) for early input and discussions. We thank members of the Taylor and Fineran laboratories for input.

## Author contributions

R.D.F. and P.C.F. conceived the study. T.M.M., E.A.S., D.W.T., P.C.F. and R.D.F. designed experiments. T.M.M. and R.D.F. performed the biochemical and genetic experiments. E.A.S. performed the structural studies. A.K. performed the mass spectrometry experiments. All authors analyzed the data. R.D.F., D.W.T. and P.C.F. secured funding and supervised the work. T.M.M., E.A.S., P.C.F. and R.D.F. wrote the manuscript with input from the other authors.

## Data and materials availability

The map for wild-type type I-D Cascade has been deposited in the EMDB with accession code EMD-21607. All other data are available in the main text or the supplementary materials.

## SUPPLEMENTARY MATERIALS

Materials and Methods

Figures S1-S2

Tables S1-S5

References (*42–56*)

Data S1 and S2

