## Supplementary Information 1 for "Diverse CRISPR-Cas complexes require independent translation of small and large subunits from a single gene"

Supplementary Information S1

### Internal translation of large subunit transcripts drives small subunit synthesis in type I CRISPR-Cas interference complexes

Tess M. McBride, Evan A. Schwartz, Abhishek Kumar, David W. Taylor, Peter C. Fineran, and Robert D. Fagerlund

#### Methods

##### Culture conditions

All strains and plasmids used in this study are listed in Supplementary Table 3 and 4, respectively, and the oligonucleotides are shown in Supplementary Table 5. Unless otherwise noted, *Escherichia coli* strains were grown at 37°C in Lysogeny Broth (LB), or on LB-agar (LBA) plates with 1.5% (w/v) agar. Media were supplemented with antibiotics when required as follows: ampicillin (Ap; 100 µg/mL), chloramphenicol (Cm; 25 µg/mL), and kanamycin (Km; 50 µg/mL). Growth was monitored as the optical density at 600 nm ($OD_{600}$) in a Jenway 6300 Spectrophotometer or in a Varioskan Flash Microplate Reader (Thermo Fisher Scientific). *Synechocystis* was grown at 30°C in bubbled BG-11 liquid medium supplemented with 5 mM glucose or BG-11 Agar supplemented with 1.5% agar, 5 mM glucose, 10 mM TES-NaOH (pH 8.2), and 3 g/L sodium thiosulfate. When appropriate, the medium was supplemented with kanamycin (25 µg/mL). Cultures were maintained at 30°C under constant illumination at 30 µE $m^{-2}s^{-1}$. Growth was monitored as the optical density at 730 nm ($OD_{730}$).

##### Generation of plasmids for type I-D expression and purification

A plasmid (pPF1549) for expression of Cas10d, Cas7d and Cas5d was constructed by PCR-amplifying their genes (primers PF2927+PF2932) using *Synechocystis* genomic DNA as template and cloning the product into pQE-80LoriT via BamHI and PstI restriction sites. The *cas10d* gene was cloned to incorporate an N-terminal $His_6$ tag followed by TEV protease recognition sequence. A plasmid (pPF1589) for expression of Cas3', Cas10d, Cas7d and Cas5d was constructed by PCR-amplifying *cas3'* (primers PF2933+PF2934) using *Synechocystis* genomic DNA as template, digesting with PstI and NsiI and cloning the product into pPF1549 via PstI restriction site. Overlap extension PCR was used to generate a plasmid (pPF1758) for expression of Cas10d, Cas7d and Cas5d, but with site-directed mutations to *cas10d* to interrupt the internal *cas11d* RBS and start codon in the Cas10d-Cas7d-Cas5d expression construct. The upstream PCR fragment was generated using pPF1549 with primers PF2927 and PF3420. The downstream PCR fragment was generated using pPF1549 with primers PF3421 and PF2932. Both fragments were purified and used as template in an overlap extension PCR with primers PF2927 and PF2932 and the resulting product was digested with EcoRI and HindIII and ligated into pQE-80LoriT, previously digested with the same enzymes.

A plasmid (pPF1538) for expression of the first spacer and flanking repeat sequences from the type I-D associated CRISPR array was constructed by PCR-amplifying this region from *Synechocystis* genomic DNA (primers PF2937+PF2938) and cloning the product into pACYCDuet-1 via NdeI and KpnI restriction sites. Three plasmids (pPF1539, pPF1552, and pPF1926) were constructed for expression of Cas6d with the first spacer and flanking repeat sequences. Plasmid pPF1539 expresses Cas6d with no fusion tag and was constructed by PCR-amplifying *cas6d* (primers PF2935+PF2936) using *Synechocystis* genomic DNA as template and cloning the product into pPF1538 via NcoI and HindIII restriction sites. Plasmid pPF1552 expresses Cas6d with an N-terminal $His_6$ tag and TEV protease recognition sequence and was constructed by PCR-amplifying *cas6d* (primers PF3027+PF2936) using *Synechocystis* genomic DNA as template and cloning the product into pPF1538 via BamHI and HindIII restriction sites. Plasmid pPF1926 expresses Cas6d with an N-terminal StrepII-tag and TEV protease recognition sequence and was constructed by cloning annealed oligonucleotides PF3760 and PF3761 into pPF1552 via restriction sites NcoI and BamHI.

A plasmid (pPF2034) for separate expression of Cas11d was constructed by PCR-amplifying the appropriate region of *cas10d* (primers PF3935+PF3721) using *Synechocystis* genomic DNA as template and cloning the product into pPF1720 via EcoRI and SpeI restriction sites. Plasmid pPF1720 was constructed by ligating the T5/LacO promoter fragment resulting from digestion of pSEVA184f with PacI and EcoRI with a similarly digested plasmid pSEVA251.

A plasmid (pPF1963) for expression of Cas5 fused with an N-terminal $His_6$ tag in *Synechocystis* was constructed by PCR-amplifying gBlock PF3648 (Integrated DNA Technologies) with primers PF3771 and PF3772 and cloning the product into EcoRI and PstI sites of pPF1720 using NEBuilder HiFi DNA Assembly (NEB).

**Expression and purification of recombinant type I-D complexes**

Type I-D Cascade with N-terminal $His_6$-tagged Cas10d was expressed in BL21(DE3) cells containing plasmids pPF1549 and pPF1539. Five hundred mL cultures were induced with 1 mM IPTG at $OD_{600}$ = 0.6 and grown overnight at 18°C. Cells were harvested at 10,000 x g for 10 min. The cell pellet was resuspended in 20 mL of 300 mM KCl lysis buffer (50 mM HEPES-NaOH, pH 7.5, 300 mM KCl, 5% Glycerol, 1 mM DTT, 0.02 mg/mL DNaseI, cOmplete EDTA free protease (Roche) and 0.1 mM of PMSF) supplemented with 10 mM imidazole. Cells were lysed by a French pressure cell press (American Industry Company) at 10,000 psi, and the lysate was clarified by centrifugation at 15,000 x g for 15 min. The complex was affinity purified using a HisTrap™ HP (GE Healthcare) column equilibrated in 300 mM KCl binding buffer (50 mM HEPES-NaOH, pH 7.5, 300 mM KCl, 5% Glycerol, 1 mM DTT) and eluted using a gradient against binding buffer containing 500 mM imidazole. The $His_6$ tag on Cas10d was removed by cleavage with TEV protease during overnight dialysis at 4°C in binding buffer. The liberated $His_6$ tag and non-specific *E. coli* proteins were removed using a second HisTrap affinity column and the flow through was collected. The sample was concentrated with a centrifugal concentrator (Amicon; 100 kDa molecular weight cut off (MWCO)) and further purified from free Cas10d by size exclusion chromatography (SEC) on a HiLoad 16/600 Superdex 200 (GE Healthcare) column equilibrated in SEC Buffer (10 mM HEPES-NaOH, pH 7.5, 100 mM KCl, 5% Glycerol, 1 mM DTT).

Type I-D Cascade with N-terminal StrepII-tagged Cas6d was expressed in LOBSTR cells containing plasmids pPF1549, pPF1926 and pPF1720. Cultures were induced with 1 mM IPTG at $OD_{600}$ = 0.6 and grown overnight at 18°C. Cells were harvested at 10,000 x g for 15 min. The cell pellet was resuspended in 20 mL of 100 mM KCl lysis buffer (lysis buffer as above but with 100 mM KCl) and cells lysed by French press at 10,000 psi. The lysate was clarified by centrifugation at 15,000 x g for 15 min and the lysate was applied to two 1 mL columns of Strep-tactin resin (IBA) equilibrated in 100 mM KCl binding buffer (lysis buffer as above but with 100 mM KCl). The complex was eluted with binding buffer supplemented with 2.5 mM desthiobiotin. SEC was used to separate free StrepII-Cas6d from the complex on a HiLoad 16/600 Superdex 200 (GE Healthcare) column equilibrated in SEC Buffer. The same approach was used for cells containing plasmids pPF1758, pPF1926 and pPF1720 (complex with arrested Cas11d expression), and pPF1758, pPF1926 and pPF2034 (complex with arrested Cas11d expression with Cas11d expressed from a separate plasmid).

Type I-D Cascade with N-terminal $His_6$-Cas6d and N-terminal $His_6$-Cas10d was expressed in LOBSTR cells containing plasmids pP1549, pPF1552 and pPF1720. Cultures were induced with 1 mM IPTG at OD600 = 0.6 and grown overnight at 18°C. Cells were harvested at 10,000 x g for 15 min. The cell pellet was resuspended in 20 mL of 100 mM KCl lysis buffer supplemented with 20 mM imidazole and cells lysed by French press at 10,000 psi. The lysate was clarified by centrifugation at 15,000 x g for 15 min and the lysate was applied to a HisTrap affinity column equilibrated in 100 mM KCl binding buffer and eluted using a gradient against binding buffer containing 500 mM imidazole. SEC separated free $His_6$-Cas6d and $His_6$-Cas10d from the complex on a HiLoad 16/600 Superdex 200 column equilibrated in SEC Buffer. The same approach was used for cells containing plasmids pPF1758, PF1552 and pPF1720 (complex with arrested Cas11d expression), and pPF1758, pPF1552 and pPF2034 (complex with arrested Cas11d expression with Cas11d expressed from a separate plasmid). Purified complexes were typically concentrated to 2 mg/mL using a centrifugal concentrator (Amicon; 100 kDa MWCO), aliquoted and stored at -80˚C.

##### *In vivo* crRNA isolation

The crRNA was isolated from the purified type I-D complex via a phenol/chloroform extraction, ethanol precipitation, and resolved on a denaturing gel containing 15% (v/v) 19:1 polyacrylamide, 7 M urea, and 0.5 x TBE (45 mM Tris, 45 mM borate, 1 mM EDTA, pH 8.3). The gel was stained with SYBR gold (Thermo Fisher Scientific) and crRNA was visualized using the Odyssey Fc imaging system (LICOR).

##### *Synechocystis* transformation and type I-D protein purification

To express N-terminal $His_6$-tagged Cas5 in *Synechocystis*, pPF1963 was introduced by electroporation. Wild type *Synechocystis* was grown to exponential phase ($OD_{730}$ ~0.5) and washed three times by centrifugation at 2,760 x g for 10 min and resuspension in 1 mM HEPES-NaOH (pH 7.5). Cells (60 µL) were then mixed with 3 µg of plasmid and electroporated at 12 kV/cm, 25 µF, 400 Ω using a gene pulser (Bio-Rad) with a 0.1 cm electroporation cuvette. A volume of 400 µL BG-11 media was immediately added. Cells were incubated on BG-11 agar without kanamycin for 24 h and then transferred to BG-11 agar with kanamycin. Single colonies were streaked onto BG-11 plates with kanamycin.

A .5 L BG11 liquid culture was inoculated with a starter culture of *Synechocystis* containing pPF1963 and was grown to an $OD_{730}$ of 0.3, $His_6$-Cas5 protein expression was induced with 1 mM IPTG, and cells were grown for approximately 24 hrs to a final $OD_{730}$ of 1.5. Cells were harvested at 10,000 x g for 10 min. The cell pellet was resuspended in 20 mL of 100 mM KCl lysis buffer supplemented with 10 mM imidazole. Cells were lysed by a French pressure cell press at 10,000 psi, and the lysate was clarified by centrifugation at 15,000 x g for 15 min. The complex was affinity purified using a HisTrap$^{TM}$ HP column equilibrated in 100 mM KCl binding buffer supplemented with 20 mM imidazole and eluted with binding buffer containing 300 mM imidazole. The eluted sample was concentrated with a centrifugal concentrator (Amicon; 100 kDa MWCO) and further purified by SEC on a HiLoad 16/600 Superdex 200 (GE Healthcare) column equilibrated in SEC Buffer. Fractions that eluted between 240 – 740 kDa (46 – 67 mL elution volume) were pooled and concentrated with a centrifugal concentrator (Amicon; 100 kDa MWCO).

**Label-Free shotgun mass spectrometry**

Purified type I-D and I-C Cascade were separated on an SDS-PAGE gel and protein bands were excised for mass spectrometry identification. For type I-D Cascade, an entire lane was excised for stoichiometry calculation between the different Cas subunits. The excised lane and bands were subjected to reduction, alkylation and digested with either trypsin or chymotrypsin using a DigestPro liquid handling workstation (Invatis AG, Germany). The extracted peptides were dried with a centrifugal vacuum concentrator (Savant, France). The dried peptides were reconstituted in 5% acetonitrile (ACN), 0.1% Formic acid (FA) in water and loaded into the UltiMate™ 3000 RSLC nanoflow uHPLC system (Thermo Scientific, USA) inline coupled to an LTQ Orbitrap XL hybrid MS system (Thermo Scientific, USA). The peptides were separated on an emitter-tip column ((New Objective, USA), 75 µM ID, 20 mm length) filled with Luna C18 bead material ((Phenomenex, USA), 3 µM particle size) using a reverse phase gradient between mobile phase A (0.1% FA in water) and B (0.1% FA in ACN). The gradient comprised of the following steps: 1) 5% B to 25% B over 136 min; 2) 25% B to 45% B over 20 min; 3) 45% B to 99% B over 4 min. The column was washed at 99% B for 1 min and re-equilibrated to 5% B.

In the mass spectrometer, MS1 precursor ion scan was performed in the Orbitrap FTMS mass analyzer at a resolution of 60,000 at 400 m/z value. The MS1 scan was performed in a scanning range of 400 to 2,000 m/z. The 11 strongest precursor ions from the MS1 scan were selected for a MS2 scan in the LTQ ion-trap mass analyzer at a normalized collision energy of 35%. Dynamic exclusion was enabled for the MS2 scans allowing for 2 repeat counts within 90 s with an exclusion duration of 120 s.

Raw MS data were processed through the analysis pipeline in the Proteome Discoverer software (version 2.4, Thermo Scientific, USA). Database dependent searches were carried out with the Sequest HT (Thermo Scientific, USA) search engine node in Proteome Discoverer against an in-house database containing the sequences of type I-D and I-C Cascade proteins. Two different cleavage settings were used in the SequestHT: 1) tryptic or chymotryptic, for identification and determining protein stoichiometry; 2) non-specific cleavage, for detecting the N-terminus of Cas11d and Cas11c. Individual Cas subunits were searched with either trypsin or chymotrypsin, depending on the cleavage enzyme used for band processing. Further common search settings allowed for carboxyamidomethyl cysteine, oxidized methionine, and deamidated asparagine and glutamine as variable modifications. The mass tolerance threshold was 10 ppm and 0.8 Da for precursor and fragment ions, respectively. Type I-D Cascade subunit stoichiometry was determined by comparing the average peptide peak area or abundance of each protein relative to the other proteins. The abundance of Cas10d was determined from peptides mapping to residues 1-830, while the abundance of Cas11d was determined from peptides mapping to residues 830-975 of Cas10d. The stoichiometry of Cas11d requires subtraction of one copy to account for the contribution from the C-terminus of Cas10d.

**Parallel reaction monitoring (PRM) mass spectrometry**

Purified type I-D Cascade from *Synechocystis* was separated on an SDS-PAGE gel and protein bands of interest were excised and digested as described in the label-free MS procedure. The dried peptides were reconstituted in 5% acetonitrile (ACN) and 0.1% Formic acid (FA) in water and loaded into an Eksigent ekspert 415 nanoflow uHPLC system (AB Sciex, USA) inline coupled to a Triple TOF 5600+ MS system (AB Sciex, USA). The peptides were separated on a Luna C18 capillary column (similar to the label-free MS procedure) using a reverse phase gradient between mobile phase A (0.1% FA in water) and B (0.1% FA in ACN) with the following steps: 1) 3% B to 22% B over 18 min; 2) 22% B to 38% B over 5 min; 3) 38% B to 97% B over 5 min. The column was washed at 97% B for 1 min and re-equilibrated to 3% B. The mass spectrometer was operated in the parallel reaction monitoring

(PRM) mode, selecting for 8 to 10 precursor mass targets at an accumulation time of 0.12 s resulting in a cycle time of 1.5 s. Suitable precursor target peptides were selected based on the results of *in silico* tryptic digestion and manual evaluation of peptide sequences and as well as the information retrieved from previous shotgun proteomics analyses of the same set of proteins.

The PRM raw data was converted to the mgf file format and initially searched with the Mascot search engine, version 2.5 (Matrix science, UK)[42] with a peptide mass tolerance of +/- 50 PPM and a fragment mass tolerance of +/- 0.1 Da. Carbamidomethylation (57.02) was selected as a static modification for cysteine. The data was searched on an in-house database containing the Cas protein sequences. Further, quantitative analyses were performed with the Skyline software, version 19.1 (https://skyline.ms, University of Washington, USA)[43]. The extracted ion chromatogram and the MS/MS spectra were manually evaluated for their fragment peak alignment, expected retention time and the correct fragment ion peak assignment in the MS/MS spectrum.

#### Electrophoretic mobility shift assays

A plasmid (pPF1836) carrying the protospacer of the type I-D CRISPR array spacer 1, flanked by a 5´-GTT-3´ PAM was constructed by ligating annealed oligonucleotides PF3029 and PF3030 into pPF1720 via KpnI and SphI restriction sites.

A 134-bp fluorescently labeled dsDNA probe containing the protospacer sequence complementary to the crRNA spacer of purified type I-D Cascade was amplified by PCR using primers PF4095 and PF4096 from template plasmid pPF1836. The non-fluorescent specific and non-specific probes were amplified using primers PF4092 and PF4093 from plasmid templates pPF1836 and pPF1720, respectively. Binding assays were performed with 250 nM type I-D Cascade variants (WT, ΔCas11d, or ΔCas11d with the Cas11d complement). Cascade complexes were incubated with 10 nM fluorescently labeled dsDNA at 30°C for 20 minutes in a total volume of 10 µL (final conditions: 10 mM HEPES-NaOH, pH 7.5, 100 mM KCl, 5% v/v glycerol, 1 mM DTT, 0.01% v/v triton X-100, 1 µg BSA, and 0.1 µg poly(dI.dC)). For the specific and non-specific controls, WT type I-D Cascade was incubated with 250 nM non-labeled probes (25x excess compared to labeled probe) at 30°C for 20 min prior to incubation with the fluorescently labeled specific probe. Final reactions were separated on 4% polyacrylamide (19:1 acrylamide:bisacrylamide) native gel containing 0.5x TBE at 4˚C.

Fluorescent probe was imaged using the Odyssey Fc imaging system (LICOR) and results were analyzed with Image Studio Lite software.

#### Negative stain electron microscopy

Three hundred mesh Cu continuous carbon grids (Ted Pella Inc.) were glow discharged for 2 minutes. Four µL of sample at ~100 nM was applied to the grid. After 1 minute, the sample was blotted and immediately washed in six ~30 µL droplets of 2% UA at pH 4.5 for 5-6 seconds each. The grid was then blotted and air dried. Images were taken using an FEI Talos transmission electron microscope operated at 200 keV. Images were collected using a CETA[44] detector at a nominal magnification of 73,000x (2.05 Å/pixel) with a defocus range of –1.5 µm to –2.5 µm.

#### 3D reconstruction

Micrographs were uploaded to Leginon[45] and Appion pre-processing software[46] was used to manually pick particles in 10 micrographs. Initial 2D class averages were used as templates for automatic particle picking using FindEM[47] with a threshold of 0.35, resulting in an initial data set of 18,696 particles. Box files were imported into RELION 3.0[48] for particle extraction with a box size of 200 pixels (2.05 Å/pixel). Reference-free 2D alignment and classification using 100 classes was used to separate unassembled complexes. After particle filtering, a new subset of 5,598 particles was extracted and uploaded into cryoSPARC v1[49]. Initial 3D models were reconstructed *ab initio* and one model (2,284 particles) was selected for further refinement. The final model obtained via homogeneous refinement was estimated to be at ~19 Å resolution using the 0.143 gold-standard criterion. This model was segmented using Segger[50] and X-ray crystal structures and homology models were fit into the map using "Fit-in-map" in UCSF Chimera v1.13[51].

#### Difference map creation

Newly purified complexes were used to reconstruct 3D models of the wild type (WT) complex, the complex lacking Cas11d (ΔCas11d), and the complex lacking Cas11d supplemented with Cas11d expressed from a separate plasmid (ΔCas11d + Cas11d). To create difference maps, the WT structure and the ΔCas11d structure were aligned in UCSF Chimera using the "Fit-in-map" feature. The VOP command in UCSF Chimera was used subtract the ∆Cas11d structure from the WT structure. The resulting difference map was then overlaid on the initial WT structure using the UCSF Chimera "Fit-in-map" feature[51].

#### Protein modeling

Homology modeling of *Synechocystis* Cas11d and *D. vulgaris* Cas11c was performed using the Robetta web server (http://robetta.bakerlab.org/) using *Synechocystis* Cas10d residues 830-975 (NCBI: WP_011153680.1) and *D. vulgaris* Cas8c residues 489-612 (NCBI: YP_009171.1) as a template, respectively.

#### Phylogenetic analysis

Protein sequences were collected from NCBI GenBank. The number of unique selected sequences analyzed of the large subunits for each system were (search criteria; selected protein length): 288 Cas10d sequences (Cas10d and manual addition of CscA from *Sulfolobus islandicus*; between 500 and 1,200 amino acids) for type I-D; 258 Cas8b1 sequences (Cst1; between 300 and 650 amino acids) for type I-B, 1769 Cas8c sequences (Cas8c; between 450 and 750 amino acids) for type I-C, 1548 Cas8f sequences (Csy1; between 300 and 600 amino acids) for type I-F1, and 93 selected Cas8u2 sequences obtained from a PSI-BLAST using Cas8u2 from *Geobacter sulfurreducens* (NCBI: GSU0052) for type I-G (formally I-U). Nucleotide sequences of selected *cas10d*, *cas8b,* and *cas8c* genes were collected from NCBI GenBank. Refer to Supplementary Dataset for complete list of sequences analyzed and alignments. Protein and nucleotide analyses were executed within Geneious Prime 2020.0.5 (https://www.geneious.com). Protein sequences were aligned by MUSCLE[52] with default parameters. A phylogenetic tree was constructed using the Neighbor-Joining method[53] and Jukes-Cantor distance model[54] with 1,000 replicates of bootstrap test.

#### In silico prediction of translation initiation rates

Translation initiation rates were predicted using the RBS Calculator (https://salislab.net/software/)[55] with 100 bp of the interested sequence, including 60 bp upstream of the potential start codon.

The following species were abbreviated in Figure 4: *Synechocystis, Synechocystis* sp. PCC 6803; *Synechococcus*, *Synechococcus* sp. PCC 7002; *Cyanothece*, *Cyanothece* sp. PCC 8802; *Calothrix*, *Calothrix* sp. PCC 7507; *Nostoc* 2107, *Nostoc carneum* NIES-2107; *Fischerlella*, *Fischerella muscicola* CCMEE 5323; *Mastigocladus*, *Mastigocladus laminosus* UU774; *Microcystis*, *Microcystis aeruginosa* PCC 9807; *Nostocales*, *Nostocales cyanobacterium*; *Leptolyngbya*, *Leptolyngbya* sp. PCC 7375; *Anabaena*, *Anabaena* sp. WA102; *Clostridium*, *Clostridium botulinum* strain 51714-DC; *Laceyella*, *Laceyella sediminis* strain RHA1; *Methanocalculus*, *Methanocalculus* sp. 52_23; *Sulfolobus*, *Sulfolobus islandicus*; *Enterococcus*, *Enterococcus* sp. 9E7_DIV0242; *Listeria*, *Listeria seeligeri* strain 7KSM; *Methanosarcina*, *Methanosarcina* sp. WH1; *Proteiniborus*, *Proteiniborus* sp. DW1; *Desulfovibrio*, *Desulfovibrio vulgaris* RCH1 plasmid pDEVAL01; *Bacillus*, *Bacillus halodurans* C-125; *Vibrio*, *Vibrio cholerae* strain RFB05; *Nitrosospira*, *Nitrosospira* sp. Nsp22.

**Generation of translation reporter plasmids**

Plasmids for reporter assays to characterize the *Synechocystis* Cas11d translational initiation site (pPF1914 – pPF1919 and pPF2116 – pPF2119) were constructed by fusing sequence relating to the translational initiation site with *eyfp* by PCR with forward primer (PF3722 – PF3727, PF4120 – PF4122, and PF4140, respectively) and reverse primer (pPF3721) using pSEVA237Y as template and cloning the product into pPF1720 via EcoRI and SpeI restriction sites. Similarly, plasmids for reporter assays to test translation from potential *cas11* translation initiation sites of other type I systems (pPF2120 – pPF2135) were constructed by amplifying forward primer (PF4124, PF4125, PF4129 – PF4131, PF4133, PF4134, PF4138, PF4143, PF4163, PF4164, PF4253, PF4313 and PF4314, respectively) and reverse primer (pPF3721) using pSEVA237Y as template and cloning the product into pPF1720 via EcoRI and SpeI restriction sites.

**Fluorescent reporter assays**

Plasmids to determine the translational start site efficiencies were comprised of potential *cas11* RBS and translational start sites fused to *eYFP*, under the control of a T5 promotor (pPF1914 – pPF1919, pPF2116 – pPF2123, and pPF2125 – pPF2134). Starter cultures of *E. coli* DH5$\alpha$ with the appropriate plasmids were grown at 37°C overnight. The cells were pelleted and resuspended to an $OD_{600}$ of 0.05 in LB containing Km and 50 µM IPTG. The cultures were grown at 37°C in the Varioskan LUX multimode microplate reader and the $OD_{600}$ and florescence intensity (excitation 513 nm, emission 531 nm) was measured after 24 h. Seven replicates of each culture were measured with scrambled (pPF2119) and/or empty vector (pPF1720) as the negative control(s). All measurements were of distinct samples. All fluorescence values were normalized to the $OD_{600}$ and plotted relative to the average fluorescence of the *Synechocystis cas11d* translation initiation reporter.

**Purification of the type I-C complex**

The type I-C complex was expressed and purified using a method similar to that described by Hochstrasser and colleagues[56]. Briefly, the type I-C interference complex with an N-terminal $His_6$-MBP(maltose-binding protein)-tagged Cas5 was expressed in LOBSTR cells containing plasmids expressing *D. vulgaris* Cascade/I-C (Cas5c-Cas8c-Cas7)/pHMGWA (Addgene plasmid # 81185 ; http://n2t.net/addgene:81185 ; RRID:Addgene_81185) and *D. vulgaris* sp2 CRISPR/pACYCDuet-1 (Addgene plasmid # 81186 ; http://n2t.net/addgene:81186 ; RRID:Addgene_81186), both gifts from Jennifer Doudna. Cultures were induced with 1 mM IPTG at $OD_{600}$ = 0.6 and grown overnight at 18°C. Cells were harvested at 10,000 x g for 15 min. The cell pellet was resuspended in 20 mL of 100 mM KCl lysis buffer and lysed by French press cells at 10,000 psi. The lysate was clarified by centrifugation at 15,000 x g for 15 min and the lysate was applied to a HisTrap affinity column equilibrated in 100 mM KCl binding buffer and eluted using a gradient against binding buffer containing 300 mM imidazole. The $His_6$-MBP tag on Cas5 was removed by cleavage with TEV protease during overnight dialysis at 4°C in binding buffer. The liberated $His_6$-MBP tag and non-specific *E. coli* proteins, were removed using a second HisTrap affinity column and the flow-through was collected. The sample was concentrated with a centrifugal concentrator (Amicon; 100 kDa MWCO) and further purified from free Cas5 by SEC on a HiLoad 16/600 Superdex 200 (GE Healthcare) column equilibrated in SEC Buffer.

**Data availability statement**

The data that support the findings of this study are available from the corresponding author upon request.

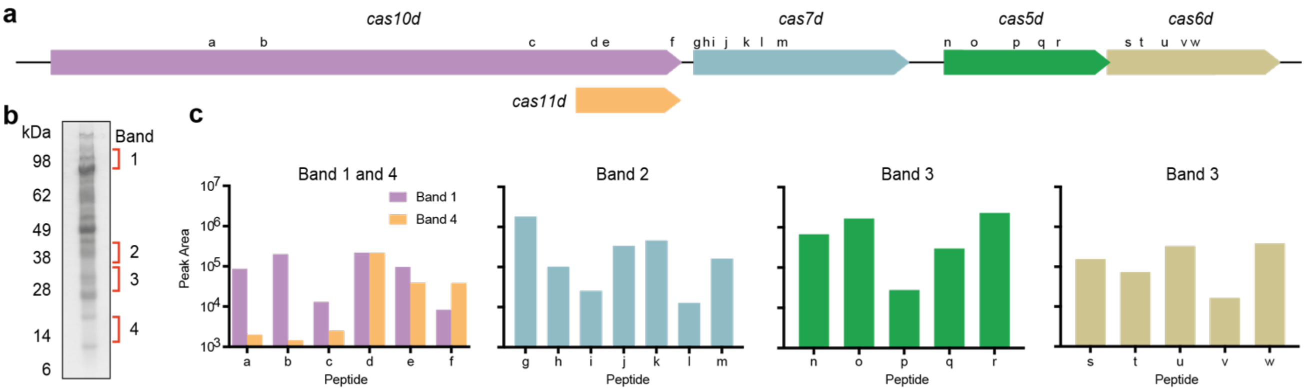


d

| ID | PEPTIDE SEQUENCE | ID | PEPTIDE SEQUENCE | ID | PEPTIDE SEQUENCE | ID | PEPTIDE SEQUENCE |
|---|---|---|---|---|---|---|---|
| a | DTTGILSNAIHNVILR | g | SQFQPSFPR | n | ALHSQVLEWLK | s | LTLHDNVFFASR |
| b | VSPQTSELLAAADIVR | h | LASGHYVHFLMLR | o | IGEPSLAEELHQSQISPFSISPLIGK | t | GTIIPHPYGLVGQNAQTPAYLDR |
| c | VEDITPALNK | i | HSQSFPVFQTDGVLNTTR | p | ISLLNGSLLQPLLK | u | GATKNYPINYGR |
| d | FYQVELSK | j | TQAGLLEKTDQLSR | q | LCHIHILPGSHSLAR | v | IRVEVEAIAPDQIK |
| e | KPSTHAILLPLSK | k | NLGLTSADK | r | ASSYALTTQAPTSSK | w | VEVEAIAPDQIK |
| f | LLALEAQQNQN | l | QCPDCILYGFAIGDSGSER | | | | |
| | | m | VYSDSAFSLGAYEQSHR | | | | |

**Supplementary Figure 1. Parallel reaction monitoring (PRM) MS on type I-D Cascade from *Synechocystis*. a,** The *Synechocystis* sp. PCC 6803 type I-D locus and location of targeted peptides. **b,** SDS-PAGE of type I-D Cascade after affinity and size-exclusion chromatography. Bands excised for MS are indicated. **c,** Peak area of each peptide was derived from extracted ion chromatograph with Skyline software. The band extracted at ~100 kDa contained all peptides specific to Cas10d at similar abundance, whereas the ~15 kDa extracted band was enriched for peptides from the C-terminus of Cas10d and consistent with the observed Cas11d subunit. **d**, Peptides relating to the mass targets searched for during PRM.

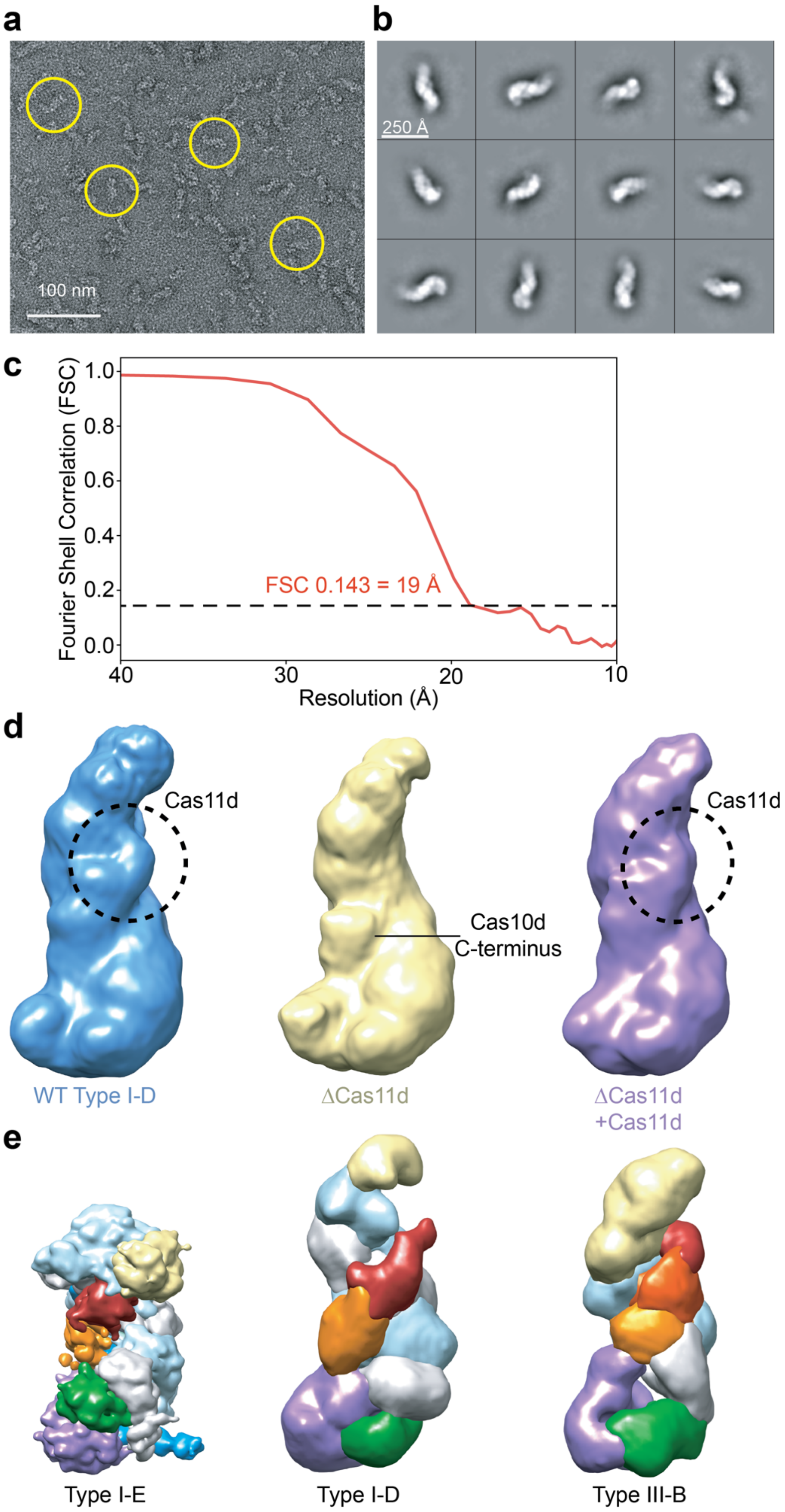


**Supplementary Figure 2. Single particle EM of wild type and ΔCas11d type I-D Cascade. a,** Representative raw micrograph of negatively stained WT type I-D Cascade. Representative particles are circled in yellow. **b,** Reference-free 2D class averages of WT type I-D Cascade. Particles are ~250 Å in the longest dimension. **c,** Fourier Shell Correlation (FSC) curve for the final refinement of the WT type I-D Cascade structure, indicating a final resolution of ~19Å using the 0.143 gold-standard criterion from two independent half-maps. **d,** Comparison of WT type I-D Cascade, ΔCas11d type I-D Cascade, and ΔCas11d type I-D Cascade complemented with Cas11d from a separate plasmid. **e,** Comparison between the structures of *E. coli* type I-E Cascade (left)[57], *Synechocystis* type I-D Cascade (middle), and *T. thermophilus* type III-B Cmr complex (right)[58].

**Supplementary Table 1. Shotgun MS on type I-D Cascade.**

| Protein name | Unique Peptides | Peptides | PSMs | Molecular mass, kDa | Length, aa | Coverage, % |
|---|---|---|---|---|---|---|
| Cas10d | 104 | 104 | 548 | 113.6 | 993* | 91 |
| Cas7d | 32 | 32 | 359 | 36.5 | 329 | 95 |
| Cas5d | 23 | 23 | 206 | 28.9 | 254 | 100 |
| Cas6d | 37 | 37 | 360 | 29.4 | 263 | 87 |
| Cas11d | 25 | 25 | 165 | 17.0 | 146 | 94 |
| (Cas10d) | | | | (113.6) | (993*) | (31) |

*Length refers to the His6-tagged Cas10d protein and is 18 amino acids longer than the native version.

**Shotgun MS output on protein bands excised from SDS-PAGE gel.** Protein name: protein name contained within group. Unique peptides: The number of peptide sequences that are unique to a protein group. These are the peptides that are common to the proteins of a protein group, and which do not occur in the proteins of any other group. Peptides: The total number of distinct peptide sequences identified in the protein group. PSMs: The number of PSMs (peptide spectrum matches) is the total number of identified peptide spectra matching the protein/protein group. The PSM value may be higher than the number of peptides identified when the mass spectrometer acquired multiple spectra for the same peptide. Molecular mass, kDa: molecular mass of the protein sequence. Length, aa: length of the protein sequence. Coverage, %: The percentage of the protein sequence covered by the identified peptides.

**Supplementary Table 2. Shotgun MS on type I-D Cascade for stoichiometry estimation.**

| Protein name | Peptide count | Abundance | Average abundance |
|---|---|---|---|
| Cas10d (1-847)* | 95 | $4.2\times10^{10}$ | $2.1\times10^{08}$ |
| Cas7d | 66 | $9.7\times10^{10}$ | $1.5\times10^{09}$ |
| Cas5d | 38 | $8.2\times10^{09}$ | $2.2\times10^{08}$ |
| Cas6d | 65 | $1.6\times10^{10}$ | $2.4\times10^{08}$ |
| Cas11d (848-993)*† | 30 | $2.0\times10^{10}$ | $1.4\times10^{09}$ |

*Length refers to the His6-tagged Cas10d protein and is 18 amino acids longer than the native version.
†Amino acid range relative to Cas10d

**Shotgun MS on entire lane excised from SDS-PAGE gel.** Protein name: protein name contained within group. Peptide count: total number of peptides observed. Abundance: sum of the peak area of all peptide ion signals matching the identified protein sequence. Average abundance: abundance divided by peptide count and is roughly proportional to the molar quantities of each protein. A proportion of the Cas11d (approximately equal to the value from Cas10d N-terminus) is contributed from the full length version of Cas10d.

#### Supplementary Table 3. Bacterial strains used in this study.

| Strain | Genotype/Phenotype/Description | Reference |
|---|---|---|
| BL21(DE3) | *E. coli* B $F^-$ *ompT, gal, dcm, lon, hsdS*$_B$(*r*$_B^-$*m*$_B^-$), λ(DE3 [*lacI lacUV5-T7p07 ind1 sam7 nin5*]) [*malB*$^+$]$_{K-12}$($λ^S$) | Promega |
| DH5α | *E. coli* $F^-$, ϕ80d*lacZ*ΔM15, Δ(*lacZYA–argF*)U169, *endA1*, *recA1*, *hsd*R17 ($r_K^-m_K^+$), *deoR*, *thi-1*, *sup*E44, $λ^-$, *gyr*A96, *rel*A1 | Gibco/BRL |
| LOBSTR | *E. coli* B $F^-$ *ompT, gal, dcm, lon, hsdS*$_B$(*r*$_B^-$*m*$_B^-$), λ(DE3 [*lacI lacUV5-T7p07 ind1 sam7 nin5*]) [*malB*$^+$]$_{K-12}$($λ^S$) *arnA slyD* | Kerafast |
| *Synechocystis* sp. PCC 6803 | Glucose tolerant laboratory wild-type strain GT-01 | 59, 60 |

#### Supplementary Table 4. Plasmids used in this study.

| Plasmid | Description | Reference |
|---|---|---|
| pACYCDuet-1 | Two T7/LacO promoters with P15A replicon, $Cm^R$ | Novagen |
| pQE-80LoriT | T5/LacO promoter with RP4 oriT, ColE1 replicon, $Ap^R$ | Qiagen |
| pSEVA184f | T5/LacO, pMB1 replicon, $Ap^R$ | 61 |
| pSEVA237Y | eYFP, pBBR1, $Km^R$ | 61 |
| pSEVA251 | Replicative plasmid, RSF1010 replicon, $Km^R$ | 61 |
| pPF1538 | Spacer1 of *Synechocystis* CRISPR array, pACYCDuet-1 | This study |
| pPF1539 | Cas6d, Spacer1 of CRISPR array, pACYCDuet-1 | This study |
| pPF1549 | N-$His_6$-tagged Cas10d, Cas7d, Cas5d, pQE80LoriT | This study |
| pPF1552 | N-$His_6$-tagged Cas6d, Spacer1 of CRISPR array, pACYCDuet-1 | This study |
| pPF1589 | N-$His_6$-tagged Cas10d, Cas7d, Cas5d, Cas3', pQE80LoriT | This study |
| pPF1720 | pSEVA251 with T5/LacO promoter | This study |
| pPF1758 | N-$His_6$-tagged Cas10d (mutated *cas11d* RBS and ATG), Cas7d, Cas5d, pQE80LoriT | This study |
| pPF1836 | Spacer 1 of the type I-D array flanked by GTT PAM, T5/LacO promoter, pSEVA251 | This study |
| pPF1914 | Translation reporter with *Synechocystis cas11d* WT RBS and ATG start codon | This study |
| pPF1915 | Translation reporter with *Synechocystis cas11d* RBS1 mut and CTG start codon | This study |
| pPF1926 | N-StrepII-tagged Cas6d, Spacer1 of CRISPR array, pACYCDuet-1 | This study |
| pPF1963 | N-$His_6$-tagged Cas5d, T5/LacO promoter, pSEVA251 | This study |
| pPF2034 | Cas11d, T5/LacO promoter, pSEVA251 | This study |
| (Cas5c-Cas8c-Cas7)/pHMGWA | *D. vulgaris* Cascade/I-C (N-$His_6$-MBP-Cas5c,Cas8c,Cas7), pHMGWA | 56 |
| CRISPR/pACYCDuet-1 | Three copies of Spacer2 from *D. vulgaris* CRISPR array, pACYCDuet-1 | 56 |
| pPF2116 | Translation reporter with *Synechocystis cas10d* WT RBS and ATG codon | This study |
| pPF2117 | Translation reporter with *Synechocystis cas11d* RBS1 mut and ATG start codon | This study |
| pPF2118 | Translation reporter with *Synechocystis cas11d* WT RBS and CTG start codon | This study |
| pPF2119 | Translational reporter with scrambled sequence and ATG | This study |
| pPF2120 | Translation reporter with *Cyanothece* potential *cas11d* RBS and start codon | This study |
| pPF2121 | Translation reporter with *Synechococcus* potential *cas11d* RBS and start codon | This study |
| pPF2122 | Translation reporter with *Listeria* potential *cas11b* RBS and start codon | This study |
| pPF2123 | Translation reporter with *Proteiniborus* potential *cas11b* RBS and start codon | This study |

| Plasmid | Description | Reference |
|---|---|---|
| pPF2125 | Translation reporter with *Mastigocladus* potential *cas11d* RBS and start codon | This study |
| pPF2126 | Translation reporter with *Microcystis* potential *cas11d* RBS and start codon | This study |
| pPF2127 | Translation reporter with *Nostocales* potential *cas11d* RBS and start codon | This study |
| pPF2128 | Translation reporter with *Anabena* potential *cas11d* RBS and start codon | This study |
| pPF2129 | Translation reporter with *Clostridium* potential *cas11d* RBS and start codon | This study |
| pPF2130 | Translation reporter with *Enterococcus* potential *cas11b* RBS and start codon | This study |
| pPF2131 | Translation reporter with *Bacillus* potential *cas11c* RBS and start codon | This study |
| pPF2132 | Translation reporter with *Vibrio* potential *cas11c* RBS and start codon | This study |
| pPF2133 | Translation reporter with *Nitrosospira* potential *cas11c* RBS and start codon | This study |
| pPF2134 | Translation reporter with *Desulfovibrio* potential *cas11c* RBS and start codon | This study |

**Supplementary Table 5. Oligonucleotides used in this study.**

| Name | Sequence (5'-3') | Notes | Restriction site (underlined) |
|---|---|---|---|
| PF2927 | ATCGGGATCCGAAAACCTGTATTTTCAGGGCACAACACTTCTTCAAACTTTGCTG | F for *cas10d*, N-His$_6$ | BamHI |
| PF2932 | ATTGCTGCAGGATTAATTAGTATCTAATAGTGATAATTGCGGTG | R for *cas5d* | PstI |
| PF2933 | ATAGCTGCAGATTAAAGAGGAGAAATTAACTATGTCATTGCCCAAAGTAGGTGCTATG | F for *cas3′d* | PstI |
| PF2934 | ATGCATGCATGTCCCTTAAAAAATTAAAGATTGACTGGTGG | R for *cas3′* | NsiI |
| PF2935 | GTACCCATGGCGTTTGATGATCGCTACAGTTTGTATTCCG | F for *cas6d* | NcoI |
| PF2936 | CATGAAGCTTTCATCCATGATTGTTAACTGAAACCTGTC | R for *cas6d* | HindIII |
| PF2937 | AGTCCATATGTTTCCCCGTAAGGGGTCGGAGG | F repeats+spacer1 | NdeI |
| PF2938 | ATGCGGTACCAAGATGGCACTAGATACTAACTCAAACC | R repeats+spacer1 | KpnI |
| PF3027 | ATCGGGATCCCGAAAACCTGTATTTTCAGGGCATGTTTGATGATCGCTACAGTTTG | F for *cas6d*, N-His6 | BamHI |
| PF3029 | CGTTGATTGTTGTGCCCCTGGCGGTCGCTTTCAATGCCTGCATG | F GTT protospacer | KpnI*, SphI* |
| PF3030 | CAGGCATTGAAAGCGACCGCCAGGGGCACAACAATCAACGGTAC | R GTT protospacer | KpnI*, SphI* |
| PF3420 | CAATGTTACCCTGGGAAAAAAGTTCAGCAAATTG | R for *cas11d* to mutate RBS | |
| PF3421 | CTTTTTTCCCAGGGTAACATTGTTCTGACCGAAAAATTGAAAC | F for *cas11d* to mutate internal RBS and ATG | |
| PF3648 | AATAAGGAGATATACTATGGCACATCACCATCACCATCACGCATCCGAAAACCTGTATTTTCA<br>GGGCACAAAAATTTATCGCTGTAAATTAACTCTCCATGACAATGTTTTTTTTGCCAGTCGAGAG<br>ATGGGAATTCTCTATGAAACAGAAAAGTATTTCCATAATTGGGCATTAAGTTATGCTTTTTTTAA<br>AGGAACAATTATTCCCCATCCTTATGGCTTAGTCGGACAGAATGCCCAAACACCTGCTTATTT<br>AGACCGAGATCGTGAGCAAAATTTACTCCACCTTAATGATTCAGGAATTTATGTTTTTCCTGCT<br>CAACCTATCCATTGGTCTTATCAAATTAATACCTTTAAAGCTGCTCAATCTGCCTATTATGGTC<br>GTTCTGTTCAATTTGGTGGGAAAGGTGCAACCAAAAACTATCCGATTAACTATGGTCGTGCCA<br>AAGAATTAGCAGTAGGTAGTGAGTTTCTGACTTATATCGTGAGTCAAAAAGAGCTAGATCTTC<br>CAGTATGGATTCGTTTAGGAAAATGGTCTTCTAAAATTCGGGTTGAAGTGGAGGCGATCGCC<br>CCTGATCAGATCAAAACTGCATCCGGAGTCTATGTCTGCAATCATCCCCTGAATCCTCTGGAT<br>TGCCCAGCAAATCAACAAATTTTGCTCTATAACCGGGTTGTTATGCCCCCATCGAGTTTATTTA<br>GCCAATCTCAATTACAGGGTGATTATTGGCAGATTGATCGTAATACGTTTTTGCCTCAAGGAT<br>TTCACTATGGAGCAACGACGGCGATCGCCCAAGATTCACCGCAATTATCACTATTAGATACTA<br>ATTAACTGCAGGTCGACAAGCTTGCGGCCG | gBlock *cas5d*, N-His$_6$ | |
| PF3721 | CAAGACTAGTTTACTTGTACAGCTCGTCCATG | R for eYFP | SpeI |
| PF3722 | GCGCGAATTCTTTTTTCCCAAGGAAATATTGTTATGACCGTGAGCAAGGGCGAGGAGCTG | F for *cas11d* promoter, WT RBS + ATG | EcoRI |
| PF3723 | GCGCGAATTCTTTTTTCCCAGGGTAACATTGTTCTGACCGTGAGCAAGGGCGAGGAGCTG | F for *cas11d* promoter, RBS1 + CTG | EcoRI |
| PF3760 | CATGGGCTGGAGCCACCCGCAGTTCGAAAAATCAGCTGCG | F StrepII tag with SAA linker | NcoI*, BamHI* |

| Name | Sequence (5'-3') | Notes | Restriction site (underlined) |
|---|---|---|---|
| PF3761 | GATCCGCAGCTGATTTTTCGAACTGCGGGTGGCTCCAGCC | R StrepII tag with SAA linker | NcoI*, BamHI* |
| PF3771 | ACCTAGGCCGCGGCCGCGCGAATTCAATAAGGAGATATACTATGGCACATCAC | F for Cas5d gBlock, with N-His$_6$ | |
| PF3772 | CGGCCGCAAGCTTGCATGCCTGCAGTTAATTAGTATCTAATAGTGATAATTG | R for Cas5d gBlock | |
| PF3935 | ACCTAGGCCGCGGCCGCGCGAATTCAATAAGGAGATATACTATGACCGAAAAATTGAAACTG ACTAAACG | F for *cas11d* | EcoRI |
| PF4092 | CTCTCTACTGTTTCTCCCCTAG | F for GTT protospacer | |
| PF4093 | TCGCCCTTGCTCACCATATG | R for GTT protospacer | |
| PF4095 | /5IRD700/CTCTCTACTGTTTCTCCCCTAG | F for GTT protospacer with IRDye700 | |
| PF4096 | /5IRD700/TCGCCCTTGCTCACCATATG | R for GTT protospacer with IRDye700 | |
| PF4121 | GCGCGAATTCTTTTTTCCCAGGGTAACATTGTTATGACCGTGAGCAAGGGCGAGGAGCTG | F for *cas11d* promoter RBS1 + ATG | EcoRI |
| PF4122 | GCGCGAATTCTTTTTTCCCAAGGAAATATTGTTCTGACCGTGAGCAAGGGCGAGGAGCTG | F for *cas11d* promoter, WT RBS + CTG | EcoRI |
| PF4124 | GCGCGAATTCTTTTTGCTCAAGGAGATACAATCATGACCGTGAGCAAGGGCGAGGAGCTG | F *Cyanothece cas11d* reporter | EcoRI |
| PF4125 | GCGCGAATTCTTTTTTCCCAAGGAAATATTGTTATGACCGTGAGCAAGGGCGAGGAGCTG | F *Synechococcus cas11d* reporter | EcoRI |
| PF4129 | GCGCGAATTCATTTTGAACCAGGAGGAGAAACCGTGACCGTGAGCAAGGGCGAGGAGCTG | F *Mastigocladus cas11d* reporter | EcoRI |
| PF4130 | GCGCGAATTCACCGAACCAATTTAACTATCGATATGACCGTGAGCAAGGGCGAGGAGCTG | F *Microcystis cas11d* reporter | EcoRI |
| PF4131 | GCGCGAATTCATTCTACTAAGGAGGAATTAATTGTGACCGTGAGCAAGGGCGAGGAGCTG | F *Nostocales cas11d* reporter | EcoRI |
| PF4133 | GCGCGAATTCTAAATACATTAATGGAGAGCCTTATGACCGTGAGCAAGGGCGAGGAGCTG | F *Anabaena cas11d* reporter | EcoRI |
| PF4134 | GCGCGAATTCATGTAGAGGAGGATGAGAAAATAGTGACCGTGAGCAAGGGCGAGGAGCTG | F *Clostridium cas11d* reporter | EcoRI |
| PF4138 | GCGCGAATTCTTGAACAAAAGGAGGAATGGGGCGTGACCGTGAGCAAGGGCGAGGAGCTG | F *Bacillus cas11c* reporter | EcoRI |
| PF4140 | GCGCGAATTCCAAAGCCCATCGTGGTCCTTAGAATGACCGTGAGCAAGGGCGAGGAGCTG | F *Scrambled cas11* reporter | EcoRI |
| PF4143 | GCGCGAATTCTTTTTCTAGGAGGATTGAATCATATGACCGTGAGCAAGGGCGAGGAGCTG | F *Enterococcus cas11b* reporter | EcoRI |
| PF4163 | GCGCGAATTCTAACTATTCAAAAGGAATACCCAATGACCGTGAGCAAGGGCGAGGAGCTG | F *Vibrio cas11c* reporter | EcoRI |
| PF4164 | GCGCGAATTCATCACAAAGAAAGAGGTTACCAAATGACCGTGAGCAAGGGCGAGGAGCTG | F *Nitrosospira cas11c* reporter | EcoRI |
| PF4253 | GCGCGAATTCGAAACAGAAAGCTGGAGGTGCCCATGACCGTGAGCAAGGGCGAGGAGCTG | F *Desulfovibrio cas11c* reporter | EcoRI |
| PF4313 | GCGCGAATTCGATTTTTAAAGGGGGTAGGATGTATGACCGTGAGCAAGGGCGAGGAGCTG | F *Proteiniborus cas11b* reporter | EcoRI |
| PF4314 | GCGCGAATTCAATTTTTGAAGGAGATAGATGTGATGACCGTGAGCAAGGGCGAGGAGCTG | F *Listeria cas11b* reporter | EcoRI |

*Partial restriction enzyme recognition site
