## Supplementary Information 2 for "Diverse CRISPR-Cas complexes require independent translation of small and large subunits from a single gene"

Supplementary Information S2

### Internal translation of large subunit transcripts drives small subunit synthesis in type I CRISPR-Cas interference complexes

Tess M. McBride, Evan A. Schwartz, Abhishek Kumar, David W. Taylor, Peter C. Fineran, and Robert D. Fagerlund

#### Phylogenetic analysis of the large subunit protein from type I CRISPR-Cas systems lacking the small subunit gene.

Refer to Supplementary information S1 for methods involved in phylogenetic analysis. Briefly, protein sequences for the large subunit from Class 1 CRISPR-Cas type I-B, -C, -D, -F and -G systems were obtained from NCBI Genbank. Protein sequences were aligned by MUSCLE and a maximum likelihood tree tree of Cas10d proteins from type I-D systems was constructed using the Neighbor-Joining method. Analysis was executed within Geneious Prime and exported in Nexus format.

The analysis of Cas10d is presented in three parts: 1) Taxa, including NCBI accession number and description of species for each protein analysed; 2) Protein alignment used for tree reconstruction for Cas10d; 3) Maximum likelihood tree in Nexus format. Only the protein alignment for the other large subunits are presented.

#### Cas10d

```
#NEXUS
begin taxa;
	dimensions ntax=288;
	taxlabels
	WP_015580876[&Description="Sulfolobus islandicus (CscA)",Modified=Mon Nov 04 14:23:59 NZDT 2019,"# Nucleotide Sequences With Quality"=0,Topology="linear","Original Location"="LocalDocuments/Rob Fagerlund/Cyanobacteria 2/Alignment 20190814 b/I-D Archeal and bacteria","Molecule Type"="AA",Created=Thu Oct 31 11:13:46 
NZDT 2019]
	'OKY78805.1'[&Description="Candidatus Methanohalarchaeum thermophilum"]
	'WP_099434783.1__2_'[&Description="Cyanobacterium aponinum"]
	'KST67896.1'[&Description="Mastigocoleus testarum BC008"]
	'WP_054464356.1__2_'[&Description="Planktothricoides sp. SR001"]
	'WP_087541441.1__2_'[&Description="Nostocales cyanobacterium HT-58-2"]
	'WP_041033210.1__2_'[&Description="Tolypothrix campylonemoides"]
	'WP_096561519.1'[&Description="Scytonema sp. NIES-4073"]
	'ALB41708.1__3_'[&Description="Anabaena sp. WA102"]
	'WP_039205098.1__2_'[&Description="Aphanizomenon flos-aquae"]
	'WP_096666728.1'[&Description="Dolichospermum compactum"]
	'WP_096540653.1'[&Description="Nostoc linckia"]
	'WP_096579980.1'[&Description="Anabaenopsis circularis"]
	'WP_094347302.1__2_'[&Description="Nostoc sp. 'Peltigera membranacea cyanobiont' 210A"]
	'WP_094328850.1__2_'[&Description="Nostoc sp. 'Peltigera membranacea cyanobiont' 213"]
	'AFY33389.1__2_'[&Description="Calothrix sp. PCC 7507"]
	'AFY42102.1__2_'[&Description="Nostoc sp. PCC 7107"]
	'WP_102178995.1__2_'[&Description="Fischerella thermalis"]
	'WP_102182618.1__2_'[&Description="Fischerella thermalis"]
	'WP_053957666.1'[&Description="Inediibacterium massiliense"]
	'OOV51320.1__4_'[&Description="Clostridium botulinum D/C"]
	'WP_078209760.1'[&Description="Clostridium botulinum"]
	'WP_039217383.1'[&Description="Clostridium novyi"]
	'WP_039231880.1'[&Description="Clostridium novyi"]
	'WP_124997487.1'[&Description="Clostridium tagluense"]
	'WP_115640522.1__2_'[&Description="Clostridium putrefaciens"]
	'WP_029452495.1'[&Description="Clostridium algidicarnis"]
	'KPA14430.1'[&Description="Candidatus Magnetomorum sp. HK-1"]
	'OQX03915.1'[&Description="Desulfobacteraceae bacterium IS3"]
	'WP_109206371.1'[&Description="Moorella sp. Hama-1"]
	'RMF76097.1'[&Description="Chloroflexi bacterium"]
	'WP_110515527.1'[&Description="Herpetosiphon llansteffanense"]
	'RMD54757.1'[&Description="Candidatus Thermofonsia bacterium partial"]
	'WP_129674530.1'[&Description="Candidatus Chloroploca sp. Khr17"]
	'PZS00723.1'[&Description="Chloroflexi bacterium"]
	'WP_083814595.1'[&Description="Ktedonobacter racemifer"]
	'WP_112428311.1__2_'[&Description="Thermogemmatispora tikiterensis"]
	'WP_052888244.1'[&Description="Thermogemmatispora carboxidivorans"]
	'WP_096675563.1'[&Description="Fischerella sp. NIES-4106"]
	'WP_007353275.1'[&Description="Kamptonema"]
	'TDA68045.1'[&Description="Clostridia bacterium"]
	'WP_013277798.1__2_'[&Description="Acetohalobium arabaticum"]
	'SNY41161.1__2_'[&Description="Orenia metallireducens"]
	'WP_084731779.1'[&Description="Seinonella peptonophila"]
	'WP_091837104.1__2_'[&Description="Marininema halotolerans"]
	'WP_113657839.1'[&Description="Thermoflavimicrobium sp. FBKL4.011"]
	'WP_124726785.1'[&Description="Thermoactinomycetaceae bacterium SCSIO 07575"]
	'WP_106346592.1__2_'[&Description="Planifilum fimeticola"]
	'WP_092035352.1__2_'[&Description="Planifilum fulgidum"]
	'WP_089967302.1__2_'[&Description="Lihuaxuella thermophila"]
	'AUS08811.1__2_'[&Description="Laceyella sacchari"]
	'WP_106342374.1'[&Description="Laceyella sediminis"]
	'PRZ14250.1'[&Description="Laceyella sediminis"]
	'WP_100060061.1'[&Description="Cylindrospermopsis raciborskii"]
	'WP_071247784.1'[&Description="Cylindrospermopsis raciborskii"]
	'PNJ97268.1__4_'[&Description="Cylindrospermopsis raciborskii C03"]
	'WP_057177813.1'[&Description="Cylindrospermopsis sp. CR12"]
	'WP_102947488.1'[&Description="Cylindrospermopsis raciborskii"]
	'WP_102943904.1'[&Description="Cylindrospermopsis raciborskii"]
	'WP_040007793.1'[&Description="Cylindrospermopsis raciborskii"]
	'WP_124956293.1__2_'[&Description="Halomarina oriensis"]
	'ABD41581.1__2_'[&Description="Methanospirillum hungatei JF-1"]
	'WP_048041820.1'[&Description="Methanosarcina mazei"]
	'KUL02466.1__2_'[&Description="Methanomicrobiales archaeon 53_19"]
	'ACL15519.1'[&Description="Methanosphaerula palustris E1-9c"]
	'WP_048144973.1'[&Description="Methanosphaerula palustris"]
	'KUK70137.1'[&Description="Methanocalculus sp. 52_23"]
	'WP_083523368.1'[&Description="Methanofollis ethanolicus"]
	'WP_048169909.1'[&Description="Methanosarcina sp. 2.H.A.1B.4"]
	'PXF57954.1'[&Description="ANME-2 cluster archaeon"]
```

'AUV82901.1__2_'[&Description="Salinigranum rubrum"]
'WP_092664863.1__2_'[&Description="Halorientalis persicus"]
'CCC39327.1__2_'[&Description="Haloquadratum walsbyi C23"]
'WP_121538498.1__2_'[&Description="Haloarcula sp. Atlit-120R"]
'CAI49723.1__2_'[&Description="Natronomonas pharaonis DSM 2160"]
'WP_124176898.1__2_'[&Description="Natrarchaeobius halalkaliphilus"]
'ACM59080.1__2_'[&Description="Halorubrum lacusprofundi ATCC 49239"]
'WP_121578412.1__2_'[&Description="Haloarcula sp. Atlit-47R"]
'WP_049920476.1'[&Description="Haloferax sulfurifontis"]
'WP_049896054.1'[&Description="Haloferax alexandrinus"]
'WP_049904832.1'[&Description="Natrialba asiatica"]
'WP_021780687.1'[&Description="Halarchaeum acidiphilum"]
'WP_020221130.1'[&Description="Halarchaeum acidiphilum"]
'WP_119814692.1__2_'[&Description="archaeon AArc-Sl"]
'WP_128478967.1'[&Description="Halorussus sp. RC-68"]
'WP_083905969.1'[&Description="Desulfurispora thermophila"]
'RME81351.1'[&Description="Caldilineae bacterium partial"]
'RLC62851.1'[&Description="Chloroflexi bacterium partial"]
'ARD84828.1'[&Description="Ferroplasma acidiphilum"]
'RLG21299.1'[&Description="Methanosarcinales archaeon partial"]
'WP_129891831.1__2_'[&Description="Ktedonobacterales bacterium SCAWS-G2"]
'WP_134440371.1'[&Description="Methylacidiphilum sp. Phi"]
'WP_126601694.1'[&Description="Dictyobacter aurantiacus"]
'WP_126218931.1'[&Description="tThermus scotoductus"]
'WP_126206860.1'[&Description="Thermus scotoductus"]
'WP_096572089.1'[&Description="Sphaerospermopsis kisseleviana"]
'WP_028091379.1'[&Description="Dolichospermum circinale"]
'WP_016949546.1'[&Description="Anabaena sp. PCC 7108"]
'WP_096554834.1__2_'[&Description="Nostoc sp. NIES-4103"]
'WP_103123815.1__2_'[&Description="Nostoc cycadae"]
'AFZ11795.1__2_'[&Description="Crinalium epipsammum PCC 9333"]
'RPH89551.1'[&Description="Chroococcales cyanobacterium metabat2.561"]
'WP_110579692.1'[&Description="Microcystis aeruginosa"]
'REJ40815.1'[&Description="Microcystis flos-aquae DF17"]
'WP_016872949.1'[&Description="Chlorogloeopsis fritschii"]
'RCJ15252.1'[&Description="Nostoc sp. ATCC 43529"]
'RAM48337.1'[&Description="Hapalosiphonaceae cyanobacterium JJU2"]
'WP_016863179.1'[&Description="Fischerella muscicola"]
'TFI55732.1__2_'[&Description="Mastigocladus laminosus UU774"]
'WP_131120682.1'[&Description="Westiellopsis prolifica"]
'PIG90759.1__2_'[&Description="Chroococcales cyanobacterium IPPAS B-1203"]
'AFZ33067.1__2_'[&Description="Gloeocapsa sp. PCC 7428"]
'CUR22982.1'[&Description="Planktothrix serta PCC 8927"]
'WP_019490246.1'[&Description="Calothrix sp. PCC 7103"]
'WP_127081050.1'[&Description="Calothrix desertica"]
'AFZ27299.1__2_'[&Description="Cylindrospermum stagnale PCC 7417"]
'WP_096625304.1'[&Description="Calothrix sp. NIES-3974"]
'WP_017311492.1'[&Description="Fischerella sp. PCC 9339"]
'WP_026731172.1'[&Description="Fischerella sp. PCC 9605"]
'PLZ99320.1__2_'[&Description="Fischerella thermalis CCMEE 5268"]
'PLZ83552.1__2_'[&Description="Fischerella muscicola CCMEE 5323"]
'WP_016865348.1'[&Description="Fischerella muscicola"]
'WP_062244597.1'[&Description="Fischerella sp. NIES-3754"]
'PMB45484.1__2_'[&Description="Fischerella thermalis CCMEE 5330"]
'PLZ91839.1__2_'[&Description="Fischerella thermalis CCMEE 5194"]
'PMB11907.1__2_'[&Description="Fischerella thermalis CCMEE 5282"]
'PLZ19049.1__3_'[&Description="Fischerella thermalis WC157"]
'WP_016871137.1'[&Description="Fischerella thermalis"]
'PMB36720.1__2_'[&Description="Fischerella thermalis BR2B"]
'WP_102166500.1__2_'[&Description="Fischerella thermalis"]
'PLZ39861.1__6_'[&Description="Fischerella thermalis WC538"]
'PLZ70284.1__3_'[&Description="Fischerella thermalis WC245"]
'WP_102173699.1__5_'[&Description="Fischerella thermalis"]
'WP_114707795.1__2_'[&Description="Fischerella thermalis"]
'WP_102175309.1__2_'[&Description="Fischerella thermalis"]
'WP_009458809.1__2_'[&Description="Fischerella thermalis"]
'WP_102147067.1__2_'[&Description="Fischerella thermalis"]
'WP_102147845.1__2_'[&Description="Fischerella thermalis"]
'PMB30020.1__2_'[&Description="Fischerella thermalis CCMEE 5208"]
'PLZ05371.1__2_'[&Description="Fischerella thermalis WC119"]
'PMB02183.1__2_'[&Description="Fischerella thermalis CCMEE 5273 partial"]
'WP_082065505.1__2_'[&Description="Aliterella atlantica"]
'PSB52452.1__2_'[&Description="filamentous cyanobacterium Phorm 6 partial"]
'EKQ70621.1__2_'[&Description="Oscillatoriales cyanobacterium JSC-12"]
'PSB54396.1__2_'[&Description="Chamaesiphon polymorphus CCALA 037"]
'WP_023173330.1'[&Description="Gloeobacter kilaueensis"]
'WP_066345914.1__2_'[&Description="Geminocystis sp. NIES-3708"]
'WP_083260496.1'[&Description="Cyanobacterium sp. IPPAS B-1200"]
'WP_085436906.1'[&Description="unicellular cyanobacterium SU2"]
'WP_002784665.1'[&Description="Microcystis aeruginosa"]
'WP_079208699.1__2_'[&Description="Microcystis aeruginosa"]
'WP_002800252.1__2_'[&Description="Microcystis aeruginosa"]
'CUR32202.1'[&Description="Planktothrix tepida PCC 9214"]
'KPQ39684.1'[&Description="Phormidium sp. OSCR"]
'PPT09156.1'[&Description="Geitlerinema sp. FC II"]
'WP_017662193.1'[&Description="Geitlerinema sp. PCC 7105"]
'WP_103112787.1__2_'[&Description="Microcystis aeruginosa"]
'WP_104395902.1'[&Description="Microcystis aeruginosa"]
'WP_002791883.1'[&Description="Microcystis aeruginosa"]
'WP_002766200.1'[&Description="Microcystis aeruginosa"]
'WP_002788626.1'[&Description="Microcystis aeruginosa"]
'WP_002797932.1'[&Description="Microcystis aeruginosa"]
'REJ40820.1'[&Description="Microcystis flos-aquae DF17"]
'WP_002742886.1__2_'[&Description="Microcystis aeruginosa"]
'WP_002755926.1'[&Description="Microcystis aeruginosa"]
'AKV70201.1__2_'[&Description="Microcystis panniformis FACHB-1757"]
'REJ60002.1'[&Description="Microcystis aeruginosa DA14"]
'WP_130756400.1'[&Description="Microcystis aeruginosa"]
'REJ44384.1'[&Description="Microcystis flos-aquae TF09"]
'WP_024970397.1'[&Description="Microcystis aeruginosa"]
'WP_125730344.1'[&Description="Microcystis viridis"]
'KXS91774.1'[&Description="Microcystis aeruginosa NIES-88 partial"]
'WP_002778661.1'[&Description="Microcystis aeruginosa"]
'WP_002762261.1'[&Description="Microcystis aeruginosa"]
'EPF21995.1__4_'[&Description="Microcystis aeruginosa SPC777"]
'AFZ42916.1__2_'[&Description="Halothece sp. PCC 7418"]
'PNW54205.1'[&Description="Euhalothece sp. KZN 001"]
'WP_018399541.1'[&Description="filamentous cyanobacterium ESFC-1"]
'WP_017306104.1'[&Description="Spirulina subsalsa"]
'WP_075891453.1__2_'[&Description="Limnothrix rosea"]
'WP_024547051.1'[&Description="Synechococcus sp. NKBG15041c"]
'PZU97760.1'[&Description="Pseudanabaena sp."]
'OYQ61999.1'[&Description="Pseudanabaena sp. SR411"]
'WP_094536065.1'[&Description="Pseudanabaena sp. SR411"]
'WP_086766415.1__2_'[&Description="Nostoc sp. 106C"]
'WP_096597869.1'[&Description="Calothrix sp. NIES-2100"]
'WP_048869210.1__2_'[&Description="Scytonema tolypothrichoides"]
'KYC39370.1__2_'[&Description="Scytonema hofmannii PCC 7110"]
'AFZ37990.1__2_'[&Description="Stanieria cyanosphaera PCC 7437"]
'WP_019503799.1'[&Description="Pleurocapsa sp. PCC 7319"]
'WP_008312261.1'[&Description="Leptolyngbya sp. PCC 6406"]
'WP_121971064.1'[&Description="Leptolyngbya sp. BC1307"]
'WP_023075354.1__2_'[&Description="Leptolyngbya sp. Heron Island J"]
'EKV02086.1__2_'[&Description="Leptolyngbya sp. PCC 7375"]
'WP_017326896.1'[&Description="Synechococcus sp. PCC 7336"]
'PSN19223.1__2_'[&Description="filamentous cyanobacterium CCP5"]
'RMH16295.1'[&Description="Gammaproteobacteria bacterium"]
'WP_054467001.1__2_'[&Description="Planktothricoides sp. SR001"]
'WP_094678459.1'[&Description="Hydrocoleum sp. CS-953"]
'WP_094678461.1'[&Description="Hydrocoleum sp. CS-953"]
'WP_052128200.1'[&Description="Neosynechococcus sphagnicola"]
'PSB20197.1__2_'[&Description="Phormidesmis priestleyi ULC007"]
'RNJ65627.1'[&Description="Leptolyngbya sp. IPPAS B-1204"]
'WP_106171784.1__2_'[&Description="filamentous cyanobacterium CCP2"]
'PSB56979.1'[&Description="filamentous cyanobacterium CCP1 partial"]
'EDX77106.1__2_'[&Description="Coleofasciculus chthonoplastes PCC 7420"]
'WP_017323372.1'[&Description="cyanobacterium PCC 7702"]
'WP_046276749.1__2_'[&Description="Limnoraphis robusta"]
'WP_023065083.1__2_'[&Description="Lyngbya aestuarii"]
'WP_009786738.1'[&Description="Lyngbya sp. PCC 8106"]
'WP_079680991.1'[&Description="Planktothrix sp. PCC 11201"]
'SKB14935.1'[&Description="Planktothrix sp. PCC 11201"]
'AFY52833.1__2_'[&Description="Rivularia sp. PCC 7116"]
'WP_088889711.1'[&Description="Leptolyngbya ohadii"]
'WP_017652463.1'[&Description="Fortiea contorta"]
'WP_089129578.1'[&Description="Tolypothrix sp. NIES-4075"]
'WP_086689580.1__2_'[&Description="Nostoc sp. T09"]
'WP_109006821.1'[&Description="Nostoc commune"]
'PSB01849.1__2_'[&Description="Merismopedia glauca CCAP 1448/3 partial"]
'AFZ00743.1__2_'[&Description="Calothrix sp. PCC 6303"]
'PAX59910.1__2_'[&Description="Calothrix elsteri CCALA 953"]
'AFZ57018.1__2_'[&Description="Anabaena cylindrica PCC 7122"]
'AFW97301.1__2_'[&Description="Anabaena sp. 90"]
'WP_028084539.1'[&Description="Dolichospermum circinale"]
'TAF08895.1'[&Description="Nostocales cyanobacterium"]
'WP_127055411.1'[&Description="Trichormus variabilis"]
'WP_045870814.1'[&Description="Nostocales"]
'WP_096577327.1'[&Description="Nostocales"]
'WP_096642022.1'[&Description="Calothrix brevissima"]
'WP_099066180.1__14_'[&Description="Nostoc linckia"]
'AFY46833.1__2_'[&Description="Nostoc sp. PCC 7524"]
'WP_011320851.1__2_'[&Description="Trichormus variabilis"]
'WP_096682794.1'[&Description="Nostoc sp. NIES-2111"]
'WP_104900394.1__2_'[&Description="Nostoc sp. 'Peltigera membranacea cyanobiont' N6"]
'WP_089091838.1'[&Description="Nodularia sp. NIES-3585"]
'RCJ39405.1'[&Description="Nostoc punctiforme NIES-2108"]
'WP_086836537.1__2_'[&Description="Nostoc sp. RF31YmG"]
'WP_114082632.1__2_'[&Description="Nostoc sp. ATCC 53789"]
'WP_038296476.1'[&Description="Scytonema hofmanni UTEX B 1581"]
'WP_118166690.1'[&Description="Nostoc sphaeroides"]
'TAE22780.1__3_'[&Description="Oscillatoriales cyanobacterium"]
'WP_104546987.1__2_'[&Description="Chroococcidiopsis sp. TS-821"]
'WP_105218136.1'[&Description="Gloeocapsopsis sp. AAB1 = 1H9"]
'TEU17163.1'[&Description="Anaerolineales bacterium"]
'RRR68494.1'[&Description="Candidatus Viridilinea halotolerans"]
'WP_129679138.1'[&Description="Chloroflexales bacterium ZM16-3"]
'WP_044199283.1'[&Description="Oscillochloris trichoides"]
'WP_029215466.1'[&Description="bacterium JKG1"]
'WP_129632988.1'[&Description="Oscillochloris sp. Chuk17"]
'WP_097645710.1'[&Description="Candidatus Viridilinea mediisalina"]
'WP_087711939.1'[&Description="Phormidium sp. HE10JO"]
'WP_012599091.1__2_'[&Description="Cyanothece sp. PCC 7424"]
'WP_011153680.1__3_'[&Description="Synechocystis sp. PCC 6803"]
'SMQ86523.1__3_'[&Description="Synechococcus sp. 7002"]
'WP_017720464.1'[&Description="Oscillatoria sp. PCC 10802"]
'WP_080810414.1'[&Description="Halomicronema hongdechloris"]
'ADN17499.1__2_'[&Description="Cyanothece sp. PCC 7822"]
'WP_046279385.1__2_'[&Description="Limnoraphis robusta"]
'WP_012627888.1__2_'[&Description="Cyanothece sp. PCC 7425"]
'WP_015783166.1__2_'[&Description="Cyanothece sp. PCC 8802"]
'WP_012593877.1__2_'[&Description="Cyanothece sp. PCC 8801"]
'ELR97243.1__2_'[&Description="Gloeocapsa sp. PCC 73106"]
'WP_035152549.1__2_'[&Description="Calothrix sp. 336/3"]
'KYC42933.1__2_'[&Description="Scytonema hofmannii PCC 7110"]
'WP_096727546.1__2_'[&Description="Nostoc carneum"]
'WP_053457730.1__2_'[&Description="Hapalosiphon sp. MRB220"]
'PHJ60297.1__14_'[&Description="Nostoc linckia z3"]
'AFY31721.1__2_'[&Description="Calothrix sp. PCC 7507"]
'KST66746.1__2_'[&Description="Mastigocoleus testarum BC008"]
'WP_088240873.1'[&Description="Calothrix rhizosoleniae"]
'WP_036842433.1'[&Description="Planktothrix agardhii"]
'CUR14078.1'[&Description="Planktothrix paucivesiculata PCC 9631"]

'WP_043938760.1'[&Description="Planktothrix"]
'WP_043937175.1'[&Description="Planktothrix rubescens"]
'WP_040957323.1'[&Description="Planktothrix agardhii"]
'WP_036834484.1'[&Description="Planktothrix"]
'WP_042156684.1'[&Description="Planktothrix agardhii"]
'BBD53976.1'[&Description="Planktothrix agardhii NIES-204"]
'WP_088889829.1'[&Description="Leptolyngbya ohadii"]
'WP_036011813.1'[&Description="Leptolyngbya sp. JSC-1"]
'OBQ22644.1__3_'[&Description="Anabaena sp. AL93"]
'AUT00479.1__2_'[&Description="Nostoc sp. CENA543"]
'WP_096680032.1__2_'[&Description="Nostoc sp. NIES-2111"]
'WP_096656782.1__2_'[&Description="Calothrix parasitica"]
'WP_018398404.1'[&Description="filamentous cyanobacterium ESFC-1"]
'WP_017659639.1'[&Description="Geitlerinema sp. PCC 7105"]
'AFY84409.1__2_'[&Description="Oscillatoria acuminata PCC 6304"]
'RMH78832.1'[&Description="Cyanobacteria bacterium J007"]
;
end;

begin characters;
dimensions nchar=2247;
format datatype=protein missing=? gap=-;
matrix
WP_015580876 [illegible]

KSWMAVDVPYKLKIQNFSN--KI------NAGVSEP-KRN-------VCSICNIEYLLHKVTYTSN------------MEVSRRYLSI-LP---RGFNTR----A-YI [illegible]
---------------NET------SKW--ISTLLYALTFNIYFDSKVVISEFPVPILNQD-EMEE-----IY--VGDVPVIFKNIFEK-----SW--------TKE [illegible]
IKKLAIFANENYIRGSINGGFI-----KDNAINKPLNIIIDIL-CRWDKNLYT--K-EDITALCKREIQKYFERVD----------------PYY [illegible]
'WP_029452495.1' [illegible]
-------------------KGEYINILRE-CEIL-NIK----EFL--PE-------YKEYIEDIRAIIAE-HSAHMSSFTESLIFNPE-------------RY--KLGK-EKME--IL [illegible]
DFKIYIKNVQ----------SG-IKVDKKCLGI-ASLEDIV----------------YEIKNRCE-------------------KSKIP------------------D------IN [illegible]
EQRVSKYILERKENETYFIDNNEDKLRFSEFIRSFYTFINMHVVVKDTTLTWNKVYDF----FDINEIDR-DYLD---------------------------VF [illegible]
SSWMAMDVPYKLKIQNFSN--KI-NA----GV-SEP-KRN-------VCSICNIEYLLHKVTYTSN------------MEVSRRYLSI-LP---RGFNTR----A-YI [illegible]
----------------NET------SKW--ISALLYALTFNIYFDSKVVISEFPVPILNQD-EMEE-----IY--VGDVPVVFKNIFEK-----SW--------TKE [illegible]
IKELAIFANENYIRGSINGGFI-----KDNAINKPINIIMDIL-CRWNKNLYT--K-EDITALCKREIQKYFERVD----------------PYY [illegible]
'KPA14430.1' [illegible] MAENTETLILK [illegible]
---------------------------------------------YDEYKNLSYKDVATQENIS-KHLNFI--FE--DSSDDFDICITTIQTIMKGHSGH---MHCNAAGLDAN [illegible]
---------------DKLSS--KDT--S---RLVKDSKGTA------------MNFIKNPFALKFSPEAVM--------------DILVKAIL---------------------AP [illegible]
NTSLTIDNIDLSNHKPETLFSSDIEVMHMGMLVGSFAYLLINYVTIDKFPLNSSTAWNLTLEA----AGVD-----PNDS---------------PELM [illegible]
SSGQG----HELMAGNLPKGIKPQLFSN--RL-KG---GG--GEP-KRN-------VCIICNQGFFWEKILHESYDN--------------HYYLHL-FADGGEH [illegible]
---------NP-----------PGKES--------NES------IKF--IFALFHLLMLTDYFHLRGILSRSSIPPLKAN-EFDS-----LY--IDHVPLAFTALI-Q--SN-NI [illegible]
IKKLAEIALNNHF--HGQSF-------KETSQAKPLDLAFDAL-SKHQPPESK----EDLKMVMLHDVSRGLERLS----------------RFGQ [illegible]
'OQX03915.1' [illegible] MTDTKELFLK [illegible]
-------------FPEYEGKSYGSIAN-------------------SENVQKIIKIMLSEG-GH-----EFD--LD----------IERIVSLIRA-HSGHYHADGDSLFAEQH [illegible]

[illegible]

'WP_109206371.1' [illegible]

'RMF76097.1' [illegible]

'WP_110515527.1' [illegible]

'RMD54757.1' [illegible]

'WP_129674530.1' [illegible]

'PZS00723.1' [illegible]

'WP_083814595.1' [illegible]

'WP_112428311.1_2_' [illegible]

'WP_052888244.1' [illegible]

'WP_096675563.1' [illegible]

'WP_007353275.1' [illegible]

'TDA68045.1' [illegible]

'WP_013277798.1_2_' [illegible]

'SNY41161.1_2_' [illegible]

'WP_084731779.1' [illegible]

'WP_091837104.1_2_' [illegible]

'WP_113657839.1' [illegible]

'WP_124726785.1' [illegible]

'WP_106346592.1_2_' [illegible]

'WP_092035352.1_2_' [illegible]

'WP_089967302.1_2_' [illegible]

'AUS08811.1_2_' [illegible]

'WP_106342374.1' [illegible]

'PRZ14250.1' [illegible]

'WP_100060061.1' [illegible]

'WP_071247784.1' [illegible]

TGSPPK--LCSLC-NRVITVKMKNCSIKADII-EDGIKEFSN--RL--L---PKSEDVS-SRL-------WCPQCYLEWMARKLSGLGY--------APGAEK[illegible]
GNYLISSRSSFDEDLDDDYSDDDEDLEMNNQYYRLASNFLILTLESSTYK---EPI---------------------KKT------ELW--MRGILTTLVLQDLL[illegible]
--------IFTLA--CRIILDS-F----------------------------------------EKKDDAYSMTMKNLAETIATQSLDLFLPRWERDGKG-----KANRYEKTYR[illegible]
'PNJ97268.1__4_' ---------------------------------------------------------------------------------MFMS[illegible]
TESDFRRVCALFALHDLHK----------------------DDYLAREVKQKE-----------------------TSIQPVLMLEIANQV-SLT----EWL--NN--EDLNGY[illegible]
FYPLIFFENGTIYIGE--KDTQS-F-DKD---SFIS--SVYNEF---------------NESLSQLGKV--S-N-DELQYNA------------KT-QRFEKYIFAF-SGI[illegible]
-FA---EKWKAVSS--YLG-------GLL-------NLLRD-----------------VYLAPDD-------TWQKTTEFTGKFLDIP-----RDIMSNI-------------FDNIST[illegible]
ADSLTVITAKN--KTRSPAK-LCSLC-NRVITS-KMKSSLIKADMIEDGIKEFSN--RL-LPKTKD---KDVSSRL-------WCPQCYLEWMIRKLSGL[illegible]
--------------------------------------------------EG---------------RIW--LEEVLNSLAEETSRQTKNH-QRVPGNYLLGSNSYSDD----EY--LDD[illegible]
--------------------------SDLV-----------TKPQLFDLIASAL--------------------------------------------------PTRLLENGKYLVS-TRLLEVKFADD[illegible]
'WP_057177813.1' ---------------------------------------------------------------------------MNLNQIIL[illegible]
TESDFRRVCALFALHDLHK----------------------DDYLAREVKQKE-----------------------TSIQPVLMLEIANQV-SLT----EWL--NN--DDLNGY[illegible]
FYPLIFFENGTIYIGE--KDTQS-F-DKD---SFIS--SVYNEF---------------NESLSQLGKV--S-N-DELQYNA-TY------------QIFEKYIFAF-SGIK[illegible]
FA---EKWKAVSS--YLG-------GLL-------NLLRD-----------------VYLAPDNT--------WQKTIEFMGNFLDIP-----RDII---------------SNIFDNVST[illegible]
-KTGSPAK-LCSLC-NRVITSKMKSSPIKADMIEDGIKE-FSN--RLLPK----T--KDVSSRL-------WCPQCYLEWMIRKLSGLGY--------APQAE[illegible]
LLGSNSY---SDDEYLDDDDYSDDDPEINNDHHRLASNFLIFSL------ESSSYN-------------EP------IKKS------ELW--MRGLLTALVLQDLLG[illegible]
--VFTLA--CHIILDS-F----------------------------------------EKKDNAYSMTLKNL-AETIAIQSLDLFLPFSESDGKG-----KANRYEKTYRAAIST[illegible]
'WP_102947488.1' ---------------------------------------------------------------------------------[illegible]
YFL------------------------------------------------------SIKS--------------------TQEPKNI-H----------------------------------------------------FYYH[illegible]

[illegible]

'WP_119814692.1__2_'

[illegible]

'WP_128478967.1'

[illegible]

'WP_083905969.1'

[illegible]

'RME81351.1'

[illegible]

'RLC62851.1'

[illegible]

'ARD84828.1'

[illegible]

'RLG21299.1'

[illegible]

'WP_129891831.1__2_'

[illegible]

'WP_134440371.1'

[illegible]

'WP_126601694.1'

[illegible]

'WP_126218931.1'

[illegible]

'WP_126206860.1'

[illegible]

'WP_096572089.1'

[illegible]

'WP_028091379.1'

[illegible]

'WP_016949546.1'

[illegible]

'WP_096554834.1__2_'

[illegible]

'WP_103123815.1__2_'

[illegible]

'AFZ11795.1__2_'

[illegible]

'RPH89551.1'

[illegible]

'WP_110579692.1'

[illegible]

'REL40815.1'

[illegible]

'WP_016872949.1'

[illegible]

'RCJ15252.1'

[illegible]

'RAM48337.1'

[illegible]

'WP_016863179.1'

[illegible]

'TFI55732.1__2_'

[illegible]

'WP_131120682.1'

[illegible]

'PIG90759.1__2_'

[illegible]

'AFZ33067.1__2_' [illegible]

'CUR22982.1' [illegible]

'WP_019490246.1' [illegible]

'WP_127081050.1' [illegible]

'AFZ27299.1__2_' [illegible]

'WP_096625304.1' [illegible]

'WP_017311492.1' [illegible]

'WP_026731172.1' [illegible]

'PLZ99320.1__2_' [illegible]

'PLZ83552.1__2_' [illegible]

'WP_016865348.1' [illegible]

'WP_062244597.1' [illegible]

'PMB45484.1__2_' [illegible]

'PLZ91839.1__2_' [illegible]

'PMB11907.1__2_' [illegible]

'PLZ19049.1__3_' [illegible]

'WP_016871137.1' [illegible]

'PMB36720.1__2_' [illegible]

'WP_102166500.1__2_' [illegible]

'PLZ39861.1__6_' [illegible]

'PLZ70284.1__3_' [illegible]

'WP_102173699.1__5_' [illegible]

'WP_114707795.1__2_' [illegible]

'WP_102175309.1__2_' [illegible]

'WP_009458809.1__2_' [illegible]

'WP_102147067.1__2_' [illegible]

'WP_102147845.1__2_' [illegible]

'PMB30020.1__2_' [illegible]

'PLZ05371.1__2_' [illegible]

-----PE--E--------F---YTPLLYLPDGVVYLAN--TNAPT-I-TTD---TLPE--GVVDKI-----------------KSLCA--EKL--G-E-RQTGFNR-DG----------KG-LKFADYYWLF-FDVVGLM-----------------KVSIDAAC-----------------------RL-LP----------------------D------SK---TASSG--------------------------------------------KR------GE---SLQ--------SYQ------------TQGELPT-------NL---NLQF------PN----EIRI-----DR--------
LA--------------------E----FG---DILC--RG--IWN-------SWC-------DRVKEAQKDIPKAKRK----------VPPELD-------LTQKLAEY----LELS-----DEIP-AI--------------KQIQSLK----------KTGGVPLDWYYLAAQYF----R--K---HP-GK-DFA-QILE----VMRGM-VDYAASLIQP------------ILQ----------EF---QD--I--P-DG-WED-------------LKTYVKRVI---------SLP-----------------------------------TGAVFAPETEPF-
-LLELKRYNAAKV-TGRGRES-VCAMS-SSAYTV---TEQMESATLFAPQV--YSN--RQ-IL--FNA--QAA-KRQ-------ICSIWSIEIMLRQILMNQT-N------AVGGDFESRKYRYLYL-YP---TYFFTP----E-TNKF---LQL-----AYNQF-ARTRF-----DA------ELR-----KHFI-T-------------------------------------------------------DKQIA---------KFS------IQNY--QQVDTL-------L-IKENLNPDD------------------DR-------TF-
--KI---SF---PEK---------------------ETLTFFFLGL-----------PP------G----R--EP---------TDT------ESW--VTPAWLALTLPLVLDVKVVASESPVPPFISGADFEET----VL--LDGEHQAIRSLI-K--KD-TY---------RLD----SILPSS------------SEKR-EFSPLNA-LTAAYCIH-LEV------NRKKDG---DPDWG-KLSDLARDL-ETSPLYVFHYLTKWLRKPKKDKDKQKT-------------LDA--V---PV----AKIR----------
LY-MD--LYKYFEP-G---------------------------E----------------------------ET-MNQ-LRKLTELYRRFYR--A-KSS--YA---KANAILKPIDEAADVI-LKIDKALVA--NTESLTDVVAARLAKLMNNVR-----R----RTAEG-----KPTL-AFVDGKWKPAL------TSEEE----RQAVYDFANYFV--KEIFEGNFKCDRARLAGTQLNLI-----RDTCEYLYR------LADDE--ERK--NLPQ-AEPED----IPDLD-VDDAA------

'PMB02183.1__2_' ----------------------------------------------MTDFEDNF----DDLPEEVP-----DFDST-DGD---------DDE--SPIKKELLTIRLFKEA---VKKSKGNQG---------DA--ILEK--FAEYVL--PNLI-QQ-LAGATAK----G---------GKF-FDV-TVP--KINA-------KRA--------SE---------GKP--------P----VD-RRNA-GDQSIVAHLLNGLFPTYRIL--RK-------------------------
LQSSK-E-TNPVKRNC---EDLQACIFITAYLLHDYEK-----F-----------------P-D--YQAWLIENN----------IAERDWELDTPKKEDAPNLGRGYITKKILDF-GLY----YLL--GD------DWQDFIDDIIEISNN-SGV---KHDSDLGLATR------GL-KTLDDERI----DS-RIRQ--VLIDLVSLSDLFASV------VKHPRDV-E-----TGR-----------LPNLI-S-RL---SN-HQL----K-------LTYHSLCENRGVLTNILNNSLIEAH---
-----PE--E--------F---YTPLLYLPDGVVYLAN--TNAPT-I-TTD---TLPE--GVVDKI-----------------KSLCA--EKL--G-E-RQTGFNR-DG----------KG-LKFADYYWLF-FDVVGLM-----------------KVSIDAAC-----------------------RL-LP----------------------D------SK---TASSG--------------------------------------------KR------GE---SLQ--------SYQ------------TQGELPT-------NL---NLQF------PN----EIRI-----DR--------
LA--------------------E----FG---DILC--RG--IWN-------SWC-------DRVKEAQKDIPKAKRK----------VPPELD-------LTQKLAEY----LELS-----DEIS-AI--------------KQIQSLK----------KTGGVPLDWYYLAAQYF----R--K---HP-GK-DFA-QILE----VMRGM-VDYAASLIQP------------ILQ----------EF---QD--I--P-DG-WED-------------LKTYVKRVI---------SLP-----------------------------------TGAVFAPETEPF-
-LLELKRYNAAKV-TGRGRES-VCAMS-SSAYTV---TEQMESATLFAPQV--YSN--RQ-IL--FNA--QAA-KRQ-------ICSIWSIEIMLRQILMNQT-N------AVGGDFESRKYRYLYL-YP---TYFFTP----E-TNKF---LQL-----AYNQF-ARTRF-----DA------ELR-----KHFI-T-------------------------------------------------------DKQIA---------KFS------IQNY--QQVDTL-------L-IKENLNPDD------------------DR-------TF-
--KI---SF---PEK---------------------ETLTFFFLGL-----------PP------G----R--EP---------TDT------ESW--VTPAWLALTLPLVLDVKVVASESPVPPFISGAD-------------------------------------------------------------------------------------------------------------------------------------------------------------------------------------------------------------------
---------------------------------------------------------------------------------------------------------------------------------------------------------------------------------------------------------------------------------------------------------------------------------------------

'WP_082065505.1__2_' -----------------------------------MNLFMADNNF---------DDLPLETL-----EFDAT-QNL---------DDE--TPIKAELLTIKLFKQA---ILDTEGNKS---------DR--ILEN--LAENVV--PNLF-SQ-LGGATAK----G---------GQF-TE-------------------DRR-------AE---------GKN--------V------ERSK-HDQSFISHLLNGLLPTYRIL--KQ------------------------LRLEQ-A-
TNPVKRYC---NDIVICLFIVSYLLHDFDK-----F-----------------P-D--YPAWLNANDNEG-------KFKNRHWQKDQPHKSDAPNFGREYIAKKITDL-GLN----YLL--GD------DWEKHIDDIVWLSNN-AGV---KYDADLGLEIR------GL-NP----KL-----DG-RTRS--ILASLVRLSDLFASV------IKRPGDV-D-----NNG-----------LFEEL-T-SL---SN-GQL----K-------FSYHSLSDNRGVITNILNNALIEAH--------
PQ--D---------F---YTALLYLPDGVVYLTK--PNAPV-I-DIS---NLPN--LVIAKI-----------------KILCG--DRL--K-R-DQTGFGR-DG----------KG-FKFADYYWLF-FDAIQLM-----------------EVSVNAAS-----------------------RL-IP----------------------S-------TK---SSSAK--------------------------------------------KR------SD---SLV--------AFQ-------------NAGDLPT-------NL---NLEF------GE----DYRI-----DR--------LA--------
------------E----FG---DTLC--RG--IWS-------EWC-------DRLNTWQKQQPKANRK----------EIPNLN-------LTHKLAEY----LGLS-----TEIS-AI--------------QAIQTLK----------KTGGVPLDWYYIAAQYF----Q--R---NL-GL-DEA-QIRE----RMTEM-VNYAATLIKP------------TLE----------QF----T--I--S-DG-WDD-------------IRTYVNRVV---------CLP-----------------------------------AGAVVTPEIQPF--
LTEISRYSAAKV-TGRGREN-VCAMS-SSSYTV---TEQMESATLFAPQV--YSN--RQ-IL--FNA--QAA-KRQ-------ICSIWSIEIMLRQILMNQT-N------ATGGDFEGRKYRYLYL-YP---AYFFTP----E-TNKF---LQK-----AYTWI-ERSRF-----DA------DIR-----KYLI-T-------------------------------------------------------EKQEA---------NFT------LDNY--QKVDAL-------L-IKEDLQPEN------------------DR-------TF--
-KI---SY---PEN---------------------QPLTFFFLAL-----------PP------G----K--DA---------TDI------ESW--VMPAWLALALPLVLDIKTVASESPVPPFISGADFEQT----VL--LDGEHQAIHSLI-K--QA-EY---------RLD----SIVPRS------------LNQR-KFSPLNA-LSAAYCIH-LEV------NRKKDG---NPDWG-KLAALARDL-ETSPLYVFHYLSKWLRKPKSNSV-QIPE-------------IKS--V---PI----SKIK----------LY-
FD--FYYYFEP-K---------------------------G-----------------------------DE-MTK-LENLTELYRKFYR--A-KSQ--YA---KANAVLKPIDEAANVV-LKIDKALAS--DNESLTQLVAARLSTLMKNVR-----R----RTAEG-----KPTL-TFIDGKWKPAL------NSEEE----RQAIYEFAKYFV--EVIFDETFKSDRARLAGMQLNLI-----RDTCDYLYR------LADDT--ERQ--QKPQ-DEPDE----IIELE-AEATI------

'PSB52452.1__2_' [illegible]

[illegible]

'EKQ70621.1__2_' [illegible]

[illegible]

'PSB54396.1__2_' [illegible]

[illegible]

'WP_023173330.1' [illegible]

[illegible]

'WP_066345914.1__2_' [illegible]

[illegible]

'WP_083260496.1' [illegible]

[illegible]

'WP_085436906.1' [illegible]

[illegible]

'WP_002784665.1' [illegible]

[illegible]

'WP_079208699.1__2_' [illegible]

[illegible]

'WP_002800252.1__2_' [illegible]

[illegible]

'CUR32202.1' [illegible]

[illegible]

'KPQ39684.1' [illegible]

[illegible]

'PPT09156.1' [illegible]

[illegible]

'WP_017662193.1' [illegible]

[illegible]

'WP_103112787.1__2_' [illegible]

[illegible]

'WP_104395902.1' [illegible]

[illegible]

'WP_002791883.1' [illegible]

[illegible]

'WP_002766200.1' [illegible]

[illegible]

'WP_002788626.1' [illegible]

[illegible]

'WP_002797932.1' [illegible]

[illegible]

'REJ40820.1' [illegible]

[illegible]

'WP_002742886.1__2_' [illegible]

[illegible]

'WP_002755926.1' [illegible]

[illegible]

'AKV70201.1__2_' [illegible]

[illegible]

[illegible]

'REJ60002.1' [illegible]

'WP_130756400.1' [illegible]

'REJ44384.1' [illegible]

'WP_024970397.1' [illegible]

'WP_125730344.1' [illegible]

'KXS91774.1' [illegible]

'WP_002778661.1' [illegible]

'WP_002762261.1' [illegible]

'EPF21995.1_4_' [illegible]

'AFZ42916.1_2_' [illegible]

'PNW54205.1' [illegible]

'WP_018399541.1' [illegible]

'WP_017306104.1' [illegible]

'WP_075891453.1_2_' [illegible]

'WP_024547051.1' [illegible]

'PZU97760.1' [illegible]

'OYQ61999.1' [illegible]

'WP_094536065.1' [illegible]

'WP_086766415.1_2_' [illegible]

'WP_096597869.1' [illegible]

'WP_048869210.1' [illegible]

'KYC39370.1_2_' [illegible]

'AFZ37990.1_2_' [illegible]

'WP_019503799.1' [illegible]

'WP_008312261.1' [illegible]

'WP_121971064.1' [illegible]

--TNLNSPE--E------E-H---YFPLLYLPTGVIYLTV--RAAPP-I-NPA---VLPD--RVVASI-----------------KSLCA--GQL--R-L-RQTGFSR-DG----------KG-MKYADYYSLF-FDTPGLM------------------EVALDATR----------------------RI-LN---------------------P------TK---GSVAS-----------------------------------KR------SE---NLV--------AFQ------------QKNVLSA-------DY---DFKF------GDDTTGDIRI-----
DQ--------LA-------------------E----FG---DLIS--RK--IWG-------EQI-------NQLEKIVKERAKARKK-QKD----LPELPDIPADVLPASLIGKIAES----WQLS-----DYLP-QV-------------REIQRINETLKSLKLKGNTGGVPYEWYFLAAKFL----E--K---NP-GI-E---DLRE----TCQNM-VAYVTALTQP-----------VSA---------QY----E--I--S-DG-WDD------------IRLWVGQVV---------MLP-----------------------
----EQTTPL--SGQADIF--LKELAAYSAAKK-SGRGRQL-ICSIS-HSPYSV---SEQMESAVLFTPQV--YTN--KQ-HL---GG--SNA-KRN-------ISSIAGIEMMLRQILMNQT-Q------AVGKRFEDGKYRYLYF-YP---TYYFTP----E-TNAF---LHK-----AYSNI-AQTRF-----DT------SIR-----NHFV-S---------------------------------------------KSLEA---------DFS------RSRY--QTVDAF-------L-
MDEELEQRK-------QLPADDPNFKRDR-------TF---KL---TY---PDD--------------------KPLTFYFMAL----------PP------G----R--DS--------TDT------ESW--VMPSWLALTFPMILDVKTVVSESPIPPFTDGAEFEES----VF--LDSAPQAFRTLV-G--QD-RF--------RLD---AILEGW------TDA---SGLK-RPAPLNT-LTAAYAIH-MDV------NAKQGKNGYDANWG-KLTEFAKDI-
ETSPLYVFSYLAKWSR---------GQK-------------ADA--P---SP----AKIR----------LYAHY--FYPCFDP-Y------------------------A-TYSFAQET--------W-NLK-TQ--SA-LNH-PKKLTDLYRSFYR--A-NKA--YNP--KANAVLKPVDIASETI-LKAESTMFQ--G-DALISVVAAEVFKLMDRVH-----N---STAEG-----RWVV-S--------D--------REKE----RQLVLDFARYFV--VDVFEEAFSGDRARLAGRQINLI--
---RDTCEFLYR------LEQDK--ENQ--ERQA-RKDSE--------A-DEAE-------
'WP_023075354.1__2_'    ----MEHPANSDVSKQQPS---------------QNNSAAEINTD---TFDASFLDG-------DFGAEEESN--------RTL--EPTAKELLTLKLLQEA---IQ--TQNSD--------DI--VMQD--FATMVL--PKLL-KY-TVGVTAK----G---------GKF-FAEYLDR--VVNP-------KRL-------AE---------GKS-------P---IR-RDNA-ADQSLNTHLLNGLFPANLIE--KR----------
-------------LEQL--D-TT-VKRVV---QEQERRLLISGFILHDFEKFDYDLI----------------P-G--MPEAFRAIRREP------AQDIRK-GS--------ITEHRAVVTVLVETL-GLA----DFI--CP--NRSEEWTQYLDDLLFLAYN-AQR---KNDTNLNLSEY------GL-NP----TL----RD-RTLR--CLADLTCLADRLASV------IKHPQDA-E-----NTG-----------INGII-H-SV---SD-GKL----K-------
FTYHAIAENRGVLTNVVNNALMEGH---TALNTAE--Q------E-I---YTPLLYLPTGVIYLAE--RDAPT-I-DAD---TLPN--RVVDTI-----------------KKLCA--GQL--R-L-RQTGFGR-DG----------KG-MKYADYYNLF-FDDLGLM------------------QVALDATL--------------------RI-LN---------------------P------NK---GSVAS--------------------------------------KR------SE---NLR--------AFQ------------QRNVLSA-----
--DY---DFTF------TD----DIRI-----DQ--------LA-------------------E----FG---DLIS--RK--IWG-------ERV-------KQLEQVIKERTKARKK-QKD----LPELPAIDDAVLPVNIVRQVAES----WQLS-----DYLP-QI-------------REIQRINETLKELKLKGNTGGVPYEWYFLAAKFL----E--Q---NP-GI-E---DLRQ----TCLNL-INHVTQLTQP-----------VVE---------QY----Q--V--S-DG-WDD-------------
IRQWVSNVV---------ILP-------------------------GYAQETST----QTDIF--LKELAYYTAAKK-PKRGVQK-ICSIS-HSPYSV---NEQMESAVLFTPQV--YTN--KQ-HL---GG--SNA-KRN-------ISSIAGIEMMLRQILMNQT-Q------AVGKRFEDGKYRYLYF-YP---TYYFTP----E-TNSF---LHK-----AYSNI-AQTRF-----DT------SIR-----NHFI-S-----------------------------------------------KQLKA---------NFS-
-----RERY--QTVDAF-------L-MEEDLVQRK-------QLPDDDPNFKRDR-------TF---KL---AY---PED--------------------KPLTFYFMAL----------PP------G----R--DS--------TDT------ESW--VMPSWLAFAFPMILDVKTVVSESPIPPFTDGAEFEES----VF--LDSAPHAFRTLI-G--QD-RF--------RLD---SILDGW------TDE---SGIK-RPAPLNT-LTAAYAIH-MDV------NAKQGKGGYDANWG-
RFTELAKDI-ETSPLYVFSYLAKWSR---------GQK-------------ADS--P---SA----QKIR----------LYAHH--FYPCFDP-Y------------------------A-PYNLTKET--------W-DLK-PE--SS-LNH-PKKLTDLYREFYR--A-NKR--YNP--KANAVLKPIDIAADTI-LKAESSMFH--D-EALVAVVAAEVFKLMDRVH-----A---STAEG-----RWII-S--------K--------REEE----RQKVLDFARYFV--
IDVFEKAFAGDRARLAGRQINLI-----RDTCEFLYR------LEQDK--ENQ--ELKA-KEDNN--------G-AAD--------
'EKV02086.1__2_'    ----MEHPANSDVSEQPSS---------------QDNSVDEINTD---TFDTSFLEG-------DFGAEEESN--------RTL--EPTAKELLTLKLLQEA---IQ--TQNSD--------DA--VMQD--FATMVL--PNLL-KY-TVGVTAK----G---------GKF-FAEYLDR--VVNP-------KRL-------AE---------GKA-------P---IR-RDNA-ADQSLNTHLLNGLFPANLIE--KR----------
-------------LEQL--D-TT-VKRVV---REQERRLLVSGFILHDFEKFDYDLI----------------P-G--MPEAFKAIRRDP------AQDIRK-GS--------IEDHRAVVTVLVETL-GLA----NFI--CP--NRPEEWTQYLDDLLFLAYN-AQR---KNDTNLNLSED------GL-NP----TL----RD-RTLR--CLADLTCLADRLASV------IKHPQDA-E-----NRG-----------INGII-H-SV---SD-GKL----K-------
FTYHAIAENRGVLTNVVNNALMEAH---TALNKAE--Q------E-I---YTPLLYLPTGVIYLAE--RDAPS-V-DAD---TLPN--RVVDTI-----------------KSLCA--GQL--C-L-RQTGFGR-DG----------KG-MKYADYYNLF-FDDLGLM------------------QVALDATL---------------------RI-LN---------------------P------NK---GSVAS--------------------------------------KR------SE---NLR--------AFQ------------QRNVLSA----
---DY---DFTF------TD----DIRI-----DQ--------LA-------------------E----FG---DLMS--RK--IWG-------ERV-------KQLEQIIRNRTKARKK-QKD----LPELPAIDDAVLPTNIVRQIAES----WQLS-----GYLP-QI-------------REIQRINETLKELKLKGNTGGVPYEWYFLAAKFL----E--K---NP-GI-E---DLRQ----TCLNL-ISHVTQLTQP-----------VVE---------QY----Q--V--S-DG-WDD-------------
IRQWVSNVV---------ILP-------------------------GYEQATAT----QTDIF--LKELAYYTAAKK-PKRGVQK-ICSIS-HSPYSV---NEQMESAVLFTPQV--YTN--KQ-HL---GG--SNA-KRN-------ISSIAGIEMMLRQILMNQT-Q------AVGKRFEDGKYRYLYF-YP---TYYFTP----E-TNSF---LHK-----AYSNI-AQTRF-----DT------SIR-----NHFI-S----------------------------------------------KQLEA---------NFS-
-----RERY--QTVDAF-------L-MEEDLVQRK-------QLPDDDPNFKRDR-------TF---KL---AY---PED--------------------KPLTFYFMAL----------PP------G----R--DS--------TDT------ESW--VMPSWLAFAFPMILDVKTVVSESPIPPFTDGAEFEES----VF--LDSAPHAFRTLI-G--QD-RF--------RLD---SILDGW------TDE---SGIK-RPAPLNT-LTAAYAIH-MDV------NAKQGKGGYDANWG-
RFTELAKDI-ETSPLYVFSYLAKWSR---------GQK-------------ADS--P---SA----QKIR----------LYAHH--FYPCFDP-Y------------------------A-SYNLIKET--------W-DLK-PE--SS-LNH-PKKLTDLYREFYR--A-NKR--YNP--KANAVLKPIDIAADTI-LKAESSMFH--D-EALVAVVAAEVFKLMDRVH-----A---STAEG-----RWIT-S--------K--------REEE----RQKVLDFARYFV--
IDVFEKAFSGDRARLAGRQLNLI-----RDTCEFLYR------LEQDK--ENQ--ELKA-KEDNN--------G-AAD--------
'WP_017326896.1'    ----------------ML---------------NEISDEGLGF----------LDS-------DYGFDEESD--------RTI--EPTARELLTLKLLREA---IQ--TQHPD--------DA--VMQD--FAEIVL--PNLL-KY-TVGVTAK----G---------GKF-FAEYLDR--VVNP-------RRI-------AE---------GKN-------P---IR-RDNA-ADQSLNTHLLNGLFPANLIE--QR-----------------------LERL--
D-TT-VKRLV---KARERRLLVSGFILHDFEKFDYDLI----------------P-E--MPEVYRAIRRDP------EQDIRK-RS--------IAEHRETIAVLMKAL-ELD----RFI--CP--TQPNEWQQYLDDLLFLAYN-AQR---RNDTNLNLSED------GL-NP----VL----RD-RTLR--CLADLTCLADRLASV------IKHPQDA-K-----HSG-----------LNEII-H-AL---SD-GQL----R--------FSYHAISENRGVLTNVVNNALMDAH---
TSLNGPE--E--------H---YTPLLYLPTGVIYLAA--RSAPT-V-SPN---SLPN--RVIASI-----------------KELCA--GQL--R-L-RQTGFGR-DG----------KG-MKYADYYNLF-FDDLGLM----------------QVALDATL---------------------RI-LN---------------------P------NK---LSVAT-------------------------------------KR------SE---NLI--------AFQ-------------QRGVLSL-------DY---SFTF------TD----DIRI-----DQ--------
LA-------------------E----FG---DLMS--RK--IWG-------ERV-------KQLETVIKERAKARKK-QKD----LAELPAIDEAVLPDNIIRKVAEF----WQLS-----EYLA-QI-------------REIQRINETLKQLKLKGNTGGVPYEWYFLAAKFL----E--Q---NP-GI-E---DLRE----TCRNA-IAYITQLTQP-----------IVA----------QY----E--L--S-DG-WDD-------------LRQWVSNVV---------MLP---------------------------
GQSQSSSV--SEQAEIF--LKELVYYNAAKK-PGRGRQM-ICSIS-HSPYSV---SEQMESAVLFTPQV--YTN--KQ-HL---GG--SNA-KRN-------ISSIAGVEMMLRQILMNQT-Q------TVGKRFEDGKYRYLYF-YP---TYYFTP----E-TNSF---LHR-----AYTNI-AQTHF-----DT------RIR-----NHFI-S-----------------------------------------------KQLEA----------DFS------RSRY--RAVDAF-------L-
MDENLDQRK-------QLPDDDPNFKRDR-------TF---KL---SY---PED--------------------KPLTFYFMAL----------PP------G----R--DS--------TDT------ESW--VMPSWLALAFPMILDVKTVVSESPIPPFTDGAEFEES----VF--LDSAPHAFRTLM-G--RD-RF--------RLD---TILDGW------TDN---SGTK-RSAPLNT-LTAAYAIH-MDV------NAKQGKGGYDANWG-RFTELAKDI-
ETSPLYVFSYLAKWSR---------GQK-------------ADA--P---SP----QKIK----------LYAQH--FYPCFDP-Y------------------------T-NYNFDKET--------W-DLK-PE--SL-LNH-PKQLTELYRQFYR--A-NKR--YNP--KANAVLKPIDIAADTI-LKAESSMFH--D-EALVTVVAAEVFKLMDRVH-----A---STAEG-----RWVV-S--------K--------REEE----RQKVLDFARYFV--
VDVFEQAFAGDRARLAGRQLNLI-----RDTCEFLYR------LEQDR--ENQ--ASDS-TSNTS--------E-EDG--------
'PSN19223.1__2_'    ----MDRPNQSGNPEQLSL---------------LDTQNEDSEFE---ELDDSWLEG-------DFGLDEESD--------RTV--EPVARELLTLKLLREA---IQ--AQNPG--------DA--VMQD--FAEIVL--PNLL-KY-TVGVTAK----G---------GKF-FAEYLDR--VVNP-------QRI-------AV---------GKD-------P---IR-RDNA-ADQSLNTHLLNGLFPANLIE--RR---------
--------------LQTL--D-TT-VRRQV---KERERRLLVSGFILHDFEKFDYDLI----------------P-R--MPEEFRAIRRDP------EGDIRK-RS--------IEEHREIITVLVETL-ELD----HFI--CP--ATPDDWQQYLDDLLFLAYN-AQR---RNDTNLNLSED------GL-NP----VL----RD-RTLR--CLADLTCLADRLASI------VKHPQDA-E-----NSG-----------LNDII-H-GL---SD-GQL----R-------
FTYHAISENRGVLTNVVNNALMDAY---ESLNIDP--E------NPH---YTTLLYLPTGAIYLAS--RDAPQ-I-SSD---SLPN--RVVDGI-----------------KALCA--GQL--S-L-RQTGFGR-DG----------KG-MKYADYYNLF-FDDPGLM------------------QVALDATL---------------------RI-VN---------------------P------NK---GSVAT-------------------------------------KR------SE---NLM--------AFQ------------QRGVLSG--
-----DY---DFSF------SD----DIRI-----DQ--------LA-------------------E----FG---DLMN--RK--IWG-------ERV-------NQLETVIKERAKERKK-QKD----LPELSEIKPEVLPENIIHKVAEF----WQLS-----EYSP-QI-------------REIQRINEKLKELKLKGNTGGVPYEWYFLAAKFL----E--K---NP-GI-E---DLRE----TCSNV-MAYVTQLTQP-----------VVA----------QY----P--V--A-DG-WDD-------------
VREWVGKVV---------MLP---------------------------GQTQETST--AEQVDVF--LKELAYYKAAKK-PGRGRQM-ICSIS-HSPYSV---NEQMESAVLFTPQV--YTN--KQ-HL---GG--SNA-KRN-------ISSIAGIEMMLRQILMNQT-Q------AVGKRFEDGKYRYLYF-YP---TYYFTP---------------------------------------------------------------------------------------------------------------------------------------------------------------------
----------------------------------------------------------------------------------------------------------------------------------------------------------------------------------------------------------------------------------------------------------------------------------------------------------------------------------------
------------------------------------------------------------------------------------------------------------------------------------------------------------------------------------------
'RMH16295.1'    ----MERPNQSGNPEQLSL---------------LDTQNEDSEFE---KLDDSWLEG-------DLGLDEESD--------RTV--EPVARELLTLKLLREA---IQ--AQNPG--------DA--VMQD--FAEIVL--PNLL-KY-TIGVTAK----G---------GKF-FAEYLDR--VVNP-------QRI-------AA---------GKE-------P---IR-RDNA-ADQSLNTHLLNGLFPANLIE--RR----------
--------------LQTL--D-TT-VRRQV---KERERRLLVSGFILHDFEKFDYDLI----------------P-D--MPESFRAIRRDP------EGDIRK-RS--------IEEHREIITVLVKTL-ELD----RFI--CP--ETPEDWKQYLDDLLFLAYN-AQR---RNDTNLNLSED------GL-NP----VL----RD-RTLR--QLADLTCLADRLASI------IKHPQDA-E-----SSG-----------LDEII-H-SL---SN-EQL----R-------
FTYHAIAENRGVLTNVVNNALMDAY---ESLNSDP--E------NPH---YTTLLYLPTGAIYLSR--RDAPQ-I-SLE---DLPD--RVVDGI-----------------KALCA--GQL--S-L-RQTGFGR-DG----------KG-MKYADYYNLF-FDDPGLM------------------QVALDATL---------------------RI-LN---------------------P------NK---GSVAT-------------------------------------KR------SE---NLI--------AFQ------------QRGVLSA---
----DY---DFSF------SD----DIRI-----DQ--------LA-------------------E----FG---DLIN--RK--IWG-------ERV-------NQLETVIKERVKARKK-QKD----LPELPAIKTEVLPEHIIHKVAEF----WKLS-----EYSP-QI-------------REIQRINEKLKELKLKGNTGGVPYEWYFLAAKFL----E--K---NP-GI-E---DLRE----TCSNV-IAYVTQLTQP-----------VIA----------QY----A--I--A-DG-WDD-------------
VREWVGKVV---------MLP---------------------------GQAQESST--AEQVDVF--LKELAYYNAAKK-PGRGRQM-ICSIS-HSPYSV---NEQMESAVLFTPQV--YTN--KQ-HL---GG--SNA-KRN-------ISSIAGIEMMLRQILMNQT-Q------AVGKRFEDGKYRYLYF-YP---TYYFTP----E-TNSF---LHR-----AYSNI-AQTRF-----DT------SIR-----NHFI-S-----------------------------------------------KQLEA--------
-DFS------REKY--QTVDAF-------L-MDEDLAQRK-------QLPDDDPNFKRDR-------TF---KL---SY---PED--------------------KPLTFYFMAL----------PP------G----R--DS--------TDT------ESW--IMPSWLAFAFPMILDVKTVVSESPIPPFTDGAEFEES----VF--LDSAPHAFRTLV-G--RD-RF--------RLD---FILEGW------TDE---SGIQ-RPAPLNT-LTAAYAIH-MDV------NAKQGKGGYDANWG-
RFTELAKDI-ETSPLYVFSYLAKWSR---------GQK-------------ADA--P---SP----QKIR----------LYAHH--FYPCFDP-Y------------------------A-TYNFDQED--------W-ILT-PD--SS-LNH-PKRLTDLFRQFYR--A-NKR--YNA--KANAILKPIDIAADTI-LKAESSMFH--D-DALVTVVAAEVFKLMDRVH-----N---STAEG-----RWVI-S--------D--------REKE----RQLVLDFARYFV--
VDVFEKAFAGDRARLAGRQINLI-----RDTCEFLYR------LEQDK--ENQ--KAKE-NELPK--------S-KSGEEE-----
'WP_054467001.1__2_'  --MAKKRKQSATETEQLSL-FDLNQATENPASNYIEAEIEAEIEAEIEDYDEVEIDD---IGYDELGITEESS-------DRTI--ETPS-ELLTLKLLREA---IA--AQNPN--------DL--VMAD--FAEYVL--PNLL-RV-AIGVTAK----G---------GKF-FDK-------IDQ-------QRE-------AA---------GKD-------K---IR-RDNA-GDQSLNTHLLNGLFPANLIE--QR-
----------------------LQQF--D-TT-VRRVV---KERERRLLVAGFILHDFEKFNYDLF----------------P-E--MPEKYRAIRRYF------TQDIRK-LS--------LEEHREIISVIVREL-GIN----QLI--DS--TNPESWSEYLDDLLFVAYN-AQA---KNDTNLNLSVH------GL-QP----KL----RD-KTLR--SLADLTRIADLMASV------IKHPQDA-E-----HRT-----------LNDIL-H-NL---SD-GNL----K-------
FTYHGIAENRGVLTNVVNNALIERY---EAINTAE--C------C-Y---YKPLLYLPTGVIYLAM--RQAPA-I-AET---DIPE--RVVNKI-----------------KQLCG--GQL--Q-R-RQTGFGR-DG----------KG-MKYAEYYSQF-FDEFGLM------------------QVGLAATL---------------------RI-LH---------------------G------TK---ASVAK-------------------------------------SR------SD---NLV--------KFQ------------EKQILSP------
-EY---DFRF------DD----DIRI-----DQ--------IA-------------------E----FG---DLVT--RK--IWG-------DRV-------TEIEK-------LCKQ-NKKTQQQLPELPD---------IVAEVAKL----WQLS-----AYLN-PI-------------REIQRINESLKEHKLKGNTGGVPYEWYYLAAKYV----Q--Q---NP-GL-E---DVTE----TCVAL-INHIAGLIQP-----------IVN----------QY----K--L--P-DG-WDD-------------LRDWVKRVV---------
ILP-------------------------GKTSPI----TQNAEVF--LDELDRYQLAKK-PGRGRQL-ICSIS-HSAYTV---TEQMESAVLFTPQV--YTN--KQ-ML---GG--SNA-KRN-------ISSIAGVEMMLRQILMNQT-Q------AVGKRFEHGNYRYLYF-YP---TYYFTP----E-TNKF---LQA-----AYTNI-SQTRF-----TA------PLR-----NHFI-T--------------------------------------------KEWQA---------KFD------RTQY--
QTVDSF-------L-IDEQLDKEK-------------------DR-------TF---KL---SY---PED--------------------QPLTFYFIAL----------PP------G-----K--DA--------KDT------ESW--VMPSWLAFAFPMILDVKTVVSESPIPPFNDGAEFEET----VF--LDSPPQAIQALT-Q--RD-RF--------RLD---YILEGW------EEGWEENQKK-YPAPLNV-LTAAYAIH-LDV------NAKQGKGGYDANWG-KLVELATNF-
QTSPLYVFNYLNRWVR---------RQG-------------VET--A---RI----EKIK----------LYAYA--FYPCFDP-Y------------------------V-EYQEDLMS--------LEIKS-PE--SP-LNH-PKKLTELYRQFYR--A-NKQ--FSP--KANAVLKPIDIAADTI-LNADPS-FQ--G-EALVDAVAAAVCKLMDRVH-----S---STAEG-----RWVF-N--------SK------ERDIE----RQKILEFARYFV--EEVFEKSFAGDRGRLAGRQINLI----
-RDTCEFLYR------LAQDK--ENQ--ENKE-NQPLN--------E-DD---------
'WP_094678459.1'    --MPRKKKKSDDDFEQLSL-FD------------MTENQENINPEAIDDSDDDWLED---E----ELGLSFEQT-------NRDI--EATHKELLTLQLLREV---II--SENKD--------DL--VIRE--FAEYVL--PNLL-KF-AIGVTAK----G---------GKF-FDR-------IDS-------RRE-------AE---------GKA-------K---VK-RNNA-ADQSLNTHILNGLFPANLIA--KR-----------
------------LEEL--D-TT-VKREMARDKEIKRRILIAGFLLHDFEKFKYNLL----------------P-G--MPQKYVDIDKNP------DQDIRK-LS--------IAEHKEIFRVIVPLL-GLD----KLI--NP--EKPDGWIEYLEDLLFIAYN-TQK---RNDTNLNLSES------GL-NP----KL----RD-RSLI--CLSNLTLLADLLSSI------IKHPQDA-E-----NSS-----------LHNII-H-QL---SD-GKL----E-------
LSYHSIAENRGVLTNIVNNAVMAAY---KNLNTEE--E------I-Y---YQPLLYLPTGVIYLIQ--KNSPP-I-SEE---DLPE--LVVSKI-----------------KELCA--KQL--R-L-RQTGFGR-DG----------KG-MKYAEYYDLF-FDTVGLM------------------EVALEATL---------------------II-LH---------------------S------GK---KSVAK-------------------------------------SR------SD---NLV--------KFQ------------QQQVLSA-------
DY---NFRF------DD----DLRI-----DH--------IA-------------------E----FG---DVLT--RN--IWG-------EVV-------NNIDK-------ARKQ-NKD----LPELPA-N--LETKNLVHIIAEY----WELS-----AYIP-----------------------------------------------------------------QIRE-----------------------------------------------------------------------------------------
-------------------------------------------------------------------IQK------------------------------------------------------------------------------------INESLKAXN--------------------------------------------------------------------------------------------------------------
----------------------LEM---------------------------------------------------------------------------------------------------------------------------------------------------------------------------------------------------------------------------------------C-------------------------------------------------
'WP_094678461.1'    ----------------------------------------------------------------------------------------------------------ML----------G--------------------------------------------------------------------------------------II-----------------------------------------------------------------------------------------------------GIH------
---SP--------------------N-SGN------------------------------PKNQ-----------------RKSQSA-X-------------------FGDVL-T----------RNI-----------------------------------------------------------------------------W---------G--------------------------EVVNNI--------------------------------------------------DK-----------------------------------------------------------------AR---------
KQ------NK----------------------DLPELPA---------------------NLET------------------------KN----------------------------------------------LVHIIAEY----WELS-----AYIP-QI-------------REIQKINESLKAQKLKGNTGGVPYDWYFLAAKYL----E--S---NP-GI-E---DVKE----ICQKT-ISHISQLIKP-----------ILE----------QY----K-----FADG-WDD-------------
LREWVKQVV---------MLP---------------------------GKTEITKTSAEVF--LDELSNYQLAKK-PGRGRKL-ICSIS-HSPYSV---TEQMESAVLFTPQV--YTN--KQ-ML---GG--SNA-KRN-------ISSIAGIEMMLRQILMNQT-Q------ATGKKFEDSKYRYLYF-YP---TYYCTP----E-TNKF---LQK-----AYNDI-AQTRF-----DT------SIR-----
NHFIDRDFQANFTREKYQSVDTFLIXETNKFLQKAYNDIAQTRFDTSIRNHFIDRDFQA---------NFT------REKY--QSVDTF-------L-IDENLEPEK--------------------DR-------TF---KL---SY---PED--------------------QPLTFYFMAM----------PP------G-----K--GD--------TDT------ESW--VMPSWLAFAFPIILDVKTVVSESPIPPFNDGSEFEET----VL--LDTAPQAFRSLV-R--GD-RF--------RLD---
YILEGWQ-----------ENGKEYPAPLNV-LTAAYAIH-LDV------NAKVGKGGYDANWG-KLTELAKDF-ETSPLYVFSYLNKWVR---------NQK-------------VET--V---RI----EKIK----------LYAYY--FYPCFDP--------------------------KVRYNQ------------EEDKLEIMGKESSLNH-PQKLTELYREFYRANKKYNP-------TTNSVLKPIDIAKDVI-LNNDPSFQN-----ETLVNNVAAKVYKLMDKIH-----A---
STAEG-----RWVLSN-----------------REEE----RKAVWKFANYFV--KDVFYETFGGDRARLAGRQINLI-----RDTCEFLYR------LENDK--ENR--EIPK-----------------------------
'WP_052128200.1'    ----MVKQIQNHDQAQLSI-FK-----------VEETIVESDGD---LSDDSWLEG-------DDGLDGETS-------DRTI--APPERELLTLKLLREA---IQ--AENPA--------DL--VMRD--FAEYVL--PNLL-QV-AIGVTAK----G---------GRF-FDD-------LDQ-------QRE-------AE---------GKT-------K---VR-RDNA-ADQSLNTHLLNGLFPANSIA--RR-----------
------------LERL--N-TT-VKRVV---RERERRLLAGFILHDFEKFNYARF----------------R-G--IPQKYQDIQIAG------EEKIRQ-LS--------LPEHRELVDSICKEL-GID----HLV--NP--GEPSKYQEYLDDLLVVIYN-AQR---RWDTNWNFSTF------GLSNL-----VL----PD-RTLG--CLADLSYLADLLSSI------IKHPQNV-E-----KNT-----------LFEIM-H-KL---SD-GQL----R-------
LTYHSIAENRGVLTNVVNNAVMEAY---TSLNTEN--Y------T-H---YEPLLYLPTGVIYLAD--RNAPP-L-NPA---GLPD--RVVAKM-----------------KQLCF--EQL--R-L-RQTGFGR-DG----------KG-MKYADYYTLF-CDDRGLM------------------TIALDATL---------------------RV-LS---------------------A------NK---SSVAP-------------------------------------SR------TE---NLL--------RFQ------------QQGTLSA--
-----DY---DFHM------KN----DIRI-----DQ--------LA-------------------E----FG---DVIS--RK--IWG-------DRY-------ENLELARKGRSQARKK-NKA----LPELPAID--LHPSDLIHRIAES----WGLA-----SYSA-PI-------------REIQRINETLKEQGLKGNTGGVPYEWYFLAATYL----D--H---HP-GL-E---TIRE----TGETI-INVVANLIEP-----------IIA----------QY----K--L--P-DG-WDD-------------
LRQWVNQVV---------MLP---------------------------GSNQPPPDQEQEQADIF--LKELAHYTAAKK-PGRGRQL-ICSIS-HSAYSV---TEQMESAVLFTPQV--YTN--KQ-ML---GG--SNA-KRN-------ISSIAGLEMMLRQILMNQT-Q------AVGKRFEDGKYRYLYF-YP---TYYFTP----E-TNKF---LQK-----AYSSI-AQTRF-----DT------SIR-----NHFI-S-----------------------------------------------KDLEA-------
--NFE------RSRY--QTVDAF-------L-MDEGLDRKK-------KLDEADPEYKRDR-------TF---KL---AY---PDD--------------------QPLTFYFMAL----------PP------G-----K--DP--------TDT------ESW--VMPAWLAFAFPMILDVKTVVSESPIPPFNDGAEFEES----VF--LDSAPQAFRVLT-G--RD-RF--------RLD---SILEGW------QNP---DGTK-QAAPLTV-LTAAYAIH-LDV------NAKQGKSGYDANWG-
KLTELAKDF-ETSPLHVFSYLSRWVR---------NQK-------------LDA--P---ST----SKMK----------LYAYH--FYPCFDP-Y------------------------T-KFDPNLEA--------L-TVE-PE--SA-INH-PQKLTDLYRQFYR--A-KKS--FNP--KSNAVLKPIDLAAETI-LKAESTVFQ--G-ETLVMAVAAEIFKLMDRVH-----A---STADG-----RWII-S--------D--------REDE----RQAVLSFAHYFV--
KDVFEGAFAGDRARLAGRQLNLI-----RDTCEFLYR------LAQDK--ENQ--EWRK-NNPDK----VKSKS-PEDELEDD---
'PSB20197.1__2_'    -----------------------------------------MANLT--NDEFDFLEG-------DFGFDEDSD--------RTL--SSTQRELLTLKLLREA---IQ--SQNPD--------DA--VMQD--FAEYVL--PNLL-RV-AIGVTAK----G---------GKF-FDE-------LDR-------KNE-------AA---------GKA-------K---VR-RDNA-ADQSLNTHLLNGLFPANLIE--RR-----------------------LAKL--
D-TT-IKRVV---RDRERRLVIAGFILHDFEK-----F----------------P-D--APDD-----------------CRK-LP--------LDAHRQIIDQKVRQL-GLD----RFI--DS--SDLTTYKAYVDDLLCMAYN-AQR---RWDTNWNFSEF------GL-NP----TL----KD-RVLR--SLSDLTCLADSLASI------VKHPYDA-D-----NTR-----------LNDLL-H-NL---SD-GQL----H-------FTYHSISENRGVLTNVLNNALVEAH---TSLNTAD--R-
-----T-Y---YEPLLYLPTGVIYIAQ--KDAPK-V-ERS---GLSD--RVVVSI-----------------KALCS--GQL--R-Q-RQTGFGR-DG----------KG-MKYADYYNLF-FDDPGLM------------------QVALDATL---------------------RI-LR---------------------V------GK---PSVAT-------------------------------------SR------SE---NLV--------KFQ------------QQNVLPA-------DF---DFKF------SD----DIRI-----DQ--------IA-----------
--------E----FG---DVIS--RK--IWG-------ERL-------AKIET-------ARKQ-NKK----LPAPVAID-------LVQKVAEF----WNLS-----EYLP-QI-------------RAIERINESLRELKLKGNTGGVPYEWYYLAAKYL----E--H---HP-GI-E---SVRK---TCEKV-ITQIAAFTRP-----------IAD----------AY----Q--L--P-DG-WND-------------LREWCDRVI---------MLP---------------------------SATLEQSI--AKHADIF--
LKELSNYNNAKK-SGRGRQL-ICSIS-HSAYTV---TEQMESAVLFTPQV--YTN--KQ-TL---GG--SNA-KRN-------ISSISGIEMMLRQILMNQT-Q------TVGKRFEDGKYRYLYF-YP---TYYFTP----E-TNRF---LQK-----AYSGI-AQTRF-----DT------SIR-----NHFI-S---------------------------------------------KDLDA----------NFE------RDRY--QTVDAF-------L-IDEDLDRKK-------NLDETDPEFKRDR--
-----TF---KL---AY---PDD--------------------KPLTFYFMAL----------PP------G----K--DP--------TDT------ESW--VMPAWLALAFPMILDVKTVVSESPIPPFNDGAEFEES----VF--LDSAPQAFRMLL-G--ND-RF--------RLD---HILEGW------KTA---DGVQ-RSAPLNT-LTAAYAIH-LDV------NAKQGKGGYDANWG-RLTELAKDL-DTSPLYVFSYLVKWAK---------KQS-------------SDT--P---SP----
QKIR----------LYTHH--FYPCFDP-Y------------------------V-KFDANQEM--------L-IVN-GE--SP-LNH-PQKLTELYRKFYR--A-KQR--YNP--KANAVLKPIDVAAEVL-LKADTSVFQ--G-EVLVMVVAAEIFKLMDRVR-----N---SRAEG-----FTQI-R--------D--------REAE----RQAILEFARYFV--EDVFEQAFAGDRARLSGRQLNLI-----RDTCELLYR------LEQDN--DNR--DLKA-KGEVI----EESET-
EN---------
'RNJ65627.1'    ------MNPVNQEPDQLSI-------------------LSSLDGTS--TLESDWLED---DTDLE-----AVA-------NRTI--EPAQRELLTLKLLRAA---IQ--AQNPD--------DA--VMND--FANYVL--PNLL-EI-AIGVTAK----G---------GNF-FDE-------LDR-------QRT-------AA---------DQA-------K---VR-RDNA-ADQSLNTHLLNGLFPANLVE--QR-------------------
----LERL--N-TT-IRRVV---RERERRLVIAGFILHDFEK-----F----------------P-D--VPND-----------------CRK-LT--------VEEHRQIIQQKVQQL-GLD----YLI--SP--KTPSAYTEYIDDLLCIAYN-AQR---RWDTNWNFSEY------GL-NP----SL----ND-RTLR--NLSNLTCLADSFASI------VKHPQDA-E-----HTR-----------LKELI-H-EL---SD-GQL----R-------FTYHSIAENQGVLTNIVNNAVMEAH---
TQLNTED--C------I-Y---YEPLLYLPTGVIYLAD--RNAPP-I-DPG---GLPN--RVVAKI-----------------KQLCF--EQL--K-L-RQTGFGR-DG----------KG-MKYAYRYLF-CDDRGLM------------------EIALNATL---------------------RI-LS---------------------S------NK---SSVAP-------------------------------------NR------SA---NLK--------KFQ------------QVGALSA-------DS---DFSL------KD----DIRI-----DQ-------
-LA-------------------E----FG---DVIS--RK--IWG-------NRV-------EQIDQVVKARAQARKK-QKT----LAELPPVE--LTPSNLVHRVAEF----LGLA-----AYLP-QI-------------REIQAINTTLKENKLPGNTGGVPYEWYFLATRYL----D--G---S-GI-E---NVRE----VGEKI-IEYVADLIEP-----------IVA----------QY----Q--L--A-DG-WDD-------------LRDWIQRVV---------MLP----------------------
TVTKTQSA--TEQAEMF--LKEFAGYSAAKR-TGRGKQA-ICSIS-HSAYSV---KEQMEAEVLFTPQV--YTN--KQ-ML---GG--SNA-KRS-------ISSIAGLEIMLRQILMNQT-Q------ATGKRFEDGKYRYLYF-YP---TYYFTP----E-TNKF---LQK-----AYRSI-AQTRF-----DT------SLR-----NHFI-S-----------------------------------------KDLQA---------DFS------QFQY--QTVDAF-------L-
MDEDLEQRM-------ALPETDPNYRRDR-------SF---KL---SY---PDD--------------------QPLTFYFMAL----------PP------G-----K--DP--------TDT------ESW--VMPTWLAFAFPLILDVKMVVSESPIPPFNDGAEFEES----VF--LDSAPQAFRMLC-G--QE-RF--------RLD---AILQGW------RTA---DGTQ-RSAPLNR-LTAAYAIH-LDV------NAKQGKGGYDPNWS-KLTELARDF-
ETSPLYVFNYLNRWVR---------KQE-------------SET--A---RI----EKIR----------LYTYH--FYPCFDP-Q------------------------V-NFDSSLEQ--------L-IVQ-ES--SP-INH-PQKLTDLYRQFYR--A-GKR--YNP--KANAVLKPIDVAAEVI-LKAESSVFY--G-ETLVMAVASEVFKLMDRVR-----A---SRAEG-----FAVI-S--------D--------REAE----RQAILAFARYFV--ENLFEQAFAGDRARLAGRQLNLI-----
RDTCEFLYR------LAQDK--ENQ--ELAA-KGEPV----QPEED-DG---------
'WP_106171784.1__2_'  ----MPKKIQNHDSAQLPL-------------------FEIEETPV--AIDLDQLDD---LLEDEFGFDGEPA-------DRTI--ESTQRELLTLKLLREA---IR--SQNPD--------DV--VMRD--FAEYVL--PNLL-KV-AIGVTAK----G---------GKF-FDD-------LDQ-------QRE-------AQ---------GKA-------R---VR-RDNA-ADQSLNTHLLNGLFPANLIE--QR--------------
---------LEKL--D-TT-IKRVV---RECERRLVIAGFILHDIEK-----F----------------P-D--SPDN-----------------CRK-LP--------LEQQRQLFEQKIQQL-GID----RFI--NP--DALEVYKEYLDDLMCIAFN-AQR---RWDTNWNFSEY------GL-NP----AL----KD-RSLR--QLADLTCLADSLASI------VKHPQDA-E-----HYR-----------LTELL-H-NL---SD-GQL----K-------FTYHSISENRGVLTNVVNNAVMEAY---
VSLNIEN--C------T-Y---YEPLLYLPTGVIYLAH--KDAPP-I-DSQ---VLPD--RVVTKI-----------------KQLCF--EQL--R-L-RQTGFGR-DG----------KG-MKYADYYTLF-CDDRGLM------------------QIALDATL---------------------RI-LT---------------------P------NK---SSVAP-------------------------------------SR------SA---KLV--------EFQ------------QQGVLSA-------DD---DFRL------KD----DIRV-----DQ-------
--FA-------------------E----FG---DVIS--RK--IWG-------DRV-------ERIEQFRKEQLQARKK-NKS----LLGLPPVE--LAAKDLVHKVAEF----LGLA-----QYSP-QI-------------REIQHINETLKEMKLKGNTGGVPYEWYFLATKFI----E--G---S-GI-E---NIRE----TGEKI-IAYLAELIEP-----------IAA----------QY----S--L--P-DG-WDD-------------LREWVKRVV---------MLP----------------------
GNAQKQNA--TEQADIF--LKELTSYNAAKK-IGRGKLA-ICSIS-HSAYSV---KEQMESEVLFTPQV--YTN--KQ-ML---GG--SNA-KRN-------ISSIAALEMMLRQILMNQT-Q------AVGKRFEDGKYRYLYF-YP---TYYFTP----E-TNKF---LQK-----AYSGI-AQTRF-----DT------NIR-----NHFI-P------------------------------------------KDLEA---------NFE------RSRY--QTVDAF-------L-IDEDLDRKK--
-----ALPENDPQYKRDR-------TF---KL---SY---PDD--------------------QPLTFYFMAL----------PP------G-----K--DP--------TDT------ESW--VMPTWLAFAFPMILDVKTVVSESPIPPFNDGAEFEES----VF--LDSAPQAFRVLT-G--HD-RL--------RLD---YILEGW------KAP---DGSQ-HPAPLNV-LTAAYAIH-LDV------NAKQGKGGYDANWG-KLTELAKDF-ETSPLYVFSYLNRWVR---------NQA---
----------SET--A---RI----EKIR----------LYAYR--FYPCFDP-Y------------------------V-EFDVVSER--------L-TVK-DEGKSS-INH-PQKLTDLYRKFYR--A-NKR--YNP--KANAVLKPIDVAADVI-LKAESTVFQ--G-GTLVVAVASEVFKLMDRVR-----A---ARAEG-----FAMI-P--------D--------REEE----RQAILDFARYFV--VEVFEKSFSGDRARLAGRQLNLL-----RDTCEFLYR------LEQDK--ENR--
ERKE-KGEPG----NDENG-DD---------
'PSB56979.1'    ------------------------------------------------------------------------------------REA---IR--SQNPD--------DV--VMRD--FAEYVL--PNLL-KV-AIGVTAK----G---------GKF-FDD-------LDQ-------QRE-------AQ---------GKA-------R---VR-RDNA-ADQSLNTHLLNGLFPANLIE--QR-----------------------LEKL--D-TT-IKRVV---
RECERRLVIAGFILHDIEK-----F----------------P-D--SPDN-----------------CRK-LP--------LEQQRQLFEQKIQQL-GID----RFI--NP--DALEVYKEYLDDLMCIAFN-AQR---RWDTNWNFSEY------GL-NP----AL----KD-RSLR--QLADLTCLADSLASI------VKHPQDA-E-----HYR-----------LTELL-H-NL---SD-GQL----K-------FTYHSISENRGVLTNVVNNAVMEAY---VSLNIEN--C------T-Y---
YEPLLYLPTGVIYLAH--KDAPP-I-DSQ---VLPD--RVVTKI-----------------KQLCF--EQL--R-L-RQTGFGR-DG----------KG-MKYADYYTLF-CDDRGLM------------------QIALDATL---------------------RI-LT---------------------P------NK---SSVAP-------------------------------------SR------SA---KLV--------EFQ------------QQGVLSA-------DD---DFRL------KD----DIRV-----DQ--------FA-------------------E---
-FG---DVIS--RK--IWG-------DRV-------ERIEQFRKEQLQARKK-NKS----LLGLPPVE--LAAKDLVHKVAEF----LGLA-----QYSP-QI-------------REIQHINETLKEMKLKGNTGGVPYEWYFLATKFI----E---------GS-GIE-NIRE----TGEKI-IAYLAELIEP-----------IAA----------QY----S--L--P-DG-WDD-------------LREWVKRVV---------MLP---------------------------GNAQKQNATEQADIF--
LKELTSYNAAKK-IGRGKLA-ICSIS-HSAYSV---KEQMESEVLFTPQV--YTN--KQ-ML---GG--SNA-KRN-------ISSIAALEMMLRQILMNQT-Q------AVGKRFEDGKYRYLYF-YP---TYYFTP----E-TNKF---LQK-----AYSGI-AQTRF-----DT------NIR-----NHFI-P------------------------------------------KDLEA---------NFE------RSRY--QTVDAF-------L-IDEDLDRKK-------ALPENDPQYKRDR--
-----TF---KL---SY---PDD--------------------QPLTFYFMAL----------PP------G-----K--DP--------TDT----------------------------------------------------------------------------------------------------------------------------------------------------------------------------------------------------------------
------------------------------------------------------------------------------------------------------------------------------------------------------------------------------------
'EDX77106.1__2_'    --MTKKPHQSEQQIQQLSL-FD-----------ITP-----------DTDDDWLDN---D---DLGFNTEPS-------NRTL--ETPQ-SLLTLTLLREA---IQ--AQNPD--------DP--VMQD--FGKYVL--PNLL-QF-AIGVTAK----G---------GKF-FDH-------LDQ-------QRE-------AE---------GKE-------K---VR-RDNA-TDQSLNTHLLNGLFPANLIQ--KR----------------
--------LKTL--N-TT-VKRVV---GERERRLVTAAFILHDFEK-----F----------------P-D--VPKD-----------------CRK-LS--------LDEHRQIIDHKVRQL-GLD----SLI--NP--ENPDTYSEYIDDLLCMAYN-AQR---RWDTNWNFSEY------GL-NP----VL----RD-RTLR--CLSELTCLADSLSSI------VKHPQDA-T-----NPR-----------LKELI-H-SL---SD-GQL----T-------FTYHSIAENRGVLTNVVNNALIQIH-------
-QT--H------S-D---IEPLLYLPTGVIYLAH--HHAPG-I-SLA---GLSD--QVVARI-----------------KSLCR--EQL--R-L-RQTGFGR-DG----------KG-MKYADYYQLF-FDESGLM------------------EVALDATL---------------------RI-LN---------------------P------NK---ASVAK-------------------------------------SR------SD---NLV--------KFQ------------QQGILSA-------DY---DFKF------GD----DIRI-----DQ--------LA---
------------------E----FG---DLVS--RK--IWG-------ERV-------NQIEA-------ARKQ-DKQ----LPELPHLD-------LVHKLAEY----WHLG-----EYLP-PI-------------REIQQINQRLKQKKLKGNTGGVPYEWYFLAAKYL----Q--K---HP-GL-E---TIRD----TGEEL-IAYATALIQP-----------LIS----------EY----Q--V--P-DG-WDD-------------LRLWLERVV---------MLP---------------------------SYRDD-----
KATVDVF--LAELATYQGAKK-SGRGRQL-ICSIS-HSPYTV---TEQMESAVLFTPQV--YTN--KQ-ML---GG--SNA-KRN-------ISSIAGIEMMLRQILMNQT-Q------AVGKRFEEGKYRYLYF-YP---TYYFTP----E-TNLF---LQK-----AYSGI-AQTRF-----NT------QIR-----NHFI-S------------------------------------------QELKA----------DFS------RRRY--QTVDMF-------L-IQNTLEPGR------------------
DR-------TF---KL---AY---PDD--------------------QPMTFYFMAI----------PP------G----R--DS--------TDT------ESW--VMASWLGLAFPMILDVKTVVSESPIPPFTDGAAFEET----VF--LDSAPPAFRALV-K--RD-RF--------RLD---YILEGW------TE----NRIE-YPAPLNV-LTAAYAIH-LDV------NAKSGKSGYDPNWG-KLTELANEF-ETSSLSVFSYLNRWVR---------RHS-------------QDT--A---PL----
NKIR----------LYAYQ--FYPCFDP-Y------------------------T-YYDSDREE--------L-IVT-EQ--SS-LNH-PKQLMELYRQFYR--A-NKR--YNP--KANAVLKPIDIAADTI-LKADTTVFQ--G-EALVMAVAAEVFKLMDRVH-----G---STAEG-----RWVI-S--------N--------REKE----REAIIEFARYFV--GEVFEKSFRCDRARLAGRQLNLI-----RDTCEFLYR------LEDDK--ENR--QRQE-ELTED---------G-
IKEG-------
'WP_017323372.1'    -------------------------------------------------------MDYLDN---H---DFGFEGNSD--------RTI--ETQE-ELLTLRLLREA---IE--AQNPG--------DT--VMKD--FANYVV--PNLL-RI-ATGVTAK----G---------GKF-FDR-------IDQ-------QRQ-------TE---------GLT-------K---VR-RNNA-ADQSLNAHLLNGLFPANLIE--RR-----------------------LQQL--D-TT-
VRRAV---EEKERRIAIAGFILHDFEK-----F----------------P-D--VPEN-----------------SRT-LS--------PAKHRQIIDEKIRQL-GLD----KFI--DP--KHPETYQNYLDDLLCLAYN-AQR---RWDTNWNFSES------GL-RP----AL----KD-RTLR--SLSDLTCLADSLASI------VKRPQDA-E-----NPR-----------LKEII-H-NL---SD-GQL----K-------LTYHSIAENRGVLTNIVNNALIQAH---ITLNTDT--Q------T-Y---
YEPLLYLPTGVIYLAT--RHAPP-V-STE---NLPE--LVVNKI-----------------KSLCT--EKL--K-L-KQTGFNR-DG----------KG-IKYAEYYDLF-FDPPSLM------------------QVALDATL---------------------RI-VN---------------------S------QK---ESYAR-------------------------------------SR------SEKLVNLV--------KRQ------------KQGNLPV-------K----NFEF------SD----DIRI-----DQ--------IA-------------------E----
FC---DLVS--RK--IWN-------AIV-------NRIRA-------VSKK-NKN----SPSIPTLD-------LTDEIVKF----WNLE-----QYTQ-EI-------------RHIQDLS----------NTGGTPYEWYYIAAKYL----Q--K---NP-GI-E---DVRQ----TCTQL-INYLTNIISP-----------ILA----------QY----K--F--P-DG-WED-------------LRQWVLQVV---------MLP---------------------------TVDQQLSP--HTRTENF--LKELNNYNAAKK-
SGRGRQL-ICSIS-HSAYTV---TEQMESAVLFTPQV--YTN--KQ-ML---GG--SDA-KRH-------ISSIAGLEIMLRQILMNQT-Q------AVGKKFEDSNYRYLYL-YP---TYYFTP----E-TNKF---LHK-----VYSRI-AQTRF-----DT------SIR-----NHFI-T--------------------------------------------QDLQA---------HLE------KENY--QSVDAF-------L-INENPD-------------------SK-------TF---KL---SY---PED--------
-------------EPLTFYFMAL----------PP------EQKNTK--PP----EQKNTNT------ESW--IMPCWLALTFPIILDVKTVVSESPIPPFNDGAEFEET----VF--FDSAPYAFRTLV-K--RD-KF--------RLD---YILEGW------NE----NDIQ-YPAPLNV-LTAAYAIH-LDV------NARREKSVYNPNWG-KFTELAQDF-ETSPLYVFSYLNRWVR---------KQK-------------LET--A---RI----EKIR----------LYTYY--
FYPCFDP-Y------------------------A-KFDPMREE--------L-IIQ-QE--SS-LNH-PQKLTELYRRFYR--A-NRR--YNP--KSNAILKPINIAAETI-LKAESSVFH--G-ETLIAAVAAEIFKLMDRVR-----A---STAEG-----RWVI-S--------K--------REEE----RQAILDFARYFV--VEVFEKSFAGDRARLAGRQLNLI-----RDTCEFLYR------LEDDK--ENA--QNHE-NSPES---------D-RENE-------
'WP_046276749.1__2_'  --MAKKRQKSDPEGQQLSL-FD-----------QPQVINTEIDE-----DDSWLEN---E----EFGLNFEFS-------DRTI--EITS-ELLSLKLLKEA---IQ--AQNSD--------DV--VMGD--FAEYVL--PNLL-RV-AIGVTAK----G---------GKF-FDE-------LDR-------RR--------------GRE-------N---VR-RDNA-GDQSLNTHLLNGLFPANLIE--QR--------------
---------LEKL--D-TT-VKRVI---KEKERRLGVAGFILHDFEKFNYRLF----------------P-S--MPEKYQEIRQDF------DKDIRK-LS--------VEDHREIIEVIVLEL-RLD----RFI--CP--ENPESWREYIDDLLFIAYN-AQE---RNDTNLNLSEH------GL-NV----IL----SD-RKLR--QLIDLTRLADLLASV------IKHPQDA-E-----HTK-----------LNELL-H-SL---SD-GQL----K-------
FTYHSISENRGVLTNVVNNALMEIH---KNLNTED--I------I-Y---YEPLLYLPTGVIYLKH--REAPP-V-SIK---ELSN--RIVDNI-----------------KKLCS--QRL--L-D-TQTGFGR-DG----------KG-MKYAEYYEQF-FDEVGLM------------------KVALKATL---------------------RI-LH---------------------P------KK---ASVAK-------------------------------------SR------SE---NLI--------QFQ------------RREVLSQ-------
DY---DFSF------AD----DIRI-----DQ--------IA-------------------E----FC---DVIS--RK--VWG-------EKV-------NQIEDLIKERNLECKK-NKD----LPVLPAVT--LKPDDLVIKVAEY----WNLT-----EYLR-QV-------------REIQRINESLKEKKLKGNTGGVPYEWYYLAAKYI----Q--K---NP-GI-E---DVRE----NCEEL-IQYIAELIQP-----------IVE----------KY----Q--L--S-DG-WED-------------LKLWVNRVV----
-----MLP---------------------------EKTTEENQVNSQQTQIF--LDELERYQLAKK-SGRGRQL-ICSIS-HSAYTV---TEQMESAVLFTPQV--YTN--KQ-ML---GG--SNA-KRN-------ISSIAGTEMMLRQILMNQT-Q------AVGKRFEDGKYRYIYF-YP---TYYFTP----E-TNKF---LQK-----AYSNI-AQTRF-----NA------GIR-----NHFI-S-------------------------------------------EDLKA---------NFE------REKY--
QSVDSF-------I-IDEDLQTKK-------NLPEDDPNRKIDR-------TF---KL---SY---PED--------------------QPLTFYFMAL----------PP------G----R--DG--------TDT------ESW--VMPTWLAFAFPMILDVKTVVSESPIPPFNDGAEFEET----VF--LDSAPQAFRILT-K--RD-RF--------RLD---YILEGW------IE----SGQS-YSAPLNV-LTAAYAIH-LDV------NARQGKSGYDANWG-KLTELANNI-
ETSSLSVFSSLKEWIR---------RQG-------------VET--A---RI----EKIK----------LYAYD--FYPCFDP-Y------------------------V-TYKPDLED--------L-IVE-EK--SP-LNH-PKKLTELYRKFYR--A-SKR--YNP--KSNAVLKPIDISAETI-LKAESSVFH--G-EALVNAVAAEIFKLMDRVH-----A---STAEG-----RWVF-S--------K--------REEE----RQAVLEFARYFV--EDVFYGSFKGDRARLAGRQINLI-----
RDTCEFLYR------LEDDK--ENK--QSAN-ESTDK----PAKTE-EE---------
'WP_023065083.1__2_'  --MAKKRQKSEPEGKQLSL-FD-----------QPEVINTETDE-----DDSWLED---E----EFGLNFEPY-------DPFF--EPETSELLTLKLLREA---IQ--SQNPD--------DA--VMAD--FAEYVL--PNLL-RV-AIGVTAK----G---------GKF-FDE-------LDE-------KRE-------AE---------GKT-------K---VR-RDNA-GDQSLNTHLLNGLFPANLIE--QR----------
-------------LEKL--N-TT-VKRVV---GEKERRLVIAGFILHDFEK-----F----------------P-D--APEN-----------------CRQ-LP--------IEEHRQIITDKIKQL-GLD----YFI--NP--DDSNAYQDYIDDLLCVAYN-AQR---RWDTNWNFSVF------GL-NP----QL----RDHRTLV--CLSDLTCLADSLASI------IKHPQDA-E-----NTR-----------LKELI-H-SL---SD-GQL----Q-------FTYHSIAENRGVLTNIVNNALMEAH-
--TALNTED--K------T-Y---YQPLLYLPTGVIYLTQ--KNSPK-I-SAS---DIPQ--RVIDKI-----------------KSFCS--RRL--I-D-SQMGFGR-DG----------KG-MKYADYYEQF-FDIADLM------------------KVALKATL---------------------RI-LT---------------------P------KK---SSVAK-------------------------------------SR------SD---NLV--------KFQ------------RQNVLSS-------DY---NFIF------ED----DIRI-----DQ--------
IA-------------------E----FG---DVIT--RK--IWG-------EQV-------NKIEK-------ERKQ-NKK----LPELPHLD-------LIHEVAKF----WKLE-----QYSP-QI-------------REIQKINESLKANKLKGNTGGVPYEWYFLAAKYI----Q--D---K--GI-E---DVRE----HCEQV-IDYINKLIEP-----------IVK----------QY----T--L--P-DG-WDD-------------LREWVRRVV---------MLP---------------------------GKPTTDSL--SADI--
F--LKELNHYQLAKK-SGRGRQL-ICSIS-HSPYSV---TEQMESAVLFTPQV--YTN--KQ-ML---GG--SNA-KRN-------ISSIAGTEMMLRQILMNQT-Q------AVGKRFEDGKYRYLYF-YP---TYYFTP----E-TNKF---LQQ-----AYQNI-AQTRF-----DT------SIR-----NHFI-S------------------------------------------KDLKA---------NFE------RSRY--QSVDTF-------L-IDEALQYKK-------
ELPETDSNHKKDR-------TF---KL---SY---PED--------------------QPLTFYFMAL----------PP------G----R--DP--------TDT------ESW--VMPTWLAFAFPMILDVKTVVSESPIPPFNDGAEFEES----VF--LDSAPQAFRVLT-K--RD-RF--------RLD---YILEGW------EE----AGKK-YSSPLNV-LTAAYAIH-LDV------NAKQGKSGYDANWG-KLSELANDF-ETSPLSVFAYLNQWVR---------RQS-----------
---VET--A---RI----EKIR----------LYAYY--FYPCFDP-Y------------------------V-VYNLKEDK--------L-IVE-KE--SS-LNH-PKRLTELYRKFYR--A-NKR--YNP--KANAVLKPIDIAAETI-LKAESSIFN--G-EALINAVAAEIFKLMDRVH-----N---STAEG-----RWIF-S--------K--------REEE----RQAILEFARYFV--DDVFYESFKGDRARLAGRQINLI-----RDTCEFIYR------LEDDK--ENS--ELEA-SKQSK--
------K-HE---------
'WP_009786738.1'    --MAKKRHKSEPEGQQLSL-FD-----------QPEVINTEIDE-----DDSWLED---E----EFGLNFESY-------DPIF--EPKISELLTLKLLREA---IH--SQNPD--------DA--VMTD--FAEYVL--PNLL-RV-AIGVTAK----G---------GKY-FDE-------LDE-------KRE-------AE---------GKT-------K---VR-RDNA-GDQSLNTHLLNGLFPANLIE--QR------------
----------LEKL--D-TT-IKRVV---GEQERRLVIAGFILHDFEK-----F----------------P-D--APEN-----------------CRQ-LP--------IEEHRQIITDKVQQL-GLD----YFI--NP--DDPNAYQDYIDDLLCVAYN-AQR---RWDTNWNFSVF------GL-NP----QL----RD-RTLV--CLSELTCLADSLASI------IKHPQDA-E-----NTR-----------LKELI-H-SL---SD-GQV----Q-------FTYHSIAENRGVLTNIVNNALMEAH---
TALNTEN--K------T-Y---YEPLLYLPTGVIYLTQ--KNSPQ-I-SAS---EIPQ--RVIDKI-----------------KSFCS--RRL--V-D-SQMGFGR-DG----------KG-MKYADYYEQF-FDTVGLM------------------KVALKATL---------------------RV-LN---------------------P------KK---SSVAK-------------------------------------SR------SD---NLV--------RFQ------------QQNVLPA-------DY---NFIF------ED----DIRI-----DQ------
--IA-------------------E----FG---DVIS--RK--IWG-------EQV-------NKIEK-------ERKQ-NKK----LPELPHLD-------LIHEVAKF----WKLE-----QYSP-QI-------------REIQKINESLKANKLKGNTGGVPYEWYFLAAKYI----Q--N---NP-GI-E---DVRE----HCEQV-IDYINQLIEP-----------IVK----------QY----T--L--P-DG-WDD-------------LREWVMRVV---------MLP---------------------------GKPTTDSL--
SADI--F--LKELNHYQLAKK-SGRGRQL-ICSIS-HSPYSV---TEQMESAVLFTPQV--YTN--KQ-ML---GG--SNA-KRN-------ISSIAGTEMMLRQILMNQT-Q------AVGKRFEDGKYRYLYF-YP---TYYFTP----E-TNKF---LQQ-----AYQNI-AQTRF-----DT------SIR-----NHFI-S------------------------------------------KDLQA---------NFE------RSRY--QSVDSF-------L-IDEALQYKK-------
ELPETDSNHKKDR-------TF---KL---SY---PED--------------------QPLTFYFMAL----------PP------G----R--DP--------TDT------ESW--VMPTWLAFAFPMILDVKTVVSESPIPPFNDGAEFEES----VF--LDSAPQAFRVLT-K--RD-RF--------RLD---YILEGW------EE----AGKK-YSSPLNV-LTAAYAIH-LDV------NAKQGKSGYDANWG-KLSELANDF-ETSPLSVFAYLNQWVR---------RQS-----------
---VET--A---RI----EKIR----------LYAYY--FYPCFDP-Y------------------------V-VYNLKEDK--------L-IVE-KE--SS-LNH-PKTLTELYRQFYR--A-NKR--YNP--KANAVLKPIDIASKTI-LEADPS-FR--G-EALISVVAAEVFKLMDRVH-----A---STAEG-----RWIY-S--------H--------REEE----RQAVLVFAKYFV--EDVFYGSFKGDRARLAGRQINLI-----RDTCEFLYR------LEDDK--ENS--KLEA-
SKQSK--------K-NE---------
'WP_079680991.1'    -------------------------------------------MNNS--ENDDNLFND-------DFGFEGDSD--------RTI--EPSQRELLTLKLLREA---IQ--AENPD--------DV--VMRD--FAEYVL--PNLL-KI-AIGVTAK----G---------GKF-FDE-------IDQ-------KRE-------AE---------GKE-------K---VR-RDNA-ADQSLNTHLLNGLFPANLIE--KR-----------------------LQQL--N-
TT-IQRIV---GEKERRLAIAAFILHDFEK-----F----------------P-D--APEN-----------------CRQ-LS--------LDQHRLIIDQKVKQL-GIN----YFI--NP--NHPEDYREYIDDLLCMAYN-AQR---RYDTNWNFSEF------GL-NP----LL----KD-RVLR--NLSDLTCLADSLASI------VKHPQDA-D-----NTR-----------LQELI-Y-VL---SD-GQL----R-------FTYHSITENRGVLTNVVNNAVMEAY---TSLNTED--C------I-
Y---YQPLLYLPTGVIYLAL--KNAPE-I-NIL---DLPN--RVVTKI-----------------KQLCG--GQL--R-L-RQTGFGR-DG----------KG-MKYAEYYNLF-CTDRDLM------------------HIALDATL---------------------RI-LS---------------------A------NK---KSVAK-------------------------------------SR------SD---NLI--------DFQ------------KKGVLSP-------NY---NFQF------DD----DIRV-----DQ--------IA-------------------E-
--LG---DVVS--RK--IWG-------DRV-------DKINQ-------ARKQ-NKK----LPEITDLD-------ITYQVAEF----WHLT-----EYLP-AI-------------RDIQSINESLKDKGLKGNTGGVPYEWYFLAAKYL----E--R---NP-GV-E---NVRE----ICNNI-IDHIADLIEP-----------IIE----------QY----Q--L--P-DG-WDD-------------LRNWVQQVI---------ILP---------------------------TQNQL-NL--SEQATVF--
LRELTNYNAAKK-SGRGRQL-ICSIS-HSPYTV---KEQMESEVLFTPQV--YTN--KQ-ML---GG--SNA-KRN-------ISSIAGLELMLRQILMNQT-Q------AVGKRFEDGKYRYLYF-YP---TYYFTP----E-TNKF---LQK-----AYNGI-AQTRF-----DT------SLR-----NHFI-S------------------------------------------KDWVA---------NLD------KEHY--QTVDTF-------L-MDEQLDHKK-------
TLPETDPNYKRDR-------TF---KL---SY---PED--------------------QPLTFYFMAL----------PP------G-----K--DP--------TDT------ESW--VMPTWLSFAFPIILDVKTVVSESPIPPFKDGSEFEES----VF--LDSAPQAFRVLV-G--RD-RF--------RLD---YILEGW------DDA---NGIK-HPSPLNS-LTVAYAIH-LDV------NAKQGKGGYDANWG-KLSELAKDF-ETSPLYVFNYLNRWVR---------HQG-----------
--AET--A---RV----EKIK----------LYTYH--FYPCFDP-Y------------------------T-TFNFEKET--------L-IVS-EK--SP-LNH-PKNLTELYRKFYR--A-NKR--YSP--KSNAVLKPIDIAAETI-LKADSTVFQ--G-ETLVMAVAAEVFKLMDRVH-----A---STAEG-----RWVF-S--------N--------REEE----RQAVLNFAQYFV--VDVFEKSFASDRARLAGRQLNLL-----RDTCEFLYR------LEDDK--DNQ--NKPE-
LIAQE--------N-TIS--------
'SKB14935.1'    ------------------------------------------MMNNS--ENDDNLFND-------DFGFEGDSD--------RTI--EPSQRELLTLKLLREA---IQ--AENPD--------DV--VMRD--FAEYVL--PNLL-KI-AIGVTAK----G---------GKF-FDE-------IDQ-------KRE-------AE---------GKE-------K---VR-RDNA-ADQSLNTHLLNGLFPANLIE--KR-----------------------LQQL--
N-TT-IQRIV---GEKERRLAIAAFILHDFEK-----F----------------P-D--APEN-----------------CRQ-LS--------LDQHRLIIDQKVKQL-GIN----YFI--NP--NHPEDYREYIDDLLCMAYN-AQR---RYDTNWNFSEF------GL-NP----LL----KD-RVLR--NLSDLTCLADSLASI------VKHPQDA-D-----NTR-----------LQELI-Y-VL---SD-GQL----R-------FTYHSITENRGVLTNVVNNAVMEAY---TSLNTED--C----
--I-Y---YQPLLYLPTGVIYLAL--KNAPE-I-NIL---DLPN--RVVTKI-----------------KQLCG--GQL--R-L-RQTGFGR-DG----------KG-MKYAEYYNLF-CTDRDLM------------------HIALDATL---------------------RI-LS---------------------A------NK---KSVAK-------------------------------------SR------SD---NLI--------DFQ------------KKGVLSP-------NY---NFQF------DD----DIRV-----DQ--------IA-------------------
E----LG---DVVS--RK--IWG-------DRV-------DKINQ-------ARKQ-NKK----LPEITDLD-------ITYQVAEF----WHLT-----EYLP-AI-------------RDIQSINESLKDKGLKGNTGGVPYEWYFLAAKYL----E--R---NP-GV-E---NVRE----ICNNI-IDHIADLIEP-----------IIE----------QY----Q--L--P-DG-WDD-------------LRNWVQQVI---------ILP---------------------------TQNQL-NL--SEQATVF--
LRELTNYNAAKK-SGRGRQL-ICSIS-HSPYTV---KEQMESEVLFTPQV--YTN--KQ-ML---GG--SNA-KRN-------ISSIAGLELMLRQILMNQT-Q------AVGKRFEDGKYRYLYF-YP---TYYFTP----E-TNKF---LQK-----AYNGI-AQTRF-----DT------SLR-----NHFI-S------------------------------------------KDWVA---------NLD------KEHY--QTVDTF-------L-MDEQLDHKK-------
TLPETDPNYKRDR-------TF---KL---SY---PED--------------------QPLTFYFMAL----------PP------G-----K--DP--------TDT------ESW--VMPTWLSFAFPIILDVKTVVSESPIPPFKDGSEFEES----VF--LDSAPQAFRVLV-G--RD-RF--------RLD---YILEGW------DDA---NGIK-HPSPLNS-LTVAYAIH-LDV------NAKQGKGGYDANWG-KLSELAKDF-ETSPLYVFNYLNRWVR---------HQG-----------
--AET--A---RV----EKIK----------LYTYH--FYPCFDP-Y------------------------T-TFNFEKET--------L-IVS-EK--SP-LNH-PKNLTELYRKFYR--A-NKR--YSP--KSNAVLKPIDIAAETI-LKADSTVFQ--G-ETLVMAVAAEVFKLMDRVH-----A---STAEG-----RWVF-S--------N--------REEE----RQAVLNFAQYFV--VDVFEKSFASDRARLAGRQLNLL-----RDTCEFLYR------LEDDK--DNQ--NKPE-
LIAQE--------N-TIS--------
'AFY52833.1__2_'    ---MTKKINDLEEHKQLSL-FD-----------IE---DNSQSE---DSDYDWLDD---E----DVDFGFETS-------DRTV--ETAQ-ELLTYKLLREA---IE--AQNPD--------DE--VMAD--FAKYVL--PNLL-QI-AIGVTAK----G---------GKY-FDK-------LDR-------EKA-------AA---------GKA-------K---VR-RDNA-ADQSLNTHLLNGLFPANLIE--RR---------------
---------LVKL--N-TT-VMRVV---GEVERRLLVAGFILHDFEK-----F----------------P-D--VPDD-----------------CRK-LA--------IEEHRRIIDQKVRQL-GLD----KFI--NP--SQSEAYQEYLDDLLCMAYN-AQR---RWDTNWNFSEF------GL-NP----QL----RD-RTLR--CLSELTCLADSLASI------IKHPQDA-Q-----SYK-----------FKELL-S-AL---SD-GEL----Q-------FTYHSIAENRGVLTNVVNNALIAAH---
TSLNTEN--N------I-C---YEPLLYLPTGVIYLAS--RDAPA-I-SQE---DLPD--RVISTI-----------------KSLCA--SQL--R-L-RQTGFGR-DG----------KG-MKYADYYNLF-FDDVALM------------------QVGLDATL---------------------RI-LH---------------------P------KK---DSVAK-------------------------------------SR------SD---NLV--------KFQ------------EKDVLPS-------DF---DLKF------SD----DIRI-----DQ--------
IA-------------------E----FG---DLIA--RK--IWG-------EKV-------KQIES-------LRKK-DKK----LPELPQLD-------LIHKVAEF----WNLT-----EYLP-EV-------------KAIQRINESLKENKLKGNTGGVPYEWYYLAAKYL----E--H---NP-GI-E---DIRE----VCQPV-ITFVQSLIEP-----------IVS----------QY----Q--I--A-DG-WDD-------------LRLWVQRVI---------MLP---------------------------GSNREAN---
ENQVETF--LDELSNYNTAKK-SGRGKQL-ICSIS-HSAYTV---TEQMESAVLFTPQV--YTN--KQ-ML---GG--SNA-KRN-------ISSIAGVEMMLRQILMNQT-Q------AVGKRFEDGKYRYLYF-YP---TYYFTP----E-TNKF---LQI-----AYSGI-AQTRF-----DT------SIR-----NHFI-P------------------------------------------KNLQA---------NLG------KDKY--QSVDSF-------L-IDEDLQSQK-------
NLPEDDPNYKKDR-------TF---KL---SY---PED--------------------QPLTFYFMAL----------PP------G----R--DS--------SDT------ESW--IMPSWLAFAFPMILDVKTVVSESPIPPFNDGAEFEES----VF--LDSAPHAFRALV-K--QD-KF--------RLD---YILEGW------QD----NNKT-YSAPLNV-LTAAYAIH-LDV------NAKQGKSGYDANWG-KFTELAKDF-ETSPLYVFSYLNRWVR---------KQG-----------
---VET--A---RI----EKIK----------LYAYD--FYPCFDA-Y------------------------V-KYDNIQEK--------L-IVE-SE--SS-LCH-TQKLTELYRRFYR--A-NKK--YNP--KSNAVLKPIDIAAETI-LKAELSVHT--G-ECLVLVVAAEVFKLMDRVH-----A---STAEG-----RWVF-N--------HK------EREEE----RQAVLEFARYFV--EDIFEKSFAGDRARLAGRQINLI-----RDTCEFLYR------LENDK--ENQ--LQGN-
DSKKE--------N-DSETLISS---
'WP_088889711.1'    -------------------------------------------MTSN---NDEIDWLED-------DLGLNEGST-------DRTV--ESPKRELLTLKLLREA---IR--SQNPD--------DA--VMQD--FAEYVL--PNLL-KM-AIGVTAK----G---------GKF-FDE-------LDQ-------QRE-------AA---------GKD-------R---VR-RDNA-ADQSLNTHLLNGLFPANLIE--KR-----------------------LEKL--
N-TT-VQRVV---REQERRLVIAAFILHDFEK-----F----------------P-D--APEN-----------------CRK-LS--------LEEHRQIIDEKVHQL-GLD----RFI--DP--NHPETYKAYLDDLLCMAYN-AQR---RWDTNWNFSEY------GL-NP----SL----KD-RVLR--SLSDLTCLADSLASI------VKHPQDA-E-----HPR-----------LKELI-H-SL---SD-GQL----K-------LTYHSIAENRGVLTNVVNNALMEAH---TSLNTSE--L-
-----I-Y---YEPLLYLPTGVIYLAD--RNAPQ-V-DPK---ELPD--RVVASI-----------------KSLCA--GQL--R-L-RQTGFGR-DG----------KG-MKYAEYYNLF-FDAPGLM------------------QVALDATL---------------------RI-LN---------------------P------NK---GSVAK-------------------------------------SR------SQ---NLV--------KFQ------------QQNVLPA-------NY---SFEF------ED----DIRI-----DQ--------LA-----------
--------E----FG---DLVS--RK--IWG-------ERFAQLAEARKRYEA-------DRKK-NKQ----LPELKPLD--LLPDQLVHKVAEF----WGLG-----EHSP-QI-------------EKIQRINENLKANKLTGNTGGVPYEWYFLAAKYL----E--R---HP-GI-E---NLRD----VCSQV-VAYLANLIQP-----------LVE----------QY----Q--L--P-DG-WDD-------------LRLWVERIV---------MLP---------------------------GNIQNQMA--
TEQVDTF--LKELAGYSAAKK-PGRGRQL-ICSIS-NSAYTV---TEQMESAVLFTPQV--YTN--KQ-HL---GG--SNA-KRN-------ISSIAGIEMMLRQILMNQT-Q------AVGKRFEDGKYRYLYF-YP---TYYFTP----E-TNRF---LQK-----AYSGI-AQTRF-----DT------SVR-----NHFI-T------------------------------------------KDMRA---------DLG------RDRY--QTVDAF-------L-IDEDLERKK-------
QLPENDPQYRRDR-------TF---KL---SY---PDD--------------------QPLTFYFMAL----------PP------G----R--DP--------TDT------ESW--VMPAWLAFAFPMILDVKTVVSESPIPPFTDGAEFEES----VF--LDSAPQAFRVLT-G--RD-RF--------RLD---YILDGW------NAP---DGTK-QPAPLNV-LTAAYAIH-FDV------NAKQGKGGYDANWG-RLTELAKDF-ETSSLYVFSYLNRWVR---------NQG------
-------VET--A---RI----EKIR----------LYAYH--FYPCFDP-Y------------------------T-QFDFNSEV--------L-IVG-EE--SP-LNH-PQKLTELYRRFYR--A-GKR--YNP--KSNAVLKPIDVAAETI-LKAESTVFQ--G-ETLVAAVAAEVSKLMDRVH-----A---STAEG-----RWVI-S--------K--------REEE----RQIILDFARYFV--KDVFEVAFAGDRARLAGRQLNLI-----RDTCEFLYR------LEDDK--ENQ--ANGE-
KPAEK--------D-VES--------
'WP_017652463.1'    ---MAKKSKKSDEYQQLSI-LE-----------AETENDST-SE---LSDDDWMSG-------DDSDFQEIS-------NLKV--ETQLAELLTLKLLQDA---IK--SENDD--------DL--IMQE--FSDYVL--PNLL-RV-AVGVTAK----G---------GKF-FDY-------LDK-------KKE-------KEIENFHQNNLGKSHKIKEPKK---LD-RRNA-GDQSLNTHLLNGLLPANLIE--
RC-----------------------LNQL--D-TT-AKRVI---QEQERRLLIAGFILHDFEKFDYRLF----------------P-K--MPDNYKAIGQDK------EQDIRK-LS--------VDAHREIIDVIVCEL-GLD----KFV--SP--DEPTAWQEYRDDLLFIAYN-AQR---RNDTNLNVSEH------GL-QP----II----DD-YILI--CLADLTCLADLLSSI------IKHPQDA-E-----HQT-----------LNGIL-H-SL---SD-GQL----K-------
FTYHSIAENRGVLTNVVNNALMEAH---TSLNTED--K------T-H---YKRLLYLPTGVIYLAR--KDAPK-I-SRQ---GLDD--RVIDTI-----------------KKLCA--NQL--Q-L-RKVGFSR-GN----------IG-MKYADYYNLF-LDIKGLM------------------QFTVDA--------------------------V-PS---------------------K------VK---TSKAA-------------------------------------DK------AT---NLA--------QFQ------------RKGVLAK-------
NI---EIDF------AE----DIRI-----DH--------LA-------------------E----LG---ALIT--GK--FWK-------EYL-------DRVKA-------LIKN-NKK----LPPIPTIN-------LTEIAIDF----LDLS-----EYKS-EI-------------IAFQQLKD----------TGGIAYDWYYIATQYL----K--R---NP-GQ-E---DVRE----ICQSL-VNRWIEVIEP-----------IIN----------QY----K--L--P-DE-WKD-------------LRDWCDRVI---------MLP---------------------
---------TETQQKSA--TDQAEIF--FKELANYNAAKK-AGRGKQL-ICSIS-HSAYTV---TEQMESAVLFTPQV--YTN--KQ-IL---GG--SNA-KRN-------ISSIAGVEMMLRQILMNQT-Q------AVGKRFEDGKYRYLYF-YP---TYYFTP----E-TNKF---LQK-----AYNGI-AQTRF-----DT------SVR-----NHFI-S--------------------------------------------------KDLQA---------NLK------RENY--QSVDSF-------L-
IDENLQRDK------------------DR-------TF---KL---SY---PED--------------------QPLTFYFMAL----------PP------G----R--DS--------TDT------ESW--VMPTWLAFAFPMILDVKTVVSESPIPPFNDGAEFEES----VF--LDSAPHAFRALV-R--RD-RF--------RLD---YILEGW------NE----NSIQ-YPAPLNV-LTAAYAIH-LDV------NARQGKSGYDANWG-RFTELAKDF-ETSPLYVFSYLNRWVR---------
NQG-------------IET--A---RI----EKIR----------LYAYH--FYPCFDP-Y------------------------V-KYDNNMET--------L-NVK-EE--SS-LNH-PKKLTDLYRRFYR--A-NKR--YNP--KSNAVLKPIDIAAETI-LKAESSVFQ--G-DTLVAAVAAEVFKLMDRVH-----S---STAEG-----RWII-S--------K--------REEE----RQAVLDFARYFV--TEVFEKSFAGDRARLAGRQLNLI-----RDTCEFLYR------LEDDK--
ENA--ARDD-KSTES--------V-DNENQ------
'WP_089129578.1'    -----------------------------------------------MSNDDWLSG-------DFGFDGDSS-------DRTI--ETE-GELLTLKLLREA---IQ--AQNPD--------DK--VMAD--FAEYVL--PNLL-RI-AIGVTAK----G---------GKF-FDE-------IDQ-------QRE-------AE---------GKT-------K---VR-RDNA-ADQSLNTHLLNGLFPANLIE--KR-----------------------LEKL--N-TT-
VQRVV---KERERRLVIAGFILHDFEK-----F----------------P-D--VPEN-----------------CRK-LP--------LAEHRKIIDKKVHQL-GLD----NFI--NP--ENPEGYREYLDDLLCMAYN-AQR---RWDTNWNFSEF------GL-NP----VL----KD-RTLR--SLSDLTCLADSLASI------VKHPQDA-E-----NSR-----------LQEVI-H-NL---SD-GQL----K-------FTYHSIAENRGVLTNVVNNALMEAY---TSLNTED--Q------
T-Y---YEILLYLPTGVIYLAR--KDAPK-I-SRE---GLDD--RVIETI-----------------KQLCA--NQL--Q-L-RKVGFSR-GN----------IG-MKYADYYNLF-LDIKGLM------------------QFTVDA--------------------------V-PS---------------------K------VK---TSKAA-------------------------------------DK------AT---NLA--------QFQ------------RKGVLAK-------NI---EIDF------AE----DIRI-----DH--------LA-------------------E----
LG---ALIT--GK--FWK-------EYL-------DRVKA-------LIKN-NKE----LPSIPTIN-------LTEIAIDF----LNLS-----EYKS-EV-------------IALQQLKD----------TGGIAYDWYYIATQYL----K--R---NP-GQ-E---DVRE----ICQSL-VNRWIEVIEP-----------IIN----------QY----K--L--P-DE-WKD-------------LRDWCDRVI---------MLP---------------------------NETRQKSP--TDQAEIF--FKELANYNAAKK-
PGRGKQL-ICSIS-HSAYTV---TEQMESAVLFTPQV--YTN--KQ-ML---RG--ANA-KRN-------ISSIAGMEMMLRQILMNQT-Q------AVGKRFEDGKYRYLYF-YP---TYYFTP----E-TNKF---LQK-----AYNGI-AQTRF-----DT------SVR-----NHFI-S------------------------------------------KDLQA---------DLG------RDRY--QSVDSF-------L-IDENLQRDK------------------DR-------TF---KL---SY---
PED--------------------QPLTFYFMAL----------PP------G----R--DS--------TDT------ESW--VMPTWLAFAFPMILDVKTVVSESPIPPFNDGAEFEES----VF--LDSAPHAFRALV-R--RD-RF--------RLD---YILEGW------NE----NGIQ-YPAPLNV-LTAAYAIH-LDV------NARQGKSGYDANWG-RFTELAKDF-ETSPLYVFSYLNRWVR---------NQG-------------SET--A---RI----EKIR----------LYAYH--
FYPCFDP-Y------------------------V-KYETNMEQ--------L-IVG-EA--SS-LNH-PKRLTELYRRFYK--A-NKM--FNP--KANAVLKPIDIAAETI-LKAELSVFH--G-ETLVMAVAAEVFKLMERVH-----S---STAEG-----RWVF-K--------RN--------QREEE----RQAVLDFARYFV--EEVFEKSFAGDRARLAGRQLNLI-----RDTCEFLYR------LEDDK--ENA--GKNE-KSAEI---------D-NENE--------
'WP_086689580.1__2_'  -----------------------------------------------MSNDDWLSG-------DFGLDGDSS-------DRTI--ETE-GELLTLKLLREA---IQ--AQNPD--------DK--VMAD--FAEYVL--PNLL-RI-AIGVTAK----G---------GKF-FDE-------IDQ-------QRE-------AE---------GKT-------K---VR-RDNA-ADQSLNTHLLNGLFPANLIE--KR-----------------------LEKL--N-TT-
VQRVV---KERERRLVIAGFILHDFEK-----F----------------P-D--VPEN-----------------CRK-LP--------LAEHREIIDKKVHQL-GLN----NFI--NP--ENPEAYREYLDDLLCMAYN-AQR---RWDTNWNFSEF------GL-NP----VL----KD-RTLR--SLSDLTCLADSLASI------VKHPQDA-E-----NSR-----------LQEII-H-NL---SD-GQL----K-------FTYHSIAENRGVLTNVVNNALMEAH---TSLNTED--K------T-
Y---YEILLYLPTGVIYLAR--KDAPK-I-SRE---GLGD--RVIDTI-----------------KQLCA--NQL--Q-L-RKVGFSR-GN----------IG-MKYADYYNLF-LDIRGLM------------------QFTVDA--------------------------I-PS---------------------K------VK---TSKAA-------------------------------------DK------AT---NLA--------QFQ------------RKGVLAK-------NI---EIDF------AE----DIRI-----DH--------LA-------------------E----
LG---ALIT--GK--FWK-------EYL-------DRVKA-------LIKN-NKE----LPPIPTIN-------LTEIAVDF----LDLS-----EYKS-EI-------------TALQQLKD----------TGGIAYDWYYIATQYL----K--H---NP-GQ-E---DVRE----ICQSL-VNQWIKVIEP-----------IIN----------QY----K--L--P-DE-WKD-------------LRDWCDRVI---------MLP---------------------------TETRQKSA--TDQAEIF--FKELANYNAAKK-
LGRGKQL-ICSIS-HSAYTV---TEQMESAVLFTPQV--YTN--KQ-ML---RG--ANA-KRN-------ISSIAGMEMMLRQILMNQT-Q------AVGKRFEDGKYRYLYF-YP---TYYFTP----E-TNKF---LQK-----AYNGI-AQTRF-----DT------SVR-----NHFI-S------------------------------------------KDLQA---------DLG------RDRY--QSVDSF-------L-IDENLQRDK------------------DR-------TF---KL---SY---
PED--------------------QPLTFYFMAL----------PP------G----R--DS--------TDT------ESW--VMPTWLAFAFPMILDVKTVVSESPIPPFNDGAEFEES----VF--LDSAPHAFRALV-K--RD-RF--------RLD---YILEGW------NE----NGIQ-YPAPLNV-LTAAYAIH-LDV------NARQSKTGYDANWG-RFTELAKDF-ETSPLYVFSYLNRWVR---------NQG-------------SET--A---RI----EKIR----------LYAYH--
FYPCFDP-Y------------------------V-KYDTNMEQ--------L-IVG-EA--SS-LNH-PKKLTDLYRRFYR--A-NKR--YNP--KSNAVLKPVDIAAETI-LKAESSVFQ--G-ETLVAAVAAEVFKLMDRVH-----S---STAEG-----RWIM-S--------K--------REEE----RQAVLDFAKYFV--VEVFDNSFAGDRARLAGRQLNLI-----RDTCEFLYR------LEDDK--ENA--GRNE-NISVI---------D-DVNEPE-----
'WP_109006821.1'    -----------------------------------------------MSNDDWLSG-------DFGFDGDSS-------NRTI--ESE-GELLTLKLLREA---IE--AQNPD--------DK--VMAD--FGEYVL--PNLL-RI-AIGVTAK----G---------GKF-FDE-------IDQ-------QRE-------AE---------GKT-------K---VR-RDNA-ADQSLNTHLLNGLFPANLIE--KR-----------------------LEKL--N-TT-
VQRVV---KERERRLVIAGFILHDFEK-----F----------------P-D--VPEN-----------------CRK-LP--------LAEHREIIDKKVHQL-GLD----NFI--NP--ENPEAYPEYLDDLLCMAYN-AQR---RWDTNWNFSEF------GL-NP----LL----KD-RTLR--SLSDLTCLADSLASI------VKHPQDA-E-----NSR-----------LQEVI-H-NL---SD-GQL----K-------FTYHSIAENRGVLTNVVNNALMEAH---TSLNTED--Q------
T-Y---YEILLYLPTGVIYLAR--KDAPK-I-SRE---GLGD--RVIDTI-----------------KQLCA--NQL--Q-L-RKVGFSR-GN----------IG-MKYADYYNLF-LDIKGLM------------------QFTVDA--------------------------V-PS---------------------K------VK---TSKAA-------------------------------------DK------AT---NLA--------QFQ------------RKGILAK-------NI---EIDF------AE----DIRI-----DH--------LA-------------------E----
LG---ALIT--GK--FWK-------EYL-------DRVKA-------LIKN-NKE----LPSIPTIN-------LTEIAIDF----LNLS-----EYKS-EI-------------IALQQLKD----------TGGIAYDWYYIATQYL----K--C---NP-GQ-E---DVRE----ICQSL-VNRWIEVIEP-----------IIN----------QY----K--L--P-DE-WKD-------------LRDWCDRVI---------MLP---------------------------NETRQKSP--TDQAEIF--FKELANYNAAKK-

AGRGKQL-ICSIS-HSAYTV---TEQMESAVLFTPQV--YTN--KQ-ML---GG--SNA-KRN-------ISSIAGVEMMLRQILMNQT-Q------AVGKRFEDGKYRYLYF-YP---TYYFTP----E-TNKF---LQK-----AYNGI-AQTRF-----DT------SVR-----NHFI-S-------------------------------------------------KDLQA---------DLG------RDRY--QSVDSF-------L-IDENLQRDK-----------------DR-------TF---KL---SY---
PED--------------------QPLTFYFMAL----------PP------G----R--DS--------TDT-------ESW--VMPTWLAFAFPMILDVKTVVSESPIPPFNDGAEFEES----VF--LDSAPHAFRALV-R--RD-RF--------RLD---YILEGW------NE----NGIQ-YPAPLNV-LTAAYTIH-LDV------NARQGKSGYDANWG-RFTELAKDF-ETSPLYVFSYLNRWVR---------NQG------------SET--A---RI----EKIR---------LYAYH--
FYPCFDP-Y------------------------V-KYDTNMEQ--------L-IVG-EA--SS-LNH-PKKLTDLYRRFYR--A-NKR--YNP--KSNAVLKPVDIAAETI-LKAESSVFQ--G-ETLVAAVAAEVFKLMDRVH-----A---STAEG-----HWIM-S---------K---------REEE----RQAVLDFARYFV---TEVFEKSFAGDRARLTGRQVNLI-----RHTCEFLYR------LEDDK--ENS--ARNE-QSTES--------N-DDGNQ------
'PSB01849.1__2_' ---MAKKTKNSKESKQLSL-FD-----------SALDSNSEITD---QADDSWLEG-------DFGFDGDSD--------RTI--ETKS-ELLTLKLLREA---IQ--SQNPD---------DQ--VMTD--FAEYVL--PNLL-SI-AIGVTAK----G---------GKF-FDE-------LDR-------KNE-------AE---------GKT-------K---VR-RDNA-ADQSLNTHLLNGLFPANLIE--RR-------------
----------LHKL--D-TS-VQRVV---GERERRLVIAGFILHDFEK-----F----------------P-D--VPNN-----------------CRK-LA--------LEEHRQIIDKKVRQL-GLD----TFI--DS--ENPEAYKEYLDDLLCMAYN-AQR---RWDTNWNFSEF------GL-NP----LL----KD-RTLR--SLSDLTCLADSLASI------IKHPQDA-D-----NPR-----------LNEII-H-SL---SD-GKL----K-------FTYHRISENRGVLTNVVNNALIQVH---
TQINTDE--N------I-Y---YEPLLYLPTGVIYLAR--RDAPA-I-SLD---DVPN--QVIENI-----------------KSLCS--GQL--K-L-RQMGFGR-DG----------KG-MKYAEYYNLF-FDEIGLM----------------KVALDATL-----------------------RV-LN---------------------A-------NK----NSVGK---------------------------------------------SR-------SD---NLV---------KFQ------------QQNVLSA-------DY---DFNF-------TN-----DIRI-----DQ-------
-IA---------------------E----FG---DLIC--RK--IWG--------ESV---------NRIEA--------ACKK-DKN----LPKLPAID---------LTQKVAGF-----WEIT------EYLP-QI--------------REIQRINESLKENKLKGNTGGVPYEWY------------------------------------------------------------------------------------------------------------------------------------------------------------------------------------------------
-----------------------------------------------------------------------------------------------------------------------------------------------------------------------------------------------------------------------------------------------------------------------------------------------------------------------
'AFZ00743.1__2_' ---MAKKTKKSEESKQLSL-FE-----------SGLDLNSEITD---EADDNWLEG-------DFGFDGDSD--------RTI--ETKS-ELLTLKLLREA---IK--AQNPD--------DQ--VMAD--FGEYVL--PNLL-SI-AIGVTAK----G---------GKF-FDE-------IDE-------KRE-------AE---------GKA-------K---VR-RDNA-ADQSLNTHLLNGLFPANLIE--RR-------------
---------LHKL--D-TT-VQRVV---RERERRLVIAGFILHDFEK-----F----------------P-D--VPND-----------------CRK-LP--------LEEHRQIIDEKVRQL-GLD----KFI--DS--ENTEAYKEYIDDLLCMAYN-AQR---RWDTNWNFSEF------GL-NP----LL----KD-RTLR--SLSDLTCLADSLASI------IKHPQDA-E-----NPR-----------LNEII-H-SL---SD-GKL----K-------FTYHRISENRGVLTNVVNNALIQVH---
TKLNTDE--N------I-Y---YEPLLYLPTGVIYLAR--RDAPE-I-SLD---DVPN--QVIENI-----------------KSLCS--GQL--K-L-RQTGFGR-DG----------KG-MKYAEYYNLF-FDEIGLM----------------KVALDATL-----------------------RI-LN-----------------------P------NK---DSVAK---------------------------------------------SR-------SD---NLV---------KFQ------------QQKVLSV-------DY---DFKF------TD-----DIRI-----DQ-------
IA----------------------E----FG---DLIC--RK--IWD--------ESA--------NRIEV--------ACKK-DKN----LPKLTQLD---------LTYKVAEF-----WELA------EYLP-QI--------------REIQRINESLKENKLKGNTGGVPYEWYYLAAKFL----E--K---HP-GI-E---NVRE----SCQQV-IEYINNLISP-----------IIS----------QY----E--L--P-DG-WND-------------LRLWVKQVV----------MLP-----------------------GNQETQTP------
DNF--LNELANYNAAKK-SGRGRQL-ICSIS-HSAYTV---TEQMESAVLFTPQV--YTN--KQ-ML---NG--SNA-KRN-------ISSIAGLEMMLRQILMNQT-Q------AVGKRFEDGKYRYLYF-YP---TYYFTP----E-TNKF---LQK-----AYTGI-SQTRF-----DT------RIR-----NHFI-S-------------------------------------------------KDLQA---------NFD------KDTY--QNVDAF-------L-IDENLDSDK---------------------DR---
----TF---KL---SY---PDD--------------------QPLTFYFMAM----------PP------G----R--DS--------SDT-------ESW--VMPTWLAFAFPMILDVKTVVSESPIPPFNDGAEFEES----VF--LDSAPHAFRTLV-G--RD-RF--------RLD---YILEGW------TE----NNKS-YPAPLNV-LTAAYAIH-IDV------NAKSGKSGYDANWG-RFTELAKDF-ETTPLYVFAYLNRWVR---------SQG------------VET--A---RI----
EKIR---------LYAHK--FYPCFDP-Y------------------------A-QYDFESKE--------W-KLM-PE--SQ-LNH-PKKLTDLYRKFYR--A-NKR--YNP--KSNAVLKPIDIAAETI-LKAEVSVFS--G-ETLVAAVAAEVFKLMDRVH-----A---NTADG-----RWVM-S---------K---------REEE----RQAVLDFARYFV---MDVFENSFSSDRARLAGRQLNLI-----KDTCEFLYR------LEDDK--ENA--NKDV-KEEEN--------D-
KNED-------
'PAX59910.1__2_' ---MAKKTKDSSESKQLSL-FE-----------SGLDLNSEITD---EADDNCLEG-------NFGFDGDSD--------RTI--ESKS-ELLTLKLLREA---VR--SQNPD--------DQ--VMAD--FCEYVL--PKLL-SI-AIGVTAK----G---------GKF-FDE-------IDE-------KRK-------AK---------GET-------K---VR-RDNA-ADQSLNTHLLNGLFPANLIE--RR-------------
---------LHKL--D-TT-VQRLV---KERERRLVIAGFILHDFEK-----F----------------P-D--VPND-----------------CRK-LP--------LKEHQQIIDEKVRQL-GLD----KFI--DS--ENLEAYKEYIDDLLCMAYN-AQR---RWDTNWNFSEF------GL-NP----IL----KD-RTLR--SLSDLTCLADSLASI------IKHPQDA-E-----NPR-----------LNEII-H-SL---SD-GKL----K-------FTYHRISENRGVLTNVVNNALIQVH---
TKLNTDE--N------I-Y---YEPLLYLPTGVIYLAQ--RNAPA-I-SLE---DVPN--QVIENI-----------------KSLCS--RQL--E-L-RQTGFGR-DG----------KG-MKYAEYYNLF-FDEIGLM----------------KVGLDATL-----------------------RI-LN-----------------------P------NK---GSVAK---------------------------------------------SR-------SD---NLV---------KFQ------------QQNVLSA-------DY---DFKF------AD-----DIRI-----DQ-------
IA----------------------E----FG---DLIS--RK--ILG--------ESV--------NRIEA--------VCKK-DKN----LPKLTQFD---------LTHKVAEF-----WNLT------EYLP-QI--------------REIQRINESLKENKLKGNTGGVPYEWYYLAAKYL----E--K---HP-GI-E---NVRE----SCQKV-IGYINNLISP-----------IIS----------QY----E--L--A-DG-WND-------------LRLWVKQVV----------MLP-----------------------GNQEMQTA------
DNF--LNEFNNYNAAKK-SGRGRQL-ICSIS-HSAYSV---TEQMESAVLFTPQV--YTN--KQ-ML---NG--SNA-KRN-------ISSIAGLEMMLRQILMNQT-Q------AVGKRFEDGKYRYLYF-YP---TYYFTP----E-TNKF---LQK-----AYTGI-SQTRF-----DT------SIR-----NHFI-S-------------------------------------------------KDLQA---------DFD------KVKY--QNVDAF-------L-IDENLDSEK---------------------DR---
----TF---KL---SY---PDD--------------------QPLTFYFMAM----------PP------G----R--DS--------SDT-------ESW--IMPTWLAFAFPMILDVKTVVSESPIPPFNDGAEFEES----VF--LDSAPHAFRTLV-G--RD-RF--------RLD---YILEGW------KE----NNKS-YPAPLNV-LTAAYAIH-LDV------NAKSGKSGYDANWG-RFTELAKDF-ETTPLYVFAYLNRWVR---------NQG------------VET--A---RI----
EKIR---------LYAYQ--FYPCFDP-Y------------------------A-QYDFESKE--------W-KLM-AE--SQ-LNH-PKKLTDLYRKFYR--A-NKR--YNP--KSNAVLKPIDIAAETI-LKAETSVFA--G-ETLVAAVAAEVFKLMDRVH-----A---NTADG-----RWIM-N---------K---------REEE----RQAVLDFARYFV---MEVFEKSFSCDRARLAGRQLNLI-----KDTCEFLYR------LEDDK--ENA--NKNI-QENEQ--------E-
KAEA-------
'AFZ57018.1__2_' ---MAKKSKDTNKHQQLSL-------------LEIEK--NSN-SA---TSDDDWLSG-------DFGFDGDND--------RTI--ETK-GQLLTLKLLREA---IQ--AENPH---------DQ--VMAD--FSEYVL--PNLL-KI-AIGVTAK-----G---------GKF-FDE-------IDE-------KRE-------AE---------GKT-------K---VR-RDNA-ADQSLNTHLLNGLFPANLIE--QR-------------
-------LEKL--D-TT-VKRLV---KDKERRLVIAGFILHDFEK-----F----------------P-D--APED-----------------CRK-LP--------LDEHRKIIDEKIKQL-GLD----KLI--NP--HNPEDYQSYIDDLLCMAYN-AQR---RWDTNWNFSEF------GL-NP----VL----KD-RTIR--SLSDLTCLADSLASI------IKHPQDA-E-----NTR-----------LNEIL-H-SL---SD-GKL----K-------FTYHHIAENRGVLTNVVNNAVIEAH---
TNLNAAE-NT--------Y---YQPLLYLPTGVIYLAL--RDAPR-I-SPE---ILPD--LVVNKI-----------------KSLCA--GQL--K-L-RQTGFGR-DG----------KG-MKYAEYYNLF-FESPGLM----------------QVALDATL-----------------------RI-LN-----------------------T------GK---TSVGK---------------------------------------------SR-------SE---NLV---------KFQ------------KQNVLSA-------DY---DFKF------DD-----DIRI-----DQ-------
-IA---------------------E----FG---DLIT--RK--IWE--------ETV--------DRIET--------ARKK-DKK----LPEVPNFD---------LTHKVAEF-----WNLT------EYLP-QV--------------REIQRINESLKEQKLKGNTGGVPYEWYYLAAKYL----E--K---HP-GI-E---DVRE----NCQQV-IYYLTDLISP-----------ILA----------QY----Q--L--P-DG-WDD-------------LRLWVSRVV----------MLP-----------------------NTNKQTSL--
TTEVDNF--INELSNYNAAKK-PGRGKQL-ICSIS-HSAYTV---TEQMESAVLFTPQV--YTN--KQ-ML---GG--SNA-KRN-------ISSIAGVEMMLRQILMNQT-Q------AVGKRFEDGKYRYLYF-YP---TYYFTP----E-TNKF---LQK-----AYMNI-AQTRF-----DT------SIR-----NHFI-S-------------------------------------------------KDLQV---------DLG------KENY--QSVDSF-------L-IDENLDGEK------------------
DR-------TF---KL---SY---PED--------------------QPLTFYFMAL----------PP------G----R--DS--------SDT-------ESW--VMPSWLAFAFPMILDVKTVVSESPIPPFNDGAEFEET----VF--LDSAPHAFRTLV-K--RD-RF--------RLD---YILEGW------EE----YGIQ-YPAPLNV-LTAAYAIH-LDV------NARQGKTGYDANWG-KFTELAKDF-ETSPLHVFAYLNKWVR---------NQG------------IET--A---RI----
EKIK---------LYTYH--FYPCFDP-Y------------------------T-KFDSTKEK--------L-IVG-EV--ST-LNH-PKKLTELYRRFYR--A-NKR--FNP--KANAVLKPIDIAAETI-LKADSSVFD--G-ETLISAVAAEVFKLMDRVH-----S---STAEG-----RWIM-S---------K---------REEE----RQAVLEFANYFV---VEVFEGCFKSDRARLAGRQLNLI-----RDTCEFLYR------LEDDK--ENA--EKSK-SEIDE--------N-E-------
--
'AFW97301.1__2_' ---MPKKSKDANEHQQLSL-------------LETEI--YSN-DE---ISDDVFSSS-------DFGFDSDSD--------RTI--ETK-GQLLTLKLLKEA---IQ--SQNPH---------DQ--VMAD--FSEYVL--PNLL-KI-AIGVTAK-----G---------GKF-FDE-------IDE-------KRE-------AE---------GKT-------K---VR-RDNA-ADQSLNTHLLNGLFPANLIE--QR-------------
----LEQL--D-TT-VKRVV---KDKERRLVIAGFILHDFEK-----F----------------P-D--APEE-----------------CRK-LP--------LDEHRKIIDEKVKQL-GLD----KFI--NP--DNPELYQEYIDDLLCMAYN-AQR---RWDTNWNFSEF------GL-NP----VL----KD-RTIR--SLCDLTCLADSLASI------IKHPQDA-E-----NTR-----------LNEIL-H-SL---SD-GKL----K-------FTYHHIAENRGVLTNVVNNAVIEAH---
TNLNTAE-NT--------Y---YQPLLYLPTGVIYLAL--RDAPP-I-SLE---ILPD--MVVNKI-----------------KSLCS--GQL--K-L-RQTGFGR-DG----------KG-MKYAEYYNLF-FESPALM----------------QVALDATL-----------------------RI-LN-----------------------T------NK---PSIAK---------------------------------------------SR-------SE---NLV---------KFQ------------KQNVLSS-------DY---DFKF------DD-----SIRI-----DQ-------
IA----------------------E----FG---DLIT--RK--IWD--------EIV--------NKIET--------ERKK-DKK----LPEVPNFD---------LTHKVAEF-----WNLT------EYLP-QV--------------REIQRINESLKEQKLKGNTGGVPYEWYYLAAKYL----E--K---HP-GI-E---DVRE----NCQQV-IYYLTDLISP-----------ILA----------QY----Q--L--P-DG-WDD-------------LRLWVSRVV----------MLP-----------------------HTNKLTSV--
TTEIDNF--INELTNYNAAKK-SGRGKQL-ICSIS-HSAYTV---TEQMESAVLFTPQV--YTN--KQ-ML---GG--SNA-KRN-------ISSIAGVEMMLRQILMNQT-Q------AVGKRFEDGKYRYLYF-YP---TYYFTP----E-TNKF---LQK-----AYMNI-AQTRF-----DT------SIR-----NHFI-S-------------------------------------------------KDLQV---------DLT------KENY--QCVDSF-------L-IDENLDSEK------------------
DR-------TF---KL---SY---PED--------------------QPLTFYFMAL----------PP------G----R--DS--------TDT-------ESW--VMPSWLAFAFPMILDVKTVVSESPIPPFNDGAEFEET----VF--LDSAPHAFRTLV-K--RD-RF--------RLD---YILEGW------EE----SGIE-YPAPLNV-LTAAYAIH-LDV------NAKQGKTGYDANWG-KFTELSKDF-ETSPLHVFSYLNKWVR---------NQK------------LET--A---RI----
EKIK---------LYTYH--FYPCFDP-Y------------------------A-KFDSTQEK--------L-IVE-EK--ST-LNH-PKNLTELYRRFYR--A-NKR--FNP--KANAVLKPIDIAADII-LKADLCMFG--G-EVLTSTVAAEVFKLMDRVH-----S---STAEG-----RWIM-S---------K---------REEE----RQAVLEFARYFV---EDVFEGCFKSDRARLAGRQLNLI-----RDTCEFLYR------LEDDK--ENS--QNSA-NTDNE--------S--------
----
'WP_028084539.1' ---MAKKSKKSDDYQQLSL-------------LEMEIKDNST-SE---LSDDDWMTG-D-----DSDFEYRSN---------LKV--ETKTGAILTLKLLQNA---IK--SENHD---------DL--IMQE--FSEYVL--PNLL-KL-AIGVTAK-----G---------GKF-FDY-------LDN-------KKQ-------EEIENYHQNNLGKNQKLKEPKK---ID-RRNA-GDQSLNTHLLNGLLPANLIE--
RR------------------------LKKL--N-TT-VKRVI---QEQERRLLIAGFILHDFEK-----FDYQLFLKMPDKYKAIYQ-D--KEQD----------------IRK-LS--------VDEHREIIDVMVCEL-GLD----KFV--SP--DEPSAWLEYRDDLLFIAYN-AQR---RNDTNLNLSEY------GL-QA----KI----ED-YILT--CLADLTYLADLLSSI------IKHPQDA-E-----NQT-----------LNGIL-H-SL---SD-GKL----K-------
FTYHHIAENRGVLTNVVNNAVIEAH---TNLNTDE-NR--------Y---YQPLLYLPTGVIYLAL--RDAPP-I-SPE---ILPN--LVVNKI-----------------KSLCS--GQL--K-L-RQTGFGR-DG----------KG-MKYAEYYNLF-FDSPGLM----------------QVALDATL-----------------------RI-LN-----------------------T------GK---TSVGK---------------------------------------------SR-------SE---NLV---------KFQ------------KQNVLSI-------
DY---DFNF------DD-----DIRI-----DQ--------IA---------------------E----FG---DLIT--RK--IWE--------ETV--------ERIET--------ARKK-DKK----LPELPNFD---------LTHKVAEF-----WNLT------EYLS-QV--------------REIQRINESLKEQKLKGNTGGVPYEWYYLAAKYL----E--K---HP-GI-E---DVRE----NCQQV-IYYLTDLISP-----------ILA----------QY----Q--L--P-DG-WDD-------------LRLWVSRVV---------
MLP------------------------------TTKQETSVKTEVDNF--INELNNYNAAKK-SGRGKQL-ICSIS-HSAYTV---TEQMESAVLFTPQV--YTN--KQ-ML---GG--SNA-KRN-------ISSIAGVEMMLRQILMNQT-Q-----AVGKRFEDGKYRYLYF-YP---TYYFTP----E-TNKF---LQK-----AYMNI-AQTRF-----DT------SIR-----NHFI-S-------------------------------------------------KDLQV---------DLG------KENY--
QSVDSF-------L-IDENLDSEK------------------DR-------TF---KL---SY---PED--------------------QPLTFYFMAL----------PP------G----R--DS--------TDT-------ESW--VMPSWLAFAFPMILDVKTVVSESPIPPFNDGAEFEET----VF--LDSAPHAFRTLV-K--RD-RF--------RLD---YILEGWEE----------YGIQ-YPAPLNV-LTAAYAIH-LDV------NAKQGKTGYDANWG-KFTELAKDF-
ETSPLHVFAYLNKWVR---------NQK------------LDA--P---SI----NKVK---------LYAYH--FYPCFDP-Y------------------------------------------TKFDSNEEKLIVEEKSTLNH-PQKLTELYRRFYRANKRFNP-------KANAVLKPIDIAADTI-LKADSSVFD--G-ETLIAAVSAKIFKLMERVD-----S---STAEG-----RWVFKRN----------------QREEE----RQAILEFAEYFV---EEVFEKSFASDRARLAGRQLNLI-----
RDTCEFLYR------LADDE--ERK--ARPQ-DEPDEITEIETEEN------------
'TAF08895.1' ---MAKKSKKSDAHQQLSL------------------LEVEITDNSNPELSDDDWMSGD------DSDFEYTSN----------LKVETKTGAILTLKLLQNA---IK--SENFD---------DL--IMQE--FSDYVL--PNLL-RV-AVGVTAK-----G---------GKF-FDY-------LDQKKQEEIENYH-----------------QNNLGINQKLKEPKKIDRRNA-GDQSLNTHLLNGLLPANLIE--
RR------------------------LKKL----DTTAKRVI---QEQERRLLIAGFILHDFEKFDYRLF-----------------P-K--MPDKYKAICQDR------EQDIRKLS---------VDEHREIIDVMVCEL-GLN----KFV--SP--DEPSAWQEYRDDLLFIAYN-AQR---RNDTNLNLSEY------GL-QP----LI----DD-NILI--CLADLTCLADLLSSI------IKHPQDA-D-----NQT-----------LNGIL-H-SL---SD-GKL----K-------
FTYHHIAENRGVLTNVVNNAVIEAH---TNLNTDENRY-------------YQPLLYLPTGVIYLAL--RDAPP-I-SPE---ILPD--LVVNKI---------------KSLCA--GQL----KLRQTGFGR-DG----------KG-MKYAEYYNLF-FESPELM-----------------KVALDATL----------------------RI-LN-----------------------T------NK---PSVAK---------------------------------------------SR-------SQ---NLV---------KFQ------------KQNVLSA-------
DY---DFNF------DD----NIRI-----DQ--------IA---------------------E----FG---DLIT--LK--IWD--------ETV--------EKIET--------ARKKDKK-----LPEVPSFD---------LTHKVAEF-----WNLT------EYLP-QI--------------REIQRINESLKEQKLKGNTGGVPYEWYYLAAKYL----E--K---NP-DI-E---DVRE----NCQQV-IYYLTDLISP-----------ILA----------QY----Q--L--P-DG-WDD-------------LRLWVSRVV---------
MLP------------------------------NTNKSTSVTTEIDNF--INELSNYNAAKK-PGRGKQL-ICSIS-HSAYTV---TEQMESAVLFTPQV--YTN--KQ-ML---GG--SNA-KRN-------ISSIAGVEMMLRQILMNQT-Q------AVGKRFEDGKYRYLYF-YP---TYYFTP----E-TNKF---LQK-----AYTNI-AQTRF-----DT------SIR-----NHFI-S-------------------------------------------------KDLQV---------DLG------KENY--
QSVDSF-------L-IDENLDSEK------------------DR-------TF---KL---SY---PED--------------------QPLTFYFMAL----------PP------G----R--DS--------TDT-------ESW--VMPAWLAFAFPMILDVKTVVSESPIPPFNDGAEFEET----VF--LDSAPHAFRTLV-K--RD-RF--------RLD---YILEGWEE----------YGIQ-YSAPLNV-LTAAYAIH-LDV------NARQGKTGYDANWG-KFTELAKDF-
ETSPLYVFAYLNKWVR---------NQK------------VET--A---RI----EKIK---------LYTYH--FYPCFDP-Y------------------------------------------AKFDSTKEELIVGEISTLNH-PKKLTELYRRFYRANKRFNP-------KANAVLKPIDIAAETI-LKADSCVFD--G-ETLTLAVAAEIFKLMDRVH-----S---STAEG-----RWMMSK------------------REEE----RQAVLEFAKYFI---EEVFEGSFKGDRARLAGRQLNLI-----
RDTCEFLYR------LEDDK--ENA--GKTK-SEGNE----NE-------------
'WP_127055411.1' ---MAKKSKKSNDHQQLSL------------------LEVEITDNSNPELSDDDWMSGD------DSDFEYTSN----------LKVETKTGAILTLKLLQNA---IK--SENHH---------DL--IMQE--FSDYVL--PNLL-QV-AIGVTAK-----G---------GKF-FDY-------LDQ-------KKE-------KEIEDYHKNNLGRN-----QKTKEPKKFDRRNAGDQSLNTHLLNGLLPANLIE--
RR------------------------LKKL--N-TT-VKRVI---QEQERRLLIAGFILHDFEKFDYRLF-----------------P-K--MPDVYKAICQDK------EQDVRKLS---------VDDHREIIDVICEL-GLD----KFV--SP--DEPAAWIEYRDDLLFIAYN-AQR---RNDTNLNLSEY------GL--------QA--KIED-YVLT--CLVDLTYLADLLSSI------IKHPQDA-D-----NQT-----------LNGIL-H-GL---SD-GKL----K-------
FTYHHIAENRGVLTNVVNNAVIEAH---TNLNTEE-NR--------Y---YQPLLYLPTGVIYLAL--RDAPP-I-SPE---TLPD--LVVNKI-----------------KSLCA--GQL--K-L-RQTGFGR-DG----------KG-MKYAEYYNLF-FESPGLM----------------QVALDATL-----------------------RI-LN-----------------------S------GK---TSVAK---------------------------------------------SR-------SE---NLV---------KFQ------------KQNVLSA-------
DY---DFNF------DD-----SIRI-----DQ--------IA---------------------E----FG---DLIT--RK--IWD--------ETV--------EKIET--------ARKKDKK-----LPEVPNFN---------LTHKVAEF-----WNLT------EYLP-QV--------------REIQRINESLKEQKLKGNTGGVPYEWYYLAAKYL----E--K---HP-GI-E---DVRE----NCQQV-IYYLTDLISP-----------ILA----------QY----Q--L--P-DG-WDD-------------LRLWVSRVV---------
MLP------------------------------NTNKPTSVKTEVDNF--INELSNYNAAKK-PGRGKQL-ICSIS-HSAYTV---TEQMESAVLFTPQV--YTN--KQ-ML---GG--SNA-KRN-------ISSIAGVEMMLRQILMNQT-Q------AVGKRFEDGKYRYLYF-YP---TYYFTP----E-TNKF---LQK-----AYMNI-AQTRF-----DT------SIR-----NHFI-S-------------------------------------------------KDLQV---------DLG------KENY--
QSVDSF-------L-IDENLDSEK------------------DR-------TF---KL---SY---PED--------------------QPLTFYFMAL----------PP------G----R--DS--------TDT-------ESW--VMPSWLAFAFPMILDVKTVVSESPIPPFNDGAEFEET----VF--LDSAPHAFRTLV-K--RD-RF--------RLD---YILEGWEE----------YGIQ-YPAPLNV-LTAAYAIH-LDV------NARQGKTGYDANWG-KFTELAKDF-
ETSPLYVFAYLNKWVR---------NQG------------VEN--A---RI----EKIK---------LYTYH--FYPCFDP-Y------------------------------------------TKFDSTKEELIVGEKSTLNH-PQKLTELYRRFYRANKRFNP-------KANAVLKPIDIAADTI-LKADSCVFD--G-ETLISAVAAEVFKLMDRVH-----S---STAEG-----RWIMSK------------------REEE----RQAVLEFANYFV---VDVFEGSFKSDRARLAGRQLNLI-----
RDTCEFLYR------LEDDK--ENA--GKTK-SETEE----NEESN-------------
'WP_045870814.1' MAKKSKAAKASNEYQQLSL-LD------------IE--NTAN-SE---SSDEDWLSG-------DFGFDSDDN---------RTI--ITKQ-KLLTLKLLQEA---IT--AQNPD---------DI--VMAD--FADYVL--PNLL-KI-AIGVTAK-----G---------GKF-FDE-------IDQ-------QRE-------AA---------GKN-------K---VR-RDNA-ADQSLNTHLLNGLFPANLIE--KR-----------------
-------LEKL--D-TT-IKRVV---KERERRLVIAGFILHDFEK-----F----------------P-D--VPEN-----------------CRK-LA--------LAEHRQIIDEKVKQL-GLD----QFI--NP--DNPEDYTEYLDDLLCLAYN-AQR---RWDTNWNFSEF------GL-NP----VL----KD-RTLR--SLADLTCLADSLASI------IKHPQDA-E-----NPR-----------LQEII-H-NL---SD-GQL----K-------FTYHSIAENRGVLTNVVNNAVIQAY---
TSLNTDE--Q------I-Y---YEPLLYLPTGVIYLAA--RNAPS-L-SAK---DLPD--LVVNNI-----------------KSLCA--GQL--K-R-RQTGFGR-DG----------KG-MKYAEYYNLF-FDSPGLM----------------KVALDATL-----------------------RI-LS-----------------------S------TK---PSVAK---------------------------------------------SR-------SD---NLV---------KFQ------------QQGVLPA-------EF---DFKF------TD-----DIRI-----DQ-------
--IA---------------------E----FG---DLVS--RK--IWD--------EAV--------NRIEA--------IRKK-DKK----LPAIPAFD---------LTHKVAEF-----WNLT------QHLP-EI--------------RHIQRINESLKENKLKGNTGGVPYEWYYLAAKYL----E--A---HP-GV-E---DVRE----VCQQV-IEYLTGLISP-----------ILA----------QY----Q--L--P-DG-WDD-------------LRQWVSRVV----------MLP-----------------------NTSQQQST--
QTQTENF--LNELGNYNAAKK-PGRGRQL-ICSIS-HSAYTV---TEQMESAVLFTPQV--YTN--KQ-ML---GG--SNA-KRN-------ISSIAGVEMMLRQILMNQT-Q------AVGKRFEDGKYRYLYF-YP---TYYFTP----E-TNKF---LQK-----AYSGV-AQTRF-----DT------SIR-----NHFI-T-------------------------------------------------KNFQA---------HLD------KENY--QNVDSF-------L-IDENLDSEK------------------
--DR-------TF---KL---AY---PED--------------------QPLTFYFMAL----------PP------G----R--DS--------TDT-------ESW--VMPTWLAFAFPMILDVKTVVSESPIPPFNDGAEFEES----VF--LDSAPHAFRALV-K--RD-RF--------RLD---YILEGW------QE----NGIQ-YPAPLNV-LTAAYAIH-LDV------NARQGKSGYDANWG-RFSELAKDL-ETSPLYVFYYLNRWAR---------NQG------------VDS--A---
RV----EKIR---------LYAYH--FYPCFDP-Y------------------------A-KFDYTKEE--------L-IVG-EE--SS-LNH-PKKLTELYRRFYR--A-NKR--YNP--KANAVLKPIDIASETI-LKAELSVFR--G-ETLVTAVAAEVFKLMDRVH-----S---STAEG-----RWVM-S---------K---------REEE----RQAILDFAQYFV---VEVFEKSFAGDRARLAGRQLNLI-----RDTCEFLYR------LEDDK--ENL--EKNQ-KSSES--------
D-KE---------
'WP_096577327.1' MAKKSKAAKASDEYQQLSL-LD------------IE--NTAN-AE---SSDEDWLSG-------DFGFDSDGD---------RTI--ETKQ-QLLTLKLLQEA---IT--AQNPD---------DI--VMAD--FAQYVL--PNLL-KI-AIGVTAK-----G---------GKF-FDE-------IDQ-------QRE-------AA---------GKT-------K---VR-RDNA-ADQSLNTHLLNGLFPANLIE--KR-----------------
---------LENL--D-TT-IRRVV---KERERRLVIAGFILHDFEK-----F----------------P-D--VPEN-----------------CRK-LA--------LAEHRQIIDEKVKQL-GLN----KFI--NP--DNPEDYKEYLDDLLCLAYN-AQR---RWDTNWNFSEF------GL-NP----VL----KD-RTLR--CLADLTCLADSLASI------VKHPQDA-E-----NPR-----------LQEII-H-NL---SD-GQL----K-------FTYHRIAENRGVLTNVVNNAVIQAY---
TNLNTNE--Q------I-Y---YEPLLYLPTGVIYLTA--RNAPS-L-STK---DLPD--LVVNNI-----------------KSLCQ--GQL--K-L-RQTGFGR-DG----------KG-MKYAEYYNLF-FDSPGLM----------------EVALDATL-----------------------RI-LS-----------------------S------TK---PSVAK---------------------------------------------SR-------SE---NLV---------KFQ------------QQGVLPA-------EF---DFKF------TD-----DIRI-----DQ-------
-IA---------------------E----FG---DLVS--RK--IWD--------EAV--------NRIEA--------IRKK-DKK----LPAIPAFD---------LTHKVAEF-----WNLT------EYLP-EI--------------RHIQRINESLKENKLKGNTGGVPYEWYYLAAKYL----E--A---HP-GV-E---DVRE----ICQQV-IEYLTGLISP-----------ILS----------QY----Q--L--P-DG-WDD-------------LRQWVSRVV----------MLP-----------------------NTNQQSSS--
ETQTENF--LNELSNYNAAKK-PGRGQL-ICSIS-HSAYTV---TEQMESAVLFTPQV--YTN--KQ-ML---GG--SNA-KRN-------ISSIAGVEMMLRQILMNQT-Q------AVGKRFEDGKYRYLYF-YP---TYYFTP----E-TNKF---LQK-----AYSGV-AQTRF-----DT------SIR-----NHFI-S-------------------------------------------------KDFQA---------NLD------KENY--QNVDSF-------L-IDENLDNEK------------------
--DR-------TF---KL---TY---PED--------------------QPLTFYFMAL----------PP------G----R--DS--------TDT-------ESW--VMPTWLAFAFPMILDVKTVVSESPIPPFNDGAEFEES----VF--LDSAPHAFRALV-K--RD-RF--------RLD---YILEGW------QE----NGIQ-YPAPLNV-LTAAYAIH-LDV------NARQGKSGYDANWG-RFSELAKDL-ETSPLYVFYYLNRWAR---------NQG------------VES--A---
RV----EKIR---------LYTYH--FYPCFDP-Y------------------------A-KFDYTKEE--------L-IVG-EE--SS-LNH-PKKLTELYRRFYR--A-NKR--YNP--KANAVLKPIDIASETI-LKAELSVFR--G-ETLVTAVAAEVFKLMDRVH-----S---STAEG-----RWVM-S---------K---------REEE----RQAILDFAQYFV---VEVFEKSFAGDRARLAGRQLNLI-----RDTCEFIYR------LEDDK--ENI--EKNQ-KSPES--------D-
NQ---------
'WP_096642022.1' MAKKSKAAKPSDEYQQLSL-LD------------IE--NIAN-SE---SSDEDWLSG-------DFGFDSDSD---------RTI--VTKQ-QLLTLKLLQEA---IT--VQNPD---------DI--VMAD--FAEYVL--PNLL-KI-AIGVTAK-----G---------GKF-FDE-------IDQ-------QRE-------AA---------GKT-------R---VR-RDNA-ADQSLNTHLLNGLFPANLIE--KR-----------------
-------LENL--D-TT-IRRVV---KERERRLVIAGFILHDFEK-----F----------------P-D--VPEN-----------------CRK-LA--------LAEHRQIIDEKVKQL-GLD----QFI--NP--NNPEAYTEYLDDLLCLAYN-AQR---RWDTNWNFSEF------GL-NP----VL----KD-RTLR--CLADLTCLADSLASI------VKHPQDA-E-----NLR-----------LQEII-H-NL---SD-GQL----K-------FTYHRIAENRGVLTNVVNNAVIQAY---
TNLNTNE--Q------I-Y---YEPLLYLPTGVIYLAA--RNAPY-L-STK---ELPD--LVVNNI-----------------KFLCA--GQL--K-L-RQTGFGR-DG----------KG-MKYAEYYNLF-FDSPGLM----------------KVALDATL-----------------------RI-LS-----------------------S------TK---PSVAK---------------------------------------------SR-------SE---NLV---------KFQ------------QQGVLPA-------EF---DFKF------TD-----DIRI-----DQ-------
-IA---------------------E----FG---DLVS--RK--IWD--------EAV--------NRIEV--------IRKK-DKK----LPAIPAFD---------LTHKVAEF-----WNLT------EYLP-EI--------------RHIQRINESLKENKLKGNTGGVPYEWYYLAAKYL----E--A---HP-GV-E---DVRE----ICQQV-IEYLTGLISP-----------LLA----------QY----Q--L--P-DG-WDD-------------LRQWVSRVV----------MLP-----------------------NTNQQSSP--
ETQIENF--LNELGNYNAAKK-PGRGRQL-ICSIS-HSAYTV---TEQMESAVLFTPQV--YTN--KQ-ML---GG--SNA-KRN-------ISSIAGVEMMLRQILMNQT-Q------AVGKRFEDGKYRYLYF-YP---TYYFTP----E-TNKF---LQK-----AYSGV-AQTRF-----DT------SIR-----NHFI-T-------------------------------------------------KDFQA---------HLD------KENY--QSVDSF-------L-IDENLDNEK------------------
-DR-------TF---KL---AY---PED--------------------QPLTFYFMAL----------PP------G----R--DS--------TDT-------ESW--VMPTWLAFAFPMILDVKTVVSESPIPPFNDGAEFEES----VF--LDSAPHAFRALV-K--RD-RF--------RLD---YILEGW------QE----NGIQ-YPAPLNV-LTAAYAIH-LDV------NARQGKSGYDANWG-RFSELAKDL-ETSPLYVFYYLNRWAR---------NQG------------VDS--A---RV-
---EKIR---------LYTYH--FYPCFDP-Y------------------------A-KFDYTKEE--------L-IVG-EE--SS-LNH-PKKLTELYRRFYR--A-NKR--YNP--KANAVLKPIDIASETI-LKAELSVFR--G-ETLVTAVAAEVFKLMDRVH-----S---STAEG-----RWVM-S---------K---------REEE----RQAILDFAQYFV---VEVFEKSFAGDRARLAGRQLNLI-----RDTCEFLYR------LEDDK--ENL--DKNQ-KSSES--------D-
KE---------
'WP_099066180.1__14_' ---MIKKSKIANKVQQLSL-LE------------TE--NDIN-SQ---VMDEDWLSG-------DFGFDGDSS-------DRTI--EIEA-ELLTLKLLREA---IQ--TQNPG---------DK--VMAD--FGEYVL--PNLL-RI-AIGVTAK-----G---------GKF-FDE-------IDQ-------KRE-------AE---------GKS-------K---VR-RDNA-ADQSLNTHLLNGLFPANLIE--KK-----------------
-----LENL--D-TT-VRRVV---KERERRLVIAGFILHDFEK-----F----------------P-D--APEE-----------------CRK-LP--------LEKHRQIVEEKVQEL-GLD----KFI--NS--EHPEAYKEYIDDLLCMAYN-AQS---RWDTNWNFSEF------GL-NP----VL----KD-RTIR--SLCDLTRLADSLLASV------IKHPQDA-E-----HRA-----------LNDIL-H-SL---SD-GQL----K-------FTYHRIAENRGVLTNVVNNAVIQAY---
TSLNTDE--H------T-Y---YEPLLYLPTGVIYLAA--RNAPS-I-SAT---DLPD--LVVNNI-----------------KSLCA--GQL--R-L-RQTGFGR-DG----------KG-MKYAEYYNLF-FDSPGLM----------------QVALDATL-----------------------RI-LN-----------------------S------NK---PSVAK---------------------------------------------SR-------SE---NLV---------KFQ------------QQGVLPP-------EF---DFNF------AD-----DIRI-----DQ-------
--IA---------------------E----FG---DLVS--RK--IWE--------ETV--------NRLEA--------VRKK-DKK----LPEIPSLN---------LIHQVAEF-----WNLT------EYIP-QI--------------RYIERINESLKENNLKGNTGGVPYEWYYLAAKYL----E--K---HP-GI-E---NVRE----SCQEV-INYLTNLISP-----------ILA----------QY----E--L--P-DG-WDD-------------LREWVTRVV----------MLP-----------------------STNQQLSS--
KIQSENF--FQELDNYNAAKK-PGRGRQL-ICSIS-HSAYTV---TEQMESAVLFTPQV--YTN--KQ-ML---GG--SNA-KRN-------ISSIAGVEMMLRQILMNQT-Q------AVGKRFEDGKYRYLYF-YP---TYYFTP----E-TNKF---LQK-----AYGGI-AQTRF-----DT------SVR-----NHFI-T-------------------------------------------------KDLQA---------HLA------KKDY--QSVDSF-------L-IDENLEDDK------------------
-DR-------TF---KL---SY---PED--------------------QPLTFYFMAL----------PP------G----R--DS--------TDT-------ESW--VMPSWLAFAFPMILDVKTVVSESPIPPFNDGAEFEES----VF--LDSAPHAFRALV-R--RD-RF--------RLD---YILEGW------NE----NGIE-YPAPLNV-LTAAYAIH-LDV------NARQGKSGYDANWG-RFTELAKDF-ETSPLYVFSYLNRWVR---------NQG------------VET--A---RI-
--EKIR---------LYTYH--FYPCFDP-Y------------------------A-KFDRINEE--------L-IVG-EE--SS-LNH-PKKLTELYRRFYR--A-NKR--YNP--KSNAVLKPIDIAAETI-LKAELSVFH--G-ETLVAAVAAEVFKLMDRVH-----S---STAEG-----RWIM-S---------K---------REEE----RLAVLEFARYFV---EEVFDKSFAGDRARLAGRQLNLI-----RDTCEFLYR------LEDDK--ENL--ARNE-KSPES--------D-
NENE-------
'AFY46833.1__2_' ---------------------------------------M----------TDIDWLSG-------DFGFDGDGS--------RTI--ETQG-ELLSLKLLREA---IQ--AQNPD---------DS--VMAD--FAEYVL--PNLL-RI-AIGVTAK-----G---------GKY-FDA-------IDQ-------QRE-------AE---------GKT-------K---VR-RDNA-ADQSLNTHLLNGLFPANLIE--RR---------------------------LQQL--D-TT-
VRRVV---KERERRLVIAGFILHDFEK-----F----------------P-D--VPND-----------------CRK-LP--------LAEHRPIIDTKVRQL-GLD----KFI--NP--ENPEAYQEYIDDLLCMAYN-AQS---RWDTNWNFSEF------GL-NP----VL----RD-RTLR--QLSGLTLLADSLASI------VKHPQDA-E-----HPK-----------LKEVL-H-FL---SD-GQL----K-------LTYHSIAENRGVLTNVVNNALIQAH---TSLNTDE--H------T-
Y---YEPLLYLPTGVIYLAA--RYAPP-I-SPQ---DLPD--RVVDNI-----------------KSLCA--GQL--R-L-RQTGFGR-DG----------KG-MKYAEYYNLF-FDSPGLM----------------QVALDATL-----------------------RI-LN-----------------------P------NK---VSVAK---------------------------------------------SR-------SD---NLI---------KFQ------------KQGVLPA-------DY---DFKF------DD-----DIRI-----DQ--------IA-------------------
E----FG---DLVS--RK--IWD--------ETV--------HRIAD--------IRKK-DKK----LPEVPALD---------LTHKVAEF-----WNLT------GYLP-QI--------------IKIQRINESLKENKLKGNTGGVPYEWYYLAAKYL----E--N---HP-GI-E---SVRQ----TCQEV-IQYLTGLISP-----------IIS----------QY----E--L--P-DG-WDD-------------LRLWVSRVV----------MLS-----------------------NTNQEASAKTQVDTF--
LAELNNYNAAKK-PGRGKQL-ICSIS-HSAYTV---TEQMESAVLFTPQV--YTN--KQ-ML---GG--SNS-KRN-------ISSIAGVEMMLRQILMNQT-Q------AVGKRFEDGKYRYLYF-YP---TYYFTP----E-TNKF---LQK-----AYNGI-AQTRF-----DT------SVR-----NHFI-S-------------------------------------------------KDLQA---------NLE------RENY--QSVDAF-------L-IDENLERDK------------------DR-------
TF---KL---SY---PDD--------------------QPLTFYFMAL----------PP------G----R--DS--------TDT-------ESW--IMPSWLAFAFPMILDVKTVVSESPIPPFNDGAEFEES----VF--LDSAPHAFRALV-R--RD-RF--------RLD---YILEGWED----------NGIQ-YAAPLNV-LTAAYAIH-LDV------NARQGKSGYDANWG-KFTELAKDF-ETSPLYVFSYLNRWVR---------NQK------------LDA--L---SL----SKIK-----
-----LYAHH--FYPCFDP-Y------------------------AKYDFQLEKWTMAEESHLHH-PKKLTDLYRKFYRAKSSKGKPI-------KSNAILKPIDEAAETI-IKADKSCFQ--G-EVLVDAVAAKIFKLMERVR-----N---STAEG-----RWVITH-------------------RDEE----RDAILDFARYFV---LEVFEKCFAGDRARLAGRQLNLL-----KDTCEFLYR------LENDK--ERR--DKAN-KSNAE-----IYEDEDGDEDEK-
----
'WP_011320851.1__2_' ---------------------------------------M----------TDHDWLSG-------DFGFDGDGS--------RTI--ETEG-ELLSLKLLREA---IQ--AQNPD---------DA--VMTD--FAEYVL--PNLL-RI-AIGVTAK-----G---------GKY-FDA-------IDQ-------QRE-------AA---------GKT-------K---VR-RDNA-ADQSLNTHLLNGLFPANLIE--RR---------------------------LQQL--D-TT-
VRRVV---KERERRLVIAGFILHDFEK-----F----------------P-D--VPND-----------------CRK-LP--------LAEHRPIIDAKIRQL-GLD----KFI--NP--ENSEAYQEYIDDLLCMAYN-AQS---RWDTNWNFSEF------GL-NP----VL----RD-RTLR--QLSGLTLLADSLASI------VKHPQDA-E-----HSK-----------LKEVL-H-FL---SD-GQL----K-------LTYHSIAENRGVLTNVVNNALIQAH---TSLNTDE--H------T-
Y---YEPLLYLPTGVIYLAA--RHAPP-I-STE---DLPD--RVVDNI-----------------KSLCA--GQL--R-L-RQTGFGR-DG----------KG-MKYAEYYNLF-FESPGLM----------------QVALDATL-----------------------RI-LN-----------------------P------NK---DSVAK---------------------------------------------SR-------SE---NLI---------KFQ------------QQGVLCA-------DY---DFKF------DD-----DIRI-----DQ--------IA-------------------
E----FG---DLVS--RK--IWD--------ETV--------NRIED--------IRKK-DKK----LPAVPALD---------LTHKVAEF-----WNLT------AYLP-QI--------------REIQHINESLKENKLKGNTGGVPYEWYYLAAKYL----E--H---HP-GI-E---SVRE----TCQQV-IQYFTGLISP-----------IIS----------QY----K--L--P-DG-WDD-------------LRLWVSRVV----------MLP-----------------------NTNQEASA--KTQVDTF--
LAELNNYNAAKK-PGRGKQL-ICSIS-HSAYTV---TEQMESAVLFTPQV--YTN--KQ-ML---GG--SNA-KRN-------ISSIAGVEMMLRQILMNQT-Q------AVGKRFEDGKYRYLYF-YP---TYYFTP----E-TNKF---LQK-----AYNGI-AQTRF-----DT------SVR-----NHFI-S-------------------------------------------------KDLQA---------NLE------REKY--QSVDAF-------L-IDENLEREK------------------DR-------
TF---KL---SY---PDD--------------------QPLTFYFMAL----------PP------G----R--DS--------TDT-------ESW--IMPSWLAFAFPMILDVKTVVSESPIPPFNDGAEFEES----VF--LDSAPHAFRALV-R--RD-RF--------RLD---YILEGW------DD----NGIQ-YPAPLNV-LTAAYAIH-LDV------NARQGKSGYDANWG-RFTELAKDF-ETSPLYVFSYLNRWVR---------NQG------------IET--A---RI----EKIR-----
-----LYAYH--FYPCFDP-Y------------------------V-QYDMDTKT--------L-TVK-SE--SS-LNH-PKNLTELYRRFYR--A-NKR--YNP--KSNAVLKPIDIAAETI-LKAESSVFH--G-ETLVVGVAAEVFRLMDRVH-----A---STAEG-----RWVM-S---------K---------REEE----RQAVLDFAKYFV---IEVFEKAFAGDRARLAGRQLNLI-----RDTCEFLYR------LEDDK--ENA--AKED-KNPES--------D-DD-------
'WP_096682794.1' ---------------------------------------M----------SDNDWLLG-------DFGFGGDGN--------RTI--ERQG-ELLSLKLLREA---IQ--AQNPD---------DT--VMAD--FAEYVL--PNLL-RI-AIGVTAK-----G---------GKY-FDA-------IDQ-------QRE-------AE---------GKN-------K---VR-RDNA-ADQSLNTHLLNGLFPANLIE--RR---------------------------LQQL--D-
TT-VRRVV---KERERRLVIAGFILHDFEK-----F----------------P-D--VPND-----------------CRK-LP--------LAEHRPIIDTKVRQL-GLD----KFI--YP--KNPEAYQEYIDDLLCMAYN-AQS---RWDTNWNFSEF------GL-NP----VL----KD-RTLR--QLSGLTLLADSLASI------VKHPQDA-E-----HSK-----------LKEVL-H-FL---SD-GQL----K-------LTYHSIAENRGVLTNVVNNALIQAH---TSLNTDE--H------
I-Y---YEPLLYLPTGVIYLAA--RHAPP-I-STK---DLPD--RVVDNI-----------------KSLCA--GQL--R-L-RQTGFGR-DG----------KG-MKYAEYYNLF-FDSPGLM----------------QVALNATL-----------------------RI-LN-----------------------P------NK---DSVAN---------------------------------------------SR-------SE---NLI---------KFQ------------QQGVLSA-------DY---DFKF------DD-----DIRI-----DQ--------IA-------------------
E----FG---DLVS--RK--IWD--------ETV--------NRIED--------IRKK-DRK----LPEVPALD---------LTHKVAQF-----WNLT------AYLP-QI--------------REIQRINESLKENKLKGNTGGVPYEWYYLAAKYL----E--H---NP-GI-E---SVRE----TCQQV-IQYLTGLISP-----------IIA----------QY----Q--L--P-DG-WDD-------------LRLWVSRIV----------MLS-----------------------NTNQETSD--KTQVNTF--
LAELNNYNAAKK-PGRGRQL-ICSIS-HSAYTV---TEQMESAVLFTPQV--YTN--KQ-ML---GG--SNA-KRN-------ISSIAGVEMMLRQILMNQT-Q------AVGKRFEDGKYRYLYF-YP---TYYFTP----E-TNKF---LQK-----AYSGI-AQTRF-----DT------SVR-----NHFI-G-------------------------------------------------KDLQA---------DLG------RENY--QSVDAF-------L-IDENLERDK------------------DR-------
TF---KL---SY---PED--------------------QPLTFYFMAL----------PP------G----R--DS--------TDT-------ESW--IMPSWLAFAFPMILDVKTVVSESPIPPFNDGAEFEES----VF--LDSAPHAFRALV-R--RD-RF--------RLD---YILEGW------QD----NGIQ-YPAPLNV-LTAAYAIH-LDV------NARQGKSGYDANWG-RFTELAKDF-ETSPLYVFSYLNRWVR---------NQG------------IET--A---RI----EKIR-----
-----LYAYH--FYPCFDP-Y------------------------V-QYDMNTKT--------L-NVQ-PE--SS-LNH-PQNLTELYRRFYR--A-NKR--YNP--KSNAVLKPIDIAAETI-LKAESRVFQ--G-ETLVVAVAAEVFKLMDRVH-----A---STAEG-----RWVM-S---------K---------REEE----RQAVLDFAKYFV---IEVFEKAFAGDRARLAGRQLNLI-----RDTCEFLYR------LADDK--ENA--GKDD-ESTES--------D-DD-------
'WP_104900394.1__2_' ---MAKKSKKSDEYQQLSL-LE------------AETENDST-SE---LSDDDWMSG-------DDSDFQETS--------NLKV--ETQSAALLTLKLLQDA---IK--SENDD---------DL--IMQE--FSDYVL--PNLL-RV-AVGVTAK-----G---------GKF-FDY-------FDE-------KRG-------KEIENFHKNNLGKSHKIKEPKK---LD-RRNA-GDQSLNTHLLNGLLPANLIE--
RC--------------------------LNKL--D-TT-AKRVI---QEQERRLLIAGFILHDFEKFDYRLF-----------------P-Q--MPDNYKAIGQDK------EQDIRK-LS--------VDAHREIIDVICEL-GLD----KFV--SP--DEPTAWQEYRDDLLFIAYN-AQR---RNDTNLNVSEH------GL-HP----II----DD-YILI--CLADLTCLADLLSSV------IKHPQDA-E-----HQA-----------LNGIL-H-SL---SD-GQL----K-------
FTYHSIAENRGVLTNVVNNALIQAH---TSLNTDE--H------T-Y---YEPLLYLPTGVIYLAS--RNAPA-I-SPE---DLPD--RVVNSI-----------------KSLCA--GQL--R-L-RQTGFGR-DG----------KG-MKYAEYYNLF-FDDIGLM----------------KVALDATL-----------------------RI-LN-----------------------P------NK---GSVAK---------------------------------------------SR-------SE---NLN---------KFQ------------QQNVLSA-------
DY---DFKF------ED-----DIRI-----DQ--------IA---------------------E----FG---DLVS--RK--IWE--------ETV--------NRVDS--------ARKK-DKK----LPAVPDFD---------LTHKVAEF-----WNLA------ECLP-QI--------------REIQRINESLKENKLKGNTGGVPYEWYYLAAKYL----E--H---HP-GI-E---DVRE----ACQQV-IDYLTTLISP-----------IIS----------QY----K--L--P-DG-WDD-------------LRLWVQRVV---------
MLP----------------------GTNQVSSV--KAQVETF--LDELGNYNAAKK-AGRGKQL-ICSIS-HSAYTV---TEQMESAVLFTPQV--YTN--KQ-ML---GG--SNA-KRN-------ISSIAGVEMMLRQILMNQT-Q------AVGKRFEDGKYRYLYF-YP---TYYFTP----E-TNKF---LQK-----AYNGI-AQTRF-----DT------SIR-----NHFI-S-------------------------------------------------KDLQA---------DLG------RDRY--
QSVDSF-------L-IDENLQRDK------------------DR-------TF---KL---SY---PED--------------------QPLTFYFMAL----------PP------G----R--DS--------TDT-------ESW--VMPTWLAFAFPMILDVKTVVSESPIPPFNDGAEFEES----VF--LDSAPHAFRALV-R--RD-RF--------RLD---YILEGW------NE----NGIQ-YPAPLNV-LTAAYAIH-LDV------NARQGKSGYDANWG-RFTELAKDF-
ETSPLYVFSYLNRWVR---------NQG------------LET--A---RI----EKIR---------LYAYH--FYPCFDP-Y------------------------V-KYDTNMEQ--------L-IVG-EA--SS-LNH-PKKLTDLYRRFYR--A-NKR--YNP--KSNAVLKPVDIAAETI-LKAESSVFQ--G-ETLVVAVAAEVFKLMDRVH-----S---STAEG-----RWIM-S---------K---------REEE----RQAVLDFAKYFV---VEVFDNAFAGDRARLAGRQLNLI---
--RDTCEFLYR------LEDDK--ENA--GINE-KSVEI--------D-NELMQEFIDNL
'WP_089091838.1' ---MTKKTKPSEENLPPSL-LK------------NNTED----------LEEDFFNQ-------DFGFDGDSS-------DRTI--ETEG-ELLTLKLLREA---IH--SQNPD---------DQ--VMAD--FAEYVL--PNLL-KI-AIGVTAK-----G---------GKF-FDE-------IDQ-------QRE-------AE---------GKT-------K---VR-RDNA-ADQSLNTHLLNGLFPANLIE--KR-----------------
---LEKL--N-TT-VRRVV---KERERRLVIAGFILHDFEK-----F----------------P-D--APED-----------------CRK-LP--------LTEHRQIIDQKVSQL-GLD----KFI--NP--DDSEAYREYLDDLLCMAYN-AQR---RWDTNWNFSEF------GL-NP----VL----KD-RTLR--SLSDLTCLADSLASI------VKHPQDA-E-----NTR-----------LQEII-H-NL---SD-GQL----K-------FTYHSIAENRGVLTNVVNNALIKAH---
TSLNTDE--N------I-Y---YEPLLYLPTGVIYIAL--RNAPP-I-AQA---DLSD--RVVDSI-----------------KSLCA--GQL--R-L-RQTGFGR-DG----------KG-MKYAEYYNLF-FDDLGLM----------------RVALDATL-----------------------RI-LN-----------------------P------NK---PSVAK---------------------------------------------SR-------SE---NLI---------KFQ------------QKNVLSP-------DY---DFEF------AD-----DIRI-----DQ-------
IA----------------------E----FG---DLIS--RK--IWE--------ETV--------KRIDV--------ARKK-DKK----LPAVTDLY---------LTQKVAEF-----WNLA------EYLP-PI--------------REIEDINESLKENKLKGNTGGVPYEWYYLAAKYL----E--L---HP-GI-E---NVRE----TCQQV-IDYLATLISP-----------IIS----------QY----E--L--P-DG-WDD-------------LRLWVNRVV----------MLP-----------------------NTNQELST--
ETQVKTF--LEELSNYNAAKK-PGRGRQL-ICSIS-HSAYTV---TEQMESAVLFTPQV--YTN--KQ-ML---GG--SNA-KRN-------ISSIAGVEMMLRQILMNQT-Q------AVGKRFEDGKYRYLYF-YP---TYYFTP----E-TNKF---LQK-----AYNGI-AQSRF-----DT------SVR-----NHFI-S-------------------------------------------------KDLQA---------NLA------KDNY--QSVDSF-------L-IDENLQADK------------------
--DR-------TF---KL---SY---PED--------------------QPLTFYFMAL----------PP------G----R--DS--------TDT-------ESW--VMPSWLAFAFPMILDVKTVVSESPIPPFNDGAEFEES----VF--LDSAPHAFRALV-K--RD-RF--------RLD---YILEGW------QE----NGIQ-YPAPLNV-LTAAYAIH-LDV------NARQGKTGYDANWG-KFTELAKDF-ETSPLYVFAYLNRWVR---------NQG------------VET--A---RI-
---EKIR---------LYVYH--FYPCFDP-Y------------------------V-KYEKNMEQ--------L-IVG-EA--SS-LNH-PKKLTDLYRKFYR--A-NKR--YNP--KSNAVLKPIDIAAETI-LKAESSVFQ--G-ETLVAAVAAEVFKLMDRVH-----S---STAEG-----RWVM-S---------R---------REEE----RQAVLAFASYFV---QEVFEKSFAGDRARLAGRQLNLI-----RDTCEFLYR------LEDDQ--ENA-----E-RVAEA--------E-----
-------
'RCJ39405.1' ---------------------------------------M----------SNDDWLSG-------DFGFDGDSS-------DRTI--ETEG-ELLTLKLLREA---IQ--SQNPD---------DK--VMAD--FGEYVL--PNLL-RI-AIGVTAK-----G---------GKF-FDE-------IDQ-------QRE-------AE---------GKT-------K---VR-RDNA-ADQSLNTHLLNGLFPANLIE--KR---------------------------LEKL--N-TT-
VQRVV---KERERRLVIAGFILHDFEK-----F----------------P-D--VPEN-----------------CRK-LP--------LTEHREIIDKKVHQL-GLD----NFI--NP--ENPEAYREYLDDLLCMAYN-AQR---RWDTNWNFSEF------GL-NP----LL----KD-RTLR--SLSDLTCLADSLASI------VKHPQDA-E-----HPR-----------LKEII-H-SL---SD-GQL----K-------FTYHSIAENRGILTNVVNNALIQAH---TSLNTDE--H------T-Y-
--YEPLLYLPTGVIYLSS--RNAPA-I-SPE---DLPD--RVVNSI-----------------KSLCA--GQL--R-L-RQTGFGR-DG----------KG-MKYAEYYNLF-FDDIGLM----------------KVALDATL-----------------------RI-LN-----------------------P------NK---GSVAK---------------------------------------------SR-------SE---NLN---------KFQ------------QQKVLSA-------DY---DFKF------ED-----DIRI-----DQ--------IA-------------------E----
FG---DLVS--RK--IWE--------ETV--------NHVNS--------ARKK-DKK----LPAVPDFD---------LTHKVAEF-----WNLA------ECLP-QI--------------REIQRINESLKENKLKGNTGGVPYEWYYLAAKYL----E--H---HP-GI-E---DVRE----ACQQV-INYLTTLIYP-----------IIS----------QY----Q--L--P-DG-WDD-------------LRLWVKRVV----------MLP-----------------------GTNQEPSV--QTQVETF--
LNELDNYNAAKK-AGRGKQL-ICSIS-HSAYTV---TEQMESAVLFTPQV--YTN--KQ-ML---GG--SNA-KRN-------ISSIAGVEMMLRQILMNQT-Q------AVGKRFEDGKYRYLYF-YP---TYYFTP----E-TNKF---LQK-----AYNGI-AQTRF-----DT------SVR-----NHFI-S-------------------------------------------------KDLQA---------NLG------RDRY--QSVDSF-------L-IDENLQRDK------------------DR-------
TF---KL---SY---PED--------------------QPLTFYFMAL----------PP------G----R--DS--------TDT-------ESW--VMPTWLAFAFPMILDVKTVVSESPIPPFNDGAEFEES----VF--LDSAPHAFRALV-R--RD-RF--------RLD---YILEGW------NE----NGIQ-YPAPLNV-LTAAYAIH-LDV------NARQGKTGYDANWG-RFTELAKDF-ETSPLYVFSYLNRWVR---------NQG------------SET--A---RI----EKIR-----
------LYAYH--FYPCFDP-Y------------------------V-KYDNNMEQ--------L-IVG-EA--SN-LNH-PKKLTDLYRKFYR--A-NKR--YNP--KSNAVLKPIDIAAETI-LKAESSVFQ--G-ETLVVAVAAEVFKLMDRVH-----A---STAEG-----HWII-S---------K---------REEE----RQAVLDFARYFV---TEVFEKSFAGDRARLTGRQVNLI-----RHTCEFIYR------LEDDK--ENS--ARNE-QSTES--------N-DDGNQ------
'WP_086836537.1__2_' ---------------------------------------M----------SNDDWLSG-------DFGFDGDSS-------DRTI--ETES-ELLTLKLLREA---IE--AQNPD---------DR--VMAD--FGEYVL--PNLL-RV-AIGVTAK-----G---------GKF-FDE-------IDQ-------QRE-------AE---------GKT-------K---VR-RDNA-ADQSLNTHLLNGLFPANLIE--KR---------------------------LEKL--N-
TT-VQRVV---KEREQRLVIAGFILHDFEK-----F----------------P-D--VPEN-----------------CRK-LP--------LTEHREIIDKKVHQL-GLD----NFI--NP--ENPEAYREYLDDLLCMAYN-AQR---RWDTNWNFSEF------GL-NP----LL----KD-RTLR--SLSDLTCLADSLASI------VKHPQDV-E-----HPR-----------LKEII-H-SL---SD-GQL----K-------FTYHSIAENRGVLTNVVNNALIQAH---TSLNTDE--H------
T-Y---YEPLLYLPTGVIYLAS--RNAPA-I-SPE---DLPD--RVVNSI-----------------KSLCA--GQL--R-L-RQTGFGR-DG----------KG-MKYAEYYNLF-FDDIGLM----------------KVALDATL-----------------------RI-LN-----------------------P------NK---GSVAK---------------------------------------------SR-------SE---NLN---------KFQ------------QQNVLSA-------DY---DFKF------ED-----DIRI-----DQ--------IA-------------------
-E----FG---DLVS--RK--IWE--------ETV--------NRVDS--------ARKK-DKK----LPAVPDFD---------LTHKVAEF-----WNLA------ECLP-QI--------------REIQRINESLKENKLKGNTGGVPYEWYYLAAKYL----E--H---HP-GI-E---DVRE----ACQQV-IDYLTTLISP-----------IIS----------QY----E--L--P-DG-WDD-------------LRLWVKRVV----------MLP-----------------------GTNQEPSV--KAQVETF--
LDELSNYNAAKK-AGRGKQL-ICSIS-HSAYTV---TEQMESAVLFTPQV--YTN--KQ-ML---GG--SNA-KRN-------ISSIAGVEMMLRQILMNQT-Q------AVGKRFEDGKYRYLYF-YP---TYYFTP----E-TNKF---LQK-----AYNGI-AQTRF-----DT------SVR-----NHFI-S-------------------------------------------------KDLQA---------DLG------RDRY--QSVDSF-------L-IDENLQRDK------------------DR-------
TF---KL---SY---PED--------------------QPLTFYFMAL----------PP------G----R--DS--------TDT-------ESW--VMPTWLAFAFPMILDVKTVVSESPIPPFNDGVEFEES----VF--LDSAPHAFRALV-R--RD-RF--------RLD---YILEGW------NE----NGIQ-YLAPLNV-LTAAYAIH-LDV------NARQGKSGYDANWG-RFTELAKDF-ETSPLYVFSYLNRWVR---------NQG------------SET--A---RI----EKIR-----
------LYAYH--FYPCFDP-Y------------------------V-KYDTNMEQ--------L-IVG-EA--SS-LNH-PKKLTDLYRRFYR--A-NKR--YNP--KSNAVLKPVDIAAETI-LKAESSVFQ--G-ETLVAAVAAEVFKLMDRVH-----S---STAEG-----RWIM-S---------K---------REEE----RQAVLDFAKYFV---VEVFDNSFAGDRARLAGRQLNLI-----RDTCEFLYR------LEDDK--ENA--GKND-KSAEI--------D-NEND-------
'WP_114082632.1__2_' ---------------------------------------M----------SNDDWLSG-------DFGFDGDSS-------DRTI--ETKG-ELLTLKLLREA---IQ--SQNPD---------DK--VMAD--FGEYVL--PNLL-RI-AIGVTAK-----G---------GKF-FDE-------IDQ-------QRE-------AE---------GKI-------K---VR-RDNA-ADQSLNTHLLNGLFPANLIE--KR---------------------------LEKL--N-TT-
VQRVV---KERERRLVIAGFILHDFEK-----F----------------P-D--VPEN-----------------CRK-LP--------LAEHREIIDKKVHQL-GLD----NFI--NP--ENTEAYREYLDDLLCMAYN-AQR---RWDTNWNFSEF------GL-NP----LL----KD-RTLR--SLSDLTCLADSLASI------VKHPQDA-E-----HPR-----------LKEII-H-SL---SD-GQL----K-------FTYHSIAENRGVLTNVVNNALIQAH---TSLNTDE--Y------T-
Y---YEPLLYLPTGVIYLAS--RNAPA-I-SPE---DLPD--RVVNSI-----------------KSLCA--GQL--R-L-RQTGFGR-DG----------KG-MKYAEYYNLF-FDDIGLM----------------KVALDATL-----------------------RI-LN-----------------------P------NK---GSVAK---------------------------------------------SR-------SE---NLN---------KFQ------------QQNVLSA-------DY---DFKF------ED-----DIRI-----DQ--------IA-------------------
E----FG---DLVS--RK--IWE--------ETV--------NRVDY--------ARKK-DKK----LPAVPDFD---------LTHKVAEF-----WNLA------ECLP-QI--------------REIQRINESLKENKLKGNTGGVPYEWYYLAAKYL----E--H---HP-GI-E---DVRE----ACQQV-IEYLTTLISP-----------IIS----------QY----Q--L--P-DG-WDD-------------LRLWVKRVV----------MLP-----------------------GTNQEPSV--KTQVETF--
LNELDNYNAAKK-AGRGKQL-ICSIS-HSAYTV---TEQMESAVLFTPQV--YTN--KQ-ML---GG--SNA-KRN-------ISSIAGVEMMLRQILMNQT-Q------AVGKRFEDGKYRYLYF-YP---TYYFTP----E-TNKF---LQK-----AYNGI-AQTRF-----DT------SVR-----NHFI-S-------------------------------------------------KDLQA---------NLG------RDRY--QSVDSF-------L-IDENLQRDK------------------DR-------
TF---KL---SY---PED--------------------QPLTFYFMAL----------PP------G----R--DS--------TDT-------ESW--VMPTWLAFAFPMILDVKTVVSESPIPPFNDGAEFEES----VF--LDSAPHAFRALV-R--RD-RF--------RLD---YILEGW------NE----NGIQ-YPAPLNV-LTAAYAIH-LDV------NARQGKSGYDANWG-RFTELAKDF-ETSPLYVFSYLNRWVR---------NQG------------SET--A---RI----EKIR-----
-------LYAYH--FYPCFDP-Y------------------------V-KYDTNMEQ--------L-IVG-EA--SS-LNH-PKKLTDLYRRFYR--A-NKR--YNP--KSNAVLKPVDIAAETI-LKAESSVFQ--G-ETLVVAVAAEVFKLMDRVH-----S---STAEG-----RWIM-S---------K---------REEE----RQAVLDFAKYFV---VEVFDNAFAGDRARLAGRQLNLI-----RDTCEFLYR------LEDDK--ENA--GKNE-NIPVI--------D-DVKEPE---
-
'WP_038296476.1' ---------------------------------------M----------SNDDWLSG-------DFGFDGDSS-------DRTI--ETEG-ELLTLKLLREA---IQ--SQNPD---------DK--VMAD--FGEYVL--PNLL-RI-AIGVTAK-----G---------GKF-FDE-------IDQ-------QRE-------AE---------GKT-------K---VR-RDNA-ADQSLNTHLLNGLFPANLIE--KR---------------------------LEKL--N-TT-
VQRVV---KERERRLVIAGFILHDFEK-----F----------------P-D--VPEN-----------------CRK-LP--------LTEHREIIDKKVHQL-GLD----NFI--NP--ENPEAYREYLDDLLCMAYN-AQR---RWDTNWNFSEF------GL-NP----LL----KD-RTLR--SLSDLTCLADSLASI------VKHPQDA-E-----HPR-----------LKEII-H-SL---SD-GQL----K-------FTYHSIAENRGVLTNVVNNALIQAH---TSLNTDE--D------T-
Y---YEPLLYLPTGVIYLAS--RNAPA-I-SPE---DLPD--RVVNSI-----------------KSLCA--GQL--R-L-RQTGFGR-DG----------KG-MKYAEYYNLF-FDDIGLM----------------KVALDATL-----------------------RI-LN-----------------------P------NK---GSVAK---------------------------------------------SR-------SE---NLN---------KFQ------------QQKVLSA-------DY---DFKF------ED-----DIRI-----DQ--------IA-------------------
E----FG---DLVS--RK--IWE--------ETV--------NRVNS--------ARKK-DKK----LPAVPDFD---------LTHKVAEF-----WNLA------ECLP-QI--------------REIQRINESLKENKLKGNTGGVPYEWYYLAAKYL----E--H---HP-GI-E---DVRE----ACQQV-IDYLTTLISP-----------IIS----------QY----E--L--P-DG-WDD-------------LRLWVKRVV----------MLP-----------------------GTNQESSV--KTQVETF--
LNELDNYNAAKK-SGRGKQL-ICSIS-HSAYTV---TEQMESAVLFTPQV--YTN--KQ-ML---GG--SNA-KRN-------ISSIAGVEMMLRQILMNQT-Q------AVGKRFEDGKYRYLYF-YP---TYYFTP----E-TNKF---LQK-----AYNGI-AQTRF-----DT------SVR-----NHFI-S-------------------------------------------------KDLQA---------DLG------RDRY--QSVDSF-------L-IDENLPRDK------------------DR-------
TF---KL---SY---PED--------------------QPLTFYFMAL----------PP------G----R--DS--------TDT-------ESW--VMPTWLAFAFPMILDVKTVVSESPIPPFNDGAEFEES----VF--LDSAPHAFRALV-R--RD-RF--------RLD---YILEGW------NE----NGIQ-YPAPLNV-LTAAYAIH-LDV------NARQGKSGYDANWG-RFTELAKDF-ETSPLYVFSYLNRWVR---------NQG------------SET--A---RI----EKIR-----
-------LYAYH--FYPCFDP-Y------------------------V-KYDTNMEQ--------L-IVG-EV--SS-LNH-PKKLTDLYRTFYR--A-NKR--YNP--KSNAVLKPVDIAAETI-LKAESSVFQ--G-ETLVAAVAAEVFKLMDRVH-----S---STAEG-----RWIM-S---------K---------REEE----RQAVLDFAKYFV---VEVFDNAFAGDRARLAGRQLNLI-----RDTCEFLYR------LEDDK--ENA--GKNE-KSAET--------D-NENE-------
-
'WP_118166690.1' ---------------------------------------M----------SNDDWLSG-------DFGFDGDSS-------DRTI--ETEG-ELLTLKLLREA---IQ--AQNPD---------DK--VMAD--FGEYVL--PNLL-RI-AIGVTAK-----G---------GKF-FDE-------IDQ-------QRE-------AE---------GKT-------R---VR-RDNA-ADQSLNTHLLNGLFPANLIE--KR---------------------------LEKL--N-
TT-VQRVV---KERERRLVIAGFILHDFEK-----F----------------P-D--VPEN-----------------CRK-LP--------LAEHREIIDKKVHQL-GLD----NFI--NP--ENTDAYREYLDDLLCMAYN-AQR---RWDTNWNFSEF------GL-NP----LL----KD-RTLR--SLSDLTCLADSLASI------VKHPQDA-E-----HPR-----------LKEII-H-SL---SD-GQL----K-------FTYHSIAENRGVLTNVVNNALIQAH---TSLNTDE--H------
T-Y---YEPLLYLPTGVIYLAS--RNAPA-I-SPE---DLPD--RVVNSI-----------------KSLCA--GQL--R-L-RQTGFGR-DG----------KG-MKYAEYYNLF-FDDIGLM----------------KVALDATL-----------------------RI-LN-----------------------P------NK---GSVAK---------------------------------------------SR-------SE---NLN---------KFQ------------QQNVLSA-------GY---DFKF------ED-----DIRI-----DQ--------IA-------------------
-E----FG---DLIS--RK--IWE--------ETV--------NRVDS--------ARKK-DKK----LPAIPDFD---------LTHKVAEF-----WNLA------ECLP-QI--------------REIQRINESLKENKLKGNTGGVPYEWYYLAAKYL----E--H---HP-GI-E---DVRE----ACQQV-INYLTTLIYP-----------IIS----------QY----E--L--P-DG-WDD-------------LRLWVKRVV----------MLP-----------------------GTNQESSV--KTQVETF--
LNELDNYNAAKK-SGRGKQL-ICSIS-HSAYTV---TEQMESAVLFTPQV--YTN--KQ-ML---GG--SNA-KRN-------ISSIAGVEMMLRQILMNQT-Q------AVGKRFEDGKYRYLYF-YP---TYYFTP----E-TNKF---LQK-----AYNGI-AQTRF-----DT------SVR-----NHFI-S-------------------------------------------------KDLQA---------DLG------RDRY--QSVDSF-------L-IDENLQRDK------------------DR-------
TF---KL---SY---PED--------------------QPLTFYFMAL----------PP------G----R--DS--------TDT-------ESW--VMPTWLAFAFPMILDVKTVVSESPIPPFNDGAEFEES----VF--LDSAPHAFRALV-R--RD-RF--------RLD---YILEGW------NE----NGIQ-YPAPLNV-LTAAYAIH-LDV------NARQSGKSGYDANWG-RFTELAKDF-ETSPLYVFSYLNRWVR---------NQG------------SET--A---RI----EKIR-----
------LYAYH--FYPCFDP-Y------------------------V-KYDTNMEQ--------L-IVG-AA--SS-LNH-PKKLTDLYRRFYR--A-NKR--YNP--KSNAVLKPVDIAAETI-LKAESSVFQ--G-ETLVVAVAAEVFKLMDRVH-----S---STAEG-----RWIM-S---------K---------REEE----RQAVLDFAKYFV---VEVFDNAFAGDRARLAGRQLNLI-----RDTCEFLYR------LEDDK--ENA--SKNE-KSAEI--------D-NEND-------
'TAE22780.1__3_' MAKK--RQKPDQEYQQLSLNFD------------IPVNLNSERSD---DADDDWLQD---E---ELGANFETS-------DRTI--EPQTRDLLTLKLLREA---IQ--SQNPG---------DR--VMGD--FAEYVL--PNLL-RV-AIGVTAK-----G---------GKF-FDE-------LDR-------KQ------------------GKE-------N---VR-RDNA-GDQSLNTHLLNGLFPANLIE--QR-------------
----------------LEKL--D-TT-VRRSV---GEKERRLVIAGFILHDFEK-----F----------------P-D--VPND-----------------CRK-LP--------LEKHRQIIDEKVRQL-GLD----RFI--NP--DDPDAYREYIDDLLCMAYN-AQR---RWDTNWNFSEF------GL-NP----VL----KD-RTLR--SLSDLTCLADSLASI------IKHPQDA-E-----NTR-----------LNELI-H-SL---SD-GQL----K-------
FTYHSIAENRGVLTNVVNNALMEVH---TALNTDD--C------T-Y---YEPLLYLPTGVIYLAR--SDAPA-I-SIA---DLPE--RVVSKI-----------------KQLCA--GQL--Q-R-RQTGFGR-DG----------KG-MKYADYYNLF-FDASELM----------------LVAVDATL-----------------------RI-LK-----------------------I------GK---NSVAK---------------------------------------------SR-------SD---NLI---------KFQ------------RQDVLSA-------
DY---NFKF------DD-----DIRI-----DQ--------IA---------------------E----LG---DLVS--RK--IWG--------EQV--------NTIEA--------ARKK-DKK----LPALTHLD---------IVDEIAKF-----WNLS------EYLP-AL--------------REIQRINERLKELKLKGNTGGVPYEWYYLAAKYL----E--H---HP-GI-E---DVRE----VYEQA-IAHISQMIKP-----------ILS----------QY----K--L--P-DG-WDD-------------LRLWVTRVV---------
MLP------------------------GNTPLDTA--SSNV--F--LDELERYQLAKK-QGRGRQL-ICSIS-HSAYTV---TEQMESAVLFTPQV--YTN--KQ-ML---GG--SNA-KRN-------ISSIAGLEMMLRQILMNQT-Q------AVGKKFEDGKYRYLYF-YP---TYYFTP----E-TNKF---LQK-----AYTSI-AQTRF-----DT------SIR-----NHFI-S-------------------------------------------------KDLQA---------DFG------RDRY--
QSIDAF-------L-IDENIEPGK------------------DR-------TF---KL---SY---PED--------------------QPLTFYFMAL----------PP------G----R--DA--------TDT-------ESW--VMPTWLAFAFPMILDVKTVVSESPIPPFNDGAEFEES----VF--LDSVPQAFRVLT-K--RD-RF--------RLD---YILEGW------EE----NGKK-YPAPLNA-LTAAYAIH-LDV------NAKQGKSGYDANWG-KLSELAKDF-
ETSLLYVFAYLNKWVR---------KQK------------LDA--P---SL----SKIK---------LYAHS--FYPCFDP-Y------------------------V-TYKPESED--------L-IVE-AE--SP-LNH-PKKLTELYRKFYR--A-KSTPGKPT-------KSNAILKPIDEAADVI-LKAYKE--E--G-FNLVQTVAAKIFKLMERVH-----N---NSTAEG-----RWVI-A---------K---------RDEE----REAIMDFANYFV---VEVFEKSFAGDRARLAGRQLNLI-----
RDTCEFLYR------LEGDK--KYR--ARKE-QASEN-----DLEDS-DSDNDD-----
'WP_104546987.1__2_' MAKKLKDTDPDEQYKQLSL-FE------------SENDLTSEQFDN--SDDDDWLQD-------DSGFDFEPS-------DRTV--EPQSHELLTLKLLREA---IQ--TQNPN---------DR--VMSD--FAEYVL--PNLL-RV-AIGVTAK-----G---------GKF-FDE-------LDQ-------QR------------------GRE-------N---GR-RNHA-ADQSLNTHLLNGLFPANLIE--QR-------------
----------------LEKL--D-TT-VRRVV---QERERRIVIAGFILHDFEK-----F----------------P-D--VPNE-----------------CRK-LP--------LEQHRQIINETVRQL-GLN----HFI--NP--DDPEAYREYLDDLLCVAYN-AQR---RWDTNWNFSKF------GL-NP----VL----RD-RPLG--CLSDLSCLADLLASI------IKHPQDA-E-----NTR-----------LKELI-H-SL---SN-GQL----K-------
FTYHSIAENRGVLTNVLNNALMEAH---ISLNTQD--C------T-Y---YEPLLYLPTGVIYLAR--RDAPC-V-RIA---DLPE--RVVKKI-----------------KELCA--SQL--R-L-RQTGFGR-DG----------KG-IKYADYYNLF-FDATSLM----------------QVALSATL-----------------------RI-LN-----------------------T------NK---SSVAN---------------------------------------------SR-------SK---NLV---------KFQ------------QQNILSV-------
GY---DFNF------KD-----DIRI-----DQ--------IA---------------------E----FG---DLIS--RK--IWG--------EQV--------NRIEN--------ERKK-NKK----LPELPYLD---------IVEEIAKF-----WNLY------EYLP-AI--------------REIQRINESLKEHKLKGNTGGVPYEWYYLAAKYL----E--H---HP-GV-E---DVRK----ICEEA-IAHVSQLIQP-----------IIT----------KY----Q--L--P-DG-WDD-------------LRLWVTRVV---------MLP--
---------------GTATP-TA--STEV--F--LQELSSYQLAKK-SGRGRQL-ICSIS-HSPYTV---TEQMESAVLFTPQV--YTN--KQ-ML---GG--SNA-KRN-------ISSIAGIEMMLRQILMNQT-Q------AVGKKFEDGKYRYLYF-YP---TYYFTP----E-TNKF---LQK-----AYSSI-AQTRF-----DT------SIR-----NHFI-S-------------------------------------------------KDLQA---------NLG------RDRY--QSVDAF--
-L-MNENIEPGK------------------DR-------TF---KL---SY---PED--------------------QPLTFYFMAL------PPEKPP-------G----K--DP-----EKKITNT------ASW--IMPSWLAFAFPMILDVKTVVSESPIPPFNDGAEFEES----VF--LDSAPQAFRVLT-K--RD-RF--------RLD---YILEGW------QE----NGKS-YSAPLNV-LTAAYAIH-LDV------NARQSESGYDSNWS-KFAELAKDF-ETSPLYVFNYLNRWVR--
-------NQD------------SET--A---RI----EKIK---------LYAYH--FYPCFDP-Y------------------------A-KYNPSLEE--------L-SVL-DEQKSS-LNH-PKKLTELYRRFYR--A-NKQ--YNP--KANAVLKPIDIASKTI-LEADPS-FQ--G-EALVAAVAAEVFKLMERVH-----A---STAEG-----RWIM-S---------R---------REEE----REAILEFARYFV---VEVFEKSFAGDRARLAGRQINLI-----KDTCEFLYR------LEDDK-
-ENR--TINN-NKPTN----SFQNK-AEE--------
'WP_105218136.1' ---------MGEPDSPLLQ----------GKGLGERLTEQ--DSDDDWLQD---E---DLDSDFEPS------NRTI--EPQSRELLTKLLREA---IQ--TQNPH--------DR--VMSD--FAEYVL--PNLL-RV-AIGVTAK---G--------GKF-FDE------LDR------KNE------AE---------GKD------R---VR-RDNA-GDQSLNTHLLNGLFPANLIE--QR---------
-----------LEKL--N-TT-VRRVV---RERERRIVIAGFILHDFEK-----F----------------P-D--VPAD-----------------CRK-LP----------LEQHRQIIDEKVRQL-GLD----RFI--NP--DDSDAYREYLDDLLCVAYN-AQR---RWDTNWNFSEF------GL-NP----VL----KD-RTLR--ALSDLTCLADSLASI------IKHPQDA-E------NTR-------------LNELI-H-SL---SD-GKL----K--------FTYHSIAENRGVLTNVVNNALMEAH---
SVLNTQN-C------A-Y---YEPLLYLPTGVIYLAQ--RDAPS-V-QVE---DLPE--RVVNKI------------KQFCA--SQL-R-L-RQTGFGR-DG------------KG-MKYAEYYNLF-FDAPDLM-----------------AVALDATL------------------RI-LR-------------------T-------GK---SSVAK---------------------------------------------SR------SD---NLI---------KFQ----------QQNLVSA--------DY---NFEF------TD-----DIRI-----DQ------
----IA-------------------E-----FG---DLIS--RK--IWG---------EQV------KQIEA--------ERKK-DKK----LPTIPHLD---------IVEEIAKF-----WNLS-------EYLP-AL------------REIQRINESLKEHKLKGNTGGVPYEWYYLAAKYL----E--K---HP-GI-E---DVRK-----ICQDA-IAYISQLIQP-----------TVV----------QY----K--L--P-DG-WDD------------LRLWVTRVV---------MLP-----------GKATSVTT---
SAEV--F--LQELSGYQAKK-PGRGQKL-ICSIS-HSPYTV---TEQMESAVLFTPQV--YTN--KQ-ML---GG--SNA-KRN---------ISSIAGIEMMLRQILMNQT-Q------AVGKKFEDGKYRYLYF-YP---TYYFTP----E-TNKF---LQK-----AYNGI-AQTRF------DT-----SIR------NHFI-S------------------------------------------------------KDLQA----------NLE------RDRY--QSVDAF-------L-INENIESGK--------
-----DR--------TF---KL----SY---PED--------------------QPLTFYFMAL----------PP-----G----R--DS-----------TDT-----ESW--VMPTWLAFAFPMILDVKTVVSESPIPPFNDGAEFEES-----VF--LDSAPQAFRVLT-K--RD-RF--------RLD---YILEGW------QE------NGKS-YSAPLNL-LTAAYAIH-LDV------NARQGKSGYDSNWG-RFTELAKDF-ETSPLYVFSYLNRWVR--------NQG-------------VET--A---RI----
-EKIK----------LYAHH--FYPCFDP-Y-----------------------A-KFNPEMEE---------L-IVG-EE--SS-LNH-PKKLTELYRQFYR--A-NKR--YNP--KSNAVLKPIDIASKTI-LKAESFFQ--G-EALVAAVAEVFKLMERVH------A---STAEG------HWIF-K---------N---------REEE-----RQAVLDFARYFV--VEVFEKSFAGDRARLAGRQLNLI------KDTCEFLYR------LEDDR--ENR--ARNS-DQSTD-------NSPNE-
AEE--------
'TEU17163.1' ------------------------------MTMTDDEQRVQ--E---DYLLEDEEEG----AAEQET--ISLAPEPLFSALLRDV---IV--KLWPD-------DT--VMAD--FVDHVA--APLS-DY-LGHRGAK-----G---GEF-VEE---------R---------RAK--------------GLQ---------V---DE--RYI-YDQTMRAHLVNGLLFPLHVA--RT----
LQAWG-------APQFRA-YDDSVRRLFIAGYVLHDYLK----E--P--VEKQLEEAGFSH-------EQAIGPAG---------LSVTEQIFRQWICATL-GLE-----EFL--AP-----VGGVEEVLHDLIYIAN-TQT---RWGTLRNLSVL---P--RL---------TL---PA-VQRD--LAEGLRLSRDLAYLAYI-----ARTPQEA-A------TNAS-----------IQREL-T-TL---SN-HIA---R----------FTYHHLADNRGVLSNFIHNAALSAM---
----SD--E---------H--RVPLLYAPSGVVYLEH--KEAPP-PPAVA----EIAE--ATIQNI------------QKAIV--RRL-H-R-SLDGIKR-DG------------KG-LKYADYYYLF-FDLPTFV------------------KLGVQAAF------------------GN-IH------------------E-------K---GK---KPSSG------------------------------------SR--------SE---NLS-----------KMK----------DKGWLDP--------EV---DLDL-----PD---DVRV------DQ--------
-IA----------------E----WC---YLAE---KQ-I----------------AEKLPDFD----------VAGFLLNE------MSLQ------DLRE-PF---------EAASRIK--------------RAGGVAYYWWFAAGHYL----E--K---HP-GL-DPA-AWQE----QVEGF-ARQLVQAVRE----------------AAQ---------SADVPPP------ESGADWDD--------------LRAYISRTL-------------TLG------------------------------AALVAGEDRTVF-----
AAELHRYQNAKR-RGRGTTH-MCALC-SSTRVV---DKQADPAVLFAPQV--YSN--KL-QL---GG--SQL-RRD--------ICSICGLEMMLRQNLMNRS-A-------ATGKDFENRRVRYLYF-YP---TYYFTP-----E-TLDL---FRR------AYTNI-RRISF----------T--------ELR-----RQLV-G----------------------------------------------------EEGTV---------DLS-------PETL--QRLEPL-------LLTPEEEMEPEE------------DR---------

[illegible]

'RRR68494.1' [illegible]

'WP_129679138.1' [illegible]

'WP_044199283.1' [illegible]

'WP_029215466.1' [illegible]

'WP_129632988.1' [illegible]

'WP_097645710.1' [illegible]

'WP_087711939.1' [illegible]

'WP_012599091.1_2_' [illegible]

'WP_011153680.1_3_' [illegible]

'SMQ86523.1_3_' [illegible]

'WP_017720464.1' [illegible]

'WP_080810414.1' [illegible]

'ADN17499.1_2_' [illegible]

'WP_046279385.1_2_' [illegible]

'WP_012627888.1_2_' [illegible]

'WP_015783166.1_2_' [illegible]

'WP_012593877.1_2_' [illegible]

'ELR97243.1_2_' [illegible]

'WP_035152549.1_2_' [illegible]

'KYC42933.1_2_' [illegible]

'WP_096727546.1_2_' [illegible]

'WP_053457730.1_2_' [illegible]

'PHJ60297.1_14_' [illegible]

'AFY31721.1_2_' [illegible]

'KST66746.1_2_' [illegible]

'WP_088240873.1' [illegible]

'WP_036842433.1' [illegible]

'CUR14078.1' [illegible]

'WP_043938760.1' [illegible]

'WP_043937175.1' [illegible]

'WP_040957323.1' [illegible]

'WP_036834484.1' [illegible]

'WP_042156684.1' [illegible]

'BBD53976.1' [illegible]

'WP_088889829.1' [illegible]

'WP_036011813.1' [illegible]

'OBQ22644.1__3_' [illegible]

'AUT00479.1__2_' [illegible]

'WP_096680032.1__2_' [illegible]

'WP_096656782.1__2_' [illegible]

'WP_018398404.1' [illegible]

'WP_017659639.1' [illegible]

'AFY84409.1__2_' [illegible]

'RMH78832.1' [illegible]

;
end;
begin trees;

tree consensus = [&U]

[illegible]

end;

### Cas8b1 (Cst1)

#NEXUS

begin characters;

dimensions nchar=1198;

format datatype=protein missing=? gap=-;

matrix

ADI02112 ---------------------MPKGVEQGKGGEVEFMPHV--HW----TGNPFVDAGLAALAAA------AR-VST-----------------------------------------------------LE---ELGAEHIEQAVKE-----------LERVLLSDQALGIGVDKAFSGPKK------------------------ALRQVFPN-------------------------------SELDNPSNWKGRTPKEKEDNARRK---------
FQDALRSELNKAKQCL--ELEEAG-----------------------ETCFACGE-RRPKIAMET-R-----RK--DK------M--PLFE-----GIVNFYPGF-SHGVRVCG------------------------------------------------------------------------LCSLAVRFLPLSTM---------------LAGNQNRLWFLHSQHLGLAATVARRYGWEHFDRLITGNKALDFYGSW-----E--TA-GEAGTVVYLFCELLNSFPDQLRA-----I----Y--------
RS-G--L----------PSTAYVF----SNDNR----GGFVQAIH---VPNE-----LLA--FLA----------KLQIES--KNAFKDFWEGLLKVRGDLR-EDER-------KTRAGFVRSISNRMIA-GENI--VGMCLTSDPPN--LQGGWLGHRL--YLQEV---RGMAPEK-------------LA--ILERLG----I-----------SIAASEDAKKLVT------------ELVTAQYH-QLYGLLLGY----VRKGWLSHQE-FYFLL------PPN--------------AHKV-
AGE--------IRDVLLAVV-------YEWQ---------HCQ-----------QEGKEFPRSTF------DKAEASPDETLRRIQEIGRRLTNNL--------------------------------PNLQRWTKELT-SARYAERIRGVYL-NAVRHGAV---------------GFEDY-VFLAPLQ----D--QRA---MW-LLRDYLLAFLFDVGRV--ELPED-------EY----PEFG-A---DE---VREIETF---------

WP_132779593 ------------------------------MDDNNGKKNNT------------KYETKEDGTTTI---------------------------------------------------------------------------NNKEVFSKIKVEE------------------------------------------------------------------------------------------------NYFENGKVINLKGKNQY-----------------------------------------------------------------YPNFID------
-VG--GN------QK-------------EVFKEF-IAAFSDLN---------------------------------------------------------------------KTEQY-------------NCELCESGLLVKKDNLDNGIDNDQKEKFFSKISTLNMAHNK--ILGPSEK-----FPNSY---------------W-----N--FN-SG--LKICHFCTFLLIHHHLAFT---------KL--------SDGS-----------------EIF----INAP------SFKLMYE---LNKL-----VRELFGKT-------------------
DVDSTKKREILAMSVIEYT-RRLQ-------TTLGQWNAM-NIEIVIKKGDE---------------IDFYTLPYETVNIISDR-----AIASI-------------LS--DIGEF----------------------------------------------------AIFDSVISG--------------------------------------------------------------NYSF---------------------------------------------------------------------------------------------------LQEISYKIIRASFKNESISAFIK-----MYKN----
-------------QNNSI-LTAQKIL--------------K-LYATIIDRRRLYARTK-----------------------------------------------

```
RKX65388 ------------------------MSDMEHKIKLFPAS---W--------LYNAGVIGFLRV----------------------------------------L--AYGMEEEK-----IE---SWLKDDGTVEVDR----------GVFRKELGGQKLPICMKYLVEFLVKDEDLEAWKEKKD----------------------------------------NKGKKNKDKYADFAKDMGDFGYKFI------------------------
------------------------------------RAGNKLFAS-------QTPYQN------LV-QFKE---------WQKFEFAKLIAGL---------------------------------------------------ENYKFSCS--DI------SCSLCGNRAVKRP---------HPKSKLENRLFVFQEPHLR--ILAPSKG--E--FPNSF---------------W-----N--LN-TS--FYLCPFCVYLLIHHHIPFI----------K--------TQHG-----------------DIF----INAP------
SFKIMWY---LNKL-----AKE-------------------FLQ-RGKEFTLREILGTSLMELT-QRIY-------LTLGVWSLM-QIEMVI-KRYN-TK------TKNTE-VAYYSLPFNTARILLNR----------------------NV-ANLVSAT----------------------------------------------QEPYILQCVLNE---------------------------------------------------------DFEALLFLSHCILGVL---------
--------------------------QGRKDQSLEGKIKNQNRSHLST-------LA---------------QILPT-LY-----------------------VKINSLIKGA----------------------------------------------

WP_028841375 ---------------------------MPDGSITLYPSN---W--------LYNAGVIGFLRV------IA-EAR-----------------------------------------GESVVENW-----IQ---ENGTVTIDVEIFR-----------NAMVGDKEIPIPKCMKYLVEYLVNDEDVKEWLYKKD---------------------------------------------------------KKGITNNEK-------------YKDIANQLGDFG-
YKFIRAG------------------------NMLFASNM-PYQNLVQLS-------EW--SS--------------------LEFARL-VSNLIKYP-----------------------------------------------------------KIKSN--I-------TCGLCSNNKVLNP---------KVDSKLENRLFVFQEPHMR--LLGPSIG--K--FPNAF---------------W-----N--FN-HS--ILICPLCAYLIIHHHVTFE-----S--ART--------QN-G------------------QIF----INAP------
SFKLMWY---LNKY-----ASE--ILN-------------------KRKTYQLKEILGISFIELA-QKVF-------TTLGAWSMI-NIEMVI-KNKD--S------------IDYYSLPYEVSKVILNK-----EVASL-------------IS--QTNEEF----I------------------------------------------LE-IILDGKFDE------LLTLNQKI-LRYAI------------------------TNSG-AYD---------------D-------KYLS---------KLR-----------NK-------------------------
DAYSLKKLSTILPELY-----------------------------------------------------------------------VKIDN-ILKRGEL---------------------------------------------------------------------

PJJ76158 ---------------------------MTNQNTLQLYPSN---W--------LYNAGVIGFLRI---------------------------------------------------LE---PINKKSVEEFINN-------------KTESIISIPPNSEDMLLNKWKE-------------------------------------------------------------LSIESKKNTSPLYGRKGY---------------------------------------------------------
YANQTDNS-IKNKIALLT--------RK--ET------SE-N-----------KNKNKT-RKSLPEY-------------------------------------------------------------SCIFCTNKVKVG-----------------KTNATFLNQSFGKLLLGSEKT-----FRNMY---------------W-----N-----NEAHDFVCEKCEFIIICHHLAFT-----N----L--------SD-K--S--------------EIF----INAP------SFQIMYY---LNKF-----TQE--IFGSSP-------------
---IQDARSKREILAMSVIEYA-TKIQ-------ANLASWVGM-NIEIIS-RHGD--K------------IEFFSLPYEVTKLLADR-----RIAAI-------------LS--QIGEYS----I-----------------------------------------LN-LVLDQDFSS------LMEMGYRL-LRIGL-------------------------------------------------------------------------------IPKQY--------------------------------------------------RGKNEKDFVN-
QILQLEKN----------------RWDPA-LVAQQIFQLCAY--IHE------KHKTNAYGYHKH-----------------------------------------------------------

WP_095416655 ------------------------------MCSLTLYPSN---W--------LYNAGVIGLLRV------LE-RTGAD------------------------------------------PLRCLSEA----------AVQIRFSILCSSV---------------------------KDVELEQDDKEFTIPLLARE-------------------------------------------------------WLLESWETLTEKKEKTE-------------CGKIKEVWGKLF-NTWYRGF-
----------------------------------FNADTRYLF-K-----ES--KK------TK-PLII-----QFGDFVRSF-GHPWGNGD--------------------------------------------------------------TCSFCFRNSRF------------------SYKNIFTSEHSK--LLGASAG--ERGVPNSF---------------W-----N--LN-PEQSLRICDFCNFVLLNHHLALT-----R----L--------SD-G--S--------------EIF----INAP------SFQVMWY---LNKF-----
ARE--VLGPAS-----------------SEEMRSKRNILAMSVIEYA-TKIQ-------ATLGVWTGM-NIEVVS-KRGG--E------------IEFFSLPYEVIQLLADR-----RIASL-------------LS--QIGEFS----V-----------------------------------------LN-LVLAQDFSR------LLELGYRL-LRI------------------------------------------------------------------------------------------------GLKTVVEWGKSERDFV-N------------------------
---------------------QTLRLEKNKQHPA------------------------RVADQ-LFQLSAL----I--EEK--------RQRRETYERSCIF-------------------------------------------------------

PMP93669 ---------------------------MSDNSIILYPSN---W--------LYNAGVIGLLLV---------------------------------------------------RE----KAGE--AVEKDLMS------DGSCQIPDDYFAPLQIN------------------TRRIPKAIINLVNELVSNEDLKEWLNNDN-------------------------------------------------------------QTKYEKYYTELGDFGFKFI-----------------------------------------------
--------------------------RAGNKLFAS-------KTPYQN------LV-QLSE-----WQSYEYAEL-VREIPRLI----------------------------------------------------------AVDTGS------------PCNLCGRYAILVQ---------TPQSKLQQRLCKLQSTHLK--DLGPSIG--E--FPNGF---------------W-----G--LS-QS--MGICYLCSFLIIHHHIALI----------RL---------SDGS-------------------EIF----INAP------SFQVMYY---LNKF-----
ARQ------------------VFSVLS-FEEMRSKRNILAISIIEYA-TKIQ-------TTLGMWTGM-NIEIVS---------------RHGSQ-VDFFSLPYNVIQLISDR-----RIAGL-------------LS--QIGEFS----I-----------------------------------------LN-HVLDEDFSQ------LVEIGYRL-LRI-----------------------------------------------------------------------------------------GLKRYSERGKAENDYV-------------------------------
-----------NRML----HLNENKNNPA-------------------------RVAEQ-VFKLCAL-------IEE----K-QKRRENYEYIRNTRG-------------------------------------------------------

WP_031505422 ------------------------------MEYIELRPKN---W--------FYNLSVTGFLEV----------------------------------------------LADQNSEE-----IE---KLIRDDGTVLIDR-----------KIFAKHRELDIPEALVRYVNYLVKGENLDEWLEEKK-------------------------------------------------------KNDQKTNKEKYKKYHDMFGEFGYKLA----------------------------------
---------------------------RSFNKL-FRSNTPYQN-------LV--QE--------------------NEWSHF-IELVKNLE---------------------------------------------------------KIS--EKKDTE------------KCEICGGKWYLDL----------DKATEFTRRLFRFASAHSS--EFGSSVG--D--FPNAF---------------W-----N-----NNSSLLVCPLCVYLIIHSHVAWT-----R----L--------SD-S--T--------------SIF----INAP------SFKVMWH---
LNKY-----AKE--LYG-------------------AGKVEGVKELFGMSLIELA-LKLN-------LQLGKWTMM-NIEVVI-KYKD--K-------KGKDK-VEFFSIPHEITMLLLDK-----SVASL-------------LF--DIGRT-------------------------------------------------------EVLSWFLDG------------------------------------------------------------------------------------------------------------------EFDKILRDGERHFR------------------------------
--------------------EALKSSDAKTL--------------------------SYSQK-LFELYAL-------VNE--------KRKGGAL--------------------------------------------------------------

WP_011915919 ------------------------------MDKLILYPGN---W--------LYNASVIGFLNV------I-------------------------------------------------SRRLGEDV-----IE---SWLNEDGGLTIDK---------AIFDPVKIVKEDIPKSLVYYVEYLTEHEDIEEWLGKNS------------------------------------------------------NSEKLKEYYDRMGEFGYKLV-----------------------------------
-------------------RAFNKL-FSKNMPYQN-------LV--QQ------------------GDRKEF-ITFLLNLF---------------------------------------------------------------------------SVT--NNKITY-----------RCDFCGIERAIEP----------RETSKLEKRLFKFDLMHSS--LLGASAG--E--FPNSF---------------W-----N-----NSASLRICPICSYFIIHHHLALT-----Q----L--------ED-G--W--------------EIF----INAP------SFQVMWY---LNKY---
IKAIN--EKV-----------------------KTDRVRRLLGMSVMEMA-LKTN-------IQLGKWTMM-NIEVVS-KLKI--RIKGK--KEYYK-IGFYSLPHEVISLLLNR-----KISSI-------------LT--EIGETS----I------------------------------------------LN-LVLSQNYER------IMEIGEKV-FKIAL-----------------------------------------------------K-----------------------------------------------KLSKESNKAKNDKSDE-----------------------
-----------DDKFIAHYFQLV------KEKSDLV--------------------------KISQK-LFELYLL-------IME--------KTKKEALI--------------------------------------------------------------

WP_013431324 ------------------------------MDKLILYPGN---W--------LYNASVIGFLSV----------------------------------------------ISRELGEDV-----VE---SWLNEDGSLAIDK---------AIFEPVKIGKEDIPKSLVYYVEYLTENDDIEEWLEKKN-----------------------------------------------------------KKNDEKKEQSNNEKHKEYFAAMGRFGYKLI--------------------------------
-------------------------------KAFDKL-FFNEMLYRN-----------LV------QQ------------GDRKEF-ITILLNLF-------------------------------------------------------------SITNNRVTH---------RCDFCGIERAVEP----------DENSKLEKRLFKFDLMHSS--LLGSSVG--K--FPNSF---------------W-----N-----NSTSLRICPICLYFIIHHHLALT----------KL--------EDGS-----------------EIF----INAQ------SFQVMWY---
LNKY-----IKAVNE-------------------KAKTADVRRLLGMSVMEMA-LKTN-------IQLGKWTMM-NIEAVSKLMFK-------IKGKPKYKIDFYSLPHEVISLLLNR-----KISSI-------------LS--EIGETS-----------------------------------------------ILNLVLNQ-------------------------------------------------------------------------------------------------------------------------NYGRIMEIGEKVFKIA--------------------------
------LKKLSGESNKG--NDDESDEDDKFIA-HYFRLAKE-----------------KNDLI-KISQNLF-----------------------ELYLLIMEKQ----------------------KRRL-----------------------------

PNV80237 ------------------------------MKESIRLYPGN---W--------LYNAGVIGFLEV------------------------------------------------ITYGECN---NVVE---EWLKNDGSVEIEK----------RVFDFKRLGNREVPFCLYYYVEFLTKDEDIENWRKRTD--------------------------------------------------------KKNISYEEKYKQIAEEFGDVL--------------------WGYKYV--
------------------------------------RAMNKL-FYSDMPYQN-------LF---Q------KG-DWSE--------NSLSNF-INKIPESI-------------------------------------------------------LKPGR-------------ICGICNNFYSSSF---------PYLSEKTVKRIEKFSEMYVHKLNIGSSLA--E--FPNSF---------------W-----N-----NSQSLNVCPLCAYLIIHHHIALT----------SL--------SDKS------------------GIF----VNAP------SFKVMWY-
--LNRY-----MRE------------------VYG-KEKIGRVRELLGISIIEMA-LRLN-------IQLGKWTMM-NIEVVS-KYKE----------GDNDK-IDFFNLPYEAVLLLSDR-----DIASL-------------LN--EIGELN-----------------------------------------------ILDLILDG----------------------------------------------------------------------KYKE----------ILD--------------------------------------------LAERLFKIGLKHKEDR-----------------------
--------------GKQENEFINNKVKLTKNRENLI---------------------------SYSQK-LFKLYAL-------IEE------KIRKEVYV-------------------------------------------------------

TYP58398 ------------------------------MENDVIVLYPGN---W--------LYNAGVIGFLEV-------------------------------------------------M--AYGEDSKE-----VE---KWFKDDGSVEIKK----------RVFDFRKVGNKFLPLCLYYYVELLTKDEDIHEWIKRTD-------------------------------------------------------KNKVSYEQKYKDIAKEFGGEW--------------------GYKYV-
------------------------------------RAMNKL-FFSNMPYQN-------LF--QK------AD----------WLSDSLINF-IRKIPEII-----------------------------------------------------DIKKDV-------------LCGLCNNFYHSEF---------PYLSEQTINRIMKFSEMHVH--NLKIGPSIFQ--FPNGF---------------W-----N-----NSQSFYICPLCTYLIIHHHIALV-----E----L--------SDKS-------------------KIF----VNAP------SFKVMWH---
LNRY-----VKE--IYG-------------------REKIEVIRELLGMSIIELS-LKFH-------IQLGKWTMM-NIEVVN-QYRE----------GDNDK-IEFFSLPYETILLLLDR-----NIASL-------------IN--DVGEIN-----------------------------------------------VLNLVLDG-------------------------------------------------------------------KYRE----------ILD------------------------------------------FGERVFKVGIKNKNDW-----------------------
--------GKQESDFINKKI----KLDKNKNNLV---------------------------TFSQK-LFKLYAL-------IEE----------KIGKGVHV-------------------------------------------------------

CUT04172 ------------------------------MEFIPSN---W--------LYNAGVIGFLRV------LE-------------------------------------------------SAGENKVN-----VE---NFLKEDGGVEIDD-----------FDK-------------KFLKEVKKVDNYEIPKLAYY------------------------------------------------------------YLIFNFNEIVNGEASSD-------------DEKIKKVWGKLF-NTYYRGF------
-------------------------------FNANTNYLF-------QH--SK------RS-PALI--------QQFSDF-VRNMTNSN----------------------------------------------------------------KNS--SI---------------KCSFCLKSDYNFF-----------------YKNKFTSEHNK--ILGASIG--E--VPNTF---------------W-----N--LD-KKNSLSICDFCSYILLHYHLGII-----K----L--------SN-N--S--------------EIF----INAP------SFKIMWH---LNNH-----ART--
LYE---------------------KQRAKEVKEILGMSLIEMT-SRLQ-------LQLSKWTKM-NIEVIV-KYKV--KVNE---KWEDK-IDFFSLPYEVVDVLSDR-----EVASL-------------LN--KIGEFK----I------------------------------------------LN-MVLDGKFNE------ILKFGERV-FRIAI-------------------------------------------------------KPRN---------------------------------------------------EWGKQERDFI-N--------------------------
------------------ENIKLERNNSNLI-----------------------SFSQN-IFKLYAL-------IND------KIKKEVLIWT----------------------------------------------------

WP_068747788 ---------------------------MEKDDVITLYPGN---W--------LYNAGVIGLLRS-------------------------------------------------VEEVEK--------LP-----LDCCFNFYDDG---------------------------------------------------------------------------------------SLSVKKIIFNELDVERRYFSEEEK-------------ISSIVGKSNLYR-------------------------------
---NYLQPSWIS----------------------------------SFPYF-VKNLSKIK------------------------------------------------------------ENDVS--F--------YCNLCYRRFALPN-SEVEDLKSRGLKKFLESIKKFDIRHNA--ILGPSLG--E--FPNSF---------------W-----N-----NNQSLYICPLCAFLIIHHHLALT-----R----L---------RN-G--S--------------EIF----INTP------SFEIMWH---LNKY-----ARQ--IFE--------------------
KENVATTKELLGMSLIEMV-LKLN-------IQLGKWTIM-NIEVVS-KYKI--D------NKDR--IDFFSLPYEIVYLLSDR-----EIASL-------------LN--DIGEFN----V------------------------------------------LN-MVLNSNFKG------ILEFAERI-FRIAL--------------------------------------------------------KSER-------------------------------------------------------------EKQENKFIND---
NVKLERNKGNNLI--------------------------AFSQK-LFKLYTL----I--EEK------VIRRFLYE-----------------------------------------------------------

WP_144399156 ----------------------------------MLDELYLID----------NFDNLSLPRLFVF------------------------------------------------ILEETAR-----IE---GDIRKAYNYFKGN----------------------------------------------------------------------------------------------------VGYYENYVAQVKEKQ------------------------------------------------------------FMENLSKI--
-------------------------------------VHYIFRKP---------------------------------------------------SLEGTD------------RCFFCGNSLKTDEDRSI------------LQRKQFSRILFK--DFSSSEG--K--FPNSF---------------W-----N--MS-NE--FYLCDVCSQMALFRHFVFP-------------------KSGE-----------------SIF----VNVP------SFKAIWY---LNEA-----IKAYRDLS--------------------LELSVSRAI----SEVH-
LKLQ-------RLLGIWERQ-NVEVIY-YNSK-------------S-YRIYYMPQRVINLLLNT-----RISSI-------------LN--SLNSIK----I------------------------------------------FEAVIYQERLNE------ILECEYLL-LKYLFDVSSSGRSH----LAENVKQYVGDVKRNAGYY--------MNTLPDLFQ--------------------EVR-----SVMGVKQMV-----------------------PVWRMRELGKEASQLF-----------------------------------------
ERTRYRLL-ELIRLAKKDEVYH-LILKTYMA----NN----------KPFPE-DLNRVFS-------AKD----E-DFKNLMFAYVSGMISG-----------------G-----EENE---------------------------

WP_148186878 ---------------------------MMEEIVKREGS--------------------IEKA----------------------------------------------------------YNYFKGNI----------GYYENYIANVSLT--------------------NKEIDELLPLVSNEQQREILLQVKEKKIK--------------------------------------------------------IPQIKKKSKELLSKFDDRD-----------------------------------------------------
---LEELRNFLL-------QSEKEK------FR----------VYLSEI-IRSIFRKP-------------------------------------------------------DLSGSE-------------RCFLCGNPLRLSENTSI------------RQRKQFSRILFK--EFSSSEG--E--FPNSF---------------W-----N--IN-NE--YHLCDLCSQFALFRHFVFP-----------K---------NSKK-----------------EIF----VNVP------SFKAIWY---LNEA-----IKAYKD---------------------
LNIELSVSRAV----AEVH-MKLQ-------RLLGVWERQ-NIEIVN-YDGK-------------S-YSVYYVPQRTVDLLLNT-----RISSI-------------LN--SINSIK----I------------------------------------------FETVIYPERLNK------LIEVEYFL-LKYFLNPGDDSKIK-----------EYIEVFDPKKKRYY--------AWMIPELFQ-------EVRN----------TLG-----VETVV-----------------------PVWKMRELGREVSQLF-------------------------
----------------ERTRYRLL-ELIRLAKKDEVYH-LLLKTYMA----NK------------KPFPE-ELNRVFS-------END----E-NFKNLMFAYVSGIISG----------------G-----EENE-------------------------

ABQ47101 ---------------------------------MPLVSNE------------QQREILLQVKEK----------------------------------------------------------KIKIPQIK------------KKSKELLSKFDDR--------------------------------------------------------------------------------------------------DLEELRNFLLQSEKEK----------------------------------------------------------------------FR-
--------------------------VYLSEI-IRSIFRKP--------------------------------------------------------DLSGSE-------------RCFLCGNPLRLSENTSI------------RQRKQFSRILFK--EFSSSEG--E--FPNSF---------------W-----N--IN-NE--YHLCDLCSQFALFRHFVFP-----------K---------NSKK-----------------EIF----VNVP------SFKAIWY---LNEA-----IKAYKD---------------------LNIELSVSRAV----AEVH-
MKLQ-------RLLGVWERQ-NIEIVN-YDGK-------------S-YSVYYVPQRTVDLLLNT-----RISSI-------------LN--SINSIK----I------------------------------------------FETVIYPERLNK------LIEVEYFL-LKYFLNPGDDSKIK-----------EYIEVFDPKKKRYY--------AWMIPELFQ-------EVRN----------TLG-----VETVV-----------------------PVWKMRELGREVSQLF-------------------------------------ERTRYRLL-
ELIRLAKKDEVYH-LLLKTYMA----NK------------KPFPE-ELNRVFS-------END----E-NFKNLMFAYVSGIISG----------------G-----EENE-----------------------

WP_065212803 -------------------------------MQRVDNEQQR---------------------EI-------------------------------------------------------LLQVKEKKIKIPQIK---KKSKELLSKFDGR-------------------------------------------------------------------------------------------------------DLEEIRNLLLQFEKEKF---------------------------------------------------------------
RS--------------------------HLSEI-ICSIFRKP-------------------------------------------------------NLSGSE-------------RCFLCGNPLEVSESISI------------RQRKQFSRILFK--EFSSSEG--K--FPNSF---------------W-----N--IS-NE--YHLCDLCSQLALFRHFVFP-----------K---------NSEK-----------------EIF----VNVP------SFKAIWY---LNEA-----IKAYRD--------------------LNVELSVSRAV----AEVH-
MKLQ-------RLLGVWERQ-NVEIVN-YDGK-------------S-YSVYYMPQRTIDLLLNT-----RIASI-------------LN--FINSIK----I------------------------------------------FEAVIYPERLSK------LIEVEYLL-LKYILKPGDNSKIK-------------EYIEVFDPKK----KGDYARVIPELFQ-------EARN----------TLG-----VETVV-----------------------PVWKMRELGREVSQLF--------------------------------------ERTKYRLL-
ELIRLAKKDEVYH-LLLKTYMA----NK------------KPFPE-ELNRVFS-------EDD----E-NFKNLMFSYVSGIISG----------------G-----EENE-----------------------

WP_081980988 ---------------------------MKIPQIKKKSKE---L----------------------------------------------------------------LSKFDGRD-----LE---EIRNLLLQFEKEK---------------------------------------------------------------------------------------------------------------------------------------------------------------------------------------------------FR------------
-----SHLSEI-ICSIFRKP-------------------------------------------------NLSGSE------------RCFLCGNPLEVSESISI------------RQRKQFSRILFK--EFSSSEG--K--FPNSF---------------W-----N--IS-NE--YHLCDLCSQLALFRHFVFP-----------K---------NSEK-----------------EIF----VNVP------SFKAIWY---LNEA-----IKAYRDLNVEL--------------------SVSRAV----AEVH-MKLQ-------
RLLGVWERQ-NVEIVN-YDGK-------------S-YSVYYMPQRTIDLLLNT-----RIASI-------------LN--FINSIK----I------------------------------------------FEAVIYPERLSK------LIEVEYLL-LKYILKPGDNSKIK-------------EYIEVFDPKK----KGDYARVIPELFQ-------EARN----------TLG-----VETVV-----------------------PVWKMRELGREVSQLF--------------------------------------ERTKYRLL-
ELIRLAKKDEVYH-LLLKTYMA----NK------------KPFPE-ELNRVFS-------EDD----E-NFKNLMFSYVSGIISG----------------G-----EENE-----------------------

WP_073164255 ------------------------------MVEPVTLYPAS---W--------YYNACAYGFLKV------------------------------------------------LAYG----LGEE--YVENNVLQ-----RDGTIRLTPGLVTALFTT-------------PEEDPSPRVPVKEDRHPPVSDIKRLAWWWVEMNR-----------------------------------------------------------EMIAWKEDELPNNLPPEEVI------------------------
------------------------------EKVCSKVITGRPLRVG-------TY--QG------NF-KKYGNLFQPSVHRDKKTF-LNKWFSCN-----------------------------------------------------------RDSGRE------------KCSFCFGQLTGQSGDT--------------FEQRLYHRFFMS--VLSDDLASSTGEFPNSF---------------W-----D--FK-PN--LLLCPLCRSIFLCAHFTIV----------------------Q-----------------VLF----ANTS------
NFLLTWH---FNKF-----LAQ-------------SQKNLFRTVK-NYQEELFYHFL-----HFN-NQMS-------RSMGAWELQ-GLEFLN-FTGE-------------G-IEHFVLPPVVCNLLLNP-----KVSSL-------------LA--RIRNRA----V----------------------------------------LK-YILEERFEL------LPNISYKS-LRVAL---------------------GTAPGNVKDPEAELSLREIINLCHLYA-------EIKK---------LFE-----RREPVVV--------------
--------------KTEEVLRCGQEAPAVL--------------------------------AKEGSSLIYRLL-ELVRLGKKDEAYY-LLLRSYIA----ES----------RPFPP-VL-TKVF-------QLQ--NMN-LFKNFMFSYIAGVQSN-------------TG----GGEI-------------------------

WP_088554534 -----------------------------MTKDVAIYPST---W--------YYNACVQGFLET------LAWGLGEDAV-----------------------------------------RSFLHDDG-----TV---KIPAKVMEAVFST------------RDVVMPENYPCLDGREVPDELKELKRITWW----------------------------------------------------------WVEKSKKKDGANVRET-------------------------------
--------------------------IDITCNSL-FGSNKTFYP------------NL------LT-HNTG--------ISRIDF-LNSWFEIA----------------------------------------------------------------NKGKREI-----------QCAFCGEIFSFDE-----------QMNRLLDRFFTLSLSK--DLGSAPK--K--VPNYF---------------W-----D--NK-PN--LPVCQQCRSYFLCFHIVYQ---------------------K----------------KYF----INSD------SLKVNWF---LNHL--
---VWN---------------------E-TRGEGSYPEAL-FTAVPYN-SQLR-------SALGSWGLQ-NMEVIS-FGGG-------------K-IDYYPLSDRLAELFLIP-----VIASL-------------LG--KLYHRK----L-------------------------------------------WD-AILQEKFEY------LLTIVYKS-LHVSLTGENRNEDP---------EVIITKGDKNDMKR--------VFVVIRLFY-------EIRK---------YLK----------KKGGVFVSTL----------------
DLKEIRRAAEESPFSL-D-------------------------------DNSKKGLIFRLL-ELTRLNKRSEVYH-LLMRTYVA-----------------CGKPF-PYCLAHL----FEVRDD----E-LFKTGIYAYISGIKTS--------------EE----SNEE-N------------------------

WP_073092678 ------------------------------MKDIEIYPSS---W--------YYNACVQGFMEI------IAHGIGEDG------------------------------NR----IIEE--EILHKDGR-----AVIPGDLANAIFNPCTVP------------------MPARYRNLEYNEVPNEIADLKRIAWW------------------------------------------------------------WIQKSQALLKKSSSNF--------------SNSHELIKATC----
---------------------------NSYLGSNKTFYPNLLPHN-------DK---------------------TSRIQF-LNNWFSLA--------------------------------------------------------NGGNV-------------HCSFCNALCQLDVHAK-------------EKEQFFIREISA--HFGSSIT--E--FPNGF---------------W-----D--GN-PN--FIMCRHCRSYFLCFHIVYS----------Y-------------------------------GFF----VNSD------SLMLNWH---LNSL--
---LTA---------------------KT-RKQKTSHYTLL-LNAIRYD-PQLR-------SGVGSWGLQ-NLEVIV-FMNN-------------K-IKHHPVSPRLARLFMVP-----KISSL-------------IS--RIAEPI----I--------------------------------------WE-IILEERFDY------LLTIIYKS-LRVLRTGKNESNDP---------EVIVKEKNKDSLTF--------VVDLIEFYA--------EVK----------KLD-----LNSSEERRYTNML-------------------
DLWKIRNEAKNAPLAL------------------------------------DNNLVFRLL-ELTRLNKKAEVYH-LLLRTYVV----NE----------MPFPG-VL-ADLFNI-----EDA----N-LFKSGIYAYISGLKA-----------------PE----NHQD---------------------------

WP_141264023 ------------------------------MQDVEIYPTS---W--------YYNACVQGFLEV--------------------------------------------------LAW--GLDVDEKKGEEIVVN---EILKKDGRAVIPG-------DIMTLAFSPRGNPHKDYRLKEVPEELPYLKRIGWWWVEKS---------------------------------------------------------YKDGFIPSKEKKAERN------
KLSEAEKIDMVFRRLFHFKGGYYP--------------------------------NLANLAYLK-------ST--------------------------TSKADF-LNGWFERD--------------------------------------------------NAATVSTL----------ICSFCGEETKLDINSS-------------IYNTFFTRPVSI--YLGNSYE--E--FPNLF---------------W-----D--GN-PN--LIMCKRCRSYFLCFHLVSK------------------------N-------------------RFF----
INSD------SFFVNWY---LNRL-----LAG--------------------------KVDNIRHQSLL--DAMQYD-QQLR-------KAVGSWGLQ-NMELLI-FERN-------------G-VEYYPISANLARLLLIP-----EITLL-------------IG--QIANNR----V---------------------------------------WE-FILQERFDY------LPTMVYKS-LRALINMKNDGKDP-------------EVVFEGPGN-----TLPVNNLIQLYY-------AIKQ---------QIA-----
SNNGGGKMSHI---------------------NLKEIREAAAEAPINPA------------------------------RNADKNFVYRLL-ELTRLNKKTEVYH-LLLRIYLT----SEEK---------KEFPP-PL-ARLF-------ETN--DCE-LFKTGIYAFIAGLIRS---------------E----KITN------------------------

WP_011393233 ------------------------------MQDIEIYPAS---W--------YYNACVQGFLEV----------------------------------LAYGLGEKGEE------------IVAG--KILQEDGR-----AVIPGDLMDAAFKPRGTP------------FPDGYVFREV---------PAELPALKRIGWWWVEKS-----------------------------------------------------------YKEGFIPSKDKKAKRN------------------
NPTNAEKIDMVFGRL-------------------------FHFKGG-YYPNLANLG-------------------HL--------KSTASKAEF-LNIWFDLN-----------------------------------------------------NIHTLRTP------------ICSFCGRKTELEINSS-------------TYNTFFTRTVSI--YLGNSYE--E--FPNLF---------------W-----D--GN-PN--LIMCKQCRSYFLCFHLISK---------------------------------------FF----INSD------
SLLVNWH---LNRL-----LAG-------------------------RTGKGGHKTLL--SAMQYD-RQLR-------RALGSWGLQ-NMELLV-FEQD-------------S-VEYYPISARLARLLLIP-----RVSSL-------------IG--QIANDR----I--------------------------------------WE-IILRERFDY------LPTVIYKS-LRALL--------------------KGGNNEKDPEVVYKDVTNVAPVTNLIHLYY-------MIRQ---------QLA-----RNEGGTSMGYL----
----------------NLKAIQAAAAEAPVDLN--------------------------------SNLDKNLVYRLL-ELTRLNKKTEVYH-LLLRLYLT----------------SGDNK-QF-PASLASLFATSDSE------LFKTGIYTFIAGLKANENDEPQ----------------GEND-----------------------

WP_062283506 ------------------------------MQDIEIYPAS---W--------YYNACVQGFLEV------LAYGLGEKG-------------------------------------EE----IVAG--KILREDGR-----AVIPGNLMDAAFKPRGTP------------FPDGYVFR---------EVPGELPALKRIGWW-----------------------------------------------------------WVEKSYDMGFMRKEDR-------------
GKSLNEVEKIE---------------------------TVARSL-FHKNANYPN-------LA--QL------TW-DLTK-----------KLSF-INNWFALE-------------------------------------------------------------EARDPSL-----------SCSFCGRQCNPDL-------------DGSVYETSFTRSLSI--HLGNSPA--V--FPNLF---------------W-----D--TR-PD--LIMCKQCRSYFLCFHLVSK-----------------------N-----------------KFF----INSD----
--SLLVNWH---LNRL-----LTG------------------------RTGRIGHKTLL--SAMQYD-RQLR-------RAVGSWGLQ-NMELLI-FTQD-------------S-VEYYPISAQLAHLLLIP-----RVSSL-------------IG--QIANDR----I-----------------------------------------WE-IILRERFDY------LPTVIYKS-LRALLKGGNNEKDP---------------EVVYKDVT-----NVAPVINLIHLYY-------TIRQ---------QLA-----RNEGGTSMGYL-----
----------------NLKAIQAAAAEAPVDL------------------------N------SNLDKNLVYRLL-ELTRLNKKTEVYH-LLLRLYLT----------------SGDNK-QF-PASLASLFATSDSE------LFKTGIYTFIAGLKAN-----------ENDE----PQGEND------------------------

WP_069590142 ------------------------------MQDIEIYPAS---W--------YYNACVQGFLEV------LAYGLGEKG---------------------------------------EE----IVAG--KILQEDGR-----AVIPGDLMDAAFKPRGTP------------FPDGYVFR---------EVPAELPGLKRIGWW-----------------------------------------------------------WVEKSYDMGFMRKEDR-------------
GKSLNEVEKIE---------------------------TVARSL-FHKNANYPN-------LA--QL------TW-DLTK-----------KLSF-INNWFALE-------------------------------------------------------------EARDPSL-----------ICSFCGRQCNPDL-------------DGSVYETSFTRSLSI--YLGNSPA--V--FPNLF---------------W-----D--TR-PD--LIMCKQCRSYFLCFHLVSK------------------------N-------------------RFF----INSD-----
-SLLVNWH---LNRL-----LAG------------------------RTGKGGHKTLL--SAMQYD-RQLR-------RALGSWGLQ-NMELLV-FEQD-------------S-VEYYPISARLARLLLIP-----RVSSL-------------IG--QIANDR----I-----------------------------------------WE-IILQERFDY------LPTVIYKS-LRALLKGGNNEKDP---------------EVVFQDPT----NVAPVINLIHLYY-------TIRQ---------QLA-----RNEGGKGMMGYL-
-------------------DLKAIQAAAAEAPVDL------------------------N------SNLDKNLVYRLL-ELTRLNKKTEVYH-LLLRLYLT----------------SGDNK-QF-PASLASLFATSDSE------LFKTGIYTFIAGLKAN-----------ENDE----PQGEND------------------------

WP_071520615 ------------------------------MQDIEIYPAS---W--------YYNACVQGFLEV------LAYGLGEKG--------------------EEIV-------AG----------KILQEDGR-----AVIPGDLMDAAFKPRGTP------------FPDGYVFR---------EVPAELPALKRIGWW-----------------------------------------------------------WVEKSYDMGFMRKEDR-------------
GKSLNEVEKIE---------------------------TVARSL-FHKNANYPN-------LA--QL------TW-DL----------TKKLSF-INNWFALE-------------------------------------------------------------EARDPSL-----------ICSFCGRQCNPDL-------------DGSVYETSFTRSLSI--YLGNSPA--V--FPNLF---------------W-----D--TR-PD--LIMCKQCRSYFLCFHLVSK------------------------N-------------------RFF----INSD-----
-SLLVNWH---LNRL-----LTG------------------------RTGRNGHKTLL--SAMQYD-RQLR-------RAVGSWGLQ-NMELLV-FEQD--S------------VEYYPISARLARLLLIP-----RVSSL-------------IG--QIANDR----I-----------------------------------------WE-IILQERFDY------LPTVIYKS-LRALL----KGGNN------EKDPEVVFQDPTN-VAP--------VINLIHLYY-------TIRQ---------QLA-----------
RNEGGKGMMGYL--------------DLKAIQAAAAEAPVDL-N------------------------------SNLDKNLVYRLL-ELTRLNKKTEVYH-LLLRLYLT----------------SGDNK-QF-PASLASLFATSDSE------LFKTGIYTFIAGLKAN-------------END----EPQG-E---ND-------------------

WP_071541950 ------------------------------MQDIEIYPAS---W--------YYNACVQGFLEV------LAYGLGEKG---------------------------------------EE----ILAG--KILQEDGR-----AVIPGDLMDAAFKPRGTP------------FPDGYVFR---------EVPAELPGLKRIGWW-----------------------------------------------------------WVEKSYDMGFMRKEDR-------------
GKSVNEVEKIE---------------------------TVARSL-FHKNANYPN-------LA--QL------TW-DLTK-----------KLSF-INNWFALE-------------------------------------------------------------EARDPSL-----------SCSFCGRQCNPDL-------------DGSVYETSFTRSLSI--YLGNSPA--V--FPNLF---------------W-----D--TR-PD--LIMCKQCRSYFLCFHLVSK------------------------N-------------------RFF----INSD---
---SLLVNWH---LNRL-----LTG------------------------RTGRNGHKTLL--SAMQYD-RQLR-------RAVGSWGLQ-NMELLI-FTQD-------------S-VEYYPISALLARLLLIP-----RVSSL-------------IG--QIANDR----I-----------------------------------------WE-IILQERFDY------LPTVIYKS-LRALLKGGNNEKDP---------------EVVFQDPT----NVAPVINLIHLYY-------TIRQ---------QLA-----RNEGGKGMMGYL-
-------------------DLKAIQAAAAEAPVDL------------------------N------SNLDKNLVYRLL-ELTRLNKKTEVYH-LLLRLYLT----------------SGDNK-QF-PASLASLFATSDSE------LFKTGIYTFIAGLKAN-----------ENDE----PQGEND------------------------

WP_071554045 ------------------------------MQDIEIYPAS---W--------YYNACVQGFLEV------LAYGLGEKG--------------------EEIV-------AG----------KILQEDGR-----AVIPGDLMDAAFKPRGTP------------FPDGYVFR---------EVPAELPGLKRIGWW-----------------------------------------------------------WVEKSYDMGFMRKEDR-------------
GKSLNEVEKIE---------------------------TVARSL-FHKNANYPN-------LA--QL------TW-DL----------TKKLSF-INNWFALE-------------------------------------------------------------EARDPSL-----------SCSFCGRQCNPDL-------------DGSVYETSFTRSLSI--YLGNSPA--V--FPNLF---------------W-----D--TR-PD--LIMCKQCRSYFLCFHLVSK------------------------N-------------------RFF----INSD----
--SLLVNWH---LNRL-----LTG------------------------RTGRNGHKTLL--SAMQYD-RQLR-------RAVGSWGLQ-NMELLI-FTQD--S------------VEYYPISARLASLLLIP-----RVSSL-------------IG--QIANDR----I-----------------------------------------WE-IILQERFDY------LPTVIYKS-LRALL----KGGNN------EKDPEVVFQDPTN-VAP--------VITLIHLYY-------TIRQ---------QLA-----------
RNEGGKGMMGYL--------------DLKAIQAAAAEAPVDL-N------------------------------SNLDKNLVYRLL-ELTRLNKKTEVYH-LLLRLYLT----------------SGDNK-QF-PASLASLFATSDSE------LFKTGIYTFIAGLKAN-------------END----EPQG-E---ND-------------------

SCY27864 ------------------MKQTTDFSYGGKSYNTLLQPTD---W--------RYSAAITGLVNF-----------------------------------------------------------------FTEFGIDYLVLENIS---DKPKTAISGFDGI---------------------------VYNQESITEDKYLEFAQSHF-------------------------------------------------------RDSMTHLTIENLLKSDEFNQDEIK-------------------RINELI----------
---------------------KSKTVFKE-VFGKTKFEG-----------SN------AD-FFLN-----KISKNRAKI-IKSTFRYG-----------------------------------------------------------KNM---------------------YTNFCNQNL------------------------LFTDSNKHCR--INGYNVDEGR--KTKYL-----------------------G--YCFNK--DSFVTNDIPEFDFIPFAFSNPS-MY--------------E----------------TYF----VN-NN----YSISTLVE---TNRE-----
ISR----------------RLDDIND-TDAKNKLLTVL-------------------KN-SKYFIDYDVEIIS-KNRD-----------EDFY-KTLFVRSQRLKRLQKLK------DQKFK-------------FS-YKIGENY----YFN--------------------------------------------------LDKEIYEKCLND----EVLDDSILFM-LKVLC----SDENS-----------------SQYT-SYL--------IKKMIEINL---------EWK-----------GQD-----------DMLAN-----------------------
DIRIARGAGYHTALKL-KED-KA---------------------------ENKISSFQQKII-GALIAHDYDRVTE-ILLNLSSY------------------AETEF-GFIYKFL-------ENPEENPE-ENKDLVFAFASAFNKN--------------SNDKG-----DNTN-E----------------------------

SKC65079 -----------------MINKIEIKYDIDGRSISFNRKIDPSD---W--------RYSAAIVGMIRF-------------------------------------------------------------------------------FQYRNIAY-----------KIEGRYLYYNIED-----------ISGDINDDRDYLYFAEKWFSDKMHHKEVDIRLNKSD-----------------------------------------------------------FSKDEISEINKKLEANTVMKK--------------------------------
-----------------------------------VFKGVKFDG-------SN--NK--------------------LIKDLI-EENRFEII------------------------------------------------------------------------SNTFCNSKNGYAK------------FTNNSKYRSKKSDVCR--LLGFYVDTGR--KTKSI------------------------G--FCFDK--DARTYNDEIEFDFIPFAFS-----------R---------SG-T------------------SIF----INNNT-----SIDYLMN----SNNE-----
MEDYLEKQ-------------------FENQKTWNSIF----YSYL-------------GGFEFIDFDVEVIIKN-METD--F--------------YESMFIRKAAIEIFKII-AKNKGFSES-----------LN--NIFKRY-----I--KIGENFYINV---------------------------------------------------ME-EVTKSILNE----LSLDDLIE---LIMKVEYESQSNTP-------------------D-TYC----------LSRLIYINE----------LIYK-----------NME---------DNMEKSA-----------------------
ELKGSSISAAEVTRYF-IAN-GQ-------------------------ENKIKGYQQRLA-NTLISKDYNRFIE-IMLQLSSY-----------------TQVPF-VFMHKLI----E--DFE------SNKNLAYNFINSLIIK--SNVKE---------GG----TSNE-K------------------------

SJZ63808 -------------------------MTLVRVEPFD---W--------RWSASIVGLLKY----------------------------------------------------LENYNIDYTLTED--YLEFDKEEITEERYY---NFVELYFENNMHH------------RTIEKIIKNMDNEELNSEELKLLNEKMKGNQILKKV--------------------------------------------------------FSNFKYEIGKEKDILKLIDENRREII-----------------------------------
---------------------------ENTYKN-----------GK---------------------SLYANF-GNPNALLK------------------------------------------------------------------------EKGK------------ICRLNGYYM-------------------------DMPKKGRSI----TYGFNSK-----ALSFT---------------------D------------------NIYFDFIPFGFS-------------------RSRE------------------GFF----INCNL-----DLRELER---VNRV----EFNE-------------------
-----IGTEENRTTFK----SNLLNNTFR----------SARKLNYDMEVIV-KNME-----------KDY-YETLYIRKDAINILRKISEEIKEII--------------LRPCKVGRNN----YSVNEWIP----------------------------------------VSEIVVDSILNL----KKLDDFIEFL------------------------------LKVDGKKF----LRGALIKINSLIY-------KEER---------NMNE------------------------------------------NFSKTVSDARRIKEFFKSK----------------------
-------PNKLRTYEQKLI-SAIGLKDYDRVKE-IMIHMSST---------------SQLPI-RTLILLFE------DFE------KNKNLAYTFINEIGEV--------------------KNEK----------------------------

SDB45648 ------------------MKHSIDPKVYSGFDTAISASD---W--------RYSAAIVGLQYY----------------------------------------------------LE----KMKK--TYKIEKNI----EID---KILDDFFLYHSQD------------------ITEKEYLQFVERFYGEELAHKALENKLNSKS----------TFHEEEVKWIK-----------------------------------EKMAANTTLKKIFSKVKFIGENKQ----------------
EILNLIEENRD-----------------------------------------FMIKETFRN-------KK--------------------------NLYDNY-CQSGVLLT---------------------------------------------------------EASKDS-----------VCRVKGYYI-------------------------------------------DTGKK-------GKSMA--------------Y-----N--FK-TD---TIVYEDDLIFDFIPFAFN----------G--------SSFE------------------TVF----LN-DN-----VDLELLIK---MNYN-----
VQV------------------------LW-KRKEEERIEIP----RNLMELLQS----------KTHQLKYGMEIIY-KDRE-----------KTH-FDSWYLRNESIEIFQKV-----NISKI-------------QL--NLKEGE----I---------------------------------------------YR-NILKETFKNIMNLSRVDDTIHFL-LKEKE-------------------------KKINTIGI-----NLAIEELLEINN-------RIKN---------GGE-----KMTG------------------------------
NIKAAKACAREVVKEFKKKNI--------------------------------EKKLDSYRQKLI-SSLVFRDYKGTLD-ILMQLSNY-----------------SGVSF-GFVFDFI-------ENP----S-QNDDLVRAFVFKLNSTEYEPNGKE------KK----TEDK------------------------

SDP18862 -----------------------MKKENLENQKFNTSMETSD---W--------RYSAAIVGLVK-----------------------------------------------------------------------------------YFDYLVER----------GYDEEYNFYKIDD-------DVIEYNSEEITEERYLMFVESYFESEMQHRDIEKMLQSDE---------------------------------------------------------FTEEQIKLINIKLDTG-------------------------
-------------------------AAIKKV-FFGTKF----------SK--EN------KD-GILR-----EINEKRQYL-IKETYKNR-----D--------------------------------------------------------------ELY--KKFSNR------------KCLFTKKNKI---------------------CRLLNYDI------DLGRKSK-----SISFN------------------W-----D--FK------NYRFEDESEFDFIPFAFS----------K--------SF-D------------------SFF----INNNY-----SIKQLVN---TNKL-----------ICI---
------------------DENPRATLL---------ENLK-------KS-GNYINF-DVEVIS-KQKI--EKGKKFENKEY--YETIFIRKRVIKIFEAI----DDYKGI-------------PK-KYRVNEN----Y--NIN--------------------------------------------VEKEVTDCIINN----IKLDYLIELL-LKPKQ----DTKTY-----------------KYPN-YWY----NIKTLIDINVLIY--------------------------GGD------------KMNE---------------------------KMKSACESARRVKVKI-ND------------
-----------------------EKKVNSYKQKLI-SAITFKDYGRFCE-ILLQLSSY-----------------SGVIF-YFAYDLF----S--DFE------EKNNIAYTFINALNNE--SN-----------KD----NGGN-K---DE---------------------

SFU69127 -----------------------MKTSLENRNYDTMLEPSD---W--------RFSAAIVGLMQY------FD-YHNI------------------------------------------------QYEMNDDAILYNQID---ITEDRYLNFVENK-----------YGERFHH-----------KVIENILKESELSDEQIKTV-----------------------------------------------------------NEKLKANTVMKNVFGKTKFNGSNKE-------------------------
------------------------------------MILNLLNEN--------RD--SL------IK-ETFR------LKSDMYADY-ANTNQIFN-------------------------------------------------------------ESQ--G----------------YCRLLGYYI--------------------------------------DSGKK-----G--KSTAY------------------------N--FVMNN----FVSQDIKEFDFIPFAFY-------------------------GDRE-------------TFF----INDNF-----NISRLKI---TSKI-----LEK--SLK-----
--------NKDEKV-KNSRQAFFKGI-----------------------IESSDFIKR-DVEVIV-KNVD--N-------DF-FETLYIRKKNIDILKSLSNNNVDYEAL---NI----------LF--KVTEKF----Y--RN---------------------------------------------------IQCEVMNDILND-IYLDDLIELFLKE-KRAYI---------------------------------------------------VSQLIKINI--------LMR---------GEG-----------NMND-------------------------KLKGAYACAKQVAGKI-E------------------
----------NNKIESYRQKLT-SSVIFKDYDRACQ-ILLQLSNY-----------------SNVEF-GFVYDLF-------D---SFE------DNKDLVYTFINALSKD------------------TN------RNDN-A---KN-----------------------

SJZ90361 -----------------------MKTSIQNEKYDTMLEPSD---W--------RFSATIVGLLQY------------------------------------------------------------------LNYHDLDY-----------KLEEDYILYNSSG----------------------INEERYLDFVEYKYGEELHHRLVENI------------------------------------------------------------LSNEEITEEQLKLINEKLVANT--------------------------------
-----------------IMKK-TFGKIKFDN-------TN--KK------EILDIIN---------KNRYEL-IRETFRRK-------------------------------------------------------------------SNMYANYGNT---------------------NQLFNDSQDHCR--LLGYCIDTGKKGKSTGF-------------------------N--FMMST----FVGSDIKEFDFIPFAFE----------------------GSRE-----------------AFF----INDNY-----TIQRLKI---SNEI-----LSK-------------------
KIEDDLEGENKRKDARQTL-----FKAI-------------METSDFIKR-DVEVIL-KDIS------------KEY-FETLYIRKESIDIFKEFINEKIEYKSF-------------CFS-HKVTDKY----YIN---------------------------------------------------IQKKVTESILNN---------VLLDEL-IEIFL----------------------------KEKNRSYL---------VLQLIKINV--------LIRR--------------------DKTMKD---------------------------RLKGAFACAKQVSKAIE---------------------------
-----SNKLDSYKQKLT-SSIIFKDYDRVCQ-ILLQLSNY-----------------SGIEF-GFVYDLY-------DDF----E-ENKDLAYTFINALSKK---------------------SENN----------------------------

RKY04506 ---------------------------------MRLVPSN---W--------LYNAGLVGFLRI------------------------------LRSKGIDVS-------SR----LEGG--AFELEAED-----LE---GFADAYFNHIIRT---------APEGFLLPRRKLGSRELKLLDKAGVNYDELRNHHLERL----------REKLNKLPSDC--------------------------------------------------
KAEEAEEVVREAAVSFADKISESLTNDPAAQRAAGKLRDKIIKRCMDKLEKQK-----------------------YAVDLLKP-FYLNKKIVG--------NYSLGG-------RE-TRLQ--------GFEREY-VIPARERL---------------------------------------------------------RTSGSPAN--GV-------RCKLCCLFP--------------------GRRSFSEGMFS--ITGVSET--K--FPNFF----------------Y-----N--LT-PD--
FYICEVCELILLCAWAGFN-----RIPFAL----RSGPQDTE----------------YIF----VNLP------SLPLLFE---ENEA-----LRK-------------------FYQIPQ-TVQNTAYEIVM-----EDLF-LKER--------EKKGEWVLQ-NVLFVE-IRPS-MRKD----RDKPI-FKYFHIGRDIARLFIDK-VAIESIRRV------------YG--SLFISNQTINL----------------------------------RR-EAVKRFVEG----DQVYDLIYQV-LRQNV-------------
--------DPGTRGNSNLPPLKSAFELSLLHSIRRVIW------EGYRK-----------GLR-----REAMESRQVYE----------------------ILSKFYEMGASLGKEID-----------------------------------LERRLRMSYRLL-SAVRSGRREEFYEMMMMKLFID----MQ-----------KPIPQ-DL-LSLLN-------PAD----PI-EFESKAYALLSGFLGE--------------------------EKEG----------------------------
```

```
WP_121510825 ----------------------------------MKVYASN---W--------LINMGIVGFLRI----------------------------LKYSGVDIAK-------LF----NADG--SIELTEDL-----LD---NFGHKYLAYSL-------------YWYTKELIDNLPYNEDLEEQTKIAIKSLEEALKDVE-----------------------------------------------------------TQLAEKPKNKDSLSERKKQVT---------------
KTIEKLSKFPTEFNLLLN----------------------------------YYANLPLIA-----------NP------SS-KTSK-----RAESIDEEF-VHPAIDSL--------------------------------------------------------------------DGTQDNSQ--P-------LCSMCSRVRSKPLE---------------KGNSIFSKSLFS--PLGVSLN--K--FVNYF---------------Y-----N--LE-PH--MYVCPVCRFLLVCAYAGLT----------R----KPYDHVDKTN---------------EVF----
VNLAA----LGIEELFN---LNEG-------------------------ISLVYSL-EKGETLYERVIELSLSGHI------------ERKSQWSLE-NVQFIE-MLAK-PQ------KDTPN-LKTYSLSPDVARLFADPKTVDTNLKNI---TGRIL-----LT-EGTNEEG----P------------------------------------------------MKQLVLKNLING----KDLMEPVFLA-LRTFV--------------------------KTDGYF--------VWLLISAFN-------TILLSARRNSIRSEID-----
GGAKNMGERGNYG--------------------ILRKYQEIGQRDFTHFE--------------------------------LDKKKRMSYVLL-SLIHNNKAEEFYE-NLLKIYIN----SD----------RSIPE-EI-VGLL-------SES--PTA-TFKEKAFAFMTGFLMGKGKDDNE-------DK----GGNQ--------------------------

ADC88653 --------------------------------MVVRFYVRD---W--------VSASAAVGLLKA--------------------------------------------------------------LEKLGQEP------K---EFIDGNSLIFEEK--------------------DIIEIFPSAFTKKLIDVSSQKNPKKDSF--------------------------------------------------------------VDRAYSVLSFLQKN---------------------------------------
---------FYSNSPLSNPSSVPKRLEEKSK------ELEDLYRSDPDKAVFELVKRF-VGDTFKKI---------------------------------------------------------------LERETSDH----------TCFFCGERKASIHN---------------GEPKTFEATNFT--PLSASPN--T--VENLF---------------Y------N--GK-NN--LYLCSVCEIVFYFAGFGFT---------------KVDKMDNR-----------------YIF----AYVS------DIKETLR---LNNT-----L----
-----------------------EYYKGFNKEFV----TSFL-KDLE-------HHKSRWILE-NIYIVE-VEKV---------SKVQSNIYSFSIPPRVAKAMKEY---LPKYPKT-------------LE--PV-----------------------------------------------------------FN-EFLKYIYFG----RSLYQLLFEI-LYGYFHEEELPDSK----------DSKKITDALRLGMRLAKGNKGLYPSLLFLS-------KFQE---------VVE----MKDKSFVEK--------------------------
QVGWAYREGLQLKNALLKELGDKA-------------------------LDRIRSISYSIL-EAIRRRDIDAFQQ-NLIRAYLS----VE----------REIPR-VF-VDAL-------KDE------SFSRIAYAFLIGLNGTERREEDG-------GE----AEETMG--------------------------

WP_010880193 --------------------------------MGVKFYIDN---W--------LTASSGVGLIEV------------------------------LQDAEVEYKD-------------LVKDSRKLELPEEL-----FE---KLPELYADFLTKG--------------------IDLTMKEQILKSKKLSIETLKNRLENPY--------------------------------------------------------TFINGYDIIKSFYRNSIFANNNP---------
YKDIIKQENSKLLNSILNDIHRKE-------------------------DMDYESIID-VLENKGYFE----------------------------------NIRNVIKYYLIETLKLII-----S----------------------------------------------------------------------NQEDKNAP----------LCFFCRERHTYVYK---------------GKYRVFGAEHFT--PLSASED--T--LPNLF---------------W-----N--GR-NK--MYLCPYCEFYLFFAAFGFT----------KV---------GNNR------------------FLF-
---VYIPD-----DLDSLIS---INSQ-----LKS------------------------KEKVEKNILGELF--RVVKFL-RNVE-------TQKARWILE-NIYFVE-IEKV---------SEATANIYSFSISPRLAKVLKNY---IDKYPPN--------------FE--SV-------------------------------------------------------------FP-KFVEYVYSG----RSLYEFLFKI-LSGFFFPKRYQNPK---------------GYDADLIKKGMSFKEFLPKKLLYFI-------KFQE----------ELI------
MGESFEK---------------------------QINFAYREGLNLKKMYEKELGKEKA-------------------------KDRIKTLSYRIL-EAVRRKDLDAFEQ-NLIRAYMQ----VE----------REIPY-IF-VQAL-------KDE------NFNRIVYAFLIGLNGR--DWSEGEPEGTSEES----LGQE---------------------------

SHK15454 --------------------------------MGIVFYVND---W--------LTAAAGVGLLRV-----------------------------------------------LENVKNSVPIEGN--RIDVPEDV-----FE---RLADYYAEYLLKD-------------------------------------------------------------------------------------------------------ITVEKLKDIIEKERQKKKEST-----------INVYNTLV------------------------------------------------
-LSKLKS-FYSNSPLAN-------PSSASKAQEVQIEI-SQDTEKLYKDAKQAIANF-IKKQFDTI---------------------------------------------------------------RSYKTSDK----------TCFFCGERKAYIKN---------------KEVKVFDATNFT--PLSASPK--T--VENFF---------------Y------N--GR-SN--QYLCPECEIFLYFSIFGFL-----------------PTGNKK-----------------YTF----VYMP------DLKKTYD---MNTI-----
LKGE--------------------------KGISTKYLKML---FTEYL-KELE-------GRKIEWTLK-NIYIVE-IELV---------GNAKANIHSFSIPPKVAIVLNER---IDSYPKF--------------LE--SVLG-----------------------------------------------------FFLDYIYSE----RSLYEFLLKL-LSGFF---------------------HKNTYTKVNVQTLEKVGAGLKSTAGLIYFV-------KFQE----------SLN-----NDGKNLDE---------------------------
PIEKVYKEGKSLGKSYD----------------------------------KEKLNAISLNLF-EAVRMRNVDLFYQ-TLIRAYID----ID----------KEIPK-VL-IHGL-------TEE------GFNWTAYSFLIGLGAV-------------TDEG----RGTS---------------------------

SNZ16151 --------------------------------MVVRFYADN---W--------LTASAGVGLLKA------LE-HKGIEW--------------------EPFLK--------GN-----------VIELPEDL-----FK---EIGKIYADYLLSD------------------------------------------------------------------------------------------------------FNQDTLKNLLEKERKKE--------------FNLYNSIAYPKI-GDFYSNS---------------
----------PIVNPSKT-GRANAEFKK----FYEKE--KN------IE-LLYS-----KAVDVVRNF-IKEQFDEL----------------------------------------------------LG--HERSQK------------VCFFCRERKAYIKD---------------KKVKVFEATNFT--PLAASPN--T--VENFF---------------W------N--GK-SN--MYMCPECEIFMYFSAFAFQ-----R-----T---------LR-G--T----------------YMF-----VYTP------DLRETFS---LNDI---
LAKEG--SVG-------------------------------IISKTI-----LEVA-KTVQ-------QRKSEWVLR-NIYIVE-IEKV--G-------DAQAN-IHTLSLSPRLASALKDL---IKDYPKY--------------ME--GL-------------------------------------------------------------FD-LFIEYVYSG----RSLYEFLWSV-MSGFF-----FKNRYKNLQGRKSLLVKKGTSFK-DYL---------PYGLLYFVK-------FQEVL----------NME-----------DKDKVNK-------------------
QVNWAYAEGKSLWEAY-TLG-MGEERA------------------------KKKVEVLSYRIL-DAIRRKDIDAFQQ-NIIRAYLE----------------VEKEIPY-VF-VEAL-------RDR------SFNRVAYAFLIGLNSK---------------GK-----QEEQ-E---TI--------------------

KJU82839 --------------------------------MRHRVYMRD---W--------YFNAGIVGFLTI-------------------------------VADGRSLDKLEHLHVGEN-----------FIEFDDAI-----FD---GFGDKFVKHAFLK------------FFSRQAYYHRFKKQKETQGKKKQMTKKQIAKSLDDL--------------------------------------------------------------------EKNPYLNFLSLLGCLTDNIE-----
THGDYMAFLDSTAGKIKGLTKHAIFQEL----------------------------NKTTDG-RHSVQSFVL------------SK------LK-GLISYKKGACAHDKVVGY-IQDLKQEG----------------------------------------------------------------------------SSKTIKNND----------LCVSCQVKK---------------------AEYEFSNAISN--IIGFNKD------NSNWI----------------W------G--FKAQN--VKVCPLCALIYNSAFESLI------------Y-----
VLKRVDSEYLNY--------------FYF-----VNQNA-----DLQTLYD---STVS-----FNTQLDII--------------------ADGTNVVIAMI-----KETV-RHIR-------TRQFNDTSG-NINFIE-VSDN--PIAGGQSTKGYN-TYNYNISVDIARFLDEQFTGDKIPKGF--------------YK-IKKTFYS----------------------------------------------------------IEEELLLYAINR-----QIGYATIGKY-FSYYL------------------------------DKSGKYN-----
LGKVTNYIISYI--------QTTR-----------GQK-------DMDNSQK----------------------------IVKKGYRSGITLREKLHGRKK-----------------------------DNQINGIAYGFL-NDLKLADREKFLD-KYMRVMMS-----------------NDLSL-AFGQDEM--------LND-----D-SFMQFGYSFLNGLLASKKDDNTD--------NS-----VQGG------------------------------

YP_002249829__2_ ----------------------MVNDSIKLYPSTVKHKIYMRD---W--------YFNAGIVGFLNI-------------------------------ITDGKELDSIPYLTIGEN-----------YIEFSDEI-----FE---GFEEKFSKYAFLK-------------FFNVNAYLQRLQKAQKDLTEKKTKIKPEQIRKKLEE-------------------------------------------------------------------IEKPPYKDFLKLLNIPVWEYK-----
SAEDFIDNLEKAQSKLKNLSKNQI------------------------------------FEILNKTPEGKESIQI--------FI--NQ------KLKGVCS--------HENVSEY-VSRIKAMK-----------------------------------------------------------------------YSKKLKNNE----------LCPSCQEKK------------------------AEYEFNNAVSN--IIGFNKD------NSNWI----------------W------G--FKVSK--LQICPLCAIIYSCAFASFA------------Y-----VLKIAEGEYLNY----------
---FYF-----PNENT------NLKTLYE-----IVKT------FNLLIDNI---------------------EENSNLLYVMI-----KQIV-DYIT---------KKQINRISE-NINFIE-IADN--PILAGQGTKGYN-IYNYNISPDIAMFLDLQFKNESVPKGY------------------YVLNKN----YCS--------------------------------------------------------IEEELLKLAISY-----QIDYSTLYRY-FAYYL--------------------------DTDNYSTKYN---------LNKVMNFVI--------KYIQ---------WVR-----
GESMEKSQK-------------------------------IINKGYLSGLSLRNELIKKGK-----------------------------ENQINGLVYGFL-NDLKIGDREKFLD-KYLRIVMS-----------------HNQPN-RFGKDEM--------LDD-----D-YFLQFGYSFVSGLISKSGIENNTGT------ND------IEEE------------------------------

TCS93601 ------------------------MGDDRGDSTLNKRIYLDE---W--------FINAGVLGFLTV-------------------------------LQRSASPSQ--------YQ-----VGQH--YIEFDPAV------LD----SFAVHFFDILIEK------------FSKYEQDRLRFQKQLEFAKREDYLKANKDQIAKTIK-----------STRDKLKKAKI---------------------------------------DKATIDQIEDLLGQLKTIKTYQD---------
-----YEALEAVTHSYLMVLEDTNI-----------NNVLTINYI-------RSVLSA-LFGQPSFLN--------PSFQGT------KE-QFIE---------KFHDDY-IKPVIAEC------KLMDWVRTVPTDQLEVELKKKKKEKETQGHLKK-------------------------------------LVNEAIKQ--TQSIERILPCSVQPDFP--------------------------SSIQFEEKMFF--PLGLSLE------NQNMA----------------W------D------GRSTILISNMTRFLLFCSSISFT----------
Y---------YLKRESHFGSTEYTPT-----FGF-----VNLYT-----SIKHLVA---ENKV------FEN------------------------K-RNESNPFQELI-----YDLL-LKTT--------EDLSIYTLQ-NILFLE-FSVE-----------KKNSK-LHYFHIPKKLALFFKED--------ASD----------------LK--RMKNKS--------------------------------------------------------------MQHLFVEEVLKR-----RDTIYFIEKS-LRNLI-----------------------------SDKHQAQVV---------FWAILARH------
--KLEH----------RLR------GESTLSDK--------------------------KLTKLFFEGKELAALFHLQQK--------------------------------DNQLQGISYRLL-NACKAGNKQLFFD-TLLRVYMT-----SN------------RIVPS-LF-LNILH-------EKD-----L-DFVEVGYSFISGLQAK---------------PE-----GGDRYE----------------------------

WP_040306841 -----------------------------------------M---------IIFFGQMSFLQK----------------------------------------------------------------GLSNKS------LD----EHKQIMFKDFIKP------------------------------------------------------------------------------------------------------------LIIDIKVEEY-------------------------------------------------------------------------------LQNNDLNSLSSFLE---------KE--
-----------------------------QGKEEN-VKAYKNYL-------------------------------------------------KKSKKDMNKFNESLCANSYCSICNIYP---------------------SFKEDFNESRFL--ILGVSNE--N--ASNFF-------------------W------N--FI-TK--YPICNLCKLVILCAPAGVV--------------K------FDKFYMEKEEWWKRNF-----------YTF-----VNLDT-------NLEDLFK----KNES-------LKKK------------------------------ISSDNPYKEFI--
--VDIL-EQAR----------EESIWTLQ-NILFVE-FNGQ--------YQSKKSN-LKYMHISKFIAKYFKTY-------SRET----------------IG--RIKNRK-----------------------------------------------------------------------------LKVELIDKILQN-----EDLNRVIFIR-LKEYI-----------------------------KNNANGADIYY-------------STLSKALL---------NCIK------------MK-------GDENKVKD------------------------------KVYRAFFTGKEMREYLKKNKS----------------------------------
ENKISSLAYRLL-NTAKVGNKKDFMD-TVLRLYIN-----SE----------MEMPR-IFLEAFY--------EDN-----I-DFETLAQSFIAGLITE--------------TK-----EVEVHE---------------------------

SEF76638 --------------------------------MQVRVYLND---W--------FYNMCIVGMIRI----------------------------LEHAKRKGEDIS--------FV-----LGED--FLEIDDKT------IN----NFHNYYFDYFYDE-----------YTEKEYIKIKDRFNQLKLRSDGDIIKEIEKFLSETK--------------------------------------------------------------LADKATKKGFESEVKELLDIIKKVKKNHD--------------
LNTLAILINKIE-SLLNKND-----------IVEKYALDNIR-------SMMYDN-FFGQMSFLQ--------KG--LS-------NK-SLDE------HKQIMFKDF-IKPLIVDI------KVKEYLQNNDFNNLSSFLENEQGKEENIKAYKNYL--KK----------------------------------SKKDVNKF--NESLCANSYCSICNIYP---------------------SFKEDFNESRFL--ILGVSNE--N--AFNFF-------------------W------N--FI-TK--YPICNLCKLVILCAPAGVV-----------
K----FDKFYMEKEEWWKRNF-----------YTF-----VNLDT-------NLEDLFK----KNES-------LKK---------------------------K-ISSDNPYKEFI------VDIL-EQAR--------EE-SIWTLQ-NILFVE-FNGQ-YQ-------SKKSN-LKYMHISKSMAKYFKTY--------SRGT----------------IG--RIKNRK-----------------------------------------------------------------------LKVELIDKILQN-----EDLNRVIFIR-LKEYI-----------------------------KNNANGADI-----
YYSTLSKALLNCI-----------------------KMK------GDENKVKD-------------------------KVYRAFFTGKEMREYLKKNKS--------------------------------ENKISSLAYRLL-NTAKVGNKKDFMD-TVLRLYVN-----SE-----------MEMPR-IFLEAFY--------EDN-----I-DFETLAQSFIAGLITE---------------TK-----EVEVHE---------------------------

SFF89372 ----------------------------------MRIYNSD--QW--------VYKMGIVGFDRI-------------------------------IQFNFENYGLDKEDYGYK-----IEDD--YIEFDVGL------LN----ELPKAFFQYFYHE-----------------YDVAENINYKFNYLLNRASKESNFKNSINEIK------------KLIKDKNARIQ----------------------------------------KLQLDNFNEFDEIYNDVCNIKFSEL--------------
EELKEGLEKYVILFRDDKV--------------------NYPQALNLFKSVLQTSFFGQNSFLN--------VC---NN-------SK-SIEE------QQDIMFKDY-VKPVIQSE------KLTKIKEDMNEGELKQYIDNNMDSSKQATEVDKYM--KDIC---------------------------KNLWGKKRKFESL--SIMDGTYNKCCLCDENL-----------------------SYGSDYSDGNFI--PLAISND--G--CKNLF-------------------W------D--MN-SK--
FPICPLCRLILLCTAAGTT------KVYKNY-----LESNYSFNDKLY-------------YGF-----ISMEG------NLGDLIK----ENNT-------FKN--------------------------L-SSKDISFEEWL------YDSLIQE-----------RKISKWQLQ-NVLYVE-FNAD-YR--------SKNSR-MNYYNIPNYIASFLCNK--------DADI-------------LQ--EIKSST----------------------------------------------------------------ERMRIFDWIMGR-----KDLNILIDRE-LRKKI----------------------
KNEFVGTNII-----NVVKLKNILESYKAEGGYSKVNDA----------GLQ-------------------------------------------TIDWLYKDGIMIAKIYHDNKM-------------------------------ENKLSGISYRLL-NSIKAGNKNEFMD-SILRIYLS-----VD-----------LPVPK-NM-LESI--------REE----KL-SFSQVGHSFLTGLNGYYKKEK-----------EG-----AISE-----------------------------

AGF58568 --------------------------------MKIYNSE--QW--------FYKLGIVGFNRI-------------------------------INYNKDYYGLDVDKYNYN-----VSSD--FIEFDSEL------LE----DFPTYYYNYFINE----------YNVSESMNEKLN-----KKLGAAKDKDKFQERLNDIK-----------KVIGERNKKIE---------------------------------------KLQVAEFDKCKDIHSKIGKVKFEQL-------------
AEVEELVKKYEDIVQLEHLNHP--------------MTVINFK-------SILYNS-FFGQISFLN--------LTHNNK-------SI-DYQK-------EMFFKDY-IKPVLDGE------KFSKIIKANDEVELKKYIEIKKEDAAKCPKNELNN--IDRVVSIINTALFG-------------KKRKAESV--NDVLENYNICRICDDNI--------------------------SFGDDYTDGNFI--PLAISNA--N--SKNLF-------------------W------M--FD-SR--
YPICPLCKLVLFCTAAGCT------KIFKSY-----LNDKYDYNDKLY-------------YGF-----VSIEG------DLLELIQ----GNDN-------FRN----------------------------V-TKKDGTFEEFI------LDSV-KQNR-----------KISSWQLE-NILYVE-FNAD-YR--------SRNSK-LNYFNIPTYLARFLRDK-----DTYTE----------------LD--NMKNSR--------------------------------------------------------------EKMKIFDLMIQR-----KDLKHVIDLE-LRALV---------------------
-------NEEAYANR-----DILRTIRIRNYLK---------QYKG-----------GCK------KVDELAKK---------------------------NLNYLYFQGVDISKIYRSNNS------------------------------ENKITGLSYRLL-NSAKAENKKDFMD-TIIRIYMS-----VD-----------KEIPM-IF-LEVM--------KES----KL-DFEEIAHSFIAGLNSKGKENDD----------RV-----EGEN-----------------------------

WP_077853371 --------------------------------MKIKVYLND---W--------FYNMGIIGFMRI-------------------------------LENSEMLNE--------VS-----RKDN--YIEFDSSI------LK----EFGKYYFDYFMNR-----------YDVPKDRLNRYQSQKAYVKKKNDYKDGIKYLKKTVI-----------------------------------------------------------DIANKFKGNENEIDLKNIADEISKVKNEEQ-------------
FEILNEQYNRFKEVLEREEVSEK---------------LTINKVR-------SILSDQ-YFGQASFLQ--------KT---VA-------AK-RLDE------QRKIAEKDY-IMPIYEMG------KLIDAYSNKSFDEVKKCIDEYLRLDKKCKDKNIDK--IYNKINK--------------KISKTKDINLISE--ELNSPKLYNCSLCGEFK---------------------------SLGDDFEEKIFV--PLGISNS--N--ANNLF-------------------W------N--FN-SK--
FPICNVCKLILFCTPAGAT------DLFKGY-----MEGDYKDKVY-------------FGF-----VNYDT------SISKLYK----ENEN-------FYN------------------------------R-KDKESPYNQII------LDIV-EEIG--------VK-SEWELQ-NILFVE-FNVN-----------GKNCK-LNYFNIPRFLAKFFVNK--------GKE-------------LD--NMKNEK--------------------------------------------------------------FKSALIELILKN-----IDIKFAIDER-LRQCI----------------------
KNGYIALDCY----------IACKIRYYL--------NMYR-----------EGK-------NDMNSK---------------------------RLFFIYKNGQDINKYFTEHSG------------------------------ENKISGIAYRML-NSVRTGNKNDFMD-AVLRTYMA-----AGQ-----------KQVPQ-VF-SEIFA-------DDD-----S-EFESVAQTFIAGLISNATPKKEE---------EA-----NKNE-----------------------------

ERI93854 --------------------------------MKIKVYLAD---W--------FYNMGIVGFRRI------------------------------LKFQEDYRSLKLSDFEYQEADNYIEFNPDL------LK----EFHNYYFDYFLGR----------------YNMARFQSIQLDKYLNLSKKEGKFNESIKVIK-----------DIVKANNNKIK-------------------------------------KIDETVFTKAEEIYKSLGDIKRQEQ-------------
ISELEELLLKYKDILELKNI------------NEKITLNKF---------KSIMSNSFYGQVSFFN--------VIHTSK-------TIVEQKD-----------IMFKDY-ISPIITME------ELKTEINENRAGDLEERISKALSDKATPKNIGKLYKNINSKFIK---------------------KKKSLEEIEQYIGGKDFALCQMCGEYK----------------------------SIGSEYGEGDFV--PLAVSTD--N--SRNMF-------------------W------N--FN-TS--VPLCDICKLILICSAAGAT-----
DIFKGYMNANLDNKYKQY-------------------FAF-----VNMDT-------TFKDLYK----TNES-------FNLK-----------------------------KDKENPFKELI------FDIV-SEAK---------ERSTWQLQ-NILYVE-FNSD--------YGSKNCK-MNYFNIPKYIAQFFKAK--------AEV-------------VN--SIKDEK--------------------------------------------------------------FKAEIVDDILSN-----KDIKFSIDRK-LREVL------------------------ANNYGNPYD------
CYRAVVIRYYL--------KVSK----------GGIK------KMATEVDDK---------------------------RLKFIYMRGLELNKHYTKNGG---------------------------ENKISSIAYRLL-NAAKSNNKKEFMD-TLIRIHLN-----AQ-----------MEVPT-IF-LDIM--------SEK-----ELAFEEIAHSYVAGLISKASSKKD----------SE-----ENEN-----------------------------

WP_069978076 --------------------------------------MND---W--------FINMGLIGFIRI---------------------------LKEAGKIDDNDI--------KK-----MKNN--YIEFNSEL------LE----DFHKNYFDYFLKR------------YDISVRECEKID-----RYLLIAQKEDKFKNATDWIK-----------GVVDNNRKKIK--------------------------------GKLENKVFEQQFDEIYKNLSKIKKADQ--------------
IEELKDSVSQFKKLMTVKD------------VNNKLSINFVR-------SIFSKQ-YFGQASFLQ--------KP--CA-------AK-SVEE------QSEIMFRDY-IKPVIGLS------ELMDQSKDIDALKMNLENELKDSSLSKNY-----------------------------EKMLKTISGLAKT--RDSLEAVKKCFYEDMLYCSVW-------------ENFRATNDFTEGVFA--PLAVSNN--N--AKNYM-------------------W------D--FC-TF--
YPISDIAKLILLCTPAGTV-----D---M---------QDEY-------------------FGF-----VNLDT-------DVYELFR----QNEN-------LRL----------------------------K-KEEDNVFESLI------YDIV-SEAR---------EK-SKWMLQ-NILFIE-FKMD-ID-------SKNCK-LNYFNIPKNVASYFKKH-------AQND----------------LG--EIKDKT----------------------------------------------------------------FKKDLVALMLDN-----KTIKTITYQY-NEKSKGKEQIANNI-------
ERLIEVKLRSIISSEKY-----SAYDALLATIAKH--------KFNR-----------TKE------GCEKVDSK----------------------------KIWVIYKSGQELNTYF-KRN-QS--------------------------ENKIQGIAYRLL-NASKAGNKKEFLD-SLLRIYMT-----AG-----------KEVPS-IF-LNVM-----H--EQD-----L-EFETVAHAFVSGLISS--------------ENHG-----VKED-K----------------------------

WP_006305869 ----------------------------------------------------------MGIVGFIRI--------------------------------LEDAGLKKE--------LD-----ICDN--YIEFDSKL------LE----KFDEYYFDYFLKR-----------YDISKREIKKVDRYL-GIARKEEKFKDAVNWIKSVI-----------DTNRKKVKGKF--------------------------------------EDKKFEYEFNGLFKEIKKVKKYDQ----------
IEKLEELVDRFKKLITIYEVNEK----------------LTLNYIR-------NILSTQ-FFGQASFLQ--------KI--CA-------RK-SLEE------QREIMHKDY-IKPILEEV------RLYKELNMSETVDDLLQFIDTELNKDCSKTYPKLL--RNIK------------------------KLLKKKKV--LDNLKKIPHCSIWEEYL---------------------ATSDFTEAVFV--PLAVSNN--N--AKNFM-------------------W------Q------GNTSYPICSLVKFILFCTPAGVT------
DMKDGY-------------------------------LGF-----VNMDT-------SVEELYR----HNEN-------LLL---------------------------M-QDEENPFESLI------YDIL-SETR---------KKSIWTLQ-NILFIE-LNAN-YD-------TKNCK-LNYFNINKNTARYFKSY-------TKNN----------------LM--KIRDTR---------------------------------------------------------------FKGELVSWILNN-----KDIKYLIHDK-LIETI------------------------------KSGKSSYD---------SYLATKARF-------
TLKK---------VLK-----GCEKVDSN--------------------------RLWVIYKEGERLNTYFKENES------------------------------ENKIQGIAYRLL-NASKARNKKEFMD-SLLRIYMA-----AG-----------KEVPS-LF-LNVLH-------EKD-----L-DFESIAHSFISGLISN-------------FDNNE-----KEEK------------------------------

CUP17261 --------------------------------MKLRVYQND---W--------FLNMGIVGFMTI-------------------------------LEKADKLHDS--------VI-----IKDN--YLEFDSKL------LQ----DFSGYYFQYFIDE-----------YRISNRLKGTLQNYKGVIKKNPDKIKEYLKRVKDTV-----------KHQADKVKKFD---------------------------------------KENYEILKTNMDLLGKIKNYEQ-------------
FEDVEKYMDDILEIFKKQHI----------DEKLTLNLYK-------YVVGDN-YFGQVSFFN--------VS--KS-------AY-DLEG------LKKVMYNDY-LLSIVSYG------ELIDVIESDDIERVKKEVSEKLDFFKEEVESKNIS--KNSIKNIEKIYKEVFKYIN--------KKKGIKEI--KEYIESLSTCKMCGDYS---------------------GIVSNYTESNFA--PLAVSAA--N--STNLY-------------------W------N-----
MDTTFEICDICKLMLFCTPAGAT-----------I----VNKRYLTNEENEF-------------YSY-----LNVDT------SLKELYE---KNQS-------LRV--------------------------Q-RDKENPFSETI------LDIA-EENQ---------KK-SEWQLK-NILFVE-FKGS-IE-------AKKSK-LNYFDMPVYLAKFISKQ--------YEL----------------IK--FIYDIN---------------------------------------------------------------LKNSVMDYLLNE-----KDLKSLIFLK-LRDYLDEESRGIIK---
----------SNRNTIDVFK---------TVKIRALLQ--------VYKK-----------GGN------DMNKKVLHF--------------------------RLDEIRKMGLQVREKYNERKE------------------------------DKKIASIGYRLL-NLIKSGNKVDFMN-DFFKIIVG-----FE-----------LIVPK-SV-INVMS-------ERD-----F-DFETFGYAFISGLLGEEDEDKKGNNDQKINEN-----KQNG-Q-----------------------------

WP_054274893 --------------------------------MKLRVYRGD---W--------FFNMGMVGFLNI------------------------------VKKADMKEK--------IF-----IMED--YIEFDSSF------LE----NFHEYYFNYFMDE-----------YDVSKRIRKSVDYSINFIKFKPDKIKDGIKKIKDNI-----------KQQNDKVKKFD---------------------------------------EEKFNLIKEKLDSMSKIKSYDE-------------
FEILETLVDEIV--DIFKIK---------SINDRLTANLYK-------YVVQDN-YFGQVSFLN--------VV--KA-------KL-DLDG------LKQVMFNDY-LRQIIYFG------ELEDLLEENDYDKLKNYLNDRLDSISKDVSEKKVS--KPGVKFSIKTIDKIMKEINKNFIK--KNKSIEDI--REYMDSLEVCEMCGSYK---------------------GILDEYSESNFA--PLGVSTN--N--ARNMF-------------------W------N-----
QEYTSSICDVCKLILFCTPAGAT------YTRKNY---------LSNEENEF-------------YLF-----VNMDT-------SINEIFE---RNNE------LKL-----------------QKSKLSD-SKDESFFNKFI-----KSIV-EENK---------LK-SEWQLR-NILFVE-FKAS-ID-------AKKCK-INYFNIPTYLVKIFTNG-------KASKK----------------IQ--DIYDYK---------------------------------------------------------------LKSNVLDLLLKN-----RDLKHLINNT-
LREKVKNDMEGSNK-------------SNISGTD-CFR--------LVQLRTIVN--------NYKR----------GVY------------KMDYK-------------------------KLKKVYESGCEIHDYY-IDK-NA---------------------------KNKIEGTVYKLL-NLIKVGNKNDFFQ-TVLRIFSP-----SE-----------KTPPE-EF-LHML-----S--EED-----L-DFEAIGYAFTMGLMSEKCDRKNKI--------VD-----NKEE-K-----------------------------

SKA34667__9_ --------------------------------MKLRVYRGD---W--------FFNMGIVGFLNI-------------------------------IKKSNKQAE--------IF-----IMED--YIEFDSSF------LE----NFHEYYFNYFMDE-----------YDVSKRIKKSIDYSINFIKSKPDKVKDGIKKIKDTV-----------KQQNDKVKKFD---------------------------------------EEKFNLIKEKLDSISKIKSYDE-------------
FDSLEILVDETIDIFKIKS-----------INDRLTANLYK-------YVVQDN-YFGQVSFFN--------VA--KA-------KL-DLDG------LKQVMFNDY-LRQIIYFG------ELEDLL-EENDYDKLKNYLNDRLNSIAKDISEKKV--SKSSVNTIEKIMKEINKNFIKKNK--SIENIKEY--MDSLE---ICEMCGLYK--------------------GILDEYSESNFA--PLGVSTN--N--ARNMF-------------------W------K-----
QEYVSSICDVCKLILFCTPAGAT-----------Y---------TR-K--DY-LTNEENEF-----YLF-----VNMDT-------SINELFD---RNNS------LKA-----------------QKSESSN-SKDENPFNQLI-----KSIV-EENK---------LK-SEWQLR-NILFVE-FKAS-ID-------SKKCK-INYFNIPTYLAKFFVKE--------NKK-------------IQ--SIYDYK---------------------------------------------------------------LKSNVLDLILKS-----KDLKHLINNT-LRNIV-----K-SNL---
---ESNNKSNI-SGLS-CFN----------VVQLRTILN--------NYKR-----------GVY------KM---DYK-------------------------KLRKIYESGCEIHDYY-IDK-NA---------------------------KNKIEGITYKIL-NLIKVGNKNDFMQ-TILRIFSP-----SE-----------KTPPE-EI-LQIFV-------EED-----L-DFESVGYAFVTGLISGKYEGKDKL-------SN-----EKEE-K-----------------------------

SJU95103__4_ --------------------------------MKLRVYRGD---W--------FFNMGIVGFLNI-------------------------------IKKADKQAE--------IF-----IMED--YIEFDSLF------LE----NFHEYYFNYFMDE-----------YDVSKRIKKSIDYSINFIKSKPDRIKDGIKKIKDSV-----------KPQNDKIKKFD---------------------------------------EEKFNLIKEKLDSMSKIKSYDE-------------
FDSLEALVDETIDIFKIKS-----------INDRLTANLYK-------YVVQDN-YFGQVSFFN--------VA--KA-------KL-DLDG------LKQVMFNDY-LRQIIYFG------ELEDLL-EENDYDKLKNYLNDRLNSIAKDINEKKV--SKSSVNTIEKIMKEINKNFIKKNK--SIEDIKEY--MDSLE---VCEMCGLYK--------------------GILDEYSESNFA--PLGVSTN--N--ARNMF-------------------W------N-----
QEYVSSICDVCKLILFCTPAGAT-----------Y---------TR-K--NY-LTNEENEF-----YLF-----VNMDT-------SINELFE---RNNS------LKA-----------------QKSESSD-SKDENPFNQLI-----KSIV-EENK---------LK-SEWQLH-NILFVE-FKAS-ID-------SKKCK-INYFNIPTYLAKFFVKE--------NKK-------------IQ--SIYDYK---------------------------------------------------------------LKSNVLDLILKN-----KDLKHLINNT-LRNIV-----K-SNL---
---ESNNKSNI-SGLS-CFN----------VVQLRTILN--------NYKR-----------GVY------KM---DYK-------------------------KLRKIYESGCEIHDYY-IDK-NA---------------------------KNKIEGITYKIL-SLIKVGNKNDFMQ-TILRIFSP-----SE-----------KTPPE-EI-LQIFV-------EED-----L-DFESVGYAFVTGLISGKYEGKDKL-------PN-----EKEE-K-----------------------------

SUY15783__24_ --------------------------------MKLRVYRGD---W--------FFNMGIVGFLNI-------------------------------IKKADKQAE--------IF-----IMED--YIEFDSLF------LE----NFHEYYFNYFMDE-----------YDVSKRIKKNIDYSINFIKSKPDRIKDGIKKIKDSV-----------KQQNDKIKKFD---------------------------------------EEKFNMIKEKLDSMSKIKSYDE-------------
FDSLEALVDETIDIFKIKS-----------INDRLTANLYK-------YVVQDN-YFGQVSFFN--------VA--KA-------KL-DLDG------LKQVMFNDY-LRQIIYFG------ELEDLL-EENDYDKLKNYLNDRLNSIAKDINEKKV--SKSSINTIEKIMKEINKNFIKKNK--SIEDIKKY--MDSLE---VCEMCGLYK--------------------GILDEYSESNFA--PLGVSTN--N--ARNMF-------------------W------N-----
QAYVPSICDICKLILFCTPAGAT-----------Y---------TR-K--NY-LINEENEF-----YLF-----VNMDT-------CINELFE---RNNS------LKA-----------------QKSEYSD-SKDENPFNQLI-----KSIV-EENT---------LK-SEWQLR-NILFVE-FKAS-ID-------SKKCK-INYFNIPTYLAKFFVKE--------NKK-------------IQ--SIYDYK---------------------------------------------------------------LKSNVLDLILKS-----KDLKHLINNT-LRNIV-----K-SNL-----
-ESNNKSNI-SGLS-CFN----------VVQLRTILN--------NYKR-----------GVY------KM---DYK-------------------------KLRKIYESGCEIHDYY-IDK-NA---------------------------KNKIEGITYKIL-NLIKVGNKNDFMQ-TILRIFSP-----SE-----------KTPPE-EI-LQIFV-------EED-----L-DFESVGYAFVTGLISGKYEGKDKL-------PN-----EKEE-K-----------------------------

VTR06361 --------------------------------MKLRVYRGD---W--------FFNMGIVGFLNI-------------------------------IKKADKQAE--------IF-----IMED--YIEFDSLF------LE----NFHEYYFNYFMDE-----------YDVSKRIKKNIDYSINFIKSKPDRIKDGIKKIKDSV-----------KQQNDKIKKFD---------------------------------------EEKFNLIKEKLDSMSKIKSYDE-------------
FDSLEALVDETIDIFKIKS-----------INDRLTANLYK-------YVVQDN-YFGQVSFFN--------VA--KA-------KL-DLDG------LKQVMFNDY-LRQIIYFG------ELEDLL-EENDYDKLKNYLNDRLNSIAKDINEKKV--SKSSINTIEKIMKEINKNFIKKNK--SIEDIKKY--MDSLE---VCEMCGLYK--------------------GILDEYSESNFA--PLGVSTN--N--ARNMF-------------------W------N-----
QEYVSSICDVCKLILFCTPAGAT-----------Y---------TR-K--NY-LINEENEF-----YLF-----VNMDT-------SINGLFE---RNNS------LKA-----------------QKSESSD-SKDENPFNQLI-----KSIV-EENT---------LK-SEWQLR-NILFVE-FKAS-ID-------SKKCK-INYFNIPTYLAKFFVKE--------NKK-------------IQ--SIYDYK---------------------------------------------------------------LKSNVLDLILKS-----KDLKHLINNT-LRNIV-----K-SNL---
--ESNNKSNI-SGLS-CFN----------VVQLRTILN--------NYKR-----------GVY------KM---DYK-------------------------KLRKIYESGCEIHDYY-IDK-NA---------------------------KNKIEGITYKIL-NLIKVGNKNDFMQ-TILRIFSP-----SE-----------KIPPE-EI-LQIFV-------EED-----L-DFESVGYAFVTGLISGKYEGKDKL-------PN-----EKEE-K-----------------------------

SJV83804__3_ ----------------------------MFNDYLRQIIYFGE---------------------LEDL-----------------------------------------------------------LEENDYDK------LK----NYLNDRLNSIAKD---------------------------------------------------------------------------------INEKKVSKSSINTIEKIMKEIN---------------------------------------------------------------------------------KNFIK-----
------KN------KS---------------------IEDIKEYM---------------------------------------------------------------DSLE---------------VCEMCGLYK--------------------GILDEYSESNFA--PLGVSTN--N--ARNMF-------------------W------N------QEYVSSICDVCKLILFCTPAGAT----------Y-----TRKNYLINEENEF-------------YLF-----VNMDT-------SINGLFE---RNNS------LKA------------------QKSESSDSKDENPFNQLI-----
KSIV-EENT---------LKSEWQLR-NILFVE-FKAS--------IDSKKCK-INYFNIPTYLAKFFVKE--------NKK-------------IQ--SIYDYK--------------------------------------------------------LKSNVLDLILKS-----KDLKHLINNT-LRNIVKSNLESNNK----------------SNISGLSCF-----NVVQLRTILNNYK--------RGVY------------KMD--------------YK-------------------------------KLRKIYESGCEIHDYYIDKNA---------------------------
KNKIEGITYKIL-NLIKVGNKNDFMQ-TILRIFSP--------------------SEKIPP-EEILQIFI-------EED-----L-DFESVGYAFVTGLISGKYEGKDKLP------NE-----KEEK------------------------------

SJW93676__8_ --------------------------------MKLRVYRGD---W--------FFNMGIVGFLNI-------------------------------IKKADKQAE--------IF-----IMED--YIEFDSLF------LE----NFHEYYFNYFMDE-----------YDVSKRIKKNIDYSINFIKSKPDRIKDGIKKIKDSV-----------KQQNDKIKKFD---------------------------------------EEKFNMIKEKLDSMSKIKSYDE-------------
FDSLESLVDETIDIFKIKSI-----------NDRLTANLYK-------YVVQDN-YFGQVSFLN--------VL--KA-------KL-DLDG------LKQVMFNDY-LRQIIYFG------ELEDLLEENDYDKLKNYLNDRLNSIAKDINEKKV--SKSSINTIEKIMKEINKNFIKKNK--SIEDIKEY--MDSLE---VCEMCGLYK--------------------GILDEYSESNFA--PLGVSTN--N--ARNMF-------------------W------N-----
QEYVSSICDVCKLILFCTPAGAT-----------Y-----TRKNYLINEENEF-------------YLF-----VNMDT-------SINGLFE---RNNS------LKA-----------------QKSESSD-SKDENPFNQLI-----KSIV-EENT---------LK-SEWQLR-NILFVE-FKAS-ID-------SKKCK-INYFNIPTYLAKFFVKE--------NKK-------------IQ--SIYDYK---------------------------------------------------------------LKSNVLDLILKS-----KDLKHLINNT-
LRNIVKSNLESNNK----------------SNISGLSCF-----NVVQLRTILNNYK--------RGVY------------KMD------YK------------------------------------KLRKIYESGCEIHDYYIDKNA---------------------------KNKIEGITYKIL-NLIKVGNKNDFMQ-TILRIFSP-----S-----------EKIPP-EEILQIFI-------EED-----L-DFESVGYAFVTGLISGKYEGKDKLP------NE-----KEEK------------------------------

SUY82599 -------------------------------------------------------------------------------------------------------------------MKEINKNF-----IK--------------------------------------------------------------------------------------------------------------------------------------------------------------------------------------------------------------KN--KS-------I----------------EDIKEY-
MDSLE-----------------------------------------------------------------------------------------VCEMCGLYK----------------------GILDEYSESNFA--PLGVSTN--N--ARNMF---------------W------NQAYV-------PSICDICKLILFCTPAGAT-------YTRKNY---------LI-NEENEF--------------YLF-----VNMDT-------CINELFE---RNNS------LKV---QKS----------------ESSD-SKDENPFNQLI-----KSIV-EENT---------LK-SEWQLR-
NILFVE-FKAS--IDS-----KKCK--INYFNIPTYLAKFFTDK-------YANKK----------------IQ--SIYDYK-----L-------------------------------------------------------KS-NVLDLLLKN-----RDLKHITNNI-LRNKV-------------------------------------------------------KNDMESNNK---------SNIS-----------GID------------CF----------------------------RVVQLR--------------------------------------------------------------------------VLINSYKK-----------------GYIKW-IV---------K-----
------IWRRMMKS-----------------------------------------------------------------

SJV46102 --------------------------------MKLRVYRGD---W--------FFNMGIVGFLNI-------------------------------IKKADKQAE--------IF-----IMED--YIEFDSLF------LE----NFHEYYFNYFMDE-----------YDVSKRIKKNIDYSINFIKSKPDRIKDGIKKIKDSV-----------KQQNDKIKKFD---------------------------------------EEKFNLIKEKLDSMSKIKSYDE-------------
FDSLESLVDETIDIFKIKS-----------INDRLTANLYK-------YVVQDN-YFGQVSFFN--------VA--KA-------KL-DLDG------LKQVMFNDY-LRQIIYFG------ELEDLLEENDYDKLKNYLNDRLNSIAKDINEKKV--SKSSINTIEKIMKEINKNFIKKNK--SIEDIKEY--MDSLE---VCEMCGLYK--------------------GILDEYSESNFA--PLGVSTN--N--ARNMF-------------------W------N-----
QEYVSSICDVCKLILFCTPAGAT-----------Y-----TRKNYLINEENEF-------------YLF-----VNMDT-------SINELFE---RNNS------LKA-----------------QKSESSD-SKDENPFNQLI-----KSIV-EENT---------LK-SEWQLR-NILFVE-FKAS-ID-------SKKCK-INYFNIPTYLAKFFTDK-------YANKK----------------IQ--SIYDYK---------------------------------------------------------------LKSNVLDLLLKN-----RDLKHLTNNI-
LRNKVKNDMESNNK----------------SNISGIDCF-----RVVQLRVLINSYK--------KGVY------------KMD-----SKNLEKNDE---------------------------KLKIIYYLGCDIHDYFVNKNS---------------------------KNKIDGVSYKLL-NSVKVGNKSDFMD-TIIRVFMS-----AE-----------KQIPA-FILDIEI--------EKD-----L-DFESIGHAFISGLISGKYEGKDKLP------NE-----KEEK------------------------------

SUY19348__3_ --------------------------------MKLRVYRGD---W--------FFNMGIVGFLNI-------------------------------IKKADKQAE--------IF-----IMED--YIEFDSLF------LE----NFHEYYFNYFMDE-----------YDVSKRIKKNIDYSINFIKSKPDRIKDGIKKIKDSV-----------KQQNDKIKKFD---------------------------------------EEKFNLIKEKLDSMSKIKSYDE-------------
FDSLEALVDETIDIFKIKS-----------INDRLTANLYK-------YVVQDN-YFGQVSFFN--------VA--KA-------KL-DLDG------LKQVMFNDY-LRQIIYFG------ELEDLLEENDYDKLKNYLNDRLNSIAKDINEKKV--SKSSINTIEKIMKEINKNFIKKNK--SIEDIKKY--MDSLE---VCEMCGLYK--------------------GILDEYSESNFA--PLGVSTN--N--ARNMF-------------------W------N-----
QAYVPSICDICKLILFCTPAGAT-----------Y-----TRKNYLINEENEF-------------YLF-----VNMDT-------CINELFE---RNNS------LKA-----------------QKSEYSD-SKDENPFNQLI-----KSIV-EENT---------LK-SEWQLR-NILFVE-FKAS-ID-------SKKCK-INYFNIPTYLAKFFTDK-------YANKK----------------IQ--SIYDYK---------------------------------------------------------------LKSNVLDLLLKN-----RDLKHLTNNI-
LRNKVKNDMESNNK----------------SNISGIDCF-----RVVQLRVLINSYK--------KGVY------------KMD-----SKNLEKNDE---------------------------KLKIIYYLGCDIHDYFVNKNS---------------------------KNKIDGVSYKLL-NSVKVGNKSDFMD-TIIRVFMS-----AE-----------KQIPA-FILDIEI--------EKD-----L-DFESIGHAFISGLISGKYEGKDKLP------NE-----KEEK------------------------------

CKG87449 --------------------------------MKLRVYRGD---W--------FFNMGIVGFLNI-------------------------------IKKADKQAE--------IF-----IMED--YIEFDSLF------LE----NFHEYYFNYFMDE-----------YDVSKRIKKNIDYSINFIKSKPDRIKDGIKKIKDSV-----------KQQNDKIKKFD---------------------------------------EEKFNLIKEKLDSMSKIKSYDE-------------
FDSLEALVDETIDIFKIKSI-----------NDRLTANLYK-------YVVQDN-YFGQVSFFN--------VA--KA-------KL-DLDG------LKQVMFNDY-LRQIIYFG------ELEDLLEENDYDKLKNYLNDRLNSIDKDINEKKV--SKSSINTIEKIMKEINKNFIKKNK--SIEDIKEY--MDSLE---VCEMCGLYK--------------------GILDEYSESNFA--PLGVSTN--N--ARNMF-------------------W------N-----
QAYVPSICDICKLILFCTPAGAT-----------Y-----TRKNYLINEENEF-------------YLF-----VNMDT-------CINELFE---RNNS------LKV-----------------QKSESSD-SKDENPFNQLI-----KSIV-EENT---------LK-SEWQLR-NILFVE-FKAS-ID-------SKKCK-INYFNIPTYLAKFFTDK-------YANKK----------------IQ--SIYDYK---------------------------------------------------------------LKSNVLDLLLKN-----RDLKHITNNI-
LRNKVKNDMESNNK----------------SNISGIDCF-----RVVQLRVLINSYK--------KGVY------------KMD-----SKNLEKNDE---------------------------KLKIIYYLGCDIHDYFVNKNS---------------------------KNKIDGVSYKLL-NSVKVGNKSDFMD-TIIRVFMS-----AE-----------KQIPA-FILDIEI--------EKD-----L-DFESIGHAFISGLISGKYEGKDKLP------NE-----KEEK------------------------------

SJV33434__7_ --------------------------------MKLRVYRGD---W--------FFNMGIVGFLNI-------------------------------IKKADKQAE--------IF-----IMED--YIEFDSLF------LE----NFHEYYFNYFMDE-----------YDVSKRIKKNIDYSINFIKSKPDRIKDGIKKIKDSV-----------KQQNDKIKKFD---------------------------------------EEKFNMIKEKLDSMSKIKSYDE-------------
FDSLEALVDETIDIFKIKSI-----------NDRLTANLYK-------YVVQDN-YFGQVSFFN--------VA--KA-------KL-DLDG------LKQVMFNDY-LRQIIYFG------ELEDLLEENDYDKLKNYLNDRLNSIDKDINEKKV--SKSSINTIEKIMKEINKNFIKKNK--SIEDIKEY--MDSLE---VCEMCGLYK--------------------GILDEYSESNFA--PLGVSTN--N--ARNMF-------------------W------N-----
QAYVPSICDICKLILFCTPAGAT-----------Y-----TRKNYLINEENEF-------------YLF-----VNMDT-------CINELFE---RNNS------LKV-----------------QKSESSD-SKDENPFNQLI-----KSIV-EENT---------LK-SEWQLR-NILFVE-FKAS-ID-------SKKCK-INYFNIPTYLAKFFTDK-------YANKK----------------IQ--SIYDYK---------------------------------------------------------------LKSNVLDLLLKN-----RDLKHITNNI-
LRNKVKNDMESNNK----------------SNISGIDCF-----RVVQLRVLINSYK--------KGVY------------KMD-----SKNLEKNDE---------------------------KLKIIYYLGCDIHDYFVNKNS---------------------------KNKIDGVSYKLL-NSVKVGNKSDFMD-TIIRVFMS-----AE-----------KQIPA-FILDIEI--------EKD-----L-DFESIGHAFISGLISGKYEGKDKLP------NE-----KEEK------------------------------

SUY82598 ------------------------------------------M-------------------------------------------------------------------DSKN------LE------------------------------------------------------------------------------------------------------------------------------------------------------------------------------------------------------------------------------------------------------
--------------------------------------------------------------------------------------------------K--------------------------N-----DE--------------------------------------------------------------------------------------------KLKIIYY-------------------------------------------------------------------------------------------LGCDIHDYFVNK------------------------------------------------------------------------------------------NS-------
-----------------------------------------------------------------------------------------------------------------------------------------------KNKIDGVSYKLL-NSVKVGNKSDFMD-TIIRVFMS---------------AEKQIPA-FI-LDIE-----I--EKD-----L-DFESIGHAFISGLISG--KYEGK--------DK-----LPNE-K----EEK--------------------

SJV07142__2_ --------------------------------MKLRVYRGD---W--------FFNMGIVGFLNI-------------------------------IKKADKQAE--------IF-----IMED--YIEFDSLF------LE----NFHEYYFNYFMDE-----------YDVSKRIKKNIDYSINFIKSKPDRIKDGIKKIKDSV-----------KQQNDKIKKFD---------------------------------------EEKFNMIKKKLDSMSKIKSYDE-------------
FDSLESLVDETIDIFKIKSI-----------NDRLTANLYK-------YVVQDN-YFGQVSFFN--------VA--KA-------KL-DLDG------LKQVMFNDY-LRQIIYFG------ELEDLLEENDYDKLKNYLNDRLNSIAKDINEKKV--SKSSINTIEKIMKEINKNFIKKNK--SIEDIKEY--MDSLE---VCEMCGLYK--------------------GILDEYSESNFA--PLGVSTN--N--ARNMF-------------------W------N-----
QAYVPSICDICKLILFCTPAGAT-----------Y-----TRKNYLINEENEF-------------YLF-----VNMDT-------SINELFE---RNNS------LKD-----------------QKSEYSD-SKDENPFNQLI-----KSIV-EENT---------LK-SEWQLR-NILFVE-FKAS-ID-------SKKCK-INYFNIPTYLAKFFTDK-------YANKK----------------IQ--SIYDYK---------------------------------------------------------------LKSNVLDLLLKN-----RDLKHLINNI-
LRNKVKNDMESNNK----------------SNISGIDCF-----RVVQLRALINSYK--------KGVY------------KMD-----SKNLEKNDE---------------------------KLRIIYYLGCDIHDYFVNKNS---------------------------KNKIGGVSYKLL-NSVKVGNKSDFMD-TIIRVFMS-----AE-----------KQMPA-FILDIEI--------EKD-----L-DFESIGHAFISGLISGKYEGKDKLP------NE-----KEEK------------------------------

SJV64910__5_ --------------------------------MKLRVYRGD---W--------FFNMGIVGFLNI-------------------------------IKKADKQAE--------IF-----IMED--YIEFDSLF------LE----NFHEYYFNYFMDE-----------YDVSKRIKKNIDYSINFIKSKPDRIKDGIKKIKDSV-----------KQQNDKIKKFD---------------------------------------EEKFNLIKEKLDSMSKIKSYDE-------------
FDSLEALVDETIDIFKIKSI-----------NDRLTANLYK-------YVVQDN-YFGQVSFFN--------VA--KA-------KL-DLDG------LKQVMFNDY-LRQIIYFG------ELEDLLEENDYDKLKNYLNDRLNSIAKDINEKKV--SKSSINTIEKIMKEINKNFIKKNK--SIEDIKKY--MDSLE---VCEMCGLYK--------------------GILDEYSESNFA--PLGVSTN--N--ARNMF-------------------W------N-----
QAYVPSICDICKLILFCTPAGAT-----------Y-----TRKNYLINEENEF-------------YLF-----VNMDT-------CINELFE---RNNS------LKA-----------------QKSEYSD-SKEENPFNQLI-----KSIV-EENT---------LK-SEWQLR-NILFVE-FKAS-ID-------SKKCK-INYFNIPTYLAKFFTDK-------YANKK----------------IQ--SIYDYK---------------------------------------------------------------LKSNVLDLLLKN-----RDLKHLINNI-
LRNKVKNDMESNNK----------------SNISGIDCF-----RVVQLRALINSYK--------KGVY------------KMD-----SKNLEKNDE---------------------------KLRIIYYLGCDIHDYFVNKNS---------------------------KNKIGGVSYKLL-NSVKVGNKSDFMD-TIIRVFMS-----AE-----------KQMPA-FILDIEI--------EKD-----L-DFESIGHAFISGLISGKYEGKDKLP------NE-----KEEK------------------------------

SUY11630__2_ --------------------------------MKLRVYRGD---W--------FFNMGIVGFLNI-------------------------------IKKADKQAE--------IF-----IMED--YIEFDSLF------LE----NFHEYYFNYFMDE-----------YDVSKRIKKNIDYSINFIKSKPDRIKDGIKKIKDSV-----------KQQNDKIKKFD---------------------------------------EEKFNLIKEKLDSMSKIKSYDE-------------
FDSLEALVDETIDIFKIKSI-----------NDRLTANLYK-------YVVQDN-YFGQVSFFN--------VA--KA-------KL-DLDG------LKQVMFNDY-LRQIIYFG------ELEDLLEENDYDKLKNYLNDRLNSIAKDINEKKV--SKSSINTIEKIMKEINKNFIKKNK--SIEDIKEY--MDSLE---ACEMCGLYK--------------------GILEEYSESNFA--PLGVSTN--N--ARNMF-------------------W------N-----
QAYVPSICDICKLILFCTPAGAT-----------Y-----TRKNYLINEENEF-------------YLF-----VNMDT-------SINELFE---RNNS------LKA-----------------QKSEYSD-SKDENPFNQLI-----KSIV-EENT---------LK-SEWQLR-NILFVE-FKAS-ID-------SKKCK-INYFNIPTYLAKFFTDK-------YANKK----------------IQ--SIYDYK---------------------------------------------------------------LKSNVLDLLLKN-----RDLKHLINNI-
LRNKVKNDMESNNK----------------SNISGIDCF-----RVVQLRALINSYK--------KGVY------------KMD-----SKNLEKNDE---------------------------KLRIIYYLGCDIHDYFVNKNS---------------------------KNKIGGVSYKLL-NSVKVGNKSDFMD-TIIRVFMS-----AE-----------KQMPA-FILDIEI--------EKD-----L-DFESIGHAFISGLISGKYEGKDKLP------NE-----KEEK------------------------------

CED93815__2_ --------------------------------MKLRVYKND---W--------FFNMGIVGFLNI-------------------------------LNKAEVKNQ--------VA-----ISED--YIEFESCL------LQ----NFHHYYFDYFMDE-----------YDVYTRVKNGIEYSIKYVKAHPDKIKDSAKKIKEIL-----------KKQNDKVKKID---------------------------------------EKNYEIIKEKLDLIGKLKKEDE-------------
VEELENISKECLDIFKLKHI-----------NDRLTVNLYK-------FIIGDN-YFGQTSFFN--------VN--KS-------KY-DLDG------LKNVMYNDY-LISIVYFG------ELQSLLNEGCIETLEKYITEKLDYINKELESKRI--SKPSIKIIEKIMKDINSKFIKKKK--NIEDIKMY--LESLE---TCEMCGTYK--------------------GLINDYSESNFA--PLGVSND--N--AKNMF-------------------W------N-----
QDSTYSICDLCKLILFCTPAGAT-----------Y-----IRKNYITDENNEF-------------YSF-----VNIDT-------SIFDIYN---TNIN------LKD------------------------L-KDRENPFNDLV-----IDIV-TENK---------DKSIWKLQ-NILFVE-FKAS-IE-------AKKCK-MNYFNMPTYLAKFFVNE---DGKNSKL----------------IQ--SIYNQK--------------------------------------------------------------FKGNVVDILLKN-----RDLKHLINIS-LRERIKENLEDSKK---
----------IKISTSDCYK-----AIKVRALINSYK--------KGVY----------GMN-----------DK--------------------------KLKVVKYAGHEIHDYYVNNNA----------------------------KNKINGVAYKLL-NSIKVGNKKDFMD-TVLRIFMS-----AE-----------KSVPS-VF-IEIMA-------EKD-----L-DFESIGHAFITGLISEKYEA-----------KN-----DDNK-----------------------------

WP_029878088 --------------------------------MKLRIYKGD---W--------FFNMGIVGFLNI-------------------------------LHEAEVKNQ--------VI-----INED--YIEFDSLL------LE----NFHNHYFDYFMKE-----------YDVSKRVENSINYSLNYIKANPEKIKESIKKIKDNL-----------KKQNDKVKKFD---------------------------------------EINYINIKDKLDTIGKIKKSNE-------------
IEQLEQLTRECVDILKIKSI-----------NDRLTANLYK-------FIVGDN-YFGQVSFFN--------VA--KS-------KL-ELEG------LKQVMFNDY-LISIIHMG------ELESLIDKEDLDSLEKYITEKLEYLNKETIEKRI--SKASIKTIEKIMKDIKSKFIKKKK--SIEDIKAY--MGGLE---TCEMCGTYK--------------------GLINDYSESNFA--VLGVSND--N--AKNMF-------------------W------N-----
QDSTYSICDICKIILFCTPAGAT-----------Y-----TKKNYIENEDNEF-------------YSF-----VNMDT-------SIDDIYN---TNIN------LKK-------------------------L-KDRDNPFNDLV-----IDIV-SENK---------EKSEWQLQ-NILFVE-FKAS-IE-------AKKCK-MNYFNMPTYLAKFFVNE------KNSKL----------------IQ--SIYNQR--------------------------------------------------------------FKANVVDILLKN-----KDLKHLISKV-LRDKI--------------------
KDSLEDGYKNNISLNDIFKAIKIRALIDSYK--------EGVS-----------KMN------------DK---------------------------KLQVVKFAGHEIHDYYVNTNS----------------------------KNKINGVAYKLL-NAVKVGNKKDFMD-TVLRVFMS-----AE-----------KSVPS-VF-IEIMA-------EKD-----L-DFESIGHAFITGLISE----------KY-----EANK-----------------------------

WP_057567909 --------------------------------MKLRVYKSD---W--------FFNMGIVGFLNI-------------------------------LYKAEVKKQ--------VT-----INED--YIEFDSFL------LN----DFHNYYFDYFINE-----------YDVSKRVESSIGYSLKYVKSNPEKIKDGAKRIKDNL-----------KKQNDKIKKFD---------------------------------------EKNYNIIKEKVDIIGKLKKAEE-------------
VEQLETLAKECIDIIKIKSI-----------NDRLTANLYK-------FIIGDN-YFGQVSFFN--------VA--KS-------KL-DLEG------LKQVMYNDY-LISIVYMG------ELKDLIENEDLNNLEKCITEKLDYLSKETSKKRI--SKASIKTIEKIMKDINSKFIKKKK--SIEDIKKY--FDSLE---VCEMCGTYK--------------------GLINDYSESNFA--PLGVSND--N--AKNMF-------------------W------N-----
```

QDSTYSICDICKLILFCTPAGAT----------Y----TKKNYIENEDNEF-----------YSF----VNTDT-----SIDDIYD---TNMN-----LKK-------------------L-KDKDNPFNDLV----IDIV-SENK--------EKSEWQLQ-NILF[illegible]
LRTKIKGNLEEEHK--------------TNISSSDCF----KSIKIRALIDSYK-------KGVY---------EMN----------DK----------------------KLQVVKFAGHEIHDYYINTNS-------------------------KNKINSVAYKLL-NAIKV[illegible]

SIT17694 ------------------------------MKVRIEPND---W--------TMLMGIVGVHNI---------------------------VQFAKKNNIMNAKDTRRIK----VEHD--ALVLDPDA-[illegible]
-----QAEQLEELQQLTDQYLKVLAEPLINRK-------------LTVNYFK------TAVMQP-FFGQPSFLQ-------AT--AN------SL-SYEE-----HIDRFQQDY-ILPVLLEA-----KMRTIVEQASTSNEVLAFLDEYQDYSP[illegible]
KDETIPISSLAKLILFCAPAGAA---VYLRKDGF--------GEQGEYRTY-----------MGF----VQSDS-----PLPDVIR---KNRY-----FQIA---------------------KQGMEPFDRVV----SRMI-EDVK--------QKAGFVAD-[illegible]
--------------ESGYSGWSAY----VATRERYRMRQYA-------KHLG---------EGG-----SAMDAKEKRVR--------------------AVYRVYRSGADIREKFNQRGPREGAPEGQYAASG--------------AKKISGIAYRLL-N[illegible]

SDH76205 ------------------------------MVLRLYASD---W--------LMSMALVGLQRL--------------------------YIYGEKHNIIPDEHRGTVR----ETDS--GLEIEPVA--[illegible]
---LEELQVCIHDYYNVVKQKEI----------NDKLTLNYFK------SVILSP-FFGQPSFLN-------VA--KN------SL-DLSG-----HIDLFYKDY-VLPVQNDI-----QLLEVINKTETGEDVLAFLDEHADYSPFRTLKKAW-[illegible]
-MYYRKDGF--------DDQGEVRLY-----------AGF----VHTDE-----KFKEIFQ---KNEH-----FKQL---------------------KKQSQSFEKIV----SRLV-RGIR-------KE-SEYIMQ-HLFFLE-MSAN--------YQSKRTLLD[illegible]
IAVRERHRIECYK-------RGEK---------DMK---------KQDN-----------------------RVAAFFNSGRDIRKAF-EQAGKARESEDTYSASA--------------GKKVAGIAYRLL-NATKANNKDSFMD-TLLRVYMS----VD[illegible]

KGP59893__2_ ------------------------------MKVELHMGG---W--------MINMGLVGLYRV--------------------------FEYGRKHEIISEEHKHSVT----VKPW--GIELDTEV[illegible]
VHQLEEHIRTLK-QVYSEPK---------INEKLTFNYFK------HAILKPLFFGQVSFLN-------VS--KN------NL-DFEG-----HVEEFYNSY-IFPVVNDL-----YLESVINDSHSMEEISSLLEKQSEYQLFRQLKRAW-[illegible]
GEQPRPLSSLAKLVLLFAPIGSTIYY-K--NEGI--------NEQTERRLY-----------AGF----VQTDA-----TFAEVLQ---KNNH-----FRE---------------------L-KKKREPFNKIV----SQLV-QSVK-------KE-SEYTVD-H[illegible]
----------KNERSTNG-PYI-------AVRERNRIL--------QFKK---------GVR---DMA---TTDK-----------------------RVTALFMQGREIRDAI-IR-----------SSGS--------------EKKVTSIAYRLL-NAAKAGNRKSFMD-T[illegible]

SFA58177 ------------------------------MKVELRMGE---W--------MVNMGLVGLYRV-------------------------FEYGQKNGIISDEYKDSVT----VKPW--GIELDTD[illegible]
----VHQLEEHIRTLK-QVYSETK---------INEKLTLNYFK------HAILKPLFFGQVSFLN-------VS--KN------DL-DFEG-----HVKEFYNSY-VFPVMNDL-----HLESVLNDSHSIEEISSLLEKKGEYQPFKQLKRE[illegible]
SQQPKPLSSLAKLVLLFAPIGVAIYL-K--NDGI--------GDQKERRPY-----------AGF----VQTDG-----NFAEILQ---KNNH-----FKQ--------------------L-KKKREPFNKIV----SQLV-QSVR-------RE-SEYAVD-[illegible]
----------KNERSSIG-PYI-------AVRERNRIL--------QFKK---------GVR---DMAMTDKTDK-----------------------RVSALFMQGREIRDAI-MR-----------SSGS--------------EKKVTSIAYRLL-NAAKAGNRKNFMD[illegible]

KFZ42528 [illegible]

WP_044731094__2_ [illegible]

ADI26429__2_ [illegible]

WP_025948828 [illegible]

KLR73406__2_ [illegible]

ATA60324 [illegible]

WP_047758566 [illegible]

KMY62737__3_ [illegible]

WP_044746560 [illegible]

AEH47542__2_ [illegible]

TCL52009 [illegible]

ACS24737__2_ [illegible]

SFA50051 [illegible]

ADP74262__2_ [illegible]

SMG36792 [illegible]

SFE52423 [illegible]

EGL83969 [illegible]

SFA87700 [illegible]

ERI07860 [illegible]

SCL93935 [illegible]

KMT48371 [illegible]

SCN37353 [illegible]

ABS22299 [illegible]

KFN04732 [illegible]

SFQ91018 [illegible]

TCS77239 [illegible]

ADG71319 [illegible]

TCW02978 [illegible]

WP_118757835 [illegible]

EDO58922 [illegible]

WP_008118711 [illegible]

AFLCPACAFVYALSPLGFT---------L--------LGNR---------------FAF----INTND-----SIKGLLT---ANDK-----ERR [illegible]
----DNPGCISS-------AFWVYMVMLWS------NIVT---------GNK-----ENGGNVEM---------------------NRVAVMNSGFA [illegible]

CDB20969 -------------------------MEKYAITPGD---F--------LKNAGIV [illegible]
---------MTAEELLERLSKLQ-VFLEQEK---------CKETFIMKSVV------YNYINR-FWDGKSFLL-------RS--NA------KK-D [illegible]
K-------------FVF----VNNNS-----DMFTLLA---ANSK-----NRK--SSLEGE--------------KEENQRYSQWF----AETL-NII [illegible]
--------KEE----------EKEVLM-----------------YCISMKKSGSELRKAI-MTA-KKDTT-------------------GDSLRGIEYQ [illegible]

CBL20527 [illegible]
VFEKDF-SEPFRKYL-----------------------------------------------------SAD--HKKAKD-----------LCIDCGDPI [illegible]
KQKELS-NVQVIL-RGVS--ADDR---------YVLSIIHRRIIEIIREP-----RISDA------------LT--ALAKAP----V [illegible]
NALAVRNVNKFMD-VVMRLYSA----YGSKRNEDGHE-LLIPT-GF-VQML----N--DKQ------KFTEYGYAFVMGLVG [illegible]

CDC93207 [illegible]

EEG37130 [illegible]

EEX23204__2_ [illegible]

AGI39866 [illegible]

ADU97573 [illegible]

EEP60017 [illegible]

ACD66304 [illegible]

WP_029521652 [illegible]

WP_029520046 [illegible]

ACN98336 [illegible]

SMC08868 [illegible]

ACO02971 [illegible]

RMA96971 [illegible]

ACI19672 [illegible]

ACM59280__3_ [illegible]

ABP67977 [illegible]

ERM91451 [illegible]

ADL41486 [illegible]

KUJ90573 [illegible]

ADH59814 [illegible]

KUK34308 [illegible]

KHO63186 [illegible]

ADQ08188 [illegible]

ADQ41858 [illegible]

KHO63184 [illegible]

KLA44239__3_ [illegible]

EEW67155__2_ [illegible]

WP_050612262 [illegible]

WP_089145121 [illegible]

AGI39652 [illegible]

PZX05806 [illegible]

KOY83702 [illegible]

EKU43637 [illegible]

-------------GLF----INSNQ-----SIVELLA---YNKI-----QLRN-------------DRGEEYTRD-KLEEMAYSKII----NEMY-QHAE--------NK-KNVEID-NIQIVK-YDKN----------NESRP-YTFNMLSRDKAQQLYDNRNDFHYLTGK-------------------FVKENGDYISL----------------------------------------YQ-EVIRRFYNG----QRFSDLLYRL-IRLVI--------------------------DDKYTDTFS---------
IYKILKISNYVLSGGSHYMT-----------------------DK-----------------------ELRSIRISGYELRKVY-RGQ--------------------------ETKLNAISHRLL-NALKVKNPNKFME-TIVQAYSY----KRHEK---------LGIPL-FFTTTLS-------DIE-------KFQTIGYAFLIGLNGKDFKKEDMAK-----EE----QVNE-------------------------

EFI68562 --------------------------------MEKIQPQIGD---------------AVDALLQI-------------------------------------------------------------VE---HLKHKDAKRFIIP-------------------------------------------------------------------------------------------RVLT---------------------------------YQVIQG-FWSNVSFLN-------SN--
AS------KK-DLYE---------EYFNYF-IKPAQDFL---------------------------------------------------TSKKDEKKYGKNKL----TCACCDNKMR-------------------SISEAFDLTWVQ--NLGVDGA--R--KSSHY---------------W------N------HQRNLFICPICNIVYSCLPLGFT-----------L---------KNGI------------------GLF----INSNQ-----SIVELLA---YNKI-----QLRN-------------DRGEEYTRDKLEEMAYSKII--
--NEMY-QHAE---------NKKNVEID-NIQIVK-YDKN----------NESRP-YTFNMLSRDKAQQLYDNRNDFHYLTGK-------------------FVKENGDYISL----------------------------------------YQ-EVIRRFYNG----QRFSDLLYRL-IRLVI--------------------------DDKYTDTFS---------IYKILKISN-----YVLS------GGS--------HYMTDK-----------------------ELRSIRISGYELRKVYRGQ-----
ETKLNAISHRLL-NALKVKNPNKFME-TIVQAYSY-----------KRHEKLGIPL-FFTTTLS-------DIE-------KFQTIGYAFLIGLNGKDFKKEDMAK-----EE----QVNE-------------------------

EES51331 ----------------------------MSLVKIEIGD---W---------LKNASIVGLLKV-------------------------------LEHKKQSINE--------VN-----IQKN---YIEFDTSM------LE---GFEEAYFQVLFKE-------------HGKSISWFKLVSYM--DEVNFMAENETTEKSIDRLN------------KIIDDMKSKSL------------INSNSYKSAYELL-------
DDENDKIWIANKEKELIKIKHKKKYNPLDFKEEVLEQIAIIKDIINWFKKPE-----------------------IKKFIMAKNVM--------YEVIQP-FWNNVSILL--------KT--NN--------KS-NMYE---------LYKQDF-IDNFINYS------------------------------------------------------------ESDTSKNK--Y---------NCFTCNNKIKKL----------------------SKPEAFDLTWLV--KTGVDMS--R--KSSHF---------------W------N--MN-GD--
AYICPICNLVYSCLPLGFI-----------V-----------KKKK----------------------GIF----INNNQ------NIKALKQ----CNIV-------ATN-------------------YEDKKYE-EIENLSYNIF----NSMENVSIE-----------NIDKEFE----NIQIVK-IDGN---------NDRRP-YSFNILSLRLMKILYFHRKNLESLIK------------------IR-VKITEKY-----YLNL-------------------------------------------------YD-EVIRRIYDG-------KNLLDLLIHKL-LIMYL-----------------------
DDKKFKGINI---------IYKILKINS-----------SLMG------------GKD-----------MRQE----------------------DINNFRNYGIRLKEAY-EKK-NA-----------------------------KSKLSGISYRLL-NAIKTKDTGKFMD-TILNAHMY-----MR-----------MELPK-ELGLALR---------DND--------KLQNIGYAFVLGLQSE---------------SLKKE-----HEGE-N----------------------

KPU26824 ----------------------------MSKIRLELSD----W---------LYNAGIVGANI-------------------------------LEYSEKKFEKKLNYIEFDEKL-------LE---NFEEMYFKYFIDK-------------YLKFTSWYKIVSFEDYIMNFNEDEIEKEDIKEKINSY------------IEYTKKRLTSS------------------------------SYKSGYSIIKFGRLDLLKEEKKLKKIKKTKKQTI-------
KDIIPNIKGQLDIISNIIYFLKEDEV----------KRIILAKNII-------YDVIAK-FLKDVSFLH--------KN--AN-------KN-DMYE----------EYKNYF-INSIISYI------------------------------------------------------------------------HEDKNNFSY------------NCFNCNRKISKL--------------------SKPASYDLTWLN--KIGVDMS--R--KTSHF---------------W------N--FY-GD--TFICPICNLVYSCIPAGFT----------V---------VRDK--------------------------------
GLF----INQNS-------DVKTLIS-----INSI------AID-------------------HNTKFEELEEKSYYAVS------ENLK-QSVV---------ENMEKEID-NIQIVK-LDSN----------NDRRP-YTFNVLSKNKLKIIYENKKLLRYLVNN------------------YVKTDAGDYIN------------------------------------------VYSEVIKRLYDN------KNQFDLINKL-LYLNM---------------------------------NEKFTRIGN-----------IKIIIKINN--------DYLG-----------GIM-----
KGKMISYN---------------------TIDKLQQLGFELRKAYEVKNS---------------------KNKINGISYRLL-NALKTKNTNKFMD-TLINAYMY-----LN--------------KEIPI-EF-VDGL-------KDE-----D-RFQTLGYSFLLGLQGESEKRDSNKENE-----GD-----EINE--------------------------------

SHH83075 ---------------------------MREISRIRLELSD---W---------LYNAGIVGIANI-------------------------------LEHSGKFEKKSNYIEFDEEL-----LE---NFEEMYFKYFIDK-------------YLKFTSWYRIVSFEDYIINFNEGEITEKDIEKINNH-----------------------------------------IQYTKKKLTSSSYKSGYSIIKSSKIDLLKEEKKLKKNKKQTI--------
KDIIPNIKEQLDVISDIIDFLKADEV------------KRVILAKNII-------YDVIAK-FIKDVSFLH--------KN--AN-------KN-NMYE----------EYKNYF-INSIIEYI------------------------------------------------------------------------NNEDNSNSNY---------NCFNCNRKISKL-------------------SKPASYDLTWLN--KIGVDMS--R--KTSHF---------------W------N--FY-GD--AFVCPICNLVYSCIPAGFT---------V------------VRDR---------------------------
--GLF----INQNS-------DVKTLIS-----INNI-------AID---------------HNTRFEELEEKSYYAVS------ENLK-QSVV----------ENMEKEID-NIQIVK-LDSN----------NDRRP-YTFNVLSKNKLKIIYENKKLLRYLVNS-----------------YVKTDIGDYINI-------------------------------------------------YS-EVIKRLYDN-------KNQFDLINKL-LYLNM-----------------------------------NEKFAKIGN-------------IKIIIKINN---------DYLG------------GIM-----
KGKMISYN-------------------------TIDKLQQLGFELKKAYEAKNS-------------------KNKISGISYRLL-NALKTKNTDKFMD-TLINAYMY-----LN--------------KEIPI-EF-IEGL--------KDE-----D-RFQTLGYSFLLGLQGENGKKELDKENE----GD-----EING--------------------------------

KGG79724 ----------------------------MSRIRLELSD----W---------LYNAGIVGIANI-------------------------------LE------HSGKKFEKKSNYIEFDEEL----LE---NFEEMYFKYFIDK-------------YLKFTSWYKIVSFE--DYIINFNEGEITEKDIEKIN-------------NHIEYTKKKL-------------TSSSYKSGYSII----------KSSKIDLLKEEKKLKKNKKQTI-------
KDIIPNIKEQLDVISDII--DFLKAD---------EVKRVILAKNII-------YDVIAK-FIKDVSFLH--------KN--AN-------KN-DMYE----------EYKNYF-INSIIEYINNEDNSN-------------------------------------------------------------------------------YSNY-----------NCFNCNRKISKL--------------------SKPASYDLTWLN--KIGVDMS--R--KTSHF---------------W------N--FY-GD--AFVCPICNLVYSCIPAGFT---------V----------VRDR----------------------
--GLF----INQNS-------DVKTLIS-----INNI-------VID--------YNTRFE----ELEEKSYYAVS------ENLK-QSVV---------EN-MEKEID-NIQIVK-LDSN----------NDRRP-YTFNVLSKNKLKIIY-------KKL-------------LR--YLVNSY-------VKTDTGDYINV--------------------------------------------YS-EVIKRLYDN-------KNQFDLINKL-LYLNI-------------------------------------NDNFSK-ISN---------IKIIIKINN--------DYLG-----------GIM-----
KGKMISYN-------------------------TIDKLQQFGFELKKAY-EAK-NS----------------KNKISGISYRLL-NALKTKNTDKFMD-TLINAYMY-----------------------LN-KEIPS-EFIEGLK-------------SGKKELDKENEG-----DEING------------------------------

SHK58591 ----------------------------MKKIEKVRLELSD----W---------LYNAGIVGLCKI-------------------------------------------------------FDTSDVNYKADDNYIEFDETA------LE---NFEEKYFKYFIDK-------------YISFTSWYKIVKFEDKLIRFDENEVDERYVKDLNEQIEYMKSKLSASYKSAYQIID---------------------------------DKEIDLLKEEKKLKKVSKTKKQSI--------
KDILPKIKKQNEVIGGIIKYLKKDNV-----------KKIILGKNVI-------YDIITK-FWSDVSFLH--------KNSSKN-------DMYL----------EYKNYF-LDNMINYL----------------------------------------------------------------------------KSKKDNFKY------------SCFSCDRLISNL---------------------SKPNSYDLTWIN--KMGVDMS--R--KTSHF---------------W------N--FY-GD--IFICPICNLVYSCIPAGFT------------------IRDK--------------------------
--GIF----INQNS------DINTLIK----VNNT------AID-------------------NNTSFQELEEKTYFAIV----DNLNQSQIE----------RAEKEID-NIQIVK-LDSS----------NSTRP-YTFNIMSKKKLRIIKNNKKLLRGLVNV-----------------------SV--KIGRKNGRDIYIN----------------------------------------------LYTEVINRLYDD------KKLFDLISKL-FYLNL-------------------------------------DDKFKKLWN-----------LKTIIKIND-----------SYIG-------------GIM------
GRGIYYK----------------------EIDQIQQIGFDLRKAYKEKNA----------------------KNKINGISYRLL-NALKTKNSSKFME-TLFNAYMY-----LN------------KAIPV-KF-IDVAG-----------DED-------EKFQTIGYSFLLGLQGEIGKLDFDEKECSDNE-----------------------------

SUY64392 ----------------------------MSKIANIKLELSD----W---------LYNAGVVGICKI------------------------------------------------------FDQNDDYYEKHQNYIVFDESL-----LE---NFEKKYFNYFINK-------------YEKFTSWYKIVSIEDKLINFSKDTVDEKYIEKFNKQ-------------IEYVKTKLKSA---------------------------SYKSGYEVIENKELDLLKEEQNLKKIKKTKKQSI--------
EDVIPMIEEQNELLQKIIEYLKKEEV-------------KRIIVAKNII-------YDVIAK-FWSDVSFLH--------SS--NN-------KK-SMYK----------EYKNYF-LDDCINYI-------------------------------------------------------------------------HEDKEKFKH-----------TCFSCNKIAKL-------------------SKPASYDLTWVN--KMGVDMS--R--KSSHF---------------W------N--FY-GD--TFICPICNLIYSCIPAGFT-----------I------LKDK----------------------
----GIF-----INENS------NIDTLIK----VNNT------AID---------------NNTSFQELEEKSYFTIA------ENMKQSEFE---------RSNKEID-NIQIVK-LDSN----------NERRP-YSFNVISKNKLRIIYNNRKNLRLLISK-----------------------NA--KIGDGS-----YINL-------------------------------------------YS-EVIKRLYNN------ENLFDLISEL-FYLKL-----------------------------------------------------------------------------------------------------------------------
------------------------------GDKFKGLSY--KNIIKINN-------------------------------------------------------------------------SFLGGIMGKKVYYKK---------------------------------------------------------------------

EDU38639 ----------------------------MKKIANIKLELSD----W---------LYNAGVVGICKI------------------------------------------------------FETNDDYYEKHSNYIVFDEKL-----LE---DFEEKYFNYFINK-------------YEKFTSWYKIVSIEDKLINFSEDTVDEKYIEKFNKQIEYVKTKLKSASYKSGYEVIE------------------------------------NKEIDLLKEEQKLKKIKKTKKQSI--------
QDVIPMIEEQNKFLQEIIEFLKKEEV--------------KRIIAKNVI-------YDVIAK-FWSDVSFLY-------KKSNKK-------DMYN----------EYKKYF-IDDCIDYI--------------------------------------------------------------------------HQNKEKFKY-------------TCFSCNNKISKL---------------------SKPASYDLAWIN--KIGVDMSPTR--KMSHF------------------W------N--FN-GD--TFICPICNIIYSCIPAGFT------------I-----------LKDK-----------------------
---GIF----INENS-------DIDTLVK---VNNT-----AID----------------NNTSFQELEEKSYFTIG-------DNMKQSEFE-----------RTNKEID-NIQIVK-LDSN----------NERRP-YSFNIISKNKLRIIYNNRGNLKLLISK------------------NA--KIGDGS------YINL----------------------------------------YS-EVIKRLYNN------KNLFDLISQL-FYLKL-------------------------------------GDKFKGLSY--------LKNIIKINN------SFLG-----------GIM-------
GKKIYYK----------------------KIDEHLKGLFCLRKEYEIKKA-----------------------DNKINGITYRLL-NALKTKNSNKFMD-TLLNAYMY----IR-----------KPIPT-EF-IEGL-----------KDEA-------KLQTIGYAFLLGLQGDTSKLESNEEGA-----------VNNE------------------------

STA92272__3_ ------------------------------MKKIAKIKLEKSD----W---------LYNAGIVGICNI------------------------------------------------------LKYNECDYDDSNINYVEFESEC-------LG---NFEEKYFNYFIGT-------------YEKFTSWYRLISFEDYMDNLDLNIISSKDLKYINKYIENLKNKLKSNSYKSAYLVIN---------------------------------------------------DKEIDLLVKKEKEIKKIKITKKQNI---------
NDVMVDIKKQIEIIKEIIEYLKRDEI-------------KRYILAKNII-------YDVIQK-FWSDVSFLH--------QKSNNK-----------DMYE----------EFKRYF-LDSTKTYL-----------------------------------------------------------------------------NSDKEKAKY-------------YCFTCGRKVS-----------------------SLKETFGLAWIN--KIGVDMS--R--KSSHF---------------W------N---MN-GD--TFICPICNLVYSCVPAGFT------------T----------LNGR-----------------
----GIF-----INENS------SIKNLIR----IKH-------SLDTTTSID------------------ELEEESYFNIA----DSI--GQVS-----------IDKSERETE-NIQIIK-LNSE-------NERRP-YTFNIISKEKLNIICKHRNMLKSMIKI-----------------------HA--KVGTNE-----YINI-----------------------------------------YR-EVLDRIYKG------KNLFDLISKL-SYLKL-----------------------------------------DGKFNGLQF-----------IEMMLKINN----------NLLE----------RGE-
EIGMVSGSINDK---------------------AIYSFKCRGAELRKGYENRKA-------------------LNKLGGVSYRLL-NSLKTKNVSRFMD-TIINSYMY----LN----------KEIPV-ESFINVVL-----------NDK-----N-KFQTIGYAFVLGLQGEEKFKKSGEEV-------------NNDK-----------------------------

KGM94848 ------------------------------MSKIQIELSD---W---------LYNAGVVGLINI--------------------------------LTHSEKKVKIVNKQ--CIEIDSEQ-------LV---DFQNDYFKYFIK-------------YVKFTSWYKVTSFQGYIESLDTENITKEDIKKINDY--------------IEYTKNKLKSN--------------------------------SYKSAYLLANDNSLDILKEEKNLKKINITKKQTL------
QDKEVKENINSTINRINKVIEYLCKETV-----------KKYVLTQNII-------YDVIAK-FWSDVSFLH----------KS-SN---------GC-NMYS-----------EYKKYF-IDCVLNYL------------------------------------------------------------------------DLDKEKDKY-------------NCFTCNNTLSKL------------------SKPIAYELAWIN--KTGVDMS--R--KSSHF---------------W------N--MK-GD--SYICPICNLVYSCIPAGFT------------V-----------FKGK------
----------GLF----INKNI-----GVKTLTK----INNQ-----SLDNVTSIE-------------------ELEQYSYFNIA---DNME-QGVI---------SQSSKEID-NIQIVK-LDSQ-------NESRP-YTFNIISKKKLEIIDKNKERLKSMIKI------------------HI--KISKDE----YLNL--------------------------------------------YN-EVLNRIYKG-----HNLFDLMYKL-VYLNL-----------------------------------NESFKRIGY---------LDMILKINN---------DLIG-----------GGK-----
MSTNKIDDK----------------------LINNCRSYGYYLRKAYVEKGG---------------------KNKLGGISYRLL-NALKTKNVSRFMD-TLFNAYMY-----------LKDKEIPS-TF-IEVL--------KDK-----E-NFQTIGYAFLLGLQGGDGKADKENK------EG-----VVNE-----------------------------

KEH93517 ----------------------------MSKIQIELSD---W---------LYNAGVVGLINI------------------------------------LTHSEKKVKIVNKQ--CIEIDSEQ-------LV---DFQNNYFKYFIDK-------------YVKFTSWYKVTSFEGYIESLDTENITKEDIKKVNDY--------------IEYTKNKLKSN--------------------------------SYKSAYLLANDNSLDILNEEKSFKKINITKKQTL------
QDEEVKENIKNTINTINKVIEYLCKETV-----------KKYVLTNQII-------YDVIAN-FWGGVSFLH----------TS--KN-------NS-DMYK----------EYKDYF-VDNTLNYV-----------------------------------------------------------------------ELDKSKDKY-------------NCFTCNNTLSKL------------------SKPIAYELAWIN--KTGVDMS--R--KSSHF---------------W------N--MK-GD--SYICPICNLIYSCIPAGFT------------V------------FKGK------
-------------GLF-----INKNI------GVKTLTK----INNQ----SLDNVTSIE----------------ELEQYSYFNIA---DNMEQGAIS---------QSNKEID-NIQIVK-LDSQ---------NESRP-YTFNIISKKKLEIINKNKERLKSMIKI--------------------HI--KISKDE-----YLNL-------------------------------------------YN-EVLNRIYKG------HNLFDLMYKL-VYLNL-----------------------------------------NESFKRIGY----------VDMILKINN-----------DLIG----------GGK------
MSTNKIDDK-----------------------LINNCRSYGYYLRKSYVEKGG---------------------KNKLGGISYRLL-NALKTKNVSRFMD-TLFNAYMY-----LS--------------NPIPS-TF-IEAL-----------KDK-----E-NFQTIGYAFLLGLQGG--DGKDGKADKENKED------VVNE--------------------------

EDS77895 ------------------------------MSKIQIELSD---W---------LYNAGVVGLINI------------------------------------LKHSEKKVKILNKQ--CIEIDSEQ-------LV---DFQNNYFKYFIDK-------------YVKFTSWYKITSFQGYIESLDTENITKEDIKKVNDY--------------IEYTKNKLKSN--------------------------------SYKSAYLLANDNSLDILNEEKSLKKINITKKQTL------
QDEEVKENIKNTINTINKVIEYLCKETV-----------KKYVLTNQII-------YDVIAN-FWGGVSFLH----------TSKNNS---------DMYK----------EYKDYF-VDNTLNYV-----------------------------------------------------------------------ELDKSKDKY-------------NCFTCNNTLSKL------------------SKPIAYELAWIN--KTGVDMS--R--KSSHF---------------W------N--MK-GD--SYICPICNLIYSCIPAGFT------------V-----------FKGK------
-------------GLF-----INKNI------GVKTLTK----INNQ----SLDNVTSIE-----------------ELEQYSYFNIA---DNMEQGSIS----------QSNKEID-NIQIVK-LDSQ---------NESRP-YTFNIISKKKLEIINKNKERLKSMIKI--------------------HI--KISKDE-----YLNL--------------------------------------------YN-EVLNRIYKG------HNLFDLMYKL-IYLNL---------------------------------------NDSFKRIGY-----------VDMILKINN-----------DLIG----------GGK------
MSTTKIDDK----------------------LINNCRSYGYYLRKAYIEKGG--------------------KNKLGGISYRLL-NALKTKNVSRFMD-TVFNAYMY------------LKDKEIPS-TF-IEALK--------DKE-----N-FQTIGYAFLLGLQGGDGKADKENK------ED------VVNE--------------------------

KOC30898 ------------------------------MSKIQIELND---W---------MYNAGVVGLINI--------------------------------LEYNEKKVKIVNNQ--CIEIDSEQ-------LQ---DFEKKYFDYFIDK-------------YLKFTSWYKITSFEDYLEKIDVEKITKEEIEKLNDY--------------IKYIKKLKSN--------------------------------SYKSGYLLANDNSLDILKEEKSLKEINITKKQTL------
QDENVKIKKNNIDGINKINKVIEYLCRENV------------KKYVLTQNII-------YDLICK-FWGGVSFLN----------SSNIKN---------KC-NMYV----------QYRKDF-IESVVSYI------------------------------------------------------------------------NLDKSKDKY-------------NCFTCNNTLSKL------------------SKPIAYELAWIN--KTGVDMS--R--KSSHF---------------W------N--MK-GD--AYICPICNLIYSCIPAGFT-------------V-----------LKGR------
-------------GLF-----INKNT------GINTLIK----INNQ----SLDNVSSIE-------------------ELEENSYFNIV----DNMEQGSIS----------QVSKEID-NIQIVK-LNSQ---------NERRP-YTFNIISKKDLKIINKYKEQLKSMIKI---------------------HI--KISKDD-----YLNL----------------------------------------------YN-EVLNRLYNG-----HNLFDLIYKL-IYLNL---------------------------------------NGNFKNTRY----------AEKILEINN-----------DLIG-----------GGK-----
MSTNKIDDK---------------------VISNCRSYGYYLREEYVQKGA------------------KNKLGGISYRLL-NALKTKNVARFMD-TLFNSYMY-----LN-----------KQIPA-RF-AEVL---------NDK-----V-KFQTIGYAFLLGLQGSGKNNNENEKK----ED-----ISNE-----------------------------

KGN04942__2_ ------------------------------MSKIQIELND---W---------LYNAGVVGLLNI-------------------------------LKHNEEKIKIVSKQ--CVEIDTDQ-------LK---DFEKKYFDYFIDK-------------YLKFTSWYKITSFQGYLDSLDIENMTKEDIKKINDY--------------IEYTKNKLKSN--------------------------------SYKSAYLLANDNSLDILKEEKNLKKINITKKQTL------
QDEEVKENINNTINGINKVIEYLYRKEV------------KKYVLTKNII-------YDIIAN-FWGGVSFLN----------TS--KN-------NS-EMYS----------EYKEYF-IEGVVSYI------------------------------------------------------------------------DLDKSKDKY-------------NCFTCNNTLSKL------------------SKPIAYELAWIN--KTGVDMS--R--KSSHF---------------W------N--MK-GD--AYICPICNLVYSCIPAGFT-------------V-----------LKGK------
-------------GLF-----INKNT------GISSLTK----INNQ----SLDNVASIE-----------------ELEENSYFNIV----DNMEQGSIS----------QASKEID-NIQIVK-LNSK---------NERRP-YTFNIISKKDLKIINKHKEQLKNMIKI--------------------HI--KISKDD-----YLNL-----------------------------------------------YN-EVLNRLYNG-----HNLFDLIYKL-FYLNL------------------------------------NGNFKNTGY----------IDMILKINN-----------DLIG-----------GGK-----
MSTNKIDDK---------------------VISNCRSYGYYLREEYVQKGA---------------------KNKLGGISYRLL-NALKTKNVARFMD-TLFNSYMY-----LN-----------KQIPS-RF-TEVL---------NDK-----V-KFQTIGYAFLLGLQGSGKNNNENEKK----ED-----ISNE-----------------------------

KOC53218__4_ ------------------------------MSKIQIELND---W---------LYNAGVVGLINI-------------------------------LRHNEEKVKILNKQ--CIEIDSDQ-------LQ---DFEKKYFDYFIDK-------------YLKFTSWYKITSFEDYLEKIDVEKITKDDIDKLNKY--------------IEYLKPKVKLN--------------------------------SFKSAYLLANDASVDILSEEKHLKKISIGKKQTL------
QDEKVKENIINNLNSIHKIDNLCKENL-------------KRYVLSKNII-------YDVIAN-FWGGVSFLH----------TS--KN-------NS-DMYS----------EYKEYF-VDGVVSYT------------------------------------------------------------------------DLDKSKDKY-------------NCFTCNNTLSKL------------------SKPIAYELAWIN--KTGVDMS--R--KSSHF---------------W------D--MK-GD--AYICPICNLVYSCIPAGFT-------------V-----------LKGK------
-------------GLF-----INKNT------GINTLTK----INNQ----SLDNVSSIE-----------------ELEENSYFNIV----DNMEQGSIS----------QVSKEID-NIQIVK-LNSQ---------NERRP-YTFNIISKKDLKIINKHKEQLKSMIKI--------------------HI--KISKDD-----YLNL-----------------------------------------------YN-EVLNRLYNG-----HNLFDLIYKL-IYLNL------------------------------------NGNFKNTVY----------IDMILKINN-----------DLIG-----------GGK-----
MSTNKIDDK---------------------VISNCRSYGYYLREEYVQKGA---------------------KNKLGGISYRLL-NALKTKNVARFMD-TLFNSYMY-----LN-----------KQIPA-RF-AEVL---------NDK-----V-KFQTIGYAFLLGLQGSGKNNNENEKK----ED-----ISNE-----------------------------

KGM99198 ------------------------------MSKIQIELND---W---------LYNAGVVGLINI-------------------------------LRHNEEKVKILNKQ--CIEIDSDQ-------LQ---DFEKKYFDYFIDK-------------YLKFTSWYKITSFEDYLEKIDVEKITKDDIDKLNKY--------------IEYLKPKVKLN--------------------------------SFKSAYLLANDASVDILSEEKHLKKISIGKKQTL------
QDEKVKENIINNLNSIHKIDNLCKENL-------------KRYVLSKNII-------YDVIAN-FWGGVSFLH----------TSKNNS---------DMYS----------EYKEYF-VDGVVSYT------------------------------------------------------------------------DLDKSKDKY-------------NCFTCNNTLSKL------------------SKPIAYELAWIN--KTGVDMS--R--KSSHF---------------W------D--MK-GD--AYICPICNLVYSCIPAGFT-------------V-----------LKGK------
-------------GLF-----INKNT------GINTLTK----INNQ----SLDNVSSIE-----------------ELEENSYFNIV----DNMEQGSIS----------QVSKEID-NIQIVK-LNSQ---------NERRP-YTFNIISKKDLKIINKHKEQLKSMIKI--------------------HI--KISKDD-----YLNL-----------------------------------------------YN-EVLNRLYNG-----HNLFDLIYKL-IYLNL------------------------------------NGNFKNTVY----------IDMILKINN-----------DLIG-----------GGK-----
MSTNKIDDK---------------------VISNCRSYGYYLREEYVQKGA---------------------KNKLGGISYRLL-NALKTKNVARFMD-TLFNSYMY-----LN-----------KQIPA-RF-AEVL---------NDK-----V-KFQTIGYAFLLGLQGSGKNNNNEN------EK-----KEDI-----------------------------

WP_026737448 ----------------------------MSEKIEIRMSN---W---------LFNAGLLGFYNV-------LGGKVAEEK----------------------GEIE---------IS---EDGQ--SLIFDSGM-----LE---NFEYKYFDFFLKR-------------YNKILPYGKILDFE--DYIDDFLSGKENRSLKEVD------------YVNSQIELVKK-----------SLKSNSYQSTYDFV-------
ENNGKEKMLNLEKEIKKVKKPKDNVT---------SETVEEIKEVFGQLKEVM--NFFKKT------------VKNEDGYEKKYLAAKNIVYLINN-SWNGVSFLN-----------RA--NA---------AK-DVYE----------EYKSYF-VIPAIEYI-------------------------------------------------------------------------NKDKSKFK--Y--------KCSISNMSM-------------------------ENYKNTLGFLN--QTGFDIA--R--KPSHV---------------W------N--FT-ND--
IAITPLIILILSCIPAGFI-------------Y-----------EINK---------------------------GMF-----VNANF------NFDQLER---INNG-----IAN-SIF--------------VQEQE-EKINLYRNLL-----KEFETNTND------------TKYEIS-DIQVVR-LENE--T------------YRFSLLSKNILYLLHENKDKLDRLLGK--------------------------YY--KDYDKY-----FNL-----------------------------------------------YD-ETIRELLNN-----QNLFSLINKL-CYYKI-----------------------
SKNKSAYY----NGLNMSDLVKINM---------NYIR-----------RLR------------KM-------ENNEDKRT--ELSEK---DIEMIRTNAYYFRKDY-IGK-SGN------------------------DKKIGGLLYRLQ-NALRINDVDMFMD-ALISAHAY-----AG-------------KGVHK-LFNRALI---------DDE------DFQTLGHAFLIGLLEDGKKDDENKD------NK-----KEGN-E-----------------------------

ALA96009__2_ --------------------------------MDEKIELKLKD---W---------LFNAGLIGLMNI-------LGEE-AKDN----------------------GKLE---------ID----DKNR--LIRFSPKV-----LE---NFEYKYFDFFIKR-------------YKGTLTYGKILEFE--KYIDEFEEGKEIRSINELK------------MINDKITFFKKA-------------KIKSESYKKAYDFI---------EKNGTDKILGLEKELKKIEKPKENVD-----
--EISMENVKNNFKIMKEII--DFFKKK------------IMDEEGNIKNYLAAKNIAYVIINN-AWSSVSFLN---------KA--NA--------VK-DIYE----------EYKSYF-VEPALEYI--------------------------------------------------------------------------NANKSKFK--Y--------KCTISNMPM---------------------------KDYKNTLGFLN--DTGFDVN--R--KPSHV---------------W------N--FV-ND--IAVTPLVTLIYSCVPAGFI----------------------Y---------GADK------
-------------GIF-----VNANH---------NIDQLYK-----IAY---NIL---------------EDESE-EKNINLYKNLL-------KEIKKEKDN-----------TKYELS-DIQIVK-FEEG--H--------YKFTLLSRNILLLLSENKEKLDDLLDK----------------------WY--LIDRRY-----FYL----------------------------------------------YD-TAITELLNN-----QNLFSLINKL-CYYKI----------------------------------SKTK-LYC-----KLKNIEDLLKINL-------DYIR-----------RLK---------
--KMDKQEIEVEKEESKKISEELTEK----DIFYIRRDAMIFREEY-IRK-SGN--------------------DKKIGSLLYRLQ-NALRINNVDMFMD-ALISAHAY-----AG-----------KNISS-LFAKALL-----------NDE------NFQTLGHGFLLGLLGEDKSKENKT-------DK-----KEGN-E-----------------------------

ACV38707__2_ --------------------------------MSDEKIELKLKD---W---------LFNAGLLGFINI-------LGEE-ARNN----------------------GELE---------ID----DKNR--LIRFSPKV-----LE---NFEYKYFDFFIKR-------------YGKTLTYGKILEFE--KYIDEFEEGKEIRSINELK------------MINDKITFFKKA-------------KIKSASYKKAYDFI---------EKDGTNKILGLEKELKKIEKPKENID-----
-EISKDDVKNNIKMKEII--DFFKKK------------ITDKNGNVKNYLAAKNIAYVIINN-AWSSVSFLN---------RA--NA--------AK-DIYE----------EKHSYF-VEPALEYI--------------------------------------------------------------------------NADKSKFK--Y--------KCVLSNMLM---------------------------KDYKNTLGFLN--DTGFDVN--R--KPSHV---------------W------N--FV-ND--IAVTPLVTLIYSCVPAGFI----------------------Y---------GADK------
-------------GIF-----VNANH---------NIDQLCK-----IAY---NIL---------------EDESE-EKNINLYKNLL-------KEIKKEKDN-----------TKYELS-DIQIVK-FEEG--H--------YKFTLLSRNILKLLSENKEKLDALLDK----------------------WY--LIDRRY-----FYL----------------------------------------------YD-TTITELLNN-----QNLFSLINKL-CYYKI----------------------------------SKTK-LSC-----KLKNIEDLLKINL-------DYIR-----------RLK---------
----KMDKQEIIEKKENKKTSEELTEK----DIFYIRRDAMIFREEY-IRK-SGN--------------------DKKIGSLLYRLQ-NALRINNVDMFMD-ALISAHAY-----AG-----------KNISS-LFAKALL-----------NDE------NFQTLGHGFLLGLLGEDKSKENKT-------DK-----KEGN-E-----------------------------

WP_026749139 --------------------------------MSNEKIELKLKD---W---------LFNAGLLGFINI-------LGEE-AKSS----------------------GELE---------ID----NKNR--LIRFSPKV-----LE---NFEYKYFDFFIKR-------------YGKTLTYGKILEFE--KYIDEFEEGKEIKSINELK------------MINDKITFFKKA-------------KIKSESYKKAYDFI---------EKNGTNKILGLEKELKKIEKPKENIA-----
EISNDDIKNNLKIMKEII--DFFKKK------------ITDKEGNVKNYLAAKNIAYVIINN-AWSSVSFLN---------KA--NA--------AK-DVYE----------EYKSYF-VEPALEYV--------------------------------------------------------------------------NADKSKFK--Y--------KCVISNMQM---------------------------KNYKNTLGFLN--DTGFDVS--R--KPSHV---------------W------N--FV-ND--IAVTPLVTLIYSCVPAGFI----------------------Y---------GADK------
-------------GIF-----VNANH---------NIDQLCN-----IAY---NIL---------------EDESE-EKNINLYKNLL-------KEIKKEKDN-----------TKYELS-DIQIVK-FEEG--H--------YKFTLLSRNILKLLSENKEKLDDLLDK----------------------WY--LIDKRY-----FYL----------------------------------------------YD-TTITELLNN-----QNLFSLINKL-CYYKI----------------------------------SKAK-LSC-----KLKNIEDLLKINL-------DYIR-----------RLK---------
KMDKQEIEKKENKTSEELTEK----DVFYIRRDAMIFREEY-IRK-SGN--------------------DKKIGSLLYRLQ-NALRINNVDMFMD-ALISAHAY-----AG-----------KNISS-LFAKALL-----------NDE------NFQTLGHGFLLGLGEDKSKENKT-------DK-----KEGN-E-----------------------------

AEV69187 ---------------------------------MSFIKIQLSD---W---------LYNAGVIGLVNI-------LK--------------------------HANDQ---------VD----INGQ--EVTIKTNV-----LA---NFEAKYKKYFVDT-------------YGKTMTFTKLINRIDKLLNIELDESKEKEIGDDIDF-------------------------------------------IKSKVSNNSAYKGIKDKIDLKKLEKPSKK------------
VTSINKEILEEL-KYLLLEN-------KKEILKSECIG-------------YYDQKSKTS-K------AP--NA--------II-DKYI-------NTNMLDVGLGVNEVKEYL-----------------------------------------------------------------------SNE--EKLNF-----------------SCFQCGRGI-----------------------QKIDKGLSFLN--RMFFDTA--R--KTSHV-------------W------N--FV-SD--IDICPICKIVYYCVPAGFT-----T----V---------FG-N-------------------GIF-----
VNYNQ-----SVERAEK---INLN-----IRH--EIL--------------HSDES-KRIYNAFGTMI-----RSLH-EKIN------DS-KKYDLA-DIQIVR-CEDD--K--------------YKFNLLSKNVLYALYDS----------RDE-------------LN--SLLNTG----F--QEVNTYFRI----------------------------------------------------YD-EVMKNLLNN-----VNLFILIHKV-LVIKI-----SSPKD------------------AKYN-ESH----------IRNLIKINF--------KYMK----------GVG--------
YMESTEKD---------------------IVKISSSAGYYLREAY-KSK-NS-----------------ENKLNGICYRLL-NALKTSNKSMFMD-TILNCYLY-------------TNKPVPT-FI-TECL-------------NDE------TKFQTIGYAFVTGLIDG--NKNE-----------NT-----DGGD-N-----------------------------

WP_073073948 ---------------------------MAKTITLKLSN---W---------LYNAGLVGIINV---------------------------------------LGRDN---------VE---LNDE--ELIIDLDK-----LE---NYDELYFNYFINT-------------YKNLFFNFRLIEKKKIIEKKDAAEISASEINEFIRLIKDKNS-----------------------------------------------------------------
GLLRSSYRKLLPLIEEKTGKKFFIEEHLDKLDLKVSEENLSILKFIFEFLDNDI------------VKKYLPVADVS---------YNVINK-FWSNVSFLN----------KN---NK-------EL-DFYK----------AYKEYF-IEPLFEYI-------------------------------------------------------------------------KSDKSGYK--H--------NCFVCNNK-------------------------NKQFSNSISFIN--NIGFDIT--R--KTSNV---------------W------N--FN-ND--
IFLCPVCTFVYSNIPAGFI-----------Y-----------TLGK-----------------------------GIF-----VNINT-----SIEDMYT-----TNLR-----IKM--------------EVFSEYR-ERFEASYLALL------KNIY-QNLN--------KT-ARLEYS-EIQIVK-LDNL-------------A-YKFNFISKETLKILN--------------EKR---------------LN--ALLNNY----YVISNSDNKKITNSI----------------------------------------YE-EALYKIINK------SPLEPLIYKL-LYLRI--------------------------
NSENEGFNYN----TINNLLNIMYYIT----------SEVR-----------KLN-----------GKEE--------------LVRKANYAKGYWLKEY-KKR-DS--------------------------VNKIPGIAYRLL-NSLKVEKFDQFLD-ILLNCYLY------VS-----------KIPK-LI-IDTL------TE---D-K-LKFIVAYSFIAGMINE--------------------------------------------KV------EKEE-IV---------------------------

EUJ57885 -------------------------------------MLEFEISD---W---------GYNASLAGFINI-----------------------------------IGEENV---------TY-----LDHQ--CVQIEEEM-------LK---DFVTKYFNYFIDT-------------YEKVLPWYRIVSYKDSMEATLESKFDNFTEKDLKEM-------------NYYVKNVLKYY----------FTSNSYKAAYPLI----------DADFNPEVEVKKLKTINLKKKDDF---
NEIKAELISQVKENFEIIAKLI-DYAKSE---------KGRKYLAGKNVV------YNLIKN-SWEGVSFLN-----------RN--TK-----------EK-DFYI------------DFEKHF-IEPANKYI-------------------------------------------------------------------------EADKTKYK--Y---------KCTNCERPI---------------------------KDLELDYSFLT--NTGFDVS--R--KKSHV---------------W------N--NDNDLAMCPICRLVYACMPAGFT------------------------------------------------IQRE------
-----GIF----INSNA-----NIQEMVR---INRM------MKN------------------DLSSKLD-DQNKSVYRTLV-------RMA-QESD------------VEKHRYELA-DIQVVR-HENE--------------S-YRFNLLAKPVLTLMNRF-------EKELKSL---------------IG-ANFKEGNERQYI------------------------------------------------------FQ-EVVKNILDN------QNLFLLAHKL-IYYKV----------------------------------ANYEQTYY----------------------------
----------------------------------YSYHIM-NVLKINSYFTGGT-----------------GEMTQ-TITLDEL----------------------E-KVRANAYYFKEELKNRKT-------------------------------------------------------------------------------------------

WP_088816243 ------------------------MSRVYLGSKKWRWGRMLEFETSD---W---------AFNSALSGFINI-----------------------------------VGVESV-----------AY-----TDSN--RVSLNEDA-------LK---DFTGKYFDYLIDT-------------YEKVLPWYRIVSYKRKIVEFEENQFDNFSEKDLENL-----------NLYMRNVVKHY----------FSSASYKAVFSLI----------
DADFDPEVELKALKPINLKKEDIF---SEVKADILPQVRENFVILRKLIDYGASEKG------------RKYLAGKNVV-------YNLIKN-SWEGVSFLN-----------RN--TK----------EK-DFYI------------DFENHF-VEPAKAYL-----------------------------------------------------------------------EADKTKYK--Y---------HCAICNKI--------------------------KDLKLDYSFLT--NTGFDVS--R--KKSHV---------------W------N--EN-ND--
LAMCPVCRLVYTCMPAGFT-----------F---------IQRE--------------------------GIF-----INIGK-----DIKEMVR---TNKV--------MKN------------------DLNMLE-ARNKSVYRALV-------SLA-QEKK-----------YEKHRYELV-NIQVVV-YEDG-------------V-YRFNLLSKRALELMNRFEKELKSL-IGM-------------------------------------------HFKEGNVRVYL------------------------------------------------------FK-EVVKNLQDN------QNLFLLIHKL-VYYKV----------------------------
------ANYGQLHY-----YPQHIMNMLKINS---------YFTE------------GVG-------------KMSQKIMPE-------------ELEKVRAGAYYFKEEL-KSR-KR-----------------------EQKLPGMAHQLL-NAVKTVDRGLFMD-IMLRGYSE------LD---------------KQIPK-IF-LRIF---------ESD-----E-NFKIIGYAYLIGLVGD--NKENN----------KEE----EENE-N-----------------------------

WP_088825424 ---------------------------------------MLEFETSD---W---------AFNSALSGFINI-----------------------------------VGEENV---------TY----IDNQ--RVSVNREL------LE----DFSKKYFDYFIDT-------------YEKVLPWHRIVSYKEKITEFEENQFDNFSEKDLENL-----------NLYIKNVVKYY----------FSSASYKAAFSLI----------DTDFDPEIEIKTLKTINLKKKDVF---
SEVKADILPQVKENFEVMTKLI-DYASSE--------KGRKYLAGKNVV-------YTLIKN-NWEGVSFLN----------RN--TK---------EK-DFYI----------DFENHF-VEPAKAYL-----------------------------------------------------------------------EADKTKYK--Y--------HCTNCDAKI----------------------------KDLKLDYSFLT--NTGFDVA--R--KASHI--------------W------N--QD-ND--LAMCPVCRLVYACMPAGFT----------F-------
IQRE--------------GIF----INSNA-----NIKEMVH---VNQA------MKN------------------DLNTLLN-DRNKSVYRALV------SLA-QKKS-----------AEKHRYELA-DIQVVR-YENE--------------N-YRFNLLSKPVLELMNRF--EDELKSL---------------IG-MNFKDGNDRLYI----------------------------------------------------------FQ-EVVKNLLDN------QNLFLLIHKL-VYYKI-------------------------------------ANNGQLFY--------YSHHIMIVLKMNS----------
YFTG----------GAG-----NMSQIITLE---------------------DLEKVRANAYYFKEDL-KSR-KR-------------------------DQKIQGIAHQML-NAVKTVNRGLFMD-ILLKNYSA-------VD---------------KQVPK-IF-LRIF---------DND-----D-NFRTIGYAYLTGLIGE----------------------TN------VNNK-EEAK--------------------------------

WP_088838977 ---------------------------------------MLEFETSD---W---------AFNSALSGFINI-----------------------------------VGEGNV---------TY----IDNQ--RVSVNREL------LE----DFSKKYFDYLIDT-------------YEKVLPWHRIVSYKEKITEFEENQFDNFSEKDLENL-----------NLYIKNVVKYY----------FSSASYKAAFSLI----------DTDFDPEIEIKTLKPINLKKKDVF---
SEVKADILPQVKENFEVLTKLI-DYASSE--------KGRKYLAGKNVV-------YTLIKN-NWEGVSFLN----------RN--TK---------EK-DFYI----------DFENHF-VEPAKAYL-----------------------------------------------------------------------EADKTKYK--Y--------HCANCDAKI----------------------------KDLKLDYSFLT--NTGFDVA--R--KASHI--------------W------N--QD-ND--LAMCPVCRLVYACMPAGFT----------F-------
IQRE--------------GIF----INSNA-----NIKEMVH---VNQA------MKN------------------DLNTLLN-DRNKSVYRALV------SLA-QKKS-----------AEKHRYELA-DIQVVR-YENE--------------N-YRFNLLSKPVLELMNRF--EDELKSL---------------IG-MNFKDGNDRLYI----------------------------------------------------------FQ-EVIKNLLDN------QNLFLLIHKL-VYYKI-------------------------------------ANNGQLFY--------YSHHIMIVLKMNS----------
YFTG----------GAG-----NMSQITLE---------------------DLEKVRANAYYFKEDL-KSR-KR-------------------------DQKIQGIAHQML-NAVKTVNRGLFMD-ILLKNYSA-------VD---------------KQVPK-IF-LRIF---------DND-----D-NFRTIGYAYLTGLIGE----------------------TN------VNNK-EEAK--------------------------------

WP_088840559 ---------------------------------------MLEFETSD---W---------AFNSALSGFINI-----------------------------------VGEENV---------TY----IDNQ--RVSVNREL------LE----DFSKKYFDYLIDT-------------YEKVLPWHRIVSYKEKITEFEENQFDNFSEKDLENL-----------NLYIKNVVKYY----------FSSASYKAAFSLI----------DTDFDPEIEIKTLKPINLKKKDVF---
SEVKADILPQVKENFEVLTKLI-DYASSE--------KGRKYLAGKNVV-------YTLIKN-NWEGVSFLN----------RN--TK---------EK-DFYI----------DFENHF-VEPAKAYL-----------------------------------------------------------------------EADKTKYK--Y--------HCANCDAKI----------------------------KDLKLDYSFLT--NTGFDVA--R--KASHI--------------W------N--QD-ND--LAMCPVCRLVYACMPAGFT----------F-------
IQRE--------------GIF----INSNA-----NIKEMVH---VNQA------MKN------------------DLSTLD-DRNKSVYRVLV------SLV-QEKQ-----------VEKHRYELA-DIQVVR-YENE--------------N-YRFNLLSKPVLELMNRF--EDELKSL---------------IG-MNFKDGNDRLYI----------------------------------------------------------FQ-EVIKNILDN------QNLFLLIHKL-VYYKI-------------------------------------ANYGQLFY--------YSHHIMIVLKMNS----------
YFTG----------GAG-----NMSQTITLE---------------------DLEKVRANAYYFKEDL-KSR-KR-------------------------DQKIQGIAHQML-NAVKTVNRGLFMD-ILLKNYSA-------VD---------------KQVPK-IF-LRIF---------DND-----D-NFRTIGYAYLTGLIGE----------------------TN------VNNK-EEAK--------------------------------

WP_088809478 ---------------------------------------MLEFETSD---W---------AFNSALSGFINI-----------------------------------VGEGNV---------TY----IDNQ--RVSVNREL------LE----DFSKKYFDYLIDT-------------YEKVLPWHRIVSYKEKITEFEENQFDNFSEKDLENL-----------NLYIKNVVKYY----------FSSASYKAAFSLI----------DTDFDPEIEIKTLKPINLKKKDVF---
SEVKADILPQVKENFEVLTKLI-DYASSE--------KGRKYLAGKNVV-------YTLIKN-NWEGVSFLN----------RN--TK---------EK-DFYI----------DFENHF-VEPAKAYL-----------------------------------------------------------------------EADKTKYK--Y--------HCANCDAKI----------------------------KDLKLDYSFLT--NTGFDVA--R--KASHI--------------W------N--QD-ND--LAMCPVCRLVYACMPAGFT----------F-------
IQRE--------------GIF----INSNA-----NIKEMVH---VNQA------MKN------------------DLSTLD-DRNKSVYRVLV------SLV-QEKQ-----------VEKHRYELA-DIQVVR-YENE--------------N-YRFNLLSKPVLELMNRF--EDELKSL---------------IG-MNFKDGNDRLYI----------------------------------------------------------FQ-EVIKNILDN------QNLFLLIHKL-VYYKI-------------------------------------ANYGQLFY--------YSHHIMIVLKMNS----------
YFTG----------GAG-----NMSQTITLE---------------------DLEKVRANAYYFKEDL-KSR-KR-------------------------DQKIQGIAHQML-NAVKTVNRGLFMD-ILLKNYSA-------VD---------------KQVPK-IF-LRIF---------DND-----D-NFRTIGYAYLTGLIGE----------------------TN------VNNK-EEAK--------------------------------

SFE21040 -----------------------------MEDIIKITLGD---W---------QYNAGIVGMYNI-------------------------------LKHSGKDIVEFTSNYVKGENKENVNSNFEICFPIEY-----LE---NFANDYFDYLIDT-------------YINQISYGKIVGFKVDAEIISIEYEDFSDEKIVEA-------VEKINKYIKNCLHYYI---------------KSASYKAALEIPNSDVLVKEIIEIKSIKISKK----
---NKPSISEIKNICNKLIKVISSLEGPMY-----------RKYIGGKNVI-------YTLIKR-NWDKVSVLN----------AQ--VK---------EK-NILK----------EFDNYF-VFPAIEYN--------------------------------------------------------------------------KTDKVKYKY-------------NCFLCDSRI-------------------------KDMNNNLSFLT--NTGFDVA--R--KSSHI---------------W------N--FQ-ND--IAICDVCKLVYACMPAGFV-----------Y-----------GFDK-------
--------------GVF-----INSNL--------NIKALTH----VNSK------LKN---------------NVLEGFNNNKKSMTYKGVV-----EAII-YERN----------NQIIKYELS-DIQVVF-FEND---------------K-YVFSILSRNILNVIYESRTQLNYIKNG-----------------------YFVENKYYYYL----------------------------------------------YE-EVLKKLLNN-----QNLFSLINRL-MHYYI-----------------------------------SNKRSCFF---------SLTSLECMLRVNL-----------NYLK--------
GVG-----LMSELNNS----------------DISIRNGYWHVRMGYDSKGA-----------------------VKKIDGISYSML-NALKTSNVNAFMD-LLIRCYMY----ID----------KSIPK-KF-IECL--------STE-----E-EFKTIGYAFILGLFNGANFEK-----------NE------EKDNNKLSWEE-----------------------------

KXI12129 ----------------------------MEDKKLNFINFH------------------------------------------EIIFPSVY--------LE---NFTEKYYTYLIDT-------------YIEQISYGRIIRFKEYANRVLDMEDDEIDEKVFEII---------NDYIKNVLKYY----------------LKSASYKSAFDLIDGSANVVNKAMSIEQVKKTNKDID--------
LDVKNKIKDVCEKIIDIIDTLEENQY----------KKYLAGKNVI-------YTYIKR-NWNGVSILN----------AQ--VK----------EK-DIYR---------EFDNYF-VKPAILYL--------------------------------------------------------------------------NEEKSKFKY-------------TCYTCGSKI-----------------------KDLKTNLSFVN--GMGFDVA--R--KSSHV-----------------W------N--FQ-ND--IGICELCKLIYACMPAGFV-----------Y-----------GFDK-------
---GIF----INSNT-----SIYALIE---TNKK-----LTI------------NMQEEINKNEGSSTYKSIV-----EAITSRGIE--------GSKFELA-DIQVVF-YEKE------------------K-YVFNMLSRNILNIIYESRKKLNYIKNG------------------------CFKENKSYYYL----------------------------------------------YE-TTVRQILDN------KNMFLWIDKL-IRYSI---------------------------------TNKADCYY-----QMGHVLNILEINF--------EYLK--------------EVG-----
EMGNSKY----------------------LIYKMKDSGNDIRVAYYGKGV------------------------EHKLLGISHQIL-NALKTSNTSQFMD-VMIRMYMY-----VK-----------KEIPK-EV-INCL---------KTD-----E-EFKVYGYAFLLGLNGSLSEKQENL------ES-----KDEN-----------------------------

KXB69171 ------------------------------------MEDVIRISLGD---W---------QYNAGIVGMYNI-------------------------------LKHAGFDIIEYTDKYNNMEDKKLNR-----FINH---EIIFPSVY--------LE---NFTEKYYTYLIDT-------------YIEQISYGRIIRFKEYANRVLDMEDDEIDEKVFERI---------NDYIKNVLKYY-------------------------------
LKSASYKSAFDLIDGSANVVNKAMSIEQVKKTNKDID---------LDVKNKIKDVCEKIIDIIDTLEENQY----------KKYLAGKNVI-------YTYIKR-NWNGVSILN----------AQ--VK----------EK-DIYR---------EFDNYF-VKPAILYL-----------------------------------------------------------------NEEKSKFK--Y-------TCYTCGSKI--------------------KDLKTNLSFVN--GMGFDVA--R--KSSHV-----------------W------N--FQ-ND--
-IGICELCKLIYACMPAGFV----------Y--------GFDK--------------------------GIF-----INSNT-----SIYALIE---TNKK-----LTI-------------NMQEEINKNEGSSTYKSIV-----EAITSRGIE--------GSKFELA-DIQVVF-YEKE------------------K-YVFNMLSRNILNIIYESRKKLNYIKNG---------------------CFKENKSYYYL-----------------------------------------------------YE-TTVRQILDN------KNMFLWIDKL-IRYSI-----------------
TNKADCYY-----QMGHVLNILEINF--------EYLK--------------EVG-----EMGNSKY----------------------LIYKMKDSGNDIRVAYYGKGV------------------------EHKLLGISHQIL-NALKTSNTSQFMD-VMIRMYMY-----VK-----------KEIPK-EV-INCL---------KTD-----E-EFKVYGYAFLLGLNGSLSEKQENL------ES-----KDEN-----------------------------

WP_075780185 -------------------------------MEEIKNNFLIFRTSD---W---------LYNAGLVGLIEI-------------------------------VRYR----------TN------EENE--DLDFDEVKISINAFE---NFEEKFFNYFIET-------------YNLPWHQLVSYKEAVETLNEKNEYNKDLELLNK-----------------------MIDYFKNNLKKNSFKSVYPLI---------EEKENIKLDILNEIKKLKKIKKVEK---
--------IEDIKNNLLILENLINFLNLKSS---------KKYIVAQNIR--------YSNLIRN-AWDGVSFLN---------RQ--TK-------EK-DVYI----------DYKKHF-IDTFFNYI--------------------------------------------------------------------NDDEEFKKYKY---------KCFSCNRPI-----------------------KSLEYNMSFLK--NIGFDDA--R--KPSNV-------------------W------N--FV-ND--TYICPICNLVYSCMPAGFT-----Y-----V---------YD-K-------
-------GIF----INLNH-----SIEELVR---INKK------IKM--QVL-----------------NDPH-NNLNNSYSELL-----KSIY--ENK-------NKTASYEYS-DIQIVK-LKNE-------------K-YEFNILSKKILSVLKNSEEELDFLIAQ----------------------FY-FPSNNKN-----WNEKISI-----------------------------------------YE-EVIYRISNN------ISLYSLINTL-LRLKI----------------------SNVNNGYF---------NTFTIKQLINLNL--------NYLK----------EVK--------
-NMEFPEN---------LYGNANYQGYLLKKEY-NRK-NS-------------------------EIKLLGISHKLL-NALKVNNKGDFMD-VILNCYLY------VD-------------KPVPK-VI-IEAL----------ENDE------E-VFKTAGYSFVAGLIGE---------------------SN------TKGE-E-----------------------------

EFS20741__2_ ---------------------------------METIVIRSED---W---------LKNAGIIGLYRI----------------------------------LEEEESDEKSS----------IS---LEED--QIRFSAEL-----LQ---NFSEKYFRYFIKQ-------------YKNVLSLYRTLNFKENISQFKEKNEYNFGKEDLEKL-----------NEHVENVKKYL-------------KSNSYRAMYPLIRCPFDPLEKEKLKKVHLRKKESI-----
---EDGISDVIKKLITHLEEIYDFLQEDS---------QKYIGAKNVI-------YNIIKN-AWDGISILN-------------PQ--VK----------EQ-NMYF----------EFDKYF-VQTTQEYL-----------------------------------------------------------------------KQEKTKFK--Y---------RCFSCGESI--------------------------KDTNIDLSFMN--HIGFDVA--R--KTSHV---------------W------N--FN-NY--VHICPLCRLIYACVPAGFT------Y--------YD-K-------
----------GIF----VNANT--------HLKEMLR---INDL------IFK--NVLGEK---------------KSGKISYIGALV-----SGMS-KEMN-----------EHVEYDLS-DIQVVR-LEKK--------------R-YTFSILSRKFLSIIKKCRTDLEYIRKS-----------------------SFKEGNDIYYI---------------------------------------------------YE-ETIKRLMQG------ENLFNLIHKL-VRLKI------------------------------------SNGEDCYY-------KMGTVGTIIEINDL---------IFLK-----------EVG-----
YMKDEKKEYN------------------PLERARIIGHHLQEAY-GGF-EEGKY--------------------NKKLDGIAYRML-NALKTKNVNKFMD-SLINAHMY-----VQ-------------KPIPS-LF-SDYL---------HQE-----L-AFKELGYAFVTGMLGEEWKN-------------EG-----NTSK-N----------------------------

KDE67615 ---------------------------------MVEKIVIRSED---W---------LNKAGIVGLYRI--------------------------------LKARDERAD-----------IF------VEED--QISFSADL-----LQ---NFSEKYFHYFIKR-------------YKNVLSLYRILNFTANISQYEEKNYETFHEEDLEKL-----------NHDVEDVKKYL-------------KSNSYRAMYPLIRCPFDPLQKERELKKVNLKKTESL---
---KDRISDIQKLLADLKEIHDFLRQEDS---------QKYIGAKNAM-----YGIIQN-AWKGISILN-------------PQ--VK-----------EQ-NMYL----------EFDKYF-VQTAREYL-----------------------------------------------------------------------EQEKTKFK--Y---------RCFSCGEAI--------------------------KDTRIDLSFMN--HIGFDVA--R--KTSHV---------------W------D--FN-NY--VHICPLCRLIYACVPAGFT-------------Y--------YD-R-------
-------------GIF----INANT-------DLEEMLR---INNL-----VFE--NVWAEN--------------KDGKSLYAALV-----LGML-KEMN-----------EHAEYELS-DIQVVR-LEKE--------------R-YSFSILSRKFLNIIKKC----------------------------------------------------------------------------------------------------------------RTDA-----------------
-----------------------------LLLNSHWY----------------DTQSL-PL-------------------------------------------------------------------------------------------------------------------

EYD69295 ---------------------------------MVEKIVIRSED---W---------LNKAGIVGLYRI--------------------------------L---KERDERAD-----------IF------VEED--QISFSADL-----LQ---NFSEKYFQYFIKR-------------YKNVLSLYRILNFTANISQYEEKNYETFHEEDLEKL-----------NDHVEDVKKYL-------------KSNSYRAMYPLIRCPFDPLQKERELKKVNLKKTESL---
----KDRISDIQKLLADLKEIHDFLRQEDS---------QKYIGAKNAM-----YGIIQN-AWKGISILN-------------PQ--VK-----------EQ-NMYL----------EFDKYF-VQTAREYL-----------------------------------------------------------------------EQEKTKFK--Y---------RCFSCGEAI--------------------------KDTRIDLSFMN--HIGFDVA--R--KTSHV---------------W------D--FN-NY--VHICPLCRLIYACVPAGFT-------------Y------------YD-R-------

---------GIF----INANT------DLEEMLR---INNL------VFE--NVWAEN-----------------KDGKSLYAALV-----LGML-KEMN----------EHTEYELS-DIQVVR-LEKE--------------R-YSFSILSRKFLNIIKKCRTDLEYIRKS-------------------SFKEGRDIYYL--------------------------------------------------YN-ETMKRLMQG----ENLFLLIHKL-IQLRI----------------------------SNSENCYY-----KMGTVSTIIKIND--------IFLK-----------EVG---
--YMQDEKKEYN----------------------PLERARIIGHHLQEAY-GGF-EEGKY-------------------------NKKLDGIAYRML-NALKTNNKTAFMD-SLINAHMY----VQ-----------KPIPS-LF-SDYL--------NQD-----L-VFKELGYAFVTGMLGEEWKNKDNQS-------KN-----EKEG-R-----------------------------

KDE61658 -------------------------------------------------------------------------------------------------------------------YLEQEKTK------FK-----------------------------------------------------------------------------------------------------------------------------------------------------------------Y----------
------------------------------------------------RCFSCGEAI-------------------KDTRIDLSFMN--HIGFDVA--R--KTSHV-------------------W------D--FN-NY--VHICPLCRLIYACVPAGFT------------Y---------LYDR----------------------GIF----INANT------DLEEMLR----INNL------VFE--NVWAEN---------------KDGKSLYAALV-----LGML-KEMN---------EHAEYELS-DIQVVR-LEKE---------------R-
YSFSILSRKFLNIIKKCRTDLEYIRKS--------------------SFKEGSDIYYI----------------------------------------------YN-ETMKRLMQG----ENLFLLIHKL-IQLRI----------------------------SNSENCYY-----KMGTVSTIIKIND--------IFLK---------EVG-----YMQDEKKEYN--------------------PLERARIIGHHLQEAYGGFEEGKY-------------------------NKKLDGIAYRML-NALKTNNKTAFMD-SLINAHMY------VQ-------
KPIPS-LF-SDYL--------NQD-----L-VFKELGYAFVTGMLGE----EWKNKDNQSKNE-----KEGR-------------------------------

PWV46879 ------------------------------------MGEIITIRSDD---W--------LKNAGICGLYNI-------------------------------------LEKAGYKDQ--------IE-----VSSD--EIKFPVEL-----LK---GFSDMFFNYLIET-----------------YKDTFSLFRIINTEKLLDNWIENDFEKFEKKELDLL-------------NLNIENFKKYI-------------KSNSYKSAYKLI------------GRNVDPELLEKKLQKIKLKKKEKV-------
LDRIEDIKSQIATMKEII--NYFKSYD--------AIKYIGAKNAI-------YNIIRN-AWDGVSFLN---------RQ--CK-------NP-DMYE---------EYQDYF-VNPAIEYI----------------------------------------------------------------ENF--DKNKAKYK--Y---------QCLACSSKI-----------------------KDLNSDIGFIR--EMGFDTN--R--KTSHV--------------------W------D--FN-NY--VAICPICRLVYSCVPAGIN-----------Y---------AFNK-----------
-----GLF----INYSM------DLNNLIK---INKK------INF--ELHENS--------------------YSGISIYKSIQ-----DNMN-ENLN--------DN-SQYELS-DVQVVR-FYRD-VD--------KDSIK-YNFNIINKVVLTISSCKTELNTIKQG--------------------KYSENSNDKYI---------------------------------------------------------YN-EVMEKLLNN----QNQFLLVQKL-LHYKL---------------------------SMPDSTYY-----NAGIVSSILKINI----------KYLE-----------EAG--
--YVKNLENN-----------------------LVDEARIAGYYLQKAYNISGKGKVEIDEVLQQESSESEDNKKNNNLAYNKKLNGIAYRML-NALKTNNKHSFMD-TLINAYMY-----VG-----------KSIPK-IF-AENL--------EDD-----L-VFKNIGYAFITGMIGY----------IKEE-----NKDK-----------------------------

EFL54064 ------------------------------------MGEIITIKSDD---W--------LKNAGICGLYNI-------------------------------------LEKAGYKDQ--------IE-----VRSD--EIEFPVEL-----LK---GFSDMFFNYLIET-----------------YKDTFSLFRIINAEKLLDNWIENDFEKFEKKELDLL-------------NLHIENFKKYI-------------KSNSYKSAYKLI------------GGNVDPELLEKNLQKIKLKKKEKV-------
LDRIEDIKSQLATMKEII--SYFKSDD--------AIKYIGAKNAI-------YNIIRN-AWDGVSFLN---------RQ--CK-------NP-DMYE---------EYQDYF-VNPAIEYI----------------------------------------------------------------EDI--DKNKTKYK--Y---------QCLACSSKI-----------------------KDLNSDIGFIR--EMGFDTN--R--KTSHV--------------------W------D--FN-NY--VAICPICRLVYSCVPAGIN-----------Y---------AFNK-----------
-----GLF----INYSM------DLNDLIK---INKK------INF--EIHENS--------------------YSGISIYKSIQ-----DNMN-ENLN--------EN-SQYELS-DVQVVR-FYRD-VD--------RDSIK-YNFNIINKVVLTISSCKTELNTIKQG--------------------KYSENSNDKYI---------------------------------------------------------YN-EVMEKLLNN----QNQFLLVQKL-LHYKL---------------------------SIPDSTYY-----NAGIVSSILKINI----------KYLE-----------EAG--
YVKKLENN-----------------------LVDEARIAGYYLQKAYNISGKGKVEIDEILQQESSESEDNKRNNNLAYNKKLNGIAYRML-NALKTNNKHSFMD-TLINAYMY-----VG-----------KSIPK-IF-AENL--------EDD-----L-VFKNIGYAFITGMIGY----------IKEE-----NKDK-----------------------------

EFK93763 ------------------------------------MGEIITIKSDD---W--------LKNAGICGLYNI-------------------------------------LEKAGYKDH--------IE-----VRSD--EIEFPVEL-----LK---GFSDMFFNYLIET-----------------YKDTFSLFRIINAEKLLDNWIENDFEKIEKKELDLL-------------NLHIENFKKYI-------------KSNSYKSAYKLI------------GGNVDPELLEKNLQKIKLKKKEKV-------
LDRIEDIKSQLATMKEII--SYFKSDD--------AIKYIGAKNAI-------YNIIRN-AWDGVSFLN---------RQ--CK-------NP-DMYE---------EYQDYF-VNPAIEYI----------------------------------------------------------------EDI--DKNKTKYK--Y---------QCLACYSKI-----------------------KDLNSDIGFIR--EMGFDTN--R--KTSHV--------------------W------D--FN-NY--VAICPICRLVYSCVPAGIN-----------Y---------AFNK-----------
-----GLF----INYSM------DLNDLIK---INKK------INF--EIHENS--------------------YSGISIYKSIQ-----DNMN-ENLN--------EN-SQYELS-DVQVVR-FYRD-VD--------RDSIK-YNFNIINKVVLTISSCKTELNTIKQG--------------------KYSENSNDKYI---------------------------------------------------------YN-EVMEKLLNN----QNQFLLVQKL-LHYKL---------------------------SIPDSTYY-----NAGIVSSILKINI----------KYLE-----------EAG--
YVKKLENN-----------------------LVDEARIAGYYLQKAYNISGKGKVEIDEILQQESSESEDNKRNNNLAYNKKLNGIAYRML-NALKTNNKHSFMD-TLINAYMY-----VG-----------KSIPK-IF-AENL--------EDD-----L-VFKNIGYAFITGMIGY----------IKEE-----NKDK-----------------------------

WP_009643660 --------------------------------MNGITIKNSD-----W---------LFNCGILGLYNI-----------------------------------LKHNNIDDSED---------IS----LSQD--ELTFPVSR-------LE---NFEEKYFSYLIDT------------YEKTFSHIRITSFEDFLQNHEDSNFENFNTHSLDRL--------------NSQIEQVKRYL--------------KSNSYVSAYKML------------DFGFDPIAKEKELKKVSLKKNETV---
----FDKQSEIKKQMDILKETV--EFFKRKE----------VRKYIGGKNAM--------YSIIKN-AWSNISIMN----------RQ--NK-------NP-DIYE---------EFSNYF-VEPVKEYL------------------------------------------------------------DEI---DSSRLKSK--Y---------RCVTCNAKI---------------------KNLDIDLSFMR--EIGFDVN--R--KTSHV---------------------W------D--FN-NY--VAICPVCRLVYACVPAGFT------------YL--------YN-R----
----GLF----VNSSM-------DLQEIVR---INQI------IKK--DVLQSS--------------EGGTVIYRALQ----KQMS-RKIN-----------ET-PGYELS-DVQVIR-FSHD-QD-------SDNVS-YTFNIITKFILKTIENCDKCFERIQNG--------------------RLSEGNATMYL---------------------------------------------------------YD-EVMNRLVNN----ENQFLLIHKL-IHYKL----------------------------SIPDKAMY-----SMSTIENILKINA--------EYLR---------
EVG-----VMEKDDKN---------------------MIRIAKSHGYYLRDAYGDEKKDNSGNKADVKSNDISNETNVKAKGTGYNKKLGSIAYRML-NALKTNNKYAFMD-TMINAHMY------VQ--------------KPIPA-IF-AENL-------END-----L-VFKNIGYAFVTGMIGY----TDEKKVD---EKG-----NEEN-------------------------------

SMB94280 --------------------------------MSKNIELKVSD--W---------IYNVGLLGLYNI---------------------------------------LEYKDEF----------VE-----VKKD--GISFELEA-----LE----GFEEYYFKYLIKK------------YYSTLSINKINSFENIISNWEDEDFKVFDEKSLETL------------NLQIESVKKYI------------TSNSYKSAYQLI------------DSEFNPLIKEKQLKKIKLSKKDTI-------
EGRLEDIKNQVSILNEII--EYFKRKD----------SIKYIGAKNVM--------YTLIRN-AWDNVSILN----------RQ--NK-------NP-DMYD---------EVANYF-VKPAIEYL---------------------------------------------------------DGY---EKSKNKDK--Y---------LCMSCGSRI-------------------------PSLNNDICFIR--ELGFDTK--K--KNSHV-----------------------W------D--FN-NY--VGICPMCKLVYTCIPAGFT------------Y----------AYNR--------
---------GLF----VNYSN------NFKSLIE---INNN------IKE---DVFKET------------------SKNTSIYYAIQ-----KNMD-EKSN------------ED-MKYELS-DVQIVR-FFRN-ID--------SDQVK-YSFNILNRPIQRVINECSGSLKFIRGA--------------------YFEEGKKNVYV-----------------------------------------------------YN-EVMNNLFNN----QNDFLIIHKM-LHYKI---------------------------SNPDKTRY-----SSEHIQNVIRINN----------SYLR--------
EVG-----YMNDKSNGK-------------------YLYFSRVSGENLRASYKKMDA----------------------------VEKLNGIAYRML-NALKTSNKHAFMD-TLIKASMY-----AN-----------EAIDD-IF-SNTM--------EDD-----L-KFKNAGYSFITGMLGK----QNNSKNDS-EDN-----GGEI-----------------------------

EGY80091 --------------------------------MEKRIELKTGD---W----------LYNTGLLGLYNI-----------------------------------LKYKDEY----------VE-----VSKE--GISFELEA-----LE---KFEEYYFKYLIDT------------YYLTLSINRIVSFENSILNWESEDFKNFNDKSLENL--------------NIQIDNVKKYM-------------ISNSYKKAYSLIDSEFNPLVKEKELKKIKLSKKDTV----------
ESKMSEIKYQISLLKETIDYFKFKDT----------KKYIGAQNII-------YNIIRN-AWDNVSILN----------RQ--NK-------NP-DMYD---------EIANYF-VKPAIEYL------------------------------------------------------------------------------------------------DEYEEKKNKSR--Y---------LCMSCGSRI-------------------------SSLNNDICFIR--ELGFDTK--K--KNSHV--------------------Y------D--FN-NY--VGICPMCKLVYTCIPAGFT------------Y---------AYNR------
----GLF----VNYSS------NFKNLIE---INNN------IKE----------DVFKEI-NKNTSIYYAIQ-----KNMD-EKSN------------EDMKYELS-DVQIVR-FFRN-ID-------SDQVK-YSFNILNRPI----------------------------------------------------------------------------------------------------------------------------------------------------------------------------------------------------------

EMS70152 --------------------------------MIYLENEIIRLALGD---W---------SWNASVAGFINI------------------------------------IGEEN---------VT-----YDGA--SAQIPVNC-------LE---DFEDRYFRYFIDT------------YEKALPWYRIVSFKNTIGFYNENGFEGLDLKKLKSI------------NDYIKDTVKRY-------------------LTSNSFKSAFELMGEAERMAELEKNLSKVREPKDDTD-
-FEKVKEEVIGELKKQFEIINKII--EYCGNE----------NGRRFIGAKNVI--------YTIIRN-GWNGISFLN--------PQ--TK-------IK-DIYE----------DYRTYF-VDEAREYL--------------------------------------------------------AADKQKFR--Y---------KCFTCGAPI------------------------RDMNNDMSFLN--QSGFDTA--R--KPSHV-------------------------W------D--FV-ND--IALCPICRLIYSCLPAGFV-------------Y----------VNNT--
-------------GLY-----INANV------NMRYNVR----VNRS-------ARD----------------SVLKEGRN-ETASRRIYKVL------VEAM-ERER--------IEKSRYELA-DVQVVR-YENE--------------R-FKFNLLPESTIKLIHKY---HDEISGL------------------IR-TGYKEGKESYSI---------------------------------------------------------------YE-QVLENIFNN-----QNLFLLVYKL-LHFKT---------------------------------ASNKDCFF-----HMGHVEDILSINA----------GLIK-----
----SLG-------GMENMDENRN--------------------------YINEAKNSGYRLRKAFMKNNEK----------------------TGKIPGISYRLL-NALKTGNRNMFMD-VLLNSYLY-----TG---------NKVPA-VI-SEIF-------RSD-----E-LFSTIGYSFVTGFIAERSDDSGNEANNEKTVN-----GEEN-------------------------------

HCT96597 --------------------------------MTDIRVRLDD---W---------QKNAGCVGFINI-------------------------------------VGKEN---------VQ-----LTEN--GFCFNSTV-------LE---DFATKYFDYLIET--------------YKPTLSWYKIIAYKDTLEKHERENFENFDEKDLERL-------------NIYIIDVCKRY-----------LTSASYKASYPLI-----------KQDFDMLEAGKQSKAIGRLKKRET--
FEEKKTEILEEVREAYERIKQII--AYCESP-----------EGRKYLAAKNVI--------YTIVKH-AWDSVSFLN--------PQ--TK-------EK-DVYK------------DFEQYF-IAPVLEYL-----------------------------------------------------------------------ESDKSRYK--L---------TCCSCENPV---------------------------KNASETLSFLN--KTGFDVN--R--KNSHA--------------------W------D--FV-ND--LIICPVCKLIYACIPAGFT------Y----V---------YN-Q--
------------GMF-----VNANS------SIEELLR---TNNL-----IKM--EVL----------------TLNRAS-VREVNTYSALV-------HAF--QKQD------------RDQFRLQTE-DVQVVR-YQNE----------------T-YRFNILSKNTLRILNQS---EAELSQL--------------------LH-TWVKENGETYSL---------------------------------------------------------------YQ-KVVEKLFNN-----ENMLTLIHKL-LVYKA-------------------------------------TKTENPGY-----QIRHIDALIKINQ---------HFLG---
------GIL-----TMNEVTQD---------------------EIKKIKGAAWYFKKGYDN-------------------------PNKVNGVSYKLL-NALKIGNRDEFMN-VILNCYAY-----LN----------QQVPQ-FF-NRVF---------ENE-----E-TFKTIGYSFVAGIIGPVEMN-----------EY----GGKA-------------------------------

WP_126762559 --------------------------------MTDIRVRLDD---W---------QKNAGCVGFINI-------------------------------------VGKEN---------VQ-----LTEN--GFRFNSTV-------LE---DFATKYFDYLIET--------------YKPTLSWYKIIAYKDTLEKHERENFENFDEKDLERL-------------NTYIIDVCKRY-----------LTSASYKASYPLI-----------KHDFDMLEAGKQSKAIGRLKKRET--
FEEKKTEILEEVREAYGRIKQII--AYCESP-----------EGRKYLAAKNVI--------YTIVKH-AWDSVSFLN--------PQ--TK-------EK-DVYK------------DFEQYF-IAPVLEYL-----------------------------------------------------------------------ESDKSRYK--L---------TCCSCENPV---------------------------KNASETLSFLN--KTGFDVN--R--KNSHA--------------------W------D--FV-ND--LIICPVCKLIYACIPAGFT------Y----V---------YS-Q--
------------GMF-----VNANS------SIEELLR---TNNL-----IKM--EVL----------------TLNRAS-VREVNTYSALV-------HAFQ-KQDR----------DQ-FRLQTE-DVQVVR-YQNE--T-------------YRFNILSKNTLRILNQS---EAELSQL--------------------LH-TWVKENGETYSL---------------------------------------------------------------YQ-KVVEKLFNN-----ENMLTLIHKL-LVYKA-------------------------------------TKTEN-PGY-----QIRHIDALIKINQ---------HFLG---
------GIL-----TMNEVTQD---------------------EIKKIKGAAWYFKKGY-DN-------------------------PNKVNGVSYKLL-NALKIGNRDEFMN-VILNCYAY-----LN----------QQVPQ-FF-NRVF---------ENE-----E-TFKTIGYSFVAGIIGP----------VEMNE-----YGGK-A-------------------------------

WP_023606046 --------------------------------MTDIRVRLDD---W---------QKNAGCVGFINI-------------------------------------VEKEN---------VQ-----LTEN--GFRFNSTV-------LE---DFATKYFDYLIET--------------YKPTLSWYKIIAYKDTLEKHERENFENFDEKDLERL-------------NTYIIDVCKRY-----------LTSASYKASYPLI-----------KHDFDMLEAGKQSKAIGRLKKRET--
FEEKKTEILEEVREAYGRIKQII--AYCESP-----------EGRKYLAAKNVI--------YTIVKH-AWDSVSFLN--------PQ--TK-------EK-DVYK------------DFEQYF-IAPVLEYL-----------------------------------------------------------------------EDSKSLYK--L---------TCCSCENPV---------------------------KNASETLSFLN--KTGFDVN--R--KNSHA--------------------W------D--FI-ND--LIICPVCKLIYACIPAGFT------Y----V---------YS-Q--
------------GMF-----VNANS------SIEELLR---TNNL-----IKM--EVL----------------TLNRAS-VREVNTYSALV-------QAFQ-KQDR-----------DQ-FRLQTE-DVQVVR-YQNE--T-------------YRFNILPTNSLRTIELA---NKQLEVL--------------------LH-TWVKENGETYSL---------------------------------------------------------------YQ-KVVEKLFNN-----ENMLTLIHKL-LVYKA-------------------------------------TKTEN-PGY-----QIRHIDALIKINQ---------HFLG---
----GIL-----TMNEVTQD---------------------EIKKIKGAAWYFKKGY-DN-------------------------PNKVNGVSYKLL-NALKIGNRDEFMN-VILNCYAY-----LN----------QQVPQ-FF-NRVF---------ENE-----E-TFKTIGYSFVAGIIGP----------VEMNE-----YGGK-A-------------------------------

WP_086348589 --------------------------------MEGKVSLYLGD---W---------QFNAGLIGLVNV-------------------------------------LGREN---------VE-----LAYD--HIVFDLNQ-------LD---RFEEAYFAYFIKT-------------YKKLLSWHKIVSYKQRLVQFESDDFEQFTETDLENL-----------NKYIKDILKYY------------LKSASYKAAYSLI-----------PSDTDVLALEKEIKTVGKMKKGET--
FADKKPEIIQEIKEQLPKLKEAI--DFCESS-----------QGKKYLAAKNVI--------YTIVKH-AWDSVSFLN--------PQ--TK-------IP-DMYV--------DYAATF-VOPAKVYL----------------------------------------------------------EEN--EEEQTKYK--Y----------HCANCNRKI---------------------------KDLKNDVSFLN--ATGFDVS--R--KAGHV---------------------W------D--SF-ND--TAVCPLCKLVYSCVSAGFT-----Y---------V---------
YN-D--------------GMF-----INAST------NLDDLYR---MNYT-------LKH--ETL----------------NAGGEN-ISEVSPYRALI-------QNLQ-KKDLQ-------EKQQLE--DVTLVR-YENE--------------T-YRFNILPTNSLRTIELA---NKQLEVL--------------------IP-TGFKEINTNFRI---------------------------------------------------------------YK-LVLQSLFNQ-----ENLFYLIHKL-LYFKL----------------------------------TNVGNLYY-----QPFHVRNIIEINS--------
IFLG----------GLN-----HMTEEKTKTLPGD---------------------ISWRVNHLGEKFKAEY-SAR-FN------------------------ENKLITIAHQML-GALKINNRDRFMD-VLLNCYSY----IN----------KPVPK-TL-LDVF--------SSD-----E-NFKTIGYSFVAGIIGK-------------TEKT-----TEEE-K-------------------------------

KHS60163__2_ --------------------------------MQTEIEVRADD---W---------LINASIVGFLKI------------------------------------VGKEN-------VR-----IDGQ--SIYFTTDL-------LE---NFETKFFNFFINE-------------YKETLVWSKIISYKDSMERFRTDDFASFDEKALTDF-------------NNYIRDVVKYN--------------LKKDNYIKVFKLI-----------DESANVTKWVGNLKPINLKKAEKW---
NDVKSEKIELIKESFKHLDSII--NFCASE-----------KGLKYLGARSVS-------YTVINK-GWSDVSFLN--------K--QA-------------SYEGYESYF-LKPLLEYL---------------------------------------------------------------------DTDLSKAK--Y---------KCFNCNQPI---------------------------KTLASGLSFMN--DTGFDVG--K--KASHV-----------------------W------D--FN-ND--VAICPVCRLIYSCVPAGFT--------Y----V--------YG-E---
--------------GMF-----VN-DS---YGIDELYR---VNER------IRN--SIL-----------------RFNRDS-INSANTYRALV-----ESIT-IEHE----------NN-RRYELA-DIQLVR-YENE--------------H-YRFNLLSKKMLHIVNDSKGILKSLIRC---------------------GYKEGNNNL-------------------------------------------------------YK-EVIQHLMNN-----ENLFSLIHKL-IYYKQ---------------------------------TSVNGLYY-----NMGNVSGILDINT---------KFLK------
-EIE-------VMKNVTQS--------------------ELWLAKSCGTKFKEGYVKKKS-------------------------ENKISGITYKLL-NSLKINDKDGFMD-TLLNSYSY-----LA----------IPVPN-VF-MHAF--------SNT-----E-AFKSVGYAFMLGVGGEATNNKDG---------GT-----TDEK-------------------------------

EHN61774 --------------------------------MQTEIEVRADD---W---------LINASIVGFLKI------------------------------------VGKEN-------VR-----IDGQ--SIYFTTDL-------LE---NFETKFFNFFINE-------------YKETLVWSKIISYKDSMERFRTDDFASFDEKALTDF-------------NNYIRDVVKYN--------------LKKDNYIKVFKLI-----------DESANVTKWVGNLKPINLKKAEKW---
NDVKSEKIELIKESFKHLDSII--NFCASE-----------KGLKYLGARNVS-------YTVINK-GWSDVSFLN--------KQ-----------APPD-------SYEGYESYF-LKPLLEYL---------------------------------------------------------------------DTDLSKAK--Y---------KCFNCNQPI---------------------------KTLASGLSFMN--DTGFDVG--K--KASHV-----------------------W------D--FN-ND--VAICPVCRLIYSCVPAGFT--------Y----V--------YG-E---
--------------GMF-----VN-DS---YGIDELYR---VNER------IRN--SIL-----------------RFNRDS-INSANTYRALV-----ESIT-IEHE----------NN-RRYELA-DIQLVR-YENE--H-------------YRFNLLSKKMLHIVNDS----------KGI---------------LK--SLIRCG-----Y-----------KEGNLNL-------------------------------------------------------YK-EVIQHLMNN-----ENLFSLIHKL-IYYKQ---------------------------------TSVNG-----------------LYYN-MGH---------VSGILDINT---------KFLK------
-EIE-------VMKNVTQS--------------------ELWLAKSCGTKFKEGY-VKK-KS-------------------------ENKISGITYKLL-NSLKINDKDGFMD-TLLNSYSY-----LA----------IPVPN-VF-MHAF--------S--NTE-----AFKSVGYAFMLGVGGEATNNKD---------GG-----TTDE-K-------------------------------

KNX95742__9_ --------------------------------MQTEIEVRSND---W---------LINAGLVGFLKI-------------------------------------VGKDN-------VK-----IDGQ--SIYFTADL-------LE---DFETKYFNFFIQE-------------YKETLAWSKIVSYKEAMERFRADGFASFDEKDALNNL-------------NKYVKNVVKFY-------------LKKANYIKVFKLI-----------NSDANVNKWLENLKPINLTNLTKKQKW---
EEVKPDIVEKVEQIYTQLDVII--DFCTSE---------KGLRYLGAKNLI-------YSVINK-GWSGVSFLF-------KQ--TK-------FI-DPYL---------DYKTTF-VDPVIEHL----------------------------------------------------------------------DTDLSKAK--Y---------NCFNCNQPI---------------------------KNLKLDLSFMN--EVGFDTA--R--KTSHV-----------------------W------D--FN-ND--VATCPICRLVYSCVPAGFT----------Y---------YG-E---
-------------GMF-----VN-DS---YGVNELHH---VNER------MRK--SIL-----------------RFNKDG-INSTNTYQALV-----ESIT-SEHE----------NN-RRYELA-DIQLIR-YENE-------------H-YHFNLLSKKMLHIMNDS---EKILKSL-----------------------IR-CGYNEGNLNL-------------------------------------------------------YK-EVIQHLMNN-----ENLFTLIHKL-IFYKQ---------------------------------SSVNGLYY-----NMWNVSGVLEINT----------KFLK------
-----EIE-----VMTNIEER-------------------DLTNAQKSGGDFKQAY-VDK-NS--------------------------ENKVSSITYKLL-NALKLNDKNGFMD-ILLNSYSY-----LG----------KPVPT-VF-MSSF---------SSK-----E-TFKSIGYAFMIGVSANKTKSKNKD-----------GG-----NTDE-K-------------------------------

EFR88692 --------------------------------MQTEMAVRADD---W---------LINAGLVGFLNI--------------------------------------VGKEN-------VR-----IDGQ--SIYFTADL-------LE---DFETKYFNFFIQE-------------YKETLAWSKIVSYKETMERFRVDGFASFDEDALDNL-------------NKYVKDVVKFY-------------LKKSNYIKVFPLI-----------DPNADVNKWLENLKTINLTKKQEW--
-EDVKADILEQVKQIYTQLDAII--DFCASD-----------KGLRYLGAKNLI-------YSVINK-GWSGLSFLS-------KQ--TK-------FI-DPYL---------DYKTSF-VEPVIEYL----------------------------------------------------------------------NTDLSKAK--Y---------NCFNCNQPI---------------------------KNLKLDLSFMN--EVGFDTA--R--KTSHV-----------------------W------D--FN-ND--VATCPICRLVYSCVPAGFT-----------Y---------YG-E---
-----------GIF-----VS-DS---YGVNELYR---VNER------MRN--SIL-----------------RFNKDG-INSANTYRALV-----ESIT-TEHE----------NV-RRYELA-DIQLVR-YENE--H-------------YHFNLLSEKMLHIVNDS---------KGI------------------LK-SLIRCG-----Y---------EEGSLNINL-------------------------------------------------------YK-EVIQRLMNN-----ETLYLFIHKL-IY-------------------------------------------------------------------------------------------------------------

ACK40458__3_ --------------------------------MQTEIELRADD---W---------LINAGLVGFLNI-------------------------------------VGKEN-------VR-----INGQ--INQGYKADL-----LV---GFEDKYFNFFIQE--------------YKETLAWFKIVSYKDTMERFRAEDFASFDEKALNDL-------------NKYMKDVVKFY-------------LKKSNYIKVFSLI-----------DAKANVTKWLENLNAINLTKKQQW---
EDVKVDIVEKVKRNYIQLDFII--DFCLGE-----------KGLKYLGAKNLI-------YSVINK-GWTGTSFLY-------TK--TK-------FA-DPYL---------DYKTSF-LDPVLDYL----------------------------------------------------------------------DSDLSKAK--Y---------NCFNCNLPI---------------------------KNLKTGLSFMN--DIGFDIE--K--KTSHV-----------------------W------D--FN-ND--VATCPICRLIYSCVPAGFT----------Y----V--------YG-E---
-------------GMF-----VN-DS---YGVEELHR---VNER------MRN--SIL-----------------RFNKDG-INSTNTYRALV-----ESIT-IEHE----------NN-RRYELA-DIQLVR-YENE--H-------------YRFNLLSKKMLHIVNDS---------KGI------------------LK-SLIRCG-----Y---------KEGNLNVNL-------------------------------------------------------YK-EVIQHLMNN-----ENLFTLIHKL-IYYKQ----------------------------------TSVNG-----------------LYYN-MGH---------VSGILDINT---------KFLK------
----EME------VMTNISQN-----------------------QLWFVQNCGKFKEGY-VKK-KS-------------------------ENKISGITYKLL-NSLKVNDKDGFMD-TLLNSYSY-----LS----------MPIPD-VF-IKMF------S--NNE-----AFKSVGYAFMLGVGGEKTKKEN-----------GG-----NTDE-K-------------------------------

KKD51591 --------------------------------MQTEIEVRADD---W---------LINAGLTGFLNI-------------------------------------VGKEN-------VR-----INGQ--SIYFTADL-------LE---DFETKYFNFFIQE--------------YKETLSWYKIVSYKETMEQFRIDEFASFDEVALGNL-------------NKYMKDVVKFY------------LKKANYIKVFPLI-----------DPSANVTKWLEKLNTINLTKKQKW---
DEVKADIIEEVKQTYLQLDLII--DFCASE-----------KGLKYLGAKNLI-------YSVINK-GWSGVSFLF-------KQ--TK-------FI-DPYL---------DYKTTF-VDPVLEYL----------------------------------------------------------------------DTDLSKAK--Y---------NCFNCNQPI---------------------------KNLKLDLSFMN--DVGFDTA--R--KTAHV-----------------------W------D--FN-ND--VATCPICRLIYSCVPAGFT----------Y----V--------YG-E---
-------------GMF-----VN-DS---YGVEELHR---VNER------MRN--SIL-----------------RFNKDG-INSTNTYRALV-----ESIT-MEHE----------NN-RRYELA-DIQLVR-YENE--H-------------YRFNLLSKKMLHIVNDS---------KGI------------------LK-SLIRCG-----Y---------KEGNLNINL-------------------------------------------------------YK-EVIQHLMNN-----ENLFSLIHKL-IYYKQ----------------------------------TSVNG-----------------LYYN-MGH---------VSGILDINT---------KFLK------
--EIE------VMKNVTQS--------------------ELWLAKSCGTKFKEGY-VKK-KS-------------------------ENKISGITYKLL-NSLKINDKDGFMD-TLLNSYSY-----LA----------MPIPD-VF-IKMF------S--NNE-----AFKSVGYAFMLGVGGEKTKKED-----------GG-----NTDE-K-------------------------------

CBY03036__21_ --------------------------------MRTEIEVRAND---W---------LINAGIGFLNI-------------------------------------VGKEN-------VR-----INGQ--SIYFTADL-------LE---DFETKYFNFFIQE--------------YKETLSWYKIVSYKETMEQFRIDEFASFDEVALGNL-------------NKYMKDVVKFY------------LKKVNYIKVFPLI-----------DPSANVTKWLENLNTINLTKKQRW---
DEVKADIIEEVKQTYLQLDLII--DFCASE-----------KGLKYLGAKNLI-------YSVINK-GWSGVSFLF-------KQ--TK-------FI-DPYL---------DYKTTF-VDSVLDYL----------------------------------------------------------------------DTDLSKAK--Y---------NCFNCNQPI---------------------------KNLKLDLSFMN--DVGFDTA--R--KTSHV-----------------------W------D--FN-ND--VATCPICRLIYSCVPAGFT----------Y----V--------YG-E---
-------------GMF-----VN-DS---YGVEELHR---VNER------MRN--SIL-----------------SFNQDG-INSTNTYRALV-----ESIT-MEHE----------NN-RRYELA-DIQLVR-YENE--H-------------YRFNLLSKKMLHIVNDS---------KGI------------------LK-GLIRCG-----Y---------KEGNLNVNL-------------------------------------------------------YK-EVLQHLMNN-----ENLFTLIHKL-IYYKQ----------------------------------TSVNG-----------------LYYN-MGH---------VSGILDINT---------KFLK------
-------EME------VMTNISQN-----------------------QLWFVQNCGKEFKEGY-VKK-KS-------------------------ENKISGITYKLL-NSLKVNDKDGFMD-TLLNSYSY-----LA----------MPIPD-VF-IKMF------S--DNE-----AFKSVGYAFMLGVGGEKTKKED-----------GG-----NTDE-K-------------------------------

KTA57795__2_ --------------------------------MRTEIEVRAND---W---------LINAGLTGFLNI-------------------------------------VGKEN-------VR-----INGQ--SIYFTADL-------LE---DFETKYFNFFIQE--------------YKETLSWYKIVSYKETMEQFRIDEFASFDEVALGNL-------------NKYMKDVVKFY------------LKKANYIKVFPLI-----------DPSANVTKWLENLNTINLTKKQKW---
DEVKADIIEEVRQTYTQLDLII--DFCASE-----------KGLKYLGAKNLI-------YSVINK-GWSGVSFLF-------KQ--TK-------FI-DPYL---------DYKTTF-VDPVLEYL----------------------------------------------------------------------DTDLSKAK--Y---------NCFNCNQPI---------------------------KNLKLDLSFMN--DVGFDTA--R--KTAHV-----------------------W------D--FN-ND--VATCPICRLIYSCVPAGFT----------Y----V--------YG-E---
-------------GMF-----VN-DS---YGVEELHR---VNER------MRN--SIL-----------------RFNKDG-INSTNTYRALV-----ESIT-MEHE----------NN-RRYELA-DIQLVR-YENE--H-------------YRFNLLSKKMLHIVNDS---------KGI------------------LK-SLIRCG-----Y---------KEGNLNVNL-------------------------------------------------------YK-EVIQHLMNN-----ENLFTLIHKL-IYYKQ----------------------------------TSVNG-----------------LYYN-MGH---------VSGILDINT---------KFLK------
-------EME------VMTNISQN-----------------------QLWFVQNCGKEFKEGY-VKK-KS-------------------------ENKISGITYKLL-NSLKVNDKDGFMD-TLLNSYSY-----LS----------MPIPD-VF-IKMF------S--NNE-----AFKSVGYAFMLGVGGEKTKKED-----------GG-----NTDE-K-------------------------------

CUK78258 --------------------------------MQTEIEVRAND---W---------LINAGLTGFLNI-------------------------------------VGKEN-------VR-----INGQ--AIYFTADL-------LE---DFETKYFNFFIQE--------------YKETLSWYKIVSYKETMEQFRIDKFASFDEVALGNL-------------NKYMKDVVKFY------------LKKANYIKVFPLI-----------DPSANVTKWLENLNTINLTKKQKW---
DEVKADIIEEVKQTYLQLDLII--DFCASE-----------KGLKYLGAKNLI-------YSVINK-GWSGVSFLF-------KQ--TK-------FI-DPYL---------DYKTTF-VDPVLEYL----------------------------------------------------------------------DTDLSKAK--Y---------NCFNCNQPI---------------------------KNLKLDLSFMN--DVGFDTA--R--KTAHV-----------------------W------D--FN-ND--VATCPICRLIYSCVPAGFT----------Y----V--------YG-E---
-------------GMF-----VN-DS---YGVEELHR---VNER------MRN--SIL-----------------RFNKDG-INSTNTYRALV-----ESIT-MEHE----------NN-RRYELA-DIQLVR-YENE--H-------------YRFNLLSKKMLHIVNDS---------KGI------------------LK-SLIRCG-----Y---------KEGNLNVNL-------------------------------------------------------YK-EVIQHLMNN-----ENLFTLIHKL-IYYKQ----------------------------------TSVNG-----------------LYYN-MGH---------VSGILDINT---------KFLK------
-------EME------VMTNISQN-----------------------QLWFVQNCGKEFKEGY-VKK-KS-------------------------ENKISGITYKLL-NSLKVNDKDGFMD-TLLNSYSY-----LS----------MPIPD-VF-IKMF------S--NNE-----AFKSVGYAFMLGVGGEKTKKED-----------GG-----NTDE-K-------------------------------

CUL11411 --------------------------------MQTEIEVRADD---W---------LINTGLTGFLNI-------------------------------------VGKEN-------VQ-----INGQ--AIYFTADL-------LE---DFETKYFNFFIQE--------------YKETLSWYKIVSYKEKMEQFRIDEFASFDEVALGNL-------------NKYMKDVVKFY------------LKKGNYIKVFPLI-----------DPSANVTKWLENLNTINLTKKQKW---
DEVKADIIEEVKQTYIQLDLII--DFCASE-----------KGLKYLGAKNLI-------YSVINK-GWSGVSFLF-------KQ--TK-------FI-DPYL---------DYKTTF-VDPVLEYL----------------------------------------------------------------------DTDLSKAK--Y---------NCFNCNQPI---------------------------KNLKLDLSFMN--DVGFDTA--R--KTAHV-----------------------W------D--FN-ND--VATCPICRLIYSCVPAGFT----------Y----V--------YG-E---
-------------GMF-----VN-DS---YGVEELHR---VNER------MRN--SIL-----------------RFNKDG-INSTNTYRALV-----ESIT-MEHE----------NN-RRYELA-DIQLVR-YENE--H-------------YRFNLLSKKMLHIVNDS---------KGI------------------LK-SLIRCG-----Y---------KEGNLNVNL-------------------------------------------------------YK-EVIQHLMNN-----ENLFTLIHKL-IYYKQ----------------------------------TSVNG-----------------LYYN-MGH---------VSGILDINT---------KFLK------
-------EME------VMTNISQN-----------------------QLWFVQNCGKEFKEGY-VKK-KS-------------------------ENKISGITYKLL-NSLKVNDKDGFMD-TLLNSYSY-----LS----------MPIPD-VF-IKMF------S--NNE-----AFKSVGYAFMLGVGGEKTKKED-----------GG-----NTDE-K-------------------------------

CUK62856__2_ --------------------------------MQTEIEVRADD---W---------LINTGLTGFLNI-------------------------------------VGKEN-------VR-----INGQ--SIYFTADL-------LE---DFETKYFNFFIQE--------------YKETLSWYKIVSYKEKMEQFRIDEFASFDEVALGNL-------------NKYMKDVVKFY------------LKKANYIKVFPLI-----------DPSANVTKWLENLNTINLTKKQKW---
DEVKADIIEEVKQTYLQLDLII--DFCASE-----------KGLKYLGAKNLI-------YSVINK-GWSGVSFLF-------KQ--TK-------FI-DPYL---------DYKTTF-VDPVLEYL----------------------------------------------------------------------DTDLSKAK--Y---------NCFNCNQPI---------------------------KNLKLDLSFMN--DVGFDTA--R--KTAHV-----------------------W------D--FN-ND--VATCPICRLIYSCVPAGFT----------Y----V--------YG-E---
-------------GMF-----VN-DS---YGVEELHR---VNER------MRN--SIL-----------------RFNKDG-INSTNTYRALV-----ESIT-MEHE----------NN-RRYELA-DIQLVR-YENE--H-------------YRFNLLSKKMLHIVNDS---------KGI------------------LK-SLIRCG-----Y---------KEGNLNVNL-------------------------------------------------------YK-EVIQHLMNN-----ENLFTLIHKL-IYYKQ----------------------------------TSVNG-----------------LYYN-MGH---------VSGILDINT---------KFLK------
-------EME------VMTNISQN-----------------------QLWFVQNCGKEFKEGY-VKK-KS-------------------------ENKISGITYKLL-NSLKVNDKDGFMD-TLLNSYSY-----LS----------MPIPD-VF-IKMF------S--NNE-----AFKSVGYAFMLGVGGEKTKKED-----------GG-----NTDE-K-------------------------------

PIL17074__8_ --------------------------------MQTEIEVRADD---W---------LINTGLTGFLNI-------------------------------------VGKEN-------VR-----INGQ--SIYFTADL-------LE---DFETKYFNFFIQE--------------YKPTLSWYKIVSYKETMEQFRIDEFASFDEVALGNL-------------NKYMKDVVKFY------------LKKANYIKVFPLI-----------DPSANVTKWLENLNTINLTKKQKW---
DEVKADIIEEVKQTYLQLDLII--DFCASE-----------KGLKYLGAKNLI-------YSVINK-GWSGVSFLF-------KQ--TK-------FI-DPYL---------DYKTTF-VDPVLEYL----------------------------------------------------------------------DTDLSKAK--Y---------NCFNCNQPI---------------------------KNLKLDLSFMN--DVGFDTA--R--KTAHV-----------------------W------D--FN-ND--VATCPICRLIYSCVPAGFT----------Y----V--------YG-E---
-------------GMF-----VN-DS---YGVEELHR---VNER------MRN--SIL-----------------RFNKDG-INSTNTYRALV-----ESIT-MEHE----------NN-RRYELA-DIQLVR-YENE--H-------------YRFNLLSKKMLHIVNDS---------KGI------------------LK-SLIRCG-----Y---------KEGNLNVNL-------------------------------------------------------YK-EVIQHLMNN-----ENLFTLIHKL-IYYKQ----------------------------------TSVNG-----------------LYYN-MGH---------VSGILDINT---------KFLK------
-------EME------VMTNISQN-----------------------QLWFVQNCGKEFKEGY-VKK-KS-------------------------ENKISGITYKLL-NSLKVNDKDGFMD-TLLNSYSY-----LS----------MPIPD-VF-IKMF------S--NNE-----AFKSVGYAFMLGVGGEKTKKED-----------GG-----NTDE-K-------------------------------

KID17666 --------------------------------MRTEIEVRADD---W---------LINAGLTGFINI-------------------------------------VGKEN-------VR-----IDGQ--AIYFTADL-------LE---DFETKYFNFFIQE--------------YKETLSWYKIVSYKETMEQFRIDKFASFDEVALGNL-------------NKYMKDVVKFY------------FKKANYIKVFPLI-----------DPSANVTKWVENLNTINLTKKQKW---
DEVKADIIEEVKQTYIQLDLII--DFCASE-----------KGLKYLGAKNLI-------YSVINK-GWSGVSFLF-------KQ--TK-------FI-DPYL---------DYKTSF-LDPVIDYL----------------------------------------------------------------------DTDLSKAK--Y---------NCFNCNQPI---------------------------KNLKLDLSFMN--DVGFDTA--R--KTAHV-----------------------W------D--FN-ND--VATCPICRLIYSCVPAGFT----------Y----V--------YG-E---
-------------GMF-----VN-DS---YGIEELHR---VNER------MRN--SIL-----------------RFNKDG-INSTNTYRALV-----ESIT-MEHE----------NN-RRYELA-DIQLVR-YENE--H-------------YRFNLLSKKMLHIVNDS---------KGI------------------LK-SLIRCG-----Y---------KEGNLNVNL-------------------------------------------------------YK-EVIQHLMNN-----ENLFTLIHKL-IYYKQ----------------------------------TSVNG-----------------LYYN-MGH---------VSGILDINT---------KFLK------
-EME------VMTNISQN-----------------------QLWFVQNCGKEFKEGY-VKK-KS-------------------------ENKISGITYKLL-NSLKVNDKDGFMD-TLLNSYSY-----LS----------MPIPD-VF-IKMF------S--NNE-----AFKSVGYAFMLGVGGEKTKKED-----------GG-----NTDE-K-------------------------------

KTA56466__13_ --------------------------------MRTEIEVRADD---W---------LINAGLTGFLNI-------------------------------------VGKEN-------VR-----IDGQ--AIYFTADL-------LE---DFETKYFNFFIQE--------------YKETLSWYKIVSYKETMEQFRIDEFASFDEVALGNL-------------NKYMKDVVKFY------------LKKANYIKVFPLI-----------DPSANVTKWLENLNTINLTKKQKW---
DEVKADIIEEVKQTYLQLDLII--DFCASE-----------KGLKYLGAKNLI-------YSVINK-GWSGVSFLF-------KQ--TK-------FI-DPYL---------DYKTSF-LDPVIDYL----------------------------------------------------------------------DTDLSKAK--Y---------NCFNCNQPI---------------------------KNLKLDLSFMN--DVGFDTA--R--KTAHV-----------------------W------D--FN-ND--VATCPICRLIYSCVPAGFT----------Y----V--------YG-E---
-------------GMF-----VN-DS---YGVEELHR---VNER------MRN--SIL-----------------RFNKDG-INSTNTYRALV-----ESIT-MEHE----------NN-RRYELA-DIQLVR-YENE--H-------------YRFNLLSKKMLHIVNDS---------KGI------------------LK-SLIRCG-----Y---------KEGNLNVNL-------------------------------------------------------YK-EVIQHLMNN-----ENLFTLIHKL-IYYKQ----------------------------------TSVNG-----------------LYYN-MGH---------VSGILDINT---------KFLK------
-------EME------VMTNISQN-----------------------QLWFVQNCGKEFKEGY-VKK-KS-------------------------ENKISGITYKLL-NSLKVNDKDGFMD-TLLNSYSY-----LS----------MPIPD-VF-IKMF------S--NNE-----AFKSVGYAFMLGVGGEKTKKED-----------GG-----NTDE-K-------------------------------

KTA38410 --------------------------------MRTEIEVRAND---W---------LINAGLTGFLNI-------------------------------------VGKEN-------VR-----INGQ--AIYFTADL-------LE---DFETKYFNFFIQE--------------YKETLSWYKIVSYKETMEQFRIDEFASFDEVALGNL-------------NKYMKDVVKFY------------LKKANYIKVFPLI-----------DPSANVTKWLENLNTINLTKKQKW---
DEVKADIIEEVKQTYIQLDLII--DFCASE-----------KGLKYLGAKNLI-------YSVINK-GWSGVSFLF-------KQ--TK-------FI-DPYL---------DYKTSF-LDPVIDYL----------------------------------------------------------------------DTDLSKAK--Y---------NCFNCNQPI---------------------------KNLKLDLSFMN--DVGFDTA--R--KTAHV-----------------------W------D--FN-ND--VATCPICRLIYSCVPAGFT----------Y----V--------YG-E---
-------------GMF-----VN-DS---YGVEELHR---VNER------MRN--SIL-----------------RFNKDG-INSTNTYRALV-----ESIT-MEHE----------NN-RRYELA-DIQLVR-YENE--H-------------YRFNLLSKKMLHIVNDS---------KGI------------------LK-SLIRCG-----Y---------KEGNLNVNL-------------------------------------------------------YK-EVIQHLMNN-----ENLFTLIHKL-IYYKQ----------------------------------TSVNG-----------------LYYN-MGH---------VSGILDINT---------KFLK------
-------EME------VMTNISQN-----------------------QLWFVQNCGKEFKEGY-VKK-KS-------------------------ENKISGITYKLL-NSLKVNDKDGFMD-TLLNSYSY-----LS----------MPIPD-VF-IKMF------S--NNE-----AFKSVGYAFMLGVGGEKTKKED-----------GG-----NTDE-K-------------------------------

AIS61706 --------------------------------MQTEIEKLASD---W---------LVNAGIVGFLNI-------------------------------------VGKEN-------VR-----IDGQ--TMYFSTDV-------LA---DFETNYFNYFIRE--------------YKETLAWHKIVSYKEKMEYYRAENFASFDEKALEDL-------------NKKYKDVVKFY------------LKKANYIKVFPLI-----------DPKANVEEWLGNLSLKSLKKQIL---
DDGKVELLEEVKETYDQLDVII--DFCASE-----------KGLKYLGAKNLI-------YSVINN-GWSGVSFLF-------RQ--TK-------FI-DPYE---------DYKTTF-LDPVFEYL----------------------------------------------------------------------ETDLSKAK--Y---------NCFICNQPI---------------------------KNLKLDLSFMN--DVGFDTA--R--KTSHV-----------------------W------D--FN-ND--VVTCPICRLVYSCVPAGFT----------Y----V--------YG-E---
-------------GMF-----VN-DS---VSIDELYD---VNEH------MRE--SIL-----------------HFNNDG-INSVNPYRALV-----ESIT-MEHE----------KQ-RRFDLA-DIQLVR-YENE--H-------------YRFNLLSKKMLHIILIDSKTILQSLIKC-------------------GYKEGNLNIL-------------------------------------------------------YK-EVIQHLMNN-----ENLFTLIHKL-IFYKQ---------------------------------TNVSGLYY-----QTGHVAGILEINT----------KFLK------
EVN------VLMSEKI-----------------------DLGKIQFFGKLFKEGY-YNK-KS-------------------------ENKIPGITYKLL-NALKVNDKDGFMD-TLLNSYSY-----LA----------KPIPR-DF-IEVF--------SNK-----E-TFKTIGYAFMLGVSGNRLKQED-----------GG-----NSDE-K-------------------------------

EFR88687 -------------------------------------------------------------------------------------------------------------------------------------------------------------------------------------------------------------------------------------------------------------------------------------------------------------------------
------------------------------------------------------------------------------------------------------Y--------------CKLIYKQTSVNG----------------------------------------LYY--------NM----GHVSGILD----INTK--------------------------------------------------------------------------------------FLKE-------------------------------------------------I--------------------
EMMATISQN-----------------------QLWFVQNCGKEFKEGY-INK-KS-------------------------ENKIAGITYKLL-NSLKVNDKDGFMD-TLLNSYSY-----LA----------KPIPS-VF-MNVF--------SND-----E-AFKTIGYAFMLGVGGELIKKND-----------GG-----NTDE-K-------------------------------

KKD44966__2_ --------------------------------MQAEIEVRAND---W---------LINSGITGFLNI-------------------------------------VGKEN-------VR-----IDGQ--SMYFSTDI-------LE---DFETKYFNYFIKE--------------YKETLAWHKIVSYKEKMEYYRAEEFASFDEKALDDL-------------NKYTKDVVKFY------------LKKPNYIKVFPLI-----------DPEANITEWLGNLTTITISKKQKF---
EEVKVEILESVKSTYNQIDAII--DFCASE-----------KGLKYLGAKNLI-------YSVINK-GWSGVSFLF-------KQ--TK-------FI-DPYE---------DYKTTF-LDPVFEYL----------------------------------------------------------------------DTDLSKAK--Y---------NCFNCNQPI---------------------------KTLKLDLSFMN--DVGFDTE--R--KTSHV-----------------------W------D--FN-ND--VATCPVCRLIYSCVPAGFT----------Y----V--------YG-E---
-------------GMF-----VN-DS---FSIDKLLD---VNVH------MRE--SIL-----------------HFNNEG-INSNNPYRALV-----ESITMEK----------EDKRRYELA-DIQLVR-YENE--H-------------YRFNILSKKMLHILNDSKTILKSLIRC-------------------GYKEGNLNVNL-------------------------------------------------------YK-EVIQHLMNN-----ENLFTLIHKL-IFYKQ---------------------------------TNANGLYY-----NMGHVAGILEINT----------KFLK------
EID------VMTNISQK-----------------------QLWFVQSCGTEFKEGYYGKKS--------------------------ENKIAGITYKLL-NSLKVNDKDGFMD-TLLNSYSY-----LA----------KPIPS-VF-MKVF--------SND-----E-AFKTIGYAFMLGVGGEKTTKEKDG-----------GN-----TDEK-------------------------------

WP_036076884 --------------------------------MREIECRASD---W---------LFNSGMVGFINI-------------------------------------VGREN---------IR-----LEGR--SIFFDEGL-------FD---AFEDKYFAYFIKE-------------YKEMLSWYKIVSFKEEIKRYQKDDFLSFDEDALERL-------------NDYIANVAKRY------------LKSASYISVYPLI-----------DVTAAPLSWEKGLTKVGVLKKKET--
FDEKRTDIIQEVKQVYLKLEQII--SFCDSE-----------KGKKYLAAKNVI-------YTVINK-GWNGVSFLN--------PQ--TK--------EK-DVYE---------DYRTKF-IAPAQEYF------------------------------------------------------------EIDKVDSE--Y---------KCSACNKPI---------------------------KKEKYNFSFLN--YAGFDIA--R--KSSHV-----------------------W------D--FY-ND--IEICAVCKLIYSCVPAGFT-----Y---------AFGN--
--------GIF-----ID-DS---FSIDTLYE---VNEK------IRF--AIL-----------------HDPAQT-LAQASPYRALV-----EATT-NQVE----------GN-RHYELA-DIQVVR-YENE----------------T-YRFNILSRKALNIFRESKNALKAIQRC-------------------GFKEGNLSVNF-------------------------------------------------------YQ-ETVRRLMNN-----ENLFTFIHKT-MYLKV---------------------------------TNVSNTYY-----HIGHIGSLLEINT----------NFLK------
-EIE------AMTEISKG-----------------------RIWHINKCGNEFKKAY-P--------------------------------ENKIAGFNYKVL-NALKTNNRDAFMD-IILNSYSY-----LG----------KVIPD-FF-LESF--------TSD-----E-TFKTVGYAFMVGVNGAIEQDDKDKD-----------GG-----NNDE-K-------------------------------

KGL39779 --------------------------------MMREIECRTGD---W---------LFNAGLVGFLNI-------------------------------------VGKEN-------VR-----IEGQ--SLFFNAAL-------LD---GFEESYFAYFIKA-------------YEKTLSWYKIISYKEEMERHREADFSDFDELSLDRL-------------NDYIANVAKYY------------LKSSSYVSAYPLI-----------DATKDPVEWGKNLAKVGQLKKKET-
-FEEKRTEIIQNVKQVYVQLDEII--DFCQKG-----------KGKKYLAAKNVI-------YTVIKN-GWNGISFLN--------PQ--TK--------EK-DCYI---------DYRTNF-VDPVLEHL-----------------------------------------------------------ETDLEKAK--F---------KCTNCENPI---------------------------KNMKLDLSFMN--ETGFDTA--R--KNSHV-----------------------W------D--FN-ND--IAVCPMCKLIYSCVPAGFT-----Y---------YG-N--
-------------GIF-----IN-DS---INIDTLHA---VNKK------VHN--EIL-----------------HHSTDN-LTSTSPYRALI-----EAIT-NQTE----------DK-RHYELA-DIQVVR-YENE----------------T-YRFNILSRKALKIFRESKEALKTIQKC-------------------GLKEGNLNINF-------------------------------------------------------YQ-ETVRRLMNN-----ENLFTFIHKT-MYLKV---------------------------------TNVSNTYY-----HMGHVGALLSINT----------NFLK------
-EIE------AMTDISKG-----------------------QIWHINKCGTEFKKAY-P--------------------------------ENKLSGFNYKIL-NALKTNNRDAFMD-IILNSYSY-----LG----------KAIPD-FF-LKAF--------TND-----E-TFKTVGYAFMVGVNGPLENNEN-----------GG-----NDDE-E-------------------------------

WP_036065862__2_ --------------------------------MSEIECRAND---W---------LFNAGLVGFINI-------------------------------------VGKKN-------VR-----MEGQ--SVFFEESL-------LD---GFEEKYFDYFIRE-------------YEKTLSWYKIVSFKVEMEQHRKTDFADFDEKALDRL-------------NEYIANVAKYY------------LKSASYASVYPLI----------
DASVDPAQWGKNLAKVGQLKKKET--FAEKRAEIIQEVKQVYEKLDQII--AFCESD-----------KGKKYLAAKNVI-------YTVIKN-GWNGVSFLN--------PQ--TK--------EK-DCYV---------DYRDNF-VAPVVEYL---------------------------------------------------------ETDLEKAK--F---------HCFTCNEPI---------------------------KNLKLDLSFMN--ETGFDTA--R--KTSHV--------------------W------D--FN-ND--
IAVCPLCKMIYSCVPAGFT-----Y---------YG-N-------------------GIF-----IN-DS---VSAEALHE---VNEK------IRN--SIL-----------------HNHEDS-LTQTSPYRALI-----EAIT-NQVE----------DK-RHYELA-DIQVVR-YENE-----------K-YRFNILSRKALNIFHDSKNALKTIQKC-------------------GFKEGTLNVNF-------------------------------------------------------YQ-ETVRRLMNN-----ENLFTFIHKA-MYLKV--------------
--TNVSNTYY-----HMGHVGSLLEINT----------NFLK------------EIE------AMTEISKG-----------------------RIWHINKCGNEFKKAY-P--------------------------------EHKRGGFNYKML-NALKTNNRNDAFMD-IILNSYSY-----LG----------KTIPD-FF-LESF--------TSD-----E-TFKTVGYAFMVGVNGAIEQDDKD-----------GG-----NDNE-K-------------------------------

AQY50207__2_ --------------------------------MREIECRASD---W---------LFNAGLVGFINI-------------------------------------VGKEN-------VR-----MEGQ--SVFFDADL-------LD---SFEDKYFAYFIEE-------------YEKTLSWYKIVSFKKEMDRYRKDDFASFDEKALEEL-------------NNYIANVAKYY------------LKSASYVSVYPLI-----------DATADPVEWGKNLAKVGALKKKET-
-FDEKRADIIQEVKQVYLKLDQII--AFCNSE-----------KGERYLAAKNVI-------YTVIKN-AWNGVSFLN--------PQ--TK--------EK-DTYV---------DYRTNF-IDPVKEHL-----------------------------------------------------------ETNLEKAK--F---------KCTNCEKPI---------------------------KNLKLDLSFMN--ETGFDTA--R--KTSHV--------------------W------D--FN-ND--IAVCAVCKFIYSCVPAGFT-----Y---------
VHNN-------------------GIF-----IN-DS---VSIDTLHA---VNEK------IRS--AIL-----------------HDPAQT-LTHTSPYRALV-----EAIT-NQVE----------DK-RHYELA-DIQVVR-YENE----------------T-YRFNILSRKALNIFRESKNALTALQRC-------------------GFKEGNVNVNF-------------------------------------------------------YQ-ETVRRLMNN-----ENLFTFIHKA-MYLKV-------------------------------TNASNTYY-----HMGHIGSLLEINT---------
NFLK------EIE------AMTEISKG-----------------------RIWHINKCGNEFKKAY-------P---------DHKRSGFNYKVL-NALKTNNRNDAFMD-IILNSYSY-----LG----------KTVPD-FF-LESF--------TSD-----E-TFKTVGYAFMVGVNGAIEQEDKDKD-----------GG-----NDNE-K-------------------------------

ADY55689 --------------------------------MEEEILRLRLGD---W---------QWNGAVIGFIRI-------------------------------------VGREN---------VL-----FENE--SIKFSSKL-------LE---NFEEKYFAYFIKA-------------YEKTLSWYKIVSFKEWLEAQEDDEFGTFVLKDLNAL-----------NEQIKTVKSLI--------------KSNSYKKAAYELM-----------ALGEDILVWEKELVSIKEPKSEQA--
FAADKPEIAIKVRETFSTLRKII--SYCESP------------GKRYIRAKNII-------YTVIKN-AWNGVSFLN--------PQ--TK--------EK-DVYS---------DYKAHF-VDGAIAYL----------------------------------------------------------KQNKDKFK--Y---------HCFICDEPI---------------------------KSMSLDLSFMN--ATGFDVA--R--KSSHV-----------------------W------D--FQ-ND--IAVCPLCKLIYSCLPAGFT-----Y---------VD-R--
---------------GIY-----ININT-----SVEDAYE---VNYKIHHEVLRK-------------------EGNLRSIYPVL-----IGAL-HEQE------------NAGASYELA-DIQVVR-YERE----------------A-YRFNILSRKMLQLIFDS--KDRLDSL-----------------------IN-TSFAENKLTVRI---------------------------------------------------------------YD-EAISRIFNN-----QNLFLLIHRM-LYDLL--------------------------------SDPSKCRF-----NGAHITHLLRINQ-----------RIYQ------
TLG--------GMKMGCEDKDNA-------------------LVREARTAGEQLKRYY-YGK-NA--------------------------QHKLPGICYRLL-NALKTSNQEMFLD-VTLNCYLY-----VG----------FPVQR-VH-TDSL--------GQE-----K-DFSTVGYAFVAALIDN-SKSEDKGSSPVGEM-----EEEK-K-------------------------------

```
KJS22467                ---------------------------MSSDTIRLTLGD---W--------QWNAAIIGFINI-------------------------------VGQENV-------RF----VGED--AVEFSVKI-----LD---GFEDRYFSCMIQA-----------YEKTLSWYKIVSFKDKLYAYEENSFEGFDLKALQGL----------NTYIKDVKRYI-----------KSNSYKAAYELI---------GLDVDMLSLEKQLLTSKEPKSQQQ--
FDLNKNGIIAEVKESFTILRQII--DYCDCA--------QSKRYIGAKNVI------YTVIKN-AWNGVSFLN-------AQ--SK------EK-DVYV--------DYKTYF-VDTVTDYR--------------------------------------------------------------------MSNESKYK--Y-------NCVVCDAPI---------------------KNMSSDLSFLN--ATGFDVA--R--KSSHV---------------W-----D--FQ-ND--IAICPVCKLIYSCLSAGMT-----Y----V--------YD-
R-----------------GIY----VNANI-----RLKDALN---INFR-----IRQ--DFLRSN----------------ERGMRSIYHAL----VGAL-HEKE-------NNTAKYELA-DVQVVR-YENE-------------T-YRFNILSRKMLEIIVAS---EQELNSL-------------IK-TGFSENGVSIQI------------------------------------------YD-EVIARIFNN----RNLFTLIHRM-LYYKL------------------------SDSSKCYF----NGAHVNHLLKINQ-------RIYL-------
--SLG-----GMKMEKTKEQEQENSD----------------LVRYARAAGYHLRGRY-VNK-GS-------------------------DNKLTGICYRLL-NALKTSNTDMFMD-VTLNCYLY----VN----------CQVPK-II-TDVL-------GKE----K-SFSTMGYAFISGLIDG-----NDTNGSLEKKA----KGEE-Q------------------------

SFG73304                ---------------------------MSNNNIQLTLGD---W--------QWNAAVVGFINI--------------------------------VGKEN-------VN----IEAD--TVEFSQEV-----LD---EFENKYFAYFIKT-----------YERTLSWYKIVNYQDNLYSYEENNFEEFDLKALEGL----------NTYIKDVKRYI-----------KSNSYKAAYELI---------KSEVNLLSLEKQLTTIKEPKNQQK--
FDEDKPKIANESKQRFHLLWQII--DYCASP--------EGKRYIGAKNVI------YTIINN-AWNGVSFLN-------PQ--TK------EK-DVYA--------DYKNYF-VDSAVAYL--------------------------------------------------------------------QSEKSKFK--Y-------NCFVCNAPI---------------------KDMSNDLSFMN--ATGFDVA--R--KASHV---------------W-----N--FQ-ND--TAICPLCKLIYSCLPAGIT-----Y----T--------YD-
R-----------------GIY----INQNI-----SLKDAIR---INSK-----IKH--KILSSQ----------------ESGMRSIYHAL----VGAL-HEQE-------NDSAKYELA-DVQVVR-FENE-------------S-YRFNILAKPMLQIIVNSKKELDSLIRV-----------------NFIENGGNINV-------------------------------------------YD-EVIGRIFNN----QNLFTLIHRM-LYGKL------------------------SNPSKCYF----YGSHVRNALIINQ-------QICN--------
-RLG-----GMNMENAKGGVQVNNE----------------LVKNARTAGYLLRKKY-VDK-DA-------------------------NHKLAGICYRLL-NALKTSNENMFMD-VALNCYLY----VN----------SQVPK-VI-TDVL-------GSE----K-DFNTMGYAFVSGLIEGQEGSGNK-------DK----KAEG-E------------------------

ACV61808                ---------------------------MKSDKIRLTLGD---W--------QWNASVIGFINI--------------------------------VGEENV-------CH----IEAD--TVEFWPEA-----LD---GFENKYFAYFINT-----------YEKTLSWYKIVNFQDRLNTYEENNFEEFDLKALEFL----------NVYIKDVKRYI-----------KSNSYKAAYELI---------DSEVELLSFEKQLITIKEPKNQQR--
FDVDKSEIVNEVKRRFLLLQQII--NYCASS--------EGRRYIGAKNAI------YNIINN-AWNGVSFLN-------SQ--TK------EK-NIYS--------DYKSYF-VDTATGYL--------------------------------------------------------------------QSEKGKYK--Y-------NCFVCDAPI---------------------KDMSNDLSFLN--ATGFDVA--R--KSSHV---------------W-----N--FQ-ND--ITICPLCKLIYSCLPAGII-----Y----T--------YN-R--
--------------GIY----INQTI-----KLKDAVR---INNK-----IKS--RILRSQ----------------DGSMRPIYHAL----VSVL-HEQE-------NDSAKYELA-DVQVVR-YENE-------------S-YRFNILAKPMLQIIVNS---KKELDSL-------------IR-VNFIENGRNINV-------------------------------------------YD-EVIGRIFNN----QNLFTLIHKM-LYGKL------------------------SDASKCYF----NGSHVRNVLIINQ-------LICS---------
RLG-----GMKMDNTY-------------------------LVKKSREAGESLRNKY-FNK-DS-------------------------NHKLAGICYRLL-NALKTTNENMFMD-VTLNCYLY----VN----------SQVPK-VI-TDAL-------GRG----K-DFSTMGYAFVSGLID-----GQDGSGNKEKKA----EGE---------------------------

RBP38226                ---------------------------MEEKIKIYLSD---W--------QFNAGIVGLYNI-------------------------------LEYAKDE-------VA----INDQ--YIEIELEM-----LK---NFEEKYFNYFIEK-----------YKETTPWYRIVSYKPIMETHENNNFENFNKKSLDYL----------NDYIKMVKDYL-----------KRPNYKKVYSFV---------ENDKDIIFLEKELKTIYIKKSDKV-------
EDKIDEVKNRFKTIKEII--DYCNSE--------DGKKYMAAKGVI------YNIINN-GWNGVCFLN-------AQ--TK------EH-DVYK--------DYKEYF-VDETINYL--------------------------------------------------------------------KSDKEKFK--Y-------NCFVCDRKI---------------------KNMNNDLSFLN--ETGFDVA--R--KTSHV---------------W-----N--FT-ND--VAICDLCKFVYSCLPAGFV-----Y----A--------YD-K--------
---------GIF----INSNT-----SVEDLLK---INRK-----LRD--DIL-------------------NKDDSDYRSLTYRALVKAI-NEQH-------SSKIQYELQ-DIQIVR-YENE-------------R-YRFNLLSKEVLKVIRES---EKELKSL-------------MN-TGYRELRTNFSI------------------------------------------YD-ITIDRLLNN----YNMFTLIHKL-LIYKI------------------------TNSQSITTYY----HMGHVMNLIIINV-------NYLK---------GVE----
-GMDKVKENKE-----------------------LVRKARSYGYYLRQEY-LNK-DRDAD----------------------KNKIRGVSYRLL-NALKTRNTEMFMH-NVITSYMY----VG----------ETIPK-KLTIALE-------NEE------ILGIIGYAFVTGLNGGEYKDKNQN------GG----ENNE-N------------------------

SJZ74431                ---------------------------MGEKIKIYLSD---W--------QFNAGIVGLYNI-------------------------------LEYAKDE-------VA----INDQ--YIEIELEM-----LK---NFEKKYFNYFIEK-----------YKETTPWYRIVSYKPIMETHENNNFENFDKKSLDYL----------NDYIKMVKDYL-----------KRPNYKKVYSFV---------ENDKDIISLEKELKTIYIKKSDKV-------
EDKIDEVKNRFKTIKEII--DYCNSE--------DGKKYMAAKGVI------YNIINN-GWNGVCFLN-------AL--TK------EH-DVYK--------DYKEYF-VDETINYL--------------------------------------------------------------------KSDKEKFK--Y-------NCFVCDRKM---------------------NNMNNDLSFLN--KTGFNVS--N--KPSHV---------------W-----Y--FA-ND--VAICDLCKFVYSCLPAGFV-----Y----A--------YD-K--------
---------GIF----INSNT-----SVEDLLK---INRK-----LRD--DIL-------------------NKDDSDYRSLTYRTLVKAI-NEQH-------SSKIQYELQ-DIQIVR-YENE-------------R-YRFNLLSKEVLKVIRES---EKELKSL-------------MN-TGYRELRTNFSI------------------------------------------YD-ITIDRLLNN----YNMFTLIYKL-LIYKI------------------------TNSQSITTYY----HMGHVMNLIIINV-------NYLK---------GVE---
--DMDKVKENKE-----------------------LVRKARSYGYYLRQEY-LNK-DRDAD----------------------KNKIRGVSYRLL-NALKTRNAEMFMH-NVITSYMY----VG----------ETIPK-KLTIALE-------NEE------ILGIIGYAFVTGLNGGEYKDKNQN------GG----ENNE-N------------------------

KLE16527                ---------------------------MENLIRIDIGD---W--------MINAGIVGICNI-------------------------------LENAGDK-------VT----KKAN--YITIDRSY-----LD---NFEEKYFKYFIDT-----------YRGTLSWNKIISYKGILDNFENSDFKNFTEESLENL----------NVYIKDTVKYY----------LKSNSYIAAYELI---------DNNIDILALERKLTTINLKKKENL-------
EDKLPQIKETIKVIREIIDYFSMEKS----------KKYLAGKNVI------YTIIKN-AWNGISFLN-------AQ--TK------EK-DMYV--------DYKNYF-VNPAIDEL--------------------------------------------------------------------EANKEKYK--Y-------HCFICNNEM---------------------KDLNNDLSFLN--GIGFDVN--R--KPSHV---------------W-----N--FN-ND--VAICNFCKLVYSCIPAGIS----------Y--------GAGS-------------
---GIF----INANI-----NAENLIK---VNEN-----IKE--SIL-------------------RNSDGIKSLSYKSLISAI-QKQE-------HDSFKYELA-DIQVIR-YEDE-------------R-YKFNILSRNSLKIIKNSKEQLDRLIKS-----------------GYIENKKFYSI------------------------------------------YE-LVIERLLDN----SNLITLIHRL-VLYKL------------------------SNPDDCRY----SVKNLIDMLRINY-------KYMK---------EIG-----
YMQDIKNENDKD--------------------IVDLAKNCGYYLRLAY-KAK-GS--------------------------EEKLSGIAYRLL-NCLKTNNASMFMD-TILNCYLY----TK----------KTVPM-EL-LGVL-------KDS----D-VFKHIGYAFVASLIGNPENDNKEN------GG----KKDD-K------------------------

KJF26764                ---------------------------MGEKIRLEMND---W--------LYNAGLVGLYNV-------------------------------LKHSGDK-------VD----YAEQ--YVELDVSL-----LE---NFEEKYFRYFIDK-----------YQAYLSYFKIISYERTIKYHEDNDFETFEEEDLESL----------NNYIKNVVKYY----------LSSNSYRAAYEII---------
NDEVDMLELAKELQVIKIKKKESIKDKVGEIKNIFQQLKVMINCCKKDN-----------------YRKYLAAKNVI------YTIIKH-SWDGVSFLN-------PQ--TK------EK-DIYL--------DYQNYF-IEPVNKYM--------------------------------------------------------------------TKKATSEK--F-------NCFTCGRSI---------------------KDLKNDLSFLN--NIGFDVS--R--KSSHT---------------W-----N--FS-ND--
VAVCPICKLVLSCTPAGIT-----Y----V--------YD-K-----------------GIY----INDNN-----SFQDAVN---INNK-----VKS--EIL-------------KEFR-TDKLLTYRALV----NSIQ-EQFQ-------DK-MKYELA-DIQVVR-YEEG-------------Q-YKFNILSKQALYIIRKS---QGDLNNI-------------IH-SGFKEINTYFNI------------------------------------------YE-LVINRILNN----HNLFTLIQKL-LVYKL------------------------
SNPKDCRF----NALQLISILNINF-------RCLE---------GMG-----YMKSSDKD------------------------IIKNANISGYYLREQY-KEK-GA-------------------------QDKLNGISYRLL-NALKTNNKDMFMD-TLLNCYLY----SQ----------KTVPS-IF-LEAL-------KDD----E-KYKTIGYAFVTGLIEG----------KESMKK----NGGD-E------------------------

ABR48349__2_            ---------------------------MAQRIRLEMND---W--------LYNAGLVGLYNI-------------------------------LVHSGEK-------VK----MNEQ--GIEMDQSQ-----LE---NFEERYFTYFIEK-----------YEVYLSWYKIISYKDVLTYHQKQNFNSFGEEELDSL----------NQYIKDVKHYL--------------------SSNSYKAAYELIGSQVTLLELAKKLQTIKLKKKETI---
----GEKLIEIKETVNKLNEIIGHCMEEDY----------KKYLAGKNVI------YTIIKN-GWNGVSILN-------PQ--TK------EK-DIYM--------DYKNYF-VKPVEDYM--------------------------------------------------------------------GDDKSTKK--Y-------NCFTCDREM---------------------KDLKNDLSFLN--QVGFDVS--R--KASHT---------------W-----D--FV-ND--IAICPICKLVFSCVPAGMT----------Y--------VYSK-----
------------GIY----INDNN-----SFQRAVN---VNQR-----IKS-----------------EILKEHE-DNRLLTYRALV----KSIQ-EQIH-------EKIKYELA-DIQVIR-YEEE-------------K-YRFNILTRQMIKIIQQSKHSLGYIHNS-----------------GFKEVNTYFNI------------------------------------------YE-LVIERVLNN----QNLFTLIHKL-LSYKL------------------------SSPKNCRF----TTSQVISVLEINF-------RFLE---------GMG---
--YMQNADQD--------------------------MIKKANVSGYYLREQYKVKGA--------------------------KDKLNGVSYRLL-NALKTNKKDMFMD-TLLNCYLY----AQ----------KTVPS-IF-LDAL-------EDE----E-KYKTIGYAFVAGLIEGKDHQ----------NK----EGDQ------------------------

AFS77693                ---------------------------MGSTIKLSMND---W--------LFNAGLVGIYNI-------------------------------LKESNFD-------VK----ISGQ--DIEFDYSY-----LE---NFEESYFYYFINK-----------YEKSLSWYKIVSYEERIEYHEESNFENFNEKSLEDL----------NNYIKNVAKYY----------IKSASYKSTYGLF---------NNEIDILALEKELSTITLRKKESI-------
EDKIVEVKDRFEKLKVII--NFCKSE--------KGKKYLASKNVI------YNIIKN-SWNGISFLN-------PQ--TK------EK-DVYI--------DYKNYF-VVPTKEYL--------------------------------------------------------------------DADKSKYK--Y-------DCFICDSKI---------------------KDLKDDLSFLN--MIGFDVN--R--KSSHV---------------W-----N--FV-ND--IAICSLCKLIYSCVPAGIT-----Y----V--------YD-K----------------
-GIY----INDNS-----SMKNAID---INQK-----IKS--EVL-------------------KTDD-NTRTLTYKALV----TSI--QEQS-------NDKIKYELA-DIQLIR-YEDS-------------K-YKFNILSKKSLKVMRDS---RDNLDSL-------------IK-CGFKEINTYFNI------------------------------------------YE-LVIDSLLNS----QNLFTLIHRL-FMQKL------------------------SSPKDSYF----NTFHLISMLKINI-------RFLK---------GVG-----
YMKDIDKDA-----------------------IVREGNASGYYLRKKY-DSK-GS--------------------------VDKLNGISYRLL-NALKTNNVDSFMD-TLLNCYLY----AK----------ISVPP-IM-VDSL-------KSE----E-EFKTIGYAFVAGLIEGKDEIKEN-------GG----KSDE-N------------------------

WP_074349147            ---------------------------MGEKIRLYMED---W--------LYNSGLVGFYNI-------------------------------LKHAENE-------VV----INQN--YLEFDSDN-----LV---DFEKKYFSYLMYK-----------YKDILSLNKITSFEDFILYYEESNFENFDEKSLEIT----------NKYISDVAKKQ----------IKSNSYKSAYELI---------KSTVDILGLEKSLKTINLKKKQKI-------
EDILPEVKDKFKLLMQIIGYMKLEDA----------QKYIGAKNAM------YTVIKN-GWNGVCFLN-------PQ--TK------EK-DMYI--------DFKDYF-VDPTIEYL--------------------------------------------------------------------KIDKSRFR--F-------SCFSCNRSM---------------------KDLTNDLSFLN--STGFDVS--R--KSSHV---------------W-----D--FQ-ND--IAVCPICKLIYSCVPAGIS-----Y----L--------YD-K----------
------GIY----INDNS-----SMKNAID---INNK-----IYM-----------------EIYKQSK-EDKKLTYKALV----KSIN-EEYN-------DK-IKYELA-DIQLVR-YEDE-------------K-YRFNILSKDSLRVIKGS---EDDLNKL-------------IN-CGFKEINTYFNV------------------------------------------YE-LVVDRLLNS----QNMFTLVQKM-LHYKL------------------------SQPKDSHY----NSYHVIRILRINT-------RFLK---------GVG----
-CMKEKEID-------------------------IVDLGNKAGYFLRKDY-GDS-----------------------------VDKLNGIAYRLL-NSLKTNNKDSFMD-TLLNCYLY----VK----------SPVPK-VF-LEVF-------SSD----E-DFKTIGYAFVAGLIEGKKENINDNG-----NG----GNDN-E------------------------

SHE48919                ---------------------------MGEKIRLYMED---W--------LYNSGLVGFYNI-------------------------------LEHAEDN-------VK----IDKN--YIEFELEN-----LI---RFEEKYFKYFTDK-----------YKDIFSLNKILSFEEFIDKQEENNFEEFDGKKLESM----------NKYISDVVKKQ----------IKSNSYKSAYELI---------DSPVDVLELEKSLKTIKLKKKQEI-------
SEILPEIKDRFTILKEIIEYMKLEKS----------QKYIGAKNAM------YTIIKN-GWNGVCFLN-------PQ--TK------EK-DMYI--------DYKNYF-IEPAIEYL--------------------------------------------------------------------NIDKSKFK--Y-------NCFSCDKSM---------------------KDFSNDLSFLN--STGFDVS--R--KSSHV---------------W-----E--FQ-ND--IAVCPICKLVYSCVPAGIS-----Y----L--------YD-K-------------
--GIY----INDNS-----SMRNAID---INNK-----IYM-----------------EIYKQNR-DDKKLTYKALV----ESIN-EEYN-------DK-IKYELA-DIQLIR-YEDE-------------K-YRFNILSRKSLEIIKAS---EDDLNKL-------------IN-CGFKEINTYFNV------------------------------------------YE-LVIDRLLNS----QNMFTLVQKM-LHYKL------------------------SKPKDSHY----NSFHVIRILRINT-------RFLK---------GVG-----
CMKGVYKD-------------------------IVRDGNDEGKKLREKYRSKGA---------------------------IDKLSGISYRLL-NSLKTNNVDSFMD-TLLNCYLY----VK----------SSVPE-IF-LDAL-------KNE----E-RFKTIGYAFVAGLIEGKKENNNDNEN----GG----KDNE------------------------

TCU77296                ---------------------------MGEKIRLYMED---W--------LYNSGLVGFYNI-------------------------------LEHAEDN-------VK----IDKN--YIEFELEN-----LI---RFEEKYFKYFTDK-----------YKDIFSLNKILSFEEFIDKQEENNFEEFDGKKLESM----------NKYISDVVKKQ----------IKSNSYKSAYELI---------DSPVDVLELEKSLKTIKLKKKQEI-------
SEILPEIKDRFTILKEIIEYMKLEKS----------QKYIGAKNAM------YTIIKN-GWNGVCFLN-------PQ--TK------EK-DMYI--------DYKNYF-IEPAIEYL--------------------------------------------------------------------NIDKSKFK--Y-------NCFSCDKSM---------------------KDFSNDLSFLN--STGFDVS--R--KSSHV---------------W-----E--FQ-ND--IAVCPICKLVYSCVPAGIS-----Y----L--------YD-K-------------
--GIY----INDNS-----SMRNAID---INNK-----IYM-----------------EIYKQNR-EDKKLTYKALV----ESIN-EEYN-------DK-IKYELA-DIQLIR-YEDE-------------K-YRFNILSRKSLEIIKAS---EDDLNKL-------------IN-CGFKEINTYFNV------------------------------------------YE-LVIDRLLNS----QNMFTLVQKM-LHYKL------------------------SKPKDSHY----NSFHVIRILRINT-------RFLK---------GVG-----
CMKGVYKD-------------------------IVRDGNDEGKKLREKYRSKGA---------------------------IDKLSGISYRLL-NSLKTNNVDSFMD-TLLNCYLY----VK----------SSVPE-IF-LDAL-------KNE----E-RFKTIGYAFVAGLIEEKKENNNDNEN----GG----KDNE------------------------

SFS35353                ---------------------------MKHSYNKIHYD---WLYRPTGDPFADVGGFALKEF------------------------------------------------------SKRYPDLD-----IL---EIIMKATDIYVDR----------------------------------------------------------------------WGGKINPFFLNSKITQ------------------------------------
PAFSL-------ER--KK--------------------EETKKYF-LGLITEEV------------------------------------------------------PGQLG------------ICRITGEK--------------------TKLYPAGRDNTV--LSGSG----T--LVNFH----------------H-----T--FQ-EG--IMLSKEIIIRYHFLPLGCE-----------L--------LL-G--K---------------VAV----IHSNN----SEITELFA---SGCC-----SRNLSAIGSNH-------SDGILKSQA-
RSAGTALFRFA----GQVV-TNTK---------KNIDEESSQYTISLYH-FTNF---------GASPD-VQIYVLPFEIFTFYRII-QSNDTLKKQ-------------WN--EFTSRY----Y--GNSEYKKAVYNEVERVYIYNDKNSSIKIQGDDFKF-------WRNTIYERLING----QSIVPHLLRY-IREHEL---------------------------------------------SWDLIKLYE--------LNIR----------KMK-----------KE---------------------------TLSKIEQMAEYILASN-
N-------------------------------EHGIKKAIKKLD-GVKSSFDLRRFVLRDIVAKYYN----------------EENNEA-IVTIEDYAEYLF--PDT----N-SWQETRDVLLISIYQK--MHERKMHVETELSE-----DDEL-N----EEN--------------------

AEE48384                ------------------------MATIDYKQDYLTQRTGD-----------PFADTGAHVIQLM-------------------------------------------------------QEKSPEKT-----IL---QLIEEATNIYVRN---------------------------------------------------------------------------WGGGINAFFLNSKITQ-----------------------------------
PAFLG-------ER--KK----------------------EESMAY-FQRLL--------------------------------------------------------SNE--EPYEEG-----------YCRILGQK----------------------TKLFSTGRDNHI--LAGSG----T--FINFH----------------H-----G--YQ-EG--LKVAKEVIIRMFFVPLGAI-----Q----L---------GK-N----------------IAV----IGSNL----EKVEALFV---EDIF------KANMVEVSHKQ-------AKGVLKSDF-
TNPANALFEFA----SKSL-NTSK-----------VVIEEDDQVEIHL-YHFT--NF-----GASPE-VNLYCFSARLFAFYRKA--MKINYRSI-------------WQ--SFVNDF----Y--RNSKNSGAVYNNATNQYELEVKKEKKNFEYQEYRVW----FN-VIYHKLLRN----QSILKNFLYW-SSNKY-------------------------KQQA-VFN--------IWPVVELYQ-------TQLH---------NMK-----------KT----------------------------
TLDKIRQIADYLIED----------------------------------EDKIKRRLNRLRLEVSNVYQLKRFLIQTIGDHYQE----------------KQPARP-LITLEEFVDDLF--PEG-----T-YGLEIKDLLMIALYER--LCEKNIPVELEGED-----VLED-D---RPNFN------------------

SEA90530                ---------------------------MTEEKNWLTSYTGD-----------PFADAGGFVIKQL------WQ-------------------------------------------------DEQFKDKS-----IL---ELIKYVTKVYVNN----------------------------------------------------------------------WGAKLHSFFLNSKITQ--------------------------------------------------------------AA-
FVGDRKIEEAI---------------------------------KYYTAL-LNDDLPYK-------------------------------------------------------EG-------------YCRISGEK---------------------TKLYLAGRDNHV--LSGSG----T--YINFH----------------P-----N--FE-PG--LYLSKEMLIRMFFLPLGVV-----Q----L---------SD-K----------------IAL----LACNV----PDIAEYFV---KENL-----EKNLLEIRSGI------SQGVLKSPF-
NFPANAIFSFI----DHYL-REIKQLTDYREEEIEEKTNA-SITLFH-FTNF---------GASPD-ASIYSLDSKVFQFYAAC---QFKFPKV--------------WN--QFLRSN----Y-----------RNYKLKGSIFDVDTETWKSKDEQLDYDAYKTWRNEVLDNLMNG----ESILPNIVKWLVDHPF-----------------------------------------------PSEITDYYL-------NYIR-----------NMD------------KK---------------------------TIQIVRKIADFISNGS-
--------------------------------EDEIAKDIKRLR-GHRSAYELRTFTVGLLDKNYKA-----------------EAQDP-LFTLNEYADYLF--PDG----G-FWRETKNLLLVGIYEN--LHKQGKKVDTPAQD----DSEE-I----ALDTENNN-------------

TDW82906                ---------------------------MVNKKYLIEYTGD-----------PFADVGGYVIETL---------------------------------------------------------WE----------DPRLKDKD-----IL---GLIEYVANIYISQ-------------------------------------------------------------------------WNANLHAFFLNSKITQ----------------------------
PAFDS-----------KR------KL--------------EETLTY-YRSIFDET---------------------------------------------------APAQVG------------YCRISGRKT----------------------NLFAAGRDNHI--LSGSST------FINFH----------------H-----G--FE-NG--LYLSKEIIIRFFFVPLGVT-----------Q--------LGDK-----------------IGL----ISSNN-----NVVAKYY---VQQL-----
LTGPDGHLTALGRGLASGVARSDYGIVANALFAFADSCIKNIRTAIFAQLD-------ENVVEVRDT-TLSLYH-FTNF---------GAKPE-VVIYQLPATVFGFYAEC--HTPNYASA-------------WR-KFIAAHYRNSKYKDAKYDPDSDTWVSAKEELAYNSYST-----------------WRNVILERLLNG----ETLIPYFKEW-IRKHK-------------------------------------------------------------FTFQIVENYL-------FYVK----------NMD-------------
KR---------------------------TLQKITDVAGFIVSRD-----------------------------------EDSIKKAILRLN-RAKYVQEVRQYLLKLIDDNYKA-----------------NPTTP-LIRLNDVEYLF---PET-----I-SWREIRDLLLIAIYEKLHDNHMIVPTEIEEQEEVETPEND------------------------------

PTN09608                ---------------------------MKQNYKTVDYE---WLTKATGDPFADAGGYAIEYL-----------------------------------------------------------SEKFPEKD-----IL---ELIDYITTIYVKK------------------------------------------------------------------------WGGKLDAFFLNSKITN----------------------------------------------------------------NA-
FKGNKKIEETK--------------------------------KYFKSL-VDESESIE-------------------------------------------------------FG-------------YCRVTGRE-----------------------AKLFFAGRNNSV--MTGSG----K--LINFH----------------H-----F--FQ-RG--ISLSKEALIRIHFVPFACQ-----Q----L---------QG-K----------------ISL----MQSSN-----RTLNNLFVKLTVEKN-----LHE-IGVGLSE---------GIAKSDY-
KKPANAIFDFV----DSAL-SQLA--------EFREGNILP-SLTLYH-FTNF--------GATPE-IQLYQLPAKVFLFYRTC--LQPRFKED-------------WQ--KFVRSH----Y------YDGKHKGAKYNLQSGKFELVKSKETEVIDFDDYKQWFN-RILNKLLID----ENIRPEILRW---------------------------------SRKH-PF----------NFEIVSIYQ-------QNII----------GMK-----------KE---------------------------TILKIKELAAFLVREE-D-
-------------------------------ADKIKKRIKALD-GAKNASALRRFILKDVVAANYV----------------A-MNENP-IVTVDDYVNYLF--PDG-----S-YWAEIRDILLIAIYQE--LHERDLISEELKVE-----LDSE-V----EEEVIND-------------------

EAY25594                ---------------------MGFQSDLVSFLTLTKMKKTIENIAYK---WLTKPTGDPFVDVGGVVMDYF-----------------------------------------------ARLHPGKD-----VL---GLIESMTDIYVNN----------------------------------------------------------------------------WSGKLNAFFLNSTITQ--------------------------------------------
---------PAFKGQKKIDE------------TL----------------------KYFKEL-IQDKRAFQ--------------------------------------------------EG-----------------YCRITGRKT-----------------------KLFYAGRDNHI--LSGSGT-------FINFH----------------S------A--FE-SG--LTLSKEVLIRIFFVPFGLL------------Q----------VSDK-----------------IAL----IQSND-----LEVSKYFV----LKNC-----QENIQGIATGA-------SEGVLKSSL-
KNPASALFGFIDDCVSDLK-TVVD-----------DRSKLK-DIAFNI-YHFT-------NFGASPE-VQLYHLHSTAFRFYLTC--LNPRYRDD-------------WQ-GFVSAHYHNSKFKNIKFNRTSAVWEDKTKVAKYGEYKN-----------------WSNSIYNKLLLN----QPILAEILNW---------------------------------------SKKYPFN---------FKIVEIYQ-------THIR-----------NMD------------KH--------------------------
TLNKIKELADFVIRQKE-------------------------------EDKIKRTITRLR-GCKSTHEVRQVLLKWVGQHYQN----------------GEQTP-LITLEEYTQYLF--PEG-----I-NWREIRDLLLIGIYQKLHEHQMQIALESSEGD-----KEEETETINQ---------------------

EHR79718                ------MKNIKKERRLNEFLTRLEIQNTQKGRTDSHAISPIFTW----TGHPFTDAGLVALLLL-----------SGKEK-----------------------------------------PEELTEED-----VK---RAVKFVAKLYSQE------------------------------------------------------------------------------------GWAQLLARIYRNNNPIL----------------MINPSMRKYA-------
------------------------TPDKLEFS-LWGLYLQLIL---------------------------------------------------------------STR--KHKDRK------------VCQICGRYPVVSG-----LELRSIINTKERERPKEITGDIFP--LLGTG----D--LRNYF----------------P------H--GNEMG--ADICAHCLFLAQLSSVGSYVIT-TKNGKTQ----------G-----------------------VLI-----VHAYP-----YEKMVELN----KAAV---
NSAKK--AYL-----------YNTAEGF-KQPVNFLFELI-----RKVA-ENVE-----------TGYWE---DVSISA-YYYL--NGNR----TGEQW-IEIFYIPNPILRFVAFA----ARKDGI---G----------WK--KIVLMG-----W--RKRPSEKEFEDFKKR-------------------------YPNDVYSKLLSG----ESILPYFLDI--------------------------------QNRK-VNA---------GWNLLSFYC-------SEVL-----------GLD------------KE----------------------
ALEFIKGVADRIVETV-EKL-P-----------------------------DNKLSRRIRELE-RAEKLYQFESFFI-RVEKDRQE---------------F-GIPEA-LMSFDDFARILT--SYGEDLNV-SWKTVKNLLLFRIYEK--LHDRLMKASLEKAE----VEEA-E----EEFEEGVEK------------

ADG13584                ---------------------------MGNKTFY------W----TGHPFFADAGLSAILLL-----------NNKQN-------------------------------------------PDEIDEND-----IK---KAIEFASNLYAKN-----------------------------------------------------------------------------------EWVRYLSGKIFPNNGFI---------------------------------------------
MTNPSMS---------KK--RS-------PE-NIAK-----NLWKLYENI-PEILEPSS--------------------------------------------------KER-------------RCIICGRRV---------------------PYGKGDRSIFP--LLGTG-----K--MLNFF----------------H------S--GNPKG--EDICAHCIFLAQFMPLVAY-----------N--------LS-S-----------------VLV----IHTYP-----YSKMLRFY---DESL----RQVRI---SSL-----------VPDAKGF-
RKPENFFFKLL----IDIT-KNVE--------SS-KYWKNT-VVTLYY-FIND---------NRSQN-IDIIYIPNDILRFVAFA---GEVDPEG-------------WR--NIVNMG-----W--SKKDNLNFEEMEKTR--------------------------VN-EVYKKLLND----ESILRYFYDG-INKKV-------NA------------------------------------SWELLEFYC-------LEVL-----------KLD------------KE----------------------------ALDFIKDVGDRIIETI-EKL-N----------------
----------DNDLRKIVGELE-RSTKLYEFENFFI-RVEKIRQK---------------NGIPRS-LLTFDEFARLLM--GYGEDITM-SWKVVKDLLLFRIYEK--LHDRL---IKTSEE-----NYEG-D---EK--------------------

AIY90677                ---------------------------MEFY------W----TGHPFVDAGLAAILLI-----------AGKSK------------------------------------------PEELQKED-----VS---KAVDFAAELYATK---------------------------------------------------------------------------------DWSSSYLHGMIFPNSGI------------------------------------------------
VMSNPSMSK-------KR--SP-----------------------ENI-AQNLWSIL-----------------------------------------------------EKT---NATNPD------------SPTCIICGRKPA---------------------YTGKEIYLSVFP--LLGTG-----G--VTNYF----------------H------S--ANPKG--ADICAHCIFLTQFLPLVSY-----------R--------LS-R------------------VLV----IHAYP-----YELMLELS---KGAL----DEIKK---SKL-----------ASQAKGF-
RIPENFLFHLI----GEIT-RKVE--------RD-EFWENA-SVTLYY-FICN---------LQNQV-LDIINVPNPILRFVAYA---NKVDYVG-------------WR--RIVAMG-----W--KGKPSEAEIENFERER--------------------------RN-AVYSRLLNY----ESILPYFYNL----------------------------------KERK-VNA--------SWNLLAFYC-------KEVA-----------GLD------------KE----------------------------ALEFIKSVGDRIVESI-ERL-P------------
---------------NHKLDDEVRSLE-RAEKLHQFEAFFI-RIEKVRQN---------------L-GIEKP-LMTFDEFTKILT--AYGEDVNV-SWKTVRNLLLFRIYEK--LHDRLIKSDKSSEI-----SEVL-E----ENEYAIFGGEGE---------

AKG92216                ---------------------MSGVNLTREQSEKRTQSGSLFYW----TGHPFVDAGLVAILLI-----------AGKKK-----------------------------------------PEDLTVED-----VE---KAISFAAELYATK--------------------------------------------------------------------------------------EWSTKYIHGMIFPNSGI-----------------------------------------------------
-------LMANPSMGK-------KR--SP-----------------------TSIAEN-LQSIFEEL----------------------------------------------------KS--SDNDSP------------TCVLCGRRPAYSK---------------------KDIYLSVFP--LLGTG----G--VPNFF----------------H------S--ANLRG--ADICAHCVFLAQFMPLISY-----------R--------LS-R------------------VLV----IHAYP-----YELMLELS---KEAL----EDIKK---SKL-----------
ASQARGF-RRPENFLFHLV----GEVT-RKME--------SS-EFWENA-SVTLYY-FICN---------NQNQI-LDLIHIPTPVLRFVAYA----NYIDP---MG----------WK--RIVAMG-----W--RVLPDENDFDKFERER---------------------------RN-VVYSRLLNN----ESILSYFYNS---------------------------------------KERK-VNT--------NWNLLAFYC-------QEVL-----------GLD------------EK----------------------------ILEFVKDVGDRIVETI-
GML-P---------------------------DHKLDDEVRSLE-RAEKLYQFEAFFI-RIEKLRQN---------------L-GIDKP-LMTFDEFARILM--AYGEDVSI-SWKTVKNLLLFRIYEK--LHDRLMKSAELDKI-----EEFG-A-----DESEYSIYGGGNE--------

AEA46792                ---------------------MSSSNLVEDHTEEEKQVDGIEFYW----TGHPFVDAGLAAILLI-----------AGKNK-----------------------------------------PEELNEKD-----IE---KAIDFASKLYATK--------------------------------------------------------------------------------------EWSSSYLHGMIFPNSGI-----------------------------------------------------
--------LMANPSMTK-------KR---------------------SPEA-----IAENLWSIL-EEFKVNNR---------------------------------------DSP-----------------KCVLCGRRPA--------------------YTKKEVYLSVFP--LLGTG-----G--VPNFF----------------H------S--ANTRG--ADICAHCIFLTQFMPLASY-----------R--------LS-KPPR--------------VLV----IHAYP-----YELMFELS---KEAL----EDIKK---SKL-----------
ASQARGF-RRPENFLFHLV----GEIT-RKIE--------RD-EVWENS-SVSLYY-FICN---------NQSQI-LDVIYVPTPVLRFVAYA-----NLVDE----KG---------WK--HIVSMG-----WGRQPSDEEFEKFERSN---------------------------RN-EVYSRLLNN----ESILPYFINR---------------------------------TSRQ-TNT---------VWSLLAFYC-------SEVL-----------GLD------------EK----------------------------VLDFIKDIGDRIVETI-
GKL-P---------------------------EHKLDDEVRSLE-RAEKLYQFEAFFI-RVEKLRQN---------------L-GIERP-LMTFDEFARILT--AYGEDINI-SWRTVKNLLLFRIYEK--LHDRLMKSVESKKI-----EEMR-VEESEYAIYGGGEE-----------

KUK00291                ---------------------------MKLIPLFT-----W----TGHPFTDAGLVALLLL-----------NDKEK--------------------------------------------PEELTAED-----IK---RAVKFASELYARK-----------------------------------------------------------------------------EWSSGYIHAMMMPNSGI---------------------------------------------
LMANPSMAK-------KR--TP---------------------------EAV-ENSLLSLL-------------------------------------------------NEP--EDENAP-------------ICEVCGRRH-------------------AREKPVYRSDFP--LVGTG----G--IPNYF----------------P------S--GK-DG--ANICSHCVFLTQLMPLVAY-----------R--------LS-R------------------ILI----IHAYP-----YEFMVELH----RDAF----DDVVK--TQL-----------ASNAKNF-
RRPENFFFHLI----SDLM-RKIE--------HS-ELWKNT-SVTLYY-FVNN---------NQGQE-LDVIHIPNQILRFVAYA-----SLVDS----SG---------WK--RIVSMG-----W--RKRPSDEEFAEYERR---------------------------YSNEVYSRLING----QSILRYFIDK----------------------------------TSRE-VNS---------KWELLQFYC-------SEVL-----------GLD------------EK----------------------------ALEFVKDVGDRVVETL-EKL-P------
--------------------DNKLKRRVRELE-NATKLYQFEAFFV-GLEKLRQE---------------L-NIEKP-LMTFDEFARILT--SYGEDFNV-SWKSVKNLLLFRIYEK--LHGRLIREEEIGEE-----ETEE-V----EIYGFGGEEA------------

AMM54964                ---------------------------MGKGSPNFPTFE---W----TGHPFVDAGLIAILLL--------------------------------------------------SKKSRPQDLTWDD-----IQ---KAVEFASELYARK-----------------------------------------------------------------------------EWSSGYIHAMVLPNSGI---------------------------------------------
LMANPSMSK------------NR------------------------TSEKIKENL-LSLLKEKE-------------------------------------------------DKSAP------------MCEICGRRH-------------------ARSKPVYRSDFP--LVGTGG-------VPNYF----------------P------S--GK-DG--ANICSHCLFLAQFMPLIAY--------------------RMQR----------------LLI----IHTNP-----PEFMLRFHEEALNDV-----RKT-------------
QLASNARGFRRPENFLFEKL----MEIG-RDLE---------LGKLDGV-SITLYY-FVSN---------NQGQE-IDIIYIPNPVLRFIAFAGPEFRKIANM--------------GWAKGCTRRRSEEEVVRE------------------------------CRNEVYRRLLEN----ESILPYFVDV-QNRRV-----------------------------------------NSSWRLLEFYC-------LEVL-----------SLE------------KD--------------------------
ALEFVRDVADRIVETLEKLP------------------------------DNKLARRVRELE-RADKLYQFEAFFI-RIEKDRQE---------------LGIPKALITFDEFATLLTSYGED---LNVSWRTVKHLILFRIYEK--LHDKLMKVKEEEKE-----VEEE---------------------------

AJC72472                MKEIGRERPLDEFLTGQPKEEPSHSSGKAPEEKRNESLFY---W----TGHPFVDAGLVALLLI------SR-KNS--------------------------------------------PEELTEED-----IE---KAVDFAAKLYARK------------------------------------------------------------------------------------EWSSGYIHAMMLPNSGI-------------------------------------------
----------------------------LMANPSMSK-------KR--TP----------------------------EAI-AQNLKDLL-----------------------------------------------KEP--EDPNAP-------------ICEICGRRH-------------------SRPKPVYRSDFP--LVGTG----G--VPNYF----------------P-----S--AK-DG--LNVCSHCLFLVQFLPLVAY-----K----V--------GR-R------------------VLV----MHTYP-----YELMLELH---GEAL----NDVRK--
NFL-----------ASNARDF-KRPENFLFHLL----GELT-RKTE--------RD-DLWVNA-SITLYY-FVVN---------NKSPD-ITIIHVPTPVLRFIAHA----SREDPTG---------------WK--RIVAMG-----W--RKRPSEEEFEKYERS--------------------------YANEVYARLLAE----ESILPYFIDV---------------------------------KNRR-ANA--------KWSLLSFYC-------SEVL-----------GLD------------KE----------------------
ALEFIKDVGDRIVETL-EKL-P---------------------------DNQLSRRVRELE-RAEKLYQFEAFFV-NLEKLRQR---------------L-GIEKP-LMTFDEFATVLT--SYGEDLNV-SWRTVKHLLLFRVYER--LHDRL-----MKAS----EEVE-E---VGEAEEEVEDVGIYGLGEVEE

AFL94587                ----------------------------------------------------------------------------------------------MEDIEKAI----------DFASKLYAR---------------------------------------------------------------------------------KEWSSGYIHAMMLPNSGI----------------------------------------------------------------LMANPSMAR-------KR--TP---
-----------------------EAI-AKNLKALL------------------------------------------------NEQ--EDESAP-------------ICEICGRRH-------------------SRPRPVYRSDFP--LVGTG----G--VPNYF----------------P-----S--AK-DG--ANVCSHCLFLVQFLPLVAY-----K-----V--------GG-R------------------VLV----MHTYP-----HELMLELH---GEAI----NDVRK--SFL-----------ASNARDF-KRPENFLFHLL----GELT-
RKTE-------RD-DLWGRA-SITLYH-FVNN---------NQGQD-LQIIHVPTPVLRFVAHA----ARYDRA---G---------WR--RIVSMG-----W--RKRPSEEEFEKYERS--------------------------YANEVYARLLAE----ESILPYFIDI---------------------------------KNRR-ANA--------KWSLLAFYC-------SEVL-----------GLD------------KE----------------------------ALEFIKDVGDRIVETL-EKL-P---------------------------
DNQLSKRVRELE-RAERLYQFEAFFV-NLEKLRQR---------------L-GIEKP-LMTFDEFATVLT--SYGEDLNV-SWRTVKHLLLFRVYER--LHDRLMKASEEAGE----EVEE-N----EEVEIFGMGVEE---------

AEH24640                ------------MWIIGKRFPDPSLHESSKEDNEKVSLFD---W----TGHPFVDAGLVALLIL------NK-KRS-------------------------------------------PEELTEED-----IR---KATEFASKLYARK------------------------------------------------------------------------------------EWGSGYIHAMMLPNSGI-------------------------------------------
---------------LMANPSMSK-K-----------------------------------RTPEAI-AQNLRDLL------------------------------------------------QEP--EDPNAP-------------ICEICGRRH-------------------SRPKPVYRSDFP--LVGTG----G--VPNYF----------------P-----S--AK-DG--LNVCSHCLFLVQFLPLVAY-----K----V--------GG-R------------------VLV----MHTYP-----YELMLELH---EEAI----NDVRK--SFL-----------
ASNARDF--KRPENFLFRLL----GELT-RKTE--------RD-DLWGKA-SITLYH-FVNN---------NQGQD-LQIIHVPTPVLRFIAHA----SREDPKG-------------WK--RIVAMG-----W--RKRPSEEDFEKYERS-------------------------YANEVYSRLLAE----ESILPYFIDM---------------------------------KNRR-ANA--------KWSLLSFYC-------SEVL-----------GLD------------KE----------------------------ALEFIKDVGDRIVETL-
EKL-P---------------------------DNQLSRRVRELE-RAEKLYQFEAFFV-NLEKLRQR---------------L-GIEKP-LMTFDEFATVLT--SYGEDLNV-SWRTVKHLLLFRIYER--LHDKL--MKASEE----EEVE-E----NEEVGIFGMGVEG--------

SNY03438                ---------------------------MDGFVGD-----------PIITSGLLAIK----------------------------------------------LNTGFSISECPEDK-----LK---QASEQLVELYLTP--------------------------------------------------------------------------------------AWSKELQSIFPNSTYIQ--------------------------------------------
SAKNYDK-------KG--KS------------------------IEF-LYELIEGL------------------------------------------------------NSENNNDE----------YCVFCGSPAHQR------------------NDGKPFVKTQVP--LIGSS----D--FTNFF----------------P------S--FK-NG--MDLCAKCALAIQFSPLLYY-----------K---------SGGK-----------------PCC----ISCNNLLVLEKFGQECIEYITQKH-----AAGSFNSKETS---------
GILDEGF-KNPQNALFNLA----YKFI--NYE--------REEEAIDKE-EIVLYL-MDNY---------NQNPAGVSIYKLPNNVFSFVASV-MHSPELKDV---------------WY--SLLSKH----YRGKKDKKSGLPIWKTN---------------------------YN-QIHDYLLKN----RSILRFFRDD-VNKKT---------------------------------------------TVPWILVENYM-------QRVK----------NVN-----------QN--------------------------
RINAIKDYADRIAQCIT---------------------------ESKKTKRINDLI-SAKDFPTFKNQLR-MIGKDWQT------------------LGKQEP-LITFDDYVYSIMPDDYS-------NWRDVQNLITIRLYEE--LHPILANTDEEMTD----IDEERAEGEQI--------------------

PQV42764                ---------------------------MDGFVGD-----------PFITSGLLAI-KL-----------------------------------------------NTGFNIGECPEDKLK---QASEQLVELYLTP-------------------------------------------------------------------------------------AWSKELQSIFPNSTYIQ-----------------------------------------------
SAKNYDK-------KG--KS-------------------------IEF-LHDLIEGL------------------------------------------------------NSENNNDG----------YCVFCGSPAHQR-------------------NDGKPFVKTQVP--LIGSS----D--FTNFF----------------P------S--FK-NG--MDLCAKCALAIQFSPLLYY-----------K---------SGGK-----------------PCC----ISCNNLFVLEKFGQECIEYITQKN-----AAGSFDSKETS--------
GIFNEGF-KNPQNALFNLA----YKFI--NYE--------REEEAIDKE-EIVLYL-MDNY---------NQNPAGVSIYKLPNNVFSFVASV-MHSPELKDV--------------WY-SLLSKHY----WGKKDKKSGLPIWKTN-------------------------YN-QIHDYLLKN----RSILRFFRDD-VNKKT---------------------------------------------TVPWILVENYM-------QRVR----------NVN-----------QN--------------------------
RINAIKDYADRIAQCIT-----------------------------ESKKTKRINDLI-AAKDFPTFKNQLR-MVGKDWQT-----------------LGKEEP-LISFDDYVYSIMPDDYS-------NWRDVQNLITIRLYEE--LHPILANTDEERTE----GEQI--------------------------

AVX29687                ---------------------------MKYFTYTGN-----------PFIDTGLNVVLSW-----------HNKKA--------------------------------------------PEELIVED-----LE---KMIEISARLYGSE-----------------------------------------------------------------------------KWLKNLYSVFPNN--------------------------------------------
PFTNPSIKN-------KE--SR------------------------YLEF-MNDLKNNI-------------------------------------------------------K--PLDTGR-------------GCVGCGKR-------------------EPYKKFTKTDIP--LTGTK----D--LRNLF----------------P------A--AE-AG--ESYCALCAMLIQFMPMALF-----A----C---------GG-R------------------FML----VHSAN-----SEINGIYV---RRCL---KEIKQQIALGDMS---------
GVFDQGY-KNPINALFFLA----EEIV-RDIN---------DFFDEPVMVRIYS-FTNY---------NQSPD-LNIYDLPAPVFYFIVQA--QRKGFTQA-------------WN--ELKNKG----Y--TGKKAENTSKEKRKN--------------------------LNNEVLLKLLNE----ESVVPYFLNI-----------------------------------STRS-IVG---------NFEFLSLYL-------KEVR-----------RYM------------QR-------------------------RIESIKRLGDELAIFI-R-----
-------------------------ENNDQRRLYLLE-TVDKFDDFRNQLR-IIYRKRLD---------------M-GISTP-LFTVDEYVENLF--PDG---PV-NWRETRDLILFRIYET--LSGWLAENAGEENE----EEEE-E------------------------

CUU09222                ---------------------------MMGKIFDITGN-----------PFVDAGIYIIAEL------AE-KE-------------------------------------------VKDVDFAD-----LE---KLYPKLIDVYFTA---------------------------------------------------------------------------------GWKKMLESI-------------------------------------------------------
FTGNHPVTHKK-----SG--SQ-----------------------SYYRKF-LEGWISDI-------------------------------------------------------G--APLPAG-----------TCFACGRRNKSNV-------------------ERKYGIGRELVP--LTGSG----D--FVNFF----------------P---------MASLG--LEMCSACLFAIQFMPIFLH-----Y-----C----------GG-K------------------FLL----LHSNS-----EKVMKYWA---KNCV----TNLYK--QITLNN-----
FTGCYSNNI--NDPVNSFFQSI----EDMV-IRYD---------ENWIDE-DVSIRF-YYFS--NY-----GQSPY-VEVFDVPTPVFRFLAYV--KQTEAYDE-------------WR--KIVNKA----S--GG------------------------------------------RRNEVYERLLRG----ESIVRYFFKD-KREIFG----------------------------------------------NWELLKFYL-------REVK-----------EMD------------EK----------------------------RINTIREFADRIANLI------------
--------------------QSLDSVKRLEQLERAKDYAEFRNVLR-FLIKDGIR---------------A-GLREP-LLRFDEYVNLIF--PEG---AM-GWSEVRDLMLFRIYEV--LHNWL------VER----KVEE-P---TE-------------------

CUS92795__7_            ---------------------------MAQIYIITGN-----------PFVDAGIYVISEL------------ARKE-----------------------------------------IEDVDVDD-----LR---NLSGKIIDVYSND--------------------------------------------------------------------------------KWGRMLDLIFTKNHPMR-------------------------------------------
----------------------HPSV------GGREYYEKF-LLGLLSDL-----------------------------------------------------S--PLSDSG-------------ACVSCGKRNKSRV-------------------ERKFGIGKELVP--LTGSG----D--FVNFF----------------P---------MASLG--LEMCSACLLAIQFMPIFLH-----Y-----C----------AG-K------------------LLL----LHSNS-----GKVMRYWA---KIGV----SNFDA--QIVFGN------FTGCYSNNI--
NDPVSSFFKSI----EDVV-MKYE----------ENWIEE-DAFIRF-YYFV--NP-----GQSPY-VEIFDVPSPVFRFIAYV-KQADAYAE-------------WK--EMVGRA----Y--DG----------------------------------------------------RKNKVYLRLLSG----ESIVRYFFRN------------------------------------KGERR-VFG--------NWNLLKFYL-------KEVK-----------GMN------------EE----------------------------RINAIREFSDRIATVI-K-----------------------
-----SLGSIKRLNQLE-TAENYAQFRNVLR-FLIKDSLR---------------A-KFEKP-ILEFDEYVKLIL--PEG---AT-GWNEVRDLILFRIYEV--LHSWL--IEIEGEK----EASE-Q----I-------------------

CUS87614                ---------------------------MAQIYIITGN-----------PFVDAGIYVISEL------------------------------------------------ARKE--IEDVDVDD-----LR---NLSGKIIDVYSND------------------------------------------------------------------------------------------KWGRMLDLI-----------------------------------------------------------FTKNHPMRH--
-----PSVGGR-------------------EYYEKF-LLGLLSDL-----------------------------------------------------SLLSDSG-----------ACVSCGKRNKSRV-------------------ERKFGIGKELVP--LTGSG----D--FVNFF----------------P---------MASLG--LEMCSACLLAIQFMPIFLH-----------Y---------CAGK-----------------LLL----LHSNS-----GKVMRYWAKIGVSNFDAQIVFGNFTGCYSNNI----------------
NDPVSSFFKSI----EDVV-MKYE----------ENWIEE-DAFIRF-YYFV-------NPGQSPY-VEIFDVPSPVFRFIAYV-KQADAYAE-------------WK--EMVGRA----YDG--------------------------------------------------RKNKVYLRLLSG----ESIVRYFFRN----------------------------------KGERRVFG--------NWNLLKFYL-------KEVK-----------GMN------------EE----------------------------RINAIREFSDRIATVI----------------------------
----KSLGSIKRLNQLETAENYAQFRNVLR-FLIKDSLR----------------AKFEKP-ILEFDEYVKLILPEGAT------GWNEVRDLILFRIYEV--LHSWLIEIEGEKEA----SEQI----------------------------

ACD89534                ---------------------------MSSLFQYTGN-----------PFVDAGISALTNW-----------CDKKT------------------------------------------PQELTEAD-----IK---KALPEIANLFSQG-------------------------------------------------------------------------------------------AWVKTFYT---------------------------------------------------
TFSNGVMVQ-------PS--NK-------------G-----KEREKWLEF-IGDLVKEL------------------------------------------------------Q--PLADHG------------SCVACGCRNAIKI-----------------KKEKRGLLRSEV--PMASG-------SLNYY----------------------S--FASTG--ADYCGTCAIAIQVSPLVLY-----R----S---------GG-K------------------MIL----VHSSS-----EKAMCSWAKMAINEVRSQISLRNYTGCF---------
----TENF-TNPQNALFRIA----KILI---------QDKDDWKSD-PITIRI-YYFT--NY-----GQGAE-LKYYDLPNRVFHFLNEV-----HHSEE---LKD-------WD--KIIGST-----Y--FFKKNNSKIYLNTDDKSEEEYKN-----------------NNNVIYEGLLKD----EWIVKYFYNF--------------------------------LQRK-AYA---------KWELVQLYL-------KEVR-----------QMD------------KQ-------------------------RTEVIKRVADEISLVI-
QRD------------------------------ESHNPKRLWQLE-RANSYGTFRNVLR-LIIKDRIK----------------NGAERP-LFSIEDYTERLF--PDG---AL-CWRETQDLILFRLYEM--LHGWLKERDIVIDE-----VEEN-S----TTEIE-----------------
```

```
SHF19325          [illegible]
[illegible]

SNX53177          [illegible]
[illegible]

AKB85836          [illegible]
[illegible]

AKB49944__2_          [illegible]
[illegible]

AKB22698          [illegible]
[illegible]

AKB19482          [illegible]
[illegible]

AEF93612          [illegible]
[illegible]

KXK23069          [illegible]
[illegible]

RCK75356          [illegible]
[illegible]

GAP08575          [illegible]
[illegible]

ABX07704          [illegible]
[illegible]

YP_001636534__4_          [illegible]
[illegible]

WP_028460058          [illegible]
[illegible]

WP_031460739          [illegible]
[illegible]

ABQ90254          [illegible]
[illegible]

ACL23511          [illegible]
[illegible]

ABU59349          [illegible]
[illegible]

;

end;
```

### Cas8c

```
#
#NEXUS
begin characters;
        dimensions nchar=3805;
        format datatype=protein missing=? gap=-;
        matrix
        'TCO63493.1'          [illegible]
[illegible]
        'KRM12369.1__2_'          [illegible]
[illegible]
        'WP_037327719.1__2_'          [illegible]
[illegible]
        'KGF08220.1'          [illegible]
[illegible]
        'EFA89355.1'          [illegible]
[illegible]
        'EFR33734.1'          [illegible]
[illegible]
        'KXB68630.1'          [illegible]
[illegible]
        'KXA28915.1'          [illegible]
[illegible]
        'SFH49362.1'          [illegible]
[illegible]
```

[illegible]

'SFE14385.1'

[illegible]

'KEY48540.1'

[illegible]

'KIS20160.1__4_'

[illegible]

'AGS46823.1'

[illegible]

'KIS09040.1'

[illegible]

'KIS13996.1'

[illegible]

'ACG61922.1'

[illegible]

'KIS15331.1'

[illegible]

'KIS21048.1'

[illegible]

'KIS12402.1'

[illegible]

'AIA68819.1'

[illegible]

'KIS09059.1'

[illegible]

'EFR44838.1'

[illegible]

'SQI47933.1'

[illegible]

'SQF48102.1'

[illegible]

'WP_019314884.1'

[illegible]

'SQG92789.1'

[illegible]

'SUN02755.1'

[illegible]

'CCW39408.1'

[illegible]

'EPU94207.1'

[illegible]

[illegible]

KI-K-PYEE----PLK--L---NRRGSWMK---L---EKE-K--E---EIL---E----LL----N---P---L---LE-TE-T---MEKPLDYRFIFGYY---A---E---K---NY---Y---Y---T---K---
---QNTEVTES---E---E---
'SQF06910.1' ---MDF---FTSLLKTYE---K---AE---LADL---V---DHQKR---SNESV---LLPI--YH---TSLKS-NGKNI-ISVKLDK--D--GQ-FH---K-A---E---FM---A---DK---QM---I---I-F-P---VTAD-SV--AR---S-G--
S---H---P--A---P---H---P-L---VDKFAYY---S---AE---M---G---Q---S---Q---Y-D---SF-HK-QLNN--W--IDY---C-EEG--
DVKRF--LTI---VQQFILK---P---EFL---I---L---I---L---D---S---L---IG---P--D-Y---QNN---Q---L---KVTFCDATG---KEK---LID---LSA---C---F---LEF---S-I---DQ---FQG---F---KNES---VS---T---FK---T---L-HQ-SY---I---
S---F---V---EA---NREK--LG---I---CN---ISG---RE-E-Q---LTDK-H--R---GL--M---G-N---A---KIIS---VSN---NDSRLDIVSP---K---R---E-AY--K---G---R---FRE---RE---D---VFS---
--VGYETSEKIHLMLKY-LL--EN---K---N---T---S---TW---L---G---S--SQY---LIN--W---FSDDLT---
DGLEEDDDDDTPPVITL---A---T---E---DNKRI---GKS---F---I-K-G---Q---KL---FA---N---D---A---T---Y--YVAILNK-TSN---GRIALKYFRQL-QASQLLTNLNKWQ--ET---YS--WE---S---RS---K---FGKSRLRTP---T--
---FHDIL---N--V---S---YGVDRD---R-FLELD---NDNFK---S---D---QIQKLVASLID--G-K--P---MPQSIVKK-LG---N---N--V---KE---RHRY---HKH---W---
YQVEQVCLAIL---HKQ---N---G---E---E---FSP---MLD---HT---NQ---N---RSYLF---GRLLAIFE---LIE---T---L---RYG---LDGNN---ND---R---ITNA---ERYWTAYT---GQPTKL---MMLLEN--
KI-K-PYEE----PLK--L---NRRGSWMK---L---EKE-K--E---EIL---E----LL----N---P---L---LE-TE-T---MEKPLDYRFIFGYY---A---E---K---NY---Y---Y---T---K---
---QNTEVTES---E---E---
'VGU61232.1__11_' ---MDF---FTSLLKTYE---K---AE---LADL---V---DHQKR---SNESV---LLPI--YH---TSLKS-NGKNI-ISVKLDK--D--GQ-FH---K-A---E---FM---A---DK---QM---I---I-F-P---VTAD-SV--AR---S-G--
S---H---P--A---P---H---P-L---VDKFAYY---S---AE---M---G---Q---T---Q---Y-D---SF-HK-QLNN--W--IDY---C-EEG--
DVKRF--LTI---VQQFILK---P---EFL---T---L---I---L---D---S---L---IG---P--D-Y---QNN---Q---L---KVTFCDATG---KEK---LID---LSA---C---F---LEF---S-I---DQ---FQG---F---KNES---VS---T---FK---T---L-HQ-SY---I---
S---F---V---EA---NREK--LG---I---CN---ISG---RE-E-Q---LTDK-H--R---GL--M---G-N---A---KIIS---VSN---NDSRLDIVSP---K---R---E-AY--K---G---R---FRE---RE---D---VFS---
--VGYETSEKIHLMLKY-LL---E---N---K---N---T---S---TW---L---G---S---SQ---Y---LIN--W---FSD---D--LTNDS---RL--D---I---VSP---I---F---D---
---D---TPPVITL---A---T---E---DNKRI---GKS---F---I-K-G---Q---K-LF---AN---D---A---T---Y--YVAILNK-TSN---GRIALKYFRQL-QASQLLTNLNKWQ--ET---YS--WE---S---RS---K---FGKSRLRTP---T---FHDIL---
---N--V---S---YGVDRD---R-FLELD---NDNFK---S---D---QIQKLVASLID--G-K--P---MPQSIVKK-LG---N---N--V---KE---RHRY---HKH---W---YQVEQVCLAIL---
HKQ---N---G---E---E---FSP---MLD---HT---NQ---N---RSYLF---GRLLAIFE---LIE---T---L---RYG---LDGNN---ND---R---ITNA---ERYWTAYT---GQPTKL---MMLLEN--KI-K-PYEE---PLK--L---
NRRGSWMK---L---EKE-K--E---EIL---E----LL----N---P---L---LE-TE-T---MEKPLDYRFIFGYY---A---E---K---NY---Y---Y---T---K---QNTEVTES---
----E---E---
'VGT80484.1' ---MDF---FTSLLKTYE---K---AE---LADL---V---DHQKR---NNVPV---LLPI--YH---TSLKS-NGKNI-ISVKLDK--D--GQ-FH---K-A---E---FM---A---DK---QM---I---I-F-P---VTAD-SV--AR---S-G--
S---H---P--A---P---H---P-L---VDKFAYY---S---AE---M---G---Q---S---Q---Y-D---SF-HK-QLNN--W--IDY---C-EEG--
DVKRF--LTI---VQQFILK---P---EFL---T---L---I---L---D---S---L---IG---P--D-Y---QHN---Q---L---KVTFCDATG---KEK---LID---LSA---C---F---LEF---S-I---DQ---FQG---F---KNES---VS---T---FK---T---L-HQ-SY---I---
S---F---V---EA---NREK--LG---I---CN---ISG---RE-E-Q---LTDK-H--R---GL--M---G-N---A---KIIS---VSN---NDSRLDIVSP---K---R---E-AY--K---G---R---FRE---RE---D---VFS---
--VGYETSEKIHLMLKY-LL--EN---K---N---T---S---TW---L---G---S--SQY---LIN--W---FSDDLT---
DGLEEDDDDDTPPVITL---A---T---E---DNKRI---GKS---F---I-K-G---Q---KL---FA---N---D---A---T---Y--YVAILNK-TSN---GRIALKYFRQL-QASQLLTNLNKWQ--ET---YS--WE---S---RS---K---FGKSRLRTP---T--
---FHDIL---N--V---S---YGVDRD---R-FLELD---NDNFK---S---D---QIQKLVASLID--G-K--P---MPQSIVKK-LG---N---N--V---KE---RHRY---RKH---W---
YQVEQVCLAIL---HKQ---N---G---E---E---FSP---MLD---HT---NQ---N---RSYLF---GRLLAIFE---LIE---T---L---RYG---LDGNN---ND---R---ITNA---ERYWTAYT---GQPTKL---MMLLEN--
KI-K-PYEE----PLK--L---NRRGSWMK---L---EKE-K--E---EIL---E----LL----N---P---L---LE-TE-T---MEKPLDYRFIFGYY---A---E---K---NY---Y---Y---T---K---
---QNTEVTES---E---E---
'VGQ26711.1' ---MDF---FTSLLKTYE---K---AE---LADL---V---DHQKR---SNEPV---LLPI--YH---TSLKS-NGKNI-ISVKLDK--D--GQ-FH---K-A---E---FM---A---DK---QM---I---I-F-P---VTAD-SV--AR---S-G--
S---H---P--A---P---H---P-L---VDKFAYY---S---AE---M---G---Q---S---Q---Y-D---SF-HK-QLNN--W--IDY---C-EEG--
DVKRF--LTI---VQQFILK---P---EFL---I---L---I---L---D---S---L---IG---P--D-Y---QHN---Q---L---KVTFCDATG---KEK---LID---LSA---C---F---LEF---S-I---DQ---FQG---F---KNES---VS---T---FK---T---L-HQ-SY---I---
S---F---V---EA---NREK--LG---I---CN---ISG---RE-E-Q---LTDK-H--R---GL--M---G-N---A---KIIS---VSN---NDSRLDIVSP---K---R---E-AY--K---G---R---FRE---RE---D---VFS---
--VGYETSEKIHLMLKY-LL--EN---K---N---T---S---TW---L---G---S--SQY---LIN--W---FSDDLT---
DGLEEDDDDDTPPVITL---A---T---E---DNKRI---GKS---F---I-K-G---Q---KL---FA---N---D---A---T---Y--YVAILNK-TSN---GRIALKYFRQL-QASQLLTNLNKWQ--ET---YS--WE---S---RS---K---FGKSRLRTP---T--
---FHDIL---N--V---S---YGIDRD---R-FLELD---NDNFK---S---D---QIQKLVASLID--G-K--P---MPQSIVKK-LG---N---N--V---KE---RHRY---RKH---W---
YQVEQVCLAIL---HKQ---N---G---E---E---FSP---MLD---HT---NQ---N---RSYLF---GRLLAIFE---LIE---T---L---RYG---LDGNN---ND---R---ITNA---ERYWTAYT---GQPTKL---MMLLEN--
KI-K-PYEE----PLK--L---NRRGSWMK---L---EKE-K--E---EIL---E----LL----N---P---L---LE-TE-T---MEKPLDYRFIFGYY---A---E---K---NY---Y---Y---T---K---
---QNTEVTES---E---E---
'VGT31614.1__13_' ---MDF---FTSLLKTYE---K---AE---LADL---V---DHQKR---NNEPV---LLPI--YH---TSLKS-NGKNI-ISVKLDK--D--GQ-FH---K-A---E---FM---A---DK---QM---I---I-F-P---VTAD-SV--AR---S-G--
S---H---P--A---P---H---P-L---VDKFAYY---S---AE---M---G---Q---S---Q---Y-D---SF-HK-QLNN--W--IDY---C-EEG--
DVKRF--LTI---VQQFILK---P---EFL---T---L---I---L---D---S---L---IG---P--D-Y---QHN---Q---L---KVTFCDATG---KEK---LID---LSA---C---F---LEF---S-I---DQ---FQG---F---KNES---VS---T---FK---T---L-HQ-SY---I---
S---F---V---EA---NREK--LG---I---CN---ISG---RE-E-Q---LTDK-H--R---GL--M---G-N---A---KIIS---VSN---NDSRLDIVSP---K---R---E-AY--K---G---R---FRE---RE---D---VFS---
--VGYETSEKIHLMLKY-LL--EN---K---N---T---S---TW---L---G---S--SQY---LIN--W---FSDDLR---
DGLEEDDDDDTPPVITL---A---T---E---DNKRI---GKS---F---I-K-G---Q---KL---FA---N---D---A---T---Y--YVAILNK-TSN---GRIALKYFRQL-QASQLLTNLNKWQ--ET---YS--WE---S---RS---K---FGKSRLRTP---T--
---FHDIL---N--V---S---YGVDRD---R-FLELD---NDNFK---S---D---QIQKLVASLID--G-K--P---MPQSIVKK-LG---N---N--V---KE---RHRY---RKH---W---
YQVEQVCLAIL---HKQ---N---G---E---E---FSP---MLD---HT---NQ---N---RSYLF---GRLLAIFE---LIE---T---L---RYG---LDGNN---ND---R---ITNA---ERYWTAYT---GQPTKL---MMLLEN--
KI-K-PYEE----PLK--L---NRRGSWMK---L---EKE-K--E---EIL---E----LL----N---P---L---LE-TE-T---MEKPLDYRFIFGYY---A---E---K---NY---Y---Y---T---K---
---QNTEVTES---E---E---
'VGT88492.1__2_' ---MDF---FTSLLKTYE---K---AE---LADL---V---DHQKR---NNEPV---LLPI--YH---TSLKS-NGKNI-ISVKLDK--D--GQ-FH---K-A---E---FM---A---DK---QM---I---I-F-P---VTAD-SV--AR---S-G--
S---H---P--A---P---H---P-L---VDKFAYY---S---AE---M---G---Q---S---Q---Y-D---SF-HK-QLNN--W--IDY---C-EEG--
DVKRF--LTI---VQQFILK---P---EFL---T---L---I---L---D---S---L---IG---P--D-Y---QNN---Q---L---KVTFCDATG---KEK---LID---LSA---C---F---LEF---S-I---DQ---FQG---F---KNES---VS---T---FK---T---L-HQ-SY---I---
S---F---V---EA---NREK--LG---I---CN---ISG---RE-E-Q---LTDK-H--R---GL--M---G-N---A---KIIS---VSN---NDSRLDIVSP---K---R---E-AY--K---G---R---FRE---RE---D---VFS---
--VGYETSEKIHLMLKY-LL--EN---K---N---T---S---TW---L---G---S--SQY---LIN--W---FSDDLT---
DGLEEDDDDDTPPVITL---A---T---E---DNKRI---GKS---F---I-K-G---Q---KL---FA---N---D---A---T---Y--YVAILNK-TSN---GRIALKYFRQL-QASQLLTNLNKWQ--ET---YS--WE---S---RS---K---FGKSRLRTP---T--
---FHDIL---N--V---S---YGVDRD---R-FLELD---NDNFK---S---D---QIQKLVASLID--G-K--P---MPQSIVKK-LG---N---N--V---KE---RHRY---RKH---W---
YQVEQVCLAIL---HKQ---N---G---E---E---FSP---MLD---HT---NQ---N---RSYLF---GRLLAIFE---LIE---T---L---RYG---LDGNN---ND---R---ITNA---ERYWTAYT---GQPTKL---MMLLEN--
KI-K-PYEE----PLK--L---NRRGSWMK---L---EKE-K--E---EIL---E----LL----N---P---L---LE-TE-T---MENPLDYRFIFGYY---A---E---K---NY---Y---Y---T---K---
---QNTEVTES---E---E---
'VGR61333.1' ---MDF---FTSLLKTYE---K---AE---LADL---V---DHQKR---NNEPV---LLPI--YH---TSLKS-NGKNI-ISVKLDK--D--GQ-FH---K-A---E---FM---A---DK---QM---I---I-F-P---VTAD-SV--AR---S-G--
S---H---P--A---P---H---P-L---VDKFAYY---S---AE---M---G---Q---S---Q---Y-D---SF-HK-QLNN--W--IDY---C-EEG--
DVKRF--LTI---VQQFILK---P---EFL---T---L---I---L---D---S---L---IG---P--D-Y---QHN---Q---L---KVTFCDATG---KEK---LID---LSA---C---F---LEF---S-I---DQ---FQG---F---KNES---VS---T---FK---T---L-HQ-SY---I---
S---F---V---EA---NREK--LG---I---CN---ISG---RE-E-Q---LTDK-H--R---GL--M---G-N---A---KIIS---VSN---NDSRLDIVSP---K---R---E-AY--K---G---R---FRE---RE---D---VFS---
--VGYETSEKIHLMLKY-LL---E---NK---N---T---S---TW---L---G---S---S---QYLINW---FSD---D---L---TN-D---S---RLD---IV---S---PI---F---D-G-LEEDD---
---D-D---TPPVITL---A---T---E---DNKRI---GKS---F---I-K-G---Q---K-LF---AN---D---A---T---Y--YVAILNK-TSN---GRIALKYFRQL-QASQLLTNLNKWQ--ET---YS--WE---S---RS---K---FGKSRLRTP---T---FHDIL---
---N--V---S---YGVDRD---R-FLELD---NDNFK---S---D---QIQKLVASLID--G-K--P---MPQSIVKK-LG---N---N--V---KE---RHRY---RKH---W---YQVEQVCLAIL---
HKQ---N---G---E---E---FSP---MLD---HT---NQ---N---RSYLF---GRLLAIFE---LIE---T---L---RYG---LDGNN---ND---R---ITNA---ERYWTAYT---GQPTKL---MMLLEN--KI-K-PYEE---PLK--L---
NRRGSWMK---L---EKE-K--E---EIL---E----LL----N---P---L---LE-TE-T---MEKPLDYRFIFGYY---A---E---K---NY---Y---Y---T---K---QNTEVTES---
----E---E---
'AIW25134.1__32_' ---MDF---FTSLLKTYE---K---AE---LADL---V---DHQKR---SNEPV---LLPI--YH---TSLKS-NGKNI-ISVKLDK--D--GQ-FH---K-A---E---FM---A---DK---QM---I---I-F-P---VTAD-SV--AR---S-G--
S---H---P--A---P---H---P-L---VDKFAYY---S---AE---M---G---Q---S---Q---Y-D---SF-HK-QLNN--W--IDY---C-EEG--
DVKRF--LTI---VQQFILK---P---EFL---T---L---I---L---D---S---L---IG---P--D-Y---QNN---Q---L---KVTFCDATG---KEK---LID---LSA---C---F---LEF---S-I---DQ---FQG---F---KNES---VS---T---FK---T---L-HQ-SY---I---
S---F---V---EA---NREK--LG---I---CN---ISG---RE-E-Q---LTDK-H--R---GL--M---G-N---A---KIIS---VSN---NDSRLDIVSP---K---R---E-AY--K---G---R---FRE---RE---D---VFS---
--VGYETSEKIHLMLKY-LL--EN---K---N---T---S---TW---L---G---S--SQY---LIN--W---FSDDLT---
DGLEEDDDDDTPPVITL---A---T---E---DNKRI---GKS---F---I-K-G---Q---KL---FA---N---D---A---T---Y--YVAILNK-TSN---GRIALKYFRQL-QASQLLTNLNKWQ--ET---YS--WE---S---RS---K---FGKSRLRTP---T--
---FHDIL---N--V---S---YGVDRD---R-FLELD---NDNFK---S---D---QIQKLVASLID--G-K--P---MPQSIVKK-LG---N---N--V---KE---RHRY---RKH---W---
YQVEQVCLAIL---HKQ---N---G---E---E---FSP---MLD---HT---NQ---N---RSYLF---GRLLAIFE---LIE---T---L---RYG---LDGNN---ND---R---ITNA---ERYWTAYT---GQPTKL---MMLLEN--
KI-K-PYEE----PLK--L---NRRGSWMK---L---EKE-K--E---EIL---E----LL----N---P---L---LE-TE-T---MGKPLDYRFIFGYY---A---E---K---NY---Y---Y---T---K---
---QNTEVTES---E---E---
'VHJ63082.1__3_' ---MDF---FTSLLKTYE---K---AE---LADL---V---DHQKR---SNEPV---LLPI--YH---TSLKS-NGKNI-ISVKLDK--D--GQ-FH---K-A---E---FM---A---DK---QM---I---I-F-P---VTAD-SV--AR---S-G--
S---H---P--A---P---H---P-L---VDKFAYY---S---AE---M---G---Q---S---Q---Y-D---SF-HK-QLNN--W--IDY---C-EEG--
DVKRF--LTI---VQQFILK---P---EFL---T---L---I---L---D---S---L---IG---P--D-Y---QNN---Q---L---KVTFCDATG---KEK---LID---LSA---C---F---LEF---S-I---DQ---FQG---F---KNES---VS---T---FK---T---L-HQ-SY---I---
S---F---V---EA---NREK--LG---I---CN---ISG---RE-E-Q---LTDK-H--R---GL--M---G-N---A---KIIS---VSN---NDSRLDIVSP---K---R---E-AY--K---G---R---FRE---RE---D---VFS---
--VGYETSEKIHLMLKY-LL--EN---K---N---T---S---TW---L---G---S--SQY---LIN--W---FSDDLT---
DGLEEDDDDDTPPVITL---A---T---E---DNKRI---GKS---F---I-K-G---Q---KL---FA---N---D---A---T---Y--YVAILNK-TSN---GRIALKYFRQL-QASQLLTNLNKWQ--ET---YS--WE---S---RS---K---FGKSRLRTP---T--
---FHDIL---N--V---S---YGVDRD---R-FLELD---NDNFK---S---D---QIQKLVASLID--G-K--P---MPQSIVKK-LG---N---N--V---KE---RHRY---HKH---W---
YQVEQVCLAIL---HKQ---N---G---E---E---FSP---MLD---HT---NQ---N---RSYLF---GRLLAIFE---LIE---T---L---RYG---LDGNN---ND---R---ITNA---ERYWTAYT---GQPTKL---MMLLEN--
KI-K-PYEE----PLK--L---NRRGSWMK---L---EKE-K--E---EIL---E----LL----N---P---L---LE-TE-T---MEKPLDYRFIFGYY---A---E---K---NY---Y---Y---T---K---
---QNTEVTES---E---E---
'VGT70874.1' ---MDF---FTSLLKTYE---K---AE---LADL---V---DHQKR---SNEPV---LLPI--YH---TSLKS-NGKNI-ISVKLDK--D--GQ-FH---K-A---E---FM---A---DK---QM---I---I-F-P---VTAD-SV--AR---S-G--
S---H---P--A---P---H---P-L---VDKFAYY---S---AE---M---G---Q---S---Q---Y-D---SF-HK-QLNN--W--IDY---C-EEG--
DVKRF--LTI---VQQFILK---P---EFL---T---L---I---L---D---S---L---IG---P--D-Y---QHN---Q---L---KVTFCDATG---KEK---LID---LSA---C---F---LEF---S-I---DQ---FQG---F---KNES---VS---T---FK---T---L-HQ-SY---I---
F---F---V---EA---NREK--LG---I---CN---ISG---RE-E-Q---LTDK-H--R---GL--M---G-N---A---KIIS---VSN---NDSRLDIVSP---K---R---E-AY--K---G---R---FRE---RE---D---VFS---
--VGYETSEKIHLMLKY-LL--EN---K---N---T---S---TW---L---G---S--SQY---LIN--W---FSDDLT---
DGLEEDDDDDTPPVITL---A---T---E---DNKRI---GKS---F---I-K-G---Q---KL---FA---N---D---A---T---Y--YVAILNK-TSN---GRIALKYFRQL-QASQLLTNLNKWQ--ET---YS--WE---S---RS---K---FGKSRLRTP---T--
---FHDIL---N--V---S---YGVDRD---R-FLELD---NDNFK---S---D---QIQKLVASLID--G-K--P---MPQSIVKK-LG---N---N--V---KE---RHRY---RKH---W---
YQVEQVCLAIL---HKQ---N---G---E---E---FSP---MLD---HT---NQ---N---RSYLF---GRLLAIFE---LIE---T---L---RYG---LDGNN---ND---R---ITNA---ERYWTAYT---GQPTKL---MMLLEN--
KI-K-PYEE----PLK--L---NRRGSWMK---L---EKE-K--E---EIL---E----LL----N---P---L---LE-TE-T---MEKPLDYRFIFGYY---A---E---K---NY---Y---Y---T---K---
---QNTEVTES---E---E---
'AAX72403.1__8_' ---MDF---FTSLLKTYE---K---AE---LADL---V---DHQKR---SNEPV---LLPI--YH---TSLKS-NGKNI-ISVKLDK--D--GQ-FH---K-A---E---FM---A---DK---QM---I---I-F-P---VTAD-SV--AR---S-G--
S---H---P--A---P---H---P-L---VDKFAYY---S---AE---M---G---Q---S---Q---Y-D---SF-HK-QLNN--W--IDY---C-EEG--
DVKRF--LTI---VQQFILK---P---EFL---T---L---I---L---D---S---L---IG---P--D-Y---QNN---Q---L---KVTFCDATG---KEK---LID---LSA---C---F---LEF---S-I---DQ---FQG---F---KNES---VS---T---FK---T---L-HQ-SY---I---
S---F---V---EA---NREK--LG---I---CN---ISG---RE-E-Q---LTDK-H--R---GL--M---G-N---A---KIIS---VSN---NDSRLDIVSP---K---R---E-AY--K---G---R---FRE---RE---D---VFS---
--VGYETSEKIHLMLKY-LL---E---NK---N---T---S---TW---L---G---S---S---QYLINW---FSDDLT---
DGLEEDDDDDTPPVITL---A---T---E---DNKRI---GKS---F---I-K-G---Q---KL---FA---N---D---A---T---Y--YVAILNK-TSN---GRIALKYFRQL-QASQLLTNLNKWQ--ET---YS--WE---S---RS---K---FGKSRLRTP---T--
---FHDIL---N--V---S---YGVDRD---R-FLELD---NDNFK---S---D---QIQKLVASLID--G-K--P---MPQSIVKK-LG---N---N--V---KE---RHRY---RKH---W---
YQVEQVCLAIL---HKQ---N---G---E---E---FSP---MLD---HT---NQ---N---RSYLF---GRLLAIFE---LIE---T---L---RYG---LDGNN---ND---R---ITNA---ERYWTAYT---GQPTKL---MMLLEN--
KI-K-PYEE----PLK--L---NRRGSWMK---L---EKE-K--E---EIL---E----LL----N---P---L---LE-TE-T---MEKPLDYRFIFGYY---A---E---K---NY---Y---Y---T---K---
---QNTEVTES---E---E---
'VGQ56204.1__2_' ---MDF---FTSLLKTYE---K---AE---LADL---V---DHQKR---NNEPV---LLPI--YH---TSLKS-NGKNI-ISVKLDK--D--GQ-FH---K-A---E---FM---A---DK---QM---I---I-F-P---VTAD-SV--AR---S-G--
S---H---P--A---P---H---P-L---VDKFAYY---S---AE---M---G---Q---S---Q---Y-D---SF-HK-QLNN--W--IDY---C-EEG--
DVKRF--LTI---VQQFILK---P---EFL---T---L---I---L---D---S---L---IG---P--D-Y---QHN---Q---L---KVTFCDATG---KEK---LID---LSA---C---F---LEF---S-I---DQ---FQG---F---KNES---VS---T---FK---T---L-HQ-SY---I---
S---F---V---EA---NREK--LG---I---CN---ISG---RE-E-Q---LTDK-H--R---GL--M---G-N---A---KIIS---VSN---NDSRLDIVSP---K---R---E-AY--K---G---R---FRE---RE---D---VFS---
--VGYETSEKIHLMLKY-LL--EN---K---N---T---S---TW---L---G---S--SQY---LIN--W---FSDDLT---
DGLEEDDDDDTPPVITL---A---T---E---DNKRI---GKS---F---I-K-G---Q---KL---FA---N---D---A---T---Y--YVAILNK-TSN---GRIALKYFRQL-QASQLLTNLNKWQ--ET---YS--WE---S---RS---K---FGKSRLRTP---T--
---FHDIL---N--V---S---YGIDRD---R-FLELD---NDNFK---S---QIQKLVASLID--G-K--P---MPQSIVKK-LG---N---N--V---KE---RHRY---RKH---W---
YQVEQVCLAIL---HKQ---N---G---E---E---FSP---MLD---HT---NQ---N---RSYLF---GRLLAIFE---LIE---T---L---RYG---LDGNN---ND---R---ITNA---ERYWTAYT---GQPTKL---MMLLEN--
KI-K-PYEE----PLK--L---NRRGSWMK---L---EKE-K--E---EIL---E----LL----N---P---L---LE-TE-T---MEKPLDYRFIFGYY---A---E---K---NY---Y---Y---T---K---
---QNTEVTES---E---E---
'SUO61003.1' ---MDF---LTSLLKTYE---K---AE---LADL---V---DHQKR---NNEPV---LLPI--YH---TSLKS-NGKNI-ISVKLDK--D--GQ-FH---K-A---E---FM---A---DK---QM---I---I-F-P---VTAD-SV--AR---S-G--
S---H---P--A---P---H---P-P---VDKFAYY---S---AE---M---G---Q---S---Q---Y-D---SF-HK-QLNN--W--IDY---C-EEG--
DVKRF--LTI---VQQFILK---P---EFL---T---L---I---L---D---S---L---IG---P--D-Y---QHN---Q---L---KVTFCDATG---KEK---LID---LSA---C---F---LEF---S-I---DQ---FQG---F---KNES---VS---T---FK---T---L-HQ-SY---I---
S---F---V---EA---NREK--LG---I---CN---ISG---RE-E-Q---LTDK-H--R---GL--M---G-N---A---KIIS---VSN---NDSRLDIVSP---K---R---E-AY--K---G---R---FRE---RE---D---VFS---
--VGYETSEKIHLMLKY-LL--EN---K---N---T---S---TW---L---G---S--SQY---LIN--W---FSDDLT---
DGLEEDDDDDTPPVITL---A---T---E---DNKRI---GKS---F---I-K-G---Q---KL---FA---N---D---A---T---Y--YVAILNK-TSN---GRIALKYFRQL-QASQLLTNLNKWQ--ET---YS--WE---S---RS---K---FGKSRLRTP---T--
---FHDIL---N--V---S---YGVDRD---R-FLELD---NDNFK---S---D---QIQKLVASLID--G-K-S---MPQSIVKK-LG---N---N--V---KE---RHRY---RKH---W---
YQVEQVCLAIL---HKQ---N---G---E---E---FSP---MLD---HT---NQ---N---RSYLF---GRLLAIFE---LIE---T---L---RYG---LDGNN---ND---R---ITNA---ERYWTAYT---GQPTKL---MMLLEN--
KI-K-PYEE----PLK--L---NRRGSWMK---L---EKE-K--E---EIL---E----LL----N---P---L---LE-TE-T---MEKPLDYRFIFGYY---A---E---K---NY---Y---Y---T---K---
---QNTEVTES---E---E---
'VGR14664.1' ---MDF---LTSLLKTYE---K---AE---LADL---V---DHQKR---NNEPV---LLPI--YH---TSLKS-NGKNI-ISVKLDK--D--GQ-FH---K-A---E---FM---A---DK---QM---I---I-F-P---VTAD-SV--AR---S-G--
S---H---P--A---P---H---P-L---VDKFAYY---S---AE---M---G---Q---S---Q---Y-D---SF-HK-QLNN--W--IDY---C-EEG--
DVKRF--LTI---VQQFILK---P---EFL---T---L---I---L---D---S---L---IG---P--D-Y---QHN---Q---L---KVTFCDATG---KEK---LID---LSA---C---F---LEF---S-I---DQ---FQG---F---KNES---VS---T---FK---T---L-HQ-SY---I---
S---F---V---EA---NREK--LG---I---CN---ISG---RE-E-Q---LTDK-H--R---GL--M---G-N---A---KIIS---VSN---NDSRLDIVSP---K---R---E-AY--K---G---R---FRE---RE---D---VFS---
--VGYETSEKIHLMLKY-LL---E---N---K---N---TSTW---L---G---S---S---QYLINW---FSDDLT---
DGLEEDDDDDTPPVITL---A---T---E---DNKRI---GKS---F---I-K-G---Q---KL---FA---N---D---A---T---Y--YVAILNK-TSN---GRIALKYFRQL-QASQLLTNLNKWQ--ET---YS--WE---S---RS---K---FGKSRLRTP---T--
---FHDTL---N--V---S---YGVDRD---R-FLELD---NDNFK---S---D---QIQKLVASLID--G-K--P---MPQSIVKK-LG---N---N--V---KE---RHRY---RKH---W---
YQVEQVCLAIL---HKQ---N---G---E---E---FSP---MLD---HT---NQ---N---RSYLF---GRLLAIFE---LIE---T---L---RYG---LDGNN---ND---R---ITNA---ERYWTAYT---GQPTKL---MMLLEN--
KI-K-PYEE----PLK--L---NRRGSWMK---L---EKE-K--E---EIL---E----LL----N---P---L---LE-TE-T---MEKPLDYRFIFGYY---A---E---K---NY---Y---Y---T---K---
---QNTEVTES---E---E---
'VGQ98935.1__5_' ---MDF---FTSLLKTYE---K---AE---LADL---V---DHQKR---NNEPV---LLPI--YH---TSLKS-NGKNI-ISVKLDK--D--GQ-FH---K-A---E---FM---A---DK---QM---I---I-F-P---VTAD-SV--AR---S-G--
S---H---P--A---P---H---P-L---VDKFAYY---S---AE---M---G---Q---S---Q---Y-D---SF-HK-QLNN--W--IDY---C-EEG--
DVKRF--LTI---VQQFILK---P---EFL---T---L---I---L---D---S---L---IG---P--D-Y---QHN---Q---L---KVTFCDATG---KEK---LID---LSA---C---F---LEF---S-I---DQ---FQG---F---KNES---VS---T---FK---T---L-HQ-SY---I---
S---F---V---EA---NREK--LG---I---CN---ISG---RE-E-Q---LTDK-H--R---GL--M---G-N---A---KIIS---VSN---NDSRLDIVSP---K---R---E-AY--K---G---R---FRE---RE---D---VFS---
--VGYETSEKIHLMLKY-LL---E---NK---N---T---S---TW---L---G---S---S---QYLINW---FSDDLT---
DGLEEDDDDDTPPVITL---A---T---E---DNKRI---GKS---F---I-K-G---Q---KL---FA---N---D---A---T---Y--YVAILNK-TSN---GRIALKYFRQL-QASQLLTNLNKWQ--ET---YS--WE---S---RS---K---FGKSRLRTP---T--
---FHDIL---N--V---S---YGVDRD---R-FLELD---NDNFK---S---D---QIQKLVASLID--G-K--P---MPQSIVKK-LG---N---N--V---KE---RHRY---RKH---W---
YQVEQVCLAIL---HKQ---N---G---E---E---FSP---MLD---HT---NQ---N---RSYLF---GRLLAIFE---LIE---T---L---RYG---LDGNN---ND---R---ITNA---ERYWTAYT---GQPTKL---MMLLEN--
KI-K-PYEE----PLK--L---NRRGSWMK---L---EKE-K--E---EIL---E----LL----N---P---L---LE-TE-T---MEKPLDYRFIFGYY---A---E---K---NY---Y---Y---T---K---
---QNTEVTES---E---E---
'KKC22640.1' ---MDF---FTSLLKTYE---K---AE---LADL---V---DHQKR---NNEPV---LLPI--YH---TSLKS-NGKNI-ISVKLDK--D--GQ-FH---K-A---E---FM---T---DK---QM---I---I-F-P---VTAD-SV--AR---S-G--
S---H---P--A---P---H---P-L---VDKFAYY---S---AE---M---G---Q---T---Q---Y-D---SF-HK-QLNN--W--IDY---C-EEG--
DVKKF--LTF---VQQFILK---P---EFL---T---L---I---L---D---S---L---IG---P--D-Y---QNN---Q---L---KITFCDVTG---KEK---LID---LSA---C---F---LEF---S-I---DQ---FQG---F---KNES---VS---T---FK---A---L-HQ-SY---I---
S---F---V---EA---NREN--LG---I---CN---ISG---RE-E-Q---LTDK-H--R---GL--M---G-N---A---KIIS---VSN---NDSRLDIVSP---K---R---E-AY--K---G---R---FRE---RE---D---VFS---
--VGYETSEKIHLMIKY-LL--EN---K---N---T---S---TW---L---G---S--SQY---LIN--W---FSDDLT---
DGLEEDDDDDTPPVIAL---A---T---E---DNKRI---GKS---F---I-K-G---Q---KL---FA---N---D---A---T---Y--YVAILNK-TSN---GRIALKYFRQL-QASQLLTNLNKWQ--ET---YS--WE---S---RS---K---FGKSRLRTP---T--
---FHDIL---N--V---S---YGVDRD---R-FLELD---NDNFK---S---D---QIQKLVASLID--G-K--P---MPQSIVKK-LG---N---N--V---KE---RHRY---RKH---W---
YQVEQVCLAIL---HKQ---N---G---E---E---FSP---MLD---HT---NQ---N---RSYLF---GRLLAIFE---LIE---T---L---RYG---LDGNN---ND---R---ITNA---ERYWTAYT---GQPTKL---MMLLEN--
KI-K-PYEE----PLK--L---NRRGSWMK---L---EKE-K--E---EIL---E----LL----N---P---L---LE-TE-T---MEKPLDYRFIFGYY---A---E---K---NY---Y---Y---T---K---
---QNTEVTES---E---E---
'VGU05487.1__2_' ---MDF---FTSLLKTYE---K---AE---LADL---V---DHQKR---NNEPV---LLPI--YH---TSLKS-NGKNI-ISVKLDK--D--GQ-FH---K-A---E---FM---A---DK---QM---I---I-F-P---VTAD-SV--AR---S-G--
S---H---P--A---P---H---P-L---VDKFAYY---S---AE---M---G---Q---T---Q---Y-D---SF-HK-QLNN--W--IDY---C-EEG--
DVKKF--LTF---VQQFILK---P---EFL---T---L---I---L---D---S---L---IG---P--D-Y---QHN---Q---L---KVTFCDVTG---KEK---LID---LSA---C---F---LEF---S-I---DQ---FQG---F---KNES---VS---T---FK---A---L-HQ-SY---I---
S---F---V---EA---NREN--LG---I---CN---ISG---RE-E-Q---LTDK-H--R---GL--M---G-N---A---KIIS---VSN---NDSRLDIVSP---K---R---E-AY--K---G---R---FRE---RE---D---VFS---
--VGYETSEKIHLMIKY-LL--E---N---K---N---T---S---TW---L---G---S---SQ---Y---LIN--W---FSDDLT---
DGLEEDDDDDTPPVIAL---A---T---E---DNKRI---GKS---F---I-K-G---Q---KL---FA---N---D---A---T---Y--YVAILNK-TSN---GRIALKYFRQL-QASQLLTNLNKWQ--ET---YS--WE---S---RS---K---FGKSRLRTP---T--
---FHDIL---N--V---S---YGVDRD---R-FLELD---NDNFK---S---D---QIQKLVASLID--G-K--P---MPQSIVKK-LG---N---N--V---KE---RHRY---RKH---W---
YQVEQVCLAIL---HKQ---N---G---E---E---FSP---MLD---HT---NQ---N---RSYLF---GRLLAIFE---LIE---T---L---RYG---LDGNN---ND---R---ITNA---ERYWTAYT---GQPTKL---MMLLEN--
KI-K-PYEE----PLK--L---NRRGSWMK---L---EKE-K--E---EIL---E----LL----N---P---L---LE-TE-T---MEKPLDYRFIFGYY---A---E---K---NY---Y---Y---T---K---
---QNTEVTES---E---E---
'VGT00038.1__5_' ---MKPEF---LTLILDS---L---IAPD--YQ---HNQLKV-T---FCDATGK---EK-LI---DL---
--F-L---EFS---I---D---F-Q---GF-KN---E---SVS---TFKTLHQ---S---A---C---
---S-Y---I---E---A---N-RE-KL---G---
---E---LIN--W---FSDDLT---CN---ISG--R---E--E-Q---LTDK-H--R---G---L---MGNA---KIIS---V---S---NKRE---A---Y-KGR--F---R---ER---ED---VFS---VGYETSEKIHLMLKY-LL--EN---K---N---T---S---TW---
--L---G---S---SQY---LIN--W---FSDDLT---NDSRLDIVSP---I---F---D---DGLEEDDDDDTPPVITL---A---T---E---DNKRI---GKS---F---
---I-K-G---Q---KL---FA---N---D---A---T---Y--YVAILNK-TSN---GRIALKYFRQL-QASQLLTNLNKWQ--ET---YS--WE---S---RS---K---FGKSRLRTP---T---FHDIL---N--V---S---YGVDRD---R-FLELD---NDNFK---S---
---D---QIQKLVASLID--G-K--P---MPQSIVKK-LG---N---KE---RHRY---RKH---W---YQVEQVCLAIL---HKQ---N---
---E---FSP---MLD---HT---NQ---N---RSYLF---GRLLAIFE---LIE---T---L---RYG---LDGNN---ND---R---ITNA---ERYWTAYT---GQPTKL---MMLLEN--KI-K-PYEE---PLK--L---NRRGSWMK---L---EKE-K--E---EIL---E---
--LL----N---P---L---LE-TE-T---MEKPLDYRFIFGYY---A---E---K---NY---Y---Y---T---K---QNTEVTES---E---E---
'VGS32794.1__2_' ---MKTYE---K---AE---LADL---V---DHQKR---SNEPV---LLPI--YH---TSLKS-NGKNI-ISVKLDK--D--GQ-FH---K-A---E---FM---A---DK---QM---I---I-F-P---VTAD-SV--AR---S-G--S---
H---P--A---P---H---P-L---VDKFAYY---S---AE---M---G---Q---T---Q---Y-D---SF-HK-QLNN--W--IDY---C-EEG-DVKKF-
-LTF---VQQFILK---P---EFL---T---L---I---L---D---S---L---IG---P--D-Y---QHN---Q---L---KVTFCDATG---KEK---LID---LSA---C---F---LEF---S-I---DQ---FQG---F---KNES---VS---T---FK---A---L-HQ-SY---I---
F---V---EA---NREN--LG---I---CN---ISG---RE-E-Q---LTDK-H--R---GL--M---G-N---A---KIIS---VSN---NDSRLDIVSP---K---R---E-AY--K---G---R---FRE---RE---D---VFS---
VGYETSEKIHLMIKY-LL--EN---K---N---T---S---TW---L---G---S--SQY---LIN--W---FSDDLT---
DGLEEDDDDDTPPVIAL---A---T---E---DNKRI---GKS---F---I-K-G---Q---KL---FA---N---D---A---T---Y--YVAILNK-TSN---GRIALKYFRQL-QASQLLTNLNKWQ--ET---YS--WE---S---RS---K---FGKSRLRTP---T--
---FHDIL---N--V---S---YGVDRD---R-FLELD---NDNFK---S---D---QIQKLVASLID--G-K--P---MPQSIVKK-LG---N---N--V---KE---RHRY---RKH---W---
YQVEQVCLAIL---HKQ---N---G---E---E---FSP---MLD---HT---NQ---N---RSYLF---GRLLAIFE---LIE---T---L---RYG---LDGNN---ND---R---ITNA---ERYWTAYT---GQPTKL---MMLLEN--
KI-K-PYEE----PLK--L---NLRGSWMK---L---EKE-K--E---EIL---E----LL----N---P---L---LE-TE-T---MEKPLDYRFIFGYY---A---E---K---NY---Y---Y---T---K---
---QNTEVTES---E---E---
'VGT87343.1__2_' ---MDF---FTSLLKTYE---K---AE---LADL---V---DHQKR---SNEPV---LLPI--YH---TSLKS-NGKNI-ISVKLDK--D--GQ-FH---K-A---E---FM---A---DK---QM---I---I-F-P---VTAD-SV--AR---S-G--
S---H---P--A---P---H---P-L---VDKFAYY---S---AE---M---G---Q---T---Q---Y-D---SF-HK-QLNN--W--IDY---C-EEG--
DVKKF--LTF---VQQFILK---P---EFL---T---L---I---L---D---S---L---IG---P--D-Y---QHN---Q---L---KVTFCDATG---KEK---LID---LSA---C---F---LEF---S-I---DQ---FQG---F---KNES---VS---T---FK---A---L-HQ-SY---I---
S---F---V---EA---NREN--LG---I---CN---ISG---RE-E-Q---LTDK-H--R---GL--M---G-N---A---KIIS---VSN---NDSRLDIVSP---K---R---E-AY--K---G---R---FRE---RE---D---VFS---
--VGYETSEKIHLMIKY-LL---E---N---K---N---T---S---TW---L---G---S---SQ---Y---LIN--W---FSD---D--LTNDS---RL--D---I---VSP---I---F---D---D-G-LEEDDDD
---D---TPPVITL---A---T---E---DNKRI---GKS---F---T-K-G---Q---K-LF---AN---D---A---T---Y--YVAILNK-TSN---GRIALKYFRQL-QASQLLTNLNKWQ--ET---YS--WE---S---RS---K---FGKSRLRTP---T---VHDIL---
---N--V---S---YGVDRD---R-FLELD---NDNFK---S---D---QIQKLVASLID--G-K--P---MPQSIVKK-LG---N---N--V---KE---RHRY---RKH---W---YQVEQVCLAIL---
HKQ---N---G---E---E---FSP---MLD---HT---NQ---N---RSYLF---GRLLAIFE---LIE---T---L---RYG---LDGNN---ND---R---ITNA---ERYWTAYT---GQPTKL---MMLLEN--KI-K-PYEE---PLK--L---
NLRGSWMK---L---EKE-K--E---EIL---E----LL----N---P---L---LE-TE-T---MEKPLDYRFIFGYY---A---E---K---NY---Y---Y---T---K---QNTEVTES---
---E---E---

```
'VGQ38155.1' -----MDF------FTSLLKTYE--------K---AE-------LADL----V-------DHQKR-----------NNEPV------------LLPI--YH----TSLKS-NGKNI-ISVKLDK--D---GQ-FH----K-A--------E--------FM------T-----DK-----QM---I-----I-F-P----VTAD-SV--AR----------S-G---
S---------H--------P--A----P---H---------P-L-----VDKFAYY-------S---AE-----M-----G-----Q---------T----------Q-----Y-D---------SF-HK-QLNN--W--IDY-----------C-EEG--
DVKKF--LTF---VQQFILK-----P------EFL-----T------L------I-----L--------D----S------L-----IG---------P--D-Y---QHN----Q---L------KITFCDATG----------KEK----------LID----------LSA------C------F---------LEF-----S-I----------DQ----------FQG---F-------KNES-----VS---------T----FK-----A---------L-HQ-SY-----I-----
S---------F-----------V------EA-----NREN--LG-----I------CN-----ISG------RE-E-Q---LTDK-H--R-----GL--M------G-N---------A-------------KIIS-------VSN----K----R------E-AY---K------G-------R------FRE-----RE-----D--------VFS----
--VGYETSEKIHLMIKY-LL--EN-----K------N------T------S-----TW-----L-----G--------S--SQY-----LIN---W----FSDDLT-----NDSRLDIVSP----------I-------F----------D-----
DGLEEDDDDDTPPVITL-----A--------T----------E----DNKRI-----GKS-----F-----I-K-G----Q------KL--------FA-----N---------D----A-------T-------Y--YVAILNK-TSN----GRIALKYFRQL-QASQLLTNLNKWQ--ET------YS--WE--------S------RS-----K-----FGKSRLRTP----------T--
--------FHDIL-----N---V--------S------YGVDRD-------R-FLELD-------NDNFK------S-----D-------QIQKLVASLID--G-K--P---MPQSIVKK-LG-----------N-----------N--V-----------KE-----RHRY------RKH------W--------
YQVEQVCLAIL----------HKQ-----------N-----G--------E--------E-----FSP----------MLD--------HT-----------NQ----N----------RSYLF------GRLLAIFE-----LIE-------T--------L--------RYG------LDGNN----------ND-----R-------ITNA---ERYWTAYT-----GQPTKL-----MMLLEN--
KI-K-PYEE----PLK--L---NRRGSWMK-----L--------EKE-K--E--------EIL--------E-----LL------N-----P-----L---LE-TE-T---MEKPLDYRFIFGYY---A----E-----K-----NY---------Y-------Y-----T-----K----
-----------QNTEVTES--------E--------E-----
'VGR06631.1__3_' -----MDF------FTSLLKTYE--------K---AE-------LADL----V-------DHQKR-----------NNEPV------------LLPI--YH----TSLKS-NGKNI-ISVKLDK--D---GQ-FH----K-A--------E--------FM------A-----DK-----QM---I-----I-F-P----VTAD-SV--AR----------S-G---
S---------H--------P--A----P---H---------P-L-----VDKFAYY-------S---AE-----M-----G-----Q---------T----------Q-----Y-D---------SF-HK-QLNN--W--IDY-----------C-EEG--
DVKKF--LTF---VQQFILK-----P------EFL-----T------L------I-----L--------D----S------L-----IG---------P--D-Y---QHN----Q---L------KVTFCDATG----------KEK----------LID----------LSA------C------F---------LEF-----S-I----------DQ----------FQG---F-------KNES-----VS---------T----FK-----A---------L-HQ-SY-----I-----
S---------F-----------V------EA-----NREN--LG-----I------CN-----ISG------RE-E-Q---LTDK-H--R-----GL--M------G-N---------A-------------KIIS-------VSN----K----R------E-AY---K------G-------R------FRE-----RE-----D--------VFS----
--VGYETSEKIHLMLRY-LL--EN-----K------N------T------S-----TW-----L-----G--------S--SQY-----LIN---W----FSDDLT-----NDSRLDIVSP----------I-------F----------D-----
DELEEEDDDDTPPVITL-----A--------T----------E----DNKRI-----GKS-----F-----I-K-G----Q------KL--------FA-----N---------D----A-------T-------Y--YVAILNK-TSN----GRIALKYFRQL-QASQLLTNLNKWQ--ET------YS--WE--------S------RS-----K-----FGKSRLRTP----------T--
--------FHDIL-----N---V--------S------YGVDRD-------R-FLELD-------NDNFK------S-----D-------QIQKLVASLID--G-K--P---MPQSIVKK-LG-----------N-----------N--V-----------KE-----RHRY------RKH------W--------
YQVEQVCLAIL----------HKQ-----------N-----G--------E--------E-----FSP----------MLD--------HT-----------NQ----N----------RSYLF------GRLLAIFE-----LIE-------T--------L--------RYG------LDGNN----------ND-----R-------ITNA---ERYWTAYT-----GQPTKL-----MMLLEN--
KI-K-PYEE----PLK--L---NRRGSWMK-----L--------EKE-K--E--------EIL--------E-----LL------N-----P-----L---LE-TE-T---MEKPLDYRFIFGYY---A----E-----K-----NY---------Y-------Y-----T-----K----
-----------QNTEVTES--------E--------E-----
'VGQ84280.1' -----MDF------FTSLLKTYE--------K---AE-------LADL----V-------DHQKR-----------NNEPV------------LLPI--YH----TSLKS-NGKNI-ISVKLDK--D---GQ-FH----K-A--------E--------FM------A-----DK-----QM---I-----I-F-P----VTAD-SV--AR----------S-G---
S---------H--------P--A----P---H---------P-L-----VDKLSYY-------S---AE-----M-----G-----Q---------T----------Q-----Y-D---------SF-HK-QLNN--W--IDY-----------C-EEG--
DVKKF--LTF---VQQFILK-----P------EFL-----T------L------I-----L--------D----S------L-----IG---------P--D-Y---QHN----Q---L------KVTFCDVTG----------KEK----------LID----------LSA------C------F---------LEF-----S-I----------DQ----------FQG---F-------KNES-----VS---------T----FK-----A---------L-HQ-SY-----I-----
S---------F-----------V------EA-----NREN--LG-----I------CN-----ISG------RE-E-Q---LTDK-H--R-----GL--M------G-N---------A-------------KIIS-------VSN----K----R------E-AY---K------G-------R------FRE-----RE-----D--------VFS----
--VGYETSEKIHLMLKY-LL--EN-----K------N------T------S-----TW-----L-----G--------S--SQY-----LIN---W----FSDDLTN-----D--SRLDI---VS--P-----I------F--------D-----DGL-
EEDDDDDDTPPVITL-----A--------T----------E----DNKRI-----GKS-----F-----I-K-G--------QK-------LFA-----N----D----A------T-----Y--YVAILNK-TSN----GRIALKYFRQL-QASQLLTNLNKWQ--ET------YS--WE-----S-----RS-----K------FGKSRLRTP------T-----
----FHDIL-----N---V--------S------YGVDRD-------R-FLELD-------NDNFK------S-----D-------QIQKLVASLID--G-K--P---MPQSIVKK-LG-----------N-----------N--V-----------KE-----RHRY-----RKH-----W--------
YQVEQVCLAIL----------HKQ-----------N-----G--------E--------E-----FSP----------MLD--------HT-----------NQ----N----------RSYLF------GRLLAIFE-----LIE-------T--------L--------RYG------LDGNN----------ND-----R-------ITNA---ERYWTAYT-----GQPTKL-----MMLLEN--
KI-K-PYEE----PLK--L---NRRGSWMK-----L--------EKE-K--E--------EIL--------E-----LL------N-----P-----L---LE-TE-T---MEKPLDYRFIFGYY---A----E-----K-----NY---------Y-------Y-----T-----K----
-----------QNTEVTES--------E--------E-----
'VGR04224.1__2_' -----MDF------FTSLLKTYE--------K---AE-------LADL----V-------DHQKR-----------SNEPV------------LLPI--YH----TSLKS-NGKNI-ISVKLDK--D---GQ-FH----K-A--------E--------FM------A-----DK-----QM---I-----I-F-P----VTAD-SV--AR----------S-G---
S---------H--------P--A----P---H---------P-L-----VDKSAYY-------S---AE-----M-----G-----Q---------T----------Q-----Y-D---------SF-HK-QLNN--W--IDY-----------C-EEG--
DVKKF--LTF---VQQFILK-----P------EFL-----T------L------I-----L--------D----S------L-----IG---------P--D-Y---QHN----Q---L------KVIFCDATG----------KEK----------LID----------LSA------C------F---------LEF-----S-I----------DQ----------FQG---F-------KNES-----VS---------T----FK-----A---------L-HQ-SY-----I-----
S---------F-----------V------EA-----NREN--LG-----I------CN-----ISG------RE-E-Q---LTDK-H--R-----GL--M------G-N---------A-------------KIIS-------VSN----K----R------E-AY---K------G-------R------FRE-----RE-----D--------VFS----
--VGYETSEKIHLMLKY-LL--EN-----K------N------T------S-----TW-----L-----G--------S--SQY-----LIN---W----FSDDLT-----NDSRLDIVSP----------I-------F----------D-----
DGLEEDDDDDTPPVITL-----A--------T----------E----DNKRI-----GKS-----F-----I-K-G----Q------KL--------FA-----N---------D----A-------T-------Y--YVAILNK-TSN----GRIALKYFRQL-QASQLLTNLNKWQ--ET------YS--WE--------S------RS-----K-----FGKSRLRTP----------T--
--------FHDIL-----N---V--------S------YGVDRN-------R-FLELD-------NDNFK------S-----D-------QIQKLVASLID--G-K--P---MPQSIVKK-LG-----------N-----------N--V-----------KE-----RHRY------RKH------W--------
YQVEQVCLAIL----------HKQ-----------N-----G--------E--------E-----FSP----------MLD--------HT-----------NQ----N----------RSYLF------GRLLAIFE-----LIE-------T--------L--------RYG------LDGNN----------ND-----R-------ITNA---ERYWTAYT-----GQPTKL-----MMLLEN--
KI-K-PYEE----PLK--L---NRRGSWMK-----L--------EKE-K--E--------EIL--------E-----LL------N-----P-----L---LE-TE-T---MEKPLDYRFIFGYY---A----E-----K-----NY---------Y-------Y-----T-----K----
-----------QNTEVTES--------E--------E-----
'VGR68296.1' -----MDF------FTSLLKTYE--------K---AE-------LADL----V-------DHQKR-----------NNEPV------------LLPI--YH----TSLKS-NGKNI-ISVKLDK--D---GQ-FH----K-A--------E--------FM------A-----DK-----QM---I-----I-F-P----VTAD-SV--AR----------S-G---
S---------H--------P--A----P---H---------P-L-----VDKFAYY-------S---AE-----M-----G-----Q---------T----------Q-----Y-D---------SF-HK-QLNN--W--IDY-----------C-EEG--
DVKKF--LTF---VQQFILK-----P------EFL-----T------L------I-----L--------D----S------L-----IG---------P--D-Y---QHN----Q---L------KVTFCDATG----------KEK----------LID----------LSA------C------F---------LEF-----S-I----------DQ----------FQG---F-------KNES-----VS---------T----FK-----A---------L-HQ-SY-----I-----
S---------F-----------V------EA-----KREN--LG-----I------CN-----IGG------RE-E-Q---LTDK-H--R-----GL--M------G-N---------A-------------KIIS-------VSN----K----R------E-AY---K------G-------R------FRE-----RE-----D--------VFS----
--VGYETSEKIHLMLKY-LL--EN-----K------N------T------S-----TW-----L-----G--------S--SQY-----LIN---W----FSDDLT-----NDSRLDIVSP----------I-------F----------D-----
DGLEEDDDDDTPPVITL-----A--------T----------E----DNKRI-----GKS-----F-----I-K-G----Q------KL--------FA-----N---------D----A-------T-------Y--YVAILNK-TSN----GRIALKYFRQL-QASQLLTNLNKWQ--ET------YS--WE--------S------RS-----K-----FGKSRLRTP----------T--
--------FHDIL-----N---V--------S------YGVDRD-------R-FLELD-------NDNFK------S-----D-------QIQKLVASLID--G-K--P---MPQSIVKK-LG-----------N-----------N--V-----------KE-----RHRY------RKH------W--------
YQVEQVCLAIL----------HKQ-----------N-----G--------E--------E-----FSP----------MLD--------HT-----------NQ----N----------RSYLF------GRLLAIFE-----LIE-------T--------L--------RYG------LYGNN----------ND-----R-------ITNA---ERYWTAYT-----GQPTKL-----MMLLEN--
KI-K-PYEE----PLK--L---NRRGSWMK-----L--------EKE-K--E--------EIL--------E-----LL------N-----P-----L---LE-TE-T---MEKPLDYRFIFGYY---A----E-----K-----NY---------Y-------Y-----T-----K----
-----------QNTEVTES--------E--------E-----
'VGT96714.1' -----MDF------FTSLLKTYE--------K---AE-------LADL----V-------DHQKR-----------NNEPV------------LLPI--YH----TSLKS-NGKNI-ISVKLDK--D---GQ-FH----K-A--------E--------FM------A-----DK-----QM---I-----I-F-P----VTAD-SV--AR----------S-G---
S---------H--------P--A----P---H---------P-L-----VDKFAYY-------S---AE-----M-----G-----Q---------T----------Q-----Y-D---------SF-HK-QLNN--W--IDY-----------C-EEG--
DVKKF--LTF---VQQFILK-----P------EFL-----T------L------I-----L--------D----S------L-----IG---------P--D-Y---QHN----Q---L------KVTFCDATG----------KEK----------LID----------LSA------C------F---------LEF-----S-I----------DQ----------FQG---F-------KNES-----VS---------T----FK-----T---------L-HQ-SY-----I-----
S---------F-----------V------EA-----NREK--LG-----I------CN-----ISG------RE-E-Q---LTDK-H--R-----GL--M------G-N---------A-------------KIIS-------VSN----K----R------E-AY---K------G-------R------FRE-----RE-----D--------VFS----
--VGYETSEKIHLMLKY-LL--EN-----K------N------T------S-----TW-----L-----G--------S--SQY-----LIN---W----FSDDLT-----NDSRLDIVSP----------I-------F----------D-----
DGLEEDDDDDTPPVITL-----A--------T----------E----DNKRI-----GKS-----F-----I-K-G----Q------KL--------FA-----N---------D----A-------T-------Y--YVAILNK-TSN----GRIALKYFRQL-QASQLLTNLNKWQ--ET------YS--WE--------S------RS-----K-----FGKSRLRTP----------T--
--------FHDIL-----N---V--------S------YGVDRD-------R-FLELD-------NDNFK------S-----D-------QIQKLVASLID--G-K--P---MPQSIVKK-LG-----------N-----------N--V-----------KE-----RHRY------RKH------W--------
YQVEQVCLAIL----------HKQ-----------N-----G--------E--------E-----FSP----------MLD--------HT-----------NQ----N----------RSYLF------GRLLAIFE-----LIE-------T--------L--------RYG------LDGNN----------ND-----R-------ITNA---ERYWTAYT-----GQPTKL-----MMLLEN--
KI-K-PYEE----PLK--L---NRRGSWMK-----L--------EKE-K--E--------EIL--------E-----LL------N-----P-----L---LE-TE-T---MEKPLDYRFIFGYY---A----E-----K-----NY---------Y-------Y-----T-----K----
-----------QNTEVTES--------E--------E-----
'VGQ45781.1' -----MDF------FTSLLKTYE--------K---AE-------LADL----V-------DHQKR-----------NNEPV------------LLPI--YH----TSLKS-NGKNI-ISVKLDR--D---GQ-FH----K-A--------E--------FM------A-----DK-----QM---I-----I-F-P----VTAD-SV--AR----------S-G---
S---------H--------P--A----P---H---------P-L-----VDKFAYY-------S---AE-----M-----G-----Q---------T----------Q-----Y-D---------SF-HK-QLNN--W--IDY-----------C-EEG--
DVKKF--LTF---VQQFILK-----P------EFL-----T------L------I-----L--------D----S------L-----IG---------P--D-Y---QHN----Q---L------KVTFCDATG----------KEK----------LID----------LSA------C------F---------LEF-----S-I----------DQ----------FQG---F-------KNES-----VS---------T----FK-----T---------L-HQ-SY-----I-----
S---------F-----------V------EA-----NREK--LG-----I------CN-----ISG------RE-E-Q---LTDK-H--R-----GL--M------G-N---------A-------------KIIS-------VSN----K----R------E-AY---K------G-------R------FRE-----RE-----D--------VFS----
--VGYETSEKIHLMLKY-LL---------E-----NK-----N-----T-----S-----TW-----L-----G--------S--------S-------QYLINW-------FSD-----------D-------L----TN--D------S------RLD------IV------S---------PI------F----------D-----D-G-LEEDDD-------
------D-D------TPPVITL-----A---------T-----------E----DNKRI-----GKS-----F-----I-K-G-----Q-----K-LF--------AN-----D-----A----T-----Y--YVAILNK-TSN----GRIALKYFRQL-QASQLLTNLNKWQ--ET------YS--WE-------S------RS------K--------FGKSRLRTP----------T-------FHDIL-----
-----N---V--------S------YGVDRD-------R-FLELD-------NDNFK------S-----D-------QIQKLVASLID--G-K--P---MPQSIVKK-LG------N------N--V------KE-----RHRY-----RKH-----W---------YQVEQVCLAIL-------
HKQ--------N-----G-------E--------E-------FSP------MLD------HT---------NQ----N---------RSYLF-----GRLLAIFE-----LIE-------T--------L----------RYG-----LDGNN---------ND----------R--------ITNA---ERYWTAYT-----GQPTKL----------MMLLEN--KI-K-PYEE----PLK--L---
NRRGSWMK-----L-------EKE-K--E--------EIL---------E-----LL------N---------P----------L---LE-TE-T---MEKPLDYRFIFGYY---A----E--------K--------NY------Y-------Y--------T-------K---------QNTEVTES----
----E--------E-----
'VGQ27769.1__3_' -----MDF------FTSLLKTYE--------K---AE-------LADL----V-------DHQKR-----------SNEPV------------LLPI--YH----TSLKS-NGKNI-ISVKLDK--D---GQ-FH----K-A--------E--------FM------A-----DK-----QM---I-----I-F-P----VTAD-SV--AR----------S-G---
S---------H--------P--A----P---H---------P-L-----VDKFAYY-------S---AE-----M-----G-----Q---------T----------Q-----Y-D---------SF-HK-QLNN--W--IDY-----------C-EEG--
DVKKF--LTF---VQQFILK-----P------EFL-----T------L------I-----L--------D----S------L-----IG---------P--D-Y---QHN----Q---L------KVTFCDATG----------KEK----------LID----------LSA------C------F---------LEF-----S-I----------DQ----------FQG---F-------KNES-----VS---------T----FK-----A---------L-HQ-SY-----I-----
S---------F-----------V------EA-----KREN--LG-----I------CN-----ISG------RE-E-Q---LTDK-H--R-----GL--M------G-N---------A-------------KIIS-------VSN----K----R------E-AY---K------G-------R------FRE-----RE-----D--------VFS----
--VGYETSEKIHLMLKY-LL--EN-----K------N------T------S-----TW-----L-----G--------S--SQY-----LIN---W----FSDDLT-----NDSRLNIVSP----------I-------F----------D-----
DGLEEDDDDDTPPVITL-----A--------T----------E----DNKRI-----GKS-----F-----I-K-G----Q------KL--------FA-----N---------D----A-------T-------Y--YVAILNK-TSN----GRIALKYFRQL-QASQLLTNLNKWQ--ET------YS--WE--------S------RS-----K-----FGKSRLRTP----------T--
--------FHDIL-----N---V--------S------YGVDRD-------R-FLELD-------NDNFK------S-----D-------QIQKLVASLID--G-K--P---MPQSIVKK-LG-----------N-----------N--V-----------KE-----RHRY------HKH------W--------
YQVEQVCLAIL----------HKQ-----------N-----G--------E--------E-----FSP----------MLD--------HT-----------NQ----N----------RSYLF------GRLLAIFE-----LIE-------T--------L--------RYG------LDGNN----------ND-----R-------ITNA---ERYWTAYT-----GQPTKL-----MMLLEN--
KI-K-PYEE----PLK--L---NRRGSWMK-----L--------EKE-K--E--------EIL--------E-----LL------N-----P-----L---LE-TE-T---MEKPLDYRFIFGYY---A----E-----K-----NY---------Y-------Y-----T-----K----
-----------QNTEVTES--------E--------E-----
'VHK21194.1' -----MDF------FTSLLKTYE--------K---AE-------LADL----V-------DHQKR-----------SNEPV------------LLPI--YH----TSLKSNGKNI--ISVKLDK--D---GQ-FH----K-A--------E--------FM------A-----DK-----QM---I-----I-F-P----VTADSVA--R---------S---G-
--------S---------HPA----P---H---------P-L-----VDKFAYY---SAE-M---G------Q---------T---------Q---------Y-D---------SF-HK-QLNN--W--IDY-----------CEEG--
DVKKF--LTF---VQQFILK-----P------EFL------T------L------IL--------DS--L-IG-----PD--Y-Q--HNQLK----VTFC---------DATGK--E-K---------L---I------D--------LSA---------C------F---------LEF-----S-I-----------D-----------QFQ---G-----------F---KNESVS---------T----FK------A---------L-HQ-SY------I------
S---------F-------V----E-----ANR----------EN------LG--I-----------CN----------ISG-RE-------E-Q---LTDK-H-----R--GL--M------G----------N--A----------KIIS----V-------------S-------N---KRE-----A------Y-KGR--F----------R-------E------RED-------VFS----
--VGYETSEKIHLMIKY-LL--EN-----K------N------T------S-----TW-----L-----G--------S--SQY-----LIN---W----FSDDLT-----NDSRLDIVSP----------I-------F----------D-----
DGLEEDDDDDTPPVITL-----A--------T----------E----DNKRI-----GKS-----F-----I-K-G----Q------KL--------FA-----N---------D----A-------T-------Y--YVAILNK-TSN----GWIALKYFRQL-QASQLLTNLNKWQ--ET------YS--WE--------S------RS-----K-----FGKSRLRTP----------T-
--------FHDIL-----N---V--------S------YGVDRD-------R-FLELD-------NDNFK------S-----D-------QIQKLVASLID--G-K--P---MPQSIVKK-LG-----------N-----------N--V-----------KE-----RHRY------RKH------W--------
YQVEQVCLAIL----------HKQ-----------N-----G--------E--------E-----FSP----------MLD--------HT-----------NQ----N----------RSYLF------GRLLAIFE-----LIE-------T--------L--------RYG------LDGNN----------ND-----R-------ITNA---ERYWTAYT-----GQPTKL-----MMLLEN--
KI-K-PYEE----PLK--L---NRRGSWMK-----L--------EKE-K--E--------EIL--------E-----LL------N-----P-----L---LE-TE-T---MEKPLDYRFIFGYY---A----E-----K-----NY---------Y-------Y-----T-----K----
-----------QNTEVTES--------E--------E-----
'VGS03764.1__5_' -----MDF------FTSLLKTYE--------K---AE-------LADL----V-------DHQKR-----------SNEPV------------LLPI--YH----TSLKSNGKNI--ISVKLDK--D---GQ-FH----K-A--------E--------FM------A-----DK-----QM---I-----I-F-P----VTADSVA--R---------S---G-
--------S---------HPA----P---H---------P-L-----VDKFAYY---SAE-M---G------Q---------T---------Q---------Y-D---------SF-HK-QLNN--W--IDY-----------CEEG--
DVKKF--LTF---VQQFILK-----P------EFL------T------L------IL--------DS--L-IG-----PD--Y-Q--HNQLK----VTFC---------DATGK--E-K---------L---I------D--------LSA---------C------F---------LEF-----S-I-----------D-----------QFQ---G-----------F---KNESVS---------T----FK------A---------L-HQ-SY------I------
S---------F-------V----E-----ANR----------EN------LG--I-----------CN----------ISG-RE-------E-Q---LTDK-H-----R--GL--M------G----------N--A----------KIIS----V-------------S-------N---KRE-----A------Y-KGR--F----------R-------E------RED-------VFS----
--VGYETSEKIHLMIKY-LL--EN-----K------N------T------S-----TW-----L-----G--------S--SQY-----LIN---W----FSDDLT-----NDSRLDIVSP----------I-------F----------D-----
DGLEEDDDDDTPPVITL-----A--------T----------E----DNKRI-----GKS-----F-----I-K-G----Q------KL--------FA-----N---------D----A-------T-------Y--YVAILNK-TSN----GRIALKYFRQL-QASQLLTNLNKWQ--ET------YS--WE--------SR-----SK------F------G--------
-KSRLR------TP-----TFLID--G-K--P---MPQSIVKK-LG-----N-----N--V------KE-----RHRY-----RKH-----W---------YQVEQVCLAIL-------
H---------K-----Q--------NGEEF----S-----P------MLD------HT---------NQ----N---------RSYLF-----GRLLAIFE-----LIE-------T------LRYGLDG-------NN---ND-----R-------ITNA---ERYWTAYT-----GQPTKL---------MMLLEN--KI-K-PYEE----PLK--L-------
-------------NRRGSWMK---------L-------EKE---------EIL---------E-----LL------N-----P---------L---LE-TE-T---MEKPLDYRFIFGYY---A----E-------K--------NY---------Y-------Y-----T-------K---------QNTEVTESEE
-----------
'VGU48602.1__7_' -----MDF------FTSLLKTYE--------K----AELA----DLVDH---QK-------RSNEPV-----LLPI--YHTSLKSNGKN-I-----ISVKLDK--D---GQ-FH----K---------A--------EF------M--------AD-----------KQ---M-----I-IFP----VTAD-SV--AR---------SGSHP--
-A----------P---H---------P-L-----VDKFAYY----SAEM----G------Q-----T-----Q-----Y-D---------SF-HK-QLNN--W--IDY-----------CEEGD--
---VKK---FLTFVQQ--FILK-----------P-------EFL-----T-------L----------ILDSLIGPDYQHNQL-----------K-VTFCDATGKEKLIDL------SAC---------F-----------LEF-----S-I--------D---Q--------FQGFKNESVS---------T----FK------A---------L-HQ-SY---------I------S--
--------F-----VE-----ANRENLG--------I---------CN-----ISG------R--EEQ---LTDK-H--R--------G----L------MGNA--------KIIS------V------S--------N---KREAY---------K---------G-----RFRER----------ED---------VFS-------
VGYETSEKIHLMIKY-LL---------E-------N------K------N---------TSTW-----L----G--------S--------S--------QYLINW-------FSD----------D-------L----TN--D------S------RLD------IV-----S--------PI------F---------D-----D-G-LEEDDD-------
-------D-D------TPPVITL-----A------T-------E----DNKRI-----GKS-----F------I-K-G----Q------K-LF--------AN-----D-----A----T------Y--YVAILNK-TSN----GRIALKYFRQL-QASQLLTNLNKWQ--ET------YS--WE------S-----RS-----K-----FGKSRLRTP--------T--------FHDIL-------
-----N---V--------S------YGVDRD-------R-FLELD-------NDNFK------S-----D----QIQKLVASLID--G-K--P---MPQSIVKK-LG----N-----N--V------KE-----RHRY-----RKH-----W---------YQVEQVCLAIL-------
HKQ--------N-----G-------E--------E-------FSP------MLD------HT---------NQ----N---------RSYLF-----GRLLAIFE-----LIE-------T--------L----------RYG-----LDGNN---------ND----------R--------ITNA---ERYWTAYT-----GQPTKL----------MMLLEN--KI-K-PYEE----PLK--L---
NRRGSWMK-----L------EKE-K--E--------EIL---------E-----LL------N-----P----------L---LE-TE-T---MEKPLDYRFIFGYY---A----E-------K--------NY------Y-------Y-----T-------K-------QNTEVTES----
----E--------E-----
'VGU23424.1__18_' -----MDF------FTSLLKTYE--------K---AE-------LADL----V-------DHQKR-----------SNEPV------------LLPI--YH----TSLKSNGKNI--ISVKLDK--D---GQ-FH----K-A--------E--------FM------A-----DK-----QM---I-----I-F-P----VTADSVA--R---------S---G-
--------S---------HPA----P---H---------P-L-----VDKFAYY---SAE-M---G------Q---------T---------Q---------Y-D---------SF-HK-QLNN--W--IDY-----------CEEG--
DVKKF--LTF---VQQFILK-----P------EFL------T------L------IL--------DS--L-IG-----PD--Y-Q--HNQLK----VTFC---------DATGK--E-K---------L---I------D--------LSA---------C------F---------LEF-----S-I-----------D-----------QFQ---G-----------F---KNESVS---------T----FK------A---------L-HQ-SY------I------
S---------F-------V----E-----AKR----------EN------LG--I-----------CN----------ISG-RE-------E-Q---LTDK-H-----R--GL--M------G----------N--A----------KIIS----V-------------S-------N---KRE-----A------Y-KGR--F----------R-------E------RED-------VFS----
--VGYETSEKIHLMLKY-LL--EN-----K------N------T------S-----TW-----L-----G--------S--SQY-----LIN---W----FSDDLT-----NDSRLDIVSP----------I-------F----------D-----
DGLEEDDDDDTPPVITL-----A--------T----------E----DNKRI-----GKS-----F-----I-K-G----Q------KL--------FA-----N---------D----A-------T-------Y--YVAILNK-TSN----GRIALKYFRQL-QASQLLTNLNKWQ--ET------YS--WE--------S------RS-----K-----FGKSRLRTP----------T--
--------FHDIL-----N---V--------S------YGVDRD-------R-FLELD-------NDNFK------S-----D-------QIQKLVASLID--G-K--P---MPQSIVKK-LG-----------N-----------N--V-----------KE-----RHRY------RKH------W--------
YQVEQVCLAIL----------HKQ-----------N-----G--------E--------E-----FSP----------MLD--------HT-----------NQ----N----------RSYLF------GRLLAIFE-----LIE-------T--------L--------RYG------LDGNN----------ND-----R-------ITNA---ERYWTAYT-----GQPTKL-----MMLLEN--
KI-K-PYEE----PLK--L---NRRGSWMK-----L--------EKE-K--E--------EIL--------E-----LL------N-----P-----L---LE-TE-T---MEKPLDYRFIFGYY---A----E-----K-----NY---------Y-------Y-----T-----K----
-----------QNTEVTES--------E--------E-----
'VGQ38168.1__2_' -----MDF------FTSLLKTYE--------K---AE-------LADL----V-------DHQKR-----------SNEPV------------LLPI--YH----TSLKSNGKNI--ISVKLDK--D---GQ-FH----K-A--------E--------FM------A-----DK-----QM---I-----I-F-P----VTADSVA--R---------S---G-
--------S---------HPA----P---H---------P-L-----VDKFAYY---SAE-M---G------Q---------T---------Q---------Y-D---------SF-HK-QLNN--W--IDY-----------CEEG--
DVKKF--LTF---VQQFILK-----P------EFL------T------L------IL--------DS--L-IG-----PD--Y-Q--HNQLK----VTFC---------DATGK--E-K---------L---I------D--------LSA---------C------F---------LEF-----S-I-----------D-----------QFQ---G-----------F---KNESVS---------T----FK------A---------L-HQ-SY------I------
S---------F-------V----E-----AKR----------EN------LG--I-----------CN----------ISG-RE-------E-Q---LTDK-H-----R--GL--M------G----------N--A----------KIIS----V-------------S-------N---KRE-----A------Y-KGR--F----------R-------E------RED-------VFS----
--VGYETSEKIHLMLKY-LL---------E-----NK-----N-----T-----S-----TW-----L-----G--------S--------S-------QYLINW-------FSD-----------D-------L----TN--D------S------RLD------IV------S---------PI------F----------D-----D-G-LEEDDD-------
------D-D------TPPVITL-----A---------T-----------E----DNKRI-----GKS-----F-----I-K-G-----Q-----K-LF--------AN-----D-----A----T-----Y--YVAILNK-TSN----GRIALKYFRQL-QASQLLTNLNKWQ--ET------YS--WE-------S------RS------K--------FGKSRLRTP----------T-------FHDIL-----
-----N---V--------S------YGIDRD-------R-FLELD-------NDNFK------S-----D-------QIQKLVASLID--G-K--P---MPQSIVKK-LG------N------N--V------KE-----RHRY-----RKH-----W---------YQVEQVCLAIL-------
HKQ--------N-----G-------E--------E-------FSP------MLD------HT---------NQ----N---------RSYLF-----GRLLAIFE-----LIE-------T--------L----------RYG-----LDGNN---------ND----------R--------ITNA---ERYWTAYT-----GQPTKL----------MMLLEN--KI-K-PYEE----PLK--L---
NRRGSWMK-----L-------EKE-K--E--------EIL---------E-----LL------N---------P----------L---LE-TE-T---MEKPLDYRFIFGYY---A----E--------K--------NY------Y-------Y--------T-------K---------QNTEVTES----
----E--------E-----
'VGU09813.1' -----MDF------FTSLLKTYE--------K---AE-------LADL----V-------DHQKR-----------NNEPV------------LLPI--YH----TSLKSNGKNI--ISVKLDK--D---GQ-FH----K-A--------E--------FM------A-----DK-----QM---I-----I-F-P----VTADSVA--R---------S---G-
--------S---------HPA----P---H---------P-L-----VDKFAYY---SAE-M---G------Q---------I---------Q---------Y-D---------SF-HK-QLNN--W--IDY-----------CEEG--
DVKKF--LTF---VQQFILK-----P------EFL------T------L------IL--------DS--L-IG-----PD--Y-Q--HNQLK----VTFC---------DATGK--E-K---------L---I------D--------LSA---------C------F---------LEF-----S-I-----------D-----------QFQ---G-----------F---KNESVS---------T----FK------A---------L-HQ-SY------I------
S---------F-------V----E-----ANR----------EN------LG--I-----------CN----------ISG-RE-------E-Q---LTDK-H-----R--GL--M------G----------N--A----------KIIS----V-------------S-------N---KRE-----A------Y-KGR--F----------R-------E------RED-------VFS----
--VGYETSEKIHLMLKY-LL--EN-----K------N------T------S-----TW-----L-----G--------S--SQY-----LIN---W----FSDDLT-----NDSRLDIVSP----------I-------F----------D-----
DGLEEDDDDDTPPVITL-----A--------T----------E----DNKRI-----GKS-----F-----I-K-G----Q------KL--------FA-----N---------D----A-------T-------Y--YVAILNK-TSN----GRIALKYFRQL-QASQLLTNLNKWQ--ET------YS--WE--------S------RS-----K-----FGKSRLRTP----------T--
--------FHDIL-----N---V--------Y------YGVDRD-------R-FLELD-------NDNFK------S-----D-------QIQKLVASLID--G-K--P---MPQSIVKK-LG-----------N-----------N--V-----------KE-----RHRY------RKH------W--------
YQVEQVCLAIL----------HKQ-----------N-----G--------E--------E-----FSP----------MLD--------HT-----------NQ----N----------RSYLF------GRLLAIFE-----LIE-------T--------L--------RYG------LDGNN----------ND-----R-------ITNA---ERYWTAYT-----GQPTKL-----MMLLEN--
KI-K-PYEE----PLK--L---NRRGSWMK-----L--------EKE-K--E--------EIL--------E-----LL------N-----P-----L---LE-TE-T---MEKPLDYRFIFGYY---A----E-----K-----NY---------Y-------Y-----T-----K----
-----------QNTEVTES--------E--------E-----
'VGU15888.1__15_' -----MDF------FTSLLKTYE--------K---AE-------LADL----V-------DHQKR-----------NNEPV------------LLPI--YH----TSLKSNGKNI--ISVKLDK--D---GQ-FH----K-A--------E--------FM------A-----DK-----QM---I-----I-F-P----VTADSVA--R---------S---G-
--------S---------HPA----P---H---------P-L-----VDKFAYY---SAE-M---G------Q---------I---------Q---------Y-D---------SF-HK-QLNN--W--IDY-----------CEEG--
DVKKF--LTF---VQQFILK-----P------EFL------T------L------IL--------DS--L-IG-----PD--Y-Q--HNQLK----VTFC---------DATGK--E-K---------L---I------D--------LSA---------C------F---------LEF-----S-I-----------D-----------QFQ---G-----------F---KNESVS---------T----FK------A---------L-HQ-SY------I------
S---------F-------V----E-----ANR----------EN------LG--I-----------CN----------ISG-RE-------E-Q---LTDK-H-----R--GL--M------G----------N--A----------KIIS----V-------------S-------N---KRE-----A------Y-KGR--F----------R-------E------RED-------VFS----
--VGYETSEKIHLMLKY-LL---------E-----NK-----N-----T-----S-----TW-----L-----G--------S--------S-------QYLINW-------FSD-----------D-------L----TN--D------S------RLD------IV------S---------PI------F----------D-----D-G-LEEDDD-------
------D-D------TPPVITL-----A---------T-----------E----DNKRI-----GKS-----F-----I-K-G-----Q-----K-LF--------AN-----D-----A----T-----Y--YVAILNK-TSN----GRIALKYFRQL-QASQLLTNLNKWQ--ET------YS--WE-------S------RS------K--------FGKSRLRTP----------T-------FHDIL-----
-----N---V--------S------YGVDRD-------R-FLELD-------NDNFK------S-----D-------QIQKLVASLID--G-K--P---MPQSIVKK-LG------N------N--V------KE-----RHRY-----RKH-----W---------YQVEQVCLAIL-------
HKQ--------N-----G-------E--------E-------FSP------MLD------HT---------NQ----N---------RSYLF-----GRLLAIFE-----LIE-------T--------L----------RYG-----LDGNN---------ND----------R--------ITNA---ERYWTAYT-----GQPTKL----------MMLLEN--KI-K-PYEE----PLK--L---
NRRGSWMK-----L-------EKE-K--E--------EIL---------E-----LL------N---------P----------L---LE-TE-T---MEKPLDYRFIFGYY---A----E--------K--------NY------Y-------Y--------T-------K---------QNTEVTES----
----E--------E-----
'VGR63267.1__2_' -----MDF------FTSLLKTYE--------K---AE-------LADL----V-------DHQKR-----------SNEPV------------LLPI--YH----TSLKSNGKNI--ISVKLDK--D---GQ-FH----K-A--------E--------FM------A-----DK-----QM---I-----I-F-P----VTADSVA--R---------S---G-
--------S---------HPA----P---H---------P-L-----VDKFAYY---SAE-M---G------Q---------T---------Q---------Y-D---------SF-HK-QLNN--W--IDY-----------CEEG--
DVKKF--LTF---VQQFILK-----P------EFL------T------L------IL--------DS--L-IG-----PD--Y-Q--HNQLK----VTFC---------DATGK--E-K---------L---I------D--------LSA---------C------F---------LEF-----S-I-----------D-----------QFQ---G-----------F---KNESVS---------T----FK------A---------L-HQ-SY------I------
S---------F-------V----E-----ANR----------EN------LG--I-----------CN----------ISG-RE-------E-Q---LTDK-H-----R--GL--M------G----------N--A----------KIIS----V-------------S-------N---KRE-----A------Y-KGR--F----------R-------E------RED-------VFS----
--VGYETSEKIHLMLKY-LL--EN-----K------N------T------S-----TW-----L-----G--------S--SQY-----LIN---W----FSDDLT-----NDSRLDIVSP----------I-------F----------D-----
DGLEEDDDDDTPPVITL-----A--------T----------E----DNKRI-----GKS-----F-----I-K-G----Q------KL--------FA-----N---------D----A-------T-------Y--YVAILNK-TSN----GRIALKYFRQL-QASQLLTNLNKWQ--ET------YS--WE--------S------RS-----K-----FGKSRLRTP----------T--
--------FHDIL-----N---V--------S------YGVDRD-------R-FLELD-------NDNFK------S-----D-------QIQKLVASLID--G-K--P---MPQSIVKK-LG-----------N-----------N--V-----------KE-----RHRY------RKH------W--------
YQVEQVCLAIL----------HKQ-----------N-----G--------E--------E-----FSP----------MLD--------HT-----------NQ----N----------RSYLF------GRLLAIFE-----LIE-------T--------L--------RYG------LDGNN----------ND-----R-------ITNA---ERYWTAYT-----GQPTKL-----MMLLEN--
KI-K-PYEE----PLK--L---NRRGSWMK-----L--------EKE-K--E--------EIL--------E-----LL------N-----P-----L---LE-TE-T---MEKPLDYRFIFGYY---A----E-----K-----NY---------Y-------Y-----T-----K----
-----------QNTEVTES--------E--------E-----
'VHJ64728.1__5_' -----MDF------FTSLLKTYE--------K---AE-------LADL----V-------DHQKR-----------NNEPV------------LLPI--YH----TSLKSNGKNI--ISVKLDK--D---GQ-FH----K-A--------E--------FM------A-----DK-----QM---I-----I-F-P----VTADSVA--R---------S---G-
--------S---------HPA----P---H---------P-L-----VDKFAYY---SAE-M---G------Q---------I---------Q---------Y-D---------SF-HK-QLNN--W--IDY-----------CEEG--
DVKKF--LTF---VQQFILK-----P------EFL------T------L------IL--------DS--L-IG-----PD--Y-Q--HNQLK----VTFC---------DATGK--E-K---------L---I------D--------LSA---------C------F---------LEF-----S-I-----------D-----------QFQ---G-----------F---KNESVS---------T----FK------A---------L-HQ-SY------I------
S---------F-------V----E-----ANR----------EN------LG--I-----------CN----------ISG-RE-------E-Q---LTDK-H-----R--GL--M------G----------N--A----------KIIS----V-------------S-------N---KRE-----A------Y-KGR--F----------R-------E------RED-------VFS----
--VGYETSEKIHLMLKY-LL--EN-----K------N------T------S-----TW-----L-----G--------S--SQY-----LIN---W----FSDDLT-----NDSRLDIVSP----------I-------F----------D-----
DGLEEDDDDDTPPVITL-----A--------T----------E----DNKRI-----GKS-----F-----I-K-G----Q------KL--------FA-----N---------D----A-------T-------Y--YVAILNK-TSN----GRIALKYFRQL-QASQLLTNLNKWQ--ET------YS--WE--------S------RS-----K-----FGKSRLRTP----------T--
--------FHDIL-----N---V--------S------YGVDRD-------R-FLELD-------NDNFK------S-----D-------QIQKLVASLID--G-K--P---MPQSIVKK-LG-----------N-----------N--V-----------KE-----RHRY------RKH------W--------
YQVEQVCLAIL----------HKQ-----------N-----G--------E--------E-----FSP----------MLD--------HT-----------NQ----N----------RSYLF------GRLLAIFE-----LIE-------T--------L--------RYG------LDGNN----------ND-----R-------ITNA---ERYWTAYT-----GQPTKL-----MMLLEN--
KI-K-PYEE----PLK--L---NRRGSWMK-----L--------EKE-K--E--------EIL--------E-----LL------N-----P-----L---LE-TE-T---MEKPLDYRFIFGYY---A----E-----K-----NY---------Y-------Y-----T-----K----
-----------QNTEVTES--------E--------E-----
'VGS85735.1__8_' -----MDF------FTSLLKTYE--------K---AE-------LADL----V-------DHQKR-----------NNEPV------------LLPI--YH----TSLKSNGKNI--ISVKLDK--D---GQ-FH----K-A--------E--------FM------A-----DK-----QM---I-----I-F-P----VTADSVA--R---------S---G-
--------S---------HPA----P---H---------P-L-----VDKFAYY---SAE-M---G------Q---------T---------Q---------Y-D---------SF-HK-QLNN--W--IDY-----------CEEG--
DVKKF--LTF---VQQFILK-----P------EFL------T------L------IL--------DS--L-IG-----PD--Y-Q--HNQLK----VTFC---------DATGK--E-K---------L---I------D--------LSA---------C------F---------LEF-----S-I-----------D-----------QFQ---G-----------F---KNESVS---------T----FK------A---------L-HQ-SY------I------
S---------F-------V----E-----ANR----------EN------LG--I-----------CN----------ISG-RE-------E-Q---LTDK-H-----R--GL--M------G----------N--A----------KIIS----V-------------S-------N---KRE-----A------Y-KGR--F----------R-------E------RED-------VFS----
--VGYETSEKIHLMLKY-LL--EN-----K------N------T------S-----TW-----L-----G--------S--SQY-----LIN---W----FSDDLT-----NDSRLDIVSP----------I-------F----------D-----
DGLEEDDDDDTPPVITL-----A--------T----------E----DNKRI-----GKS-----F-----I-K-G----Q------KL--------FA-----N---------D----A-------T-------Y--YVAILNK-TSN----GRIALKYFRQL-QASQLLTNLNKWQ--ET------YS--WE--------S------RS-----K-----FGKSRLRTP----------T--
--------FHDIL-----N---V--------S------YGVDRD-------R-FLELD-------NDNFK------S-----D-------QIQKLVASLID--G-K--P---MPQSIVKK-LG-----------N-----------N--V-----------KE-----RHRY------HKH------W--------
YQVEQVCLAIL----------HKQ-----------N-----G--------E--------E-----FSP----------MLD--------HT-----------NQ----N----------RSYLF------GRLLAIFE-----LIE-------T--------L--------RYG------LDGNN----------ND-----R-------ITNA---ERYWTAYT-----GQPTKL-----MMLLEN--
KI-K-PYEE----PLK--L---NRRGSWMK-----L--------EKE-K--E--------EIL--------E-----LL------N-----P-----L---LE-TE-T---MEKPLDYRFIFGYY---A----E-----K-----NY---------Y-------Y-----T-----K----
-----------QNTEVTES--------E--------E-----
'VGT66131.1' -----MDF------FTSLLKTYE--------K---AE-------LADL----V-------DHQKR-----------NNEPV------------LLPI--YH----TSLKSNGKNI--ISVKLDK--D---GQ-FH----K-A--------E--------FM------A-----DK-----QM---I-----I-F-P----VTADSVA--R---------S---G-
--------S---------HPA----P---H---------P-L-----VDKFAYY---SAE-M---G------Q---------T---------Q---------Y-D---------SF-HK-QLNN--W--IDY-----------CEEG--
DVKKF--LTF---VQQFILK-----P------EFL------T------L------IL--------DS--L-IG-----PD--Y-Q--HNQLK----VTFC---------DATGK--E-K---------L---I------D--------LSA---------C------F---------LEF-----S-I-----------D-----------QFQ---G-----------F---KNESVS---------T----FK------A---------L-HQ-SY------I------
S---------F-------V----E-----ANR----------EN------LG--I-----------CN----------ISG-RE-------E-Q---LTDK-H-----R--GL--M------G----------N--A----------KIIS----V-------------S-------N---KRE-----A------Y-KGR--F----------R-------E------RED-------VFS----
--VGYETSEKIHLMLKY-LL--EN-----K------N------T------S-----TW-----L-----G--------S--SQY-----LIN---W----FSDDLT-----NDSRLDIVSP----------I-------F----------D-----
DGLEEDDDDDTPPVITL-----A--------T----------E----DNKRI-----GKS-----F-----I-K-G----Q------KL--------FA-----N---------D----A-------T-------Y--YVAILNK-TSN----GRIALKYFRQL-QASQLLTNLNKWQ--ET------YS--WE--------S------RS-----K-----FGKSRLRTP----------T--
--------FHDIL-----N---V--------S------YGVDRD-------R-FLELD-------NDNFK------S-----D-------QIQKLVASLID--G-K--P---MPQSIVKK-LG-----------N-----------N--V-----------KE-----RHRY------RKH------W--------
YQVEQVCLAIL----------HKQ-----------N-----G--------E--------E-----FSP----------MLD--------HT-----------NQ----N----------RSYLF------GRLLAIFE-----LIE-------T--------L--------RYG------LDGNN----------ND-----R-------ITNA---ERYWTAYT-----GQPTKL-----MMLLEN--
KI-K-PYEE----PLK--L---NRRGSWMK-----L--------EKE-K--E--------EIL--------E-----LL------N-----P-----L---LE-TE-T---MENPLDYRFIFGYY---A----E-----K-----NY---------Y-------Y-----T-----K----
-----------QNTEVTES--------E--------E-----
'VGR83021.1__19_' -----MDF------FTSLLKTYE--------K---AE-------LADL----V-------DHQKR-----------NNEPV------------LLPI--YH----TSLKSNGKNI--ISVKLDK--D---GQ-FH----K-A--------E--------FM------A-----DK-----QM---I-----I-F-P----VTADSVA--R---------S---G-
--------S---------HPA----P---H---------P-L-----VDKFAYY---SAE-M---G------Q---------T---------Q---------Y-D---------SF-HK-QLNN--W--IDY-----------CEEG--
DVKKF--LTF---VQQFILK-----P------EFL------T------L------IL--------DS--L-IG-----PD--Y-Q--HNQLK----VTFC---------DATGK--E-K---------L---I------D--------LSA---------C------F---------LEF-----S-I-----------D-----------QFQ---G-----------F---KNESVS---------T----FK------A---------L-HQ-SY------I------
S---------F-------V----E-----ANR----------EN------LG--I-----------CN----------ISG-RE-------E-Q---LTDK-H-----R--GL--M------G----------N--A----------KIIS----V-------------S-------N---KRE-----A------Y-KGR--F----------R-------E------RED-------VFS----
--VGYETSEKIHLMLKY-LL---------E-----NK-----N-----T-----S-----TW-----L-----G--------S--------S-------QYLINW-------FSD-----------D-------L----TN--D------S------RLD------IV------S---------PI------F----------D-----D-G-LEEDDD-------
------D-D------TPPVITL-----A---------T-----------E----DNKRI-----GKS-----F-----I-K-G-----Q-----K-LF--------AN-----D-----A----T-----Y--YVAILNK-TSN----GRIALKYFRQL-QASQLLTNLNKWQ--ET------YS--WE-------S------RS------K--------FGKSRLRTP----------T-------FHDIL-----
-----N---V--------S------YGVDRD-------R-FLELD-------NDNFK------S-----D-------QIQKLVASLID--G-K--P---MPQSIVKK-LG------N------N--V------KE-----RHRY-----RKH-----W---------YQVEQVCLAIL-------
HKQ--------N-----G-------E--------E-------FSP------MLD------HT---------NQ----N---------RSYLF-----GRLLAIFE-----LIE-------T--------L----------RYG-----LDGNN---------ND----------R--------ITNA---ERYWTAYT-----GQPTKL----------MMLLEN--KI-K-PYEE----PLK--L---
NRRGSWMK-----L-------EKE-K--E--------EIL---------E-----LL------N---------P----------L---LE-TE-T---MEKPLDYRFIFGYY---A----E--------K--------NY------Y-------Y--------T-------K---------QNTEVTES----
----E--------E-----
'SQF67432.1' -----MDF------FTSLLKTYE--------K---AE-------LADL----V-------DHQKR-----------DNEPV------------LLPI--YH----TSLKS-NGKNI-ISVKLDK--D---GQ-FH----K-A--------E--------FM------A-----DK-----QM---I-----I-F-P----VTAD-SV--AR----------S-G---
S---------N--------P--A----P---H---------P-L-----VDKFAYY-------S---AE-----V-----D-----Q---------S----------Q-----Y-D---------SF-HK-QLNN--W--IDY-----------C-EEG--
```

DVKRF--LTI---VQQFILK [illegible]

[illegible]

'SUN51496.1__2_' [illegible]

'SUN55799.1' [illegible]

'SUN85468.1__2_' [illegible]

'KXT78394.1' [illegible]

'KXT78402.1' [illegible]

'SUN40056.1' [illegible]

'BAL68715.1' [illegible]

'KIC78181.1' [illegible]

'EHO85750.1' [illegible]

'EFE27917.1' [illegible]

'WP_118017620.1__4_' [illegible]

'WP_002577811.1' [illegible]

'WP_117623301.1__2_' [illegible]

'SCB97534.1__2_' [illegible]

'SCC79659.1' [illegible]

'AGH40905.1' [illegible]

'KFJ02827.1' [illegible]

'AOL10810.1' [illegible]

'AEF28113.1' [illegible]

'EFT83496.1' [illegible]

[illegible]

'EIT88136.1'

[illegible]

'SPU23726.1__2_'

[illegible]

'EEG71147.1'

[illegible]

'WP_094637789.1'

[illegible]

'EDT45691.1'

[illegible]

'ADB10233.1'

[illegible]

'BAN13313.1'

[illegible]

'WP_019673747.1'

[illegible]

'CDI43445.1'

[illegible]

'CDR85526.1'

[illegible]

'WP_096494231.1'

[illegible]

'WP_135293496.1__2_'

[illegible]

'WP_100184380.1__2_'

[illegible]

'WP_012391427.1'

[illegible]

'WP_132014876.1'

[illegible]

'WP_109730132.1'

[illegible]

'WP_081034262.1'

[illegible]

'WP_002594035.1'

[illegible]

'WP_080664529.1'

[illegible]

'WP_118366911.1__2_'

[illegible]

[illegible]

'WP_106764291.1'

[illegible]

'WP_055040323.1'

[illegible]

'WP_007881872.1'

[illegible]

'RKJ63703.1'

[illegible]

'WP_120393510.1__2_'

[illegible]

'WP_051142866.1'

[illegible]

'WP_014826497.1__2_'

[illegible]

'WP_013841448.1'

[illegible]

'WP_007785240.1__2_'

[illegible]

'WP_014184259.1__2_'

[illegible]

'WP_104370683.1'

[illegible]

'WP_106799039.1'

[illegible]

'WP_118682532.1__2_'

[illegible]

'WP_102289221.1'

[illegible]

'WP_083492992.1'

[illegible]

'ERI68076.1'

[illegible]

'WP_135558055.1__2_'

[illegible]

'WP_080926233.1'

[illegible]

'WP_046922791.1'

[illegible]

'WP_117644168.1__2_'

[illegible]

'WP_087152013.1' [illegible]

'WP_087174029.1' [illegible]

'WP_118697434.1__2_' [illegible]

'WP_117760999.1__5_' [illegible]

'WP_117509537.1__2_' [illegible]

'WP_118750568.1__2_' [illegible]

'WP_118666379.1__5_' [illegible]

'WP_117801504.1__2_' [illegible]

'WP_075669168.1' [illegible]

'WP_055303659.1__2_' [illegible]

'WP_118309820.1__2_' [illegible]

'WP_118689190.1__2_' [illegible]

'WP_118724883.1__2_' [illegible]

'WP_117990070.1__2_' [illegible]

'WP_119209846.1__2_' [illegible]

'WP_118397483.1__2_' [illegible]

'WP_118364161.1__2_' [illegible]

'WP_117962135.1__2_' [illegible]

'WP_117890189.1__3_' [illegible]

'WP_117916924.1__2_' [illegible]

'WP_055161652.1__2_' [illegible]

--VRA---IYEYLKK----------S------TLI----------------------------Q------D------L----------------V--------A-H-----K-----I-----IKL----------------N--E-E----NQ-----I---D-------E-K-ENIQ----------------GIP-----------QPK--------A--------F-----------------VRF------I-I-R-------SFDEK---------------IYQ----E--------TPDE-------------------------CW---------K-----DR------T-----------L-QE-CY------------------------------------------E--------------
Y---------V--------------------------------------------------------RS---------------------------------------Q-E--------K--E---KG-----------------------------CY-------------LTG--------NM-E-S---ISYL-H--S--K--KI--R--------------------N-E-G-------DG-A-------------KLIS-----A------------NDS---------------------------------Q-----N------F-TY---R---------G----------------------------R--------FTS------KE------E----------AFA------
VGNETSQKAHNALKW-II-----------R-----------K-----Q-------G------------TF----------------------------F-----------------D------------T---------MTIVTW---------ESN--------Q-------L-------------------S---------------------------MP--------------K----------W--------E----------R---------DTEA----IVS-------------------EYERE-------------YVGNA-ED-D-WED--DW--
-------------------STEEI-V-----SD--------------G-----------N--------P-----ITADK------FYK------A---------------------L-R-G-----Y---------GK--------K-----I---D--------N-------T----S----N--------------M--ILLAFD---AAT-P---GRLALIENQEL-NSVRYLKNIEKWH---ND------CN--WI-----------H-------------------TK------WK---------------D------------GERIQFAGM--------IGVKD---------
IADIL-------------------YG---------------I---------EN-----KGTVSI-------V-DANGK-------KLYAQ--------------------------------------------------------VAQRLLPCIWN---G-T--R---VPYDYVML-AV------------S----------------K--A--------------SN---------------PLAYKERRNWE-------------------RVLT------------------R---------VTNA----KRYMSTFS------QRPYET---------ACSF-------VKKDKKDRY-
PYLN-------K---L--------DIAKRT----------------------------------F-----------------------Y---------------ENL-L---H--------EID--------------EL---------------------------F----------D----K-NTFT-DN-S----RLEGLYLLGFH---N---Q-----------------------------------------------------------------------------------A-------------------YD--------L----K-----------L-------------------------------K-------------------------K-------------
----EK-----------------------------------------------SEEKEESSEE---------------
'WP_117569983.1__4_'    ------------------------------------------------------------------------------------------------MSW---------INELIDLYE------------------K-------NSDK-----I--------GVIEY-R------------GDI------------PYVLLPT---FH-----TTVTA-Q------ITVTIDQ---D---GN-FV----R-A----------E----------QV----DQN-------------------------DK-----------------L----T-------I-I-P-----VTEK-SG--SR---------T-A-----------
---G--------K--E----P---H------------P-L-----------------CDNLRYL--------A------G-------------------------------------------------------------------------------------------------------------------------------------------------------------------D-YKK-----YY---------------------K-----------------------------D---------D-------------GVC--------------S-K----------LY-IS-QLEK---W--VES-----------D-YCH-EK--
--VRA---IYEYLKK-------------S------TLI--------------------------Q------D------L------------------V------A-Y----K-----I-----IKL----------------N--E-E----NQ------I---D-------E-K-ENIQ----------------GIP------------QPK---------A--------F------------------VRF------I-I-R-------SFDEK--------------IYQ----E---------TPDE---------------------CW----------K-----DR------T-----------L-QE-CY--------------------------------------E-----------
Y--------------V----------------------------------------------RS-----------------------------------------Q-E--------K--E---KG------------------------------CY-------------LTG------NM-E-S----ISYL-H--S--K--KI--R---------------------N-E-G---------DG-A-------------KLIS-----A-------------NDS-----------------------------------Q-----N------F-TY----R----------G----------------------------R--------FTS-------KE-------E-----------AFA------
VGNETSQKAHNALKW-II-----------R-----------K------Q--------G----------TF--------------------------F---------------------D---------T---------MTIVTW--------ESN--------Q--------L---------------------S--------------------------MP--------------K-----------W--------E---------R----------DTEA----IVS-------------------EYERE-------------HVGNA-ED-D-WED--DW--
-------------------STGEI-V-----SD-------------G------------N---------P-----ITADK------FYK-------A-----------------L-R-G------Y---------GK--------K-----I----D--------N-------T----S----N---------------M--ILLAFD---AAT-P---GRLALIENQEL-NSVRYLKNIEKWH---ND------CN--WI-------------H-----------------TK-------WK-------------D------------GERIQFAGM-----------VGVKD--------
IADIL-------------------YG----------------I----------EN-----KGTVSI-------V-DANGK--------KLYAQ------------------------------------------------------------VAQRLLPCIWN---G-T--R---VPYDYVML-AV----------S----------------K--A-----------SN---------------PLVYKEQRNWE------------------RVLT-----------------R---------VTNA----KRYMSTFS--------QRPYET------------ACSF------VKK---------
----------D-------------K-------------------------------------------------K--------------DRYK-------E-----------------------------EWN-----------V-------ALD-------------KK----E------TD---RNYLY------GRLLAVAD------RIE-------------YLTYDEK----------------D-----------------------------------------------------NG-------------R----------VTNA----KRYMSTFS--------QRPYET------------WKVIEE----NI-R-
PYLN-------K---L--------DIAKRT----------------------------------F-------------------Y-------------------ENL-L---H--------EID--------------EL---------------------------F----------D----K-STFT-DN-S----RLEGLYLLGFH---N---Q-----------------------------------------------------------------------------------A-------------------YD--------L---K--------L---------------------------------K-------------------------K----------
----EK-----------------------------------------------SEEKEESSEE---------------
'WP_117540731.1__3_'    -----------------------------------------------------------------------------------------------MSW---------INELIDLYE------------------K-------NSDK-----I--------GVIEY-R-------------GDI-------------PYVLLPT---FH-----TTVTA-Q------ITVTIDQ---D---GN-FV----R-A----------E----------QV----DQN---------------------------DK-----------------L----T-------I-I-P-----VTEK-SG--SR---------T-A-----------
---G--------K--E----P---H------------P-L-----------------CDNLRYL--------A------G-------------------------------------------------------------------------------------------------------------------------------------------------------------------D-YKK-----YY---------------------K-----------------------------D---------D-------------GVC--------------S-K----------LY-IS-QLEK---W--VES-----------D-YCH-EK--
--VRA---VYEYLKK-------------S------TLI--------------------------Q------D------L------------------V------A-H----K-----I-----IKL----------------N--E-E----NQ------I---D-------E-K-ENIQ----------------GIP------------QPK---------A--------F------------------VRF------I-I-R-------SFDEK--------------IYQ----E---------TPDE---------------------CW----------K-----DR------T-----------L-QE-CY--------------------------------------E-----------
Y--------------V----------------------------------------------RS-----------------------------------------Q-E--------K--E---KG------------------------------CY-------------LTG------NM-E-S----ISYL-H--S--K--KI--R---------------------N-E-G---------DG-A-------------KLIS-----A-------------NDS-----------------------------------Q-----N------F-TY----R----------G----------------------------R--------FTS-------KE-------E-----------AFA------
VGNETSQKAHNALKW-II-----------R-----------K------Q--------G----------TF--------------------------F---------------------D---------T---------MTIVTW--------ESN--------Q--------L---------------------S--------------------------MP--------------K-----------W--------E---------R----------DTEA----IVS-------------------EYERE-------------HVGNA-ED-D-WED--DW--
IADIL-----------YGI---------ENKGT-------V-----SIVDAN-------GKKLYA----------------L-R-G------Y---------GK--------K-----I----D--------N-------T----S----N---------------M--ILLAFD---AAT-P---GRLALIENQEL-NSVRYLKNIEKWH---ND------CN--WI-------------H-----------------TK-------WK--------DGERIQF--------------------------------ACSF------VKKDKKDRY-
-----------------------------K------------------------------------------------------------------------------E-------------------W------------------N------V------ALD------KK---------ET-----D--------RNYLY------GRLLAVAD------RIE-----------Y-----LTYDEKD-----------------------------------------NG-------------R-----ITNA----KRYMNSFS--------QRPYET------------WKMIEE----NI-R-
PYLN-------K---L-------------------------------DIAKRTF-------------------Y-------------------ENL-L---H--------EID--------------EL---------------------------F----------DKSTFT-DN-S----RLEGLYLLGFH---N---Q-----------------------------------------------------------------------------------A-------------------YD--------LK---L-----------------------------------------------------------------------------------
----EK-----------------------------------------------SEEKEESSEE---------------
'WP_118297498.1__2_'    -----------------------------------------------------------------------------------------------MSW--------VNELIELYE------------------K-------NSDK-----I--------GVIEY-R-------------GDI-------------PYTLLPP---FH-----TTVTA-Q------IVTIDQ---D---GN-FI----N-A----------E----------RV----DPA-------------------------DK-----------------M----T-------I-I-P-----VTEK-SG--SR---------T-A-----------
---G--------K--E----P---H------------P-L-----------------CDNLRYL--------A------G-------------------------------------------------------------------------------------------------------------------------------------------------------------------D-YTK-----YC--------------------K---------------------------D---------D-------------GTC--------------S-Q----------LY-IS-QLEK---W--KES-----------E-YCH-EK--
--VSA---IYSYLKK-----------N------ILI--------------------------R------D------L---------V------E-K-----Q-----V----IKL-------------------D--D-K----QQ------I---D--------E-K-ESIQ----------------GIV-------------QTK--------A--------F-------------------VRF-------I-V-R-------KSALS-----------AVD-----D-------SPDE-----------------------CW---------K-----DR------T-----------L-QE-CY------------------------------------------E-----------
-Y-----------T------------------------------------------------RS-------------------------------------------Q-E--------K--E---KD-------------------------------CY------------LTG-------NI-E-A----ISYL-H--S--K--KI--R--------------------N-E-G---------DG-A-------------KLIS-----A-------------NDS-----------------------------------Q-----N------F-TY----R----------G-----------------------------R--------FAN-------RE-------E-----------AFA------
VGNETSQKIHNALKW-II-----------R-----------K-----Q--------G----------TF-------------------------F--------------------D-----------T---------LTIVTW--------ESN--------Q--------L--------------------S--------------------------MP--------------E-----------W---------N--------E---------DTDK-----IAS-------------------EYDWD---------------ED-D-WDK--GD--
----------------TEGKY-V-----SD-----------G-----------N--------G------ITAEK------FYK-------A-----------------L-R-G------Y---------GK--------K-----V----E--------N-------T----S----N----------------M--ILLFGD---AAT-P---GRLAMVEEKAL-DSAKYLENIGKWH---ES------CS--WI-------------H----------------TK------WK-------------DGKRHTF----------------LGMVGVK--------D-----------
IADIL-----------------FG-------------V----ENNGKLTIMDANGKKLYA--------------------------------------------------------------------------------------E------VAERLLPCIWN---AGYR---IPYDYVER-AV--------------------------L-------------KA---------------------------------TPLAYKDPQNWKCVLT-----------------------------------------L-----------ACSL-------VKK-----Y----
-----------------K------------------------------------------------------------D----------KEQE-----E-------------------------W-----------------N-------V------SLD---------TK---------QK------N--------RNYLY------GRLLAVAD------RIE-----------Y-----------------------------------------------------------------------RTYDEKDRTR-------VTNA----KRYMSTFS--------QRPYET------------WKVIEE----NV-Q-
PYMS-------K---L--------------------------------------------GIAERLF-------------------Y------------------ENL-L---N-------EIY--------------DL-------------------------F----------E----IEQFQ-D-----NSRLDGYLLGFHSQAY--------------------------------------------------------------------------------------------------LK------------M----------------------------------------------------------------K------------
EDAADKKEEE------------------------------------------------------------------------
'WP_118530584.1__2_'    -----------------------------------------------------------------------------------------------MSW--------VNELIDLYE------------------------NSDK-----V--------GVIEY-R-------------GDM--------------PYVLLPP---FH-----TTVTA-Q------IEVTIDR---N---GN-FK-----E-A----------K-----------LV-----SPE----------------------------DK----------------L----T-------V-I-P-----VTEK-SG--SR---------T-A-----------
---G--------K--E----P---H------------P-L-----------------CDNLRYL--------A------E-------------------------------------------------------------------------------------------------------------------------------------------------------------------D-YTK-----YY-------------------------K-----------------------------D---------D-------------GVC--------------Y-Q----------LY-IS-QLKE---W--AES-----------E-FSH-EK-
---VLA---VYQYLKK-----------N------SLI--------------------------R------D------L---------S-------D-Q----K-----I-----IKL-------------N--E-Q----GK----I---D-------D-K-EKLQ----------------GIE-----------QAK---------A--------F-------------------VRF------I-I-R-------PESSG-------------VFE---T--------CPDE---------------------CW----------K-----DR------S-----------L-QE-RY---------------------------------------------------------
Y--------------V----------------------------------------------RS-----------------------------------------R-E--------K--E---QG-------------------------------L-------------CY-------------LTG-------NT-E-A----ISYL-H--S--R--KI--R--------------------N-E-G---------DG-A-------------KLIS-----A-------------NDS-----------------------------------Q-----N------F-TY----R----------G----------------------------R--------FVC-------KE-------E-----------AFA------
VGNETSQKIHNALKW-II-----------R-----------K------Q--------G----------TF--------------------------F---------------------D---------T---------LVIVTW--------ESN--------R--------S-----------------------------------------MP--------------K-----------W--------N---------A---------DTED-----IAS-------------------EYDEE---------------QD-D-WDD--EE--
---------------NDESA-V-----SD---------G-----------N----------A-----ITAEK------FYK-------A-----------------I-R-G------Y---------GQ--------K-----I----E--------N-------T----S----S----------------M--ILLSFD---AAT-P---GRLSMVEEATL-DSAKYLENIKKWH---ES------CG--WI-------------H-----------------TK------WK--------------D-----------KKRVQFYGM--------IGVKD-----------IADIL---
------------FG------------------ES------KGTLTI------V-DANGK-------KMYAE-------------------------------------------------------------------------------VAKRLLPCIWS---G-H--R---IPYDYVNR-AV----------------------K--A---------------------------PLAYKDRNNWE-----------------RVLT----------------------------------------ACSL-------VKK-----W-----------
----------------------K------------K---------------------------------------------------------E-----------------NEQE------E------------------WT--------V------ALD------KE---------ET-----N---------RSYLY------GRLLAVAD------RIE-----------Y--------R-------TYDEKDKA----------------------------R--------MTNA----KRYMSTFS--------QRPFET------------WKIIEE----NI-Q-PYMN----
-K--L-----EFPERR----------------------------F-----------------Y-------------------ENL-L---H--------EIN--------------GL-----------------------F----------D----V-DSFG-DN-S------KLDGLYLLGFH----H---Q-----------------------------------------------------------------------------------------------------YE--------L---T----SKKTKF--------------------N-----------------------------E---------------
KE-------EE---------
'WP_117470372.1__3_'    -----------------------------------------------------------------------------------------------MSW--------VNELIDLYE------------------I-------NSDK-----V--------GIIEY-R-------------GDM--------------PYVLLPP---FH-----TTVTA-Q------IEVTIDR---N---GN-FM-----E-A----------K-----------LV-----SPE----------------------------DK----------------L----T-------V-I-P-----VTEK-SG--SR---------T-A-----------
---G--------K--E----P---H------------P-L-----------------CDNLRYL--------A------G-------------------------------------------------------------------------------------------------------------------------------------------------------------------D-YTK-----YY-------------------------K-----------------------------D---------D-------------GVC--------------Y-Q----------LY-IS-QLKE---W--AES-----------E-FSH-EK-
---VLA---VYQYLKK-----------N------SLI--------------------------R------D------L---------S-------D-Q----K-----I-----IKL-------------N--E-Q----GK----I---D-------D-K-EKLQ----------------GIE-----------QAK---------A--------F-------------------VRF------I-I-R-------PESSG-------------VFE---T--------CPDE---------------------CW----------K-----DR------S-----------L-HE-RY---------------------------------------------------------
Y--------------V----------------------------------------------RS-----------------------------------------R-E--------K--E---QG-------------------------------L-------------CY-------------LTG-------NT-E-A----ISYL-H--S--R--KI--R--------------------N-E-G---------DG-A-------------KLIS-----A-------------NDS-----------------------------------Q-----N------F-TY----R----------G----------------------------R--------FVS-------KE-------E-----------AFA------
VGNETSQKIHNALKW-II-----------R-----------K------Q--------G----------TF--------------------------F---------------------D---------T---------LVIVTW--------ESN--------R--------S-----------------------------------------MP--------------K-----------W--------N---------A---------DTED-----IAS-------------------EYDEE---------------QD-D-WDD--EE--
---------------NDESA-V-----SD---------G-----------N----------A-----ITAEN------FYK-------A-----------------I-R-G------Y---------GQ--------K-----I----E--------N-------T----S----S----------------M--ILLSFD---AAT-P---GRLSMVEEATL-DSAKYLENIKKWH---ES------CG--WI-------------H-----------------TK------WK--------------DKKRVQFY--------------------------G------MVGVKDIA-
-------------------D-------IL-------FGIESK------G-TLTIV------DANGK------RM----------------------------------------------------------------------------VAKRLLPCIWS---G-H--R---IPYDYVNR-AV----------------Q------K--A--------ST-----------PLAYKDRNNWE-----------------RVLT--------------------------------------------ACSL-------VKK-----W-----------
---K--L-----EFPERR---------------------------------------------E------NEQE------E--------------WT--------V------ALD------KE---------ET-----N---------RSYLY------GRLLAVAD------RIE-----------Y--------R-------TYDEKDKA-------------------------------------R------VTNA----KRYMSTFS--------QRPFET------------WKMIEE----NI-Q-PYMY--
-K--L-----EFPERR----------------------------F-----------------Y-------------------ENL-L---H--------EIN--------------GL-----------------------F----------D----V-DSFG-DN-S------KLDGLYLLGFH----H---Q-----------------------------------------------------------------------------------------------------YE--------L---T----SKKTKF--------------------N-----------------------------E---------------
KE-------EE---------
'WP_117484356.1__3_'    -----------------------------------------------------------------------------------------------MSW--------VNELIDLYE------------------I-------NSDK-----V--------GIIEY-R-------------GDM--------------PYVLLPP---FH-----TTVTA-Q------IEVTIDR---N---GN-FM-----E-A----------K-----------LV-----SPE----------------------------DK----------------L----T-------V-I-P-----VTEK-SG--SR---------T-A-----------
---G--------K--E----P---H------------P-L-----------------CDNLRYL--------A------G-------------------------------------------------------------------------------------------------------------------------------------------------------------------D-YTK-----YY-------------------------K-----------------------------D---------D-------------GVC--------------Y-Q----------LY-IS-QLKE---W--AES-----------E-FSH-EK-
---VLA---VYQYLKK-----------N------SLI--------------------------R------D------L---------S-------D-Q----K-----I-----IKL-------------N--E-Q----GK----I---D-------D-K-EKLQ----------------GIE-----------QAK---------A--------F-------------------VRF------I-I-R-------PESSG-------------VFE---T--------CPDE---------------------CW----------K-----DR------S-----------L-QE-RY---------------------------------------------------------
Y--------------V----------------------------------------------RS-----------------------------------------R-E--------K--E---QG-------------------------------L-------------CY-------------LTG-------NT-E-A----ISYL-H--S--R--KI--R--------------------N-E-G---------DG-A-------------KLIS-----A-------------NDS-----------------------------------Q-----N------F-TY----R----------G----------------------------R--------FVS-------KE-------E-----------AFA------
VGNETSQKIHNALKW-II-----------R-----------K------Q--------G----------TF--------------------------F---------------------D---------T---------LVIVTW--------ESN--------R--------S-----------------------------------------MP--------------K-----------W--------N---------A---------DTED-----IAS-------------------EYDEE---------------QD-D-WDD--EE--
---------------NDESA-V-----SD---------G-----------N----------A-----ITAEN------FYK-------A-----------------I-R-G------Y---------GQ--------K-----I----E--------N-------T----S----S----------------M--ILLSFD---AAT-P---GRLSMVEEATL-DSAKYLENIKKWH---ES------CG--WI-------------H-----------------TK------WK--------------DKKRVQFY--------------------------GMV--------GVKDIA-
-------DIL-----FGIES-------K---GTLTIV---DANGKRMYA---------------------------------------------------------------------VAKRLLPCIWS---G-H--R---IPYDYVNR-AV----------------Q-------------KA---------------------S-----------TPLAYKDRNNWERVLT--------------------------------------------ACSL-------VKK-----W-----------
----------------------K------------------------------------------------------------E--------NEQE------E--------------W-----------------------------V------ALD------KE---------ET-----N---------RSYLY------GRLLAVAD------RIE-----------Y--------RTYDEKD----------------------------------------------KA------------R------VTNA----KRYMSTFS--------QRPFET------------WKMIEE----NI-Q-PYMY--
---K--L-----EFPERR---------------------------F-----------------Y-------------------ENL-L---H--------EIN--------------GL-----------------------F----------DVDSFG-DN-S------KLDGLYLLGFH----H---Q-----------------------------------------------------------------------------------------------------YELTSKKTKF--NE-----------------------------------------------------------------------------------
--EE---------
'WP_118600353.1__3_'    -----------------------------------------------------------------------------------------------MSW--------VNELIDLYE------------------I-------NSDK-----V--------GIIEY-R-------------GDM--------------PYVLLPP---FH-----TTVTA-Q------IEVTIDR---N---GN-FK-----E-A----------K-----------LV-----SPE----------------------------DK----------------L----T-------V-I-P-----VTEK-SG--SR---------T-A-----------
---G--------K--E----P---H------------P-L-----------------CDNLRYL--------A------G-------------------------------------------------------------------------------------------------------------------------------------------------------------------D-YTK-----YY-------------------------K-----------------------------D---------D-------------GIC--------------Y-Q----------LY-IS-QLKE---W--AES-----------E-FSH-EK-
---VLA---VYQYLKK-----------N------SLI--------------------------R------D------L---------S-------D-Q----K-----I-----IKL-------------N--E-Q----GK----I---D-------D-K-EKLQ----------------GIE-----------QAK---------A--------F-------------------VRF------I-I-R-------PESSG-------------VFE---T--------CPDE---------------------CW----------K-----DR------S-----------L-QE-RY---------------------------------------------------------
--Y--------------V----------------------------------------------RS-----------------------------------------R-E--------K--E---QG-------------------------------L-------------CY-------------LTG-------NT-E-A----ISYL-H--S--R--KI--R--------------------N-E-G---------DG-A-------------KLIS-----A-------------NDS-----------------------------------Q-----N------F-TY----R----------G----------------------------R--------FVC-------KE-------E-----------AFA------
VGNETSQKIHNALKW-II-----------R-----------K------Q--------G----------TF--------------------------F---------------------D---------T---------LVIVTW--------ESN--------R--------S-----------------------------------------MP--------------K-----------W--------N---------A---------DTED-----IAS-------------------EYDEE---------------QD-N-WDD--EE--
---------------NDESA-V-----SD---------G-----------N----------A-----ITAEK------FYR-------A-----------------I-R-G------Y---------GQ--------K-----I----E--------N-------T----S----S----------------M--ILLSFD---AAT-P---GRLSMVEEATL-DSAKYLENIKKWH---ES------CG--WI-------------H-----------------TK------WK--------------DKKRVQFY--------------------------G------MVGVKDIA-
------------------D-------IL-------FGIESK------G-TLTIV------DANGK------RM----------------------------------------------------------------YA--E----------VAKRLLPCIWS---G-H--R---IPYDYVNR-AV----------------Q------K--A--------ST-----------PLAYKDRNNWE-----------------RVLT--------------------------------------------ACSL-------VKK-----W-----------
---K--L-----EFPERR---------------------------------------------E------NEQE------E--------------WT--------V------ALD------KE---------ET-----N---------RSYLY------GRLLAVAD------RIE-----------Y--------R-------TYDEKDKA-------------------------------------R------MTNA----KRYMSTFS--------QRPFET------------WKMIEE----NI-Q-PYVN--
-K--L-----EFPERR----------------------------F-----------------Y-------------------ENL-L---H--------EIN--------------GL-----------------------F----------D----V-DSFG-DN-S------KLGGLYLLGFH----H---Q-----------------------------------------------------------------------------------------------------YE--------L---T----SKKTKF--------------------N-----------------------------E---------------
-KE-------EE---------
'WP_118736696.1__2_'    -----------------------------------------------------------------------------------------------MSW--------VNELIDLYE------------------T-------NSDK-----V--------GIIEY-R-------------GDM--------------PYVLLPP---FH-----TTVTA-Q------IEVTIDR---N---GN-FK-----E-A----------K-----------LV-----SPE----------------------------DK----------------L----T-------V-I-P-----VTEK-SG--SR---------T-A-----------
---G--------K--E----P---H------------P-L-----------------CDNLRYL--------A------G-------------------------------------------------------------------------------------------------------------------------------------------------------------------D-YTK-----YY-------------------------K-----------------------------D---------D-------------GVC--------------Y-Q----------LY-IS-QLKE---W--AES-----------E-FSH-EK-
---VLA---VYQYLKK-----------N------SLI--------------------------R------D------L---------S-------D-Q----K-----I-----IKL-------------N--E-Q----GK----I---D-------D-K-EKLQ----------------GIE-----------QAK---------A--------F-------------------VRF------I-I-R-------PESSG-------------VFE---T--------CPDE---------------------CW----------K-----DR------S-----------L-QE-RY---------------------------------------------------------
Y--------------V----------------------------------------------RS-----------------------------------------R-E--------K--E---QG-------------------------------L-------------CY-------------LTG-------NT-E-A----ISYL-H--S--R--KI--R--------------------N-E-G---------DG-A-------------KLIS-----A-------------NDS-----------------------------------Q-----N------F-TY----R----------G----------------------------R--------FVS-------KE-------E-----------AFA------
VGNETSQKIHNALKW-II-----------R-----------K------Q--------G----------TF--------------------------F---------------------D---------T---------LVIVTW--------ESN--------R--------S-----------------------------------------MP--------------K-----------W--------N---------A---------DTED-----IAS-------------------EYDEE---------------QD-D-WDD--EE--
---------------NDESA-V-----SD---------G-----------N----------A-----ITAEK------FYK-------A-----------------I-R-G------Y---------GQ--------K-----I----E--------N-------T----S----S----------------M--ILLSFD---AAT-P---GRLSMVEEATL-DSAKYLENIKKWH---ES------CG--WI-------------H-----------------TK------WK--------------DKKRVQFY--------------------------GMV--------GVKDIA-
-------DIL-----FGIES-------K---GTLTIV---DANGKRMYA---------------------------------------------------------------------VAKRLLPCIWS---G-H--R---IPYDYVNR-AV----------------Q-------------KA---------------------S-----------TPLAYKDRNNWERVLT--------------------------------------------ACSL-------VKK-----W-----------
----------------------K------------------------------------------------------------E--------NEQE------E--------------W-----------------------------V------ALD------KE---------ET-----N---------RSYLY------GRLLAVAD------RIE-----------Y--------RTYDEKD----------------------------------------------KA------------R------VTNA----KRYMSTFS--------QRPFET------------WKMIEE----NI-Q-PYVN--
---K--L-----EFPERR---------------------------F-----------------Y-------------------ENL-L---H--------EIN--------------GL-----------------------F----------DVDSFG-DN-S------KLGGLYLLGFH----H---Q-----------------------------------------------------------------------------------------------------YELTSKKTKF--NE-----------------------------------------------------------------------------------
--EE---------
'WP_119245992.1__2_'    -----------------------------------------------------------------------------------------------MSW--------VNELIDLYE------------------K-------NSDK-----I--------GVIEY-R-------------GEI--------------PYVLLPP---FH-----TTVTA-Q------ITVTIDE---K---GN-FI----N-A----------E-----------SV----DSD--------------------------DK----------------L----T-------I-I-P-----VTEK-SG--SR---------T-A-----------
---G--------K--E----P---H------------P-L-----------------CDNLRYL--------A------G-------------------------------------------------------------------------------------------------------------------------------------------------------------------D-YAK-----YY-------------------------K-----------------------------D---------D-------------GVC--------------N-A----------LY-IS-QMEK---W--VKS-----------D-FCH-DK-
--VRA---IYLYLKK-----------G------TLI--------------------------R------D------L---------V-------D-R----R-----I-----IKL-------------N--E-S----NQ----I---D-------D-K-ENIQ----------------GIV-------------QTK--------A--------F-------------------VRF------I-V-R-------FSGIQ-------------SLQ---N--------VPDE---------------------CW----------K-----DR------T-----------L-QE-CY---------------------------------------------------------
Y--------------V----------------------------------------------RS-----------------------------------------Q-E--------K--E---KG-------------------------------M-------------CY-------------LTG-------NT-E-S----ISYL-H--S--K--KI--R--------------------N-E-G---------DG-A-------------KLIS-----A-------------NDS-----------------------------------Q-----N------F-TY----R----------G----------------------------R--------FVS-------RE-------E-----------AFA------
VGDETSQKVHNALKW-II-----------R-----------K------Q--------G----------AF--------------------------F---------------------D---------T---------LAVVTW--------ESN--------R--------L-----------------------------------------MP--------------R-----------W--------N---------A---------DTYS-----ITA----------------EYEDI-----------QDTC-DD-E-WEDDWDE-
----------------KEDNA-V-----SD-----------G------N-------E-----ITAQK------FYR-------A---------------L-R-G------Y---------GR--------K-----I----D--------N-------T----S----S----------------M--ILLGFD---AAT-P---GRLSMVEENTL-TTARYLENIKKWH---ES------CE--WI--------------------HAKWNGEKWIC----------------------------------------------F-------S------GMVGVKD------
VADML----------------FGIENKGKLEIVDKN-----------------------------------------------------------------------------------SKKLYAE-------VARRLLPCIWN---G-H--K---IPYDYVNR-AV------------------------------------I------------KA--------------------------FPLTYKERKNWEKVLT--------------------------------------------ACSL-------VKK-----Y-----------
-----------------K------------K---------------------------------------------------------Q---------ENEQE------E--------------W-----------------------------V------ALD------KK---------AK-----D---------RNYLY------GRLLAVAD------RIE-----------Y--------R-------TYDESD-----------------------------------------------RA------------R------VTNA----KRYMSTFS--------QRPFET------------WKVIEE----SI-Q-
PYMN-------K---L-----------------------------GIAERWF-----------------------Y-------------------ENL-L---N--------DIY--------------DL-----------------------F----------E----KETFC-DN-S-----RLDGLYLLGFH---S---Q-----------------------------------------------------------------------------------------------------Y-DLKVKK-------EN-SEDK-------------------K-------------------------E----------
----------EE-----------
'WP_026649891.1'    -----------------------------------------------------------------------------------------------MSW--------VNELIDLYE------------------K-------NSDK-----I--------GVIEY-R-------------GEI--------------PYVLLPP---FH-----TTVTA-Q------ITVTIDE---K---GN-FI----N-A----------E-----------SV----DSD--------------------------DK----------------L----T-------I-I-P-----VTEK-SG--SR---------T-A-----------
---G--------K--E----P---H------------P-L-----------------CDNLRYL--------A------G-------------------------------------------------------------------------------------------------------------------------------------------------------------------D-YAK-----YY-------------------------K-----------------------------D---------D-------------GVC--------------N-A----------LY-IS-QMEK---W--VKS-----------D-FCH-DK-
--VRA---IYLYLKK-----------G------TLI--------------------------K------N------L---------V-------D-R----R-----I-----IKL-------------N--E-S----NQ----I---D-------D-K-ENIQ----------------GIV-------------QTK--------A--------F-------------------VRF------I-V-R-------FSGIQ-------------SLQ---N--------VPDE---------------------CW----------K-----DR------T-----------L-QE-CY---------------------------------------------------------
Y--------------V----------------------------------------------RS-----------------------------------------Q-E--------K--E---KG-------------------------------M-------------CY-------------LTG-------NT-E-S----ISYL-H--S--K--KI--R--------------------N-E-G---------DG-A-------------KLIS-----A-------------NDS-----------------------------------Q-----N------F-TY----R----------G----------------------------R--------FVS-------RE-------E-----------AFA------
VGDETSQKVHNALKW-II-----------R-----------K------Q--------G----------AF--------------------------F---------------------D---------T---------LAVVTW--------ESN--------R--------L-----------------------------------------MP--------------R-----------W--------N---------A---------DTAS-----ITA----------------EYENV-----------QDEDEWDD-E-
WEDDWDE---------------KEDNK-V-----SD-----------G------N-------E-----ITAQK------FYR-------A---------------L-R-G------Y---------GK--------K-----I----D--------N-------T----S----F----------------M--MLLGFD---AAT-P---GRLSMVEEKTL-DTARYLENIKKWH---ES------CE--WI--------------------HEKKIGEKWI----------------------------------------RFSGMV-------
-G--------VKDVA-------------------DI---------FGIENK----GKLEIVDKN-------------------------SKKLYAE-------AAERLIPCIWN---G-H--K---IPYDYVNR-AV------------------------------------I------------KA--------------------------SPLSYKERKNWERVLT--------------------------------------------ACSL-------
VKK-----Y-----------K------------K--------------------------------------------------ENEQE------E--------------W---------------------S--------V------ALD------KK---------ET-----D---------RSYLY------GRLLAVAD------RIE-----------Y--------R-------TYDEGDKA-------------------------------------R------VTNA----KRYMSTFS--------QRPFET------------
WKVIEE--SI-Q-PYMN-----K---L------------------------------------GIAERRF-----------------------Y-------------------ENL-L---D--------EIC--------------DL---------------------------F----------E----KETFLD-NN-A-----NKRLDGLYLLGFH---S---Q-----------------------------------------------------------------------------------------------------------YD---L------------------K--------------------------------------------
----K------IKKENSEDKKEEE-
'WP_092072106.1'    -----------------------------------------------------------------------------------------------MSW--------VNELIDLYE------------------I-------NSDK-----I--------GVIEY-R-------------GEM--------------PYVLLPP---FH-----TTVTA-Q------IEVTIDQ---D---GN-FM-----N-A----------E-----------PV----DLN--------------------------DK----------------L----T-------V-I-P-----VTEK-SG--SR---------T-A-----------
---G--------K--E----P---H------------P-L-----------------CDNLRYL--------A------G-------------------------------------------------------------------------------------------------------------------------------------------------------------------D-YTK-----YY-------------------------K-----------------------------D---------D-------------GIC--------------H-E----------LY-IS-QLKE---W--VES-----------E-YCH-EK-
--VRA---IYLYLKK-----------N------TLI--------------------------S------D------L---------V-------D-K----K-----I-----IKL-------------D--E-Q----NQ----I---D-------D-K-ENIQ----------------GIV-------------QTK--------A--------F-------------------VRF------I-I-R-------SSNSD-------------LFE---L--------KTDE---------------------CW----------K-----DR------S-----------L-QD-CY---------------------------------------------------------
Y--------------V----------------------------------------------RS-----------------------------------------R-E--------K--E---KG-------------------------------L-------------CY-------------LTG-------NM-E-A----ISYL-H--S--K--KI--R--------------------N-E-G---------DG-A-------------KLIS-----A-------------NDS-----------------------------------Q-----N------F-TY----R----------G----------------------------R--------FVS-------KE-------E-----------AFA------
VGNETSQKIHNALKW-II-----------R-----------K------Q--------G----------TF--------------------------F---------------------D---------T---------LAVVTW--------ESN--------R--------L-----------------------------------------MP--------------R-----------W--------N---------A---------DTEM-----ITS----------------EYA------------WDD-EE--
---------------PEEET-I-----SD-----------G------N-------A-----VTAEK------FYK-------A---------------L-R-G------Y---------GK--------K-----V----D--------N-------T----S----S----------------M--ILLGFD---AAT-P---GRLSMVEEMTL-DSARYLENIKKWH---ES------CE--WI---------------------------EK--------------------------------------------KGKERIEFS------GMVGVKD--------VAEIL-----
---------------FG------------I--------ES------KGGLAI------A-DANGK------KMYAE--------------------------------------------------------------------------VARRLLPCIWN---G-T--E---IPHDYVNR-AV----------------L------K--A--------SN-----------PLTYKDRKNWE-----------------RVLT--------------------------------------------ACSL-------VKK-------------
--------------D-------------R------------------------K---------------------------------------MEEW------N--------------------------------V------ALD------KN---------Q---------KD--------RNYLY------GRLLAVAD------RIE-----------Y--------TYDDGDKA-------------------------------------R------VTNA----KRYMSTFS--------QRPFET------------WKVIEE----NI-Q-PYMN----K--
-L-------------------------------------GIAERR--------------F-----------------Y-------------------ENL-L---N--------DIY--------------EL-----------------------F----------E----I-DSFK-DN-K------KLDGLYLLGFH---S---Q-----------------------------------------------------------------------------------------------------YD--------L---K--------------------------------------------------------EDSKEKEE-
-----EE-------------
'WP_118254297.1__2_'    -----------------------------------------------------------------------------------------------MSW--------VNELIDLYE------------------K-------NSDK-----I--------GVIEY-R-------------RDM--------------PYVLLPP---FH-----TTVTA-Q------ITVTIDQ---D---GN-FM-----N-A----------E-----------PV----DSN--------------------------DK----------------L----T-------I-I-P-----VTEK-SG--SR---------T-A-----------
---G--------K--E----P---H------------P-L-----------------CDNLRYL--------A------G-------------------------------------------------------------------------------------------------------------------------------------------------------------------D-YTK-----YY-------------------------K-----------------------------D---------D-------------GIC--------------H-E----------LY-IS-QLKE---W--VES-----------E-YCH-EK-
--VRA---IYLYLKK-----------N------TLI--------------------------S------N------L---------I-------E-K----K-----I-----IKL-------------N--E-Q----NR----I---D-------D-K-ENIQ----------------GIV-------------QTK--------A--------F-------------------VRF------I-I-R-------LPNTD-------------FFE---A--------PPDE---------------------CW----------K-----DK------T-----------L-QD-CY-------------------------------------------------------Y-
Y--------------V----------------------------------------------RS-----------------------------------------Q-E--------K--E---IG-------------------------------L-------------CY-------------LTG-------NT-E-A----ISYL-H--S--K--KI--R--------------------N-E-G---------DG-A-------------KLIS-----A-------------NDS-----------------------------------Q-----N------F-TY----R----------G----------------------------R--------FAT-------KE-------E-----------AFA------
VGNETSQKIHNALKW-II-----------R-----------K------Q--------G----------TL--------------------------F---------------------D---------T---------LAVVTW--------ESH--------R--------L-----------------------------------------MP--------------R-----------W--------N---------A---------DTDT-----IAS----------------EYNE-----------DD-D-WDD--EE--
---------------SGEES-G-----CD-----------G------S-------A-----ITAEK------FYS-------A---------------L-R-G------Y---------GK--------K-----V----D--------N-------T----S----E----------------M--ILLGFD---AAT-P---GRLSMVEEMTL-DSARYLENIKKWH---ES------CS--WI-------------H----------------EK------WK---------EKDKKRIRF--------------GMVGVKD-------
IADIL------------FGIENKGSLMIVDANGKKL-------------------------------------------------------------------------------YA--E----------VAKRLIPCIWN---G-N--K---IPCDYVDR-AV----------------------K--A-----------S---------------MPLTYKDWKNWERVLT--------------------------------------------ACSL-------VKKDKSERY-
-----------------K------------K----------------------------------------------------E-----------------E-------------------W-----------------------------V------ALD------KK---------QK-----D---------RNYLY------GRLLAVAD------RIE-----------Y--------TYDDGD------------------------------------------------------------R-------VTNA----KRYMCTFS--------QRPFET------------WKVIEE----SI-Q-
PYVN-------K---L-----------------------------GIAERRF-----------------------Y-------------------ENL-L---N--------DIY--------------DL-----------------------F----------E----VESFS-DN-S------KLDGLYLLGFH---S---Q-----------------------------------------------------------------------------------------------------Y-DLKQKK-------ED-SKEK-------------------K-------------------------E----------
----------EE-----------
'WP_117942785.1__2_'    -----------------------------------------------------------------------------------------------MSW--------VNELIDLYE------------------I-------NSDK-----I--------GVIEY-R-------------RDM--------------PYVLLPP---FH-----TTVTA-Q------ITVTIDQ---D---GN-FM-----N-A----------E-----------LV----DLN--------------------------DK----------------L----T-------I-I-P-----VTEK-SG--SR---------T-A-----------
---G--------K--E----P---H------------P-L-----------------CDNLRYL--------A------G-------------------------------------------------------------------------------------------------------------------------------------------------------------------D-YTK-----YY-------------------------K-----------------------------D---------D-------------GIC--------------H-E----------LY-IS-QLKE---W--VES-----------G-YCH-EK-
--VRA---IYLYLKK-----------N------TLV--------------------------S------D------L---------V-------K-K----R-----I-----IKL-------------N--E-Q----NQ----I---D-------D-K-ENIQ----------------GIV-------------QTK--------A--------F-------------------VRF------I-I-R-------SSNTD-------------LSE---I--------IPDE---------------------CW----------K-----DK------T-----------L-QD-CY---------------------------------------------------------
Y--------------V----------------------------------------------RS-----------------------------------------Q-E--------K--E---KG-------------------------------L-------------CY-------------LTG-------NT-E-A----ISYL-H--S--K--KI--R--------------------N-E-G---------DG-A-------------KLIS-----A-------------NDS-----------------------------------Q-----N------F-TY----R----------G----------------------------R--------FVS-------KE-------E-----------AFA------
VGNETSQKIHNALKW-II-----------R-----------K------Q--------G----------TF--------------------------F---------------------D---------T---------LAVVTW--------ESH--------R--------L-----------------------------------------MP--------------R-----------W--------N---------A---------DTDT-----IAS----------------EYAG-----------DD-D-WDD--EE--
---------------SEEES-A-----CD-----------G------N-------A-----ITAEK------FYS-------A---------------L-R-G------Y---------GK--------K-----V----D--------N-------T----S----K----------------M--ILLGFD---AAT-P---GRLSMVEEMTL-DSARYLENIKKWH---ES------CN--WI-------------H----------------EK------WK---------E---KD-----------------KRIQFSGMV----GVKD-----------
VKDVA----------------D-------ILFGTE-----S-KGGLA-------IADTN------G-----KKLYA--------------------------------------------E----------VARRLLPCIWN---G-N--R---IPYDYVNRAV------------------K--------AS----M------------PL-----------IYKDRKNWE-----------------------------------------------------RVLT---------ACSL-------VKK--------
-----------------K------------S----------------------------------------------------E-----------------R---G-------------------EWN-----------------------------V------ALD------KK---------QK-----D---------RNYLY------GRLLAVAD------RIE-----------Y--------R-------TYDDGDKA-------------------------------------R------VTNA----KRYMSTFS--------QRPFET------------WKVIEE----SI-
Q-PYMN-----K---L-----GIAERR--------------F-----------------Y-------------------ENL-L---N--------DIY--------------E-----------------------L-----F----------E----S-SFK-DN-K------KLDGLYLLGFH---S---Q-----------------------------------------------------------------------------------------------------YD--------L---K--------------------------------------------------------QKKEDSKE-
----------------KE------EE-----------
'WP_118648142.1__2_'    -----------------------------------------------------------------------------------------------MSW--------ADELCELYE------------------K-------NKHH-----A--------GVMEY-K------P-VN-TKKGVEMI-------PYVLLPP---FH-----TTVTA-Q------ITVTIDE---H---GN-FL-----Q-A----------E-----------PV----DNG--------------------------DK----------------L----T-------I-I-P-----VTEK-SG--SR---------T-A-----------T-
A-----------S-------K--E----P---H------------P-F-----------------CDNLKYV--------A------G-------------------------------------------------------------------------------------------------------------------------------------------------------------------D-YVK-----YS-------------------------D-----------------------------E---------D-------------S---T----GDAAY-------------Y-E----------LY-IQ-ALEK---W--HTS-----------
L-YTH--KK-----VDA---LYLYLEK-------------G------CLI---------------------------R------D------L---------A-------A-A-H----V-----LKL-------------D--E-N----GK----L--S-------K-ENIQ------------------NIP---------QSK--------A--------F----------------VRF------R-I-METFS---DVTSV-------------AAG---Q-------YRSE-------------------------------CW----------D-----DQ------T-----------L-QN-SF---------------------------------------------------------
I-----------E------------------Y---------------------RS-----------------------------------------L-A--------D--S---KG----------------------------CY-------------LSG-------HE-E-Q----ISYL-H--S--K--KI--R--------------------N-E-G---------DG-A-------------KLIS-----C-------------NDS-----------------------------------E-----N------F-TY----R----------G----------------------------R--------FST-------KE-------
Q----------AFA-------IGNETSQKVHNALKW-II---R---K-Q-----------------------------------------------F---------------------D---------T---------L-T--------IVT-W--------ETN-----------------------------------------------MP--------------R-----------W--------N---------A---------DTDE-----ICS--------------------DY-----------D-G-FE-
FWGMV----------------GIRD-------IADIL---YGTESN---GMLTILDRS--------------------------------------------------------------------------------------------------------------------------------------------------------------------------------------
--------------------------------------------------------------------------------HEKNHE----E------------V----------WT------------V------ALD------KD---------QK-----D---------RNYLY------GRLLAVAD------RVE-----------Y--------N----------------------------------------------------------------TFDPEKDDSR----TTNA---RRYMTAFS--------QRPYET----------------
WKVVEE--NL-L-PYYN------K--L-----------------------------KRSERLH------------------------Y-------------------QRL-M---D--------EIC--------------QL--------------------------F-----------D----VQSFS-DN-S----NSRLDPLYLLGFHSQSH-----------------------------------------------------------------------------------------------------------KE------------Y--------------K-----------------------------------------
--------N--------------TKSEED---------------------------------------------------------------------
'WP_095174792.1'    -----------------------------------------------------------------------------------------------MDW--------VSELCDLYE------------------K-------NKYL-----A--------GEMRY-K------E-QI-RKNGVEKT-------PLVLLPP---FH-----TTVTA-Q------ITVTISE---N---GE-FQ-----D-A----------K-----------LV----ENE--------------------------DK----------------L----T-------I-I-P-----VTEN-SG--SR---------T-
---------S----------T--------V---P---H------------P-L-----------------CDNIKYL--------S------G---------------------------------------------------------------------------------------------------------------------------------------------------------------------D-YKD-----YC---------------------------------------------E---------G-----------------T---K----NAEEC--------------Y-Q----------LY-IE-ALRR---W--HIS-----------P-
YSH--KK-----VDA---LYLYLSK-------------G------CLI---------------------------H------D------L---------V-------D-A-G----K-----LKL-------------G--E-N----GK----L--S-------G-S-VKIQ------------------GIE---------QDK---------S--------F-------------------VRF------R-I-LEGYQENEDILSD-------------TTG---K-------YYSE-------------------------------CW----------L-----DR------T-----------L-QE-SF---------------------------------------------------------
----Y------------Y----------V-------------------------------------------------------RS------------------------L-M------T--E---KD----------------------------CY-------------LSG-------NT-E-Q----PTYL-H--S--K--KI--R--------------------N-E-G---------DG-A-------------KLIS-----S-------------NDS-----------------------------------Q-----N------Y-T-----R----------G----------------------------R--------FHN-------KE-------
Q--------AFA----IGNETSQKVHNALKW-II---R---K-Q------------------------------------------------------------------------------D---------T---------FC--------IVS-W--------ESD---------------------------------------------------------------------------------------------------------------------------DTDS-----ICS----------------QY------------EE-D-
WG--EE------------EEGIP-V-----WD-----------G------N-------Q-----AGALR------FNA-------A---------------L-L-G------Y---------GR--------K-----P-----G--------N-------M----S-----R------------M--VLMAFD---AAT-T---GRLALVEYKSL-ESTRYLDNIKYWH---RE------CE--WL-------------D--------------------LK-------------------------------LKDGGAYY--------------------YKGMA-
-G--------VKEIA-------------------DI---------I---YGTEQNGILSLGNKSKLY--------------------------------------------------------IYRRLIPCIWD---R-M-D---VPFDMVDL-AV------------------------------------A-------------------S-----------SPVSYEKRYNWERVLTS------------------------------------------------ACSL-------
VKK----------------K-------------QQNPK------E--------------------------------------------------------------V--------------------------------------WT------------L-------ALN------KK---------SK-----D---------RNYLY------GRLLAVAD------RVE-----------Y--------------------------------------------------------------------------------------RT-----FDKRDRDANR-------ETNA----RRYMSMFS--------QRPYET------
--WKIIEE--SL-A-PYFP------K---L-------------------------------------------------------------KYQEKIY---------------Y-------------------CKQ-L---E--------EIC--------------EL-----------------------F----------DE-ETFKDNSKLDGLYLLGFH---S---E-----------------------------------------------------------------------------------------------------YE--------L----------------------------------------------------------
-N--------KEADKEHDIISEEE---------------------------------------------------------------------------------------------------------------------------------------------------------------------------------------------------------------------------
'WP_115624284.1'    -----------------------------------------------------------------------------------------------MDW--------VSELCDLYE------------------K-------NKYL-----A--------GEMRY-K------E-QI-RKNGLEKI-------PLVLLPP---FH-----TTVTA-Q------ITVTISE---S---GE-FQ-----D-A----------K-----------LV----ENE--------------------------DK----------------L----T-------I-I-P-----VTEN-SA--SR---------T-S-
---------T-------V---V---P---H------------P-L-----------------CDNIKYL--------S------G---------------------------------------------------------------------------------------------------------------------------------------------------------------------D-YKD-----YC---------------------------------------------E---------G-----------------T---K----NAEEC--------------Y-Q----------LY-IE-ALRR---W--HIS-----------
YSH--KK-----VDA---LYLYLSK-------------G------CLI---------------------------W------D------L---------V-------N-E-G----K-----LKL-------------G--E-N----GK----L--S-------E-S-VKIQ------------------GIE---------QDK---------S--------F-------------------VRF------R-I-LEEYQEDEDILSD-------------ITG---Q-------YYSE-------------------------------CW----------L-----DR------T-----------L-QE-RF---------------------------------------------------------
----Y----------------V-------------------------------------------------------RS-----------------------------L-M------T--E---KD----------------------------CY-------------LSG-------NI-E-Q----PTYL-H--S--K--KI--R--------------------N-E-G---------DN-N-------------KLIS-----S-------------NDK-----------------------------------Q-----N------Y-T-----R----------G----------------------------R--------FHN-------NE-------
Q--------AFA----IGNETSQKVHNALKW-II---R---R-Q------------------------------------------------------------------------------D---------T---------FC--------IVS-W--------ESD---------------------------------------------------------------------------------------------------------------------------DTDS-----ICS----------------QY------------EE-D-
WG--EE------------EEEVQ-V-----WD-----------G------N-------Q-----AGAFQ------FSA-------A---------------L-L-G------Y---------RR--------K-----P-----G--------N-------M----S-----R------------M--VLMAFD---AAT-T---GRLALVEYKNL-ETSRYLDNIKYWH---KE------CE--WL-------------D--------------------LK-------------------------------KEGGSYYYKGMV-
-G--------VKEIA-------------------D------IIYGTE----QNGILSLGN------------------------------------------------------------KSKLYGQ-------IYRRLIPCIWD---Q-M-D---VPFDMVNL-AV------------------------------------A-------------------S-----------PVSYEKRYNWERTLS------------------------------------------------ACSL-------
VKK----------------M--------------K---------------------------QNPK---------E--------------------------------------------------WT------------L-------ALN------RK---------SR-----D---------RNYLY------GRLLAVAD------RVE-----------Y--------------------------------------------------------------------------------------RT-----FDKKDRDANR-------ETNA----RRYMSMFS--------QRPYET------
--WKIIEE--SL-A-PYFS------K---L-------------------------------------------------------------KYQEKIY---------------Y-------------------CKQ-L---E--------EIC--------------EL-----------------------F----------DE-ETFKDNSKLDGLYLLGFH---S---E-----------------------------------------------------------------------------------------------------YE--------L----------------------------------------------------------
-N--------KEADKAQDITSEEE---------------------------------------------------------------------------------------------------------------------------------------------------------------------------------------------------------------------------
'WP_021636853.1'    -----------------------------------------------------------------------------------------------MNW--------IAELCDLYD------------------K-------NQHL-----A--------GRMGEDH-----------IVLLPP---YH-----TTVAA-Q------ITVTIDG---D---GN-FL-----R-A----------E-----------TV----PDE--------------------------DK----------------L----T-------I-I-P-----VTEK-SA--SR---------T-A-----------
---G--------I--E----P---H------------P-L-----------------CDNLKYL--------A------A-------------------------------------------------------------------------------------------------------------------------------------------------------------------D-YNQ-----YV---------------------------------------------------------------------KG-K----DCSKN--------------H-L----------LY-ME-QLKK---W--ADS-----------S-YCH-
SK--------VKA-----VYQYLEK-----------G------TLM-----------------------------------K------N-------N--------N----Q--G----I---FIQ-------------D--E-S----GK----I---S-------E-K-AKIQ------------------IVA----------QTD--------A--------F-------------------IRF------Q-I-DTGEVLPEDALMD-------------DSG---S-------LSVK-------------------------------CW----------K-----DC------S-----------L-QQ-NY---------------------------------------------------------
---------E----------Y----------------------------------------------RS------------------------------------------Q-Q-----K--E---TG-------------------------------------------------------------SY-------------LTG-------DL-T-Q----ISWL-Q--P--K--KI--R--------------------N-E-G---------DG-A-------------KLIS-----A-------------NDE-----------------------------------S-----N------N-TY----R----------G----------------------------R--------FAN-------KE-------E-----------AFA------
--AFS-------IGYEDSQKAHNALKW-LL-----------R-------Q-------G----------TT-------------------------------------------------------------LVT-W-----ESD------------------------------------------------------------------------------------------------------------------------------------------SSDE---LCI-----QAE-----------ALPDEVSEEE---------
E-FSDEVWD-------------EEEEQ-Y-----KG-----------T------E-------G------AGAMR------FRQ-------A---------------I-K-G------Y---------RK--------N-----L-----T--------Q-------T----S-----S----------------T--MIMAFD---AAT-T---GRLAMVECRSY-DSSRYLDSLQSWY---ER------CE--WR-------------------------------------------------------QPKSSKERGR--------------------IYYGMV--
-----G--------IKDAA---------------------------D----LLYGTE---QNGYFSLSG-----------------------------------------------------------------KEERYKD--------VVKRWMPCILD---K-K-E---VPADMVRL-AV------------------------------------------------------------------S-----------SPVSFESRFLWERILA-------------
ACSL-------VKKQQIEKRY-----------------------------------------------------------------------------G----E-----------------------------------------------------------------------W---------------V-------ALD------KK---------CR-----K---------RNYLY------GRLLAVAD------RIE-----------Y-------------------------------------------------------------TFDKE-----DG-------ETNA----KRYMSAFS--------QHPYRT-----
--WKVLEE--KL-E-PYFM------K---L------------------------------------------------------------KAPERLV--------------Y-----------------QRL-L---D--------EIC--------------EL-----------------------F----------E----IEDFE-N-------DTALNGLYLLGFH---S---Q--------------------------------------------------------------------------------------------YA--------------------L---------------------------------------------------------------------
----------E----------------KKKEEEE-----------------------------------------------------------------------------------------------------------------------------------------------------------------------------------------------------
'WP_130433894.1'    -----------------------------------------------------------------------------------------------MNW--------VRELCDLYE----------KNQTLAGKVERGRFGEPL-----I----------------------------------------LLPL---FH-----TTV-TA-Q------ITVTINH---E---GV-FL-----AERVA-----------DE--------------------------DK----------------L----T-------II-P-----VTEK-SA--AR---------T-A---G-----
-P-----------E----A----H--------P-L-----------------CDNLKYLSGDYNDYTVSEKG-------------------------------------------------------------------------------------------------------------------------------------------------------------------------Y---------S------------E----------------------------HHR---------CY-IE-GLEA---W--AKS-----------EFSHE-
-K-------VRA----VYKYLEK-----------G------NLM----------------------------K------------------------------------------------------------------IACGVL--------------------------TGEDGCVLPKEKIQNVAQQDAFV------------------RF-------C-V-----------------------------E------E-----------------------------------EW----------DE------GRNLLREGEDRIPNE-CW--------------------------------------------------------------L
--DLTLQEAYIAY--------------------G-----------CR--------------------------------------------------------------------------------------------------ASVTQVG----------------L----------------------SY-------------LTG------EWTS----ISYL-H--P--K--KI--R--------------------IRNE-G---------DG-A-------------KLIS-----A-------------NDS-----------------------------------DETD-----Y-TF----R----------G----------------------------RFINK-------
--AFA-----IGYEDSQKAHNALKW-II-----------R-------R----------G------------C-------------------------------------------------------------------------------------------------------------------------------------------------------------------------------------------------------------------------------------------------DNYEAWE-G-
WGDEEEDRR--YAG-------------------------T-------------D-------P--REAAR----FQR-------A---------------I-A-G------Y---------EK--------MMD---------K----S----M--Q-----T--ILMAFD---AAT-T---GRLAMMECQQL-STSAYLKNLKRWH---ED-------CG--WL-----------------------------------------------------------------------------PKYKDKSYYR------------------------YGVAGVR-
---G-------IANLL------------------------T-------ESRGDM---ALKGSAEKM-------------------------------------------------------------------------------------------------------------------------VCKRLLPCILS--G-R-P---VPQDMVNL-AV-----------------------------------------------------------------------------S-----------SPVSYESRMNWDRTLS---------------------------------------------ACSL-
VKK---------------------------------------------E--------------IERKEE------E---------------------------------------------W-----------------------------V------ALD------KE---------CK-----D---------RNYLY------GRLLAVAD------RIE-----------Y--------RTQDKEE------------------------------------------------------------R-----ETNA----KRFMNAFS--------QQPFRT-----
WKVIEE--RI-Q-PYKM------K---L------AAPERRH------------------------------------------------------------------Y-------------------EHM-I---E--------EIC--------------WL-----------------------F----------QEGDFE-KG-D-----SLNGRYLLGFH---N---Q-----------------------------------------------------------------------------------------------------E---------------------------------------------------------------------
--------R-----------N----------RTEEEKDHE----------------------------------------------------------------------------------------------------------------------------------------------------------------------------------------------------
'WP_025486472.1'    -----------------------------------------------------------------------------------------------MNW--------VRELCDLYE------------K--------N---------A--------NLAGQYEKGRFGEPLI------------LLPV---FH-----STV-AA-Q------ITVTINE---K---GE-FL-----R-AVPVED---------------------------------------------------EE------------------KL----T-----V-I-P----VTEK-SA--SR-----------T-
AGV------------------EAH-------P-L-------------------CDNLKYL--------A------G-------------------------------------------------------------------------------------------------------------------DY------MDYVVPDKGKDYTEN----------------------------------------------------------------------------------------------H-K-------LY-MD-GLKK--W--AES-------------
AFSHE---------K-VHA---IWNYLNQ-----------E------CLM----------------------R------D---------LVSYGCLKTDGQGKVLKTIKIQNVSQVDSF---V---RFRV--------------------------------------------------------------------------------EK-----------------PW-------------DA-----EENLLEEKDGTE-RPECW---------------------------------------------------------
L---------------------------------------------------------------------------------------------------DVSLENAYIAYCRASWQQRGL-----------SY-------------LTG------NQ-E-Q----ISYL-H--P--K--KI--R--------------------N-E-G---------DG-A-------------KLIS-----A-----------------------------------------------NDETYYTY-----R-----------G---------------------------------RFSNKEE-
---------AFA------IGYEDSQKAHNALKW-II-----------R-----R-Q-----------G-------N----------N---------------------------S--LC-------VVT-W------E----------------------------------------------------D---LIPLPDWCLDTD--------------M--------V----------C-------------------------------------
DIYESED-I-WGDEEESKKVPEYMG-------------------------------------D----------S----------RGAAR----FQA-------A---------------I-A-G------Y---------GA-------------RHD-------K----S----T--VLMAFD---AAT-T---GRLAMMEYESL-DSSRYLKRLKEWH---AG------CG--WL-------------------------------------------------------------------PKFKDKKYFL-------------------YGMAGVR-
------D---------IADIL-------------YG-------------S----EQNGML---TLKGGNEKM-----------------------------------------------------------------------------------VLKRLLPCIIS--G-R-P---VPEDMVHL-AY--------------------------------------------------------------------------------------KA----------------SPVSYDSRYNWERVLA-----------
---------ACSFEKKKRMEQ--------------------------------------------------------------------------------------E---------------------------------------------------------------W-------------WK--------------V---------A--------------V-------ALN------TE---------SQ---D-------RNYLY------GRLLAVAD------RVE-----------Y--------------------------------------------------------------------------------------------TFEK---DE------V----R------ETNA---KRFMNAFS-----
QQPFRT---------WKVIEE--RL-E-PYFV--------KLPL-------------------------------------------------------------PERLY--------------------EHM-I---E-------DIS--------------W-----------------------KF------------KEGEFE-KN-D------ALNGLYLLGYH---N---Q--------------------------------------------------------------------------------------------------------------------------------------------------------------------------
-----------------------R-----------------N-----------------YAEEEKK------------------------------------------------------------------------------------------------------------------------------------------------------------------------------
'WP_006442419.1'    -----------------------------------------------------------------------------------------------MNW--------TNELLDLYE------------------K-------NQDI-----A--------GEVTY-K------T-IK-GKKGEEHI-------PLILLPV---FH-----TTVTA-Q------IEVTIDE---S---GE-FL-----S-A----------K-----------AV----DKA--------------------------EN-------------------L----T-------M-I-P-----VTEK-SL--SR---------T-A---
-----G-------T--E----P---H------------P-L-----------------CDNLKYL--------A------G-------------------------------------------------------------------------------------------------------------------------------------------------------------------D-YMV-----YY-------------------------------------------------------------------------------------GKQK----DYSEN-------------Y-R----------LY-ME-ALQR---W--HES-----
P-FTH--KK-----VDA---IYAYVCK-------------G------CMI---------------------------K------D------L---------A-------K-----------------------------------------S--E-----------T-----------I-KQ--------------------------------------------------------VRF-------R-V-ESILDAGDILAD-------------TSG---Q-------YCAE-------------------------------CW----------L-----DK------T-----------L-QE-AY---------------------------------------------------------
---------------------TS---------------------------------------------------M-L---P--E--KD---------------------------------------------------------------------------------------------------------------------CY-------------LTG------KR-E-A----ISYL-Q--P--K--KI--R--------------------N-E-G---------DG-A-------------KLIS-----A-------------NDE-----------------------------------S-----N------F-TF----R----------G-----------------------------R--------FSD------KT-

[illegible]

'WP_117771473.1__2_'

[illegible]

'WP_049729887.1'

[illegible]

'WP_004604883.1'

[illegible]

'WP_089982255.1'

[illegible]

'WP_103228660.1__2_'

[illegible]

'WP_092365221.1'

[illegible]

'RKJ33296.1'

[illegible]

'WP_019662283.1'

[illegible]

'WP_008974804.1'

[illegible]

'WP_028520265.1'

[illegible]

'WP_082335043.1'

[illegible]

'WP_009984821.1'

[illegible]

'WP_109726611.1'

[illegible]

'WP_092983467.1'

[illegible]

'WP_092983072.1'

[illegible]

'WP_002849082.1'

[illegible]

'WP_093039827.1'

[illegible]

'WP_013497833.1'

[illegible]

'WP_062413140.1'

[illegible]

'WP_051652299.1'

[illegible]

[illegible]

'WP_019543033.1'

[illegible]

'WP_006689561.1'

[illegible]

'WP_006695539.1'

[illegible]

'WP_009350456.1'

[illegible]

'WP_009729419.1'

[illegible]

'WP_009438895.1'

[illegible]

'WP_009658389.1__2_'

[illegible]

'WP_026762241.1'

[illegible]

'WP_009441727.1'

[illegible]

'WP_006693577.1'

[illegible]

'WP_021685845.1'

[illegible]

'WP_009657051.1__2_'

[illegible]

'WP_106626745.1'

[illegible]

'WP_050343674.1'

[illegible]

'WP_020224350.1'

[illegible]

'WP_117894803.1__2_'

[illegible]

'EEF66037.1'

[illegible]

'WP_006905219.1'

[illegible]

'WP_027113660.1'

[illegible]

'WP_101698636.1'

[illegible]

[illegible]

'WP_085833907.1'

[illegible]

'WP_020074667.1'

[illegible]

'WP_009062363.1__2_'

[illegible]

'WP_042431769.1'

[illegible]

'WP_120440002.1__3_'

[illegible]

'WP_016217546.1'

[illegible]

'WP_055151092.1__2_'

[illegible]

'WP_025653943.1__2_'

[illegible]

'WP_108981830.1'

[illegible]

'WP_087372535.1'

[illegible]

'WP_087233389.1'

[illegible]

'WP_016323950.1'

[illegible]

'WP_033116946.1'

[illegible]

'WP_075704675.1'

[illegible]

'WP_058118515.1'

[illegible]

'WP_116722325.1'

[illegible]

'WP_022786160.1'

[illegible]

'WP_050618386.1'

[illegible]

'PWL50276.1'

[illegible]

'WP_081579648.1'

[illegible]

'WP_006354664.1'

[illegible]

'WP_079699059.1__3_'

[illegible]

'WP_132085060.1'

[illegible]

'WP_090164510.1'

[illegible]

'WP_090472240.1'

[illegible]

'WP_036379586.1__2_'

[illegible]

'WP_027441235.1'

[illegible]

'AUO19654.1'

[illegible]

'EFB90881.1'

[illegible]

'WP_120146848.1__3_'

[illegible]

'WP_118002338.1__2_'

[illegible]

'WP_109769883.1'

[illegible]

'WP_002606245.1'

[illegible]

'WP_044904679.1'

[illegible]

'WP_117444014.1__3_'

[illegible]

'WP_065530107.1__2_'

[illegible]

'WP_008816402.1'

[illegible]

'WP_013759293.1__2_'

[illegible]

'TCU63078.1'

[illegible]

'WP_102715653.1__2_'

[illegible]

'WP_102739675.1__2_'

[illegible]

[illegible]

'QBH17563.1'

[illegible]

'WP_011833220.1'

[illegible]

'AKL98464.1'

[illegible]

'PKK39971.1'

[illegible]

'TCL55900.1'

[illegible]

'WP_077610084.1'

[illegible]

'WP_081721282.1'

[illegible]

'SDD52781.1'

[illegible]

'PWK15200.1'

[illegible]

'WP_108306869.1'

[illegible]

'WP_036146971.1'

[illegible]

'WP_042475427.1'

[illegible]

'WP_062109030.1'

[illegible]

'WP_129678401.1__2_'

[illegible]

'WP_026770863.1'

[illegible]

'WP_053217377.1'

[illegible]

'WP_135110974.1__2_'

[illegible]

'WP_090241460.1'

[illegible]

'WP_036146668.1'

[illegible]

'WP_012291755.1'

[illegible]

[illegible]

'KZA03726.1__3_'

[illegible]

'WP_015694583.1'

[illegible]

'WP_013774548.1'

[illegible]

'SER55234.1'

[illegible]

'WP_010766146.1'

[illegible]

'GAK05752.1'

[illegible]

'GAK11818.1'

[illegible]

'WP_107943245.1'

[illegible]

'WP_053994641.1'

[illegible]

'SEM82097.1__3_'

[illegible]

'PZX02418.1'

[illegible]

'RLK36865.1__3_'

[illegible]

'PLT47752.1'

[illegible]

'EHB62323.1'

[illegible]

'SDM16821.1'

[illegible]

'PYE47157.1'

[illegible]

'PXW11321.1__2_'

[illegible]

'SDD50387.1'

[illegible]

'ACB59636.1'

[illegible]

'TCT21776.1'

[illegible]

[illegible]

'SFO69105.1'

[illegible]

'TCP29034.1'

[illegible]

'SFK00111.1'

[illegible]

'RAP14431.1__2_'

[illegible]

'SES27298.1'

[illegible]

'SER92646.1'

[illegible]

'SFS81954.1'

[illegible]

'TDQ37158.1'

[illegible]

'GAF23122.1'

[illegible]

'GAE28763.1'

[illegible]

'SDZ39440.1'

[illegible]

'EES74001.1'

[illegible]

'SES30003.1'

[illegible]

'SEG63853.1'

[illegible]

'PRX55995.1'

[illegible]

'SDJ32211.1'

[illegible]

'AKN52807.1'

[illegible]

'AJH78187.1'

[illegible]

'EJL40221.1'

[illegible]

'RED75764.1'

[illegible]

L----------------------------------G----------------------------T-------------KGTYL--S-------KII---------D-----EV [illegible]
KDNPEGSGNIQIRGRDRDDDFGS----------------------------------------------------------------------------------

'KEF36269.1' [illegible]

------S-----------KI--AP---Y---------P-L------------------HDKLSYV------A---G [illegible]
---VKS---IYTYLSK--KHLI-----------K-----DLV--------------------E-----R----------------------------RILFLDDKGNLIEKWD [illegible]
R---------Y--YKDLLGD---------------------------------------------------------------ED [illegible]
VAS------ISYEVSQKAHNALKW-LI---------N---------------R-----Q-----G----------KI------------------------------V [illegible]
----------------------------------------------------T---------N-----------Q---EFANE----VAK----A-----------------------L-D-G----F [illegible]
-----------------FA---------------------T---YGPKASDKV [illegible]
------------K---------------------------------------------------------------------------E-------------G---------------Y-------------K [illegible]
---------------------G---------------------------T-------------KGLYL--S-------KMI---------D-----EV [illegible]
[illegible]

'TDW03919.1' [illegible]

G---------S---------KVA----P---Y---------P-L------------------HDKLSYV------A---G [illegible]
PYATS--K------VKS---IYHYLKK---------------R------TLI-------------------K-----------------------------DLVEEGILYLDAAGRL [illegible]
-----I--------------Q---------FYQDFLGG------------------------------------------------------ED [illegible]
-----------------AAA------ISYEVSQKAHNALKW-LI---------N---------------K-----Q-----G----------KI [illegible]
----------------------------------------------------G---------V------------EEDSFDISLLDPAPAVSQ----------------------------K-K-G [illegible]
--------TKDIA-----------------------FA------------------------A---YGSKAGEKV [illegible]
[illegible]

'TCW53147.1__2_' [illegible]

[illegible]

'AJK37991.1__2_' [illegible]

[illegible]

'SFM21720.1' [illegible]

[illegible]

'AFU17710.1' [illegible]

[illegible]

'OSY02406.1' [illegible]

[illegible]

'OSY06810.1' [illegible]

[illegible]

'ABB28270.1' [illegible]

[illegible]

'SKB36654.1' [illegible]

[illegible]

'WP_013484590.1__4_' [illegible]

[illegible]

'ERJ00272.1' [illegible]

[illegible]

'RCX20103.1' [illegible]

[illegible]

'AEY65500.1' [illegible]

[illegible]

'EGD49242.1' [illegible]

[illegible]

'ABR47841.1' [illegible]

[illegible]

'ACV64359.1' [illegible]

[illegible]

'SNS94721.1' [illegible]

[illegible]

'PRR82654.1' [illegible]

[illegible]

'WP_019604446.1' [illegible]

[illegible]

'WP_015818588.1'

[illegible]

'WP_018276374.1'

[illegible]

'WP_081042193.1'

[illegible]

'WP_015406748.1'

[illegible]

'WP_078409561.1'

[illegible]

'WP_058292009.1'

[illegible]

'RPE95721.1'

[illegible]

'TCP10414.1'

[illegible]

'ERH28584.1'

[illegible]

'AHG79312.1'

[illegible]

'ABR74646.1'

[illegible]

'TDQ57408.1'

[illegible]

'SOD68624.1'

[illegible]

'EKY02930.1'

[illegible]

'ANB91333.1'

[illegible]

'AKG18643.1'

[illegible]

'KDN25753.1'

[illegible]

'EGT75761.1'

[illegible]

'SFG94234.1'

[illegible]

'AHG83783.1__2_'

[illegible]

'TCT17821.1'

[illegible]

```
----VSA---VLHYVKK-----------------G------KVV---------------------E------D------L-----------I------T-A---G------V----FQL--------------G--A-D----GK----V---L------S-KWVEKGDA------PAIFSTLPK---------TKG---------EIE---FGSA-------L-----------------VCW-----R-V----------E-IK-----------G---D-------PQSD------------------TW---------T-----DV-----T-----------V-QQ-SW------------------------------I-------
---------D---------Y-----------L----------------------------------------------------AL----------------------------A-D--------S--Q---TG------------------------------F--------------CF---------IQG-------KE-S-P---VSNM-H--P--A--KL--R-----------------H-T-G------DK-A------------KLIS-----S-------------NDT-----------------------A----G-----Y-TF---R---------G------------------------------R-----------FET------AE-------E----
-----AAS------ISTEVSAKAHGALRW-LI---------S----------------------R------Q-----------GI----------------------------R-----------N-----G----------E----Q----V--------TVA--W----------A----ISGEKVPS-PLQDPF----DECYD------------------------------------YD--------------------------------L--------E---------------------------------------------------------------
--------------------------------------------------------E----------I-------------S----AVENN------VESEMPSET----------------------------R-G------K--IDHSVDLGKNAAEMLKKK---------YQGYKAKLKA-------H-----E-----------Q-------I--SLLMLDSATP-------GRMALTYYQEF-LPADYFANLDAWI--DD------FS--WY----------Q-------------------------------RH---------------------------ISETKNGKKN---------D-
-------KRLVWAIV-----------------------------------P---P----------SP------FAIGNA-------V-YSKSL-------SDSQK---------------K-----------------------------------------------------------------------------------------------LYARLLPVIAG--G-KSVP----IPYDLVQQ-SF-----------------------------------------Q--V-----------------------AC----------------------NPHGCENWE------------------------------------------------------W-QRNIG-----V-------------------ACAL--------
YKG------W-------------------------------------R--------------------ARHHNES-----------------------------------------------E-----------RRTY----D------------------------------------------------------M------SLD------KE----N---S-----RDYLY------GRLLAVAE-------NIE---------------------A--------------------ALYLAGEK----------------------------------------R--------STNA---ERYMPKFA-----NRPFYT------------
-WRNIE---IALK-PYQE------R--L-------------------R--NHNKDTG---------------------------------------------SQA-L--A-------EIT-----------------DL-------------------------------F--------------V----T-EDYT-ND-S------PLSAEFLLGYH---C----Q-------R---------------------------------------------------------K-----------------------ME-----------------I--ARQLAE-------LT-AKKS-----------K-----
-------------T-------------------------TE-------TE------------------------------------------------------------------------------------------------------------------------------
                              'AHG81512.1'                 -------------------------------------------------------------------------------------MQKLYRTYE-------------A---------A--L----Q--------KA----S--N---------LSEEP-------------LTPI--GH-----TQQNA-H------IVIVLNG--D---GE-FR----T-A---------Q--------VM-----P---------------------------------PK---------------------TA-----I-----M-L-P-----ATES-SE--NR----------T---S---G----------
E-------------A----P---H-----------P-L------------------ADKIQYV---------A----R-------------------------------------------------------------------------------------------------------------------------------------------------------------------------D-YSA---Y----------------G-------------------E----------K-KAY--------------------------------F-E------------GY-LN-QLQA--W--CDS--------------A-VSH--PK-------
VSA---VLHYVKK-----------------G------KVV---------------------E------D------L-----------I------T-A---G------V----FQL--------------G--A-D----GK----V---L------S-KWVEKGDA------PAIFSTLPK---------TKG---------EIE---FGSA-------L-----------------VCW-----R-V----------E-IK-----------G---D-------PQSD------------------TW---------T-----DV-----T-----------V-QQ-SW------------------------------I-------
------D---------Y-----------L----------------------------------------------------AL----------------------------A-D--------S--Q---TG------------------------------F--------------CF---------IQG-------KE-S-P---VSNM-H--P--A--KL--R-----------------H-T-G------DK-A------------KLIS-----S-------------NDT-----------------------A----G-----Y-TF---R---------G------------------------------R-----------FET------AE-------E----
-AAS------ISTEVSAKAHGALRW-LI---------S----------------------R------Q-----------GI----------------------------R-----------N-----G----------E-----------------------------------------------------------------------------------------------------------------------------------YE------------------F-------E-------------------------------------------------------------
--------------------------------------------------------E----------I-------------S----VVESD------VESEVPSEI-----------------------------E-G------K--IDHSVDLGKNAAEMLKKK---------YQGYKAKLKA-------H-----E-----------Q-------I--SLLMLDSATP-------GRMALTYYQEF-LPADYFANLDAWI--DD------FS--WY---------------------------------------QR----------------------------------------------H-------SIETKNGKKNEKRP------
-------IWAVV-------------------------PP-------------S--PFAIGNAVYGKSLSDSLKKQ---------------------------------------------------------------------------------------------------------------------LYARLLPVIAG--G-KNIP----IPYDLVQQ-SF-----------------------------------------Q--V-----------------------AC-------------------N-------------PHGCENWEWQRNID----------------------------------------------V---------------ACAL-------YKG-----
--W-------------------------------------------R--------------------------------------------------------ARHHNESERRTYD-----------------------------------------------------------M--------SLD---------KE-----------N---S-----RDYLY------GRLLAQAE-------RLE--------WYALYLQNGKK-------------T---------------N----------------------------------------------PS-----------R--------ATNA---ERYFQRFA-------QLPYST----------
WLNFERERLVPYKNYLT------S--L---------------------------------------------------------GKDF------------Y--------------KQA-I--G--------EII-------------------EL----------------------------------------QNNDFT-CD-D-------KLSGEFLLGYY---C----Q-------------------------------------------------------------K--------------------------ME-------------------I--------------------------------------------------------------
---R--------QIAARKAAKQKTTETD----------------------------------------------------------------------------------------------------------------------------------------------------
                              'AHG86671.1'                 -------------------------------------------------------------------------------------MQKLYRTYE-------------A---------A--L----Q--------KA----S--N---------LSEEP-------------LTPI--GH-----TQQNA-H------IVIVLNG--D---GE-FR----T-A---------Q--------VM-----P---------------------------------PK---------------------TA-----I-----M-L-P-----ATES-SE--NR----------T---S---G----------
E-------------A----P---H-----------P-L------------------ADKIQYV---------A----R-------------------------------------------------------------------------------------------------------------------------------------------------------------------------D-YSA---Y----------------G-------------------E----------K-KAY--------------------------------F-E------------GY-LN-QLQA--W--CDS--------------A-ASH--PK-------
VSA---VLHYVKK-----------------G------KVV---------------------E------D------L-----------I------T-A---G------V----FQL--------------G--A-D----GK----V---L------S-KWVEKGDA------PAIFSTLPK---------TKG---------EIE---FGSA-------L-----------------VCW-----R-V----------E-IK-----------G---D-------PQSD------------------TW---------T-----DV-----T-----------V-QQ-SW------------------------------I-------
------D---------Y-----------L----------------------------------------------------AL----------------------------A-D--------S--Q---TG------------------------------F--------------CF---------IQG-------KE-S-P---VSNM-H--P--A--KL--R-----------------H-T-G------DK-A------------KLIS-----S-------------NDT-----------------------A----G-----Y-TF---R---------G------------------------------R-----------FET------AE-------E----
-AAS------ISTEVSAKAHGALRW-LI---------S----------------------R------Q-----------DI----------------------------R-----------N-----G----------E----Q----V--------TVA--W----------A----ISGEKVPS-PLQDPF----DECYD------------------------------------YD--------------------------------L--------E---------------------------------------------------------------
--------------------------------------------------------E----------I-------------S----AVENN------VESEMPSET----------------------------R-G------K--IDHSVDLGKNAAEMLKKK---------M-----YQGYKAKLKA-------H-----E-----------Q-------I--SLLMLDSATP-------GRMALTYYQEF-LPTDYFANLDAWI--DD------FS--WY----------Q-------------------------------RH------SIET------K-----NGKKNEKR------------------------------------P-
-------IWAVV------------------PP-------------SPFAI------GN-----AVYGKS-----------------L-------S----DSLKK------------------------------------------------------------------------------LYARLLPVIAG--G-KNVP----IPYDLVQQ-SF-----------------------------------------Q-------VA-C-------------------NP-------------------------------HGCENWE------------------------------------------------------W-Q--------RNIG-----V-------------ACAL--------
YKG------W-------------------------------------R--------------ARHHNES-------------------------------------------------------------------------------------TYD--------M-----SLD------------KE-------------NR---S-------------RDYLY------GRLLAQAE-------RLE---------------------W----------------Y------------------------ALYLQNGKKT----PS----------------------R------ATNA---ERYFQRFA-------QLPYST-----
--------WLNFER--ERLV-PYKN-----YLT-SL-------------------GKD------------------------------F-----------------Y---------------------KQA-I--G-----------EII--------------EL-----F----G-----------------------N----N-D-FT-CD-D-------KLSGEFLLGYY---C----Q-------------------------------------------------------------K--------------------------ME---------------------I------------------------------------------------------------------
---R------------QIAARKAAKQKTTET-------D---------------------------------------------------------------------------------------------------------------------------------------------------------
                              'TCP90140.1'                 -------------------------------------------------------------------------------------MQKLYKTYE------------Y-------V--Q------E------LE------------------IDDED---------------LALP--FH----MPKAV-H------LKVVLND--K---GE-LV-----Q-A---------E-----------RF------D---------------------------------GK-----------------KQ----V------P-I-Q----VTEK-SS--KR---------S-G---S----
----T-----------I--A----A---Y----------A-L--------------HDSLQYI----------A----K---------------------------------------------------------------------------------------------------------------------T-AGQ----YLTIEYLSKVAEKES----G----------E-------------------------K----------K----------W-KAFLSDTEDGKKQFVDEEKAKYQKSF-E----------FY-EQ-QLKK--W----
AEF------------S-NVN----E----IKV-----VSEYVQK---------------G------SLI---------------------------------------------------------E-------D------L---------------L--------S-K---N-----I-----FSI-----------------------------K-E------------------N-ILLSGNDD---------PFSLT---------------------------IVW-------A-V-------------E-MP----------------H-------D-------PHSD------------------LW-------S------KS------S----------I-KT-EW--
-----------------R---------Y--------QESL-D--------------------------------------------EL----------------------------------S-F-------N---Q---KD-------------------L---------CY----------ATG-------EL-D-F------PATA-Y--P--------------------K-I-E-------GN-A----------KLVS------A--------------NDT------------------------E-----G-----L-TF--L---------G--------------------------------R----------FQT-------DK------
-Q--------AAS------LGRDVSQKAFNMLKW-LI--------K-----------------------R-------Q-------------GI---------------------------------R------------N-----G--------E-----S------Q---V---------TVA--W--------A-ISGKEIPSPMKDNFDEIDWDNLDTSAAE------------------------------------------------TEAEISYENE-----------------------------------------------------------------------------------------------------------
---------------------------------------------------------------------------------------------------------G-------K-----AAADI-------IKK-----------------KLH-G---Y------------------QA----------------N------LEA-----H----E-----------Q-------I--SLLMLDSATP-------GRMALTYYQEF-LPADYFANLDAWI--DD------MS--WY------------Q-----------------------------------QR-------------------------------------------------Y------SIEKPNAKKSNKKQ-------
-----T---------VWAFI-------------PPSP-------FAIAQ-------T----I----LSRAER---EDKEKKAQK-----------------------------------------------------------------------------------------------------------------LYARLLPVIAG--G-HSVP----IPFDLVQK-SF-----------------------------------------Q-----------------------TA-----------------------------CNPNGCENWEWQRNIG---------------------------------------------------------------ACAL-------
--YKG------W-------------------H-------------------------------------------------------------------------------------------------------------------------H-----YNSSPRRTY----S----------------------------------------NR------S-------------RDYLF------GRLLAVAE-------SIE--------------------S-----------Y---------ALYLA------------------------------------------------------------DE------K-----R-------ATTA---ERYMQHFA-------EHPFAT-------
--WRNIEL-A-K-PYQE------R--L------------------RNRGKDTG---------------------------------------------------------SQT-I--S-----------EIM-----------------EL------------------------------------F------------ELADFT-CD-D-------KLSGEFLLGYH---C----Q----------------------------------------------------------------------------------------K--------------------------ME---------------------T----------I-------------------------------------------------------
---Q---------------IAERKAAKAKSTDTIE----------------------------------------------------------------------------------------------------------------------------------------------
                              'KYK82925.1__2_'             -------------------------------------------------------------------------------------MSW-------MQKLCRTYE-------------Y--------V---Q----E--------QG------------------LEDED---------------LALP--FH----MSKAV-H------LKVILNN--K---AE-LV-----G-V-----------E--------------RF------E---------------------------------GK-----------------KQ----V------P-I-Q----VTEK-SS--KR---------A-G---S-----
----T----------I--A----P---Y-------------A-L--------------HDGLQYI-----------A---K------------------------------------------------------------------------------------------------------------------T-AKE----YLTIEYLSKVAEKEN-----G-----------------------------K---------K--------W-KNFLAGTDEDKRNFADTEKTRYKTSF-K--------AY-EE-QLSA--W-
AEA-------------S-HLQ----E----IKI--VLQYIQK---------------G------SLI---------------------------------E-------D------L---------------L-------E-K-------K----V------FTL-----------------E--N---------------N-ILLAGKDD------PFNFT------------------------------IVW-------A-V-------------E-IP----------N-----D-------LHSN------------------LW-------S-----KS------S-----------I-KE-QW------
----------K---------Y---------QESQ-S---------------------------------------------------GQ------------------------------------------------E-D----------E----K--PE----------------L----------CY------------ITG------ES-S-Y---LANA-Y--P-----------------K-I-E-------GN-A-----------KLVS------A--------------NDT-----------------------D----G------F-TF--L---------G---------------------------------------------R------FLI------DK------
Q---------AAA-------LGREVSQKAFNMLKW-LI--------K-----------------------------R-------Q-----------GI--------------------------------R------------N-----G--------D---Q------V--------TVA--W--------A-ISGKAVPSPMKD---------------------------------------------------------------------------LSDEIDWDNLDISADESAVENSDR---------------------------F----------S-----------------------------
-------------------------------------------------S--------PTSSETPDWSTNIGK------PAAQI----IKK-----------------KLH-G-----Y----------------QE---------E------LKN-------H-----E-----------Q-------I--SLIMLDSATP-------GRMALTYYQEF-LPADYFANLDAWV--DD------FS--WY----------------------------------------------------------------------------------------------------------
YSVGVQNGKKTDKRTQWRGVPP---------------S-----------PYSVA------------------------------DA-------------------------V----------YGKSLSDTL---------------------------------------------------------------------KK-----Q-----------LYARLLPVIAG--G-KSVP----IPDDLMQK-SF-----------------------------------------Q--V------------------------AC--------------------N-----------------------PNGSENWEWQRNIG-----------
---------------V---------------ACAL--------YKG--------W-------------------------------------------R--------------------------------------------------------------------------------------------------------------ARHHDLSERRTYSM-----ALD---------KE-------------NR---S-------------RDYLY------GRLLAVAE-------NTE------------------------------S--------------------------------------------------YALYLVGEKR---------
ATTA---ERYMQRFA-------EHPFAT-------WRNIEL-A-L-K-PYQD------R--L------------------RNNGKDTG---------------------------------------------------------------AQA-I--G-------EIM-------------------DL----------------------------------F------------ENQDFS-SD-D-------KLSGEFLLGYH---C----Q----------------------------------------------------K-------------------------------------------MD----------I-----
----A------------------R-------------------R----------IAEISANKSKTDE-----------------------------------------------------------------------------------------------------------------------
                              'KYK95002.1'                 -------------------------------------------------------------------------------------LSAWAEASH-------------------------------------------------------------------------------------------LQEI--------------KI--------VLQYIQK-------GS-LI-------------------E---------D---------------------------------------------L-E-----------KK-------------------------------------------V----------------F-T-
-----------------LENNILL-------A----G-----------------------------------------------------------------------------------------------------------K--------------------------------------------------------------------------------------------------------------------------------------------NF-TI-V------W---AVE----------I-----P-----------------------------------------------------
--------------------------------------------------N-D-L-----------------------------HSN--------------------------------------------------------------------------LW-------------S--------KS--------S--------------I-KE-QW-----------------------------------------------------------K------------------------------Y-------QESQ-S-----------------------------------------------------------GQE---------DE---------
-------------------K----P-E----------L-----------------CY------------ITG-----E---S--S-Y---LANA-Y--P--K----I--E---------------------GN---A------------------KLVS-----A------------------------------------------------DTD-------G---F-TF--L-------------G------------RFLID-------------------------KQ-------------------AAA-------LGREVSQKAFNMLKW-LI---------K-------------------------------------R------------Q---------------------G----------------
-R-----------N----G-------D----Q----V--------TVA--W-------A----ISGKAVPSPMKDLSDEIDWDNLDI-----------------------------------------------------------------------------------------------------------------------------------------------------------------------------------------------------------------------------------------------------------------S-------PTSSETPDWSTNIGK----PAAQI-----
IKK-------------------KLH-G-----Y-------------------QE---------E------LKN-------H-----E------I--SLIMLDSATP-------GRMALTYYQEF-LPADYFANLDAWV--DD------FS--WY----------------------------------------------------------------------------------------------------------ADESAVENSD------------R-----------F----------S------------------------------------------------------------------------------------------------------------------------------------------------------
DTLKK------------------------------Q------------LYARLLPVIAG--G-KSVP----IPDDLMQK-SF-----------------------Q--V-----------------QRYSV-------------GVQNGKKT----DKRTQWRGVPP-----------------------------------PNG-------SENWEWQ-R-----NIGV-----ACAL-------YKG------W---------------------------------------------------
-------------------RARHHDLSE----R--R--TY-----------S----------M----ALD---------------KE-------N---------RS--RDYLY------GRLLAVAE-------NTESY-ALYLVGEK-----------------------------------------------------------------------------------------R--------ATTA---ERYMQRFA-------EHPFAT------------WRNIE---LALK-PYQD------R--L-------------------------RNNG-KDT--G-AQA-------
--------------IGE-I--M----------D--------L----FE----------------------------N-QD--------------FS-SD-D-------KLSGEFLLGYH---C----Q------------------------------------------------------------------K-------------------------MD-----------------IA---------------------------------------R------------R-IAEISANKSKTDE-----------------------------------------------------------------
                              'AFI86060.1'                 -------------------------------------------------------------------------------------MSW-------MQKLCRTYE-------------Y--------V---Q----E--------QG------------------LEDED---------------LALP--FH----MSKAV-H------LKVILNN--K---AE-LV-----G-V-----------E--------------RF------E---------------------------------GK-----------------KQ----V------P-I-Q----VTEK-SS--KR---------A-G---S-----
----T----------I--A----P---Y-------------A-L--------------HDGLQYI-----------A---K------------------------------------------------------------------------------------------------------------------T-AKE----YLTIEYLSKVAEKEN-----G-----------------------------K---------K--------W-KNFLAGTDEDKRNFADTEKTRYKTSF-K--------AY-EE-QLSA--W-
AEA-------------S-HLQ----E----IKI--VLQYIQK---------------G------SLI---------------------------------E-------D------L---------------L-------E-K-------K----V------FTL-----------------E--N---------------N-ILLAGKDD------PFNFT------------------------------IVW-------A-V-------------E-IP----------N-----D-------LHSN------------------LW-------S-----KS------S-----------I-KE-QW------
----------K---------Y---------QESQ-S---------------------------------------------------GQ----------GV-------------------------------------E-D----------E----K--PE----------------L----------CY------------ITG------ES-S-Y---LANA-Y--P-----------------K-I-E-------GN-A-----------KLVS------A--------------NDT-----------------------D----G------F-TF--L---------G---------------------------------------------R------FLI------DK------
Q---------AAA-------LGREVSQKAFNMLKW-LI--------K-----------------------------R-------Q-----------GI--------------------------------R------------N-----G--------D---Q------V--------TVA--W--------A-ISGKAVPSPMKD---------------------------------------------------------------------------LSDEIDWDNLDISADESAVENSDR---------------------------F----------S-----------------------------
-------------------------------------------------S--------PTSSETPDWSTNIGK------PAAQI----IKK-----------------KLH-G-----Y----------------QE---------E------LKN-------H-----EQ--------I--SLIMLDSATP-------GRMALTYYQEF-LPADYFANLDAWV--DD------FS--WY-------------------------------------------------------------QR-----------------------------------------------
YSVEVQNGKKTDKRTQWRGVPP---------------S-----------PYSVA------------------------------DA-------------------------V----------YGKSLSDTL---------------------------------------------------------------------KK-----Q-----------LYARLLPVIAG--G-KSVP----IPDDLMQK-SF-----------------------------------------Q--V------------------------AC--------------------N-----------------------PNGSENWEWQRNIG-----------
---------------V---------------ACAL--------YKG--------W-------------------------------------------R--------------------------------------------------------------------------------------------------------------ARHHDLSERRTYSM-----ALD---------KE-------------NR---S-------------RDYLY------GRLLAVAE-------NTE------------------------------S--------------------------------------------------YALYLVGEKR---------
ATTA---ERYMQRFA-------EHPFAT-------WRNIEL-A-L-K-PYQD------R--L------------------RNNGKDTG---------------------------------------------------------------AQA-I--G-------EIM-------------------DL----------------------------------F------------ENQDFS-SD-D-------KLSGEFLLGYH---C----Q----------------------------------------------------K-------------------------------------------MD----------I-----
----A------------------R-------------------R----------IAEISANKSKTDE-----------------------------------------------------------------------------------------------------------------------
                              'KYK88993.1'                 -------------------------------------------------------------------------------------MSW-------MQKLCRTYE-------------Y--------V---QE----------------QGLEDED---------------LALP--FH----MSK-AV-H------LKVILNN--K---AE-LV-----G-----------VERFE------------------------------------GK-----------------KQ--------------V-PIQVTEKSS------------------------------------K-R-----A--G-----
-----S---------TIA----P---Y-------------A-L--------------HDGLQYI-----------A---------------------------------------------------------------------------------------------------------------------------------------------------------------------K----------------------TAKEYLTIEYLSKVAEKENGKKWKNFLAGTDEDK--------RNFADTEKTRYKTS-------------------F-K---------AY-EE-QLSA--W-
AEA-------------SHLQE-------IKI--VLQYIQK---------------G------SLI----------------------------------------E-D-L----------------------------------EKK------------------------------------------------------------VF-----T-LENNILLAGKDDPFNFTIVWAVEIPNDLHS---N--------------------LW-------S-----KS------S-----------I-KE-QW-----
---------K-----------Y---------------------------------------------QESQSGQEDEKPE---------------------------------------------------L------------CY------------ITG--E---S--S-Y---LANA-Y--P--K----I---------------------EGNA-----------KLVS------A------------------------------DTDS------F-TF--L------------G---------RFLID-----------------------KQ--------
--------AAA-------LGREVSQKAFNMLKW-LI--------K-----------------------------R-------Q-----------GI--------------------------------R------------N-----G--------D---Q------V--------TVA--W--------A-ISGKAVPSPMKD---------------------------------------------------------------------------LSDEIDWDNLDISADESAVENSDR---------------------------F----------S-----------------------------
-------------------------------------------------S--------PTSSETPDWSTNIGK------PSAQI----IKK-----------------KLH-G-----Y----------------QE---------E------LKN-------H-----E-----------Q-------I--SLIMLDSATP-------GRMALTYYQEF-LPADYFANLDAWV--DD------FS--WY-------------------------------------QR-----------------------------------------------
YSVEVQNGKKTDKRTQWRGVPP---------------S-----------PYSVA------------------------------DA-------------------------V----------YGKSLSDTL---------------------------------------------------------------------KK-----Q-----------LYARLLPVIAG--G-KSVP----IPDDLMQK-SF-----------------------------------------Q--V------------------------AC--------------------N-----------------------PNGSENWEWQRNIG-----------
---------------V---------------ACAL--------YKG--------W-------------------------------------------R--------------------------------------------------------------------------------------------------------------ARHHDLSERRTYSM-----ALD---------KE-------------NR---S-------------RDYLY------GRLLAVAE-------NTE------------------------------S--------------------------------------------------YALYLVGEKR---------
ATTA---ERYMQRFA-------EHPFAT-------WRNIEL-A-L-K-PYQD------R--L------------------RNNGKDTG---------------------------------------------------------------AQA-I--G-------EIM-------------------DL----------------------------------F------------ENQDFS-SD-D-------KLSGEFLLGYH---C----Q----------------------------------------------------K-------------------------------------------MD----------I-----
----A------------------R-------------------R----------IAEISANKSKTDE-----------------------------------------------------------------------------------------------------------------------
                              'KOE70404.1'                 -------------------------------------------------------------------------------------MSW-------MQKLCRTYE-------------Y--------V---QE----------------QGLEDED---------------LALP--FH----MSK-AV-H------LKVILNN--K---AE-LV-----G-----------VERFE------------------------------------GK-----------------KQ--------------V-PIQVTEKSS------------------------------------K-R-----A--G-----
-----S---------TIA----P---Y-------------A-L--------------HDGLQYI-----------A---------------------------------------------------------------------------------------------------------------------------------------------------------------------K----------------------TAKEYLTIEYLSKVAEKENGKKWKNFLAGTDEDK--------RNFADTEKTRYKTS-------------------F-K---------AY-EE-QLSA--W-
AEA-------------SHLQE-------IKI--VLQYIQK---------------G------SLI----------------------------------------E-D-L----------------------------------EKK------------------------------------------------------------VF-----T-LENNILLAGKDDPFNFTIVWAVEIPNDLHS---N--------------------LW-------S-----KS------S-----------I-KE-QW-----
---------K-----------Y---------------------------------------------QESQSGQEDEKPE---------------------------------------------------L------------CY------------ITG--E---S--S-Y---LANA-Y--P--K----I---------------------EGNA-----------KLVS------A------------------------------DTDS------F-TF--L------------G---------RFLID-----------------------KQ--------
--------AAA-------LGREVSQKAFNMLKW-LI--------K-----------------------------R-------Q-----------GI--------------------------------R------------N-----G--------D---Q------V--------TVA--W--------A-ISGKAVPSPMKD---------------------------------------------------------------------------LSDEIDWDNLDISADESAVENSDR---------------------------F----------S-----------------------------
-------------------------------------------------S--------PTSSETPDWSTNIGK------PSAQI----IKK-----------------KLH-G-----Y----------------QE---------E------LKN-------H-----E-----------Q-------I--SLIMLDSATP-------GRMALTYYQEF-LPADYFANLDAWV--DD------FS--WY-------------------------------------QR-----------------------------------------------
YSVEVQNGKKTDKRTQWRGVPP---------------S-----------PYSVA------------------------------DA-------------------------V----------YGKSLSDTL---------------------------------------------------------------------KK-----Q-----------LYARLLPVIAG--G-KSVP----IPDDLMQK-SF-----------------------------------------Q--V------------------------AC--------------------N-----------------------PNGSENWEWQRNIG-----------
---------------V---------------ACAL--------YKG--------W-------------------------------------------R--------------------------------------------------------------------------------------------------------------ARHHDLSERRTYSM-----ALD---------KE-------------NR---S-------------RDYLY------GRLLAVAE-------NTE------------------------------S--------------------------------------------------YALYLVGEKR---------
ATTA---ERYMQRFA-------EHPFAT-------WRNIEL-A-L-K-PYQD------R--L------------------RNNGKDTG---------------------------------------------------------------AQA-I--G-------EIM-------------------DL----------------------------------F------------ENQDFS-SD-D-------KLSGEFLLGYH---C----Q----------------------------------------------------K-------------------------------------------MD----------I-----
----A------------------R-------------------R----------IAEISANKSKTDE-----------------------------------------------------------------------------------------------------------------------
                              'KYK73531.1__3_'             -------------------------------------------------------------------------------------MSW-------MQKLCRTYE-------------Y--------V---QE----------------QGLEDED---------------LALP--FH----MSK-AV-H------LKVILNN--K---AE-LV-----G-----------VERFE------------------------------------GK-----------------KQ--------------V-PIQVTEKSS------------------------------------K-R-----A--G-----
-----S---------TIA----P---Y-------------A-L--------------HDGLQYI-----------A---------------------------------------------------------------------------------------------------------------------------------------------------------------------K----------------------TAKEYLTIEYLSKVAEKENGKKWKNFLAGTDEDK--------RNFADTEKTRYKTS-------------------F-K---------AY-EE-QLSA--W-
AEA-------------SHLQE-------IKI--VLQYIQK---------------G------SLI----------------------------------------E-D-L----------------------------------EKK------------------------------------------------------------VF-----T-LENNILLAGKDDPFNFTIVWAVEIPNDLHS---N--------------------LW-------S-----KS------S-----------I-KE-QW-----
---------K-----------Y---------------------------------------------QESQSGQEDKKPE---------------------------------------------------L------------CY------------ITG--E---S--S-Y---LANA-Y--P--K----I---------------------EGNA-----------KLVS------A------------------------------DTDS------F-TF--L------------G---------RFLID-----------------------KQ--------
--------AAA-------LGREVSQKAFNMLKW-LI--------K-----------------------------R-------Q-----------GI--------------------------------R------------N-----G--------D---Q------V--------TVA--W--------A-ISGKAVPSPMKD---------------------------------------------------------------------------LSDEIDWDNLDISADESAVENSDR---------------------------F----------S-----------------------------
-------------------------------------------------S--------PTSSETPDWSTNIGK------PAAQI----IKK-----------------KLH-G-----Y----------------QE---------E------LKN-------H-----E-----------Q-------I--SLIMLDSATP-------GRMALTYYQEF-LPADYFANLDAWV--DD------FS--WY-------------------------------------QR-----------------------------------------------
YSVEVQNGKKTDKRTQWRGVPP---------------S-----------PYSVA------------------------------DA-------------------------V----------YGKSLSDTL---------------------------------------------------------------------KK-----Q-----------LYARLLPVIAG--G-KSVP----IPDDLMQK-SF-----------------------------------------Q--V------------------------AC--------------------N-----------------------PNGSENWEWQRNIG-----------
---------------V---------------ACAL--------YKG--------W-------------------------------------------R--------------------------------------------------------------------------------------------------------------ARHHDLSERRTYSM-----ALD---------KE-------------NR---S-------------RDYLY------GRLLAVAE-------NTE------------------------------S--------------------------------------------------YALYLVGEKR---------
ATTA---ERYMQRFA-------EHPFAT-------WRNIEL-A-L-K-PYQD------R--L------------------RNNGKDTG---------------------------------------------------------------AQA-I--G-------EIM-------------------DL----------------------------------F------------ENQDFS-SD-D-------KLSGEFLLGYH---C----Q----------------------------------------------------K-------------------------------------------MD----------I-----
----A------------------R-------------------R----------IAEISANKSKTDE-----------------------------------------------------------------------------------------------------------------------
                              'KYK90791.1__4_'             -------------------------------------------------------------------------------------MSW-------MQKLCRTYE-------------Y--------V---QE----------------QGLEDED---------------LALP--FH----MSK-AV-H------LKVILNN--K---AE-LV-----G-----------VERFE------------------------------------GK-----------------KQ--------------V-PIQVTEKSS------------------------------------K-R-----A--G-----
-----S---------TIA----P---Y-------------A-L--------------HDGLQYI-----------A---------------------------------------------------------------------------------------------------------------------------------------------------------------------K----------------------TAKEYLTIEYLSKVAEKENGKKWKNFLAGTDEDK--------RNFADTEKTRYKTS-------------------F-K---------AY-EE-QLSA--W-
AEA-------------SHLQE-------IKI--VLQYIQK---------------G------SLI----------------------------------------E-D-L----------------------------------EKK------------------------------------------------------------VF-----T-LENNILLAGKDDPFNFTIVWAVEIPNDLHS---N--------------------LW-------S-----KS------S-----------I-KE-QW-----
---------K-----------Y---------------------------------------------QESQSGQEDEKPE---------------------------------------------------L------------CY------------ITG--E---S--S-Y---LANA-Y--P--K----I---------------------EGNA-----------KLVS------A------------------------------DTDS------F-TF--L------------G---------RFLID-----------------------KQ--------
--------AAA-------LGREVSQKAFNMLKW-LI--------K-----------------------------R-------Q-----------GI--------------------------------R------------N-----G--------D---Q------V--------TVA--W--------A-ISGKAVPSPMKD---------------------------------------------------------------------------LSDEIDWDNLDISADESAVENSDR---------------------------F----------S-----------------------------
-------------------------------------------------S--------PTSSETPDWSTNIGK------PAAQI----IKK-----------------KLH-G-----Y----------------QE---------E------LKN-------H-----E-----------Q-------I--SLIMLDSATP-------GRMALTYYQEF-LPADYFANLDAWV--DD------FS--WY-----------------------------------------------------------------------------------------------------------
YSVEVQNGKKTDKRTQWRGVPP---------------S-----------PYSVA------------------------------DA-------------------------V----------YGKSLSDTL---------------------------------------------------------------------KK-----Q-----------LYARLLPVIAG--G-KSVP----IPDDLMQK-SF-----------------------------------------Q--V------------------------AC--------------------N-----------------------PNGSENWEWQRNIG-----------
---------------V---------------ACAL--------YKG--------W-------------------------------------------R--------------------------------------------------------------------------------------------------------------ARHHDLSERRTYSM-----ALD---------KE-------------NR---S-------------RDYLY------GRLLAVAE-------NTE------------------------------S--------------------------------------------------YALYLVGEKR---------
ATTA---ERYMQRFA-------EHPFAT-------WRNIEL-A-L-K-PYQD------R--L------------------RNNGKDTG---------------------------------------------------------------AQA-I--G-------EIM-------------------DL----------------------------------F------------ENQDFS-SD-D-------KLSGEFLLGYH---C----Q----------------------------------------------------K-------------------------------------------MD----------I-----
----A------------------R-------------------R----------IAEISANKSKTDE-----------------------------------------------------------------------------------------------------------------------
                              'BAS49155.1'                 -------------------------------------------------------------------------------------MSW-------MQKLCRTYE--Y---QE------QGLE------DEDL----------------------------------VLPF-------HMSKAV-H------LKVILNN--K---AE-LV-----------------------GV--------------E------RFEGK-----------------KQ--------------V-PIQVTEKSS------------------------------------K-R-----A--G-----
---------STI----AP-------------YAL--------------HDGLQYI----A--KTA------------------------------------------------------------------------------------------------------------------------------------------------KEYLTIEYLSKVAEKE---------------NG-------------K-------KWKNFLAGTDEDKRNFADTEKTRYKTSF-K---------AY-EE-QLSA--W-
AEA-------------SHLQE-------IKI--VLQYIQK---------------G------SLI----------------------------------------E-D-L----------------------------------EKK------------------------------------------------------------VF------T-L-------ENN---ILLAG--KD-----DPFNFTIVW-AVEIPND--LHSNLW------------S-----KS------S-----------I-KE-QW-----
---------K-----------Y---------------------------------------------QESQSGQEDEKPE---------------------------------------------------L------------CY------------ITG--E---S--S-Y---LANA-Y--P--K----I--E-----------G---N---A------------KLVS------A-------------------------DTD-------S--F-TF--L-------G-RFLID-----------------------------KQ--------
--------AAA-------LGREVSQKAFNMLKW-LI--------K-----------------------------R-------Q-----------GI--------------------------------R------------N-----G--------D---Q------V--------TVA--W--------A-ISGKAVPSPMKD---------------------------------------------------------------------------LSDEIDWDNLDISADESAVENSDR---------------------------F----------S-----------------------------
-------------------------------------------------S--------PTSSETPDWSTNIGK------PAAQI----IKK-----------------KLH-G-----Y----------------QE---------E------LKN-------H-----E-----------Q-------I--SLIMLDSATP-------GRMALTYYQEF-LPADYFANLDAWV--DD------FS--WY-------------------------------------------------------------------------------------------------------------
---------------QWRGV----------------------------------P--------------------SP--YSVADAVYGKSLSDTLK----------K--------------Q--------------------------LYARLLPVIAG--G-KSVP----IPDDLMQK-SF----------------------------------------Q--V------------------------AC---------NPNGSENWE----------------------------QR----------------------------------------Y-------SVEVQNGKKTDKRT-
ACAL------YKG------W-------------------R------------------------------------R----RHHDLSERR---T-----------------Y------------------------------S-----M-------ALD---------KE-------NR---S-------------RDYLY------GRLLAVAE-------NTE-------S-------YALYLVG------------------------------------------EK------------------R-------ATTA---ERYMQRFA-------------
EHPFAT-------WRNIE---LALK-PYQD------R--L----------------------RNNGKDTG------------------------------------AQA-I--G-------EIM----------DL--------F----------ENQDFS-SD-D-------KLSGEFLLGYH---C----Q-------------------------------------------------K-----------------------------MD------------I--ARRIAE------------IS-
ANKS-------------K------------T------------------DE------------------------------------------------------------------------------------------------------------------------------------------
                              'KYK92371.1'                 ----------------------------------------------------------------------------------------------------------------------------FKAYEEQLS------------------------------AW---AEASHLQE----IKIVLQYIQK----GS-LI-------------------E---------DL---------------------------------------------L-E-----------KK-------------------------------------------V----------------
-FTL------------ENNILLA---------G-----------------------------------------------------------------------------------------------------------------------------------------K--------------------------------------------------------------------NFTIV------W-----------A-----------VEI-----------P-----------------------------------------------------
--------------------------------------------------N-D-L-----------------------------HSN--------------------------------------------------------------------------LW-------------S--------KS--------S--------------I-KE-QW-----------------------------------------------------------K------------------------------Y----QESQSGQEDEK----------------------------------------------PE-----------
---------------------------L-----------------------CY------------ITG-----E---S--S-Y---LANA-Y--P--K----I--E------------------------------------------G-----N---A-----------KLVS------A--------------------------------------------DTDG-------F-TF--L----------------G-RFLID-------------------------------KQ-----------AAA-------LGREVSQKAFNMLKW-LI--------K------------R--------------------------Q-------GI------
---------R----------N----G-----D---Q----V-------TVA--W--------A----ISGKAVPSPMK---------------------------------------------------------------------------------------------------------------------------------------------------------------------DLSDEIDWDNLDISADESAVENSD-----------------R---------F-----------S--------------PTSSETPDWSTNIGK------
PAAQI----IKK-----------------KLH-G-----Y----------------QE---------E------LKN-------H-----E-----------Q-------I--SLIMLDSATP-------GRMALTYYQEF-LPADYFANLDAWV--DD------FS--WY--------------QR-----------------------------------------Y-------SVEVQNGKKTDKRT-------------------QWRGV------------------------------P--------------------------P----------SP-----
YSVADAVYGKSLSDTLK--------------------------------K----Q-------------LYARLLPVIAG--G-KSVP----IPDDLMQK-SF-----------------------Q--V--------------------AC------------------NPNGSENWE-------------------------------------------EK---------------R------ATTA---ERYMQRFA-------EHPFAT---------WRNIE---LALK-PYQD------R--L-------------RNNGKDTG---
--------------------------AQA-I--G-------EIM-------------------DL----------------------F----------ENQDFS-SD-D-------KLSGEFLLGYH---C----Q----------------------------K-------------------MD-------I-ARRIAE------------IS-ANKS--------K-----------T------------------------DE---------------------------------------------------------------------------------
                              'SDZ22575.1'                 -------------------------------------------------------------------------------------MSW--------IQKLYDTYE---------T-----NKEL-----A--------GRI-M-G--------EGSPV-------------LLPI--CH----TTQRA-H------ITVTLDK--D--GN-YK-----R-A-------T---------LV-----AP-----------------ED-------------------------SR----T-----I-I-P---CTEK-SA--FR-----------V-G------T--
----K---------P---Q----N----H-------------P-L------------------ADSLQYV----------A----G-------------------------------------------------------------------------------------------------------------------------------------------------------------D-FVD---F------------G------------------F-------------V-TSG-----------------F-S--KKPQEPYLSY-RE-ELGK--W--CCS-------------E-
HAN--PK------VIN---VFKYVQK----------------------G------TLI----------------------------V-------N-D----G----L---CI------------E---N-N----GM----L----L-----E-TWA--KDDKD--NMPELLRIIGN--------DQP-------------QSK-------A----------IRW------A-V--------E-IP-------------G----------------RQSE-------------V-----------------PW-------N-----DP------Q------------I-LQ-AW-------
--------------E---H--------------------------TT----------------------------------------------ID-----------------N-Q---T---L--QD--------------------CY----------VTG-----ET-S-N--LAQG-H--P--A--KI--R---------N-Q-G-G-------DS-A------------KLIS------S-------------------------NDT-----------G------S--Y-TY--R-------G----------------------------------R--------FKG------AS-----
---E---------AAG-------VSFEVTQKAHNALRW-LI-----------E---------------------------------K-------Q-----------GV------------------------------------------------R---------S-----N-----D--------VLILTW------------------------------QK-------------------------------------------------------------------------------------------------------------------------------------------------VE-------G------F-----------------------------------------------------------------
NPPNPADDTASFMAVGIEE-------------------------------------------------------SYTADEFSES-----LRK-----S--------------L-Y-G----Y----------------------QK------------------Q------LE-----S------S-----RN------I-MIMMMDSATP-------GRLSIRLYREM-LPDSFVKRVENWH---------IKTAWI--------------------------------------------------HR---------------------------------------
YRGAKWTDRIKEAVPEEYVSAP---------------S----------AFDIA--------------------------------------------EA-------------------V-----YGKRIDDNL---------------------------------------------------------------------RK-----K------------ALRLVRCLLD--G-Q-K---IPRDMMEA-VI---------------------------R----------RA---------------------------------------------------CNPVGFEASWEWMKALT----------
---------------------V------------ACSM----YRN---V----------------------------------------------------------------------------------------------------------------------------------------EGYALALE-----EK-----------RR----S--------RDYLY------GRLLALAD------SLE------------------E------------------------------------------------------------------------------WALREGGEKR---------QTNA---
LRLMQRFS------ERPFST-------WRNIEL--SL-A-PYKA------R--L-------------------------------GGKGKG-----------------------------------------L----------------------TNR-I--S--------EVM--------------EL--------------------------------F----------------DSEDFV-SD-K-------KLSGEFLLGYH---C----Q-----------------------------------------R--------------------L-------------------------------W-----
-------------K-------------------K-----------------------EE----------------------VEQEQTETIEGGDM----------------------------------------------------------------------------------------------------------------------------------------------------------------
                              'ABQ92671.1'                 -------------------------------------------------------------------------------------MNW------MQSLYETYE-------------Q-------CAKG-----G--------LVGYV-S----------GTKRP----------------LLPI--CH----ITIQA-H------IEVVIDG--C---GN-FR-----R-A---------Q--------LV------SK-------------------------------DD-----------------AV-----T-----I-I-P-----ATES-SA--SR--------S--G--S-
------K---------P--E---C----H---------P-L------------------CDKLQYL----------A-----G------------------------------------------------------------------------------------------------------------------------------------------------------------------D-FVK---Y----------------------------------------V-TSG-----------------F-Q--RNPAEPHQNF-VT-LLSG--W--NS-----------K-
CLH--PK------VSA---ILKYIQK----------------------G------TLM----------------------------A-----D--D-------L----------V-----K-E-G-----G----V-----L-----D-H-R----RL----L--A-----K-DDI--EPEK--RRGTIFDVL---------ND--------QGD------A------V----------VRW------V-V--------E-IP-----------------G-------D---TESR--------------------VW-------R-----DT------K--------I-WD-SW-------
I-------D-------Y-----------Y----------------------------LN-----------------------------------------H-R-------S--GT-SE--------------------L------------CF--------------VIG-------EE-R-T---LTNN-H--P--K--YI--R--------------R-E-G-------DN-A------------KLIS-----S-------------------------NDT------------Q---G----F-TF---R---------G------------------------------R-----------FIT------DE-----
Q---------ACS------VSLEASHKAHYALMW-LI---------E--------------------------------------------K-------Q------------------GY-----------------------------------R-----N---G-----D----L--A--------IVA--W--------AISGATVTVPQPTD----------------------------------------------------------------------------------------------------------------------------------------------------------------
PSDDKSID----------------------------------------------------------------------------------------------------------KIA-G----Y----------------RQ------------------E-------IGD-----H-------E-----------Q-----V--VVMGVDSATP-------GRLAIVYYRTL-KGSDFLDRIETWH--DT-----CA--WR-----------------------------------------------------------------------------------TYRVI--
-------------------MYG------------------------------------------------------------------------------------------------------------------------Q------A---------------TIKRLLPCIVD--G-Q--P---LPRDLVES-AI-----------------------------R-----------A--------------------------------------SN----------------------PVVLSD----------------DQKEKVLT-----------
--------EERDTNGKLYRRRIPFIGA------PA-----PADIAE--------A-AYGRR-------DDRLK---------------------------------------------------------------------------------------------------------------------------------------------------------------------------------------------------------------------ACAL----YRK----Y-
PYLM--------R--L--------------------------------------------------------------G---------------------------------------------------------------------------------------------------------------------ET--------RT--RDYLY------GRLLAVAE-------VIE--------------E--------------R----------------------------ALYKAEQN-------------R-------PTNA---IRYMQQFS-------QRPFRT--------WKQIHD--LL-M-
------RA----------DDREDDNDGTSI----------------------------------------------------------------------------------------------------------------------------------------------------------------
                              'WP_011715062.1__2_'         -------------------------------------------------------------------------------------MSW-------MQMLHTTYE-------------C--------C--------------KDQ------------------LGPI--YH-----TTQQA-H------LEVVLDH--S---GT-FL-----R-A---------H--------VP------SSK--------------------------TD-----------ST---T-----L-I-P----CSEK-SA--GR----------T-----S---------
G---------L--M---N----H-------------P-L------------------ADKLQYV----------A----G-------------------------------------------------------------------------------------------------------------------------------------------------------------D-YLS---F-------------------A----------------------L-SEK---------------E-Q--TKMMESHRLY-KE-ALAT--W--ADS----------P-QSH-
PT-----VKA---IYAYVAQ----------------------G-------RLV----------------------------------D-----D--L-------------V----N-A--G----L----LPC----------V-E-G---SI---P---P-----K-WQ--GPKE-ETPAIYKAV---------TGM--------PLD------A-----F------------VRW------A-V--------E-EPAP------------------G------------CEIE---------------PW-------N-----DP------Q------------I-LQ-AW------
-------D---------H----------Y--------------------------------------------------------------------------------L-----L-----E--ESAPRG---------------------------------------------T-------------------------CM-------VSG--E--ES-V-L--LAAS-H--P--A--KL--R---------H-A-A-------DK-A------------KLIS-----S---------------------------NDS----------------------S-----G-----Y-TY--R----------G---------------------------------------R-------FLS-----AD-----
E---------VCG------VGLETTQQVHNTLRW-LI--------E-----------------------K---------Q-----------G-----------------------------------------R-----N---G-------D----L-----R-----IVA--W-----AVDGRTT----------------------------------------------------------------------------------------------------------------------------------------------------------------------------------------
DNVPAETEEEDEL-LP----TSSMGHSVD-----------------------------------------------------------------------------Q--------S----------E-------F----GLR-L----GKK-----------------S-G---Y----------------YS--QLG-----------D------L------------T--D-------I--VVMGIDSATP-------GRMSIIYYREL-QQTEFLQRLEKWH--SQ-----MA--WP------Q-----------------------------------------------NF----------------------------
KDSQFTGAP---------------------------------EA-------------------------------------------------------------------------Y---GRRLD--------DKLKS----------------------------------------------------------------------------TLQRILPCIID--E--Q--P----FPEDLIHA-VL----------------------------------------------------------------------------------------------------------------KEKCLG----
----------ACALVRGSNLRA-----H------------------------------------------------------K---------------------------------------------------------------------------------------------------------------------------------------------------------------------------------------------------------------------------------------------------------------------------------------------------------------
------------ACALVRGSNLRA-----------H-------------------------------------------------------------------------A---------------M------ALE----------------PD---Y-----TS--RDYLY------GRLLALAD-------YLE---------E-----------------------------------------R----------------------------------------ALHAANEH---------------R-------ATNA---ARLMQRFS--------
QRPNAT---------WLTIYD--AL-L-PYKN-------R--L--------------QVN--KPGLYIN--------------------------------------M---------------MRE-L--D----NVM------------------NL----------------------F----------------I-ADYQ-ND-A-------PLKGEYLLGYH---C----Q--------------------------------------------------------------------------------------------------------------------------------------------R---------------------------AV----------L--------W--AKKGDK------
TG-TQAT--------------------------------------------TE--------------------------------------------------------------------------------------------------------------------------------------------------
                              'CRH05230.1'                 -------------------------------------------------------------------------------------MSW-------MQMLHATYQ-------------Q--------C--------------QEK------------------IPPI--YH-----TTQQA-Q------IEIFIDG--L---GN-FH-----R-A---------E--------VI------TDK-------------------------------KE--------------AT---T-----L-I-P----CTEK-SS--GR--------S--G--S--
-------P----I----N---H-------------P-L------------------CDKLQYV----------A----K-------------------------------------------------------------------------------------------------------------------------------------------------------------D-FLQ---F------------------------------------------------------V-TSG------------------------Y-A--KDPVVPHQLF-TA-DLNR--W--ANS---------A-QTH--PK-
---LKA--IQAYVQK------------------G------RVI----------------------------------E-----D--L-------------L-----I---H-A---G----I-----LPV---------------D-D-A-------PS----R-----N-AWE--GEKE--QEPAIFKMV-------------AKT--------PMD-------------------E-----F----------VRW------V-V--------E-VI----------------G----------------GEST--------------TW-------N-----DP------------I-LQ-CW--------
------E-------------------------------------------------------------S----------L-----------L-----E--S--EIALRG---------------------------------------T-------------------------CM-----VSG--E--ES-V-L--LAAS-H--P--A--KL--R---------H-A-A-------DK-A------------KLIS------S-----------------------------NDS-------------------------S------G------F-TY--R-------G-------------------------------------R-------FTQ------AE----Q--
----VCG-------VGMEATQQTHNTLRW-LI---------E----------------------R--------Q-----------AY--------------------------------------R----------N-----G--------D------L-----Q-----IVA--W--------A-V--NGLHT-------------PN--P---------------------------------------------------------------------------------------------------------------------------L-VNSHDLMMDEED-------D----------------------------ELP-
EPVKPK----------------------------------------------Q--------------QAFGQRLARK------IRG------Y-------------------------------------------------RTQLG----------S------LS-------------------------------I-VVMGLNSATP-------GRMSVIFYREL-KSAEFLDRLERWH--TQ-----TS--WQ---------------------------------------------QN-----------------------------------------F------------GKKLQFEGAP----------
-PRDIA-----------------EA-------------A--YGARLDDKL-------------------------------------------------------------------------------------------KS--S----------S------TVQRLLPCIMD--N-Q--P---LPKDLVDS-VV----------------------------H-------------RVSN---------------------------RV---------AYERGRNGYQWEWEKHLG-----------------------------------------------------ACAL----YRG-----F--
----------MQL--------------------------------------------E-------------------------------------------------------------------Y-----------------------------------------------------------------------------------------------------------------------------------------------------------------------------------------------------------------------------------------------------------------------------------------------------
------------MQL---------------------------E---------------------E--------------Y-------------T---M--ALD----KS--------------------R-----------RDYLY------GRLLALAE-------HIE-----------------S--------------------VLNDA-----------------------------------------------EK--------S---------R-------PTNA---VRLMQRFS--------QRPNAS-----------WLTIHQ--AL-L-
PYKN---------R--L-------------QVNHPGQLVH-----------------------------------------------------------NRE-I--Q---------EVM---------D-L------F------------------------N---H-DEYLD-D-------SPLSGEYLLGYH---C----Q---------------------------------------------------------------------------------------------------------------------------------------------------------------------------------------
-------------KSDKNASGDKA------------------------------------------------------------------------------------------------------------------------------------------------------------------------
                              'TDY62978.1'                 -------------------------------------------------------------------------------------MSW-------MQRLYETYE-------------L------A-----M-------KLPEQ-G------------------ENPRP--------------I-PT--SH-----ISQQA-Y------IEIVLDE--K---GQ-LL------N-A-----------S-----------VV------NK------------------------ED-------------------T------L-L-P----ATED-SA--TR---------S-S-G--G--
---A-----------P-----------H------------P-L------------------CDKIQYV----------A----A------------------------------------------------------------------------------------------------------------------------------------------------------D-YPT---Y----------------------------------K-PSY--------------------F-EDFKSGSENKLGY-RS-LLVK--W--HEF-------------S--RH-
PK-----LKA-----IRTFIEE------------------G------RVV--------------------------------------------------V-------------G-Q----G-----L------VHL-----------------D-E-K-----GK-----L---C----T-SWP---DAK-TMPPLFKLLTKK--------KEG--------DDTVQDQGDA-------M-----------VRW------K-V--------R-QP----------G----------------LEDR-------------TW-------K-----DK-----------L-IT-SW--------
-------K-------------------------------------------------------ET-----------------------------------L-Q-------G--E--NG-----------------------------------------CM-------VTG-----EL-L-P--LGTK-H--P--A--SI--R---------H-S-G-------DK-A------------KLIS-----S-------------------------NDT----------T------G-----Y-TF---R--------G--------------------------------------R----------FTD------
GG------Q---------ACT-------VGYEVSQKAHNALRW-LI--------E-------------------R--------Q----------AF-------------------------------------L-Q------G---E--NG---------------Q---V---------IVS--W-----------------A---------------------------------------------------------------------------------------------------------------------------------------------------------------------
GT----------------------------------------------------------------A-----------------N--SLEAF------FEEETGESL-------------------------------A-N-G------TVIGDIGQEYALR-----LRKKIAGWSAKLGD-------A----D-----N----K----I--VVLGLDSTIPG--G-RMAITLYREL-KGSEFLRRLENWH--SS-----FA--WP--------------------------------------------------Q-------------------------SKDIQFVGAP----
----------PKDIA-----------------------------------------AFGRRI---------------------------------------------------------------------------DPKLSKS-----------------------TVNRILPCIVD--G-S--P----IPRDLVRSVCI-----------------------R----------------A------------------------------ST-------------------------------------------GRAGLEFWEWFKVLG----------------------ACGL------FRG--S-
PYMT--------R--L-----------------------------------------------------------------------------------------------------------------M----ALE-------TE------------------RR---T---------RDYLY------GRLLAIAE-------NIE--------S------------------------------------------------------R--------------------------------R------AFLS------------------GE-----------K------DTNA---AKLMQRFS-------ERPNST------------WLLIRQ--SL-T-
KETDAQGPDTDITVQEEE-------------------------------------------------------------------------------------------------------------------------------------------------------------------------------
                              'CUS33063.1'                 -------------------------------------------------------------------------------------MSW-------IQKLNETYE-------------L-----------C--------A--------GLPQF-S------------SN--P------------LLPI--SH----TTQQA-H------IEIVIDG--S---GN-FR-----R-A---------N--------VV------SK--------------------------AD-------------------QT----T-----L-I-P------CTEE-SG--GR-----------A-G---S---
------K--------P--K---N----H------------P-L------------------CDKLQYV----------A----G-------------------------------------------------------------------------------------------------------------------------------------------------------------D-YWE--L----------------G-------------------------------V-TIG-------------------------F-A--KEKEEPHCSY-LK-DLEA--W--GAS---------P-EGH-
-YK-----LAA---ILTYVRK-----------------G------K--------HVV---------------------------------------------V--------V--R-E--R------R--L--LPL-----------D-S-Q-----GK---L---L-----K-KWN--GDKK--NVPAIFKVIPA----------GSS-------------PED----------A-----F------------IRW---------H-V--------E-LA----------------N-----D-----PETA------------------------------IW-------K-----ED-------E-------I-ID-AW-------
-------K---------Y-----------------------------------------------------------------AS------------------------------------Q-G-Q-Q----S--K-RG-----------------------------------------CM---------VSG-----EE-R-T---LAEQ-H--P--A--KL--R---------N-A-G---DK-A--------KLIS-----S-------------------------NDT---------------------Q-----G-----Y-TF---R----------G------------------------------R--------FSD-----AV-----E---
---------ACG-----VGFVVTQQAHKALQW-LL-----------D---------F-----E-----------R-----Q---------AF----------------------------------R-------N---Q--------V------------------IVA--W---AV--VSGKPL---------------------------------------D---------------------------------------------------------------------P---------------------AN-----SAQ--------M-------F----------LESREE-
DA----TPVYQGDAG------------------------------------------------------------------------------------------------------------------------------------------------------A-G----Y-----------------RT--RLG-----------------D---------------A--------I--VVMGLDSATP-------GRMAITYYREL-TGSEFLERIQLWH--ES-----CA--WH------------QS----------------------------------------F------------GKNLKFVGVP--------
----PRDIT--------------EA-------------A---YGRQVE--GKSGEKLRK-------------------------------------------------------------------------------------------------------------TVERLLPCIID--G-R--P----IPRDLVES-TV----------------------------------------------------------RA-------------------------------------------VNRLGLEKWEWEKNLG------------------------------------ACAL------FRG---H--
---------K---------------------------------------------------------------------------------------------------------------------------------------------------------------------------------------------------------------------------------------------------------------------------------------------------------------------------------------------------------------------------------------------------------------------
PYKA---------R--L-----------------------RTRLPGFLVN--------------------------------------------------------EKL-L--D----------GII-----------------CT---------------------------F------------G-------G-QDFMS-E-------ARLSGEFLLGYH---C----Q--------------------------------------------------------------------------------------------------------------------------------------------------------------------------------------------------------------------------------------
---PESTNAEESTETQTDQGE--------------------------------------------------------------------------------------------------------------------------------------------------------------------------------------
                              'PTM63281.1'                 -------------------------------------------------------------------------------------MSW-------IQKLYETYE-------------Q----------A--------------SNPAF-L--------------NDKIP-----------LLPI--SH-----ITQQA-H------IEIVIDG--Q---GN-FL-----R-A---------S--------SV------PK-------------------------------------------------------EV---T-----I-L-P-----ATEK-SM--GR--------R-T--S-
G------V-----A----P---Y-------------P-L------------------CEKIQYC----------A-----G-------------------------------------------------------------------------------------------------------------------------------------------------------------D-YAK---Y----------------------------------------------------------K-KSH-----------------F-A--A-------Y-SE-LLES--W--CTS---------R-FSH--PK-
AKA----VFEYISK-----------------E------TVV-----------------------------------------------A-----D-----L----V-----H-E--H----V---LYL-----------D-G-D------EH------L---L-----T-SWK--SGTP--APDIFRYL------------TP----------KEGLRDQGDA------L----------VRW---------R-V--------E-MP------------------G-----D------ASPA-----------------TW-------D-----DH-------------T-----L-MD-AW--------
-----------K---------Y-------------------------TS---------------------------------------------------------K-E-------A----V--SG-------------------------------L--------CM------------VTG-------EE-R-T---LAEQ-H--P--A--KL--R---------H-S-G-------DG-A------------KLIS-----S-------------------------NDT----------T----G-----F-TY--L---------R----------------------------------------------R------FTDTKDNAK--------
G-------Q-----ACT----VGAEVSQKAHNALRW-LV----------G------------------R---------R--------Q-----------AY--------------------------R-------------Q---A------VVA--W---A---VSGKRIPD---------------------------------------------------------------------------P--------------------LA-------N-----S--------N---F------------------------DL---FGI------
GTEELEPRQGQVYQGD------------------------------------------------------------------------AGQ---AFGQR------LAK----I-----------------------L-A-G------Y---------------------QAELGS-------------------I----E---I--VVMGL--DSA--TDGRIAITFYREL-TGSEFVERLVWH--KN-------HA--WR---------------------------------------------------------------------------------------------------------
SRDIKFIGAP----------------A------PKDIA----------------EA-------------A--FGRRLDEKL--------------------------------------------------RK---A-----------TVERLLPCIID--G-Q-K---IPRDLVES-TV---------------------------R------------------A--------------SNRVGLEHWEWEKYLG---------------------------------------------------------------------------------------------
--ACAL------FSG------Y-------------------H------------K---------------------------------------------------------------------------------------------------------------------------------------------------------------------------------------------------------------------FALDMTKESR---------DTSA---AKLMQRFA-------DRPNST---------
---------WRNIEL--SL-T-PYKS------R--L-----------------------RSRAPGFLYT------------------------------------------------------------------R---------ERQ-L--D-------EVT---------------CL----------------F----------G-------G--DDFMD-E---------RKLSGEFLLGYH---C----Q---------------------------------------------------------------------------------------------------------------------
----------------KPETEQDENIEATVAETN-----------------------------------------------------------------------------------------------------------------------------------------------------------------------------------
                              'PXW81496.1'                 -------------------------------------------------------------------------------------MSW-------IQKLYDTYE-------------Q----------C-------A-------GQADL-S-----------NAESS----------LEPI--CH------TSQQA-H------IEVTIDG--K---GN-FL------R-A-----------N----------VI-------SK----------------------------------------GQ---------------------GV------T------L-V-P-----CTEA-SA--GR-----------A-G---S-----
------K--------P---T----N----H------------P-L------------------CDKLQYL----------A-----G-------------------------------------------------------------------------------------------------------------------------------------------------------------D-FLK---F------------------------------G----------------------V-TSG--------------------------F-A--NEPTEPHKSY-LS-LLSD--W--ANS-----------S-
FTH--PK-----IIA---IQAYVQK-----------------G-------HLV----------------------------------S-------D--L-------------V-----R-E--R------R--I--LLT-------------D-E-K-----NQ----L---L----T-EWG--GDKK--EAPSVFGVMPN---------GNS--------------PDG------A-----V----------------IRW----------R-V--------E-IP----------------G-------D-----PLSE---------------------------TW-------K-----DR------G--------L-MD-AW------
-----I-------A------F---------Y----------------------------------------SS----------------------------------------------Q-K------D--S--QG---------------------------------------------CL-----------VTG-----EI-L-S--LAEQ-H--P--A--KL--R---------H-G-A-------DK-A------------KFIS------S-------------------------NDT-----------S----G--------F-TF---R-----G-------------------------------------------R-----------FID------AN-----
```

[illegible]

'PTR08404.1__3_'

[illegible]

'SCX54583.1'

[illegible]

'RAR82230.1'

[illegible]

'TDP13375.1'

[illegible]

'SDX66139.1'

[illegible]

'ABD71602.1'

[illegible]

'ACM21757.1'

[illegible]

'TEB12458.1'

[illegible]

'EIW25544.1__3_'

[illegible]

'ADY55248.1'

[illegible]

'KGP75382.1'

[illegible]

'KPL71271.1__2_'

[illegible]

'REG10371.1__2_'

[illegible]

'WP_008910924.1'

[illegible]

'WP_058798750.1'

[illegible]

'WP_046007366.1'

[illegible]

'WP_061899414.1'

[illegible]

'SEG74290.1'

[illegible]

'WP_094144966.1'

[illegible]

'WP_100754455.1'

[illegible]

[illegible]

---L---DKK-L--Q---KLC---E---GI---EIKNF---P---V---T---FT-LE-E---QGQFAVGYY---H---Q---K---QA---Q---F---N---R---TELRETVEEE
[illegible]

'SMC05801.1' ---M---IRELVDYAR---T-H---NL [illegible]
G---QPR---C---H---F-L---IETVAVV---T---N---W---P [illegible]
LAE---WERCLQD---P---WIL---E---K---I---R---A---DLK---H--Q [illegible]
-L---QSP---GE---E---S-I---G---T [illegible]
IGRHVAVAYQNALNA-LI---Q---R---G---L---TL---S---G---I [illegible]
---R-Q---A---L---F---EAERG---LRQ---V---L-T-G---G---E--P [illegible]
---RQ---L---TVRKDEEVP---AP [illegible]
R---N---RGGNRIM---N---R [illegible]
---ESNQLKQW---F---RNH-L--M---EVT---N---RI---TAI [illegible]
[illegible]

'ABF44540.1' ---M---LAA [illegible]
P---KALGQT--QA---A---H---F-L---ADSCGVI---A---L---L---P [illegible]
-Q--LLP---VLEVLRD---P---AE---L---A---ELR---A--EL--A---QRG [illegible]
---F---GQP---TT---T---G-T---M---L [illegible]
VSEENAAAYRAALEH-LL---A---Q---A---PV---L---G---Q---M [illegible]
PEQQA---A-V---A---H---T---RAQQL---LSA---L [illegible]
FFSLL---MN---I--Q---RPKPQS---TSMDDYLKP [illegible]
[illegible]

'WP_114798653.1__2_'
[illegible]

'WP_119277146.1'
[illegible]

'ADH65195.1'
[illegible]

'ADD28249.1__3_'
[illegible]

'WP_027888790.1'
[illegible]

'RKX23015.1'
[illegible]

'RLI08120.1'
[illegible]

'AFM23520.1'
[illegible]

'WP_119629531.1'
[illegible]

'WP_086135264.1'
[illegible]

'WP_077731139.1'
[illegible]

'WP_092052304.1'
[illegible]

'BBJ22385.1'
[illegible]

'WP_093032550.1'
[illegible]

'WP_043753350.1'
[illegible]

'WP_114446098.1__2_'
[illegible]

'WP_120682443.1__2_'
[illegible]

'CQR65546.1'
[illegible]

'WP_100606285.1__2_' [illegible]

'AIA06578.1' [illegible]

'SDT83407.1' [illegible]

'WP_078950659.1' [illegible]

'BAK19890.1' [illegible]

'RAJ47990.1' [illegible]

'WP_078552483.1' [illegible]

'WP_079100778.1' [illegible]

'SKA36544.1' [illegible]

'SDQ60927.1' [illegible]

'RAS59733.1' [illegible]

'WP_015099451.1' [illegible]

'ROP21220.1' [illegible]

'ROO60289.1' [illegible]

'SDZ24145.1' [illegible]

'SCF19379.1' [illegible]

'ABD12700.1__2_' [illegible]

'KDA41608.1' [illegible]

'OAA19957.1' [illegible]

'SFO06681.1' [illegible]

'RCW45125.1' [illegible]

[illegible]

'SDP96777.1' [illegible]

'SDQ77537.1' [illegible]

'SDZ50827.1' [illegible]

'WP_121221044.1' [illegible]

'SER77926.1' [illegible]

'WP_075975713.1' [illegible]

'SES36449.1' [illegible]

'ACO32176.1__2_' [illegible]

'EPR34125.1' [illegible]

'EPR43457.1' [illegible]

'WP_090718871.1' [illegible]

'WP_101908721.1' [illegible]

'PWM53492.1' [illegible]

'WP_006599002.1' [illegible]

'WP_016280412.1' [illegible]

'WP_117939087.1__2_' [illegible]

'WP_117897007.1__5_' [illegible]

'WP_117876534.1__2_' [illegible]

'WP_041337294.1' [illegible]

'WP_118158046.1__2_' [illegible]

[illegible]

'WP_118160818.1__2_'

[illegible]

'WP_118161860.1__2_'

[illegible]

'WP_118123679.1__2_'

[illegible]

'WP_117904369.1__2_'

[illegible]

'WP_117958132.1__3_'

[illegible]

'WP_118054167.1__2_'

[illegible]

'WP_117863908.1__2_'

[illegible]

'WP_117857745.1__2_'

[illegible]

'WP_117928221.1__2_'

[illegible]

'WP_117923124.1__2_'

[illegible]

'WP_117891632.1__2_'

[illegible]

'WP_019679807.1'

[illegible]

'WP_015558733.1'

[illegible]

'WP_054685123.1'

[illegible]

'WP_117968067.1__2_'

[illegible]

'WP_118446305.1__2_'

[illegible]

'WP_088109963.1'

[illegible]

'PWM22000.1'

[illegible]

'WP_091649542.1'

[illegible]

'WP_008718025.1__2_'

[illegible]

[illegible]

'RKJ34427.1' [illegible]

[illegible]

'WP_117767916.1__2_' [illegible]

[illegible]

'WP_096240537.1' [illegible]

[illegible]

'WP_087312701.1' [illegible]

[illegible]

'WP_087279829.1' [illegible]

[illegible]

'WP_087281069.1' [illegible]

[illegible]

'WP_080908073.1' [illegible]

[illegible]

'WP_076349607.1' [illegible]

[illegible]

'WP_094414332.1__3_' [illegible]

[illegible]

'WP_095738147.1__2_' [illegible]

[illegible]

'WP_009278655.1' [illegible]

[illegible]

'WP_118343116.1__2_' [illegible]

[illegible]

'WP_118110387.1__2_' [illegible]

[illegible]

'PWM50412.1' [illegible]

[illegible]

'WP_087410535.1' [illegible]

[illegible]

'WP_050340707.1' [illegible]

[illegible]

'WP_021625749.1' [illegible]

[illegible]

'WP_016478057.1' [illegible]

[illegible]

'WP_117799690.1__2_' [illegible]

[illegible]

'WP_114549443.1__2_' [illegible]

[illegible]

RK----------S--------NA-----G-----L--SI-------K------------L-------------EKE-M--D-------GIM---------P----LL------G---ES---------C----F [illegible]
---------SS---------------S--DAAS---TETS----------------------GE--------------------------------

'WP_080143417.1' ------------------------------------------------------------MI------LQELNKYYE [illegible]
--------M-----------L----P---F---------F-L------------------CDNAAYL------L---G------------------------Y---------------DE [illegible]
G--ARA---LLRFFER---EDR----------D-----AGL-------D--------E---A-----V-----R------------------------------------------A--E-I [illegible]
----------------------------------------------R---------AA---------G-----------------E-V-------------V-------G-----------------R [illegible]
ISESVAFNAGSALRQ-LC---------A-----D----------P-----G-----H----------RV-----------------------R---------I----G--------E---T---M [illegible]
-------------------------------------------T---------V-----------E---KVHKA-----LQN-----M---------------------E-L-G-----L------PL--G [illegible]
--LQ-----------T---A-------------------P-------------------M------G------A-------------------S--------D--Q------------I----PS----T [illegible]
-----R-----------K--------A---------RLGSEGSNERDGVANADDE-------------------------R--------------G---------------L---------------T [illegible]
----RK-----------F---------NP------G-----L--CV-------K-------------F--------------EKA-N--D--------EVM---------G-----MI------D----GP---------D [illegible]
----D--------DE-----------------G--EPAD---AEPA---------------------VKDN---------------------------------

'WP_083833078.1' ---------------------------------------------------------MGGGGPMI------LQELNN [illegible]
--G---------M-----------L----P---F---------F-L-----------------CDNAAYM------L---G----------------------Y--------------DE [illegible]
E---G--AHA---VLRFFER---DDR----------D-----ADL-------D--------E---D-----A-----R-----------------------------------------T--E-L [illegible]
-------------------------------------------------K---------AV----------G-----------------E-V-------------V-------G-----------------Q [illegible]
ISERVAFNAGSALRQ-LC---------A-----D----------R-----T-----H----------RV-----------------------R---------V----G--------E---T---T [illegible]
--------------------------------------------V---------V-----------E---RVHNA-----FES-----M---------------------E-R-G-----L------PL--Q [illegible]
----LQ-----------T---A-------------------P-------------------M------G------A-------------------A---------D--Q------------L----PS----T [illegible]
-------R-----------R--------A---------RLGAERANEQDGAVRVDDE-------------------------R--------------G---------------L---------------K [illegible]
--------RK-----------F---------NP------G-----L--NV-------K-------------F--------------EKA-N--D--------EIM---------G-----LL------E----GP [illegible]
DS--------E--------AD---------------A--DA-T---ADDN---------------------NETN---------------------------------

'WP_117752676.1__2_' ------------------------------------------------------------MI------LQELNNYYE [illegible]
[illegible]

'WP_080797040.1'
[illegible]

'WP_087193631.1'
[illegible]

'WP_087190837.1'
[illegible]

'WP_123673816.1__2_'
[illegible]

'WP_096226813.1__2_'
[illegible]

'WP_052832434.1'
[illegible]

'WP_101432863.1'
[illegible]

'WP_110452643.1__2_'
[illegible]

'WP_029678125.1'
[illegible]

'KJY63744.1__2_'
[illegible]

'WP_110413425.1__2_'
[illegible]

'WP_121913646.1__2_'
[illegible]

'WP_100512974.1__2_'
[illegible]

'WP_034526776.1'
[illegible]

'WP_051917393.1'
[illegible]

'WP_094730565.1'
[illegible]

'WP_117675351.1__3_'
[illegible]

'WP_118262061.1__2_'
[illegible]

'WP_115769061.1__4_'
[illegible]

'WP_101027957.1'
[illegible]

'WP_080828919.1'
[illegible]

'WP_116868643.1__2_'
[illegible]

'WP_074740653.1__2_'
[illegible]

'RHH02629.1'
[illegible]

'WP_025221978.1'
[illegible]

'WP_051994509.1'
[illegible]

'WP_113017463.1__2_'
[illegible]

'WP_131012511.1__2_'
[illegible]

'WP_094049066.1'
[illegible]

'WP_020041696.1'
[illegible]

'WP_014094599.1'
[illegible]

'WP_005806400.1'
[illegible]

'WP_042294562.1'
[illegible]

'WP_014017910.1'
[illegible]

'OYV04502.1'
[illegible]

'WP_134329556.1'
[illegible]

'WP_022636788.1'
[illegible]

'PIE28683.1__2_'
[illegible]

[illegible]

'WP_013253668.1'

[illegible]

'WP_099521097.1'

[illegible]

'WP_056042046.1'

[illegible]

'WP_134754760.1'

[illegible]

'WP_090781650.1'

[illegible]

'WP_051237221.1'

[illegible]

'WP_094045368.1'

[illegible]

'WP_051318558.1'

[illegible]

'WP_019007803.1'

[illegible]

'WP_106766165.1'

[illegible]

'WP_119734915.1__2_'

[illegible]

'WP_035319231.1'

[illegible]

'WP_025698336.1'

[illegible]

'WP_052769938.1'

[illegible]

'WP_087796888.1__2_'

[illegible]

'WP_099448714.1__2_'

[illegible]

'WP_040197366.1'

[illegible]

'WP_081406721.1'

[illegible]

'WP_109121609.1'

[illegible]

'WP_114859321.1'

[illegible]

VSARVAHGYTAALNH-LL [illegible]

[illegible]

'WP_085555889.1'

[illegible]

'WP_109905104.1__2_'

[illegible]

'WP_097278400.1'

[illegible]

'WP_008104116.1'

[illegible]

'WP_089408953.1'

[illegible]

'WP_102741617.1__2_'

[illegible]

'WP_102734061.1__7_'

[illegible]

'WP_102734693.1__6_'

[illegible]

'WP_102736643.1__2_'

[illegible]

'WP_102725968.1__2_'

[illegible]

'WP_102722930.1__4_'

[illegible]

'WP_102727518.1__2_'

[illegible]

'RMF16839.1'

[illegible]

'WP_008664695.1'

[illegible]

'WP_014436071.1'

[illegible]

'WP_105332124.1__3_'

[illegible]

'WP_013630167.1'

[illegible]

'REK29138.1__3_'

[illegible]

'RLT05969.1'

[illegible]

'WP_007363024.1__2_'

[illegible]

[illegible]

'WP_029134845.1'

[illegible]

'WP_007042344.1'

[illegible]

'WP_015256024.1__2_'

[illegible]

'WP_122962942.1__2_'

[illegible]

'WP_116043347.1__2_'

[illegible]

'WP_052147155.1'

[illegible]

'WP_083574374.1'

[illegible]

'WP_109430684.1'

[illegible]

'WP_047071596.1__2_'

[illegible]

'WP_087346068.1'

[illegible]

'WP_092373224.1'

[illegible]

'WP_051658238.1__2_'

[illegible]

'WP_114582889.1__2_'

[illegible]

'WP_031587722.1'

[illegible]

'RMG06604.1__2_'

[illegible]

'WP_013559648.1'

[illegible]

'PIY22877.1__2_'

[illegible]

'WP_013969347.1'

[illegible]

'WP_061914945.1'

[illegible]

'WP_132779610.1__2_'

[illegible]

[illegible]

'PWH13943.1'

[illegible]

'WP_082174620.1__2_'

[illegible]

'RJQ41526.1'

[illegible]

'WP_048513299.1'

[illegible]

'WP_118287108.1__2_'

[illegible]

'WP_075580441.1'

[illegible]

'WP_012938135.1'

[illegible]

'WP_087477553.1'

[illegible]

'PWL92658.1'

[illegible]

'WP_051127501.1'

[illegible]

'RKJ31574.1'

[illegible]

'WP_114620951.1__2_'

[illegible]

'WP_123197687.1__2_'

[illegible]

'WP_039687910.1'

[illegible]

'WP_123208911.1__2_'

[illegible]

'WP_102373465.1'

[illegible]

'WP_015760874.1__7_'

[illegible]

'WP_086414482.1__3_'

[illegible]

'WP_092196989.1__2_'

[illegible]

'WP_087196499.1'

[illegible]

'WP_009304158.1__8_' [illegible]

'WP_103265261.1__2_' [illegible]

'WP_102410316.1' [illegible]

'WP_009646998.1__2_' [illegible]

'WP_006194036.1__2_' [illegible]

'PWM55983.1' [illegible]

'WP_101909151.1' [illegible]

'WP_099204777.1' [illegible]

'WP_090303951.1' [illegible]

'WP_090247217.1' [illegible]

'WP_118476386.1__2_' [illegible]

'WP_014016932.1__2_' [illegible]

'WP_054336445.1' [illegible]

'WP_027895263.1__2_' [illegible]

'WP_095340745.1__2_' [illegible]

'WP_118022269.1__2_' [illegible]

'WP_020310786.1' [illegible]

'WP_009213700.1' [illegible]

'WP_021774881.1' [illegible]

'WP_118336222.1__2_' [illegible]

'WP_087165200.1' [illegible]

-AEA---VKAYFET----WN----------P------GQA--------S-------------E------N------P-------------------V----L--QP--------HL--------------D--E-M----------M---A-----------------------------------GAN------------------L--------------------VF-----L-W-----------------------K--G-------------R------Y-------VQ------E----DS------E----------V-QN-AW--------E--------SY---------R----------------------Q-----------Q----
----------------------------------------------------D----------EQ----------E---------------------A-E--------------------------G------------------I----------------CL-----------VTG--K---R--S-S---IARI-H--S--N---I--K----G-----L--Q--G--A--QS--SG-A---------------ALVS-----F-----------N----------A---------P-----A-----F-ES--Y--------G------K------------------E--------Q-------SY------------------NAP------
VGNYAVYAYTTALNY-LL---------K------D-----------R------K------K-----------MS--------------------------Y----------V-----G---------D----T----A--------VVY--W----------A------------E--------------N---------G-------------------------------------------------------E-------------N--------E--------YR------S--------V------F--------A------------K----------S--------NL--------------------NAP-------
------------------------------------------N---------Q------------E----LVHGV-----FHN-----L--------------------------E-L-G-----R-------AV--D--L---EA-----V---QD----KL-D------P-----E-----Q-------K------F--CILGI---APN--A-ARIAVRFFYTD-SFGNILRLHKEHY-DR------ME--IV---------R-----------------PA-------TD------T-----LR----Y--------L----------G-----------------VWRMM---------
------QE------------T---V---------N----------Q--------------------------K------S--------K-------------D---------K----------K------------------P-----AA----G------------------LTAQVYQSIIS--G-G--R---YPDGLYQA-VL---------------G--------------R-I-------------RA-------------------D-----------QDNS----DQRIYKIT---------R-----G---------KAAV-------IKA-------Y-------------------L----------L-------
------R------------N---------T-----S--Y-E---------------R----------------------------E----------------V-------------------L------------------T------------V----ALN--------E--D---------CK-----I------------PAYVA-----GRVFAVLE-----EIQ------------E------------D------------------A-------------N----------PG------I----------N--------STIK----DRYFNSAC------ATPGVV-------------FPILMK--LK-N-SHIR------K--L------ES----------
-K---------------G------M--RI--------Y-------------F---------------EQK-L--T--------ELE----------G-----KL------E----IQEDGT------I-------W-----P-------R-----R---LS-LE-E---------QGAFILGYY---H----Q-------------------------------------------------------------QK-----------R------Y----------E--------K-----------------K----------E-------------EQ------
-------------E---------------------------------------------------------

'WP_120473328.1__2_' ---------------------------------------------------------MI------LQSLVEYYG---------T-----L-----------A----K------------D-------------GE-------------ITEP--GW---CKA-KV-S-----YALDIGE--N--GE-LK----G------L---I------------PL------K---EEEK---------K-G--------KK---------T-VM--V---P--KN---I------K-V-P-----E--M-DT--TR-----------A----S--G--------
-----V------------K----A----N----------F-L---------------------CDNSSYF--------L----G----------------------------I-------------------DN-----------KG---------------------------------------------------------K-P-------------E-----------R------------------T---------------------------K----------E---------C----------------F-E--------DA-KK-KHKI--V--LED-------I---K---S--K--A-
AIS---VLKYFET----WN----------P------EQA---------M--------------E------H------T-----------------------E-----L---KE---------DL-------------K--E-L-----------M---T--------------------------------------SN---------------------I-------------------VF-----Y-V--------------------D--G-------------I-----Y------AH---------E---DE-----E---------V-RK-AW---------E--------NY---------L-------------I------------T---------
---------------------------------------------H---------GE------------E-----------------E-E-----------------------G---------------------I----------------CL-----------VTG--Q---I--T-K---IAKK-H--G--V---I--K----G-------V--Q--G--G--QQ--NG-G--------------VFVS-----F-----------N----------E----------N----S--F-ES---Y---------G------K-----------------K--------S---------NL------------------NAP-------
IGTYAVHAYTTALND-LL----------Q-------D-----------R------S------H--------------TT-----------------------------I-----------I------G-----------D-----T------T-------IVY---W-----------S------------E-------------R-----------G--------------------------------------------------E--------------YK----------N--------V------F--------A----------------AV-------S----E--------PT---------G--------D---------
---------------------------------------N-------------Q-----------K-----VVDGV------FKN------L----------------------------Q-A-G-----K-------GV---A---V---ED------V----------SL-D-------L-----N-----Q-------R-------F--YILGI---APN--A-ARLAVRFFYQD-SFGNILNNLKNHY--ER------MN---IV-----------R-------------------PA--------SD---------T------WE----Y---------------L-----------G-----------IWRML---------
QE------------T---V------------N-----------K--------------------------K--------S--------R--------------D-----------K------------K------------------P-----VS------N--------------MAGAVYRGIIT--G-G--Q---YPISLYQS-VL----------------L------------------R-I---------------RA---------------------E-----------QDDS------DSHIYKIT-------------R------G-----------------RAAI-----IKA----F-------------L--------------M-------------
K------------N-----------L-----H-----------------------K---------------------------------E-----------------E-----------------L------------------T-----------M-----ALN---------E--K----------SQ------N-----------IAYTL-----GRAFAVLE-----AIQ--------------E---------------D------------------A-------------N----------PG------M---------N-----ATIK----DKYFNSAC-------ATPAAI------------FPILFK--LK-N-SHIR-------K--L--------DN-------K-------
------------G-----K--EI---------Y----------------------Y--------------------EKM-L---I----------ELQ----------E-----KL-------S------LEEGQV--------T--A-----Y-------P-------R------R---LS-LE-E---------QGMFILGYY----H----Q------------------------------------------------------V-----------------QK-------------R-------Y----------E------------K--------------------K-------------TK-------------E-----------EE-------
-----K----------------------------------------------------------------------

'WP_028530336.1' ---------------------------------------------------------MI------LQSLVQYYE---------A-----L-----------V-----E-----------R-------------GE--------------VTKP--GW---CRA-KV-S-----FALNLSA--E--GD-LT----G------V----L-----------PL------R---VSEE----------R-G--------KK--------T-VM---I---P--QQ----M------V-V-P-----E--M--I--SR-----------S----S--G------
-------V-------------A----A---N-----------F-L---------------------CDNSSYF--------L----G----------------------------I-------------------DN-------------KG-------------------------------------------------------K-P------------------------D----------------------R----------------S---------------------------E---------------------------------F-L----------CA-AQ-KHQD--V--LGQ----------A------D-----S--K-
-A--AQA---VIQYFEK-----WQ----------P-------EKA---------A-------------E------S-------D-------------------------V-----L---KE-----------EL----------------E---D-I-----------M---A-------------------------------------GAN------------------L----------------IF------S-V-------------------------E---------V------------YR----------S-------------V-----F-----------A-----------------K---------------S--------NL---------------NAP-----
--------------------------------------------D-----------EH------------V--------------------------R-S-----------------------G------------------T--------------CL----------VTG--E---K--T-Q---IART-H--G--L---I--K----G-----V--P--G--A--QS--SG-A-----------------SLVS-----F-------------N-----------A-----------P-----A-----F-ES--Y--------G------K---------------E---------Q-------SY---------------NAP------
VGNYAVYAYTTALNY-LL-----------A------D-------------R------Q------H---------AT--------------------------------------------------G---------D-----T-----M---------IVY---W----------S--------------E-------------N---------G-------------------------------------------------E----------------YQ----------N--------V-----F----------A----------------CA-------T-----E--------PT---------M------D---------
-----------------------------------------------N-------------Q---------E----IVDGV------FKN-----L------------------------T-M-G-----R-------AI---V---V---DE------V--------AL-S-------P------N-----Q-------R-------F--YILGV---SPN--A-ARIAVRFFYQD-SFGNIMKNIKAHY--GR------ME---IA----------R-------------------PE-------AD---------Q-----LE-----Y-----------------L------------G-----------IWRML----------
-------QE------------T---V-----------N------------K-----------------------K------S----------R--------------D----------K------------K-------------------P------VL------S-----------LPGAVYQAVIS--G-G--R---YPASLYQA-VL-------------------G--------------R-I------------------------RA--------------------------E------------QDDS-----DSRIYKIT-----------------R------G---------------RAAI-----IKA-----Y------------L---------L---------
--K-------------N-------G-----H---Q-N------------K-------------------------------E-----------E----------I-------------------L------------------T---------------M-----ALN---------E--N-----------ST----N----------VAYTL-----GRTFAVLE-----AIQ--------------E--------------D------------------A--------------N----------PG-----I----------N-------ATIK----DRYFNSAC-------ATPASI-------------FPVLFK--LE-N-SHIK-------K---I-------SS----------P---
--------G-----T--KI--------H-------------Y------------------EKM-L---G--------ELQ---------E-----KI--------Y-----VSDEQR------D--A-----Y-------P-------R------R---LS-LE-E----------QGMFVLGYY----H----Q----------------------------------------------------------------T--------------------RK-----------R--------------E--------------K--------------------K---------------AK---------------E----------E----------
-------------------------------------------------------------------------------------

'WP_110324406.1' -----------------------------------------------------------MI------LQSLVTYYE---------S-----L-----------E-----R------------K---------------GK--------------ITSP--GW---CSA-KV-S-----FGLELSE--E--GE-LI----R-------V--I-----------PL------R---RSDE---------R-G---------KK--------T-VE---V----P--SI-----W------R-V-P------Q--M--V--AR-----------S----S--G--------
---V---------------S---A---N-------------F-L---------------------CDNSSYL--------L----G----------------------------I-------------------DN---------------KG-------------------------------------------------------K-P------------------------E----------------------R----------------S--------------------------E------------C-------------------F-E-------AA-KE-KHLE--L--LED---------V--D----S--R--A-
ARA---VILYFKT-----WK----------P-------EKA--------M----------------E------H------A-------------------------A-----L---SE-------VL--------------------E---E-I-------T--A--------------------------------------------------GGN------------------L----------------IF------F-V---------------------------------------------------Y-------E----------D----EP------A--------I-KA-AW--------E---------Q------------------R-------------Q----Q----
-----------------------------------------------------P-----------GE-----------------G-------------------T-E--------------------------G--------------I--------------CL----------VTG--K---Q--A-E---IARI-H--G--T---I--K----G--------V--P--G--A--QS--SG-A--------------ALVS-----F-----------------N-------------A-------------S--A-----F-ES--Y---------G------K-------------------E--------Q-------SY-----------------NAP-------
VSTYAAYAYTTALNY-LL-----------A------D-------------R------D------H----------MT-------------------------------M---------------------I--------D-----T------M-------IVY---W----------S---------------E--------------R------------G-----------------------------------------------------E-------------Y----------N-----------------F-----------------------FM-------M-----E----------PT-----------A-------D-----------
----------------------------------N-----------------Q-------------E----IVAGV------FKN-----L-------------------------A-A-G------K-------AV---D---V---SD------V----EE-------SL-S-------L-----N-----Q-------R------F--YILGL---SPN--A-ARLSVRFFYQD-SFGNILRHVKDHY--DR------LR---IV----------------R-----------------------PA--------GD-----------R------ME-----Y-----------------L-------------G-----------VWRLL-----------
-----SE------------T---V-------------N------------K--------------------K-----------S------------R--------------D---------K--------------K-------------------P---AP----N-----------MAGSLYRAVIS--G-S--N---YPESMYQA-VI--------------------G----------------R-I--------------------RS--------------------------E--------------QDDS------DSHSYKIT----------------R-------G-----------------RAAM-----IKA-----F----------------L----------L---
----K------------N-----------R------G-G-S-------------------E---------------------------E--------------------I-------------------L------------------T-----------M-----AVN---------E--N---------SN-----D------------VAYIL-----GQEFAVLE-----AIQ--------------E---------------D------------------A---------------N----------KE---------I-----------N-------ATIK-----DKYFNSAC-------ATPASI--------------FPILFK--LK-N-SHLR--------K---I-------------NN------
--K----------------G-----K--EI-------Y------------------------Y-------EKM-L---G--------ALQ---------S------KI--------T-----EI-----------------------P----------K-------R------R---LN-LD-A----------QGRFILGYY----H----Q-----------------------------------------------------T-------------------RK----------R-----------------E-------------K----------------------K--------------SK----------E----------EV--------
-----------------------------------------------------------------------

'WP_117621567.1__2_' -------------------------------------------------------------MI------LQSLVTYYE----------S------L-----------E-----R-----------K------------------GK--------------ITSP---GW---CSA-KI-S-----FALELSE---A---GE-LL----R-------I---I----------PL-------K---ESVL----------R-G---------KK-----------T-AL---V----P--TI----R------K-V-P------Q--M--V--AR-----------S------S--G----------
V---------------S---A---N------------F-L---------------------CDNSSYL--------L----G----------------------------I-------------------DN---------------KG------------------------------------------------------K-P-------------------------E-------------------------R---------------S---------------------------V----------E-----------C------------------F-E------------AA-KE-KHLE--I---LKE----------T----G--G--K--A-
ARA---VVLYFKT-----WK----------P------EKA--------M----------------E------H------T-------------------------A----L---SE-------GL----------------------E----E-I---------T---A--------------------------------------GGN------------------L----------------IF------F-I-----------------------G--D-----------E---------E------F--------------AQ-------D----DP-----A---------V-KA-AW----------E--------TY---------S---------------------Q-------------K--------
-----------------------------------------------P---------GD------------G------------------V-E--------------------------------G--------------T--------------CL---------VTG--K---R--A-E---IARI-H--G--T---I--K----G--------V--P--G--A--QS--SG-A--------------ALVS-----F------------N-----------A-------------P-------A------F-ES---Y---------G-------K----------------E--------Q------SY----------------NAP--------
VSTYAAYAYTTALNY-LL-----------A------D-------------R-----D-------H----------MT-------------------------------M------------------I------G----------D-----T------M------IVY---W-----------S----------------------E---------------R----------G-----------------------------------------------------------V-------------YN-----------R-----------T---------F--------------------S---------------FM-------M-----E--------PT----------A-------D-------------
------------------------------------N-----------------Q----------E-----IVDGV------FKN------L-------------------------A-A-G------K-------AV---D---E---NG------T-----QE-------SL-S-------L-----D-----Q-------K-------F--YILGL---SPN--A-ARLSVRFFYQD-SFGNILRHVKEHY--DR------LR---IV------------------R------------------PS--------GD------------H----ME-----Y----------------------L-----------G-----------VWRLL--------
-----SE------------T---V-------------N-------------K--------------------K-------------S-----------------R----------------D-----------K------------------K------------------P--AP---N------------MAGSLYRAIIS--G-S--N---YPESMHQA-VL-------------------G------------------R-I-----------------RS------------------------------E---------------QDDS----DSRIYKIT------------R-----------G-------------RAAI-----IKA---------F----------------L--------------L-----
-----K--------------N-----------R------G--C-S-------------------E--------------------------E---------------------I----------------L--------------T----------------M------EMN----------E--N-----------SN-----D--------------VAYIL------GQEFAVLE------AIQ----------E--------------D--------------------A-------------N----------PG----------I------------N-------ATIK----DRYFNSAC-------ATPSSI------------FPILFK--LK-N-SHLR--------K--M-------NN------------
K----------------------G-----R--EI-------Y-------------------------------EKM-L---G--------ALQ----------S------KI---------T-----EV------------------------P------------K-------R------R---LN-LD-A-----------QDRFILGYY----H----Q-------------------------------------------------------T-------------------------RK------------R-----------------E---------------K-------------------K-----------------SK---------------E-----------EV---------
-------------------------------------------------------------------

'WP_117630905.1__6_' --------------------------------------------------------------MI------LQSLVTYYE-----------S------L------------E-----R----------K------------------GK--------------ITSP---GW---CSA-KI-S------FALELSE---A---GE-LL----R-------I---I-----------PL-------K----ESVL---------R-G----------KK---------T-AL---V----P--TI----R------K-V-P------Q--M--V--AR-----------S------S--G-----------
-V------------S---A---N------------F-L---------------------CDNSSYL--------L----G----------------------------I-----------------------DN-----------------KG----------------------------------------------------K-P-------------------------E-----------------------R-------------S---------------------------------V---------E--------C-----------------F-E-------------TA-KE-KHLE--I--LKE--------T----G--G--K--A-
ARA---VVLYFKT-----WK----------P------EKA--------M----------------E------H------T-------------------------A-----L---SE--------GL---------------------E----E-I----------T---A-------------------------------------GGN-----------------L-----------------IF------F-I----------------------G--D-----------E----------E------F-----------------AQ-------E----DP-----A----------V-KA-AW-----------E---------TY----------S------------------------Q-----------K------
-----------------------------------------------P----------GD-----------G-----------------V-E--------------------------------G--------------T--------------CL---------VTG--K---R--A-E---IARI-H--G--T---I--K----G---------V--P--G--A--QS--SG-A--------------ALVS-----F------------N-----------A--------------P-------A------F-ES---Y---------G-------K-----------------E--------Q------SY----------------NAP---------
VSTYAAYAYTTALNY-LL-----------A------D-------------R------D-------H-----------MT------------------------------M----------------I------G-----------D-----T------A------IVY---W-----------S---------------E--------------R----------G---------------------------------------------------V--------------YN------------R--------------T--------F---------------S---------------FM-------M-----E--------PT---------A--------D---------------
------------------------------------N-----------------Q----------E------IVDGV------FKN-----L------------------------A-A-G-------K-------KV---D---E---NG------T----QE-------SL-S-------L-----D------Q------K-------F--YILGL---SPN--A-ARLSVRFFYQD-SFGNILRHVKEHY--DR------LR---IV-------------------R-----------------PS---------GD-------------H----ME-----Y------------------L----------G-----------VWRLL---------
-----SE------------T---V--------------N-------------K---------------------K------------S-----------------R---------------D----------K---------------K------------------P---AP---N------------MAGSLYRAIIS--G-S--N---YPESMHQA-VL---------------------G-----------------R-I----------------RS----------------------------E--------------QDDS----DSRIYKIT------------R------------G-------------RAAI-----IKA-------F---------------L------------L------
----K--------------N-----------R------G--C-S-----------------------E-------------------------E--------------------I---------------L--------------T----------------M-----EMN-----------E--N-----------SN-----D--------------VAYIL------GQEFAVLE------AIQ----------E------------D-----------------A---------------N----------PG---------I-----------N-------ATIK----DRYFNSAC-------ATPSSI------------FPILFK--LK-N-SHLR---------K--M--------NN------------
K----------------------G-----R--EI-------Y-----------------------------EKM-L---G--------ALQ----------S------KI---------T-----EV------------------------P------------K-------R------R---LN-LD-A-----------QDRFILGYY----H----Q-----------------------------------------------------------T---------------------------QK------------R-------------------E---------------K--------------------K----------------SK----------------E----------EA----------
-------------------------------------------------------------------

'WP_117502778.1__2_' ------------------------------------------------------------MI------LQSLVTYYE-----------S------L------------E-----R-----------K-----------------GK--------------ITSP---GW---CSA-KI-S-----FALELSE---A---GE-LL-----R------I---I-----------PL-------K----ESVL----------R-G----------KK-------T-AL---V----P--TI----R-------K-V-P------Q--M--V--AR-----------S------S--G------------
-V---------------S---A---N------------F-L---------------------CDNSSYL--------L----G---------------------------I-----------------------DN----------------KG-------------------------------------------------K-P--------------------------E-----------------------R-------------S-------------------------------V--------E---------C------------------F-E-------------AA-KE-KHLE--I--LKE---------T----G--G--K--A-
ARA---VVLYFKT-----WK----------P------EKA--------M----------------E------H------T------------------------A-----L---SE-------GL-----------------------E----E-I----------T---A---------------------------------------GGN--------------------L-----------------IF------F-I-----------------------G--D---------E----------E-------F-----------------AQ------D----DP-----A--------V-KA-AW------------E--------TY----------S------------------------Q---------------K------
-----------------------------------------------P----------GD-----------G-----------------V-E--------------------------------G----------------T--------------CL---------VTG--K---R--A-E---IARI-H--G--T---I--K----G---------V--P--G--A--QS--SG-A--------------ALVS-----F------------N-----------A--------------P-------A-------F-ES---Y---------G-------K-----------------E--------Q------SY----------------NAP---------
VSTYAAYAYTTALNY-LL-----------A------D-------------R------D--------H-----------MT------------------------------M-----------------I------G-----------D-----T------A------IVY---W-----------S----------------E---------------R----------G------------------------------------------------V-------------YN------------R---------------T-------F--------------S----------------FM-------M-----E--------PT---------A--------D------------
------------------------------------N-----------------Q------------E------IVDGV------FKN-----L-----------------------A-A-G-------K-------KV---D---E---NG------T-----QE------SL-S-------L-----D-----Q-------K------F--YILGL---SPN--A-ARLSVRFFYQD-SFGNILRHVKEHY--DR------LR---IV------------------R-------------------PS----------GD------------H----ME-----Y----------------L----------G----------VWRLL-----------
-----SE------------T---V--------------N-------------K-------------------K------------S----------------R----------------D-----------K--------------K-------------------P---AP----N------------MAGSLYRAIIS--G-S--N---YPESMHQA-VL----------------------G-----------------R-I----------------RS----------------------------E---------------QDDS----DSRIYKIT-------------R------------G-------------RAAI-----IKA------F----------------L--------------L------
----K--------------N-----------R------G--C-S------------------------E-------------------------E------------------I-----------------L---------------T--------------M-----EMN-----------E--N------------SN------D--------------VAYIL------GQEFAVLE------AIQ----------E------------D-----------------A-----------------N-----------PG---------I-----------N-------ATIK----DRYFNSAC--------ATPSSI------------FPILFK--LK-N-SHLR----------K--M---------NN----------
K----------------------G-----R--EI-------Y-----------------------------EKM-L---G--------ALQ-----------S------KI---------T-----EV-------------------------P-------------K-------R------R---LN-LD-A-----------QDRFILGYY----H----Q--------------------------------------------------------T-------------------------QK-------------R--------------------E---------------K--------------------K----------------SK----------------E----------EA---------
-------------------------------------------------------------------

'WP_118706500.1__2_' -------------------------------------------------------------MI------LQSLVQYYE----------T-----L------------E-----R-----------K-----------------GK-------------ISSP---GW---CTA-KV-S-----FALELSG--E---GK-LK----R-------V---I------------PL------K---VEKE---------R-G---------KK--------I-VL--E---P--QS---M------K-V-P----Q--M--V--TR-----------S------S--G---------
---V---------------A---S---N-----------F-L--------------------CDNSSYL--------L----G---------------------------I----------------------DN-----------------KG-------------------------------------------------K-P--------------------------E---------------------R-----------------S-------------------------------E---------------------------------------F-Q---------CA-KE-KHIK--I--LQG------T----G--S--R--T-
-AKA---IIQYFET----WN------------P-------ETA--------D-----------------E---S---------------------V-----L---SE------VL--------------------E---E-I-------T---A------------------------------------------GAN--------------------L--------------------VF-----Y-V---------------------D---N------------E-----------Y--------AQ----------L----DG-----S-------------I-IS-AW-----------E--------DY------------R--------------A------------D-------
---------------------------------------A---------SA----------G----------------P-E----------------------------------------T-------------------------CL--------VTG--K--K--T-G---ISRI-H--G--T---I--K----G----------V--V--G--A--QS--SG-A--------------ALVS-----F-----------N-----------A-------------P-------A------F-ES---Y--------G------K------------------E--------Q-------SY----------------NAP--------
VGEYAAYAYTTALNH-LL-----------A------D----------R------K------H------------VT---------------------------------T-----------------------I---------------G----------------D-----V------T------IVY---W-----------S----------------E----------------D-----------G-----------------------------------------------------------------CQ-----------N--------V---------F---------A-----------GV-------A-------E------PT---------L---------D----------
------------------------------------F--------------------E----IVDGV------FKN-----L---------------------------------E-A-G------K-------AI---H---I---QG-------V----EE-------SL-S-------L-----E-----Q-------R-------F--FILGL---APN--A-ARIAVRFFYQD-SFGNILRHLKEHY--DR------ME---IV--------------R-----------------PA-------KD-------S----IQ------Y--------------L-----------G-----------VWRML-----------
ME-------------T---V-------------N-------------K--------------------K------------S-----------------R--------------D----------K------------K---------------------P--VS----N------------MAGAVYRSIIS--G-D--R---YPSNLYMA-VL------------------G------------------R-I------------------RA------------------------------E------------QDDG------DNGIYKIT---------------R-------------G----------RAAI-----IKA------F-----------------L---------L--------
R-------------N-----------G-----T--C-D-------------------------E--------------------------E---------------------I-------------------L--------------T-----------------V-----ALN---------E--N------------SK------N------------VAYTL-----GREFAVLE------AVQ------------------Q-------------E---------------A----------------N------PG---------I-----------N--------ATIK----DRYFNAAC-------ATPASI-----------FPILFK--LK-N-SHIR-----------K--M-------NN----------K-
--------------G-----K--EI-------Y-----------------------Y---------EKI-L---V----------EACEGV--------P--S------Y---------P-----------K--------R------R---LS-LE-E-------------QGIFNLGYY----F---Q-----------------------------------------------------------T-------------------QK--------------R----------------------E-------------------K------------------------K-------------KM-------------------E--------------ED-----------
-----------------------------------------------------------------------

'WP_118683102.1__2_' -------------------------------------------------------------MI------LQSLVQYYE----------A-----L-------------E-----R-------------R--------------GE--------------ITRP---GW---CNA-RV-S------FALELSP--D---GE-LK-----R-------S------V---I------------SL------K---EEKQ----------K-G---------RK--------T-VW--E---P--QN---L------K-V-P-----Q--M--V--TR-----------S------S--G----------
-----V--------------A---A---N-----------F-L---------------------CDNSSYF--------L----G----------------------------I--------------------DN----------------KG---------------------------------------------------K-P--------------------------E--------------------R--------------S----------------------------------------------------------------F-L-------------CA-KE-KHLD--I--LKE----------A---D--C--P-----
A--ALA---VIRFFER-----WD------------P------DQA---------A-----------------E-------N-------P-------------------------Y----V--SE---------IL------------------D--E-I------------T---A-------------------------------------GSN--------------------L--------------------VF------W-A--------------------------E----D------------E--------Y------AH-------T----DI-----S---------I-IR-AW------------E-------DH--------R-----------------------S------------N--------
---------------------------------------------A-------------------------A-T---------------------------------------G-------------------T-------------CL----------VTG--Q--K--T-E---IARI-H--G--N---I--K----G-------V--Q--G--A--QP--SG-A--------------ALVS-----F------------N-----------A-------------P------A-------F-ES--Y--------G------K-------------------E--------Q-------SY--------------------NAP-------
VGSYAVYAYTTALNH-LL-----------A------D----------R------K------H----------VT---------------------------------I----------------------I--------------G------------D-----T------T--------IVY---W-----------S------------------E---------D----------G-------------------------------------------------------------------------YQ-------------D--------------F---------------------L---------------NM-------S------E---------PT-----------A---------D----------
---------------------------------------C-------------------E----IVDGV------FKN-----L------------------------------Q-E-G-------K-------AV---H---V---SG------V----AE-------NI-S-------L-----S-----Q-------R-------F--YILGL---APN--A-ARIAVRFFYQD-SFGSILQNLKAHY--DR-------MD---IV-----------------R------------------PQ--------MD------------K----NR------Y-------------------L----------G-----------VWRML-----------
---ME------------T---V-------------N------------K--------------------K------------S-----------------R-----------------D----------K-------------K--------------------P---AP----N-------------MAGSLYRAIIS--G-D--R---YPSNLFQA-VM------------------G------------------R-I-----------------RS----------------------------------E---------------QDDS----SSGIYKIT-----------------R-----------G------------------RAAI-----IKA----F-----------------L--------L--------
R------------N--------------T---T-C-D---------------K--------------------E-------------------------E-----------I-----------------L-------------------T----------------M------ALN----------E--D-----------SN------N-----------TAYTL-----GRMFAVLE-------AIQ-----------E---------------D------------------A-------------N-----------PG----------I---------------N-------ATIK----DRYFNSAC-------ATPASI------------FPILLK--LK-N-SHTR-------------K--L----------NN-----------
R-------------G-----K--EI--------Y----------------------Y-----------EKL-L---V-----------GLQ----------D-----KI---------K-----APDEMS-------S--A-----Y------P-------R------R---LS-LE-E--------------QGMFILGYY----H----Q------------------------------------------------------------------------T---------------------------QK-----------R-------Y--------------E---------------K-------------------K--------------TK-------------E----------EQ----------
-------------------------------------------------------------------

'WP_009255682.1' --------------------------------------------------------MI------LQSLVQYYE---------A-----L-----------E-----Q-----------K-------------GE--------------ITRP---GW---CIA-RV-S------FALELSP--N---GD-LK-----R-------I-----I-----------SL-------K---NEVT----------K-G----------KK---------T-VR--E---P--QN---L------K-V-P-----Q--M--L--TR-----------S------S--G---------
-----V-------------T---A---N-----------F-L--------------------CDNSSYF--------L----G-------------------------I--------------------DN----------------KG-------------------------------------------------K-P--------------------------E---------------------R-------------S----------------------------------E-------------C-------------------------F-L--------------CA-KE-KHLE--I--LRE---------T---D--C--P--A-
AEA---VKRYFES-----WD-----------P--------DKA---------E-----------------E------N-----C----------------------L---V--SE-----------VL------------------D--E-I-------------T---A------------------------------------GAN-------------------L--------------------VF------W-V------------------------E--D------------E----------Y-------VH--------T---DT-------A------------V-VE-AW-----------E----------NY--------R---------------------A------------N-------
------------------------------------------EE-------------G--------------------I-T------------------------------------T----------------------CL--------VTG--E--K--T-E---IARI-H--G--N---I--K----G---------V--Q--G--A--QS--SG-A--------------ALVS-----F------------N-----------A-------------P------A-------F-ES--Y--------G------K-------------------E--------Q-------SY-------------------NAP--------
VGNYAVYAYTTALNH-LL-----------A------D----------R------R------H-------VS-----------------------------------M-----------------------I------G------------D-----T------T------IVY---W-----------S---------------E-----------------D----------G------------------------------------------------------------------------YQ--------------D--------------F-----------------L---------------SM-------S------E--------PT-------------M------D--------
----------------------------------------N-----------C------------E----IVDGV------FKN-----L-------------------------E-S-G------K-------AV---N---I---SG------V-----AE------NL-S-------L-----S-----Q-------K-------F--YILGL---APN--A-ARIAVRFFYQD-SFGNILKNLKAHY--DR------MD---II------------------R-------------------PQ--------TD------------G-----IK-----Y------------------L-----------G---------------VWRML--------
ME---------------T---V--------------N--------------K--------------------K--------------S-----------------R------------------D----------K---------------K-----------------------P---VP----N-------------MAGAVYRAIIS--G-D--R---YPSDLFRS-VM---------------------G---------------R-I----------------RS-------------------------------E---------------QDDS----SIGIYKIT--------------R---------------G-------------RAAI-----IKA----F----------------L-----------L----------
R-------------N---------T---T-C-D---------------K----------------------E---------------------------E---------I----------------------L---------------T---------------M-----ALN-----------E--D-----------SN------N------------TAYTL------GRMFAVLE------AIQ-------------E------------D--------------------A-------------N-----------PG---------I------------N--------ATIK-----DRYFNSAC-------ATPASI----------FPVLLK--LK-N-SHIR---------------K--L---------NN-----------
------------G-----K--EI---------Y-----------------------Y---------EKL-L--G--------GLQ----------D-----KI-------T-----AAEDMN------A--A-----Y-------P-------R------R---LT-LD-E---------------QGMFILGYY----H----Q-------------------------------------------------------------------------T-------------------------QK----------------R-------Y--------------E---------------K--------------------K----------------DK------------------E----------EQ-----------
-------------------------------------------------------------------

'WP_089981720.1' ------------------------------------------------------MI------LQSLVQYYE---------A-----L-----------E-----Q-----------K-------------GE--------------ITRP---GW---CIA-RV-S-----FALELSP--N---GD-LK-----R-------I-----I------------SL-------K---NEVS---------K-G----------KK---------T-VW--E---P--QK---L------K-V-P-----Q--M--L--TR----------S------S--G--------
---V---------------T---A---N-----------F-L-------------------CDNSSYF--------L----G-------------------------I---------------------DN------------------KG--------------------------------------------------K-P---------------------------E--------------------R--------------S---------------------------------E--------------C---------------------------F-L-------------CA-KE-KHLE--I--LRE-----------T---D--C--P--A-
AEA---VKRYFEN-----WD-----------P--------DKA---------E------------------E------N------C--------------------L---V--SE-----------VL--------------------D--E-I-----------T---A------------------------------------GAN-------------------L--------------------VF------W-V-----------------------E----D-----------E-----------Y--------VH--------T---DT-----A----------V-VE-AW-------------E---------NY--------R-----------------------A---------------N--------
------------------------------------------EE--------------G----------------------I-T---------------------------------------T-------------------CL--------VTG--K--K--T-E---IARI-H--G--N---I--K----G--------V--Q--G--A--QS--SG-A-------------ALVS-----F------------N----------A--------------P------A-------F-ES--Y--------G------K-------------------E--------Q-------SY-------------------NAP---------
VGNYAVYAYTTALNH-LL-----------A------D----------R------R------H------VS------------------------------------M------------------------I------G-------------D-----T------T--------IVY---W-----------S-----------------E-----------------D----------G-----------------------------------------------------------------------YQ------------D---------------F-----------------L----------------SM-------S-----E--------PT-------------M------D--------
---------------------------------------N------------C----------E----IVDGV------FKN-----L----------------------------E-S-G------K-------AV---N---I---SG------V-----AE------NL-S-------L-----S-----Q-------K-------F--YILGL---APN--A-ARIAVRFFYQD-SFGNILKNLKAHY--DR------MD---II---------------------R---------------------PQ-------TD-------------G-----IK-----Y------------------L-----------G----------------VWRML---------
ME---------------T---V--------------N--------------K-------------------K--------------S-----------------R------------------D----------K---------------K-----------------------P---VP----N-------------MAGAVYRAIIS--G-D--R---YPSDLFQS-VM---------------------G---------------R-I----------------RS-------------------------------E---------------QDDS----SIGIYKIT--------------R---------------G-------------RAAI-----IKA----F----------------L-----------L----------
R-------------K---K--KN-----------D----------------Y----------------ETL-L-G--------DLQ--------D-----KI-------T-----ASEDMN-------A--A-----Y------P-------R------R--LT-LD-E---------------QGMFILGYY----H----Q-----------------------------------------------------------------------T-------------------QK-------------R-------Y--------------E------------------K-----------------------K--------------DK-----------------E-----------EQ------------
-------------------------------------------------------------------

'WP_069432348.1__2_' --------------------------------------------------------MI------LQSLVQYYE---------A-----L-----------E-----Q-----------K-------------GE--------------ITRP---GW---CIA-RV-S-----FALELSP--N---GD-LK-----R-------I-----I-----------SL-------K---NEVP---------K-G----------KK---------T-VW--E---P--QK---L------K-V-P-----Q--M--L--TR-----------S------S--G----------
---V----------------T---A---N-----------F-L---------------------CDNSSYF--------L----G---------------------------I-----------------------DN------------------KG-------------------------------------------------------K-P-------------------------------E-----------------------R--------------S-------------------------------------E---------C------------------------F-Q---------------CA-KE-KHLE--I--LRE-----------T---D--C--P--A-
AEA---VKRYFES-----WD-----------P--------DNA----------E-----------------E------N-------C---------------------F---V--SE-----------VL--------------------D--E-I------------T---A------------------------------------GAN--------------------L----------------------VF------W-V-------------------------E---D-----------E-----------Y---------VH---------T---DT-----A----------V-AE-AW-------------E---------NY--------R----------------------A-------------N--------
------------------------------------------EE------------G----------------------I-T---------------------------------------T---------------------CL--------VTG--K--K--T-E---IARI-H--G--N---I--K----G---------V--Q--G--A--QS--SG-A-------------ALVS-----F------------N----------A--------------P-----A--------F-ES--Y-------G-------K-------------------E--------Q-------SY----------------------NAP--------
VGNYAVYAYTTALNH-LL-----------A------D----------R------R------H--------VS-----------------------------------M----------------------------I-------G------------D-----T-------T-------IVY---W-----------S----------------E------------------D----------G------------------------------------------------------------------------YQ-------------D--------------F----------------L-----------------SM-------S------E-------PT----------M---------D----------
------------------------------------------N-------------C------------E----IVDGV------FKN-----L-------------------------E-S-G------K-------AV---N---I---SG------V-----AE------NL-S-------L-----S------Q-------K------F--YILGL---APN--A-ARIAVRFFYQD-SFGNILKNLKAHY--DR------MD---II--------------------R---------------------PQ-------TD------------G-----IK-----Y------------------L------------G-----------------VWRML---------
ME---------------T---V---------------N--------------K-------------------K------------S-------------------R-------------------D----------K---------------K----------------------P---VP----N-------------MAGAVYRAIIS--G-D--R---YPSDLFQA-VM---------------------G---------------R-I----------------RS-------------------------------E---------------QDDS----SIGIYKIT------------R---------------G-------------RAAI-----IKA----F----------------L-----------L----------
R-------------N---------T---T-C-D---------------K---------------------E--------------------------E---------I----------------------L----------------T--------------M------ALN-----------E--D-----------SN------N------------TAYTL------GRMFAVLE-----AIQ--------------E-----------D--------------------A---------------N----------PG----------I------------N--------ATIK-----DRYFNSAC-------ATPASI----------FPVLLK--LK-N-SHIR-------------K--L---------NN-----------
---------G-----K--EI---------Y----------------------Y------------EKL-L--G---------GLQ----------D-----KI-------T-----ASEDMN-------A--A-----Y------P------R------R--LT-LD-E---------------QGMFILGYY----H----Q-------------------------------------------------------------------T-------------------QK-----------------R-------Y--------------E--------------K--------------------K--------------DK-----------------E-----------EQ-------------
-------------------------------------------------------------------

'WP_120094084.1__3_' -------------------------------------------------------MI------LQSLVQYYE---------A-----L-----------E-----Q-----------K-------------GE--------------ITRP---GW---CIA-RV-S-----FALELSP--N---GD-LK-----R-------I-----I------------SL-------K---NEVS---------K-G----------KK---------T-VW--E---P--QK---L------K-V-P-----Q--M--L--TR-----------S------S--G--------
---V---------------T---A---N-----------F-L--------------------CDNSSYF--------L----G-------------------------I-----------------------DN-----------------KG---------------------------------------------------K-P--------------------------E---------------------R------------------S--------------------------------E----------------C-------------------------F-L------------CA-KE-KHLE--I--LRE------------T---D--C--P--A-
AEA---VKRYFEN-----WD-----------P--------DKA---------E----------------E------N-------C--------------------L---V--SE----------VL---------------------D--E-I-----------T---A------------------------------------GAN--------------------L-----------------------VF------W-V--------------------------E---D-----------E-----------Y--------VH---------T---DT-----A----------V-VE-AW-----------E--------NY---------R------------------------A------------N--------
------------------------------------------EE------------G----------------------I-T------------------------------------T-----------------------CL--------VTG--K--K--T-E---IARI-H--G--N---I--K----G---------V--Q--G--A--QS--SG-A--------------ALVS-----F------------N----------A--------------P-----A--------F-ES---Y--------G-------K------------------E--------Q-------SY-------------------NAP----------
VGNYAVYAYTTALNH-LL-----------A-----D---------R------R------H-------VS----------------------------------------M-------------------------I-------G------------D-----T------T------IVY---W-----------S---------------E----------------D---------G--------------------------------------------------------------------YQ------------D---------------F-----------------L----------------SM-------S------E--------PT-----------M--------D--------
------------------------------------------N------------C------------E----LVDGV------FKN-----L-------------------------E-S-G------K-------AV---N---I---SG------V-----AE------NL-S-------L-----S------Q------K------F--YILGL---APN--A-ARIAVRFFYQD-SFGNILRNLKAHY--DR-------MD---II-------------------R---------------------PQ-------TD------------G-----IK-----Y------------------L-----------G----------------VWRML---------
ME---------------T---V-------------N---------------K-------------------K--------------S------------------R-----------------D----------K--------------K-----------------------P---VP----N-------------MAGAVYRAIIS--G-D--R---YPSDLFQS-VM---------------------G---------------R-I----------------RS------------------------------E---------------QDDS----SIGIYKIT-------------R---------------G------------RAAI-----IKA-----F----------------L------------L----------
R-------------N---------T---T-C-D---------------K--------------------E-------------------------E---------I---------------------L---------------T----------------M-----ALN-----------E--D-----------SN------N------------TAYTL------GRMFAVLE-----AIQ-------------E------------D--------------------A--------------N-----------PG----------I------------N--------ATIK-----DRYFNSAC-------ATPASI----------FPVLLK--LK-N-SHIR-------------K--L---------NN------------
---------G-----K--EI---------Y-----------------------Y------------EKL-L--G---------GLQ----------D-----KI-------T-----ASEDMN------A--A-----Y------P-------R------R--LT-LD-E--------------QGMFILGYY----H----Q----------------------------------------------------------------T-------------------QK-----------R-------Y--------------E--------------K--------------------K--------------DK-----------------E-----------EQ-------------
-------------------------------------------------------------------

'WP_044938556.1' -------------------------------------------------------MI------LQSLVNYYE--------A-----L-----------E-----R----------K-------------GV--------------VTSP---GW---CRA-KV-S-----SALEISA--E--GE-LL----R-------I-----I------------PL------K---VEEQ---------G-G---------KK---------T-VL--R---P--QT---L------K-V-P-----E--M--V--TR------------S------S--G---------
---I--------------S---A---N-----------F-L-------------------CDTSSYF--------L----G-------------------------I---------------------DN---------------KG-----------------------------------------------K-P------------------------E------------------R-----------------S------------------------E---------------C-----------------F-Q----------SA-RK-KHEE--I--LEG-------V----T--S--E--A-
ARA---VLHFFEK-----WN------------P-------HIA---------A---------------E-----------------------------V---L---KE----------SW------------------D--E-V-----------S---A------------------------------------GGN----------------L---------------------IF-----E-V--------------------------E-------------E---------Y---------AH--------E---DE-----E------------V-TE-AW-----------E-------HY--------R------------------------K------------N--------
----------------------------------------Q-------ET-------G-------------V-E------------------------------------------G-----------------T--------------CL--------VTG--A--Q--T-E---IARI-H--G--T---I--K----G--------V--Q--G--A--QS--SG-A--------------ALVS-----F------------N----------A--------------Q-----A--------F-ES---Y--------G--------K--------------E--------Q-------SY------------------NAP----------
VGKYAAYAYTTALNY-LI-----------S------D----------R------R------N------H--------IS--------------------------------R----------------------I------G-------------D-----T-----T------VVY---W-----------A--------------E-------------------D-------G---------------------------------------------------------------------------YQ-----------N---------------F--------------------------AA-------V-----E----------PA--------T------D----------
-------------------------------------N--------------Q----------E----IVDGV------FKN-----L----------------------------E-Q-G------K-------SV---D---V---EG------V-----AE------SL-S-------L-----D------Q-------R------F--YILGL---APN--A-ARLSVRFFYQD-SFGNILRNIKKHY--DR-------ME---IV------------------R---------------------PA------AD----------A------VE-------Y----------------L------------G----------------IWRML--------
QE--------------T---V-------------N--------------K-------------------K------------S---------------R----------------D----------K-----------K----------------------P---VP--G------------MAGATYRAVIS--G-S--R---YPAALYQG-VM-----------------G--------------R-I------------------RA---------------------------------E----------QEDK-----DTHSYKIT--------------------R---------------G--------------RAAI-----IKA------Y-----------------L-----------L----------
--R-----------N--------G---H-C-E---------------K-----------------------E-------------------------E-----------I--------------------L-------------------T-----------------V-----SLN-----------T--D-------------SK-----N-------------VAYTL-----GREFAVLE-----AIQ--------------E---------------D-----------------A---------------N----------PR-------I---------------N-------ATIK-----DKYFNSAC-------ATPAAI-----------FPILFK--LK-N-SHIR-------------K--I-----------NN-------------
-----------A-----T--EV------------Y----------------------F-----------EKI-L--G------------ELQ---------D------KI-------E----VSDTQV-------I--S-----Y------P-------R------R--LS-LE-E---------------QGMFILGYY----H----Q-------------------------------------------------------------------T------------------------QK-------------R-------Y-----------E-------------K----------------------K----------------VK--------------------E-------------N-------------
-------------------------------------------------------------------

'WP_033141979.1' --------------------------------------------------------MI------LQSLVRYYE---------A-----L-----------E-----K----------K-------------EK--------------ITSP---GW---CTA-KI-S-----FALDISE--K--GE-LL----R-------V-----I------------PL------K---LEKQ---------K-G---------KK---------T-VM--E---P--QL---L-----R-V-P-----E--M--V--TR------------S------S--G---------
---V---------------S---S---N-----------F-L--------------------CDNSGYF--------L----G-------------------------I-------------------DN-----------------KG-------------------------------------------------K-P--------------------------E--------------------R-----------------S-------------------------------E------------------C--------------------------F-A-------------AA-KK-KHLD--I--LDG-------V----E--G--N--A-
AAA---VRNFFET----WN-----------P------GGA--------A------------------E------H------P-------------------------A----L---AD--------NL-----------------------E--E-M-----------I---A----------------------------------------GGN------------------L---------------------IF------C-V--------------------------D-G---------D-----------W---------VQ--------N---DY-----D-----------V-RE-AW------------E-------IY-------R--------------------------A-----------D-------
--------------------------------------A---------EA-------D------------------N-E-------------------------------------------G----------------------------CL---------VTG--Q--K--T-E---ISRI-H--G--L---I--K----G----------V--P--G--A--QS--SG-A--------------ALVS-----F------------N------------A--------------P----A---------F-ES--F---------G-------K-----------------E--------Q-------GY--------------------NAP---------
VGTYAAYAYTTALNH-LL-----------A------D-----------R------K------S------H-------TS---------------------------------------------I--------------G------------D-----T------T-------VVY---W-----------A--------------E-------------------D-----G---------------------------------------------------------------------------YQ-----------N-------------F-----------------------------GA-------V------E-----------PT-------------V------D----------
-------------------------------------N----------Q--------A---IVAGV-----FNS-----L----------------------------E-M-G------K-------PI---V---V---EG------V-----EG-------SL-S-------L-----D------Q-------R------F--YILGL---APN--A-ARIAVRFFYQD-SFGNILRLHKEHY--DR-------ME---IV------------------R---------------------PA-------AD---------E-------MR-----Y------------------L------------G-------------------VWRML--------
--QE-------------T---V---------------N-------------K--------------------K-----------S-------------------R-------------------D-----------K--------------K--------------------P---TA----N--------------MAGAVYRAIIS--G-G--Q---YPASLFQG-VM--------------------G-------------R-I-------------------RA---------------------------E-----------QDDK------DSHIHKIT----------------------K-------------G----------------RASI-----IKA------Y----------------L------------L----------
--R-------------N--------G---Y-D-K------------------------------------E-----------------------------E-----------------I------------------L---------------T------------V-----SLN-----------E--N-------------SK------N-------------VAYTL------GREFAVLE------AIQ--------------E--------------D--------------------A----------------N----------PG--------I-------------N--------ATIK----DKYFNSAC-------ATPAAI------------FPVLFK--LK-N-SHIR--------------K--I-------------SN---------------H-------
----------G-----K--VV-----------Y----------------------F-----------EKL-L--G--------ELQ-----------D----KM----P---VTEEQM-------V-P-----C------P------R------R---LS-LE-E------------QGMFILGYY----H----Q--------------------------------------------------------------------------N---------------------QK-------------R-----------Y----------E-------------K-----------------------K---------------DK-----------------E----------------EA-------
-----------Q--DVRD--N---------------------------------

'WP_103732078.1__3_' --------------------------------------------------------MI------LQSLVRYYE---------A-----L-----------E-----K----------K-------------EK--------------ITSP---GW---CTA-KV-S-----FALDISE--K--GE-LL----R-------V-----I------------PL------K---LEKQ---------K-G---------KK---------T-VM--E---P--QL---L-----R-V-P-----E--M--V--TR------------S------S--G---------
---V---------------S---S---N-----------F-L--------------------CDNSGYF--------L----G-------------------------I-------------------DN-----------------KG-------------------------------------------------K-P--------------------------E--------------------R-----------------S-------------------------------E------------------C--------------------------F-A-------------AA-KK-KHLD--I--LDG-------V----E--G--N--A-
AAA---VRNFFET----WN-----------P------GGA--------A------------------E------H------P-------------------------A----L---AD--------SL-----------------------E--E-M-----------I---A----------------------------------------GGN------------------L---------------------IF------C-V--------------------------D-G---------D-----------W---------VQ--------N---DY-----D-----------V-RE-AW------------E-------IY-------R--------------------------A-----------D-------
--------------------------------------A---------ET-------D------------------N-E-------------------------------------------G----------------------------CL---------VTG--Q--K--T-E---ISRI-H--G--L---I--K----G----------V--P--G--A--QS--SG-A--------------ALVS-----F------------N------------A--------------P----A---------F-ES--F---------G-------K-----------------E--------Q-------GY--------------------NAP---------
VGTYAAYAYTTALNH-LL-----------A------D-----------R------K------S------H-------TS---------------------------------------------I--------------G------------D-----T------T-------VVY---W-----------A--------------E-------------------D-----G---------------------------------------------------------------------------YQ-----------N-------------F-----------------------------GA-------V------E-----------PT-------------V------D----------
-------------------------------------N----------Q--------A---IVAGV-----FNS-----L----------------------------E-M-G------K-------PI---V---V---EG------V-----EG-------SL-S-------L-----D------Q-------R------F--YILGL---APN--A-ARIAVRFFYQD-SFGNILRLHKEHY--DR-------ME---IV------------------R---------------------PA-------AD---------E-------MR-----Y------------------L------------G-------------------VWRML--------
--QE-------------T---V---------------N-------------K--------------------K-----------S-------------------R-------------------D-----------K--------------K--------------------P---TA----N--------------MAGAVYRAIIS--G-G--Q---YPASLFQG-VM--------------------G-------------R-I-------------------RA---------------------------E-----------QDDK------DSHIHKIT----------------------K-------------G----------------RASI-----IKA------Y----------------L------------L----------
--R-------------N--------G---Y-D-K------------------------------------E-----------------------------E-----------------I------------------L---------------T------------V-----SLN-----------E--N-------------SK------N-------------VAYTL------GREFAVLE------AIQ--------------E--------------D--------------------A----------------N----------PG--------I-------------N--------ATIK----DKYFNSAC-------ATPAAI------------FPVLFK--LK-N-SHIR--------------K--I-------------SN---------------H-------
----------G-----K--VV-----------Y----------------------F-----------EKL-L--G--------ELQ-----------D----KM----P---VTEEQM-------V-P-----C------P------R------R---LS-LE-E------------QGMFILGYY----H----Q--------------------------------------------------------------------------N---------------------QK-------------R-----------Y----------E-------------K-----------------------K---------------DK-----------------E----------------EA-------
-----------Q--DVRD--N---------------------------------

'WP_118743661.1__2_' --------------------------------------------------------MI------LQSLVRYYE---------A-----L-----------E-----K----------K-------------EK--------------ITSP---GW---CTA-KV-S-----FALDISE--K--GE-LL----R-------V-----I------------PL------K---LEKQ---------K-G---------KK---------T-VM--E---P--QL---L-----R-V-P-----E--M--V--TR------------S------S--G---------
---V---------------S---S---N-----------F-L--------------------CDNSGYF--------L----G-------------------------I-------------------DN-----------------KG-------------------------------------------------K-P--------------------------E--------------------R-----------------S-------------------------------E------------------C--------------------------F-A-------------AA-KK-KHLD--I--LDG-------V----E--G--N--A-
AAA---VRNFFES----WN-----------P------GGA--------A------------------E------H------P-------------------------A----L---AD--------SL-----------------------E--E-M-----------I---A----------------------------------------GGN------------------L---------------------IF------C-V--------------------------D-G---------D-----------W---------VQ--------N---DY-----D-----------V-KE-AW------------E-------IY-------R--------------------------A-----------D-------
--------------------------------------A---------ET-------D------------------N-E-------------------------------------------G----------------------------CL---------VTG--Q--K--T-E---ISRI-H--G--L---I--K----G----------V--P--G--A--QS--SG-A--------------ALVS-----F------------N------------A--------------P----A---------F-ES--F---------G-------K-----------------E--------Q-------GY--------------------NAP---------
VGTYAAYAYTTALNH-LL-----------A------D-----------R------K------S------H-------TS---------------------------------------------I--------------G------------D-----T------T-------VVY---W-----------A--------------E-------------------D-----G---------------------------------------------------------------------------YQ-----------N-------------F-----------------------------GA-------V------E-----------PT-------------V------D----------
-------------------------------------N----------Q--------A---IVAGV-----FNS-----L----------------------------E-M-G------K-------PI---V---V---EG------V-----EG-------SL-S-------L-----D------Q-------R------F--YILGL---APN--A-ARIAVRFFYQD-SFGNILRLHKEHY--DR-------ME---IV------------------R---------------------PA-------AD---------E-------MR-----Y------------------L------------G-------------------VWRML--------
--QE-------------T---V---------------N-------------K--------------------K-----------S-------------------R-------------------D-----------K--------------K--------------------P---TA----N--------------MAGAVYRAIIS--G-G--Q---YPASLFQG-VM--------------------G-------------R-I-------------------RA---------------------------E-----------QDDK------DSHIHKIT----------------------K-------------G----------------RASI-----IKA------Y----------------L------------L----------
--R-------------N--------G---Y-D-K------------------------------------E-----------------------------E-----------------I------------------L---------------T------------V-----SLN-----------E--N-------------SK------N-------------VAYTL------GREFAVLE------AIQ--------------E--------------D--------------------A----------------N----------PG--------I-------------N--------ATIK----DKYFNSAC-------ATPAAI------------FPVLFK--LK-N-SHIR--------------K--I-------------SN---------------H-------
----------G-----K--VV-----------Y----------------------F-----------EKL-L--G--------ELQ-----------D----KM----P---VTEEQM-------V-P-----C------P------R------R---LS-LE-E------------QGMFILGYY----H----Q--------------------------------------------------------------------------N---------------------QK-------------R-----------Y----------E-------------K-----------------------K---------------DK-----------------E----------------EA-------
-----------Q--DVRD--N---------------------------------

'WP_029470697.1' --------------------------------------------------------MI------LQSLVEYYE---------A-----L-----------E-----K----------K-------------GE--------------ITRP---GW---CKA-KI-S-----FALDISE--T--GV-LL----R-------V-----L------------PL------K---IEKQ---------R-G---------KK---------T-VM--E---P--Q---Q------E-V-P-----E--M--V--TR------------S------S--G---------
---V---------------S---A---N-----------F-L--------------------CDNSGYF--------L----G-------------------------I-------------------DN-----------------KG-------------------------------------------------K-P--------------------------E--------------------R-----------------S-------------------------------E------------------C--------------------------F-V-------------AA-KK-KHLE--I--LEN-------A----K--G--L--A-
ARA---VCSFFEA----WD-----------P------ACA--------S------------------E------H------P-------------------------A----V---KD--------SI-----------------------E--E-I-----------I---S----------------------------------------GGN------------------L---------------------IF------C-V--------------------------N-G---------D-----------W---------AQ--------D---EQ-----E-----------I-RA-AW------------E-------NS-------R--------------------------A-----------K-------
--------------------------------------A---------EK-------E------------------N-E-------------------------------------------G----------------------------CL---------VTG--Q--R--T-E---ISRI-H--G--L---I--K----G----------V--P--G--A--QS--SG-A--------------ALVS-----F------------N------------A--------------P----A---------F-ES--Y---------G-------K-----------------E--------Q-------SY--------------------NAP---------
VGTYAAYAYTTALNH-LL-----------A------D-----------R------K------S------H-------TS---------------------------------------------I--------------G------------D-----T------T-------VVY---W-----------A--------------E-------------------D-----G---------------------------------------------------------------------------YQ-----------N-------------F-----------------------------GT-------V------E-----------PT-------------V------D----------
-------------------------------------N----------Q--------D---IIDGV-----FKS-----L----------------------------E-A-G------K-------PI---A---V---DE------V-----EE-------SL-S-------L-----E------Q-------K------F--YILGL---APN--A-ARIAVRFFYQD-SFGNILRLHKEHY--DR-------ME---IV------------------R---------------------PA-------AD---------E-------MR-----Y------------------L------------G-------------------IWRML--------
QE-------------T---V---------------N-------------K--------------------K-----------S-------------------R-------------------D-----------K--------------K--------------------P---VS----N--------------MAGSVYRAVIS--G-G--R---YPASLFQG-VM--------------------G-------------R-I-------------------RA---------------------------E-----------QDDK------DSHIRKIT----------------------K-------------G----------------RAAI-----IKA------Y----------------L------------L----------
R-------------N--------G---Y-D-K------------------------------------E-----------------------------E-----------------I------------------L---------------T------------V-----SLN-----------E--N-------------SK------N-------------VAYTL------GREFAVLE------AIQ--------------E--------------D--------------------A----------------N----------PG--------I-------------N--------TTIK----DKYFNSAC-------ATPAAI------------FPVLFK--LK-N-SHIR--------------K--I-------------SN---------------H-------
----------G-----K--VV-----------Y----------------------F-----------EKL-L--G--------ELQ-----------N----RI----P---VTEGQM-------V-P-----C------P------R------R---LS-LE-E------------QGMFILGYY----H----Q--------------------------------------------------------------------------N---------------------QK-------------R-----------Y----------E-------------K-----------------------K---------------VK-----------------E----------------EE-------
-----------Q--DVRG--N---------------------------------

'WP_018596858.1' --------------------------------------------------------MI------LQSLVEYYE---------A-----L-----------E-----K----------K-------------GE--------------ITRP---GW---CKA-KI-S-----FALDISE--T--GV-LL----R-------V-----L------------PL------K---IEKQ---------R-G---------KK---------T-VM--E---T--QQ---M------E-V-P-----E--M--V--TR------------S------S--G---------
---V---------------S---A---N-----------F-L--------------------CDNSGYF--------L----G-------------------------I-------------------DN-----------------KG-------------------------------------------------K-P--------------------------E--------------------R-----------------S-------------------------------E------------------C--------------------------F-A-------------AA-KN-KHLE--I--LED-------T----K--G--T--A-
ARA---VCGFFES----WD-----------P------ACA--------S------------------E------H------P-------------------------A----V---KD--------SI-----------------------E--E-I-----------I---S----------------------------------------GGN------------------L---------------------IF------C-V--------------------------N-G---------D-----------W---------AQ--------D---EQ-----E-----------I-RA-AW------------E-------NS-------R--------------------------A-----------K-------
--------------------------------------A---------EK-------E------------------N-E-------------------------------------------G----------------------------CI---------VTG--Q--R--T-E---ISRI-H--G--L---I--K----G----------V--P--G--A--QS--SG-A--------------ALVS-----F------------N------------A--------------P----A---------F-ES--Y---------G-------K-----------------E--------Q-------SY--------------------NAP---------
VGTYAAYAYTTALNH-LL-----------A------D-----------R------K------S------H-------TS---------------------------------------------I--------------G------------D-----T------T-------VVY---W-----------A--------------E-------------------D-----G---------------------------------------------------------------------------YQ-----------N-------------F-----------------------------GT-------V------E-----------PS-------------V------D----------
-------------------------------------N----------Q--------D---IIDGV-----FKS-----L----------------------------E-A-G------K-------PI---A---V---GE------V-----EE-------SL-S-------L-----E------Q-------K------F--YILGL---APN--A-ARIAVRFFYQD-SFGNILRLHKEHY--DR-------ME---IV------------------R---------------------PA-------AD---------E-------MR-----Y------------------L------------G-------------------IWRML--------
QE-------------T---V---------------N-------------K--------------------K-----------S-------------------R-------------------D-----------K--------------K--------------------P---VS----N--------------MAGSVYRAVIS--G-G--R---YPASLFLG-VM--------------------G-------------R-I-------------------RA---------------------------E-----------QDDK------DSHIRKIT----------------------K-------------G----------------RAAI-----IKA------Y----------------L------------L----------
R-------------N--------G---Y-D-K------------------------------------E-----------------------------E-----------------I------------------L---------------T------------V-----SLN-----------E--N-------------SR------N-------------VAYTL------GREFAVLE------AIQ--------------E--------------D--------------------A----------------N----------PG--------I-------------N--------TTIK----DKYFNSAC-------ATPAAI------------FPVLFK--LK-N-SHIR--------------K--I-------------SN---------------H-------
----------G-----K--VV-----------Y----------------------F-----------EKL-L--G--------ELQ-----------N----RI----P---VTEGQM-------V-P-----C------P------R------R---LS-LE-E------------QGMFILGYY----H----Q--------------------------------------------------------------------------N---------------------QK-------------R-----------Y----------E-------------K-----------------------K---------------VK-----------------E----------------EE-------
-----------Q--DVRG--N---------------------------------

'WP_065543455.1__2_' --------------------------------------------------------MI------LQSLVEYYE---------A-----L-----------E-----K----------K-------------GK--------------ITRP---GW---CKA-KI-S-----FALDISE--T--GD-LL----G-------V-----I------------PL------K---IEKQ---------R-G---------KK---------T-VM--E---P--RQ---M------E-V-P-----E--M--V--TR------------S------S--G---------
---V---------------S---A---N-----------F-L--------------------CDNSGYF--------L----G-------------------------I-------------------DN-----------------KG-------------------------------------------------K-P--------------------------E--------------------R-----------------S-------------------------------E------------------C--------------------------F-A-------------AA-KN-KHLE--I--LEG-------E----G--T--T-
ARA---VRSFFGS----WN-----------P------DSA--------A------------------E------H------A-------------------------A----I---KD--------SI-----------------------E--E-I-----------I---S----------------------------------------GGN------------------L---------------------IF------C-V--------------------------N-G---------D-----------W---------AQ--------D---EH-----E-----------I-RE-AW------------E-------NC-------R--------------------------G-----------N-------
--------------------------------------A---------EW-------E------------------N-E-------------------------------------------G----------------------------CL---------VTG--Q--K--T-E---ISRI-H--G--L---I--K----G----------V--P--G--A--QS--SG-A--------------ALVS-----F------------N------------A--------------P----A---------F-ES--Y---------G-------K-----------------E--------Q-------SY--------------------NAP---------
VGTYAAYAYTTALNH-LL-----------A------D-----------R------K------S------H-------TS---------------------------------------------I--------------G------------D-----T------T-------VVY---W-----------A--------------E-------------------D-----G---------------------------------------------------------------------------YQ-----------N-------------F-----------------------------GA-------V------E-----------PS-------------V------D----------
-------------------------------------N----------Q--------D---IIDGV-----FRS-----L----------------------------E-A-G------K-------PT---A---V---DE------V-----EE-------SL-S-------L-----E------Q-------K------F--YILGL---APN--A-ARIAVRFFYQD-SFGNVLRLHKEHY--DR-------MG---IV------------------R---------------------PA-------AD---------E-------MR-----Y------------------L------------G-------------------IWRML--------
--QE-------------T---V---------------N-------------K--------------------K-----------S-------------------R-------------------D-----------K--------------K--------------------P---VS----N--------------MAGSVYRAVIS--G-G--Q---YPASLFLG-VM--------------------G-------------R-I-------------------RA---------------------------E-----------QDDK------DSHIRKIT----------------------K-------------G----------------RAAI-----IKA------Y----------------L------------I----------
--R-------------N--------G---Y-D-K------------------------------------E-----------------------------E-----------------I------------------L---------------T------------V-----SLN-----------E--D-------------SR------N-------------VAYTL------GREFAVLE------AIQ--------------E--------------D--------------------A----------------N----------PG--------I-------------N--------TTIK----DKYFNSAC-------ATPAAI------------FPVLFK--LK-N-SHIR--------------K--I-------------SN---------------------
R---------G-----K--VV-----------Y----------------------F-----------EKL-L--G--------ELQ-----------D----RI----P---VTEGQM-------V-P-----C------P------R------R---LS-LE-E------------QGMFILGYY----H----Q--------------------------------------------------------------------------N---------------------QK-------------R-----------Y----------E-------------K-----------------------K---------------VK-----------------E----------------EE-------
-----------Q--DVRG--N---------------------------------

'WP_104803918.1' --------------------------------------------------------MI------LQSLVEYYE---------A-----L-----------E-----K----------K-------------GE--------------ITRP---GW---CKA-KI-S-----FALDISE--T--GD-LL----G-------V-----I------------PL------K---IEKQ---------K-G---------KK---------T-VM--E---P--RQ---M------E-V-P-----E--M--V--TR------------S------S--G---------
---V---------------S---A---N-----------F-L--------------------CDNSGYF--------L----G-------------------------I-------------------DN-----------------KG-------------------------------------------------K-P--------------------------E--------------------R-----------------S-------------------------------E------------------C--------------------------F-A-------------AA-KN-KHLE--I--LEG-------M----E--G--T--T-
ARA---VCSFFES----WN-----------P------DSA--------A------------------E------H------A-------------------------A----V---KE--------SL-----------------------E--E-I-----------I---S----------------------------------------GGN------------------L---------------------IF------C-V--------------------------N-G---------D-----------W---------VQ--------D---EH-----E-----------I-RE-AW------------E-------NC-------R--------------------------G-----------N-------
--------------------------------------V---------EW-------E------------------N-E-------------------------------------------G----------------------------CL---------VTG--Q--K--T-E---ISRI-H--G--L---I--K----G----------V--P--G--A--QS--SG-A--------------ALVS-----F------------N------------A--------------P----A---------F-ES--Y---------G-------K-----------------E--------Q-------SY--------------------NAP---------

[illegible]

'WP_107434545.1__2_'

[illegible]

'WP_118756608.1__2_'

[illegible]

'WP_007708673.1__2_'

[illegible]

'WP_024737271.1'

[illegible]

'WP_002589368.1__2_'

[illegible]

'WP_002577347.1__2_'

[illegible]

'WP_118017194.1__4_'

[illegible]

'WP_119205515.1__2_'

[illegible]

'WP_002568438.1__3_'

[illegible]

'WP_117625707.1__2_'

[illegible]

'WP_002570353.1'

[illegible]

'WP_027644257.1'

[illegible]

'WP_065548865.1'

[illegible]

'WP_002585644.1'

[illegible]

'CUP29249.1'

[illegible]

'WP_002595865.1'

[illegible]

'WP_022512574.1'

[illegible]

'WP_089776479.1'

[illegible]

'WP_022202383.1'

[illegible]

'WP_118600813.1__3_'

[illegible]

[illegible]

'WP_118679154.1__3_'

[illegible]

'WP_118752145.1__2_'

[illegible]

'WP_008398233.1'

[illegible]

'WP_118571933.1__2_'

[illegible]

'WP_118610770.1__2_'

[illegible]

'WP_119253980.1__2_'

[illegible]

'WP_118586011.1__3_'

[illegible]

'WP_118629569.1__2_'

[illegible]

'WP_118738801.1__2_'

[illegible]

'WP_117944370.1__2_'

[illegible]

'WP_004853850.1'

[illegible]

'WP_118017416.1__3_'

[illegible]

'WP_117549122.1__2_'

[illegible]

'WP_117831401.1__2_'

[illegible]

'WP_117718533.1__5_'

[illegible]

'RGW35038.1'

[illegible]

'WP_015517329.1__9_'

[illegible]

'WP_012743317.1'

[illegible]

'WP_118535307.1__2_'

[illegible]

'WP_117852404.1__2_'

[illegible]

[illegible]

'WP_117603849.1__2_' [illegible]

'WP_117889180.1__2_' [illegible]

'WP_118384375.1__2_' [illegible]

'WP_118212687.1__2_' [illegible]

'WP_117687678.1__2_' [illegible]

'WP_005610783.1__2_' [illegible]

'WP_016441068.1' [illegible]

'WP_116441475.1' [illegible]

'WP_119988372.1__2_' [illegible]

'WP_016439664.1' [illegible]

'WP_004612294.1__8_' [illegible]

'WP_117994974.1__4_' [illegible]

'WP_101880310.1__3_' [illegible]

'WP_101882989.1__2_' [illegible]

'WP_117501528.1__3_' [illegible]

'WP_118349045.1__2_' [illegible]

'WP_117787131.1__2_' [illegible]

'WP_101695987.1' [illegible]

'WP_117822901.1__2_' [illegible]

'WP_118550047.1__2_' [illegible]

'WP_118538667.1__2_'

[illegible]

'WP_118669565.1__2_'

[illegible]

'WP_118731108.1__2_'

[illegible]

'WP_132277627.1__2_'

[illegible]

'WP_025642756.1'

[illegible]

'WP_117451872.1__2_'

[illegible]

'WP_083425209.1'

[illegible]

'WP_045091752.1'

[illegible]

'WP_007863415.1'

[illegible]

'WP_048929553.1'

[illegible]

'WP_004068710.1'

[illegible]

'WP_006860504.1'

[illegible]

'WP_118570334.1__2_'

[illegible]

'WP_118599779.1__3_'

[illegible]

'WP_118297976.1__2_'

[illegible]

'WP_117567095.1__3_'

[illegible]

'WP_118496165.1__2_'

[illegible]

'WP_118762720.1__2_'

[illegible]

'WP_117960134.1__2_'

[illegible]

'WP_055056879.1__3_'

[illegible]

'WP_118513667.1__3_'

[illegible]

[illegible]

'WP_059085217.1__4_'

[illegible]

'WP_118752445.1__2_'

[illegible]

'WP_118721368.1__2_'

[illegible]

'WP_118706048.1__2_'

[illegible]

'WP_118540410.1__2_'

[illegible]

'WP_047776666.1'

[illegible]

'WP_009002351.1'

[illegible]

'WP_025655204.1__2_'

[illegible]

'WP_055154389.1__2_'

[illegible]

'WP_050641462.1'

[illegible]

'WP_118620563.1__2_'

[illegible]

'WP_109733669.1'

[illegible]

'WP_087153300.1'

[illegible]

'WP_089822784.1'

[illegible]

'WP_087203017.1'

[illegible]

'WP_087168563.1'

[illegible]

'WP_087417156.1'

[illegible]

'WP_087271881.1'

[illegible]

'WP_087175543.1'

[illegible]

'WP_087177636.1'

[illegible]

VGRYAEFAYTTALNY-LL [illegible]

[illegible]

'PWL47312.1' [illegible]

[illegible]

'WP_081483129.1' [illegible]

[illegible]

'EIM57402.1' [illegible]

[illegible]

'WP_029201956.1' [illegible]

[illegible]

'WP_031550423.1' [illegible]

[illegible]

'WP_045085855.1' [illegible]

[illegible]

'WP_120483804.1__2_' [illegible]

[illegible]

'WP_016314946.1' [illegible]

[illegible]

'WP_050698727.1' [illegible]

[illegible]

'WP_135546087.1__2_' [illegible]

[illegible]

'WP_110940145.1' [illegible]

[illegible]

'WP_049756823.1' [illegible]

[illegible]

'WP_080063951.1' [illegible]

[illegible]

'WP_083908970.1' [illegible]

[illegible]

'EMS72167.1' [illegible]

[illegible]

'WP_085532120.1' [illegible]

[illegible]

'WP_105206035.1' [illegible]

[illegible]

'WP_114174658.1' [illegible]

[illegible]

'WP_082671629.1' [illegible]

[illegible]

'WP_075704387.1' [illegible]

[illegible]

[illegible]

'WP_086035854.1'

[illegible]

'WP_118478535.1__2_'

[illegible]

'WP_093373729.1'

[illegible]

'WP_014080658.1'

[illegible]

'WP_016229738.1'

[illegible]

'WP_087020869.1'

[illegible]

'PWM23190.1'

[illegible]

'WP_120395771.1__2_'

[illegible]

'WP_087392525.1'

[illegible]

'WP_016148215.1'

[illegible]

'WP_101693221.1'

[illegible]

'WP_044940447.1'

[illegible]

'WP_054338281.1'

[illegible]

'WP_006570571.1'

[illegible]

'WP_087287308.1'

[illegible]

'WP_021751294.1'

[illegible]

'WP_025543475.1'

[illegible]

'WP_119311097.1'

[illegible]

'WP_009257010.1'

[illegible]

'WP_087358649.1'

[illegible]

```
[illegible]
'WP_007489325.1' [illegible]
[illegible]
'WP_024724181.1' [illegible]
[illegible]
'WP_021631529.1' [illegible]
[illegible]
'WP_089413496.1' [illegible]
[illegible]
'CUQ04037.1' [illegible]
[illegible]
'WP_050617233.1' [illegible]
[illegible]
'WP_118480575.1__3_' [illegible]
[illegible]
'WP_110435577.1' [illegible]
[illegible]
'WP_119194871.1__2_' [illegible]
[illegible]
'WP_118730148.1__3_' [illegible]
[illegible]
'WP_118761654.1__2_' [illegible]
[illegible]
'WP_118361303.1__2_' [illegible]
[illegible]
'WP_119189827.1__2_' [illegible]
[illegible]
'WP_118564365.1__3_' [illegible]
[illegible]
'WP_117751870.1__2_' [illegible]
[illegible]
'PWL87004.1' [illegible]
[illegible]
'WP_118622540.1__2_' [illegible]
[illegible]
'PWM95101.1' [illegible]
[illegible]
'PWM76894.1' [illegible]
[illegible]
'WP_097793567.1__2_' [illegible]
[illegible]
```

'WP_118553829.1__2_'

[illegible]

'AXA81336.1__2_'

[illegible]

'RGB90652.1'

[illegible]

'RAW51766.1'

[illegible]

'WP_097801867.1__2_'

[illegible]

'WP_118657552.1__2_'

[illegible]

'WP_097780407.1__2_'

[illegible]

'WP_071427506.1__2_'

[illegible]

'WP_114000281.1__2_'

[illegible]

'WP_058965937.1'

[illegible]

'WP_087183597.1'

[illegible]

'WP_087180256.1'

[illegible]

'WP_031477054.1'

[illegible]

'PWM55851.1'

[illegible]

'WP_087324200.1'

[illegible]

'WP_087249184.1'

[illegible]

'WP_087365976.1'

[illegible]

'WP_087385543.1'

[illegible]

'WP_091351050.1'

[illegible]

'WP_124032145.1'

[illegible]

'ACF43896.1'

[illegible]

[illegible]

'WP_006966954.1__2_' [illegible]

'RLC13468.1' [illegible]

'RLB93060.1' [illegible]

'WP_014747986.1' [illegible]

'WP_062767987.1' [illegible]

'WP_106755092.1' [illegible]

'WP_082676055.1' [illegible]

'WP_056106059.1' [illegible]

'WP_101326945.1' [illegible]

'WP_015921089.1' [illegible]

'WP_085377863.1' [illegible]

'WP_028719258.1__2_' [illegible]

'WP_019960727.1' [illegible]

'PIP96388.1' [illegible]

'AQS41297.1' [illegible]

'WP_111352721.1' [illegible]

'WP_116684517.1__2_' [illegible]

'WP_078706396.1' [illegible]

'WP_056503906.1' [illegible]

'WP_092815989.1__2_' [illegible]

[illegible]

'WP_026381853.1'

[illegible]

'WP_112874288.1'

[illegible]

'WP_039004710.1'

[illegible]

'WP_089235483.1'

[illegible]

'WP_110806044.1'

[illegible]

'WP_013384679.1'

[illegible]

'WP_107674404.1__2_'

[illegible]

'WP_108643981.1'

[illegible]

'WP_130991708.1__2_'

[illegible]

'OYU17141.1'

[illegible]

'WP_105516362.1'

[illegible]

'WP_076363198.1'

[illegible]

'WP_097031955.1'

[illegible]

'WP_013067033.1'

[illegible]

'WP_023921140.1'

[illegible]

'WP_093039033.1'

[illegible]

'PKQ10625.1'

[illegible]

'WP_074971179.1'

[illegible]

'WP_074991370.1__2_'

[illegible]

'WP_085503168.1'

[illegible]

[illegible]

'WP_112062299.1'

[illegible]

'WP_035597547.1'

[illegible]

'WP_131308532.1__2_'

[illegible]

'WP_135049828.1__2_'

[illegible]

'WP_016922097.1'

[illegible]

'WP_123194635.1'

[illegible]

'WP_050472648.1'

[illegible]

'WP_058899464.1'

[illegible]

'WP_018289541.1'

[illegible]

'WP_076401182.1'

[illegible]

'WP_132010398.1'

[illegible]

'WP_038304726.1'

[illegible]

'WP_023786786.1'

[illegible]

'WP_020087240.1'

[illegible]

'WP_008387836.1'

[illegible]

'WP_055037664.1'

[illegible]

'WP_054161002.1'

[illegible]

'WP_092684710.1'

[illegible]

'WP_027276965.1'

[illegible]

'WP_110782087.1'

[illegible]

[illegible]

'TAH67549.1'

[illegible]

'WP_107355497.1'

[illegible]

'WP_047307681.1'

[illegible]

'WP_013502809.1'

[illegible]

'WP_012817718.1'

[illegible]

'WP_013418890.1'

[illegible]

'WP_019403924.1'

[illegible]

'WP_009796705.1'

[illegible]

'WP_075632907.1'

[illegible]

'WP_024710163.1'

[illegible]

'WP_065008529.1_2_'

[illegible]

'WP_122085453.1'

[illegible]

'WP_056234651.1'

[illegible]

'WP_011315192.1'

[illegible]

'WP_035026870.1'

[illegible]

'WP_130514478.1_2_'

[illegible]

'WP_023514069.1'

[illegible]

'WP_109459769.1_2_'

[illegible]

'WP_112691434.1'

[illegible]

'WP_109705819.1_2_'

[illegible]

'WP_027669084.1'
[illegible]

'WP_117193999.1'
[illegible]

'WP_029060025.1'
[illegible]

'THF50753.1'
[illegible]

'RMF12111.1'
[illegible]

'WP_011388585.1'
[illegible]

'WP_021133647.1'
[illegible]

'WP_002725660.1'
[illegible]

'WP_056407584.1'
[illegible]

'WP_036262256.1'
[illegible]

'WP_051949378.1'
[illegible]

'WP_086054694.1'
[illegible]

'WP_108783898.1'
[illegible]

'WP_134725989.1'
[illegible]

'WP_081904700.1'
[illegible]

'WP_083848267.1'
[illegible]

'WP_082429920.1'
[illegible]

'WP_084629238.1'
[illegible]

'WP_071469720.1'
[illegible]

'WP_066710854.1__2_'
[illegible]

'KPQ01758.1'
[illegible]

'WP_103966102.1'
[illegible]
'WP_080522118.1'
[illegible]
'WP_054107356.1'
[illegible]
'WP_066710485.1__3_'
[illegible]
'WP_083956843.1'
[illegible]
'WP_086640456.1'
[illegible]
'WP_035379699.1'
[illegible]
'WP_061487988.1'
[illegible]
'WP_045541641.1'
[illegible]
'WP_095351568.1'
[illegible]
'WP_006116165.1__3_'
[illegible]
'WP_025828082.1'
[illegible]
'WP_132296049.1__2_'
[illegible]
'WP_012553887.1'
[illegible]
'WP_012222309.1'
[illegible]
'WP_029356790.1'
[illegible]
'WP_026873905.1'
[illegible]
'WP_075822409.1__2_'
[illegible]
'WP_019462121.1'
[illegible]
'WP_093566088.1'
[illegible]

VSAEAGFRYGAALNR-ML---------D-----RN-----------S------R------N-----------RL------------------------RRP--------L-----G---------D---A----T--------VVF--W---------A-------------D-------------T---------S [illegible]
----P---------------------D---Q-D------------A----------A----E----------A-------------A----LIRDA-----LAT-----L------------------------A-A-G-----R-------PV--E--A---VA-----P---K------V-A------P----G----T-------R [illegible]
IGRLL--------------------------VK------------T---T-------------------A-------------L-----------Q-------E--------K----------------------F----------D--N----------------I----PP----L---------------LAGEVARAVLA--G-T--R---Y [illegible]
L-------------T-------------R-------------D----------L-------------R-------------L----------------N-GDR-----------G------------------E-----------------I------------------P------------V----SLD---------K--D----------DP----S [illegible]
R--L----------RK--D----------R-K---------D------Y------------------W---------------I----------------ERE-I--G--------EIV-----------E-----RL-------------P------P-----V-------L-----P-------R-----V---LR-LE-A---------QGRFAIGYY [illegible]
AL---------A----------AA-----------------V--EADE---TKDS-------------------------QNQDSPIEDSQD-DV-----------------------
'WP_082167656.1' ----------------------------------------------------------MTV------LQALDRYYD---------R----M-----------A----A-----------R-G------------E [illegible]
-------I-----------L------P---N-----------R-L--------------------WDKTAYV-------L----G--------------------------------R------------------T---------------------A [illegible]
G--LLA---LARFLES----WD------------P------AAF---------D------------------A------A------P----------------------------------------F-------------E--P--E-M-----------L [illegible]
------------------------AG---------------------AP-----------AA----------E-----------------------A-T------------------G---------P------------------V-----------------CL-----------VTG--R---P--A-A---PLRL-H--P--T---I--K----G [illegible]
VSAEAGFRYGAALNR-ML---------D-----RT-----------S------R------N-----------RL------------------------RRP--------L-----G---------D---A----T--------VVF--W---------A-------------D-------------T---------S [illegible]
---P---------------------D---Q-D------------A----------A----E----------A-------------A----LIRDA-----LAA-----L------------------------A-A-G-----R-------PI--A--A---VA-----P---A------V-S------P----G----T-------R [illegible]
---------------------VK------------T---T-------------------A-------------L-----------Q-------E--------K----------------------F----------E--N----------------I----PP----L---------------LAGEVARAVLA--G-T--R---YPRSWLA [illegible]
----M-------------R-------------D----------L-------------R-------------L----------------T-GDR-----------G------------------E-----------------I------------------P------------V----SLD---------K--D----------DP----S-------------PAYQL [illegible]
-----RK--D----------G-K---------D------Y------------------W---------------I----------------ERE-I--G--------EIV-----------E-----RL-------------P------S----A------L-----P-------R-----V---LR-LE-A---------QGRFAIGYY---H [illegible]
--------AS-----------------V--EADG---TEAG-------------------------PTDDTPIEDSQD-DV-----------------------
'WP_082742535.1' ----------------------------------------------------------MTI------LQALDRYYD---------R----M-----------A----A-----------R-G------------E [illegible]
----I-----------L------P---N-----------R-L--------------------WDKTAYV-------L----G--------------------------------R------------------T---------------------A [illegible]
-LLA---LVRFLDA----WD------------P------AAF---------D------------------A------A------P----------------------------------------F-------------E--P--E-M-----------L [illegible]
[illegible]
'WP_091888966.1' [illegible]
[illegible]
'WP_091752732.1' [illegible]
[illegible]
'WP_013300568.1' [illegible]
[illegible]
'WP_021228872.1' [illegible]
[illegible]
'WP_109922984.1__2_' [illegible]
[illegible]
'WP_086642592.1' [illegible]
[illegible]
'WP_066704078.1' [illegible]
[illegible]
'WP_010543172.1' [illegible]
[illegible]
'WP_014076066.1' [illegible]
[illegible]
'WP_027443944.1' [illegible]
[illegible]
'WP_022681041.1' [illegible]
[illegible]
'WP_018262008.1' [illegible]
[illegible]
'WP_012331627.1' [illegible]
[illegible]
'WP_037014812.1' [illegible]
[illegible]
'WP_034999601.1' [illegible]
[illegible]
'WP_113037777.1' [illegible]
[illegible]
'PCI23199.1' [illegible]
[illegible]
'WP_040727079.1' [illegible]
[illegible]

[illegible]

'WP_010603693.1' [illegible]

[illegible]

'WP_004397038.1' [illegible]

[illegible]

'WP_039425122.1' [illegible]

[illegible]

'WP_116963181.1' [illegible]

[illegible]

'PIQ01579.1' [illegible]

[illegible]

'WP_103855782.1__2_' [illegible]

[illegible]

'WP_103853157.1__2_' [illegible]

[illegible]

'WP_115249802.1' [illegible]

[illegible]

'WP_039081831.1' [illegible]

[illegible]

'WP_039091921.1' [illegible]

[illegible]

'WP_039157479.1' [illegible]

[illegible]

'WP_018346674.1' [illegible]

[illegible]

'WP_039163963.1' [illegible]

[illegible]

'WP_039086327.1' [illegible]

[illegible]

'WP_021462180.1' [illegible]

[illegible]

'WP_039168445.1' [illegible]

[illegible]

'RKV68694.1' [illegible]

[illegible]

'WP_040354990.1' [illegible]

[illegible]

'WP_079539609.1' [illegible]

[illegible]

'WP_048714866.1' [illegible]

[illegible]

[illegible]

'WP_090654484.1' [illegible]

'WP_100319891.1__2_' [illegible]

'WP_085364203.1' [illegible]

'WP_009117700.1' [illegible]

'WP_115134129.1' [illegible]

'WP_107855297.1' [illegible]

'WP_004284051.1' [illegible]

'WP_004285513.1' [illegible]

'WP_107860791.1' [illegible]

'WP_027074457.1' [illegible]

'WP_003781698.1' [illegible]

'WP_109427111.1' [illegible]

'WP_049335859.1' [illegible]

'WP_003762755.1__2_' [illegible]

'WP_107825054.1' [illegible]

'WP_049228929.1' [illegible]

'WP_077582705.1' [illegible]

'WP_077586450.1' [illegible]

'WP_131978509.1' [illegible]

'WP_018652891.1' [illegible]

'WP_115589918.1'
[illegible]

'WP_131928837.1'
[illegible]

'WP_006250670.1'
[illegible]

'WP_040080916.1__2_'
[illegible]

'WP_006252735.1'
[illegible]

'WP_031206576.1'
[illegible]

'WP_031192850.1__9_'
[illegible]

'WP_015484658.1__3_'
[illegible]

'AGK02095.1'
[illegible]

'WP_025247935.1'
[illegible]

'WP_025217638.1'
[illegible]

'WP_025289164.1'
[illegible]

'WP_034612302.1'
[illegible]

'WP_100052054.1'
[illegible]

'WP_117471492.1'
[illegible]

'WP_123805651.1__2_'
[illegible]

'WP_089036626.1__2_'
[illegible]

'WP_115611710.1'
[illegible]

'WP_006986820.1'
[illegible]

'WP_049254122.1'
[illegible]

'WP_100325810.1__2_'
[illegible]

[illegible]

'WP_107879708.1'

[illegible]

'WP_107929152.1'

[illegible]

'WP_085391319.1'

[illegible]

'WP_085418202.1'

[illegible]

'WP_054618559.1'

[illegible]

'WP_009117236.1'

[illegible]

'WP_047761588.1'

[illegible]

'WP_077465020.1'

[illegible]

'WP_123804953.1__3_'

[illegible]

'WP_096295766.1__2_'

[illegible]

'WP_077476501.1'

[illegible]

'WP_108923676.1__2_'

[illegible]

'WP_003797746.1'

[illegible]

'WP_109079179.1'

[illegible]

'WP_042508494.1'

[illegible]

'WP_118795991.1'

[illegible]

'WP_118845927.1'

[illegible]

'WP_118997288.1'

[illegible]

'WP_005703627.1__2_'

[illegible]

'WP_114922176.1'

[illegible]

VSEQSAFAYTTALNY-LL---------R-----RE-----------N------N------H------------CL------------------------T----------I-----G---------D----A----S--------TVF--W---------A------------E--------------A----------D----------D-------------------------D---------N-A---T-A---------------------Q---------------A-------------TE-------G-----------F-------F-----------A--------------------QV-----F-------------------------T-------P---------------P--------
-----------------D-D-----------E----------Q----E----------S-------------A----KVFNV-----LEQ-----I-----------------------G-K-G-----R-------PL--Q--E---IA-----P---E-------L-S------A----N----T-------R-------F--YILGL---APN--A-ARISVRFWLDT-TFGQLAENLAQHW--QD-------LA--LE----------P-------------------------CA------WK--------------T-----P--------------------P------------S----------IWRLL-------------------
------LQ-------------T---A--------------------V----------------------L-------G--------K-----------------------S----------E--N----------------I-----SP-----V--------------LAGEMTRAVIC--G-T--P---YPLSLLSQ-LI-------------T--------------R--I-----------------RA-------------------------D-------G-------------D-----VN-------------------------G----------L--------------RVAM------MKA------V--------------------L-------------E-------------R-
-----------R---------F------------R-------------K-G-----------------FIE-----------E-----------------G-----------------V-----------------P-------------M----SLN--------N--E---------SP-----N-------------RAYLL------GRLFAVLE-----RIQ----------------Y-------------Q----------A------L-------------G-------------E-------L---------N--------AGIA---DRYYGSAS-----AVPFSV-------------FPRLLS--GA-K-HHLS-------R--L----------RK-------------
D-K---------A------G--------------------M--AV-N----------L-----------------DKD-L--G---------EII-----------A-----KL-------------P------E-----T-------F------P--------R-----H---LS-ID-E---------QGRFAIGYY---H----Q----------------------------------------------------------------------------------K--------------------QS-------------Y---------F---------------------A-------------K------------------K------------------------ET--------A----------ET-----------
---------I------------EN--------------------------------------------------------------

'WP_101810652.1__2_' ---------------------------------------------------------------------------------MI------LASLARYYR---------R-----L--------------A----A---------------Q-I--------DDMGNPK-----------------VPPY--GF---SEE-KI-G------WILVLDK--E---GR-LK-----T-------A---V----------PN-------L---TA--------------D--------K----------KP--Q---P--KL---M------S-V-P-----Q--S--F--KR------------P---G---T----
-------T---------P--K-------P----F-------------F-L------------------------WDKTSYA-------L----G-------------------------------------V--------------------E---------------------ANKNKA-------------------------------------------------------------------------EA---------------------K---------------------------E--------------------------K-PFTS---S-------------------------E-----------K--------------T-----------------------------F-D------------AF-KQ-YHLD--L--LQN--------S-----E----
D--E---G--LQA---LCRFLQN----WQ-----------P------AHF---------A--------------A-------E-------N----------------------------------------------L--------------P--A--E-M-----------L-------------------------------------------------DAN------------------------I----------------------------AF------S-L--------------E----------------K---P-----------T----A-------L-------IH----------K----RE------A------------A-QT-LW--------------------------------------A------------------G------------Y----
---------L-----------------------------K-------------------------S-----------DE-------------A--------------------------L-------------E--------G--------------------L----------------------CL-----------ISG--D---T--A-P---IARL-H--P--A---I--K----G--------V--F--G---G--QS--SG-G----------------SIIS-----F---------------N------KE-------------------------A-------F-AS---F-------------G-----K---E--------------Q--------------------GA------------------------NAP--------
VSEQSAFAYTTALNY-LL---------R-----RE-----------N------N------H------------CL------------------------T----------I-----G---------D----A----S--------TVF--W---------A------------E--------------A----------D----------D-------------------------D---------N-A---T-A---------------------Q---------------A-------------AE-------G-----------F-------F-----------A--------------------QV-----F-------------------------T-------P---------------P--------
-----------------D-D-----------E----------Q----E----------S-------------A----KVFNV-----LEQ-----I-----------------------G-K-G-----R-------PL--Q--E---IA-----P---E-------L-S------A----N----T-------R-------F--YILGL---APN--A-ARISVRFWLDT-TFGQLAENLAQHW--QD-------LA--LE----------P-------------------------CA------WK--------------T-----P--------------------P------------S----------IWRLL-------------------
------LQ-------------T---A--------------------V----------------------L-------G--------K-----------------------S----------E--N----------------I-----SP-----V--------------LAGEMTRAVIC--G-T--P---YPLSLLSQ-LI-------------T--------------R--I-----------------RA-------------------------D-------G-------------D-----VN-------------------------G----------L--------------RVAM------MKA------V--------------------L-------------Q-------------
R-------------R---------F------------R-------------K-G-----------------FIE-----------E-----------------G-----------------V-----------------P-------------M----SLN--------N--E---------SP-----N-------------RAYLL------GRLFAVLE-----RIQ----------------Y-------------Q----------A------L-------------G-------------E-------L---------N--------AGIA---DRYYGSAS-----AVPFSV-------------FPRLLS--GA-K-HHLS-------C--L----------RK-------------
-D-K---------A------G--------------------M--AV-N----------L-----------------DKD-L--G---------EII-----------A-----KL-------------P------E-----T-------F------P--------R-----H---LS-ID-E---------QGRFAIGYY---H----Q----------------------------------------------------------------------------------K--------------------QS-------------Y---------F---------------------A-------------K------------------K------------------------ET--------A----------ET-----------
----------I------------EN--------------------------------------------------------------

'WP_111279113.1__2_' ---------------------------------------------------------------------------------MI------LASLARYYR---------R-----L--------------A----A---------------E-N--------DEMGNPK-----------------VPPY--DF---SEE-KI-G------WILVLDK--E---GC-LK-----T-------V---V----------PN-------L---TA--------------D--------K----------KP--Q---P--KL---M------S-V-P-----Q--S--F--KR------------P---G---T----
-------T---------P--K-------P----F-------------F-L------------------------WDKTSYA-------L----G-------------------------------------V--------------------E---------------------ANKNKA-------------------------------------------------------------------------EA---------------------K---------------------------E--------------------------K-PFTS---S-------------------------E-----------K--------------T-----------------------------F-D------------AF-KQ-YHLD--L--LQN--------S-----E----
D--E---G--LQA---LCRFLQN----WQ-----------P------AHF---------A--------------A-------E-------N----------------------------------------------L--------------P--A--E-M-----------L-------------------------------------------------DAN------------------------I----------------------------AF------S-L--------------E----------------K---P-----------T----A-------L-------IH----------K----RE------A------------A-QS-LW--------------------------------------A------------------G------------Y----
---------L-----------------------------K-------------------------S-----------DE-------------A--------------------------L-------------E--------G--------------------L----------------------CL-----------ISG--D---T--A-P---IARL-H--P--A---I--K----G--------V--F--G---G--QS--SG-G----------------SIIS-----F---------------N------KE-------------------------A-------F-AS---F-------------G-----K---E--------------Q--------------------GA------------------------NAP--------
VSEQSAFAYTTALNY-LL---------R-----RE-----------N------N------H------------CL------------------------T----------I-----G---------D----A----S--------TVF--W---------A------------E--------------A----------D----------D-------------------------D---------N-A---T-A---------------------Q---------------A-------------AE-------G-----------F-------F-----------A--------------------QV-----F-------------------------T-------P---------------P--------
-----------------D-D-----------E----------Q----E----------S-------------A----KVFNV-----LEQ-----I-----------------------G-K-G-----R-------PL--Q--E---IA-----P---E-------L-S------A----N----T-------R-------F--YILGL---APN--A-ARISVRFWLDT-TFGQLAENLAQHW--QD-------LA--LE----------P-------------------------CA------WK--------------T-----P--------------------P------------S----------IWRLL-------------------
------LQ-------------T---A--------------------V----------------------L-------G--------K-----------------------S----------E--N----------------I-----SP-----V--------------LAGEMTRAVIC--G-T--P---YPLSLLSQ-LI-------------T--------------R--I-----------------RA-------------------------D-------G-------------D-----VN-------------------------G----------L--------------RVAM------MKA------V--------------------L-------------Q-------------
R-------------R---------F------------R-------------K-G-----------------FIE-----------E-----------------G-----------------V-----------------P-------------M----SLN--------N--E---------SP-----N-------------RAYLL------GRLFAVLE-----RIQ----------------Y-------------Q----------A------L-------------G-------------E-------L---------N--------AGIA---DRYYGSAS-----AVPFSV-------------FPRLLS--GA-K-HHLS-------C--L----------RK-------------
-D-K---------A------G--------------------M--AV-N----------L-----------------DKD-L--G---------EII-----------A-----KL-------------P------E-----T-------F------P--------R-----H---LS-ID-E---------QGRFAIGYY---H----Q----------------------------------------------------------------------------------K--------------------QS-------------Y---------F---------------------A-------------K------------------K------------------------ET--------A----------ET-----------
----------I------------EN--------------------------------------------------------------

'WP_003777776.1' ---------------------------------------------------------------------------------MI------LASLARYYR---------R-----L--------------A----A---------------E-N--------DEMGNPK-----------------VPPY--GF---SEE-KI-G------WILVLDK--E---GR-LK-----T-------V---V----------PN-------L---TA--------------D--------K----------KP--Q---P--KL---M------S-V-P-----Q--S--F--KR------------P---G---T----
[illegible]

'WP_111302649.1__3_'
[illegible]

'WP_005642817.1'
[illegible]

'WP_107863166.1'
[illegible]

'WP_118800701.1'
[illegible]

'WP_118813523.1'
[illegible]

'WP_107810318.1'
[illegible]

'WP_107768950.1'
[illegible]

'WP_003680670.1'
[illegible]

'WP_118783216.1'
[illegible]

'WP_013448701.1'
[illegible]

'WP_021615275.1'
[illegible]

'WP_107867200.1'
[illegible]

'WP_109128002.1'
[illegible]

'WP_115002826.1'
[illegible]

'WP_109843264.1'
[illegible]

'WP_118775608.1'
[illegible]

'WP_118785320.1'
[illegible]

[illegible]

'WP_049344176.1'

[illegible]

'WP_003744750.1'

[illegible]

'WP_003758690.1'

[illegible]

'WP_106013917.1__2_'

[illegible]

'WP_045072974.1'

[illegible]

'WP_122254860.1__2_'

[illegible]

'WP_111300978.1__2_'

[illegible]

'WP_111711287.1'

[illegible]

'WP_046938673.1'

[illegible]

'WP_134079369.1'

[illegible]

'WP_111328360.1__2_'

[illegible]

'WP_107848845.1'

[illegible]

'WP_118792312.1'

[illegible]

'WP_107843122.1'

[illegible]

'WP_109860522.1'

[illegible]

'WP_005646984.1__2_'

[illegible]

'WP_109059347.1'

[illegible]

'WP_107971324.1'

[illegible]

'WP_049376143.1'

[illegible]

'WP_004520755.1'

[illegible]

[illegible]

'WP_012771280.1'

[illegible]

'WP_111407980.1__2_'

[illegible]

'WP_109083086.1'

[illegible]

'WP_107853752.1'

[illegible]

'WP_109859483.1'

[illegible]

'WP_109129734.1'

[illegible]

'WP_107810905.1'

[illegible]

'WP_107976851.1'

[illegible]

'WP_049353497.1'

[illegible]

'WP_049351751.1'

[illegible]

'WP_107820401.1__2_'

[illegible]

'WP_049328693.1'

[illegible]

'WP_050693402.1'

[illegible]

'WP_111402351.1__2_'

[illegible]

'WP_005702613.1'

[illegible]

'WP_083016776.1__2_'

[illegible]

'WP_003685597.1'

[illegible]

'WP_049225765.1'

[illegible]

'WP_046942529.1'

[illegible]

'WP_118843464.1'

[illegible]

'WP_108700278.1' [illegible]

'WP_003746232.1' [illegible]

'WP_046952760.1' [illegible]

'WP_114892516.1' [illegible]

'WP_005565917.1__2_' [illegible]

'BAS49431.1' [illegible]

'WP_005579127.1' [illegible]

'WP_005556489.1' [illegible]

'WP_061885970.1' [illegible]

'WP_011478970.1' [illegible]

'WP_013647393.1' [illegible]

'WP_013819641.1' [illegible]

'WP_062559737.1' [illegible]

'WP_130414895.1' [illegible]

'WP_037453459.1' [illegible]

'WP_081486769.1' [illegible]

'WP_102114222.1__2_' [illegible]

'WP_029009608.1' [illegible]

'WP_119507757.1__2_' [illegible]

'WP_035671734.1' [illegible]

'WP_098734979.1__2_' [illegible]

[illegible]

'WP_108547602.1__2_' [illegible]

[illegible]

'WP_111067557.1__2_' [illegible]

[illegible]

'WP_121938840.1' [illegible]

'WP_018575261.1' [illegible]

'WP_134243495.1__2_' [illegible]

'WP_109793955.1__2_' [illegible]

'WP_090486341.1' [illegible]

'WP_087666955.1' [illegible]

'WP_087631866.1' [illegible]

'WP_116560916.1' [illegible]

'WP_086382392.1' [illegible]

'WP_056358515.1' [illegible]

'WP_089166540.1' [illegible]

'WP_060817499.1' [illegible]

'WP_062168642.1' [illegible]

'WP_021031848.1' [illegible]

'WP_062374244.1' [illegible]

'WP_018916769.1' [illegible]

'WP_096922675.1__2_' [illegible]

'WP_081138323.1' [illegible]

[illegible]

'WP_113270886.1__2_'

[illegible]

'WP_096650223.1__2_'

[illegible]

'WP_019950945.1'

[illegible]

'WP_090135724.1'

[illegible]

'WP_010626368.1'

[illegible]

'WP_095620338.1__2_'

[illegible]

'WP_089683184.1'

[illegible]

'WP_133481786.1'

[illegible]

'WP_087718753.1'

[illegible]

'WP_102651739.1__2_'

[illegible]

'WP_110647215.1'

[illegible]

'WP_110060898.1'

[illegible]

'WP_058523910.1'

[illegible]

'OWQ84878.1'

[illegible]

'WP_135480430.1__2_'

[illegible]

'WP_077529552.1'

[illegible]

'WP_038271236.1'

[illegible]

'WP_110265304.1'

[illegible]

'WP_056763342.1'

[illegible]

'WP_028239961.1'

[illegible]

[illegible]

'WP_028079637.1'

[illegible]

'WP_122102149.1__2_'

[illegible]

'WP_024888940.1'

[illegible]

'WP_057637114.1'

[illegible]

'WP_014161866.1'

[illegible]

'WP_028920493.1'

[illegible]

'WP_045728617.1'

[illegible]

'WP_045770660.1'

[illegible]

'WP_045761072.1'

[illegible]

'WP_045764763.1'

[illegible]

'WP_012917358.1'

[illegible]

'WP_087785558.1'

[illegible]

'WP_113255644.1__2_'

[illegible]

'WP_113306785.1__2_'

[illegible]

'WP_044757099.1__17_'

[illegible]

'RBK06776.1'

[illegible]

'RBK21110.1'

[illegible]

'WP_075244868.1'

[illegible]

'QBN93702.1__232_'

[illegible]

'RBH73706.1__2_'

[illegible]

[illegible]

'REI38999.1'

[illegible]

'RBC33562.1'

[illegible]

'RBL01553.1'

[illegible]

'RBD00282.1'

[illegible]

'RBF88823.1'

[illegible]

'RBD06771.1'

[illegible]

'WP_017911788.1'

[illegible]

'WP_131082938.1__3_'

[illegible]

'WP_017907376.1'

[illegible]

'WP_053836965.1'

[illegible]

'WP_053840473.1'

[illegible]

'WP_053834102.1'

[illegible]

'WP_064521678.1'

[illegible]

'WP_058360755.1'

[illegible]

'WP_104603865.1__2_'

[illegible]

'WP_065468758.1__2_'

[illegible]

'WP_010372414.1'

[illegible]

'WP_017113403.1__10_'

[illegible]

'WP_010381903.1__5_'

[illegible]

'WP_017119023.1__28_'

[illegible]

'WP_017117513.1'

[illegible]

'WP_017116852.1'

[illegible]

'WP_042823989.1__2_'

[illegible]

'WP_029221384.1'

[illegible]

'WP_059032304.1'

[illegible]

'WP_005927103.1'

[illegible]

'WP_015472884.1'

[illegible]

'WP_040154518.1'

[illegible]

'PPT89228.1'

[illegible]

'WP_046977474.1__2_'

[illegible]

'WP_039006211.1'

[illegible]

'WP_003465420.1'

[illegible]

'WP_058195594.1'

[illegible]

'WP_058363849.1'

[illegible]

'WP_058364362.1'

[illegible]

'WP_058358538.1'

[illegible]

'WP_058363095.1'

[illegible]

'WP_003480731.1'

[illegible]

'WP_009605933.1'

[illegible]

'WP_114074393.1__2_'

[illegible]

'WP_011134781.1'

[illegible]

LRA---LRRFLQD----WR-----------P------EQF--------D-------------A------F-----P--------------------PE---------------L----------Q--E--S-L---------R
L-------------------------A--------------------E---------GG----------G-----------------D-----------------T-------G------------------M-------------CL-----------VSG--E---T--A-P
ISEFAAFAYTAALNH-LL---------R---REED-----------N------R------Q------------RL--------------------------Q---------I-----G---------D----A----T---------VVF--W---------A
----------------A---D-D------------A---------S----E-----------A-------------Q---RVLAA-----LQD-----I-------------------------A-D-G-----R-------PA--A--E---LG-----L----R-------L-E
-----------YA------------V---A-------------------A---------Q-----------------------G--------K----------------------A----------E--N-----------------V----PP----Q-------------LAGELARSILG
------R------------D----------Q------------R--------------L-G-------V------A-GIN----------L----------------E----------------L-----------------P------------V----SLD--------I--A---------NR
-------D-K---------P------G-------------------L--WK-T---------L-----------------ERD-L--A---------DII----------H------GL-------------D------A-----P-------L-----P-------R----N----LG-IA-A
------------------------DAA-------------------------A--------SEET-EE-----------------------
'WP_011396662.1' -------------------------------------------------------------------MI-------LSALNEYYQ---------R----L-----------A-----E
I-----------E------P----C-----------F-L-------------------WDKTSYV-------L-----G-------------------------------V-----------------S-------------------AK
LKA---VNLFLKN----WS-----------P------DQF--------C----------------------------P---------------------PH---------------F-----------K--E--E-M---------R
K--------------------------K--------------------N-----------DD------------A--------------------T-----------------E---------G-------------------R-----------------CL-----------VTG--V---Y--G-P
VSEQAAFAYTTVLNH-LL---------R---RDEQ-----------N------R------Q------------RL--------------------------Q---------I-----G---------D----A----S---------VVF--W---------A
----------------T---D-D-------------G-----------Q----E----------T---------------A----KLHSI-----LEK-----I---------------------------A-Q-G-----R-------PL--S--D---VN-----P---D-------L-D
-----------KA--------------L----A-----------------------------------V-----------L-------G---------K-----------------------T----------E--N-----------------I----PS-----K-------------LAGEMTRAILN
------R------------E----------Y-------------R--------------L-N-----------AN--R-SQT-------------E----------------E-----------------V---------------------P------------V----SLD---------T--E----------NS
--------E-K---------V-------G-------------------L--AK-A---------L-----------------ERD-V--S---------EII-----------D-----GL--------------G------S-----S---------F-----P-------R----S----LR-LD-D
------------------------GEIE-----------------------ASH------NEGE-IA-----------------------
'WP_040574346.1' -------------------------------------------------------------------MI-------LQALNSYYQ---------R----L-----------A-----D
--I------------R-------P----H-----------F-L--------------------WDKSSYV-------L-----G------------------------------V-----------------T---------------------A
[illegible]
'WP_093387232.1'
[illegible]
'WP_104449536.1__2_'
[illegible]
'WP_119952524.1__2_'
[illegible]
'WP_108215233.1'
[illegible]
'WP_104729694.1__2_'
[illegible]
'WP_012701708.1'
[illegible]
'WP_022654261.1'
[illegible]
'WP_011914755.1__3_'
[illegible]
'RCL51599.1'
[illegible]
'WP_128701110.1'
[illegible]
'WP_062877662.1'
[illegible]
'WP_094954022.1__4_'
[illegible]
'WP_031631553.1__2_'
[illegible]
'WP_116806645.1__2_'
[illegible]
'WP_089035046.1__16_'
[illegible]
'WP_121395709.1'
[illegible]
'WP_023103414.1__13_'
[illegible]
'WP_023119728.1__2_'
[illegible]

[illegible]

'WP_023104936.1' ------MI------LSALNDYYQ [illegible]

[illegible]

'WP_025981579.1__22_' ------MI------LSALNDYYQ [illegible]

[illegible]

'PXB81918.1' [illegible]

[illegible]

'WP_134612090.1' [illegible]

[illegible]

'WP_033985464.1' [illegible]

[illegible]

'WP_033993299.1__2_' [illegible]

[illegible]

'WP_121411920.1' [illegible]

[illegible]

'WP_095541598.1__4_' [illegible]

[illegible]

'WP_095549874.1__2_' [illegible]

[illegible]

'WP_132228792.1' [illegible]

[illegible]

'WP_005221618.1' [illegible]

[illegible]

'WP_020651078.1' [illegible]

[illegible]

'WP_028009467.1' [illegible]

[illegible]

'WP_096448291.1__2_' [illegible]

[illegible]

'WP_107221944.1__2_' [illegible]

[illegible]

'WP_018230090.1' [illegible]

[illegible]

'WP_133595157.1' [illegible]

[illegible]

'WP_006786410.1' [illegible]

[illegible]

'WP_004299431.1' [illegible]

[illegible]

'WP_002938511.1' [illegible]

[illegible]

[illegible]

'WP_004260234.1'

[illegible]

'WP_108972067.1__2_'

[illegible]

'WP_088602634.1__2_'

[illegible]

'WP_119647622.1__2_'

[illegible]

'WP_119536702.1__2_'

[illegible]

'WP_019703287.1'

[illegible]

'WP_027996905.1'

[illegible]

'WP_053244138.1'

[illegible]

'WP_103271225.1__2_'

[illegible]

'WP_097203033.1'

[illegible]

'WP_034391367.1'

[illegible]

'WP_046240458.1'

[illegible]

'WP_061289319.1'

[illegible]

'WP_016453850.1'

[illegible]

'WP_038210600.1'

[illegible]

'WP_092938605.1'

[illegible]

'WP_110088506.1__2_'

[illegible]

'WP_013596237.1'

[illegible]

'WP_013902815.1'

[illegible]

'WP_091816924.1'

[illegible]

[illegible]

'WP_092949865.1'

[illegible]

'WP_092840232.1'

[illegible]

'WP_026434161.1'

[illegible]

'WP_053842815.1'

[illegible]

'WP_107161118.1__2_'

[illegible]

'AVT07065.1'

[illegible]

'AVS78836.1'

[illegible]

'WP_036944117.1'

[illegible]

'WP_094184777.1'

[illegible]

'WP_090942187.1'

[illegible]

'WP_094606372.1'

[illegible]

'WP_134214738.1'

[illegible]

'WP_020990709.1'

[illegible]

'WP_009303812.1'

[illegible]

'WP_078695410.1'

[illegible]

'WP_077835923.1'

[illegible]

'AWK52121.1'

[illegible]

'PWL61154.1'

[illegible]

'WP_019177386.1'

[illegible]

'WP_048149154.1'

[illegible]

'WP_015286682.1__2_'
[illegible]

'PKL68306.1'
[illegible]

'PKL56999.1'
[illegible]

'WP_048111877.1'
[illegible]

'WP_130647022.1__2_'
[illegible]

'WP_004040744.1__2_'
[illegible]

'WP_048040842.1'
[illegible]

'WP_048045507.1'
[illegible]

'WP_048156522.1'
[illegible]

'WP_048121235.1'
[illegible]

'WP_048128075.1'
[illegible]

'WP_048131821.1'
[illegible]

'WP_048170471.1'
[illegible]

'WP_078684207.1'
[illegible]

'WP_028588589.1'
[illegible]

'WP_028318028.1'
[illegible]

'WP_092349385.1'
[illegible]

'WP_051958128.1'
[illegible]

'WP_020586896.1'
[illegible]

'WP_031388068.1'
[illegible]

'WP_092123893.1'
[illegible]

[illegible]

'WP_028571952.1'

[illegible]

'ADP88373.1__2_'

[illegible]

'ABM29987.1'

[illegible]

'WP_040099922.1'

[illegible]

'WP_040199761.1'

[illegible]

'WP_103114551.1__2_'

[illegible]

'WP_092080806.1'

[illegible]

'WP_054651141.1'

[illegible]

'WP_008870466.1'

[illegible]

'WP_027370387.1'

[illegible]

'WP_014323747.1'

[illegible]

'WP_118230345.1__3_'

[illegible]

'WP_092153981.1'

[illegible]

'WP_103080087.1__4_'

[illegible]

'WP_011639920.1'

[illegible]

'WP_085755348.1'

[illegible]

'WP_053111379.1'

[illegible]

'WP_044352468.1'

[illegible]

'WP_038261397.1'

[illegible]

'WP_031514027.1'

[illegible]

[illegible]

[illegible]

'PID74152.1' [illegible]

'WP_124330973.1' [illegible]

'WP_083761995.1' [illegible]

'GBF08184.1' [illegible]

'AEB11037.1' [illegible]

'EKP95535.1' [illegible]

'SHF51564.1' [illegible]

'OIQ61453.1' [illegible]

'SDF44684.1' [illegible]

'AEG34529.1' [illegible]

'AAS82464.1' [illegible]

'WP_051105287.1' [illegible]

'CCQ93019.1' [illegible]

'KLR73783.1' [illegible]

'KMY60017.1__2_' [illegible]

'KJE29173.1' [illegible]

'WP_077303197.1' [illegible]

'WP_018664638.1' [illegible]

'WP_035198648.1__2_' [illegible]

'WP_034772367.1' [illegible]

```
-------------------------Y------------------QQR-I--Q----------LIT-----------G------QL------------------------------KEF------P-------A-----V---LD-LQ-G---------QAEFALGYY---H----E------------------------------------------------------------------------------R------------------------QD-----------I--------Y--------------------------------K-----------------------K---------------KENMSADAANISDLNKTVVEGE-
'WP_026684802.1'
-----------------------------------------------------------------------------------------------------------------------------------------MY-------MRRLVEFAE---------Q----HA---------------N-----D------------------------------------------------------LPPV---GY----ARK-SY-Q-------WFVVITH---D---DR-FE------F-----------IQ----------AE-------R-------------------G-------EQ--------R------V-----------I-P---------NR----SR--------Q----S----G---------V----
----S--------P---I----------L-L-------------TDKAEYV-------F---Q------------------------------------L-----------------P----------------KNP---------------------------------------------------------ANA-------------------K-----------------------------------------N------------------------------A--------E--------R--------------------------------H-F---------AY-LK-LLKE--C--CEA----------T-----R---NE------V---VKK----
IVRILES--------------------------------------------------------------------K-------SW------------Q--L-P----------D-----G------M-------------------------KPD---------------------D--------F-------------ILF--------R-T---------E---------------A------G----------D-------F-----------PH---------E------DN-----K------------V--KK-FW-----------E-------DFL-----------L---------K---------N---------S----
----------------SKE----------EN-------------------------------------------------D-S-----------------------V--------------CF--------------ICG--R---S---A-P---AVKR-H---T---I----E---F-----S---L-----------G----K-ER--------T-------KLIS-------A---------------N------------K-----------S------A------Y-FS----Y-----------G-------------------L-E---N--------SE---------------------------------IAP--------TCFACEQKYGQALSY-LL----
--R---KYPN-----------K-------K-QHGGPH------------MM--------------------------------E---------I-------G----------N------V---T----------YYY--W--------T--------------V----Q------N----------------DP-ET---------------------------------------T---------------------SM--------------F------T---------S--------L-----S--------------------------------------D----------P---
--------D----------P---------D-----AVQRI-----LSS------P--------------------------F-R-R-------T-----E--------N-----YA-------------------S---L-D---------R-------I---CILAL----SAN--K-ARMVVRDYAEP-PVWEVKTQVRRYF---DA-----QQ---VV----------G-------------------AP--------------------K---------------P------------A------------SIYLL-------------------------------------------------------A-
----AC-MYRDG------RKEMRK---------------------------------------------------E---------------DVLDWMNWALY--G-K--R----LSERIVAK-IV--------------------R-I--------------QA------------------------RGGMT-------------------W-----------I------------HAAA------IRS------W--------------L-------------ISQN------------K------------
------------------------E---------------E--------------W---------------T--------V----DLD----------HA----R-----IT---PPYLC------GRLFAVLE-------AVQ-----------Y-------E------------------------AV----RG------------N------ETIA---SRFYAAAS-------STPRTI----------FGLLMR---NS-K-WHLS---------K---I-----------------SKENKGYEVR-----
--------Y-----------QQR-I--Q------LIT-----------G------QL-------KE----------F-----P-------A-----V---LD-LQ-G---------QAEFALGYY---H----E-------------------------------------R------------------QD---------I--------Y-------------------------K-----------------------KE-------------DMPADSANVSHLNKTVVEGEK----
'WP_061565738.1'
-----------------------------------------------------------------------------------------------------------------------------------------MY-------MRRLVEFAE---------Q----HA---------------N-----D------------------------------------------------------LPPV---GY----ARK-SY-Q-------WFVVITH---D---DC-FE------F-----------IQ----------AE-------R-------------------G-------EQ--------R------V-----------I-P---------NR----SR--------Q----S----G---------V----
----S--------P---I----------L-L-------------TDKAEYV-------F---Q------------------------------------L-----------------P----------------KNP---------------------------------------------------------ANA-------------------N-----------------------------------------R------------------------------A--------E--------R--------------------------------H-L---------AY-LK-LLKE--C--CEA----------T-----R---NE------I---VQK----
IVRILES--------------------------------------------------------------------K-------SW------------R-L-P----------D-----G------M-------------------------KPD---------------------D--------F-------------ILF--------R-T---------E---------------A------G----------D-------F-----------PH---------E------DN-----K------------V--KK-FW-----------E-------DFL-----------L---------K---------N---------S----
----------------SKE----------EN-------------------------------------------------D-S-----------------------V--------------CF--------------ICG--R---T---A-P---AVKR-H---T---I----E---F-----S---L-----------G----K-ER--------T-------KLIS-------A---------------N------------K-----------S------A------Y-FS----Y-----------G-------------------L-E---N--------SE---------------------------------IAP--------TCFACEQKYGQALSY-LL----
--R---KYPN-----------K-------K-QHGGPH------------MM--------------------------------E---------I-------G----------N------V---T----------YYY--W--------T--------------V----Q------N----------------DP-ET---------------------------------------T---------------------SM--------------F------S---------S--------L-----S--------------------------------------D----------P---
--------D----------P---------D-----AVQRI-----LSS------P--------------------------F-R-R-------T-----E--------N-----YA-------------------S---L-D---------R-------V---CILAL----SAN--K-ARMVVRDYAEP-PVWEVKAQIRRYF---DA-----QQ---VV----------G-------------------AP--------------------K---------------P------------A------------SIYLL-------------------------------------------------------A-
----AC-MYRDG------RKEMRK---------------------------------------------------E---------------DVLDWMNWALY--G-K--R----LSERIVAK-IV--------------------R-I--------------QA------------------------RGGMT-------------------W-----------I------------HAAA------IRS------W--------------L-------------ISQN------------K------------
------------------------E---------------E--------------W---------------T--------V----DLD----------HA----R-----IT---PPYLC------GRLFAVLE-------AVQ-----------Y-------E------------------------AV----RG------------N------ETIA---SRFYAAAS-------STPRTI----------FGLLMR---NS-K-WHLS---------K---I-----------------GKENKGYEVR-----
--------Y-----------QQR-I--Q------LIT-----------G------QL-------KE----------F-----P-------A-----V---LD-LQ-G---------QAEFALGYY---H----E-------------------------------------R------------------QD---------I--------Y-------------------------K-----------------------KE-------------DMSADSANVNHLNKTVVEGDK----
'WP_061577663.1'
-----------------------------------------------------------------------------------------------------------------------------------------MY-------MRRLVEFAE---------Q-H----AN-----------------D------------------------------------------------------LPPV---GY----ARK-SY-Q-------WFVVITH---D---DR-FE------F--IQ----------AE-------R-------------------GE-----------QR---V--------I-P---------NR----------S-RQS-----G---------V----
----S--------P---I----------L-L-------------TDKAEYV-------F----------------------------------------------------------------------------------------------------------------------------------------Q-LPKNPANANRA------------A------E--------R--------------------------------H-L---------AY-LK-LLKE--C--CEA----------T-----R---NE------V---VQK----
IVRILES--KSWQ------L-----P------DGM-----------K-------P-------------------------------------------D-D-F-------------------------------------------------------------------------LF--------R-T---------EAG--D--------F----------PH---------E------DN-----K------------V--KK-FW-----------E-------DFL---------------E------D----------F------LLKN-S----
----------------SK------------------------------E-E----------N--------D-S-----------------------V--------------CF--------------ICG--R---T---A-P---AVKR-H---T---I----E---------FSLG-------------KERT---------KLIS----A----------N--------------KS-----A------Y-FS----Y-----------G--------------------LE------NSE-----------------------IAP--------TCFACEQKYGQALSY-LL----
--R---KYPN-----------K-------K-QHGGPH------------MM--------------------------------E---------I-------G----------N------V---T----------YYY--W--------T--------------V----Q------N----------------DP-ET-------------------------Q--------NDP----------ET-------T---------SM----------F---------T-------------SL----SDP---------------------------------------------------P---
--------D----------P---------D-----AVQRI-----LSS------P--------------------------F-R-R-------T-----E------------TE-------------------NYASLD------R------V---SILAL----SAN--K-ARMVVRDYAEP-PVWEVKAQIRRYF---DA-----QQ---VV----------G-------------------AP--------------------K---------------P------------A------------SIYLL-------------------------------------------------------A-
----AC-MYRDG------RKEMRK---------------------------------------------------E---------------DVLDWMNWALY--G-K--R----LSERIVAK-IV--------------------R-I--------------QA------------------------RGGMT-------------------W-----------I------------HAAA------IRS------W--------------L-------------ISQN------------K------------
------------------------E---------------E--------------W---------------T--------V----DLD----------HA----R-----IT---PPYLC------GRLFAVLE-------AVQ-----------Y-------E------------------------AV----RG------------N------ETIA---SRFYAAAS-------STPRTI----------FGLLMR---NS-K-WHLS---------K---I-----------------GKENKGYEVR-----
--------Y-----------QQR-I--Q------LIT-----------G------QL-------KE----------F-----P-------A-----V---LD-LQ-G---------QAEFALGYY---H----E-------------------------------------R------------------QD---------I--------Y-------------------------K-----------------------KE-------------DMSADSANVSHLNKTVVEGEK----
'WP_104851189.1__2_'
-----------------------------------------------------------------------------------------------------------------------------------------MY-------MRRLVEFAE---------Q----HA---------------N-----D------------------------------------------------------LPPV---GY----ARR-SY-Q-------WFVVITH---D---DR-FE------F-----------IQ----------AD-------R-------------------G-------EQ--------R------V-----------I-P---------NR----SR--------A----S----G---------V----
----S--------P---I----------L-L-------------TDKAEYV-------F---K------------------------------------L-----------------P----------------KNP---------------------------------------------------------ANA-------------------N-----------------------------------------R------------------------------A--------E--------C--------------------------------H-L---------AY-MK-LLKE--C--CEA----------T-----R---NE------L---VQK----
IVQILER--------------------------------------------------------------------Q------DW------------Q--I-P----------D-----G------M-------------------------KST---------------------D--------V-------------ILF--------R-T---------E---------------A------G----------D-------F-----------PH---------E------DN-----K------------V--KK-FW-----------E-------DFL-----------I---------K---------N---------S----
----------------SKE----------EN-------------------------------------------------D-S-----------------------V--------------CF--------------ICG--C---S---A-P---AVKR-H---T---I----E---F-----S---V-----------G----K-ER--------T-------KLIS-------A---------------N------------K-----------S------A------Y-LS----Y-----------G-------------------L-E---N--------SE---------------------------------IAP--------TCFECEQKYGQALSY-LL----
--R---KYAN-----------K-------K-QHGGPH------------MM--------------------------------E---------I-------G----------N------V---T----------YYY--W--------A--------------V----Q------N----------------DP-ET---------------------------------------T---------------------SM--------------F------T---------S--------L-----S--------------------------------------D----------P---
--------D----------P---------V-----AVRRI-----LSS------P--------------------------F-K-R-------A-----E--------N-----HG-------------------S---L-D---------R-------I---CILAL----SAN--K-ARMVVRDYAEP-PVWKVKNQIRQFFYAQQ-----------IV---------G-------------------AP--------------------K---------------P------------A------------SIYLL-----------------------------------------------------------
AACMYRDGRKEMQ----------------------------------------------KE---------------DVQDWMNWAIY--G-K--R----LSGRIIAK-MV--------------------R-I--------------QA-------------------------------RGGMA-------------------W-----------I------------HAAA------IKS------W--------------L-------------ISQK------------K------------
------------------------E---------------E--------------W---------------T--------V----DLD----------AQ-----RV---T------PAYLC------GRLFAVLE-------AVQ-----------Y-------E------------------------AV----KG------------N------ETIA---SRFYAAAS-------STPRTI----------FGLLMR---NS-K-WHLS---------K---I-----------------SKENKGYEVR-----
--------Y-----------QQR-I--Q------LIT-----------G------QL-------KEF-------P-------A-----V---LD-LQ-G---------QAEFALGYY---H----E-------------------------------------R------------------QD---------I--------Y-------------------------K-----------------------KE-------------KENMSADAANVSDFNNTVVEGE-----
'AJH77393.1__2_'
-----------------------------------------------------------------------------------------------------------------------------------------MY-------MRRLVEFAE---------Q-H----AN-----------------D------------------------------------------------------LPPV---GY----ARK-SY-Q-------WFVVITH---D---DR-FE------F--IQ----------AE-------R-------------------GE-----------QR---V--------I-P---------NR----------S-RAS-----G---------V----
----S--------P---I----------L-L-------------TDKAEYV-------F---KLP--------------------------------------------------------------------------------------------------------------------------------------------------------PANAKRA---AE------C--------------------------------H-L---------AY-MK-LLKE--C--CEA----------T-----R---NE------L---VQK----
IVQILER---QDWQ------I-----P------DGM-----------K-------S---------------------------------------------T-D-V-------------------------------------------------------------------------LF--------R-T---------EAG--D--------F----------PH---------E------DN-----K------------V--KK-FW-----------E-------DFL---------------I------D----------F------LLKN-S----
----------------SK------------------------------E-E----------N--------D-S-----------------------V--------------CF--------------ICG--R---S---A-P---AVKR-H---T---I----E---------FSVG-------------KERT---------KLIS----A----------N--------------ES-----A------Y-LS----Y-----------G--------------------LENSE-----------------------IAP--------TCFECEQKYGQALSY-LL----
--R---KYAN-----------K-------K-QHGGPH------------MM--------------------------------E---------I-------G----------N------V---T----------YYY--W--------A--------------V----Q------N----------------DP-ET-------------------------T-----------SM----------F---------T-------------S-------------------L-------------------------------------------------------------P---
--------D----------P---------V-----AVRRI-----LSS------P--------------------------F-K-R-------A-----E--------N-----HG-------------------S---L-D---------R-------I---CILAL----SAN--K-ARMVVRDYAEP-PVWKVKNQIRQFF---DA-----QQ---IV----------G-------------------AP--------------------K---------------P------------A------------SIYLL-----------------------------------------------------------C---
MYRDGRKEM----------------------------------------------------------QK-----E-----DVQDWMNWALY--G-K--R----LSGRIIAK-MV--------------------R-I--------------QA-------------------------------RGGMT-------------------W-----------I------------HAAA------IRS------W--------------L-------------ISQN------------K------------
------------------------E---------------E--------------W---------------T--------V----DLD----------HA----RI-----T------PPYLC------GRLFAVLE-------AVQ-----------Y-------E------------------------AV----KG------------N------ETIA---SRFYAAAS-------STPRTI----------FGLLMR---NS-K-WHLS---------K---I-----------------SKENKGYEVR-----
--------Y-----------QQR-I--Q------LIT-----------G------QL-------KEF-------P-------A-----V---LD-LQ-G---------QAEFALGYY---H----E-------------------------------------R------------------QD---------I--------Y-------------------------K-----------------------K-------------KENMPADAANVSDLNKTVVEGE-----
'WP_133537158.1'
-----------------------------------------------------------------------------------------------------------------------------------------MY-------MRRLVEFAE---------Q----HA---------------N-----D------------------------------------------------------LPPV---GY----ARK-SY-Q-------WFVVITH---D---DR-FE------F-----------IQ----------AE-------R-------------------G-------EQ--------R------V-----------I-P---------NR----SR--------Q----S----G---------V----
----S--------P---I----------L-L-------------TDKAEYV-------F---Q------------------------------------L-----------------P----------------KNP---------------------------------------------------------ANA-------------------N-----------------------------------------R------------------------------A--------E--------R--------------------------------H-L---------AY-LK-LLKE--C--CEA----------T-----R---NE------V---VQK----
IVRILES--------------------------------------------------------------------K-------SW------------R-L-P----------D-----G------M-------------------------KPD---------------------D--------F-------------ILF--------R-T---------E---------------A------G----------D-------F-----------PH---------E------DN-----K------------V--KK-FW-----------E-------DFL-----------L---------K---------N---------S----
----------------SKE----------EN-------------------------------------------------D-S-----------------------V--------------CF--------------ICG--R---N---A-P---AVKR-H---T---I----E---F-----S---L-----------G----K-ER--------T-------KLIS-------A---------------N------------K-----------S------A------Y-FS----Y-----------G-------------------L-E---N--------SE---------------------------------IAP--------TCFECEQKYGQALSY-LL----
--R---KYAN-----------K-------K-QHGGPH------------MM--------------------------------E---------I-------G----------N------V---T----------YYY--W--------A--------------V----Q------N----------------DP-ET---------------------------------------T---------------------SM--------------F------T---------S--------L-----S--------------------------------------D----------P---
--------D----------P---------V-----AVRRI-----LSS------P--------------------------F-K-R-------A-----E--------N-----HG-------------------S---L-D---------R-------I---CILAL----SAN--K-ARMVVRDYAEP-PVWKVKNQIRQFF---DA-----QQ---IV----------G-------------------AP--------------------K---------------P------------A------------SIYLL-----------------------------------------------------------
----AC-MYRDG------RKEMQK---------------------------------------------------E---------------DVQDWMNWALY--G-K--R----LSERIVAK-IV--------------------R-I--------------QA------------------------RGGMT-------------------W-----------I------------HAAA------IRS------W--------------L-------------ISQN------------K------------
------------------------E---------------E--------------W---------------T--------V----DLD----------HA----R-----IT---PAYLC------GRLFAVLE-------AVQ-----------Y-------E------------------------AV----RG------------N------ETIA---SRFYAAAS-------STPRTI----------FGLLMR---NS-K-WHLS---------K---I-----------------GKENKGYEVR-----
--------Y-----------QQR-I--Q------LIT-----------G------QL-------KE----------F-----P-------A-----V---LD-LQ-G---------QAEFALGYY---H----E-------------------------------------R------------------QD---------I--------Y-------------------------K-----------------------KE-------------DMPADSANVSHLNKTVVEGEK----
'WP_017552737.1'
-----------------------------------------------------------------------------------------------------------------------------------------MY-------MRRLVEFAE---------Q-H----AN-----------------D------------------------------------------------------LPPV---GY----ARK-SY-Q-------WFVVITH---D---DR-FE------F--IQ----------AE-------R-------------------GE-----------QR---V--------I-P---------NR----------S-RQS-----G---------V----
----S--------P---I----------L-L-------------TDKAEYV-------F---QLP--------------------------------------------------------------------------------------------------------------------------------------------------------PANANRA---VE------R--------------------------------H-L---------AY-LK-LLKE--C--CEA----------T-----R---NE------V---VQK----
IVRILES---KSWQ------L-----P------DGM-----------K-------P---------------------------------------------D-D-F-------------------------------------------------------------------------LF--------R-T---------EAG--D--------F----------PH---------E------DN-----K------------V--KK-FW-----------E-------DFL---------------E------D----------F------LLKN-S----
----------------SK------------------------------E-E----------N--------D-S-----------------------V--------------CF--------------ICG--R---S---A-P---AVKR-H---T---I----E---------FSVG-------------KERT---------KLIS----A----------N--------------ES-----A------Y-LS----Y-----------G--------------------LENSE-----------------------IAP--------TCFECEQKYGQALSY-LL----
--R---KYAN-----------K-------K-QHGGPH------------MM--------------------------------E---------I-------G----------N------V---T----------YYY--W--------A--------------V----Q------N----------------DP-ET-------------------------T-----------SM----------F---------T-------------S-------------------L-------------------------------------------------------------P---
--------D----------P---------V-----AVRRI-----LSS------P--------------------------F-K-R-------A-----E--------N-----HG-------------------S---L-D---------R-------I---CILAL----SAN--K-ARMVVRDYAEP-PVWKVKNQIRQFF---DA-----QQ---IV----------G-------------------AP--------------------K---------------P------------A------------SIYLL-----------------------------------------------------------C---
MYRDGRKEM----------------------------------------------------------QK-----E-----DVQDWMNWAIY--G-K--R----LSGRIIAK-MV--------------------R-I--------------QA-------------------------------RGGMA-------------------W-----------I------------HAAA------IKS------W--------------L-------------ISQK------------K------------
------------------------E---------------E--------------W---------------T--------V----DLD----------AQ-----RV---T------PAYLC------GRLFAVLE-------AVQ-----------Y-------E------------------------AV----KG------------N------ETIA---SRFYAAAS-------STPRTI----------FGLLMR---NS-K-WHLS---------K---I-----------------SKENKGYEVR-----
--------Y-----------QQR-I--Q------LIT-----------G------QL-------KEF-------P-------A-----V---LD-LQ-G---------QAEFALGYY---H----E-------------------------------------R------------------QD---------I--------Y-------------------------K-----------------------K-------------KENMPADAANVSDLNNTVVEGE-----
'WP_039963358.1'
-----------------------------------------------------------------------------------------------------------------------------------------MI-------LQALAQLAE----------Q------E--------G-----L---------------------------------------------------------VARP---DY---EPK-RV-S------WVVLTE---D---GS-FV------N--------LV----------SH--------R--------------I--------NLAE---G-TKKK-PV-YVG---KW-----V-------Q-V-P------K------QP--TR--------T-----S----G--
--------A---------K-----A------F--------I-------------IDKSEYV-------F-----G---------------------------L-----------D------PQSK----------------------------------------------------------QKQ-------------------K--------------------------------------L---------------------E--------------------------------------------F-G--------LF-RA-QVNE--A--AKS------S-----S---AM--
P-------LQA--VAKFLNR-----------------LA-------DDP---------S------P---I-----------K--------------------T----H--------A---NF-------D----S-I---------------------------------------IGF-----------K-I--------------------------L---------------------G-----T--------Q------------W----------VQ-------E------AP-------R-----V-IA-YF------------DSR----------K------------S-------
----------------------------PEA--------------SD----------A----------------------------------------------P-F--------------------------------------------------------------------D---------E-----S-----------------------NDL----------------------------------------------------------------F-LS---H-------G-----------------L--S--G--------ND--------------------------NAP---------
ISRTAAEASAAALTR-LL--D----P-------NFPDPKNLDDVLP---L------R-------------QI-------------------------------G--------AN-------TI-------VVF--W---------A-------------K---NSAQE---IA-S------------------------ALVS-------H---------------N------A--------S------A--------S--------------------D--------MLNPLL---GGE-----------------------------
-------------------------------------------------------------S-----------------E--------E---TVQEA-----YRS-----V-----------------------W-H-G--------K---PV--------PL------------N-----------E---S---E------F-YALTL---SGT-Q-GRVVIRDWIET-SVEVVNENLAMHF---DD-------LK-VC-------------------------------E-------------------------NT--------------------E--------------KKGSKPSETIPMSWLL-------
-----------KS----------------------L----AAEGRSEPV---------------------------------------------------PPL--------------------------VESGFVRAAFQ--G-S--E----YPFQLLQR-AL----------------------------------------------------------------------------------A------------------------------------------------RAAI----LRA------V----------------
---N--------R----------------------------------------------------K--------------RRN---D---------------------PL------------IPKYPEVD------FE-------------MNPNVQS---------------DGYSC------GMLMAVLE---RLQ-------------------------------------------------------------------AL----DD--------I---------N-----ASVV---DRFFSAAS-----ASPRSV-----------FVRLLK---NS-Q-HHYR-------K---V--
------------------RDDDEKRGFARY--------------------------------------------------L----------ERI-K-D------EIL-----------------T------RF-------------DPQPSPSARSVYPGSASGI----P-------L-----H----LN-LE-Q---------QALFVLGYHQMRHWLRMS---------------------------------------------------------------------------K----------------------------DE--------------------------------------
QAWHDEHPNVPPAFGRKKVEQPEPEPANTIS-----
'WP_018106359.1'
-----------------------------------------------------------------------------------------------------------------------------------------MI-------LQSLYERDK--------A----LG--------E------K---------------------------------------------------------LPD---GF---VWK-EI-P------LIIVIDL---A---GN-FV------S-----------ME------------GL------------------------------DR-----E-P-KD-K---G-KP---Y-----M-I-P--------T---PV--TR----------T-----S----G--
-----VS------P---N--------L-F---------------WDHYGYV------L-----G---------------------------L-------------A--------------------K------------------------------------KER-------------A---------------------K---------------N-----------------------------------------------R--------------------------------M-D--------------AF-VE-QVRQ--F--AEK-------F----P----SN---S----V--
FQA----VKAFYDK-----------------------------EE----------Y----------LR-ITQ------D-------------------------E---D-------QV----------S-A-I-----------T-----K-----------------------------PGA---------------------------MTF------C-V------------S-----------------S--------P--------GDSD------------R--------LV--------T---SEQ------D-------L-KD-YQ-------Y------------A---------G----
---------Q-----------------------------------------TDG--------E---------------------------------E-A----------------------------------------------------------------------------------------------------------------------------------------------------------------------------------------------------------------------------------K---S-Q--------YV----------------NAP------
VSPQAAEGYASALND-LL----G------K-----G-------------------N-------------------F-------------------L---A----------G---------------ITG--R---KG--P---ISRL-H-P---G--I-K---L--S----------G-MP-----I--------SLVN----F-----------S--------S-----P---GDSD--------------R-----------D----------------L-KD-YQ-----------------------------------
LQIDRS-------------------------------------------------------------------------------------------------------------------------------------------------------------------------------------------------------------------------------------------------------S--------------------------------HADEDKLAGDFK-----SAT----------------------------------------------------PPLAPKEENGKKRKPTKGKGKTP-
LYALV-----------------------------------------------------G-----ALMPAA---LKADKWPKR---------------------------------------------------R-G--------------------------------PIEALVRSITT---G-A--P---YPADLYAS-CL-------------------------------------------------------------------------------------------------------RNVS------------------E-------L---------RAAY-------IKG-----Y---------
----------------KRDNPALYTI---------------------------------C---------------AKQ-L---D-----EIM------------AL------------LGGNESPY----P----M----H----FT-LE-E--------QGFFSLGYY---H---Q-----------------------------------------------------------K-----------------------------------QY--------H-----------Y----------S--------------------------------
KEEKEAIDNNSSED------------------
'WP_078737129.1'
-----------------------------------------------------------------------------------------------------------------------------------------MI-------LQALYNYYW---------K---LA-------------A------Y--------------------------QEQLD-E-------E-EGGRK-----------------IAPP---GM---EWK-TI-P----YIVVIKS---D---GT-FV-----R--------------IE-----------KT------------------------------------------------------------------------------------------------D---N---QL-Y------L-I-P--------K------DN---AR------T-----S----Q--
-------VE----A---Q-----------T-L-------------------------L----------------------------------------------P-------------KTDND---------------------------------------------------SDK-----------------------K------------------------------------------------------------------------------------------------------------F-D---------AF-NK-RVLE--L-----------------P-----DT---V----S--
IKA----VKLFYQK---------------------------EE-------Y------LK-AR-------------------------D----Q--------------------------S-E-I-------K-S--------------------------------------------KGG---------------------------------------ITF-----V-I------------------E-------------DKDR--------------I---------E-------A------SLP-------E-------V-IE-VM---------D--------PRI----------R---------E--------N---------H-
--------------------------------EPH------------G-------------------------------------------------------------------R-M---------T----------CI-----------VTG--K---QD-S-S---IAPT-H---E-K---I---A-----L------------------------GK-DS-G----------PLIG------I--------------Q-----------KA--------S-----G--F-DS-------Y---------------------------G--------K-Q-Q--------GF-------------NAP-----
ISFEASDAIGTALKE-LL--------R----KD--------------N-----G-----------------T-------------S---L-----------------------------------------T--------YIF--W-------------S-LLD----------------RE-FI------------------------------------------------------------NYQ--------KF----------A--------F-------------L----------------ET---SKQ-----SKDDSD-----TPSE-----------D--------------
P---------------E-----------K-------K----------------NLRKA-------MKM---Y-------------------A-G-G-----K-----N-----S---L--NG----I----D------Y----G----K---E------G----R---F-FMLGL---MPN--S-GRIAVKFWAEG-TVQEMTQNILTHM---ED-------FH-II------------------------------K--------------------------------------------------------------NGKIDPMPQP----------QS-------LHKII-------
-------------YS-------------I----------M---PKGKKIDKM--------------------------------------------------------------ST-----H----------------LLEELIESIVK--N-K---P---YPSLVQKL-CL--------------------------------K----N-----------RR-------------------------------------------------------E-------------------------M-------------------RAAI-----LKA-----------------N------
------------------------------------K----NRFQKRNIVM----Q-----------------------------------------V----GLD--------KN--------------QT-----D----------VGYLG----GRLLALYE-----YTQ---------------E------K----------------------------AL---GD-------V-----------TSIK----DRFYSKAS-----VVPRTV-----------FHKMTV---LH-Q-KHIR-------K---I----
---------SRNKPKTAYF-------------------------------L--------TKV-H--Q-----EIR------------NL------------------FPGSELVF------P-----Q---N----P-------H-----FT-LD-Q--------QNMFDNGYY---H---Q-----------------------------------------------------------------------KELRTLPPIDPQNDK-------------------------------------------
EEENPQKSK------------------------
'WP_081779600.1'
-----------------------------------------------------------------------------------------------------------------------------------------MQNIQESKG----------------------------------------------------------------------------------------------GR-------------ITFCIEE---N-----------------------QV-----------------------------------------------------------------------------------------------------------------------------------------------------------
--------------------------------------------------------------------------------------------------------------------------------------------------------------------------------------------------------------------------------------------------------Q------------------DF-----------V------------AS----L-----P--------------------------------------------------------------------------V---
-------------------------------------------------------------------------------------------KSN------------------------------------------------------------------------------------------------------------------------------G---------------------------------------------------------------------------------------------------------------------------------------------------------------
------------------CL-------VTG-----------A-PSRPVAIT-H--EKIA----I-------------G-----R------DN-A------------PFIG--------I-------------------KGCG------F-DS---Y-------G-------------------RIQ--------GF-----------NAP-----------ISFEASDAIGSALKE-LL-----------R----KG-----------------G-------D----T------------SM--------------------------------------------------R---------N------T---
--V-------YVF--W----------S-------------------S-LQD-------------PN-FI-----------------------K-------------NYR----------KL--------A------F-------Y--------T--------------EK---NQD-------NNSEEE------NDDD------------DTAN-----P-------------------S----------------E------------E------ELLYA-------MKM-------Y------------I-S-A---
----S----G-A-----S---RN-----F--E-------D------E----G------R----F-FMLGL---MPN--S-GRIAVKFWAEG-SISEIVQNALIHL---ED--------LN-II-------------------------------Q------------------DGNIENSALP----------KS---------LYGIL----------------------HS----------L----K------PKGKKIEKM-----------------------------------
-------PG----S-----------LLEGIIESIVK--N-K--P---YPALVQQM-CL------------------------------R-----------N---------------------------------------------------RR----------------------------------------------------------RAAI-------LKA-------Y--------------------------------K------NRHLNKTTI-----M-----------
Q-----------T----GLD-----KT---------QT------D------VGYLA-----GRLFALYE-----YTQ------------------------AL----GE------V--------SSIT---DRYFSQAS------LTPTAV-------FPKLRR--LN-M-KHLK--------K---L------------RTNKKGLAFK-----------------------------AKE-I---E-----SIQ--------LL------
IPCSGTGF-----P------Q-----R---FS-LD-Q---------QTTFGNGYY----H---Q--------------------------------------------------------------------------------------------------------------------------------------------------------------------------------------------------------------------------------------
'WP_018360775.1'
-----------------------------------------------------------------------------------------------------------------------------------------MI-------LQALYNYYW------NLSSYQ--------E-----SL------GDEGRR----------------------------------VPPP---GM---EWK-TV-P-----FLIIIKL---D---GT-FV----R--------------------NRPQKETDIVETSTTNTDLSDNENE-----------------------------------------------------------------------------
------------E----A-------QTL--------------WDHCGYL-------L------G-----------------------------------------------------------------------------------------------------------------------------------------------------------------------P------------------------KTDEKSDKENFTKQ-------------------DN------------F-D----------AF-QK-KIYS--L---PES-------D-------E----------INA-
-VKLFYKK-------------KEY--------LKA------R-------EHKYIQNI------------------------------Q-ESK--------------------------------------------------------------------------------------------------------------------------TF----C-I--------------EEN-LVQ-------------------------------------------------------------------------------------------------------F---VNPVNKSNGKE-
-------------------------DT------------------------N-M------------------------------R-------------CL------------VTG--------A-PSRPVAIT-H--E--K---I-----I-----G-----R------DN-A------------PFIGIQKGC--------------------------------------------------------------G-----F-DS--------------------------GF------------NAP-----ISFEASDAIGSALKE-LL-
--------R----KG--------G-----D-----T-----------SM-------------------------R---------I----G-----N------T----V-------YVF-W----------S-----------------S-LQD-------------PN-FI--------------K-------------NYR----------KL--------A-----F-----Y-------------------EK---NQD-------NNSEEE------NDDD------------DTAN-----P-------------------S-----------
-E-------------K--------D--------A-------E-----ELLYA------MKM-------Y-------------I-S-A------S------G-A-----S--------RN------F-----E--------D------E-----G----R---F-FMLGL---MPN--S-GRIAVKFWAEG-SISEIVQNALIHL---ED--------LN-II-------------------------------Q------------------WD-GN-IENS-A-------------LP--------KSLYG-------ILHSL------------------KP-----
-----------------------K-----G---KKIEK---------M-------------------------------MPG--S-------------LLEGIIESIVK--N-K--P----YPALVQQM-CL--------------------------R------------N--------------------------------------------RR--------------------------------------------VTEM----RAAI-----LKA--------Y-------------N-------N-RK-------
----------------L------AFKMAKEIESI----Q------LLI----P-CS-------GT------G-----------F-----P-----Q-----R----FS-LD-Q------QTTFGNGYY----H----Q----------------------------------------------------------------------VE---------L------------------------E-----------------------RT-----NKK-----
G-------------------------------------------------------------------------------------------------------------------------------------------------------------------------------------------------------------------------------------------------------------------------------------------------------------------------------------------------------------------------------------------
ETSTTNT---DLSD----NENE-----------
'WP_004331271.1'
-----------------------------------------------------------------------------------------------------------------------------------------MI-------LQALYNYYE------AMNKLQ-------------L----------------------------------------------------IAPP---GR---EYK-AI-A-----WAIIIKE---D---GS-FV----R---------LK---------PL------K-----------------------------DT-------------------SDKNSNGKE---Y------LVI-P---------AV---GR-----------T-----S----G--------I-
----------E----A----N------H-L---------------WDKKMYI------LGIDKG------------------------------------------------------------------------------------------------------SKTK-------------------------------------------------------------------------------------------------------------------------------------------------------------------------------------------------------
-VQQFYSK-----------GIHKSP-------EVL----------------------------------------SF--------------K-D-V------N----------------------------------------------------ETD-------------W--------------------------ISF-----Q-L----------------KGSTSI-------------------I----------AS----D-----ND----A-----------L-------SW--------------------------------------------------------------------------
-------------------------AE-------------------------EQLSSGKITG-----------------------L---------------------------CL---------ITG--------L-EGP---LVEK-H-T---P--I-----Q-----------------LPDSKRG------------DG-T---------PLVS----F--------------S-----------------N--------------QP---A----F-CS----Y-----------------------G--------R--------GVKDLD-----------NAP------ISKYAAFAYATAFNS-
LC--------D--------N----P--S-----T-------------HY---------------------D--------L----G-------S----N---R---YVF-W----------S---------------------------------------------SDVLN-------ES-IN------------------------------------I-----------TPR----------DV------T-----F---------V-------D----------------HR----NKQ--------DEEEES---------TIEEDKPKQRNRSKD--------KALA-----------P------
-------------T-------Q---------E------------T-----------H---KLLEQ---FKA---I------------------R-G-N---R---G-E----G-----QH------W---K--------SQKFQLN---------LVQSIVESILN--S-T----P----YPLTLQQA-CI--------D-----R----Q-------------------------PQ-------------ILRQKNEKEKKYKKRFD---------------------------------------------------TGNVDEENP---------------P---LRSLYKIV---
-----------------S--TVSPVSTSP-------------------------------------------------------------S------------------------------------------------------------------------------------------------------------------------------------------------------------------------------------------------------------------------------------------CVAI--------LKG-------------------------------------
---R----------------------------------K----ARVYKHLK--E-----L----E-------------------------------------------D-------------M-------ALD--------KQ--------ND-------N--------IAYLS-----GRLFAVFE----QVQ-----------R-----------E-------A-----------------------ALG---KG-------V------N-------ATIR---DRFYASAS------TRPNTV------MGPLFR--LS-N-HHLS---------K--------------------
-------YKETPGLAVF------------------------------F-------------------DKL-L----N---EIT--------------EL---------------ITTKEKAF-----P------Q-----R-------T----FS-LD-E-------QSLFAVGYY---H----Q--------------------------------------------------------------K-------------------VY-------------------T------------------------KEDADNPTEEIEQ-------
'WP_013760347.1__2_'
-----------------------------------------------------------------------------------------------------------------------------------------MI-------LQALYNYYE------------A------M------RKAHR-------------------------------------IATP---GL---EVK-PI-R-----WVIVIRE---D---GS-FV----R---------LKNLK------DPSDK------------------------------------SK------------------G---QD---Y-------L-V-P----------AGLKR--------------S-----G--
K------------CPPAQP----------------------F-------------WDNRKYI------L----G------------------------------------------------------------------------------------------------------------------------------------------------------------------------------------------E-------E------S-------------A-EGVSPCTDRLEDF-KR-LMAE--I--AEL---------Y-------PDNAS-
-FQA---VCRFYEH-----------YLGDEMPLP------EDA---------------------------H-------------------------------------E-R-I------L---A-----------------------------SDW-------------------M------------TF-----Q-L--------QGDT-----EL------LATQD--------D------------------------AH-------D------------------------Y-TAQAY------------------------------
---------------------------EE---------N-L-----------------------------------------------------------------T-----L----------------CL------ITG---E---K-K-S---LASF-H-------A---R-----------IVAP---G------SQ---AG-A---------TIAS------F-------E---------TFR---------TV---------F------------------KKQ------GE-------NAP------
ISKYASFAHTTALND-LL-----R------N----E-------Q----T---------RY----------T-------L-------G----D---I---S------YVF--W-----------S---------SNLDD------------PS-IN------------------------------------------E-------------------------------------DP-----A--Y-LS---Y-------------G----------------VK---D-D----DAQEPS-------DTDDEQPKRKGKSKK------
KAPN-----P--------------------T--------Y------KVLEQ-----FKA------I----------------R-G-N------R--G-E-----G-----QH-----W---K---------SQKFQLN----------LVQSIVESVLN--G-T--P---YPMTLQQA-CV------------------------------------------------------------------------------------------------------------------------DGNVDEENP----------------------
LRSLY-------------GIVKSVSSP-------------------SK-----------------------------------------------SSTFAMN---------LVQSIVESVLN--G-T--P---YPMTLQQA-CV----------R-----------Q-------------------TK------------------------------RAVS-------------------------L--------RAAI------LKG--------------------
K--L------------------------------------K---ARIYKHLK---E----L------------------D-------------------------------M--------ALD-------KQ-------ND-------N--------IAYLA-----GRLFAVLE----QIQ------------------Q------A--------------------------SLG---KG------V---------N---------ATIR---DRFYTSAS------TRPNMV-------MGRLIA--LS-N-HHLS---------
K--L------------RKEKPGLAVN----------------------EKL-L----G----EIF------------------AL---------------------------IPAEVPTF-------P-------Q----R----FS-LD-E------QSLFAVGYY----H-----Q-----------------------------------------------------------------------------------------------------------------------------------------
EDTNNSTEETEQ----------------------
'WP_007364828.1'
-----------------------------------------------------------------------------------------------------------------------------------------MI-------LQALYNYYE------------A------M------RKAHR-------------------------------------IATP---GL---EVK-PI-R-----WVIVIRE---D---GS-FV----R---------LKDLK------DPSDK------------------------------------SK------------------G---QD---Y-------L-V-P----------AGLKR--------------S-----G--
K------------CPPAQP----------------------F-------------WDNRKYI------L----G------------------------------------------------------------------------------------------------------------------------------------------------------------------------------------------E-------E------S-------------A-EGVSPCTDRLEDF-KR-VMAE--I--AEH---------Y-------------
PDNAS--FKA---VCRFYDH----------YLGAETPLP---------EEA---------------------Q----T------------------------E-R-I------L---A------------------------------------SDW-------------------M------------TF-----Q-L-----------QGDT-----EL------LATQD--------D--------AY-----------D------------------------L-AAQAY-----------------------------------------------
---------------------------ED-------------N-L-----------------------------------------------------------------------R----------CL------ITG---E---E-K-P---LAAF-H-------A---R-----------IVAP---G------SQ---AG-A---------TIAS------F-------E---------TFR---------TV---------F------------------KKQ------GE-------NAP------
ISKYASFAHTTALNE-LL-----R------N----E-------Q----T---------RY----------T-------L-------G----D---I---S------YVF--W-----------S---------SDLDD------------PS-IN------------------------------------------E-------------------------------------DP-----A--Y-LS---Y-------------G----------------VR---D-D----DTQEQS-------DTDDEQPKRKGKSKK------
KAPN-----P--------------------T--------Y------KVLEQ-----FKT------I----------------R-G-N------R--G-E-----G-----QH-----W---K---------SQKFQLN----------LVQSIVESILN--S-G--P---YPMTLQQA-CI------------------------------------------------------------------------------------------------------------------------DGFVDEENP----------------------
LRSLY-------------G------------IV-----KSVSTP-------------------SK------------------------------------SSTFATN---------LVQSIVESILN--G-N--P---YPMTLQQA-CV----------R-----------I-------------------TQ-----------------------------------RTVS-------------------------L--------RAAI------LKG-------------C-------I-
-----------------R----------------K---ARIYKHLK----E----L-------------------------D---------------------------------M--------ALD-------KQ-------ND-------N--------IAYLA-----GRLFAVLE----QIQ------------------Q------A--------------------------SLG---KG------V---------N---------ATIR---DRFYASAS------TRPNMV-------MGRLIA--LS-N-HHLS---------K-------
L---------------RKEKPGLAVN---------------------EKL-L----G----EIF------------------AL---------------------------IPAEAPTF-------P-------Q----R----FS-LD-E------QSLFAVGYY----H-----Q-------------------------------------------------------------------------------------------------------------------------------------------
-------------TTNPTEETEQ-----------
'WP_025883162.1'
-----------------------------------------------------------------------------------------------------------------------------------------MI-------LQALYNYYE------------A------M------RKAHR-------------------------------------IATP---GL---EVK-PI-R-----WVIVIRE---D---GT-FV----R---------LKDLK------DPSDK------------------------------------SK------------------G---QD---Y-------L-V-P----------AGLKR--------------S-----G--
K------------CPPAQP----------------------F-------------WDNRKYI------L----G------------------------------------------------------------------------------------------------------------------------------------------------------------------------------------------E-------E------S-------------A-EGVSPSTDRLEEF-KR-VMSE--I--AEH---------Y-------------
PDNAS--FKA---VCRFYDH----------YLGAETPLP---------EEA---------------------Q----T------------------------E-R-I------L---A------------------------------------SDW-------------------M------------TF-----Q-L-----------QGDT-----EL------LATQE--------D--------AY-----------D------------------------L-AAQAY-----------------------------------------------
---------------------------ED-------------N-L-----------------------------------------------------------------------R----------CL------ITG---E---E-K-P---LAAF-H-------A---R-----------IVAP---G------SQ---AG-A---------TIAS------F-------E---------DP-A---------L-Y-S---Y------------G--------------------KKQ------GE-------NAP------
ISKYASFAHTTALND-LL-----R------N----E-------Q----T---------RY----------T-------L-------G----D---I---S------YVF--W-----------S---------SDLDD------------PSIN-------------------T---F-------R---------T-----VTF-----EGV----RNDETQEQSDTDDEQPKRRGKSKKKAPS-
P-TQ-----P--------------------T--------Y------KVLEQ-----FKA------I----------------R-G-N------R--G-E-----G-----QG----HWE---D---E---R---Q---F-YLGL---TAG---GRITIKYWQQG-TVSEIFDRVYQHL---VD----MN-II-------------------------------------------------------------------------------------------------TGNVDEENP----------------------
LRSLY-------------G------------IV-----KSVSTP-------------------SK------------------------------------SSTFATN---------LVQSIVESILN--G-N--P---YPMTLQQA-CV----------R-----------I-------------------T-------------------------------------RTVS-------------------------L--------RAAI------LKG-------------C-------I-
-----------------R----------------K---ARIYKHLK----E----L-------------------------D---------------------------------M--------ALD-------KQ-------ND-------N--------IAYLA-----GRLFAVLE----QIQ------------------Q------A--------------------------SLG---KG------V---------N---------ATIR---DRFYASAS------TRPNMV-------MGRLIA--LS-N-HHLS---------K-------EE---
L---------------RKEKPGLAVN---------------------EKL-L----G----EIF------------------AL---------------------------IPSEAPTF-------P-------Q----R----FS-LD-E------QSLFAVGYY----H-----Q-------------------------------------------------------------------------------------------------------------------------------------------
-------------TTNPTEETEQ-----------
'WP_009751916.1__2_'
-----------------------------------------------------------------------------------------------------------------------------------------MI-------LKALYDYYQ--------A---MA----------E--------------KKDE------------------VAQF---GM---DWV-TI-S-----FVIIINE---D---GT-FK------D--------------IE-----------ST-------K------------G------------L-L-P----T---DK--TR----------
--VV----P---Y--------I-L--------------WDKASYV-------L------G-----------------------------------------------S-------DKEGG-----------------------------------------------------------------------------------------------------------------------------------------------------------------------------------------------------------------------------------------------------------------------
IHLFYEK------------EQ--------Y------LV---A-------K-----D------------------------------E-A-L--------Q-D-A--I----D----T-------------KGG---------W-----------------IAF-----R-L----------KS----------D---------------------A-------------------------L-D-------------SF-RK-EIQK--I--HKE-------N-----P----EN--I---G---FRA-----
-------------------------NID-----T-------------------------------------------------------------------C---------CM------ITG--K---RE--S---IAKT-H--K-G---I-K---I------------SR-DN-P-----------PLIS-----F----------------SDPH-------------N-------------H------------SD-E------------N-----------------SFRIA-----
---E-----WG------R----N-------T---------NY----------------K--------I-----G------D----N-------G-------T----T------FVF-W---S--------------------T-LQK----------------DD-LL-----------------------------------------------AYK--------EA-----T--------------T-----G-------LP--SDS---DFDDEE-------KEASPLPNKSN-------EKLD--------P-------
-----------K------------------------R---VLKRA-----LEV-----V-----------------R-G-Y-----Q------G---E-S-----H----D------E--K---------E----R-----F-YMLGL---TPN--K-SRLSVKLWMEG-TVREIVGNTLQHL---ED-------MN-VVS--------------------------------------------------------------------------------------DGLGDECNP---------------------------
--SIFKA-----ILPIPT----NKKESDINY-------------------------------------------------------PP----H-----------IVQGLLESIVK--G-L--P---YPIALQNA-CL--------E-----------------R--I---------------------QH------------------RSRKDAKGRRISPIT-------------------------------------T-----------------RVAL------LKA------C---
------------------------------------------------------K-----HRKNHNIK-----K---------------------------------------------M-----ALD-------LT-----NN----A---------PAYLA-----GRLFFILE------DIQ-------------------------------------------------AIG-----YE--------I-------N---SSIA---DRYFSRAA------SDPKVV------FGELLV---KA-N-YHLR-------K----L---------------------------------
ERSNGRSH---------------------F-----------ARKEI---E------AIL-------E----LL-------------------PQEIISPEDTQGQA----K--QG-LD-D-------ESLFAVGYY---H----Q-------------------------------------------------------------------------------------------------------------RRAENREKKEAENE------------------------
'WP_036850543.1'
-----------------------------------------------------------------------------------------------------------------------------------------MI-------LQALYKYYE---------A---MQ-----------S------E----------E-S-----------------IAPL---GL---EWR-EI-P---------FIVISP---E---GE-FV--D---------FI------DT--------R---------D------DK--E-N-K---A--G-RK---F-------L-V-A----K----NK--GR----------
------SHSI----S---Q---------I-L-----------WDHSGYV-------L------E------------------------------------------------TEGE--------------------------------------------------------NKA-----------------------------------------------------------------------P--------------------------------------------SF-LK-ELQT--L--KKE---------F------P----HN---E-------
FLA----VERFYEK-----------------KQ--------H-------LA----L--------S-------------------------------------------E-----E---------E------NI---------K-T-I---------Q---------------------------QGG-------------------N----------ISF-----L-L-----S-----------T-----E--------PNQS-------------------N------------K---S-----Y-NS-----Y-------E------------
-------------------------ENG-----------S-----------------------------------------------------------------------------------------CL---------VTG--K---HE---A-----IARL-H--D-G---Y-K---L-----------------N-KE--SN-T-----------PLIS------F--------------------------------------------------K-Q-Q---------GF-----------NAP------
ISTKAAKAIAAALSN-LL-----F---TG-----FGKE------------T------F-----T---T-------FVF-W-----------S----------------------IT-EQS---------------AE-LY-------------------------------------------------LYK------------KA-----T--------------------------S---TNTLNS-----DKIQETGSTKRKRGDKNTT-Q-
VPQEPN--------P---------E-----------------K-----------------R-----KVLAA---FKAN-----I-----R---G-S-----E-----G------H-----W---E----------E-----------E-------R-----F-YILGL---TTN--K-SRMCVKLWTEG-TVGEIVGNTLQHL---DD-------MN-IV-------------------------------------------------S-----------------Y-LRTPAK-------EK-GDKE-----
LHKLG---------------------------SITLA-----ITKEGI--------ADKFIS---------------------------------------------------LTEELVESIVK--G-L--P----YPAVVQKL-CL--------------------------N-------R-V--------------------------------------------------------------HKTSGKTK-----------------L---------RVAI-------LKA--------Y------
-------N--------R---------------------K---NRYYKKDIV----M-------------------------------------------S------------E-----L--GLD--------RE----------QT------N--------IGYLA-----GRLFAVLE------YTQ------------------------------------------------------R----------------AL----GE-------I---------STVS---DRFFSKAS------LTPNVV---------FPKAER--LY-M-KHIK-----K--------
-L-------------------------KSKRYYVAKT---------------------------------------LRE-Y-D-------SIC-------------IL-------FSGSETNF-----P-----Q----S----R----FS-LD-E------QSMFVIGFH----H----Q---------------------------------------------------------------------------------------------------------------------------------------
---PTEGVKETFDHNEIMGSDLDI-----------
'WP_039432739.1'
-----------------------------------------------------------------------------------------------------------------------------------------MI-------LQALYQYYQ---------R---MR----------A---A---------------------------GTN------------IAPP---GM---EWK-AI-P----FIIDLTP---E---GE-FI-----Q-------LS-------DT-------R--------D-------K-----E-H-K----K---G-GE---Y---L-V-S-----R---SA--TR-----------S-----G---K-----------
N--------AYQV---A---Q----------V-M-------------WDHLGYV---------L-----G-----------------------------------------------------------------------------------------------------------KEE-------------------K----------------------------------------------------------------------------Y-E-----------YF-VE-KVKL--L--KEL-------Y-------P----HN--L----
-Y--ISA---VNRFYQD----------------------LR-LRE------N-----------------------------------D-------E-------LI-----------K-S-I---------Q------------------------------------KSA-------------------N-----------ISF-----R-I-----P-------------D-------------LGPS-------------------LV---------A---SIQ----D-------I-ET-YM-------G--------ASM-------T------S-------E----
-----------------------------------TDE------SN----------------------------------------------------------------------------S-G------------I-----CL-------VTG---L-RS--E---IARL-H-H-P-S---I--K---L--A-----------G-AQ-AT-A------------SLIN------F-------------------------------------------------K-E-Q---------GF-----------NAP---------
VSVHAAGSIATALND-LL---------S-----KD------------K-----------------------------------------------------------------------------------------N-FTD-------------------DD-LI-----------------------------------------------------------------------------------------------------------------------------IS----GEH--------
DEEDEDNEVSNSDHSDTEPLPKRSRRNR-P-K-ATPA-------P---------------------N---------T------E--------S----Y------KVLQT-----LKA------A--------------C-G-Y-----K------G-----D-----V-----R----R----F-YILGL----APN--S-ARISIKLWAEG-TISDIVGNTLRHQ---YD-------MN-IV--------------------------------------------------------------------------------------------
----------EGNINTEVP---------LRSIYQIV----------RT--------------------------V---SSSDKSDKW-------------------------------------------------------------------------SA-----N---------IAYLA-----GRLFALLE------YTQ------------------------------------------------------SR-----------------------------------------------
------RAAI-----LKA----Y---------------------------------------------------------K-----------KQQE-----Y------------------------I----------------T-----M--ALD------TN-------------QT------N--------KAYLA-----GRLFALLE------NIQ-----------------------AL---GN-------T-----------NSTIR---DSYYCAAS-----
TTPRAV------FGRLDA--LS-K-AHLS------K--L-------------RKEKIGLAID--------------------R-----------ERQ-I--E-----EIY-----------NL---------------GNSPELPSHFT-LD-D-------QSFFAVGYY---H----Q-------------------------------------------------------------------------------------------------------------------
----------------KEKTDEDSDDSSEDASN-------------
'WP_004333880.1'
-----------------------------------------------------------------------------------------------------------------------------------------MI-------LQALYQYYQ---------Q---MH-----------A-------------------------GAD-----------------IAPP---GM---EKK-AI-P----FIIELTP---E---GE-VI-----Q------VT--------ET--------M--------D-------S--------E-N-K---K---KT---F-----L-V-S----R-----SA--TR-------S-----G---K-----------
-----AFQV---A----Q--------V-M---------------WDHLGYV-------L------G-------------------------------------V-----------------KDGTE-----------------------------------------------------------KEE-----------------------------------------------------------------------------------------------------SF-IK-KVDE--L--REL---------Y-------P----EN---P--
H--IAA---VARFYHD------------------------LR-LIE------D-----------------------------------------------K-D-I-------------Q----K--T-----------------TGA----------------------ISF-------R-V------------------D---------LEDD----------------R------LV----------A----SIQ------D-------V-QS-YM---------N-E---------T-------S---E----
-----------------------TDN----------SN------------------------------------------T-S-------------------------------I-----------CL-------VTG---E---KS---T---IARL-H-H-P-S---V--K---L--A-----------G-AQ-AT-A------------SLIS------F--------------------------------------------------K-E-Q---------GF-----------NAP---------
ISGHAAESIATALNN-LL-------A-----KD--------K-------N----T------------------------------------------------------------------------------------------------N-FTN--------------------------------------------------------------------------------------------------------------------------SSYRTATFNGISSSSEEEEENLDL-----------
ENPEEEKSKKSRRKKKNIP------------------------------------------------------------------------------------I-------------------------------------------------------------------------------------------------------------------------------------------------------------------------------------------KGNVDAETP---
----P-------LRSIY--------------------T-------------IAPDRESYRV-----LQT---L----------KAACG------Y-------NGGKVH---------E------D-------N---G-------R-----F-YILGL---APN--A-ARISIKLWAEG-SIPEIVGNTLRHQ---YD-------MN-IV----------------------------------SE-----------------------------------------------------------------
--------------------------QI---VKTVSSSDK-------------R----------------------------LIQSIVESIVE--N-K--P---YPQSLQQA-CL------------------E-----R------I---------------R----------------------------------------------------------------------------------------------------RAAI-------LKA-----Y-----------
-------K--L-------------------------------------------------------------KKQQE-------Y----------M---------SLD--------DN-------QT-------N--------KAYLA-----GRLFALFE----SIQ-------------------------------------------------------------ALG---GN-------L----------NRIT---RTIR---DAYYGTAS-----TTPRTI--------FGRLDA--LS-K-VHLS-
------K---L---------RKEKPGLAVL---------------------EKQ-L--E-----EIY-------------------NL---------------IPGGAPEM-------P---------T----H----FS-LD-D-------QSIFAVGYY---H----Q-------------------------------------------------------------------------------------------------------------------
KEETEDHLEE-------------------------
```

'WP_075564449.1'
[illegible]

'WP_009437516.1'
[illegible]

'RKV79203.1'
[illegible]

'WP_044114298.1'
[illegible]

'WP_021666377.1'
[illegible]

'WP_060936145.1'
[illegible]

'WP_052080636.1'
[illegible]

'WP_044231508.1'
[illegible]

'WP_036844792.1'
[illegible]

'WP_117302852.1__2_'
[illegible]

'WP_089838097.1'
[illegible]

'RMH23931.1'
[illegible]

'WP_008523302.1__3_'
[illegible]

'WP_008519402.1'
[illegible]

'WP_101479734.1'
[illegible]

'PJB25944.1'
[illegible]

'ABF90712.1__2_'
[illegible]

'WP_002633773.1__2_'
[illegible]

'AEI64170.1'
[illegible]

'SDF25903.1'
[illegible]

'AKF84136.1'
[illegible]

[illegible]

'AGC48887.1'

[illegible]

'WP_044241840.1'

[illegible]

'WP_082362866.1'

[illegible]

'WP_121770055.1_2_'

[illegible]

'WP_120622793.1_2_'

[illegible]

'WP_120608670.1_2_'

[illegible]

'WP_121758217.1_2_'

[illegible]

'WP_078015836.1'

[illegible]

'EHQ07661.1'

[illegible]

'WP_081476506.1'

[illegible]

'PKN43912.1'

[illegible]

'SIQ75766.1'

[illegible]

'WP_123481231.1_5_'

[illegible]

'WP_107036830.1_3_'

[illegible]

'WP_123550133.1_2_'

[illegible]

'WP_117724645.1_2_'

[illegible]

'WP_099463339.1'

[illegible]

'WP_007558740.1'

[illegible]

'WP_117463303.1_3_'

[illegible]

'WP_005938665.1'

[illegible]

MSKSAEFAIAAALKK-LI---------D-----YN----------S------H------N-----------KF----------------------T---------V----G--------L----R----T--------[illegible]
---------------------D-----------S-------------K---------L-------------E----NITQI-----FKS-----I---------------------Y-N-G-----N-----K-P------I---QT[illegible]
KA--------------V---------S---LKENIKENC-----------------------------------------------------------------------------PP----S[illegible]
------------------------------------------------------------L----NDNNNNKKL----D------------------------------------------I----MLD-------KD[illegible]
AKDKKGLANF------------------------------------F--------------DDQ-K--K--------EII--------------AL----------------------IQDF-----P------N[illegible]
[illegible]

'WP_007219521.1__2_' ---MI------LKALYDYYY[illegible]
--------A---N----------I-L--------------------WDNCAYV-------L----G---------------------------L[illegible]
SESVE--LRA---LHLFYER----YN-----------K------DQI----------------------L------E------------------------------VLQQDIKW--------------Q--D[illegible]
-------------------------------------------------------------KG---------------------------------------------------------L[illegible]
ISADAEFCYTTALKN-LL---------D-----KD-----------S------R------N-----------KF---------------------N---------I-----G---------S----R----T--------F[illegible]
---------------------------------------------D---------P-------------NARIEQARKV-----FHA-----I----------------------Y-S-G-----S-----------------LN[illegible]
---SA---------------V-------------TLRGKVADV--------------------------------------------------------------------------QP----N[illegible]
R--------------------------------------------------------------------------LSNDINKKIE--------------------------------------V----MID[illegible]
---------------------DKGVQIW--------------F---------------EQI-K--S--------EIV---------D-----KI-----------------------MAEGF-----P------C----H---L[illegible]
[illegible]

'WP_009132543.1' ---MI------LKALYDYYY[illegible]
---------A---N-----------I-L---------------------WDNCAYV-------L----G------------------------L----------------SDVDFSDAGAALDEE[illegible]
SESVE--LRA---SHLFYER----YN-----------K------DQI-------------------------L---------------------E--VLQQD----IKW--------------Q--D[illegible]
[illegible]

'WP_008143459.1'
[illegible]

'WP_106040168.1__2_'
[illegible]

'WP_118086055.1__2_'
[illegible]

'WP_094880579.1__2_'
[illegible]

'WP_094464453.1__2_'
[illegible]

'WP_089545263.1__2_'
[illegible]

'WP_119238152.1__2_'
[illegible]

'WP_009348108.1'
[illegible]

'WP_027450839.1'
[illegible]

'WP_016275320.1'
[illegible]

'WP_041525563.1__2_'
[illegible]

'REC36223.1'
[illegible]

'WP_122378810.1__2_'
[illegible]

'WP_122268054.1__2_'
[illegible]

'WP_057317108.1__5_'
[illegible]

'WP_122379689.1__2_'
[illegible]

'RGZ22660.1'
[illegible]

'WP_131225790.1__2_'
[illegible]

[illegible]

'WP_120027180.1__2_'

[illegible]

'WP_118940839.1__2_'

[illegible]

'WP_117596677.1__3_'

[illegible]

'WP_008782057.1__10_'

[illegible]

'RHA00623.1'

[illegible]

'WP_117709499.1__2_'

[illegible]

'WP_117870486.1__2_'

[illegible]

'WP_118121588.1__2_'

[illegible]

'WP_117892964.1__2_'

[illegible]

'WP_118327575.1__2_'

[illegible]

'WP_016271357.1__15_'

[illegible]

'WP_117543425.1__5_'

[illegible]

'WP_005852927.1__2_'

[illegible]

'WP_038604774.1__4_'

[illegible]

'WP_007843554.1'

[illegible]

'WP_007846891.1__3_'

[illegible]

'WP_117475216.1__2_'

[illegible]

'WP_117682092.1__4_'

[illegible]

'WP_117595864.1__4_'

[illegible]

'WP_117829889.1__3_'

[illegible]

NPGGQVF [illegible]

'CUP48124.1' [illegible]

'WP_118292476.1__2_' [illegible]

'WP_087364739.1' [illegible]

'WP_028859806.1' [illegible]

'WP_027010287.1' [illegible]

'WP_078253862.1' [illegible]

'WP_078256154.1' [illegible]

'WP_005578608.1' [illegible]

'WP_040976599.1' [illegible]

'WP_039405443.1__2_' [illegible]

'WP_003754490.1' [illegible]

'WP_016687509.1' [illegible]

'WP_101755122.1__3_' [illegible]

'WP_080614068.1__2_' [illegible]

'WP_118787163.1' [illegible]

'WP_115119316.1' [illegible]

'WP_114935547.1' [illegible]

'WP_003713924.1' [illegible]

'WP_118810668.1' [illegible]

'WP_096108547.1' [illegible]

'WP_118812146.1'

[illegible]

'WP_118816359.1'

[illegible]

'WP_003708651.1'

[illegible]

'WP_118855499.1'

[illegible]

'WP_118801882.1'

[illegible]

'WP_019273185.1'

[illegible]

'WP_118817185.1'

[illegible]

'WP_009311471.1'

[illegible]

'WP_049250273.1'

[illegible]

'WP_060974792.1'

[illegible]

'WP_109063563.1__2_'

[illegible]

'RKV79750.1'

[illegible]

'WP_054599854.1'

[illegible]

'WP_111750226.1__3_'

[illegible]

'WP_005820512.1'

[illegible]

'WP_028114978.1'

[illegible]

'WP_077246478.1'

[illegible]

'WP_038910772.1'

[illegible]

'WP_013317287.1'

[illegible]

'WP_024105343.1'

[illegible]

'WP_046830387.1'

[illegible]

[illegible]

'WP_040000510.1'

[illegible]

'WP_024108605.1__3_'

[illegible]

'WP_103415741.1__2_'

[illegible]

'WP_024109517.1'

[illegible]

'WP_029729384.1'

[illegible]

'WP_047435397.1'

[illegible]

'WP_054305177.1'

[illegible]

'WP_044387143.1'

[illegible]

'WP_100820768.1'

[illegible]

'WP_018402823.1'

[illegible]

'WP_007348312.1'

[illegible]

'WP_104415632.1'

[illegible]

'WP_114332987.1'

[illegible]

'WP_113860994.1'

[illegible]

'WP_022991922.1'

[illegible]

'WP_022987789.1'

[illegible]

'WP_109838952.1__2_'

[illegible]

'WP_099034913.1'

[illegible]

'WP_015487357.1'

[illegible]

'PCJ84479.1'

[illegible]

[illegible]

'WP_046021481.1'

[illegible]

'RJR48861.1'

[illegible]

'WP_130594125.1__2_'

[illegible]

'RLB70438.1'

[illegible]

'WP_136064930.1'

[illegible]

'PJC76766.1__3_'

[illegible]

'WP_133608424.1'

[illegible]

'WP_012346190.1'

[illegible]

'RJQ14283.1'

[illegible]

'WP_014271804.1__2_'

[illegible]

'WP_023389232.1'

[illegible]

'WP_136368669.1__2_'

[illegible]

'WP_044666150.1'

[illegible]

'WP_095266719.1__2_'

[illegible]

'WP_028563662.1'

[illegible]

'WP_046497843.1'

[illegible]

'WP_010259993.1'

[illegible]

'WP_083052190.1'

[illegible]

'WP_004093890.1'

[illegible]

'WP_082524962.1'

[illegible]

----SSLAL----QG----KLDNVP----P----R----LAGEWMRAILE--G-L--P---YPAGLLNA-AV----V [illegible]
----E----QRRA----D----RTLAIQHAHFKE----N----L----DME----QD----N----VPYRL----GRLFAVLE [illegible]
----SD---GQEA----Y----F----ERL-I--G----EIV----Q--AM-TD----F----P----R----Q---LT-LA-D----QGRFALGYY---H----Q [illegible]
-----ATASTNTTTTPQEA----

'WP_056874296.1' ----MI----LHALADLYE----R----R----S----H-D----P-DPAR-R [illegible]
----V----A----A---N----L-L----WDSAEYA----L----G--A----SA----K----A----ESP [illegible]
V---G--LNA---LLAFLRA----P----HL----I----P----L----Q----A----A----P----HW----A--E-M----L----E----TNA [illegible]
----S----AG----E----L-A----P----CL----ILQ--G---QQ-L-P---AARL-H--T--S---I--K----G--V [illegible]
VSEQAAFAYTTALND-LL----K----TD----S----R----N----RV----Q----V----G----D---A---S----TVF--W----A----D----R----P [illegible]
----D----R----G----R----D----AVRAL----YRT----V----Q-A-G----Q----Q----P---AA----E----G----S----T----R----F--FILGL---APN [illegible]
----TTCAL----Q----GKADNI----PP----R----LAGEWMRAILE--D-L--P---LPAGLLNA-AV----Q [illegible]
-R----Q----MRSQPRPNL----S----TL----P----F-KE-TLD----MS----ND----T----PAYKC----GRLFATFE [illegible]
----TKKRAEY----F----QKL-V--G----EIV----AL---PMRE----L----P----H----R---LT-LP-Q----QAEFALGYY---H----Q [illegible]
----

'WP_026240865.1' ----MI----LQSLHEYYQ----R----K----R----D-S----LP [illegible]
V----E----A---N----L-L----WGNVEYV----L----G--Q----P----DP----K----K----LEDQRKKGKEK----HY [illegible]
ST--P---E--IAA---ALAFLAS----G----HF----T----P----V----L----A----D----P----LW----S--Q-V----C----S----TGA [illegible]
G----K----AA----L-G----I----CL----ITG--N---S--D-V---VERL-H--P--P---I--R----G--V [illegible]
IGKHAAFAYTTALNH-LL----V----S----D----Q----RI----Q----V----G----D---A---S----TVF--W----A----A----S----D [illegible]
[illegible]

'WP_093187967.1' [illegible]

'WP_104936583.1' [illegible]

'PJB69687.1' [illegible]

'WP_048670071.1' [illegible]

'WP_006872858.1' [illegible]

'WP_027268492.1__33_' [illegible]

'WP_027225859.1' [illegible]

'RME60968.1' [illegible]

'WP_110390004.1' [illegible]

'TBR10907.1' [illegible]

'PZN83935.1' [illegible]

'WP_110018240.1' [illegible]

'WP_132541437.1' [illegible]

'WP_026610091.1' [illegible]

'WP_008437583.1' [illegible]

'WP_058361219.1' [illegible]

'WP_045531688.1' [illegible]

'WP_026224322.1' [illegible]

------SNKRRAVN[illegible]

'PZU53034.1'

[illegible]

'PKM03619.1'

[illegible]

'WP_028446649.1'

[illegible]

'WP_009151516.1__2_'

[illegible]

'PKO87781.1'

[illegible]

'WP_009856766.1'

[illegible]

'WP_085315143.1'

[illegible]

'WP_079435287.1'

[illegible]

'AOG22889.1'

[illegible]

'PJC18734.1__2_'

[illegible]

'WP_075585869.1'

[illegible]

'WP_028875185.1'

[illegible]

'WP_135281232.1__2_'

[illegible]

'WP_133716584.1'

[illegible]

'WP_123103864.1__2_'

[illegible]

'WP_094161757.1'

[illegible]

'WP_112487067.1'

[illegible]

'WP_012823706.1'

[illegible]

'WP_025917351.1'

[illegible]

'WP_028324876.1'

[illegible]

'WP_090657021.1'

[illegible]

'WP_046850919.1'

[illegible]

'TAN37772.1'

[illegible]

'WP_010959997.1__2_'

[illegible]

'WP_017364194.1'

[illegible]

'RPI74079.1'

[illegible]

'PJB04975.1__3_'

[illegible]

'PTM65875.1__2_'

[illegible]

'ABI59275.1__2_'

[illegible]

'WP_090284587.1'

[illegible]

'WP_090961320.1'

[illegible]

'WP_090881449.1'

[illegible]

'PJC70655.1__7_'

[illegible]

'WP_020560771.1'

[illegible]

'WP_028487915.1'

[illegible]

'EIJ33767.1'

[illegible]

'WP_134358529.1__2_'

[illegible]

'WP_013032917.1'

[illegible]

'TAN41726.1'

[illegible]

'RMG68731.1'

[illegible]

'RJP19133.1'

[illegible]

[illegible]

'WP_070066838.1'

[illegible]

'PIU50277.1'

[illegible]

'WP_108104050.1'

[illegible]

'PIS36004.1'

[illegible]

'ABB31306.1__2_'

[illegible]

'WP_006584174.1__2_'

[illegible]

'WP_002703148.1__2_'

[illegible]

'WP_021331146.1__3_'

[illegible]

'WP_021687359.1'

[illegible]

'WP_005846119.1__2_'

[illegible]

'WP_027951987.1'

[illegible]

'WP_037547374.1'

[illegible]

'WP_024266934.1'

[illegible]

'PZM83688.1'

[illegible]

'WP_129732454.1'

[illegible]

'WP_122363322.1__2_'

[illegible]

'WP_122376361.1__2_'

[illegible]

'WP_115499562.1__2_'

[illegible]

'WP_122203554.1__3_'

[illegible]

'WP_122122367.1__2_'

[illegible]

[illegible]

'WP_129943851.1'

[illegible]

'WP_075966378.1'

[illegible]

'WP_005641472.1__7_'

[illegible]

'WP_122342651.1__2_'

[illegible]

'WP_046150094.1'

[illegible]

'WP_122328479.1__2_'

[illegible]

'WP_007653591.1'

[illegible]

'PIX95277.1'

[illegible]

'WP_005032281.1'

[illegible]

'ABB24157.1'

[illegible]

'ABL65462.1'

[illegible]

'ABP37314.1'

[illegible]

'WP_076790121.1'

[illegible]

'WP_015405727.1__2_'

[illegible]

'RJQ65413.1'

[illegible]

'PIE62863.1'

[illegible]

'RKX95211.1'

[illegible]

'RKX87100.1'

[illegible]

'TAL31746.1'

[illegible]

'PIR17627.1'

[illegible]

[illegible]

'PSR01375.1'

[illegible]

'WP_112326136.1'

[illegible]

'WP_068122543.1'

[illegible]

'WP_103036656.1'

[illegible]

'PSQ86889.1'

[illegible]

'WP_133530738.1'

[illegible]

'WP_005487699.1'

[illegible]

'WP_114474231.1'

[illegible]

'WP_089719410.1'

[illegible]

'WP_133516153.1'

[illegible]

'WP_110301018.1'

[illegible]

'WP_134058977.1'

[illegible]

'WP_093692089.1'

[illegible]

'OYT70798.1'

[illegible]

'WP_119313945.1'

[illegible]

'WP_027893401.1'

[illegible]

'WP_099749198.1__2_'

[illegible]

'WP_027460536.1'

[illegible]

'WP_034405584.1'

[illegible]

'WP_056295576.1'

[illegible]

[illegible]

'WP_013178147.1'

[illegible]

'WP_097654877.1'

[illegible]

'RIK40967.1'

[illegible]

'WP_099532716.1'

[illegible]

'WP_017306222.1'

[illegible]

'PLS86080.1'

[illegible]

'WP_038681208.1'

[illegible]

'WP_119067369.1'

[illegible]

'WP_015159301.1__2_'

[illegible]

'WP_015164990.1__2_'

[illegible]

'WP_096384500.1'

[illegible]

'WP_106923778.1__2_'

[illegible]

'PZV12891.1'

[illegible]

'PKL77041.1'

[illegible]

'WP_131920242.1'

[illegible]

'PIU54223.1'

[illegible]

'OIQ61466.1'

[illegible]

'YP_429369.1__2_'

[illegible]

'PRR77634.1'

[illegible]

'RJQ46959.1'

[illegible]

'RJX23328.1' [illegible]

[illegible]

'RJX22477.1' [illegible]

[illegible]

'RJQ29546.1' [illegible]

[illegible]

;
end;

### Cas8f (Csy1)

#NEXUS
begin characters;
dimensions nchar=1430;
format datatype=protein missing=? gap=-;
matrix

WP_110816676 [illegible]

WP_019671932 [illegible]

WP_045858073 [illegible]

WP_109838093 [illegible]

WP_086931677 [illegible]

WP_110458481 [illegible]

WP_078924977 [illegible]

WP_085604419 [illegible]

WP_095457616 [illegible]

WP_055063983 [illegible]

WP_086479116 [illegible]

WP_144220941 [illegible]

WP_144128554 [illegible]

WP_144190966 [illegible]

WP_082050258 [illegible]

WP_138598477 [illegible]

WP_063379825 [illegible]

WP_069949105 [illegible]

WP_077670177 [illegible]

WP_088136878 [illegible]

WP_108315820 [illegible]

WP_140932885 [illegible]

WP_082379029 [illegible]

WP_082866156 [illegible]

WP_057625031 [illegible]

WP_077451825 [illegible]

WP_020443856 [illegible]

WP_090332120 [illegible]

WP_066976324 [illegible]

WP_080521540 [illegible]

WP_067981469 [illegible]

WP_067013514 [illegible]

[illegible]

WP_108603332
[illegible]

WP_102565865
[illegible]

WP_026879509
[illegible]

WP_004728800
[illegible]

WP_131669124
[illegible]

WP_082625690
[illegible]

WP_074905107
[illegible]

WP_107784400
[illegible]

WP_124297548
[illegible]

WP_099540775
[illegible]

WP_124296404
[illegible]

WP_054393548
[illegible]

WP_109201528
[illegible]

WP_115020443
[illegible]

WP_018025761
[illegible]

WP_081685295
[illegible]

WP_065588421
[illegible]

WP_065574526
[illegible]

WP_065561647
[illegible]

WP_086357533
[illegible]

WP_086365054
[illegible]

WP_065614518
[illegible]

WP_065618155
[illegible]

WP_110432830
[illegible]

WP_086360868
[illegible]

WP_086319913
[illegible]

WP_065734296
[illegible]

WP_065650777
[illegible]

WP_065578580
[illegible]

WP_110447304
[illegible]

WP_144189819
[illegible]

WP_086363020
[illegible]

WP_110453129
[illegible]

WP_086354136
[illegible]

WP_086362412
[illegible]

WP_110422818
[illegible]

WP_082209444
[illegible]

WP_065633479
[illegible]

WP_065556818
[illegible]

WP_065579850
[illegible]

WP_091121577
[illegible]

WP_065625321
[illegible]

[illegible]

WP_065635959 [illegible]

WP_091347974 [illegible]

WP_085163699 [illegible]

WP_065637927 [illegible]

WP_065581505 [illegible]

WP_075508109 [illegible]

WP_065586743 [illegible]

WP_065605838 [illegible]

WP_085247578 [illegible]

WP_065609892 [illegible]

WP_065652667 [illegible]

WP_049215679 [illegible]

WP_005756147 [illegible]

WP_010201103 [illegible]

WP_011927917 [illegible]

WP_135325949 [illegible]

WP_138018494 [illegible]

WP_095356844 [illegible]

WP_099808535 [illegible]

WP_072281255 [illegible]

WP_101963921 [illegible]

WP_101746500 [illegible]

WP_091356616 [illegible]

WP_008247105 [illegible]

WP_117905973 [illegible]

WP_049239681 [illegible]

WP_120764383 [illegible]

WP_049149702 [illegible]

WP_013107620 [illegible]

WP_120672612 [illegible]

WP_003672780 [illegible]

WP_120656379 [illegible]

WP_038518821 [illegible]

WP_003672252 [illegible]

WP_126739457 [illegible]

WP_120655696 [illegible]

WP_064653485 [illegible]

WP_064651698 [illegible]

WP_003667347 [illegible]

WP_003664170 [illegible]

WP_117982840 [illegible]

```
NPSTHELNKQLLWHI-------NA-----------DTACQLNELD---YINIIPLYPSVLTHGVYLK-IN----Q--L---R--Y-S [illegible]
T----ID---VR--DK----RK------DAIDRLLHTIFGIAHVLQ--N-------K-P-----A---G----WS------K-----T-----------YA-------LPL---AQKIW [illegible]
------------
WP_077655961 ----------------------------------------------------M--TTN-------DLAK [illegible]
-----------FK---KSVW-----LD---LA-----ANRW------TQ----WMT-F--------------GTHISKGIHLS---A-------------R---G-D------NV [illegible]
NPSTHELNKQLLWHI-------NA-----------DTACQLNELD---YINIIPLYPSVLTHGVYLK-IN----Q--L---R--Y-S [illegible]
T----ID---VR--DK----RK------DAIDRLLHTIFGIAHVLQ--N-----K---P-----A---G----WS------K-----T-----------YA-------LPL---AQKIW [illegible]
------------
WP_003670775 ----------------------------------------------------M--TTN-------DLAK [illegible]
-----------FK---KSVW-----LD---LA-----ANRW------TQ----WMT-F--------------GTHISKGIHLS---A-------------R---G-D------NV [illegible]
NPSTHELNKQLLWHI-------NA-----------DTACQLNELD---YINIIPLYPSVLTHGVYLK-IN----Q--L---R--Y-S [illegible]
T----ID---VR--DK----RK------DAIDRLLHTIFGIAHVLQ--N-----K---P-----A---G----WS------K-----T-----------YA-------LPL---AQKIW [illegible]
-------------
WP_120661174 ----------------------------------------------------M--TTN-------DLAK [illegible]
-----------FK---KSVW-----LD---LA-----ANRW------TQ----WMT-F--------------GTHISKGIHLS---A-------------R---G-D------NV [illegible]
NPSTHELNKQLLWHI-------NA-----------DTACQLNELD---YINIIPLYPSVLTHGVYLK-IN----Q--L---R--Y-S [illegible]
T----ID---VR--DK----RK------DAIDRLLHTIFGIAHVLQ--N-----K [illegible]
-------------
WP_003666003 [illegible]
------------FK---KSVW-----LD---LA-----ANRW------TQ----WMT-F [illegible]
NPSTHELNKQLLWHI-------NA-----------DTACQLNELD---YINI [illegible]
T----ID---VR--DK----RK------DAIDRLLHTIFGIAHVLQ--N------K [illegible]
-------------
WP_078253660 [illegible]
------------FE---RTHW-----LD---LA-----VNSM------AK----QLK-F [illegible]
NPTTHERNKQMLWVT-------ND-----------DTANHLDELQ---YIN [illegible]
N----VD---VR--DK----RK------DAIDRMLHTIFAIAHTLQ--D-----K [illegible]
------------------
WP_103034137 [illegible]
--------------FE---RTHW-----LD---VA-----VNSM------AK----QLK-F [illegible]
NPTTHERNKQMLWTV-------ND-----------NTANHLDELQ---YIN [illegible]
N----VD---VR--DK----RK------DAIDRMLHTIFAIAHTLQ--E-----K [illegible]
------------------
WP_078318338 [illegible]
------------FE---CTHW-----LD---VA-----VNSM------AK----QLK-F [illegible]
NPTTHERNKQMLWAV-------ND-----------NTANHLDELQ---YIN [illegible]
N----VD---VR--DK----RK------DAIDRMLHTIFAIAHTLQ--E-----K [illegible]
-----------------
WP_114800711 [illegible]
----------FE---KIAW-----LD---LA-----AGTM------AK----QLK-F [illegible]
NPATHERNKQMLWLT-------ND-----------NTVNHLDELQ---YIN [illegible]
I----VT---VR--DK----RK------DAIDRMLHTIFAIAHQLQ--N-----K [illegible]
----------------
WP_064618279 [illegible]
------------FE---RVHW-----LD---LA-----VNSM------AK----QLK-F [illegible]
NPATHERNKQMLWVT-------ND-----------NTVNHLDELQ---YIN [illegible]
T----IN---VR--DK----RK------DAIDRLLHTIFGIAHALQ--N-----K [illegible]
---------------
WP_042510140 [illegible]
------------FE---RVHW-----LD---LA-----VNSM------AK----QLK-F [illegible]
NPATHERNKQMLWVT-------ND-----------NTVNHLDELQ---YIN [illegible]
T----ID---VR--DK----RK------DAIDRLLHTIFGIAHALQ--N-----K [illegible]
---------------
WP_064610221 [illegible]
------------FE---CVHW-----LD---LA-----VNSM------AK----QLK-F [illegible]
NPATHERNKQMLWVT-------ND-----------NTVNHLDELQ---YIN [illegible]
T----ID---VR--DK----RK------DAIDRLLHTIFGIAHALQ--N-----R [illegible]
---------------
WP_064644643 [illegible]
------------FE---RVHW-----LD---LA-----VNSM------AK----QLK-F [illegible]
NPATHERNKQMFWVT-------ND-----------NTVNHLDELQ---YIN [illegible]
T----ID---VR--DK----RK------DAIDRLLHTIFGIAHALQ--N-----K [illegible]
---------------
WP_065249208 [illegible]
------------FE---RVHW-----LD---LA-----VNSM------AK----QLK-F [illegible]
NPATHERNKQMLWVT-------ND-----------NTVNHLDELQ---YIN [illegible]
T----ID---VR--DK----RK------DAIDRLLHTIFGIAHVLQ--N-----K [illegible]
--------------
WP_063453874 [illegible]
------------FE---RVHW-----LD---LA-----VNSM------AK----QLK-F [illegible]
NPATHERNKQMLWVT-------ND-----------NTVNHLDELQ---YIN [illegible]
T----ID---VR--DK----RK------DAIDRLLHTIFGIAHVLQ--N-------K [illegible]
--------------
WP_077451977 [illegible]
----------------YS---LEVW-----MD---NA-----ANRM------TK----QIG [illegible]
QPITSEYNKQTLWPH---------------------------ANKQ---YTCIIPLY [illegible]
VR--DL----RK------QAMDEVLTQIFDYAHYMQ--N-----E-L-P [illegible]
---------
WP_092615999 [illegible]
--------------YQ---RDAW-----LD---DA-----VNRM------AK----QLN-F [illegible]
QPSSHERNKQTLWAM-------NP-----------YDCSNIDDLK---YIA [illegible]
T----VD---VR--DT----RK------EAIDNVLHILFALVHSIR--T-----T-L [illegible]
------------
WP_090921212 [illegible]
----------------------------YS---SAIW-----LE---NA-----ANKM------AK [illegible]
-PVTHERNKQILWAT-------NS-----------NVAESIEDLS---YHNII [illegible]
-VD---IR--DA----RK------FAIDEILHILFSISTYIK--N-----T-Y-P [illegible]
---------
WP_082894355 [illegible]
--------------------YS---VANW-----IP---DA-----ATRM------AK----QL [illegible]
PVTDERNKQLLWVT-------NA-----------YQGEDIDELN---YINII [illegible]
VD---IR--DA----RK------EAMDEILKRIFEFANYMR--N-----D-L-P [illegible]
------------
WP_055123769 [illegible]
-------------YQ---LDNW-----MD---NA-----ANKM------AN----QIS-F [illegible]
PCSSEYNKQVLWPT---------------------------NHKTDAYICTIPLY [illegible]
VR--DA----RK------SAIDGVLYQIFAAAEDIR--T-----T-M-P-----A [illegible]
------
WP_102091015 [illegible]
--------------YS---VNNW-----IA---DA-----ATRM------AK----QLN-F [illegible]
PVTHERNKQTLWVI-------NA-----------YEETDIEQLN---YINIV [illegible]
VD---VR--NA----RK------RAIDEVLYQIFAAAEDIR--T-----T-L-H [illegible]
-----------
WP_060491555 [illegible]
-------------YS---VATW-----IP---DA-----ATRM------AT----QLN-F [illegible]
PITHERNKQTLWVT-------NA-----------YESTDIEALD---YINVIP [illegible]
VD---VR--DA----RK------LAIDEVLYQIFAAAEDIR--S-----S-M-P [illegible]
--------------
WP_102093766 [illegible]
-------------YS---IANW-----IA---DA-----ATRM------AK----QLN-F [illegible]
PVTHERNKQTLWVT-------NA-----------YEGADIEELD---YINIV [illegible]
VD---VR--DA----RK------LAIDKLLHKLFKAAEDIR--T-----D-M-P [illegible]
------------
WP_101211226 [illegible]
-------------YS---IANW-----IA---DA-----ATRM------AK----QLN-F [illegible]
PVTHERNKQTLWVT-------NA-----------YEGADIEELD---YINIV [illegible]
ID---VR--DA----RK------SAIDEVLYQIFAAAEDIR--M-----T-M-P [illegible]
------------
WP_126756981 [illegible]
---------------YQ---LETW-----MA---DA-----ANRM------AN----QLH [illegible]
KPATHERNKQLLWPI---------------------------AEHD---YNVIVPLY [illegible]
VN---IR--KA----RR------AILDDLLFQVFSIASSIQ--Q-----Q-Y-L [illegible]
------------
WP_034618103 [illegible]
----------------YS---LTVW-----MK---NA-----AERM------AK----QLQ [illegible]
EPTTHERNKQLLFPT---------------------------ATGD---YHVIVPLY [illegible]
IR--DA----RK------AILDDILRHLCTAAREIQ--T-----A-R-T-----A [illegible]
WP_132966259 [illegible]
----------------YS---FVDW-----LE---DA-----AERM------AK----QLK [illegible]
EQVTHERNKQLLFPL-------NH-------------------HD---YRCLVPL [illegible]
VA---IR--DA----RK------ACLDDILFRLIGAVSEIQ--Q-----Q-K-E [illegible]
------------
WP_072577007 [illegible]
-------------YQ---ADNW-----LK---E------AERM------AR----QLN-F [illegible]
QPSTHERNKQILWPI---------------------------DADNDCYHTLVP [illegible]
N----KA---LR--DS----RD------AVLDTMIDTTFLLALALQ--------A [illegible]
-----------
WP_078274255 [illegible]
-----------------YQ---KQTW-----LS---D------AKKM------AT----QL [illegible]
VPTTHERNKQLFWAL---------------------------PDDE---YHLIIPLF [illegible]
ID---VR--TA----RD------DILQNIILSVLQVAHAIR--Q-----K-E-A [illegible]
-------------------------
WP_051226021 [illegible]
------------------YQ---KQTW-----LS---D------AKKM------AT----QL [illegible]
VPATHERNKQLFWAL---------------------------PDDE---YHLIIPLF [illegible]
ID---VR--TV----RD------DILQNIILSVLQVAHAIR--Q-----K-E-A [illegible]
-------------------------
WP_115007033 [illegible]
------------------YQ---KQTW-----LS---DA-----KKMA-------T----QL [illegible]
VPATHERNKQLFWAL---------------------------PDDE---YHLIIPLF [illegible]
ID---VR--TA----RD------DILQNIILSVLQVAHAIR--Q-----K-E-A [illegible]
-------------------------
WP_009174232 [illegible]
--------------YG---LDAW-----LE---QD-----AVRF------AA----QLK-F [illegible]
TPAAHERNKQVLWPL-------GG---------------DAAAADR---YITLI [illegible]
--K----KD---TR--LL----RE------QALEQMVRQIVVVSEHLR--E [illegible]
--------------
WP_039089377 [illegible]
YQ---LDTW-----ME---KA-----ANQM------SK----QLS-F [illegible]
KPVTYERNKQLLWPN---------------------NRQAISEDN---YTCLI [illegible]
--ER--DK----RD------AAIENIIEVIFSIAQIYQ----------QKP-----A [illegible]
WP_039166343 [illegible]
YQ---LDTW-----ME---KA-----ANQM------SK----QLS-F [illegible]
KPVTYERNKQLLWPN---------------------NRQAISEDN---YTCLI [illegible]
--ER--DK----RD------AAIENIIEVIFSIAQIYQ----------QKP-----A [illegible]
WP_021461779 [illegible]
YQ---LDTW-----ME---KA-----ANQM------SK----QLS-F [illegible]
KPVTYERNKQLLWPN---------------------NRQAISEDN---YTCLI [illegible]
--ER--DK----RD------VAIENIIEVIFSIAQIYQ----------QKP-----A [illegible]
WP_039153731 [illegible]
YH---LDTW-----ME---KA-----ANQM------SK----QLS-F [illegible]
KPVTYERNKQLLWPN---------------------NRQAISEDN---YTCLI [illegible]
--ER--DK----RD------AAIENIIEVIFSIAQIYQ----------QKP-----A [illegible]
WP_018347332 [illegible]
YH---LDTW-----ME---KA-----ANQM------SK----QLS-F [illegible]
KPVTYERNKQLLWPN---------------------NRQAISEDN---YTCLI [illegible]
--ER--DK----RD------AAIENIIEVIFSIAQIYQ----------QKP-----A [illegible]
WP_039095080 [illegible]
YH---LDTW-----ME---KA-----ANQM------SK----QLS-F [illegible]
KPVTYERNKQLLWPN---------------------NRQAISEDN---YTCLI [illegible]
--ER--DK----RD------AAIENIIEVIFSIAQIYQ----------QKP-----A [illegible]
WP_013747107 [illegible]
YQ---LDTW-----ME---KA-----ANQM------SK----QLS-F [illegible]
KPVTYERNKQLLWPN---------------------NRQAISEDN---YTCLI [illegible]
--ER--DK----RD------AAIENIIEVIFSIAQIYQ----------QKP-----A [illegible]
WP_039142397 [illegible]
YQ---LDTW-----ME---KA-----ANQM------SK----QLS-F [illegible]
KPVTYERNKQLLWPN---------------------NRQAISEDN---YTCLI [illegible]
-ER--DK----RD------AAIENIIEVIFSIAQIYQ----------QKP-----A [illegible]
WP_039092599 [illegible]
YQ---LDTW-----ME---KA-----ANQM------SK----QLS-F [illegible]
KPVTYERNKQLLWPN---------------------NRQAISEDN---YTCLI [illegible]
--ER--DK----RD------AAIENIIEVIFSIAQIYQ----------QKP-----A [illegible]
WP_039160698 [illegible]
YQ---LDTW-----ME---KA-----ANQM------SK----QLS-F [illegible]
KPVTYERNKQLLWPN---------------------NRQAISEDN---YTCLI [illegible]
--ER--DK----RD------AAIENIIEVIFSIAQIYQ----------QKP-----A [illegible]
WP_094872970 [illegible]
YQ---LDTW-----ME---KA-----ANQM------SK----QLS-F [illegible]
KPVTYERNKQLLWPN---------------------NRQAISEDN---YTCLI [illegible]
--ER--DK----RD------AAIENIIEVIFSIAQIYQ----------QKP-----A [illegible]
```

WP_082987476
[illegible]

WP_066105647
[illegible]

WP_065233741
[illegible]

WP_065238557
[illegible]

WP_065238041
[illegible]

WP_143533186
[illegible]

WP_077435662
[illegible]

WP_061888233
[illegible]

WP_061887476
[illegible]

WP_053330059
[illegible]

WP_005540027
[illegible]

WP_109432129
[illegible]

WP_111305372
[illegible]

WP_111294995
[illegible]

WP_039708942
[illegible]

WP_126374978
[illegible]

WP_115587180
[illegible]

WP_115590523
[illegible]

WP_005600277
[illegible]

WP_039196338
[illegible]

WP_009875009
[illegible]

WP_005603468
[illegible]

WP_005611354
[illegible]

WP_005596018
[illegible]

WP_005616691
[illegible]

WP_005606943
[illegible]

WP_132024713
[illegible]

WP_010906577
[illegible]

WP_096743827
[illegible]

WP_115323175
[illegible]

WP_101776180
[illegible]

WP_005764761
[illegible]

WP_016534561
[illegible]

WP_014390883
[illegible]

WP_143931642
[illegible]

WP_078801852
[illegible]

WP_016504385
[illegible]

WP_143928166
[illegible]

WP_061405932
[illegible]

WP_135953059
[illegible]

WP_064973075
[illegible]

WP_046333889
[illegible]

WP_064969230
[illegible]

WP_115173242
[illegible]

WP_059246147
[illegible]

WP_005753650
[illegible]

WP_104884314
[illegible]

WP_115299549
[illegible]

WP_083005318
[illegible]

WP_108511407
[illegible]

WP_104227731
[illegible]

WP_126417400
[illegible]

WP_115098541
[illegible]

WP_134794479
[illegible]

WP_046339715
[illegible]

WP_126373729
[illegible]

WP_017350190
[illegible]

WP_005752986
[illegible]

WP_141442889
[illegible]

WP_022850861
[illegible]

WP_128466782
[illegible]

WP_132873132
[illegible]

WP_015337045
[illegible]

WP_051294727
[illegible]

WP_084167025
[illegible]

WP_127526457
[illegible]

WP_074821064
[illegible]

WP_047780694
[illegible]

WP_126467409
[illegible]

WP_061506302
[illegible]

WP_124297695
[illegible]

WP_099541353
[illegible]

WP_070325445
[illegible]

WP_136540530
[illegible]

WP_136544417
[illegible]

WP_127032758
[illegible]

WP_080155886
[illegible]

WP_107275484
[illegible]

WP_052675283
[illegible]

WP_107194538
[illegible]

WP_087853257
[illegible]

WP_065177210
[illegible]

WP_107265080
[illegible]

WP_107212779
[illegible]

WP_101769137
[illegible]

WP_107239008
[illegible]

WP_107250165
[illegible]

WP_107274991
[illegible]

WP_107197890
[illegible]

WP_107190827
[illegible]

WP_107237469
[illegible]

WP_107229117
[illegible]

WP_065194078
[illegible]

WP_107293814
[illegible]

WP_017004729
[illegible]

WP_087018363
[illegible]

WP_053394904
[illegible]

WP_107347640
[illegible]

WP_107292051
[illegible]

WP_106338479
[illegible]

WP_069530484
[illegible]

WP_138240850
[illegible]

WP_086368026
[illegible]

WP_115060687
[illegible]

WP_138242589
[illegible]

WP_087482898
[illegible]

WP_047880224
[illegible]

WP_108168061
[illegible]

WP_094955976
[illegible]

WP_038224872
[illegible]

WP_114417353
[illegible]

WP_013505553
[illegible]

WP_084624022
[illegible]

WP_098420804
[illegible]

WP_098422737
[illegible]

WP_028685357
[illegible]

WP_119393254
[illegible]

WP_044448285
[illegible]

WP_077666218
[illegible]

WP_074212616
[illegible]

WP_077674770
[illegible]

WP_077679703
[illegible]

WP_077774628
[illegible]

WP_077639453
[illegible]

WP_116645049
[illegible]

WP_077660692
[illegible]

WP_077523536
[illegible]

WP_077607839
[illegible]

WP_077648883
[illegible]

FA---TR--YR----RDH-----DFILPLTDLVFDYAARVQ--Q-----Q-V-P-----A---G----WS------K-----G---A---D--DKACR--------LKR---EHALWL-----D-------------VYN----P-------------DAQFQR--ERE--AQD-[illegible]
--------------

WP_025740058 ------------------------------------------------------------ME------------HSLY--DALL--A-YIE-QR-----K----QAK[illegible]
------AK------FE---PIHW-----VT---DA-----ATRA-------K----QVS-L----------------ATHCPKFTHGD---A---------------K---A-T------SM----L----V----T-------EV-------DKS--E--SR--Y------LTS--Y--SL------T[illegible]
LSAHNLTKQTYFPV------------------------NEGE---YLLLCPLYSSSLAHTLYNK-VR----A--T---R--F-G---------------EK-VKAIR-E-----AK------------N--KQYYH--PE-R-WEI-----FPH-TVKQAFGG-T[illegible]
FA---TR--YR----RDH-----DFILPLTDLVFDYAARVQ--Q-----Q-V-P-----A---G----WS------Q-----R---A---ENKDEGCH--------LKR---EHALWL-----D-------------VHN----P-------------DDQFQR--ERD--AQ[illegible]
-------------------

WP_077577768 ------------------------------------------------------------ME------------HRLY--DALL--A-YIE-QR-----K----QAK[illegible]
------AK------FE---PIHW-----IT---DA-----ATRA-------K----QIS-L----------------ATHCPKFTHGD---A---------------K---A-T------SM----L----V----T-------EV-------DKS--E--SR--Y------LTS--Y--SL------T[illegible]
LSAHNLTKQTYFPV------------------------NEGE---YLLLCPLYSSSLAHALYNK-VR----A--T---R--F-G---------------EK-VKAIR-E-----AK------------N--KQYYH--PE-R-WEI-----FPH-TVKQAFGG-T[illegible]
FA---TR--YR----RDH-----DFILPLTDLVFDYAARVQ--Q-----Q-V-P-----A---G----WS------Q-----G---T---D--DQACR--------LKR---EHAIWL-----D-------------VHN----P-------------DDQFQR--ERD--AQA-[illegible]
----------------

WP_069362484 ------------------------------------------------------------ME------------HSLN--EKLL--D-DIE-QN-----K----QVK[illegible]
------AK------YE---PIRW-----IT---DA-----ARRA-------K----QIS-L----------------ATHCPKFTHGD---A---------------K---A-T------SM----L----V----T-------EV-------DKS--E--SS--Y------LTS--Y--SL------T[illegible]
LSAHNLIKQTYFPV------------------------NEGE---YHLLCPLYSSSLAHTLYAK-VR----A--T---R--F-G---------------EK-VKAIR-E-----AK------------N--KQHYH--PE-S-LKA-----FPR-TVKQTFGG-S[illegible]
FA---TR--YR----RDH-----DFILPLTDVVFDYAARVQ--Q-----Q-V-S-----A---G----WS------K-----G---A---D--DKACR--------LKR---EHALWL-----D-------------VYN----P-------------DAQFQR--ERE--AQD-[illegible]
----------------

WP_077647631 ------------------------------------------------------------ME------------HSLY--DALL--E-YIE-QR-----K----EPK[illegible]
------AK------YE---PIHW-----VT---DA-----AKRA-------K----KIS-L----------------ATHCPKFVHSD---A---------------R---A-S------SM----L----V----T-------EV-------DKS--E--SR--Y------LTS--Y--SL------T[illegible]
[illegible]

WP_069586490 [illegible]

WP_077651638 [illegible]

WP_077487135 [illegible]

WP_075497108 [illegible]

WP_045110093 [illegible]

WP_045104157 [illegible]

WP_130047000 [illegible]

WP_130086579 [illegible]

WP_065595287 [illegible]

WP_069594522 [illegible]

WP_074373080 [illegible]

WP_038184014 [illegible]

WP_077338624 [illegible]

WP_007986846 [illegible]

WP_008845637 [illegible]

WP_008135459 [illegible]

WP_045978134 [illegible]

WP_130152373 [illegible]

WP_037430897 [illegible]

WP_113609795 [illegible]

WP_039422062 [illegible]

WP_012197494 [illegible]

WP_012089885 [illegible]

WP_033102944 [illegible]

WP_037396300 [illegible]

WP_072102476 [illegible]

WP_050098028 [illegible]

WP_050296085 [illegible]

WP_145480522 [illegible]

WP_145526012 [illegible]

WP_049617263 [illegible]

WP_072100569 [illegible]

WP_087816332 [illegible]

WP_032816279 [illegible]

WP_050286499 [illegible]

WP_145604028 [illegible]

WP_091343855 [illegible]

WP_008898789 [illegible]

WP_008608781 [illegible]

WP_134053647 [illegible]

WP_012016803 [illegible]

WP_141176897 [illegible]

WP_040153822 [illegible]

WP_101078659 [illegible]

WP_120131061 [illegible]

WP_120162725 [illegible]

WP_129952741 [illegible]

WP_047612529 [illegible]

WP_047609317 [illegible]

WP_015696920 [illegible]

WP_131638820 [illegible]

WP_112151831 [illegible]

WP_132963909 [illegible]

WP_112288525 [illegible]

WP_129638682 [illegible]

WP_092876333 [illegible]

WP_131488424 [illegible]

WP_061708565 [illegible]

WP_126815656 [illegible]

WP_112781227 [illegible]

WP_121527679 [illegible]

WP_133630696 [illegible]

WP_125293928 [illegible]

WP_032699662 [illegible]

WP_132730759 [illegible]

WP_126509586 [illegible]

WP_128320710 [illegible]

WP_128319033 [illegible]

WP_108415996 [illegible]

WP_032638986 [illegible]

WP_101737964 [illegible]

WP_112012290 [illegible]

WP_102891155 [illegible]

WP_006174403 [illegible]

WP_063144436 [illegible]

WP_131047977 [illegible]

WP_100683639 [illegible]

[illegible]

WP_141964469 [illegible]

WP_129996322 [illegible]

WP_045284503 [illegible]

WP_048027169 [illegible]

WP_069719518 [illegible]

WP_049136597 [illegible]

WP_063943599 [illegible]

WP_059361211 [illegible]

WP_047071858 [illegible]

WP_100122389 [illegible]

WP_059362015 [illegible]

WP_046092263 [illegible]

WP_045334123 [illegible]

WP_045352058 [illegible]

WP_039025185 [illegible]

WP_045406789 [illegible]

WP_065101864 [illegible]

WP_069598063 [illegible]

WP_023327012 [illegible]

WP_063435812 [illegible]

WP_045261781 [illegible]

WP_088204934 [illegible]

WP_069732554 [illegible]

WP_126817679 [illegible]

WP_080297786 [illegible]

WP_088555711 [illegible]

WP_040022890 [illegible]

WP_062856776 [illegible]

WP_100168432 [illegible]

WP_127869016 [illegible]

WP_058679280 [illegible]

WP_095455593 [illegible]

WP_108453384 [illegible]

WP_038981409 [illegible]

WP_063449185 [illegible]

WP_112792503 [illegible]

WP_131637889 [illegible]

WP_047354353 [illegible]

WP_126290833 [illegible]

WP_032647879 [illegible]

WP_063842579 [illegible]

PASHKLAKQGYFPV [illegible]
-FR---IR--EA----RD------GYINMLIDLLFEIASELQ [illegible]
-----------
WP_048983759 [illegible]
WP_047052339 [illegible]
WP_063959227 [illegible]
WP_047737750 [illegible]
WP_045142247 [illegible]
WP_063945028 [illegible]
WP_047637455 [illegible]
WP_047056387 [illegible]
WP_032639217 [illegible]
WP_045327286 [illegible]
WP_048994023 [illegible]
WP_087529461 [illegible]
WP_047343458 [illegible]
WP_023338204 [illegible]
WP_023337012 [illegible]
WP_048887701 [illegible]
WP_131065732 [illegible]
WP_105651152 [illegible]
WP_105745290 [illegible]
WP_105665893 [illegible]
WP_038886655 [illegible]
WP_038874595 [illegible]
WP_105663953 [illegible]
WP_038864772 [illegible]
WP_105606924 [illegible]
WP_104676799 [illegible]
WP_105576133 [illegible]
WP_043056261 [illegible]
WP_032981412 [illegible]
WP_105945035 [illegible]
WP_088277132 [illegible]
WP_105546862 [illegible]
WP_097640972 [illegible]
WP_104674236 [illegible]
WP_097569788 [illegible]
WP_105929122 [illegible]
WP_105571218 [illegible]
WP_133460144 [illegible]
WP_048282018 [illegible]
WP_048278514 [illegible]

WP_048272867
[illegible]

WP_048286806
[illegible]

WP_108474610
[illegible]

WP_113857382
[illegible]

WP_095573800
[illegible]

WP_105285199
[illegible]

WP_085460572
[illegible]

WP_000415800
[illegible]

WP_108188876
[illegible]

WP_097438612
[illegible]

WP_137517176
[illegible]

WP_001706989
[illegible]

WP_097512310
[illegible]

WP_134700638
[illegible]

WP_097726146
[illegible]

WP_001704454
[illegible]

WP_001297315
[illegible]

WP_000415803
[illegible]

WP_001714017
[illegible]

WP_000415804
[illegible]

WP_134737656
[illegible]

WP_115738859
[illegible]

WP_130261261
[illegible]

WP_107180013
[illegible]

WP_105279790
[illegible]

WP_115439792
[illegible]

WP_105266422
[illegible]

WP_061090905
[illegible]

WP_000415805
[illegible]

WP_016248534
[illegible]

WP_126511159
[illegible]

WP_001516089
[illegible]

WP_141109089
[illegible]

WP_130258203
[illegible]

WP_135492109
[illegible]

WP_130205261
[illegible]

WP_016158984
[illegible]

WP_094184957
[illegible]

WP_137326176
[illegible]

WP_059223746
[illegible]

WP_059273390
[illegible]

WP_059256517
[illegible]

WP_059257704
[illegible]

WP_064530183
[illegible]

WP_115775361
[illegible]

WP_085448907
[illegible]

WP_038355175
[illegible]

WP_016261857
[illegible]

WP_137501653
[illegible]

WP_105494077
[illegible]

WP_000415801
[illegible]

WP_141419755
[illegible]

WP_021531159
[illegible]

WP_033559618
[illegible]

WP_001552676
[illegible]

WP_052896461
[illegible]

WP_101943191
[illegible]

WP_096837812
[illegible]

WP_115735852
[illegible]

WP_137534347
[illegible]

WP_000415807
[illegible]

WP_021519928
[illegible]

WP_119110739
[illegible]

WP_021511678
[illegible]

WP_113234477
[illegible]

WP_089647070
[illegible]

WP_137504956
[illegible]

WP_136672370
[illegible]

WP_000415806
[illegible]

WP_089574392
[illegible]

WP_042020509
[illegible]

WP_097725830
[illegible]

WP_000415799
[illegible]

WP_001519631
[illegible]

WP_001546553
[illegible]

WP_112040252
[illegible]

WP_000415802
[illegible]

WP_097511793
[illegible]

WP_069371351
[illegible]

WP_016232820
[illegible]

WP_135565502
[illegible]

WP_071343263
[illegible]

```
PSSHKLAKQIYFPV--------------------------ADG---YHLLCPLFSSSLAQAMYEK-LT----A--G---R--F-S----------------EE-SKAIR-D-----AR-------------K--AGKWH--SQ-P-DIR-----FPN-LAEMHFGG-T----KPQ---------NISLLN--SVRG----------GRVWLLPSM-P----PAWG--P---LD-RAPQKM----RSIF----AL-RGDFN--R-----AA--SGIIARMTYLL------K----------V------D--------------IN---------N----
VH---IR--TA----RA------KYIDELIDLLFMQASAFQ--------Q-E-K----WQ---G----WS------A-----Q---S---------PD--------LPR---HQQLWL-----D----------------PWR--SLS--------------DETFKQ--ERE--KND-WQ-VTVADDFARWLN-----YR----L---K-------K-------S-S-F------------DV---GAVEQ------KEWRS-Q-------SL--FA-----------------Q-----RMREMEAVL-----------------QEAL-------K-----------------------
---------------
WP_130236270 ------------------------------------------------------------MEK-------------TALI--QFIT--D-YIA-SR-----R----QPK----------------------------------------------------------------------------------------------I---------D-------A-FEK--EAAK------R--L---------------------------E-QGEDAS---------------------------VI-------------------------------------A-LER-QE-----------------------LE---------
AR------YL---PRHW-----LT---DA-----AKRA-------G----QIK-L-----------------VTHGAKFSHGD---S----------------K---A-S------SF----Y----L----E-------TS---------A--T--ES--Y-------LNT--A--SL------A-------N--I--A-TD-A-IG----N----AA----A--L-----DVAKLL-Q-----T----D--V--N---------GDSL------------LASMK-RGD-Y---QVLSTF---A---E-----DK----AQLEQWVAGFSQA--W---TT--GR----
PASHKLAKQIYFPV--------------------------ADG---YHLLCPLFSSSLAQVMYEK-LT----A---V---R--F-S----------------EE-SKAIR-D-----AR-------------K--AGKWH--SQ-P-DIR-----FPN-LAEMHFGG-T----KPQ---------NISLLN--SVRG----------GRVWLLPSM-P----PTWA--T---LD-RAPQNM----LSIF----AL-RGDFN--R-----AA--SGIIARMTYLL------K----------V------D--------------IN---------N----
VH---IR--TA----RA------KYIDELIDLLFMQASSFQ--------Q-E-K----WQ---G----WS------A-----Q---S---------PD--------LPR---HQQLWL-----D----------------PWR--SLS--------------DETFKL--ERE--KND-WQ-GTVADDFARWLN-----YR----L---K-------K-------A-R-F------------DV---GAVEQ------REWRS-Q-------SL--FT-----------------Q-----RMREMEAVL-----------------QEAL-------K-----------------------
---------------
WP_131162678 ------------------------------------------------------------MEK-------------TALI--QFIT--D-YIA-SR-----R----QPK----------------------------------------------------------------------------------------------I------------------EAFEK--EAAK------R--L---------EQ-----------------------GEDAS---------------------------VI-------------------------------------A-LER-QE-----------------------LE---------
AR------YL---PRNW-----LT---DA-----AKRA-------G----QIK-L-----------------VTHGAKFSHGD---S----------------K---A-S------SF----Y----L----E------------T----SAN--E--SY---------LNT--A--SL------V-------N--V--A-TD-A-IG----N----AA----A--L-----DVAKLL-Q-----T----D--V--N---------GDSL------------LASLK-RGD-Y---QALSTF---A---E-----DK----AQLELWVAGFSQA--W---TT--GQ----
PASHKLAKQIYFPV--------------------------ADS---YHLLCPLFSSSLAQAMYEK-LT----A---V---R--F-S----------------EE-SKAIR-D-----AR-------------K--AGKWH--SQ-P-DIR-----FPN-LAEMHFGG-T----KPQ---------NISLLN--SVRG----------GRVWLLPSM-P----PTWE--T---LD-RAPQNM----RSIF----SL-RGDFN--R-----AA--SGIVARMTYLL------K----------V------D--------------IN---------N----
VH---IR--TA----RA------KYIDELIDLLFMQASSFQ--------Q-E-K----WQ---G----WS------V-----Q---S---------PD--------LPR---HQQLWL-----D----------------PWR--SLS--------------DETFKQ--ERE--KND-WQ-ETVADDFARWLN-----YR----L---K-------K-------A-R-F------------DV---GAVEQ------REWRS-Q-------SL--FT-----------------Q-----RMREMEAVL-----------------QEAL-------K-----------------------
----------------
WP_135524506 ------------------------------------------------------------MEK-------------TALI--QFIT--D-YIP-SR-----R----QPK----------------------------------------------------------------------------------------------I---------E-------A-FEK--EAAK------R--L---------------------------E-QGEDAS---------------------------VI-------------------------------------A-LER-QE-----------------------LE---------
AR------YL---PRNW-----LT---DA-----AKRA-------G----QIK-L-----------------VTHGAKFSHGD---S----------------K---A-S------SF----Y----L----E-------TS---------A--N--ES--Y-------LNT--A--SL------V-------N--V--A-TD-A-IG----N----AA----A--L-----DVAKLL-Q-----T----D--V--N---------GDSL------------LASLK-RGD-Y---QALSTF---A---E-----DK----AQLELWVAGFSQA--W---TT--GQ----
PASHKLAKQIYFPV--------------------------ADS---YHLLCPLFSSSLAQAMYEK-LT----A---V---R--F-S----------------EE-SKAIR-D-----AR-------------K--AGKWH--SQ-P-DIR-----FPN-LAEMHFGG-T----KPQ---------NISLLN--SVRG----------GRVWLLPSM-P----PTWE--T---LD-RAPQNM----RSIF----SL-RGDFN--R-----AA--SGIVARMTYLL------K----------V------D--------------IN---------N----
VH---IR--TA----RA------KYIDELIDLLFMQASSFQ--------Q-E-K----WQ---G----WS------V-----Q---S---------PD--------LPR---HQQLWL-----D----------------PWR--SLS--------------DETFKQ--ERE--KND-WQ-ETVADDFARWLN-----YR----L---K-------K-------A-R-F------------DV---GAVEQ------REWRS-Q-------SL--FT-----------------Q-----RMREMEAVL-----------------QEAL-------K-----------------------
----------------
WP_135490215 ------------------------------------------------------------MEK-------------TALI--QFIT--D-YIA-SR-----R----QPK----------------------------------------------------------------------------------------------I------------------EAFEK--EAAK------R--L---------EQ-----------------------GEDAS---------------------------VI-------------------------------------A-LER-QE-----------------------LE---------
AR------YL---PRNW-----LT---DA-----AKRA-------G----QIK-L-----------------VTHGAKFSHGD---S----------------K---A-S------SF----Y----L----E------------T----SAN--E--SY---------LNT--A--SL------A-------N--V--A-TD-A-IG----N----AA----A--L-----DVAKLL-Q-----T----D--V--N---------GDSL------------LASLK-RGD-Y---QALSTF---A---E-----DK----AQLEQWVTGFSQA--W---TI--GR----
PVSHKLAKQIYFPV--------------------------ADS---YHLLCPLFSSSLAQAMYEK-LT----A---V---R--F-S----------------EE-SKAIR-D-----AR-------------K--AGKWH--SQ-P-DIR-----FPN-LAEMHFGG-T----KPQ---------NISLLN--SVRG----------GRVWLLPSM-P----PTWE--T---LD-RAPQNM----RSIF----SL-RGDFN--R-----AA--SGIIARMTYLL------K----------V------D--------------IN---------N----
VH---IR--TA----RA------KYIDELIDLLFMQASSFQ--------Q-E-K----WQ---G----WS------A-----Q---S---------PD--------LPR---HQQLWL-----D----------------PWR--SLS--------------DETFKL--ERE--KND-WQ-ETVADDFARWLN-----YR----L---K-------K-------A-R-F------------DV---GAVEQ------REWRS-Q-------SL--FT-----------------Q-----RMREMEAVL-----------------QEAL-------K-----------------------
---------------
WP_135409816 ------------------------------------------------------------MEK-------------TALI--QFIT--D-YIA-SR-----R----QPK----------------------------------------------------------------------------------------------I---------E-------A-FEK--EAAK------R--L---------EQ------GE---------------DAS---------------------------VI-------------------------------------A-LER-QE-----------------------LE---------
AR------YL---PRNW-----LT---DA-----AKRA-------G----QIK-L-----------------VTHGAKFSHGD---S----------------K---A-S------SF----Y----L----E------------T----SAN--E--SY---------LNT--A--SL------A-------N--V--A-TD-A-IG----N----AA----A--L-----DVAKLL-Q-----T----D--V--N---------GDSL------------LASLK-RGD-Y---QALSTF---A---E-----DK----AQLEQWVTGFSQA--W---TI--GR----
PVSHKLAKQIYFPV--------------------------ADS---YHLLCPLFSSSLAQAMYEK-LT----A---V---R--F-S----------------EE-SKAIR-D-----AR-------------K--AGKWH--SQ-P-DIR-----FPN-LAEMHFGG-T----KPQ---------NISLLN--SVRG----------GRVWLLPSM-P----PTWE--T---LD-RAPQNM----RSIF----SL-RGDFN--R-----AA--SGIIARMTYLL------K----------V------D--------------IN---------N----
VH---IR--TA----RA------KYIDELIDLLFMQASSFQ--------Q-E-K----WQ---G----WS------A-----Q---S---------PD--------LPR---HQQLWL-----D----------------PWR--SLS--------------DETFKQ--ERE--KND-WQ-ETVADDFARWLN-----YR----L---K-------K-------A-R-F------------DV---GAVEQ------REWRS-Q-------SL--FT-----------------Q-----RMREMEAVL-----------------QEAL-------K-----------------------
---------------
WP_135524101 ---------------------------------------------MKAQGGN----------LLN-------------NNLV--DFIT--D-YIA-SR-----R----QPK----------------------------------------------------------------------------------------------I---------D-------A-FEK--EAAK------R--L---------------------------E-QGEDAS---------------------------II-------------------------------------A-LER-QE-----------------------LE---
------AR------YL---PRNW-----LT---DA-----AKRA-------G----QIK-L-----------------VTHGAKFSHGD---S----------------K---A-S------SF----Y----L----E-------TS---------A--N--ES--Y-------LNT--A--SL------A-------N--V--A-TD-A-IG----N----AA----A--L-----DVAKLL-Q-----T----D--V--N---------GNSL------------LASLK-RGD-Y---QALSTF---A---E-----DK----AQLEQWVAGFSQA--W---TT--GR----
PASHKLAKQIYFPV--------------------------ADG---YRLLCPLFSSSLAQAMYEK-LT----A---V---R--F-S----------------EE-SKAIR-D-----AR-------------K--AGKWH--SQ-P-EIR-----FPN-LAEMHFGG-T----KPQ---------NISLLN--SVRG----------GRIWLLPSM-P----PTWA--T---LD-HAPQNM----RSIF----AL-RGDFN--R-----AA--SGIIARMTYLL------K----------V------D--------------IN---------N----VH-
--IR--TA----RA------KYIDELIDLLFMQASSFQ--------Q-E-K----WQ---G----WS------A-----Q---S---------PD--------LPR---HQQFWL-----D----------------PWR--SLS--------------DETFKQ--ERE--KND-WQ-ETVADDFARWLN-----YR----L---K-------K-------A-R-F------------DV---GAVEQ------REWRS-Q-------SL--FT-----------------Q-----RMREMEAVL-----------------QEAL-------K-----------------------
-----------
WP_137555826 ------------------------------------------------------------MN--------------NNLV--DFIT--D-YIA-SR-----R----QPK----------------------------------------------------------------------------------------------I------------------DAFEK--EAAK------R--L---------EQ-----------------------GEDAS---------------------------VI-------------------------------------A-QER-QE-----------------------LE---------
AR------YL---PRNW-----LT---DA-----AKRA-------G----QIK-L-----------------VTHGAKFSHGD---S----------------K---A-S------SF----Y----L------------------DT----SAT--E--NY---------LNT--A--SL------A-------N--I--A-TD-A-IG----N----AA----A--L-----DVAKLL-Q-----T----D--V--N---------GDTL------------LASLK-RGD-Y---QALSTF---A---E-----DK----AQLEQWVAGFSQA--W---TT--GR----
PASHKLAKQIYFPV--------------------------ADG---YHLLCPLFSSSLAQAMYEK-LT----A---V---R--F-S----------------EE-SKAIR-D-----AR-------------K--AGKWH--SQ-P-DIW-----FPN-LAEMHFGG-T----KPQ---------NISLLN--SVRG----------GRVWLLPSM-P----PTWA--T---LD-RAPQKM----RSIF----AL-RGDFN--R-----AA--SGIIARMTYLL------K----------V------D--------------IN---------N----
VH---IR--TA----RA------KYIDELIDLLFMQASSFQ--------Q-E-K----WQ---G----WS------A-----Q---S---------PD--------LPR---HQQLWL-----D----------------PWR--SLS--------------DETFKQ--ERE--KND-WH-ETVADDFARWLN-----YR----L---K-------K-------S-S-F------------DV---GAVEQ------KEWRS-Q-------SL--FA-----------------Q-----RMREMEAVL-----------------QEAL-------K-----------------------
---------------
WP_001063627 ------------------------------------------------------------MN--------------NNLV--DFIT--D-YIA-SR-----R----QPK----------------------------------------------------------------------------------------------I------------------EAFEK--EAAK------R--L---------EQ-----------------------GEDAS---------------------------VI-------------------------------------A-QER-QE-----------------------LE---------
AR------YL---PRHW-----LT---DA-----AKRA-------G----QIK-L-----------------VTHGAKFSHGD---S----------------K---A-S------SF----Y----L----E------------T----SAN--E--SY---------LNT--A--SL------A-------N--V--A-TD-A-IG----N----AA----A--L-----DVAKLL-Q-----T----D--V--N---------GDSL------------LASLK-RGD-Y---QALSTF---A---E-----DK----AQLEQWVAGFSQA--W---TT--GR----
PASHKLAKQIYFPV--------------------------ADG---YHLLCPLFSSSLAQAMYEK-LT----A---V---R--F-S----------------EE-SKAIR-D-----AR-------------K--AGKWH--SQ-P-DIW-----FPN-LAEMHFGG-T----KPQ---------NISLFN--SVRG----------GRVWLLPSM-P----PTWA--T---LD-RAPQNM----RSIF----AL-RGDFN--R-----AA--SGIIARMTYLL------K----------V------D--------------IN---------N----
VH---IR--TA----RA------KYIDELIDLLFMQASAFQ--------Q-E-K----WQ---G----WS------A-----Q---S---------QD--------LPR---HQQLWL-----D----------------PWR--SLS--------------DEAFKL--DRE--KND-WQ-ETVADDFARWLN-----YR----L---K-------K-------S-R-F------------DV---GAVEQ------REWRS-Q-------SL--FT-----------------Q-----RMREMEAVL-----------------QEAL-------K-----------------------
---------------
WP_130215667 ------------------------------------------------------------MEK-------------TALI--QFIT--G-YIA-SR-----R----QPK----------------------------------------------------------------------------------------------I---------E-------A-FEK--EAAK------R--Q---------------------------E-QGEDAS---------------------------VI-------------------------------------A-LER-QE-----------------------LE---------
AR------YL---PRNW-----LT---DA-----AKRA-------G----QIK-L-----------------VTHGAKFSHGD---S----------------K---A-S------SF----Y----L----D-------TS---------A--T--EN--Y-------LNT--A--SL------A-------N--I--A-TD-A-IG----N----AA----A--L-----DVAKLL-Q-----T----D--V--N---------GDSL------------LASLK-RGD-Y---QALSTF---A---E-----DK----AQLEQWVAGFSQA--W---TT--GR----
PASHKLAKQIYFPV--------------------------ADG---YHLLCPLFSSSLAQAMYEK-LT----A---V---R--F-S----------------EE-SKAIR-D-----AR-------------K--AGKWH--SQ-P-DIW-----FPN-LAEMHFGG-T----KPQ---------NISLLN--SVRG----------GRVWLLPSM-P----PTWA--T---LD-RAPQNM----RSIF----SL-RGDFN--R-----AA--SGIIARMTYLL------K----------V------D--------------IN---------N----
VH---IR--TA----RA------KYIDELIDLLFMQASSFQ--------Q-E-K----WQ---G----WS------A-----Q---S---------PD--------LPR---HQQLWL-----D----------------PWR--SLS--------------DETFKQ--ERE--KND-WQ-ETVADDFARWLN-----YR----L---K-------K-------A-H-F------------DV---GAVEQ------REWRS-Q-------SL--FT-----------------Q-----RMREMEAVL-----------------QEAL-------K-----------------------
---------------
WP_135486317 ------------------------------------------------------------MN--------------NNLV--DFIT--D-YIA-SR-----R----QPK----------------------------------------------------------------------------------------------I---------E-------A-FEK--EATK------R--Q---------------------------E-QGEDAS---------------------------VI-------------------------------------A-LER-QE-----------------------LE---------
AR------YL---PRNW-----LT---DA-----AKRA-------G----QIK-L-----------------VTHGAKFSHGD---S----------------K---A-S------SF----Y----L----D-------TS---------A--T--EN--Y-------LNT--A--SL------A-------N--I--A-TD-A-IG----N----AA----A--L-----DVAKLL-Q-----T----D--V--N---------GDSL------------LASLK-RGD-Y---QALSTF---A---E-----DK----AQLEQWVAGFSQA--W---TT--GR----
PASHKLAKQIYFPV--------------------------ADG---YHLLCPLFSSSLAQAMYEK-LT----A---V---R--F-S----------------EE-SKAIR-D-----AR-------------K--AGKWH--SQ-P-DIW-----FPN-LAEMHFGG-T----KPQ---------NISLLN--SVRG----------GRVWLLPSM-P----PTWA--T---LD-RAPQNM----RSIF----SL-RGDFN--R-----AA--SGIIARMTYLL------K----------V------D--------------IN---------N----
VH---IR--TA----RA------KYIDELIDLLFMQASSFQ--------Q-E-K----WQ---G----WS------A-----Q---S---------PD--------LPR---HQQLWL-----D----------------PWR--SLS--------------DETFKQ--ERE--KND-WQ-ETVADDFARWLN-----YR----L---K-------K-------A-R-F------------DV---GAVEQ------REWRS-Q-------SL--FT-----------------Q-----RMREMEAVL-----------------QEAL-------K-----------------------
---------------
WP_135385775 ------------------------------------------------------------MN--------------NNLV--DFIT--D-YIA-SR-----R----QPK----------------------------------------------------------------------------------------------I------------------EAFEK--EAAK------R--L---------EQ-----------------------GEDAS---------------------------VI-------------------------------------A-LER-QE-----------------------LE---------
AR------YL---PRNW-----LT---DA-----AKRA-------G----QIK-L-----------------VTHGAKFSHGD---S----------------K---A-S------SF----Y----L----E------------T----SAN--E--SY---------LNT--A--SL------A-------N--V--A-TD-A-IG----N----AA----A--L-----DVAKLL-Q-----T----D--V--N---------GDSL------------LASLK-RGD-Y---QALSTF---A---E-----DK----AQLEQWVAGFSQA--W---TL--AQ----
PASHKLAKQIYFPV--------------------------DDG---YHLLCPLFSSSLAQAMYEK-LT----A---V---R--F-S----------------EE-SKAIR-D-----AR-------------K--AGKWH--SQ-P-DIR-----FPN-LAEMHFGG-T----KPQ---------NISLLN--SVRG----------GRVWLLPSM-P----PTWA--T---LD-RAPQNM----RSIF----SL-RGDFN--R-----AA--SGIIARMTYLL------K----------V------D--------------IN---------N----
VH---IR--TA----RA------KYIDELIDLLFMQASSFQ--------Q-E-K----WQ---G----WS------A-----Q---S---------PD--------LPR---HQQLWL-----D----------------PWR--SLS--------------DETFKQ--ERE--KND-WQ-ETVADDFARWLN-----YR----L---K-------K-------A-R-F------------DV---GAVEQ------REWRS-Q-------SL--FT-----------------Q-----RMREMEAVL-----------------QEAL-------K-----------------------
---------------
WP_130222131 ------------------------------------------------------------MN--------------NNLV--DFIT--D-YIA-SR-----R----QPK----------------------------------------------------------------------------------------------I------------------EAFEK--EVAK------R--L---------EQ---GEDA-------------------------------------------------SV-------------------------------------IALER-QE-----------------------LE---------
AR------YL---PRNW-----LT---DA-----AKRA-------G----QIK-L-----------------VTHGAKFSHGD---S----------------K---A-S------SF----Y----L----E------------T----SAN--E--SY---------LNT--A--SL------A-------N--I--A-TD-A-IG----N----AA----A--L-----DVAKLL-Q-----T----D--V--N---------GDSL------------LASLK-RGD-Y---QALSTF---A---E-----DK----AQLEQWVAGFSQA--W---TT--GR----
PASHKLAKQIYFPV--------------------------DDG---YHLLCPLFSSSLAQVMYEK-LT----A---V---R--F-S----------------EE-SKAIR-D-----AR-------------K--AGKWH--SQ-P-DIW-----FPN-LAEMHFGG-T----KPQ---------NISLLN--SVRG----------GRVWLLPSM-P----PTWA--T---LD-RAPQNM----RSIF----SL-RGDFN--R-----AA--SGIIARMTYLL------K----------V------D--------------IN---------N----
VH---IR--TA----RA------KYIDELIDLLFMQVSSFQ--------Q-E-K----WQ---G----WS------A-----Q---S---------PD--------LPR---HQQLWL-----D----------------PWR--SLS--------------DETFKQ--ERE--KND-WQ-ETVADDFARWLN-----YR----L---K-------K-------A-R-F------------DV---GAVEQ------REWRS-Q-------SL--FT-----------------Q-----RMREMEAVL-----------------QEAL-------K-----------------------
---------------
WP_135519353 ------------------------------------------------------------MN--------------NNLV--DFIT--D-YIA-SR-----R----QPK----------------------------------------------------------------------------------------------I------------------EAFEK--EAAK------R--L---------EQ-----------------------GEDAS---------------------------II-------------------------------------ALERQELE-----------------
AR------YL---PRNW-----LT---DA-----AKRA-------G----QIK-L-----------------VTHGAKFSHGD---S----------------K---A-S------SF----Y----L----E--------T------SAN--E--SY---------LNT--A--SL------A-------N--V--A-TD-A-IG----N----AA----A--L-----DVAKLL-Q-----T----D--V--N---------GDSL------------LASLK-RGD-Y---QVLSTF---A---E-----DK----AQLEQWVAGFSQA--W---TT--GR----
PASHKLAKQIYFPV--------------------------ADG---YHLLCPLFSSSLAQAMYEK-LT----A---V---R--F-S----------------EE-SKAIR-D-----AR-------------K--AGKWH--SQ-P-DIW-----FPN-LAEMHFGG-T----KPQ---------NISLLN--SVRG----------GRVWLLPSM-P----PTWA--T---LD-RAPQNM----RSIF----SL-RGDFN--R-----AA--SGIIARMTYLL------K----------V------D--------------IN---------N----
VH---IR--TA----RA------KYIDELIDLLFMQASSFQ--------Q-E-K----WQ---G----WS------A-----Q---S---------PD--------LPR---HQQLWL-----D----------------PWR--SLS--------------DETFKQ--ERE--KND-WQ-ETVADDFARWLN-----YR----L---K-------K-------A-H-F------------DV---GAVEQ------REWRS-Q-------SL--FT-----------------Q-----RMREMEAVL-----------------QEAL-------K-----------------------
---------------
WP_135488479 ------------------------------------------------------------MN--------------NNLV--DFIT--D-YIA-SR-----R----QPK----------------------------------------------------------------------------------------------I---------E-------A-FEK--EAAK------R--L---------EQGEDA---------------------------------------------------SV-------------------------------------IALER-QE-----------------------LE---------
AR------YL---PRNW-----LT---DA-----AKRA-------G----QIK-L-----------------VTHGAKFSHGD---S----------------K---A-S------SF----Y----L----E------------T----SAN--E--SY---------LNT--A--SL------A-------N--V--A-TD-A-IG----N----AA----A--L-----DVAKLL-Q-----T----D--V--N---------GDSL------------LASLK-RGD-Y---QALSTF---A---E-----DK----AQLEQWVAGFSQA--W---TT--GR----
PASHKLAKQIYFPV--------------------------DDG---YHLLCPLFSSSLAQAMYEK-LT----A---V---R--F-S----------------EE-SKAIR-D-----AR-------------K--AGKWH--SQ-P-DIW-----FPN-LAEMHFGG-T----KPQ---------NISLLN--SVRG----------GRVWLLPSM-P----PTWA--T---LD-RAPQNM----RSIF----SL-RGDFN--R-----AA--SGIIARMTYLL------K----------V------D--------------IN---------N----
VH---IR--TA----RA------KYIDELIDLLFMQASSFQ--------Q-E-K----WQ---G----WS------A-----Q---S---------PD--------LPR---HQQLWL-----D----------------PWR--SLS--------------DETFKQ--ERE--KND-WQ-ETVADDFARWLN-----YR----L---K-------K-------A-H-F------------DV---GAVEQ------REWRS-Q-------SL--FT-----------------Q-----RMREMEAVL-----------------QEAL-------K-----------------------
---------------
WP_103776180 ------------------------------------------------------------MTT-------------EALT--QFII--Q-YIA-SR-----R----QTK----------------------------------------------------------------------------------------------L------------------EAFDK--EAAK------R--L---------AQ-----------------------GDDTS---------------------------LV-------------------------------------A-QER-QA-----------------------LE---------
AR------YQ---PQNW-----LT---EA-----ARRA-------G----QIS-L-----------------VTQAAKFTHGD---S----------------K---S-S------SI----Y----R----E------------T----PDD--E--GY---------LST--A--AL------D-------N--M--P-AD-A-VG----N----AA----A--L-----DVAKLL-Q-----S----E--V--D---------GDSL------------LACLK-RGE-Y---GPLQTF---A---G-----NG----EQLAQWIKGFNQA--L---TS--AQ----
PTSHKLAKQIYFPV--------------------------GDR---YHLLCPLFATSLAQVMHEK-LI----A---S---R--F-G----------------EA-GKLVR-A-----AR-------------R--AGKWH--PQ-P-DVR-----YPN-LAEMHFGG-T----KPQ---------NISALN--SSRG----------GRVWLLPSQ-P----PKWK--L---LD-KVPQKM----GHLF----SL-RGAFN--Y-----AA--TGIISRMVYLL------K--EN-------A------D--------------KN---------N--
--LS---VR--NA----RA------ACVDELIDLLFMQAAAYQ--------Q-E-N----GQ---G----WT------L-----S---P---------QA--------LPL---HQQLWL-----D----------------PWR--SKT--------------DETFRL--ERE--KGD-WK-AGVADDFARWLN-----QR----L---S-------K-------A-R-L------------EV---GVVER------REWRT-Q-------RR--FD-----------------Q-----RMREMEAIV-----------------QEAL-------K-----------------------
------------------
WP_115181770 ------------------------------------------------------------MTT-------------EALT--HFIT--D-YIA-SR-----R----QPK----------------------------------------------------------------------------------------------L---------E-------A-FEK--EAAK------R--L---------------------------A-QGEDAS---------------------------TI-------------------------------------A-LER-QA-----------------------LE---------
AR------YQ---PQNW-----LT---DA-----ARRA-------G----QIS-L-----------------VTHAAKFTHGD---S----------------K---S-S------SI----Y----R----D-------TV---------G--E--EG--Y-------LST--A--SL------K-------S--L--P-TD-A-VG----N----AA----A--L-----DVAKLL-Q-----T----D--V--E---------GDSL------------LACLK-RGE-H---QPLHAF---A---E-----ND----AQLTQWVEGFNRA--L---TP--TQ----
PTSHKLAKQVYFPV--------------------------VND---YHLLCPLFATSLAQVLHEK-LV----A---S---R--F-G----------------EE-VKAAR-D-----AR-------------K--AGKWH--PQ-P-DVR-----YPD-MAEMHFGG-T----KPQ---------NISALN--SARG----------GRIWLLPSR-P----PQWL--N---LD-KAPQNM----RNLF----AL-RGAFN--R-----AA--SGIISRMVFLL------K--EN-------T------E--------------KN---------N-
---HA---IR--NA----RA------AYVDELIDLLFMHAATFQ--------Q-A-S----WQ---G----WT------R-----N---N---------TD--------LPL---HQQLWL-----D----------------PWR--GQT--------------DETFRL--EHD--KDD-WQ-ISVADDFARWLN-----YR----L---N-------K-------A-R-L------------DV---GLVER------REWRT-Q-------SL--FR-----------------Q-----RMREMEAIL-----------------REAL-------KCVI-----------------
------------------
WP_134769560 ------------------------------------------------------------MTT-------------EALT--HFIT--D-YIA-SR-----R----QPK----------------------------------------------------------------------------------------------L------------------EAFEK--EAAK------R--L---------AQ-----------------------GEDAS---------------------------TI-------------------------------------A-LER-QT-----------------------LE---------
AR------YQ---PQNW-----LT---DA-----ARRA-------E----QIS-L-----------------VTHAAKFTHGD---S----------------K---S-S------SI----Y----R------------------DT----VGE--E--GY---------LST--A--SL------K-------N--L--P-TD-A-VG----N----AA----A--L-----DVAKLL-Q-----T----D--V--E---------GDSL------------LACLK-RGE-H---QPLHAF---A---E-----ND----AQLTQWVEGFNRA--L---TP--TQ----
PTSHKLAKQVYFPV-------G-------------------ND---YHLLCPLFATSLAQVLHEK-LV----A---S---R--F-G----------------EE-AKAAR-D-----AH-------------K--AGKWH--PQ-P-DVR-----YPD-MAEMHFGG-T----KPQ---------NISALN--SARG----------GRIWLLPSQ-P----PQWL--N---LD-KAPQNM----RNLF----AL-RGAFN--R-----AA--SGIVSRMVFLL------K--EN-------T------E--------------KN---------
N----HA---IR--NA----RA------AYVDELIDLLFMHAATFQ--------Q-A-S----WQ---G----WT------R-----N---N---------TD--------LPL---HQQLWL-----D----------------PWR--GQT--------------DETFRL--ERD--KDD-WQ-ISVADDFARWLN-----YR----L---N-------K-------A-C-L------------DV---GLVER------REWRT-Q-------SL--FR-----------------Q-----RMREMEAIL-----------------REAL-------KCVI-----------------
------------------
WP_115614901 ------------------------------------------------------------MTT-------------EALT--HFIT--D-YIA-SR-----R----QPK----------------------------------------------------------------------------------------------L---------E-------A-FEK--EAAK------R--L----A-QGED-----AS---------------------------------------------------TI-------------------------------------A-LER-QA-----------------------LE---------
AR------YQ---PQNW-----LT---DA-----ARRA-------G----QIS-L-----------------VTHAAKFTHGD---S----------------K---S-S------SI----Y----R----D---------T-V------GE--E--GY---------LST--A--SL------K-------S--L--P-TD-A-VG----N----AA----A--L-----DVAKLL-Q-----T----D--V--E---------GDSL------------LACLK-RGE-H---QPLHAF---A---E-----ND----AQLTQWVEGFNRA--L---TP--TQ----
LASHKLAKQVYFPV--------------------------GND---YHLLCPLFATSLAQVLHEK-LV----A---S---R--F-G----------------EE-AKAAR-D-----AH-------------K--AGKWH--PQ-P-DVR-----YPD-MAEMHFGG-T----KPQ---------NISALN--SARG----------GRIWLLPSQ-P----PQWL--N---LD-KAPQNM----RNLF----AL-RGAFN--R-----AA--SGIVSRMVFLL------K--EN-------T------E--------------KN---------
N----HA---IR--NA----RA------AYVDELIDLLFMHAATFQ--------Q-A-S----WQ---G----WT------R-----N---N---------TD--------LPL---HQQLWL-----D----------------PWR--GQT--------------DETFRL--ERD--KDD-WQ-ISVADDFARWLN-----YR----L---N-------K-------A-C-L------------DV---GLVER------REWRT-Q-------SL--FR-----------------Q-----RMREMEAIL-----------------REAL-------KCVI-----------------
------------------
WP_115626544 ------------------------------------------------------------MTT-------------EALT--HFIT--D-YIA-SR-----R----QPK----------------------------------------------------------------------------------------------L---------E-------A-FEK--EAAK------R--L----A-QGED-----AS---------------------------------------------------TI-------------------------------------A-LER-QA-----------------------LE---------
AR------YQ---PQSW-----LT---DA-----ARRA-------G----QIS-L-----------------VTHAAKFTHGD---S----------------K---S-S------SI----Y----R----D---------T-V------GD--E--GY---------LST--V--SL------K-------N--L--P-TD-A-VG----N----AA----A--L-----DVAKLL-Q-----T----D--V--E---------GDSL------------LACLK-RGE-H---QPLHAF---A---E-----ND----AQLTQWVEGFNRA--L---TP--TQ----
PASHKLAKQVYFPV--------------------------GND---YHLLCPLFATSLAQVLHEK-LV----A---S---R--F-G----------------EE-AKAVR-D-----AR-------------K--AGKWH--PQ-P-DVR-----YPD-MAEMHFGG-T----KPQ---------NISALN--SARG----------GRIWLLSSQ-P----PQWL--N---LD-KAPQNM----RNLF----AL-RGAFN--R-----AA--SGIVSRMVFLL------K--EN-------T------E--------------KN---------
N----HA---IR--NA----RA------AYVDELIDLLFMHAATFQ--------Q-A-S----WQ---G----WT------R-----N---N---------TD--------LPL---HQQLWL-----D----------------PWR--GQT--------------DETFRL--QRD--KDD-WQ-ISVADDFARWLN-----YR----L---N-------K-------A-R-L------------DV---GLVER------REWRT-Q-------SL--FR-----------------Q-----RMREMEAIL-----------------REAL-------KCVI-----------------
------------------
WP_123907514 ------------------------------------------------------------MTT-------------EALT--HFIT--D-YIA-SR-----R----QPK----------------------------------------------------------------------------------------------L------------------EAFEK--EAAK------R--L---------AQ-----------------------GEDAS---------------------------TI-------------------------------------A-LER-QT-----------------------LE---------
AR------YQ---PQNW-----LT---DA-----ARRA-------E----QIS-L-----------------VTHAAKFTHGD---S----------------K---S-S------SI----Y----R------------------DT----VGE--E--GY---------LST--A--SL------K-------N--L--P-TD-A-VG----N----AA----A--L-----DVAKLL-Q-----T----D--V--E---------GDSL------------LACLK-RGE-H---QPLHAF---A---E-----ND----AQLTQWVEGFNRA--L---TP--TQ----
PASHKLAKQVYFPV--------------------------GND---YHLLCPLFATSLAQVLHEK-LV----A---S---R--F-G----------------EE-AKAVR-D-----AH-------------K--AGKWH--PH-P-DVR-----YPD-MAEMHFGG-T----KPQ---------NISALN--SARG----------GRIWLLSSQ-P----PQWL--N---LD-KAPQNM----LNLF----AL-RGAFN--R-----AA--SGIVSRMVFLL------K--EN-------T------E--------------KN---------
N----HA---IR--NA----RA------AYVDELIDLLFMHAATFQ--------Q-A-S----WQ---G----WT------R-----N---N---------TD--------LLL---HQQLWL-----D----------------PWR--GQT--------------DETFRL--ERD--KDD-WQ-ISVADDFARWLN-----YR----L---N-------K-------A-R-L------------DV---GLVER------REWRT-Q-------SL--FR-----------------Q-----RMREMEAIL-----------------REAL-------KCVI-----------------
------------------
WP_114848456 ------------------------------------------------------------MTT-------------EALT--HFIT--D-YIA-SR-----R----QPK----------------------------------------------------------------------------------------------L------------------EAFEK--EAAK------R--L---------AQ-----------------------GEDAS---------------------------TI-------------------------------------A-LER-QT-----------------------LE---------
AR------YQ---PQNW-----LT---DA-----ARRA-------E----QIS-L-----------------VTHAAKFTHGD---S----------------K---S-S------SI----Y----R------------------DT----VGE--E--GY---------LST--A--SL------K-------N--L--P-TD-A-VG----N----AA----A--L-----DVAKLL-Q-----T----D--V--E---------GDSL------------LACLK-RGE-H---QPLHAF---A---E-----ND----AQLTQWVEGFNRA--L---TP--TQ----
PASHKLAKQVYFPV--------------------------GND---YHLLCPLFATSLAQVLHEK-LV----A---P---R--F-G----------------EE-AKAVR-D-----AH-------------K--AGKWH--PH-P-DVR-----YPD-MAEMHFGG-T----KPQ---------NISALN--SARG----------GRIWLLSSQ-P----PQWL--N---LD-KAPQNM----LNLF----AL-RGAFN--R-----AA--SGIVSRMVFLL------K--EN-------T------E--------------KN---------
N----HA---IR--NA----RA------AYVDELIDLLFMHAATFQ--------Q-A-S----WQ---G----WT------R-----N---N---------TD--------LLL---HQQLWL-----D----------------PWR--GQT--------------DETFRL--ERD--KDD-WQ-ISVADDFARWLN-----YR----L---N-------K-------A-R-L------------DV---GLVER------REWRT-Q-------SL--FR-----------------Q-----RMREMEAIL-----------------REAL-------KCVI-----------------
------------------
WP_115609435 ------------------------------------------------------------MTT-------------EALT--HFIT--D-YIA-SR-----R----QPK----------------------------------------------------------------------------------------------L------------------EAFEK--EAAK------R--L---------AQ-----------------------GEDAS---------------------------TI-------------------------------------A-LER-QA-----------------------LE---------
AR------YQ---PQNW-----LT---DA-----ARRA-------G----QIS-L-----------------VTHAAKFTHGD---S----------------K---S-S------SI----Y----R------------------DT----VGE--E--GY---------LST--A--SL------K-------N--L--P-TD-A-VG----N----AA----A--L-----DVAKLL-Q-----T----D--V--E---------GDSL------------LACLK-RGE-H---QPLHAF---A---E-----ND----AQLTQWVEGFNRA--L---TP--TQ----
PASHKLAKQVYFPV--------------------------DND---YHLLCPLFATSLAQVLHEK-LV----A---S---R--F-G----------------EE-AKAVR-D-----AR-------------K--AGKWH--PH-P-DVR-----YPD-MAEMHFGG-T----RPQ---------NISALN--SARG----------GRIWLLPSQ-P----PQWL--N---LD-KAPQNM----RNLF----AL-RGAFN--R-----AA--SGIISRMVFLL------K--EN-------T------E--------------KN---------
N----HA---IR--NA----RA------AYVDELIDLLFMHAATFQ--------Q-A-S----WQ---G----WT------R-----N---N---------TD--------LPL---HQQLWL-----D----------------PWR--GQT--------------DETFRL--ERD--KDD-WQ-ISVADDFARWLN-----YR----L---N-------K-------A-R-L------------DV---GLVER------REWRT-Q-------SL--FR-----------------Q-----RMREMEAIL-----------------REAL-------KCVI-----------------
------------------
WP_071259522 ------------------------------------------------------------MTT-------------EALT--HFIT--D-YIA-SR-----R----QPK----------------------------------------------------------------------------------------------L---------E-------A-FEK--EAAK------R--L----A-QGED-----AS---------------------------------------------------TI-------------------------------------A-LER-QT-----------------------LE---------
AR------YQ---PQNW-----LT---DA-----ARRA-------E----QIS-L-----------------VTHAAKFTHGD---S----------------K---S-S------SI----Y----R----D---------T-V------GE--E--GY---------LST--A--SL------K-------N--L--P-TD-A-VG----N----AA----A--L-----DVAKLL-Q-----T----D--V--E---------GDSL------------LACLK-RGE-H---QPLHAF---A---E-----ND----AQLTQWVEGFNRA--L---TP--TQ----
PASHKLAKQVYFPV--------------------------GND---YHLLCPLFATSLVQVLHEK-LV----A---S---R--F-G----------------EE-AKAVR-D-----AR-------------K--AGKWH--PQ-P-DVR-----YPD-MAEMHFGG-T----KPQ---------NISALN--SARG----------GRIWLLSSQ-P----PQWL--N---LD-KAPQNM----RNLF----AL-RGAFN--R-----AA--SGIVSRMVFLL------K--EN-------T------E--------------KN---------
N----HA---IR--NA----RA------AYVDELIDLLFMHAATFQ--------Q-A-S----WQ---G----WT------R-----N---N---------TD--------LPL---HQQLWL-----D----------------PWR--GQT--------------DETFRL--ERD--KDD-WQ-ISVADDFARWLN-----YR----L---N-------K-------A-R-L------------DV---GLVER------REWRT-Q-------SL--FR-----------------Q-----RMREMEAIL-----------------REAL-------KCVI-----------------
------------------
WP_049011450 ------------------------------------------------------------MTT-------------EALT--HFIT--D-YIA-SR-----R----QPK----------------------------------------------------------------------------------------------L---------E-------A-FEK--EAAK------R--L----A-QGED-----AS---------------------------------------------------TI-------------------------------------A-LER-QA-----------------------LE---------
AR------YQ---PQNW-----LT---DA-----ARRA-------G----QIS-L-----------------VTHAAKFTHGD---S----------------K---S-S------SI----Y----R----D---------T-V------GE--E--GY---------LST--A--SL------K-------N--L--P-TD-A-VG----N----AA----A--L-----DVAKLL-Q-----T----D--V--E---------GDSL------------LACLK-RGE-H---QPLHAF---A---E-----ND----AQLTQWVEGFNRA--L---TP--TQ----
PASHKLAKQVYFPV--------------------------DND---YHLLCPLFATSLAQVLHEK-LV----A---S---R--F-G----------------EE-AKAVR-D-----AR-------------K--AGKWH--PQ-P-DVR-----YPD-MAEMHFGG-T----KPQ---------NISALN--SARG----------GRIWLLSSQ-P----PQWL--N---LD-KAPQNM----RNLF----AL-RGAFN--R-----AA--SGIVSRMVFLL------K--EN-------T------E--------------KN---------
N----HA---IR--NA----RA------AYVDELIDLLFMHAATFQ--------Q-A-S----WL---G----WT------R-----N---N---------TD--------LPL---HQQLWL-----D----------------PWR--GQT--------------DETFRL--ERD--KDD-WQ-ISVADDFARWLN-----YR----L---N-------K-------A-R-L------------DV---GLVER------REWRT-Q-------SL--FR-----------------Q-----RMREMEAIL-----------------REAL-------KCVI-----------------
------------------
WP_130028930 ------------------------------------------------------------MTT-------------EALT--HFIT--D-YIA-SR-----R----QPK----------------------------------------------------------------------------------------------L---------E-------A-FEK--EAAK------R--L----A-QGED-----AS---------------------------------------------------TI-------------------------------------A-LER-QA-----------------------LE---------
AR------YQ---PQNW-----LT---DA-----ARRA-------E----QIS-L-----------------VTHAAKFTHGD---S----------------K---S-S------SI----Y----R----D---------T-V------GE--E--GY---------LST--A--SL------K-------N--L--P-TD-A-VG----N----AA----A--L-----DVAKLL-Q-----T----D--V--E---------GDSL------------LACLK-RGE-H---QPLHVF---A---E-----ND----AQLTQWVEGFNRA--L---TP--TQ----
PASHKLAKQVYFPV--------------------------DND---YHLLCPLFATSLAQVLHEK-LV----A---S---R--F-G----------------EE-AKAVR-D-----AR-------------K--AGKWH--PQ-P-DVR-----YPD-MAEMHFGG-T----KPQ---------NISALN--SARG----------GRIWLLSSQ-P----PQWL--N---LD-KAPQNM----PNLF----AL-RGAFN--R-----AA--SGIVSRMVFLL------K--EN-------T------E--------------KN---------
N----HA---IR--NA----RA------AYVDELIDLLFMHAATFQ--------Q-A-S----WQ---G----WT------R-----N---N---------TD--------LPL---HQQLWL-----D----------------PWR--GQT--------------DETFRL--ERD--KDD-WQ-ISVADDFARWLN-----YR----L---N-------K-------A-R-L------------DV---GLVER------REWRT-Q-------SL--FR-----------------Q-----RMREMEAIL-----------------REAL-------KCVI-----------------
------------------
WP_058668947 ------------------------------------------------------------MTT-------------EALT--HFIT--D-YIA-SR-----R----QPK----------------------------------------------------------------------------------------------L------------------EAFEK--EAAK------R--L---------AQ-----------------------GEDAS---------------------------TI-------------------------------------A-LER-QA-----------------------LE---------
AR------YQ---PQNW-----LT---DA-----ARRA-------E----QIS-L-----------------VTHAAKFTHGD---S----------------K---S-S------SI----Y----R------------------DT----VGE--E--GY---------LST--A--SL------K-------N--L--P-TD-A-VG----N----AA----A--L-----DVAKLL-Q-----T----D--V--E---------GDSL------------LACLK-RGE-H---QPLHAF---A---E-----ND----AQLTQWVEGFNRA--L---TP--TQ----
PASHKLAKQVYFPV--------------------------DND---YHLLCPLFATSLAQVLHEK-LV----A---S---R--F-G----------------EE-AKAVR-D-----AR-------------K--AGKWH--PQ-P-DVR-----YPD-MAEMHFGG-T----KPQ---------NISALN--SARG----------GRIWLLSSQ-P----PQWL--N---LD-KAPQNM----RNLF----AL-RGAFN--R-----AA--SGIVSRMVFLL------K--EN-------T------E--------------KN---------
N----HA---IR--NA----RA------AYVDELIDLLFMHAATFQ--------Q-A-S----WQ---G----WT------R-----N---N---------TD--------LPL---HQQLWL-----D----------------PWR--GQT--------------DETFRL--ERD--KDD-WQ-ISVADDFARWLN-----YR----L---N-------K-------A-R-L------------DV---GLVER------REWRT-Q-------SL--FR-----------------Q-----RMREMEAIL-----------------REAL-------KCVI-----------------
------------------
WP_014228623 ------------------------------------------------------------MIT-------------EALA--HFIT--D-YIA-SR-----R----QPK----------------------------------------------------------------------------------------------L------------------EAFEK--EAEK------R--L---------AQ------GE--------------DAG---------------------------AI-------------------------------------A-QER-QD-----------------------LE---------
VK------YQ---PCNW-----LT---DA-----ARRA-------G----QIS-L-----------------VTHAAKFTHGD---S----------------K---S-S------SI----Y----R----E------------T----AGD--E--GY---------LST--A--VL------A-------N--V--A-AD-A-VG----N----AA----A--L-----DVAKLL-Q-----T----E--V--G---------GDSL------------LACLK-RGD-R---SPLKAF---A---E-----NG----AQLALWIEGFSQA--L---TP--AQ----
PASHKLAKQIYFPV--------------------------DGG---YHLLSPLFATSLAQAMHEK-MV----A---A---R--F-G----------------EE-AKAVR-E-----AH-------------R--TGKWH--AQ-P-DVR-----YPN-VAEMHFGG-T----KPQ---------NISALN--SSRG----------GRIWLLPSQ-P----PKWT--M---LD-KAPQNL----ASLF----AL-RGEFN--R-----AA--TGTVSRMVSLL------K--EK-------T------D--------------KN---------
N----RH---IR--RV----RA------GYVDELIDLLFLQAATYQ--------Q-E-A----WQ---G----WT------L-----N---S---------PD--------LPL---HQQLWL-----D----------------PWR--SKT--------------DETFRS--ERE--KGA-WQ-RNIADDFSRWLN-----YR----L---N-------K-------A-R-L------------DV---GISER------REWRT-Q-------RL--FS-----------------Q-----RMREMDAIV-----------------REAL-------K-----------------------
------------------
WP_049591102 ------------------------------------------------------------MTT-------------EALT--HFII--H-YIT-SR-----R----QPK----------------------------------------------------------------------------------------------L------------------DAFEK--EAAK------R--L---------AQ------GE--------------DAG---------------------------AI-------------------------------------A-QER-QE-----------------------LE---------
VK------YQ---PRNW-----LT---DA-----ARRA-------G----QIS-L-----------------VTHAAKFTHGD---S----------------K---S-S------SI----Y----R----E------------T----VGD--E--GY---------LST--A--VL------A-------N--V--A-AD-A-VG----N----AA----A--L-----DVAKLL-Q-----I----E--V--G---------GDSL------------LACLK-RGD-R---SPLMGF---A---E-----ND----AQLALWMEGFSQA--L---TP--VQ----
PASHKLAKQIYFPV--------------------------DGG---YHLLSPLFATSLAQAMHEK-MV----A---A---R--F-G----------------EQ-AKAAR-E-----AR-------------R--AGKWH--AQ-P-DVR-----YPN-VAEMHFGG-T----KPQ---------NISALN--SSRG----------GRVWLLPSQ-P----PKWT--M---LD-KAPQNL----TSLF----AL-RGDFN--R-----AA--TDTVSRMVSLL------K--EK-------T------D--------------NN---------
N----RH---IR--NA----RA------GYVDELIDLLFLHAATYQ--------Q-E-A----WQ---G----WT------L-----N---S---------PD--------LPL---HQQLWL-----D----------------PWR--SKT--------------DETFRS--ERE--KGD-WQ-RSVADDFARWLN-----YR----L---N-------K-------A-R-L------------DV---GIAER------REWQT-Q-------RL--FS-----------------Q-----RMREMEAIV-----------------QEAL-------K-----------------------
------------------
WP_123830263 ------------------------------------------------------------MTT-------------EALT--HFII--H-YIA-SR-----R----QPK----------------------------------------------------------------------------------------------L---------D-------A-FEK--EAAK------R--L---------AQGE-----------------------DAG---------------------------AI-------------------------------------A-QER-QE-----------------------LE---------
VK------YQ---PRNW-----LT---DA-----ARRA-------G----QIS-L-----------------VTHAAKFTHGD---S----------------K---S-S------SI----Y----R----E------------T----LGD--E--GY---------LST--A--VL------A-------N--V--A-VD-A-VG----N----AA----A--L-----DVAKLL-Q-----T----E--V--G---------GDSL------------LACLK-RGD-R---SPLMGF---A---E-----ND----AQLALWMEGFSQA--L---TP--VQ----
PASHKLAKQIYFPV--------------------------DGG---YHLLSPLFATSLAQAMHEK-MV----A---A---R--F-G----------------EQ-AKAAR-E-----AR-------------R--AGKWH--AQ-P-DVR-----YPN-VAEMHFGG-T----KPQ---------NISALN--SSRG----------GRVWLLPSQ-P----PKWT--M---LD-KAPQNL----TSLF----AL-RGDFN--R-----AA--TDTVSRMVSLL------K--EK-------T------D--------------NN---------
N----RH---IR--NA----RA------GYVDELIDLLFLHAATYQ--------Q-E-A----WQ---G----WT------L-----N---S---------PD--------LPL---HQQLWL-----D----------------PWR--SKT--------------DETFRS--ERE--KGD-WQ-RSVADDFARWLN-----YR----L---N-------K-------A-R-L------------DV---GIAER------REWQT-Q-------RL--FS-----------------Q-----RMREMEAIV-----------------QEAL-------K-----------------------
------------------
WP_115239791 ------------------------------------------------------------MTT-------------EALT--HFII--D-YIA-SR-----K----QPK----------------------------------------------------------------------------------------------L------------------DAFEK--EAAK------R--L---------AQ------GE--------------DAG---------------------------AI-------------------------------------A-QER-QE-----------------------LE---------
VK------YQ---PRNW-----LT---DA-----ARRA-------G----QIS-L-----------------VTHAAKFTHGD---S----------------K---S-S------SI----Y----R----E------------T----VGD--E--GY---------LST--A--VL------A-------N--V--A-AD-A-VG----N----AA----A--L-----DVAKLL-Q-----T----E--V--G---------GDSL------------LACLK-RGD-R---SPLMGF---A---E-----ND----AQLALWMEGFSQA--L---TP--AQ----
PASHKLAKQIYFPV--------------------------DGG---YHLLSPLFATSLAQAMHKK-MV----A---A---R--F-G----------------EQ-AKAAR-E-----AR-------------R--AGKWH--AQ-P-DVR-----YPN-IAEMHFGG-T----KPQ---------NISALN--SSRG----------GRVWLLPSQ-P----PKWT--M---LD-KAPQNL----TSLF----AL-RGDFN--R-----AA--TDTVSRMVSLL------K--EK-------T------D--------------NN---------
N----RH---IR--NA----RA------GYVDELIDLLFLHAATYQ--------Q-E-A----WQ---G----WT------L-----N---S---------PN--------LPL---HQQLWL-----D----------------PWR--SKT--------------DETFRS--ARE--KGD-WQ-GSVADDFARWLN-----YR----L---N-------K-------A-R-L------------DV---GIAER------REWQT-Q-------RL--FS-----------------Q-----RMREMEAIV-----------------QEAL-------K-----------------------
------------------
WP_064345130 ------------------------------------------------------------MTT-------------EALT--HFII--D-YIA-SR-----K----QPK----------------------------------------------------------------------------------------------L------------------DAFEK--EAAK------R--L---------AQ------GE--------------DAG---------------------------AI-------------------------------------A-QER-QE-----------------------LE---------
VK------YQ---PRNW-----LT---DA-----ARRA-------G----QIS-L-----------------VTHAAKFTHGD---S----------------K---S-S------SI----Y----R----E------------T----VGD--E--GY---------LST--A--VL------A-------N--V--A-AD-A-VG----N----AA----A--L-----DVAKLL-Q-----T----E--V--G---------GDSL------------LACLK-RGD-R---SPLMGF---A---E-----ND----AQLALWMEGFSQA--L---TP--AQ----
PTSHKLAKQIYFPV--------------------------DGG---YHLLSPLFASSLAQAMHEK-MV----A---A---R--F-G----------------EQ-AKAAR-E-----AR-------------R--AGKWH--AQ-P-DVR-----YPN-VAEMHFGG-T----KPQ---------NISALN--SSRG----------GRVWLLPSQ-P----PKWT--M---LD-KAPQNL----TSLF----AL-RGDFN--R-----AA--TDTVSRMVSLL------K--EK-------T------D--------------NN---------
N----RH---IR--NA----RA------GYVDELIDLLFLHAATYQ--------Q-E-A----WQ---G----WT------L-----N---S---------PD--------LPL---HQQLWL-----D----------------PWR--SKT--------------DETFRS--ERE--KGD-WQ-RSVADDFARWLN-----YR----L---N-------K-------A-R-L------------DV---GIAER------REWQT-Q-------RL--FS-----------------Q-----RMREMEAIV-----------------QEAL-------K-----------------------
------------------
WP_105889986 ------------------------------------------------------------MTT-------------EALT--HFII--D-YIA-SR-----K----QPK----------------------------------------------------------------------------------------------L------------------DAFEK--EAAK------R--L---------AQ------GE--------------DAG---------------------------AI-------------------------------------A-QER-QE-----------------------LE---------
VK------YQ---PRNW-----LT---DA-----ARRA-------G----QIS-L-----------------VTHAAKFTHGD---S----------------K---S-S------SI----Y----R----E------------T----VGD--E--GY---------LST--A--VL------A-------N--V--A-AD-A-VG----N----AA----A--L-----DVAKLL-Q-----T----E--V--G---------GDSL------------LACLK-RGD-R---SPLMGF---A---E-----ND----AQLALWMEGFSHA--L---TP--AQ----
PASHKLAKQIYFPV--------------------------DGG---YHLLSPLFATSLAQAMHEK-MV----A---A---R--F-G----------------EQ-AKAAR-E-----AR-------------R--AGKWH--AQ-P-DVR-----YPN-IAEMHFGG-T----KPQ---------NISALN--SSRG----------GRVWLLPSQ-P----PKWT--M---LD-KAPQNL----TSLF----AL-RGDFN--R-----AA--TDTVSRMVSLL------K--EK-------T------D--------------NN---------
N----RH---IR--NA----RA------GYVDELIDLLFLHAATYQ--------Q-E-A----WQ---G----WT------L-----N---S---------PD--------LPL---HQQLWL-----D----------------PWR--SKT--------------DETFRS--ERE--KGD-WQ-RSVADDFARWLN-----YR----L---N-------K-------A-R-L------------DV---GIAER------REWQT-Q-------RL--FS-----------------Q-----RMREMEAIV-----------------QEAL-------K-----------------------
------------------
WP_023320639 ------------------------------------------------------------MTT-------------EALT--HFII--D-YIA-SR-----K----QPK----------------------------------------------------------------------------------------------L---------D-------A-FEK--EAAK------R--L---------AQGE-----------------------DAG---------------------------AI-------------------------------------A-QER-QE-----------------------LE---------
VK------YQ---PRNW-----LT---DA-----ARRA-------G----QIS-L-----------------VTHAAKFTHGD---S----------------K---S-S------SI----Y----R----E------------T----VGD--E--GY---------LST--A--VL------A-------N--V--A-AD-A-VG----N----AA----A--L-----DVAKLL-Q-----T----E--V--G---------GDSL------------LACLK-RGD-R---SPLMGF---A---E-----ND----AQLALWMEGFSQA--L---TP--AQ----
PASHKLAKQIYFPV--------------------------DGG---YHLLSPLFATSLAQAMHEK-MV----A---A---R--F-G----------------EQ-AKAAR-E-----AR-------------R--AGKWH--AQ-P-DVR-----YPN-VSEMHFGG-T----KPQ---------NISALN--SSRG----------GRVWLLPSQ-P----PKWT--M---LD-KAPQNL----TSLF----AL-RGDFN--R-----AA--TDTVSRMVSLL------K--EK-------T------D--------------NN---------
N----RH---IR--NA----RA------GYVDELIDLLFLHAATYQ--------Q-E-A----WQ---G----WT------L-----N---S---------PD--------LPL---HQQLWL-----D----------------PWR--SKT--------------DETFRS--ERE--KGD-WQ-RSVADDFARWLN-----YR----L---N-------K-------A-R-L------------DV---GIAER------REWQT-Q-------RL--FS-----------------Q-----RMREMEAIV-----------------QEAL-------K-----------------------
------------------
WP_131867806 ------------------------------------------------------------MEK-------------HALT--DFIV--N-YVE-KR-----L----QAK----------------------------------------------------------------------------------------------L---------I-------T-FDK--EADK------A--L---------ST------LK------G---ETLALAES-------------------------EL-------------------------------------L-KKR-RE-----------------------LE----
----QL------YE---IRTW-----LT---DA-----AKQS-------Y----QIK-Q-----------------ATHILKFTHSS---A----------------K---GG-S------SV----L----N----D-------YK-------DVS--S---Q--F-------LST--A--SL------I-------N--P--V-MDTA-IS----D----AK----V--L-----GIAALL-T-----E----V--H--Q---------DDSL------------IAALK-RRD-Y---SALEKI---A---E-----GP----EQLEQWVTGFQKI--F---ID--PH----
ISSHKFAKQIFFPI--------------------------STEK---YHLISPLYSSSLAQALYQR-IT----H---A---R--F-S----------------EE-AKAIK-L-----AQ-------------K--ENRQH--EK-A-SVF-----FPN-LAVQNIGG-T----KPQ---------NISALN--SARS----------GRSYLLSCA-A----PQWK--T---VE-KPPQKT----MSIF----HE-RSEVD--F-----YT--RATMVAMHHFL------L--SV--------R-----T--------------LPN---------N----
RN---IR--RQ----RL------NYLDQLIDQLFFYISTVQ--N------M---R-----A---G----WS------V-----D-----------ST--------LSR---AQQLWL-----D----------------PHR--GMN--------------DAAFRN--ERE--AGD-WQ-AEVAYDFGRWLN-----SR----L---K-------H-------E-R-L------------LF---GEAER------REWST-A-------TY--FK-----------------R-----RLRNLESLI-----------------KVEI-------A------
---------
WP_121576227 ------------------------------------------------------------MKE-------------DGLS--RFII--D-YIE-NR-----K----RLK----------------------------------------------------------------------------------------------L---------E-------------AAEK---------K---------FA-----DD-----------------------------------------------------AL-------------------------------------ALAVERQD-------------------IA---------SR---
---YQ---VQAW-----LT---DA-----ASRA-------G----QIS-L-----------------VTHALKFTHSD---A----------------K---G-S------SV----L----S----A-----------QAP---EEN--H--EY---------LST--A--TL------E-------Q--P--T-VD-A-VG----N----AA----A--L-----DVAKLL-Q-----T----E--Y--Q---------GDSL------------VSALQ-RGD-H---SALAGL---A---D-----DP----QQLAQWVAGFNQV--L---MD--KQ----
PSSHKLVKQIYFPL-------SD-------AL-----------ADDH---YHLISPLYSSSLAQALYQR-IT----A---A---R--F-S----------------DE-AKAVR-Q-----AR-------------K--EKRWH--DG-V-EVS-----YPN-LAVQNMGG-T----KPQ---------NISFLN--SARG----------GKSYLLPCA-A----PIWE--S---IA-KPPLQH----KSIF----RP-RGEFE--H-----LS--RSTVQQMRQFL------L--SV--------A-----A--------------VGN---------
T----ME---IR--QQ----CW------AYADSIIDNLFNYVAAIQ--N------L---D-----A---G----WS------K-----Q---E---T-----CL--------LKR---SQQLWL-----D----------------PYR--AES--------------DQDFKF--ERD--GGD-WK-KEVAHDFGVWLN-----RQ----L---E-------H-------E-Q-L------------RF---SEVER------REWST-A-------AL--FK-----------------R-----RMRESEFAL-----------------TEEM-------Q------------------
--------------------
WP_095848584 ------------------------------------------------------------MEE-------------VGLS--RFII--D-YIE-NR-----R----RLR----------------------------------------------------------------------------------------------L---------E-------V-AEK--K-------------------------------------------FANDAL------------------------AL-------------------------------------A-AER-QE-----------------------IN---------DR---
---YQ---VRAW-----LT---DA-----ASRA-------E----QIS-L-----------------VTHALKFTHSD---A----------------K---G-S------SV----F----S----A-------AP-------VEE--S--PE--Y-------LST--A--AL------R-------Q--P--A-VD-A-VG----N----AA----A--L-----DVAKLL-Q-----T----E--Y--G---------GDSL------------AAALQ-RED-H---AALAVL---T---D-----DP----QQLTQWITGFKRA--L---TD--KQ----
LRSHKLAKQIYFPL-------SD-------TA-----------AGSQ---YHLLSPLYSSSLAQALHQR-IT----V---A---R--F-S----------------DE-AKAIR-Q-----AR-------------K--EQRWH--DA-I-DVS-----YPN-LAVQNMGG-T----KPQ---------NISFLN--SGRR----------GKSYLLPCA-A----PVWQ--S---IA-KPPLQH----KSIF----RP-RGEFE--H-----HA--RPAVRQMQQFL------L--SV--------A-----P--------------SGN---------
T----LE---MR--RQ----CW------AYADSIIDSLFNYAAAIQ--N------L---D-----A---G----WS------A-----N---E---C-----CE--------LKR---SQQLWL-----D----------------PWR--AES--------------DGDFKF--ARG--GGG-WK-QEIAQDFGVWLN-----RQ----L---E-------H-------E-Q-L------------RF---SEVER------REWST-A-------AL--FK-----------------R-----RMRESESAL-----------------TEAM-------R------------------
--------------------
WP_012764362 ------------------------------------------------------------ME--E----------SELT--QFIV--S-YIN-DK-----E----KSK----------------------------------------------------------------------------------------------Q---------D-------D-LDK--VADN------QR--M---------SL------SG---------EALAAMEL----------------------------KL-------------------------------------E-QER-LE-----------------------I----
--------------------VRTW-----LT---DN-----APKA-------Y----QIK-R-----------------ATHVLKFTHSS---A----------------K---G-S------SR----L----D----LF-------K-------DLS--L--PF---------LST--A--SL------S-------H--P--V-LDTA-VD----N----AA----V--L-----GIAKLL-T-----E----E--H--Q---------GDSL------------LAALQ-RGD-Y---RALEAL---A---E-----SP----EQLAQWLAGFKQV--L---SD--KQ----
PSSHSLAKQIYFPV-------GD----------------------GQ---YHLLSPLYSSSLSQALDQR-LN----A---V---R--F-G----------------EQ-AKAIR-E-----AR-------------R--QKRWH--DE-V-DVS-----YPG-IAVQNMGG-D----NKQ---------NVSALN--VKRN----------GRSYLFNCS-P----PQWR--S---QD-KPPLKH----DTIF----RN-SGELY--T-----RT--RDQLSAMRKFL------L--SV--------K-----D--------------VDN---------
-N----RD---IR--NQ----RR------RYLDQLIDTLFDYVVTVQ--N------Q-Y-P-----A---G----WS------V-----D-----------SC--------LKR---HLQLWL-----D----------------PQR--GSR--------------DSAFLR--ERD--MGD-WQ-QDVAHEFGQWLN-----AR----L---K-------H-------E-K-L------------IF---GEVER------REWST-A-------AL--FK-----------------Q-----RLRETERAL-----------------KEER-------A------------------
--------------------
WP_100850096 ------------------------------------------------------------ME--E----------SELR--QFIV--S-YIE-NR-----K----QAK----------------------------------------------------------------------------------------------L---------D-------A-FDK--AADK------QR--A---------AL------SG---------EALAAAEL----------------------------TL-------------------------------------V-QKR-RK-----------------------IE---
HA--------------YE---VRTW-----LT---DA-----ASKA-------S----QIK-R-----------------ATHVLKFTHSS---A----------------K---G-S------SR----L----D----LF-------K-------DLS--L--PF---------LST--A--SL------S-------H--P--I-LDTA-VD----N----AA----V--L-----GIATFL-T-----E----E--H--Q---------GNSL------------VAALQ-RGD-H---RSLGAL---A---E-----NP----QQLAQWLAGFKQV--L---SD--KQ----
PSSHSLAKQIYFPV-------GD----------------------GE---YHLLSPLYSSSLSQALDQR-LN----A---V---R--F-G----------------EQ-AKAIR-E-----AR-------------R--QKRWH--DE-V-DVN-----YPG-IAVQNMGG-D----NKQ---------NVSALN--VKRN----------GRSYLFNCS-P----PQWR--S---QD-KPPVEH----DTIF----RN-SGELY--T-----RT--RDQLSAMRKFL------L--SV--------K-----E--------------VDN---------
-N----RE---IR--NQ----RR------RYLDQLIDTLFGYVATVQ--N------L-R-P-----A---G----WS------V-----D-----------SR--------LNV---PLQLWL-----D----------------PLR--GLQ--------------DKDFLR--ERD--AGD-WQ-REVAYEFGRWLN-----AR----L---Q-------H-------E-K-L------------IF---GEVER------REWST-A-------AL--FK-----------------Q-----RLRETERLL-----------------KEEL-------A------------------
--------------------
WP_042872680 ------------------------------------------------------------MEE-------------SELT--QFIV--S-YIE-TR-----K----QAK----------------------------------------------------------------------------------------------L---------D-------AFDKA-ADKQR------------------AA---LSGE---------ALAAAEL----------------------------TL-------------------------------------A-QKR-RK-----------------------
IEHA--------------YE---VRTW-----LT---DA-----ASKA-------N----QIK-R-----------------ATHVLKFTHSS---A----------------K---G-S------SR----L----D----L-------F-KDL---SLP-------F---------LST--A--SL------S-------H--P--V-LDTA-VD----T----AA----V--L-----GIATFLTE----------E--H--Q---------GDSL------------VAALQ-RGD-H---RALEAL---A---E-----NA----EQLTQWLAGFGQV--L---SD--KQ----
PSSHKLAKQIYFPV-------GD----------------------GE---YHLLSPLYSSSLSQALDQR-LN----A---V---R--F-G----------------EQ-AKAIR-K-----AR-------------Y--EKRWH--DE-V-DVS-----YPG-IAVQNMGG-G----NQQ---------NVSALN--VKRN----------GRSYLLSCA-P----PQWR--S---QD-KPPLDH----DTIF----RE-RGEVD--A-----LT--YADRRDMRQFL------L--SV--------K-----D--------------VDN---------
-N----RD---IR--NQ----RR------RYLDQLIDTLFGYVATVQ--N------L-R-P-----A---G----WS------K-----D-----------SR--------LNI---PLQLWL-----D----------------PFR--CRQ--------------DDAFCY--ARE--GGD-WK-EQVAREFGLWLN-----AR----L---Q-------H-------E-K-L------------IF---GEVER------REWST-A-------AL--FK-----------------Q-----RLRETERTL-----------------KEEL-------A------------------
--------------------
WP_038916788 ------------------------------------------------------------MS--S-----------GGLQ--AFIL--S-YIE-TR-----K----QAK----------------------------------------------------------------------------------------------L---------D-------A-FDK--AADK------QR--A---------AL------SG---------EVLAAAEV----------------------------AL-------------------------------------L-QQR-RE-----------------------IE---
QA--------------YE---VRTW-----LT---DA-----ASRA-------G----QIS-L-----------------VTHALKFTHSD---A----------------K---G-S------SV----F----S---TE-----------P-------AMA--T--DV---------LST--A--SL------A-------Q--P--A-ID-A-VG----N----AA----A--L-----DVAKLL-Q-----T----E--H--Q---------GDSL------------VAALQ-RGD-H---RALEAL---A---E-----NP----EQLTQWLAGFGQV--L---SD--KQ----
PSSHKLAKQIYFPV-------GD----------------------GE---YHLLSPLYSSSLSQALDQR-LN----A---V---R--F-G----------------EQ-AKAIR-E-----AR-------------R--QKRWH--DE-V-DVS-----YPG-IAVQNMGG-T----KPQ---------NISALN--SARG----------GRSYLFNCA-A----PVWQ--G---VS-KPPQKH----KSVF----RE-RGEID--Y-----YT--RFTRTRMQQFL------L--SV--------K-----D--------------SEN---------
```

```
N----RD---IR--RQ----RL------NYLDQLIEQLFFYVISVQ--K-----L-R-P-----A---G----WS------L-----D------------SQ--------LKR---SQQLWL-----D-------------PCR--SLS-------------DTVFRR--ERD--ADD-WQ-QEVARDFGLWLN-----AR----L---Q-------H-------E-K-L-----------IF---GEVER------REWST-A-------AL--FK----------------Q-----RLRETERAL----------------EEEL-------A-----------------------
---------------
WP_024107197 ------------------------------------------------------------ME--E----------RELT--QFIV--S-YIG-TR-----K----QAK-------------------------------------------------------------------------------------------------L---------D-------A-FDK--AADK-----QR--E---------AL-----SG---------EALAAAEL---------------------------TL-----------------------------------A-QDR-RE-----------------------IE--
LA-------------YE---VRGW-----LT---DA-----ASRA-------G----QIS-L-----------------VTHALKFTHSD---A-----------------K---G-S------SV----F----S---TE---------L-------AGD--N--AV----------LST--A--SL------A-------Q--P--A-ID-A-VG----N----AA----A--L-----DVAKLL-Q-----T----E--H--H---------GDSL------------LAALQ-RGD-H---RALEAL---A---E----NS----EQLAQWLAGFRQV--L--SD--KQ----
PGSHFLAKQIYFPV-------GE---------------------GQ---YHLLSPLYSSSLSQALDQR-LN----L--A---R--F-G-----------------EQ-AKATR-E-----AR-------------R--QKRWH--DE-V-DVS-----YPS-IAVQNMGG-T-----KPQ----------NISALN--SARS-----------GRSYLLNSA-P-----PQWR--S---QI-KPPLDH----DTLF----RE-RGEVD--A------LT--YADRRDMRQFL------L--SV------K-----D-------------VEN---------
N----RE---IR--NQ----RR------RYLDQLIDTLFGYVASVQ--N-----L-R-P-----A---G----WS------K-----D------------SR--------LNT---PLQLWL-----D-------------PFR--CRQ-------------DDAFRY--ARE--GGD-WK-EQVAHEFGQWLN-----DR----L---Q-------H-------E-K-L-----------IF---GEVER------REWST-A-------AL--FK----------------Q-----RLRETERAL----------------KEEL-------A-----------------------
------------------
WP_038911898 ------------------------------------------------------------MS--S----------EGLQ--AFIL--S-YIS-NR-----L----QTK-------------------------------------------------------------------------------------------------L---------D-------A-FDK--AADK-----QR--A---------AL-----SG---------EALAEAEL---------------------------TL-----------------------------------A-QAR-RE-----------------------IE--
QA-------------YE---VRNW-----LT---DA-----ASRA-------G----QIS-L-----------------VTHALKFTHSD---A-----------------K---G-S------SV----F----S---TK---------P-------AAD--S--AV----------LST--A--TL------A-------Q--P--A-ID-A-VG----N----AA----A--L-----DVAKLL-Q-----T----E--H--H---------GDSL------------LAALQ-RGD-H---RALEAL---A---E----SP----QQLTQWLAGFGLV--L---TD--KQ----
PSSHKLAKQIYFPL-------GD---------------------GE---YHLLSPLYSSSLSQALDQR-LT----A--V---R--F-G-----------------EQ-AKAIR-E-----AR-------------R--EKRWH--GE-V-LVS-----YPG-VAVQNMGG-T-----KPQ----------NISALN--SARS-----------GRSYLLSSA-P-----PQWR--S---HI-KPPVEH----DTIF----RE-RGEVD--A------LT--YADRREMRQFL------L--SV------K-----E-------------VEN---------N-
---RE---IR--NQ----RR------RYLDQLIDTLFGYVATVQ--N-----L-R-P-----A---G----WS------V-----D------------SR--------LKR---HLQLWL-----D-------------PWR--GLQ-------------DKDFLR--ERD--AGD-WK-EQVAHEFGRWLN-----AR----L---Q-------H-------E-K-L-----------IF---GEVER------REWST-A-------AL--FK----------------Q-----RLRETERAL----------------KGEL-------A-----------------------
---------------
WP_038902018 ------------------------------------------------------------ME--E----------GELR--QFIV--S-YIE-NR-----K----QAK-------------------------------------------------------------------------------------------------L---------D-------A-FDK--AADK-----QR--A---------AL-----SG---------EALAEAEL---------------------------TL-----------------------------------A-QAR-RE-----------------------IE--
QA-------------YE---VRNW-----LT---DA-----ASRA-------G----QIS-L-----------------VTHALKFTHSD---A-----------------K---G-S------SV----F----S---TK---------P-------AAD--S--AV----------LST--A--SL------A-------Q--P--A-ID-A-VG----N----AA----A--L-----DVAKLL-Q-----T----E--H--H---------GDSL------------LAALQ-RGD-H---RALEAL---A---E----SP----QQLTQWLAGFGQV--L---TD--KQ----
PSSHKLAKQIYFPL-------GD---------------------GE---YHLLSPLYSSSLSQALDQR-LT----A--V---R--F-G-----------------EQ-AKAIR-E-----AR-------------R--ENRWH--DE-V-LVS-----YPG-VAVQNMGG-T-----KPQ----------NISALN--SARS-----------GRSYLLSSA-P-----PQWR--S---HI-KPPVEH----DTIF----RE-RGEVD--A------LT--YADRREMRQFL------L--SV------K-----E-------------VEN---------N-
---RE---IR--NQ----RR------RYLDQLIDTLFGYVATVQ--N-----L-R-P-----A---G----WS------V-----D------------SR--------LKR---HLQLWL-----D-------------PLR--GLQ-------------DTDFLR--ERD--AGD-WK-EQVAHEFGRWLN-----AR----L---Q-------H-------E-K-L-----------IF---GEVER------REWST-A-------TL--FK----------------Q-----RLRETERAL----------------KGEL-------A-----------------------
-------------
WP_039693736 ------------------------------------------------------------MS--S----------DGLQ--AFIL--S-YIE-NR-----K----QAK-------------------------------------------------------------------------------------------------L---------D-------A-FDK--AADK-----QR--A---------AL-----SG---------EALAAAEL---------------------------AL-----------------------------------A-QQR-RE-----------------------IE--
QT-------------YQ---VRTW-----LS---DA-----ASRA-------G----QIS-L-----------------VTHALKFTHSD---A-----------------K---G-S------SV----F----S---TE---------P-------AAD--S--AV----------LST--A--SL------A-------Q--P--A-ID-A-VG----N----AA----A--L-----DVAKLL-Q-----T----E--H--H---------GDSL------------LAALQ-RGD-H---RALEAL---A---E----SP----QQLAQWLAGFGQV--L---TD--KQ----
PSTHKLAKQIYFPV-------GD---------------------GQ---YHLLSPLYSSSLSQALDQR-LT----A--V---R--F-G-----------------EA-AKAIR-E-----AR-------------R--QKRWH--DE-V-DVS-----YPG-IAVQNMGG-T-----KPQ----------NISALN--SARS-----------GRSYLLSSA-P-----PQWR--S---QI-KPPLEH----DTIF----RE-RGEVD--A------LT--YVERREMRQFL------L--SV------K-----E-------------VEN---------N-
---RE---IR--NQ----RR------RYLDRLIDTLFGYVATVQ--N-----L-R-Q-----A---G----WS------K-----D------------SQ--------LNV---PLQLWL-----D-------------PFR--CRQ-------------DDAFRY--ARE--GGD-WK-EQVAHEFGRWLN-----AR----L---Q-------H-------E-K-L-----------IF---GEVER------REWST-A-------AL--FK----------------Q-----RLRENERLL----------------KEEL-------A-----------------------
---------------
WP_042860987 ------------------------------------------------------------MS--S----------EGLQ--AFIL--S-YIE-NR-----K----QAK-------------------------------------------------------------------------------------------------L---------D-------A-FDK--AADK-----QR--A---------AL-----SG---------EALAAAEL---------------------------ML-----------------------------------A-QQR-RE-----------------------IE--
QI-------------YE---VRTW-----LT---DA-----ASRA-------G----QIS-L-----------------VTHALKFTHSD---A-----------------K---G-S------SV----F----S---TE---------P-------AAD--G--AV----------LST--V--SL------A-------Q--P--A-ID-A-VG----N----AA----A--L-----DVAKLL-Q-----T----E--H--H---------GDSL------------VAALQ-RGD-H---RALEAL---A---E----SP----QQLTQWLAGFGQV--L--TD--KQ----
PSSHKLAKQIYFPV-------GD---------------------GE---YHLLSPLYSSSLSQALDQR-LT----A--V---R--F-G-----------------EA-AKAIR-E-----AR-------------R--QKRWH--DE-V-DVS-----YPG-IAVQNMGG-T-----KPQ----------NISALN--SARS-----------GRSYLLSSA-P-----PQWR--S---QL-KPPVEH----DTIF----RE-RGEVD--A------LT--YIERREMRQFL------L--SV------K-----E-------------VDN---------N-
---RE---IR--NQ----RR------RYLDQLIDTLFGYVATLQ--N-----L-R-P-----V---G----WS------V-----D------------SG--------LKR---HLQLWL-----D-------------PLR--GLQ-------------DKDFLR--ERD--AGD-WQ-REVAYEFGRWLN-----AR----L---Q-------H-------E-K-L-----------IF---GEVER------REWST-A-------AL--FK----------------Q-----RLRETERLL----------------KEEL-------A-----------------------
-------------
WP_121480586 ------------------------------------------------------------MS--S----------EGLQ--AFIL--S-YIE-TR-----K----QTK-------------------------------------------------------------------------------------------------L---------D-------A-FDK--AADK-----QR--A---------AL-----SG---------EAQAEAEL---------------------------TL-----------------------------------A-QQR-RE-----------------------IE--
QT-------------YQ---VSTW-----LT---DA-----ASRA-------G----QIS-L-----------------VTHALKFTHSD---A-----------------K---G-S------SV----F----S---TE---------P-------AAD--S--AV----------LST--A--SL------A-------Q--P--A-ID-A-VG----N----AA----A--L-----DVAKLL-Q-----T----E--Y--H---------GDSL------------LAALQ-RGD-H---RALEAL---A---E----SP----QQLIQWLAGFGQV--L--TD--KQ----
PSTHKLAKQIYFPV-------GE---------------------GQ---YHLLSPLYSSSLSQALDQR-LT----A--V---R--F-G-----------------EQ-AKAIR-E-----AR-------------R--QKRWH--DE-V-DVN-----YPG-IAVQNMGG-T-----KPQ----------NISALN--SARS-----------GRSYLLSSA-P-----PQWR--S---QI-KPPLDH----DTIF----RV-RGEVD--A------LT--YTARREMQQFL------L--SV------K-----E-------------VEN---------N-
---RD---IR--NQ----RR------RYLDQLIDTLFGYVATVQ--N-----L-R-P-----A---G----WS------V-----D------------SR--------LNV---PLQLWL-----D-------------PLR--GLQ-------------DKDFLR--ERD--AGD-WQ-REVAYEFGLWLN-----AR----L---Q-------H-------E-K-L-----------IF---GEVER------REWST-A-------AL--FK----------------Q-----RLRETERLL----------------KEEL-------A-----------------------
-------------
WP_038920151 ------------------------------------------------------------MS--S----------EGLQ--AFIL--S-YIE-TR-----K----QTK-------------------------------------------------------------------------------------------------L---------D-------A-FDK--AADK-----QR--A---------VL-----SG---------EAQAEAEL---------------------------TL-----------------------------------A-QQR-RE-----------------------IE--
QT-------------YQ---VRTW-----LS---DA-----ASRA-------G----QIS-L-----------------VTHALKFTHSD---A-----------------K---G-S------SV----F----S---TE---------P-------AAD--S--AV----------LST--A--SL------A-------Q--P--A-ID-A-VG----N----AA----A--L-----DVAKLL-Q-----T----E--Y--H---------GDSL------------LAALQ-RGD-H---RALEAL---A---E----SP----QQLTQWLAGFGQV--L--TD--KQ----
PSTHKLAKQIYFPV-------GE---------------------GQ---YHLLSPLYSSSLSQALDQR-LT----A--V---R--F-G-----------------EA-AKATR-E-----AR-------------R--QKRWH--DE-V-DVS-----YPG-IAVQNMGG-T-----KPQ----------NISALN--SARS-----------GRSYLLSSA-P-----PQWR--S---QL-KPPVEH----DTIF----RE-RGEVD--A------LT--YTERREMRQFL------L--SV------K-----E-------------VEN---------
N----RE---IR--NQ----RR------RYLDQLIDTLFGYVATVQ--N-----L-L-P-----A---G----WS------V-----D------------SR--------LNV---PLQLWL-----D-------------PLR--GLQ-------------DKDFLR--ERD--AGD-WQ-QDVAHEFGQWLN-----AR----L---K-------H-------E-K-L-----------IF---GEVER------REWST-A-------AL--FK----------------Q-----RLRENERLL----------------KEEL-------A-----------------------
---------------
WP_049855545 ------------------------------------------------------------MS--S----------EGLQ--AFIL--S-YIE-NR-----K----QAK-------------------------------------------------------------------------------------------------L---------D-------A-FDK--AADK-----QR--A---------AL-----SG---------EAQAEAEL---------------------------TL-----------------------------------A-QQR-RE-----------------------IE--
QT-------------YQ---VRTW-----LS---DA-----ASRA-------G----QIS-L-----------------VTHALKFTHSD---A-----------------K---G-S------SV----F----S---TE---------P-------AAD--S--AV----------LST--A--SL------A-------Q--P--A-ID-A-VG----N----AA----A--L-----DVAKLL-Q-----T----E--Y--H---------GDSL------------LAALQ-RGD-H---RALEAL---A---E----SP----QQLTQWLAGFGQV--L--TD--KQ----
PSTHKLAKQIYFPV-------GE---------------------GE---YHLLSPLYSSSLSQALDQR-LN----A--V---R--F-G-----------------EQ-AKATR-E-----AR-------------R--QKRWH--DE-V-DVS-----YPG-IAVQNMGG-T-----KPQ----------NISALN--SARS-----------GRSYLLSSA-P-----PQWR--S---QL-KPPVEH----DTIF----RE-RGEVD--A------LT--YVERREMRQFL------L--SV------K-----E-------------VEN---------
N----RE---IR--NQ----RR------RYLDQLIDTLFGYVATVQ--N-----L-R-P-----A---G----WS------V-----D------------SR--------LNV---PLQLWL-----D-------------PLR--GLQ-------------DKDFLR--ERD--AGD-WQ-REVAYEFGRWLN-----AR----L---Q-------H-------E-K-L-----------IF---GEVER------REWST-A-------AL--FK----------------Q-----RLRETERLL----------------KEEL-------A-----------------------
---------------
WP_038665951 ------------------------------------------------------------MS--S----------DGLQ--AFIL--S-YIE-NR-----K----QAK-------------------------------------------------------------------------------------------------L---------D-------A-FDK--AADK-----QR--A---------AL-----SG---------EALAAAEL---------------------------TL-----------------------------------A-QQR-RE-----------------------IE--
QI-------------YQ---VRTW-----LS---DA-----ASRA-------G----QIS-L-----------------VTHALKFTHSD---A-----------------K---G-S------SV----F----S---TE---------P-------AAD--S--AV----------LST--A--SL------A-------Q--P--A-ID-A-VG----N----AA----A--L-----DVAKLL-Q-----T----E--H--H---------GDSL------------LAALQ-RGD-H---RALEAL---A---E----SP----QQLAQWLAGFGQV--L--TD--KQ----
PSTHKLAKQIYFPV-------GE---------------------GQ---YHLLSPLYSSSLSQALDQR-LT----A--V---R--F-G-----------------EA-AKAIR-E-----AR-------------R--QKRWH--DE-V-DVS-----YPG-IAVQNMGG-T-----KPQ----------NISALN--SARS-----------GRSYLLSSA-P-----PQWR--S---QL-KPPVEH----DTIF----RE-RGEVD--A------LT--YAERREMRQFL------L--SV------K-----E-------------VEN---------
N----RE---IR--NQ----RR------RYLDQLIDTLFGYVATVQ--N-----L-R-P-----A---G----WS------V-----D------------SR--------LNV---PLQLWL-----D-------------PLR--GLQ-------------DKDFLR--ERD--AGD-WQ-REVAYEFGRWLN-----AR----L---Q-------H-------E-K-L-----------IF---GEVER------REWST-A-------AL--FK----------------Q-----RLRENERLL----------------KEEL-------A-----------------------
---------------
WP_035344067 ------------------------------------------------------------ME--E----------SELT--QFIV--S-YIE-TR-----K----QAK-------------------------------------------------------------------------------------------------L---------D-------A-FDK--AADK-----QR--A---------AL-----SG---------EALAAADV---------------------------AL-----------------------------------L-QQR-RE-----------------------IE--
QA-------------YE---VRTW-----LT---DA-----ASRA-------G----QIS-L-----------------VTHALKFTHSD---A-----------------K---G-S------SV----F----S---SE---------P-------ETA--T--DV----------LST--A--SL------A-------Q--P--A-ID-A-VG----N----AA----A--L-----DVAKLL-Q-----T----E--H--Q---------GDSL------------VAALQ-RGD-H---RALEAL---A---E----SP----QQLAQWLAGFGQV--L--SD--KQ----
PSSHKLAKQIYFPV-------GD---------------------GE---YHLLSPLYSSSLSQALDQR-LN----P--A---R--F-G-----------------EQ-AKAIR-E-----AR-------------R--QKRWH--DE-V-DVS-----YPG-IAVQNMGG-T-----KPQ----------NISALN--SARS-----------GRSYLLNCA-P-----PQWR--S---QI-KLPLEH----DTIF----RH-NGEVD--A------LT--YVDRRDMRQFL------L--SV------K-----E-------------VEN---------
N----RD---IR--NQ----RR------RYLDQLIDTLFGYVATVQ--N-----L-R-P-----A---G----WS------K-----D------------SR--------LNL---PLQLWL-----D-------------PFR--CRQ-------------DDAFRY--ARD--GGD-WK-EQVAHEFGQWLN-----DR----L---Q-------H-------K-K-L-----------IF---GEVER------REWST-A-------AL--FK----------------Q-----RLRETERAL----------------KEEL-------A-----------------------
-----------------
WP_102802311 ------------------------------------------------------------MS--S----------EGLQ--AFIL--S-YIE-TR-----K----QAK-------------------------------------------------------------------------------------------------L---------D-------A-FDK--AADK-----QR--A---------AL-----SG---------EALAAAEV---------------------------AL-----------------------------------L-QQR-RE-----------------------IE--
QA-------------YE---VRTW-----LT---DA-----ASRA-------G----QIS-L-----------------VTHALKFTHSD---A-----------------K---G-S------SV----F----S---TT---------P-------ATA--T--DV----------LST--A--SL------A-------Q--P--A-ID-A-VG----N----AA----A--L-----DVAKLL-Q-----T----E--H--Q---------GDSL------------VAALQ-RGD-H---RALEAL---A---E----SP----QQLTQWLAGFRQV--L--SD--KQ----
PSSHKLAKQIYFPV-------GD---------------------GE---YHLLSPLYSSSLSQALDQR-LN----P--A---R--F-G-----------------EQ-AKAIR-E-----AR-------------R--QKRWH--DE-V-DVS-----YPG-IAVQNMGG-T-----KPQ----------NISALN--SARS-----------GRSYLLNCA-P-----PQWR--S---QV-KPPLDS----DTIF----RN-NGEVD--A------LT--YVDRRDMRQFL------L--SV------K-----D-------------VEN---------
N----RD---IR--NQ----RR------RYLDQLIDTLFGYVATVQ--N-----L-R-P-----A---G----WS------K-----D------------SR--------LNV---PLQLWL-----D-------------PLR--CRQ-------------DDAFRY--ARE--GGD-WK-EQVAREFGLWLN-----AR----L---Q-------H-------E-K-L-----------IF---GEVER------REWST-A-------AL--FK----------------Q-----RLRETERAL----------------KEEL-------A-----------------------
-----------------
WP_038905566 ------------------------------------------------------------MS--S----------EGLQ--AFIL--S-YIE-TR-----K----QAK-------------------------------------------------------------------------------------------------L---------D-------A-FDK--AADK-----QR--A---------AL-----SG---------EELAAAEV---------------------------AL-----------------------------------L-QQR-RE-----------------------IE--
QA-------------YE---VRTW-----LT---DA-----ASRA-------G----QIS-L-----------------VTHALKFTHSD---A-----------------K---G-S------SV----F----S---TE---------P-------ATA--T--DV----------LST--A--SL------A-------Q--P--A-ID-A-VG----N----AA----A--L-----DVAKLL-Q-----T----E--H--Q---------GDSL------------VAALQ-RSD-H---RALEAL---A---E----NP----EQLTQWLAGFGQV--L--SD--KQ----
PSSHKLAKQIYFPV-------GD---------------------GE---YHLLSPLYSSSLSQALAQR-LN----L--A---R--F-G-----------------EQ-AKAIR-E-----AR-------------R--QKRWH--DE-V-DVS-----YPG-IAVQNMGG-T-----KPQ----------NISALN--SARS-----------GRSYLLNCA-P-----PQWR--S---QI-KLPLEH----DTIF----RN-NGEVD--A------LT--YVDRRDMRQFL------L--SV------K-----D-------------VEN---------N-
---RD---IR--NQ----RR------RYLDQLIDTLFGYVATVQ--N-----L-R-P-----A---G----WS------K-----D------------SR--------LNV---PLQLWL-----D-------------PFR--CRQ-------------DDAFCY--ARE--GGD-WK-EQVAREFGLWLN-----AR----L---Q-------H-------E-K-L-----------IF---GEVER------REWST-A-------AL--FK----------------Q-----RLRETERAL----------------KEEL-------A-----------------------
--------------
WP_019843462 ------------------------------------------------------------MSS------------EGLQ--AFIL--S-YIE-TR-----K----QAK-------------------------------------------------------------------------------------------------L---------D-------AFDKA-ADKQR-------------------AA---LSGE---------ALAAAEV---------------------------AL-----------------------------------L-QQR-RE
IEQA---------------YE---VRTW-----LT---DA-----ASRA-------G----QIS-L-----------------VTHALKFTHSD---A-----------------K---G-S------SV----F----S----------------TE---PAT--A-TDV----------LST--A--SL------A-------Q--P--A-ID-A-VG----N----AA----A--L-----DVAKLL-Q-----T----E--H--Q---------GDSL------------VAALQ-RGD-H---RALEAL---A---E----NA----EQLTQWLAGFGQV--L--SD--KQ----
PSSHKLAKQIYFPV-------GD---------------------GE---YHLLSPLYSSSLSQALDQR-LN----P--A---R--F-G-----------------EQ-AKAIR-E-----AR-------------R--QKRWH--DE-V-DVS-----YPG-IAVQNMGG-T-----KPQ----------NISALN--SARS-----------GRSYLLNCA-P-----PQWR--S---QI-KLPLEH----DTIF----RN-NGEVD--A------LT--YVDRRDMQQFL------L--SV------K-----E-------------VEN---------
N----RD---IR--NQ----RR------RYLDQLIDTLFGYVATVQ--N-----L-R-P-----A---G----WS------K-----D------------SR--------LNV---PLQLWL-----D-------------PFR--CRQ-------------DDAFCY--ARE--GGD-WK-EQVAREFGLWLN-----DR----L---Q-------H-------K-K-L-----------IF---GEVER------REWST-A-------AL--FK----------------Q-----RLRETERAL----------------KEEL-------A-----------------------
-----------------
WP_107760366 ------------------------------------------------------------MS--S----------EGLQ--AFIL--S-YIE-TR-----K----QAK-------------------------------------------------------------------------------------------------L---------D-------A-FDK--AADK-----QR--A---------AL-----SG---------EALAAAEV---------------------------TL-----------------------------------L-QQR-RE-----------------------IE--
QA-------------YE---VRTW-----LT---DA-----ASRA-------G----QIS-L-----------------VTHALKFTHSD---A-----------------K---G-S------SV----F----S---TE---------P-------ATA--T--DV----------LST--A--SL------A-------Q--P--A-ID-A-VG----N----AA----A--L-----DVAKLL-Q-----T----E--H--Q---------GDSL------------VAALQ-RGD-H---RALEAL---A---E----SP----QQLAQWLAGFGQV--L--TD--KQ----
PSSHKLAKQIYFPV-------GE---------------------GE---YHLLSPLYSSSLSQALDQR-LT----A--V---R--F-G-----------------EQ-AKATR-E-----AR-------------R--QKRWH--DE-V-DVS-----YPG-IAVQNMGG-T-----KPQ----------NISALN--SARS-----------GRSYLLNCT-P-----PQWR--S---QI-KPPLDH----DTIF----RN-NGEVD--A------LT--YVDRRDMRQFL------L--SV------K-----E-------------VEN---------
N----RD---IR--NQ----RR------RYLDQLIDTLFGYVATVQ--N-----L-R-P-----A---G----WS------K-----D------------SR--------LNV---PLQLWL-----D-------------PFR--CRQ-------------DDAFRY--ARE--GGD-WK-EQVAREFGLWLN-----AR----L---Q-------H-------E-K-L-----------IF---GEVER------REWST-A-------AL--FK----------------Q-----RLRETERAL----------------KEEL-------A-----------------------
-----------------
WP_023640697 ------------------------------------------------------------ME--E----------SELT--QFIV--S-YIE-TR-----K----QAK-------------------------------------------------------------------------------------------------L---------D-------A-FDK--AADK-----QR--A---------AL-----SG---------EALAAAEV---------------------------TL-----------------------------------L-QQR-RE-----------------------IE--
QA-------------YA---VRTW-----LT---DA-----ASRA-------G----QIS-L-----------------VTHALKFTHSD---A-----------------K---G-S------SV----F----S---TE---------P-------ATA--T--DV----------LST--A--SL------A-------Q--P--A-ID-A-VG----N----AA----A--L-----DVAKLL-Q-----T----E--H--Q---------GDSL------------VAALQ-RGD-H---RALEAL---A---E----NA----EQLTQWLAGFGQV--L--SD--KQ----
PSSHKLAKQIYFPV-------GD---------------------GE---YHLLSPLYSSSLSQALDQR-LN----P--A---R--F-G-----------------EQ-AKAIR-E-----AR-------------R--QKRWH--DE-V-DVS-----YPG-IAVQNMGG-T-----KPQ----------NISALN--SARS-----------GRSYLLNCA-P-----PQWR--S---QI-KPPLDS----DTIF----RN-NGEVD--A------LT--YVDRRDMRQFL------L--SV------K-----D-------------VEN---------
N----RD---IR--NQ----RR------RYLDQLIDTLFGYVATVQ--N-----L-R-P-----A---G----WS------K-----D------------SR--------LNV---PLQLWL-----D-------------PLR--CRQ-------------DDAFRY--ARE--GGD-WK-EQVAREFGLWLN-----DR----L---Q-------H-------K-K-L-----------IF---GEVER------REWST-A-------AL--FK----------------Q-----RLRETERAL----------------KEEL-------A-----------------------
-----------------
WP_110371414 ------------------------------------------------------------ME--E----------SELT--QFIV--S-YIE-TR-----K----QAK-------------------------------------------------------------------------------------------------L---------D-------A-FDK--AADK-----QR--A---------AL-----SG---------EALAAAEV---------------------------AL-----------------------------------L-QQR-RE-----------------------IE--
QA-------------YE---VRTW-----LT---DA-----ASRA-------G----QIS-L-----------------VTHALKFTHSD---A-----------------K---G-S------SV----F----S---TE---------P-------ATA--T--DV----------LST--A--SL------A-------Q--P--A-ID-A-VG----N----AA----A--L-----DVAKLL-Q-----T----E--H--Q---------GDSL------------VAALQ-RGD-H---RALKAL---A---E----NP----EQLTQWLAGFGQV--L--SD--KQ----
PSSHKLAKQIYFPV-------GD---------------------GE---YHLLSPLYSSSLSQALDQR-LN----A--V---R--F-G-----------------EQ-AKAIR-E-----AR-------------R--QKRWH--DE-V-DVS-----YPG-IAVQNMGG-T-----KPQ----------NISALN--SARS-----------GRSYLLNCA-P-----PQWL--S---QI-KPPLDS----DTIF----RN-NGEVD--A------LT--YVDRRDMRQFL------L--SV------K-----D-------------VEN---------
N----RD---IR--NQ----RR------RYLDQLIDTLFGYVVTVQ--N-----L-R-P-----A---G----WS------K-----D------------SR--------LNV---SLQLWL-----D-------------PFR--CRQ-------------DDAFCY--ARE--GGD-WK-EQVAREFGLWLN-----AR----L---Q-------H-------E-K-L-----------IF---GEVER------REWST-A-------AL--FK----------------Q-----RLRENERWL----------------KEEL-------A-----------------------
-----------------
WP_012886077 ------------------------------------------------------------MSS------------EGLQ--AFIL--S-YIE-TR-----K----QAK-------------------------------------------------------------------------------------------------L---------D-------AFDKA-ADKQR-------------------AA---LSGE---------ALAAAEV---------------------------TL-----------------------------------L-QQR-RE
IEQA---------------YE---VRTW-----LT---DA-----ASRA-------G----QIS-L-----------------VTHALKFTHSD---A-----------------K---G-S------SV----F----S----------------TE---PAT--A-TDV----------LST--A--SL------A-------Q--P--A-ID-A-VG----N----AA----A--L-----DVAKLL-Q-----T----E--H--Q---------GDSL------------VAALQ-RGD-H---RALEAL---A---E----NA----EQLTQWLAGFGQV--L--SD--KQ----
PSSHKLAKQIYFPV-------GD---------------------GE---YHLLSPLYSSSLSQALDQR-LN----A--V---R--F-G-----------------EQ-AKATR-E-----AR-------------R--QKRWH--DE-V-DVS-----YPG-IAVQNMGG-T-----KPQ----------NISALN--SARS-----------GRSYLLNCA-P-----PQWL--S---QI-KPPLDS----DTIF----RN-NGEVD--A------LT--YVDRRDMRQFL------L--SV------K-----D-------------VEN---------
N----RD---IR--NQ----RR------RYLDQLIDTLFGYVVTVQ--N-----L-R-P-----A---G----WS------K-----D------------SR--------LNV---PLQLWL-----D-------------PFR--CRQ-------------DDAFCY--ARE--GGD-WK-EQVAREFGLWLN-----AR----L---Q-------H-------E-K-L-----------IF---GEVER------REWST-A-------AL--FK----------------Q-----RLRENERWL----------------KEEL-------A-----------------------
-----------------
WP_123250920 ------------------------------------------------------------ME--E----------SELR--QFIV--S-YIE-NR-----K----QTK-------------------------------------------------------------------------------------------------L---------D-------A-FDK--TADK-----QR--A---------AL-----SG---------EALAAAEL---------------------------TL-----------------------------------L-QQR-RE-----------------------IE--
QT-------------YQ---VRTW-----LT---DA-----ASRA-------G----QIS-L-----------------VTHALKFTHSD---A-----------------K---G-S------SV----F----S---SE---------P-------VTD--S--AV----------LST--A--SL------A-------Q--P--A-ID-A-VG----N----AA----A--L-----DVAKLL-Q-----T----E--H--Q---------GDSL------------LAALQ-RGD-H---RALEAL---A---E----SP----QQLTQWLAGFGQV--L--TD--KQ----
PSSHKLAKQIYFPV-------GD---------------------GE---YHLLSPLYSSSLSQALDQR-LN----A--V---R--F-G-----------------EQ-AKAIR-E-----AR-------------R--QKRWH--DE-V-DVS-----YPG-IAVQNMGG-T-----KPQ----------NISALN--SARS-----------GRSYLLSSA-P-----PQWL--S---QI-KLPRDH----DTIF----RV-RGEVD--A------LT--YIARREMQQFL------L--SV------K-----E-------------VEN---------N--
--RE---IR--NQ----RR------RYLDRLINTLFDYVATVQ--N-----L-G-K-----A---G----WS------S-----Y------------SQ--------LNF---PLQLWL-----D-------------PRR--CRH-------------DDAFRY--ARD--GGD-WK-EQVAHEFGQWLN-----AR----L---H-------H-------D-K-L-----------IF---GEVER------REWST-A-------AL--FK----------------Q-----RLRESERWL----------------KEEL-------A-----------------------
--------------
WP_123252912 ------------------------------------------------------------MS--S----------EGLQ--AFIL--S-YIE-TR-----K----QTK-------------------------------------------------------------------------------------------------L---------D-------A-FDK--TADK-----QR--A---------AL-----SG---------EALAAAEL---------------------------TL-----------------------------------L-QQR-RE-----------------------IE--
QT-------------YQ---VCTW-----LT---DA-----ASRA-------G----QIS-L-----------------VTHALKFTHSD---A-----------------K---G-S------SV----F----S---SE---------P-------VTD--S--AV----------LST--A--SL------A-------Q--P--A-ID-A-VG----N----AA----A--L-----DVAKLL-Q-----T----E--H--Q---------GDSL------------LAALQ-RGD-H---RALEAL---A---E----SP----QQLTQWLAGFGQV--L--SD--KQ----
PSSHKLAKQIYFPV-------GD---------------------GE---YHLLSPLYSSSLSQALDQR-LN----A--V---R--F-G-----------------EQ-AKAIR-E-----AR-------------R--QKRWH--DE-V-DVS-----YPG-IAVQNMGG-T-----KPQ----------NISALN--SARS-----------GRSYLLSSA-P-----PQWL--S---QI-KLPREH----DTIF----RV-RGEVD--A------LT--YIARREMQQFL------L--SV------K-----E-------------VEN---------N--
--RE---IR--NQ----RR------RYLDRLINTLFDYVATVQ--N-----L-G-K-----A---G----WS------S-----Y------------SQ--------LNF---PLQLWL-----D-------------PRR--CRH-------------DDAFRY--ARD--GGD-WK-EQVAHEFGQWLN-----AR----L---H-------H-------D-K-L-----------IF---GEVER------REWST-A-------AL--FK----------------Q-----RMRENERWL----------------KEEL-------A-----------------------
--------------
WP_033569151 ------------------------------------------------------------MSS------------EGLQ--AFIL--S-YIE-NR-----K----QTK-------------------------------------------------------------------------------------------------L---------D-------AFDKT-ADKQR-------------------AA---LSGE---------ALAAAEL---------------------------TL-----------------------------------L-QQR-RE
IEQT---------------YQ---VRTW-----LT---DA-----ASRA-------G----QIS-L-----------------VTHALKFTHSD---A-----------------K---G-S------SV----F----S----------------SE---PVT--D-SAV----------LST--A--SL------A-------Q--P--A-ID-A-VG----N----AA----A--L-----DVAKLL-Q-----T----E--H--Q---------GDSL------------LAALQ-RGD-H---RALEAL---A---E----SP----QQLTQWLAGFGQV--L--TD--KQ----
PSSHKLAKQIYFPV-------GD---------------------GE---YHLLSPLYSSSLSQALDQR-LN----A--V---R--F-G-----------------EQ-AKAIR-E-----AR-------------R--QKRWH--DE-V-DVS-----YPG-IAVQNMGG-T-----KPQ----------NISALN--SARS-----------GRSYLLSSA-P-----PQWL--S---QI-KLPRDH----DTIF----RV-RGEVD--A------LT--YIARREMQQFL------L--SV------K-----E-------------VEN---------N--
--RE---IR--NQ----RR------RYLDRLINTLFDYVATVQ--N-----L-G-K-----A---G----WS------S-----Y------------SQ--------LNF---PLQLWL-----D-------------PRR--CRH-------------DDAFRY--ARD--GGD-WK-EQVAHEFGQWLN-----AR----L---H-------H-------D-K-L-----------IF---GEVER------REWST-A-------AL--FK----------------Q-----RLRESERWL----------------KEEL-------A-----------------------
--------------
WP_038908962 ------------------------------------------------------------MS--S----------EGLQ--AFIL--S-YIE-TR-----K----QAK-------------------------------------------------------------------------------------------------L---------D-------A-FDK--AADK-----QR--A---------AL-----SG---------EALAAAEV---------------------------AL-----------------------------------L-QQR-RE-----------------------IE--
RA-------------YE---VRTW-----LT---DA-----ASRA-------G----QIS-L-----------------VTHALKFTHSD---A-----------------K---G-S------SV----F----S---TE---------P-------ATA--T--DV----------LST--A--SL------A-------Q--P--A-ID-A-VG----N----AA----A--L-----DVAKLL-Q-----T----E--H--Q---------GDSL------------VAALQ-RGD-H---RALEAL---A---E----KP----EQLTQWLAGFGQV--L--SD--KQ----
PSSHKLAKQIYFPV-------GD---------------------GE---YHLLSPLYSSSLSQALDQR-LN----A--V---R--F-G-----------------EQ-AKAIR-E-----AR-------------Y--EKRWH--DG-V-DVS-----YPG-IAVQNMGG-T-----KPQ----------NISALN--SARS-----------GRSYLLSSA-P-----PQWL--S---QI-KLPLEH----DTIF----RV-RGEVD--A------LT--YIARRDMRQFL------L--SV------K-----E-------------VEN---------N--
-RD---IR--NQ----RR------RYLDRLINTLFDYVATVQ--N-----L-G-K-----A---G----WS------T-----Y------------SQ--------LNF---PLQLWL-----D-------------PRR--CQQ-------------DDAFRY--ARE--GGD-WK-EQVAHEFGQWLN-----DR----L---H-------H-------D-K-L-----------IF---GEMER------REWST-A-------AL--FK----------------Q-----RLRENERWL----------------KEEL-------A-----------------------
--------------
WP_039999707 ------------------------------------------------------------ME--E----------SELT--QFIV--S-YIE-TR-----K----QAK-------------------------------------------------------------------------------------------------L---------D-------A-FDK--AADK-----QR--A---------AL-----SG---------EALAAAEV---------------------------AL-----------------------------------L-QQR-RE-----------------------IE--
QA-------------YE---ARTW-----LT---DA-----ASRA-------G----QIS-L-----------------VTHALKFTHSD---A-----------------K---G-S------SV----F----S---TE---------P-------ETA--T--DV----------LST--A--SL------A-------Q--P--A-ID-A-VG----N----AA----A--L-----DVAKLL-Q-----T----E--H--Q---------GDSL------------VAALQ-RGD-H---RALEAL---A---E----NA----EQLAQWLAGFGQV--L--SD--KQ----
PSSHKLAKQIYFPV-------GD---------------------GE---YHLLSPLYSSSLSQALDQR-LN----A--V---R--F-A-----------------EQ-AKAIR-E-----AR-------------R--QKRWH--DE-V-DVS-----YPG-IAVQNMGG-T-----KPQ----------NISALN--SARS-----------GRSYLLSSA-P-----SQWL--S---QI-KLPLEH----DTIF----RE-RGEVD--A------LT--YTARREMQQFL------L--SV------K-----E-------------VEN---------N--
--RE---IR--NQ----RR------RYLDRLINALFDYVATVQ--N-----L-G-K-----A---G----WS------T-----Y------------SQ--------LNF---PLRLWL-----D-------------PRR--CRH-------------DDAFRY--ARD--GGD-WK-EQVAHEFGQWLN-----AR----L---H-------H-------D-K-L-----------IF---GEVER------REWST-A-------AL--FK----------------Q-----RLRETERAL----------------KEER-------A-----------------------
--------------
WP_143839526 ------------------------------------------------------------ME--E----------SELT--QFIV--S-YIE-TR-----K----QAK-------------------------------------------------------------------------------------------------L---------D-------A-FDK--AADK-----QR--A---------AL-----SG---------EALAAAEV---------------------------AL-----------------------------------L-QQR-RE-----------------------IE--
QA-------------YE---VRTW-----LT---DA-----ASRA-------G----QIS-L-----------------VTHALKFTHSD---A-----------------K---G-S------SV----F----S---TE---------P-------ATS--T--DV----------LST--A--SL------A-------Q--P--A-ID-A-VG----N----AA----A--L-----DVAKLL-Q-----T----E--Y--Q---------GDSL------------VAALQ-RGD-H---RSLEAL---A---E----NT----EQLAQWLAGFKQV--L--SD--KQ----
PSSHKLAKQIYFPV-------GE---------------------EE---YHLLSPLYSSSLSQALDQR-LN----A--V---R--F-G-----------------EQ-AKAIR-E-----AR-------------Q--QKRWH--DE-I-YVS-----YPG-IAVQNMGG-T-----KPQ----------NISALN--SARS-----------GRSYLLSSA-P-----SQWL--S---QI-KLPLEH----DTIF----RE-RGEVD--A------LT--YTARRELQQFL------L--SV------K-----E-------------VEN---------N--
RE---IR--NQ----RR------RYLDRLINALFDYVATVQ--N-----L-G-K-----A---G----WS------K-----D------------SR--------LNV---PLQLWL-----D-------------PRR--CRH-------------DDAFRY--ARD--GGD-WK-EQVAHEFGQWLN-----AR----L---R-------H-------D-K-L-----------IF---GEVER------REWST-A-------AL--FK----------------Q-----RLRETERAL----------------KEEL-------A-----------------------
------------
WP_027710957 ------------------------------------------------------------ME--E----------SELT--QFIV--S-YIE-TR-----K----QAK-------------------------------------------------------------------------------------------------L---------D-------A-FDK--AADK-----QR--A---------AL-----SG---------EALAAAEL---------------------------TL-----------------------------------L-QQR-RE-----------------------IE--
QA-------------YE---VRTW-----LT---DA-----ASRA-------G----QIS-L-----------------VTHALKFTHSD---A-----------------K---G-S------SV----F----S---NE---------P-------ATA--T--DV----------LST--A--SL------A-------Q--P--A-ID-A-VG----N----AA----A--L-----DVAKLL-Q-----T----E--H--Q---------GDSL------------VAALQ-RGD-H---RALEAL---A---E----NA----EQLTQWLAGFGQV--L--SD--KQ----
PSSHKLAKQIYFPV-------GE---------------------EE---YHLLSPLYSSSLSQALDQR-LN----A--V---R--F-G-----------------EQ-AKAIR-E-----AR-------------Q--QKRWH--DE-I-YVS-----YPG-IAVQNMGG-T-----KPQ----------NISALN--SARS-----------GRSYLLSSA-P-----SQWL--S---QI-KLPLEH----DTIF----RE-RGEVD--A------LT--YTARRELQQFL------L--SV------K-----E-------------VEN---------N--
RE---IR--NQ----RR------RYLDRLINTLFDYVATVQ--N-----L-G-K-----A---G----WS------T-----Y------------SQ--------LNV---PLQLWL-----D-------------PRR--CRH-------------DDAFRY--ARD--GGD-WK-EQVAHEFGQWLN-----DR----L---Q-------H-------K-K-L-----------IF---GEVER------REWST-A-------AL--FK----------------Q-----RLRETERAL----------------KEEL-------A-----------------------
------------
WP_125260582 ------------------------------------------------------------MS--S----------EGLQ--AFIL--S-YIE-TR-----K----QAK-------------------------------------------------------------------------------------------------L---------D-------A-FDK--AADK-----QR--A---------AL-----SG---------EALAAAEV---------------------------AL-----------------------------------L-QQR-RE-----------------------IE--
QA-------------YE---VRTW-----LT---DA-----ASRA-------G----QIS-L-----------------VTHALKFTHSD---A-----------------K---G-S------SV----F----S---NE---------P-------ATA--T--DV----------LST--A--SL------A-------Q--P--A-ID-A-VG----N----AA----A--L-----DVAKLL-Q-----T----E--H--Q---------GDSL------------VAALQ-RGD-H---RALQAL---A---E----NA----EQLAQWLAGFGQV--L--SD--KQ----
PSSHKLAKQIYFPV-------GD---------------------GE---YHLLSPLYSSSLSQALDQR-LN----A--V---R--F-G-----------------EQ-AKAIR-E-----AR-------------Y--EKRWH--DE-I-YVS-----YPG-IAVQNMGG-T-----KPQ----------NISALN--SARS-----------GRSYLLSSA-P-----SQWL--S---QI-KLPLEH----DTIF----RE-RGEVD--A------LT--YTARRELQQFL------L--SV------K-----E-------------VDN---------N--
RD---IR--NQ----RR------RYLDRLINALFDYVATVQ--N-----L-G-K-----A---G----WS------T-----Y------------SQ--------LNV---PLQLWL-----D-------------PRR--CQH-------------DDAFRY--ARE--GGD-WK-EQVAHEFGQWLN-----DR----L---Q-------H-------K-K-L-----------IF---GEVER------REWST-A-------AL--FK----------------Q-----RLRETERAL----------------KEEL-------A-----------------------
------------
WP_033577739 ------------------------------------------------------------MS--S----------EGLQ--AFIL--S-YIE-TR-----K----QAK-------------------------------------------------------------------------------------------------L---------D-------A-FDK--AADK-----QR--A---------AL-----SG---------EALAAAEV---------------------------AL-----------------------------------L-QQR-RE-----------------------IE--
QA-------------YE---VRTW-----LT---DA-----ASRA-------G----QIS-L-----------------VTHALKFTHSD---A-----------------K---G-S------SV----F----S---TK---------P-------ATA--A--DV----------LST--A--SL------A-------Q--P--A-ID-A-VG----N----AA----A--L-----DVAKLL-Q-----T----E--H--Q---------GDSL------------VAALQ-RGD-H---RSLEAL---A---E----NA----EQLAQWLAGFGQV--L--SD--KQ----
PSSHKLAKQIYFPV-------GE---------------------EE---YHLLSPLYSSSLSQALDQR-LN----A--V---R--F-G-----------------EQ-AKAIR-E-----AR-------------Q--QKRWH--DE-I-YVS-----YPG-IAVQNMGG-T-----KPQ----------NISALN--SARS-----------GRSYLLSSA-P-----SQWL--S---QI-KLPLEH----DTIF----RE-RGEVD--A------LT--YTARRELQQFL------L--SV------K-----E-------------VEN---------N--
RE---IR--NQ----RR------RYLDRLINALFDYVATVQ--N-----L-G-K-----A---G----WS------T-----Y------------SQ--------LNV---PLRLWL-----D-------------PRR--CRH-------------DDAFRY--ARD--GGD-WK-EQVAHEFGQWLN-----AR----L---R-------H-------D-K-L-----------IF---GEVER------REWST-A-------AL--FK----------------Q-----RLRETERAL----------------KEDL-------A-----------------------
------------
WP_012768459 ------------------------------------------------------------MSS------------EGLQ--AFIL--S-YIE-TR-----K----QAK-------------------------------------------------------------------------------------------------L---------D-------AFDKA-ADKQR-------------------AA---LSGE---------ALAAAEV---------------------------AL-----------------------------------L-QQR-RE
IEQA---------------YE---VRTW-----LT---DA-----ASRA-------G----QIS-L-----------------VTHALKFTHSD---A-----------------K---G-S------SV----F----S----------------AK---TAT--A-TDV----------LST--A--SL------A-------Q--P--A-ID-A-VG----N----AA----A--L-----DVAKLL-Q-----T----E--H--Q---------GDSL------------VAALQ-RGD-H---RSLEAL---A---E----NT----EQLAQWLAGFGQV--L--SD--KQ----
PSSHKLAKQIYFPV-------GE---------------------EE---YHLLSPLYSSSLSQALDQR-LN----A--V---R--F-G-----------------EQ-AKAIR-E-----AR-------------Q--QKRWH--DE-I-YVS-----YPG-IAVQNMGG-T-----KPQ----------NISALN--SARS-----------GRSYLLSSA-P-----SQWL--S---QI-KLPLEH----DTIF----RE-RGEVD--A------LT--YTARRELQQFL------L--SV------K-----E-------------VEN---------N--
RE---IR--NQ----RR------RYLDRLINALFDYVATVQ--N-----L-G-K-----A---G----WS------T-----Y------------SQ--------LNF---PLQLWL-----D-------------PRR--CQQ-------------DDAFRY--ARE--GGD-WK-ERVAHEFGQWLN-----AR----L---R-------H-------D-K-L-----------IF---GEVER------REWST-A-------AL--FK----------------Q-----RLRETERAL----------------KEEL-------A-----------------------
------------
WP_075278342 ------------------------------------------------------------ME--E----------NRLT--QFIV--S-YIN-TR-----K----QAK-------------------------------------------------------------------------------------------------L---------D-------A-FDK--EAEK-----KR--S---------TL-----SG---------EALAAEEL---------------------------EL-----------------------------------A-EKR-RE-----------------------IE--
QK-------------HE---VRAW-----LT---DA-----STQA-------H----QIK-Q-----------------ATHILKFTHSS---A-----------------K---QG-S------SV----L----T---LK---------K-------NKC--P--LF----------LST--D--AI------A-------A--P--V-VDTA-IH----D----AK----V--L-----AVAKLL-T-----E----E--S--H---------GESL------------VSALQ-RGD-H---SALEAL---A---E----NA----EQLAQWLTGFQQI--F--SD--KQ----
PSSHSLAKQIYFPI-----------------------------ANGE---YHLLSPLYSSSLAHALYQR-IN----P--I--H--F-G-----------------EE-AKTIR-K-----AR-------------K--DNLWH--DQ-L-DIS-----YPN-LAVQSMGG-T-----KPQ----------NISSLN--VTRN-----------GRSYLLNCT-P-----PQWN--S---IE-KPPQQH----DSIF----RS-NGEVD--Y------HT--RDIVRPMRHFL------L--SV------K-----K-------------VEN---------N--
LG---IR--QQ----RQ------RYLNQLIEQLFFYVTSVQ--N-----L--P-----E---G----WS------T-----E------------SE--------LKR---TQQLWL-----D-------------PYR--AET-------------DTVFRR--ERE--AGD-WQ-KAVAYDFGRWLN-----LR----L---K-------H-------E-E-L-----------IF---GEVER------REWST-A-------AL--FK----------------R-----RMREMENAL----------------KEDL-------A-----------------------
----------
WP_021017051 ------------------------------------------------------------MK--E----------NTLT--HFIL--S-YIE-KR-----R----QAK-------------------------------------------------------------------------------------------------L---------D-------A-FDK--EAEK-----KR--S---------PL-----SG---------EALAVAGL---------------------------EL-----------------------------------A-QAR-QV-----------------------IE--
QR-------------HE---VRTW-----LT---DA-----ASRA-------G----QIS-L-----------------VTHALKFTHSD---A-----------------K---G-T------SV----F----S---PE---------A-------VKN--A--TS----------LST--A--TL------T-------Q--P--A-ID-A-VG----N----AA----A--L-----DVAKLL-Q-----T----E--Y--E---------GDSL------------VAALQ-RHD-Y---SALTAL---A---E----SP----EQLAQWLAGFQQV--F--TV--RQ----
PSSHKLAKQIYFPI-----------------------------GKGE---YHLLSPLYASSLAQALYQR-II----A--V---R--F-G-----------------DE-AKAIR-L-----AQ-------------K--ENRWH--DQ-L-SIS-----YPN-LAVQNMGG-T-----KPQ----------NISALN--STRS-----------GRSYLLSCA-P-----PQWQ--G---EE-KPPQQH----QSIF----RE-RGEVD--Y------HT--RAIIPLMRQFL------L--SV------K-----E-------------VEN---------N--
RE---IR--RL----RL------RYLDQLIDQLFCYVASIQ--N-----L--R-----S---G----WS------A-----D------------SK--------LSR---IQQLWL-----D-------------PHR--SNE-------------DAVFRR--ERE--AGD-WQ-KDVAYDFGRWLN-----RR----L---Q-------H-------E-K-L-----------IF---GEAER------REWST-A-------VL--FK----------------R-----RMREMESAL----------------KEEL-------A-----------------------
---------
WP_109052523 ------------------------------------------------------------ME--E----------KTLT--QFIV--S-YIE-NR-----R----QAK-------------------------------------------------------------------------------------------------L---------D-------A-HDK--EAEK-----KR--S---------GL-----SG---------DALAAAES---------------------------EL-----------------------------------V-QKR-RE-----------------------IE--
QR-------------HE---VRAW-----LT---DA-----AGRA-------G----QIS-L-----------------VTHALKFTHSD---A-----------------K---G-S------SI----F----S---AE---------T-------VTD--A--KT----------LST--A--TL------T-------K--S--T-ID-A-VG----N----AA----A--L-----DVAKLL-Q-----T----E--H--Q---------GDSL------------VAALQ-RND-H---SALAAL---A---E----NP----EQLAQWLAGFQQV--F--TD--RQ----
PSSHKLAKQIYFPI-----------------------------GNGE---YHLLSPLYPSSLAQALYQR-IN----A--V---R--F-G-----------------DE-AKAIR-Q-----AQ-------------K--ENQWH--DQ-L-SIS-----YPN-LAVQNMGG-T-----KPQ----------NISALN--STRS-----------GRSYLLSCA-P-----PQWR--S---VE-QPPQRH----KSIF----RE-RGEAD--Y------YT--RTTLTRMRQFL------L--SV------K-----T-------------VEN---------N--
--RD---IR--HQ----RL------RYLDQVIDQLFFYVASIQ--N-----L--S-----S---S----WS------A-----E------------SE--------LNR---AQQLWL-----N-------------PHR--CVD-------------DAAFRH--ERE--SGD-WQ-KEVAHDFGRWLN-----LR----L---K-------H-------D-E-L-----------IF---GEVER------REWST-A-------AL--FK----------------Q-----RMREMESIL----------------KEEF-------A-----------------------
-----------
WP_136155399 ------------------------------------------------------------M--K-----------DGLT--AFIV--S-YIE-NR-----R----QAK-------------------------------------------------------------------------------------------------L---------D-------A-HDK--EAEK-----KR--S---------EL-----SG---------DALAAAES---------------------------EL-----------------------------------V-QKR-RE-----------------------IE--
QR-------------HE---VRAW-----LT---DA-----ASRA-------G----QIS-L-----------------VTHALKFTHSD---A-----------------K---G-S------SI----F----S---AE---------T-------VID--A--KT----------LST--A--TL------T-------K--S--T-ID-A-VG----N----AA----A--L-----DVAKLL-Q-----T----E--H--Q---------GDSL------------VAALQ-RND-H---SALAAL---A---E----NP----EQLAQWLAGFQQV--F--TD--RQ----
PSSHKLAKQIYFPI-----------------------------GNGE---YHLLSPLYPSSLAQALYQR-IN----A--V---R--F-G-----------------DE-AKAIR-Q-----AQ-------------K--ENQWH--DQ-L-SIS-----YPN-LAVQNMGG-T-----KPQ----------NISSLN--STRS-----------GRSYLLSCA-P-----PQWR--S---VE-QPPQRH----KSIF----RE-RGEAD--Y------YT--RTTLIRMRQFL------L--SV------K-----T-------------VEN---------N--
-RN---IR--HQ----RL------RYLDQVIDQLFFYVASVQ--N-----L--S-----S---S----WS------A-----E------------SE--------LNR---AQQLWL-----N-------------PHR--CVD-------------DAAFRH--ERE--SGD-WQ-KEVAHDFGHWLN-----RR----L---K-------H-------D-E-L-----------IF---GEVER------REWST-A-------AL--FK----------------Q-----RMREMESIL----------------KEEL-------A-----------------------
-----------
WP_116238045 ------------------------------------------------------------ME--D----------NRLT--PFIV--S-YIN-NR-----K----QAK-------------------------------------------------------------------------------------------------L---------D-------E-FDK--KTKK-----KL--A---------AL-----SG---------EALAVAEL---------------------------ER-----------------------------------A-QAR-RE-----------------------IE--
QK-------------YD---VRNW-----LT---DA-----ASRA-------W----QIS-L-----------------VTHALKYTHSD---A-----------------K---G-S------SI----F----S---AE---------A-------VAD--A--KT----------LST--A--TL------A-------Q--P--V-ID-V-VG----N----AA----V--L-----DVAGLL-Q-----T----E--H--E---------GDSL------------VAALQ-RGD-H---RALEAL---A---E----SP----EQLAQWLAGFQQV--F--TD--RQ----
PSSHKLAKQIYFPT-----------------------------ENGE---YHLLSPLYSSSLAQALHQR-IN----A--V---R--F-G-----------------DE-AKAIS-Q-----AQ-------------K--ANQWH--DQ-L-DIR-----YPN-LAVQNRGG-S-----NKQ----------NISSLN--VTRN-----------GRSYLLSSA-P-----PQWN--N---IE-KPPQQH----ESIF----RS-KGEVN--Y------HT--SDIVESMRSFL------L--SV------K-----E-------------KQN---------N--
-RD---IR--QQ----RL------RYLDQLIDQLFFYVASVQ--N-----L--P-----A---G----WS------A-----E------------SE--------LKR---AQQLWL-----D-------------PCR--AKT-------------DAVFRR--ERE--AGD-WQ-QVVAYDFGRWLN-----LR----L---K-------H-------E-E-L-----------IF---GEVER------REWST-A-------AL--FK----------------R-----RMREMESSL----------------KEEL-------A-----------------------
-----------
WP_005974986 ------------------------------------------------------------MED------------NRLT--KFIV--S-YIE-ER-----R----QDK-------------------------------------------------------------------------------------------------L---------D-------EHDKK-VKTAR-----TE--L---------SS-------------DALADAEL---------------------------TL-----------------------------------A-EKR-RE-----------------------IE--
-----RK------YE---VRNW-----LT---DA-----ASRA-------G----QIS-L-----------------VTHALKFTHSD---A-----------------R---G-S------SI----F----S----------------AN---TVA--D-AKT----------LST--A--TL------V-------Q--P--A-ID-A-VG----N----AA----A--L-----DVAKLL-Q-----T----E--Y--E---------GDSL------------VAALQ-RGD-H---RALEAL---A---E----SP----EQLAQWLAGFQQV--F--TD--RQ----
PSSHKLSKQIYFPL-----------------------------ANGE---YHLLSPLYSSSLAQALYQR-IT----A--V---R--F-G-----------------DE-AKAIR-R-----AQ-------------K--VSQWH--DQ-L-SIS-----YPN-LAVQNMGG-T-----KPQ----------NISALN--SSRS-----------GRSYLLSSA-P-----PQWN--H---IE-KPPQQH----ESIF----RP-RGEVD--Y------HT--RATLAQMQRFL------L--SV------K-----D-------------VEN---------N--
--RD---IR--QQ----RL------RYLDQLIDQLFFYVASVQ--N-----L--P-----V---G----WS------A-----E------------SE--------LKR---AQQLWL-----D-------------PYR--ADT-------------DTVFRR--ERE--AGD-WQ-QAVAYDFGRWLN-----RR----L---K-------H-------E-N-L-----------IF---GEVER------REWST-A-------AL--FK----------------R-----RMREVESAL----------------KEEL-------A-----------------------
-----------
WP_100018204 ------------------------------------------------------------MGS------------EKLT--EFIL--S-YIN-NR-----K----QTK-------------------------------------------------------------------------------------------------L---------D-------A-FDK--EAEK-----KR--V---------DL-----SG---------EALAAEEL---------------------------EL-----------------------------------A-EKR-RE
IEKK---------------HE---VRAW-----LT---DA-----ASRA-------G----QIS-L-----------------VTHALKYTHSY---A-----------------K---G-S------SV----F----S---D---------K-T-----LTD--A--KT----------LST--A--TL------T-------Q--P--A-ID-V-VG----N----AA----V--L-----DVAGLL-Q-----A----E--H--K---------GDSL------------VAALQ-RGD-H---RALEAL---A---E----NS----EQLAKWLTGFQQV--F--TD--RQ----
PSSHKLAKQIYFPI-----------------------------ANGE---YHLLSPLYSSSLAHALHQR-IN----A--V---R--F-G-----------------DE-AKAIR-Q-----AQ-------------K--ANQWH--DQ-R-SIS-----YPY-LAVQNMGG-S-----NKQ----------NISSLN--VTRN-----------GRSYLLNCA-P-----PQWN--S---IE-KPPQQH----ECIF----RL-NGEMD--Y------HT--RDIVRPMRHFL------L--SV------K-----D-------------VEN---------N--
--RD---IR--HQ----RL------RYLDQLIDRLFFYVASVQ--N-----L--P-----S---G----WS------A-----Q------------SE--------LKH---AQQLWL-----D-------------PYR--AKT-------------DTVFRH--ERE--AGD-WQ-KAVAYDFGRWLN-----RR----L---K-------H-------E-D-L-----------VF---GEVER------REWST-A-------AL--FK----------------R-----RMREMESAL----------------KEDL-------A-----------------------
--------------
WP_039502076 ------------------------------------------------------------MED------------SRLT--KFIV--S-YIE-ER-----R----QDK-------------------------------------------------------------------------------------------------L---------D-------KHDKT-VKTAR-------------------TE---LSGD---------ALADAEL---------------------------TL-----------------------------------A-EKR-RE-----------------------IE--
KK-------------HE---VRAW-----LT---DA-----ASRA-------G----QIS-L-----------------VTHALKYTHSY---A-----------------K---G-S------SV----F----S----------------DK---TLT--D-AKT----------LST--A--TL------T-------Q--P--A-ID-V-VG----N----AA----V--L-----DVAGLL-Q-----A----E--H--K---------GDSL------------VAALQ-RGD-H---RALEAL---A---E----NS----EQLAKWLTGFQQV--F--TD--RQ----
PSSHQLAKQIYFPI-----------------------------ANGE---YHLLSPLYSSSLAHALHQR-IN----A--V---R--F-G-----------------DE-AKAIR-Q-----AQ-------------K--ANQWH--DQ-R-SIS-----YPY-LAVQNMGG-S-----NKQ----------NISSLN--VTRN-----------GRSYLLNCA-P-----PQWN--S---IE-KPPQQH----DSIF----RP-RGEVD--Y------HT--RATLAQMQRFL------L--SV------K-----E-------------VEN---------N-
---RD---IR--QQ----RL------RYLDQLIDQLFFYVASVQ--N-----L--P-----S---G----WS------A-----E------------SE--------LKR---AQQLWL-----D-------------PYR--AET-------------DTVFRR--ERE--AGD-WQ-KAVAYDFGRWLN-----RR----L---K-------H-------E-D-L-----------VF---GEVER------REWST-A-------AL--FK----------------R-----RMREMESAL----------------KEDL-------A-----------------------
-------------
WP_094370082 ------------------------------------------------------------MED------------SRLT--KFIV--S-YIE-ER-----R----QDK-------------------------------------------------------------------------------------------------L---------D-------KHDKT-VKTAR-------------------TE---LSGD---------ALADAEL---------------------------TL-----------------------------------A-EKR-RE-----------------------IE---
-----KK------HE---VRAW-----LT---DA-----ASRA-------G----QIS-L-----------------VTHALKYTHSY---A-----------------K---G-S------SV----F----S----------------DK---TLT--D-AKT----------LST--A--TL------T-------Q--P--A-ID-V-VG----N----AA----V--L-----DVAGLL-Q-----A----E--H--K---------GDSL------------VAALQ-RGD-H---RALEAL---A---E----NS----EQLAKWLTGFQQV--F--TD--RQ----
PSSHQLAKQIYFPI-----------------------------ANGE---YHLLSPLYSSSLAHALHQR-IN----A--V---R--F-G-----------------DE-AKAIR-Q-----AQ-------------K--ANQWH--DQ-R-SIS-----YPY-LAVQNMGG-S-----NKQ----------NISSLN--VTRN-----------GRSYLLNCA-P-----PQWN--S---IE-KPPQQH----ECIF----RL-NGEMD--Y------HT--RDIVRPMRHFL------L--SV------K-----D-------------VEN---------N--
--RD---IR--HQ----RL------RYLDQLIDRLFFYVASVQ--N-----L--P-----S---G----WS------A-----Q------------SE--------LKH---AQQLWL-----D-------------PYR--AKT-------------DTVFRH--ERE--AGD-WQ-KAVAYDFGRWLN-----RR----L---K-------H-------E-E-L-----------IF---GEVER------REWST-A-------AL--FK----------------R-----RMREMESAL----------------KEDL-------A-----------------------
------------
WP_129712160 ------------------------------------------------------------ME--E----------NRLT--QFIV--S-YVT-NR-----K----QAK-------------------------------------------------------------------------------------------------L---------D-------A-FDK--EAEK-----KR--A---------AL-----SD---------EVLAVAEQ---------------------------EL-----------------------------------A-QAR-QE-----------------------IE--
QK-------------HE---VRNW-----LT---DA-----ASRA-------G----QIS-L-----------------VTHAVKFTHSD---A-----------------K---G-N------SV----F----N---AE---------T-------VAD--A--KT----------LST--A--TL------A-------Q--P--A-ID-A-VG----N----AA----A--L-----DVAKLL-Q-----T----E--H--D---------GDSL------------VAALQ-RGD-H---RALEAL---A---E----NP----EQLAPWLIGFQQV--F--TN--RQ----
PSSHKLAKQIYFPL-----------------------------ASGE---YHLLSPLYSSSLAQALHQR-IN----A--M---R--F-G-----------------DE-AKAIR-Q-----AQ-------------R--TNQWH--DQ-L-SIS-----YPN-LAVQNMGG-T-----KPQ----------NISALN--SSRS-----------GRSYLLSSA-P-----PQWN--S---IE-KPPQQH----ECIF----RL-NGEVD--Y------RT--RDIVRPMRNFL------L--SV------K-----N-------------VEN---------N--
--RD---IR--QQ----RL------RYLDQLIDQLFFYVASVQ--N-----L--P-----S---G----WS------A-----E------------SE--------LKR---AQQLWL-----D-------------PYR--AET-------------DTVFRR--ERE--AED-WQ-QAIAYDFGRWLN-----RR----L---K-------H-------E-N-L-----------IF---GEVER------REWST-A-------AL--FK----------------R-----RMREMESAL----------------KEEL-------A-----------------------
-----------
WP_129706591 ------------------------------------------------------------MS--G----------EKLK--AFIL--S-YIS-TR-----K----QAK-------------------------------------------------------------------------------------------------L---------D-------A-FDK--EAEK-----KR--A---------AL-----SD---------EVLAVAER---------------------------EL-----------------------------------A-QAR-LE-----------------------IE--
QK-------------HE---VRNW-----LT---DA-----ASRA-------G----QIS-L-----------------VTHAVKFTHSD---A-----------------K---G-S------SV----F----N---AE---------T-------VAD--A--KT----------LST--A--TL------A-------Q--P--A-ID-A-VG----N----AA----A--L-----DVAKLL-Q-----T----E--H--D---------GDSL------------VAALQ-RGD-H---RALEAL---A---E----NP----EQLAQWLIGFQQV--F--TN--RQ----
PSSHKLAKQIYFPL-----------------------------ASGE---YHLLSPLYSSSLAQALHQR-IN----A--V---R--F-G-----------------DE-AKAIR-Q-----AQ-------------R--TNQWH--DQ-L-SIS-----YPN-LAVQNMGG-T-----KPQ----------NISALN--SSRS-----------GRSYLLSSA-P-----PQWN--S---IE-KPPQQH----ECIF----RL-NGEVD--Y------RT--RDIVRPMRNFL------L--SV------K-----N-------------VEN---------N--
-RD---IR--QQ----RL------RYLDQLIDQLFFYVASVQ--N-----L--P-----S---G----WS------A-----E------------SE--------LKR---AQQLWL-----D-------------PYR--AET-------------DTVFRR--ERE--AED-WQ-QAIAYDFGRWLN-----RR----L---K-------H-------E-N-L-----------IF---GEVER------REWST-A-------AL--FK----------------R-----RMREMESAL----------------KEEL-------A-----------------------
----------
WP_039324337 ------------------------------------------------------------MSG------------EKLK--AFIL--S-YIN-NR-----K----QAK-------------------------------------------------------------------------------------------------L-----------------DVFDK-EAEKK-----RA--V---------LS--G-EA-----------LAVAER---------------------------EL-----------------------------------A-QAR-QE-----------------------IE---
------QK------HE---VCNW-----LT---DA-----ANRA-------G----QIS-L-----------------VTHAVKFTHSD---A-----------------K---G-S------SV----F----N----A---------E-T-----VAD--A--KT----------LST--A--SL------A-------Q--P--A-ID-A-VG----N----AA----A--L-----DVAKLL-Q-----T----E--H--D---------GDSL------------VAALQ-RGD-N---CALEAL---A---E----NP----EQLAQWLAGFQQV--F--TN--RQ----
PSSHKLTKQIYFPL-----------------------------ANGE---YHLLSPLYSSSLAHALHQR-IS----A--V---R--F-G-----------------DE-AKAIR-Q-----AQ-------------R--TNQWH--DQ-L-SIS-----YPN-LAVQNMGG-T-----KPQ----------NISALN--SSRS-----------GRSYLLSSA-P-----PQWN--S---IE-KPPQQH----ESIF----RP-RGEVD--Y------HT--RATLAQMQRFL------L--SV------K-----N-------------VEN---------N--
--RD---IR--QQ----RL------RYLDQLIDQLFFYVASVQ--N-----L--P-----V---G----WS------A-----E------------SE--------LKR---AQQLWL-----D-------------PYR--AET-------------DTVFRR--ERE--AGD-WQ-QAVAYDFGRWLN-----RR----L---K-------H-------E-N-L-----------IF---GEVER------REWST-A-------AL--FK----------------R-----RMREMESAL----------------KEEL-------A-----------------------
-------------
WP_039305971 ------------------------------------------------------------MSG------------EKLK--AFIL--S-YIN-NR-----K----QAK-------------------------------------------------------------------------------------------------L-----------------DAFDK-EAEKK-----RA--A---------LS----GEA-----------LAVAER---------------------------EL-----------------------------------A-QAR-QE-----------------------IE---
------QK------HE---VRNW-----LT---DA-----ANRA-------G----QIS-L-----------------VTHAVKFTHSD---A-----------------K---G-S------SV----F----N----A-----------ET-----VAD--A--KT----------LST--A--TL------A-------Q--P--A-ID-A-VG----N----AA----A--L-----DVAKLL-Q-----T----E--H--D---------GDSL------------VAALQ-RGD-N---RALEAL---A---E----NP----EQLAQWLTGFQQV--F--TN--RQ----
PSSHKLTKQIYFPL-----------------------------ANGE---YHLLSPLYSSSLAHALHQR-IS----A--V---R--F-G-----------------DE-AKAIR-Q-----AQ-------------R--TNQWH--DQ-L-SIS-----YPN-LAVQNMGG-T-----KPQ----------NISALN--SSRS-----------GRSYLLSSA-P-----PQWN--S---IE-KPPQQH----ESIF----RP-RGEVD--Y------HT--RATLAQMQRFL------L--SV------K-----N-------------VEN---------N--
--RD---IR--QQ----RL------RYLDQLIDQLFFYVASVQ--N-----L--P-----V---G----WS------A-----E------------SE--------LKR---AQQLWL-----D-------------PYR--AET-------------DTVFRR--ERE--AGD-WQ-QAVAYDFGRWLN-----RR----L---K-------H-------E-N-L-----------IF---GEVER------REWST-T-------AL--FK----------------R-----RMREMESAL----------------KEEL-------A-----------------------
-------------
```

```
WP_039293538 ---------------------------------------------------M--R---------NGLP--EFIL--S-YIN-NR-----K----QAK--------------------------------------------------------L---------D-------A-FDK--EAEK-----KR--A--------TL-----SG---------EALAVAEL-----------------------EL------------------------------A-QAR-RE---------------------IE--
LK-------------HE---VRNW-----LT---DA-----ASRA-------G----QIS-L----------------VTHALKFTHSD---A---------------K---G-S------SV----F----S---AE--------T-------VAD--A--KT----------LST--A--TL------E-------Q--P--A-ID-A-VG----N----AA----A--L-----DVAKLL-Q-----T----E--H--D---------GDSL------------VAALQ-RGD-H---SALEAL---A---E----NP----EQLAQWLTGFQQV--F---TN--RQ----
PSSHKLAKQIYFPL--------------------------ASGE---YHLLSPLYSSSLAQALYQR-IN----A--V---R--F-G-----------------DE-AKAIR-Q-----AQ------------R--TNQWH--DQ-L-SIS-----YPN-LAVQNMGG-T----KPQ----------NISALN--SSRS-----------GRSYLLSSA-P----PQWN--S---IE-KPPQQH----ESIF----RP-HGEVD--Y-----HT--RATLAQMQRFL------L--SV------K-----D-------------VEN---------N--
--RD---IR--QQ----RL------RYLDQLIDQLFFYVASVQ--N-----L---P-----V---G----WS------A-----E------------SE--------LKR---PQQLWL-----D-------------PYR--AET-------------DTVFRR--ERE--AGD-WR-QAVAYDFGRWLN-----RR----L---K-------H-------E-N-L------------IF---GEVER------REWST-A-------AL--FK-----------------R-----RMREMESAL----------------KEEL-------A----------------------
-------------
WP_039310142 -----------------------------------------------------MR-------------NGLP--EFIL--S-YIN-NR-----K----QAK--------------------------------------------------------L---------------DAFDK-EAEKK------------------RA---TLSG---------EALAVAEL-----------------------EL------------------------------A-QAR-RE---------------------IE-----
----LK------HE---VRNW-----LT---DA-----ASRA-------G----QIS-L---------------VTHALKFTHSD---A---------------K---G-S------SV----F----S---AE--------T-VAD---AKT---------------LST--A--TL------E-------Q--P--A-ID-A-VG----N----AA----A--L-----DVAKLL-Q-----T----E--H--D---------GDSL------------VAALQ-RGD-H---SALEAL---A---E----NP----EQLAQWLTGFQQV--F---TN--RQ----
PSSHKLAKQIYFPL--------------------------ASGE---YHLLSPLYSSSLAQALYQR-IN----A--V---R--F-G-----------------DE-AKAIR-Q-----AQ------------R--TNQWH--DQ-L-SIS-----YPN-LAVQNMGG-T----KPQ----------NISALN--SSRS-----------GRSYLLSSA-P----PQWN--S---IE-KPPQQH----ESIF----RP-RGEVD--Y-----HT--RATLAQMQRFL------L--SV------K-----D-------------VEN---------N--
--RD---IR--QQ----RL------RYLDQLIDQLFFYVASVQ--N-----L---P-----V---G----WS------V-----E------------SE--------LKR---AQQLWL-----D-------------PYR--AET-------------DTIFRR--ERE--AGD-WQ-QAVAYDFGRWLN-----RR----L---K-------H-------E-N-L------------IF---GEVER------REWST-A-------AL--FK-----------------R-----RMREMESAL----------------KEEL-------A----------------------
------------
WP_011095181 ---------------------------------------------------M--R---------NGLP--EFIL--S-YIN-NR-----K----QAK--------------------------------------------------------L---------D-------A-FDK--EAEK-----KR--A--------TL-----SG---------EALSVAEL-----------------------EL------------------------------A-KAR-RE---------------------IE--
QK-------------HE---VRNW-----LT---DA-----ASRA-------G----QIS-L----------------VTHALKFTHSD---A---------------K---G-S------SV----F----N---AE--------T-------VED--A--TT----------LST--A--TL------A-------Q--P--A-ID-A-VG----N----AA----A--L-----DVAKLL-Q-----T----E--H--D---------GDSL------------VAALQ-RGD-N---RALEAL---A---E----NP----EQLAQWLTGFQQV--F---TN--RQ----
PSSHKLAKQIYFPL--------------------------ANGE---YHLLSPLYSSSLAHALHQR-IS----A--V---R--F-G-----------------DE-AKAIR-Q-----AQ------------R--TNQWH--DQ-L-SIS-----YPN-LAVQNMGG-T----KPQ----------NISALN--SSRS-----------GRSYLLSSA-P----PQWN--S---IE-KPPQQH----ESIF----RP-RGEVD--Y-----HT--RATLAQMQRFL------L--SV------K-----D-------------VEN---------N--
--RD---IR--QQ----RL------HYLDQLIDQLFFYVASVQ--N-----L---P-----V---G----WS------A-----E------------SE--------LKR---AQQLWL-----D-------------PYR--AET-------------DTVFRR--ERE--AGD-WQ-QAVAYEFGRWLN-----RR----L---K-------H-------E-N-L------------IF---GEVER------REWST-A-------AL--FK-----------------R-----RMREMESAL----------------KEEL-------A----------------------
-------------
WP_048262191 ---------------------------------------------------ME--E---------NRLT--QFIV--S-YVT-NR-----K----QAK--------------------------------------------------------L---------D-------A-FDK--EAEK-----KR--A--------AL-----SG---------EALAVAER-----------------------EL------------------------------T-QAR-RE---------------------IE--
QK-------------HE---VRNW-----LT---DA-----ASRA-------G----QIS-L----------------VTHALKFTHSD---A---------------K---G-S------SV----F----N---AE--------T-------VED--A--TT----------LST--A--TL------A-------Q--P--A-ID-A-VG----N----AA----A--L-----DVAKLL-Q-----T----E--H--D---------GDSL------------VAALQ-RGD-N---RALEAL---A---E----NP----EQLAQWLTGFQQV--F---TN--RQ----
PSSHKLAKQIYFPL--------------------------ANGE---YHLLSPLYSSSLAHALHQR-IS----A--V---R--F-G-----------------DE-AKAIR-Q-----AQ------------R--TNQWH--DQ-L-SIS-----YPN-LAVQNMGG-T----KPQ----------NISALN--SSRS-----------GRSYLLSSA-P----PQWN--S---IE-KPPQQH----ESIF----RP-RGEVD--Y-----HT--RATLAQMQRFL------L--SV------K-----D-------------VEN---------N--
--RD---IR--QQ----RL------HYLDQLIDQLFFYVASVQ--N-----L---P-----V---G----WS------A-----E------------SE--------LKR---AQQLWL-----D-------------PYR--AET-------------DTVFRR--ERE--AGD-WQ-QAVAYEFGRWLN-----RR----L---K-------H-------E-N-L------------IF---GEVER------REWST-A-------AL--FK-----------------R-----RMREMESAL----------------KEEL-------A----------------------
-------------
WP_113626666 ---------------------------------------------------ME--E---------NKLA--QFIV--S-YIS-TR-----K----QAK--------------------------------------------------------L---------D-------A-FDK--EAEK-----KR--A--------AL-----SG---------EALAVAER-----------------------EL------------------------------A-QAR-QE---------------------IE--
QK-------------HE---ACNW-----LT---DA-----ASRA-------G----QIS-L----------------VTHALKFTHSD---A---------------K---G-S------SV----F----N---AE--------T-------VAN--A--KT----------LST--A--TL------A-------Q--P--A-ID-A-VG----N----AA----A--L-----DVAKLL-Q-----T----E--H--D---------GDSL------------VAALQ-RGD-N---RALEAL---A---E----NP----EQLAQWLTGFQQV--F---TN--RQ----
PSSHKLAKQIYFPL--------------------------ANGE---YHLLSPLYSSSLAHALHQR-IS----A--V---R--F-G-----------------DE-AKAIR-Q-----AQ------------R--TNQWH--DQ-L-SIS-----YPN-LAVQNMGG-T----KPQ----------NISALN--SSRS-----------GRSYLLSSA-P----PQWN--S---IE-KPPQQH----ESIF----RP-RGEVD--Y-----HT--RATLAQMQRFL------L--SV------K-----D-------------VEN---------N--
--RD---IR--QQ----RL------RYLDQLIDQLFFYVASVQ--N-----L---P-----V---G----WS------A-----E------------SE--------LKR---AQQLWL-----D-------------PYR--AET-------------DTVFRR--ERE--AGD-WQ-QAVAYEFGRWLN-----RR----L---K-------H-------E-N-L------------IF---GEVER------REWST-A-------AL--FK-----------------R-----RMREMESAL----------------KEEL-------A----------------------
-------------
WP_025919357 ---------------------------------------------------ME--D---------NRLT--QFIV--S-YIN-NR-----K----QAK--------------------------------------------------------L---------D-------A-FDK--DAEK-----KR--A--------AL-----SG---------EALAVTEL-----------------------EL------------------------------A-QVR-RE---------------------IE--
QK-------------HE---VQNW-----LT---DA-----ASRA-------G----QIS-L----------------VTHALKFTHSD---A---------------R---G-S------SI----F----S---AD--------T-------VAD--A--KT----------LST--A--TL------A-------Q--P--A-ID-A-VG----N----AA----A--L-----DVAKLL-Q-----T----E--Y--D---------GDSL------------VAALQ-RGD-H---SALEAL---A---E----SP----EQLAQWLAGFQQV--F---TD--RQ----
PSSHKLAKQIYFPL--------------------------ANGE---YHLLSPLYSSSLAQALHQR-IN----A--V---R--F-G-----------------DE-AKAIR-Q-----AQ------------K--ANQWH--DQ-L-SIS-----YPN-LAVQNMGG-T----KPQ----------NISALN--SSRS-----------GRSYLLSSA-P----PQWN--S---IE-KPPQQH----ECIF----RL-NGEVD--Y-----HT--RDIVRPMRHFL------L--SV------K-----E-------------VEN---------N--
-RD---IR--QQ----RL------RYLDQLIDQLFFYVASVQ--N-----L---P-----A---G----WS------A-----E------------SE--------LKR---AQQLWL-----D-------------PYR--ADT-------------DTVFRR--ERE--AGD-WQ-QAVAYDFGRWLN-----RR----L---K-------H-------E-N-L------------IF---GEVER------REWST-A-------AL--FK-----------------R-----RMREMEGTL----------------KEEL-------A----------------------
-------------
WP_015731171 -----------------------------------------------------MED-------------NRLT--QFIV--S-YIN-NR-----K----QAK--------------------------------------------------------L---------------DVFDK-DAEKK-------R--A--------AL---SGETLAVTEL-----------------------EL------------------------------A-QVR-RE---------------------IE---
-----QK------HE---VQNW-----LT---DA-----ASRA-------G----QIS-L---------------VTHALKFTHSD---A---------------R---G-S------SI----F----S---AD--------T-VAD---AKT---------------LST--A--TL------A-------Q--P--A-ID-A-VG----N----AA----A--L-----DVAKLL-Q-----T----E--Y--D---------GDSL------------VAALQ-RGD-H---RALEAL---A---E----NS----EQLAQWLTGFQQV--F---TD--RQ----
PSSHKLAKQIYFPL--------------------------ANGE---YHLLSPLYSSSLAQALHQR-IN----A--V---R--F-G-----------------DE-AKAIR-Q-----AQ------------K--ANQWH--DQ-L-SIS-----YPN-LAVQNMGG-T----KPQ----------NISALN--SSRS-----------GRSYLLSSA-P----PQWN--S---IE-KPPQQH----ECIF----RL-NGEVD--Y-----HT--RDIVRPMRHFL------L--SV------K-----E-------------VEN---------N--
-RD---IR--QQ----RL------RYLDQLIDQLFFYVASVQ--N-----L---P-----V---G----WS------A-----E------------SE--------LKR---AQQLWL-----D-------------PYR--ADT-------------DTVFRR--ERE--AGD-WQ-QAVAYDFGRWLN-----RR----L---K-------H-------E-N-L------------IF---GEVER------REWST-A-------AL--FK-----------------R-----RMREMEGTL----------------KEEL-------A----------------------
-------------
WP_121296313 ---------------------------------------------------ME--D---------NRLT--QFIV--S-YIN-NR-----K----QAK--------------------------------------------------------L---------D-------V-FDK--DAEK-----KR--A--------AL-----SG---------EALAVTEL-----------------------EL------------------------------A-QVR-RE---------------------IE--
QK-------------HE---VQNW-----LT---DA-----ASRA-------G----QIS-L----------------VTHALKFTHSD---A---------------R---G-S------SI----F----S---AD--------T-------VAD--A--KT----------LST--A--TL------A-------Q--P--A-ID-A-VG----N----AA----A--L-----DVAKLL-Q-----T----E--Y--D---------GDSL------------VAALQ-RGD-H---SALEAL---A---E----SP----EQLAQWLAGFQQV--F---TD--RQ----
PSSHKLAKQIYFPL--------------------------ANGE---YHLLSPLYSSSLAQALHQR-IN----A--V---R--F-G-----------------DE-AKAIR-Q-----AQ------------K--ASQWH--DQ-L-SIS-----YPN-LAVQNMGG-T----KPQ----------NISALN--SSRS-----------GRSYLLSSA-P----PQWN--S---IE-KPPQQH----ESIF----RP-RGEVD--Y-----YT--RATLAQMQRFL------L--SV------K-----E-------------VEN---------N--
--RD---IR--QQ----RL------RYLDQLIDQLFFYVASVQ--N-----L---P-----A---G----WS------A-----E------------SE--------LKR---AQQLWL-----D-------------PYR--ADT-------------DTVFRR--ERE--AGD-WQ-QAVAYDFGRWLN-----RR----L---K-------H-------E-N-L------------IF---GEVER------REWST-A-------AL--FK-----------------R-----RMREMEGTL----------------KEEL-------A----------------------
-------------
WP_033072110 ---------------------------------------------------MR--G---------EKLK--TFIL--S-YIS-TR-----K----QAK--------------------------------------------------------L---------D-------A-FDK--DAEK-----KR--A--------AL-----SG---------EALAVTEL-----------------------EL------------------------------A-QVR-RE---------------------IE--
QK-------------HE---VQNW-----LT---DA-----ASRA-------G----QIS-L----------------VTHALKFTHSD---A---------------R---G-S------SI----F----S---AD--------T-------VAD--A--KT----------LST--A--TL------A-------Q--P--A-ID-A-VG----N----AA----A--L-----DVAKLL-Q-----T----E--Y--D---------GDSL------------VAALQ-RGD-H---SALEAL---A---E----SP----EQLAQWLAGFQQV--F---TD--RQ----
PSSHKLAKQIYFPL--------------------------ANGE---YHLLSPLYSSSLAQALHQR-IN----A--V---R--F-G-----------------DE-AKAIR-Q-----AQ------------K--ASQWH--DQ-L-SIS-----YPN-LAVQNMGG-T----KPQ----------NISALN--SSRS-----------GRSYLLSSA-P----PQWN--S---IE-KPPQQH----ESIF----RP-RGEVD--Y-----YT--RATLAQMQRFL------L--SV------K-----E-------------VEN---------N--
--RD---IR--QQ----RL------RYLDQLIDQLFFYVASVQ--N-----L---P-----A---G----WS------A-----E------------SE--------LKR---TQQLWL-----D-------------PYR--AET-------------DTVFRR--ERE--AGD-WQ-QAVAYDFGRWLN-----RR----L---K-------H-------E-N-L------------IF---GEVER------REWST-A-------AL--FK-----------------R-----RMREMEGTL----------------KEEL-------A----------------------
-------------
WP_014701307 -----------------------------------------------------MRG-------------EKLK--TFIL--S-YIS-TR-----K----QAK--------------------------------------------------------L---------D-------AFDKD-AEKKR-----AA--L--------SG---------------EALAVTEL-----------------------EL------------------------------A-QVR-RE---------------------IE---
-----QK------HE---VQNW-----LT---DA-----ASRA-------G----QIS-L---------------VTHALKFTHSD---A---------------R---G-S------SI----F----S--------------AD---TVA--D-AKT----------LST--A--TL------A-------Q--P--A-ID-A-VG----N----AA----A--L-----DVAKLL-Q-----T----E--Y--D---------GDSL------------VAALQ-RGD-H---SALEAL---A---E----SP----EQLAQWLAGFQQV--F---TD--RQ----
PSSHKLAKQIYFPL--------------------------ANGE---YHLLSPLYSSSLAQALHQR-IN----A--V---R--F-G-----------------DE-AKAIR-Q-----AQ------------K--ASQWH--DQ-L-SIS-----YPN-LAVQNMGG-T----KPQ----------NISALN--SSRS-----------GRSYLLSSA-P----PQWN--S---IE-KPPQQH----ESIF----RP-RGEVD--Y-----YT--RATLAQMQRFL------L--SV------K-----E-------------VEN---------N--
--RD---IR--QQ----RL------RYLDQLIDQLFFYVASVQ--N-----L---P-----A---G----WS------A-----E------------SE--------LKR---AQQLWL-----D-------------PYR--ADT-------------DTVFRR--ERE--AGD-WQ-QAVAYDFGRWLN-----RR----L---K-------H-------E-N-L------------IF---GEVER------REWST-A-------AL--FK-----------------R-----RMREMEGTL----------------KEEL-------A----------------------
-------------
WP_103807965 -----------------------------------------------------MRG-------------EKLK--TFIL--S-YIS-TR-----K----QAK--------------------------------------------------------L---------------DAFDK-DAEKK-------R--A--------AL---SGEALAVTEL-----------------------EL------------------------------A-QVR-RE---------------------IE---
-----QK------HE---VQNW-----LT---DA-----ASRA-------G----QIS-L---------------VTHALKFTHSD---A---------------R---G-S------SI----F----S---AD--------T-VAD---AKT---------------LST--A--TL------A-------Q--P--A-ID-A-VG----N----AA----A--L-----DVAKLL-Q-----T----E--Y--D---------GDSL------------VAALQ-RGD-H---SALEAL---A---E----SP----EQLAQWLAGFQQV--F---TD--RQ----
PSSHKLAKQIYFPL--------------------------ANGE---YHLLSPLYSSSLAQALHQR-IN----A--V---R--F-G-----------------DE-AKAIR-Q-----AQ------------K--ASQWH--DQ-L-SIS-----YPN-LAVQNMGG-T----KPQ----------NISALN--SSRS-----------GRSYLLSSA-P----PQWN--S---IE-KPPQQH----ESIF----RP-RGEVD--Y-----YT--RATLAQMQRFL------L--SV------K-----E-------------VEN---------N--
--RD---IR--QQ----RL------RYLDQLIDQLFFYVASVQ--N-----L---P-----V---G----WS------A-----E------------SE--------LKR---AQQLWL-----D-------------PYR--ADT-------------DTVFRR--ERE--AGD-WQ-QAVAYDFGRWLN-----RR----L---K-------H-------E-N-L------------IF---GEVER------REWST-A-------AL--FK-----------------R-----RMREMEGTL----------------KEEL-------A----------------------
-------------
WP_095700531 ---------------------------------------------------ME--D---------NCLT--QFIV--S-YIN-NR-----K----QAK--------------------------------------------------------L---------D-------A-FDK--EAEK-----KR--A--------AL-----NG---------EALAAEEL-----------------------DL------------------------------A-EKR-RE---------------------IE--
QK-------------HE---VRAW-----LT---DA-----ASRA-------G----QIS-L----------------VTHALKFTHSD---A---------------K---G-S------SI----F----S---AE--------T-------VAD--A--KT----------LST--S--TL------A-------Q--P--A-ID-A-VG----N----AA----A--L-----DVAKLL-Q-----T----E--N--D---------GDSL------------VAALQ-RGD-H---SALDPL---A---E----NP----EQLTQWLAGFQQV--F---TD--RQ----
PSSHKLAKQIYFPT--------------------------ENSE---YHLLSPLYSSSLAQALHQR-IN----A--V---R--F-G-----------------DE-AKDIR-K-----AR------------K--DNLWH--DQ-L-DIS-----YPN-LAVQNMGG-T----KPQ----------NISSLN--SSRS-----------GRSYLLSCA-P----PQWN--S---VE-KPPQQH----DSIF----RP-RGEVD--Y-----HT--RATLAQMQRFL------L--SV------K-----E-------------VEN---------N-
---RD---IR--QQ----RL------RYLDQLIDQLFFYVASVQ--N-----L---S-----A---G----WS------A-----E------------SE--------LKR---AQQLWL-----D-------------PYR--AKT-------------DTVFRR--ELE--AGD-WQ-KSVAYDFGRWLN-----RR----L---K-------H-------E-E-L------------IF---GEVER------REWST-A-------AL--FK-----------------H-----RMREMESSL----------------KEDL-------A----------------------
-------------
WP_116186940 ---------------------------------------------------ME--D---------NRLT--PFIV--S-YIN-NR-----K----QAK--------------------------------------------------------L---------D-------A-FDK--EAEK-----KR--V--------AL-----SG---------EALAAEEL-----------------------EL------------------------------A-EKR-RE---------------------IE--
QK-------------HE---VRAW-----LT---NA-----ASRA-------G----QIS-L----------------VTHALKYTHSD---A---------------K---G-S------SI----F----S---AE--------A-------AAD--A--RT----------LST--A--TL------A-------Q--P--A-ID-V-VG----N----AA----V--L-----DVAGLL-Q-----T----E--H--E---------GDSL------------VVALQ-RGD-H---RALEAL---A---E----SP----EQFALWLAGFQKV--F---TD--RQ----
PSSHKLAKQIYFPL--------------------------ESGE---YHLLSPLYSSSLAQALHQR-IN----A--V---R--F-G-----------------DE-AKAIR-Q-----AQ------------K--ANQWH--EQ-L-SIS-----YPN-LAVQNRGG-S----NKQ----------NISSLN--VTRN-----------GRSYLLSSA-P----PQWN--S---VE-KPPQQH----ESIF----RP-RGEVD--Y-----HT--RATLAQMQRFL------L--SV------K-----E-------------VEN---------N-
---RD---IR--QQ----RL------HYLDQLIDQLFFYVSSVQ--N-----L---P-----A---G----WS------A-----E------------SE--------LKR---AQQLWL-----D-------------PYR--AET-------------DTVFRR--ERE--AGD-WQ-KAVAYDFGRWLN-----LR----L---K-------H-------E-A-L------------IF---GEVER------REWST-A-------AL--FK-----------------R-----RMREMESAL----------------KEDL-------A----------------------
-------------
WP_125233096 ---------------------------------------------------MA--E---------NRLT--QFIV--S-YVT-NR-----K----QAK--------------------------------------------------------L---------D-------A-FDK--EAEK-----KC--A--------AL-----SG---------EALAAEEL-----------------------EL------------------------------A-EKR-RE---------------------IE--
QK-------------HE---VRAW-----LT---NA-----ASRA-------G----QIS-L----------------VTHALKYTHSD---A---------------K---G-S------SI----F----S---AE--------A-------AAD--A--RT----------LST--A--TL------A-------Q--P--A-ID-V-VG----N----AA----V--L-----DVAGLL-Q-----T----E--H--E---------GDSL------------VVALQ-RGD-H---RALEAL---A---E----SP----EQFALWLAGFQKV--F---TD--RQ----
PSSHKLAKQIYFPL--------------------------ESGE---YHLLSPLYSSSLAQALHQR-IN----A--V---R--F-G-----------------DK-AKAIR-Q-----AQ------------K--ANQWH--EQ-L-SIS-----YPN-LAVQNRGG-S----NKQ----------NISSLN--VTRN-----------GRSYLLSSA-P----PQWN--S---VE-KPPQQH----ESIF----RP-RGEVD--Y-----YT--RATLAHMQRFL------L--SV------K-----E-------------VEN---------N-
---RD---IR--QQ----RL------RYLDQLIDQLFFYVASVQ--N-----L---P-----A---G----WS------A-----Q------------SE--------LKR---AQQLWL-----D-------------PYR--AET-------------DTVFRR--ERE--AGG-WQ-KAVAYDFGRWLN-----RR----L---K-------H-------E-E-L------------IF---GEVER------REWST-A-------AL--FK-----------------R-----RMREMESSL----------------KEDL-------A----------------------
-------------
WP_103942333 ---------------------------------------------------MS--G---------EKLK--AFIL--S-YIN-NR-----K----QAK--------------------------------------------------------L---------D-------A-FDK--EAEK-----KR--A--------TL-----SG---------EALAEEEL-----------------------EL------------------------------A-EKR-RE---------------------IE--
QK-------------HE---VRAW-----LT---DA-----ASRA-------G----QIS-L----------------VTHALKFTHSD---A---------------K---G-S------SI----F----S---AE--------T-------ATD--A--KT----------LST--A--TL------A-------Q--P--A-ID-A-VG----N----AA----A--L-----DVAKLL-Q-----T----E--H--E---------GDSL------------VAALQ-RGD-H---SALEAL---A---E----SP----EQLAQWLAGFQQV--F---TD--RQ----
PSSHKLAKQIYFPT--------------------------ENGE---YHLLSPLYSSSLAQVLHQR-IN----A--V---R--F-G-----------------DE-AKDIR-K-----AR------------K--DNLWH--DQ-L-DIS-----YPN-LAVQNMGG-T----KPQ----------NISSLN--SSRS-----------GRSYLLSCA-P----PQWN--S---VE-KPPQQH----ECIF----RS-NGEAD--Y-----HT--SDIVKSMRRFL------L--SV------K-----K-------------VQN---------N--
--RD---IR--QQ----RL------HYLDQLIDQLFFYASSVQ--N-----L---S-----A---G----WS------A-----E------------SE--------LTR---AQQLWL-----D-------------PYR--AET-------------DTVFRR--ERE--AGG-WQ-KAVAYDFGRWLN-----RR----L---K-------H-------E-E-L------------IF---GEVER------REWST-A-------AL--FK-----------------R-----RMREMESSL----------------KEDL-------A----------------------
-------------
WP_103972276 -----------------------------------------------------MSG-------------EKLK--AFIL--S-YIN-NR-----K----QAK--------------------------------------------------------L---------------DAFDK-EAEKK-------R--A--------TL---SG-E----------ALAEEEL-----------------------EL------------------------------A-EKR-RE---------------------IE---
-----QK------HE---VRAW-----LT---DA-----ASRA-------G----QIS-L---------------VTHALKFTHSD---A---------------K---G-S------SI----F----S----A--------E-T-----ATD--A--KT----------LST--A--TL------A-------Q--P--A-ID-A-VG----N----AA----A--L-----DVAKLL-Q-----T----E--H--E---------GDSL------------VAALQ-RGD-H---SALEAL---A---E----SP----EQLAQWLAGFQQV--F---TD--RQ----
PSSHKLAKQIYFPT--------------------------ENGE---YHLLSPLYSSSLAQALHQR-IN----A--V---R--F-G-----------------DE-AKDIR-K-----AR------------K--DNLWH--DQ-L-DIS-----YPN-LAVQNMGG-T----KPQ----------NISSLN--SSRS-----------GRSYLLSCA-P----PQWN--S---VE-KPPQQH----ECIF----RS-NGEAD--Y-----HT--SDIVKSMRRFL------L--SV------K-----K-------------VQN---------N--
--RD---IR--QQ----RL------HYLDQLIDQLFFYASSVQ--N-----L---S-----A---G----WS------A-----E------------SE--------LTR---AQQLWL-----D-------------PYR--AET-------------DTVFRR--ERE--AGG-WQ-KAVAYDFGRWLN-----RR----L---K-------H-------E-E-L------------IF---GEVER------REWST-A-------AL--FK-----------------R-----RMREMESSL----------------KEDL-------A----------------------
-------------
WP_119158203 -----------------------------------------------------MSG-------------EKLK--AFIL--S-YIN-NR-----K----QAK--------------------------------------------------------L---------------DAFDK-EAEKK-------R--A--------TL---S-GE----------ALAEEEL-----------------------EL------------------------------A-EKR-RE---------------------IE---
-----QK------HE---VRAW-----LT---DA-----ASRA-------G----QIS-L---------------VTHALKFTHSD---A---------------K---G-S------SI----F----S----A---------ET----ATD--A--KT----------LST--A--TL------A-------Q--P--A-ID-A-VG----N----AA----A--L-----DVAKLL-Q-----T----E--H--E---------GDSL------------VAALQ-RGD-H---SALEAL---A---E----SP----EQLAQWLAGFQQV--F---TD--RQ----
PSSHKLAKQIYFPT--------------------------ENGE---YHLLSPLYSSSLAQALHQR-IN----A--V---R--F-G-----------------DE-AKDIR-K-----AR------------K--DNLWH--DQ-L-DIS-----YPN-LAVQNMGG-T----KPQ----------NISSLN--SSRS-----------GRSYLLSCA-P----PQWN--S---VE-KPPQQH----ECIF----RS-NGEAD--Y-----HT--SDIVKSMRRFL------L--SV------K-----K-------------VQN---------N--
--RD---IR--QQ----RL------HYLDQLIDQLFFYASSVQ--N-----L---S-----A---G----WS------A-----E------------SE--------LTR---AQQLWL-----D-------------PYR--AET-------------DTVFRR--ERE--AGG-WQ-KAVAYDFGRWLN-----RR----L---K-------H-------E-E-L------------IF---GEVER------REWST-A-------AL--FK-----------------R-----RMREMESAL----------------KEDL-------A----------------------
-------------
WP_127157172 ---------------------------------------------------ME--D---------NCLT--QFIV--S-YIN-NR-----K----QAK--------------------------------------------------------L---------D-------A-FDK--EAEK-----KR--A--------TL-----SG---------EALAEEEL-----------------------EL------------------------------A-EKR-RE---------------------IE--
QK-------------HE---VRTW-----LT---DA-----ASRA-------G----QIS-L----------------VTHALKFTHSD---A---------------K---G-S------SI----F----S---AE--------T-------VAD--A--KT----------LST--A--TL------A-------Q--P--A-ID-A-VG----N----AA----A--L-----DVAKLL-Q-----T----E--N--D---------GDSL------------VAALQ-RGD-H---SALDPL---A---E----SP----EQLAQWLAGFQQV--F---TN--RQ----
PSSHKLAKQIYFPT--------------------------ENGE---YHLLSPLYSSSLAQALHQR-IN----A--V---R--F-G-----------------DE-AKAIR-Q-----AQ------------K--ANQWH--DQ-L-AIS-----YPN-LAVQNMGG-T----KPQ----------NISSLN--SSRS-----------GRSYLLSCA-P----PQWN--S---IE-KPPLQH----DSIF----RP-RGEVD--Y-----HT--RATLTQMQRFL------L--SV------K-----E-------------VEN---------N--
--RD---IR--QQ----RL------HYLDQLIDQLFFYVSSVQ--N-----L---P-----A---G----WS------A-----E------------SE--------LTR---TQQLWL-----D-------------PYR--AET-------------DTVFHR--ERE--AGD-WQ-KAVAYDFGRWLN-----LR----L---K-------H-------E-T-L------------IF---GEVER------REWST-A-------AL--FK-----------------R-----RMREMESSL----------------KEDL-------A----------------------
-------------
WP_102117294 ---------------------------------------------------ME--D---------NRLT--PFIV--S-YIN-NR-----K----QAK--------------------------------------------------------L---------D-------A-FDK--EAEK-----KR--V--------AL-----SG---------EALAAEEL-----------------------EL------------------------------A-EKR-RE---------------------IE--
QK-------------HE---VHAW-----LT---DA-----ASRA-------G----QIS-L----------------VTHALKFTHSD---A---------------K---G-S------SI----F----S---AE--------T-------VAD--A--KT----------LST--A--TL------A-------Q--P--A-ID-A-VG----N----AA----A--L-----DVAKLL-Q-----T----E--H--D---------GDSL------------VAALQ-RGD-H---SALEAL---A---E----SP----EQLAQWLAGFQQV--F---TD--RQ----
PSSHKLAKQIYFPT--------------------------ENGE---YHLLSPLYSSSLAQALHQR-IN----A--V---R--F-G-----------------DE-AKAIR-Q-----AQ------------K--ANQWH--DQ-L-AIS-----YPN-LAVQNMGG-T----KPQ----------NISSLN--SSRS-----------GRSYLLSCA-P----PQWN--S---VE-KPPQQH----ESIF----RP-RGEVD--Y-----HT--RATLALMQRFL------L--SV------K-----E-------------VEN---------N-
---RD---IR--QQ----RL------HYLDQLIDQLFFYVTSVQ--N-----L---P-----S---G----WS------A-----E------------SE--------LKR---AQQLWL-----D-------------PYR--AET-------------DTVFRR--ERE--AGD-WQ-KAVAYDFGRWLN-----LR----L---K-------H-------E-A-L------------IF---GEVER------REWST-A-------AL--FK-----------------R-----RMREMESAL----------------KEDL-------A----------------------
-------------
WP_103861798 ---------------------------------------------------MS--G---------EKLK--AFIL--S-YIN-NR-----K----QAK--------------------------------------------------------L---------D-------A-FDK--EAEK-----KR--A--------TL-----SG---------EALAEEEL-----------------------EL------------------------------A-EKR-RE---------------------IE--
QK-------------HE---VRAW-----LT---DA-----ASRA-------G----QIS-L----------------VTHALKFTHSD---A---------------K---G-S------SI----F----S---AE--------T-------ATD--A--KT----------LST--A--TL------A-------Q--P--A-ID-A-VG----N----AA----A--L-----DVAKLL-Q-----T----E--H--E---------GDSL------------VAALQ-RGD-H---SALEAL---A---E----SP----EQLAQWLAGFQQV--F---TD--RQ----
PSSHKLAKQIYFPT--------------------------ENGE---YHLLSPLYSSSLAQALHQR-IN----A--V---R--F-G-----------------DE-AKDIR-K-----AR------------K--DNLWH--DQ-L-DIS-----YPN-LAVQNMGG-T----KPQ----------NISSLN--SSRS-----------GRSYMLSSA-P----PQWN--S---IE-KPPQQH----DSIF----RP-RGEVD--Y-----HT--LATLAQMQRFL------L--SV------K-----E-------------VEN---------N-
--RD---IR--QQ----RL------HYLDQLIDQLFFYVTSVQ--N-----L---P-----S---G----WS------A-----K------------SE--------LKR---AQQLWL-----D-------------PYR--AET-------------DTVFRR--ERE--AGD-WQ-KAVAYDFGRWLN-----LR----L---K-------H-------E-A-L------------IF---GEVER------REWST-A-------AL--FK-----------------R-----RMREMESAL----------------KEDL-------A----------------------
-------------
WP_130634398 ---------------------------------------------------MS--G---------EKLK--AFIL--S-YIN-NR-----K----QAK--------------------------------------------------------L---------D-------A-FDK--EAEK-----KR--A--------TL-----SG---------EALAAEEL-----------------------EL------------------------------A-EKR-RE---------------------IE--
QK-------------HE---VRAW-----LT---DA-----ASRA-------G----QIS-L----------------VTHALKFTHSD---A---------------K---G-S------SI----F----S---AE--------T-------VAD--A--KT----------LST--A--TL------A-------Q--P--A-ID-A-VG----N----AA----A--L-----DVAKLL-Q-----T----E--H--E---------GDSL------------VVALQ-RGD-H---RALEAL---A---E----NP----EQLAQWLAGFQQV--F---TD--RQ----
PSSHKLAKQIYFPL--------------------------ENGE---YHLLSPLYSSSLAHALYQR-IN----A--V---R--F-G-----------------DE-AKAIR-Q-----AQ------------K--ANQWH--DQ-L-SIS-----YPY-LAVQNMGG-T----KPQ----------NISSLN--SSRS-----------GRSYMLSSA-P----PQWN--S---IE-KPPQQH----DSIF----RP-RGEVD--Y-----HT--RATLAQMQRFL------L--SV------K-----E-------------VEN---------N-
--RD---IR--QQ----RL------HYLDQLIDQLFFYVTSVQ--N-----L---P-----S---G----WS------A-----K------------SE--------LKR---AQQLWL-----D-------------PYR--AET-------------DTVFRR--ERE--AGD-WQ-KAVAYDFGRWLN-----RR----L---K-------H-------E-A-L------------IF---GEVER------REWST-A-------AL--FK-----------------R-----RMREMESAL----------------KEDL-------A----------------------
-------------
WP_039541300 ---------------------------------------------------MS--G---------EKLK--AFIL--S-YIN-TR-----K----QAK--------------------------------------------------------L---------D-------A-FDK--EAEK-----KR--A--------TL-----SG---------EALAEEEL-----------------------EL------------------------------S-EKR-RE---------------------IE--
QK-------------HE---VRAW-----LT---DA-----ASRA-------G----QIS-L----------------VTHALKFTHSD---A---------------K---G-S------SI----F----S---AE--------T-------VAD--A--KT----------LST--A--TL------A-------Q--P--T-ID-A-VG----N----AA----A--L-----DVAKLL-Q-----T----E--H--E---------GDSL------------VAALQ-RGD-H---SALEAL---A---E----SP----EQLAQWLAGFQQV--F---TD--RQ----
PSSHKLAKQIYFPL--------------------------ENGE---YHLLSPLYSSSLAHALHQR-VN----A--V---R--F-G-----------------DE-AKAIR-Q-----AQ------------K--ANQWH--DQ-L-SIS-----YPY-LAVQNMGG-T----KPQ----------NISSLN--SSRS-----------GRSYLLSCA-P----PQWN--S---IE-KPPQQH----DSIF----RP-RGEVD--Y-----HT--RATLALMQRFL------L--SV------K-----E-------------VEN---------N-
--RD---IR--QQ----RL------HYLDQLIDQLFFYVTSVQ--N-----L---P-----E---G----WS------T-----E------------SE--------LKR---AQQLWL-----D-------------PYR--AET-------------DTVFRR--ERE--AGD-WQ-KAVAYDFGRWLN-----RR----L---K-------H-------E-E-L------------IF---GEVER------REWST-A-------AL--FK-----------------R-----RMREMESAL----------------KEDL-------A----------------------
-------------
WP_039507081 ---------------------------------------------------MG--S---------EKLT--EFIL--S-YIN-NR-----K----QAK--------------------------------------------------------L---------D-------A-FDK--EAEK-----KR--A--------AL-----NG---------EALAAEEL-----------------------EL------------------------------A-EKR-RE---------------------IE--
QK-------------HE---VRAW-----LT---DA-----ASRA-------G----QIS-L----------------VTHALKFTHSD---A---------------K---G-S------SI----F----S---AE--------T-------VAD--A--KT----------LST--A--TL------A-------Q--P--A-ID-A-VG----N----AA----A--L-----DVAKLL-Q-----T----E--H--E---------GDSL------------VVALQ-RGD-H---SALEAL---A---E----SP----EQLTQWLAGFQQV--F---TD--RQ----
PSSHKLAKQIYFPL--------------------------ENGE---YHLLSPLYSSSLAHALHQR-VN----A--V---R--F-G-----------------DE-AKAIR-Q-----AQ------------K--ANQWH--DQ-L-SIS-----YPY-LAVQNMGG-T----KPQ----------NISSLN--SSRS-----------GRSYLLSCA-P----PQWN--S---IE-KPPQQH----DSIF----RP-RGEVD--Y-----HT--RATLALMQRFL------L--SV------K-----E-------------VEN---------N-
---RD---IR--QQ----RL------HYLDQLIDQLFFYVTSVQ--N-----L---P-----S---G----WS------A-----E------------SE--------LKR---AQQLWL-----D-------------PYR--AEA-------------DTVFRR--ERE--AGD-WQ-KAVAYDFGRWLN-----RR----L---K-------H-------E-E-L------------IF---GEVER------REWST-A-------AL--FK-----------------R-----RMREMESAL----------------KEDL-------A----------------------
-------------
WP_040032644 -----------------------------------------------------MSG-------------EKLK--AFIL--S-YIN-TR-----K----QAK--------------------------------------------------------L---------------DAFDK--EAEK------K--L--------AA---LNGE----------ALAAEEL-----------------------EL------------------------------A-EKR-RE---------------------IE---
----QK------HE---VRAW-----LT---DA-----ASRA-------G----QIS-L---------------VTHALKFTHSD---A---------------K---G-S------SI----F----S---AE--------T-VAD---AKT---------------LST--A--TL------T-------Q--P--A-ID-A-VG----N----AA----A--L-----DVAKLL-Q-----T----E--H--E---------GDSL------------VAALQ-RGD-H---SALEAL---A---E----SP----EQLAQWLAGFQQV--F---TD--RQ----
PSSHKLAKQIYFPL--------------------------ENGE---YHLLSPLYSSSLAHALHQR-VN----A--V---R--F-G-----------------DE-AKAIR-Q-----AQ------------K--ANQWH--DQ-L-SIS-----YPY-LAVQNMGG-T----KPQ----------NISSLN--SSRS-----------GRSYLLSSA-P----PQWN--S---IE-KPPQQH----DSIF----RP-RGEVD--Y-----HT--RATLAQMQRFL------L--SV------K-----E-------------VEN---------N-
---RD---IR--QQ----RL------HYLDQLIDQLFFYVTSVQ--N-----L---P-----S---G----WS------A-----E------------SE--------LKR---AQQLWL-----D-------------PYR--AEA-------------DTVFRR--ERE--AGD-WQ-KAVAYDFGRWLN-----RR----L---K-------H-------E-E-L------------IF---GEVER------REWST-A-------AL--FK-----------------R-----RMREMESAL----------------KEDL-------E----------------------
-------------
WP_130637400 ---------------------------------------------------MA--E---------NRLT--QFIV--S-YVT-NR-----K----QAK--------------------------------------------------------L---------D-------A-FDK--EAEK-----KR--A--------TL-----SG---------EALAAEEL-----------------------EL------------------------------A-EKR-RE---------------------IE--
QK-------------HE---VRAW-----LT---DA-----ASRA-------G----QIS-L----------------VTHALKFTHSD---A---------------K---G-S------SI----F----S---AE--------T-------VAD--A--KT----------LST--A--TL------A-------Q--P--A-ID-A-VG----N----AA----A--L-----DVAKLL-Q-----T----E--H--E---------GDSL------------VAALQ-RGD-H---SALEAL---A---E----SP----EQLAQWLAGFQQV--F---TD--RQ----
PSSHKLAKQIYFPL--------------------------ENGE---YHLLSPLYSSSLAHALHQR-VN----A--V---R--F-G-----------------DE-AKAIR-Q-----AQ------------K--ANQWH--DQ-L-SIS-----YPY-LAVQNMGG-T----KPQ----------NISSLN--SSRS-----------GRSYMLSSA-P----PQWN--S---IE-KPPQQH----DSIF----RP-RGEVD--Y-----HT--RATLAQMQRFL------L--SV------K-----E-------------VEN---------
N----RD---IR--QQ----RL------HYLDQLIDQLFFYVTSVQ--N-----L---P-----S---G----WS------A-----E------------SE--------LKR---AQQLWL-----D-------------PYR--AET-------------DTVFRR--ERE--AGD-WQ-KAVAYDFGRWLN-----RR----L---K-------H-------E-T-L------------IF---GEVER------REWST-A-------AL--FK-----------------R-----RMREMESAL----------------KEDL-------A----------------------
----------------
WP_103160653 ---------------------------------------------------ME--E---------NRLA--QFIV--S-YVT-NR-----K----QAK--------------------------------------------------------L---------D-------A-FDK--EAEK-----KR--A--------AL-----NG---------EALAAEEL-----------------------EL------------------------------A-EKR-RE---------------------IE--
QK-------------HE---VRAW-----LT---DA-----ASRA-------G----QIS-L----------------VTHALKFTHSD---A---------------K---G-S------SI----F----S---AE--------T-------VAD--A--KT----------LST--A--AL------A-------Q--P--V-ID-A-VG----N----AA----A--L-----DVAKLL-Q-----T----E--H--D---------GDSL------------VAALQ-RGD-H---RALEAL---A---E----SP----EQLAQWLAGFQQV--F---TD--RQ----
PSSHKLAKQIYFPL--------------------------ENGE---YHLLSPLYSSSLAHALHQR-VN----A--V---R--F-G-----------------DE-AKAIR-Q-----AQ------------K--ANQWH--DQ-L-SIS-----YPY-LAVQNMGG-T----KPQ----------NISSLN--SSRS-----------GRSYLLSCA-P----PQWN--S---VE-KPPQQH----ESIF----RP-RGEVD--Y-----HT--RATLAQMQRFL------L--SV------K-----E-------------VEN---------
N----RD---IR--QQ----RL------RYLDQLIDQLFFYISSVQ--N-----L---P-----S---G----WS------A-----E------------SE--------LKR---AQQLWL-----D-------------PYR--AET-------------DTVFRR--ERE--AGD-WQ-KAVAYDFGRWLN-----LR----L---K-------H-------E-E-L------------IF---GEVER------REWST-A-------AL--FK-----------------R-----RMREMESAL----------------KEDL-------A----------------------
----------------
WP_103183148 ---------------------------------------------------ME--E---------NRLA--QFIV--S-YVT-NR-----K----QAK--------------------------------------------------------L---------D-------A-FDK--EAEK-----KR--A--------AL-----NG---------EALAAEEL-----------------------EL------------------------------A-EKR-RE---------------------IE--
QK-------------HE---VRAW-----LT---DA-----ASRA-------G----QIS-L----------------VTHALKFTHSD---A---------------K---G-S------SI----F----S---AE--------T-------VAD--A--KT----------LST--A--TL------A-------Q--P--A-ID-A-VG----N----AA----A--L-----DVAKLL-Q-----T----E--H--D---------GDSL------------VAALQ-RGD-H---RALEAL---A---E----SP----EQLAQWLAGFQQV--F---TD--RQ----
PSSHKLAKQIYFPL--------------------------ENGE---YHLLSPLYSSSLAHALHQR-VN----A--V---R--F-G-----------------DE-AKAIR-Q-----AQ------------K--ANQWH--DQ-L-SIS-----YPY-LAVQNMGG-T----KPQ----------NISSLN--SSRS-----------GRSYLLSCA-P----PQWN--S---VE-KPPQQH----ESIF----RP-RGEVD--Y-----HT--RATLAQMQRFL------L--SV------K-----E-------------VEN---------
N----RD---IR--QQ----RL------RYLDQLIDQLFFYISSVQ--N-----L---P-----S---G----WS------A-----E------------SE--------LKR---AQQLWL-----D-------------PYR--AET-------------DTVFRR--ERE--AGD-WQ-KAVAYDFGRWLN-----LR----L---K-------H-------E-E-L------------IF---GEVER------REWST-A-------AL--FK-----------------R-----RMREMESAL----------------KEDL-------A----------------------
----------------
WP_103198672 -----------------------------------------------------MEE-------------NRLA--QFIV--S-YVT-NR-----K----QAK--------------------------------------------------------L---------D-------AFDKE-AEKKR-----AA--L--------NG---------------EALAAEEL-----------------------EL------------------------------A-EKR-RE---------------------IE--
-------QK------HE---VRAW-----LT---DA-----ASRA-------G----QIS-L---------------VTHALKFTHSD---A---------------K---G-S------SI----F----S---AE--------T-VAD---AKT---------------LST--A--AL------A-------Q--P--A-ID-A-VG----N----AA----A--L-----DVAKLL-Q-----T----E--H--D---------GDSL------------VAALQ-RGD-H---RALEAL---A---E----SP----EQLAQWLAGFQQV--F---TD--RQ----
PSSHKLAKQIYFPL--------------------------ENGE---YHLLSPLYSSSLAHALHQR-VN----A--V---R--F-G-----------------DE-AKAIR-Q-----AQ------------K--ANQWH--DQ-L-SIS-----YPY-LAVQNMGG-T----KPQ----------NISSLN--SSRS-----------GRSYLLSCA-P----PQWN--S---VE-KPPQQH----ESIF----RP-RGEVD--Y-----HT--RATLAQMQRFL------L--SV------K-----E-------------VEN---------
N----RD---IR--QQ----RL------RYLDQLIDQLFFYISSVQ--N-----L---P-----S---G----WS------A-----E------------SE--------LKR---AQQLWL-----D-------------PYR--AET-------------DTVFRR--ERE--AGD-WQ-KAAAYDFGRWLN-----LR----L---K-------H-------E-E-L------------IF---GEVER------REWST-A-------AL--FK-----------------R-----RMREMESAL----------------KEDL-------A----------------------
----------------
WP_103183968 ---------------------------------------------------ME--E---------NRLA--QFIV--S-YVT-NR-----K----QAK--------------------------------------------------------L---------D-------A-FDK--EAEK-----KR--A--------AL-----N---------GEALAAEEL-----------------------EL------------------------------A-EKR-RE---------------------IE--
QK-------------HE---VRAW-----LT---DA-----ASRA-------G----QIS-L----------------VTHALKFTHSD---A---------------K---G-S------SI----F----S---AE--------T-VAD---AKT---------------LST--A--AL------A-------Q--P--A-ID-A-VG----N----AA----A--L-----DVAKLL-Q-----T----E--H--D---------GDSL------------VAALQ-RGD-H---RALEAL---A---E----SP----EQLAQWLAGFQQV--F---TD--RQ----
PSSHKLAKQIYFPL--------------------------ENGE---YHLLSPLYSSSLAHALHQR-VN----A--V---R--F-G-----------------DE-AKAIR-Q-----AQ------------K--ANQWH--DQ-L-SIS-----YPY-LAVQNMGG-T----KPQ----------NISSLN--SSRS-----------GRSYLLSCA-P----PQWN--S---VE-KPPQQH----ESIF----RP-RGEVD--Y-----HT--RATLAQMQRFL------L--SV------K-----E-------------VEN---------
N----RD---IR--QQ----RL------RYLDQLIDQLFFYISSVQ--N-----L---P-----S---G----WS------A-----E------------SE--------LKR---AQQLWL-----D-------------PYR--AET-------------DTVFRR--ERE--AGD-WQ-KAVAYDFGRWLN-----LR----L---K-------H-------E-E-L------------IF---GEVER------REWST-A-------AL--FK-----------------R-----RMREMESAL----------------KEDL-------A----------------------
----------------
WP_104212802 -----------------------------------------------------MED-------------SRLT--KFIV--S-YIE-ER-----R----QDK--------------------------------------------------------L---------D-------KHDKT-VKTAR-------------------TE---LSGD----------ALADAEL-----------------------TL------------------------------A-EKR-RE---------------------IE--
QK-------------HE---VRAW-----LS---DA-----ASRA-------G----QIS-L----------------VTHALKFTHSD---A---------------K---G-T------SV----F----S-------------T-GTV----TEA--K-T---------LST--A--TL------A-------Q--P--A-ID-A-VG----N----AA----A--L-----DVAKLL-Q-----T----E--H--D---------GDSL------------VAALQ-RGD-H---RALEKL---A---E----NP----EQLAQWLAGFQQV--F---TD--RQ----
PSSHKLAKQIYFPI--------------------------ANGE---YHLLSPLYSSSLAQALHQR-IN----A--V---R--F-G-----------------EE-AKAIR-Q-----AQ------------K--ANQWH--DQ-P-SIS-----YPN-LAVQNMGG-T----KPQ----------NISSLN--SSRS-----------GRSYLLSSA-P----PQWN--S---IE-KPPQQH----DSIF----RP-RGEVD--Y-----HT--RATLAQMQRFL------L--SV------K-----E-------------VEN---------N--
--RD---IR--QQ----RL------RYLDQLIDQLFFYISSVQ--N-----L---P-----S---G----WS------A-----E------------SE--------LKR---AQQLWL-----D-------------PYR--AET-------------DTVFRR--ERE--AGD-WQ-KAVAYDFGRWLN-----RR----L---K-------H-------E-D-L------------IF---GEVER------REWST-A-------AL--FK-----------------R-----RMREMESAL----------------KEDL-------A----------------------
-----------
WP_039497834 -----------------------------------------------------MED-------------SRLT--KFIV--S-YIE-ER-----R----QDK--------------------------------------------------------L---------D-------KHDKT-VKTAR-------------------TE---LSGD----------ALADAEL-----------------------TL------------------------------A-EKR-RE---------------------IE---
-----QK------HE---VRAW-----LT---DA-----ASRA-------G----QIS-L---------------VTHALKFTHSD---A---------------K---G-T------SV----F----S-----T-------G-TVT---EAK--T-------------LST--A--TL------A-------Q--P--A-ID-A-VG----N----AA----A--L-----DVAKLL-Q-----T----E--H--D---------GDSL------------VAALQ-RGD-H---RALEKL---A---E----NP----EQLAQWLAGFQQV--F---TD--RQ----
PSSHKLAKQIYFPI--------------------------ANGE---YHLLSPLYSSSLAQALHQQ-IN----A--V---R--F-G-----------------EE-AKAIR-Q-----AQ------------K--ANQWH--EQ-L-SIS-----YPY-LAVQNMGG-T----KPQ----------NISSLN--SSRS-----------GRSYLLSSA-P----PQWN--S---IE-KPPQQH----DSIF----RP-RGEVD--Y-----HT--RATLAQMQRFL------L--SV------K-----E-------------VEN---------N--
-RD---IR--QQ----RL------RYLDQLIDQLFFYISSVQ--N-----L---P-----S---G----WS------A-----E------------SE--------LKR---AQQLWL-----D-------------PYR--AET-------------DTVFRR--ERE--AGD-WQ-KVVAYDFGRWLN-----RR----L---K-------H-------E-E-L------------IF---GEVER------REWST-A-------AL--FK-----------------R-----RMREMESAL----------------KEDL-------A----------------------
-----------
WP_039465888 ---------------------------------------------------MS--G---------EKLK--TFIL--S-YIS-TR-----K----QTK--------------------------------------------------------L---------D-------A-FDK--EAEK-----KR--A--------AL-----NG---------ETLAAEEL-----------------------EL------------------------------A-EKR-RE---------------------IE--
QK-------------HE---VRVW-----LS---DA-----ASRA-------G----QIS-L----------------VTHALKFTHSD---A---------------K---G-T------SV----F----S---TG--------T-------VTE--A--KT----------LST--A--TL------A-------Q--P--A-ID-A-VG----N----AA----A--L-----DVAKLL-Q-----T----E--H--D---------GDSL------------VAALQ-RGD-H---RALEKL---A---E----NP----EQLAQWLAGFQQV--F---TD--RQ----
PSSHKLAKQIYFPI--------------------------GNGE---YHLLSPLYSSSLAQALHQR-IN----A--V---R--F-G-----------------EE-AKAIR-Q-----AQ------------K--ANQWH--DQ-P-SIS-----YPN-LAVQNMGG-T----KPQ----------NISSLN--SSRS-----------GRSYLLSSA-P----PQWN--S---IE-KPPQQH----DSIF----RP-RGEVD--Y-----HT--RATLAQMQRFL------L--SV------K-----E-------------VEN---------N--
--RD---IR--QQ----RL------RYLDQLIDQLFFYISSVQ--N-----L---P-----S---G----WS------A-----E------------SE--------LKR---AQQLWL-----D-------------PYR--AET-------------DTVFRR--ERE--AGD-WQ-KAVAYDFGRWLN-----RR----L---K-------H-------E-D-L------------IF---GEIER------REWST-A-------AL--FK-----------------R-----RMREMESAL----------------KEDL-------A----------------------
-----------
WP_039555228 ---------------------------------------------------MG--S---------EKLT--EFIL--S-YIN-NR-----K----QAK--------------------------------------------------------L---------D-------A-FDK--EAEK-----KR--A--------VL-----SG---------EALAAEEL-----------------------DL------------------------------A-EKR-RE---------------------IE--
QK-------------HD---IRNW-----LS---DA-----ASRA-------G----QIS-L----------------VTHALKFTHSD---A---------------K---G-T------SV----F----S---TR--------T-------VTE--A--KT----------LST--A--TL------A-------Q--P--A-ID-A-VG----N----AA----A--L-----DVAKLL-Q-----T----E--H--D---------GDSL------------VAALQ-RDD-H---RALEAL---A---E----SP----EQLAQWLAGFQQV--F---TD--RQ----
PSSHKLAKQVYFPL--------------------------KNGE---YHLLSPLYSSSLAQALHQR-IN----T--V---R--F-G-----------------EE-AKAIR-Q-----AQ------------K--ANQWH--EQ-L-SIS-----YPY-LAVQNMGG-T----KPQ----------NISSLN--SSRS-----------GRSYLLSSA-P----PQWN--S---IE-KPPQQH----DSIF----RP-RGEVD--Y-----HT--RATLAQMQRFL------L--SV------K-----E-------------VEN---------N-
---RD---IR--QQ----RL------RYLDQLIDQLFFYISSVQ--N-----L---P-----S---G----WS------A-----E------------SE--------LKR---AQQLWL-----D-------------PYR--AET-------------DTVFRR--ERE--AGD-WQ-KAVAYDFGRWLN-----RR----L---K-------H-------E-E-L------------IF---GEVER------REWST-A-------AL--FK-----------------R-----RMREMESAL----------------KEDL-------A----------------------
-----------
WP_119874739 ---------------------------------------------------ME--E---------SRLT--QFIV--S-YIN-NR-----K----QAK--------------------------------------------------------L---------D-------A-FDK--EAEK-----KR--A--------AL-----SG---------ETLAAEEL-----------------------EL------------------------------A-EKR-RE---------------------IE--
QK-------------HE---VHAW-----LS---DA-----ASRA-------G----QIS-L----------------VTHALKFTHSD---A---------------K---G-T------SV----F----S---TG--------T-------VTE--A--KT----------LST--A--TL------A-------Q--P--A-ID-A-VG----N----AA----A--L-----DVAKLL-Q-----T----E--H--D---------GDSL------------VAALQ-RGD-H---RALEAL---A---E----GP----EQLAQWLAGFQQV--F---TD--RQ----
PSSHKLAKQIYFPL--------------------------KNGE---YHLLSPLYSSSLAQALHQR-IN----A--V---R--F-G-----------------EE-AKAIR-Q-----AQ------------K--ANQWH--DQ-F-SIS-----YPN-LAVQNMGG-T----KPQ----------NISSLN--SSRS-----------GRSYLLSCA-P----PQWN--S---VE-KPPQQH----DSIF----RP-RGEVD--Y-----HT--LATLAQMQRFL------L--SV------K-----E-------------VEN---------N-
---RD---IR--QQ----RL------RYLDQLIDQLFFYISSVQ--N-----L---P-----S---G----WS------A-----E------------SE--------LKR---AQQLWL-----D-------------PYR--AET-------------DTVFRR--ERE--AGD-WQ-KAVAYDFGRWLN-----RR----L---K-------H-------E-E-L------------IF---GEVER------REWST-A-------AL--FK-----------------R-----RMREMESAL----------------KEDL-------A----------------------
-----------
WP_039509851 ---------------------------------------------------ME--E---------SRLT--QFIV--S-YIN-NR-----K----QAK--------------------------------------------------------L---------D-------A-FDK--EAEK-----KR--A--------AL-----SG---------EALAAEEL-----------------------EL------------------------------A-EKR-RE---------------------IE--
QK-------------HE---VRAW-----LT---DA-----ASRV-------G----QIS-L----------------VTHALKFTHSD---A---------------K---G-T------SI----F----S---TE--------T-------VAD--A--KT----------LST--A--TL------A-------Q--P--A-ID-A-VG----N----AA----A--L-----DVAKLL-Q-----T----E--H--D---------GDSL------------VAALQ-RGD-H---RALETL---A---E----NP----EQLAQWLAGFQQV--F---TD--RQ----
PSSHKLAKQIYFPT--------------------------ENGE---YHLLSPLYSSSLAQALHQR-IN----A--V---R--F-G-----------------EE-AKAIR-Q-----AQ------------K--ANQWH--EQ-L-AIS-----YPN-LAVQNRGG-S----NKQ----------NISSLN--VTRN-----------GRSYLLSSA-P----PQWN--S---IE-KSPQQH----DSIF----RP-RGEVD--Y-----HT--RATLAQMQRFL------L--SV------K-----E-------------VEN---------N--
--RD---IR--QQ----RL------RYLDQLIDQLFFYISSVQ--N-----L---P-----S---G----WS------A-----E------------SE--------LKR---AQQLWL-----D-------------PYR--AET-------------DTVFRR--ERE--AGD-WQ-KAVAYDFGRWLN-----RR----L---K-------H-------E-D-L------------IF---GEVER------REWST-A-------AL--FK-----------------R-----RMREMESAL----------------KEDL-------A----------------------
-----------
WP_080163310 -----------------------------------------------------MEE-------------SRLT--QFIV--S-YIN-NR-----K----QAK--------------------------------------------------------L---------------DAFDK-EAEKK------------------RA---ALSG---------EALAAEEL-----------------------EL------------------------------A-EKR-RE---------------------IE---
-----QK------HE---VRAW-----LT---DA-----ASRV-------G----QIS-L---------------VTHALKFTHSD---A---------------K---G-T------SI----F----S---TE--------T-VAD---AKT---------------LST--A--TL------A-------Q--P--A-ID-A-VG----N----AA----A--L-----DVAKLL-Q-----T----E--H--D---------GDSL------------VAALQ-RGD-H---RALETL---A---E----NP----EQLAQWLAGFQQV--F---TD--RQ----
PSSHKLAKQIYFPT--------------------------ENGE---YHLLSPLYSSSLAQALHQR-IN----A--V---R--F-G-----------------EE-AKAIR-Q-----AQ------------K--ANQWH--EQ-L-AIS-----YPN-LAVQNMGG-T----KPQ----------NISSLN--SSRS-----------GRSYLLSSA-P----PQWN--S---IE-KSPQQH----DSIF----RP-RGEVD--Y-----HT--RATLAQMQRFL------L--SV------K-----E-------------VEN---------N--
--RD---IR--QQ----RL------RYLDQLIDQLFFYISSVQ--N-----L---P-----S---G----WS------A-----E------------SE--------LKR---AQQLWL-----D-------------PYR--AET-------------DTVFRR--ERE--AGD-WQ-KAVAYDFGRWLN-----RR----L---K-------H-------E-D-L------------IF---GEVER------REWST-A-------AL--FK-----------------R-----RMREMESAL----------------KEDL-------A----------------------
-----------
WP_052902937 -----------------------------------------------------MEE-------------SRLT--RFIV--S-YIN-NR-----K----QAK--------------------------------------------------------L---------------DAFDK-EAEKK-----RA--A--------FS----G-EA----------LAAEEL-----------------------EL------------------------------A-EKR-RE---------------------IE---
-----QK------HE---VRAW-----LS---DA-----ASRA-------G----QIS-L---------------VTHALKFTHSD---A---------------K---G-S------SI----F----S----A--------E-T-----VAD--A--KT----------LST--A--TL------A-------Q--P--A-ID-A-VG----N----AA----A--L-----DVAKLL-Q-----T----E--H--D---------GDSL------------VAALQ-RGD-H---SALEKL---A---E----NP----EQLAQWLVGFQQV--F---TD--RQ----
PSSHKLAKQIYFPI--------------------------ANGE---YHLLSPLYSSSLAQALHQR-IN----T--V---R--F-G-----------------DE-AKAIR-Q-----AQ------------K--ANQWH--EQ-L-SIS-----YPY-LAVQNMGG-T----KPQ----------NISSLN--SSRS-----------GRSYLLSSA-P----PQWN--S---IE-KPPQQH----DSIF----RP-RGEVD--Y-----HT--RATLAQMQRFL------L--SV------K-----E-------------VEN---------N--
-RD---IR--QQ----RL------RYLDQLIDQLFFYVASVQ--N-----L---P-----S---G----WS------A-----E------------SE--------LKR---AQQLWL-----D-------------PYR--AET-------------DTVFRR--ERE--AGD-WQ-KAVAYDFGRWLN-----RR----L---K-------H-------E-D-L------------IF---GEVER------REWST-A-------TL--FK-----------------R-----RMREMESVL----------------KEDL-------A----------------------
-----------
WP_039521518 -----------------------------------------------------MEE-------------SRLT--QFIV--S-YIN-NR-----K----QAK--------------------------------------------------------L---------------DAFDK-EAEKK-----RA--A--------FS----GEA----------LAAEEL-----------------------EL------------------------------A-EKR-RE---------------------IE---
-----QK------HE---VRAW-----LS---DA-----ASRA-------G----QIS-L---------------VTHALKFTHSD---A---------------K---G-S------SI----F----S----A---------ET---VAD--A-KT----------LST--A--TL------A-------Q--P--A-ID-A-VG----N----AA----A--L-----DVAKLL-Q-----T----E--H--D---------GDSL------------VAALQ-RGD-H---RALEKL---A---E----NP----EQLAQWLVGFQQV--F---TD--RQ----
PSSHKLAKQIYFPI--------------------------ANGE---YHLLSPLYSSSLAQALHQR-IN----A--V---R--F-G-----------------DE-AKAIR-Q-----AQ------------K--ANQWH--EQ-L-SIS-----YPY-LAVQNMGG-T----KPQ----------NISSLN--SSRS-----------GRSYLLSCA-P----PQWN--S---VE-KPPQQH----DSIF----RP-RGEVD--Y-----HT--RATLAQMQRFL------L--SV------K-----E-------------VEN---------N--
--RD---IR--QQ----RL------RYLDQLIDQLFFYVASVQ--N-----L---P-----S---G----WS------A-----E------------SE--------LKR---AQQLWL-----D-------------PYR--AKT-------------DTVFRR--ERE--AGD-WQ-KAVAYDFGRWLN-----RR----L---K-------H-------E-E-L------------IF---GEVER------REWST-A-------AL--FK-----------------R-----RMREMESTL----------------KEDL-------A----------------------
------------
WP_039278461 ---------------------------------------------------ME--E---------SRLT--QFIV--S-YIN-NR-----K----QAK--------------------------------------------------------L---------D-------A-FDK--EAEK-----KR--A--------VL-----SG---------EALAAEEL-----------------------EL------------------------------A-EKR-RE---------------------IE--
QK-------------HE---VRVW-----LT---DA-----ASRA-------G----QIS-L----------------VTHALKFTHSD---A---------------K---G-T------SV----F----S---TG--------T-------VTE--A--KT----------LST--A--TL------A-------Q--P--A-ID-A-VG----N----AA----A--L-----DVAKLL-Q-----T----E--H--D---------GDSL------------VAALQ-RGD-H---RALETL---A---E----SP----EQLAQWLAGFQQV--F---TD--RQ----
PSSHKLAKQIYFPT--------------------------ENGE---YHLLSPLYSSSLAQALHQR-IN----A--V---R--F-G-----------------EE-AKAIR-Q-----AQ------------K--ANQWH--DQ-L-SVS-----YPN-LAVQNMGG-T----KPQ----------NISSLN--SSRS-----------GRSYLLSSA-P----PQWN--S---IE-KPPQQH----DSIF----RP-RGEVD--Y-----HT--RATLAQMQRFL------L--SV------K-----E-------------AEN---------N-
---RD---IR--QQ----RL------RYLDQLIDQLFFYVASVQ--N-----L---P-----S---G----WS------A-----E------------SE--------LKR---AQQLWL-----D-------------PYR--AET-------------DTVFRR--ERE--AGD-WQ-KAVAYDFGRWLN-----RR----L---K-------H-------E-E-L------------IF---GEVER------REWST-A-------AL--FK-----------------R-----RMREMESTL----------------KEDL-------A----------------------
-------------
WP_039517607 ---------------------------------------------------ME--E---------SRLT--QFIV--S-YIN-NR-----K----QAK--------------------------------------------------------L---------D-------A-FDK--EAEK-----KR--V--------DL-----SG---------EVLAAEEL-----------------------EL------------------------------A-EKR-RE---------------------IE--
QK-------------HE---VRAW-----LT---DA-----ASRA-------G----QIS-L----------------VTHALKFTHSD---A---------------K---G-T------SV----F----S---AG--------T-------VTE--A--KT----------LST--A--TL------A-------Q--P--A-ID-A-VG----N----AA----A--L-----DVAKLL-Q-----T----E--H--D---------GDSL------------VAALQ-RGD-H---RALEAL---A---E----SP----EQLAQWLAGFQQV--F---TD--RQ----
PSSHKLAKQIYFPI--------------------------ANGE---YHLLSPLYSSSLAQALHQR-IN----T--V---R--F-G-----------------DE-AKAIR-Q-----AQ------------K--ANQWH--EQ-L-SIS-----YPY-LAVQNMGG-T----KPQ----------NISSLN--SSRS-----------GRSYLLSSA-P----PQWN--S---IE-KPPQQH----DSIF----RP-RGEVD--Y-----HT--RATLAQMQRFL------L--SV------K-----E-------------VEN---------N--
-RD---IR--HQ----RL------RYLDQLIDQLFFYVSSVQ--N-----L---P-----S---G----WS------A-----E------------SE--------LKR---AQQLWL-----D-------------PYR--AET-------------DTVFRR--ERE--AGD-WQ-KAVAYDFGRWLN-----RR----L---K-------H-------E-D-L------------IF---GEIER------REWST-A-------AL--FK-----------------R-----RMREMESAL----------------KEDL-------A----------------------
----------
WP_010279272 -----------------------------------------------------MEE-------------SRLT--RFIV--S-YIN-NR-----K----QAK--------------------------------------------------------L---------------DAFDK-EAEKK-------R--A--------TL---SGED----------LAAEEL-----------------------DL------------------------------A-EKR-RE---------------------IE---
-----QK------HE---VRAW-----LS---DA-----ASRA-------G----QIS-L---------------VTHALKFTHSD---A---------------K---G-T------SV----F----S----T--------G-TVT---EAK--T-------------LST--A--TL------A-------Q--P--A-ID-A-VG----N----AA----A--L-----DVAKLL-Q-----T----E--H--D---------GDSL------------VAALQ-RGD-H---HALEKL---A---E----NP----EQLAQWLVGFQQV--F---TD--RQ----
```

[illegible]

WP_119872007
[illegible]

WP_094432323
[illegible]

WP_014916577
[illegible]

WP_039558306
[illegible]

WP_065505800
[illegible]

WP_074055397
[illegible]

WP_060431063
[illegible]

WP_101427675
[illegible]

WP_086580877
[illegible]

WP_047025098
[illegible]

WP_116286779
[illegible]

WP_115435441
[illegible]

WP_103085864
[illegible]

WP_049208621
[illegible]

WP_049198327
[illegible]

WP_110593244
[illegible]

WP_038879097
[illegible]

WP_004089534
[illegible]

WP_083321355
[illegible]

WP_115102172
[illegible]

WP_130961902
[illegible]

WP_130976116
[illegible]

WP_039187557
[illegible]

WP_064573470
[illegible]

WP_040044711
[illegible]

WP_025798009
[illegible]

WP_046360261
[illegible]

WP_130995116
[illegible]

WP_049582103
[illegible]

WP_046394379
[illegible]

WP_065390688
[illegible]

WP_112877865
[illegible]

WP_011146070
[illegible]

WP_113042067
[illegible]

WP_058589481
[illegible]

WP_036775829
[illegible]

WP_015834854
[illegible]

WP_046976569
[illegible]

WP_133814645
[illegible]

WP_036774582
[illegible]

WP_065824320
[illegible]

LSSHKLAKQLYFPV-------NS------------------ECDE---YHLLGPLFSSSLADAMYQR-II----K--T---R--F-G-----------------EL-SKEIN-K-----AK-------------K--EESWH--PG-T-RVI-----YPN-TVVQKFGG-A----KPQ---------NISYLN--SVRR-----------GNIWLLSCA-S----PNWL--N---IT-RPPMER----KSIF----ER-RSEFV--S-----LV--RETTRKMQQYL------F--IV------Q-----D-------------IEN---------N----
RK---IR--KL----RQ------EFVDQVIDILFNYVAGIQ--N-----L-F-E----IK---G----WS------A-----S---D---D-----CE--------LKR---AQQLWL-----D-------------PYR--CQL-------------DKEFRS--ERE--RGD-WQ-KEIVADFSYWLN-----QN-----L---K-------H-------E-R-L------------EM---ELSEC------HEWAS----------V--FK-----------------K-----RLREFEDEL----------------PEVL-------L-----------------------
--------
WP_054475749 ------------------------------------------------------------MNE-------------NGLS--RFIV--D-YIE-SC-----KQSKLQSK----------------------------------------------------------------------------------L---------A-------T-FDK--KAEK------K--L--------AK-----LT-----T--MEDITLVKQ----------------------------EL-------------------------------------A-EQR-KE------------------------
QE---------ER------YR---VRNW-----LD---DA-----ANRA-------K----KIS-R----------------ATHAPKYTHSD---V-----------------K---G-S------SV----F----S----V-------LP---N---ARE--N--TK--Y------LST--S--AL------H-------K--P--A-ID-I-VA----N----AQ----D--L-----DVAKLL-Q-----T----E--Y--Q---------EDSL------------MECLN-RED-Y---SALAEL---A---E----NE----QQLEQWVSGFRQA--L---SD--EM----
LSSHKLAKQLYFPV-------NS------------------ECDE---YHLLSPLFSSSLADAMHQR-II----E--A---R--F-G-----------------EL-SKEIN-K-----AK-------------K--EERWH--PE-T-RVI-----YPN-TAVWNIGG-T----KPQ---------NISYLN--NVRG-----------GRVWLLSCA-S----PNWL--S---IT-KPPMGH----RSIF----ER-RSEFV--S-----LV--REIIGKMQQYL------F--VV------Q-----H-------------IEN---------N----
EK---IR--EH----RQ------GFVDQIIDILFNYIAGIQ--N-----L-F-E----IK---G----WS------A-----S---D---D-----CE--------LKR---AQQLWL-----D-------------PYR--CQL-------------DKEFRS--ERE--RGD-WK-KEIAADFSYWLN-----QS-----L---K-------H-------E-R-L------------EM---ELSER------NEWAS----------V--FK-----------------K-----RLREFEDEL----------------PEVP-------L-----------------------
------
WP_036840434 ------------------------------------------------------------MNE-------------DGLS--RFIV--D-YIE-SC-----KQSKLQSK----------------------------------------------------------------------------------L---------A-------V-FDK--KAEK------K--L--------AK-----LT-----T--MEDITLVKQ----------------------------EL-------------------------------------A-EQR-KE------------------------
LE---------EW------YE---VRNW-----LD---DA-----ANRA-------K----KTS-R----------------ATHAPKYSHSD---A-----------------K---G-S------SI----F----S----V-------SP---N---AKE--N--TE--Y------LST--S--AL------Y-------K--P--A-ID-V-VA----D----AK----Y--L-----DVAKLL-Q-----T----E--Y--R---------GDSL------------IECLN-RRD-Y---SALAEL---A---E----NE----TQLKRWVSGFRQA--L---SD--ET----
LSSHKLAKQLYFPI-------NS------------------DGDE---YHLLSPLFSSSLASAVHHR-IV----D--A---R--F-S-----------------EL-SKEIN-K-----SR-------------K--EESWH--PG-T-RVI-----YPN-TAVQNFGG-T----KPQ---------NISYLN--SGRG-----------GSTWLLSCA-S----PNWL--S---IT-KPPMGR----RSIF----DR-RSEFV--S-----LA--RETIRKMQQYL------F--VV------Q-----D-------------IEN---------N----
RK---IR--KL----RQ------EFVDQIIDILFNYVADIQ--N-----L-F-E----LK---G----WS------A-----S---D---D-----CE--------LKR---AQQLWL-----D-------------PYR--CQL-------------DKEFRF--ERE--GGD-WK-KEIAADFSYWLN-----QS-----L---K-------H-------E-R-L------------EM---ELPER------HEWAS----------E--FK-----------------K-----RLREFEDEL----------------PEVM-------P-----------------------
--------
WP_036848911 ------------------------------------------------------------MNE-------------NGLS--RFIV--D-YIE-SC-----K----QSK-----------------------------------------------------------------------------------L---------A-------V-FDK--KAEK------K--L--------AK-----LT-----T--MEDITLVKE----------------------------EL-------------------------------------A-EQR-KE------------------------LE---
------ER------YE---VRNW-----LD---DA-----ANRA-------K----KTS-R----------------ATHAPKYSHSD---A-----------------K---G-S------SI----F----S----V--------S-SDV---KEN--T--EY---------LST--S--AL------Y-------K--P--A-ID-V-VA----D----AK----Y--L-----DVAKLL-Q-----T----E--Y--R---------GDSL------------MECLN-RRD-Y---SALAEL---A---E----NE----KQLKRWVSGFRQV--L---SD--ET----
LSSHKLAKQLYFPI-------NS------------------DGDE---YHLLSPLFSSSLASAVHHR-IV----D--A---R--F-S-----------------EL-SKEIN-K-----SR-------------K--EESWH--PG-T-RVV-----YPN-TAVQNFGG-T----KPQ---------NISYLN--SGRG-----------GSTWLLSCA-S----PNWL--S---IT-KPPMER----RSIF----DR-RSEFV--S-----LA--RETIGKMRQYL------F--VV------Q-----D-------------IEN---------N----
-RK---IK--KL----RQ------EFVDQIIDILFNYVAGIQ--N-----L-F-E----LK---G----WS------A-----S---D---D-----CE--------LKR---TQQLWL-----D-------------PYR--CQL-------------DKEFRF--ERE--GGD-WK-KEIAADFSYWLN-----QS-----L---K-------H-------E-R-L------------EM---ELPER------REWAS----------E--FK-----------------K-----RLREFEDEL----------------PEVM-------P-----------------------
---------
WP_015834867 ------------------------------------------------------------MNE-------------NRLS--RFIV--D-YIG-SC-----KQLKLQSK----------------------------------------------------------------------------------L---------E-------D-FDR--KAGE------K--L--------AK-----LT-----S--SEDITMIQQ----------------------------KL-------------------------------------A-DKR-KE------------------------
LE---------ES------YE---VRNW-----LD---NA-----ANHA-------K----KIS-R----------------ATHAPKYTHSN---A-----------------K---G-S------SI----L----L----V-------LS---D---AKE--N--TQ--Y------LST--S--VL------Y-------K--P--V-ID-I-VV----D----AK----Y--L-----EVAKLL-Q-----T----E--Y--Q---------GDSL------------IECLS-RRD-Y---SALAEL---A---E----NE----QQLERWVNGFRQA--L---SD--ET----
RSSHKLAKQLYFPV-------NG------------------ECGE---YHLLSPLFSSSLIHAMYQR-IA----E--A---R--F-S-----------------EL-SKDIN-K-----AK-------------K--EERWH--PG-T-RVI-----YPN-TAIQNMGG-T----KPQ---------NISYLN--NVLG-----------GRVWLLSCS-P----PNWQ--S---NI-KLPERQ----RSIF----EQ-RSQFV--R-----LS--YDTVRLMRKYL------F--RA------Q------------------KR---------T---
-EE---IR--QK----RQ------AYIDEIIDILFDYVASVQ--D-----Q-V-E----FK---G----W--------------S---A---N-----CD--------LKR---SQQLWL-----D-------------PYR--CQQ-------------DEIFKF--ERE--GGD-WK-KEIADDFGLWLN-----NE-----L---R-------H-------E-R-L------------EV---GLTER------NEWAS----------V--FK-----------------K-----RLREFEDEL----------------PEVP-------L-----------------------
------
WP_118984460 ------------------------------------------------------------MNE-------------NGLS--RFII--D-YIE-SR-----K----QPK-----------------------------------------------------------------------------------L---------E-------A-FDK--EAEK------K--L--------AK-----LT-----S--AEDISLVKQ----------------------------KL-------------------------------------D-EQR-NE------------------------LE---
-----KR------YE---VRNW-----LD---AA-----ANRA-------K----QIS-L-----------------VTHALKFTHSD---A-----------------K---G-S------SI----F----A----I-------SE---N---GKK--N--IE--Y------LST--T--AL------S-------K--P--A-ID-A-VG----N----AA----A--L-----DIAKLL-Q-----T----E--Y--Q---------EDSL------------VVCLN-RGD-Y---SALAEL---A---E----NE----QQLERWVNGFRQA--L---SD--ET----
LSSHKLAKQLYFPV-------DG------------------ECAE---YHLLSPLFSSSLAHAMNQR-IA----E--A---R--F-G-----------------ES-SKEIN-K-----AK-------------K--EERWH--SE-T-RVI-----YPN-TAVQNMGG-T----KPQ---------NISYLN--SVRG-----------GRVWLLSCS-A----PNWQ--S---NI-NLPERH----RSIF----EQ-RSQFV--L-----LS--YRTVRLMRQYL------F--RV------Q------------------KR---------T-
---EE---IR--RN----RQ------AYVDEIIDILFNYVASIQ--N-----Q-V-E----FK---G----W--------------S---A---N-----CD--------LKR---SQQLWL-----D-------------PYR--CQQ-------------DEIFKF--ERE--SGD-WK-KEIADDFGLWLN-----NE-----L---R-------H-------E-R-L------------EV---GLTER------NEWAS----------V--FK-----------------K-----RLREFEDEL----------------PEVP-------L-----------------------
---------
WP_036768763 ------------------------------------------------------------MNE-------------NGLS--RFIT--D-YIG-SR-----K----QSK-----------------------------------------------------------------------------------L---------E-------V-FDK--EAEK------K--L--------AK-----LT-----S--SEDITLVQQ----------------------------EI-------------------------------------A-EQR-KA------------------------LE---
------EH------YE---VRSW-----LD---DA-----ANRA-------K----QIS-L-----------------VTHALKFTHSD---A-----------------R---G-S------SI----F----A----A-------SS---N---TTE--N--IK--Y------LST--A--TL------Y-------K--P--V-ID-S-VG----N----AA----A--L-----DIAKLL-Q-----T----E--Y--Q---------EDSL------------VACLN-RGN-Y---SALAEL---A---E----NE----QQLERWVSGFRQA--L---SD--ET----
LSSHKLAKQLYFPV-------NS------------------ECDE---YHLLSPLFSSSLAHAMHQR-IA----K--A---R--F-G-----------------ES-SKEIN-K-----AK-------------K--EERWH--SE-V-RVS-----YPN-TAVQNIGG-T----KPQ---------NISYLN--SVRG-----------GRAWLLSCS-A----PTWQ--S---NI-KLPKRR----RSIF----EQ-PGQFV--R-----LS--YRTVRSMRQYL------F--RV------Q------------------KH---------T-
---EK---IR--QN----RQ------AYVDEIIDILFNYVASVQ--D-----Q-V-E----FK---G----W--------------S---A---D-----CN--------LKR---SQQLWL-----D-------------PYR--CQQ-------------DEIFKF--ERE--GGD-WK-KEIADDFSLWLN-----NE-----L---R-------H-------E-R-L------------EF---GLTER------NEWAS----------V--FK-----------------K-----RLREFEDEL----------------PEVP-------L-----------------------
----------
WP_065823625 ------------------------------------------------------------MNE-------------NGLS--RFIT--D-YIG-SR-----K----QSK-----------------------------------------------------------------------------------L---------E-------V-FDK--EAEK------K--L--------AK-----LT-----S--SEDITLVQQ----------------------------EI-------------------------------------A-EQR-KA------------------------LE---
------EH------YE---VRSW-----LD---DA-----ANRA-------K----QIS-L-----------------VTHALKFTHSD---A-----------------R---G-S------SI----F----A----A-------SS---N---TTE--N--IK--Y------LST--A--TL------Y-------K--P--V-ID-S-VG----N----AA----A--L-----DIAKLL-Q-----T----E--Y--Q---------EDSL------------VACLN-RGN-Y---SALAEL---A---E----NE----QQLERWVSGFRQA--L---SD--ET----
LSSHKLAKQLYFPV-------NS------------------ECGE---YHLLSPLFSSSLAHAMHQR-IA----E--A---R--F-G-----------------ES-SKEIN-K-----AK-------------K--EERWH--SE-V-RVS-----YPN-TAVQNIGG-T----KPQ---------NISYLN--SVRG-----------GRAWLLSCS-A----PTWQ--S---NI-KLPKRR----RSIF----EQ-PGQFV--R-----LS--YRTVRSMRQYL------F--RV------Q------------------KH---------T-
---EK---IR--QN----RQ------VYVDEIIDILFNYVASVQ--D-----Q-V-E----FK---G----W--------------S---A---D-----CN--------LKR---SQQLWL-----D-------------PYR--CQQ-------------DEIFKF--ERE--GGD-WK-KEIADDFSLWLN-----NE-----L---R-------H-------E-R-L------------EF---GLTER------NEWAS----------V--FK-----------------K-----RLREFEDEL----------------PEVP-------L-----------------------
----------
WP_036807245 ------------------------------------------------------------MNE-------------NGLS--RFII--D-YIK-SR-----K----QPK-----------------------------------------------------------------------------------L---------E-------A-FDK--EAEK------K--L--------AK-----LT-----S--SEDMTSVRE----------------------------EI-------------------------------------A-EQR-KE------------------------LE---
------EQ------YE---VRNW-----LD---DA-----ANRA-------K----QIS-L-----------------VTHALKFTHSD---A-----------------R---G-S------SI----F----A----T-------SS---N---TTG-------K--Y------LST--A--TL------Y-------K--P--V-ID-S-VG----N----AA----A--L-----DVAKLL-Q-----T----E--Y--Q---------GDSL------------VECLS-CGD-Y---SVLAEL---A---E----ND----QQLERWVNGFRQA--L---SD--ET----
LSSHKLAKQLYFPI-------NN------------------ESGE---YHLLSPLFSSSLAHAMYQR-IA----E--A---R--F-G-----------------ES-SKEIN-K-----AK-------------K--EERWH--PE-T-RVI-----YPN-TAVQNIGG-T----KPQ---------NISYLN--SVRG-----------GRAWLLSCS-A----PNWQ--S---NI-KLPERH----RSIF----EQ-RSQFV--R-----LS--YRTVRLMRQYL------F--RA------Q------------------KR---------T----
EE---IR--QN----RQ------AYIDEIIDILFSYVASVQ--D-----Q-V-E----FK---G----W--------------S---A---N-----CK--------LKR---SQQLWL-----D-------------PYR--CQQ-------------DEIFKF--ERE--DGD-WK-KEIADDFGLWLN-----NE-----L---R-------H-------E-R-L------------EF---GLTER------NEWAS----------V--FK-----------------E-----RLREFEYEL----------------PEVP-------L-----------------------
-----
WP_113026306 ------------------------------------------------------------MNE-------------NGLS--RFIT--D-YIG-SR-----K----QPK-----------------------------------------------------------------------------------L---------E-------A-FDK--EAEK------R--L--------AK-----LT-----S--SEDITLVQQ----------------------------EI-------------------------------------A-EQR-KA------------------------LE---
------EH------YE---VRNW-----LD---DA-----ANRA-------K----QIS-L-----------------VTHALKFTHSD---A-----------------R---G-S------SI----F----A----T-------SS---N---TTE--N--IK--Y------LST--A--TL------Y-------K--P--V-ID-S-VG----N----AA----A--L-----DVAKLL-Q-----T----E--Y--Q---------EDSL------------VACLN-RGD-Y---SALAEL---A---E----NE----QQLERWVNGFRQA--L---SD--ET----
LSSHKLAKQLYFPI-------NN------------------ESGE---YHLLSPLFSSSLAHAMHQR-IA----E--A---R--F-G-----------------ES-SKEIN-K-----AK-------------K--EERWH--PE-T-RVI-----YPN-TAVQNIGG-T----KPQ---------NISYLN--SVRG-----------GRAWLLSCS-A----PNWQ--S---NI-KLPERH----RSIF----EQ-RSQFV--R-----LS--YRTVRLMRQYL------F--RA------Q------------------KR---------T----
EE---IR--QN----RQ------AYIDQIIDILFNYVASVQ--D-----Q-V-E----FK---G----W--------------S---A---N-----CK--------LKR---SQQLWL-----D-------------PYR--CQQ-------------DEIFKF--ERE--GGD-WK-KEIADDFGLWLN-----NE-----L---R-------H-------E-R-L------------EF---GLTER------NEWAS----------V--FK-----------------K-----RLREFEDEL----------------PEVP-------L-----------------------
------
WP_110087106 ------------------------------------------------------------MNE-------------NGLS--RFIT--D-YIG-NR-----K----QPK-----------------------------------------------------------------------------------L---------E-------A-FDK--EAEK------R--L--------AK-----LT-----S--SEDITLVQQ----------------------------EI-------------------------------------A-EQR-KA------------------------LE---
------EH------YE---VRNW-----LD---DA-----ANRA-------K----QIS-L-----------------VTHALKFTHSD---A-----------------R---G-S------SI----F----A----T-------SS---N---TTG-------K--Y------LST--A--TL------Y-------K--P--V-ID-S-VG----N----AA----A--L-----DVAKLL-Q-----T----E--Y--Q---------EDSL------------VACLN-RGD-Y---SALAEL---A---E----NE----QQLERWVNGFRQA--L---SD--ET----
LSSHKLAKQLYFPI-------NN------------------ESGE---YHLLSPLFSSSLAHAMYQR-IA----E--A---R--F-G-----------------ES-SKEIN-K-----AK-------------K--EERWH--PE-T-RVI-----YPN-TAVQNIGG-T----KPQ---------NISYLN--SVRG-----------GRAWLLSCS-A----PNWQ--S---NI-KLPERH----RSIF----EQ-RSQFV--R-----LS--YRTVRLMRQYL------F--RA------Q------------------KR---------T----
EE---IR--QN----RQ------AYIDEIIDILFSYVASVQ--D-----Q-V-E----FK---G----W--------------S---A---N-----CK--------LKR---SQQLWL-----D-------------PYR--CQQ-------------DEIFKF--ERE--GGD-WK-KEIADDFGLWLN-----NE-----L---R-------H-------E-R-L------------EF---GLTER------NEWAS----------V--FK-----------------E-----RLREFEYEL----------------PEVP-------L-----------------------
-----
WP_040152319 ------------------------------------------------------------MNE-------------NELS--RFIT--D-YIG-SR-----K----QPK-----------------------------------------------------------------------------------L---------E-------A-FDK--EAEK------R--L--------AK-----LT-----S--SEDITLVQQ----------------------------EI-------------------------------------A-EQR-KA------------------------LE---
-----EH------YE---VRNW-----LD---DA-----ANRA-------K----QIS-L-----------------VTHALKFTHSD---A-----------------R---G-S------SI----F----A----T-------SS---N---TTG-------K--Y------LST--A--TL------Y-------K--P--V-ID-S-VG----N----AA----A--L-----DVAKLL-Q-----T----E--Y--Q---------EDSL------------VACLN-RGD-Y---SALAEL---A---E----ND----QQLERWVNGFRQA--L---SD--ET----
LSSHKLAKQLYFPI-------NN------------------ESGE---YHLLSPLFSSSLAHAMYQR-IT----E--A---R--F-G-----------------ES-SKEIN-K-----AK-------------K--EERWH--PE-T-RVI-----YPN-TAVQNIGG-T----KPQ---------NISYLN--SVRG-----------GRAWLLSCS-A----PNWQ--S---NI-KLPERH----RSIF----EQ-RSQFV--R-----LS--YRTVRLMRQYL------F--RA------Q------------------KR---------T----
EE---IR--QN----RQ------AYIDEIIDILFSYVASVQ--D-----Q-V-E----FK---G----W--------------S---A---N-----CK--------LKR---SQQLWL-----D-------------PYR--CQQ-------------DEIFKF--ERE--GGD-WK-KEIADDFGLWLN-----NE-----L---R-------H-------E-R-L------------EF---GLTER------NEWAS----------V--FK-----------------E-----RLREFEYEL----------------PEVP-------L-----------------------
-----
WP_112896710 ------------------------------------------------------------MNE-------------NGLS--QFII--D-YIE-SR-----K----HPK-----------------------------------------------------------------------------------L---------E-------A-FDK--EAEK------K--L--------AK-----LT-----S--SEDMMLVRE----------------------------EI-------------------------------------A-EQR-KV------------------------LE---
------EH------YE---VRNW-----LD---DA-----ANRA-------K----QIS-L-----------------VTHALKFTHSD---A-----------------R---G-S------SI----F----A----T-------SS---N---TTG-------K--Y------LST--A--TL------Y-------K--P--V-ID-S-VG----N----AA----T--L-----DVAKLL-Q-----T----E--Y--Q---------KDSL------------VECLT-RGD-Y---SALAEL---A---E----ND----QQLERWVNGFRQA--L---SD--ET----
LSSHKLAKQLYFPV-------NG------------------ECDE---YHLLSPLFSSSLAHAMHQR-IA----E--A---R--F-G-----------------ES-SKEIN-K-----AK-------------K--EERWH--PE-T-RVI-----YPN-TAVQNIGG-T----KPQ---------NISYLN--SVRG-----------GRSWLLSCS-A----PNWQ--S---NI-KLPERH----RSIF----EQ-RSQFV--L-----LS--YRTVRLMRQYL------F--RV------Q------------------KR---------T---
-EE---IR--RN----RQ------AYVDEIIDILFNYVASVQ--N-----Q-A-E----FK---G----W--------------S---A---N-----CD--------LKR---SQQLWL-----D-------------PYR--CQR-------------DEIFKF--ERE--GGD-WK-KEIADDFGLWLN-----NE-----L---R-------H-------E-R-L------------EV---GLAER------HEWAF----------V--FK-----------------K-----RLREFEDEL----------------PEVP-------L-----------------------
--------
WP_139657135 ------------------------------------------------------------MNE-------------NGLS--RFII--D-YIE-SR-----K----QPK-----------------------------------------------------------------------------------L---------E-------A-FDK--EAEK------K--L--------AK-----LT-----S--VEDITLVKQ----------------------------EI-------------------------------------A-EQR-KA------------------------LE----
----EY------YE---ISNW-----LD---DA-----ANRA-------K----QIS-L-----------------VTHALKFTHSD---A-----------------R---G-S------SI----F----A----T-------SS---N---TTG--N--TK--Y------LST--A--TL------Y-------K--P--V-ID-S-VG----N----AA----A--L-----DIAKLL-Q-----T----E--Y--Q---------EDSL------------VECLN-RGD-Y---SALAEL---A---E----ND----QQLECWVNGFRQA--L---SD--ET----
LSSHKLAKQLYFPI-------NS------------------ECDG---YHLLSPLFSSSLAHATHQR-IA----E--A---R--F-G-----------------ES-SKEIN-K-----AK-------------K--EERWH--PE-T-RVI-----YPN-TAVQNIGG-T----KPQ---------NISYLN--SVRG-----------GRAWLLSCA-A----PNWQ--G---NI-KLPERH----RSIF----EQ-RSQFV--L-----LS--YRTVRLMRQYL------F--RV------Q------------------KR---------T----
EE---VR--RN----RQ------AYVDEIIDILFNYVASVQ--N-----Q-V-E----FK---G----W--------------S---A---N-----CD--------LKR---SQQLWL-----D-------------PYR--CQQ-------------DEIFKF--ERE--GGD-WK-KEIADDFGLWLN-----NE-----L---R-------H-------E-Q-L------------EV---GLTER------NEWAS----------V--FK-----------------K-----RLREFEDEL----------------PEVP-------L-----------------------
--------
WP_088372075 ------------------------------------------------------------MNE-------------NGLS--RFII--D-YIE-SR-----K----QPK-----------------------------------------------------------------------------------L---------E-------T-FDK--EAEK------K--L--------AK-----LN-----S--VEDITLVKQ----------------------------EI-------------------------------------A-EQR-KA------------------------LE----
----EY------YE---ISNW-----LD---DA-----ANRA-------K----QIS-L-----------------VTHALKFTHSD---A-----------------R---G-S------SI----F----A----T-------SS---N---TTG--N--TK--Y------LST--A--TL------Y-------K--P--V-ID-S-VG----N----AA----A--L-----DIAKLL-Q-----T----E--Y--Q---------EDSL------------VECLN-RGD-Y---SALAEL---A---E----NE----QQLERWVNGFRQA--L---SD--ET----
LSSHKLAKQFYFPI-------NS------------------ECDG---YHLLSPLFSSSLAHAMHQR-IA----E--A---R--F-G-----------------ES-SKEIN-K-----AK-------------K--DECWH--PE-T-RVI-----YPS-TAVQNIGG-T----KPQ---------NISYLN--SVRG-----------GRAWLLSCS-A----PNWQ--S---NI-KLPKRH----RSIF----EQ-RSQFV--R-----LS--YRTVRLMRQYL------F--RA------Q------------------KR---------T---
-EE---IR--QN----RQ------AYVDEIIDILFNYVASVQ--D-----Q-V-E----FK---G----W--------------S---A---N-----CD--------LKR---SHQLWL-----D-------------PYR--CQR-------------DEIFKF--ERE--GGD-WK-KEIADDFGLWLN-----NE-----L---R-------H-------E-Q-L------------EV---GLTER------NEWAS----------V--FK-----------------K-----RLREFEDEL----------------PEVP-------L-----------------------
--------
WP_021325938 ------------------------------------------------------------MNE-------------SKLS--RFII--D-YIE-SR-----K----QPK-----------------------------------------------------------------------------------L---------E-------A-FDK--EAEK------K--L--------AK-----LT-----S--VEDITLVKQ----------------------------EL-------------------------------------A-EQR-NE------------------------LE----
----KH------YE---VRNW-----LD---DA-----ANRA-------K----QIS-L-----------------VTHALKFTHSD---A-----------------R---G-S------SI----F----A----T-------SS---N---AAE--N--TE--Y------LST--T--TL------Y-------K--P--V-ID-S-VG----N----AA----A--L-----DIAKLL-Q-----T----E--Y--Q---------GDSL------------VASLN-RSD-Y---SALVEL---A---E----DE----QQLERWINGFRQA--L---SD--ET----
LSSHKLAKQLYFPV----------------------------SSSE---YHLLSPLFSSSLAHAIYQR-IA----E--V---R--F-G-----------------EL-SKEIN-K-----AK-------------R--EERWH--PE-A-RVI-----YPN-TSVQNTGG-T----KPQ---------NISYLN--SIRS-----------GRTWLLSCS-A----PNWQ--S---ST-KLPKRH----RSIF----EQ-RSQFT--L-----LS--YTTVRLMRQFL------F--SA------Q------------------KR---------T----
EE---TR--RS----RQ------AYVEEIIDILFNYVASVQ--N-----Q-V-E----FK---G----WS------A-----S---D---D-----CE--------LKR---AQQLWL-----D-------------PYR--CQL-------------DKEFRF--ERE--GGD-WK-KEIAADFSYWMN-----QS-----L---K-------H-------E-R-L------------EM---ELPER------REWVS----------E--FK-----------------K-----RLREFEDEL----------------PEVI-------P-----------------------
-----------
WP_132354259 ------------------------------------------------------------MNE-------------SGLS--RFII--D-YIE-SR-----K----QPK-----------------------------------------------------------------------------------L---------E-------V-FDK--EAER------R--L--------TK-----LT-----S--AEEILVIQQ----------------------------EI-------------------------------------A-EQR-KE------------------------LE----
---EH------YE---TRNW-----LD---DA-----ANRA-------K----QIS-L-----------------VTHALKFTHSD---A-----------------K---G-S------SI----F----A----T-------S-VSK---EEN--I--EY---------LST--T--AL------Y-------K--P--A-ID-A-VG----N----AA----A--L-----DIAKLL-Q-----T----E--Y--Q---------GDSL------------VASLN-RSD-F---SALVEL---A---E----DE----QQLERWINGFRQA--L---SD--ET----
LSSHKLAKQLYFPV-------SS------------------NE---YHLLSPLFSSSLVHAMYQR-IA----E--A---R--F-G-----------------ES-SKEIN-K-----AK-------------K--EERWH--SE-A-RVI-----YPN-TAVQNIGG-T----KPQ---------NISYLN--SVRG-----------GRTWLLSCS-A----PNWQ--G---ST-KLPKRY----RSIF----EQ-RSQFT--L-----LS--YTTVRLMRQFL------F--ST------Q------------------KR---------T----
EE---TR--RS----CR------AYVEEIIEILFNYVASVQ--D-----Q-V-E----FK---G----W--------------S---K---D-----CD--------LKR---AQQLWL-----D-------------PYR--CQQ-------------DKAFQF--ERE--NGD-WK-KEIADDFGFWLN-----NE-----L---R-------H-------E-R-L------------EV---GLTER------NEWAS----------V--FK-----------------N-----RLREFEDEL----------------PEGP-------L-----------------------
---------
WP_061807483 ------------------------------------------------------------MAD-------------NGLT--ALIV--E-YIA-SR-----K----QPK-----------------------------------------------------------------------------------L---------E-------A-LEK--DLAK------K--L--------AG-----LT-----E--AEEIAVARQ----------------------------SA-------------------------------------L-EQR-QE------------------------LE---
------RN------YT---PRHW-----LT---DA-----ASRA-------G----QIS-L-----------------VTHALKFTHSD---A-----------------K---G-S------SI----F----A----A-------DS-------ESH--A--TD--R------LSS--A--SL------S-------Q--P--A-ID-A-VG----N----AA----A--L-----DVAKLL-Q-----L----E--Y--E---------GESL------------IACLS-RGD-S---TPLAAL---A---E----NP----EQLELWMAGFKQV--L---VN--KN----
LSSHTLAKQLYFPV-------GP------------------NE---YHLLSPLFSSSLAQAVHQR-IS----D--V---R--F-S-----------------DT-AKECY-Q-----AL-------------K--AGEWY--PE-P-RVI-----YPK-TAVLNMGG-T----KPQ---------NISYLN--SVRG-----------GRVWLLPCS-A----PVWL--T---VS-KLPLNQ----QSIF----QE-RKEFS--A-----LA--RDAVWQLKQHL------R--DA------N-----K-----------V-PGN---------
N----VE---IR--QQ----RS------AYVDEVIDILFSYVAGIQ--N-----R-Q-E----WQ---C----WS------Q-----R---E---A-----CQ--------LSR---AQQLWL-----D-------------PWR--MKQ-------------DEDFRF--ERE--GGG-WK-KEIALDFGVWLN-----HQ-----L---K-------S-------E-E-M------------KF---SEVER------REWST-A-------PL--FK-----------------R-----RLREFEAEF----------------KGDF-------SWEL-----------
--------------------------
WP_129937976 ------------------------------------------------------------MNK-------------NGFS--ELIV--E-YIA-SR-----K----QPK-----------------------------------------------------------------------------------L---------E-------A-LEK--DLAK------R--L--------AG-----LT-----E--AEEIAVAKQ----------------------------TA-------------------------------------L-EQR-QE------------------------LE---
------KN------YT---PRDW-----LT---DA-----ASRA-------G----QIS-L-----------------VTHALKFTHSD---A-----------------K---G-S------SI----F----A----------T-GSA---SLA--T--DR---------LSS--A--SL------S-------Q--P--A-ID-A-VG----N----AA----A--L-----DVAKLL-Q-----L----E--Y--E---------GESL------------IACLS-RGD-S---TPLAVF---A---E----NP----QQLELWVAGFKQV--L---AN--KT----
LSSHTLAKQLYFPV-------GP------------------NE---YHLLSPLFSSSLAHAMHQR-IS----D--A---R--F-S-----------------DA-AKESY-Q-----AL-------------K--AGEWS--PS-P-RVI-----YPK-TAVLNMGG-T----KPQ---------NISYLN--SVRG-----------GRAWLLPCS-A----PVWL--S---VN-KPPLNQ----HSIF----QE-RKAFS--A-----MA--RDTVWQLKQYL------R--GV------N-----Q-----------S-SEN---------
T----AE---IR--QQ----RS------AYLDEIIDILFGYVAGIQ--N-----C-E-D----WQ---C----WS------Q-----S---E---G-----CQ--------LSR---AQQLWL-----D-------------PWR--MNQ-------------DEDFRF--ERE--GDD-WK-KEIALDFGVWLN-----HQ-----L---K-------S-------E-E-M------------KF---SEVER------REWST-A-------PL--FK-----------------R-----RLREFEAEF----------------KGDF-------AWEL-----------
------------------------
WP_129937489 ------------------------------------------------------------MNK-------------NGFS--ELIV--E-YIA-SR-----K----QPK-----------------------------------------------------------------------------------L---------E-------A-LEK--DLAK------R--L--------AG-----LT-----E--AEEIAVAKQ----------------------------TA-------------------------------------L-EQR-QE------------------------LE---
------KN------YT---PRDW-----LT---DA-----ASRA-------G----QIS-L-----------------VTHALKFTHSD---A-----------------K---G-S------SI----F----A----------T-GSA---SLA--T--DR---------LSS--A--SL------S-------Q--P--A-ID-A-VG----N----AA----A--L-----DVAKLL-Q-----L----E--Y--E---------GESL------------IACLS-RGD-S---TPLAVF---A---E----NP----QQLELWVAGFKQV--L---AN--KT----
LSSHTLAKQLYFPV-------GP------------------NE---YHLLSPLFSSSLAHAMRQR-IS----D--A---R--F-S-----------------DA-AKESY-Q-----AL-------------K--AGEWS--PS-P-RVI-----YPK-TAVLNMGG-T----KPQ---------NISYLN--SVRG-----------GRAWLLPCS-A----PVWL--S---VN-KPPLNQ----HSIF----QE-RKAFS--A-----MA--RDTVWQLKQYL------R--GV------N-----Q-----------S-SEN---------
T----AE---IR--QQ----RS------AYLDEIIDILFGYVAGIQ--N-----C-E-D----WQ---C----WS------Q-----S---E---G-----CQ--------LSR---AQQLWL-----D-------------PWR--MNQ-------------DEDFRF--ERE--GDD-WK-KEIALDFGVWLN-----HQ-----L---K-------S-------E-E-M------------KF---SEVER------REWST-A-------PL--FK-----------------R-----RLREFEAEF----------------KGDF-------AWEL-----------
------------------------
WP_142009394 ------------------------------------------------------------MAD-------------DGLT--ALIV--E-YIA-SR-----K----QPK-----------------------------------------------------------------------------------L---------E-------A-LEK--EAAK------K--L--------AG-----LT-----D--PEEIAVAKQ----------------------------TV-------------------------------------L-EQL-QT------------------------LE---
-----KN------YA---PRNW-----LT---DA-----AGRA-------G----QIS-L-----------------VTHALKFTHSD---A-----------------K---G-S------SV----F----S----V-------GS-------SEQ--T--IN--T------LAT--N--TL------S-------Q--P--A-VD-A-VG----N----AA----A--L-----DVAKLL-Q-----L----E--Y--K---------GESL------------IACLS-RDD-T---APLEKL---A---E----NP----QQLEQWVAGFKQV--L---TN--KN----
PSSHSLAKQIYFPV-------GQ------------------GE---YHLLSPLFSSSLAQAMHQR-IS----E--A---R--F-S-----------------EA-SKEIY-Q-----AC-------------K--AGEWH--PE-P-RVI-----YPK-TAALNMGG-T----KPQ---------NISYLN--SVRG-----------GHVWLLPCA-A----PVWQ--N---QN-KPPLNQ----SSIF----QE-RKAFS--A-----MA--KDAVWQLKQYL------R--SL------D-----P--------------QDK---------
N----ME---IR--EQ----RA------AYVDEIIDILFCYVAGIQ--N-----R-E-D----WQ---C----WS------M-----K---E---E-----CQ--------LSR---AQQLWL-----D-------------PWR--MKH-------------DKDFQF--ERE--GGD-WK-KDIALDFGVWLN-----HQ-----L---K-------T-------D-E-L------------KF---SEVER------REWST-A-------SL--FK-----------------R-----RLREFEQEF----------------KGDF-------AWVP-----------
--------------------------
WP_024487012 ------------------------------------------------------------MSQ-------------NGLT--ALIV--E-YIA-NR-----K----QPK-----------------------------------------------------------------------------------L---------E-------A-LEK--ETAK------K--L--------AG-----LA-----D--PEEIAVAKQ----------------------------TA-------------------------------------L-EQR-QE------------------------LE---
-----KN------YA---PRNW-----LT---DA-----AGRA-------G----QIS-L-----------------VTHALKFTHSD---A-----------------K---G-S------SI----Y----S--------------S-GSQ---EQT--T--NT---------LST--S--SL------S-------Q--P--A-VD-A-VG----N----AA----A--L-----DVAKLL-Q-----L----E--Y--E---------GESL------------IACLA-RDN-T---VPLAEL---A---E----DP----QQLEQWVTGFRQV--L---ND--KH----
LSSHTLAKQIYFPI-------AP------------------GE---YHLLSPLFSSSLAHAMHSR-IS----E--A---R--F-S-----------------DA-SKEIY-Q-----AF-------------K--AGDWH--PE-P-RVI-----YPK-TAVVNMGG-T----KPQ---------NISYLN--SVRG-----------GRAWLLPCA-A----PIWQ--S---QN-KPPLNQ----NSIF----QE-RKAFS--A-----MA--RDAVWQLKQYL------R--SL------N-----P--------------QDR---------N--
--VE---IR--QQ----RA------AFVDEIVDTLFSYVAGIQ--N-----R-E-D----WQ---C----WS------M-----E---E---G-----CQ--------LSR---AQQLWL-----D-------------PWR--IKQ-------------DKDFQF--ERE--GGD-WK-KEIAIDFGVWLN-----HQ-----L---K-------T-------E-E-M------------KF---SEVER------REWST-T-------SL--FK-----------------R-----RLREFEQEF----------------KEDF-------A-----------------
------------------
WP_124232892 ------------------------------------------------------------MAD-------------EGFT--LLIT--A-YIR-AR-----G----QPK-----------------------------------------------------------------------------------L---------E-------A-FDK--AAEK------T--L--------VE-----LS-----D--SAQISLARQ----------------------------NL-------------------------------------A-SQR-RE------------------------LE---
------QR------YE---VQRW-----LS---DA-----ASRA-------G----QIS-L-----------------VTHALKFTHSD---A-----------------K---G-S------SR----F----E----A-------SS-------GQT--D--AH--Y------LST--A--TL------H-------K--P--S-ID-A-VG----N----AA----A--L-----DVAKLL-Q-----I----E--Y--Q---------GESL------------IAALS-RGD-Y---SPLAPL---A---E----SE----QQLTAWVEGFQQV--L---HD--RQ----
ASSHKLAKQLYFPV-------GD------------------NQ---YHLLGPLFSSSLAHALHQR-IV----E--A---R--F-S-----------------DN-AKEIN-A-----AF-------------R--EKRWH--SA-P-RIG-----YLN-MASQNPGG-S----RPQ---------NISYLN--SVRG-----------GRSWLLSCA-A----PVWH--S---QP-VPPVKH----TSVF----NR-YSAFS--Q-----MA--RSAVWQLREFL------L--RV------K-----G-------------QPN---------
-N----ME---IR--QR----RL------AYTDEIIDMLFNYAAGVQ--H-----D-D-M----QQ---G----WS------A-----Q---D---A-----CQ--------LKR---SQQLWL-----D-------------PGR--AKG-------------DEQFCA--ERE--RGE-WQ-KELAVDFGLWLN-----RQ-----L---K-------H-------K-D-L------------NM---SETER------REWST-A-------SL--FK-----------------Q-----RLREFEQAV----------------KEEW-------Q-----------------
--------------------------
WP_034937159 ------------------------------------------------------------MAD-------------EGFT--LLIT--S-YVK-DR-----R----EPR-----------------------------------------------------------------------------------L---------E-------A-FDK--EAEK------K--L--------TN-----LT-----D--AQQIALVRQ----------------------------ET-------------------------------------A-SQR-RE------------------------LE---
-------QR------YE---IKSW-----LT---DA-----ASRA-------E----QIN-L-----------------VTHALKFTHSD---A-----------------R---G-S------SR----F----D-----------A-SAR---EDD--A--TY---------LST--A--TL------H-------K--P--A-ID-A-VG----N----AA----A--L-----DVARLL-Q-----T----E--Y--Q---------GESL------------IETLR-RGD-Y---SPLAPL---A---E----TE----GQLMQWVNGFMQA--L---TD--KQ----
PGSHKLAKQLYFPV-------GE------------------ND---YHLLGPLFSSSLAHALHQR-IA----D--A---R--F-S-----------------DK-AKEIN-A-----AF-------------K--AKRWH--AE-P-RVS-----YLN-MASQNPGG-S----KPQ---------NISYLN--SVRR-----------GRNWLLSCA-P----PCWQ--N---QP-RAPTKH----KSIF----NH-YSEFS--Q-----MA--RSTIWQLREFL------L--RV------K-----D-------------APN---------
N----RE---IR--HR----RL------AYTDEIIDILFSYVAGVE--N-----M-A-V----EH---G----WS------A-----G---D---A-----CQ--------LKR---SQQLWL-----D-------------VGR--AKE-------------DSQFRA--ERE--SGE-WQ-REIAIDFGVWLN-----RQ-----L---K-------H-------D-E-L------------KV---SEVER------REWS-----------SL--FK-----------------Q-----RLGEFEQAV----------------REES-------Q-----------------
------------------
WP_145591464 ------------------------------------------------------------MSD-------------DGLA--AFIV--S-YIE-SR-----K----QPK-----------------------------------------------------------------------------------L---------E-------A-FDK--EAEK------K--L--------AG-----LT-----A--AEDIARVQQ----------------------------EI-------------------------------------A-LQR-QE------------------------LA---
------AR------YE---VRNW-----LT---DA-----ASRA-------G----QIS-L-----------------VTHALKYIHSD---A-----------------K---G-S------SV----F----N----L-------PA-------AEN--N--AI--Y------LSS--A--TL------A-------K--P--A-ID-A-VG----N----AA----A--L-----DVAKLL-Q-----T----E--H--N---------GDSL------------LACLH-RAD-Y---SPLAAL---A---E----NP----LQLTQWVEGFKQA--L---VD--KQ----
LTSHKLAKQLYFPV-------GP------------------NQ---YHLLSPLFSSSLAHAMHQR-IT----D--A---R--F-G-----------------DQ-PKAIS-A-----AY-------------K--AKTWH--PQ-T-RVK-----YLK-TAMQNIGG-T----KPQ---------NISYLN--SVRG-----------GKVWLLPAG-A----PQWH--S---SH-KPPTKH----TSIF----HE-YSEFT--A-----LA--RDTTWQMQRYL------L--SV------K-----H-------------LSN---------T-
---LE---IR--AS----RL------AYADEIIDILFNYVAGVQ--N-----L-K-I----AD---G----WS------N-----D---P---D-----CN--------LKL---SQQIWL--------------------------------------------DPALKQ--QRE--LGD-WK-KEVAQDFGFWLN-----RR-----L---Q-------H-------D-K-L------------VF---GEVER------REWAS----------L--FK-----------------Q-----RLREFEQGS----------LFEQDNLEAE-------S-----------------
----------
WP_042562387 ------------------------------------------------------------MSD-------------EGLA--AFIV--S-YIE-SR-----K----QPK-----------------------------------------------------------------------------------L---------E-------A-FDK--EAEK------K--L--------AG-----LT-----A--AEDIARVQQ----------------------------EN-------------------------------------A-LQR-QE------------------------LA-
-------AR------YE---TRTW-----LT---DA-----ASRA-------G----QIS-L-----------------VTHALKYIHSD---A-----------------R---G-S------SV----F----N----L-------PA-------AEN--D--AI--Y------LSS--A--TL------A-------K--P--A-ID-A-VG----N----AA----A--L-----DVAKLL-Q-----T----E--H--N---------GDSL------------LACLH-RAD-Y---SPLAEL---A---E----NP----AQLTLWVEGFKQA--L---VD--KQ----
LTSHKLAKQLYFPV-------GP------------------NQ---YHLLSPLFSSSLAHAMHQR-IT----D--A---R--F-G-----------------DQ-SKEAK-A-----AY-------------K--AKTWH--PQ-P-RVM-----YLD-TAVQNIGG-T----KPQ---------NISYLN--SVRG-----------GKVWLLPAG-A----PQWR--S---SR-KPPTKH----SSIF----ND-NSEFT--A-----LA--YSTARQMQQYL------L--SV------K-----P-------------LTN---------
T----ME---IR--AS----RL------AYSDEIIDILFNYVAEVQ--N-----L-K-I----AA---G----WS------Q-----E---D---G-----CN--------LKL---SQQIWL--------------------------------------------DPAFKQ--QRE--LGD-WK-KEVAQEFGYWLN-----RR-----L---K-------H-------D-E-L------------VF---GEVER------REWAS----------L--FK-----------------L-----RLREFEQGS----------RSEPESLEAE-------S-----------------
-------------
WP_145586643 ------------------------------------------------------------MSD-------------EGLA--AFIV--P-YIE-GR-----K----QPK-----------------------------------------------------------------------------------L---------E-------A-FDK--EAEK------R--L--------AG-----LT-----A--AEDIARVQQ----------------------------EN-------------------------------------A-LQR-QE------------------------LA-
--------AR------YE---TRSW-----LT---DA-----ASRA-------G----QIS-L-----------------VTHALKYIHSD---A-----------------R---G-S------SV----F----N----L-------PA-------AED--N--AV--Y------LSS--A--TL------A-------K--P--A-ID-A-VG----N----AA----A--L-----DVAKLL-Q-----T----E--H--N---------GDSL------------LACLH-RAD-Y---SPLAEL---A---E----NP----AQLTLWVEGFKQA--L---VD--KQ----
LTSHKLAKQLYFPV-------GP------------------NQ---YHLLSPLFSSSLAHAMHQR-IT----D--A---R--F-G-----------------DQ-SKEAK-A-----AY-------------K--AKTWH--PQ-P-RVM-----YLD-TAVQNIGG-T----KPQ---------NISYLN--SVRG-----------GKVWLLPAG-A----PQWR--S---SR-KPPTKH----SSIF----ND-NSEFT--A-----LA--YSTARQMQQYL------L--SV------K-----P-------------LTN---------
T----ME---IR--AS----RL------AYSDEIIDILFNYVAEVQ--N-----L-K-I----AA---G----WS------Q-----E---D---G-----CN--------LKL---SQQIWL--------------------------------------------DPAFKQ--QRE--LGD-WK-KEVAQEFGYWLN-----RR-----L---K-------H-------D-E-L------------VF---GEVER------REWAS----------L--FK-----------------L-----RLREFEQGS----------RSEPESLEAE-------S-----------------
-------------
WP_145587151 ------------------------------------------------------------MQE-------------NGLA--AFIV--S-YIE-GR-----K----QPK-----------------------------------------------------------------------------------L---------E-------A-FDK--EAEK------K--L--------AG-----LT-----A--AEDIARVQQ----------------------------EN-------------------------------------A-LQR-QE------------------------LA-
--------AR------YE---TRSW-----LT---DA-----ASRA-------G----QIS-L-----------------VTHALKYIHSD---A-----------------R---G-S------SV----F----N----L-------PA-------AEN--N--AV--Y------LSS--A--TL------A-------K--P--A-ID-A-VG----N----AA----A--L-----DVAKLL-Q-----T----E--H--N---------GDSL------------LACLH-RAD-Y---SPLAEL---A---E----NP----AQLTLWVEGFKQA--L---VD--KQ----
LTSHKLAKQLYFPV---------------------------EPNQ---YHLLSPLFSSSLAHAMHQR-IT----D--A---R--F-G-----------------DQ-SKEAK-A-----AY-------------K--AKTWH--PQ-P-RVM-----YLD-TAVQNIGG-T----KPQ---------NISYLN--SVRG-----------GKVWLLPAG-A----PQWR--S---SR-KPPTKH----SSIF----ND-NSEFT--A-----LA--YSTARQMQQYL------L--SV------K-----P-------------LTN---------
T----MG---IR--AS----RL------AYSDEIIDILFNYVAEVQ--N-----L-K-I----AA---G----WS------Q-----E---D---G-----CN--------LKL---SQQIWL--------------------------------------------DPAFKQ--QRE--LGD-WK-KEVAQEFGYWLN-----RR-----L---K-------H-------D-E-L------------VF---GEVER------REWAS----------L--FK-----------------Q-----RLREFEQGS----------LSEPESLEAE-------S-----------------
--------------
WP_057633516 ------------------------------------------------------------MQE-------------NGLA--AFIV--S-YIE-GR-----K----QPK-----------------------------------------------------------------------------------L---------E-------A-FDK--EAEK------R--L--------AG-----LT-----A--AEDIARVQQ----------------------------EN-------------------------------------A-LQR-QE------------------------LA-
--------AR------YE---TRSW-----LT---DA-----ASRA-------G----QIS-L-----------------VTHALKYIHSD---A-----------------R---G-S------SV----F----N----L-------PA-------AEN--N--AV--Y------LSS--A--TL------A-------K--P--A-ID-A-VG----N----AA----A--L-----DVAKLL-Q-----T----E--H--N---------GDSL------------LACLH-RAD-Y---SPLAEL---A---E----NP----AQLTLWVEGFKQA--L---VD--KQ----
LTSHKLAKQLYFPV-------GP------------------NQ---YHLLSPLFSSSLAHAMHQR-IT----D--A---R--F-G-----------------DQ-SKEAK-A-----AY-------------K--AKTWH--PQ-P-RVM-----YLD-TAVQNIGG-T----KPQ---------NISYLN--SVRG-----------GKVWLFPAG-A----PQWR--S---SR-KPPTKH----STIF----ND-NSEFT--A-----LA--YSTARQMQQYL------L--SV------K-----P-------------LTN---------
T----ME---IR--AS----RL------AYSDEIIDILFNYVAEVQ--N-----L-K-I----AA---G----WS------Q-----E---D---G-----CN--------LKL---SQQIWL--------------------------------------------DPAFKQ--QRE--LGD-WK-KEVAQEFGYWLN-----RR-----L---K-------H-------D-E-L------------VF---GEVER------REWAS----------L--FK-----------------Q-----RLREFEQGS----------LSEPESLEAE-------S-----------------
--------------
WP_100191356 ------------------------------------------------------------MQE-------------NGLA--AFIV--S-YIE-GR-----K----QPK-----------------------------------------------------------------------------------L---------E-------A-FDK--EAEK------R--L--------AG-----LT-----A--AEDIARVQQ----------------------------EN-------------------------------------A-LQR-QE------------------------LA-
--------AR------YE---TRSW-----LT---DA-----ASRA-------G----QIS-L-----------------VTHALKYIHSD---A-----------------R---G-S------SV----F----N----L-------PA-------AEN--N--AV--Y------LSS--A--TL------A-------K--P--A-ID-A-VG----N----AA----A--L-----DVAKLL-Q-----T----E--H--N---------GDSL------------LACLH-RAD-Y---SPLAEL---A---E----NP----AQLTLWVEGFKQA--L---VD--KQ----
LTSHKLAKQLYFPV-------GP------------------NQ---YHLLSPLFSSSLAHAMHQR-IT----D--A---R--F-G-----------------DQ-SKEAK-A-----AY-------------K--AKTWH--PQ-P-RVM-----YLD-TAVQNIGG-T----KPQ---------NISYLN--SVRG-----------GKVWLLPAG-A----PQWR--S---SR-KPPTKH----SSIF----ND-NSEFT--A-----MA--YSTARQMQQYL------L--SV------K-----P-------------LTS---------
T----ME---IR--AS----RL------AYSDEIIDILFNYVAEVQ--N-----L-K-I----AA---G----WS------Q-----E---D---G-----CN--------LKL---SQQIWL--------------------------------------------DPAFKQ--QRE--LGD-WK-KEVAQEFGYWLN-----RR-----L---K-------H-------D-E-L------------VF---GEVER------REWAS----------L--FK-----------------Q-----RLREFEQGS----------LSEPESLEAE-------S-----------------
--------------
WP_004389634 ------------------------------------------------------------MQE-------------NGLA--AFIV--S-YIE-GR-----K----QPK-----------------------------------------------------------------------------------L---------E-------A-FDK--EAEK------R--L--------AG-----LT-----A--AEDIARVQQ----------------------------EN-------------------------------------A-LQR-QE------------------------LA-
--------AR------YE---TRSW-----LT---DA-----ASRA-------G----QIS-L-----------------VTHALKYIHSD---A-----------------R---G-S------SV----F----N----L-------PA-------AED--N--AV--Y------LSS--A--TL------A-------K--P--A-ID-A-VG----N----AA----A--L-----DVAKLL-Q-----T----E--H--N---------GDSL------------LACLH-RAD-Y---SPLAEL---A---E----NP----AQLTLWVEGFKQA--L---VD--KQ----
LTSHKLAKQLYFPV-------GP------------------NQ---YHLLSPLFSSSLAHAMHQR-IT----D--A---R--F-G-----------------DQ-SKEAK-A-----AY-------------K--AKTWH--PQ-P-RVM-----YLD-TAVQNIGG-T----KPQ---------NISYLN--SVRG-----------GKVWLLPAG-A----PQWR--S---SR-KPPTKH----SSIF----ND-NSEFT--A-----LA--YSTARQMQQYL------L--SV------K-----P-------------LTN---------
T----MG---IR--AS----RL------AYSDEIIDILFNYVAEVQ--N-----L-K-I----AA---G----WS------Q-----E---D---G-----CN--------LKL---SQQIWL--------------------------------------------DPAFKQ--QRE--LGD-WK-KEVAQEFGYWLN-----RR-----L---K-------H-------D-E-L------------VF---GEVER------REWAS----------L--FK-----------------Q-----RLREFEQGS----------LSEPESLEAE-------S-----------------
--------------
WP_145511680 ------------------------------------------------------------MQE-------------NGLA--AFIV--S-YIE-GR-----K----QPK-----------------------------------------------------------------------------------L---------E-------A-FDK--EAEK------R--L--------AG-----LT-----A--AEDIARVQQ----------------------------EN-------------------------------------A-LQR-QE------------------------LA-
--------AR------YE---TRSW-----LT---DA-----ASRA-------G----QIS-L-----------------VTHALKYIHSD---A-----------------R---G-S------SV----F----N----L-------PA-------AEN--N--AV--Y------LSS--A--TL------A-------K--P--A-ID-A-VG----N----AA----A--L-----DVAKLL-Q-----T----E--H--N---------GDSL------------LACLH-RAD-Y---SPLAEL---A---E----NP----AQLTLWVEGFKQA--L---VD--KQ----
LTSHKLAKQLYFPV-------GP------------------SQ---YHLLSPLFSSSLAHAMHQR-IT----D--A---R--F-G-----------------DQ-SKEAK-A-----AY-------------K--AKTWH--PQ-P-RVM-----YLD-TAVQNIGG-T----KPQ---------NISYLN--SVRG-----------GKVWLLPAG-A----PQWR--S---SR-KPPTKH----SSIF----ND-NSEFT--A-----LA--YSTARQMQQYL------L--SV------K-----P-------------LTN---------
T----ME---IR--AS----RL------AYSDEIIDILFNYVAEVQ--N-----L-K-I----AA---G----WS------Q-----E---D---G-----CN--------LKL---SQQIWL--------------------------------------------DPAFKQ--QRE--LGD-WK-KEVAQEFGYWLN-----RR-----L---K-------H-------D-E-L------------VF---GEVER------REWAS----------L--FK-----------------Q-----RLREFEQGS----------LSEPESLEAE-------S-----------------
--------------
WP_100286405 ------------------------------------------------------------MQE-------------NGLA--AFIV--S-YIE-GR-----K----QPK-----------------------------------------------------------------------------------L---------E-------A-FDK--EAEK------R--L--------AG-----LT-----A--AEDIARVQQ----------------------------EN-------------------------------------A-LQR-QE------------------------LA-
--------AR------YE---TRSW-----LT---DA-----ASRA-------G----QIS-L-----------------VTHALKYIHSD---A-----------------R---G-S------SV----F----N----L-------P-AAE---NNA-----VY-------LSS--A--TL------A-------K--P--A-ID-A-VG----N----AA----A--L-----DVAKLL-Q-----T----E--H--N---------GDSL------------LACLH-RAD-Y---SPLAEL---A---E----NP----AQLTLWVEGFKQA--L---VD--KQ----
LTSHKLAKQLYFPV-------GP------------------NQ---YHLLSPLFSSSLAHAMHQR-IT----D--A---R--F-G-----------------DQ-SKEAK-A-----AY-------------K--AKTWH--PQ-P-RVM-----YLD-TAVQNIGG-T----KPQ---------NISYLN--SVRG-----------GKVWLLPAG-A----PQWR--S---SR-KPPTKH----SSIF----ND-NSEFT--A-----LA--YSTARQMQQYL------L--SV------K-----P-------------LTN---------
T----ME---IR--AS----RL------AYSDEIIDILFNYVAEVQ--N-----L-K-I----AA---G----WS------Q-----E---D---G-----CN--------LKL---SQQIWL--------------------------------------------DPAFKQ--QRE--LGD-WK-KEVAQEFGYWLN-----RR-----L---K-------H-------D-E-L------------VF---GEVER------REWAS----------L--FK-----------------Q-----RLREFEQGS----------LSEPESLEAE-------S-----------------
--------------
WP_099528622 ------------------------------------------------------------MSD-------------EGLA--AFIV--S-YIE-GR-----K----QPK-----------------------------------------------------------------------------------L---------E-------A-FDK--EAEK------K--L--------AG-----LT-----A--AEDIARVQQ----------------------------EN-------------------------------------A-LQR-QE------------------------LA-
--------AR------YE---TRSW-----LT---DA-----ASRA-------G----QIS-L-----------------VTHALKYIHSD---A-----------------R---G-S------SV----F----N----L-------PA-------AEN--N--AV--Y------LSS--A--TL------A-------K--P--A-ID-A-VG----N----AA----A--L-----DVAKLL-Q-----T----E--H--N---------GDSL------------LACLH-RAD-Y---SPLAEL---A---E----NP----AQLTLWVEGFKQA--L---VD--KQ----
LTSHKLAKQLYFPV-------GP------------------NQ---YHLLSPLFSSSLAHAMHQR-IT----D--A---R--F-G-----------------DQ-SKEAK-A-----AY-------------K--AKTWH--PQ-P-RVM-----YLD-TAVQNIGG-T----KPQ---------NISYLN--SVRG-----------GKVWLLPAG-A----PQWR--S---SR-KPPTKH----SSIF----ND-NSEFT--A-----LA--YSTARQMQQYL------L--SV------K-----P-------------LTN---------
T----ME---IR--AS----RL------AYSDEIIDILFNYVAEVQ--N-----L-K-I----AA---G----WS------Q-----E---D---G-----CN--------LKL---SQQIWL--------------------------------------------DPAFKQ--QRE--LGD-WK-KEVAQEFGYWLN-----RR-----L---K-------H-------D-E-L------------VF---GEVER------REWAS----------L--FK-----------------Q-----RLREFEQGS----------LSEPESLEAE-------S-----------------
--------------
WP_050118533 ------------------------------------------------------------MSD-------------EGLA--AFIV--S-YIE-SR-----K----QPK-----------------------------------------------------------------------------------L---------E-------A-FDK--EAEK------R--L--------AG-----LT-----A--AEDIARVQQ----------------------------EN-------------------------------------A-LQR-QE------------------------LA-
-------AR------YE---TRSW-----LT---DA-----ASRA-------G----QIS-L-----------------VTHALKYIHSD---A-----------------R---G-S------SV----F----N----L-------PA-------AEN--N--AV--Y------LSS--A--TL------A-------K--P--A-ID-A-VG----N----AA----A--L-----DVAKLL-Q-----T----E--H--N---------GDSL------------LACLH-RAD-Y---SPLAEL---A---E----NP----AQLTLWVEGFKQA--L---VD--KQ----
LTSHKLAKQLYFPV-------GP------------------NQ---YHLLSPLFSSSLAHAMHQR-IT----D--A---R--F-G-----------------DQ-SKEAK-A-----AY-------------K--AKTWH--PQ-P-RVM-----YLD-TAVQNIGG-T----KPQ---------NISYLN--SVRG-----------GKVWLLPAG-A----PQWR--S---SR-KPPTKH----SSIF----ND-NSEFT--A-----LA--YSTARQMQQYL------L--SV------K-----P-------------LTS---------
T----ME---IR--AS----RL------AYSDEIIDILFNYVAEVQ--N-----L-K-I----AA---G----WS------Q-----E---D---G-----CN--------LKL---SQQIWL--------------------------------------------DPAFKQ--QRE--LGD-WK-KEVAQEFGYWLN-----RR-----L---K-------H-------D-E-L------------VF---GEVER------REWAS----------L--FK-----------------Q-----RLREFEQGS----------LSEPESLEAE-------S-----------------
--------------
WP_049556072 ------------------------------------------------------------MSD-------------EGLA--AFIV--S-YIE-SR-----K----QPK-----------------------------------------------------------------------------------L---------E-------A-FDK--EAEK------R--L--------AG-----LT-----A--AEDIARVQQ----------------------------EN-------------------------------------A-LQR-QE------------------------LA-
-------AR------YE---TRSW-----LT---DA-----ASRA-------G----QIS-L-----------------VTHALKYIHSD---A-----------------R---G-S------SV----F----N----L-------PA-------AEN--N--AV--Y------LSS--A--TL------A-------K--P--A-ID-A-VG----N----AA----A--L-----DVAKLL-Q-----T----E--H--N---------GDSL------------LACLH-RAD-Y---SPLAEL---A---E----NP----AQLTLWVEGFKQA--L---VD--KQ----
LTSHKLAKQLYFPV-------GP------------------NQ---YHLLSPLFSSSLAHAMHQR-IT----D--A---R--F-G-----------------DQ-SKEAK-A-----AY-------------K--AKTWH--PQ-P-RVM-----YLD-TAVQNIGG-T----KPQ---------NISYLN--SVRG-----------GKVWLFPAG-A----PQWR--S---SR-KPPTKH----SSIF----ND-NSEFT--A-----LA--YSTARQMQQYL------L--SV------K-----P-------------LTN---------
T----ME---IR--AS----RL------AYSDEIIDILFNYVAEVQ--N-----L-K-I----AA---G----WS------Q-----E---D---G-----CN--------LKL---SQQIWL--------------------------------------------DPAFKQ--QRE--LGD-WK-KEVAQEFGYWLN-----RR-----L---K-------H-------D-E-L------------VF---GEVER------REWAS----------L--FK-----------------Q-----RLREFEQGS----------LSEQESLEAE-------S-----------------
--------------
WP_050111539 ------------------------------------------------------------MSD-------------EGLA--AFIV--S-YIE-SR-----K----QPK-----------------------------------------------------------------------------------L---------E-------A-FDK--EAEK------R--L--------AG-----LT-----A--AEDIARVQQ----------------------------EN-------------------------------------A-LQR-QE------------------------LA-
-------AR------YE---TRSW-----LT---DA-----ASRA-------G----QIS-L-----------------VTHALKYIHSD---A-----------------R---G-S------SV----F----N----L-------PA-------AEN--N--AV--Y------LSS--A--TL------A-------K--P--A-ID-A-VG----N----AA----A--L-----DVAKLL-Q-----T----E--H--N---------GDSL------------LACLH-RAD-Y---SPLAEL---A---E----NP----AQLTLWVEGFKQA--L---VD--KQ----
LTSHKLAKQLYFPV-------GP------------------NQ---YHLLSPLFSSSLAHAMHQR-IT----D--A---R--F-G-----------------DQ-SKEAK-A-----AY-------------K--AKTWH--PQ-P-RVM-----YLD-TAVQNIGG-T----KPQ---------NISYLN--SVRG-----------GKVWLFPAG-A----PQWR--S---SR-KPPTKH----SSIF----ND-NSEFT--A-----LA--YSTARQMQQYL------L--SV------K-----P-------------LTN---------
T----ME---IR--AS----RL------AYSDEIIDILFNYVAEVQ--N-----L-K-I----AA---G----WS------Q-----E---D---G-----CN--------LKL---SQQIWL--------------------------------------------DPAFKQ--QRE--LGD-WK-KEVAQEFGYWLN-----RR-----L---K-------H-------D-E-L------------VF---GEVER------REWAS----------L--FK-----------------Q-----RLREFEQGS----------LSEQESLEAE-------S-----------------
--------------
WP_049613699 ------------------------------------------------------------MSD-------------EGLT--AFTV--S-YIE-GR-----K----QPK-----------------------------------------------------------------------------------L---------E-------A-FDK--DAEK------R--L--------AG-----LT-----A--AADIALAQQ----------------------------EI-------------------------------------A-QQR-QE------------------------LI---
------AR------YE---VRAW-----LT---DA-----ANRA-------G----QIS-L-----------------VTHALKFTHSD---A-----------------R---G-S------SV----F----N----L-------PT-------AEN--D--AL--Y------LST--A--TL------A-------K--P--A-ID-A-VG----N----AA----A--L-----DVAKLL-Q-----T----E--H--N---------GDSL------------LASLN-RAD-Y---SSFAEL---A---E----NS----QQLALWVDGFQQV--L---VD--KQ----
LTSHKLAKQLYFPV-------GP------------------DQ---YHLLSPLFSSSLAHALHQR-IA----D--A---R--F-G-----------------DT-SKAIY-S-----AY-------------K--ADKWH--PQ-P-RVI-----YPK-TAVQNIGG-T----KPQ---------NISYLN--SVRG-----------GKVWLLPCG-A----PHWQ--R---IQ-KPPIKH----KSIF----HD-RSEFT--A-----LA--RNTLWQMQQYL------L--SV------E-----P-------------LTN---------T----
MV---IR--AS----RL------AYSDEIIDILFNYVAEIQ--N-----L-D-G----IS---G----WS------S-----S---E---D-----CE--------LKR---SQQLWL-----D-------------PGR--AKD-------------DPVFHL--ERE--KGD-WK-KEIAQEFGYWLN-----RR-----L---R-------H-------D-K-L------------VF---GEVER------QEWAS----------L--FK-----------------Q-----RLREFEQGS----------------LETG-------S-----------------
---------
WP_145501216 ------------------------------------------------------------MSD-------------EGLA--AFIV--S-YIK-SR-----E----QPK-----------------------------------------------------------------------------------L---------E-------A-FDK--EAEK------R--L--------AG-----LT-----Q--AEDIALAQQ----------------------------EI-------------------------------------A-QQR-QE------------------------LI---
-----AR------YE---VRNW-----LT---DA-----ANRA-------G----QIS-L-----------------VTHALKFTHSD---A-----------------K---G-S------SV----F----N----L-------PA-------AEN--D--AI--Y------LSS--A--TL------G-------Q--P--A-VD-A-VG----N----AA----A--L-----DVAKLL-Q-----T----E--H--N---------GDSL------------LACLN-RAD-N---SPLAAL---A---E----NP----QQLALWVEGFKQA--L---VD--KQ----
LTSHKLAKQFYFPV-------GP------------------DQ---YHLLSPLFSSSLTQAMHQR-IA----E--A---R--F-S-----------------DQ-SKEAK-V-----AH-------------K--TGKWH--SE-A-RVI-----YLK-TAVQNIGG-T----KPQ---------NISYLN--SVRG-----------GKVWLLPCG-A----PPWK--N---IQ-KPPIKY----RSIF----HD-RSEFT--V-----LA--RNNLWQMQQYL------L--GV------K-----P-------------LSN---------T--
--MD---IR--AV----RL------ACSDEIIDILFNYVAEIQ--N-----L-D-G----TN---G----WS------D-----G---K---D-----CK--------LKR---SQQLWL-----D-------------PNR--AMD-------------DPVFKL--ERE--KGD-WK-QEVSQDFGYWLN-----RW-----L---Q-------H-------D-E-L------------VF---SYVEQ------REWSS----------L--FK-----------------Q-----RLREFEQGS----------------LETW-------S-----------------
--------------
WP_032466506 ------------------------------------------------------------MSD-------------EGLT--AFIV--S-YIK-SR-----E----QPK-----------------------------------------------------------------------------------L---------E-------A-FDK--EAEK------R--L--------AG-----LT-----Q--AEDIALAQQ----------------------------EI-------------------------------------A-QQR-QE------------------------LI---
-----AR------YE---VRNW-----LT---DA-----ANRA-------G----QIS-L-----------------VTHALKFTHSD---A-----------------K---G-S------SV----F----N----L-------PA-------AEN--D--AI--Y------LSS--A--TL------G-------Q--P--A-VD-A-VG----N----AA----A--L-----DVAKLL-Q-----T----E--H--N---------GDSL------------LACLN-RAD-N---SPLAAL---A---E----NP----QQLALWVEGFKQA--L---VD--KQ----
LTSHKLAKQFYFPV-------GP------------------DQ---YHLLSPLFSSSLAQAMHQR-II----E--A---R--F-S-----------------DQ-SKEAK-V-----AH-------------K--AGKWH--SE-A-RVI-----YLK-TAVQNIGG-T----KPQ---------NISYLN--SVRG-----------GKVWLLPCG-A----PPWK--N---IQ-KPPIKY----RSIF----HD-RSEFT--V-----LA--RNNLWQMQQYL------L--GV------K-----P-------------LSN---------T---
-MD---IR--AV----RL------ACSDEIIDILFNYVAEIQ--N-----L-D-G----TN---G----WS------D-----A---E---D-----CK--------LKR---SEQLWL-----D-------------PNR--AMD-------------DPVFKL--ERE--KED-WK-QEVSQDFGYWLN-----RW-----L---Q-------H-------D-E-L------------VF---GYVEQ------REWSS----------L--FK-----------------Q-----RLREFEQGS----------------LETW-------S-----------------
--------------
WP_038823635 ------------------------------------------------------------MSD-------------EGLT--AFIV--S-YIK-SR-----E----QPK-----------------------------------------------------------------------------------L---------E-------A-FDK--EAEK------R--L--------AG-----LT-----Q--AEDIALAQQ----------------------------EI-------------------------------------A-QQR-QE------------------------LI---
-----AR------YE---VRNW-----LT---DA-----ANRA-------G----QIS-L-----------------VTHALKFTHSD---A-----------------K---G-S------SV----F----N----L-------PA-------AEN--D--AI--Y------LSS--A--TL------G-------Q--P--A-VD-A-VG----N----AA----A--L-----DVAKLL-Q-----T----E--H--N---------GDSL------------LACLN-RAD-N---SPLAAL---A---E----NP----QQLALWVEGFKQA--L---VD--KQ----
LTSHKLAKQFYFPV-------GL------------------DQ---YHLLSPLFSSSLAQAMHQR-IA----E--A---R--F-S-----------------DQ-SKEAK-V-----AH-------------K--AGKWH--SE-A-RVI-----YLK-TAVQNIGG-T----KPQ---------NISYLN--SVRG-----------GKVWLLPCG-A----PPWK--N---IQ-KPPIKY----RSIF----HD-RSEFT--V-----LA--RNNLWQMQQYL------L--GV------K-----P-------------LSN---------T--

--MD---IR--AV----RL------ACSDEIIDILFNYVAEIQ--N-----L-D-G----TN---G----WS------D-----A---E---D-----CK--------LKR---SEQLWL-----D--------------PNR--AMD--------------DPVFKL--ERE--KGD-WK-QEVSQDFGYWLN-----RW----L---Q-------H-------D-E-L------------VF---GYVEQ------REWSS----------L--FK-----------------Q-----RLREFEQGS----------------LETW------S-----------------------
---------------
WP_106441629 ------------------------------------------------------------MSD------------EGLT--AFIV--S-YIK-SR-----E----QPK----------------------------------------------------------------------------------------------L---------E-------A-FDK--EAEK------R--L--------AG-----LT-----Q--AEDIALAQQ----------------------------EI-----------------------------------A-QQR-QE------------------------LV---
------AR------YE---VRNW-----LT---DA-----ANRA-------G----QIS-L----------------VTHALKFTHSD---A-----------------K---G-S------SV----F----N----L-------PA-------AEN--D--AI--Y------LSS--A--TL------G-------Q--P--A-VD-A-VG----N----AA----A--L-----DVAKLL-Q-----T-----V--H--N---------GDSL------------LACLN-RAD-N---SPLAAL---A---E----NP----QQLALWVEGFKQA--L---VD--KQ----
LTSHKLAKQFYLPV-------GP--------------------DQ---YHLLSPLFSSSLAQAMHQR-IA----E--A---R--F-S-----------------DQ-SKEAK-V-----AH-------------K--AGKWH--SE-A-RVI-----YLK-TAVQNIGG-T----KPQ---------NISYLN--SVRG-----------GKVWLLPCG-A----PPWK--N---IQ-KPPIKY----RSIF----HD-RSEFT--V-----LA--RNNLWQMQQYL------L--GV------K-----R-------------LSN---------T--
--MD---IR--AV----RL------ACSDEIIDILFNYVAEIQ--N-----L-D-G----TN---G----WS------D-----G---E---D-----CE--------LKR---SEQLWL-----D--------------PNR--AMD--------------DPVFKL--ERE--KGD-WK-QEVSQDFGYWLN-----RW----L---Q-------H-------D-E-L------------VF---GYVEQ------REWSS----------L--FK-----------------Q-----RLREFEQGS----------------LETW------S-----------------------
---------------
WP_050093397 ------------------------------------------------------------MSD------------EGLT--AFIV--S-YIK-SR-----E----QPK----------------------------------------------------------------------------------------------L---------E-------A-FDK--EAEK------R--L--------AG-----LT-----Q--AEDIALAQQ----------------------------EI-----------------------------------A-QQR-QE------------------------LV---
------AR------YE---VRNW-----LT---DA-----ANRA-------G----QIS-L----------------VTHALKFTHSD---A-----------------K---G-S------SV----F----N----L-------PA-------AEN--D--AI--Y------LSS--A--TL------G-------Q--P--A-VD-A-VG----N----AA----A--L-----DVAKLL-Q-----T-----E--H--N---------GDSL------------LACLN-RAD-N---SPLAAL---A---E----NP----QQLALWVEGFKQA--L---VD--KQ----
LTSHKLAKQFYFPV-------GP--------------------DQ---YHLLSPLFSSSLAQAMHQR-IA----E--A---R--F-S-----------------EQ-SKEAK-V-----AH-------------K--AGKWH--SE-A-RVI-----YLK-TAVQNIGG-T----KPQ---------NISYLN--SVRG-----------GKVWLLPCG-A----PLWK--N---IQ-KPPIKY----RSIF----HD-RSEFT--V-----LA--RNNLWQMQQYL------L--GV------K-----P-------------LSN---------T--
--MD---IR--AV----RL------ACSDEIIDILFNYVAEIQ--N-----L-D-G----TN---G----WS------D-----G---E---D-----CE--------LKR---SEQLWL-----D--------------PNR--AMD--------------DPVFKL--ERE--KGD-WK-QEVSQDFGYWLN-----RW----L---Q-------H-------D-E-L------------VF---GYVEQ------REWSS----------L--FK-----------------Q-----RLREFEQGS----------------LETW------S-----------------------
---------------
WP_071991219 ---------------------------------------------------------------------------------------------------------------------------------------------------------------------------------------------------------------------------------------------------------------------------------------------------------------MRNW-----LT---DA-----
ANRA-------G----QIS-L------------------VTHALKFTHSD---A-----------------K---G-S------SV----F----N----L--------P-AAE---NDA-----IY---------LSS--A--TL------G-------Q--P--A-VD-A-VG----N----AA----A--L-----DVAKLL-Q-----T-----E--H--N---------GDSL------------LACLN-RAD-N---SPLAAL---A---E----NP----QQLALWVEGFKQA--L---VD--KQ----LTSHKLAKQFYLPV-------GP--------------------
DQ---YHLLSPLFSSSLAQAMHQR-IA----E--A---R--F-S-----------------EQ-SKEAK-V-----AH-------------K--AGKWH--SE-A-RVI-----YLK-TAVQNIGG-T----KPQ---------NISYLN--SVRG-----------GKVWLLPCG-A----PLWK--N---IQ-KPPIKY----RSIF----HD-RSEFT--V-----LA--RNNLWQMQQYL------L--GV------K-----P-------------LSN---------T----MD---IR--AV----RL------
ACSDEIIDILFNYVAEIQ--N-----L-D-G----TN---G----WS------D-----G---E---D-----CE--------LKR---SEQLWL-----D--------------PNR--AMD--------------DPVFKL--ERE--KGD-WK-QEVSQDFGYWLN-----RW----L---Q-------H-------D-E-L------------VF---GYVEQ------REWSS----------L--FK-----------------Q-----RLREFEQGS----------------LETW------S-------------------------------------------
WP_057533090 ------------------------------------------------------------MEE------------KGLT--AFIV--S-YIK-SR-----E----QPK----------------------------------------------------------------------------------------------L---------E-------A-FDK--EAEK------R--L--------AG-----LT-----Q--AEDIALAQQ----------------------------EI-----------------------------------A-QQR-RE------------------------LI----
-----AR------YE---VRNW-----LT---DA-----ANRA-------G----QIS-L----------------VTHALKFTHSD---A-----------------K---G-S------SV----F----N----L-------PA-------AEN--D--AI--Y------LSS--A--TL------G-------Q--P--A-VD-A-VG----N----AA----A--L-----DVAKLL-Q-----T-----E--H--N---------GDSL------------LACLN-RAD-N---SPLAAL---A---E----NP----QQLALWVEGFKQA--L---VD--KQ----
LTSHKLAKQFYFPV-------GP--------------------DQ---YHLLSPLFSSSLAQAMHQR-IA----E--A---R--F-S-----------------DQ-SKEAK-V-----AH-------------K--AGKWH--SE-A-RVI-----YLK-TAVQNIGG-T----KPQ---------NISYLN--SVRG-----------GKVWLLPCG-A----PPWK--N---IQ-KPPIKY----RSIF----HD-RSEFT--V-----LA--RNNLWQMQQYL------L--GV------K-----P-------------LSN---------T--
--MD---IR--AV----RL------ACSDEIIDILFNYVAEIQ--N-----L-D-G----TN---G----WS------D-----G---E---D-----CE--------LKR---SEQLWL-----D--------------PNR--AMD--------------DPVFKL--ERE--KGD-WK-QEVSQDFGYWLN-----RW----L---Q-------H-------D-E-L------------VF---GYVEQ------REWSS----------L--FK-----------------Q-----RLREFEQGS----------------LETW------S-----------------------
---------------
WP_012303963 ------------------------------------------------------------MEE------------KGLT--AFIV--S-YIK-SR-----E----QPK----------------------------------------------------------------------------------------------L---------E-------A-FDK--EAEK------R--L--------AG-----LT-----Q--AEDIALAQQ----------------------------EI-----------------------------------A-QQR-QE------------------------LV---
-----AR------YE---VRNW-----LT---DA-----ANRA-------G----QIS-L----------------VTHALKFTHSD---A-----------------K---G-S------SV----F----N----L-------PA-------AEN--D--AI--Y------LSS--A--TL------G-------Q--P--A-VD-A-VG----N----AA----A--L-----DVAKLL-Q-----T-----E--H--N---------GDSL------------LACLN-RAD-N---SPLAAL---A---E----NP----QQLALWVEGFKQA--L---VD--KQ----
LTSHKLAKQFYFPV-------GP--------------------DQ---YHLLSPLFSSSLAQAMHQR-IA----E--A---R--F-S-----------------DQ-SKEAK-V-----AH-------------K--AGKWH--SE-A-RVI-----YLK-TAVQNIGG-T----KPQ---------NISYLN--SVRG-----------GKVWLLPCG-A----PPWK--N---IQ-KPPIKY----RSIF----HD-RSEFT--V-----LA--RNNLWQMQQYL------L--GV------K-----P-------------LSN---------T--
--MD---IR--AV----RL------ACSDEIIDILFNYVAEIQ--N-----L-D-G----TN---G----WS------D-----A---E---D-----CK--------LKR---SEQLWL-----D--------------PNR--AMD--------------DPVFKL--ERE--KED-WK-QEVSQDFGYWLN-----RW----L---Q-------H-------D-E-L------------VF---GYVEQ------REWSS----------L--FK-----------------Q-----RLREFEQGS----------------LETW------S-----------------------
---------------
WP_057533851 ------------------------------------------------------------MEE------------KGLT--AFIV--S-YIK-SR-----E----QPK----------------------------------------------------------------------------------------------L---------E-------A-FDK--EAEK------R--L--------AG-----LT-----Q--AEDIALAQQ----------------------------EI-----------------------------------A-QQR-RE------------------------LI----
-----AR------YE---VRNW-----LT---DA-----ANRA-------G----QIS-L----------------VTHALKFTHSD---A-----------------K---G-S------SV----F----N----L-------PA-------AEN--D--AI--Y------LSS--A--TL------G-------Q--P--A-VD-A-VG----N----AA----A--L-----DVAKLL-Q-----T-----E--H--N---------GDSL------------LACLN-RAD-N---SPLAAL---A---E----NP----QQLALWVEGFKQA--L---VD--KQ----
LTSHKLAKQFYFPV-------GP--------------------DQ---YHLLSPLFSSSLAQAMHQR-IA----E--A---R--F-S-----------------DQ-SKEAK-V-----AH-------------K--AGKWH--SE-A-RVI-----YLK-TAVQNIGG-T----KPQ---------NISYLN--SVRG-----------GKVWLLPCG-A----PPWK--N---IQ-KPPIKY----RSIF----HD-RSEFT--V-----LA--RNNLWQMQQYL------L--GV------K-----P-------------LSN---------T--
--MD---IR--AV----RL------ACSDEIIDILFNYVAEIQ--N-----L-D-G----TN---G----WS------D-----A---E---D-----CK--------LKR---SEQLWL-----D--------------PNR--AMD--------------DPVFKL--ERE--KED-WK-QEVIQDFGYWLN-----RW----L---Q-------H-------D-E-L------------VF---GYVEQ------REWSS----------L--FK-----------------Q-----RLREFEQGS----------------LETW------S-----------------------
---------------
WP_012104945 ------------------------------------------------------------MEE------------KGLT--AFIV--S-YIK-SR-----E----QPK----------------------------------------------------------------------------------------------L---------E-------A-FDK--EAEK------R--L--------AG-----LT-----Q--AEDIALAQQ----------------------------EI-----------------------------------A-QQR-RE------------------------LI----
-----AR------YE---VRNW-----LT---DA-----ANRA-------G----QIS-L----------------VTHALKFTHSD---A-----------------K---G-S------SV----F----N----L-------PA-------AEN--D--AI--Y------LSS--A--TL------G-------Q--P--A-VD-A-VG----N----AA----A--L-----DVAKLL-Q-----T-----E--H--N---------GDSL------------LACLN-RAD-N---SPLAAL---A---E----NP----QQLALWVEGFKQA--L---VD--KQ----
LTSHKLAKQFYFPV-------GP--------------------DQ---YHLLSPLFSSSLAQAMHQR-IA----E--A---R--F-S-----------------DQ-SKEAK-V-----AH-------------K--AGKWH--SE-A-RVI-----YLK-TAVQNIGG-T----KPQ---------NISYLN--SVRG-----------GKVWLLPCG-A----PPWK--N---IQ-KPPIKY----RSIF----HD-RSEFT--V-----LA--RNNLWQMQQYL------L--GV------K-----P-------------LSN---------T--
--MD---IR--AV----RL------ACSDEIIDILFNYVAEIQ--N-----L-D-G----TN---G----WS------D-----A---E---D-----CK--------LKR---SEQLWL-----D--------------PNR--AMD--------------DPVFKL--ERE--KED-WK-QEVRQDFGYWLN-----RW----L---Q-------H-------D-E-L------------VF---GYVEQ------REWSS----------L--FK-----------------Q-----RLREFEQGS----------------LETW------S-----------------------
---------------
WP_050136751 ------------------------------------------------------------MEE------------KGLT--AFIV--S-YIK-SR-----E----QPK----------------------------------------------------------------------------------------------L---------E-------A-FDK--EAEK------R--L--------AG-----LT-----Q--AEDIALAQQ----------------------------EI-----------------------------------A-QQR-RE------------------------LI----
-----AR------YE---VRNW-----LT---DA-----ANRA-------G----QIS-L----------------VTHALKFTHSD---A-----------------K---G-S------SV----F----N----L-------PA-------AEN--D--AI--Y------LSS--A--TL------G-------Q--P--A-VD-A-VG----N----AA----A--L-----DVAKLL-Q-----T-----E--H--N---------GDSL------------LACLN-RAD-N---SPLAAL---A---E----NS----QQLALWVEGFKQA--L---VD--KQ----
LTSHKLAKQFYFPV-------GL--------------------DQ---YHLLSPLFSSSLAQAMHQR-IA----E--A---R--F-S-----------------DQ-SKEAK-V-----AH-------------K--AGKWH--SE-A-RVI-----YLK-TAVQNIGG-T----KPQ---------NISYLN--SVRG-----------GKVWLLPCG-A----PPWK--N---IQ-KPPIKY----RSIF----HD-RSEFT--V-----LA--RNNLWQMQQYL------L--GV------K-----P-------------LSN---------T--
--MD---IR--AV----RL------ACSDEIIDILFNYVAEIQ--N-----L-D-G----TN---G----WS------D-----A---E---D-----CK--------LKR---SEQLWL-----D--------------PNR--AMD--------------DPVFKL--ERE--KGD-WK-QEVSQDFGYWLN-----RW----L---Q-------H-------D-E-L------------VF---GYVEQ------REWSS----------L--FK-----------------Q-----RLREFEQGS----------------LETW------S-----------------------
---------------
WP_011192626 ------------------------------------------------------------MEE------------KGLT--AFIV--S-YIK-SR-----E----QPK----------------------------------------------------------------------------------------------L---------E-------A-FDK--EAEK------R--L--------AG-----LT-----Q--AEDIALAQQ----------------------------EI-----------------------------------A-QQR-RE------------------------LI----
-----AR------YE---VRNW-----LT---DA-----ANRA-------G----QIS-L----------------VTHALKFTHSD---A-----------------K---G-S------SV----F----N----L-------PA-------AEN--D--AI--Y------LSS--A--TL------G-------Q--P--A-VD-A-VG----N----AA----A--L-----DVAKLL-Q-----T-----E--H--N---------GDSL------------LACLN-RAD-N---SPLAAL---A---E----NP----QQLALWVEGFKQA--L---VD--KQ----
LTSHKLAKQFYFPV-------GL--------------------DQ---YHLLSPLFSSSLAQAMHQR-IA----E--A---R--F-S-----------------DQ-SKEAK-V-----AH-------------K--AGKWH--SE-A-RVI-----YLK-TAVQNIGG-T----KPQ---------NISYLN--SVRG-----------GKVWLLPCG-A----PPWK--N---IQ-KPPIKY----RSIF----HD-RSEFT--V-----LA--RNNLWQMQQYL------L--GV------K-----P-------------LSN---------T--
--MD---IR--AV----RL------ACSDEIIDILFNYVAEIQ--N-----L-D-G----TN---G----WS------D-----A---E---D-----CK--------LKR---SEQLWL-----D--------------PNR--AMD--------------DPVFKL--ERE--KED-WK-QEVSQDFGYWLN-----RW----L---Q-------H-------D-E-L------------VF---GYVEQ------REWSS----------L--FK-----------------Q-----RLREFEQGS----------------LETW------S-----------------------
---------------
WP_025384237 ------------------------------------------------------------MEE------------NGLA--AFIV--S-YIK-SR-----E----QPK----------------------------------------------------------------------------------------------L---------E-------A-FDK--EAEK------R--L--------AG-----LT-----Q--AEDIALAQQ----------------------------EI-----------------------------------A-QQR-QE------------------------LI----
-----AR------YE---VRNW-----LT---DA-----ANRA-------G----QIK-L------------------ATHAPKYTHSD---S-----------------K---S-S------GI----L----N---IE-----------LQ---SKK--K--DY---------FSS--V--DL------A-------E--Q--A-SD-V-IG----N----AA----V--F-----DLVKLL-Q-----S----E--C--E---------GGSL------------IKCLE-QGD-N---SVLKLF---S---N----DE----ELVEEWTTKFKSI--L---NN--EK----
FTSHKLAKQFYFPV-------GP--------------------DQ---YHLLSPLFSSSLAQAMHQR-IA----E--A---R--F-S-----------------DQ-SKEAK-V-----AH-------------K--TGKWH--SE-A-RVI-----YLK-TAVQNIGG-T----KPQ---------NISYLN--SVRG-----------GKVWLLPCG-A----PPWK--N---IQ-KPPIKY----RSIF----HD-RSEFT--V-----LA--RNNLWQMQQYL------L--GV------K-----P-------------LSN---------T--
--MD---IR--AV----RL------ACSDEIIDILFNYVAEIQ--N-----L-D-G----TN---G----WS------D-----G---K---D-----CK--------LKR---SQQLWL-----D--------------PNR--AMD--------------DPVFKL--ERE--KGD-WK-QEVSQDFGYWLN-----RW----L---Q-------H-------D-E-L------------VF---SYVEQ------REWSS----------L--FK-----------------Q-----RLREFEQGS----------------LETW------S-----------------------
---------------
WP_050104570 ------------------------------------------------------------MEE------------NGLA--AFIV--S-YIK-SR-----E----QPK----------------------------------------------------------------------------------------------L---------E-------A-FDK--EAEK------R--L--------AG-----LT-----Q--AEDIALAQQ----------------------------EI-----------------------------------A-QQR-QE------------------------LR-
-------AR------YE---VRNW-----LT---DA-----ANRA-------G----QIK-L------------------ATHAPKYTHSD---S-----------------K---S-S------GI----L----N---IE-----------LQ---SKK--K--DY---------FSS--V--DL------A-------E--Q--A-SD-V-IG----N----AA----V--F-----DLVKLL-Q-----S----E--C--E---------GGSL------------IKCLE-QGD-N---SVLKLF---S---N----DE----ELVEEWTTKFKSI--L---NN--EK----
FTSHKLAKQFYFPV-------GP--------------------DQ---YHLLSPLFSSSLAQAMHQR-IA----E--A---R--F-S-----------------DQ-SKEAK-V-----AH-------------K--TGKWH--SE-A-RVI-----YLK-TAVQNIGG-T----KPQ---------NISYLN--SVRG-----------GKVWLLPCG-A----PPWK--N---IQ-KPPIKY----RSIF----HD-RSEFT--V-----LA--RNNLWQMQQYL------L--GV------K-----P-------------LSN---------T--
--MD---IR--AV----RL------ACSDEIIDILFNYVAEIQ--N-----L-D-G----TN---G----WS------D-----G---K---D-----CK--------LKR---SQQLWL-----D--------------PNR--AMD--------------DPVFKL--ERE--KGD-WK-QEVSQDFGYWLN-----RW----L---Q-------H-------D-E-L------------VF---SYVEQ------REWSS----------L--FK-----------------Q-----RLREFEQGS----------------LETW------S-----------------------
---------------
WP_053010844 ------------------------------------------------------------MEE------------KGLT--AFIV--S-YIK-SR-----E----QPK----------------------------------------------------------------------------------------------L---------E-------A-FDK--EAEK------R--L--------AG-----LT-----Q--AEDIALAQQ----------------------------EI-----------------------------------A-QQR-QE------------------------LV---
------AR------YE---VRNW-----LT---DA-----ANRA-------G----QIK-L------------------ATHAPKYTHSD---S-----------------K---S-S------GI----L----N---IE-----------FQ---SKK--K--DY---------FSS--V--DL------T-------E--Q--A-SD-V-IG----N----AA----V--F-----DLVKLL-Q-----S----E--C--E---------GGSL------------IKCLE-QGD-N---SVLKLF---S---N----DE----ELVEEWTTKFKSI--L---NN--EK----
FTSHKLAKQFYFPV---------------------------EPDQ---YHLLSPLFSSSLAQAMHQR-II----E--A---R--F-S-----------------DQ-SKEAK-A-----AH-------------K--AGQWH--PT-V-RVL-----YLD-TAVQNIGG-T----KPQ---------NISYLN--SVRG-----------GKVWLLPCG-A----PPWK--N---IQ-KPPIKY----RSIF----HD-RSEFT--V-----LA--RNNLWQMQQYL------L--GV------K-----R-------------LSN---------T--
--MD---IR--AV----RL------ACSDEIIDILFNYVAEIQ--N-----L-D-G----TN---G----WS------D-----G---E---D-----CK--------LKR---SEQLWL-----D--------------PNR--AMD--------------DPVFKL--ERE--KGD-WK-QEVSQDFGYWLN-----RW----L---Q-------H-------D-E-L------------VF---GYVEQ------REWSS----------L--FK-----------------Q-----RLREFEQGS----------------LETW------S-----------------------
---------------
WP_050117236 ------------------------------------------------------------MSD------------EGLA--AFIV--S-YIK-SR-----E----QPK----------------------------------------------------------------------------------------------L---------A-------A-FDK--EAEK------R--L--------AG-----LT-----Q--AEDIALAQQ----------------------------EI-----------------------------------A-QQR-QE------------------------LV---
------AR------YE---VRNW-----LT---DA-----ANRA-------G----QIK-L------------------ATHAPKYTHSD---S-----------------K---S-S------GI----L----N---IE-----------FQ---SKK--K--DY---------FSS--V--DL------T-------E--Q--A-SD-V-IG----N----AA----V--F-----DLVKLL-Q-----S----E--C--E---------GGSL------------IKCLE-QGD-N---SVLKLF---S---N----DE----ELVEEWTTKFKSI--L---NN--EK----
FTSHKLAKQFYFPV---------------------------EPDQ---YHLLSPLFSSSLAQAMHQR-II----E--A---R--F-S-----------------DQ-SKEAK-A-----AH-------------K--AGQWH--PT-V-RVL-----YLD-TAVQNIGG-T----KPQ---------NISYLN--SVRG-----------GKVWLLPCG-A----PPWK--N---IQ-KPPIKY----RSIF----HD-RSEFT--V-----LA--RNNLWQMQQYL------L--GV------K-----R-------------LSN---------T--
--MD---IR--AV----RL------ACSDEIIDILFNYVAEIQ--N-----L-D-G----TN---G----WS------D-----G---E---D-----CE--------LKR---SEQLWL-----D--------------PNR--AMD--------------DPVFKL--ERE--KGD-WK-QEVSQDFGYWLN-----RW----L---Q-------H-------D-E-L------------VF---GYVEQ------REWSS----------L--FK-----------------Q-----RLREFEQGS----------------LETW------S-----------------------
---------------
WP_042819776 ------------------------------------------------------------MSD------------EGLA--AFIV--S-YIK-SR-----E----QPK----------------------------------------------------------------------------------------------L---------A-------A-FDK--EAEK------R--L--------AG-----LT-----Q--AEDIALAQQ----------------------------EI-----------------------------------A-QQR-QE------------------------LV--
-------AR------YE---VRNW-----LT---DA-----ANRA-------G----QIK-L------------------ATHAPKYTHSD---S-----------------K---S-S------GI----L----N---IE-----------FQ---SKK--K--DY---------FSS--V--DL------T-------E--Q--A-SD-V-IG----N----AA----V--F-----DLVKLL-Q-----S----E--C--E---------GGSL------------IKCLE-QGD-N---SVLKLF---S---N----DE----ELVEEWTTKFKSI--L---NN--EK----
FTSHKLAKQFYFPV---------------------------EPDQ---YHLLSPLFSSSLAQAMHQR-II----E--A---R--F-S-----------------DQ-SKEAK-A-----AH-------------K--AGQWH--PT-V-RVL-----YLD-TAVQNIGG-T----KPQ---------NISYLN--SVRG-----------GKVWLLPCG-A----PPWK--N---IQ-KPPIKY----RSIF----HD-RSEFT--V-----LA--RNNLWQMQQYL------L--GV------K-----R-------------LSN---------T--
--MD---IR--AV----RL------ACSDEIIDILFNYVAEIQ--N-----L-D-G----TN---G----WS------D-----G---E---D-----CE--------LKR---SEQLWL-----D--------------PNR--AMD--------------DPVFKL--ERE--KGD-WK-QEVSQDFGYWLN-----RW----L---Q-------H-------D-E-L------------VF---GYVEQ------REWSS----------L--FK-----------------Q-----RLREFEQGS----------------LETW------S-----------------------
---------------
WP_050320394 ------------------------------------------------------------MSD------------EGLA--AFIV--S-YIK-SR-----E----QPK----------------------------------------------------------------------------------------------L---------E-------A-FDK--EAEK------R--L--------AG-----LT-----Q--AEDIALAQQ----------------------------EI-----------------------------------A-QQR-QE------------------------LI----
-----AR------YE---VRNW-----LT---DA-----ANRA-------G----QIK-L------------------ATHAPKYTHSD---S-----------------K---S-S------GI----L----N---IE-----------LQ---SKK--K--DY---------FSS--V--DL------A-------E--Q--A-SD-V-IG----N----AA----V--F-----DLVKLL-Q-----S----E--C--E---------GGSL------------IKCLE-QGD-N---SVLKLF---S---N----DE----ELVEEWTTKFKSI--L---NN--EK----
FTSHKLAKQFYFPV-------GP--------------------DQ---YHLLSPLFSSSLAQAMHQR-II----E--A---R--F-S-----------------DQ-SKEAK-A-----AH-------------K--AGRWH--PT-V-RVL-----YLD-TAVQHIGG-T----KPQ---------NISYLN--SVRG-----------GKVWLLPCG-A----PPWK--N---IQ-KPPIKY----RSIF----HD-RSEFT--V-----LA--HNNLRQMQQYL------L--GV------K-----P-------------LSN---------T--
-MD---IR--AA----RL------ACCDEIIDILFNYVAEIQ--N-----L-D-G----TN---G----WS------D-----G---E---D-----CK--------LKR---SEQLWL-----D--------------PNR--AMD--------------DPVFKL--ERE--KGD-WK-QEVSQDFGYWLN-----RW----L---Q-------H-------D-E-L------------VF---GYVEQ------REWSS----------L--FK-----------------Q-----RLREFEQGS----------------LETS------S-----------------------
---------------
WP_054878224 ------------------------------------------------------------MSD------------EGLA--AFIV--S-YIK-SR-----E----QPK----------------------------------------------------------------------------------------------L---------E-------A-FDK--EAEK------R--L--------AG-----LT-----Q--AEDIALAQQ----------------------------EI-----------------------------------A-QQR-QE------------------------LI----
-----AR------YE---VRNW-----LT---DA-----ANRA-------G----QIK-L------------------ATHAPKYTHSD---S-----------------K---S-S------GI----L----N---IE-----------LQ---SKK--K--DY---------FSS--V--DL------A-------E--Q--A-SD-V-IG----N----AA----V--F-----DLVKLL-Q-----S----E--C--E---------GGSL------------IKCLE-QGD-N---SVLKLF---S---N----DE----ELVEEWTTKFKSI--L---NN--EK----
FTSHKLAKQFYFPV-------GP--------------------DQ---YHLLSPLFSSSLAQAMHQR-II----E--A---R--F-S-----------------DQ-SKEAK-A-----AH-------------K--AGRWH--PT-V-RVL-----YLD-TAVQHIGG-T----KPQ---------NISYLN--SVRG-----------GKVWLLPCG-A----PPWK--N---IQ-KPPIKY----RSIF----HD-RSEFT--V-----LA--HNNLRQMQQYL------L--GV------K-----P-------------LSN---------T--
-MD---IR--AA----RL------ACCDEIIDILFNYVAEIQ--N-----L-D-G----TN---G----WS------D-----A---E---D-----CK--------LKR---SEQLWL-----D--------------PNR--AMD--------------DPVFKL--ERE--KED-WK-QEVSQDFGYWLN-----RW----L---Q-------H-------D-E-L------------VF---GYVEQ------REWSS----------L--FK-----------------Q-----RLREFEQGS----------------LETW------S-----------------------
---------------
WP_049600451 ------------------------------------------------------------MSD------------EGLA--AFIV--S-YIK-SR-----E----QPK----------------------------------------------------------------------------------------------L---------E-------A-FDK--EAEK------R--L--------AG-----LT-----Q--AEDIALAQQ----------------------------EI-----------------------------------A-QQR-QE------------------------LI----
-----AR------YE---VRNW-----LT---DA-----ANRA-------G----QIK-L------------------ATHAPKYTHSD---S-----------------K---S-S------GI----L----N---IE-----------LQ---SKK--K--DY---------FSS--V--DL------A-------E--Q--A-SD-V-IG----N----AA----V--F-----DLVKLL-Q-----S----E--C--E---------GGSL------------IKCLE-QGD-N---SVLKLF---S---N----DE----ELVEEWTTKFKSI--L---NN--EK----
FTSHKLAKQFYFPV-------GP--------------------DQ---YHLLSPLFSSSLAQAMHQR-II----E--A---R--F-S-----------------DQ-SKEAK-A-----AH-------------K--AGRWH--PT-V-RVL-----YLD-TAVQHIGG-T----KPQ---------NISYLN--SVRG-----------GKVWLLPCG-A----PPWK--N---IQ-KPPIKY----RSIF----HD-RSEFT--V-----LA--HNNLRQMQQYL------L--GV------K-----P-------------LSN---------T--
-MD---IR--AA----RL------ACCDEIIDILFNYVAEIQ--N-----L-D-G----TN---G----WS------D-----A---E---D-----CK--------LKR---SEQLWL-----D--------------PNR--AMD--------------DPVFKL--ERE--KED-WK-QEVSQDFGYWLN-----RW----L---Q-------H-------D-E-L------------VF---GYVEQ------REWSS----------L--FK-----------------Q-----RLREFEQGS----------------LETS------S-----------------------
---------------
WP_076631715 ------------------------------------------------------------MSD------------EGLA--AFIV--S-YIK-SR-----E----QPK----------------------------------------------------------------------------------------------L---------E-------A-FDK--EAEK------R--L--------AG-----LT-----Q--AEDIALAQQ----------------------------EI-----------------------------------A-QQR-QE------------------------LI----
-----AR------YE---VRNW-----LT---DA-----ANRA-------G----QIK-L------------------ATHAPKYTHSD---S-----------------K---S-S------GI----L----N---IE-----------LQ---SKK--K--DY---------FSS--V--DL------A-------E--Q--A-SD-V-IG----N----AA----V--F-----DLVKLL-Q-----S----E--C--E---------GGSL------------IKCLE-QGD-N---SVLKLF---S---N----DD----ELVEEWTTKFKSI--L---NN--EK----
FTSHKLAKQFYFPV-------GP--------------------DQ---YHLLSPLFSSSLAQAMHQR-II----E--A---R--F-S-----------------DQ-SKEAK-A-----AH-------------K--AGRWH--PT-V-RVL-----YLD-TAVQHIGG-T----KPQ---------NISYQN--SVRG-----------EKVWLLPCG-A----PSWK--N---IQ-KPPIKY----RSIF----HD-RSEFT--V-----LA--RNNLWQMQLYL------L--GV------K-----P-------------LSN---------T--
--MD---IR--AA----RL------ACCDEIIDILFNYVAEIQ--N-----L-D-G----TN---G----WS------D-----A---E---D-----CK--------LKR---SEQLWL-----D--------------PTR--AMD--------------DPVFKL--ERE--KED-WK-QEVSQDFGYWLN-----RW----L---Q-------H-------D-E-L------------VF---GYVEQ------REWSS----------L--FK-----------------Q-----RLREFEQGS----------------LETS------S-----------------------
---------------
WP_087493427 ------------------------------------------------------------MSD------------EGLA--AFIV--S-YIK-SR-----E----QPK----------------------------------------------------------------------------------------------L---------E-------A-FDK--EAEK------R--L--------AG-----LT-----Q--AEDIALAQQ----------------------------EI-----------------------------------A-QQR-QE------------------------LI----
-----AR------YE---VRNW-----LT---DA-----ANRA-------G----QIK-L------------------ATHAPKYTHSD---S-----------------K---S-S------GI----L----N---IE-----------LQ---SKK--K--DY---------FSS--V--DL------A-------E--Q--A-SD-V-IG----N----AA----V--F-----DLVKLL-Q-----S----E--C--E---------GGSL------------IKCLE-QGD-N---SVLKLF---S---N----DD----ELVEEWTTKFKSI--L---NN--EK----
FTSHKLAKQFYFPV-------GP--------------------DQ---YHLLSPLFSSSLAQAMHQR-II----E--A---R--F-S-----------------DQ-SKEAK-A-----AH-------------K--AGRWH--PT-V-RVL-----YLD-TAVQHIGG-T----KPP---------NISYLN--SVRG-----------GKVWLLPCG-A----PSWK--N---IQ-KPPIKY----RSIF----HD-RSEFT--V-----LA--RNNLWQMQLYL------L--GV------K-----P-------------LSN---------T--
-MD---IR--AA----RL------ACCDEIIDILFNYVAEIQ--N-----L-D-G----TN---G----WS------D-----A---E---D-----CK--------LKR---SEQLWL-----D--------------PTR--AMD--------------DPVFKL--ERE--KED-WK-QEVSQDFGYWLN-----RW----L---Q-------H-------D-E-L------------VF---GYVEQ------REWSS----------L--FK-----------------Q-----RLREFEQGS----------------LETS------S-----------------------
---------------
WP_012229482 ------------------------------------------------------------MSD------------EGLA--AFIV--S-YIK-SR-----E----QPK----------------------------------------------------------------------------------------------L---------E-------A-FDK--EAEK------R--L--------AG-----LT-----Q--AEDIALAQQ----------------------------EI-----------------------------------A-QQR-QE------------------------LI----
-----AR------YE---VRNW-----LT---DA-----ANRA-------G----QIK-L------------------ATHAPKYTHSD---S-----------------K---S-S------GI----L----N---IE-----------LQ---SKK--K--DY---------FSS--V--DL------A-------E--Q--A-SD-V-IG----N----AA----V--F-----DLVKLL-Q-----S----E--C--E---------GGSL------------IKCLE-QGD-N---SVLKLF---S---N----DD----ELVEEWTTKFKSI--L---NN--EK----
FTSHKLAKQFYFPV-------GP--------------------DQ---YHLLSPLFSSSLAQAMHQR-II----E--A---R--F-S-----------------DQ-SKEAK-A-----AH-------------K--AGRWH--PT-V-RVL-----YLD-TAVQHIGG-T----KPQ---------NISYLN--SVRG-----------GKVWLLPCG-A----PSWK--N---IQ-KPPIKY----RSIF----HD-RSEFT--V-----LA--RNNLWQMQLYL------L--GV------K-----P-------------LRN---------T--
--MD---IR--AA----RL------ACCDEIIDILFNYVAEIQ--N-----L-D-G----TN---G----WS------D-----A---E---D-----CK--------LKR---SEQLWL-----D--------------PTR--AMD--------------DPVFKL--ERE--KED-WK-QEVSQDFGYWLN-----RW----L---Q-------H-------D-E-L------------VF---GYVEQ------REWSS----------L--FK-----------------Q-----RLREFEQGS----------------LETS------S-----------------------
---------------
WP_076636197 ------------------------------------------------------------MSD------------EGLA--AFIV--S-YIK-SR-----E----QPK----------------------------------------------------------------------------------------------L---------E-------A-FDK--EAEK------R--L--------AG-----LT-----Q--AEDIALAQQ----------------------------EI-----------------------------------A-QQR-QE------------------------LI----
-----AR------YE---VRNW-----LT---DA-----ANRA-------G----QIK-L------------------ATHAPKYTHSD---S-----------------K---S-S------GI----L----N---IE-----------LQ---SKK--K--DY---------FSS--V--DL------A-------E--Q--A-SD-V-IG----N----AA----V--F-----DLVKLL-Q-----S----E--C--E---------GGSL------------IKCLE-QGD-N---SVLKLF---S---N----DD----ELVEEWTTKFKSI--L---NN--EK----
FTSHKLAKQFYFPV-------GP--------------------DQ---YHLLSPLFSSSLAQAMHQR-II----E--A---R--F-S-----------------DQ-SKEAK-A-----AH-------------K--AGRWH--PT-V-RVL-----YLD-TAVQHIGG-T----KPQ---------NISYLN--SVRG-----------GKVWLLPCG-A----PSWE--N---IQ-KPPIKY----RSIF----HD-RSEFT--V-----LA--RNNLWQMQLYL------L--GV------K-----P-------------LSN---------T--
-MD---IR--AA----RL------ACCDEIIDILFNYVAEIQ--N-----L-D-G----TN---G----WS------D-----A---E---D-----CK--------LKR---SEQLWL-----D--------------PTR--AMD--------------DPVFKL--ERE--KED-WK-QEVSQDFGYWLN-----RW----L---Q-------H-------D-E-L------------VF---GYVEQ------REWSS----------L--FK-----------------Q-----RLREFEQGS----------------LETS------S-----------------------
---------------
WP_002222491 ------------------------------------------------------------MSD------------EGLA--AFIV--S-YIK-SR-----E----QPK----------------------------------------------------------------------------------------------L---------E-------A-FDK--EAEK------R--L--------AG-----LT-----Q--AEDIALAQQ----------------------------EI-----------------------------------A-QQR-QE------------------------LI----
-----AR------YE---VRNW-----LT---DA-----ANRA-------G----QIK-L------------------ATHAPKYTHSD---S-----------------K---S-S------GI----L----N---IE-----------LQ---SKK--K--DY---------FSS--V--DL------A-------E--Q--A-SD-V-IG----N----AA----V--F-----DLVKLL-Q-----S----E--C--E---------GGSL------------IKCLE-QGD-N---SVLKLF---S---N----DD----ELVEEWTTKFKSI--L---NN--EK----
FTSHKLAKQFYFPV-------GP--------------------DQ---YHLLSPLFSSSLAQAMHQR-II----E--A---R--F-S-----------------DQ-SKEAK-A-----AH-------------K--AGRWH--PT-V-RVL-----YLD-TAVQHIGG-T----KPQ---------NISYLN--SVRG-----------GKVWLLPCG-A----PSWK--N---IQ-KPPIKY----RSIF----HD-RSEFT--V-----LA--RNNLWQMQLYL------L--GV------K-----P-------------LSN---------T--
-MD---IR--AA----RL------ACCDEIIDILFNYVAEIQ--N-----L-D-G----TN---G----WS------D-----A---E---D-----CK--------LKR---SEQLWL-----D--------------PTR--AMD--------------DPVFKL--ERE--KED-WK-QEVSQDFGYWLN-----RW----L---Q-------H-------D-E-L------------VF---GYVEQ------REWSS----------L--FK-----------------Q-----RLREFEQGS----------------LETS------S-----------------------
---------------
WP_002216548 --------------------------------------------------------------------------M--RVIV--S-YIK-SR-----E----QPK----------------------------------------------------------------------------------------------L---------H-------Y-LDK--EAEK------R--L--------AG-----LF-----Q--AEDIALAQQ----------------------------EI-----------------------------------A-QQR-QE------------------------LI---------
AR------YE---VRNW-----LT---DA-----ANRA-------G----QIK-L------------------ATHAPKYTHSD---S-----------------K---S-S------GI----L----N---IE-----------LQ---SKK--K--DY---------FSS--V--DL------A-------E--Q--A-SD-V-IG----N----AA----V--F-----DLVKLL-Q-----S----E--C--E---------GGSL------------IKCLE-QGD-N---SVLKLF---S---N----DD----ELVEEWTTKFKSI--L---NN--EK----
FTSHKLAKQFYFPV-------GP--------------------DQ---YHLLSPLFSSSLAQAMHQR-IA----E--A---R--F-S-----------------DQ-SKEAK-A-----AH-------------K--AGRWH--PT-V-RVL-----YLD-TAVQHIGG-T----KPQ---------NISYLN--SVRG-----------GKVWLLPCG-A----PSWK--N---IQ-KPPIKY----RSIF----HD-RSEFT--V-----LA--RNNLWQMQLYL------L--GV------K-----P-------------LSN---------T--
-MD---IR--AA----RL------ACCDEIIDILFNYVAEIQ--N-----L-D-G----TN---G----WS------D-----A---E---D-----CK--------LKR---SEQLWL-----D--------------PTR--AMD--------------DPVFKL--ERE--KED-WK-QEVSQDFGYWLN-----RW----L---Q-------H-------D-E-L------------VF---GYVEQ------REWSS----------L--FK-----------------Q-----RLREFEQGS----------------LETS------S-----------------------
---------------
WP_002421472 ---------------------------------------------------------------------------------------------------------------------------------------------------------------------------------------------------------------------------------------------------------------------------------------------------------------MRNW-----LT---DA-----
ANRA-------G----QIK-L------------------ATHAPKYTHSD---S-----------------K---S-S------GI----L----N---IE-----------LQ---SKK--K--DY---------FSS--V--DL------A-------E--Q--A-SD-V-IG----N----AA----V--F-----DLVKLL-Q-----S----E--C--E---------GGSL------------IKCLE-QGD-N---SVLKLF---S---N----DD----ELVEEWTTKFKSI--L---NN--EK----FTSHKLAKQFYFPV-------GP--------------------DQ---
YHLLSPLFSSSLAQAMHQR-II----E--A---R--F-S-----------------DQ-SKEAK-A-----AH-------------K--AGRWH--PT-V-RVL-----YLD-TAVQHIGG-T----KPQ---------NISYLN--SVRG-----------GKVWLLPCG-A----PSWK--N---IQ-KPPIKY----RSIF----HD-RSEFT--V-----LA--RNNLWQMQLYL------L--GV------K-----P-------------LSN---------T----MD---IR--AA----RL------ACCDEIIDILFNYVAEIQ--
N-----L-D-G----TN---G----WS------D-----A---E---D-----CK--------LKR---SEQLWL-----D--------------PTR--AMD--------------DPVFKL--ERE--KED-WK-QEVSQDFGYWLN-----RW----L---Q-------H-------D-E-L------------VF---GYVEQ------REWSS----------L--FK-----------------Q-----RLREFEQGS----------------LETS------S-----
WP_016599751 ------------------------------------------------------------------------------------------------------------------------------------------------------------------------------------------------------------------------------------------------------------------------------------------------------------------T---DA-----ANRA---
----G----QIK-L------------------ATHAPKYTHSD---S-----------------K---S-S------GI----L----N---IE-----------LQ---SKK--K--DY---------FSS--V--DL------A-------E--Q--A-SD-V-IG----N----AA----V--F-----DLVKLL-Q-----S----E--C--E---------GGSL------------IKCLE-QGD-N---SVLKLF---S---N----DD----ELVEEWTTKFKSI--L---NN--EK----FTSHKLAKQFYFPV-------GP--------------------DQ---
YHLLSPLFSSSLAQAMHQR-II----E--A---R--F-S-----------------DQ-SKEAK-A-----AH-------------K--AGRWH--PT-V-RVL-----YLD-TAVQHIGG-T----KPQ---------NISYLN--SVRG-----------GKVWLLPCG-A----PSWK--N---IQ-KPPIKY----RSIF----HD-RSEFT--V-----LA--RNNLWQMQLYL------L--GV------K-----P-------------LSN---------T----MD---IR--AA----RL------ACCDEIIDILFNYVAEIQ--
N-----L-D-G----TN---G----WS------D-----A---E---D-----CK--------LKR---SEQLWL-----D--------------PTR--AMD--------------DPVFKL--ERE--KED-WK-QEVSQDFGYWLN-----RW----L---Q-------H-------D-E-L------------VF---GYVEQ------REWSS----------L--FK-----------------Q-----RLREFEQGS----------------LETS------S-----
WP_016599914 ---------------------------------------------------------------------------------------------------------------------------------------------------------------------------------------------------------------------------------------------------------------------------------------------------------------------------------------
-------------------------Q--G--------------------------D------------------------------------N---------SVLKLF-----------------------------SND----------------D-------ELVEEWTTKFKSI--L---NN--EK----FTSHKLAKQFYFPV-------GP--------------------DQ---YHLLSPLFSSSLAQAMHQR-II----E--A---R--F-S-----------------DQ-
SKEAK-A-----AH-------------K--AGRWH--PT-V-RVL-----YLD-TAVQHIGG-T----KPQ---------NISYLN--SVRG-----------GKVWLLPCG-A----PSWK--N---IQ-KPPIKY----RSIF----HD-RSEFT--V-----LA--RNNLWQMQLYL------L--GV------K-----P-------------LSN---------T----MD---IR--AA----RL------ACCDEIIDILFNYVAEIQ--N-----L-D-G----TN---G----WS------D-----A---E---D-----CK--------
LKR---SEQLWL-----D--------------PTR--AMD--------------DPVFKL--ERE--KED-WK-QEVSQDFGYWLN-----RW----L---Q-------H-------D-E-L------------VF---GYVEQ------REWSS----------L--FK-----------------Q-----RLREFEQGS----------------LETS------S-----
WP_126530780 ------------------------------------------------------------MSN------------------QG------IV--A-FIE-SR-----K----QPK----------------------------------------------------------------------------------------------L---------E-------A-FDK--DTEK------K--L--------AG-----LS-----D--AAEIAAVRQ----------------------------AA-----------------------------------A-EQR-RE------------------------LE----
----LR------YE---VRNW-----LT---DA-----ANRA-------G----QIG-L------------------VTHALKFTHSD---A-----------------K---G-S------SI----F----A----E-------QQ-------EDE--G--AR--Y-------LAT--A--TL------R-------Q--P--A-ID-A-VG----N----AA----A--L-----DVAKLL-Q-----M----E--Y--Q---------GDTL------------IAQLN-RGD-H---SALAVL---A---E----SE----QQLEQWVSGFKQV--L---TN--KQ----
PASHALAKQLYFPI-------GD--------------------NQ---YHLLSPLFSSSLAQALHQR-LA----Q--A---R--F-S-----------------EQ-AKEIS-A-----AV-------------R--AGKWH--AE-A-RVM-----YPN-LAVQSMGG-T----KPQ---------NISYLN--STRG-----------GRNWLLNCA-A----PQWV--T---QS-KPPLKH----TSIF----YR-SSEFS--A-----IA--RDAVRQLRAYL------L--SV------Q-----P-------------RDS---------T-
---ME---IR--QR----RA------AGVDEVIGMLFCYAAGIQ--N-----L-E-V----QA---G----WS------A-----E---A---E-----CR--------LKR---SQQLWL-----D--------------PLR--GRD--------------DPDFRA--ERE--AGD-WQ-REIALDFGVWLN-----RQ----L---E-------H-------E-Q-M------------LF---SEVER------REWST-A-------PL--FK-----------------Q-----RLHEFEAEW----------------AEAA-------L-----------
------------------
WP_015670635 ------------------------------------------------------------MSN------------------QGIV--A-FIE-SR-----K----QPK----------------------------------------------------------------------------------------------L---------E-------A-FDK--DTEK------K--L--------AG-----LS-----D--AAEIAAVRQ----------------------------AA-----------------------------------V-EQR-RE------------------------LE----
----QR------YE---VRNW-----LT---DA-----ASRA-------G----QIG-L------------------VTHALKFTHSD---A-----------------K---G-S------SI----F----A----A-----------QE---DND--A--RY---------LGT--A--TL------R-------Q--P--A-ID-A-VG----N----AA----A--L-----DVAKLL-Q-----M----E--Y--Q---------GDTL------------IAQLN-RGD-H---SALAVL---A---E----SE----QQLELWVSGFKQV--L---TN--KQ----
PASHALAKQLYFPI-------GD--------------------NQ---YHLLSPLFSSSLAQALHQR-LA----Q--A---R--F-S-----------------EQ-AKEIS-A-----AV-------------R--AGKWH--AE-A-RVM-----YPN-LAVQSMGG-T----KPQ---------NISYLN--STRG-----------GRNWLLNCA-A----PQWV--T---QS-KPPLKH----TSIF----YR-SSEFS--A-----MA--RDAVRQLRAYL------L--SV------Q-----P-------------REK---------
T----RE---IR--QR----RA------AGVDEVIDMLFSYAAGIQ--N-----L-E-A----QA---G----WS------A-----G---A---E-----CR--------LKR---SQQLWL-----D--------------PLR--GRD--------------DPDFRA--ERE--AGD-WQ-REIALDFGVWLN-----RQ----L---E-------H-------E-Q-M------------LF---SEVER------REWST-A-------PL--FK-----------------Q-----RLREFEAEW----------------AEAA-------L-----------
---------------------
WP_076942992 ------------------------------------------------------------MED------------NEFG--RIIV--R-YIE-SR-----R----QPK----------------------------------------------------------------------------------------------L---------E-------A-FDK--EAEK------R--L--------AG-----LT-----D--SEQLAAAQQ----------------------------EI-----------------------------------A-QQR-RE------------------------LE-
------QR------YE---VRNW-----LT---DA-----ASRA-------G----QIS-L----------------VTHALKFTHSD---A-----------------K---G-S------SV----F----L----P--------T-------VVS--D--SA--Y-------LST--A--TL------R-------N--P--A-ID-A-VG----N----AA----A--L-----DVAKLL-Q-----T----E--Y--E---------GESL------------IAAIN-RGD-D---SPLAAL---A---E----SD----QQLQQWISGFKQV--L---VD--KQ----
PSSHKLAKQIYFPV-------GD--------------------ND---YHLLSPLFSSSLAHAMHQR-IA----D--A---R--F-S-----------------EA-AKEIY-G-----AF-------------K--AGVWH--SA-P-RVS-----YPN-TAVQNIGG-T----KPQ---------NISYLN--SVRG-----------GRTWLLSCA-A----PSWE--I---TP-KPPTKH----KSIF----HE-NSDFS--R-----MA--RPIVRELRHYL------L--SV------K-----Q-------------LEN---------T----
KE---IR--DH----RL------AFTDEIIDTLFNYVAEIQ--N-----L-D-V----QV---G----WS------A-----E---E---G-----CQ--------LKR---SQQLWL-----D--------------AGR--AEK--------------DEDFKF--ERE--GGD-WQ-KEIALDFGVWLN-----RR----L---Q-------H-------E-E-L------------KF---GEVER------REWST-A-------KL--FK-----------------Q-----RLREFEQDL----------------QEGL-------V-----------
---------------
WP_024912539 ------------------------------------------------------------MED------------KTFS--QLIA--R-YIE-GR-----K----QPK----------------------------------------------------------------------------------------------L---------E-------A-FDK--EAEK------R--L--------AG-----LT-----D--SEQLAVTQQ----------------------------DI-----------------------------------A-QQR-RE------------------------LE-
-------QR------YE---VRNW-----LT---DA-----ASRA-------G----QIS-L----------------VTHALKFTHSD---A-----------------K---G-S------SV----F----L----S-----------TA---EGD--L--AY---------LAT--A--TL------R-------N--P--A-LD-A-VG----N----AA----A--L-----DCAKLL-Q-----M----E--Y--E---------GESL------------IAVLD-RGD-D---SPLAAL---A---E----SD----RQLQQWVNGFKQV--L---TD--KQ----
PSSHKLAKQIYFPI-------GD--------------------NE---YHLLSPLFSSSLAHAMHQR-IA----D--A---R--F-S-----------------DA-AKEIY-S-----AF-------------K--AGEWH--PA-A-RVS-----YPN-TAVQNIGG-T----KPQ---------NISYLN--SVRG-----------GRTWLLSCA-A----PSWK--I---IS-KPPTRH----KSIF----HD-YGDFS--R-----MA--YPIVRELRHYL------L--SV------K-----D-------------LEN---------T---
KD---IR--DS----RL------AFTDEIIDTLFNYVAMVQ--D-----L-E-V----QA---G----WS------A-----K---E---G-----CQ--------LKR---SQQLWL-----D--------------AGR--AKD--------------DADFQF--ERE--GGD-WQ-KEIALDFGVWLN-----RR----L---Q-------H-------E-E-L------------KF---GEVER------REWST-A-------KL--FK-----------------Q-----RLREFEQDL----------------KEGL-------A-----------
---------------
WP_123337114 ------------------------------------------------------------MLQ------------ETLA--SFIT--D-YIN-ER-----K----TEK----------------------------------------------------------------------------------------------L---------D-------V-FEK--KKKK------Q--L--------SA-----LS-----E--QEGSASVVQ----------------------------DL-----------------------------------N-QQR-AE------------------------IE--
-------KK------YQ---LEVW-----LT---DA-----AILT-------G----QIS-Q----------------ATHALKFTHGK---A-----------------R---G-S------SV----F----Y----E-------EA-----------TS--S--TS--Y-------LST--A--TL------G-------R--R--K-LD-I-VG----N----AA----Y--F-----YIAKVL-L-----R----E--V--Q---------GDSL------------IAGLN-RGD-Y---SVLESL---T---D----NK----ELSDAWIEGFKQV--L---ID--KQ----
PVSHNLAKQIYFPI-------AP--------------------QQ---YHLISPLFASSLTHVLHQR-IA----D--A---R--F-S-----------------EE-AKNAR-A-----AM-------------K--AKTWH--RE-P-VVV-----FPG-TAVANFGG-T----KPQ---------NISYLN--SLRG-----------GKVWLLPCL-P----PAWQ--T---VS-KPPLKN----QSLF----SY-S-EFS--R-----QA--WPVILRMKRYL------H--QV------K-----P-------------LDS---------T---
ME---IR--QQ----RL------AFTDEIIDTLFNYAAGIQ--N-----L-T-Q----VS---G----WS------A-----D---R---E-----CR--------LKQ---SQQLWL-----D--------------PLR--SKT--------------DLQFRL--ERE--KGD-WK-QEIARDFGWLN-----HQ----L---R-------D-------E-T-L------------KL---GAVEH------QHFSA-S-------PL--LN-----------------K-----RLRELEDDL----------------AEDL-------P-----
---------
WP_137269073 ------------------------------------------------------------MSR------------ETLT--DFIT--T-YIS-GR-----K----TVK----------------------------------------------------------------------------------------------L---------E-------A-HDK--DAEK------K--L--------AG-----VT-----A--QDAAQT-QA----------------------------EL-----------------------------------N-QKR-AE------------------------LE-
-------SQ------YQ---VTEW-----LT---NA-----ASRA-------G----QIS-L----------------VTHALKFTHSD---A-----------------R---G-S------SV----F----T----S-------NS-------GQ--Q--NA--Y-------LST--S--TL------N-------K--P--A-LD-A-VG----N----AA----A--L-----DIAKLL-Q-----T----E--V--E---------GDSL------------IASLQ-RGD-T---SALDAF---T---D----DS----QLLESWVNGFKQV--L---VD--KQ----
PASHKLAKQLYFPI-------GD--------------------SQ---YHLLSPLFSSSLAHAVNQR-IT----A--A---R--F-S-----------------DE-SKAAR-A-----AM-------------S--ARHWS--AD-P-VVS-----YPA-TAVINMGG-T----KPQ---------NISYLN--SVRG-----------GKVWLLSCA-S----PGWS--N---LS-KPPLKH----KSIF----HF-NSEIG--R-----QA--RPVIARLKRFL------H--CV------Q-----P-------------LAN---------T---
MN---IR--QQ----RL------ALTDEIIDILFNYVLDVQ--N-----Q-T-E----HQ---G----WS------A-----R---P---E-----CQ--------LKR---SQQLWL-----D--------------PYR--SLT--------------DDDFRF--ERE--GGE-WK-NEVARDFGHWLI-----RQ----L---Q-------D-------E-A-L------------HM---GAVER------REFST-A-------PL--FK-----------------Q-----RLRELEQDL----------------QEDL-------S-----
-------------
WP_023655989 ------------------------------------------------------------MLR------------ETLA--SFIT--S-YIA-AR-----K----TAK----------------------------------------------------------------------------------------------L---------E-------A-FDK--ESAK------K--L--------AA-----LA-----S--EDAISVLRQ----------------------------QL-----------------------------------Q-QQR-AE------------------------LE---
-----QK------YQ---PQAW-----LS---DA-----ASRA-------G----QIK-L----------------VTHAVKFTHSD---A-----------------R---G-S------SI----F----T----S-------GS-------GQH--E--AY---------LST--V--TL------Q-------K--S--A-LD-I-AC----N----AA----A--L-----DVAKLL-Q-----S----E--V--E---------GDSL------------IASLQ-RGD-Y---SALESL---T---D----DP----ELCASWVSGFKQV--L---VD--RQ----
PASHKLAKQIYFPI---------------------------ADGQ---YHLLSPLFSSSLAHALNQR-II----S--T---K--F-S-----------------EE-TKAAR-A-----AL-------------K--AKSWH--GE-A-LVT-----FPG-TAITQFGG-T----KPQ---------NISYLN--SVRG-----------GKVWLLPCA-P----PVWQ--T---QN-RPPSQH----KSIF----NS-SNDFS--R-----QA--WPVIQRLGRFL------R--RV------K------------------------------P----LD-
--IR---QQ----RL------AMTDEIIDILFNYVAGVQ--N-----Q-T-E----LR---G----WS------A-----H---P---D-----CV--------LKR---SQQLWL-----D--------------PWR--SQD--------------DTEFQF--ERE--GGD-WK-NEVARDFGHWLS-----RH----L---H-------S-------D-R-L------------NM---GETER------RHFST-A-------LL--FK-----------------Q-----RLRELEKDL----------------AEDL-------P-----
---------
WP_012440590 ------------------------------------------------------------MLR------------ETLA--SFIT--S-YIA-AR-----K----TAK----------------------------------------------------------------------------------------------L---------E-------A-FDK--ESAK------K--L--------AV-----LA-----S--EDEISVLRQ----------------------------QL-----------------------------------Q-QQR-AE------------------------LE---
-----QK------YQ---PQAW-----LS---DA-----ASRA-------G----QIK-L----------------VTHAAKFTHSD---V-----------------R---G-S------SI----F----S----S-------GS-------GQH--E--TY---------LST--A--TL------Q-------K--P--A-LD-A-VG----N----AA----A--L-----DIARLL-Q-----S----E--V--E---------GDSL------------IASLQ-RGD-Y---SALESL---T---D----NP----ELCASWISGFKQV--L---VD--RQ----
PASHKLAKQIYFPI---------------------------ADGQ---YHLLSPLFSSSLAHALNQR-IT----E--A---K--F-S-----------------EQ-AKTAR-A-----AL-------------K--AKSWH--DA-P-VVA-----YPD-TAITQFGG-T----KPQ---------NISYLN--SVRG-----------GKVWLLPCA-P----PVWQ--T---LS-KPPAKH----KSIF----NS-SNDFS--R-----QS--WPVIQRMSRFL------R--RV------E-----R-------------LDS---------T---
LD---IR--QQ----RL------AMTDEIIDILFNYVAGIQ--N-----Q-T-E----SI---G----WS------A-----H---P---D-----CV--------LKR---SQQLWL-----D--------------PWR--G-----------------DKEFQF--ERE--GGD-WK-SEVARDFGHWLS-----RH----L---H-------S-------D-K-L------------NM---GETER------RHFST-A-------PL--FK-----------------Q-----RLRELEKDL----------------AEDL-------P-----
---------
WP_012667183 ------------------------------------------------------------MLR------------ETLA--SFIT--S-YIA-AR-----K----TAK----------------------------------------------------------------------------------------------L---------E-------A-FDK--ESAK------K--L--------AA-----LA-----S--EDQISVLRQ----------------------------QL-----------------------------------Q-QQR-AE------------------------LE---
------QK------YQ---PQAW-----LS---DA-----ASRA-------G----QIN-L----------------VTHAAKFTHSD---A-----------------R---G-S------SI----F----A----S-----------SSV---QHE--M--Y----------LST--A--AL------Q-------K--P--A-LD-V-VG----N----AA----I--L-----DVAKLL-Q-----T----E--V--E---------GNSL------------IASLQ-RGD-Y---SALESL---T---D----NP----ELCASWVNGFKQV--L---VD--RQ----
PASHKLAKQIYFPI---------------------------ADGQ---YHLLSPLFSSALAHALNQR-II----S--A---K--F-S-----------------EE-AKTAR-A-----AL-------------K--AKSWH--DG-P-VVA-----YPG-TAITQFGG-T----KPQ---------NISYLN--SVRG-----------GKVWLLPCA-P----PVWQ--T---QS-QPPSQH----KSIF----NF-SNDFS--R-----QS--WPVIQRLKHFL------R--RV------K-----T-------------LDS---------T---
LD---IR--QQ----RL------AMTDEIIDILFNYVAGLQ--N-----Q-T-D----LR---G----WS------A-----H---P---D-----CV--------LKR---SQQLWL-----D--------------PWR--SQD--------------DAEFQF--ERE--GGD-WK-NEVARDFGHWLS-----RH----L---H-------S-------D-K-L------------NM---GETER------RHFST-A-------PL--FK-----------------Q-----RLRELEKDL----------------AEDL-------S-----
WP_101828210 ------------------------------------------------------------------------MDIT--S-WIN-DR-----K----QSK----------------------------------------------------------------------------------------------L---------A-------A-FDK--DAKK------R--R--------EA-----AS-----A---ADLSALDQ----------------------------AL-----------------------------------A-EAR-RE------------------------LE--------
-LK------YE---PRTW-----LT---NA-----AQRA-------G----QIS-L----------------VTHAAKYTHGD---A-----------------Q---S-S------SL----Y----S----C-------IR---------Q--Q--EG--Y------LST--A--SL------A-------A--P--S-LD-A-VG----N----AA----A--L-----DVAKLL-Q-----T----R--F--G---------KDSL------------LDCLR-RKE-Y---SLLAEL---A---E----DS----EQLAAWVTGFSRV--L---DT--GQ----
PTSHTLSKQIYFPV---------------------------GDG---YHLLSPLFSSSLAQALHQK-IV----A--Q---R--F-S-----------------EE-SKAAR-Q-----AR-------------R--NKQWH--PL-P-VVS-----FPN-TAVMNFGG-T----KPQ---------NISALN--SSRG-----------GRMWLLSSQ-P----PQWQ--V---QE-RKPLAI----TSLF----AP-KGPFE--Y-----LT--ANTLTSFTALL------T--HS------G-----N-------------YN---------N---
-AA---IK--RR----RE------HYVDELIDLLFNVAAGIQ--------R-P-E----WC---N----WT------T-----E---S---S-----CL--------LRP---HQALWL-----D--------------PWR--TQD--------------DPAFAL--DRE--RGD-WQ-AAVADDFAQWLN-----TQ----L---N-------A-------A-K-C------------AV---GMVER------KAWRT-Q-----------CQ-----------------Q-----RLREMEHVI----------------KEAQ-------S-----
---------------
WP_101855123 ------------------------------------------------------------------------MDIT--S-WIN-DR-----K----QSK----------------------------------------------------------------------------------------------L---------A-------A-FDK--DAKK------R--R--------EA-----AS-----A---ADLSALDQ----------------------------AL-----------------------------------A-EAR-RE------------------------LE--------
-LK------YE---PRTW-----LT---NA-----AQRA-------G----QIS-L----------------VTHAAKYTHGD---A-----------------Q---S-S------SL----Y----S----C-------IR---------Q--Q--EG--Y------LST--A--SL------A-------A--P--S-LD-A-VG----N----AA----A--L-----DVAKLL-Q-----T----R--F--G---------KDSL------------LDCLR-RKE-Y---SLLAEL---A---E----DS----EQLAAWVTGFSRV--L---ET--GQ----
PTSHTLAKQIYFPV---------------------------GDG---YHLLSPLFSSSLAQALHQK-IV----A--Q---R--F-S-----------------EE-SKAAR-Q-----AR-------------R--NKQWH--PL-P-VVS-----FPN-TAVMNFGG-T----KPQ---------NISALN--SSRS-----------GRIWLLSSQ-P----PQWQ--A---QE-RKPLAI----TSLF----AP-KGPFE--Y-----LT--ANTLTNFTALL------T--HS------G-----N-------------YN---------N---
AA---IK--RR----RE------HYVDELIDLLFNVAAGIQ--------R-P-E----WC---N----WT------T-----E---P---S-----CL--------LKP---HQALWL-----D--------------PWR--TQD--------------DPAFAL--ERE--RGD-WQ-AAVADDFAQWLN-----TQ----L---N-------A-------A-K-C------------AM---GMVER------KAWRT-Q-----------CQ-----------------Q-----RLREMEHVI----------------KEAQ-------S-----
---------------
WP_072926965 ------------------------------------------------------------------------MDIT--S-WIN-DR-----K----QSK----------------------------------------------------------------------------------------------L---------A-------V-FDK--DAKK------R--R--------EA-----AS-----A---AGLSALDQ----------------------------VL-----------------------------------A-EAR-RE------------------------LE--------
-LK------YE---PRTW-----LT---NA-----AQRA-------G----QIS-L----------------VTHAAKYTHGD---A-----------------Q---S-S------SL----Y----S----C-------IR---------Q--Q--EG--Y------LST--A--SL------A-------A--P--S-LD-A-VG----N----AA----A--L-----DVAKLL-Q-----T----R--F--G---------KDSL------------LDCLR-RKE-Y---SLLAEL---A---E----DS----EQLAAWVTGFSRV--L---DT--GQ----
PTSHTLAKQIYFPV---------------------------GDG---YHLLSPLFSSSLAQALHQK-IV----A--Q---R--F-S-----------------EE-SKAAR-Q-----AR-------------R--NKQWH--PL-P-VVS-----FPN-TAVMNFGG-T----KPQ---------NISALN--SSRG-----------GRMWLLSSQ-P----PQWQ--A---QE-RKPLAI----TSLF----AP-KGPFE--Y-----LT--ANTLTSFTALL------T--HS------G-----N-------------YN---------N---
-AA---IK--RR----RE------HYVDELIDLLFNVAAGIQ--------R-P-E----WC---N----WT------T-----E---P---S-----CL--------LRP---HQALWL-----D--------------PWR--TQD--------------DTAFAL--ERE--RGD-WQ-AAVADDFAQWLN-----TQ----L---N-------A-------A-K-C------------AM---GMVER------KAWRT-Q-----------CQ-----------------Q-----RLREMEHVI----------------KEAQ-------S-----
---------------
WP_101857821 ------------------------------------------------------------------------MDIT--S-WIN-DR-----K----QSK----------------------------------------------------------------------------------------------L---------A-------A-FDK--DAKK------R--R--------EA-----AS-----E---AELSALDQ----------------------------AL-----------------------------------A-EAR-RE------------------------LE--------
-LK------YE---PRTW-----LT---NA-----AQRA-------G----QIS-L----------------VTHAAKYTHGD---A-----------------Q---S-S------SL----Y----S----C-------IR---------Q--Q--EG--Y------LST--A--SL------A-------A--P--S-LD-A-VG----N----AA----A--L-----DVAKLL-Q-----T----R--F--G---------EDSL------------LDCLR-RKE-Y---SLLAEL---A---E----DS----EQLAAWVTGFSRV--L---DT--GQ----
PTSHTLAKQIYFPV---------------------------GDG---YHLLSPLFSSSLAQALHQK-IV----A--Q---R--F-S-----------------EE-SKAAR-Q-----AR-------------R--NKQWH--PL-P-EVS-----FPN-TAVMNFGG-T----KPQ---------NISALN--SSRG-----------GRMWLLSSQ-P----PQWQ--A---QE-RKPLAI----TSLF----AP-KGPFE--Y-----LT--ANTLTSFTALL------T--HS------G-----N-------------YN---------N---
-AA---IK--RR----RE------HYVDELIDLLFNIAADIQ--------R-P-E----WC---N----WT------T-----E---P---S-----CL--------LRP---HQALWL-----D--------------PWR--TQD--------------DPAFAL--ERE--RGD-WQ-AAVADDFAQWLN-----TQ----L---N-------A-------A-K-C------------AM---GMVER------KAWRT-Q-----------CQ-----------------Q-----RLREMEHVI----------------KEAQ-------S-----
---------------
WP_101836100 ------------------------------------------------------------------------MDIT--S-WIN-DR-----K----QGK----------------------------------------------------------------------------------------------L---------A-------A-FDK--DAKK------R--R--------EA-----AS-----E---AELSALDQ----------------------------AL-----------------------------------A-EAR-RE------------------------LE--------
-LK------YE---PRTW-----LT---NA-----AQRA-------G----QIS-L----------------VTHAAKYTHGD---A-----------------Q---S-S------SL----Y----S----C-------IR---------Q--Q--EG--Y------LST--A--SL------A-------A--P--S-LD-A-VG----N----AA----A--L-----DVAKLL-Q-----T----R--F--G---------KDSL------------LDCLR-RKE-Y---SLLAEL---A---E----DS----EQLAAWVTGFSRV--L---ET--GQ----
PTSHTLAKQIYFPV---------------------------GDG---YHLLSPLFSSSLAQALHQK-IV----A--Q---R--F-S-----------------EE-SKAAR-Q-----AR-------------R--NKQWH--PL-P-VVS-----FPN-TAVMNFGG-T----KPQ---------NISALN--SSRG-----------GRMWLLSSQ-P----PQWQ--A---QE-RKPLAI----TSLF----AP-KGPFE--Y-----LT--ANTLTSFTALL------T--HS------G-----N-------------YN---------N---
-AA---IK--RR----RE------HYVDELIDLLFNIAAGIQ--------R-P-E----WC---N----WT------T-----E---P---S-----CL--------LRP---HQALWL-----D--------------PWR--TQD--------------DTAFAL--ERE--RGD-WQ-AAVADDFAQWLN-----TQ----L---N-------A-------A-K-C------------AM---GMVER------KAWRT-Q-----------CQ-----------------Q-----RLREMEHVI----------------KEAQ-------S-----
---------------
WP_128179519 ------------------------------------------------------------MPE------------AELI--NYIV--D-FIA-GR-----R----QAK----------------------------------------------------------------------------------------------L---------E-------A-FDK--EARK------R--L--------AS-----VE-----A---EESDVLQQ----------------------------AL-----------------------------------L-QQR-TV------------------------LE---
------DR------YT---PGNW-----LT---DA-----ANRA-------G----QIS-L----------------VTHAAKYTHGD---S-----------------K---S-S------SV----Y----S----Q-------IT---------N--N--EG--Y------VST--A--SL------R-------N--T--V-VD-A-VG----N----AA----M--L-----DVAKLL-Q-----M----E--H--D---------GDSL------------LACLQ-RKD-Y---RVLEAL---A---E----DP----QQLEQWITGFNQA--L---TP--HQ----
PASHKLAKQVYFPV---------------------------GDG---YHLLSPLFSSALAQALYQR-II----D--A---R--F-S-----------------EG-SKAAR-Q-----AR-------------R--DGEWC--AE-P-VVY-----FPA-TAIMNFGG-T----NSQ---------NISLLN--SRRG-----------GRVFLLSAQ-P----PLWK--G---RE-KPPLGM----KTLF----GH-RGPFD--H-----LA--RNVISQLSHLL----------T------R-----Q-------------EN---------N---

[illegible]

WP_144812048 [illegible]

WP_046449893 [illegible]

WP_130985587 [illegible]

WP_130971823 [illegible]

WP_043494060 [illegible]

WP_059308552 [illegible]

WP_032616959 [illegible]

WP_144883571 [illegible]

WP_102103906 [illegible]

WP_124024297 [illegible]

WP_121265582 [illegible]

WP_037588610 [illegible]

WP_061275474 [illegible]

WP_126358484 [illegible]

WP_080801368 [illegible]

WP_007293313 [illegible]

WP_013163749 [illegible]

WP_036503038 [illegible]

WP_123016005 [illegible]

WP_133564012 [illegible]

WP_067036487 [illegible]

WP_114411031 [illegible]

WP_139758236 [illegible]

WP_100187842 [illegible]

WP_115897803 [illegible]

WP_113916936 [illegible]

WP_044618898 [illegible]

WP_121468036 [illegible]

WP_028886984 [illegible]

WP_018414916 [illegible]

WP_084088883 [illegible]

WP_012729574 [illegible]

WP_127958201 [illegible]

WP_100295269 [illegible]

WP_116727003 [illegible]

WP_118881659 [illegible]

WP_113069879 [illegible]

WP_017788587 [illegible]

WP_099358605 [illegible]

WP_139730391 [illegible]

WP_139740171 [illegible]

WP_052441382 [illegible]

WP_042038594
[illegible]

WP_087755246
[illegible]

WP_144714573
[illegible]

WP_076573269
[illegible]

WP_136613941
[illegible]

WP_107864571
[illegible]

WP_070070703
[illegible]

WP_130803135
[illegible]

WP_038342050
[illegible]

WP_113997528
[illegible]

WP_067670944
[illegible]

WP_086187542
[illegible]

WP_075174487
[illegible]

WP_045796730
[illegible]

WP_104472260
[illegible]

WP_087536693
[illegible]

WP_005163439
[illegible]

WP_086196694
[illegible]

WP_122902223
[illegible]

WP_099339711
[illegible]

WP_004752521
[illegible]

WP_068975385
[illegible]

WP_130130897
[illegible]

WP_018125463
[illegible]

WP_011996910
[illegible]

WP_131359022
[illegible]

WP_006365989
[illegible]

WP_081491843
[illegible]

WP_058069158
[illegible]

WP_014848626
[illegible]

WP_014500608
[illegible]

WP_023593282
[illegible]

WP_011240574
[illegible]

WP_141931626
[illegible]

WP_012817199
[illegible]

WP_086650437
[illegible]

WP_048838277
[illegible]

WP_086657827
[illegible]

WP_086552796
[illegible]

WP_122102140
[illegible]

WP_100313076
[illegible]

WP_100318839
[illegible]

WP_100320368
[illegible]

R----MQ---QD--QK----RR------AMEQRLGSILAAFAVTVR--D-----S-Y-P-----P---G----WT------R-----D---A---A-----CR--------LPL---CEQLWL-----D-------------PER--ARLPNRDQAQDPDGYAADEIFKA--AWQ--RQD-WP-DEIAERFAQWLN-----QW----L---R------DN-------G-F-A------------V----GSVEL------RHFAR-Q-------AL--VDT----------------D-----WPNPIRRDL-----------------PSSR-------
DWHTEVSA------------------------------
WP_100310800 ------------------------------------------MNL---------------RA-------------SNFR--QAIA--R-FIK-ER-----R----DAK------------------------------------------------------------------------------------L---------K-----------------G----------A---------EA------AD----SA---------QA-----------------------------A------------------------------------K--------------------
YD---YDTW-----LA---DA-----ARRV-------T----QIQ-M----------------VTHVLKATHPD---A----------------K---G-S------SL----F----V----P---------P-ANL---PAR--D---E---------LGS--H--VL------G-R-----Q--F--A-ED-V-VG-----N----AA----A--L-----DVYKLL-K-----L----E--V--E---------GKRL------------LDWLL-QED-A---DLLAAL---S---D-----NP-----DTARAYADAFCGL--Q---RE--AD-S--
YASHTYAKQLYWCA-------GE-----DA------CD----DAQ---YILLQPMFPSSLVQAVHEA-IQ----Y--S---R--F-E-----------------VA-IVEAR-R-----AR-------------R--EKKPS--DM-Q-IPE-----YPE-LAIRKLGG-T-----KPQ----------NISQLN--SERG------------GSNYLLASF-P----PIWD--G-Y-RP-RSFLNL----DTAF-----MR-----LL--H-F--HDM--QQQLDAFARHL----------G-------A-----Y--------------KQR----------
R----MQ---QD--QK----RR------AMEQRLGNTLVAFAATVR--D-----S-Y-P-----P---G----WT------R-----D---A---A-----CQ--------LPL---CEQLWL-----D-------------PER--TRLPNRDQAQDPDGYAADETFKV--AWQ--RQD-WP-DEVAERFAQWFN-----QW----L---R------DN-------G-F-A-------------V----GSVEL------RHFAR-Q-------AL--VDT----------------D-----WPNPIRRDL-----------------PSNR-----
---DWHTEVSA------------------------------
WP_100317018 ------------------------------------------MNL---------------RA-------------SNFR--QAIA--R-FIK-ER-----R----DAK------------------------------------------------------------------------------------L---------K-----------------G----------A---------EA------AD----SA---------QA-----------------------------A------------------------------------K--------------------
YD---YDTW-----LA---DA-----ARRV-------T----QIQ-M----------------VTHVLKATHPD---A----------------K---G-S------SL----F----V----P---------P-ANL---PAR--D---E---------LGS--H--VL------G-R-----Q--F--A-ED-V-VG-----N----AA----A--L-----DAYKLL-K-----L----E--V--E---------GKRL------------LDWLL-QAD-T---DLLAAL---S---D-----NP-----DTARAYADAFCSL--Q---RE--AD-S--
YASHTYAKQLYWCT-------GE-----DA------CD----DAQ---YILLQPMFPSSLVQAVHEA-IQ----Y--S---R--F-D-----------------VA-IVEAR-R-----AR-------------R--EKKPS--DM-Q-IPE-----YPE-LAIRKLGG-T-----KPQ----------NISQLN--SERG------------GSNYLLASF-P----PIWD--G-Y-RP-RSFLNL----DTAF-----MR-----LL--H-F--HDM--QQQLEAFARHL----------G-------A-----Y--------------KQR----------
R----MQ---QD--QK----RR------AMEQRLGGTLTAFAATVR--D-----S-Y-P-----P---G----WT------R-----D---A---A-----CR--------LPL---CEQLWL-----D-------------PER--ARLPNRDQAQDPEGYAADETFKA--AWE--KQD-WP-DEIAERFAQWLN-----QW----L---R------DN-------G-F-A-------------V----GSVEL------RHFAR-Q-------AV--VDA-------------------N-----WPNPIRRDL-----------------PSSR------
-DWHTEVSA------------------------------
WP_100308822 ------------------------------------------MDS---------------RK-------------RKFY--GAIA--R-FIK-ER-----R----DAK------------------------------------------------------------------------------------L---------K-----------------D----------A---------EA------AD----SA---------QA-----------------------------A------------------------------------K--------------------
YD---YDTW-----LA---DA-----ARRV-------A----QIQ-M----------------VTHVLKATHPD---A----------------K---G-S------SL----F----V----P---------P-ANL---PAR--D---E---------LGS--H--VL------G-M-----Q--F--A-ED-V-VG-----N----AA----A--L-----DVYKLL-K-----L----E--V--E---------GQRL------------LDWLL-QED-A---DLLAAL---S---D-----NP-----DTARAYADAFCSL--Q---RE--AD-S--
YASHTYAKQLYWCA-------GE-----DA------CD----DAQ---YILLQPMFPSSLVQAVHEA-IQ----Y--S---R--F-D-----------------VA-IVEAR-R-----AR-------------R--EKKPS--DM-Q-IPE-----YPE-LAIRKLGG-T-----KPQ----------NISQLN--SERG------------GSNYLLASF-P----PIWD--G-Y-RP-RSFLNL----DTAF-----MR-----LL--H-F--HDM--QQQLEAFARHL----------G-------A-----Y--------------KQH----------
Q----MQ---QD--QK----RR------AMEQRLGGTLTAFAATVR--D-----S-Y-P-----P---G----WT------R-----D---A---A-----CR--------LPL---CEQLWL-----D-------------PER--AQLPNRDQAQDPEGYAADETFKA--AWE--KQD-WP-DEIAERFAQWLN-----QW----L---R------DN-------G-F-A-------------V----GSVEL------RHFAR-Q-------AV--VDA-------------------N-----WPNPIRRDL-----------------PSSR------
--DWHTEVSA------------------------------
WP_100326660 ------------------------------------------------------------MD--SRK---------RKFY--EAIA--R-FIK-ER-----R----DAK------------------------------------------------------------------------------------L-----------------------K------D--A-------------------------------------EA--------------------------------A-DSA-QA--------------------AK--------------------
YD---YDTW-----LA---DA-----ARRV-------A----QIQ-M----------------VTHVLKATHPD---A----------------K---G-S------SL----F----V----P---------P-ANL---PAR--D---E---------LGS--H--VL------GR-------Q--F--A-ED-V-VG-----N----AA----A--L-----DVYKLL-K-----L----E--V--E---------GQRL------------LDWLL-QAD-T---DLLAAL---S---D-----NP-----DTARAYADAFCSL--Q---REA--DS----
YASHTYAKQLYWCT-------GE-----DA------CD----DAQ---YILLQPMFPSSLVQAVHEA-IQ----Y--S---R--F-D-----------------VA-IVEAR-R-----AR-------------R--EKKPS--DM-Q-IPE-----YPE-LAIRKLGG-T-----KPQ----------NISQLN--SERG------------GSNYLLASF-P----PIWD--G-Y-RP-RSFLNL----DTAF-----MR-----LL--H-F--QDM--QQQLEAFARHL----------G-------A-----Y--------------KQH----------
Q----MQ---QD--QK----RR------AMEQRLGGTLTAFAATVR--D-----S-Y-P-----P---G----WT------R-----D---A---A-----CR--------LPL---CEQLWL-----D-------------PER--AQLPNRDQAQDPEGYAADETFKA--AWE--KQD-WP-DEIAERFAQWLN-----QW----L---R------DN-------G-F-A-------------V----GSVEL------RHFAR-Q-------AV--VDA-------------------N-----WPNPIRRDL-----------------PSSR------
--DWHTEVSA------------------------------
WP_054021653 ------------------------------------------MELP-NALS----------PRS-------------GAFR--AAIV--A-FVD-AR-----R----EAK------------------------------------------------------------------------------------L---------K-------------------------------------R-----DE-----KA----------GT--------------------------AA-------------------------------------SR--------------------
--YD---PDVW-----LA---DA-----ARRA-------P----QLQ-A----------------ATHVAKATHPD---A----------------R---G-S------NL-----H----V----A---------P-DKL---PQH--A---E---------IGS--H--SL------G-H------A--F--A-ED-V-VG---------AG-----A--L-----GVFKFL-Q-----L----E--V--E---------GRRL------------LDWML-LGD-E---DLQHAL---H----P-----DP-----AVARAWMAAFCAL--V---RE--AG-P--
PRSHGLAKQVYWLV-------GD-----DP------SN-----DDD---FHLLQPMFSSSLAQALHAS-VQ-----E--I------F-G-------------------DD-AKAAR-Q-----AK-------------R--DRAPH--EG-T-FFE------YRA-LAMRKLGG-S-----KPQ----------NISQLN--SERG------------GNNLLLASL-P----PLWR--E-L-AG-LSMFGQ----RSAI-----EC-----LP--R-L--AEV--RTLLGRLVTCL------------R-------D------A-----------------AGK---------
-T-----ME---TR--IE-----RE------RIERALGWQLPVFFEMVQ--A------R-H-E------A---G-----WT------R------S----P----L-----CE--------LPL---CEQLWL-----D-------------PER--TELPPREDSL------EEDLAFNE--AFA--SGH-WA-DEVAERFSRWLN-----GS-----L---R------DH-------G-I-T--------------AV----GDVEF-------RHWAR-Q-------AL--VD-V---------------------A-----WPMSMQRRA---------------------SGKA-------TA--
---------------------------------
WP_067646235 ------------------------------------------MTEL-----T---------DRG---------------KAFR--SAVT--T-FIE-AR------R----EVD------------------------------------------------------------------------------R----------K------------------K-----------------AK------HD------KA----------KA----------------------------AS-----------------------------------K--------------------
YD---YDTW-----LA---DA-----ASRA-------H-----NLQ-V-----------------VTHPIKFTHGS----I------------------K---GAS------SI-----H-----A------R----------T-KRI---FVH--R---E---------IGT--H--SI------D-E------L--K--E-DF-A-IS-----D-----SK-----H--L-----DVYSLL-K-----E-----V--V--D---------GRSL------------LDWLR-EDD-A---DFISVL---S---S-----DP-----VKAAALADSLNKV--L---RQ--DY-E--VVSHPLAKQIYWLK--
-----DA-----SP------AD-----NAH-----YHLLQPMFSSSLEHVVNAD-IR-----A--V----R-----------------------EF-AFAAR-G------TK-------------K--QKPTF---AD----HST------YSG-LLGRALGG-S-----NAQ----------NISPQN--KVRG------------GMNYLLASL-P-----PTWA--R-E-LP-RNLLHL------ESVL-----ER-----FT---Y-F--EGV--QSLVQSLAGFL-------Q--SV------A------S--------------RQR-----------N-----RK----IN--DR-----
RE-------DIEQALGFQLPVFAASIQ--A------S-L-E------P---G-----WT------R------D----P----E-----CR--------LPL---CEQLWL-----D-------------PER--AGLPIREHPESPEWTQQDLEFNA--AYE--FGD-WP-DQVAGRFANWVN-----AQ-----L---R------EA-------G-L-T--------------AV----GDAEY-------KHWAK-Q-------AI--VD-A---------------------A-----WPVSLQRRA-------------------PPGG-------QT-------------
------------
WP_052191963 ------------------------------------------MEDR-----L----------TRS-------------QRFQ--IAIE--R-FIT-ER------R----DTK------------------------------------------------------------------------------L----------N-----------------G-------------------KD-----DP----------VT---------------------------AA---------------------------------------K--------------------
YD---YSTW-----LA---DA-----ARRV-------S----QIQ-A----------------VTHVLKATHPD---A----------------R---G-S------SL----Y-----I-----D----------P-TSL---PGR--D---E---------IGS--H--SL------G-A------D--F--A-ID-I-VG-----N-----AA-----A--L-----DVYKFL-K-----L-----Q--V--D---------NRSL------------LDWLQ-DDD-A---DLLAAL---S----P-----QP----ERASTWATAFKGL--I---RQ--DE-T--
VTSHERAKQLYWCV-------DG-----DP------AR-----TEA---FHLLEPLFSSSLVHAVHLE-VN-----D--A---R--F-G-------------------EA-NKEAR-L------AW-------------R--GGRAH--PN-G-YRE------YRG-LLVRKLGG-T-----KPQ----------NISQLN--SERG------------GVNYLLSSL-P-----PAWQ--R-Q-PS-KPLLLL-----DSSF-----KR-----FQ---R-F--DGV--QRQLNALYQYF-----------E-------Q--------A-------------------PDP-----
-----T-----FA---TR---KR-----RE------RIERALGLALTAFGVSIR--T-----Q-F-P------A---G-----WT------R------D----P----D-----CQ--------LPL---CEQLWL-----D-------------PGR--AEMAIREGYE------DMDGAFQQ--AAE--RQQ-WP-EEVAKHFALWLN-----AM-----L---S------RK-------G-F-A--------------V-----SDAEH-------MQWAR-K-------AL--AHTL----------------------G------FAVSTREVS----------------HG-----------
---------------------------------
WP_145470975 ------------------------------------------MEIL-----T----------ERA-------------QAFR--QAIT--A-FID-GR------R----DAK------------------------------------------------------------------------------L----------K-----------------G-------------------D------DN-----DG-----------DI--------------------------AA---------------------------------------K--------------------
YE---YGIW-----LA---DA-----ARRV-------N----QIQ-A----------------VTHVLKATHPD---A----------------R---G-S------SL----H-----L-----A----------P-INL---PQH--R---E---------IGS--H--LL------G-D------T--F--A-ED-I-VG-----N-----AA-----A--L-----DVFKFL-K-----T-----E--V--E---------GKRL------------FDWMQ-EAD-A---DLQAAL---S-----A-----DA----ETAQEWMQAFSSL--I---RA--ES-Q--
PVSHVAAKQLYWLV-------GE-----EP------TD-----NAE---YQLLQPLFSSALAHKVHAD-IQ-----A--A---R--F-G-------------------EE-NKLAR-Q------AR-------------R--ENSAY--DG-S-YHD------YRG-LVVRKLGG-T-----KPQ----------NISQLN--SERG------------GVNYLLASL-P-----PHWN--T-S-DR-IKVLNV-----DSVF------HS-----FY---F-F--DDV--AELLKALSSLL-----------L-------S--------D------------------PDA---------
V-----KA-----TR---EK-----RE------ALAQALGQQLALFGESIR--F------A-Q-E------P---G-----WS------Q------H----P----D-----CR--------LPQ---CQRLWL-----D-------------PER--IELLPREEHQ------AEDEQFNA--AYQ--WGD-WP-DEIAGYFASWLN-----AQ-----L---H------KT-------G-L-T--------------TV----GDAEY-------TYCAK-Q-------AI--VD-A---------------------D-----GPVSLQHRA--------------------P-GG----------VA-----
---------------------------------
WP_129142165 ------------------------------------------MNETTEND-G--V----------SRI------------ERFR--LAIA--T-FID-ER------R----VAK------------------------------------------------------------------------------L----------K-----------------G-------------------R------ED-----DA----------AA-----------------------------AA-----------------------------------K--------------------
--YE---YDTW-----LA---DA-----ARRV-------G----QIQ-A----------------VTHVLKASHPD---A----------------R---G-T------SL----Y-----I-----P----------P-QTL---PQH--D---E---------IGS--H--SL------A-D------D--F--A-ED-V-VG-----N-----AA-----A--L-----DVFKFL-K-----V-----E--V--E---------GRSL------------LEWVK-ARD-A---DLLQAL---H-----A-----DE-----AIAQSRMDAFGNL--V---RS--GT----
FSSHALAKQVYWLT-------QG-----NA-------AT-----DDN---FHLLQPLFSSALAHRVHAD-IQ-----S--A---R--F-G-------------------EE-NRDVR-A------AV-------------R--TKQAD--TR-V-YRD------YRN-LAVRKMGG-T-----KPQ----------NISQLN--SERG------------GANYLLPSL-P-----PNWS--Q-Q-RP-PNLLGL-----ESST-----ER-----FT---F-F--AGV--RPLVRELARLV-----------L-------S-------K-----------------RSK---------
--T-----ME---TR--QE-----RE------ALEQRLGAKLPFFAAAIH--S------Q-L-S------P---G-----WT------R------D----D-----R-----CE--------LSL---CECLWL-----D-------------PER--LTLAPRPLGEGG----EEDQAFVE--AYH--WGD-WP-DQVAGIFASWLN-----DQ-----L---R------RA-------G-A-T--------------AV----GDSEY-------RHWAR-Q-------AI--ID-A---------------------A-----WPVPMQRRA----------------------PEER--------A-
---------------------------------
WP_123659456 ------------------------------------------MSETTDHP-TGTA---------ARR-------------DRFR--QAIA--D-FID-AR------R----TAK------------------------------------------------------------------------------L----------K-----------------G-------------------R------ED-----DT----------AA-----------------------------AA-----------------------------------K--------------------
------YE---YETW-----LA---SA-----ARRV-------G----QIQ-A----------------VTHVLKASHPD---A----------------R---G-S------SL----Y-----I-----P----------P-TDL---PTR--D---E---------IGS--H--SL------G-S------D--F--T-DD-V-VG-----N-----AA-----A--L-----DVFKFL-K-----V-----E--V--D---------GRAL------------LDWVK-QDD-P---DLKHAL---H-----A-----DA-----NTADERMQAFGNL--V---RS--GA----
PSSHALAKQVYWLI-------DG-----RP------DD-----NRN---FHLLQPLFSSALAHRVHAD-IQ-----A--A---R--F-G-------------------EE-NKALR-Q------AK-------------K--SDKPH--DG-I-YRD------YRQ-LAVRKLGG-S-----NTQ----------NVSQLN--AERR------------GTNYLFASL-P-----PTWS--R-M-RA-ITLLEL-----ESST-----RQ-----FF---S-F--EGV--RDLLRRLVQLL-----------Q-------S--------D-----------------PKR---------
T-----ME---TR--QK-----RE------AIEQALGARLPLFAATIR--D-----Q-L-S------A---G-----WT------R------D----P-----T-----CE--------LPL---CEQLWL-----D-------------PDR--LTLNKRPAGEGG---EADHAFEQ--AYH--WGD-WP-DDVAGIFANWLN-----EQ-----L---R------RV-------G-V-T--------------SV----GDMEY-------RHWAR-Q-------AI--VD-A---------------------A-----WPIPMQRRA-------------------PEDR--------S--
---------------------------------
WP_037336895 ------------------------------------------MPDTTDNQ-TGTA---------ARS-------------ERFR--EAIA--D-FIN-AR------R----VAK------------------------------------------------------------------------------L----------K-----------------G-------------------R------ED-----DA----------AA-----------------------------AA-----------------------------------K--------------------
------YE---YETW-----LA---SA-----AHRV-------S----QIQ-A----------------VTHVLKASHPD---A----------------R---G-S------SL----Y-----I-----P----------P-TDL---PLR--D---E---------IGS--H--SL------G-N------E--F--T-ED-V-VG-----N-----AA-----A--L-----DVFKFL-K-----V-----E--V--D---------GRSL------------LDWVK-QDD-T---DLKQAL---H-----T-----DE-----DIAHRRMQAFGNL--V---RS--GT----
PSSHALAKQVYWLV-------DG-----RP------DD-----NHN---FHLLQPLFSSALAHRVHAD-IQ-----A--A---R--F-G-------------------EE-NKAGR-Q------AK-------------R--SKKPH--DG-I-YQD------YRQ-LAVRKMGG-T-----KPQ----------NISQLT--SERG------------GTNYLLASL-P-----PTWS--L-M-RA-TTLLGR-----ESST-----GR-----FF---Y-F--EGV--RGLVRSLGRLL-----------R-------S-------D-----------------PKK---------
---T-----LE---TR--QK-----RE------ELEKALGAQLPLFAATIR--D-----Q-L-S------P---G-----WT------R------D----P-----A-----CE--------LPL---CEQLWL-----D-------------PDR--LTLDARLLGEGG---EDDRAFEQ--AYH--WGD-WP-DQVAGIFANWLN-----DR-----L---R------HA-------G-I-T--------------SV----GDIEY-------RHWAR-Q-------AI--VE-A---------------------A-----WPVPMQRRA-------------------AEDR--------S--
---------------------------------
WP_109061065 ------------------------------------------MEDPMSNR-SERS---------PRS-------------ERFR--TTIS--D-FIN-TR------R----DAK------------------------------------------------------------------------------L----------K-----------------G-------------------K------ED-----DA----------EA-----------------------------AA-----------------------------------K--------------------
------YD---YATW-----LG---DA-----ARRV-------G----QIQ-A----------------VTHVLKATHPD---A----------------R---G-S------SL----H-----V-----H----------P-ETL---PQH--P---E---------IGS--H--SL------G-E------N--F--A-ED-V-VG-----N-----AA-----A--L-----DVFKFL-K-----L-----A--V--D---------GRKL------------LDWMQ-VAD-A---DLLAAL---H-----T-----DP-----TTAQEWATAFSGL--V---RT--ET-Q--
AVSHAKAKQLYWLT-------GN-----DP------SD-----NSQ---YLLLQPLFSSSLAHAVHDD-IQ-----E--A---R--F-G-------------------EP-NKLAR-Q------AR-------------R--SKEAH--DS-P-YRD------YRN-LIVRKLGG-T-----KPQ----------NISQLN--SERG------------GINYLLASLPP-----QRWQ--D-Q-DK-IKLHNV-----KSAL-----DG-----LR---N-L--PQV--RSLIQELAAFL-----------L-------T--------N-----------------PPP---------
T-----MK---TR--LK-----RE------AIEQALGAQLPVFVERIK--V-----Q-M-Q------HP---G-----WT------R------T----D-----CE--------LAL---CEQLWL-----D-------------ADR--TGLDLRETPEEQ---KKDQDFNA--AYH--WGD-WP-DQVAGRFANWVN-----EQ-----L---H------VA-------G-L-T--------------IV----GTSEA-------KHWAR-Q-------AI--ID-A---------------------A-----WPVPMQRRM---------------------GE-------TA--
---------------------------------
WP_109719119 ------------------------------------------MREE-ND----------RV-------------RVFR--EAIT--T-FIH-ER------R----DAK------------------------------------------------------------------------------L----------K-----------------G-------------------DD-----DA----------DK------------------------------AA-----------------------------------K--------------------
YE---YATW-----LA---DA-----ARRV-------S----QIQ-A----------------VTHVLKATHPD---A----------------R---G-S------SL----H-----A-----A----------P-QSL---PSH--S---E---------IGS--H--TL------A-A------D--F--A-ED-I-VG-----N-----AA-----A--L-----DVFKFL-K-----M-----E--V--D---------GRRL------------FDWVL-EKD-V---DLQRAL---S-----P-----DA-----ETAAEWMEAFGGL--L---RT--AV-Q--
PKSHALAKQVYWCV-------GD-----AP------AD-----DGH---YQLLQPLFSSSLMHAIHAD-IN-----D--A---R--F-G-------------------EQ-NKVAR-K------AH-------------R--EKTAS--DG-P-YRD------YRG-VVARKLGG-T-----KPQ----------NISQLN--SERG------------GVNYLLASV-P-----PRWH--K-E-TS-PRLLNT-----DSVL-----DR-----IG---Q-L--GSV--RHLIRVLTDFL-----------L-------K-------D-----------------PPN---------
-N-----EA---TR--NR-----RE------AIEQALSQELPVFAASIR--A------G-L-A------P---G-----WT------R------D----P-----A-----CR--------LPR---CEQLWL-----D-------------SDR--VELPLREDHA------EIDQAFID--DYH--QGD-WR-DEVAGRFANWIN-----AR-----L---R------DA-------G-L-S--------------GV----GDAEY-------HHWAR-Q-------AV--IE-A---------------------Q-----WPVPMQARA-----------------------SA-----------
---------------------------------
WP_123212514 ------------------------------------------MEV-TN----------RS-------------LKFR--SAIK--A-FID-AR------R----DDK------------------------------------------------------------------------------L----------R-----------------G-------------------K-----EG-----GD----------NA-----------------------------AA-----------------------------------K--------------------
YE---YGVW-----LA---DA-----ARRV-------L----QIQ-A----------------VTHVLKATHPD---A----------------R---G-S------SL----H-----A-----P----------P-SAL---SVR--T---E---------IGT--H--SL------G-A------E--F--V-DD-I-VG-----N-----AA-----A--L-----DVFKFL-K-----L-----E--V--D---------GRRL------------LDWMQ-DGD-A---DLLAAL---H-----E-----DE-----TVAVGWVNAFKGL--V---RS--DV-Q--
ASSHQAAKQLYWLV-------GD-----DP------AD-----DAH---YQLLQPLFSSSLAHAVHAR-IQ-----D--A---R--F-S-------------------EA-SVAAR-K------AR-------------R--EKQPH--AK-G-YSD------YRN-LVARKLGG-T-----KPQ----------NISQLN--SERG------------GVNYLLASL-P-----PKWD--R-D-NR-PAVLGV-----ESAM-----KR-----FA---R-F--EGV--AELVDQLAKFL-----------L-------S-------N-----------------PDR---------
----N-----AE---TR--DK-----RE------AIEQALGLQLPIFAASVF--A------H-M-E------P---G-----WT------R------D----E-----R-----CM--------LPL---CEQLWL-----D-------------PER--TQLAVREDHR------QDDEDFKR--AYT--WAD-WP-DEVAGRFANWLN-----EQ-----L---R------SK-------G-L-P--------------AL----GDAEF-------KHWGK-Q-------AI--IE-A---------------------D-----WPVPIQRRA-----------------------EVAS-----------
---------------------------------
WP_071230851 ------------------------------------------MSKE-SK----------RS-------------GHFR--SAIA--A-FIE-AR------R----EVK------------------------------------------------------------------------------L----------K-----------------G-------------------K------ED-----DA----------EA-----------------------------AA-----------------------------------K--------------------
YE---YAAW-----LA---DA-----ARRV-------T----QIQ-A----------------VTHVLKASHPD---A----------------R---G-S------SL----H-----A-----E----------P-ETL---PSH--A---E---------VGS--H--LL------R-D------S--Y-S-ED-I-VG-----N-----AA-----A--L-----DVFKFL-K-----L-----E--V--E---------NRRL------------LDWMR-DGD-P---DLRDAL---S-----A-----DA-----ELASGWMEAFSSL--V---RS--NT-A--
PVSHEMAKQVYWLI-------GN-----EP------AR-----NDQ---YHLLQPMFSSSLAHAVHAD-IQ-----D--A---R--F-G-------------------EA-NKLAR-Q------AF-------------R--AKETY--NE-E-YRD------YRN-LIARKLGG-T-----KPQ----------NVSQLN--SERG------------GVNYLLASL-P-----PKWQ--Q-E-RK-PRLLNL-----DSAV-----ER-----FA---W-Y--EGV--PFLLRKLAAFL-------K--SV-------A--------D-----------------KPR---------
-----N-----MD---LV---EK-----RE------AIEQALGQQLALFAVSIR--A-----S-L-E------P---G-----WT------R------D----P-----D-----CT--------LPL---CEKLWL-----D-------------PER--TELPLRRDPEQA---DDDRAFNA--AYT--FGD-WP-DEVAGRFANWVN-----ER-----L---R------EA-------G-L-T--------------AV----SEVEY-------RHWAK-Q-------AI--ID-A---------------------A-----WPVPMQRRA-------------------SAGG---------
EA---------------------------------
WP_055773113 ------------------------------------------MQEL-TD----------RG-------------KGFR--MAIA--G-FIE-AR------R----EAK------------------------------------------------------------------------------L----------K-----------------G-------------------K-----EE-----EG----------DI-----------------------------ST-----------------------------------K--------------------
YD---YGAW-----LA---DA-----ARRV-------G----QIQ-A----------------VTHVLKATHPD---A----------------R---G-S------SL----H-----V-----R----------P-DAL---PRH--A---E---------IGT--H--VL------G-D------D--F--A-ED-V-VG-----N-----AA-----A--L-----DVFKFL-K-----V-----E--V--E---------GRRL------------LDWMQ-AGD-A---DLRSAL---H-----A-----DS-----ETAANWMKAFSGL--V---RG--DT-H--
PTSHEAAKQLYWLV-------DG-----AP------TE-----DEA---YHLLQPLFSSSLAHAVHAE-IQ-----D--A---R--F-G-------------------ED-NKAAR-Q------AF-------------R--AGEAH--EA-P-YRD------YRN-LVARKLGG-T-----KPQ----------NVSQLN--SERG------------GINYLLASL-P-----PSWD--I-E-QS-RKLLNL-----ESAM-----DR-----LA---W-F--PGV--RDTVDQLASFL-----------L-------S-------N-----------------PPP---------
--T-----EA---TR--RK-----RT------ELAQELGAQLAIFAVETQ--V------R-F-E------P---G-----WT------R------N----E-----S-----CT--------LPL---HQQVWL-----D-------------PER--VELDPRPEPE---LAAEDEAFVA--IWE--FGD-WP-DQVAGDFANWLN-----DR-----L---R------SA-------G-L-I--------------AV----GDAEY-------RHWAR-H-------AV--AD-A---------------------T-----WPVPVRRRV-------------------PKGG---------
TA---------------------------------
WP_056932447 ------------------------------------------MQEL-TD----------RG-------------KAFR--VAIA--G-FIE-AR------R----EAK------------------------------------------------------------------------------L----------K-----------------G-------------------K-----EE-----EG----------DI-----------------------------SA-----------------------------------K--------------------
YD---YNAW-----LA---DA-----ARRV-------G----QIQ-A----------------VTHVLKATHPD---A----------------R---G-S------SL----H-----V-----R----------P-DAL---PRR--A---E---------IGT--H--VL------G-D------D--F--A-ED-V-VG-----N-----AA-----A--L-----DVFKFL-K-----I-----E--V--E---------GRRL------------LDYMQ-AGD-A---DLRSAL---H-----P-----DA-----GTATAWMKAFSGL--V---RG--DT-H--
PASHEAAKQLYWLV-------DG-----PP------TE-----DEA---YHLLQPLFSSSLAHAVHAE-IQ-----G--A---R--F-G-------------------ED-NKAAR-Q------AF-------------R--AKEAH--ET-S-YRD------YHN-LVVRKLGG-T-----KPQ----------NVSQLN--SERG------------GINYLLASL-P-----PSWD--I-E-QP-RKLLNL-----ESAM-----DR-----LG---W-F--PDV--RDTVDQLASFL-----------L-------S-------N-----------------PPP---------
--T-----EA---TR--RK-----RT------ELAQELGAQLAIFAVETQ--V------R-F-E------P---G-----WT------R------N----E-----S-----CT--------LPL---HQQVWL-----D-------------PER--VELDQRPQPE---LAAEDEAFVA--VWE--FGD-WP-DQVAGDFANWLN-----DR-----L---R------SA-------G-L-V--------------AV----GDAEY-------RHWAR-H-------AV--TD-A---------------------A-----WSAPEQRRK-------------------PKGG---------
IA---------------------------------
WP_128421554 ------------------------------------------MTEE-SGRS---------ARS-------------EHFR--TAIA--T-FID-TR------R----EAK------------------------------------------------------------------------------L----------K-----------------G-------------------K-----ED-----GA----------DT-----------------------------AS-----------------------------------K--------------------
-YA---YTTW-----LA---DA-----ARRV-------G----QIQ-A----------------VTHVLKATHPD---A----------------R---G-S------SL----H-----V-----S----------P-GAL---PRH--V---E---------IGS--H--VL------G-E------D--Y--A-ED-V-VG-----N-----AA-----A--L-----DVFKFL-K-----L-----E--V--E---------GRRL------------LDWMQ-AGD-G---DLRDAL---S-----A-----DA-----AVATDWMSAFGSL--V---RR--DP-Q--
PSSHEAAKQVYWLA-------GE-----EP------AD-----DIQ---YHLLQPLFSSSLAHAVHAD-IR-----E--A---R--L-G-------------------DG-NKGAR-Q------AY-------------R--DKRPF--DG-V-YRD------YRN-LVARKLGG-T-----KPQ----------NISQLN--SDRG------------GVNYLLASL-P-----PHWT--Q-E-RP-RSLLHL-----DSAL-----DR-----FA---W-F--EGV--RELIKALGDFL-----------L-------T-------D-----------------PPK---------
-N-----DA---TR--HR-----RE------AIEQALGLQLAYFASTIH--A------R-F-E------P---G-----WT------R------D----P-----D-----CR--------LPL---CERLWL-----D-------------SVR--IDLPNRQDLQHPEWERDDGDFKA--AYD--VGD-WP-DEVAGRFANWIN-----DQ-----L---R------EA-------G-V-S--------------GL----GDDQY-------RHWAK-Q-------AI--VD-A---------------------A-----WPVPLQRHA-------------------PAGG---------
AA---------------------------------
WP_046978243 ------------------------------------------MKE-----LN---------DRG-------------KAFR--TAIA--A-FID-TR------R----EAK------------------------------------------------------------------------------L----------K-----------------G-------------------K-----ED-----GA----------DA-----------------------------AS-----------------------------------NK-------------------
YE---YGTW-----LA---DA-----ARRV-------G----QIQ-A----------------VTHVLKATHPD---A----------------R---G-S------SL----H-----V-----S----------P-AAL---PRH--V---E---------IGS--H--VL------G-E------D--Y--A-ED-V-VG-----N-----AA-----A--L-----DVFKFL-K-----L-----E--V--A---------GRRL------------RDWMQ-AGD-D---DLRDAL---N-----A-----DA-----AVATDWMSAFGSL--V---RS--DP-Q--
PSSHEAAKQIYWLA-------GE-----AP------AD-----DTQ---YHLLQPLFSSSLAHAVHAE-IR-----E--A---R--F-G-------------------EG-NKTAR-Q------AY-------------R--DKQPF--DG-I-YRD------YRH-LVARKLGG-T-----KPQ----------NISQLN--SDRG------------GVNYLLASL-P-----PRWT--R-E-RP-RSLLNL-----ESAQ-----ER-----FA---W-F--EGV--RHLLKALGDFL-----------L-------S-------N-----------------PPQ---------
-N-----QE---TR--NT-----RE------AIEQALGLQLAYFASTIH--A------R-F-E------P---G-----WT------R------D----P-----N-----CE--------LPL---CERLWL-----D-------------PAR--IDLPNREDSEHPEWQQQDRDFKA--AYD--FGD-WP-DEVAGRFANWLS-----DQ-----L---R------EA-------G-I-A--------------GL----GDDQF-------RHWAA-Q-------AI--VD-A---------------------A-----WPVPLRRRA-------------------PTGD---------
AA---------------------------------
WP_045737741 ------------------------------------------MTED-PQHL---------SRS-------------ARFR--SAIA--A-FID-AR------R----EAK------------------------------------------------------------------------------L----------K-----------------G-------------------N------EG-----DS----------DT-----------------------------DA-----------------------------------K--------------------
--YD---YATW-----LS---DA-----ARRV-------G----QIQ-A----------------VTHVLKATHPD---A----------------R---G-S------SL----H-----V-----A----------P-ARL---QTH--T---E---------IGT--H--ML------G-T------D--Y--A-ED-V-VG-----N-----AA-----A--L-----DVFKLL-K-----L-----E--V--D---------GRRL------------LDWMQ-EGD-E---DLRAAL---H-----D-----NT-----KVASDWIGAFCNL--V---RH--DA-Q--
PSSHQAAKQVYWLI-------GD-----EP------QD-----DTH---YHLLQPLFSSSLAHAVHGE-----------I------F-G-------------------ES-NKQAR-Q------AY-------------R--DRRPF--DG-T-YRD------YRN-LVARKLGG-T-----KPQ----------NISQLN--SERG------------GINYLLASL-P-----PRWT--QEE-RP-RTLLKL-----DSAL-----DR-----FA---Y-F--DEV--PTLIKKLADFL-----------A-------T--------D-----------------PPK---------N--
-DA---TR--QK-----RE------GMENDLINKLALFAAETA--V------R-F-E------P---G-----WS------R------N----P-----D-----CV--------LPL---SQQLWL-----D-------------AER--VELPMRIDPQHPEWEQQDRNFVE--TYH--LGD-WQ-DEIAGQFATWLN-----DQ-----L---R------TA-------G-I-V--------------GL----GDDQY-------QHWAS-L-------TI--VD-A---------------------A-----WPVPMQRRA-------------------PAGG-------AA--
---------------------------------
WP_045770745 ------------------------------------------MTEN-PQHA---------IRS-------------ERFR--SAIA--A-FID-AR------R----EAK------------------------------------------------------------------------------L----------K-----------------G-------------------N------EG-----DS----------DT-----------------------------DA-----------------------------------K--------------------
-YD---YDTW-----LS---DA-----ARRV-------S----QIQ-A----------------VTHVLKATHPD---A----------------R---G-S------SL----H-----V-----D----------P-TRL---RAH--A---E---------VGT--H--ML------G-T------D--Y--A-ED-V-VG-----N-----AA-----A--L-----DVFKLL-K-----L-----E--V--D---------GRRL------------LDWMQ-DDD-A---DLRAAL---H-----D-----NA-----KVASDWMGAFCNL--V---RQ--DA-Q--
PSSHQAAKQVYWLV-------GD-----EP------ND-----ETH---YHLLQPLFSSSLAHAVHAD-IQ-----D--A---R--F-G-------------------ER-NKQAR-Q------AY-------------R--DKDAF--DG-T-YRD------YRN-LVARKLGG-T-----KPQ----------NISQLN--SERG------------GINYLLASL-P-----PRWT--QEE-RP-RTLLKL-----DSAL-----DR-----FA---Y-F--DEV--PKLFKKLADFL-----------A-------T--------D-----------------PPK---------
--N-----DA---TQ--KN-----RR------HIEQELVNKLVLFAAETT--V------R-F-E------P---G-----WS------R------N----P-----D-----CV--------LPL---AQQLWL-----D-------------AER--VELPIRIDPEHPEWEQQDRNFVE--TYH--LGD-WQ-DEVAGQFATWVN-----DR-----L---R------AA-------G-I-V--------------GL----GDDQY-------RHWAS-L-------TI--ID-A---------------------A-----WPVPMQRRA-------------------PAGG---------
VT---------------------------------
WP_012917520 ------------------------------------------MTED-PQHL---------TRS-------------ERFR--SAIA--A-FID-AR------R----EAK------------------------------------------------------------------------------L----------K-----------------G-------------------N------EG-----DS----------DT-----------------------------DS-----------------------------------K--------------------
--YD---YATW-----LS---DA-----ARRV-------S----QIQ-A----------------VTHVLKATHPD---A----------------R---G-S------SL----H-----V-----A----------P-TRL---QAH--T---E---------VGT--H--VL------G-T------D--Y--A-ED-V-VG-----N-----AA-----A--L-----DVFKLL-K-----L-----E--V--D---------GRRL------------LDWMQ-DDD-A---DLRVAL---H-----D-----NA-----KVASDWMGAFCNL--V---RH--DA-L--
PSSHQAAKQVYWLV-------GD-----EP------RE-----DTH---YHLLQPLFSSSLAHAVHAD-IQ-----D--A---R--F-G-------------------ER-NKQAR-Q------AY-------------R--DKHPF--DG-T-YHD------YRN-LVARKLGG-T-----KPQ----------NISQLN--SERG------------GINYLLASL-P-----PRWT--QEE-RP-RTLLKL-----DSAL-----DR-----FA---Y-F--DEV--PTLIKKLADFL-----------A-------T--------D-----------------PPK---------
-N-----DA---TQ--KK-----RE------GMENDLINKLALFAAETA--V------R-F-E------P---G-----WS------R------N----P-----D-----CV--------LPL---SQQLWL-----D-------------AER--VELPIRIDPEHAEWEQQDRNFVD--TYH--LGD-WQ-DEIAGQFATWLN-----DR-----L---R------AA-------G-I-T--------------GL----GDDQY-------RHWAK-Q-------AI--VD-A---------------------A-----WPVPMRRRA-------------------PAGG---------
VT---------------------------------
WP_045762057 ------------------------------------------MTEN-PQPA---------IRS-------------ERFR--SAIA--A-FID-AR------R----EAK------------------------------------------------------------------------------L----------K-----------------G-------------------N------EG-----DS----------DT-----------------------------DA-----------------------------------K--------------------
-YD---YDTW-----LS---DA-----ARRV-------S----QIQ-A----------------VTHVLKATHPD---A----------------R---G-S------SL----H-----V-----D----------P-TRL---RAH--A---E---------VGT--H--ML------G-T------D--Y--A-ED-V-VG-----N-----AA-----A--L-----DVFKLL-K-----L-----E--V--D---------GRRL------------LDWMQ-DDD-A---DLRAAL---H-----D-----NA-----KVASDWMGAFCNL--V---RH--DA-Q--
PSSHQAAKQVYWLV-------GD-----EP------RE-----DTH---YHLLQPLFSSSLAQVVHAE-IE-----D--A---R--F-G-------------------ER-NKQAR-Q------AY-------------R--DKDPF--DG-S-YRD------YRN-LVARKLGG-T-----KPQ----------NISQLN--SERR------------GTNYLLASL-P-----PRWT--QEE-RP-RTLLKL-----DSSL-----EY-----FN---W-F--KGV--RHLLKTLSDFL-----------L-------S--------N-----------------PPQ---------
---N-----KD---TR--ER-----RE------AIEQALGLQLSYFASTIH--A------S-F-E------P---G-----WT------R------D----P-----D-----CR--------LPT---CEQLWL-----D-------------PER--TALTDREDAEHPEWEQQDLAFKA--AYD--FGD-WP-NEVAGSFATWVN-----DR-----L---R------AA-------G-I-T--------------GL----GDDQY-------RHWAK-Q-------AI--VD-A---------------------A-----WPVPMRRRA-------------------PAGG---------
--VT---------------------------------
WP_045764807 ------------------------------------------MTEN-PQPA---------IRS-------------ERFR--SAIA--A-FID-AR------R----EAK------------------------------------------------------------------------------L----------K-----------------G-------------------N------EG-----DS----------DT-----------------------------DA-----------------------------------K--------------------
-YD---YDTW-----LS---DA-----ARRV-------S----QIQ-A----------------VTHVLKATHPD---A----------------R---G-S------SL----H-----V-----D----------P-TRL---RAH--A---E---------VGT--H--ML------G-T------D--Y--A-ED-V-VG-----N-----AA-----A--L-----DVFKLL-K-----L-----E--V--D---------GRRL------------LDWMQ-DDD-A---DLRAAL---H-----D-----NA-----KVASDWMGAFCNL--V---RH--DA-Q--
PRSHQAAKQVYWLV-------GD-----EP------RE-----DTH---YHLLQPLFSSSLAQVVHAE-IE-----D--A---R--F-G-------------------ER-NKQAR-Q------AY-------------R--DKDPF--DG-S-YRD------YRN-LVARKLGG-T-----KPQ----------NISQLN--SERR------------GTNYLLASL-P-----PRWT--QEE-RP-RTLLKL-----DSSL-----EY-----FN---W-F--KGV--RHLLKTLSDFL-----------L-------S--------N-----------------PPQ---------
---N-----KD---TR--ER-----RE------AIEQALGLQLSYFASTIH--A------S-F-E------P---G-----WT------R------D----P-----D-----CR--------LPT---CEQLWL-----D-------------PER--TALTDREDAEHPEWEQQDLAFKA--AYD--FGD-WP-NEVAGSFATWVN-----DR-----L---R------AA-------G-I-T--------------GL----GDDQY-------RHWAK-Q-------AI--VD-A---------------------A-----WPVPMRRRA-------------------PAGG---------
---VT---------------------------------
WP_045728733 ------------------------------------------MTED-PQHL---------TRS-------------ERFR--SAIA--A-FID-AR------R----EAK------------------------------------------------------------------------------L----------K-----------------S-------------------N------EG-----DS----------DT-----------------------------DA-----------------------------------K--------------------
--YD---YATW-----LS---DA-----ARRV-------G----QIQ-A----------------VTHVLKATHPD---A----------------R---G-S------SL----H-----I-----A----------P-ARL---QTH--A---E---------VGT--H--TL------G-A------D--Y--A-ED-V-VG-----N-----AA-----A--L-----DVFKLL-K-----L-----E--V--D---------GRRL------------LDWMQ-EDD-A---DLRAAL---H-----E-----NA-----KVASDWIGAFCNL--V---RH--DA-Q--
PSSHQAAKQVYWLV-------GD-----EP------KD-----DTH---YHLLQPLFSSSLAHAVHAD-IQ-----D--A---R--F-G-------------------ET-NKQAR-Q------AY-------------R--EKDPF--DG-T-YRD------YRN-LVARKLGG-T-----KPQ----------NISQLN--SERG------------GINYLLASL-P-----PHWT--QEE-RP-RSLLKL-----DSAL-----DR-----FY---R-F--EGV--RHLLKTLSDFL-----------L-------S--------N-----------------PPQ---------
--N-----QD---TR--ER-----RE------AIEQALGLHLSYFASTIH--A------R-F-E------P---G-----WT------R------D----P-----D-----CQ--------LPT---CEQLWL-----D-------------PER--TALADREDAEHPEWTQQDRAFKA--AYD--FGD-WP-DEVAGSFATWVN-----DR-----L---R------AA-------G-I-V--------------GL----GDDQY-------RHWAK-Q-------AI--VD-A---------------------A-----WPVPMRRRA-------------------PAGG---------
--AA---------------------------------
WP_045766478 ------------------------------------------MTED-PQHL---------TRS-------------ERFR--SAIA--A-FID-AR------R----EAK------------------------------------------------------------------------------L----------K-----------------G-------------------N------EG-----DS----------DT-----------------------------DS-----------------------------------K--------------------
--YD---YATW-----LS---DA-----ARRV-------G----QIQ-A----------------VTHVLKATHPD---A----------------R---G-S------SL----H-----V-----A----------P-ARL---KTH--T---E---------VGT--H--ML------G-T------D--Y--A-ED-V-VG-----N-----AA-----A--L-----DVFKLL-K-----L-----E--V--D---------GRRL------------LDWMQ-DDD-A---DLRAAL---H-----D-----NA-----KVASDWMGAFCNL--V---RH--DA-Q--
PSSHQAAKQVYWLV-------GD-----EP------KD-----DTH---YHLLQPLFSSSLAHAVHAD-IQ-----D--A---R--F-G-------------------ER-NKQAR-Q------AY-------------R--DKDAF--DG-T-YRD------YRN-LVARKLGG-T-----KPQ----------NISQLN--SERG------------GINYLLASL-P-----PRWT--QEE-RP-RTLLKL-----NSAL-----DR-----FY---R-F--EGV--RHLLKTLSDFL-----------L-------S--------N-----------------PPD---------
--N-----KD---TR--ER-----RE------AIEQALGLQLSYFASTIH--A------S-F-E------P---G-----WT------R------D----P-----D-----CQ--------LPM---CEQLWL-----D-------------PER--TALADREDAAHPEWTQQDRAFKA--AYD--FGD-WP-DEVAGSFATWVN-----DQ-----L---R------KA-------G-L-T--------------GL----GDDQY-------RHWAK-Q-------AI--ID-A---------------------A-----WPVPMRRRA-------------------PAGG---------
--AA---------------------------------
WP_045768079 ------------------------------------------MTED-PQHL---------TRS-------------ERFR--SAIA--A-FID-AR------R----EAK------------------------------------------------------------------------------L----------K-----------------G-------------------N------EG-----DS----------DT-----------------------------DS-----------------------------------K--------------------
--YD---YDTW-----LS---DA-----ARRV-------G----QIQ-A----------------VTHVLKATHPD---A----------------R---G-S------SL----H-----V-----A----------P-ARL---KTH--T---E---------VGT--H--ML------G-T------D--Y--A-ED-V-VG-----N-----AA-----A--L-----DVFKLL-K-----L-----E--V--D---------GRRL------------LDWMQ-DDD-A---DLRAAL---H-----D-----NA-----KVASDWMGAFCNL--V---RH--DA-Q--
PSSHQAAKQVYWLV-------GD-----EP------KD-----DTH---YHLLQPLFSSSLAHAVHAD-IQ-----D--A---R--F-G-------------------ER-NKQAR-Q------AY-------------R--DKDAF--DG-T-YRD------YRN-LVARKLGG-T-----KPQ----------NISQLN--SERG------------GINYLLASL-P-----PRWT--QEE-RP-RTLLKL-----NSAL-----DR-----FY---R-F--EGV--RHLLKTLSDFL-----------L-------S--------N-----------------PPD---------
--N-----KD---TR--ER-----RE------AIEQALGLQLSYFASTIH--A------S-F-E------P---G-----WT------R------D----P-----D-----CQ--------LPM---CEQLWL-----D-------------PER--TALADREDAAHPEWTQQDRAFKA--AYD--FGD-WP-DEVAGSFATWVN-----DQ-----L---R------KA-------G-L-T--------------GL----GDDQY-------RHWAK-Q-------AI--ID-A---------------------A-----WPVPMQRRA-------------------PAGG---------
--AA---------------------------------
WP_104588949 ------------------------------------------MAKP-TE----------RA-------------EVFR--AAIA--A-FIE-AR------R----QAK------------------------------------------------------------------------------L----------K-----------------D----------D----------AA------ED------AT----------GA-----------------------------AS-----------------------------------K--------------------
YD---YATW-----LA---DA-----ARRV-------G----QIQ-A----------------VTHVLKATHPD---A----------------R---G-S------SL----H-----V-----P----------P-GDL---PRH--A---E---------IGS--H--LL------G-E------D--Y--A-ED-V-VG-----N-----AA-----A--L-----DVFKFL-K-----I-----E--V--E---------GRRL------------LDWMH-AGD-S---DLRAAL---H-----T-----DA-----DVADGWMQAFGAL--V---RS--DP-Q--
PVSHAQAKQVYWLV-------GD-----QP------TD-----DTQ---YHLLQPMFSSTLAHAVHAD-IQ-----D--A---R--F-G-------------------ET-NKLAR-Q------AW-------------R--AREPH--PG-S-YRD------YRN-LVARKLGG-T-----KPQ----------NISQLN--SERG------------GVNYLLASL-P-----PTWT--Q-D-RP-RNLLKI-----GSAI-----DR-----LG---Y-F--DGV--RAMVKRLADFL-----------L-------T-------E-----------------PPS---------
---N-----QQ---TR--DT-----RE------RIEQELGAQLALFAAETH--A------R-F-P------P---G-----WT------R------D----A-----D-----CQ--------LPL---YERLWL-----D-------------PER--TELPARSDPQHPEWTQDDEAFNA--AYT--FGD-WP-DAVADHFANWVN-----AR-----L---H------AA-------G-L-T--------------SV----GDSEH-------RHWAR-Q-------AI--VD-A---------------------A-----WPVPLQRHA-------------------HAGS---------
---HP---------------------------------
WP_104602918 ------------------------------------------MTEV-SE----------RS-------------QRFR--SAIA--D-FIE-AR------R----LAK------------------------------------------------------------------------------L----------K-----------------D-------------------G-----KD-----EA----------GA-----------------------------AT-----------------------------------K--------------------
YD---YTTW-----LA---DA-----ARRV-------G----QIH-A----------------VTHVLKATHPD---A----------------R---G-S------SL----H-----V-----P----------P-DSL---PRH--A---E---------VGS--H--LL------G-S------G--Y--T-PD-V-VG-----N-----AA-----A--L-----DVFKFL-K-----V-----E--V--D---------GQGL------------LAWMQ-TGD-S---DLRAAL---H-----A-----DT-----KVATGWMQAFGTL--V---SS--EP-Q--
LSSHAMAKQVYWLV-------GE-----EP------TD-----DTH---YHLLQPMFSSTLAQVVHAD-IQ-----D--A---R--F-G-------------------ER-NKLAR-Q------AH-------------R--ARQAH--AD-D-YRD------YRN-LVARKLGG-T-----KPQ----------NISQLN--SERG------------GVNYLLASL-P-----PSWA--H-D-HP-RSLLKS-----ESAM-----ER-----FG---Y-G--KEV--RAIMKALADFL-----------R-------G-------A-----------------PPA---------
---N-----DR---TR--QT-----RD------RIAQELGAQLTVFAAQTQ--A------S-L-A------P---G-----WT------R------D----A-----D-----CM--------LPL---DEQLWL-----D-------------PER--TELPIRHDPQHGQWTQDDEAFNA--AYL--FGD-WP-DQVAGRFANWIS-----AQ-----L---R------KA-------G-L-T--------------GI----GDSEY-------RHWAK-Q-------AI--VE-A---------------------A-----WPVPLQRRA-------------------PGGA---------
---RP---------------------------------
WP_104615308 ------------------------------------------MTEP-TE----------RV-------------ERFR--SAIA--D-FIE-TR------R----LAK------------------------------------------------------------------------------L----------K-----------------D----------K-----------KE------ED------DT----------GA-----------------------------AS-----------------------------------K--------------------
YD---YATW-----LA---DA-----ARRV-------G----QIQ-A----------------VTHVLKATHPD---A----------------R---G-S------SL----H-----V-----P----------P-DSL---PQH--S---D---------IGS--H--LL------G-A------H--C--A-QD-V-VG-----N-----AA-----A--L-----DVFKFL-K-----I-----E--V--E---------GQGL------------LAWMQ-AGD-S---ALRDAL---H-----Q-----DP-----DIAVSWMQAFGAL--A---RS--EP-Q--
LRSHGMAKQVYWLV-------GT-----DP------AD-----DTH---YHLLQPMFSSALAHVVHAE-IQ-----D--A---R--F-G-------------------ET-NKLGR-Q------AL-------------R--AKQPH--VG-S-YRD------YRN-LVARKLGG-T-----KPQ----------NISQLN--SERG------------GVNYLLASL-P-----PSWT--L-E-HP-RSLLKT-----ESAM-----ER-----FG---Y-G--KEV--RALVKQLGDFL-----------L-------G-------D-----------------PTP---------
---N-----DK---TR--QT-----RE------RIDQALGAQLAVFASQTH--A------R-F-A------P---G-----WT------R------D----A-----D-----CM--------LPL---DEQLWL-----D-------------PER--TELPVRRDPQHPEWTHDDDAFNA--AYT--FGD-WP-DQVAARFANWVN-----AR-----L---H------TM-------G-L-T--------------GV----GDSEY-------RHWAK-Q-------AI--VE-A---------------------T-----WPVPLQRRA-------------------PTGG-
------KA---------------------------------
WP_116905142 ------------------------------------------MTEP-TE----------RP-------------ERFR--SAIA--E-FIQ-AR------R----LAK------------------------------------------------------------------------------L----------K-----------------E-------------------D------ED-----DA----------GV-----------------------------AS-----------------------------------K--------------------
YD---YTTW-----LA---DA-----ARRV-------G----QIQ-A----------------VTHVLKATHPD---A----------------R---G-S------SL----H-----V-----P----------P-DSL---PQH--A---E---------IGS--H--LL------G-A------H--Y--A-QD-V-VG-----N-----AA-----A--L-----DVFKFL-K-----I-----E--V--E---------GQGL------------LAWMQ-AGD-S---DLRDAL---H-----P-----DI-----DIATGWMQAFAAL--A---RS--ES-Q--
LSSHAMAKQVYWLV-------GD-----QP------SD-----DTQ---YHLLQPMFSSALAHVVHAD-IR-----D--A---R--F-G-------------------ET-NKLAR-Q------AQ-------------R--TKQPH--EG-S-YRD------YRN-LVARKLGG-T-----KPQ----------NISQLN--SKRG------------GVNYLLASL-P-----PSWT--L-E-HP-HSLLKI-----DSAM-----DR-----FG---Y-G--KEL--RAVVKALANFL-----------L-------A-------D-----------------PEA---------
---N-----DK---TR--QT-----RQ------RIEQELGAQLAVFAAETQ--A------R-F-A------P---G-----WT------R------D----A-----D-----CT--------LPL---YEQLWL-----D-------------PER--TELPVRQDLQHPQWTQDDEAFNA--AYT--FGD-WP-EQVADRFANWVN-----AQ-----L---R------KV-------G-L-T--------------TV----GDNEY-------RHWAK-Q-------AI--VE-A---------------------A-----WPVPQQRRA-------------------PGGG---------
-----QA---------------------------------
WP_064631674 ------------------------------------------MTEP-TE----------RP-------------ERFR--SAIA--E-FIQ-AR------R----LAK------------------------------------------------------------------------------L----------K-----------------E-------------------D------ED-----DA----------GV-----------------------------AS-----------------------------------K--------------------
YD---YTTW-----LA---DA-----ARRV-------G----QIQ-A----------------VTHVLKATHPD---A----------------R---G-S------SL----H-----V-----P----------P-DSL---PQH--A---E---------IGS--H--LL------G-A------H--Y--A-QD-V-VG-----N-----AA-----A--L-----DVFKFL-K-----I-----E--V--E---------GQGL------------LAWMQ-AGD-S---DLRDAL---H-----P-----DI-----DIATGWMQAFAAL--A---RS--ES-Q--
LSSHAMAKQVYWLV-------GD-----QP------SD-----DTQ---YHLLQPMFSSALAHVVHAD-IR-----D--A---R--F-G-------------------ET-NKLAR-Q------AQ-------------R--TKQPH--EG-S-YRD------YRN-LVARKLGG-T-----KPQ----------NISQLN--SKRG------------GVNYLLASL-P-----PSWT--L-E-HP-HSLLKI-----DSAM-----DR-----FG---Y-G--KEL--RAVVKALANFL-----------L-------A-------D-----------------PEA---------
---N-----DK---TR--QT-----RQ------RIEQELGAQLAVFAAETQ--A------R-F-A------P---G-----WT------R------D----A-----D-----CT--------LPL---YEQLWL-----D-------------PER--TELPVRQDLQHPQWTQDDEAFNA--AYT--FGD-WP-EQVADRFANWVN-----AQ-----L---R------KV-------G-L-T--------------TV----GDNEY-------RHWAR-Q-------AI--VE-A---------------------A-----WPVPQQRRA-------------------PGGG---------
-----QA---------------------------------
WP_002802608 ------------------------------------------MTEP-TE----------RA-------------KRFR--SAIV--D-FIE-AR------R----LAK------------------------------------------------------------------------------L----------K-----------------D-------------------G-----KG-----DA----------GA-----------------------------AS-----------------------------------K--------------------
YD---YATW-----LA---DA-----ARRV-------G----QIQ-A----------------VTHVLKATHPD---A----------------R---G-S------SL----H-----V-----P----------P-DSL---PQH--A---E---------IGS--H--LL------G-A------H--Y--A-QD-V-VG-----N-----AA-----A--L-----DVFKFL-K-----I-----E--V--E---------GQGL------------LAWMQ-AGD-S---DLRAAL---H-----P-----DT-----DVATGWMQAFAAL--A---RS--EP-Q--
LSSHGMAKQVYWLV-------GD-----QP------TD-----DTQ---YHLLQPLFSSTLAHVVHAD-IQ-----D--A---R--F-G-------------------ET-NKLAR-Q------AQ-------------R--TKQPH--DG-S-YRD------YRN-LVARKLGG-S-----NQQ----------NVSQLN--SERG------------GVNYLLASL-P-----PSWT--L-D-HP-RSLLKT-----DSVI-----DR-----FG---Y-G--KDV--RAIVKTLADFL-----------L-------D-------D-----------------PPP---------
---N-----DK---TR--RI-----RQ------RIEQELSAQLAVFAAETQ--A------R-F-A------P---G-----WT------R------D----A-----N-----CT--------LPL---YEQLWL-----D-------------PER--TVLPVRRDPQHPQWTQDDDAFNA--AYA--FGD-WP-DQVAGRFANWVN-----AQ-----L---H------KA-------G-L-T--------------SV----GDNEY-------RHWAR-Q-------AI--IE-A---------------------T-----WPVPLQRRA-------------------SAGG---------
--QA---------------------------------
WP_108817071 ------------------------------------------MTEP-TE----------RA-------------KRFR--SAIV--D-FIE-AR------R----LAK------------------------------------------------------------------------------L----------K-----------------D-------------------G-----KG-----DA----------GA-----------------------------AS-----------------------------------K--------------------
YD---YATW-----LA---DA-----ARRV-------G----QIQ-A----------------VTHVLKATHPD---A----------------R---G-S------SL----H-----V-----P----------P-DSL---PQH--A---E---------IGS--H--LL------G-A------H--Y--A-QD-V-VG-----N-----AA-----A--L-----DVFKFL-K-----I-----E--V--E---------GQGL------------LAWMQ-AGD-S---DLRAAL---H-----P-----DT-----DVATGWMQAFAAL--A---RS--EP-Q--
LSSHGMAKQVYWLV-------GD-----QP------TD-----DTQ---YHLLQPLFSSTLAHVVHAD-IQ-----D--A---R--F-G-------------------ET-NKLAR-Q------AQ-------------R--TKQPH--DG-S-YRD------YRN-LVARKLGG-S-----NQQ----------NVSQLN--SERG------------GVNYLLASL-P-----PSWT-LD---HP-RSLLKT-----DSVI-----DR-----FG---Y-G--KDV--RAIVKTLADFL----------------------------------------------
--LD--------------------------------------------------------------------------------------------D---------------PPP------------------NDKTRR--IRQ--RIE--------------------------------------------------------------------------------------------------QE------------------------------------------------
WP_014509415 ------------------------------------------MTEP-TE----------RI-------------DRFR--SAIA--D-FID-GR------R----LAK------------------------------------------------------------------------------L----------K-----------------D-------------------G-----DD-----DA----------GA-----------------------------AS-----------------------------------R--------------------
YD---YATW-----LA---DA-----ARRV-------G----QIQ-A----------------VTHVLKATHPD---A----------------R---G-S------SL----H-----V-----P----------P-DSL---PQH--A---E---------VGS--H--LL------G-A------H--Y--A-QD-V-VG-----N-----AA-----A--L-----DVFKFL-K-----I-----E--V--E---------GRGL------------LAWMQ-AGD-S---DLRTAL---H-----P-----DT-----DIATGWMQAFAAL--A---RS--EP-Q--
LSSHAMAKQVYWLV-------GD-----QP------TD-----DTH---YHLLQPLFSSALAQVVHAD-IQ-----D--A---R--F-G-------------------ET-NKQAR-Q------AQ-------------R--TKQPH--TE-T-YRD------YRN-LVARKLGG-T-----KPQ----------NISQLN--SERG------------GVNYLLASL-P-----PSWT--L-E-HP-RSLLNT-----DSAM-----DR-----FG---Y-G--KQV--RAIVKTLADFL-----------L-------G-------D-----------------PPR---------
---T-----DK---TR--RI-----RE------RIEQELGAQLAVFGAQTQ--A------R-F-A------P---G-----WT------R------D----A-----D-----CR--------LPL---DEQLWL-----D-------------PER--AELPVRRDPQDPEWTQDDDAFNA--AYT--FGD-WP-DQVAGRFANWVN-----AR-----L---H------KA-------G-L-T--------------SI----GDSEY-------RHWAK-Q-------AL--VE-A---------------------A-----WPAPLQQRA-------------------PAGD---------
---QA---------------------------------
WP_076053182 ------------------------------------------MNEV-SEHA---------VRS-------------ERFR--SAIA--D-FID-GR------R----LAK------------------------------------------------------------------------------L----------K-----------------D-------------------G-----DD-----DA----------GA-----------------------------AS-----------------------------------R--------------------
---YD---YATW-----LA---DA-----ARRV-------G----QIQ-A----------------VTHVLKATHPD---A----------------R---G-S------SL----H-----V-----P----------P-DSL---PQH--A---E---------VGS--H--LL------G-A------H--Y--A-QD-V-VG-----N-----AA-----A--L-----DVFKFL-K-----I-----E--V--E---------GRGL------------LAWMQ-AGD-S---DLRTAL---H-----P-----DT-----DIATGWMQAFAAL--A---RS--EP-Q--
LSSHAMAKQVYWLV-------GD-----QP------TD-----DTH---YHLLQPLFSSALAQVVHAD-IQ-----D--A---R--F-G-------------------ET-NKQAR-Q------AQ-------------R--TKQPH--TE-T-YRD------YRN-LVARKLGG-T-----KPQ----------NISQLN--SERG------------GVNYLLASL-P-----PSWT--L-E-HP-RSLLNT-----DSAM-----DR-----FG---Y-G--KQV--RAIVKTLADFL-----------L-------G-------D-----------------PPR---------
---T-----DK---TR--RI-----RE------RIEQELGAQLAVFGAQTQ--A------R-F-A------P---G-----WT------R------D----A-----D-----CR--------LPL---DEQLWL-----D-------------PER--AELPVRRDPQHPEWTQDDDAFNA--AYT--FGD-WP-DQVAGRFANWVN-----AR-----L---H------KA-------G-L-T--------------SV----GDSEY-------RHWAK-Q-------AL--VE-A---------------------A-----WPAPLQQRA-------------------PAGD---------
----QA---------------------------------
WP_116922741 ------------------------------------------MNEV-SEHA---------VRS-------------ERFR--SAIA--D-FID-GR------R----LAK------------------------------------------------------------------------------L----------K-----------------D-------------------G-----DD-----DA----------GA-----------------------------AS-----------------------------------R--------------------
---YD---YATW-----LA---DA-----ARRV-------G----QIQ-A----------------VTHVLKATHPD---A----------------R---G-S------SL----H-----V-----P----------P-DSL---PQH--A---E---------VGS--H--LL------G-A------H--Y--A-QD-V-VG-----N-----AA-----A--L-----DVFKFL-K-----I-----E--V--E---------GRGL------------LAWMQ-AGD-S---DLRTAL---H-----P-----DT-----DIATGWMQAFAAL--A---RS--EP-Q--
LSSHAMAKQVYWLV-------GD-----QP------TD-----DTH---YHLLQPLFSSALAQVVHAD-IQ-----D--A---R--F-G-------------------ET-NKQTR-Q------AQ-------------R--TKQPH--TE-T-YRD------YRN-LVARKLGG-T-----KPQ----------NISQLN--SERG------------GVNYLLASL-P-----PSWT--L-E-HP-RSLLNT-----DSAM-----DR-----FG---Y-G--KQV--RAIVKTLADFL-----------L-------G-------D-----------------PPR---------
---T-----DK---TR--RI-----RE------RIEQELGAQLAVFGAQTQ--A------R-F-A------P---G-----WT------R------D----A-----D-----CR--------LPL---DEQLWL-----D-------------PER--AELPVRRDPQHPEWTQDDDAFNA--AYT--FGD-WP-DQVAGRFANWVN-----AR-----L---H------KA-------G-L-T--------------SV----GDSEY-------RHWAK-Q-------AL--VE-A---------------------A-----WPAPLQQRA-------------------PAGD---------
----QA---------------------------------
WP_141483322 ------------------------------------------MNKD-PERG---------SRS-------------EQFR--MAIA--A-FIE-AR------R----ELK------------------------------------------------------------------------------L----------K-----------------G-------------------K-----ED-----DV----------DA-----------------------------VS-----------------------------------K--------------------
---YD---YGTW-----LA---DA-----VRRV-------G----QIQ-A----------------VTHVLKATHPD---A----------------R---G-S------SL----H-----V-----V----------P-DRL---PPH--A---E---------IGS--H--LL------G-E------D--F--A-ED-V-VG-----N-----AA-----A--L-----DVFKFL-K-----V-----E--I--D---------GRRL------------LDWMH-AGD-S---DLGRAL---H-----P-----DP-----AIASAWMEAFGSL--V---RS--DP-V--
PVSHGSAKQIYWLV-------GD-----QP------ED-----DAH---YHLLQPLFSSSLAHAVHAD-IQ-----D--A---R--F-G-------------------ET-NKLAR-Q------AF-------------R--AREAH--DA-D-YRD------YRN-LVARKLGG-T-----KPQ----------NISQLN--SERG------------GINYLLASL-P-----PTWD--V-E-RP-RKLLKL-----ESAM-----DR-----LA---Y-I--RGM--RHRVGDLADFL-----------L-------A-------N-----------------PPN---------
-N-----DA---TR--TR-----RR------RMEQELGGQLALFAAETR--A------R-F-E------P---G-----WT------R------D----P-----D-----CL--------LAE---CEQLWL-----D-------------PGR--AE--------------EDEDFKA--AYH--RGD-WP-DEVAGRFANWLN-----AQ-----L---H------EA-------G-L-A--------------TV----GDAEH-------AHWAR-Q-------AI--IE-A---------------------A-----WPVPTRRSA-------------------PEGA-------TA--
---------------------------------
WP_140727501 ------------------------------------------MSEE-PGRG---------SRS-------------ERFR--SAIT--D-FID-ER------R----EAK------------------------------------------------------------------------------L----------K-----------------G-------------------K-----ED-----DA----------DG-----------------------------AS-----------------------------------K--------------------
---YE---YATW-----LA---DA-----ARRV-------G----QIQ-A----------------VTHVLKATHPD---A----------------R---G-S------SL----H-----V-----A----------P-AEL---PRH--A---E---------VGS--H--LL------G-E------D--F--A-ED-V-VG-----N-----AA-----A--L-----DVFKFL-K-----I-----E--V--D---------GRRL------------LDWMQ-AGD-D---DLRSAL---H-----P-----DP-----TVADGWMEAFGGL--V---RS--DS-A--
PSSHEAAKQIYWLV-------GE-----QP------SN-----DAD---YHLLQPLFSSSLAHAVHAD-IQ-----D--A---R--F-G-------------------ET-NKLAR-Q------AF-------------R--AKEAH--GA-P-YRD------YRN-LVARKLGG-T-----KPQ----------NISQLN--SERG------------GINYLLASLPP-----PKWD--Q-D-RF-RQLLNV-----DTAF-----DR-----FA---H-F--KGV--RFLVRTLADFL-----------H-------S-------N-----------------PDQ---------
--N-----ME---TR--DR-----RE------AIEQALGFQLPVFAASIQ--A------N-S-G------P---G-----WT------R------D----P-----D-----CA--------LPL---CEQLWL-----D-------------PER--VDLPPRTGHE------QEDEEFIA--AYH--HSD-WP-DEVASRFANWIN-----AQ-----L---R------DA-------G-L-P--------------TV----GDAEY-------KHWAR-Q-------AI--ID-A---------------------A-----WPVPMQRRI-------------------PAGV-------GA---
---------------------------------
WP_090728916 ------------------------------------------MTDL-TDRG---------KAFR--SAIA--A-FIE-AR------R----EAK------------------------------------------------------------------------------L----------K-----------------G-------------------K-----DD-----DA----------GV-----------------------------AS-----------------------------------K--------------------
YD---YATW-----LA---DA-----ARRV-------G----QIQ-A----------------VTHVLKATHPD---A----------------R---G-S------SL----H-----V-----A----------P-NNL---PQR--T---E---------IGT--H--VL------G-D------D--Y--A-ED-V-VG-----N-----AA-----A--L-----DVFKFL-K-----I-----E--V--D---------DRRL------------LDWMQ-AGD-A---DLCSAL---H-----P-----DT-----TVATGWMDTFGKL--V---RS--DP-T--
PSSHEAAKQVYWLV-------GE-----EP------SN-----DVD---YHLLQPLFSSSLAHAIHAD-IQ-----D--A---R--F-G-------------------EG-NKLAR-Q------AF-------------R--TKEAY--EA-P-YRD------YRN-LIARKLGG-T-----KPQ----------NISQLN--SERG------------GINYLLASL-P-----PTWD--L-D-RP-RKLLKL-----DSAM-----DR-----LA---Y-F--KSV--RKQVKTLADFL-----------L-------A-------D-----------------PPA---------
N-----DA---TR--TL-----RR------KIEQELGNQLALFAAETH--A------R-F-E------P---G-----WT------C------D----P-----D-----CM--------LPL---CERLWL-----D-------------PER--TELPLREGHE------QEDADFKA--AYE--RGD-WP-DEVAGRFANWIN-----AQ-----L---R------EA-------G-L-P--------------TV----GDAEH-------KHWAR-Q-------AI--VD-A---------------------A-----WPVPLQRRA-------------------PAGG-------EA--
---------------------------------

WP_131301750 ----------------------------------------MTEL-TDRG------------------------KAFR--SAIA--A-FIE-AR-----R----EAK--------------------------------------------------------------------------L---------K-----------------G-------------------K-----DD----DA---------GV----------------------------AS-----------------------------------------K----------------------------------------------
YD---YGTW-----LA---DA-----AHRV-------G----QIQ-A-----------------VTHVLKATHPD---A----------------R---G-S------SL----H----V----A---------P-NSL---PQR--P---E---------IGT--H--VL------G-D-----D--Y--A-ED-V-VG----N----AA----A--L-----DVFKFL-K-----I----E--V--D---------DRRL------------LDWMQ-AGD-A---DLCSAL---H---P-----DT----TVATGWMDAFGKL--V---RS--DP-T--
PSSHEAAKQVYWLI-------GK-----EP------SN----DVD---YHLLQPLLSSSLAHTVHAD-IQ----D--A---R--F-G-----------------EA-NKLAR-Q-----AF-------------R--TKEAH--EA-P-YRD-----YRN-LVARKLGG-T-----KPQ----------NISQLN--SERG-----------GINYLLASLPP----PRWD--Q-D-RF-RNLLTL----DSAL----DR----FA--Y-F--TGV--RHQVKTLADFL----------L------T------N--------------PDR---------
N----MQ---TR--DR----RE------AIEQALSLQLPVFAASIQ--A------G-S-E------P----G-----WT-------R------D----P----D------CR---------LPL----CEQLWL------D--------------PER--TELPPREGHE------QEDADFKA--AYE--RGD-WP-DEVAGRFANWVN------AQ-----L----R-------EA---------G-L-T-------------AV---GDAEY-------KHWAR-Q--------AI--VD-A--------------------A-----WPVPLQRRA-------------------PAGG-------EA-
--------------------------
WP_136888952 ----------------------------------------MTEL-TDRG------------------------KAFH--SAIA--A-FIE-AR-----R----EAK--------------------------------------------------------------------------L---------K-----------------G-------------------K-----DD----DA---------GV----------------------------AS-----------------------------------------K----------------------------------------------
YD---YATW-----LA---DA-----ARRV-------G----QIQ-A-----------------VTHVLKATHPD---A----------------R---G-S------SL----H----V----A---------P-NSL---PQR--T---E---------IGT--H--ML------G-D-----D--Y--A-ED-V-VG----N----AA----A--L-----DVFKFL-K-----I----E--V--D---------DRRL------------LDWMQ-AGD-A---DLRTAL---H---P-----DA----AIAIGWMDAFGKL--V---RS--DP-T--
PSSHEAAKQVYWLI-------GE-----EP------SN----DVD---YHLLQPLFSSSLAHAIHAD-IQ----D--A---R--F-G-----------------ED-NKLAR-Q-----AF-------------R--TKEVH--EA-P-YRD-----YRN-LAARKLGG-T-----KPQ----------NISQLN--SERG-----------GINYLLASLPP----PKWD--Q-D-RF-RNLLTL----DSAL----DR----FA--Y-F--TGV--RHQVKTLADFL----------L------T------N--------------PDR---------
N----MQ---TR--DR----RE------AIEQTLGLQLPVFAASIQ--A------G-S-E------P----G-----WT-------R------D----P----N------CR---------LPL----CERLWL------D--------------PER--TELPPREGHE------QEDADFKA--AYE--RGD-WP-DEIAGRFANWVN------AQ-----L----R-------EA---------G-L-T-------------AV---GDAEY-------KHWAR-Q--------AI--VD-A--------------------A-----WLVPLQRRA-------------------PAGG-------EA---
--------------------------
WP_027491656 ----------------------------------------MSEL----T---------DRG-------------KAFR--TAIT--G-FIE-AR-----R----EAK--------------------------------------------------------------------------L---------K-----------------G-------------------K-----DD----DA---------DT----------------------------AS-----------------------------------------K----------------------------------------------
YD---YATW-----LA---DA-----ASRA-------H-----NLR-V-----------------VTHPIKFTHSG----I----------------K----GAS------SV-----H-----M-----G---------T-AEL---TAR--D---E---------IGT--H--SL------A-V------Q--Q--F-DF-A-IS-----D-----AK-----H--L------DVYSFL-K-----E-----V--V--D---------GKRL------------LEWLH-EDD-A---DLTNAL---D---D-----NP----AAAAALMKSFRQV--L---QP--DD-K--
LVSSTLAKQIYWLK-------GE-----DP------TD----DGQ---YHLLQPMFSSSLEQVVHED-IR----S--V---R-----------------EG-AFAAR-G-----TR-------------K--QKPTF--AD---HST-----YLD-LVARTIGG-S-----NAQ----------NVSPQN--KARG-----------GVNYLLASLPP----PAWK--P-R-DG-VNLFKL----ESVF----DDKRGAFI--L-F--EEV--RQLVRGLADFL----------K------A------N--------------PDQ---------
---N----KE---TR--ER----VT-------GFLQDIAEALSLFGADAR--A------R-G-G------A----G-----WT-------R------N----A----D------FQ---------LPP----CEQLWL------D--------------PER--TELPLRDDPEHPHWREDDVAFNQ--AYQ--RGD-WA-DEVAMRFGLWLN------GQ-----L----R-------KR---------S-D-K-------------LAML-GEAEM-------RHFAR-H--------AI--LD-V--------------------A-----WPIPLQRRA-------------------K-AG---
----AA--------------------------
WP_028840496 ----------------------------------------MSNL----T---------DRG-------------TAFR--AAIA--G-FID-AR-----R----EAK--------------------------------------------------------------------------L---------K-----------------G-------------------K-----ED----DA---------EA----------------------------AS-----------------------------------------K----------------------------------------------
YD---YAGW-----LA---DA-----ARRV-------G----QIQ-A-----------------VTHVLKATHPD---A----------------R---G-S------SL----H----V----Q---------P-SAL---SQH--A---E---------VGS--H--LL------G-T-----D--Y--A-ED-V-VG----N----AA----A--L-----DVFKLL-K-----L----E--V--D---------GKRL------------LNWMQ-AGD-A---DLAAAL---S---G-----DV----DESRRWMQAFAAL--V---HA--DT-R--
PATHPMAKQLYWLA-------GD-----TP------YD----DSH---YQLLQPMFSSALAHAVHAE-IQ----D--A---R--F-G-----------------EI-NKAAR-Q-----AR-------------R--ENKPA--ET-T-YRD-----YRD-LVVRKLGG-T-----KPQ----------NISQLN--SERG-----------GVNYLFTSLPP----RAWK--S-R-EF-TSLLKL----DSVF----DDPHGALK--R-F--GGI--RELLDTLSDFL----------R------E------N--------------PAP---------
-----V----ME---TR--QH----RE------RIELAIGQELATFGAAIR--G------S-Q-P------A----G-----WS-------R------E----D----S------CR---------LPL----CERLWL------D--------------PER--TELEPRNDPEHPHWQADDLAFNE--AYE--HGD-WA-EEVAGRFGRWLN------DE-----L----R-------KR---------S-D-K-------------LVML-GEAEM-------RHFAR-Q--------AV--LD-V--------------------A-----WPIPLRRRS-------------------K-AG---
----AA--------------------------
WP_056390880 ----------------------------------------MHTS----T---------DRK-------------LLFQ--SAIT--G-FIE-VR-----R----SAK--------------------------------------------------------------------------L---------K-----------------G-------------------K-----DD----DA---------DT----------------------------AS-----------------------------------------K----------------------------------------------
YD---YATW-----LA---DA-----ARRV-------G----QIQ-A-----------------VTHVLKATHPD---A----------------R---G-S------SL----H----I----V---------P-SHL---PQH--T---E---------IGS--H--IL------G-M-----D--Y--A-ED-V-VG----N----AA----A--L-----DVFKLL-K-----L----D--V--E---------GKRL------------LDWMQ-AGD-A---DLQEAL---S---D-----DA----EQAEAWMKAFASL--V---RE--EP-Q--
PSSHPMAKQLYWLT-------GE-----DP------QD----DAH---YHLLQPMFSSSLAHAVHAD-IQ----N--A---R--F-G-----------------EA-NKEAR-Q-----AK-------------R--DNKPA--DA-T-YHD-----YRD-LVARKLGG-T-----KPQ----------NISQLN--SERG-----------GMNYLLASLPP----PAWK--P-R-DG-VNLLKL----ESVF----DEKRGTFL--V-F--DEV--RQLVRGLADFL----------K------A------D---------------
PDQ---------N----KE---TR--DR----VM------AFLQDITEALSLFGADVR--G------R-G-G------A----G-----WT-------R------N----A----D------CH---------LLL----CEQLWL------D--------------PER--TELPERDDPEHPDWREADLAFND--AYA--RGD-WA-DEVATRFGLWLN------GQ-----L----R-------KR---------S-D-K-------------LAML-GEAEM-------RHFAS-N--------AI--LD-V--------------------A-----WPIPRQRRA---------------------
-K-AG-------AA--------------------------
WP_123769970 ----------------------------------------MTEEVITPS---------SRK-------------GRFR--SAIA--G-FIE-ER-----R----VAK--------------------------------------------------------------------------E---------K-----------------D-------------------K-----NY----E-------------------------------------PA-----------------------------------------K----------------------------------------------
YE---YNAW-----LA---DA-----VRRV-------G----QIQ-A-----------------VTHVLKATHPD---A----------------R---G-S------SL----H----V----A---------P-NAL---PQR--A---E---------IGS--H--VL------G-G-----N--F--A-ED-V-VG----N----AA----A--L-----DVFKFL-K-----I----E--V--D---------GRRL------------LDWMQ-ASD-A---DLKAAL---C---N-----DQ----DTAATWMSAFASL--V---RE--ET-E--
PASHPLAKQLYWLT-------GE-----DP------CD----DAQ---YHLLQPMFSSTLAQAVHAD-IQ----D--A---R--F-G-----------------EA-NVQAR-T-----AF-------------R--EKKPS--GT-P-YRD-----YRN-LAFRKLGG-T-----KPQ----------NISQLN--SERG-----------GLNYLLPSLPP----PAWK--P---SG-THLLKQ----ETAF----AG-----LL--R-F--GGM--RELVAELAGFL----------A------A------N--------------PDP---------
--T----AQ---TR--NR----RE------ALEQAIGQELALFGAAVR--G------Q-Y-P------A----G-----WT-------R------A----P----D------CR---------LPR----HQQLWL------D--------------PDR--TTLAVRSDPAHPEWQADDEAFNH--AYE--SGG-WA-DEVARDFGLWLN------AQ-----L----R-------KR---------S-D-K-------------LIEL-GDVEM-------RHFAR-Q--------AI--LD-V--------------------A-----WPAPMQRRA-------------------A-GR---
---AA--------------------------
WP_125165848 ----------------------------------------------MN--PDT--GRA--------ARFR--EAIS--G-FIE-ER-----R----SAK-----------------------------------------------------------------------------------EQDR------N--Y--------DP-----AK--------------------------------------------------------------------
YE---YRAW-----LA---DA-----ARRV-------G----QIQ-V-----------------VTHVLKATHPD---A----------------R---G-S------SL----H----V----P---------P-SNL---PPH--P--D---------IGS--H--VL--------------DGGF--A-ED-V-VG----N----AA----A--L-----DVFKFL-R-----T----E--V--E---------GRRL------------LDWMR-AGD-A---DLKAAL---S---D-----DP----AEAEAWTATFASL--V---RQ--GE-A--
PASHPLAKQVYWLV-------GD-----DP------RN----DGN---YHLLQPLFSSSLAHAIHAD-IQ----S--A---R--F-G-----------------EA-NVEAR-K-----AF-------------R--DNRPW--DK-P-YRE-----YRD-LAFRKLGG-T-----KPQ----------NISQLN--SERG-----------GLNYLLPSLPP----PAWK--A---RG-VRLLRH----ASAF----TD-----LI--R-F--GAM--RERVATLAEFL----------A------S------N--------------PDP---------
-T----IE---TR--NR----RE------DLEQAICLELAEFGATIR--S------Q-Q-Q------A----G-----WT-------R------D----P----Q------CE---------LPR----HQQLWL------D--------------PDR--TTLEVRTDPGHPEWQADDEAFIR--EYE--KGD-WA-DEVATDFGRWLN------QQ-----L----R-------ER---------S-P-R-------------LVAS-AEVEM-------RHFAC-Q--------AI--LD-V--------------------A-----WPTPMRRRA-------------------AVGG---
-AA--------------------------
WP_125077314 ----------------------------------------------MN--PDT--RRA--------ARFR--EAIS--G-FIE-ER-----R----SAK-----------------------------------------------------------------------------------EQDR------N--Y--------DP-----AK--------------------------------------------------------------------
YE---YRAW-----LA---DA-----ARRV-------G----QIQ-V-----------------VTHVLKATHPD---A----------------R---G-S------SL----H----V----P---------P-SNL---PPH--P--D---------IGS--H--VL--------------DGGF--A-ED-A-VG---------TA----A--L-----DVFKFL-R-----T----E--V--E---------GRRL------------LDWMR-AGD-A---DLKAAL---S---D-----DP----IEAESWMAAFASL--V---RQ--DE-A--
PASHPLAKQVYWLV-------GD-----DP------RD----DGH---YHLLQPLFSSSLAHAVHAD-IQ----S--A---W--F-G-----------------EA-NMEAR-K-----AF-------------R--ENRPS--EV-P-CRE-----YRD-LAFRKLGG-T-----NSQ----------NISQLN--SERG-----------GLNYLLPSLPP----PAWK--A---RG-VRLLRH----ASAF----TG-----LI--H-F--GAM--RERVAALAEFL----------A------S------N--------------PDP---------
---T----IE---TR--NR----RE------DLEQAICLELAEFGATIR--S------Q-Q-Q------A----G-----WT-------R------D----P----Q------CE---------LPR----HQQLWL------D--------------PDR--TTLEVRTDPGRPEWQADDEAFIR--EYE--KGD-WA-DEVATDFGRWLN------QQ-----L----R-------ER---------S-P-R-------------LVAS-AEVEM-------RHFAC-Q--------AI--LD-V--------------------A-----WPTPMRRRA-------------------AVGG---
---AA--------------------------
WP_136385918 ----------------------------------------MNEE-TESG---------ARS-------------NRFR--TTIA--A-FIE-AR-----R----EAK--------------------------------------------------------------------------L---------K-----------------G-------------------K-----DD----DA---------GA----------------------------AS-----------------------------------------K----------------------------------------------
--YE---YATW-----LA---DA-----ARRV-------G----QIQ-A-----------------VTHVLKATHPD---A----------------R---G-S------SL----H----V----D---------P-ATL---PRH--R---E---------IGS--H--LL------G-E-----D--Y--A-ED-V-VG----N----AA----A--L-----DVFKFL-K-----L----D--V--D---------GKRL------------LDWMQ-AGD-A---DLMAAL---S---P-----DS----KLAKAWMEAFASL--V---RE--GA-K--
PATHPMAKQVYWLV-------GE-----EP------QD----DSQ---YHLLQPMFSSSLAHAVHAD-IQ----D--A---R--F-G-----------------DA-NKEAR-Q-----AF-------------R--DKRPS--DA-P-YRD-----YRN-LVARKLGG-T-----KPQ----------NISQLN--SERG-----------GINYLLPSLPP----PAWQ--A-D-GR-SSVSKR----ESVL----EV-----FL--W-F--EDV--RRLTRALSDYL----------L------S------V--------------QDK---------
-----TGN--KD---IR--DA----RD------DLIKKIAEQLALFGVAIR--G------R-Y-P------A----G-----WT-------R------D----G----D------CR---------LTR----CERLWL------D--------------PER--TELPLRDDPEHPEWKAGDEAFNA--EYD--HGD-WA-DEVATRFGLWLN------AQ-----I----R-------KA---------G-L-V-------------TMAL----PEL-------NHWVS-H--------AI--VD-V--------------------A-----WPTPLRRRA-------------------KVEE---
--IA--------------------------
WP_131658731 ----------------------------------------MTEE-LHPT---------SRA-------------GRFR--QAIA--A-FIE-TR-----R----AAK--------------------------------------------------------------------------L---------K-----------------G-------------------Q-----ED----AA---------GA----------------------------AS-----------------------------------------R----------------------------------------------
--YD---YATW-----LA---DA-----ARRV-------G----QIQ-A-----------------VTHVLKATHPD---A----------------R---G-S------SL----H----V----A---------P-GGL---PAH--A---E---------IGS--H--VL------G-K-----D--Y--A-ED-V-VG----N----AA----A--L-----DVFKFL-K-----I----E--V--E---------GRRL------------LDWMQ-AGD-A---DLLAAL---S---G-----DA----AQAAGWMQAFASL--V---RE--DQ-R--
PASHPMAKQVYWLV-------GD-----DP------LD----DGG---YHLLQPMFSSTLAHAVHAE-IQ----D--A---R--F-G-----------------EA-NSAAR-Q-----AF-------------R--AGAPA--EA-V-YRD-----YRG-LVVRKLGG-T-----KPQ----------NISQLN--SERG-----------GVNYLLPSLPP----PAWA--P---RG-SNLFKR----ESAF----DE-----LL--R-F--GGV--RELVYALADFL----------K------A------N--------------PDP---------
----V----AD---TR--EQ----RD------TLAQAIADELGMFGATIR--G------G-Y-A------P----G-----WS-------R------E----A----E------CK---------LPD----CEQLWL------D--------------PGR--YDLDIRQDPQHPEWTTQDEAFRR--NYD--WGD-WP-DELATRFGLWLN------GQ-----L----R-------KR---------G-G-D-------------ALAGLNEPEM------RHWAR-Q--------AV--LD-V--------------------A-----WPVPMQRRA-------------------N-
RG-------AA--------------------------
WP_082582859 ----------------------------------------MTEE-MEPQ---------SRS-------------GRFR--TAIA--S-FIE-AR-----R----QAK--------------------------------------------------------------------------L---------K-----------------G-------------------K-----DD----DA---------EM----------------------------AS-----------------------------------------K----------------------------------------------
---YA---YATW-----LA---DA-----ARRV-------G----QIQ-A-----------------VTHVLKATHPD---A----------------R---G-S------SL----H----V----A---------P-ASL---PSR--P---E---------IGS--H--LL------G-E-----D--Y--A-ED-V-VG----N----AA----A--L-----DVFKLL-K-----V----V--D---------GRRL------------LDWMQ-AGD-A---DLRAAL---S---D-----DA----ELAEGWMSAFASL--V---RK--EA-Q--
PSTHPMAKQIYWLV-------GE-----EP------QD----DGQ---YHLLQPMFSSTLAHAVHAD-IQ----E--A---R--F-G-----------------EA-NKEAR-Q-----AF-------------R--DKKSS--RA-S-YRD-----YRN-LVARKLGG-T-----KPQ----------NISQLN--SERG-----------GVNYLLPSLPP----PAWE--Q-G-RA-SNLLKR----ESAL----EV-----LH--W-Y--GGV--RDLIRTFAKFL----------L------D------N--------------PDP---------
----T----ME---TR--ER----RE------AIAQAIGQELAMFGVAVR--A------R-Y-P------A----G-----WT-------R------D----T----A------CG---------LPL----YEQVWL------D--------------PDR--IDLPVRNDPEHQDWTTDDQAFRE--AYE--FGD-WP-DQVAGRFGLWLN------QQ-----L----R-------DK---------G-L-A-------------TMAG----PEM-------RHWAR-Q--------AI--LD-V--------------------A-----WPIPLQRRA-------------------Q-GD---
---AA--------------------------
WP_051235641 ----------------------------------------MDNS-FRQP---------HRV-------------AAFY--QAIS--E-FIE-ER-----R----DAK--------------------------------------------------------------------------L---------K-----------------G---------Q---------EG-----EA----AE---------KL----------------------------AA-----------------------------------------K----------------------------------------------
----YD---YRTW-----LA---DA-----ARRV-------S----QIQ-A-----------------VTHVLKATHPD---A----------------K---G-S------SL----H----I----P---------P-ETL---GQR--T---E---------VGS--H--LL------G-E-----A--F--A-ED-V-VG----N----AA----A--L-----DVYKLL-K-----L----E--V--D---------GKRL------------LDWLQ-EDD-P---SLLAAL---D---T-----DE----SVAQEWAEAFKAL--V---RP--PA-A--
PVSHVLAKQVYWCV-------SG-----EP------TD----NAG---FHLLQPMFSSSLAHEVHGR-IN----E--A---R--F-G-----------------ES-NKEAR-Q-----AY-------------Y--KRQEH--PV-P-YAE-----YRG-LAARKLGG-T-----KPQ----------NISQLN--SERG-----------GVNYLFASL-P----PQWR--Q-S-GE-RRLMSS----EAVM----RH-----FR--R-Y--EGV--NGLLKALIKLL----------K------N------D--------------PDS---------
-----T----ME---TR--KK----RE------EIERALGESLVAFGCTVR--H------S-R-E------P----G-----WT-------R------D----P----D------CQ---------LVE----CEQLWL------D--------------PER--ANLPPRPE--SE--QADMTFQQ--ALE--WKD-WP-QEVANRFGSWLN------NI-----L----H-------NA---------G-V-P-------------V----GDTEA-------AHWAR-Q--------AL--VD-S--------------------D-----WPSRWRRPA-------------------AQVT---
-ATQEVSNG--------------------------
WP_042120857 ----------------------------------------MNEQ-ILSS---------PRS-------------TLFR--STIA--N-FIN-ER-----K----EAK--------------------------------------------------------------------------L---------K-----------------G---------V---------EP-----DD----DA---------PT----------------------------AA-----------------------------------------K----------------------------------------------
--YD---YATW-----LA---DA-----ARRV-------A----QIQ-A-----------------VTHVLKATHPD---A----------------R---G-T------SL----F----V----D---------P-STL---VQH--K---E---------IGS--H--VL------G-D-----D--H--A-TD-I-VG----N----AA----A--L-----DVYKFL-K-----V----E--V--E---------DRRL------------LDWLR-EDD-T---DLLAAL---H---D-----DP----QLASEWAEAFKGL--I---RP--SD-T--
VSSHEMAKQVYWSV-------SN-----DP------VD----DSG---YHLLQPLFPSSLLHAVHLD-IN----D--A---R--F-G-----------------ER-NKEVR-Q-----AW-------------R--NKAQH--AD-T-YRD-----YRD-LVARKLGG-T-----KPQ----------NISQLN--SDRG-----------GVNYLLSSL-P----PAWG--S---AT-RNPLLA----IDSVL----RR-----FR-GF----DGV--NRQVTALCTLL-----------------E-----N--
DPDQCFD---TR--LQ----RE------RIEKALGRSLAAFGHGIR--N------Q-Y-S------P----G-----WT---TDSN-----------------CQ---------LPL----CEQLWL------D--------------PER--ALLPPREGCEEE------DLEFIS--AFG--RQT-WP-DEVAKRCGNWLN------EI-----L----K-------KR---------D-L-------------PV----STVEH-------AYWSR-E-----------LS--------------------R-----YVIPLGRDL-------------------ETHH------E----
-------------------------
WP_129831723 ----------------------------------------MTED-TERT---------TRS-------------ALFR--SAIT--Q-FIH-ER-----R----DAK--------------------------------------------------------------------------L---------K-----------------A---------T---------SE-----DT----NE---------TT----------------------------SS-----------------------------------------K----------------------------------------------
--YE---YGAW-----LA---DA-----ARRV-------N----QIQ-A-----------------VTHVLKATHPD---A----------------R---G-S------NL----H----A----A---------P-ETL---PAH--A---E---------IGT--H--VL------G-A-----S--Y--A-DD-I-VG----N----AA----A--L-----DVYKLL-K-----I----E--V--D---------GKRL------------LDWMQ-VDD-T---DLLTAL---H---D-----DV----AVAKTWANAFKGL--L---RP--VE-S--
WSSHEKAKQLYWCV-------KG-----EP------TD----NAS---YHLLQPMFSSTLTHAVHAE-IN----D--A---R--F-G-----------------ES-NKLAR-Q-----AW-------------R--ERKPH--EG-F-YRD-----YRQ-LLARKLGG-T-----KPQ----------NISQLN--SERG-----------GVNYLLASL-P----PSWD--L-T-TP-KTLLKL----DTAL----SR-----FA--R-Y--EGV--GDLVDALCRLL----------T------S------N--------------PGK---------
--T----ME---PR--VK----RE------FIEQALGQSLAAFGAEIS--A------R-F-D------P----G-----WT-------R------D----D----Q------CE---------LPL----CERLWL------D--------------PDR--AELPIRDE--YA---EEDRAFIA--ARE--WGD-WP-DEVAHRFGNWLN------AI-----L----L-------EK---------G-L-S-------------V----GDAEH-------AHWAR-Q--------AI--VD-A--------------------A-----WPVTMQRRA-------------------VAPT-------
KAVESTHE--------------------------
WP_144569979 ----------------------------------------M-QDAN---------QRQ-------------QLFR--STIR--G-FIQ-QR-----K----DEK--------------------------------------------------------------------------L---------K-----------------D---------S---------DD-----------F---------DK----------------------------AA-----------------------------------------K----------------------------------------------
YD---YHTW-----LA---DA-----ARRV-------T----QIQ-A-----------------VTHVLKATHPD---A----------------R---G-S------SL----Y----L----V---------P-TEL---PSH--A---E---------IGS--H--TL------G-A-----E--Y--A-DD-V-VG----N----AA----A--L-----DVYKFL-K-----L----D--V--E---------QRRL------------LDWFL-ADD-P---DLVAAL---H---P-----DS----ATAQDWSAAFKGL--I---RS--AD-A--
DYASHPMAKQVYWCI-------GD-----DP------TQ----DEH---FHLLQPLFSSSLAHAVHHD-IQ----A--A---R--F-G-----------------EA-NKEAR-Q-----AR-------------R--DNQPH--DA-P-YVE-----YRG-LAVRKLGG-T-----KPQ----------NISQLN--SERG-----------GVNYLLSSA-P----PSWA--Q-V-QP-RQLLFI----DSAL----QR-----FQ--S-Y--EEV--PSLLKQLGEFL----------T------S------N--------------PPP---------
-----T----ME---TR--TK----RE------RIEQALGQSLAAFGLAMQ--Q------A-F-P------A----G-----WT-------R------D----T----N------CK---------LPL----CEQLWL------D--------------PER--TELAPRPD--YQ---DEDIAFAQ--AYE--WKD-WP-DEVAGRFANWAN------NY-----L----S-------AK---------Y-P-L-P-------------V----GDVEY-------RHWAK-Q--------AI--VEVT--------------------E-----WPATMRRRA-------------------PCSQ---
GGAL--------------------------
WP_122469690 ----------------------------------------MDEG-SKPT---------PRS-------------LLFQ--TTIS--T-FIN-DR-----K----EAK--------------------------------------------------------------------------L---------K-----------------G---------S---------ED-----------L---------EA----------------------------AS-----------------------------------------K----------------------------------------------
YG---YAAW-----LA---DA-----ARRV-------A----QIQ-A-----------------VTHVLKATHPD---A----------------R---G-S------SL----H----V----A---------P-HTL---PPH--T---E---------IGS--H--LL------G-D-----R--Y--A-DD-I-VG----N----AA----A--L-----DVYKFL-K-----L----E--V--E---------GRRL------------LDWLQ-DND-A---DLLQAL---S---H-----DP----ATANEWASAFKSL--V---RP--AA-A--
LSSHELAKQVYWCV-------SG-----EP------AD----NSG---FHLLQPLFPSSLVHSVHQD-VN----D--A---R--F-G-----------------EI-NKAAR-Q-----AK-------------R--NDKQH--DA-P-YRD-----YRN-LVARKLGG-T-----KPQ----------NISQLN--SERG-----------GVNYLLASK-P----PQWG--Q-K-QS-PGLLRI----DTAF----DR-----FK--Y-H--GTV--PQELKALCLLI----------K------N------D--------------PPK---------
--I----KP---TR--SA----RQ------AIEQALGQSLAEFGLVNR--E------L-F-E------P----G-----WT-------R------H----P----D------CK---------LAL----CEQLWL------D--------------PGR--YDLAVRDG--HQ---EEDLAFMA--AFD--WGD-WP-AQVAQRFAQWLN------AI-----L----R-------EQ---------G-L-P-------------V----SDVEQ-------KHWAR-Q--------AI--VE-A--------------------D-----SPVPA-GLL-------------------ALAS-------
HREETRHE--------------------------
WP_122320947 ----------------------------------------MDEG-SVLT---------PRA-------------ITFY--NAIT--A-FIS-NR-----K----NEK--------------------------------------------------------------------------L---------K-----------------G---------K---------HD-----------P---------KI----------------------------AE-----------------------------------------K----------------------------------------------
YE---YTTW-----LA---NA-----ARKV-------K----QIH-A-----------------VTHTLKATHSD---A----------------R---G-S------SW----K----V----I---------P-GSL---PKH--R---E---------IGT--H--VL------G-D-----V--Y--E-ID-V-DG----N----AA----A--L-----DVYKFL-K-----E----E--V--H---------SRRL------------LDWLL-DDD-A---DLLKAL---S---F-----DK----DQANEWASAFKGL--L---AP--VN-N--
STTHTLAKQLYWSV-------SG-----KP------VD----SHG---FHLLQPMFASSLAHAVHKE-ID----D--A---R--F-G-----------------DA-NKAAR-Q-----AR-------------Q--QERTH--HN-L-YRD-----YRN-LVIRKLGG-T-----KKQ----------NISQLN--SERD-----------GVNYLLDSI-P----PTWK--R-D-RA-PNFLNI----ETAF----DR-----FR--Y-F--EGV--PEQIEALCTLL----------K------K------D--------------PPK---------I-
---LP---TR--DA----RK------SIEKALGLSLVAFGLTSR--E------Q-L-K------Q----G-----WS-------R------H----P----D------CD---------LPL----CEQLWL------D--------------PGR--IDLPLRDD--HL---EDDQAFIA--AFE--RND-WP-DEVAKRFGQWLN------AI-----L----R-------DI---------G-L-P-------------V----GDVEL-------RHWTK-Q--------AI--IE-A--------------------D-----SFMPI-------------------TGRESANG---
-------------------
WP_122317474 ----------------------------------------MDEG-SRLT---------PRA-------------KLFR--SAIA--A-FIT-ER-----K----EAK--------------------------------------------------------------------------L---------K-----------------G---------N---------DD-----DG----HD---------QA----------------------------AS-----------------------------------------K----------------------------------------------
----YD---YATW-----LA---DA-----ARRV-------A----QIQ-A-----------------VTHVLKATHPD---A----------------R---G-S------SL----H----M----V---------P-QAL---HQH--A---E---------IGT--H--AL------G-E-----A--Y--A-DD-I-VG----N----AA----A--L-----DVYKFL-K-----L----E--V--D---------GRRL------------LDWLQ-AND-A---DLLKAL---S---P-----DE----ATAHEWATAFKGL--I---RP--AA-A--
LTSHTMAKQVYWNV-------SG-----NP------TD----DSG---FHLLQPLFASSLAHAVHQD-IN----D--S---R--F-G-----------------EA-NKAAR-Q-----AK-------------R--EEKLH--DD-Q-YSD-----YRN-LVTRKLGG-T-----KPQ----------NISQLN--SERG-----------GVNYLLASI-P----PSWK--Q-D-RP-RYFLHI----ESAL----DR-----FR--R-F--EGV--DEQIKALCDLL----------G------R------D--------------PPR---------
-T----KP---TR--DA----RK------PLEQSLGASLAAFGLASR--E------L-F-E------P----G-----WT-------R------D----P----D------CE---------LAL----CEQLWL------D--------------PGR--IALPLRDD--HL---VEDQTFVT--AFH--NGD-WP-EQVAKRFGQWLN------DI-----L----R-------KT---------G-L-P-------------V----GDVEL-------KHWSR-Q--------AI--IE-A--------------------D-----SDLPI-------------------TSLEASNG---
-------------------------
WP_095067107 ----------------------------------------MDEG-SRPT---------PRS-------------TLFR--AAIT--A-FIN-ER-----K----EAK--------------------------------------------------------------------------L---------K-----------------G---------N---------DE-----GE----GG---------QA----------------------------AG-----------------------------------------K----------------------------------------------
-----YD---YATW-----LA---DA-----ARRV-------A----QIQ-A-----------------VTHVLKATHPD---A----------------R---G-S------SL----H----A----V---------P-QSL---HQH--A---E---------VGT--H--VL------G-E-----V--Y--A-DD-I-VG----N----AA----A--L-----DVYKFL-K-----V----E--V--E---------GRRL------------LDWLQ-AND-A---DLLSAL---S---D-----DE----ATAHEWSAAFKGL--V---RP--AS-T--
LTSHTMAKQVYWSV-------SG-----EP------TD----NSG---FHLLQPLVASSLAHAVHQD-IN----D--A---R--F-G-----------------EA-SKEAR-Q-----AR-------------R--EEKLY--DG-Q-YRD-----YRN-LVTRKLGG-T-----KPQ----------NISQLN--SERG-----------GVNYLLASI-P----PTWK--R-Y-RQ-PGFLNI----ETAF----DR-----FR--Y-F--EGV--PEQIEALCTLL----------K------K------D--------------PPR---------
T----LP---TR--DA----RK------SIEKALGLSLVTFGLTNR--E------Q-L-E------P----G-----WS-------R------H----P----D------CD---------LPL----CEQLWL------D--------------PGR--IGLPLRDD--YL---DEDQAFIA--AFE--RND-WP-DEVAKRFGQWLN------AI-----L----R-------NI---------G-L-P-------------V----GDVEL-------RHWTR-Q--------AI--IE-A--------------------D-----SFMPI-------------------TGRETSNG---
-------------------------
WP_117186403 ----------------------------------------MDEE-SRPT---------PRS-------------TLFR--TAIT--A-FIN-ER-----K----EAK--------------------------------------------------------------------------L---------K-----------------G---------N---------DE-----GD----GG---------QA----------------------------AG-----------------------------------------K----------------------------------------------
----YD---YATW-----LA---DA-----ARRV-------A----QIQ-A-----------------VTHVLKATHPD---A----------------R---G-S------SL----H----V----V---------P-QSR---HQH--V---E---------VGT--H--VL------G-E-----V--Y--A-DD-I-VG----N----AA----A--L-----DVYKFL-K-----V----E--V--E---------DRRL------------LDWLQ-AND-A---DLLNAL---S---D-----DE----ATAHEWASAFKGL--I---RP--AS-T--
LTSHTMAKQMYWSV-------SG-----NP------TD----NSG---FHLLQPLFASSLAHAVHQN-IN----D--A---R--F-G-----------------EA-SKEAR-Q-----AR-------------R--EEKLY--DG-Q-YRD-----YRN-LVTRKLGG-T-----KPQ----------NISQLN--SERG-----------GVNYLLASI-P----PNWK--Q-Y-RQ-LNFLNI----ETAL----DR-----FR--Y-F--EGV--PEQVEALCTLL----------K------K------D--------------PPK---------
-I----LP---TR--DA----RK------SIEKALNHSLAAFGLTSR--E------L-L-E------P----G-----WS-------R------H----P----D------CD---------LPL----CEQLWL------D--------------PGR--IDLPLRDD--HI---DEDQAFIA--AFK--RDD-WP-DQVAQRFGQWLN------AI-----L----R-------NT---------G-L-P-------------V----GDVEQ-------RHWTK-Q--------AI--IE-A--------------------D-----SFMPI-------------------TGRETSNG---
-------------------------
WP_122371790 ----------------------------------------MDEG-SRLT---------PRS-------------TLFR--TAIT--A-FIN-ER-----K----EAK--------------------------------------------------------------------------L---------K-----------------G---------N---------DE-----GD----GD---------QA----------------------------AG-----------------------------------------K----------------------------------------------
----YD---YATW-----LA---DA-----ARRV-------A----QIQ-A-----------------VTHVLKATHPD---A----------------R---G-S------SL----H----V----V---------P-QSR---HQH--V---E---------VGT--H--VL------G-E-----V--Y--A-DD-I-VG----N----AA----A--L-----DVYKFL-K-----V----E--V--E---------GRRL------------LDWLQ-ADD-V---DLLNAL---S---D-----DE----ATAHEWATAFKAL--V---RA--AS-T--
LTSHTMAKQVYWSI-------SG-----EP------TE----NSG---FHLLQPLFASSLAHAVHQD-IN----D--A---R--F-G-----------------EA-SKEAR-Q-----AR-------------R--EEKLY--DG-Q-YRD-----YRN-LVTRKLGG-T-----KPQ----------NISQLN--SERG-----------GVNYLLASI-P----PNWK--Q-Y-RQ-LNFLNI----ETAL----DR-----FR--Y-F--EGV--PEQVEALCTLL----------K------K------D--------------PPK---------
I----LP---TR--DA----RK------SIEKALNHSLAAFGLTSR--E------L-L-E------P----G-----WS-------R------H----P----D------CD---------LPL----CEQLWL------D--------------PGR--IALPLRDD--HL---DEDQAFIA--AFE--RND-WP-DEVAKRFGQWLN------AI-----L----R-------NT---------G-L-P-------------V----GDVEQ-------RHWTK-Q--------AI--IE-A--------------------D-----SFMPI-------------------TGREASNG---
-------------------------
WP_054057723 ----------------------------------------MDEV-SDQI---------SRT-------------TRFR--SAIA--A-FID-ER-----R----EAK--------------------------------------------------------------------------L---------K-----------------D---------G---------EG-----D----A---------AQ----------------------------VA-----------------------------------------K----------------------------------------------
-YE---YSAW-----LA---DA-----ARRV-------A----QIQ-A-----------------VTHVLKATHPD---A----------------R---G-S------SL----H----I----P---------P-GSL---RPH--V---E---------IGS--H--VL------G-D-----Q--Y--A-DD-I-VG----N----AA----A--L-----DVYKFL-K-----R----E--V--D---------GKRL------------LDWFL-AND-E---DLLAAL---S---S-----DE----ATAQEWAMAFKSL--V---RP--VA-N--
LASHGLAKQVYWCV-------NG-----DP------TD----DTD---FHLLQPLFASSLTHAVHQD-VQ----S--A---R--F-G-----------------EQ-NKMAR-Q-----AW-------------R--RKEAH--DG-P-YRD-----YPS-LVIRKLGG-T-----KPQ----------NISQLN--SDRG-----------GVNYLLASL-P----PIWK--Q-D-KP-RNFLNI----DSAL----ER-----FR--Y-S--EGV--WEQLQALCKLL----------E------S------D--------------PPN---------
----V----MD---TR--DE----RK------EIEQELARLLAAFGVASR--E------L-F-E------P----G-----WS-------R------D----P----A------CR---------LPL----CEQLWL------D--------------PDR--ISLPVREE--HA---EEDQAFIA--AFH--WRD-WP-DQVAQRFGLWLN------GI-----L----R-------KR---------Q-L-P-------------V----GDVEQ-------IHWAK-Q--------AI--IE-V--------------------D-----EPIPFKAPA-------------------AVSN-------
DKQEAAHG--------------------------
WP_095051625 ----------------------------------------MDEK-SDQT---------PRA-------------TLFH--TSIA--T-FIN-KR-----K----EAK--------------------------------------------------------------------------L---------K-----------------D---------T---------DA-----N----AQ------------------------------------AA-----------------------------------------T----------------------------------------------
-YD---YGTW-----LA---DA-----ARRV-------T----QIQ-A-----------------VTHVLKATHPD---A----------------R---G-S------SL----H----I----P---------P-DSL---PQH--A---E---------IGT--H--AL------G-N-----H--Y--A-DD-I-VG----N----AA----A--L-----DVYKFL-R-----L----E--V--D---------GQRL------------LYWLL-AND-A---DLLAAL---D---R-----DE----ATALEWAVAFKGL--I---RP--TV-S--
LTSHGLAKQVYWSV-------AD-----DP------TD----DAS---FHLLQPLFASSLAHAVHQD-IN----A--A---R--F-G-----------------EI-NKLAR-Q-----AK-------------R--KEEPL--DG-V-YQD-----YRN-LVLRKLGG-T-----KPQ----------NISQLN--SDRG-----------GVNYLLASL-P----PIWN--Q-D-KP-RNLLLI----DSAL----ER-----FR--Y-F--GGV--QEQLQALCTLL----------E------N------D--------------PDA---------
T----KK---TR--DQ----RK------RIEQEIARHLSAFGLVSR--E------L-M-E------P----G-----WS-------R------W----D----D------CK---------LPP----CERLWL------D--------------PGR--VGLPLRED--HL---EEDQAFVA--AFA--RKD-WP-EQVAHRFGNWLN------AI-----L----R-------QR---------Q-L-P-------------V----GDIEQ-------AYWAR-Q--------AI--IE-A--------------------E-----SPMPIKRSA-------------------TLTD-------
SDQEPTHGR--------------------------
WP_060782316 ----------------------------------------MHMDETSDQISRI---------TRFR--SAIA--T-FID-ER-----K----EAK--------------------------------------------------------------------------L-------------------KDAD-----------------SD----GA------------------------------------QA-----------------------------------------AK---------------------------------------------
--YD---YGTW-----LA---DA-----ARRV-------T----QIQ-A-----------------VTHVLKATHPD---A----------------R---G-S------SL----H----I----S---------PDSLP---QHP--E-------------IGS--H--VL------G---------E--L--Y-ADDI-VG----N----AA----A--L-----DVYKFL-K-----R----E--V--D---------GKRL------------LDWFL-LND-A---DLLAAL---H---N-----DE----ATALEWATAFKSL--I---RP--PG--
NLTSHALAKQVYWSV-------DG-----DP------TD----DGG---FHLLQPLSASSLAHAVHQD-IQ----S--A---R--F-G-----------------ET-NKLAR-Q-----AW-------------R--KREPC--DG-V-YRD-----YRL-LVLRKLGG-T-----KPQ----------NISQLN--SDRG-----------GVNYLLASL-P----PIWD--Q-D-RP-RRFLRI----ESAL----ER-----FR--H-F--AGV--REQLQALCTLL----------E------S------D--------------PER---------
---I----KE---TR--DQ----RK------QIEQDITHSLAAFGLFSR--E------L-F-E------P----G-----WS-------R------D----P----D------CK---------LPL----CEQLWL------D--------------PER--IGLPWRDD--HL---EEDQAFVA--AFA--RND-WP-DEVAKRFGNWLN------AV-----L----R-------ER---------N-L-P-------------V----GDVEQ-------THWAK-Q--------AI--IE-A--------------------Q-----SPMPFKPTV-------------------TLMD-------
RQQEVPHG--------------------------
WP_053138528 ----------------------------------------MNQG-TNPA---------IRS-------------RLFR--HTIN--A-FIQ-ER-----R----DAK--------------------------------------------------------------------------L---------K-----------------G---------A---------EA-----S----DA---------LS----------------------------AA-----------------------------------------K----------------------------------------------
--YE---YATW-----LA---DA-----ARRV-------G----QVQ-A-----------------VTHVLKATHPD---A----------------R---G-S------SL----H----V----A---------P-QTL---PRH--R---E---------VGS--H--ML------G-S-----D--Y--A-QD-I-VG----N----AA----A--L-----DVYKFL-K-----L----E--V--E---------QRRL------------LDWLL-EDD-A---DLLAAL---S---P-----DP----ALAQSWAEAFKGL--V---RP--HQ-T--
LSSHAMAKQVFWCV-------SG-----EP------GD----DAG---FELLQPMFPSSLVHAVHAD-IN----D--A---R--F-G-----------------ER-NQAAR-Q-----AR-------------R--ERQAH--DG-I-YRD-----YPQ-LVIRKLGG-T-----KPQ----------NISQLN--SERG-----------GVNYLLASL-P----PKWR--Q-Q-HP-GNLLHL----ESVF----SR-----LR--R-F--EDM--GELIEALCLLL----------Q------A------D--------------PAR---------
--S----LR---TR--AQ----RE------RIERALGRALAAFGLSLR--A------S-L-P------P----G-----WS-------R------D----R----H------CQ---------LPL----CEQLWL------D--------------PQR--AQLASADD--HQ---DADQAFAA--AFA--QRR-WP-DEIAQRFGHWLN------AI-----L----L-------QR---------G-L-P-------------V----GDTEQ-------AHWAR-Q--------AV--VD-A--------------------E-----ISCA-TREV-------------------VE-------
QEVRHG--------------------------
WP_085596349 ----------------------------------------MNQG-TNPA---------IRS-------------RLFR--HTIN--A-FIQ-ER-----R----DAK--------------------------------------------------------------------------P---------K-----------------G---------A---------EA-----S----DA---------LS----------------------------AA-----------------------------------------K----------------------------------------------
--YE---YATW-----LA---DA-----ARRV-------G----QIQ-A-----------------VTHVLKATHPD---A----------------R---G-S------SL----H----V----A---------P-QTL---PRH--R---E---------VGS--H--ML------G-S-----D--Y--A-QD-I-VG----N----AA----A--L-----DVYKFL-K-----L----E--V--E---------QRRL------------LDWLL-EDD-A---DLLAAL---S---P-----DP----ALAQSWAEAFKGL--V---RP--HQ-T--
LSSHAMAKQVFWCV-------SG-----EP------GD----DAG---FELLQPMFPSSLVHAVHAD-IN----D--A---R--F-G-----------------ER-NQAAR-Q-----AR-------------R--QRQAH--DG-I-YRD-----YPQ-LVIRKLGG-T-----KPQ----------NISQLN--SERG-----------GVNYLLASL-P----PKWR--Q-----QHPGNL-LHLESVF----SR--------L-RRFEDM--GELIEALCLLL------Q--AD----P-A-----R-
---S----LR---TR--AQ----RE------RIERALGRALAAFGLSLR--A------S-L-P------P----G-----WS-------R------D----R----H------CQ---------LPL----CEQLWL------D--------------PQR--AQLASADDHQDA------DQAFAA--AFA--QRR-WP-DEIAQRFGHWLN------AI-----L----L-------QR---------G-L-P-------------V----GDTEQ-------AHWAR-Q--------AV--VD--------------------A-----EISCATREV-------------------VEQEVRHG---
--------------------------
WP_103325043 ----------------------------------------MNQG-TNPA---------IRS-------------RLFR--HTIN--A-FIQ-ER-----R----DAK--------------------------------------------------------------------------P---------K-----------------A---------A---------EA-----S----DA---------LS----------------------------AA-----------------------------------------K----------------------------------------------
--YE---YATW-----LA---DA-----ARRV-------G----QIQ-A-----------------VTHVLKATHPD---A----------------R---G-S------SL----H----V----A---------P-QTL---PRH--R---E---------VGS--H--ML------G-S-----D--Y--A-QD-I-VG----N----AA----A--L-----DVYKFL-K-----L----E--V--E---------QRRL------------LDWLL-EDD-A---DLLAAL---S---P-----DP----ALAQSWAEAFKGL--V---RP--HQ-T--
LSSHAMAKQVFWCV-------SG-----EP------GD----DAG---FELLQPMFPSSLVHAVHAD-IN----D--A---R--F-G-----------------ER-NQAAR-Q-----AR-------------R--QRQAH--DG-I-YRD-----YPQ-LVIRKLGG-T-----KPQ----------NISQLN--SERG-----------GVNYLLASL-P----PKWR--Q-----QHPGNL-LHLESVF----SR--------L-RRFEDM--GELIEALCLLL------Q--AD----P-A-----R-
---S----LR---TR--AQ----RE------RIERALGRALAAFGLSLR--A------S-L-P------P----G-----WS-------R------D----R----H------CQ---------LPL----CEQLWL------D--------------PQR--AQLASADDHQDA------DQAFAA--AFA--QRR-WP-DEIAQRFGHWLN------AI-----L----L-------QR---------G-L-P-------------V----GDTEQ-------AHWAR-Q--------AV--VD--------------------A-----EISCATREV-------------------VEQEVRHG---
--------------------------
WP_047335692 ----------------------------------------MPEG-SNQI---------SRT-------------RTFR--DTIA--L-FIN-ER-----W----EAK--------------------------------------------------------------------------F---------K-----------------T---------A---------AI-----E----SP---------EQ----------------------------KS-----------------------------------------K----------------------------------------------
-YE---YSTW-----LA---DA-----ARRV-------T----QIQ-A-----------------VTHVLKATHPD---A----------------R---G-S------SL----H----A----A---------P-QTL---TQH--R---E---------IGT--H--LL------A-S-----D--Y--V-DD-I-VG----N----AA----A--L-----DVYKFL-K-----L----E--V--E---------GVRL------------LDWLQ-QDD-V---DLLAAL---S---P-----DP----TLAQSWAQAFKGL--I---RS--NE-T--
LISHSLAKQVFWCV-------SG-----EP------SD----DAG---FELLQPMFPSSLVHAVHAD-IN----E--R---R--F-G-----------------EA-SKTAR-L-----AR-------------R--NRQPH--TG-T-YGD-----YPQ-LVVRKLGG-T-----KPQ----------NISQLN--SERG-----------GVNYLLASL-P----PLWK--Q-H-PG-SALHL----DSAF----TR-----LR--R-F--EDV--DELIEALCTLL----------Q------A------A--------------PQS---------
T----MH---TR--LR----RE------RIERALGRALAAFGLSFR--A------S-L-P------A----G-----WS-------R------D----G----Q------CR---------LAL----CEQLWL------D--------------PQR--AQLPPREE--HM---EADQAFTA--AFA--RQA-WA-EEVAHRFGNWLN------GI-----L----L-------QR---------G-L-P-------------V----GETEQ-------AHWAR-Q--------AV--IE-A--------------------E-----TSQSWAVDE-------------------AA-------
VEVMHG--------------------------
WP_124345732 ----------------------------------------MNEG-SNQA---------ARS-------------TLFR--NTIT--T-FIN-ER-----R----DAK--------------------------------------------------------------------------L---------K-----------------G-------------------D----EA---------GD----------------------------AT-----------------------------------------K----------------------------------------------
YE---YSTW-----LA---DA-----ARRV-------A----QIQ-A-----------------VTHVLKATHPD---A----------------R---G-S------SL----H----A----A---------P-QTL---TQH--Q---E---------IGT--H--VL------A-S-----D--Y--V-ED-I-VG----N----AA----A--L-----DVYKFL-K-----L----E--V--E---------GRRL------------LDWLQ-QDD-V---DLLAAL---S---P-----DP----ILAQSWAGAFKGL--I---RS--NE-T--
LVSHSLAKQVFWCV-------SG-----EP------SD----DTG---FELLQPMFPSSLVHAVHAD-IN----E--R---R--F-G-----------------EA-NKAAR-Q-----AR-------------R--NRQPY--TG-T-YGD-----YPQ-LVVRKLGG-T-----KPQ----------NISQLN--SERG-----------GVNYLLASL-P----PLWK--Q-Q-HP-GSALHL----DSAF----TR-----LR--R-F--EDV--DELVEALCTLL----------Q------A------D--------------SPP---------
---T----MH---TR--LR----RE------HIERALGRALAAFGLSFR--A------S-L-P------A----G-----WS-------R------D----N----Q------CR---------LAL----CEQLWL------D--------------PQR--AQLPPREE--YM---EADQAFTA--AFV--RQA-WA-EEVAHRFGNWLN------GI-----L----L-------QR---------G-L-P-------------V----GDTEQ-------AHWAR-Q--------AV--IE-A--------------------E-----ASQSWVVDE-------------------TA-------
VEVMHG--------------------------
WP_136349318 ----------------------------------------MAEP-PKRT---------PRA-------------HTFQ--QTIA--A-FIA-ER-----R----DAK--------------------------------------------------------------------------L---------K-----------------G---------L---------NE-----EA----AV---------QV----------------------------AA-----------------------------------------K----------------------------------------------
---YD---YHEW-----LT---DA-----ARRV-------A----QIQ-A-----------------VTHVLKATHPD---A----------------R---G-T------NL----H----V----V---------P-DSL---PRH--V---E---------VGT--H--LL------G-S-----D--F--A-DD-I-VG----N----AA----A--L-----DVYKFL-K-----L----E--V--E---------GRRL------------LDWLQ-ADD-A---DLLRAL---H---E-----DA----ATAWQWAAAFKGL--V---RE--PV-A--
LASHGLAKQVYWSV-------SG-----EP------AD----DGG---FHLLQPLFSSSLAHTVHQE-IN----E--A---R--F-G-----------------EA-NKAAR-Q-----AR-------------R--DKKAH--DT-P-YRD-----YRN-LAVRKLGG-T-----KPQ----------NVSQLN--SERG-----------GLNYLLASL-P----PRWD--Q-D-RP-RQFLFI----DSAL----AR-----FR--R-F--EGV--GELLEALLALL----------K------S------N--------------PAP---------
--T----LD---TR--VK----RE------RIERALGQSLAAFGLASR--E------L-F-Q------P----G-----WT-------R------D----P----A------CR---------LPL----CEQIWL------D--------------PER--AELPLREDAEHR------LEDETFNA--ALE--WKD-WP-DEVAHRFGNWLN------DI-----L----R-------HH---------D-L-P-------------V----GDAEH-------AHWAR-Q--------AI--ID-A--------------------E-----WPAPMQRRA-------------------VSGD-------
RTREATHG--------------------------
WP_027351074 ----------------------------------------MPDA-EERT---------QR-------------FR--QTIA--N-FIN-ER-----R----DAK--------------------------------------------------------------------------L---------K-----------------G---------L---------NV-----EV----VA---------LA----------------------------AD-----------------------------------------K----------------------------------------------
-YD---YHAW-----LA---DA-----ARRV-------A----QIQ-A-----------------VTHVLKATHPD---A----------------R---G-S------SL----Y----I----D---------P-QAL---PSR--S---E---------IGS--H--QL------G-Q-----N--Y--E-TD-I-VG----N----AA----A--L-----DVYKFL-K-----L----D--V--E---------GRRL------------LDWLE-ADD-G---DLLAAL---H---E-----DP----GIASQWAAAFKGL--T---RP--AD-T--
LASHGLAKQVYWSV-------SG-----EP------AD----DSG---YHLLQPLFSSSLAHAVHQE-IN----D--A---R--F-G-----------------EA-SKEAR-Q-----AK-------------G--GKKPH--EG-V-YRD-----YRN-LVVRKLGG-T-----KPQ----------NISQLN--SERG-----------GLNYLLPSL-P----PRWN--Q-D-RP-RQFLVI----DSAL----ER-----FR--R-Y--EGV--NELLEALLALL----------T------S------N--------------PKP---------
-T----LD---TR--VK----RE------RIERALGLSLAAFGLASR--Q------L-F-Q------P----G-----WS-------R------D----A----D------CR---------LPL----CERIWL------D--------------PER--SELPVREEEGPR------QDDETFIA--AFE--WKD-WP-DEVAHRFGNWLN------TI-----L----L-------QH---------E-L-P-------------V----GDAEH-------AHWAR-Q--------AL--ID-A--------------------E-----WPAPMQRRA-------------------RPTD-------
QEVAHG--------------------------
WP_131188739 ----------------------------------------MPDA-EERT---------QR-------------FR--QTIA--T-FIN-ER-----R----DTK--------------------------------------------------------------------------L---------K-----------------G---------L---------DA-----EA----AT---------LT----------------------------AG-----------------------------------------K----------------------------------------------
YD---YATW-----LA---DA-----ARRV-------A----QIQ-A-----------------VTHVLKATHPD---A----------------R---G-S------SL----Y----I----D---------P-RAL---PSR--S---E---------IGS--H--LL------G-K-----H--Y--E-TD-I-VG----N----AA----A--L-----DVYKFL-K-----L----E--V--E---------GRRL------------LDWLE-ADD-G---DLLAAL---H---E-----DP----DTASQWAAAFKGL--T---RA--GE-T--
LASHSLAKQVYWCV-------SG-----EP------AD----DSG---YHLLQPLFSSSLAHAVHQE-IN----D--A---R--F-G-----------------EA-SKEVR-Q-----AR-------------R--DKKSH--EG-V-YRD-----YRN-LVVRKLGG-T-----KPQ----------NISQLN--SERG-----------GLNYLLASL-P----PRWN--Q-D-RP-RQFLFI----DSAL----QR-----FR--R-Y--EGV--NELLEALLALL----------T------S------N--------------PEP---------
-T----FD---TR--TK----RE------RIERALGQSLAAFGLASR--Q------L-F-Q------A----G-----WS-------R------D----A----D------CR---------LPL----CERIWL------D--------------PER--TDLPLREEEGVR------QEDEDFIA--AFE--WKD-WP-DEVAHRFGNWLN------AI-----L----L-------QH---------K-L-P-------------V----GDAEH-------AHWAR-Q--------AL--ID-A--------------------E-----WPAPMQRRA-------------------RPTG-------
VEVTHG--------------------------
WP_131182813 ----------------------------------------MPDA-EERT---------QR-------------FR--QTIA--T-FIN-ER-----R----DTK--------------------------------------------------------------------------L---------K-----------------G---------L---------DA-----EA----AT---------LT----------------------------AG-----------------------------------------K----------------------------------------------
YD---YATW-----LA---DA-----ARRV-------A----QIQ-A-----------------VTHVLKATHPD---A----------------R---G-S------SL----Y----I----D---------P-RAL---PSR--S---E---------IGS--H--LL------G-K-----H--Y--E-TD-I-VG----N----AA----A--L-----DVYKFL-K-----L----E--V--E---------GRRL------------LDWLE-ADD-G---DLLAAL---H---E-----DP----GTASQWAAAFKGL--T---RA--AD-T--
LASHGLAKQVYWSV-------SG-----EP------AD----DSG---YHLLQPLFSSSLAHAVHQE-IN----D--A---R--F-G-----------------EA-SKEVR-Q-----AR-------------R--DKKPN--EG-A-YRD-----YRN-LVVRKLGG-T-----KPQ----------NISQLN--SERG-----------GLNYLLASL-P----PRWN--Q-D-RP-RQFLFI----DSAL----ER-----FR--R-Y--EGV--NELLETLLALL----------T------S------N--------------PEP---------
-T----FD---TR--TK----RE------RIERALGQSLAAFGLASR--Q------L-F-Q------A----G-----WS-------R------D----A----D------CR---------LPL----CERIWL------D--------------PER--TDLPLREEEGVR------QEDEDFIA--AFE--WKD-WP-DEVAHRFGNWLN------AI-----L----L-------QH---------K-L-P-------------V----GDAEH-------AHWAR-Q--------AL--ID-A--------------------E-----WPAPMQRRA-------------------RPTG-------
VEVTHG--------------------------
WP_131190002 ----------------------------------------------MP--DAE--ERT--------QRFR--QTIA--T-FIA-ER-----R----DAK--------------------------------------------------------------------------L---------K------G-LDA--EAAT-----L--T--------------------------------AGK----------------------------------------------
YD---YATW-----LA---DA-----ARRV-------A----QIQ-A-----------------VTHVLKATHPD---A----------------R---G-S------SL----Y----I----D---------P-RAL---PSR--S---E---------IGS--H--LL------GK--------H--Y--E-TD-I-VG----N----AA----A--L-----DVYKFL-K-----L----E--V--E---------GRRL------------LDWLE-ADD-G---DLLAAL---H---E-----DP----DTASQWAAAFKGL--T---RAA--DT----
LASHGLAKQVYWCV-------SG-----EP------AD----DSG---YHLLQPLFSSSLAHGVHQE-IN----D--A---R--F-G-----------------EA-SKEAR-Q-----AR-------------R--DKKPH--EG-A-YRD-----YRN-LVVRKLGG-T-----KPQ----------NISQLN--SERG-----------GLNYLLASL-P----PRWN--Q-D-RP-RQFLFI----DSAL----QR-----FR--R-Y--EGV--NELLEALLALL----------T------S------N--------------PEP---------
--T----FD---TR--TK----RE------RIERALGQSLAAFGLASR--Q------L-F-Q------A----G-----WS-------R------D----A----D------CR---------LPL----CERIWL------D--------------PER--TDLPLREEEGVR------QEDEDFIA--AFE--WKD-WP-DEVAHRFGNWLN------AI-----L----L-------QH---------K-L-P-------------V----GDAEH-------AHWAR-Q--------AL--ID-A--------------------E-----WPAPMQRRA-------------------RPTG-------
VEVTHG--------------------------
WP_081359893 ----------------------------------------MTEG-PDQT---------SRA-------------RLFH--NAIS--A-FIH-ER-----R----EAK--------------------------------------------------------------------------L---------K-----------------G---------S---------E-----A----NQ---------VT----------------------------AD-----------------------------------------K----------------------------------------------
--YA---YTTW-----LA---DA-----ARRV-------T----QIQ-A-----------------VTHVLKAIHPD---A----------------R---G-S------NL----H----R----V---------P-QSL---PSH--S---E---------IGT--H--VL------G-N-----E--Y--V-DD-V-VG----N----AA----A--L-----DVYKFL-K-----L----E--V--E---------GRRL------------LDWLQ-DDD-S---DLLSAL---D---S-----DP----KVAHEWATSFKGL--I---RS--SG-K--
PTSHGLAKQVYWSV-------SG-----EP------AD----DTG---FHLLQPLFPSSLVHAVHTD-IN----E--A---R--F-G-----------------EN-NKAAR-L-----AK-------------R--NNQPH--IG-V-YRD-----YRN-LVVRKLGG-T-----KPQ----------NISQLN--SERG-----------GVNYLLASM-P----PQWQ--Q-G-QL-KHFLSI----SSVF----ER-----LR--R-F--EDV--NELILALCTLL----------E------S------A--------------PKP---------
T----SA---TR--LR----RE------RIEQALAQALAAFGLTNR--E------Q-L-E------P----G-----WT-------R------H----P----D------CR---------LPL----CEQLWL------D--------------PLR--AELPWRED--HL---EEDQAFAA--AIE--SGD-WP-SQVAHRFGNWLN------AI-----L----Q-------QR---------G-L-P-------------V----GDAEH-------AHWAR-Q--------AM--ID-A--------------------D-----YLLP--RPS-------------------ASLN-------
DEKDIAHG--------------------------
WP_124309473 ----------------------------------------MTEG-PDQT---------SRA-------------RLFH--NAIS--A-FIH-ER-----R----EAK--------------------------------------------------------------------------L---------K-----------------G---------S---------E-----A----NQ---------AT----------------------------AD-----------------------------------------K----------------------------------------------
--YA---YATW-----LA---DA-----ARRV-------T----QIQ-A-----------------VTHVLKAIHPD---A----------------R---G-S------NL----H----R----V---------P-QSL---PSH--S---E---------IGT--H--VL------G-N-----D--Y--V-DD-V-VG----N----AA----A--L-----DVYKFL-K-----L----E--V--E---------GRRL------------LDWLQ-DDD-S---DLLSAL---D---S-----DP----KVAHEWATSFKGL--I---RS--SG-K--
PTSHGLAKQVYWSV-------SG-----EP------AD----DTG---FHLLQPLFPSSLVHAVHTD-IN----K--A---R--F-G-----------------EN-NKAAR-L-----AK-------------R--NNQPH--IG-V-YRD-----YRN-LVVRKLGG-T-----KPQ----------NISQLN--SERG-----------GVNYLLASM-P----PQWQ--Q-G-QL-KHFLFI----SSVL----ER-----LR--R-F--EDV--NELILALCALL----------E------S------A--------------PKP---------
T----SA---TR--LR----RE------RIEQALAQALAAFGLINR--E------Q-L-E------P----G-----WT-------R------H----P----D------CR---------LPL----CEQLWL------D--------------PLR--AELPWRED--HL---EEDQAFAA--AIE--SGD-WP-SQVAHRFGNWLN------AI-----L----Q-------QR---------G-L-P-------------V----GDAEH-------AHWAR-Q--------AM--ID-A--------------------D-----YLLP--RPS-------------------ASLN-------
DEKDIAHG--------------------------
WP_092417651 ----------------------------------------MTEG-PDQT---------LRV-------------HLFH--NAIS--A-FIH-ER-----R----EAK--------------------------------------------------------------------------L---------K-----------------G---------S---------D-----A----DP---------AT----------------------------AE-----------------------------------------K----------------------------------------------
-YA---YATW-----LA---DA-----ARRV-------A----QIQ-A-----------------VTHVLKATHPD---A----------------R---G-S------SL----H----V----I---------P-QSL---PSH--S---E---------IGT--H--VL------G-Q-----D--Y--V-DD-V-VG----N----AA----A--L-----DVYKFL-K-----L----E--V--E---------GRRL------------LDWLQ-DND-S---DLLNAL---D---G-----DP----KVAHEWAMSFKGL--I---RS--SG-K--
PTSHGMAKQVYWSV-------SG-----EP------AD----DTG---FHLLQPLFPSSLVHAVHTD-IN----E--A---R--F-G-----------------EN-NKAAR-L-----AK-------------R--NNQPH--IG-I-YRD-----YRN-LVVRKLGG-T-----KPQ----------NISQLN--SERG-----------GVNYLLASM-P----PQWQ--Q-G-QL-KHFLSI----SSVF----ER-----LR--R-F--EDV--NELILALCALL----------E------S------A--------------PKP---------
T----SS---TR--LR----RE------RIEQALAQALAAFGLTHR--E------Q-F-E------P----G-----WT-------R------H----P----D------CR---------LPL----CEQLWL------D--------------PLR--AELPLRED--HL---EEDQAFAS--AIE--SGD-WP-SQVAHRFGNWLN------AI-----L----Q-------QR---------G-L-P-------------V----GDAEH-------AHWAR-Q--------AM--ID-A--------------------D-----YLLP--RHT-------------------ALLN-------
DEKDIAHG--------------------------
WP_038576094 ----------------------------------------MNNE-SDPT---------PRS-------------TLFH--SAIT--A-FIH-ER-----R----ETK--------------------------------------------------------------------------L---------K-----------------G---------G---------DA-----GA----DA---------QT----------------------------VA-----------------------------------------K----------------------------------------------
----YD---YGTW-----LA---DA-----ARRV-------S----QIQ-A-----------------VTHVLKANHPD---A----------------R---G-S------SL----H----V----T---------P-SAL---PQH--S---E---------VGT--H--VL------A-D-----D--C--A-DD-V-VG----N----AA----A--L-----DVYKFL-K-----L----E--V--E---------ACRL------------LDWLL-KDD-Q---DLLRAL---H---P-----DP----STAQEWASAFKGL--I---RP--AE-R--
WSSHGLAKQIYWSV-------SG-----EP------SD----DTG---FHLLQPLFSSSLAHAAHAE-IN----D--A---R--F-G-----------------ES-NKATR-Q-----AK-------------R--NNVPH--DE-P-YRD-----YRN-LVIRKLGG-T-----KPQ----------NISQLN--SERG-----------GVNYLLASL-P----PQWQ--Q-T-QP-GHFLSV----PSVF----ER-----FR--Q-F--EGV--NELIQGLCALL----------E------S------D--------------PPK---------
-T----LP---TR--LQ----RERLERGRRLEQGLGHALAAFGLTSR--E------R-L-E------P----G-----WS-------R------N----S----D------CQ---------LPL----CEQLWL------D--------------PLR--SELPLRED--YL---EDDRAFVI--AFE--LKD-WP-DQVAHRFGNWLN------AI-----L----Q-------QR---------G-L-P-------------V----GDVEH-------AHWAK-Q--------AL--ID-A--------------------E-----WPAPMQRRA-------------------RPAD-------
NGQETIHD--------------------------
WP_044464872 ----------------------------------------MNNE-SAPT---------QRS-------------TLFH--SAIT--A-FIH-ER-----R----EAK--------------------------------------------------------------------------L---------K-----------------G---------D---------D-----T----DA---------QT----------------------------TA-----------------------------------------K----------------------------------------------
-YD---YGTW-----LA---DA-----ARRV-------S----QIQ-A-----------------VTHVLKATHPD---A----------------R---G-S------SL----H----V----T---------P-AGL---PRH--A---E---------VGT--H--VL------A-N-----D--C--A-DD-V-VG----N----AA----A--L-----DVYKFL-K-----L----E--V--E---------ERRL------------LDWLL-QDD-Q---DLLQAL---H---P-----DP----GTAREWASAFKGL--I---RP--AE-R--
WSSHALAKQVYWSV-------SG-----EP------GD----DTG---FHLLQPLFSSSLAHAAHAQ-IN----D--A---R--F-G-----------------ES-NKAAR-Q-----AK-------------R--ANVPH--DG-P-YRD-----YRN-LVVRKLGG-T-----KPQ----------NVSQLN--SERG-----------GVNYLLASL-P----PQWQ--Q-A-QP-GPFLSE----SSVF----ER-----FR--R-F--EGV--EELIQGLCALL----------E------S------D--------------PPK---------
----T----LA---TR--LQ----RERLERGTRLEQGLGQALAAFGLASR--E------R-L-E------P----G-----WS-------R------H----R----D------CE---------LPL----CEQLWL------D--------------PRR--TELPLRDD--HQ---EQDQAFNA--AFE--FKD-WP-DQVAHRFGNWLN------AI-----L----Q-------QR---------G-L-P-------------V----GDVEH-------AHWAR-Q--------AL--ID-A--------------------E-----WPAPMQRRV-------------------RPSS---
-----NGPEPLHD--------------------------
WP_045881456 ----------------------------------------MNNE-SAPT---------QRS-------------TLFH--SAIT--A-FIH-ER-----R----EAK--------------------------------------------------------------------------L---------K-----------------G---------D---------D-----T----DA---------QT----------------------------TA-----------------------------------------K----------------------------------------------
-YD---YGTW-----LA---DA-----ARRV-------S----QIQ-A-----------------VTHVLKATHPD---A----------------R---G-S------SL----H----V----T---------P-AGL---PRH--A---E---------IGT--H--VL------A-N-----D--C--A-DD-V-VG----N----AA----A--L-----DVYKFL-K-----L----E--V--Q---------ERRL------------FDWLL-QDD-Q---DLLQAL---H---P-----DP----GTAREWAGAFKGL--I---RP--AE-R--
WSSHALAKQVYWSV-------SG-----EP------GD----DTG---FHLLQPLFSSSLAHAAHAQ-IN----D--A---R--F-G-----------------ES-NKAAR-Q-----AK-------------R--ANIPH--DG-P-YRD-----YRN-LVVRKLGG-T-----KPQ----------NISQLN--SERG-----------GVNYLLASL-P----PQWQ--Q-A-QP-GPFLSE----SSVF----ER-----FR--R-F--EGV--EELIQGLCALL----------E------S------D--------------PPK---------
---T----LA---TR--LQ----RERLERGTRLEQGLGQALAAFGLASR--E------R-L-E------P----G-----WS-------R------H----R----D------CE---------LPL----CEQLWL------D--------------PRR--TELPLRDD--HQ---EQDQAFNA--AFE--FKD-WP-DQVAHRFGNWLN------AI-----L----Q-------QR---------G-L-P-------------V----GDVEH-------AHWAR-Q--------AL--ID-A--------------------E-----WPAPMQRRA-------------------RPSS---
---NGPEALHD--------------------------
WP_036239583 ----------------------------------------MSEEKLS---------SRA-------------KAIR--KAMH--D-FIA-DR-----L----SAR--------------------------------------------------------------------------I---------E-----------------K---------L---------AI-----D----DP---------KY----QA----------------------LQ-----------------------------------------DS----------------------------------------------
--HL---PDVW-----LA---SA-----IDRS-------P----QIQ-V-----------------VTHPLKATYPHAHIT----------------K---T-T------SL----Y----C----Q---------P-KDL---PQH--N---L---------VAT--Q--AL------K-E-----N--F--T-DD-V-TG----N----AG----A--L-----DIYALL-Q-----I----V--F--E---------GKTL------------LKLCL-AED-K---DMQAAL---H---D-----DP----ETGKTWLAALADV--T---QP--KM-Q-

[illegible]

WP_053111199 [illegible]

WP_015862798 [illegible]

WP_100118022 [illegible]

WP_027706081 [illegible]

WP_119012108 [illegible]

WP_132750361 [illegible]

WP_084267996 [illegible]

WP_081837884 [illegible]

WP_101101221 [illegible]

WP_089418409 [illegible]

WP_110401628 [illegible]

WP_105749279 [illegible]

WP_105730451 [illegible]

WP_022775606 [illegible]

WP_090826348 [illegible]

WP_025042086 [illegible]

WP_090459026 [illegible]

WP_132977564 [illegible]

WP_141290479 [illegible]

WP_093192411 [illegible]

WP_036231181 [illegible]

WP_111748304 [illegible]

WP_087718523 [illegible]

WP_091643009 [illegible]

WP_045051385 [illegible]

WP_087698007 [illegible]

WP_048403876 [illegible]

WP_011135306 [illegible]

WP_081541647 [illegible]

WP_082824068 [illegible]

WP_131862017 [illegible]

WP_012696552 [illegible]

WP_088860462 [illegible]

WP_082693582 [illegible]

WP_089085022 [illegible]

WP_062790878 [illegible]

WP_107800475 [illegible]

WP_048414290 [illegible]

WP_107732823 [illegible]

WP_043591176 [illegible]

```
------N----KD---AQ--DR----RD------DLGNALLEELLLFGKEMH--E-----G-L-P-----P---G----WS------A-----E---A---R-----CR--------LPE---YQQLWL-----D-------------PWR--ADA-------------DADFAA--RWH--GED-WP-LEVAKRFGRWLN-----QQ----F---D------AS-------P-L-E-------------M---GDAEF------RHWAR-E-------LA--GD-L--GW-QAQHDQTR--R-----QLAAMAKEQ----------------A--------
-------------------------------------
WP_118268306     ------------------------------------------MPPTLT---------PRA------------TALR--TLMA--G-FIR-QR-----R----DDK----------------------------------------------------------------------L---------D----------------K---------L--------AP-----DD----DA---------KR----EA---------------------LL-----------------------------------------------AQ-----------
-----HQ---PAAW-----LA---DA-----ARRA-------G----QLQ-A----------------ATHTLKAVHPD--A----------------R---G-S------NL----Y----A----P---------P-SSL---PAR--R---E---------VGS--H--CL------G-A-----D--F--A-DD-V-VG----N----AA----A--L-----DVYKFL-K------Q----Q--H--D---------GQSL------------LALAL-AQD-A---DLAAAL---S---D----DA----EQAQAWIVAFAAL--T---DA--GA---
DAASHSRAKQVYWLV-------GY-----EP------AK----DED---YHLLAPLYPSALAQRLYQT-IQ----E--H---R--F-G----------------EA-AKAAR-Q-----AR-------------R--EQRDH--PH-G-YCD-----YPQ-LAVQKLGG-T----KPQ----------NISQLN--SERK-----------GNNYLLASL-P-----PQWR--S-L-GL-KPPLNT----EDAL-----LR----FG--R-R--NDV--RRLVRQLRVFL-----------E------S-----Q--------------PRK---
------I----MD---TR--DR----RD------ELSDALLEELLLFGKEMH--E-----G-L-P-----P---G----WS------A-----E---A---R-----CR--------LPE---YQQLWL-----D-------------PWR--ADA-------------DADFAA--RWH--GED-WP-LEVAKRFGRWLN-----QQ----F---D------AS-------P-L-E-------------M---GDVEF------RHWAR-E-------LA--GD-L--GW-QAQHDQTR--R-----QLAAMAKEQ----------------A--------
-------------------------------------
WP_081574158     ------------------------------------------MPPTLT---------PRA------------TALR--ALMA--G-FIR-QR-----R----DDK----------------------------------------------------------------------L---------D----------------K---------L--------AP-----DD----DA---------KR----EA---------------------LL-----------------------------------------------AQ-----------
-----HQ---PAAW-----LA---DA-----ARRA-------G----QLQ-A----------------ATHTLKAVHPD--A----------------R---G-S------NL----Y----A----P---------P-SSL---PAR--R---E---------VGS--H--CL------G-A-----D--F--A-DD-V-VG----N----AA----A--L-----DVYKFL-K------Q----Q--H--D---------GQSL------------LALAL-AQD-A---DLAAAL---S---D----DA----EQAQAWIAAFAAL--T---DA--GA---
DAASHSRAKQVYWLV-------GY-----EP------AK----DED---YHLLAPLYPSALAQRLYQT-IQ----E--H---R--F-G----------------EA-AKAAR-Q-----AR-------------R--EQRDH--PH-G-YCD-----YPQ-LAVQKLGG-T----KPQ----------NISQLN--SERK-----------GNNYLLASL-P-----PQWR--S-L-GL-KPPLNT----EDAL-----LR----FG--R-R--NDV--RRLVRQLRAFL-----------E------S-----Q--------------PRK---
------I----MD---TR--DR----RD------ELSDALLEELLLFGKEMH--E-----G-L-P-----P---G----WS------A-----E---A---R-----CR--------LPE---YQQLWL-----D-------------PWR--ADT-------------DADFAA--RWH--GED-WP-LDVAKRFGRWLN-----QQ----F---D------AS-------P-L-E-------------M---GDAEF------RHWAR-E-------LA--GD-L--GW-QAQHDQTR--R-----QLAAMAKEQ----------------A--------
-------------------------------------
WP_039755802     ------------------------------------------MPPTLT---------PRA------------TALR--ALIA--D-FIR-QR-----R----DDK----------------------------------------------------------------------L---------D----------------K---------L--------AP-----DD----DA---------KR----EA---------------------LL-----------------------------------------------TQ-----------
---HQ---PAAW-----LA---DA-----ARRA-------G----QLQ-A----------------ATHTLKAIHPD--A----------------R---G-S------NL----Y----A----P---------P-SSL---PAR--R---E---------VGS--H--CL------G-A-----N--F--A-DD-V-VG----N----AA----A--L-----DVYKFL-K------Q----Q--H--D---------GQSL------------LALAL-AQD-A---DLAAAL---S---D----DA----EQAQAWIAAFAAL--T---DA--GA---
DAASHSRAKQVYWLV-------GY-----EP------AK----DED---YHLLAPLYPSALAQRLYQT-IQ----E--H---R--F-G----------------EA-AKAAR-Q-----AR-------------R--EQRDH--PH-G-YCD-----YPQ-LAVQKLGG-T----KPQ----------NISQLN--SERK-----------GNNYLLASL-P-----PQWR--S-L-GL-KPPLNT----EDAL-----LR----FG--R-R--NDV--RRLVRQLRVFL-----------E------S-----Q--------------PRK---
------I----MD---TR--DR----RD------ELSDALLEELLLFGKEMH--E-----G-L-P-----P---G----WS------A-----E---A---R-----CR--------LPE---YQQLWL-----D-------------PWR--ADA-------------DADFAA--RWH--GED-WP-LEVAKRFGRWLN-----QQ----F---D------AS-------P-L-E-------------M---GDAEF------RHWAR-E-------LA--GD-L--GW-QAQHDQTR--R-----QLAAMAKEQ----------------A--------
-------------------------------------
WP_048408883     ------------------------------------------MPPTLT---------PRA------------TALR--TLMA--G-FIR-QR-----R----DDK----------------------------------------------------------------------L---------D----------------K---------L--------AP-----DD----DA---------KR----EA---------------------LL-----------------------------------------------AQ-----------
-----HQ---PAAW-----LA---DA-----ARRA-------G----QLQ-A----------------ATHTLKAIHPD--A----------------R---G-S------NL----Y----A----P---------P-STL---PPR--R---E---------VGS--H--CL------G-A-----D--F--A-DD-V-VG----N----AA----A--L-----DVYKFL-K------Q----Q--H--N---------GQSL------------LALAL-AQD-A---DLAAAL---S---D----DA----EQAQAWIAAFAAL--T---DA--GA---
AAASHSRAKQVYWLV-------GY-----EP------AK----DEH---YHLLAPLYPSALAQRLYQT-IQ----E--Q---R--F-G----------------EA-AKAAR-Q-----AR-------------R--EQRDH--PH-G-YCD-----YPQ-LAVQKLGG-T----KPQ----------NISQLN--SERK-----------GNNYLLASL-P-----PQWR--S-Q-GL-KPPLNS----EDAL-----LR----FG--R-R--NDV--RRLVRQLRAFL-----------E------S-----Q--------------PRK---
-------I----MD---TR--DR----RD------ELSDALLEELLLFGKEMH--E-----G-L-P-----P---G----WS------A-----D---T---R-----CR--------LPE---YQQLWL-----D-------------PWR--ADT-------------DADFAA--RWH--GED-WP-LEVAKRFGRWLN-----QQ----F---D------AS-------P-L-E-------------M---GDAEF------RHWAR-E-------LA--GD-L--GW-QAQHDQTR--R-----QLAAMAKEQ----------------A--------
-------------------------------------
WP_081554780     ------------------------------------------MPPTLT---------PRA------------TALR--ALMA--D-FIR-QR-----R----DDK----------------------------------------------------------------------L---------D----------------K---------L--------AP-----DD----DA---------KR----EA---------------------LL-----------------------------------------------AQ-----------
-----HQ---PAAW-----LA---DA-----ARRA-------G----QLQ-A----------------ATHTLKAIHPD--A----------------R---G-S------NL----Y----A----P---------P-STL---PPR--R---E---------VGS--H--CL------G-A-----D--F--A-DD-V-VG----N----AA----A--L-----DVYKFL-K------Q----Q--H--D---------GQSL------------LALAL-AQD-A---DLAAAL---S---D----NP----EQAQAWIAAFAAL--T---DA--GA---
AAASHSRAKQVYWLV-------GY-----EP------AK----DEH---YHLLAPLYPSALAQRLYQT-IQ----E--H---R--F-G----------------EA-AKAAR-Q-----AR-------------R--EQRDH--PH-G-YCD-----YPQ-LAVQKLGG-T----KPQ----------NISQLN--SERK-----------GNNYLLASL-P-----PQWR--S-L-GL-KPPLNS----EDAL-----LR----FG--R-R--NDV--RRLVRQLRVFL-----------E------S-----Q--------------PRK---
-------I----MD---TR--DR----RD------ELSDALLEELLLFGKEMH--E-----G-L-P-----P---G----WS------A-----D---T---R-----CR--------LPE---YQQLWL-----D-------------PWR--ADA-------------DADFAA--RWH--GED-WP-LEVAKRFGRWLN-----QQ----F---D------AS-------P-L-E-------------M---GDAEF------RHWAR-E-------LA--GD-L--GW-QAQHDQTR--R-----QLAAMAKEQ----------------A--------
-------------------------------------
WP_043640079     ------------------------------------------MPPTLT---------PRA------------TALR--ALMA--D-FIR-QR-----R----DDK----------------------------------------------------------------------L---------D----------------K---------L--------AP-----DD----DA---------KR----EA---------------------LL-----------------------------------------------AQ-----------
-----HQ---PAAW-----LA---DA-----ARRA-------G----QLQ-A----------------ATHTLKAIHPD--A----------------R---G-S------NL----Y----A----P---------P-STL---PPR--R---E---------VGS--H--CL------G-A-----D--F--A-DD-V-VG----N----AA----A--L-----DVYKFL-K------Q----Q--H--D---------GQSL------------LALAL-AQD-A---DLAAAL---S---D----NP----EQAQAWIAAFAAL--T---DA--GA---
AAASHSRAKQVYWLV-------GY-----EP------AK----DEH---YHLLAPLYPSALAQRLYQT-IQ----E--H---R--F-G----------------EA-AKAAR-Q-----AR-------------R--EQRDH--PH-G-YCD-----YPQ-LAVQKLGG-T----KPQ----------NISQLN--SERK-----------GNNYLLASL-P-----PQWR--S-L-GL-KPPLNS----EDAL-----LR----FG--R-R--NDV--RRLVRQLRAFL-----------E------S-----Q--------------PRK---
-------I----MD---TR--DR----RD------ELSDALLEELLLFGKEMH--E-----G-L-P-----P---G----WS------A-----D---T---R-----CR--------LPE---CQQLWL-----D-------------PWR--ADA-------------DADFAA--RWH--GED-WP-LEVAKRFGRWLN-----QQ----F---D------AS-------P-L-E-------------M---GDAEF------RHWAR-E-------LA--GD-L--GW-QAQHDQTR--R-----QLAAMAKEQ----------------A--------
-------------------------------------
WP_120097117     ------------------------------------------MSE---------TQA------------KALR--STIE--R-FLG-ER-----F----AAK---------------------------------------------------------------------T---------E----------------K---------L--------ST------D----EP---------QY----QK--------------------LT-------------------------------------------DQ-----------
-FH---YETW-----LT---DA-----ARRV-------A----QLQ-V----------------VTHSLKPVHPD--A----------------K---G-S------SL----Y----A----P---------P-RDD---TVS--H---T---------IGT--H--TL------D-D-----D--F--T-VD-V-VG----N----AA----A--L-----DVFKLL-K------L----E--H--D---------GRTL------------LERVE-QGD-E---QLAAAL---S---D----DP----AVAQQLMRDFATI--T---EP--SG---
QLSSHKRGKQLYWLV-------GD-----DP------SL----NDD---FHLVSPLYATSLAHRVFRT-IN----D--D---R--F-S----------------DT-AKAAR-E-----AR-------------K--KGEPA--DH-G-YVD-----YPG-LAVQRFGG-S----KPQ----------NISQLN--SERG-----------GNNYLLSCA-P-----PTWR--S-R-HV--SPIHG----SSAF----TA----FG--R-R--PDV--RRNLRALGHML-----------A------G-----D--------------PEP---
-----N----MD---TR--DR----RD------ELTRALVDELIVFTSEMH--Q-------L-E-----P---G----WT------D-----H---W---H-----CR--------LAF---AHQCWL-----D-------------PFR--AAE-------------DNEFRA--EMA--DTD-WL-TTICADFSRWLN-----GR----L----E------YF----W--K-L-A-------------M---GDPEH------RHFAQ-S-------AA--KD-Q--NV-ATLAETYR--E-----WQADLQAEM----------------AGFA-----
EVIDDDR------------------------------
WP_123631769     ------------------------------------------MRGST---------NET------------EKLR--LTIQ--E-FLS-ER-----L----AGK---------------------------------------------------------------------V---------E----------------K---------L--------AE------D----DP---------KH----EA--------------------LV-------------------------------------------AQ-----------
--FD---YEHW-----LA---DA-----ARRV-------A----QLQ-V----------------VTHSLKPIHPD--A----------------K---G-S------SV----Y----A----P---------P-GAP---RPG--A---V---------VGT--D--LL------G-D-----Q--F--T-VD-V-VG----N----AA----A--L-----DVFKFL-K------I----D--V--D---------GRTL------------LERVE-QSD-P---DLEAAL---S---D----DP----QTGRERMQQFATI--T---EP--GG---
RYASHTRGKQLFWLV-------GD-----DP------AV----NDD---FHLLAPLYATSLAHRVFQT-IN----R--D---R--F-S----------------DE-AKAAR-D-----AR-------------R--KDQPS--ET-G-YCD-----YPG-LAVQRFGG-S----KPQ----------NISQLN--SERG-----------GNSYLLGSA-P-----PLWR--S-R-AV--APIRS----SSAF----NA----YG--R-R--RNV--RATIRELGTFL-----------A------E-----D--------------PTR---
---N----MH---TR--DY----RD------QLTAELVDELIVFTSEMH--Q-------L-P-----P---G----WT------D-----D---W---Q-----CR--------LPI---EQQCWL-----D-------------PHR--AEM-------------DREFAE--TLV--ETE-WL-PEMCQDFSRWLN-----AR----L---D------RH----R--K-L-R-------------M---GDAEH------RHFAR-A-------AE--ND-T--NVRNALESARK--D-----WQAQLEKEL----------------AALA-----
EVFDDE-------------------------------
WP_123590784     ------------------------------------------MRGST---------NET------------EKLR--LTIQ--E-FLS-ER-----L----AGK---------------------------------------------------------------------V---------E----------------K---------L--------AE------D----DP---------KH----EA--------------------LV-------------------------------------------AQ-----------
--FD---YEHW-----LA---DA-----ARRV-------A----QLQ-V----------------VTHSLKPIHPD--A----------------K---G-S------SV----Y----A----P---------P-GAP---RPG--A---V---------VGT--D--LL------G-D-----Q--F--T-VD-V-VG----N----AA----A--L-----DVFKFL-K------I----D--V--D---------GRTL------------LERVE-QSD-P---DLEAAL---S---D----DP----QTGRERMQQFATI--T---EP--GG---
RYASHTRGKQLFWLV-------GD-----DP------AV----NDD---FHLLAPLYATSLAHRVFQT-IN----R--D---R--F-S----------------DE-AKAAR-D-----AR-------------R--KDQPS--ET-G-YCD-----YPG-LAVQRFGG-S----KPQ----------NISQLN--SERG-----------GNSYLLGSA-P-----PLWR--S-R-AV--APIRS----PSAF----NA----YG--R-R--RNV--RATIRELGTFL-----------A------E-----D--------------PTR---
---N----MH---TR--DY----RD------QLTAELVDELIVFTSEMR--Q-------L-P-----P---G----WT------D-----D---W---Q-----CR--------LPI---EQQCWL-----D-------------PHR--AEM-------------DREFAE--TLV--ETE-WL-PEMCQDFSRWLN-----AR----L---D------RH----R--N-L-H-------------M---GDAEH------RHFAS-A-------AG--ND-T--NIRNALESARK--D-----WQAQLEKEL----------------AALA-----
EVFDDE-------------------------------
WP_068997101     ------------------------------------------MSDTQ----------SY------------GSLR--SLIE--H-FIR-DR-----F----ETK---------------------------------------------------------------------T---------E----------------K---------L--------EP------G----SE---------AY----LK--------------------LQ-------------------------------------------AQ-----------
-FE---PQAW-----LA---DA-----ARRV-------S----QLQ-V----------------VTHSLKAIHPD--A----------------K---G-T------NV----Y----V----T---------P-QQL---PTI--P---L---------VGS--H--CL------P-D-----A--F--E-GD-V-VG----N----AA----A--L-----DVYKFL-K------L----S--W--G---------ERSL------------LALAL-QRD-A---DLLAAL---S---P----DP----AEATQWVEAFAAI--V---EP--KG---
APTSHTRAKQVYWLV-------GD-----NP------RH----DQD---YRLLAPLYSTALTHRFYEH-VS----P--N---RWFY-G----------------DD-NKAAR-H-----AL-------------R--DRREY--EG-G-YIR-----YPD-LAVQKLGG-T----KPQ----------NISQLN--SERG-----------GQNYLLASL-P-----PHWQ--S-R-EI-YPPRR------NAL----GA----FQ--R-R--PGV--GRLVQELKAFL-----------E------A-----D--------------PVK---
-------N----MH---TR--DL----RD------DLTAMLIDELMVFTLDMH--S-------L-A-----P---G----WS------A-----D---E---D-----CE--------LAD---DEQYWL-----D-------------PGR--IHE-------------DDAFAQ--ARR--QSD-WS-SAISDRAAHWLN-----RQ----L----T------YR----S--G-L-R-------------M---GDVEH------RHWKQ-Q-------ID--ET-----------LRDFR--H-----QLDELQTAL----------------DAED------
EDTD-------AGDMKAGDANGGDVNSGDVNGDQAGALS
WP_025731946     ------------------------------------------MMS---------ASV------------ASIR--QLVD--S-FIN-DR-----F----NAK---------------------------------------------------------------------V---------E----------------K---------L--------SP------D----DP---------KY----LK--------------------LQ-------------------------------------------EQ-----------
-FE---PQTW-----IA---DA-----ARRV-------S----QLQ-V----------------VTHSLKAIHPD--A----------------K---G-T------NL----Y----V----E---------P-GKL---AQL--E---Y---------VGS--H--CL------S-D-----G--F--K-GD-V-VG----N----AA----A--L-----DVYKFL-R------L----E--W--E---------GKTL------------LERSL-TRD-Q---ELIRAL---S---P----KQ----EEGETWVNAFAGI--V---ES--KG---
TPCSHAMGKQLYWLV-------DQ-----DP------TQ----DSN---YVMLAPLYPTALIHPFHIR-LS----E--A---R--F-G----------------EH-AKQGR-K-----AR-------------R--ENKYF--EG-EEYAD-----YPD-LVVQKLGG-T----KPQ----------NISQLN--SERG-----------GQNYLLASL-P-----PQWQ--S-R-SI-YPPTR------DAF----RS----FG--R-R--PIV--RRTVHELKYFL-----------E------S-----G--------------PRK------
--N---MG---TR--DW----RD------DFTTILIDELMLFTFDMH--S-------L-P-----A---G----WS------A-----D---E---K-----NQ--------LSD---TECYWL-----D-------------PGR--VEQ-------------DSDFAE--ARR--LSD-WH-QAVSDRAALWLN-----KA----L---N------RK----N--T-L-K-------------L---ADTEH------RHWKS-Q-------FE--ET-----------MKDFR--R-----QLDDLQAAV----------------FPDE-------EH------
--TEEAL------------------------
WP_087108890     ------------------------------------------MTEPPN---------ET------------LSLK--VLIH--D-FIQ-NR-----F----ATK---------------------------------------------------------------------T---------E----------------K---------L--------AA------D----EP---------AY----LK--------------------LQ-------------------------------------------EQ-----------
FA---PQTW-----LA---DA-----ARRV-------S----QLQ-V----------------VTHSLKAVHPD--A----------------K---G-T------NL----Y----V----E---------P-GSL---DRG--S---L---------IGS--H--CL------P-S-----D--F--Q-GD-V-VG----N----AA----A--L-----DVYKFL-K------L----E--W--Q---------GRSL------------LERAL-DQD-S---ELVDAL---S---K----NS----EEAQEWMCAFANI--V---EP--KG---
TPSSHARGKQVYWLV-------GD-----EP------VD----NAN---YHLLAPLYATALTHPFHAR-LS----E--A---R--F-G----------------EH-AKAGR-K-----AR-------------R--DSAAF--EG-G-YSD-----YLN-LAVQKLGG-T----KPQ----------NISQLN--SERG-----------GQNYLLASL-P-----PSWV--S-R-DV-TPPLRT-----ESAL----PH----FG--R-R--APV--RQLVNDLVSWI-----------K------Q-----
NMQEDERRALSGKAPRK-----------T-----YQ---DR--DL-----HD-----DLTAMLSDELLMFTHELH--R-------L-A-----P---G----WS------K-----D---S---E-----CR--------LVT---AECYWL-----D-------------PGR--SEL-------------DSDFAT--ARR--QSD-WH-EEVSRRVAGWLK-----NT----L---T------KR----L--N-I-S-------------L---GDPEH------EFWAN-K-------IE--LV-----------LEKFR--R-----QLEDLEDAL---
-------------RDDD-------GSL---------LGDSP------------------------
WP_027337548     ------------------------------------------MSEPQP---------PG------------SLR--ELIE--Q-FIQ-QR-----F----ATK---------------------------------------------------------------------T---------E----------------K---------M--------AQ------D----DP---------AY----LK--------------------LQ-------------------------------------------AQ-----------
--FE---PQTW-----LA---DA-----ARRV-------S----QLQ-V----------------VTHSLKAIHPD--A----------------K---G-T------NL----Y----V----E---------P-QSL---EQG--D---L---------VGS--H--CL------P-D-----D--F--Q-GD-V-VG----N----AA----A--L-----DVYKFL-K------L----E--W--K---------GRSL------------LELAL-ENV-D---TLVDAL---S---K----ER----EEAQVWAQAFANI--V---EP--KG---
APSSHARGKQVYWLV-------ND-----DP------TD----NAN---YHLLAPLYATALTHPFHAR-LS----E--A---R--F-G----------------EH-AKVGR-K-----AR-------------R--DNAAF--EG-G-YSD-----YLN-LVVQKLGG-T----KPQ----------NISQLN--SERG-----------GQNYLLASL-P-----PNWI--S-R-DV-TPPLRT-----ESVL----PR----FG--R-R--TPV--RQLINDLVNWI-----------K------Q-----
NMKEEERRVQAGKAPRK-----------T-----YQ---DR--DL-----HD-----DLTAMLADELLMFTYELH--R-------L-R-----P---G----WS------T-----D---S---E-----CR--------LVT---SECYWL-----D-------------PGR--SEL-------------DPDFAT--ARR--RSD-WH-EDISQRVAGWLK-----TT----L---S------KR----M--K-I-S-------------L---DDPAH------DFWAD-R-------IE--IV-----------LRNFR--R-----QLDDLQDAL---
------------HDDD-------ESL---------QGEPA------------------------
WP_058578641     ------------------------------------------MTEPPH---------QS------------LSLK--ELIH--D-FIQ-QR-----F----ATK---------------------------------------------------------------------T---------E----------------K---------L--------AK------D----DP---------AY----LK--------------------FQ-------------------------------------------EQ-----------
-FA---PQTW-----LA---DA-----ARRV-------S----QLQ-V----------------VTHSLKPIHPD--A----------------K---G-T------NL----Y----V----E---------P-RTL---DQG--D---L---------VGS--H--CL------P-D-----D--F--Q-GD-V-VG----N----AA----A--L-----DVYKFL-K------L----E--W--Q---------GRSL------------LELAL-END-D---TFIDAL---S---K----NP----EEAQAWVSAFASI--V---EP--KG---
APSSHTRGKQMYWLV-------GD-----DP------TD----DAS---YHLLAPLYATALTHPFHAR-LS----E--A---R--F-G----------------EH-AKAGR-K-----AR-------------R--ENVAF--EG-V-YSD-----YLN-LAVQKLGG-T----KPQ----------NISQLN--SERG-----------GQNYLLASL-P-----PHWQ--S-H-DI-YPPMR------DAF----RA----FK--R-R--PTV--WKTVHELKRFL-----------E------T-----D--------------PAK-----
----N----MH---TR--DL----RD------DFITMLVDELMLFTFDMH--S-------L-A-----P---G----WT------A-----D---D---K-----CE--------LGP---EEQYWL-----D-------------PGR--ALQ-------------DSEFDQ--ARR--RSP-WH-EAISDRAAAWLN-----RQ----L---N------YK----S--A-L-S-------------V---GDLEH------DHWQR-Q-------IE--AV-----------LKNFR--R-----QLDDLQDAL----------------RDDD-------ESF---
-----QGESA------------------------
WP_141321810     ------------------------------------------MIE--Q-FIN-DR-----F----TNK---------------------------------------------------------------------T---------E----------------K---------L--------AP------E----DP---------AY----LK--------------------LH-------------------------------------------QQ-------------------FD---
PQTW-----LA---EA-----ARRV-------S----QLQ-V----------------VTHSLKPIHPD--A----------------K---G-T------NL----Y----V----E---------P-SQL---ADN--G---L---------VGS--H--CL------L-S-----D--F--Q-GD-V-VG----N----AA----A--L-----DVYKFL-K------L----E--W--Q---------GSSL------------LELAI-ASD-D---AFIDAL---S---Q----DR----TEAESWTRAFASI--T---EP--KG---VPSSHARGKQMYWLT---
----GD-----EP------TD-----NVN---YHLLAPLYATALAQPLYAR-LS----E--D---R--F-G----------------EH-AKAAR-K-----AR-------------R--DDAAF--DG-G-YAD-----YPN-LAIQKMGG-T----KPQ----------NISQLN--SERG-----------GQNYLLASL-P-----PDWV--S-R-DV-SPPLRT-----ESVL----PR----FG--R-R--QPV--RQLVNDLARRM-----------Q------Q-----NTREDEQRQKLGKNPRK-----------T-----MQ--
-DR--DL-----HD-----DLTASLADELLVFTFEMH--R-------L-P-----A---G----WS------A-----D---D---D-----CR--------LVI---SEQYWL-----D-------------PGR--AES-------------DLEFKE--ARH--QSD-WH-VDVAQRVAHWLK-----HT----L---D------RR----I--K-T-N-------------L---GDPDH------DFWAG-Q-------IE--TV-----------MGSFR--R-----QLDDLQDAL----------------RDAS-------DD-P-------EGEAA--
---------------------
WP_019951122     ------------------------------------------MIR-------G-FIH-DR-----F----QTK---------------------------------------------------------------------T---------E----------------K---------L--------AP-----DD----------------------------------------PA---------------------------------YLKLQ-QQ----------------------------------------FE---
PQTW-----LA---DA-----AKRV-------S----QLQ-V----------------VTHSLKAIHPD--A----------------K---G-T------NL----Y----V----E---------PGQ---LER--D--AL---------VGS--H--CL------P----------ADFE-GD-V-VG----N----AA----A--L-----DVYKFL-K------L----Q--W--Q---------GRSL------------LERAL-EGD-K---ALVEAL---S---D----DR----AMAQSWVQAFANI--V---EP--KG---
APSSHARGKQLYWLV-------GD-----EP------GD----DTN---YRLLAPLYATALAHPFHAR-LS----E--A---R--F-G----------------EH-AKAGR-K-----AR-------------R--DDVGF--ED-G-YAD-----YPN-LAIQKFGG-T----KPQ----------NISQLN--SERG-----------GQNYLLPSL-P-----PHWA--S-R-EI-TAPLRT-----ESVF----PR----FG--R-R--TPV--RHLVNDLVSWM-----------R------Q-----
NLREDEKRQRQGKAPRK-----------T-----VQ---DR--NL-----HN-----DLVASLADELLLFTFELH--R-------L-P-----A---G----WS------A-----D---A---D-----CR--------LVP---AEQYWL-----D-------------PGR--AEL-------------DTEFDS--ARR--QSQ-WH-EEIAERVARWLR-----HA----I---D------RR----V--K-A-G-------------L---GDPEH------DFWTG-K-------IE--TV-----------MESFR--R-----QLDDLQDVL--
--------------RNDD-------EDQ---------EGEAA------------------------
WP_110715867     ------------------------------------------MIE--QFIL--DRFLT---------K----TEK-------------------------------------------------------------L---------------------------------------------AP-----DD--------------------------------------------PV----------------------------------YLKLQ-QQ-------------------------FD---
PQTW-----LA---DA-----ARRV-------S----QLQ-V----------------VTHSLKAIHPD--A----------------K---G-T------NL----Y----V----E------------PGQ-LKRREL---------VGS--H--CL------P----------ADFE-GD-V-VG----N----AA----A--L-----DVYKFL-K------L----T--W--E---------ERSL------------LERAL-DGD-E---ALIEAL---S---Q----DR----VEAESWVRAFADI--V---EP--KG---TPSSHARGKQIYWLV---
----GD-----NP------AS-----DED---YRLLAPLYATALTHPVHAR-LS----E--A---R--F-G----------------DH-AKAGR-K-----AR-------------R--EDAVF--QG-G-YAD-----YLN-LSVQKLGG-T----KPQ----------NISQLN--SERG-----------GQNYLLASL-P-----PQWQ--S-R-EV-TPPLRT-----ESVF----PR----FG--R-R--APV--RQVVNDLVSWM-----------R------Q-----NSKEDEKRQKEGKRLRK-----------A-----VQ-
--DR--NY-----HK------DLVASLTDELLLFTFELH--R-------L-P-----A---G----WS------T-----D---A---E-----CR--------LVL---AEQYWL-----D-------------PGR--AAQ-------------DVDFDS--NCR--QSQ-WH-EEVALRFARWLR-----HA----I---D------RR----V--T-T-G-------------L---GDPEH------DFWAK-Q-------VE--VV-----------MESLR--R-----QLDDLQDAL----------------HDDE-------E--D-------AGDLT-
---------------------
WP_110690810     ------------------------------------------MIE--Q-FIL-DR-----F----QTK---------------------------------------------------------------------T---------E----------------K---------L--------AP-----DD--------------------------------------------PV----------------------------------YLKLQ-QQ-------------------------FD---
PQAW-----LS---DA-----ARRV-------S----QLQ-V----------------VTHSLKAIHPD--A----------------K---G-T------NL----Y----V----E---------P-------AQL--KRREL---------VGS--H--CL------P----------ADFE-GD-V-VG----N----AA----A--L-----DVYKFL-K------L----R--W--E---------ERSL------------LERAL-DGD-E---ALIEAL---S---P----DR----AEAESWVRAFADI--V---EP--KG---TPSSHARGKQVYWLV--
-----GD-----DP------AS-----DTG---YRLLAPLYATALIHPVHSR-LS----E--A---R--F-G----------------EH-AKAGR-K-----AR-------------R--EDAIF--QG-G-YAD-----YLN-LSVQKLGG-T----KPQ----------NISQLN--SERG-----------GQNYLLASL-P-----PQWQ--S-R-EV-TTPLRT-----ESVF----PR----FG--R-R--APV--RQVVNDLVSWM-----------R------Q-----NSKEDEKRQKEGKRPRK-----------T-----VH--
-DR--NY-----HK------DLVASLTDELLLFTFELH--R-------L-P-----A---G----WS------R-----D---A---E-----CR--------LVL---AEQYWL-----D-------------PGR--AAL-------------EVDFDS--NRR--QSQ-WH-EEVALRFARWLR-----HA----I---D------RR----V--T-T-G-------------L---GDPEH------DFWAK-Q-------VE--VV-----------MESLR--R-----QLDDLQDAL----------------RDDE-------K--D-------AGELT---
---------------------
WP_136065513     ------------------------------------------MSEPTN---------SPP------------PTLE--AVIR--G-FIY-ER-----F----LTK---------------------------------------------------------------------T---------E----------------K---------L--------AP------D----DP---------AY----LK--------------------LH-------------------------------------------QQ-----------
--FE---PQTW-----LA---DA-----ARRV-------S----QLQ-V----------------VTHSLKAMHPD--A----------------K---G-T------NL----Y----V----E---------P-SQL---ERN--E---L---------VGS--H--CL------P-A-----D--F--E-GD-V-VG----N----AA----A--L-----DVYKFL-K------L----K--W--E---------ERSL------------LERAL-DGD-E---ALIEAL---S---Q----DR----AEAESWVRAFADI--V---EP--KG---
TPSSHARGKQVYWLV-------GD-----DP------AS----DKD---YRLLAPLYATALTHPVHAR-LS----E--A---R--F-G----------------DH-AKAGR-K-----AR-------------R--EDAAF--QG-G-YAD-----YLN-LSVQKLGG-T----KPQ----------NISQLN--SERG-----------GQNYLLASL-P-----PNWQ--S-R-EV-YAPMR------DPF----KA----FK--G-R--PEV--RRTLSDLKRFL-----------A------S-----D--------------PTA----
-----N----RQ---TR--DL----RD------DLTAMLVDELMLFTFDMH--S-------L-T-----P---G----WT------T-----E---A---R-----CE--------LAS---EEQYWL-----D-------------PGR--AEQ-------------DAEFAE--ARR--QSN-WH-DEISARAALWLN-----RE----L---G------QK----S--P-L-Q-------------F---GDIEH------GHWKS-R-------FD--DV-----------LKRFR--R-----QLDDLQDAL----------------RDDE-------E--D-
-----AGDLT------------------------
WP_086623391     ------------------------------------------MSEPVDSPP-------PSLK--ALLH--G-FIR-DR-----F----QTK---------------------------------------------------------------------T---------E----------------K---------L--------AP-----DD--------------------------------------------PA---------------------------------YLKLQ-QQ-----------------
----FE---PQAW-----LA---DA-----ARRV-------S----QLQ-V----------------VTHSLKAIHPD--A----------------K---G-T------NL----Y----V----E------------PGQ-LERSAL---------VGS--H--CL------P----------VNFD-GD-V-VG----N----AA----A--L-----DVYKFL-K------L----Q--W--E---------GRSL------------LERAL-EGD-K---ALIEAL---S---D----DC----DEAQSWVQAFADI--V---EP--KG---
TPSSHARGKQLYWLV-------GD-----DP------CE----DAN---YRLLSPLYATALAHPFYAQ-LS----E--A---R--F-G----------------ER-AKAGR-K-----AR-------------R--DNAVF--EG-G-YAD-----YLN-LAVQKFGG-T----KPQ----------NISQLN--SERG-----------GQNYLLASL-P-----PHWQ--S-R-DV-YPPMR------DAF----QT----FG--R-R--QNV--RRTLDDLKRFL-----------E------S-----D--------------PAA---
------N----MH---TR--DL----RD------DLTAMLVDELMLFTFDIH--S-------I-Q-----A---G----WT------T-----D---E---R-----CQ--------LAP---EEQYWL-----D-------------PGR--AEQ-------------DVEFAE--ARR--KTE-WH-EDIRQRAAVWLN-----RA----L---N------RK----S--S-L-N-------------L---GDVEH------EHWKN-R-------FD--GV-----------LKSFR--R-----QLDDLQDVL----------------RDDD-------E--A-
-----EGDAA------------------------
WP_110670828     ------------------------------------------MSESIDSPP-------PTLK--ALVH--G-FIH-DR-----F----QTK---------------------------------------------------------------------T---------E----------------K---------L--------TP-----DD--------------------------------------------PV----------------------------------YLKLQ-QQ----------------
---FE---PQTW-----LA---DA-----ARRV-------S----QLQ-V----------------VTHSLKAIHPD--A----------------K---G-T------NL----Y----V----E------------PGQ---LKR--D--AL---------VGS--H--CL------------PSDF--E-GD-V-VG----N----AA----A--L-----DVYKFL-K------L----Q--W--E---------GRSL------------LERAL-EGD-D---ALVEAL---S---D----DR----AEAEIWVQAFADI--V---EP--NG---
TPSSHARGKQLYWLV-------GD-----EP------GD----DAN---YRLLAPLYATALAHRVYAQ-VS----L--N---TWFY-G----------------DH-NKAAR-K-----AL-------------K--EKTPF--EG-G-YTS-----YLN-LAIQKFGG-T----KPQ----------NISQLN--SERG-----------GQNYLLASL-P-----PHWQ--S-R-DV-YPPMR------DAF----QT----FG--R-R--QEV--RRTLDDLKRFL-----------E------S-----D--------------PAA---
------N----RH---TR--DL----RD------DLTAMLVDELMLFTFDIH--S-------V-P-----A---G----WT------T-----D---E---R-----CQ--------LAV---EEQYWL-----D-------------PGR--AEL-------------DAEFAE--ARR--QSD-WH-VAVSARAAQWLN-----RA----L---N------RD----S--P-L-N-------------L---GDVEH------EHWKG-R-------FD--GV-----------LKSFR--R-----QLDDLQDAL----------------RDDD-------E--
D-------EGAAA------------------------
WP_115853811     ------------------------------------------MSEHINSPP-------------PSLE--ALIH--G-FMH-DR-----F----QTK---------------------------------------------------------------------T-EK----------------L--------AP-----DD--------------------------------------------PV----------------------------------YLKLQ-QQ----------------
----FE---PQTW-----LA---DA-----AHRV-------S----QLQ-V----------------VTHSLKAIHPD--A----------------K---G-T------NL----Y----V----E------------------PGQLKR-NEL---------VGS--H--CL------P------AD--F--D-GD-V-VG----N----AA----A--L-----DVYKFL-K------L----Q--W--E---------GRSL------------LERAL-EGD-Q---ALIEAL---S---D----DR----AEAQSWVQAFANI--V---EP--KGT---
PSSHARGKQLYWLV-------GD-----EP------GD----NAN---YHLLAPLYATALAHPFHAR-VS----E--A---R--F-G----------------EH-AKTGR-K-----AR-------------R--DNAAF--EG-G-YAD-----YLN-LAVQKFGG-T----KPQ----------NISQLN--SERG-----------GQNYLLASL-P-----PHWQ--S-R-DV-YPPMR------DAL----QT----FG--R-R--QEV--RRTLDDLKRFL-----------E------S-----D--------------PAA----
----N----RH---TR--DR----RD------DLTAMLVDELMLFIFDIH--S-------V-P-----A---G----WT------T-----D---E---R-----CQ--------LAA---EEQYWL-----D-------------PRR--AEL-------------DAEFAE--ARR--QSD-WH-DKISARAAQWLN-----RT----L---N------RD----S--P-L-N-------------L---GDVEH------EHWKG-R-------FD--SV-----------LKSFR--R-----QLDDLQDTL----------------RDDD-------E--D---
---EGAAA------------------------
WP_112056239     ------------------------------------------MSEPLP---------P-------------TSIR--ELIE--Q-FIH-DR-----F----ATK---------------------------------------------------------------------T---------E----------------K---------L--------AP------D----DP---------AY----LK--------------------LQ-------------------------------------------QQ-----------
FE---LQTW-----LA---DA-----ARRV-------G----QLQ-V----------------VTHSLKAIHPE--A----------------K---G-T------NL----Y----V----E---------P-GQL---HGK--E---G---------VGS--H--CL------P-T-----D--F--E-GD-V-VG----N----AA----A--L-----DVYKFL-K------L----R--W--Q---------GRSL------------LELAL-END-E---SLIDAL---S---E----DR----AEAQAWVHAFAGI--V---EP--KG---
TPSSHARGKQIYWLV-------GD-----EP------TD----NAS---YHLLAPLYATALTHRFYAQ-VS----P--N---TWFF-G----------------DH-NKAAR-K-----AL-------------R--ERTAF--EG-G-YTS-----YPN-LAVQKLGG-T----KPQ----------NISQLN--SERG-----------GQNYLLASL-P-----PTWT--S-R-EV-YAPMR------DVF----RA----FN--G-R--PEV--RRIVGELKRFL-----------E------A-----E--------------PAK---
---N----MH---TR--DL----RD------DLTTMLVDELMLFTLDIH--S-------L-A-----A---G----WT------A-----D---E---K-----CQ--------LAP---EEQYWL-----D-------------PER--GLE-------------DADFDD--ARR--RAD-WQ-EAVSSRAALWLN-----RE----L---N------HK----S--Q-L-S-------------F---GDPEH------QHWKS-Q-------FE--GI-----------VTEFR--R-----QVDDLQDAL----------------RDDV-------D--
DPDDPDAPEGDAA------------------------
WP_104203293     ------------------------------------------MPEHFH---------SL------------PTIQ--ALIY--G-FIH-DR-----F----ATK---------------------------------------------------------------------T---------E----------------K---------L--------PA-----N----DP---------KY----LE--------------------LQ-------------------------------------------QQ-----------
FE---PQTW-----LA---DA-----ARRV-------S----QLQ-V----------------VTHSLKAIHPD--A----------------K---G-T------NL----Y----V----E---------P-GQL---NQG--N---L---------VGS--H--CL------P-A-----D--F--Q-GD-V-VG----N----AA----A--L-----DVYKFL-K------L----E--W--Q---------GHTL------------LERAI-DFD-E---TLIDAL---S---Q----DR----DEAESWMRAFASI--T---EP--KG---
TPSSHLMGKQLYWLV-------GD-----DP------ND----DNN---YHLLAPLYATALTHRFYAQ-VS----P--N---TWFF-G----------------DH-NKAAR-K-----AL-------------K--DKTPF--EG-G-YTS-----YPN-LAVQKLGG-T----KPQ----------NISQLN--SERG-----------GQNYLLASL-P-----PNWE--S-R-EV-YAPMR------DAF----RA----FK--G-R--PNV--RRIVGELKRLL-----------E------A-----D--------------PAK---
------N----MH---TR--DL----RD------DLTAMLVDELMLFTFAIH--S-------L-A-----P---G----WT------A-----D---E---K-----CQ--------LAL---EEQYWL-----D-------------PGR--AHE-------------DSEFAE--ARR--QSN-WH-EVISSRAANWLN-----RE----L---S------YN----S--P-L-S-------------F---GDAEH------HHWKR-R-------FE--GI-----------VTEFR--R-----QLDDLQEAL----------------RDED-------D--D---
----QGDAA------------------------
WP_064121249     ------------------------------------------M--PEP--QHH---------ASIR--DLVE--W-FIR-DR-----F----ATK---------------------------------------------------------------------A---------E---------K-LAA--DDPK-------Y--L--------------------------------------------EL-----------------------------------Q-QQ-----------
--FD---PQAW-----LA---DA-----ARRV-------S----QLQ-V----------------VTHSLKAIHPD--A----------------K---G-T------NL----Y----V----E---------P-GQL---NQ--G--GL---------VGS--H--CL----PAD-----------F--Q-GD-V-VG----N----AA----A--L-----DVYKFL-K------Q----E--W--Q---------GRTL------------LERAL-DFD-E---ALIDAL---S---L----DR----EEAENWVRAFASI--T---EP--KG-T--
PSSHVMGKQLYWLV-------GD-----DP------TD----DKN---YHLLAPLYGTALTHRFYAQ-AS----P--N---TWFF-G----------------DH-NKAAR-K-----AL-------------K--DKTFF--EG-G-YTS-----YPN-LAVQKLGG-T----KPQ----------NISQLN--SERG-----------GQNYLLASL-P-----PHWK--S-R-EI-YAPMR------DAF----RA----FK--G-R--PDV--RRTISELKRFL-----------E------T-----D--------------PAK------
--N----MH---TR--DL----RD------DLTAMLVDELMLFTFDIY--S-------L-A-----P---G----WT------A-----D---E---K-----CQ--------LAL---EEQYWL-----D-------------PGR--AEE-------------DGEFAE--ARR--QSN-WH-AAVSSRAANWLN-----RE----L---N------HK----S--P-L-S-------------F---GDAEH------HHWKR-R-------FE--GI-----------VTEFR--R-----QLDDLQETL----------------RDED-------N--D---
-QGDAA------------------------
WP_084671801     ------------------------------------------MIE--W-FIH-DR-----F----ATK---------------------------------------------------------------------V---------E---------K-LAT--GDPS-------Y--L--------------------------------------------EL----------------------------------Q-QQ-------------------------FE---
PETW-----LA---NA-----AHRV-------S----QLQ-V----------------VTHSLKAIHPD--A----------------K---G-T------NL----Y----V----EP-----------GP---LVQ--N--GL---------VGS--H--CL----PAD-----------F--Q-GD-V-VG----N----AA----A--L-----DVYKFL-K------L----E--W--Q---------GRSL------------LGLAL-DFD-E---VLIDAL---S---Q----DR----EKAKSWMRAFASI--T---EP--KG-K--PSSHVRGKQIYWLV--
-----GD-----DP------TD-----NES---YHLLAPLYATALTHRFYAQ-VS----P--N---AWFF-G----------------DH-NKAAR-K-----AL-------------K--DKAPF--EG-G-YIS-----YPN-LAAQKMGG-T----KPQ----------NISQLN--SERG-----------GQNYLLASL-P-----PNWE--S-R-EI-YAPMR------EPF----RA----FK--R-R--PAV--RKIVDELKRFL-----------E------V-----G--------------PAK-----------N-----MH---TR--DL----
RD------DLTVMLVDELMLFTFDIH--S-------L-T-----P---G----WT------T-----D---E---K-----CQ--------LAP---EEQYWL-----D-------------PGR--ADE-------------DGEFAE--ARR--RSD-WH-AAVSSRAAIWLN-----RE----L---G------HK----S--R-L-H-------------F---GDAEH------HRWER-Q-------FE--GV-----------MTEFR--R-----QLNDLQDAL----------------RDEE-------D--M-------EALA--
-----
WP_110619891     ------------------------------------------MHD---------ESV------------QPLR--QRIS--E-FLD-ER-----L----ETK---------------------------------------------------------------------L---------S----------------G---------L--------KD------D----DP---------EK----VS--------------------KE-------------------------------------------Q-ELR-EK-----------
-----FR---REAW-----IA---DA-----ARRV-------S----QLQ-V----------------VTHPTKATHPD--S----------------K---A-T------SL----Y----T----P---------P-PEL---SDH--S---L---------VGS--H--VL------G-K-----D--F--P-TD-V-VG----N----AA----S--L-----DVFKFL-K------I----E--H--E---------GRTL------------LARAL-EDE-P---ELARAF---S---D----DS----DQGREWVRSFAGI--T---EP--RG---
AVKGDPRAKQVYWLV-------GD-----EP------TR----DAD---FHLLAPLYSTVLAHRIHAV-IS----H--T---R--F-S----------------EE-SRAAR-K-----AH-------------W--NGEYS--EY-E-HRE-----YPY-LAAQTFGG-S----KPQ----------NISQLN--SERG-----------GNNYLLGSL-P-----PNWR--S-A-EV-RPLLHT-----DSAL----RA----FG--R-R--TTV--RQTLRELERFL-----------V------A-----N--------------PRP----
---N----QE---TR--QR----RD------ALVAALVDELLVFAAELQ--E-------L-E-----P---G----WS------A-----D---P---A-----CE--------LPE---EEALWL-----D-------------PER--ASV-------------DPEFAQ--RRE--RGD-WV-SEVRDRFAAWLN-----QR----L---H------K-----S-L-P-------------V---GDAEH------DYWEE-Q-------LK--HE-----------------TAALEEVL----------------THA-----
-------------------
WP_027313110     ------------------------------------------MTEDPQ---------QEA------------SRLR--AQIA--A-FIK-ER-----L----DAK---------------------------------------------------------------------L---------K----------------K---------L--------EP------N----DP---------KY----AL--------------------LI-------------------------------------------QQ-----------
--HE---YHTW-----LN---DA-----AKRA-------G----KIQ-I----------------ATHTIKAIHTS--A----------------R---G-T------NL----Y----I----E---------P-HNL---PSH--P---I---------VGS--H--LL------G-N-----N--F--K-PDAA-VE----D----AK----V--L-----DVYAFL-Q------I----Q--T-GS---------GKTL------------LELAL-LQD-E---SLKNAL---N---H----DE----EKAQHLLDAFAAP--V---KA--AR---
KAVSHTLAKQLYWHI-------GN-----DP------CQ----DDQ---YHLLAPLFASSLTQRVYEQ-IN----E--D---R--F-G----------------PA-ATSAR-E-----AR-------------K--TKSPH--HQ-G-FNE-----YPN-LAIQKLGG-E----NPQ----------NAGKLN--QARR-----------GQNYLLASL-P-----PRWD--R-S-LL-TPPLRT-----DSVF----PR----FT--R-R--KEV--WLTVRQLRALL-----------L------S-----E--------------PPK---
---N----LQ---TR--ET----RY------SLTSKLVDELLQFGAEYW--Q-------L-P-----P---G----WS------A-----Q---P---E-----CR--------LVE---AEALWL-----D-------------PYR--AHH-------------DEDFAS--KWR--QGD-WQ-QTICQRFANWLN-----RL----L---R------IV-----D-L-P-------------V---ADPEH------RQWSK-E-------VK--LA-E--DW-REWLKHLE--Q-----ELNQLQKEL----------------SNE-------
-----------------------------------
WP_110189784     ------------------------------------------MPASNK---------DES------------DKLR--ALIA--A-FIK-ER-----L----DAK---------------------------------------------------------------------L---------D----------------K---------L--------SA------D----DP---------KY----QA--------------------LT-------------------------------------------AQ-----------
--YQ---YSSW-----LE---DA-----ARRV-------G----QIQ-S----------------VTHTLKAIHPD--A----------------R---G-T------NL----Y----V----S---------P-SDL---TGH--S---V---------LGS--H--DL------P-V-----S--F--S-AD-V-VG----N----AA----A--L-----DVYKLL-K------L----A--G-DD---------GRTL------------LDRAL-AQD-T---ALKAAL---S---D----DP----EQAQQWLDAFAGL--V---AS--RG---
GSASHQRAKQVYWLT-------GN-----DP------LN----DEH---YHLLAPLYASSLSHQVFQQ-IN----S--H---R--F-G----------------EE-AKAAR-E-----AR-------------R--KGEQH--AT-G-YYD-----YPA-LAVQKLGG-T----KPQ----------NISQLN--SERG-----------GNNYLLASL-P-----PSWD--R-N-LL-TPPLRT-----DSVF----PR----FT--R-R--PEV--RLTVRQLRVLL-----------L------G-----D--------------PPK----
-----N----VQ---TR--ET----RF------SLTSRLVDELLQFGAECW--Q-------L-P-----P---G----WS------A-----Q---P---E-----CR--------LVE---AETLWL-----D-------------PYR--AKD-------------DEAFAA--RWR--QGD-WQ-QEICQRFASWLN-----RL----L---R------IA-----E-L-P-------------V---ADPEH------KQWSK-E-------VK--LT-D--DW-REWLKHLE--Q-----ELDQLQKEL----------------SYE-------
-----------------------------------
WP_104156584     ------------------------------------------MPASNK---------DES------------DKLR--ALIA--A-FIK-ER-----L----DAK---------------------------------------------------------------------L---------D----------------K---------L--------SA------D----DP---------KY----QA--------------------LT-------------------------------------------AQ-----------
--YQ---YSSW-----LE---DA-----ARRV-------G----QIQ-S----------------VTHTLKAIHPD--A----------------R---G-T------NL----Y----V----S---------P-SDL---TGH--S---V---------LGS--H--DL------P-V-----S--F--S-AD-V-VG----N----AA----A--L-----DVYKLL-K------L----E--D-DD---------GRTL------------LDRAL-AQD-A---ALKAAL---S---D----DP----EQAQQWLDAFAGL--V---AS--RG---
GSASHQRAKQVYWLT-------GN-----DP------LN----DEH---YHLLAPLYASSLSHQVFQQ-IN----S--H---R--F-G----------------EE-VKAAR-E-----AR-------------R--KGEQH--AT-G-YYD-----YPA-LAVQKLGG-T----KPQ----------NISQLN--SERG-----------GNNYLLASL-P-----PSWD--R-N-LL-LPPLRT-----DSVF----PR----FT--R-R--KEV--WLTVRQLRVLL-----------L------G-----A--------------PPK----
-----N----LL---TR--ET----RF------SLTSKLVDELLQFGAEYW--Q-------L-P-----P---G----WS------A-----Q---P---E-----CR--------LVE---AETLWL-----D-------------PYR--AKD-------------DEAFAA--RWR--QGD-WQ-QEICQRFASWLN-----RL----M---R------IA-----E-L-P-------------V---ADPEH------KQWSK-E-------VK--LT-D--DW-REWLKHLE--Q-----ELDQLQKEL----------------SYE-------
-----------------------------------
WP_115719507     ------------------------------------------MADLE---------SRS------------QKLR--ALIA--E-FIQ-GR-----L----AAK---------------------------------------------------------------------L---------E----------------K---------L--------VA------D----DP---------KY----AL--------------------LT-------------------------------------------AQ-----------
-YQ---YDAW-----LE---SA-----ARRV-------G----QIQ-S----------------VTHTLKAIHPD--A----------------K---G-S------NL----Y----V----S---------P-GTL---KAH--S---L---------LGS--H--DL------A-G-----G--F--Q-AD-V-VG----N----AA----A--L-----DVYKFL-K------L----T--D-EH---------GNTL------------LELAL-ADD-A---AFKAAL---S---K----DA----AKAEQWLAAFASL--L---GS--DG---
ALASHPQAKQIYWLT-------GE-----DP------CN----DLH---YQVLAPLYASSLAHEVFTR-IN----Q--H---R--F-G----------------EE-AKAAR-E-----AR-------------R--NGHFH--ET-G-YCD-----YPN-LAVQKLGG-T----KPQ----------NISQLN--SERG-----------GNNYLLASL-P-----PSWD--T-S-LL-TPPLKT-----DSVF----KR----FG--R-R--PAV--RQTVKALQGLL-----------K------A-----E--------------PPK----
---N----LR---TR--ET----RF------SLTSAVVDELLQFGAEFW--E-------L-N-----P---G----WS------Q-----H---P---D-----CR--------LVE---EEIFWL-----D-------------PYR--AKS-------------DLDFAE--RRQ--QAD-WP-QQIRERFANWLN-----AQ----L---H------KA-----G-L-P-------------V---GDPEF------AQWQK-E-------VK--LD-D--DW-RHWLKHLQ--Q-----ELDQLQKEL----------------SHD-------
-----------------------------------
WP_050658320     ------------------------------------------MSPPFN---------ENV------------IKLR--ALIA--D-FIQ-GR-----L----ATK---------------------------------------------------------------------L---------E----------------K---------L--------AA------D----DP---------KY----AL--------------------LT-------------------------------------------AQ-----------
YQ---YDAW-----LE---SA-----ARRV-------G----QIQ-S----------------VTHTLKAIHPD--A----------------K---G-S------NL----Y----V----S---------P-GTL---TAH--P---L---------LGS--H--DL------A-G-----G--F--Q-AD-V-VG----N----AA----A--L-----DVYKFL-K------L----T--D-ER---------GNTL------------LELAL-ADD-A---ALKAAL---S---K----DA----TEAEQWLAAFASL--L---GS--DG---
ALASHPQAKQIYWLT-------GE-----DP------GN----DLH---YQVLAPLYASSLAHEVFTR-IN----Q--H---R--F-G----------------EE-AKAAR-E-----AR-------------R--NGHFH--ET-G-YCD-----YPN-LAVQKLGG-T----KPQ----------NISQLN--SERG-----------GNNYLLASL-P-----PSWD--T-S-LL-SPPLKT-----DSVF----KR----FG--R-R--PAV--RQTVKALQGFL-----------K------A-----E--------------PPK----
---N----LR---TR--ET----RF------SLTSTLVDELLQFGAEFW--E-------L-S-----P---G----WS------Q-----H---P---D-----CR--------LVE---EEIFWL-----D-------------PYR--AKS-------------DLDFAE--RRQ--QAD-WP-QQIRERFANWLN-----AQ----L---H------KA-----G-L-P-------------V---GDPEF------AQWQK-E-------VK--LD-D--DW-RHWLKHLQ--Q-----ELDQLQKEL----------------SHD-------
-----------------------------------
WP_123421524     ------------------------------------------MADLE---------SQS------------QKLR--ALIA--D-FIQ-GR-----L----ATK---------------------------------------------------------------------L---------E----------------K---------L--------AA------D----DP---------KY----AL--------------------LT-------------------------------------------AQ-----------
-YQ---YDAW-----LE---SA-----ARRV-------G----QIQ-S----------------VTHTLKAIHPD--A----------------K---G-S------NL----Y----V----S---------P-GTL---TAH--P---L---------LGS--H--DL------A-G-----G--F--Q-AD-V-VG----N----AA----A--L-----DVYKFL-K------L----T--D-ER---------GNTL------------LELAL-ADD-A---ALKAAL---S---K----DA----TEAEQWLAAFASL--L---GS--DG---
ALASHPQAKQIYWLT-------GE-----DP------GN----DLH---YQVLAPLYASSLAHEVFTR-IN----Q--H---R--F-G----------------EE-AKAAR-E-----AR-------------R--NGHFH--ET-G-YCD-----YPN-LAVQKLGG-T----KPQ----------NISQLN--SERG-----------GNNYLLASL-P-----PSWD--T-S-LL-SPPLKT-----DSVF----KR----FG--R-R--PAV--RQTVKALQGFL-----------K------A-----E--------------PPK----
---N----LR---TR--ET----RF------SLTSTLVDELLQFGAEFW--E-------L-S-----P---G----WS------Q-----H---P---D-----CR--------LVE---EEIFWL-----D-------------PYR--AKS-------------DLDFAE--RRQ--QAD-WP-QQIRERFANWLN-----AQ----L---H------KA-----G-L-P-------------V---GDPEF------AQWQK-E-------VK--LD-D--DW-RHWLKHLQ--Q-----ELDQLQKEL----------------SHD-------
-----------------------------------
WP_037003915     ------------------------------------------MNALSS---------SRQ------------QAIR--SLIE--H-FIA-ER-----L----AAK---------------------------------------------------------------------L---------E----------------G---------L--------AD------D----DP---------KR----AT--------------------LL-------------------------------------------AQ-----------
-YT---PAAW-----LE---DA-----ARRA-------G----QIQ-A----------------VTHTLKAIHPD--A----------------K---G-T------NL----Y----C----L---------P-NSL---PQH--A---L---------VGS--H--CL------G-D-----D--F--A-GD-V-VG----N----AA----A--L-----DVYKML-K------L----E--H--D---------GQPL------------LSLLL-AED-E---GALAAL---H---D----DA----EQSRSLTAAFTGI--T---CA--RG---
GLASHTHGKQLYWLT-------GD-----DP------CN----DAD---FHLLAPLYASSLAHRVYQQ-IQ----A--D---R--F-G----------------EP-AKEAR-Q-----AR-------------R--DKQFH--PG-V-IHE-----YPD-LAVQKLGG-T----KPQ----------NISQLN--SERR-----------GDNYLLASL-P-----PSWQ--Q-R-DV-RLPIRT--RDGSIF----PA----FG--K-R--PAV--AELVRNLRDFL-----------L------S-----G--------------PPP----
-----N----VH---TR--NS----RD------ELLTELIDELLLYAAELW--A-------F-E-----P---G----WS------A-----K---P---E-----CN--------LNR---AEQLWL-----D-------------FRR--AEH-------------DAVFAS--EWR--HMD-WL-TEIRERFANWLN-----AR----L---V------S-------K-L-S-------------V---GDVEN------SYWAK-Q-------AG--SN-A--LW-LDLDWLNQ--A-----WVDQFEREQ----------------RDL-------
---------EAEHE-----------------------
WP_074676881     ------------------------------------------MHDPST---------HRQ------------SEIR--SLIQ--G-FIA-ER-----L----ATK---------------------------------------------------------------------L---------E----------------G---------L--------AD------D----DP---------RH----TT--------------------LL-------------------------------------------AQ-----------
--YA---PATW-----LE---EA-----ARRA-------G----QIQ-A----------------VTHTLKAIHPD--A----------------K---G-T------NL----Y----C----L---------P-RAL---PQH--Q---L---------VGS--H--CL------G-D-----D--F--A-GD-V-VG----N----AA----A--L-----DVYKLL-K------L----E--H--D---------GQAL------------LDLLL-AAD-A---DAIAAL---H---D----DA----EQARSLAAAFTSI--T---RA--RD---
GLASHTHGKQLYWLV-------GD-----DP------RN----DAH---FHLLAPLYASSLAHRTYQG-IQ----A--D---R--F-G----------------EP-AKEAR-Q-----AR-------------R--DKQFH--PG-V-IHE-----YPD-LAVQKLGG-T----KPQ----------NISQLN--SERR-----------GDNYLLASL-P-----PNWQ--Q-R-DV-RLPIRT--RDGSIF----PA----FG--Q-R--PAV--AELVRNLRDFL-----------L------S-----G--------------PPP----
------N----VH---TR--NS----RD------ELLTELIDELLLYAAELW--A-------F-E-----P---G----WS------T-----R---P---E-----CN--------LNR---AEQLWL-----D-------------FRR--AEH-------------DAIFAS--EWR--RME-WL-AEIRQRFANWLN-----AR----L---V------S-------K-L-S-------------V---GDVEN------SYWAK-K-------AG--SN-A--LW-LDLDWLNR--A-----WVEQFEREQ----------------RDL-------
-------EAEHE-----------------------
WP_045733459     ------------------------------------------MHDPSP---------HRQ------------SEIR--TLIQ--R-FIA-ER-----L----ATK---------------------------------------------------------------------L---------E----------------G---------L--------AD------D----DP---------KR----AT--------------------LL-------------------------------------------AQ-----------
--YA---PDVW-----LE---DA-----ARRA-------G----QIQ-A----------------VTHTLKAIHPD--A----------------K---G-T------NL----Y----C----L---------P-SAL---PQH--Q---L---------VGS--H--CL------G-D-----D--F--A-GD-V-VG----N----AA----A--L-----DVYKLL-K------L----E--H--D---------GQTL------------LDLLL-SAD-A---DAIAAL---H---I----DD----EQARSLAAAFTGI--T---RA--RD---
GLASHTHGKQLYWLV-------GD-----DP------RN----DEH---YHLLAPLYASSLAHRVYQE-LQ----A--D---R--F-G----------------EP-AKEAR-Q-----AR-------------R--GKQFH--PG-V-IHE-----YPD-LAVQKLGG-T----KPQ----------NISQLN--SERR-----------GDNYLLASL-P-----PSWQ--P-R-DV-RLPIRT--RDGSIF----PA----FG--Q-R--PAV--AELVRSLRDFL-----------L------S-----G--------------PPA----
-----N----VH---TR--NS----RD------ELLTELIDELLLYAVELW--A-------F-E-----P---G----WS------A-----K---P---E-----CN--------LNR---AEQLWL-----D-------------FRR--AEH-------------DAIFAS--EWR--RME-WL-TEIRERFANWLN-----AR----L---V------S-------K-L-S-------------V---GDVEN------SYWVK-K-------AG--SN-A--LW-LDLDWLNR--A-----WVEQFEREQ----------------RDL-------
-------EAEHE-----------------------
WP_075750822     ------------------------------------------MNAPSS---------SRQ------------QAIR--SLIE--R-FIA-ER-----L----ATK---------------------------------------------------------------------L---------E----------------G---------L--------AD------D----DP---------KR----AT--------------------LL-------------------------------------------AQ-----------
--YA---PDVW-----LE---DA-----ARRA-------G----QIQ-A----------------VTHTLKAIHPD--A----------------K---G-T------NL----Y----C----L---------P-NSL---PQH--A---L---------VGS--H--CL------G-D-----D--F--A-GD-V-VG----N----AA----A--L-----DVYKLL-K------L----E--H--D---------GQTL------------LDLLL-AAD-A---DAIAAL---H---T----DD----EQARSLAPAFTGI--T---RA--RD---
GLASHTHGKQLYWLV-------GD-----DP------RN----DEH---YHLLAPLYASSLAHRVYQA-IQ----A--D---R--F-G----------------EP-AKEAR-Q-----AR-------------R--DKQFH--PG-V-IHE-----YPD-LAVQKLGG-T----KPQ----------NISQLN--SERR-----------GNNYLLASL-P-----PSWQ--P-R-DV-RLPIRT--RDGSIF----PA----FG--K-R--PAV--AELVRSLRDFL-----------L------S-----G--------------PPA----
-----N----VH---TR--NS----RD------ELLTELIDELLLYAVELW--A-------F-E-----P---G----WS------A-----K---P---E-----CN--------LNR---AEQLWL-----D-------------FRR--AEH-------------DAIFAS--EWR--RME-WL-TEVRERFANWLN-----AR----L---V------S-------K-L-S-------------V---GDVEN------SYWVK-K-------AG--GN-A--LW-LDLDWLNR--A-----WVEQFEREQ----------------RDL-------
-------EAEHE-----------------------
WP_128578447     ------------------------------------------MNAPSS---------SRQ------------QAIR--SLIE--R-FIA-ER-----L----ATK---------------------------------------------------------------------L---------E----------------S---------L--------SA------D----DP---------KR----AT--------------------LL-------------------------------------------AQ-----------
-YA---PDVW-----LE---DA-----ARRA-------G----QIQ-A----------------VTHTLKAIHPD--A----------------K---G-T------NL----Y----C----L---------P-NSL---PQH--A---L---------VGS--H--CL------G-D-----D--F--A-GD-V-VG----N----AA----A--L-----DVYKLL-K------L----E--H--D---------GQTL------------LDRLL-AAD-A---DAIAAL---H---T----DD----EQARSLAAAFTGI--T---RA--RD---
GLASHTHGKQLYWLV-------GD-----DP------RN----DEH---YHLLAPLYASSLAHRVYQE-IQ----A--D---R--F-G----------------EP-AKEAR-Q-----AR-------------R--DKQFH--PG-V-IHE-----YPD-LAVQKLGG-T----KPQ----------NISQLN--SERR-----------GDNYLLASL-P-----PSWQ--P-R-DV-RLPIRT--RDGSIF----PA----FG--Q-R--PAV--AELVRSLRDFL-----------L------S-----G--------------PPA----
-----N----VH---TR--NS----RD------ELLTELIDELLLYAVELW--A-------F-E-----P---G----WS------A-----K---P---E-----CN--------LNR---AEQLWL-----D-------------FRR--AEH-------------DAVFAS--EWQ--RME-WL-AEVRQRFANWLN-----GR----L---V------S-------K-L-S-------------V---GDVEN------SYWIK-Q-------AG--SN-A--LW-LDLDWLNQ--A-----WVEQFEREQ----------------RDL-------
--------EAEHE-----------------------
WP_074855815     ------------------------------------------MHSPSP---------GRR------------QAIR--SLIE--R-FIA-ER-----L----TTK---------------------------------------------------------------------L---------E----------------G---------L--------AD------D----DP---------KR----AT--------------------LL-------------------------------------------AQ-----------
--YA---PDVW-----LE---DA-----ARRA-------G----QIQ-A----------------VTHTLKAIHPD--A----------------K---G-T------NL----Y----C----L---------P-NSL---PQH--A---L---------VGS--H--CL------G-D-----D--F--A-GD-V-VG----N----AA----A--L-----DVYKLL-K------L----E--H--D---------GQTL------------LDLLL-AAD-A---DAIAAL---H---T----DD----EQARSLAAAFTGI--T---RA--RD---
GLASHTYGKQLYWLV-------GD-----DP------RN----DEH---YHLLAPLYASSLAHRVYQE-IQ----A--D---R--F-G----------------EP-AKEAR-Q-----AR-------------R--DKQFH--PG-V-IHE-----YPD-LAVQKLGG-T----KPQ----------NISQLN--SERR-----------GDNYLLASL-P-----PSWQ--P-R-DV-RLPIRT--RDGSIF----PA----FG--Q-R--PAV--AELVRTLRDFL-----------L------S-----G--------------PPA----
-----N----VH---TR--NS----RD------ELLTELIDELLLYAVELW--A-------F-E-----P---G----WS------A-----K---P---E-----CN--------LNR---AEQLWL-----D-------------FRR--AEH-------------DAIFAS--EWR--RME-WL-TEIRERFANWLN-----AR----L---V------S-------K-L-S-------------V---GDVEN------SYWAK-Q-------AG--SN-A--LW-LDLDWLNR--A-----WVEQFEREQ----------------RDL-------
------EAEHE-----------------------
WP_108234999     ------------------------------------------MHSPSP---------GRR------------QAIR--SLIE--R-FIA-ER-----L----TTK---------------------------------------------------------------------L---------E----------------G---------L--------AD------D----DP---------KR----AT--------------------LL-------------------------------------------AQ-----------
--FA---PDVW-----LE---DA-----ARRA-------G----QIQ-A----------------VTHTLKAIHPD--A----------------K---G-T------NL----Y----C----L---------P-NSL---PQH--A---L---------VGS--H--CL------G-D-----D--F--A-GD-V-VG----N----AA----A--L-----DVYKLL-K------L----E--H--D---------GQTL------------LDLLL-AAD-A---DAIAAL---H---T----DD----EQARSLAAAFTGI--T---RA--RD---
GLASHTYGKQLYWLV-------GD-----DP------RN----DEH---YHLLAPLYASSLAHRVYQE-IQ----A--D---R--F-G----------------EP-AKEAR-Q-----AR-------------R--DNQFH--PG-V-IHE-----YPD-LAVQKLGG-T----KPQ----------NISQLN--SERR-----------GDNYLLASL-P-----PSWQ--P-R-DV-RLPIRT--RDGSIF----PA----FG--Q-R--PAV--AELVRTLRDFL-----------L------S-----G--------------PPA----
-----N----VH---TR--NS----RD------ELLTELIDELLLYAVELW--A-------F-E-----P---G----WS------A-----K---P---E-----CN--------LNR---AEQLWL-----D-------------FRR--AEH-------------DAIFAS--EWR--RME-WL-TEIRERFANWLN-----AR----L---V------S-------K-L-S-------------V---GDVEN------SYWAK-Q-------AG--SN-A--LW-LDLDWLNR--A-----WVEQFEREQ----------------RDL-------
------EAEHE-----------------------
```

WP_125874184 ----------MHSPSP---------GRQ-----------QAIR--SLIE--R-FIA-ER-----L----ATK----------L---------E---------------S---------L--------SA------D----DP---------KR----AT------------------LL-----------------------------------AQ------------
-YA---PDVW-----LE---DA-----ARRA-------G----QIQ-A---------------VTHTLKAIHPD--A---------------K---G-T------NL----Y----C----L--------P-NSL---PQH--A---L---------VGS--H--CL------G-D-----D--F--A-GD-V-VG----N----AA----A--L-----DVYKLL-K-----L----E--H--D---------GQTL-----------LDLLL-AAD-A---DAIAAL---H---T----DD----EQARSLAAAFTGI--T---RA--RD---
GLASHTYGKQLYWLV-------GD-----DP------RN----DEH---YHLLAPLYASSLAHRVYQE-IQ----A--D---R--F-G---------------EP-AKEAR-Q-----AR------------R--DKQFH--PG-V-IHE-----YPD-LAVQKLGG-T----KPQ---------NISQLN--SERR----------GDNYLLASL-P----PSWQ--P-R-DV-RLPIRT--RDGSIF----PA----FG--Q-R--PAV--AELVRTLRDFL----------L------S-----G------------PPA----
-----N----VH---TR--NS----RD------ELLTELIDELLLYAVELW--A-------F-E-----P---G----WS------A-----K---P---E-----CN--------LNR---AEQLWL-----D-------------FRR--AEH-------------DAIFAS--EWR--RME-WL-TEIRERFANWLN-----AR----L---V-------S-------K-L-S------------V---GDVEN------SYWAK-Q-------AG--SN-A--LW-LDLDWLNR--A-----WVEQFEREQ--------------RDL------------
------EAEHE--------------------

WP_003464677 ----------MNAPSS---------SRQ-----------QAIR--SLIE--R-FIA-ER-----L----ATK----------L---------E---------------S---------L--------SA------D----DP---------KR----AT------------------LL-----------------------------------AQ------------
-YA---PDVW-----LE---DA-----ARRA-------G----QIQ-A---------------VTHTLKAIHPD--A---------------K---G-T------NL----Y----C----L--------P-SAL---PQH--Q---L---------VGS--H--CL------G-D-----D--F--A-GD-V-VG----N----AA----A--L-----DVYKLL-K-----L----E--H--D---------GQTL-----------LDLLL-AAD-A---DAIAAL---H---T----DD----EQARSLAAAFTGI--T---RA--RD---
GLASHTYGKQLYWLV-------GD-----DP------RN----DEH---YHLLAPLYASSLAHRVYQE-IQ----A--D---R--F-G---------------EP-AKEAR-Q-----AR------------R--DKQFH--PG-V-IHE-----YPD-LAVQKLGG-T----KPQ---------NISQLN--SERR----------GDNYLLASL-P----PSWQ--P-R-DV-RLPIRT--RDGSIF----PA----FG--Q-R--PAV--AELVRTLRDFL----------L------S-----G------------PPA----
-----N----VH---TR--NS----RD------ELLTELIDELLLYAVELW--A-------F-E-----P---G----WS------A-----K---P---E-----CN--------LNR---AEQLWL-----D-------------FRR--AEH-------------DAIFAS--EWR--RME-WL-TEIRERFANWLN-----AR----L---V-------S-------K-L-S------------V---GDVEN------SYWAK-Q-------AG--SN-A--LW-LDLDWLNR--A-----WVEQFEREQ--------------RDL------------
------EAEHE--------------------

WP_116620376 ----------MNAPSS---------SRQ-----------QAIR--SLIE--R-FIA-ER-----L----ATK----------L---------E---------------S---------L--------SA------D----DP---------KR----AT------------------LL-----------------------------------AQ------------
-YA---PDVW-----LE---DA-----ARRA-------G----QIQ-A---------------VTHTLKAIHPD--A---------------K---G-T------NL----Y----C----L--------P-NSL---PQH--A---L---------VGS--H--CL------G-D-----D--F--A-GD-V-VG----N----AA----A--L-----DVYKLL-K-----L----E--H--D---------GQTL-----------LDLLL-AAD-A---DAIAAL---H---T----DD----EQARSLAAAFTGI--T---RA--RD---
GLASHTYGKQLYWLV-------GD-----DP------RN----DEH---YHLLAPLYASSLAHRVYQE-IQ----A--D---R--F-G---------------EP-AKEAR-Q-----AR------------R--DKQFH--PG-V-IHE-----YPD-LAVQKLGG-T----KPQ---------NISQLN--SERR----------GDNYLLASL-P----PSWQ--P-R-DV-RLPIRT--RDGSIF----PA----FG--Q-R--PAV--AELVRTLRDFL----------L------S-----G------------PPA----
-----N----VH---TR--NS----RD------ELLTELIDELLLYAVELW--A-------F-E-----P---G----WS------A-----K---P---E-----CN--------LNR---AEQLWL-----D-------------FRR--AEH-------------DAIFAS--EWR--RME-WL-TEIRERFANWLN-----AR----L---V-------S-------K-L-S------------V---GDVEN------SYWAK-Q-------AG--SN-A--LW-LDLDWLNR--A-----WVEQFEREQ--------------RDL------------
------EAEHE--------------------

WP_018719049 ----------MPT---------GST-----------AALR--KTIE--E-FLS-ER-----Q----QTK----------L---------D---------------R---------L--------AS------D----DP---------KR----AR------------------II-----------------------------------DA------------
HQ---RENW-----LA---DA-----TRRV-------A----QLQ-V---------------VTHALKATHPD--A---------------R---G-T------SL----F----A----P--------P-ETL---PAH--P---A---------VGT--H--VL------G-D-----G--A--P-RD-V-VG----N----AA----A--L-----DVYKFL-Q-----T----E--H--D---------GRSM-----------LDRAL-AGE-A---AFAAAL---S---D----DP----TQGQAWMEAFAGI--T---RA--GD---
DQASHTRAKQVYWLV-------GD-----DP------TD----DRG---FHLLAPLYATSLAHAAFRA-IN----E--D---R--F-S---------------EV-AKAAR-T-----AR------------R--ERRYS--AV-E-SRE-----YPK-LAVQKFGG-T----KPQ---------NISQLN--SERR----------GGNYLLASL-P----PDWR--S-A-DV-RAPLHT----DSVF----PL----FG--R-R--RPV--RSAVNALLRFL----------R------S-----E------------PKP-----
----N----VA---TR--RH----RD------ELLAELTDALLVFAAELS--A-------L-P-----P---G----WS------A-----H---A---D-----CR--------LVE---AECLWL-----D-------------PDR--AAE-------------DAEFAM--RRA--QVD-WP-REVRERFAAWLN-----AQ----L---Q-------G-------P-L-P------------V---ADPER------RFWAD-R-------LQ--AE---------------------MDELEETL---------------PHV-------------------------
------------------

WP_061333308 [illegible]

WP_076724187 [illegible]

WP_091826793 [illegible]

WP_019935460 [illegible]

WP_114572975 [illegible]

WP_141440148 [illegible]

WP_096299905 [illegible]

WP_081192040 [illegible]

WP_038480455 [illegible]

WP_062373693 [illegible]

WP_062361891 [illegible]

WP_096650755 [illegible]

WP_101147093 [illegible]

WP_022520842 [illegible]

WP_009288126 [illegible]

WP_085917641 [illegible]

WP_064235066 [illegible]

WP_007111959 [illegible]

WP_092827231 [illegible]

WP_074397795 [illegible]

WP_097021695 [illegible]

WP_137055364 [illegible]

WP_013330863 [illegible]

WP_128689682 [illegible]

WP_133697396 [illegible]

WP_031385165 [illegible]

WP_052448503 [illegible]

WP_016418866 [illegible]

WP_041717099 [illegible]

WP_116027124 [illegible]

WP_097058516 [illegible]

WP_133491512 [illegible]

WP_145953648 [illegible]

WP_145912314 [illegible]

WP_012983811 [illegible]

WP_006748945 [illegible]

WP_015259208 [illegible]

IEASHTRAKQVYWLA-------GG------DP------TV----DTD---FHLLAPLYAASLAHVVYQT-IN----D--H---R--F-S-------------------EE-AKAAR-K------AR------------R--EGEYS--ET-G-FHE-----YPN-LAVQKLGG-T-----KPQ----------NVSQLN--LERH-------------GTNYLLASL-P-----PNWA-NR---DL-RPPLFV-----DSVF----SR--SSFG--R-R--PGV--RQVVTALKRFL---------AT-------------N--------------PPP----
-----N----VT---TR--QR----RD------KHLNKLVDELLLYAAELH---------T-L-E-------P---G-----WS-------A-----S---P---D-----CS---------LNR---FEALWL-----D------------PRR--ADI-------------DPDFAT--ECE--NEE-WV-DKVSHRFGNWLN-----AR-----L--QH------------------R-L--------------PF---DDAAF---------RFWKD-------------------------------------------SLREVLLHA--------------------------------
---------
WP_021760540 --MSKPDP------------ERI-------------RGLQ--QVMD--G-FLV-VR------R----DAK-------------------------------------------------------------------------L-----------E---------K---------L----------KP------D---DP---------KR----EE--------------------VS---------------A---------------
-----YE---RPAW------LA---DA-----ARRV--------G----QIQ-A----------------ATHTVKPMHPD---A----------------------R---G-T--------NL-----Y---C----K-----------A-DAL---PEH--A---L-------------VGS--H--LL-------G-E------G--C--A-AD-V-IG----N----AA-----A--L------DVNKFL-T-----Q----E---Y--Q------------GKTL---------LAWMQ-EDE-A---AVLAAF---A--VE----DA---ALARQWLQAFLGL--T---RT--PA-V-R-
VSHTLAKQLYWLT-------GD-----DP-------RD-----DAQ---YHLLAPLYASSLAHRVYLT-IQ-----A---H---R--F-S-------------------DE-TKAAR-E-----AK---------------K--SNSPC--EH-G-LHE-----YRE-LAVQKFGG-T-----KPQ----------NISQLN--SERK-------------GVNYLLASC-P-----PVWS--P-Q-PV-RPPFGV----ESVF-----VG----FG--R-Q--RDA--KALVRVLAEFL----------A-------S-----E-----------------PAA--------
T----LA---TR--QH----RD---------ELLFQILGELHQFAERYR---A------L-P-----P---G-----WS-------A-----D-----P---R------CR---------LVE----AQQCWL-----D------------PQR--AAM------------DAEFAD--IWL--HTA-WR-DEIKRRFGNWLN-----AR-----L--G----------DK-------L--P-----------------L--GDVEH--------VYWAD---------L--FD-D--ERWRRQLQDADRRML------EKEAPHA------------------------
WP_024892113 --MVHTPT-----------GRQ--------------AMFR--ASIE--T-FIR-ER-----L---DAK---------------------------------------------------------------------------L----------D---------------K---------L----------AD-----D---DP----------RR-----DA----------------------LI----------------------------QQ---------------
---HQ---PATW------LA---DA-----ARRA--------G----QIQ-A----------------VTHSLKPIHPD---A-----------------------R---G-S--------NL-----Y----C----P-------P-DTL---PRH--A---E------------IGT--H--AL-------A-G------D--F--T-GD-V-VG----N----AA-----A--L------DVYKFL-K------R---D--V--D---------------GRPL---------LDWML-DED-A---DLAAAL---H---P---DP----DQSRDWIAAFTGL--T---RP---RE-G-
GHASHTHAKQLYWLA-------GD-----DP-------LD-----DEQ---FRILAPLYATSLAHAVFQT-LN-----E---D--R--F-G-------------------ET-AKQAR-Q-----AR-------------R--DKQAH--PV-G-YRE-----YPH-LAVQKLGG-T-----KPQ----------NISQLN--SERG-------------GNNYLLASLPP-----PPWK--S-E-LQLLPSKGQ-----GSIF-----D-----VWS--K-N--LDV--WMPAGELKRFL------------A---------S----E-------------------------PGK--
--------GVRP--
WP_106701092 --MEA--HVQ-----------QRFA--HATS--R-WIN--------R-----ADS---------------------------------------------------------VARK-------------------------------------------------------------------------------------------AI---------------A-DAR-EK------------------------------
YM---LDSI------LG---KG-----INAA----------P------GVA-V------------ATHLAKGTHPD---P------------------R---V-K-----DV------V-----N-------------P-CVS---FDALSQ-HAE----------VGS--H--NL-------A-------GQAS--R-ED-V-TG----NGAVNSA-----G--Y------ELYLLL------------ECVF--D------------GKKL------------SDWIS-DKD-E---DAYLAF---G--------GE----DQAYSCLELLSNK--S-------RT-----
VATNEKLKQLYWFV-------GD-----------EQSASDATVDAQ---YHLLAPLYATSLAHAVYGV-LQ-----E---D--R--F-G-------------------EA-NKAAR-Q-----AR------------R--DGKLH--DG-P-LRD-----YPQ-LAVQKMGG-T-----KPQ----------NISQLN--SERG-------------GNNYLLASL-P-----PQWK--S-R-DI-RQPWGV-----QSVF-----DR--MLI--R-R--EGV--RATLADFLAFL------------K----------T---------D------------
PPA---------N---VE---TR--NR-----VD------AYYSDLIDALVTLGGELQ-R---A-W-P-----A---H----WT-------Q-----D---P---R------CE---------LVR----EEQLWL-----D------------ARR--AEV--------------DEDFGA--EWQ--RMD-WP-AQIGKRFGNWLN-----TQ-----L--E-------KQ---------F-A----------V----GDAEQ---------REWAN-E---------L--VD-ESEDGWAQNLHKR---T-------ALNAPHYIP--------------------
TRLG-------DQ----------GAVA----------
WP_124077196 --MPDPISA---------GRR---------------ERIH--AKID--E-FLA-GR-----L---IVK-------------------------------------------------------------------------L----------E---------K---------L-----------AP-----E---DP---------KR----QE-------------------LI----------------------------AQ---------------
-HQ---RPAW------LE---NA-----ARRV--------Q----QIQ-L----------------VTHTLKPIHPD---A----------------------R---G-T--------NL-----Y---I----D-----------P-GVM---APR--D---E-------------VGS--H--VL-------S-P------Q--F--S-AD-V-VG----N----AA-----A--L------DVYKFL-K------L---E--I--D--------------GRSL------------LDWLH-DED-G---EAMAAL---S---D----DV----AEARAWRDAFVSV--L---AQ--RG-G-
ELSTHPLSKQVFWLA-------GN-----DA-------TLDEQ---YHLLAPLYASSLAHAVFPV-IQ-----E---D--R--Y-G-------------------EK-NKAAR-Q-----AR-------------N--ARRWH--DG-A-VRE-----YPG-LAVQKLGG-T-----KPQ----------NVSQLN--SERG-------------GVNYLLGAL-P-----PVWQ--A-Q-EL-RPPWKM-----ESIF-----EG----VYA--R-Q--AGV--REVIAALLRFL-----------E------Q-----D-------------------PPR--
--------EA----TR--KR-----VS-------RYVDKLIDELVGLAVRYH---E------R-L-E-----P---G-----WS-------R-----D---A---R------CR---------DLE----EEKVWL-----D------------PLR--ATLRD------------ETEFAR--EWM--ALD-WP-VAIGSRFARWFN-----AQ-----L--R-------TR----------I-S----------V----GDDEH---------RNWKR-E---------L--LD-ERDGGWSAFLNTLR---R------ELDVQLRVV--------------A------
-------ESKA-------
WP_077297441 --MA---------PSV-------------PVFR--SAIA--A-FIE-ER-----L---AAK-------------------------------------------------------------------------L----------A---------K--------L-----------DA-----D---DP-----------KC----AA-----------------------LR--------------------------------EQ-------------------
YQ---YLTW-----LD---DA-----ARRV--------G----QIQ-L----------------VTHTLKPIHPD---A----------------------R---G-S--------NL-----Y---A-----P-----------P-STL---ADY--P---E-------------LGS--H--AL-------A-G------M-H--A-DD-V-VG----N----AA-----A--L------DVYKLL-K------L---Q--L--D-------------GRSL------------LDWLR-DGH-P---DDAHTLAEAFVGL--A---QP--RA-G-
GPASHTYAKQVFWLV-------GD-----DP-------LDDGQ---YHLLSPLYPSSLAHALFAV-LN-----E---H--R--Y-G-------------------EA-NKLAR-E-----AK-------------R--ARRDH--DQ-P-FYS-----YPG-LAVQKLGG-T-----KPQ----------NISQLN--SERG-------------GNNYLLSSL-P-----PIWK--S-H-KV-RPPTGA-----SVF------D----LFG--R-H--DDV--RRLVHGLRHFL------------E-------S----S-
APDRADGDAPGPKN-------N----RH---TR--DI------RYYADLIDELVLFGEELQ--E------Q-L-P-----A---N-----WS-------A-----H---A---D------CV---------LHE----EEQLWL-----D------------PYR--CET---------------DPKFCA--SWM--AMD-WP-AEVGQRFANWLN-----KQ-----L--A--------QH------------L-P----------------------E-----------RHWRK-E-----------L--VD-ERDGWARHLHRQR--R-------
EDQAPAYIP---------TRAA------T---------REAT---------
WP_086060192 --M------AE------D----DP---------KR----PD------------------------------------------------------------------AQ------------------------------------------------------------------------YQ---VETW-------LA-
-AA-----ARRV-------E-----QIR-A------------ATHTLKAIHPD---A-----------------R---G-T-------NL-----F-----V-----R--------P-QEL----CEL---V---Q------------LSS--H--VL-------T-E------S--F--T-FD-V-VG----N----AG-----A---L------DIYKFL-S-------Q----Q--L--D---------------GRSL--------LDRLQ-AKD-P---TLLAAF---C---D------HP----DQAQEWASAFLGL--L---ND--GE-Q-RPVSHVLAKQLYWLM-------GT----
DG-----------QNDEH---YHLLAPLFPSSLVHAVYAK-LR-----E---Y----Q--F-D---------------KQ-VKQGR-D-----AL----------Y--GKKMH--DG-V-VHV-----YPR-LAIRKLGS-T-----KPQ----------TVSQLN--MERG-------------GVNYLLSSL-P----PRWT--S-R-SS-PSPWGV-----ESIF-----DS----LLM--Q-R--REV--RQQLGGLRRFL-----------Q------S----Q------------PKS------I----KE---TR--DR---VD-------
DYLARLIDELVSFAAELR-T-----A-W-P-----A---H----WT-------A-----D---S---R------CQ---------LAR----EEQLWL-----D------------IER--VRQEEVFST-------EADQGFRA--EWY--WMD-WP-DQIGQRFARWIN----QR----L--D--------GC---------L-P------------V----GDVEY-------LHWKK-E-------LL--VD-Q-THWADQLQTLR--R-----Q--VPLSKT---------------RMK-----------------------EEA-----
WP_121738368 --MNESVRA---------GKT-------------GRWR--ELIQ--R-FIQ-AR-----L---QAK---------------------------------------------------------------------------L----------E---------K---------L----------SE------D---DP----------KR----PD-------------------LI----------------------------AQ----------------
-----YQ---VETW----LE----DA----ARRV----------K----QIQ-A------------VTHALKPIHPD---A----------------------R---G-T---------NL-----Y---V----D-----------P-TAL---PAL--T---Q------------LSS--H--VL-------H-D------N--F--R-AD-V-VG----N----AA-----A--L------DVYKFL-R------L---E--R--D---------------GCSI-----------LDRLQ-ADD-P---AVLAAF---C---S----DL----EQAREWQEAFLGL--L---QD--NG-Q-
ATASHVLAKQLYCLV-------GE-----D------------DDAH---YHLLAPLFPTSLVHAVHAV-LQ-----E---D--R--F-G-------------------EQ-MKQAR-Q-----AQ----------R--DKVMH--EH-A-VRW-----YPN-LAVRKLGS-T-----KPQ----------NISQLN--SERG-------------GVNYLLSSL-P-----PTWS--S-R-SV-PSPWAV-----ESIF-----DS----LFM--Q-R--REV--RQQVGGLRRFL-----------Q-------S--Q----------
PKP---------I---KE---TR--DR-----VD-------DHLARLIDELVSFAAELR---M------A-W-P-----A---H----WT-------A-----D---N---R------CQ--------LNR----EEQLWL-----D------------TER--VRQDDLFSS-----EDDQAFRA--EWY--WMD-WP-DQIGQRFARWIN-----GR-----L--G---------DT----------L-P----------V----GDVEY-------FHWKK-E---------LL--ID-Q-AHWASHLRTLR--Q-----QNNAPLYIS-------
GRK-----------------GGI------
WP_125304711 --M------AE------D----DP---------KR----PD------------------------------------------------------------------AQ------------------------------------------------------------------------YK----VETW-------LA-
-DA------ARRV------K----QIQ-A-------------VTHALKPIHPD---A----------------------R---G-T--------NL-----Y-----V------D---------P-TAL-----PAL--A---Q------------LGS--H--VL-------N-E-----K--F--A-SD-V-VG-----N----AA-----A--L------DVYKFL-R------L---E--L--D---------------GLSI------------LDRLQ-ADD-P---AMLAAL---G---S----DT-----DQAREWRGAFLGL--L---QD--NG-Q-SISSHVLAKQLYWLV----------GE-----DC-
--------HDDAQ---YHLLAPLFPTSLVHAVHAV-LQ-----E---D--R--F-G-------------------DQ-MKQAR-Q-----AQ-------------R--DKVMH--EH-A-VRW-----YPD-LAVRKLGS-T-----KPQ----------NISQLN--SERG-------------GVNYLLSSL-P-----PRWT--S-R-SS-PSPWGV-----ESIF-----DS----LLM--Q-R--REV--LQQVGGLRRFL-----------Q-------S--Q-------------PKS-------I-----KE----TR--DR-----VD-------
DYLARLIDELVSFAAELR---T------A-W-P-----A---H----WT-------A-----D---S---R------CQ---------LAR----EEQLWL-----D------------IER--VRQEEVFST-----EADQGFRA--EWY--WMD-WP-DQIGQRFARWIN-----GR-----L--D---------GC-----------L-P-----------V----GDVEY--------LHWKK-E---------LL--VD-Q-THWADQLRTLR--R-----Q--VPLSKT-----------------RMK-----------------------EEA-----
-----
WP_123051312 --MAED--D----PK-----R---PDL-----------------------------------------------------------------------------------------------------------------------------------------------------------------------------AQ-----------YQ----VETW-------LA-
--DA-----ARRV------K----QIQ-A-------------VTHSLKPVHPD---A----------------------R---G-T--------NL-----Y-----V---D-------P-SAL---AAS--P---Q-------------LSS--H--VL-------HDN------F--A-SD-V-VG----N----AA-----A--L------DVYKFL-R------L---E--L--D------------GLSI------------LDRLQ-ADD-P---AVLAAF---C---S----DL-----EQAREWRDAFLGL--L---QD--NG-K-SMSSHVLAKQLYWLV----------GE-----
DC----------RDDAQ---YHLLAPLFPTSLVHAV-LQ-----E---D--R--F-G-------------------EQ-IKQAR-Q-----AQ-------------R--DKVMH--EH-A-VRW------YPD-LAVRKLGS-T-----KPQ----------NISQLN--SERG-------------GVNYLLSSL-P-----PRWT--S-R-SA-PSPWGV-----ESIF-----DS----LLM--Q-R--REV--RQQLGGLRRFL-----------Q-------S--Q---------------PKP-------I-----QE---TR--DR---VD-------
DYLARLIDELVSFAAELR-----V-----A-W-P-----A---H----WT-------A-----Q---S---G------CL---------LAR----EEQLWL-----D------------IER--VRQEEVFRT-----EADQQFRA--EWY--WMD-WP-DQIGQRFARWIN-----GR-----L--D---------GY-----------L-P-----------T----GDVEY--------RHWKK-E---------LL--ID-Q-TYWADQLRTLR--R-----Q--ASLSKT-------------------SMK----------------------EPV-----
WP_043680632 --MSKAQDLA---------RQI--------------HHIR--EVIA--A-FLQ-DR-----L---EAK---------------------------------------------------------------------------L----------S-------K---------L----------PT------D---DP----------KR----ED-------------------LI----------------------------AQ----------------
---YQ---PAAW------LE---DA-----ARRV--------R----QIQ-A----------------VTHALKAVHPD---A----------------------R---G-T--------NL-----F---V-----E-----------P-SGM---TPH--A---V-----------VGS--H--VL-------G-A------G--F--A-SD-V-VG----N----AA-----A--L------DVYKFL-R------L---E--V--D---------------GRAL---------LDWLS-EDD-P---AAVQAL---S---D----DE-----TKAREWQSAFVGL--V---QA--RD-E-
DVASHALAKQIYWLA-------GE-----DA-------ADDAQ---YHLLAPLYATSLAHAVYSV-LQ-----E---G--R--F-G-------------------EA-NKAAR-E-----AR------------R--EGREH--EG-V-FRD-----YPA-LAVRKLGG-T-----KPQ----------NISQLN--SERG-------------GVSYLLGSL-P-----PQWK--S-R-L-SVY-RQFWGV-----QSVF-----GP---VLM-R-Y--GEV--RATRALLRFL----------R------K------D-----------------PPS--
-------N----ME---TR--NR-----VD---------AYVDRLIDELVAMASEIR---R------S-L-P-----T---G-----WT-------A-----D---P---R------CR---------LVR----EEWFWL-----D------------PWR--AE--E------------DSEFRS--EWL--FMD-WP-AQIGHRFGNWLN-----GR-----L--E---------EH----------F-K----------V----GEIEH--------RQWKK-E---------LL--LD-EDEAGWARALHELR--T--------DLDAPHYIP------------------TRE--------
---------GTV--------------
WP_136406613 --MSA---------EQI--------------IHVR--SAIH--T-FLR-DR------L---ADK-------------------------------------------------------------------------L---------S---------K---------L----------AE-----D---DP---------KR----IE---------------------LE-------------------------------EQ--------------------
FQ---PGSW------LQ---DA-----ARRV--------K----QIQ-A-----------VTHSLKPIHPD---A-----------------------R---G-T--------NL-----Y-----V---P------------P-QQL----PAH--A---E-------------VGS--H--VL-------N-E------G--F--A-SD-V-VG----N----AA-----A--L------DVYKFL-K--------I----A--INPE-----------GSDL--------------LDALV-AGD-A---HAIAAL---S---D----D----EAAQQWRDALVGL--V---GT--AS-A-
NVASHQRAKQLYWMV-------GS-----DA-------SANED---YHLLAPLYASSLAHAVYQD-VQ-----E---V--R--F-G-------------------EA-NKAAR-A-----AR-------------R--EGKAH--EG-V-FRD-----YPQ-LAVQKLGG-T-----KPQ----------NISQLN--SERG-------------GSNYLLGAL-P-----PTWR--S-R-T-REPWGL-----HSVI-----DH----MLL--Q-R--DGV--KPGIKAFLDFL----------L-------T-----D-------------------PPA--
-------N----AA---TR--NR-----VD-------AHVASLIDELVLMAAELQ--R------D-W-T------SYGTNWS-------A-----D---P---K------CR--------LAP----AEQLWL-----N------------PQR--TE--S---------------DADFNA--QWQ--LMD-WP-AQIGHRFGNWLN-----AQ-----L--A---------GK----------L-P------------V-----GEAEH----------RQWKK-E----------LL--LD-ESDDGWAQLHDVR--T-----ELKAPHYIH--------------------SRK--------
-----------GKQA------------
WP_046740887 --MPT---------EQT-------------LRVR--SAIQ--T-FLH-DR------L---QDK-------------------------------------------------------------------------L---------S---------K---------L----------AI-----D---DP---------KR----VA-------------------LE-------------------------------AQ--------------------
FQ---PATW------LQ---DA-----ARRV--------K----QIQ-A-----------VTHALKPIHPD---A------------------------R---G-T--------NL-----Y---T----D-------------P-TQL---PQH--E---L-------------VGS--H--VL-------N-E------G--F--S-SD-V-VG----N----AA-----A--L------DVYKFL-K------I---D--I--D---------------GISL--------LDLLL-AGD-A---AALHAL---S---D----DA-----QQAKEWASAFISL--T---QE--RG-A-
GISSHARAKQLYWMV-------GS-----DA-------SADED---YHLLAPLYATSLAHAVYQD-IQ-----E---A--R--F-G-------------------EA-NKQAR-A-----AR--------------R--EGKMH--DG-V-FRD-----YPQ-LAVQKLGG-T-----KPQ----------NISQLN--SERG-------------GNNYLLSAL-P-----PSWR--S-R-L-RAPWGI-----RSVI-----DE----VLL--H-R--DGV--RAGIQALLTFL-----------A---------T------D-------------------PPA--
-------N----TE---TR--RQ-----VD-------EHVAGLIDDLVLMAAELQ--R------E-W-A-----A---AYGETWT-------A-----D---A---R------CK--------LVA----AEQLWL-----D------------PQR--AQ--H---------------DEEFNA--QWQ--AMD-WP-AQIGHRFGNWLN-----AQ-----L--A---------GK----------L-P------------V----GEAEH----------RQWQK-E----------LL--LD-EGEGGWASTLHSAR--T-----ANAAPQYIP---------------------SRK--------
-------------GNAP-----------
WP_122228371 --MST---------GQT-------------RSVR--SAIH--T-FLH-DR------L---QDK-------------------------------------------------------------------------L---------S---------K---------L----------AD-----D---DP---------KR----FD-------------------LK-------------------------------EQ--------------------
-FQ---PAVW------LE---DA-----ARRV--------K----QIQ-A-----------VTHSLKPIHPD---A------------------------R---G-T--------NL-----Y---V-----E-------------P-AQL---PEH--A---E-------------VGS--H--SL-------G-E------K--F--A-SD-V-VG----N----AA-----A--L------DVYKFL-K------I---D--V--D---------------GESL-----------LELLL-NGD-D---ATLHAL---S---D----DL-----TKARSWRDAFISL--T---QE--RS-A-
NVSSHPRAKQLYWMV-------GS-----DA-------SANED---YHLLAPLYATSLAHAVYQD-VQ-----D---V--R--F-G-------------------EA-NKEAR-S-----AR-------------R--ESKGH--EG-V-FRD-----YPQ-LAVQKLGG-T-----KPQ----------NISQLN--SERG-------------GNNYLLASL-P-----PSWR--S-R-V-REPWGI-----CSVI-----DE----VLL--H-R--DGV--KPGIKAFLDFL----------L-------T-----D-------------------PPA--
-------N----AD---TR--NR-----VD--------GYVAALIDELVLMGAELQ--S------D-W-A------AYGATWS-------A-----D---T---R------CK--------LVS----AEQLWL-----D------------PQR--AE--K---------------DAFNT--LWQ--AMD-WP-AQIGHRFGNWLN-----GQ-----L--E---------GK----------L-P------------V----GEAEH----------RQWKK-E----------LL--LD-ESEDGWAQALHDAR--T-----ANAAPHTIP--------------------SRK--------
-------------GTAP--------
WP_073355685 --MEQHPDS---------GRT---------------QRVR--AAIQ--A-FLQ-ER-----L---QAK--------------------------------------------------------------------------------------------------------KD-----N---KCL---------------EQ-------------------WI-------------------------------------------------
-------FQ----PAAW------LA----DA-----ARRA--------S----QIQ-A---------------VTHSLKPIHPD---A-----------------------R---G-T--------NL-----Y---K------R--------P-TEL---AHH--A---E-------------VGS--H--LL-------R-E------H--F--S-SD-V-VG----N----AA-----A--L------DVYKFL-K------Q--V--D---------------GKSL---------LEWLL-AGD-A---SAVQAL---S---D----DP-----ETAQQWQAAFVGL--T---QP--RD-G-
AIASHQRAKQLYWLI-------GN-----DP-------TDNAQ---YHLLAPLYATSLAHSVYTT-LQ-----E---D--R--F-G-------------------DD-NKAAR-T-----AR-------------K--GESH--EG-V-LRE-----YPD-LAVQKLGG-T-----KPQ----------NISQLN--SERG-------------GSNYLLASL-P-----PQWQ--S-R-AM-RQPWGI-----ASVF-----DH----MLM--A-R--EGV--RPTLAAFLRFL----------K-------S---N-------------------PPT--
------N----AD---TR--NR-----VD-------GYVAALIDELVSTLVTLAGELQ--R------G-W-P-----D---G------WT-------A-----D---A---R------CK--------LVQ----AEQLWL-----D------------PRR--AE--T---------------DETFRT--EWL--AMD-WP-AQIGHRFGNWLN-----TQ-----L--E---------GQ----------L-R----------V----GDAEL--------REWKK-E---------LL--LD-ESDDGWAQSLHRLR--S-----SQDAPQYIP--------------------ARQ--------
-------------GVVA-------
WP_127775010 --MSASDEA---------GRT---------------QRTR--AVIQ--A-FLQ-ER-----L---QAK-------------------------------------------------------------------------L---------E---------K---------L-----------AE-----D---DP----------KR----SE----------------------WI-----------------------------AQ---------------
------FQ---AERW------LE---DA-----ARRV--------R----QIQ-A-------------VTHALKPIHPD---A-----------------------R---G-T--------NL-----Y----V-----E-----------P-ADL---PAH--A---L-------------VGS--H--LL-------R-P------D--F--S-AD-V-VG----N----AA-----A--L------DVYKFL-K------L---L--V--D---------------GQTI-----------LTWLL-ADD-A---AVIAAL---H---S----DP-----AQAAQWQSAFIGL--T---QA--RE-G-
GLSSHVRAKQLYWLV-------GS-----DA-------TDDGQ---YHLLAPLYATSLAHVVFAT-IQ-----E---D--R--F-G-------------------EN-NKAAR-A-----AR---------------R--EGRVH--DG-V-LHE-----YPG-LAVQKMGG-T-----KPQ----------NISQLN--SERG-------------GNNYLLASL-P-----PQWQ--S-G-KL-RQPWGL-----QSVF-----DH----MLL--A-R--EGV--GPTVKAFLKFL---------------K-------S------D---------------------PPP-
-------N----VE---TR--DR---VE-------TYVASLIDEMVTLAGELQ-R-----G-W-P-----T---G-----WT-------A-----D---A---R------CK--------LAL----PEQLWL-----D------------PKR--AE--N---------------DESFRS--LWL--FMD-WP-AQIGQRFGNWLN-----TH-----L--S---------KE----------F-R----------V----GDIEA--------REWKK-E---------LL--VD-ESTDGWAQRLHKLR--E-----EQNVPRYIP---------------------TRA--------
-------------GGQA------------
WP_134838777 --MSASDEA---------GRT---------------QRTR--AVIQ--A-FLQ-ER-----L---QAK-------------------------------------------------------------------------L---------E---------K---------L-----------AE-----D---DP----------KR----SE----------------------WI-----------------------------AQ---------------
------FQ---AERW------LE---DA-----ARRV--------R----QIQ-A-------------VTHALKPIHPD---A-----------------------R---G-T--------NL-----Y----V-----E-----------P-ALL---PAH--A---L-------------VGS--H--LL-------R-P------D--F--S-AD-V-VG----N----AA-----A--L------DVYKFL-K------L---L--V--D---------------GQTI-----------LTWLL-ADD-A---AVIAAL---H---S----DP-----AQAAQWQSAFIGL--T---QA--RE-G-
GLSSHVRAKQLYWLV-------GS-----DA-------TDDGQ---YHLLAPLYATSLAHVVFAT-IQ-----E---D--R--F-G-------------------EN-NKAAR-A-----AR---------------R--EGRVH--DG-V-LHE-----YPG-LAVQKMGG-T-----KPQ----------NISQLN--SERG-------------GNNYLLASL-P-----PQWQ--S-G-KL-RQPWGL-----QSVF-----DH----MLL--A-R--EGV--GPTVKAFLKFL---------------K-------S------D---------------------PPS-
-------N----VE---TR--DR---VE-------TYVASLIDEMVTLAGELQ-R-----G-W-P-----T---G-----WT-------A-----D---A---R------CK--------LAL----PEQLWL-----D------------PKR--AE--N---------------DESFRS--LWL--FMD-WP-AQIGQRFGNWLN-----TH-----L--S---------KE----------F-R----------V----GDIEA--------REWKK-E---------LL--VD-ESTDGWAQRLHKLR--E-----EQNVPRYIP---------------------TRA--------
-------------GGQA------------
WP_010106219 --MAETDA---------PAY----------------SSFQ--AAVQ--A-FIQ-ER------L---KAK-------------------------------------------------------------------------L----------D---------K---------L----------PE------D---DP----------ER----ER-------------------LI----------------------------AR--------------
---YQ---PAAW------IE---DA-----AQRV--------R----QIQ-A-------------VTHSLKPIHPD---A----------------------R---G-T--------NL-----Y---V-----E-----------P-SSL--PTL--S---E-------------LGS--H--AL-------G-N------A--F--A-FD-V-VG----N----AA-----A--L------DVYKFL-R------L---E--V--N--------------GRSL-----------LTALQ-EED-A---SALRAL---H---G----DA----TQAKTLRNALLRL--T---QP--RS-D-
RTSSHTRAKQLYWLT-------GS-----DA-------CADDS---YELLAPLFATSLAHGVHAQ-VQ-----E---A--R--F-G-------------------EA-NKAAR-Q-----AR-------------R--ERRAH--DG-V-FVD-----YPA-LAVQKLGG-T-----KPL----------NISQLN--SERG-------------GVNYLLASL-P-----PIWQ--P-R-RQ-RLPVNT-----DSVF-----KH---LYP--E-R--PDV--RNTVRALRFL-----------E-------T------D-----------------------PRP--
-------N----QA---TR--ER-----IA-------ADIEKLVDELVMATKLQ--Q------S-L-A-----P---G-----WT-------R-----D---E---R------HK--------LAV----EEQLWL-----D------------PLR--AELPG-------------EDDFAK--EWR--WMD-WP-NTIGRFFANWLN-----AK-----L--A---------AK----------L-P-----------V----GDVEA--------REWQR-T---------LL--VK-D--DGWRETLAEQR--R-----RIDGQ---------------
--------RDSR-------
WP_011809225 --MAKTDT---------PSH----------------DSFK--AAVQ--A-FLQ-ER-----L---EAK-------------------------------------------------------------------------L----------E---------K---------L----------KD------D---DP----------KR----DE-------------------LE----------------------------NK----------------
---YR---PTVW------IE---DA-----ARRV--------Q----QIQ-A-------------VTHSLKPIHPD---A-----------------------R---G-T--------NL-----Y---V------E-----------P-DSL--PML--T---E-------------LGS--H--AL-------G-E------K--F--A-PD-V-VG----N----AA-----A--L------DVYKFL-K------L---Q--V--G---------------ERSL---------LTAQ-EED-P---AAMQVL--H---G----DA----AQAKALRDKLVGL--T---QP--GS-G-
GASSHSRAKQLYWLT-------GS-----DA-------AADD----YELLAPLYATSLAHRVYAD-VQ-----E---V--R--F-G-------------------EA-NKAAR-Q-----AR-------------R--ERKAH--DG-V-LVD-----YPG-LAMQKMGG-T-----KPQ----------NISQLN--SERG-------------GVNYLLASL-P-----PIWK--V-T-ER-RLPVNT-----DSVF-----KH---LYP--Q-R--PDV--QKTVRELRRFL-----------E--------T------D-----------------------PEP--
-------N----LA---TR--ER-----RA-------AYVETLVNELLNLAGELQ--Q------G-L-P-----A---G-----WT-------L-----ND--K---RF----CQ--------LAP----VEQLWL-----D------------PRR--ATLPD-------------EDDFAK--EWL--WQD-WP-AAIAKRFGNWLN-----DK-----L--A---------GR-----------L-P----------V----GDVEA--------REWQR-A---------LI--GE-D--DGWREALTEQR--K-----LDAPHYIP---------------
--------RGQ-------
WP_011803947 --MESHRETT-------ERC--------------QRIR--LAMH--E-HVA-HR-----L---AHA----------------------------------------------------------------------------------------------------------QG------K-D--DA--AAKQ---RR----IE--------------------AQ-------------------------------EK------------
---YQ---LVHI------LE---RG-----VRTV--------E----QIA-V-----------ASHIAKGVHPDLPVK------------------------K--T-T------NL---A----V------D-------F-SSL--PMV--D---V------------VGS--H--VL------S-KRDGLHD--T--T-GN-G-SY----N----SA-----A--Y-----ELYLLL-E-----C---R--V--E---------------GQKL-----------IDLLR-LED-P---DATEAV---N---S----S-----AQLDNAAALCIQL--L---EP--KC-T-
AVSANTLSKQIYWLT-------GP-----DA-------SDDTH---YTLLAPLYATAHVHAQ-VQ-----E---A--R--F-G-------------------EA-NKAAR-Q-----AR-------------R--ERKMH--DG-V-FHD-----YPG-LAVQNMGG-T-----KPQ----------NISQLN--SERR-------------GVNYLLSSL-P-----PQWQ--A-S-AV-RLPVHA-----TSVF-----DR---LFI--A-R--PEV--RRTVRALRVFL-----------E-------S------N----------------------PDA--
-------N----LA---TR--ER-----RE-------ELLDALVDEIVSLAAELQ--Q------M-L-P-----P---G-----WS-------R-----D---Q---R------FK--------LAY----EEALWL-----D------------PLR--AEIPE-------------EAQFAH--DWQ--YMD-WP-AAIGKAFAGWLN-----AK-----L--A---------DK-----------L-P----------V----GDAEA--------REWKK-V---------LL--TD-E--DGFKQQLRELR--N-----KLDAPHYIP-----------------IRKT--------
HAELV--ALR-EENP--------
WP_052952254 --MESHR-ETT-ERC-----------QRIR--LAMH--E-HVA-HR-----L---AHA----------------------------------------------------------------------------------------------------------R----K---L----QG-----KD-------------------------------------------------------------------------A-AKQ-RR------------
AQEK----------YQ---LVHI------LE---RG-----VRTV--------E----QIA-V-----------ASHIAKGVHPDLPVK------------------------K--T-T------NL---A----V------D-------F-SSL--PMV--D---V------------VGS--H--VL------S-KRDGLHD--T-TG-----N-SSYNSA-----A--Y----ELYLL-E-----C---R--V--E---------------GQKL-----------IDLLR-LED-P---DATEAV---N---S----S-----AQLDNAAALCIQL--L---EP--KC-T-
AVSANTLSKQIYWLT-------GP-----DA-------SDDTH---YTLLAPLYATAHVHAQ-VQ-----E---A--R--F-G-------------------EA-NKAAR-Q-----AR-------------R--ERKMH--DG-V-FHD-----YPG-LAVQNMGG-T-----KPQ----------NISQLN--SERR-------------GMNYLLSSL-P-----PQWQ--A-S-AV-RMPVHA-----TSVF-----DR---LFI--A-R--PEV--RRTVRALRVFL-----------E-------S------N----------------------PDA--
-------N----LA---TR--ER-----RE-------ELLDALVDEIVSLAAELQ--Q------I-L-P-----P---G-----WS-------R-----D---Q---R------FK--------LAY----EEALWL-----D------------PLR--AEIPE-------------EAQFAH--DWQ--YMD-WP-AVVGQRFAGWLN-----AK-----L--A---------DK-----------L-P----------V----GDAEA--------RAWQK-E---------LL--TD-E--DGFQQQLRTLR--Q-----RLD---------------------------RT-------
ATEVM---------P-----------
WP_103271586 --MSDVHVT---------RRK---------------ERFR--AAID--Q-FLL-ER-----L---QSK-------------------------------------------------------------------------L----------L---------K---------L----------GP------D---DS----------QR----EE-------------------II----------------------------AQ---------------
---YA---RDTW------LL---SA-----AKRV--------E----QIQ-A-------------VTHSLKPVHPD---A-----------------------R---G-T--------NL-----Y---V----D-----------P-RGL---PPL--K---E-------------LGS--H--VL-------G-G------S--F--V-GD-V-VG----N----AA-----A--L------DVYKFL-K------L---E--V--N---------------GISL-----------LTALL-TND-E---AARLAL---H---D----NP-----AQAQSLSDAFVGL--T---QP--RA-G-
APSSHTRAKQLYWLA-------AD-----DP-------CCDDAH---YHLLAPLYATSLVHELYAQ-VQ-----E---E--R--F-G-------------------EA-NKQAR-E-----AR-------------R--ERKPH--DG-V-FLD-----YPA-LAVQKLGG-T-----KPQ----------NISQLN--SERG-------------GVNYLLASL-P-----PMWS--R-T-KL-WQPAHV-----ESVF-----EK---LYI--V-R--PEV--RRAIRDLRAFL-----------E-------T------E----------------------PAP--
-------N----LQ---TR--QR-----RD-------AYIDVVVDEIVTMAGEFL--Q------T-F-P-----A---G-----WT-------R-----D---A---E------RF--------QK--------LAR----EEQLWL-----D------------PLR--GELPE-------------ESKFAR--EWL--LMT-WP-NEIGRRFANWLN-----RY-----L--S---------GG-----------L-P----------V----GDVEA--------RAWKK-E---------LL--AD-E--DGFVQQLRTLR--Q-----QLDASDRIS----------------------RQD--------
GDSPT--AGR-GDAW-----
WP_091451669 --MPLT------------EQR-------------LLWH--AVIA--A-FIA-ER-----L---DAK-------------------------------------------------------------------------L----------D---------K---------L----------KD------D---DP----------KR----HE-------------------LL----------------------------AQ---------------
HQ---YGAW------IE---NA-----AKRV--------E----QIQ-A---------------VTHSLKPIHPD---A-----------------------R---G-T--------NL-----Y---L------P-----------P-GKL---PTQ--E---L-------------LGS--H--AL-------G-S------A--F--A-SD-A-VG----N----AA-----A--L------DVYKFL-R------L---E--L--E---------------NSSL-----------LAALL-AED-E---CATQAL---H---D----NP-----MQAKALRDAFVAL--A---QP--RT-T-
GPSSHPLAKQLYWLT-------DD-----DP-------CVDSH---YALLAPLYATSLAHAVYTQ-IQ-----E---H--R--F-G-------------------EA-NKQAR-Q-----AC------------H--AHKAH--AG-V-FHD-----YSG-LAVQKMGG-T-----KPQ----------NISQLN--SERG-------------GTNYLLASL-P-----PQWQ--A-S-AA-RLPVHN-----ESVF-----DR---VFP--A-R--PEV--RKTLRQLRQFL------------E--------T------D-----------------------PTP--
-------N----QE---TR--QR-----RD-------AYLDTLLDELVSLSGELV--Q------T-M-P-----A---G-----WT-------S-----P---T---V---DSE---K---RF------QK--------LVR----AEQLWL-----D------------PLR--AELPP-------------EADFAQ--EWL--QMD-WP-AEVGKRFANWLN-----DE-----L--K---------GQ-----------W-P----------V----GDAEA--------REWKK-A---------LL--TD-E--DGFKQQLRELR--E-----RLDAPHYIP---------------------------------
IRPT---------HEKLIEPMQQRGEMP--------
WP_119110051 --MNFRFSPEGDCVKDESTPP--------AGH-------ASYR--AAIA--G-FLQ-ER-----L---QAK-------------------------------------------------------------------------L----------D---------K---------L----------KP-------D---DP----------KR----D---------------------LI----------------------------AS---------------
--------FA---YDVW------LA---SA-----AKRV--------E----QIQ-A-------------VTHSLKPIHPD---A-----------------------R---G-T-------NL-----Y---V-----E-----------P-ASL--SAL--A---E-------------LGS--H--AL-------G-E------Q--F--V-GD-V-VG----N----AA-----A--L------DVYKFL-K------L---E--V--H---------------GRSL------------LTALL-AQD-A---DALVAL---H---T----DP-----AQARALRNAFVSL--T---QP--RA-E-
GPSSHTLAKQLYWLT-------GS-----DA-------CADDS---YHLLAPLYATSLAHAVHAQ-IQ-----E---A--R--F-G-------------------EA-NKAAR-L-----AR-------------R--EKKAH--DG-V-FHD-----YPG-LAVQNMGG-T-----KPQ----------NISQLN--SERR-------------GMNYLLSSL-P-----PQWK--V-S-EK-RLPVST-----DSVF-----SH---YFG--G-R--PLV--RQTVQALRFL-----------E-------S-----K----------------------PEA--
-------N----AA---TR--SQ-----RD-------DMVYRLVDEMVAMAADLE--H------L-L-T-----P---G-----WT-------L-----SD--E---RF-----SK--------LNS----EEQLWL-----D------------PLR--AELPE-------------EADFAE--AWL--AME-WP-GAIGRRFANWLN-----DR-----L--D---------GK----------L-H----------F----GEIEF--------RAWKK-E---------LL--AD-E--DGFAQQLRERR--N-----RIRT-------------------------
-ARK-EAMP----------
WP_066471425 --MAEIEDA---------PVY---------------DSFK--AAAH--A-FIG-ER-----L---RAK-------------------------------------------------------------------------L----------D---------K---------L----------AD------D---DP----------KR----QE-------------------LI----------------------------EQ---------------
---HM---PDVW------LE---DA-----ARRV--------E----QIQ-V-------------VTHSLKPIHPD---A-----------------------R---G-T--------NL-----Y---V-----E-----------P-GQL---PVL--C---E-------------LGS--H--AL-------G-E------E--F--A-GD-V-VG----N----AA-----A--L------DVYKFL-K------L---Q--V--G---------------GRSL-----------LSALL-AEE-A---DALQAL---H---D----DP-----GQAQALREALVSL--T---AP--RA-G-
GPSSHGRAKQLYWLT-------GD-----DA-------CDDAQ---YELLAPLYATSLAHAVYGE-IQ-----E---H--R--F-G-------------------ES-NKAAR-L-----AR-------------R--DRKSH--DG-V-FHD-----YPG-LAAQKMGG-T-----KPQ----------NISQLN--SERG-------------GVNYLLSSL-P-----PAWT--P-S-AV-RLPVHA-----ESVF-----DR---MFI--A-R--PEV--RRTVRALRVFL-----------E-------T--------------------------TQS-----
-------N----AA---TR--DT-----RK-------TYTDTLLDEVLAFANELQ--Q------L-V-Q-----P---G-----WS-------Q-----D---E---R------FQ--------QAR----QEQLWL-----D------------PLR--AELPE-------------EAQFAS--DWM--DMD-WP-AEIGMRFANWLN-----AK-----L--A---------GK----------L-P----------V----GDAEA--------REWRK-E---------LL--TD-E--DGFKQQLRELR--E-----KLDAPHYIP---------------------IRKT--------
HDGLA--A--GGDE-----
WP_092839385 --MISTDPS---------K-----------------TLWP--SVIS--A-FIH-ER-----L---RAR-------------------------------------------------------------------------L----------D---------K---------L----------RP------D---DP----------RR----PE-------------------VL----------------------------AQ---------------
-HV---PGTW------LA---DA-----AWRV--------A----QIQ-A-------------VTHSLKPIHPD---A-----------------------R---G-T--------NL-----Y---V-----E-----------P-SAL---PVL--A---E-------------LGS--H--AL-------V-G------H--F--A-TD-V-VG----N----AA-----A--L------DVYKLL-K------L---E--V--N---------------GRSL-----------LTALL-ADD-A---DALTAL---H---D----DA-----ARARALRDAFVSL--T---QP--RS-E-
VPSSHTLAKQLYWLV-------GS-----DA-------CDDAH---YHLLAPLYATSLAHAVHAQ-VQ-----E---A--R--F-G-------------------EA-NKAAR-Q-----AR-------------R--ERKVH--DG-V-FHD-----YPG-LAVQNMGG-T-----KPQ----------NISQLN--SERR-------------GVNYLLSSL-P-----PQWQ--A-S-AV-RLPVHA-----ASVF-----DR---LFL--A-R--PEV--RRTVRALRVFL-----------E-------A------D----------------------PDA--
-------N----RA---TR--ER-----RE-------SMVDMLLGELVALAAELQ--Q------L-L-P-----P---G-----WS-------C-----D---A---R------FQ--------LHR----SEQLWL-----D------------PLR--AERPD-------------ETDFAH--EWL--RMD-WP-AAIGLRFANWLN-----AK-----L--A---------GQ-----------L-P----------V----GDTEA--------RAWQK-E---------LL--TD-E--DGFHQQLRELR--Q-----RLDR-------------------------------
--SAT--EVSP--------
WP_019375094 --MWK--EAIE--A-FIS-ER-----L---HAK-------------------------------------------------------------------------L----------E---------K---------L-----------AQ-------E---D--DP------HE--------------------LI--------------------------------------------------------------------------------------HQ--
PSVW-----IE---DA-----ARRA--------S----QIQ-A-------------VTHCLKPIHPD---A-----------------------R---G-T--------NL-----Y---V------E-----------P-VHL--PPL--D---E-------------LGT--H--AL-------G-E------D--F--A-SD-V-VG----N----AA-----A--L------DVYKFL-K------L---E--V--N---------------GRSL------------LTALL-AQD-L---DALAAL---H---T----DP-----ARAQALREAFVSL--T---QP--RA-E-GPSSHALAKQLYWLT-
--GD-----DA-------CADGD---YHLLAPLYATSLAHAVYAQ-VQ-----E---A--R--F-G-------------------EA-NKAAR-Q-----AR-------------R--ERKAH--DG-V-YRD-----YPG-LAVQKMGG-T-----KPQ----------NISQLN--SERG-------------GVNYLLSSL-P-----PKWM--P-S-TQ-RLPVQA-----ESVF-----DR---LFI--A-R--PEV--RRTVSALRFL-----------E-------S------E----------------------PEP--
RE-------AFVDTLLDELVSMAAEIQ--Q--S-P-Q-----A---G-----WT-------R-----DH--E---RF-----EA---------LGV--DEKLWL-----D-------------PLR--TELPE-------------ETDFAR--QWL--QMD-WP-AEIGRRFANWLN-----DQ-----L--E---------GR-----------L-G----------V----GDVEF--------REWKK-E---------LL--TE-E--DGFKQQLRELR--D-----KLNAPQYIP-------------------SRKT-------SAELV--AEE-ETDT----
-----------------
WP_103045038 --MASSLEI---------E-----------------ISWK--DVIA--T-FIN-ER-----L---KAK-------------------------------------------------------------------------L----------E---------K---------L----------KD------D---DP----------KR----PE-------------------LL----------------------------AQ---------------
HD---PVTW------IA---DA-----SRRV--------T----QIQ-A-------------VTHSLKPIHPD---A-----------------------R---G-T--------NL-----Y---V-----E-----------P-ASQ---PVL--T---E-------------LGS--H--AL-------G-E------Q--F--V-GD-V-VG----N----AA-----A--L------DVYKFL-K------L---E--V--N---------------GRSL------------LTALL-AQD-A---DALAAL---H---T----DP-----AQARALRDAFVSL--T---QP--RA-G-
EPSSHTLAKQLYWLT-------GD-----DA-------CADAD---YTLLAPLYATSLAHAVHAQ-VQ-----E---D--R--F-G-------------------DT-NKAAR-Q-----AR-------------R--ERKAH--DG-V-FHH-----YPN-LAVQKMGG-T-----KPQ----------NISQLN--SERR-------------GMNYLLSSL-P-----PQWR--T-S-AV-RLPVHA-----TSVF-----DR---LFI--A-R--PEV--RRTVQMLRFL-----------E-------S------D------------------------PEP--
-------N----LA---TR--QR-----RE-------ALLDTLLDEVLSLAAELQ--Q------L-L-P-----P---G-----WS-------R-----D---E---R------FQ--------LSA----ERQLWL-----D------------PLR--AELPE-------------EFEFAG--EWL--QMD-WP-VEIGKAFANWLN-----AQ-----L--R---------GK-----------L-P----------V----GDAEA--------REWKK-V---------LL--TD-E--DGFKQQLCELR--D-----KLDAPHYIP---------------------IRKT--------
HAELA--ALR-EENP--------
WP_103035998 --MASSLEI---------E-----------------ISWK--DVIA--T-FIN-ER-----L---KAK-------------------------------------------------------------------------L----------E---------K---------L----------KD------D---DP----------KR----PE-------------------LL----------------------------AQ---------------
HD---PVTW------IA---DA-----SRRV--------T----QIQ-A-------------VTHSLKPIHPD---A-----------------------R---G-T--------NL-----Y---V-----E-----------P-ASQ---PVL--T---E-------------LGS--H--AL-------G-E------Q--F--V-GD-V-VG----N----AA-----A--L------DVYKFL-K------L---E--V--N---------------GRSL------------LTALL-AQD-A---DALAAL---H---T----DP-----AQARALRDAFVSL--T---QP--RA-G-
EPSSHTLAKQLYWLT-------GD-----DA-------CADAD---YTLLAPLYATSLAHAVHAQ-VQ-----E---D--R--F-G-------------------DT-NKAAR-Q-----AR-------------R--ERKAH--DG-V-FHH-----YPN-LAVQKMGG-T-----KPQ----------NISQLN--SERR-------------GMNYLLSSL-P-----PQWR--T-S-AV-RLPVHA-----TSVF-----DR---LFI--A-R--PEV--RRTVQMLRFL-----------E-------S------D------------------------PEP--
-------N----LA---TR--QR-----RE-------ALLDTLLDEVLSLAAELQ--Q------L-L-P-----P---G-----WS-------R-----D---E---R------FQ--------LDP----DQKLWL-----D------------PLR--AELPE-------------EFEFAG--EWL--QMD-WP-VEIGKAFANWLN-----AQ-----L--R---------GK-----------L-P----------V----GDAEA--------REWKK-V---------------------------------------------------------------------
-------------
WP_124455073 --MSQTPGT---------GGE---------------TGYR--AAIS--A-FLQ-ER-----L---QGK-------------------------------------------------------------------------L----------D---------K---------L----------KD------D---DP----------QR----DE-------------------VM----------------------------AS---------------
------FA---YDAW------LT---SA-----AKRV--------E----QIQ-A-------------VTHSLKPIHPE---A-----------------------R---G-T--------NL-----Y---V-----E-----------P-ATL---PSL--A---E-------------LGS--H--AL-------G-S------S--F--S-SD-V-VG----N----AA-----A--L------DVYKFL-K------L---E--V--N---------------GRSL------------LAALL-AHD-S---DALTAL---H---S----DP-----AQAQALRDALVSL--T---QP--RS-E-
GPSSHTLAKQVYWLT-------GS-----DA-------CADDS---YHLLAPLYATSLAHAVHAQ-VQ-----E---A--R--F-G-------------------ES-NKAAR-Q-----AR-------------R--ERKMH--DG-M-FHE-----YPG-LAVQNMGG-T-----KPQ----------NISQLN--SERR-------------GVNYLLSSQ-P-----PRWQ--P-S-AV-RLPRAHA-----TSVF-----DR---LFI--A-R--PEV--RRAIHSLRAFL-----------E-------S------D----------------------PEA--
-------N----LA---TR--QR-----RE-------ELLDTLMDDLVLAAAELQ--Q------L-L-P-----P---G-----WS-------R-----D---E---R------FQ--------LSR----TEQLWL-----D------------PLR--AELPE-------------EAQFAH--EWL--WMD-WP-AAIGKRFANWLN-----AK-----L--R---------DG-----------L-H----------V----GDSEA--------REWKK-E---------LL--TD-E--DGFKQQLRDLR--D-----RLDAPHYIG------------------------IRQT--------
HAELV--AQR-EGNP--------
WP_087494895 --MSEAPET---------GGK---------------AGYR--AAIS--A-FLQ-ER-----L---QAK-------------------------------------------------------------------------L----------D---------K---------L----------KP------D---DP----------QR----EE-------------------VI----------------------------AE---------------
------FA---HNVW------VA---SA-----AKRV--------E----QIQ-A-------------VTHSLKPIHPD---A-----------------------R---G-T--------NL-----Y---V-----E-----------P-RTL---PPL--A---E-------------VGS--H--AL-------G-E------R--F--V-GD-V-VG----N----AA-----A--L------DVYKFL-K------L---E--V--N---------------GRSL------------LTALL-AHD-A---DALAAL---H---S----DP-----AQAQALRDALVTL--T---QP--RA-E-
GPSSHTLAKQLYWLT-------GT-----NA-------CDDAH---YHLLAPLYATSLAHAVHAQ-VQ-----E---A--R--F-G-------------------EV-NKAAR-Q-----AR-------------R--EGKMH--DG-V-FHD-----YPG-LAVQNMGG-T-----KPQ----------NISQLN--SERR-------------GVNYLLSSL-P-----PQWK--A-S-AV-RLPVHA-----ASVF-----ER---LFI--A-R--PEV--RRAVRTLRVFL-----------E-------S------N----------------------PDT--
-------N----LA---TR--ER-----RE-------ELLDALVDEIVSLAAELQ--Q------T-L-P-----P---G-----WS-------R-----D---E---R------FQ--------LAS----EEQLWL-----D------------PLR--AEQPD-------------EAQFAH--DWL--WMD-WP-AAVGQRFANWLN-----AK-----L--A---------GK----------L-P----------V----GDAEA--------REWKK-E---------LL--TD-E--DGFKQQLRELR--Q-----RLDR-------------------------------
--TAT-GATP-------
WP_099657499 --MNDAPPT---------F-----------------SNYQ--EAIH--A-FIG-ER-----L---HAK-------------------------------------------------------------------------L----------E---------K---------L----------KA------D-E--DG----------KR----TE-------------------LI----------------------------AQ---------------
---HQ---PAAW------LD---DA-----ARRV--------E----QIQ-A-------------VTHSLKPIHPD---A-----------------------R---G-T--------NL-----Y---V-----E-----------P-RTQ---PPL--A---E-------------LGS--H--AL-------G-E------H--F--V-GD-V-VG----N----AA-----A--L------DVYKFL-K------L---E--V--N---------------GRSL------------LTALL-AQD-A---DALSAL---H---T----DP-----AQTQALRDAFVSL--T---QP--RA-E-
GPSSHTLAKQLYWLT-------DT-----DA-------CDDSH---YHLLAPLYATSLAHAVHAQ-VQ-----E---A--R--F-G-------------------EA-NKAAR-Q-----AR-------------R--ERKAH--DG-V-FHD-----YPG-LAVQNMGG-T-----KPQ----------NISQLN--SERR-------------GVNYLLSSL-P-----PQWK--V-S-AV-RLPVHA-----TSVF-----DR---LFI--A-R--PEV--RRTVRALRVFL-----------E-------S------D----------------------PEA--
-------N----LA---TR--DR-----RE-------ELLDALVDEIVSLAAELQ--Q------I-L-P-----P---G-----WS-------R-----D---E---R------FK--------LAY----EEALWL-----D------------PLR--AEIPE-------------EAQFAH--DWQ--YMN-WP-AAIGKAFAGWLN-----AK-----L--A---------DK-----------L-P----------V----GDAEA--------REWKK-E---------LL--TD-E--DGFKQQLRELR--E-----KLDAPHYIP-----------------IRKT--------
HAELV--ALR-EENP--------
WP_063459909 --MSEAPAT---------GGK---------------AGYR--AAIS--A-FLQ-ER-----L---QAK-------------------------------------------------------------------------L----------D---------K---------L----------KP------D---DP----------QR----DE-------------------VI----------------------------AE---------------
------FA---HDVW------LA---NA-----AKRV--------E----QIQ-A-------------VTHSLKPIHPD---A-----------------------R---G-T--------NF-----Y---V-----E-----------P-RTL---PSL--A---E-------------VGS--H--AL-------G-E------R--F--V-GD-V-VG----N----AA-----A--L------DVYKFL-K------L---E--V--S---------------GRSL------------LAALL-AHD-A---DALAAL---H---A----DQ-----AQAQALRDAFVSL--T---QP--RA-E-
GPSSHTLAKQLYWLT-------GS-----DA-------CDDAH---YHLLAPLYATSLAHAVHAQ-VQ-----E---A--R--F-G-------------------ES-NKAAR-Q-----AR-------------R--ERKMH--DG-V-FHD-----YPG-LAVQNMGG-T-----KPQ----------NISQLN--SERR-------------GVNYLLSSL-P-----PQWK--A-S-AV-RLPVHA-----ASVF-----ER---LFI--A-R--PEV--RRAVRALRVFL-----------E-------S------N----------------------PDA--
-------K----LA---TR--ER-----RE-------ELLDALVDEIVSLAAELQ--Q------T-L-P-----P---G-----WS-------R-----D---E---R------FK--------LIY----EEALWL-----D------------PLR--AEIPE-------------EAQFAH--DWQ--YTD-WP-AAIGKAFGNWLN-----AQ-----L--R---------GK-----------L-P----------V----GDAEA--------REWKK-E---------FL--TD-D--DGFKQQLRELR--D-----KLGTPHYIP-----------------IRKT--------
HAEPL--ALR-EENP--------
WP_034388756 --MSNAPVT---------GGK---------------AGWR--AEIG--R-FLQ-ER-----L---QAK-------------------------------------------------------------------------L----------E---------K---------L----------KP------D---DP----------QR----DE-------------------VI----------------------------AS---------------
------FA---RATW------LE---DA-----ARRV--------Q----QIQ-A-------------VTHSLKPIHPD---A-----------------------R---G-T--------NL-----Y---V-----E-----------P-AQL---PML--A---E-------------LGS--H--AL-------G-A------G--F--A-GD-V-VG----N----AA-----A--L------DVYKLL-K------L---Q--V--E---------------GRSL------------LSALL-AGD-V---DALQAL---H---E----DP-----EQAQVWRQAFIGL--T---QP--RA-G-
GPSSHGRAKQLYWLT-------GD-----DA-------CDDAQ---YELLAPLYATSLAHAVHGE-IQ-----E---H--R--F-G-------------------EA-NKAAR-I-----AR-------------R--DRKPH--DG-V-FHD-----YPG-LAVQNMGG-T-----KPQ----------NISQLN--SERG-------------GVNYLLSSL-P-----PTWT--P-S-AV-RLPVQA-----KSVF-----DR---LLI--A-R--PEV--RSTLRELRRFL-----------E-------T------D-------------------------PPAP---
-------N----LA---TR--DR-----RE-------ELLDALVDEIVSLAAELQ--Q------V-L-A-----P---G-----WT-------R-----A---E---R------TR--------LDD----NEKLWL-----D------------PLR--AELPG-------------EADFAR--RWL--RMD-WP-AEIGRRFGQWLK-----AQ-----L--A---------GK-----------L-P----------V----GDAEA--------REWKK-E---------LL--TD-E--DGFKQQLRELR--D-----RLDAPHYIP-----------------IRKT--------
HAELM--AAK-GGQP--------
WP_088446640 --MPPNLRV---------TR----------------WH--DAIA--T-FIQ-DR-----L---QSK-------------------------------------------------------------------------L----------E---------K---------L----------IE------D---DP----------KR----QE-------------------LI----------------------------AQ---------------
-HQ---YDVW------LE---DA-----ARRV--------A----QIQ-A-------------VTHSLKPIHPD---A-----------------------R---G-T--------NL-----Y---V-----E-----------P-VSL---PAL--T---E-------------LGS--H--AL-------G-E------S--F--A-FD-V-VG----N----AA-----A--L------DVYKFL-K------L---E--V--E--------------GRSL------------LTALL-EED-E---AALQAL---H---E----DP-----EQARSLREAFVSL--T---QP--RA-G-
GPSSHARAKQMYWLA-------GD-----DA-------CADGG---YELLAPLYATSLAHAVHVQ-MQ-----E---D--R--F-G-------------------EA-NKAAR-K-----AW-------------F--DRKAH--EG-V-FHD-----YPG-LAVQKMGG-T-----KPQ----------NISQLN--SERG-------------GVNYLLASL-P-----PQWR--S-N-KV-RLPVHA-----HSIF-----EK---LFI--A-R--PEV--YRGVRELRACL-----------E-------T------D--------------------------PPAL--
-------N----LA---TR--RH----RD-------ALIDGLIDEMVSLAGELQ--Q------A-L-P-----P---G-----WT-------R-----D---S---E-----RF--------QK--------LTV----EEKLWL-----D------------PLR--AEIPE-------------EEDFAG--EWL--RMD-WP-AEVGKRFANWLN-----SQ-----L--L---------GR-----------L-H----------M----GDTEA--------REWKK-E---------LL--GD-E--DGFVQQLRERR--R-----VLKSTAYIP---------------------IRRT--------
---HVERA--AQRKGDAR--------
WP_126311278 --MSNAPAT---------GRK---------------AGFR--AAID--Q-FLQ-ER-----L---QSK-------------------------------------------------------------------------L----------E---------K---------L----------KP------D---DP----------QR----DE-------------------LV----------------------------AS---------------
------FL---RDTW------VA---SA-----AKRV--------D----QIQ-A-------------VTHSLKPIHPD---A-----------------------R---G-T--------NL-----Y---V-----E-----------P-ASL---PAL--Q---E-------------LGS--H--AL-------G-Q------S--F--A-GD-V-VG----N----AA-----A--L------DVYKFL-K------V---E--V--D---------------GTSL------------LAALL-AQD-P---GALQAL---H---E----DP-----QQAMGLRDAFVGL--T---QP--RA-G-
APSSHTRAKQLYWLT-------GS-----DA-------CADEG---YELLAPLFATSLAHAVHAQ-VQ-----E---D--R--F-G-------------------ES-NKAAR-Q-----AR-------------R--ERKAH--DG-V-FHD-----YPG-LAVQKMGG-T-----KPQ----------NISQLN--SERG-------------GVNYLLASL-P-----PVWR--S-S-DV-RLPVHA-----SSVF-----DK---LFI--G-R--PEV--RSTVRKLRAFL-----------A-------T------D----------------------PEP--
-------N----LQ---TR--ER-----RE-------AFVDALIDEMVSLAGEVQ--Q------A-L-P-----P---G-----WT-------R-----D---A---R------FQ--------LVR----EEQLWL-----D------------PLR--AELRD-------------EADFAR--EWL--RMD-WP-AEVGKRFANWLN-----SQ-----L--R---------GR-----------L-P----------V----GDAEA--------RVWKK-E---------LL--TD-E--DGFVQQLRELR--K-----RIE----------------------------
-DR-EDAQ--------
WP_016451883 --MSNAPAT---------GRK---------------AGFR--AAID--Q-FLQ-ER-----L---QSK-------------------------------------------------------------------------L----------E---------K---------L----------KP------D---DP----------QR----DE-------------------LV----------------------------AS---------------
------FL---RDTW------VA---SA-----AKRV--------D----QIQ-A-------------VTHSLKPIHPD---A-----------------------R---G-T--------NL-----Y---V-----E-----------P-AGL---PVL--Q---E-------------LGS--H--AL-------G-Q------S--F--S-GD-V-VG----N----AA-----A--L------DVYKFL-K------V---E--V--D---------------GTSL------------LAALL-AQD-P---GALQAL---H---E----DP-----QQAMGLRDAFVGL--T---QP--RA-G-
IPSSHTRAKQLYWLT-------GS-----DA-------CADEG---YELLAPLFATSLAHAVHAQ-VQ-----E---D--R--F-G-------------------ES-NKAAR-Q-----AR-------------R--ERKAH--DR-V-FHD-----YPG-LAVQKMGG-T-----KPQ----------NISQLN--SERG-------------GVNYLLASL-P-----PVWR--S-S-DV-RLPVHA-----SSVF-----DK---LFI--G-R--PEV--RSTVRKLRAFL-----------A-------T------D----------------------PEP--
-------N----LQ---TR--ER-----RE-------AFVDALIDEMVSLAGEVQ--Q------A-L-P-----P---G-----WT-------R-----D---A---R------FQ--------LVR----EEQLWL-----D------------PLR--AELRD-------------EADFAR--EWL--RMD-WP-AEVGKRFANWLN-----SQ-----L--R---------GR-----------L-P----------V----GDAEA--------RVWKK-E---------LL--TD-E--DGFVQQLRELR--K-----RLDAPAYIP---------------------IRKT--------
-HDELV--MHR-GGVV--------
WP_047219650 --MSNAPAT---------GRK---------------AGFR--AAID--Q-FLQ-ER-----L---QSK-------------------------------------------------------------------------L----------E---------K---------L----------KP------D---DP----------QR----DE-------------------LV----------------------------AS---------------
------FL---RDTW------VA---SA-----AKRV--------D----QIQ-A-------------VTHSLKPIHPD---A-----------------------R---G-T--------NL-----Y---V-----E-----------P-ASL---PAL--Q---E-------------LGS--H--AL-------G-Q------S--F--A-GD-V-VG----N----AA-----A--L------DVYKFL-K------V---E--V--D---------------GTSL------------LAALL-AQD-P---GALQAL---H---E----DP-----QQAMGLRDAFVGL--T---QP--RA-G-
APSSHTRAKQLYWLT-------GS-----DA-------CADEG---YELLAPLFATSLAHAVHAQ-VQ-----E---D--R--F-G-------------------ES-NKAAR-Q-----AR-------------R--ERKAH--DG-V-FHD-----YPG-LAVQKMGG-T-----KPQ----------NISQLN--SERG-------------GVNYLLASL-P-----PVWR--S-S-DV-RLPVHA-----SSVF-----DK---LFI--G-R--PEV--RSTVRKLRAFL-----------A-------T------D----------------------PEP--
-------N----LQ---TR--ER-----RE-------AFVDALIDEMVSLAGEVQ--Q------A-L-P-----P---G-----WT-------R-----D---A---R------FQ--------LVR----EEQLWL-----D------------PLR--AELRD-------------EADFAR--EWL--RMD-WP-AEVGKRFANWLN-----SQ-----L--F---------GQ-----------L-P----------V----GDAEA--------RVWKK-E---------LL--TD-E--DGFVQQLRELR--K-----RLDAPAYIP---------------------IRKT--------
--HDELV--MHR-GGAV--------
WP_034366815 --MSNAPAT---------GRK---------------AGFR--AAID--Q-FLQ-ER-----L---QSK-------------------------------------------------------------------------L----------E---------K---------L----------KP------D---DP----------QR----DE-------------------LV----------------------------AS---------------
------FL---RDTW------VA---SA-----AKRV--------D----QIQ-A-------------VTHSLKPIHPD---A-----------------------R---G-T--------NL-----Y---V-----E-----------P-ASL---PVL--Q---E-------------LGS--H--AL-------G-Q------S--F--A-GD-V-VG----N----AA-----A--L------DVYKFL-K------V---E--V--D---------------GTSL------------LAALL-AQD-P---GALQAL---H---E----DP-----QQAMGLRDAFVGL--T---QP--RA-G-
APSSHTRAKQLYWLT-------GS-----DA-------CADEG---YELLAPLFATSLAHAVHAQ-VQ-----E---D--R--F-G-------------------ES-NKAAR-Q-----AR-------------R--ERKAH--DG-V-LHD-----YPG-LAVQKMGG-T-----KPQ----------NISQLN--SERG-------------GVNYLLASL-P-----PVWR--S-S-DV-RLPVHA-----SSVF-----DK---LFI--G-R--PEV--RSTVRKLRAFL-----------A-------T------D----------------------PEP--
-------N----LQ---TR--ER-----RE-------AFVDALIDEMVSLAGEVQ--Q------A-L-P-----P---G-----WT-------R-----D---A---R------FQ--------LVR----EEQLWL-----D------------PLR--AELRD-------------EADFAR--EWL--RMD-WP-AEVGKRFANWLN-----SQ-----L--F---------GQ-----------L-P----------V----GDAEA--------RVWKK-E---------LL--TD-E--DGFVQQLRELR--K-----RLDAPAYIP---------------------IRKT--------
--HDELV--MHR-GGAV--------
WP_013804306 --MPPTPSA---------MR----------------WH--DAIA--H-FIQ-ER-----L---QSK-------------------------------------------------------------------------L----------E---------K---------L----------AE------D---DP----------KR----QE-------------------LI----------------------------AQ---------------
-HQ---RDTW------LE---DA-----ARRV--------Q----QIQ-A-------------VTHSLKPIHPD---A-----------------------R---G-T--------NL-----Y---V-----E-----------P-ASL---PAL--Q---E-------------LGS--H--AL-------G-Q------S--F--A-GD-V-VG----N----AA-----A--L------DVYKFL-K------V---E--V--D---------------GTSL------------LTALL-AQD-P---GALQAL---H---E----DP-----QQAMGLRDAFVGL--T---QP--RA-G-
APSSHTRAKQLYWLT-------GS-----DA-------CADEG---YELLAPLFATSLAHAVHAQ-VQ-----E---D--R--F-G-------------------ES-NKAAR-Q-----AR-------------R--ERKAH--DG-V-FHD-----YPG-LAVQKMGG-T-----KPQ----------NISQLN--SERG-------------GVNYLLASL-P-----PVWR--S-S-DV-RLPVHA-----SSVF-----DK---LFV--G-R--PEV--RSTVRKLRAFL-----------A-------T------D----------------------PEP--
-------N----LQ---TR--ER-----RE-------AFMDALIDEMVSLAGEVQ--Q------A-L-P-----P---G-----WT-------R-----D---A---R------FQ--------LVR----EEQLWL-----D------------PLR--TELRD-------------EEEFAR--EWL--RMD-WP-AEVGKRFANWLN-----SQ-----L--F---------GQ-----------L-P----------V----GDAEA--------RVWKK-E---------LL--TD-E--DGFVQQLRELR--K-----RLDAPAYIP---------------------IRKT--------
--HDELV--MHR-GGVV--------
WP_043789324 --MPPTPSA---------MR----------------WH--DAIA--I-FIQ-ER-----L---QAK-------------------------------------------------------------------------L----------E---------K---------L----------AE------D---DP----------KR----QE-------------------LI----------------------------AQ---------------
-HQ---RDTW------ME---DA-----ARRV--------T----QIQ-A-------------VTHSLKPIHPD---A-----------------------R---G-T--------NL-----Y---V-----E-----------P-ASL---PAL--Q---E-------------LGS--H--AL-------G-Q------S--F--A-GD-V-VG----N----AA-----A--L------DVYKFL-K------V---E--V--D---------------GTSL------------LAALL-AQD-P---GALQAL---H---E----DP-----QQAMALRDAFVGL--T---QP--RA-G-
IPSSHTRAKQLYWLT-------GS-----DA-------CADEG---YELLAPLFATSLAHAVHAQ-VQ-----E---D--R--F-G-------------------EI-NKAAR-Q-----AR-------------R--ERKAH--DG-V-FHD-----YPG-LAVQKMGG-T-----KPQ----------NISQLN--SERG-------------GVNYLLASL-P-----PVWR--S-S-DV-RLPVHA-----SSVF-----DK---LFI--G-R--PEV--RSTVRKLRAFL-----------A-------T------D----------------------PEP--

[illegible]

WP_046240415
[illegible]

WP_012203143
[illegible]

WP_065345326
[illegible]

WP_096914931
[illegible]

WP_141830498
[illegible]

WP_074920593
[illegible]

WP_071954291
[illegible]

WP_093349097
[illegible]

WP_124893433
[illegible]

WP_025101852
[illegible]

WP_052142961
[illegible]

WP_060001803
[illegible]

WP_017923024
[illegible]

WP_042624905
[illegible]

WP_055138606
[illegible]

WP_070106580
[illegible]

WP_052449854
[illegible]

WP_063814401
[illegible]

WP_086961051
[illegible]

WP_062605007
[illegible]

WP_056364028
[illegible]

WP_086382914
[illegible]

WP_061999593
[illegible]

WP_133817427
[illegible]

WP_072895104
[illegible]

WP_073104313
[illegible]

WP_029891296
[illegible]

WP_141354701
[illegible]

WP_108975556
[illegible]

WP_002923969
[illegible]

WP_107492160
[illegible]

WP_094786950
[illegible]

WP_108414479
[illegible]

WP_089953196
[illegible]

WP_051560552
[illegible]

WP_044346389
[illegible]

WP_141477475
[illegible]

WP_106738134
[illegible]

WP_059230352
[illegible]

WP_140219922
[illegible]

WP_058794400
[illegible]

PAASHGLAKQVYFPL-------------------------PDGG---YHLLAPLFPTSLMHRAQRR-LR----E--D---R--F-G----------------AA-AKAAR-E-----AR------------K--NGQPA--TQ-G-YCE-----YLD-LAIQKFGG-T----KPQ---------NISQLN--SERY-------------GENWLLASL-P-----PTWR----SLDL-RPPLRR----DSAF----------EFL--Y-RNDKAL--RARVAELQHFL-------R--VT----A-H-----N----------------------
N----WQ---IR--RK----RA------RLLGDITDAFHQYAAELL--EANRNTV-L-A-----P---G----WS------A-----H---P---D-----CR--------LDD---SERHWL-----D-------------PLR--AHT-------------DADFRS--SHL--WRD-WP-TQASQRFGNWLN-----AK-----L--D------HD-----------T-L-------------RL---GEAEA--------DHWRG-V-----------LH------------------D------ELKLFKEML-----------------DDGR--------A----------
-------------------------

WP_116425918 ------------------------------------------MLQANA------------------------LALR--QAIQ--S-FID-ER-----L----QAK------------------------------------------------------------------------------------------------------------------------------------------------------------------------------------------------------------------------------------------------------------
-HR---REAW-----LA---DA-----ARRV-------G----QIQ-M----------------ASHTLKPIHPD---A----------------R---G-S------NL----Y----V----M--------P-ESP----KP--P--GL---------VGT--H--SL------K-G-----E--L--A-HD-V-VG----N----AA-----A-L-----DVFKFL-N------L----E--L--E-----------GRSL--------------LQRLL-DDE-P--QVLAAL---S---E-----ES-----SLAREWRDAFVSI--T---RS--KG---
KTASHTLAKQLYFPL-------AG-------------------GD---YHLLAPLFPTSLAHKMQQT-LR----A--D---R--F-G----------------EA-ARAAR-E-----AR------------A--KGEGW--PH-G-YRE-----YPN-LAIQNFGG-T----KPQ---------NISQLN--SERH-------------GENWLLASL-P----PQWE--S---PEVRLPLDT----DSVF-----PF---------LARHDGTL--RRRVRELRDFL-----------AHTSH-------------N----------
-N----LA---IR--RQ----RE------RLLGDICDRLLHYAAQVR------------E-----P--SG----WS----ARQ-----D------------CR--------LHD---AERLWL-----D-------------PLR--GRQ-------------DEIFSA--QRV--RLD-WP-QQVGQRFANWLN-----QA-----L--G------GH--------E-S---------RL---GEDEA--------RQWTH-D-----------LH-------------------K------ELQLFREEL-----------------EYDP--------S----------
-------------------

WP_064581594 ------------------------------------------MLQASA------------------------LALR--QAIQ--S-FID-ER-----L----QAK------------------------------------------------------------------------------------L---------D--------------K----------L----------RP------DD---EG----------KR---QA--------------------LI----------------------------------------------------EG----------
HR---REAW-----LA---DA-----ARRV-------G----QIQ-M----------------ASHTLKPIHPD---A----------------R---G-S------NL----Y----V----M--------P-ESP----KP--P--GL---------VGT--H--SL------K-G-----E--L--A-HD-V-VG----N----AA-----A-L-----DVFKFL-N------L----E--L--E-----------GRSL--------------LQRLL-DDE-P--QVLAAL---S---E-----ES-----SLAREWRDAFVSI--T---RS--KG---
KTASHTLAKQLYFPL--------------------------AGGD---YHLLAPLFPTSLAHKVQQT-LR----A--D---R--F-G----------------EA-ARAAR-E-----AR------------A--KGEGW--PH-G-YRE-----YPN-LAIQNFGG-T----KPQ---------NISQLN--SERH-------------GENWLLASL-P----PQWE--S---PEVRLPLDT----DSVF-----PF---------LARHDGTL--RRRVRELRDFL-----------AHTSH-------------N----------
N----LA---MR--RK----RE------RLLGDICDRLLHYAAQVR------------E-----P--SG----WS----ARQ-----D------------CR--------LHD---AERLWL-----D-------------PLR--GRQ-------------DEIFSA--QRV--RLD-WP-QQVGQRFANWLN-----QA-----L--G------GH--------E-H---------RL---GEDEA--------RQWTH-D-----------LH-------------------K------ELQLFREEL-----------------EYDP--------S----------
-------------------

WP_058487834 ------------------------------------------MLQANA------------------------LALR--QAIQ--S-FID-ER-----L----QAK------------------------------------------------------------------------------------L---------D--------------K----------L----------RP------DD---EG----------KR---QA--------------------LI----------------------------------------------------EG----------
-HR---REAW-----LA---DA-----ARRV-------G----QIQ-M----------------ASHTLKPIHPD---A----------------R---G-S------NL----Y----V----M--------P-ESP----KP--P--GL---------VGT--H--SL------K-G-----E--L--A-HD-V-VG----N----AA-----A-L-----DVFKFL-N------L----E--L--E-----------GRSL--------------LQRLL-DDE-P--QVLAAL---S---E-----ES-----SLAREWRDAFVSI--T---RS--KG---
KTASHTLAKQLYFPL--------------------------AGGD---YHLLAPLFPTSLAHKVQQT-LR----A--D---R--F-G----------------EA-ARAAR-E-----AR------------A--RGEGW--PH-G-YRE-----YPN-LAIQNFGG-A----KPQ---------NISQLN--SERH-------------GENWLLASL-P----PQWE--S---PEVRLPLDT----DSVF-----PF---------LARHDGTL--RRRVRELRDFL-----------AHTSH-------------N----------
N----LA---IR--RK----RA------RLLGDICDRLLHYAAQVR------------E-----P--SG----WS----ARQ-----D------------CR--------LHD---AERLWL-----D-------------PLR--GRQ-------------DEIFSA--QRV--RLD-WP-QQVGQRFANWLN-----QA-----L--G------GH--------E-H---------RL---GEDEA--------RQWTH-D-----------LH-------------------K------ELQLFREEL-----------------EYDP--------S----------
-----------------

WP_043269890 ------------------------------------------MLQANA------------------------LALR--QAIQ--S-FID-ER-----L----QAK------------------------------------------------------------------------------------L---------D--------------K----------L----------RP------DD---EG----------KR---QA--------------------LI----------------------------------------------------EG----------
-HR---REAW-----LA---DA-----ARRV-------G----QIQ-M----------------ASHTLKPIHPD---A----------------R---G-S------NL----Y----V----M--------P-ESP----KP--P--GL---------VGT--H--SL------K-G-----E--L--A-HD-V-VG----N----AA-----A-L-----DVFKFL-N------L----E--L--E-----------GRSL--------------LQRLL-DDE-P--QVLAAL---S---E-----ES-----SLAREWRDAFVSI--T---RS--KG---
KTASHTLAKQLYFPL--------------------------AGGD---YHLLAPLFPTSLAHKVQQT-LR----A--D---R--F-G----------------EA-ARAAR-E-----AR------------A--RGEGW--PH-G-YRE-----YPN-LAIQNFGG-A----KPQ---------NISQLN--SERH-------------GENWLLASL-P----PQWE--S---PEVRLPLDT----DSVF-----PF---------LARHDGTL--RRRVRELRDFL-----------AH--TSHN-----------N----------
-----LA---IR--RK----RE------RLLGDICDRLLHYAAQVR------------E-----P--SG----WS----ARQ-----D------------CR--------LHD---AERLWL-----D-------------PLR--GRQ-------------DEIFSA--QRV--RLD-WP-QQVGQRFANWLN-----QA-----L--GGH-------------RL---GEDEA--------RQWTH-D-----------LH-------------------K------ELQLFREEL-----------------EYDP--------S----------
-------------------

WP_126869041 ----------------------------------------------MTSPLPTP-----------T--W---------QELR--QFIE--S-FIQ-ER-----L----QGK------------------------------------------------------------------------------------L---------D--------------K----------L----------HP------DE-----DD----------KR---QT--------------------LL----------------------------------------------------AT----------
-----HR---REAW-----LA---DA-----ARRV-------G----QLQ-L---------------VTHTLKPIHPD---A----------------R---G-S------NL----H----S----L--------P-QAP-----GQ--P--GL---------AGS--H--EL-------G-D-----K--L--V-SD-V-VG----N----AA-----A-L-----DVFKFL-S------L----Q--Y--Q-----------GKNL--------------LNWLK-EDS-T--EALQAL---S---D-----NA-----EQAREWRQAFIDI--T---AV--KG---
APASHSLAKQLYFPL-------PG-------------------SD---YHLLAPLFPTSLVHHVHAL-LR----E--A---R--F-G----------------EA-ARAAR-E-----AR------------S--RQESW--PH-G-FSE-----YPN-LAIQKFGG-T----KPQ---------NISQLN--SERY-------------GENWLLPSL-P----PHWQ--R-Q-DQ-RAPIRH----SSVF-----EH---DFG--R-S--PEV--SRLTRTLQRLL--------------A-----K-----------TRH-N----------
N----FT---IR--KY----RA------QLVGQICDEALQYAARLR-------E------L-E-----P--G----WS------A----T----P---G-----CQ--------LHE---TEQLWL-----D-------------PLR--AQT-------------DETFLQ--RRL--RGD-WP-AEVGNRFANWLN-----RA-----V--S------S--D-S-Q-I---------L---GSPEA--------AQWSQ-E---------LS---------------K------ELTMFKEIL-----------------EDER--------D----------
-------------------

WP_134288395 ----------------------------------------------MTSPLPTP-----------T--W---------QELR--QFIE--S-FIQ-ER-----L----QGK------------------------------------------------------------------------------------L---------D--------------K----------L----------HP------DE-----DD----------KR---QT--------------------LL----------------------------------------------------AT----------
-----HR---REAW-----LA---DA-----ARRV-------G----QLQ-L---------------VTHTLKPIHPD---A----------------R---G-S------NL----H----S----L--------P-QAP-----GQ--P--GL---------AGS--H--EL-------G-D-----K--L--V-SD-V-VG----N----AA-----A-L-----DVFKFL-S------L----Q--Y--Q-----------GKNL--------------LNWLK-EDS-T--EALQAL---S---D-----NA-----EQAREWRQAFIGI--T---AV--KG---
APASHSLAKQLYFPL-------PG-------------------SG---YHLLAPLFPTSLVHHVHAL-LR----E--A---R--F-G----------------EA-ARAAR-E-----AR------------S--RQESW--PH-G-FSE-----YPN-LAIQKFGG-T----KPQ---------NISQLN--SERY-------------GENWLLPSL-P----PHWQ--R-Q-DQ-HAPIRH----FSVF-----EH---DFG--R-S--PEV--SRLTRTLQRLL--------------A-----K-----------TRH-N----------
N----FT---IR--KY----RA------QLVGQICDEALQYAARLR-------E------L-E-----P--G----WS------A----T----P---G-----CQ--------LHD---AEQLWL-----D-------------PLR--AQT-------------DETFLQ--RRL--RGD-WP-AEVGNRFANWLN-----RA-----V--S------S--D-S-Q-I---------L---GSTEA--------AQWSQ-E---------LS---------------K------ELTMFKEIL-----------------EDER--------D----------
-------------------

WP_033993612 ----------------------------------------------MTSPLPTP-----------T--W---------QELR--QFIE--S-FID-ER-----L----QGK------------------------------------------------------------------------------------L---------D--------------K----------L----------QP------DE-----DD----------KR---QT--------------------LL----------------------------------------------------AT----------
-----HR---REAW-----LA---DA-----ARRV-------G----QLQ-L---------------VTHTLKPIHPD---A----------------R---G-S------NL----H----S----L--------P-QAP-----GQ--P--GL---------AGS--H--EL-------G-D-----K--L--V-SD-V-VG----N----AA-----A-L-----DVFKFL-S------L----Q--Y--Q-----------GKNL--------------LNWLT-EDS-A--EALQAL---S---D-----NA-----EQAREWRQAFIGI--T---TV--KG---
APASHSLAKQLYFPL-------PG-------------------SD---YHLLAPLFPTSLVHHVHAL-LR----E--A---R--F-G----------------EA-ARAAR-E-----AR------------N--RRESW--PH-G-FSE-----YPN-LAIQKFGG-T----KPQ---------NISQLN--SERY-------------GENWLLPSL-P----PHWQ--R-Q-DQ-RAPIRH----SSVF-----EH---DFG--R-S--PEV--SRLTRTLQRLL--------------A-----K-----------TRH-N----------
N----FT---IR--RY----RA------QLVGQICDEALQYAARLR-------E------L-E-----P--G----WS------A----T----P---G-----CQ--------LHD---AEQLWL-----D-------------PLR--AQT-------------DETFLQ--RRL--RGD-WP-AEVGNRFANWLN-----RA-----V--S------S--D-S-Q-I---------L---GSPEA--------AQWSQ-E---------LS---------------K------ELTMFKEIL-----------------EDER--------D----------
-------------------

WP_128687974 ----------------------------------------------MTSPLPTP-----------T--W---------QELR--QFIE--S-FIQ-ER-----L----QGK------------------------------------------------------------------------------------L---------D--------------K----------L----------HP------DE-----DD----------KR---QT--------------------LL----------------------------------------------------AT----------
-----HR---REAW-----LA---DA-----ARRV-------G----QLQ-L---------------VTHTLKPIHPD---A----------------R---G-S------NL----H----S----L--------P-QAP-----GQ--P--GL---------AGS--H--EL-------G-D-----K--L--V-SD-V-VG----N----AA-----A-L-----DVFKFL-S------L----Q--Y--Q-----------GKNL--------------LNWLK-EDS-T--EALQAL---S---D-----NA-----EQAREWRQAFIDI--T---AV--KG---
APASHSLAKQLYFPL-------PG-------------------SD---YHLLAPLFPTSLVHHVHAL-LR----E--A---R--F-G----------------EA-ARAAR-E-----AR------------S--RQESW--PH-G-FSE-----YPN-LAIQKFGG-T----KPQ---------NISQLN--SERY-------------GENWLLPSL-P----PHWQ--R-Q-DQ-RAPIRH----SSVF-----EH---DFG--R-G--PEV--SRLTRTLQRLL--------------A-----K-----------TRD-N----------
N----FT---IR--KY----RA------QLVGQICDEALQYAARLR-------E------L-E-----P--G----WS------A----T----P---G-----CQ--------LHD---AEQLWL-----D-------------PLR--AQT-------------DETFLQ--RRL--RGD-WP-AEVGNRFANWLN-----RA-----V--S------S--D-S-Q-I---------L---GSPEA--------AQWSQ-E---------LS---------------K------ELTMFKEIL-----------------EDER--------D----------
-------------------

WP_058156381 ----------------------------------------------MTSPLPTP-----------T--W---------QELR--QFIE--S-FIQ-ER-----L----QGK------------------------------------------------------------------------------------L---------D--------------K----------L----------HP------DE-----DD----------KR---QT--------------------LL----------------------------------------------------AT----------
-----HR---REAW-----LA---DA-----ARRV-------G----QLQ-L---------------VTHTLKPIHPD---A----------------R---G-S------NL----H----S----L--------P-QAP-----GQ--P--GL---------AGS--H--EL-------G-D-----K--L--V-SD-V-VG----N----AA-----A-L-----DVFKFL-S------L----Q--Y--Q-----------GKNL--------------LNWLK-EDS-T--EALQAL---S---D-----NA-----EQAREWRQAFIGI--T---AV--KG---
APASHSLAKQLYFPL-------PG-------------------SG---YHLLAPLFPTSLVHHVHAL-LR----E--A---R--F-G----------------EA-ARAAR-E-----AR------------S--RQESW--PH-G-FSE-----YPN-LAIQKFGG-T----KPQ---------NISQLN--SERY-------------GENWLLPSL-P----PHWQ--RQ--DQ-RAPIRH----SSVF-----EH---DFG--R-S--PEV--SRLTRTLQRLL-------A--KT-----R-D-----N----------
N----FT---IR--RY----RA------QLVGQICDEALQYAARLR-------E------L-E-----P--G----WS------A----T----P---G-----CQ--------LHD---AEQLWL-----D-------------PLR--AQT-------------DETFLQ--RRL--RGD-WP-AEVGNRFANWLN-----RA-----V--S------SD-------S-Q-I---------L---GSPEA--------AQWSQ-E---------LS---------------K------ELTMFKEIL-----------------EDER--------D----------
-------------------

WP_023100320 ----------------------------------------------MTSPLPTP-----------T--W---------QELR--QFIE--S-FIQ-ER-----L----QGK------------------------------------------------------------------------------------L---------D--------------K----------L----------HP------DE-----DD----------KR---QT--------------------LL----------------------------------------------------AT----------
-----HR---REAW-----LA---DA-----ARRV-------G----QLQ-L---------------VTHTLKPIHPD---A----------------R---G-S------NL----H----S----L--------P-QAP-----GQ--P--GL---------AGS--H--EL-------G-D-----K--L--V-SD-V-VG----N----AA-----A-L-----DVFKFL-S------L----Q--Y--Q-----------GKNL--------------LNWLK-EDS-T--EALQAL---S---D-----NA-----EQAREWRQAFIGI--T---AV--KG---
APASHSLAKQLYFPL-------PG-------------------SD---YHLLAPLFPTSLVHHVHAL-LR----E--A---R--F-G----------------EA-ARAAR-E-----AR------------S--RQESW--PH-G-FSE-----YPN-LAIQKFGG-T----KPQ---------NISQLN--SERY-------------GENWLLPSL-P----PHWQ--R-Q-DQ-RAPIRH----SSVF-----EH---DFG--R-S--PEV--SRLTRTLQRLL-------A--KT-----R-H-----N----------
N----FT---IR--KY----RA------QLVGQICDEALQYAARLR-------E------L-E-----P--G----WS------A----T----P---G-----CQ--------LHD---AEQLWL-----D-------------PLR--AQT-------------DETFLQ--RRL--RGD-WP-AEVGNRFANWLN-----RA-----V--S------SD-------S-Q-I---------L---GSPEA--------AQWSQ-E---------LS---------------K------ELTMFKEIL-----------------EDER--------D----------
-------------------

WP_039700635 ----------------------------------------------MSERPNG-----------D--S---------PTLR--QLIE--S-FIQ-ER-----L----QGK------------------------------------------------------------------------------------L---------D--------------K----------L----------KP------DE-----DV----------KR---QS--------------------LL----------------------------------------------------AG----------
-----HR---REAW-----LA---DA-----ARRV-------G----QLQ-L---------------VTHTLKPIHPD---A----------------R---G-S------NL----H----S----L--------P-QAP-----GQ--P--GL---------AGS--H--EL-------G-D-----K--L--V-SD-V-VG----N----AA-----A-L-----DVFKFL-S------L----Q--Y--Q-----------GKNL--------------LNWLT-EDS-A--EAVQAL---S---D-----NA-----EQAREWRQAFIGI--T---AV--KG---
APTSHSLAKQLYFPL-------PG-------------------SG---YHLLAPLFPTSLVHHVHAL-LR----E--A---R--F-G----------------EA-ARAAR-E-----AR------------S--RQESW--PH-G-FSE-----YPN-LAIQKFGG-T----KPQ---------NISQLN--SERY-------------GENWLLPSL-P----PHWQ--R-Q-DQ-RAPIRH----SSVF-----EH---DFG--R-S--PEV--SRLTRTLQRLL--------------D-----K-----------TRH-N----------
N----FT---IR--KY----RA------QLVGQICDEALQYAARLR-------E------L-E-----P--G----WS------A----T----P---G-----CQ--------LHD---AEQLWL-----D-------------PLR--AQT-------------DETFLQ--RRL--RGD-WP-AEVGNRFANWLN-----RA-----V--S------S--D-S-Q-I---------L---GSPEA--------AQWSQ-E---------LS---------------K------ELTMFKEIL-----------------EDER--------D----------
-------------------

WP_003162917 ----------------------------------------------MSERPNG-----------D--S---------PTLR--QLIE--S-FIQ-ER-----L----QGK------------------------------------------------------------------------------------L---------D--------------K----------L----------KP------DE-----DV----------KR---QS--------------------LL----------------------------------------------------AG----------
-----HR---REAW-----LA---DA-----ARRV-------G----QLQ-L---------------VTHTLKPIHPD---A----------------R---G-S------NL----H----S----L--------P-QAP-----GQ--P--GL---------AGS--H--EL-------G-D-----K--L--V-SD-V-VG----N----AA-----A-L-----DVFKFL-S------L----Q--Y--Q-----------GKNL--------------LNWLT-EDS-A--EAVQAL---S---D-----NA-----EQAREWRQAFIGI--T---AV--KG---
APTSHSLAKQLYFPL-------PG-------------------SG---YHLLAPLFPTSLVHHVHAL-LR----E--A---R--F-G----------------EA-ARAAR-E-----AR------------S--RQESW--PH-G-FSE-----YPN-LAIQKFGG-T----KPQ---------NISQLN--SERY-------------GENWLLPSL-P----PHWQ--R-Q-DQ-RAPIRH----SSVF-----EH---DFG--R-S--PEV--SRLTRTLQRLL-------A--KT-----R-H-----N----------
N----FT---IR--KY----RA------QLVGQICDEALQYAARLR-------E------L-E-----P--G----WS------A----T----P---G-----CQ--------LHD---AEQLWL-----D-------------PLR--AQT-------------DETFLQ--RRL--RGD-WP-AEVGNRFANWLN-----RA-----V--S------S--D-S-Q-I---------L---GSPEA--------AQWSQ-E---------LS---------------K------ELTMFKEIL-----------------EDER--------D----------
-------------------

WP_061196291 ----------------------------------------------MSERPNG-----------D--S---------PTLR--QLIE--S-FIQ-ER-----L----QGK------------------------------------------------------------------------------------L---------D--------------K----------L----------KP------DE-----DV----------KR---QS--------------------LL----------------------------------------------------AG----------
-----HR---REAW-----LA---DA-----ARRV-------G----QLQ-L---------------VTHTLKPIHPD---A----------------R---G-S------NL----H----S----L--------P-QAP-----GQ--S--GL---------AGS--H--EL-------G-D-----K--L--V-SD-V-VG----N----AA-----A-L-----DVFKFL-S------L----Q--Y--Q-----------GKNL--------------LNWLT-EDS-A--EAVQAL---S---D-----NA-----EQAREWRQAFIGI--T---AV--KG---
APASHSLAKQLYFPL-------PG-------------------SG---YHLLAPLFPTSLVHHVHAL-LR----E--A---R--F-G----------------EA-ARAAR-E-----AR------------S--RQESW--PH-G-FSE-----YPN-LAIQKFGG-T----KPQ---------NISQLN--SERY-------------GENWLLPSL-P----PHWQ--R-Q-DQ-RAPIRH----SSVF-----EH---DFG--R-S--PEV--SRLTRTLQRLL--------------A-----K-----------TRH-N----------
N----FT---IR--KY----RA------QLVGQICDEALQYAARLR-------E------L-E-----P--G----WS------A----T----P---G-----CQ--------LHD---AEQLWL-----D-------------PLR--AQT-------------DETFLQ--RRL--RGD-WP-AEVGNRFANWLN-----RA-----V--S------S--D-S-Q-I---------L---GSPEA--------AQWSQ-E---------LS---------------K------ELTMFKEIL-----------------EDER--------D----------
-------------------

WP_134242446 ----------------------------------------------MSERPNG-----------D--S---------PALR--QLIE--S-FIQ-ER-----L----QGK------------------------------------------------------------------------------------L---------D--------------K----------L----------KP------DE-----DV----------KR---QS--------------------LL----------------------------------------------------AG----------
-----HR---REAW-----LA---DA-----ARRV-------G----QLQ-L---------------VTHTLKPIHPD---A----------------R---G-S------NL----H----S----L--------P-QAP-----GQ--P--GL---------AGS--H--EL-------G-D-----K--L--V-SD-V-VG----N----AA-----A-L-----DVFKFL-S------L----Q--Y--Q-----------GKNL--------------LNWLT-EDS-A--EAVQAL---S---D-----NA-----EQAREWRQAFIGI--T---AV--KG---
APASHSLAKQLYFPL-------PG-------------------SG---YHLLAPLFPTSLVHHVHAL-LR----E--A---R--F-G----------------EA-ARAAR-E-----AR------------S--RQESW--PH-G-FSE-----YPN-LAIQKFGG-T----KPQ---------NISQLN--SERY-------------GENWLLPSL-P----PHWQ--R-Q-DQ-RAPIRH----SSVF-----EH---DFG--R-S--PEV--SRLTRTLQRLL--------------A-----K-----------TRH-N----------
N----FT---IR--RY----RA------QLVGQICDEALQYAARLR-------E------L-E-----P--G----WS------A----T----P---G-----CQ--------LHD---AEQLWL-----D-------------PLR--AQT-------------DETFLQ--RRL--RGD-WP-AEVGNRFANWLN-----RA-----V--S------S--D-S-Q-I---------L---GSPEA--------AQWSQ-E---------LS---------------K------ELTMFKEIL-----------------EDER--------D----------
-------------------

WP_046688914 ----------------------------------------------MSERPNG-----------D--S---------PTLR--QLIE--S-FIQ-ER-----L----QGK------------------------------------------------------------------------------------L---------D--------------K----------L----------KP------DE-----DV----------KR---QS--------------------LL----------------------------------------------------AG----------
-----HRRXWREAW-----LA---DA-----ARRV-------G----QLQ-L---------------VTHTLKPIHPD---A----------------R---G-S------NL----H----S----L--------P-QAP-----GQ--P--GL---------AGS--H--EL-------G-D-----K--L--V-SD-V-VG----N----AA-----A-L-----DVFKFL-S------L----Q--Y--Q-----------GKNL--------------LNWLT-EDS-A--EAVQAL---S---D-----NA-----EQAREWRQAFIGI--T---AV--KG---
APASHSLAKQLYFPL-------PG-------------------SG---YHLLAPLFPTSLVHHVHGL-LR----E--A---R--F-G----------------EA-ARAAR-E-----AR------------S--RQESW--PH-G-FSE-----YPN-LAIQKFGG-T----KPQ---------NISQLN--SERY-------------GENWLLPSL-P----PHWQ--R-Q-DQ-RAPIRH----SSVF-----EH---DFG--R-S--PEV--SRLTRTLQRLL--------------A-----K-----------TRH-N----------
N----FT---IR--RY----RA------QLVGQICDEALQYAARLR-------E------L-E-----P--G----WS------A----T----P---G-----CQ--------LHD---AEQLWL-----D-------------PLR--AQT-------------DETFLQ--RRL--RGD-WP-AEVGNRFANWLN-----RA-----V--S------S--D-S-Q-I---------L---GSPEA--------AQWSQ-E---------LS---------------K------ELTMFKEIL-----------------EDER--------D----------
-------------------

WP_121410065 ----------------------------------------------MSERPNG-----------D--S---------PTLR--QLIE--S-FIQ-ER-----L----QGK------------------------------------------------------------------------------------L---------D--------------K----------L----------KP------DE-----DV----------KR---QS--------------------LL----------------------------------------------------AG----------
-----HR---REAW-----LA---DA-----ARRV-------G----QLQ-L---------------VTHTLKPIHPD---A----------------R---G-S------NL----H----S----L--------P-QAP-----GQ--P--GL---------AGS--H--EL-------G-D-----K--L--V-SD-V-VG----N----AA-----A-L-----DVFKFL-S------L----Q--Y--Q-----------GKNL--------------LNWLT-EDS-A--EAVQAL---S---D-----NA-----EQAREWRQAFIGI--T---AV--KG---
APASHSLAKQLYFPL-------PG-------------------SG---YHLLAPLFPTSLVHHVHAL-LR----E--A---R--F-G----------------EA-ARAAR-E-----AR------------S--RQESW--PH-G-FSE-----YPN-LAIQKFGG-T----KPQ---------NISQLN--SESY-------------GENWLLPSL-P----PHWQ--R-Q-DQ-RAPIRH----SSVF-----EH---DFG--R-S--PEV--SRLTRTLQRLL--------------A-----K-----------TRH-N----------
N----FT---IR--KY----RA------QLVGQICDEALQYAARLR-------E------L-E-----P--G----WS------A----T----P---G-----CQ--------LHD---AEQLWL-----D-------------PLR--AQT-------------DETFLQ--RRL--RGD-WP-AEVGNRFANWLN-----RA-----V--S------S--D-S-Q-I---------L---GSPEA--------AQWSQ-E---------LS---------------K------ELTMFKEIL-----------------EDER--------D----------
-------------------

WP_023103132 ----------------------------------------------MSERPNG-----------D--S---------PTLR--QLIE--S-FIQ-ER-----L----QGK------------------------------------------------------------------------------------L---------D--------------K----------L----------KP------DE-----DV----------KR---QS--------------------LL----------------------------------------------------AG----------
-----HR---REAW-----LA---DA-----ARRV-------G----QLQ-L---------------VTHTLKPIHPD---A----------------R---G-S------NL----H----S----L--------P-QAP-----GQ--P--GL---------AGS--H--EL-------G-D-----K--L--V-SD-V-VG----N----AA-----A-L-----DVFKFL-S------L----Q--Y--Q-----------GKNL--------------LNWLT-EDS-A--EAVQAL---S---D-----NA-----EQAREWRQAFIGI--T---AV--KG---
APASHSLAKQLYFPL-------PG-------------------SG---YHLLAPLFPTSLVHHVHAL-LR----E--A---R--F-G----------------EA-ARAAR-E-----AR------------S--RQESW--PH-G-FSE-----YPN-LAIQKFGG-T----KPQ---------NISQLN--SERY-------------GENWLLPSL-P----PHWQ--RQ--DQ-RAPIRH----SSVF-----EH---DFG--R-S--PEV--SRLTRTLQRLL-------A--KT-----R-H-----N----------
N----FT---IR--KY----RA------QLVGQICDEALQYAARLR-------E------L-E-----P--G----WS------A----T----P---G-----CQ--------LHD---AEQLWL-----D-------------PLR--AQT-------------DETFLQ--RRL--RGD-WP-AEVGNRFANWLN-----RA-----V--S------SD-------S-Q-I---------L---GSPEA--------AQWSQ-E---------LS---------------K------ELTMFKEIL-----------------EDER--------D----------
-------------------

WP_034020359 ----------------------------------------------MSERPNG-----------D--S---------PTLR--QLIE--S-FIQ-ER-----L----QGK------------------------------------------------------------------------------------L---------D--------------K----------L----------KP------DE-----DV----------KR---QS--------------------LL----------------------------------------------------AG----------
-----HR---REAW-----LA---DA-----ARRV-------G----QLQ-L---------------VTHTLKPIHPD---A----------------R---G-S------NL----H----S----L--------P-QAP-----GQ--P--GL---------AGS--H--EL-------G-D-----K--L--V-SD-V-VG----N----AA-----A-L-----DVFKFL-S------L----Q--Y--Q-----------GKNL--------------LNWLT-EDS-A--EAVQAL---S---D-----NA-----EQAREWRQAFIGI--T---TV--KG---
APASHSLAKQLYFPL-------PG-------------------SG---YHLLAPLFPTSLVHHVHAL-LR----E--A---R--F-G----------------DA-AKAAR-E-----AR------------S--RQESW--PH-G-FSE-----YPN-LAIQKFGG-T----KPQ---------NISQLN--SERY-------------GENWLLPSL-P----PHWQ--R-Q-DQ-RAPIRH----SSVF-----EH---DFG--R-S--PEV--SRLTRTLQRLL--------------A-----K-----------TRD-N----------
N----FT---IR--RY----RA------QLVGQICDEALQYAARLR-------E------L-E-----P--G----WS------A----T----P---G-----CQ--------LHD---AEQLWL-----D-------------PLR--AQT-------------DETFLQ--RRL--RGD-WP-AEVGNRFANWLN-----RA-----V--S------S--D-S-Q-I---------L---GSPEA--------AQWSQ-E---------LS---------------K------ELTMFKEIL-----------------EDER--------D----------
-------------------

WP_033994843 ----------------------------------------------MSERPNG-----------D--S---------PTLR--QLIE--S-FIQ-ER-----L----QGK------------------------------------------------------------------------------------L---------D--------------K----------L----------KP------DE-----DV----------KR---QS--------------------LL----------------------------------------------------AG----------
-----HR---REAW-----LA---DA-----ARRV-------G----QLQ-L---------------VTHTLKPIHPD---A----------------R---G-S------NL----H----S----L--------P-QAP-----GQ--P--GL---------AGS--H--EL-------G-D-----R--L--V-SD-V-VG----N----AA-----A-L-----DVFKFL-S------L----Q--Y--Q-----------GKNL--------------LNWLT-EDS-A--EAVQAL---S---D-----NA-----EQAREWRQAFIGI--T---AV--KG---
APASHSLAKQLYFPL-------PG-------------------SG---YHLLAPLFPTSLVHHVHAL-LR----E--A---R--F-G----------------DA-AKAAR-E-----AR------------S--RQESW--PH-G-FSE-----YPN-LAIQKFGG-T----KPQ---------NISQLN--SERY-------------GENWLLPSL-P----PHWQ--R-Q-DQ-RAPIRH----SSVF-----EH---DFG--R-S--PEV--SRLTRTLQRLL--------------A-----K-----------TRD-N----------
N----FT---IR--RY----RA------QLVGQICDEALQYAARLR-------E------L-E-----P--G----WS------A----T----P---G-----CQ--------LHD---AEQLWL-----D-------------PLR--AQT-------------DETFLQ--RRL--RGD-WP-AEVGNRFANWLN-----RA-----V--S------S--D-S-Q-I---------L---GSPEA--------AQWSQ-E---------LS---------------K------ELTMFKEIL-----------------EDER--------D----------
-------------------

WP_034054824 ----------------------------------------------MSERPNG-----------D--S---------PTLR--QLIE--S-FIQ-ER-----L----QGK------------------------------------------------------------------------------------L---------D--------------K----------L----------KP------DE-----DV----------KR---QS--------------------LL----------------------------------------------------AG----------
-----HR---REAW-----LA---DA-----ARRV-------G----QLQ-L---------------VTHTLKPIHPD---A----------------R---G-S------NL----H----S----L--------P-QAP-----GQ--P--GL---------AGS--H--EL-------G-D-----K--L--V-SD-V-VG----N----AA-----A-L-----DVFKFL-S------L----Q--Y--Q-----------GKNL--------------LNWLT-EDS-A--EAVQAL---S---D-----NA-----EQAREWRQAFIGI--T---AV--KG---
APASHSLAKQLYFPL-------PG-------------------SG---YHLLAPLFPTSLVHHVHAL-LR----E--A---R--F-G----------------DA-AKAAR-E-----AR------------S--RQESW--PH-G-FSE-----YPN-LAIQKFGG-T----KPQ---------NISQLN--SERY-------------GENWLLPSL-P----PHWQ--R-Q-DQ-RAPIRH----SSVF-----EH---DFG--R-S--PEV--SRLTRTLQRLL--------------A-----K-----------TRH-N----------
N----FT---IR--KY----RA------QLVGQICDEALQYAARLR-------E------L-E-----P--G----WS------A----T----P---G-----CQ--------LHD---AEQLWL-----D-------------PLR--AQT-------------DETFLQ--RRL--RGD-WP-AEVGNRFANWLN-----RA-----V--S------S--D-S-Q-I---------L---GSPEA--------AQWSQ-E---------LS---------------K------ELTMFKEIL-----------------EDER--------D----------
-------------------

WP_033980640 ----------------------------------------------MSERPNG-----------D--S---------PTLR--QLIE--S-FIQ-ER-----L----QGK------------------------------------------------------------------------------------L---------D--------------K----------L----------KP------DE-----DV----------KR---QS--------------------LL----------------------------------------------------AG----------
-----HR---REAW-----LA---DA-----ARRV-------G----QLQ-L---------------VTHTLKPIHPD---A----------------R---G-S------NL----H----S----L--------P-QAP-----GQ--P--GL---------AGS--H--EL-------G-D-----R--L--V-SD-V-VG----N----AA-----A-L-----DVFKFL-S------L----Q--Y--Q-----------GKNL--------------LNWLT-EDS-A--EAVQAL---S---D-----NA-----EQAREWRQAFIGI--T---AV--KG---
APASHSLAKQLYFPL-------PG-------------------SG---YHLLAPLFPTSLVHHVHAL-LR----E--A---R--F-G----------------DA-AKAAR-E-----AR------------S--RQESW--PH-G-FSE-----YPN-LAIQKFGG-T----KPQ---------NISQLN--SERY-------------GENWLLPSL-P----PHWQ--RQ--DQ-RAPIRH----SSVF-----EH---DFG--R-S--PEV--SRLTRTLQRLL-------A--KT-----R-H-----N----------
N----FT---IR--RY----RA------QLVGQICDEALQYAARLR-------E------L-E-----P--G----WS------A----T----P---G-----CQ--------LHD---AEQLWL-----D-------------PLR--AQT-------------DETFLQ--RRL--RGD-WP-AEVGNRFANWLN-----RA-----V--S------SD-------S-Q-I---------L---GSPEA--------AQWSQ-E---------LS---------------K------ELTMFKEIL-----------------EDER--------D----------
-------------------

WP_057381967 ----------------------------------------------MTSPLPTP-----------T--W---------QELR--QLIE--S-FIQ-ER-----L----QGK------------------------------------------------------------------------------------L---------D--------------K----------L----------KP------DE-----DV----------KR---QS--------------------LL----------------------------------------------------AG----------
-----HR---REAW-----LA---DA-----ARRV-------G----QLQ-L---------------VTHTLKPIHPD---A----------------R---G-S------NL----H----S----L--------P-QAP-----GQ--P--GL---------AGS--H--EL-------G-D-----K--L--V-SD-V-VG----N----AA-----A-L-----DVFKFL-S------L----Q--Y--Q-----------GKNL--------------LNWLT-EDS-A--EAVQAL---S---D-----NA-----EQAREWRQAFIGI--T---TV--KG---
APASHSLAKQLYFPL-------PG-------------------SG---YHLLAPLFPTSLVHHVHAL-LR----E--A---R--F-G----------------DA-AKAAR-E-----AR------------S--RQESW--PH-G-FSE-----YPN-LAIQKFGG-T----KPQ---------NISQLN--SERY-------------GENWLLPSL-P----PHWQ--R-Q-DQ-RAPIRH----SSVF-----EH---DFG--R-S--PEV--SRLTRTLQRLL--------------A-----K-----------TRD-N----------
N----FT---IR--RY----RA------QLVGQICDEALQYAARLR-------E------L-E-----P--G----WS------A----T----P---G-----CQ--------LHD---AEQLWL-----D-------------PLR--AQT-------------DETFLQ--RRL--RGD-WP-AEVGNRFANWLN-----RA-----V--S------S--D-S-Q-I---------L---GSPEA--------AQWSQ-E---------LS---------------K------ELTMFKEIL-----------------EDER--------D----------
-------------------

WP_134547718 ----------------------------------------------MTSPLPTP-----------T--W---------QELR--QLIE--S-FIQ-ER-----L----QGK------------------------------------------------------------------------------------L---------D--------------K----------L----------KP------DE-----DV----------KR---QS--------------------LL----------------------------------------------------AG----------
-----HR---REAW-----LA---DA-----ARRV-------G----QLQ-L---------------VTHTLKPIHPD---A----------------R---G-S------NL----H----S----L--------P-QAP-----GQ--P--GL---------AGS--H--EL-------G-D-----K--L--V-SD-V-VG----N----AA-----A-L-----DVFKFL-S------L----Q--Y--Q-----------GKNL--------------LNWLT-EDS-A--EAVQAL---S---D-----NA-----EQAREWRQAFIGI--T---TV--KG---
APASHSLAKQLYFPL-------PG-------------------SG---YHLLAPLFPTSLVHHVHAL-LR----E--A---R--F-G----------------DA-AKAAR-E-----AR------------S--RQESW--PH-G-FSE-----YPN-LAIQKFGG-T----KPQ---------NISQLN--SERY-------------GENWLLPSL-P----PHWQ--R-Q-DQ-RAPIRH----SSVF-----EH---DFG--R-S--PEV--SRLTRTLQRLL--------------A-----K-----------TRD-N----------
N----FT---IR--RY----RA------QLVGQICDEALQYAARLR-------E------L-E-----P--G----WS------A----T----P---G-----CQ--------LHD---AEQLWL-----D-------------PLR--AQT-------------DETFLQ--RRL--RGD-WP-AEVGNRFANWLN-----RA-----V--S------S--D-S-Q-I---------L---GSP-------------------------------------------------------------------------------------------------
WP_047106005 ----------------------------------------------MTSPLPTP-----------T--W---------QELR--QLIE--S-FIQ-ER-----L----QGK------------------------------------------------------------------------------------L---------D--------------K----------L----------KP------DE-----DV----------KR---QS--------------------LL----------------------------------------------------AG----------
-----HR---REAW-----LA---DA-----ARRV-------G----QLQ-L---------------VTHTLKPIHPD---A----------------R---G-S------NL----H----S----L--------P-QAP-----GQ--P--GL---------AGS--H--EL-------G-D-----K--L--V-SD-V-VG----N----AA-----A-L-----DVFKFL-S------L----Q--Y--Q-----------GKNL--------------LNWLT-EDS-A--EAVQAL---S---D-----NA-----EQAREWRQAFIGI--T---AV--KG---
APASHSLAKQLYFPL-------PG-------------------SG---YHLLAPLFPTSLVHHVHAL-LR----E--A---R--F-G----------------DA-AKAAR-E-----AR------------S--RQESW--PH-G-FSE-----YPN-LAIQKFGG-T----KPQ---------NISQLN--SERY-------------GENWLLPSL-P----PHWQ--R-Q-DQ-RAPIRH----SSVF-----EH---DFG--R-S--PEV--SRLTRTLQRLL--------------A-----K-----------TRH-N----------
N----FT---IR--RY----RA------QLVGQICDEALQYAARLR-------E------L-E-----P--G----WS------A----T----P---G-----CQ--------LHD---AEQLWL-----D-------------PLR--AQT-------------DETFLQ--RRL--RGD-WP-AEVGNRFANWLN-----RA-----V--S------S--D-S-Q-I---------L---GSPEA--------AQWSQ-E---------LS---------------K------ELTMFKEIL-----------------EDER--------D----------
-------------------

WP_124205903 ----------------------------------------------MTSPLPTP-----------T--W---------QELR--QLIE--S-FIQ-ER-----L----QGK------------------------------------------------------------------------------------L---------D--------------K----------L----------KP------DE-----DV----------KR---QS--------------------LL----------------------------------------------------AG----------
-----HR---REAW-----LA---DA-----ARRV-------G----QLQ-L---------------VTHTLKPIHPD---T----------------R---G-S------NL----H----S----L--------P-QAP-----GQ--P--GL---------AGS--H--EL-------G-D-----K--L--V-SD-V-VG----N----AA-----A-L-----DVFKFL-S------L----Q--Y--Q-----------GKNL--------------LNWLT-EDS-A--EAVQAL---S---D-----NA-----EQAREWRQAFIGI--T---AV--KG---
APASHSLAKQLYFPL-------PG-------------------SG---YHLLAPLFPTSLVHHVHAL-LR----E--A---R--F-G----------------DA-AKAAR-E-----AR------------S--RQESW--PH-G-FSE-----YPN-LAIQKFGG-T----KPQ---------NISQLN--SERY-------------GENWLLPSL-P----PHWQ--RQ--DQ-RAPIRH----SSVF-----EH---DFG--R-S--PEV--SRLTRTLQRLL-------A--KT-----R-H-----N----------
N----FT---IR--RY----RA------QLVGQICDEALQYAARLR-------E------L-E-----P--G----WS------A----T----P---G-----CQ--------LHD---AEQLWL-----D-------------PLR--AQT-------------DETFLQ--RRL--RGD-WP-AEVGNRFANWLN-----RA-----V--S------SD-------S-Q-I---------L---GSPEA--------AQWSQ-E---------LS---------------K------ELTMFKEIL-----------------EDER--------D----------
-------------------

WP_057383734 ----------------------------------------------MTSPLPTP-----------T--W---------QELR--QLIE--S-FIQ-ER-----L----QGK------------------------------------------------------------------------------------L---------D--------------K----------L----------KP------DE-----DV----------KR---QS--------------------LL----------------------------------------------------AG----------
-----HR---REAW-----LA---DA-----ARRV-------G----QLQ-L---------------VTHTLKPIHPD---A----------------R---G-S------NL----H----S----L--------P-QAP-----GQ--P--GL---------AGS--H--EL-------G-D-----K--L--V-SD-V-VG----N----AA-----A-L-----DVFKFL-S------L----Q--Y--Q-----------GKNL--------------LNWLT-EDS-A--EAVQAL---S---D-----NA-----EQAREWRQAFIGI--T---AV--KG---
APASHSLAKQLYFPL-------PG-------------------SG---YHLLAPLFPTSLVHHVHAL-LR----E--A---R--F-G----------------DA-AKAAR-E-----AR------------S--RQESW--PH-G-FSE-----YPN-LAIQKFGG-T----KPQ---------NISQLN--SERY-------------GENWLLPSL-P----PHWQ--R-Q-DQ-RAPIRH----SSVF-----EH---DFG--R-S--PEV--SRLTRTLQRLL--------------A-----K-----------TRH-N----------
N----FT---IR--RY----RA------QLVGQICDEALQYAARLR-------E------L-E-----P--G----WS------A----T----P---G-----CQ--------LHD---AEQLWL-----D-------------PLR--AQT-------------DETFLQ--RRL--RGD-WP-AEVGNRFANWLN-----RA-----V--S------S--D-S-Q-I---------L---GSPEA--------AQWSQ-E---------LS---------------K------ELTMFKEIL-----------------EDER--------D----------
-------------------

WP_128669076 ----------------------------------------------MTSPLPTP-----------T--W---------QELR--QLIE--S-FIQ-ER-----L----QGK------------------------------------------------------------------------------------L---------D--------------K----------L----------KP------DE-----DV----------KR---QS--------------------LL----------------------------------------------------AG----------
-----HR---REAW-----LA---DA-----ARRV-------G----QLQ-L---------------VTHTLKPIHPD---A----------------R---G-S------NL----H----S----L--------P-QAP-----GQ--P--GL---------AGS--H--EL-------G-D-----R--L--V-SD-V-VG----N----AA-----A-L-----DVFKFL-S------L----Q--Y--Q-----------GKNL--------------LNWLT-EDS-A--EAVQAL---S---D-----NA-----EQAREWRQAFIGI--T---AV--KG---
APASHSLAKQLYFPL-------PG-------------------SG---YHLLAPLFPTSLVHHVHAL-LR----E--A---R--F-G----------------DA-AKAAR-E-----AR------------S--RQESW--PH-G-FSE-----YPN-LAIQKFGG-T----KPQ---------NISQLN--SERY-------------GENWLLPSL-P----PHWQ--RQ--DQ-RAPIRH----SSVF-----EH---DFG--R-S--PEV--SRLTRTLQRLL-------A--KT-----R-H-----N----------
N----FT---IR--RY----RA------QLVGQICDEALQYAARLR-------E------L-E-----P--G----WS------A----T----P---G-----CQ--------LHD---AEQLWL-----D-------------PLR--AQT-------------DETFLQ--RRL--RGD-WP-AEVGNRFANWLN-----RA-----V--S------SD-------S-Q-I---------L---GSPEA--------AQWSQ-E---------LS---------------K------ELTMFKEIL-----------------EDER--------D----------
-------------------

WP_033961478 ----------------------------------------------MTSPLPTP-----------T--W---------QELR--QLIE--S-FIQ-ER-----L----QGK------------------------------------------------------------------------------------L---------D--------------K----------L----------KP------DE-----DL----------KR---QS--------------------LL----------------------------------------------------AG----------
-----HR---REAW-----LA---DA-----ARRV-------G----QLQ-L---------------VTHTLKPIHPD---A----------------R---G-S------NL----H----S----L--------P-QAP-----GQ--P--GL---------AGS--H--EL-------G-D-----R--L--V-SD-V-VG----N----AA-----A-L-----DVFKFL-S------L----Q--Y--Q-----------GKNL--------------LNWLT-EDS-A--EAVQAL---S---D-----NA-----EQAREWRQAFIGI--T---AV--KG---
APASHSLAKQLYFPL-------PG-------------------SG---YHLLAPLFPTSLVHHVHAL-LR----E--A---R--F-G----------------DA-AKAAR-E-----AR------------S--RQESW--PH-G-FSE-----YPN-LAIQKFGG-T----KPQ---------NISQLN--SERY-------------GENWLLPSL-P----PHWQ--R-Q-DQ-RAPIRH----SSVF-----EH---DFG--R-S--PEV--SRLTRTLQRLL--------------A-----K-----------TRH-N----------
N----FT---IR--RY----RA------QLVGQICDEALQYAARLR-------E------L-E-----P--G----WS------A----T----P---G-----CQ--------LHD---AEQLWL-----D-------------PLR--AQT-------------DETFLQ--RRL--RGD-WP-AEVGNRFANWLN-----RA-----V--S------S--D-S-Q-I---------L---GSPEA--------AQWSQ-E---------LS---------------K------ELTMFKEIL-----------------EDER--------D----------
-------------------

WP_134277129 ----------------------------------------------MSERPNG-----------D--S---------PTLR--QLIE--S-FIQ-ER-----L----QGK------------------------------------------------------------------------------------L---------D--------------K----------L----------KP------DE-----DV----------KR---QS--------------------LL----------------------------------------------------AG----------
-----HR---REAW-----LA---DA-----ARRV-------G----QLQ-L---------------VTHTLKPIHPD---A----------------R---G-S------NL----H----S----L--------P-QAP-----GQ--P--GL---------AGS--H--EL-------G-D-----K--L--V-SD-V-VG----N----AA-----A-L-----DVFKFL-S------L----Q--Y--Q-----------GKNL--------------LNWLT-EDS-A--EAVQAL---S---D-----NA-----EQAREWRQAFIGI--T---AV--KG---
APASHSLAKQLYFPL-------PG-------------------SG---YHLLAPLFPTSLVHHVHAL-LR----E--A---R--F-G----------------DA-AKAAR-E-----AR------------S--RQESW--PH-G-FSE-----YPN-LAIQKFGG-T----KPQ---------NISQLN--SERY-------------GXNWLLPSL-P----PHWQ--R-Q-DQ-RAPIRH----SSVF-----EH---DFG--R-S--PEV--SRLTRTLQRLL--------------A-----K-----------TRH-N----------
N----FT---IR--RY----RA------QLVGQICDEALQYAARLR-------E------L-E-----P--G----WS------A----T----P---G-----CQ--------LHD---AEQLWL-----D-------------PLR--AQT-------------DETFLQ--RRL--RGD-WP-AEVGNRFANWLN-----RA-----V--S------S--D-S-Q-I---------L---GSPXA--------AQWSQ-E---------LS---------------K------ELTMFKEIL-----------------EDER--------D----------
-------------------

WP_025324967 ----------------------------------------------MSERPNG-----------D--S---------PTLR--QLIE--S-FIQ-ER-----L----QGK------------------------------------------------------------------------------------L---------D--------------K----------L----------KP------DE-----DV----------KR---QS--------------------LL----------------------------------------------------AG----------
-----HR---REAW-----LA---DA-----ARRV-------G----QLQ-L---------------VTHTLKPIHPD---A----------------R---G-S------NL----H----S----L--------P-QAP-----GQ--P--GL---------AGS--H--EL-------G-D-----K--L--V-SD-V-VG----N----AA-----A-L-----DVFKFL-S------L----Q--Y--Q-----------GKNL--------------LNWLT-EDS-A--EAVQAL---S---D-----NA-----EQAREWRQAFIGI--T---AV--KG---
APASHSLAKQLYFPL-------PG-------------------SG---YHLLAPLFPTSLVHHVHAL-LR----E--A---R--F-G----------------DA-AKAAR-E-----AR------------S--RQESW--PH-G-FSE-----YPN-LAIQKFGG-T----KPQ---------NISQLN--SERY-------------GENWLLPSL-P----PHWQ--R-Q-DQ-RAPIRH----SSVF-----EH---DFG--R-S--PEV--SRQTRTLQRLL--------------A-----K-----------TRH-N----------
N----FT---IR--RY----RA------QLVGQICDEALQYAARLR-------E------L-E-----P--G----WS------A----T----P---G-----CQ--------LHD---AEQLWL-----D-------------PLR--AQT-------------DETFLQ--RRL--RGD-WP-AEVGNRFANWLN-----RA-----V--S------S--D-S-Q-I---------L---GSPEA--------AQWSQ-E---------LS---------------K------ELTMFKDIL-----------------EDER--------D----------
-------------------

WP_023122499 ----------------------------------------------MTSPLPTP-----------T--W---------QELR--QFIE--S-FIQ-ER-----L----QGK------------------------------------------------------------------------------------L---------D--------------K----------L----------HP------DE-----DD----------KR---QT--------------------LL----------------------------------------------------AT----------
-----HR---REAW-----LA---DA-----ARRV-------G----QLQ-L---------------VTHTLKPIHPD---A----------------R---G-S------NL----H----S----L--------P-QAP-----EQ--P--GL---------AGS--H--EL-------G-D-----K--L--V-SD-V-VG----N----AA-----A-L-----DVFKFL-S------L----Q--Y--Q-----------GKNL--------------LNWLT-EDS-A--EAVQAL---S---D-----NA-----EQAREWRQAFIGI--T---AV--KG---
APASHSLAKQLYFPL-------PG-------------------SG---YHLLAPLFPTSLVHHVHAL-LR----E--A---R--F-G----------------DA-AKAAR-E-----AR------------S--RQESW--PH-G-FSE-----YPN-LAIQKFGG-T----KPQ---------NISQLN--SERY-------------GENWLLPSL-P----PHWQ--R-Q-DQ-RAPIRH----SSVF-----EH---DFG--R-S--PEV--SRLTRTLQRLL--------------A-----K-----------TRH-N----------
N----FT---IR--RY----RA------QLVGQICDEALQYAARLR-------E------L-E-----P--G----WS------A----T----P---G-----CQ--------LHD---AEQLWL-----D-------------PLR--AQT-------------DETFLQ--RRL--RGD-WP-AEVGNRFANWLN-----RA-----V--S------S--D-S-Q-I---------L---GSPEA--------AQWSQ-E---------LS---------------K------ELTMFKEIL-----------------EDER--------D----------
-------------------

WP_121328008 ----------------------------------------------MTSPLPTP-----------T--W---------QELR--QLIE--S-FIQ-ER-----L----QGK------------------------------------------------------------------------------------L---------D--------------K----------L----------KP------DE-----DV----------KR---QS--------------------LL----------------------------------------------------AG----------
-----HR---REAW-----LA---DA-----ARRV-------G----QLQ-L---------------VTHTLKPIHPD---A----------------R---G-S------NL----H----S----L--------P-QAP-----GQ--P--GL---------AGS--H--EL-------G-D-----K--L--V-SD-V-VG----N----AA-----A-L-----DVFKFL-S------L----Q--Y--Q-----------GKNL--------------LNWLT-EDS-A--EAVQAL---S---D-----NA-----EQAREWRQAFIGI--T---AV--KG---
APASHSLAKQLYFPL-------PG-------------------SG---YHLLAPLFPTSLVHHVHAL-LR----E--A---R--F-G----------------DA-AKAAR-E-----AR------------S--RQESW--PH-G-FSE-----YPN-LAIQKFGG-T----KPQ---------NISQLN--SERY-------------GENWLLPSL-P----PHWQ--R-Q-DQ-RAPIRH----SSVF-----EH---DFG--R-S--PEV--SRLTRTLQRLL--------------A-----K-----------TRH-N----------
N----FT---IR--RY----RA------QLVGQICDEALQYAARLR-------E------L-E-----P--G----WS------A----T----P---G-----CQ--------LHD---AEQLWL-----D-------------PLR--AQT-------------DETFLQ--RRL--RGD-WP-AEVGNRFANWLN-----RA-----V--S------S--D-S-Q-I---------L---GSPEA--------AQWSQ-E---------LS---------------K------ELTMFKEIL-----------------EDER--------D----------
-------------------

WP_073664313 -------------------------------------------------------------------------------------------------------------------------------------------------------------------------------------------------------------------------------------------------------------------------------------------------------------------------------------------------------------------------------------------------------------------------------------------------MA---DA-----ARRV-
------G----QLQ-L---------------VTHTLKPIHPD---A----------------R---G-S------NL----H----S----L--------P-QAP-----GQ--P--GL---------AGS--H--EL-------G-D-----K--L--V-SD-V-VG----N----AA-----A-L-----DVFKFL-S------L----Q--Y--Q-----------GKNL--------------LNWLT-EDS-A--EAVQAL---S---D-----NA-----EQAREWRQAFIGI--T---AV--KG---APASHSLAKQLYFPL-------PG-------------------SG---
YHLLAPLFPTSLVHHVHGL-LR----E--A---R--F-G----------------DA-AKAAR-E-----AR------------S--RQESW--PH-G-FSE-----YPN-LAIQKFGG-T----KPQ---------NISQLN--SERY-------------GENWLLPSL-P----PHWQ--R-Q-DQ-RAPIRH----SSVF-----EH---DFG--R-S--PEV--SRLTRTLQRLL--------------A-----K-----------TRH-N----N---FT---IR--RY----RA------
QLVGQICDEALQYAARLR-------E------L-E-----P--G----WS------A----T----P---G-----CQ--------LHD---AEQLWL-----D-------------PLR--AQT-------------DETFLQ--RRL--RGD-WP-AEVGNRFANWLN-----RA-----V--S------S--D-S-Q-I---------L---GSPEA--------AQWSQ-E---------LS---------------K------ELTMFKEIL-----------------EDER--------D-----------------
WP_124136638 ----------------------------------------------MTSPLPTP-----------T--W---------QELR--QLIE--S-FIQ-ER-----L----QGK------------------------------------------------------------------------------------L---------D--------------K----------L----------KP------DE-----DV----------KR---QS--------------------LL----------------------------------------------------AG----------
-----HR---REAW-----LA---DA-----ARRV-------G----QLQ-L---------------VTHTLKPIHPD---A----------------R---G-S------NL----H----S----L--------P-QAP-----GQ--P--GL---------AGS--H--EL-------G-D-----K--L--V-SD-V-VG----N----AA-----A-L-----DVFKFL-S------L----Q--Y--Q-----------GKNL--------------LNWLT-EDS-A--EAVQAL---S---D-----NA-----EQAREWRQAFIGI--T---AV--KG---
APASHSLAKQLYFPL-------PG-------------------SG---YHLLAPLFPTSLVHHVHAL-LR----E--A---R--F-G----------------DA-AKAAR-E-----AR------------S--RQESW--PH-G-FSE-----YPN-LAIQKFGG-T----KPQ---------NISQLN--SERY-------------GENWLLPSL-P----PHWQ--R-Q-DQ-RAPIRH----SSVF-----EH---DFG--R-S--PEV--SRLTRTLQRLL--------------A-----K-----------TRH-N----------
N----FT---IR--RY----RA------QLVGQICDEALQYAARLR-------E------L-E-----P--G----WS------A----T----P---G-----CQ--------LHD---AEQLWL-----D-------------PLR--AQT-------------DETFLQ--RRL--RGD-WP-AEVGNRFANWLN-----RA-----V--S------S--D-S-Q-I---------L---GSPEA--------AQWSQ-E---------LS---------------K------ELTMFKEIL-----------------EDER--------D----------
-------------------

WP_033980045 ----------------------------------------------MSERPNG-----------D--S---------PTLR--QLIE--S-FIQ-ER-----L----QGK------------------------------------------------------------------------------------L---------D--------------K----------L----------KP------DE-----DV----------KR---QS--------------------LL----------------------------------------------------AG----------
-----HR---REAW-----LA---DA-----ARRV-------G----QLQ-L---------------VTHTLKPIHPD---A----------------R---G-S------NL----H----S----L--------P-QAP-----GQ--P--GL---------AAS--H--EL-------G-D-----K--L--V-SD-V-VG----N----AA-----A-L-----DVFKFL-S------L----Q--Y--Q-----------GKNL--------------LNWLT-EDS-A--EAVQAL---S---D-----NA-----EQAREWRQAFIGI--T---AV--KG---
APASHSLAKQLYFPL-------PG-------------------SG---YHLLAPLFPTSLVHHVHAL-LR----E--A---R--F-G----------------DA-AKAAR-E-----AR------------S--RQESW--PH-G-FSE-----YPN-LAIQKFGG-T----KPQ---------NISQLN--SERY-------------GENWLLPSL-P----PHWQ--R-Q-DQ-RAPIRH----SSVF-----EH---DFG--R-S--PEV--SRLTRTLQRLL--------------A-----K-----------TRH-N----------
N----FT---IR--RY----RA------QLVGQICDEALQYAARLR-------E------L-E-----P--G----WS------A----T----P---G-----CQ--------LHD---AEQLWL-----D-------------PLR--AQT-------------DETFLQ--RRL--RGD-WP-AEVGNRFANWLN-----RA-----V--S------S--D-S-Q-I---------L---GSPEA--------AQWSQ-E---------LS---------------K------ELTMFKEIL-----------------EDER--------D----------
-------------------

WP_134321152 ----------------------------------------------MSERPNG-----------D--S---------PTLR--QLIE--S-FIQ-ER-----L----QGK------------------------------------------------------------------------------------L---------D--------------K----------L----------KP------DE-----DV----------KR---QS--------------------LL----------------------------------------------------AG----------
-----HR---REAW-----LA---DA-----ARRV-------G----QLQ-L---------------VTHTLKPIHPD---A----------------R---G-S------NL----H----S----L--------P-QAP-----GQ--P--GL---------AGS--H--EL-------G-D-----K--L--V-SD-V-VG----N----AA-----A-L-----DVFKFL-S------L----Q--Y--Q-----------GKNL--------------LNWLT-EDS-A--EAVQAL---S---D-----NA-----EQAREWRQAFIGI--T---AV--KG---
APASHSLAKQLYFPL-------PG-------------------SG---YHLLAPLFPTSLVHHVHAL-LR----E--A---R--F-G----------------DA-AKAAR-E-----AR------------S--RQESW--PH-G-FSE-----YXN-LAIQKFGG-T----KPQ---------NISQLN--SERY-------------GENWLLPSL-P----PHWQ--R-Q-DQ-RAPIRH----SSVF-----EH---DFG--R-S--PEV--SRLTRTLQRLL--------------A-----K-----------TRH-N----------
N----FT---IR--RY----RA------QLVGQICDEALQYAARLR-------E------L-E-----P--G----WS------A----T----P---G-----CQ--------LHD---AEQLWL-----D-------------PLR--AQT-------------DETFLQ--RRL--RGD-WP-AEVGNRFANWLN-----RA-----V--S------S--D-S-Q-I---------L---GSPEA--------AQWSQ-E---------LS---------------K------ELTMFKEIL-----------------EDER--------D----------
-------------------

WP_034025305 ----------------------------------------------MTSPLPTP-----------T--W---------QELR--QFIE--S-FIQ-ER-----L----QGK------------------------------------------------------------------------------------L---------D--------------K----------L----------HP------DE-----DD----------KR---QT--------------------LL----------------------------------------------------AT----------
-----HR---REAW-----LA---DA-----ARRV-------G----QLQ-L---------------VTHTLKPIHPD---A----------------R---G-S------NL----H----S----L--------P-QAP-----GQ--P--GL---------AGS--H--EL-------G-D-----K--L--V-SD-V-VG----N----AA-----A-L-----DVFKFL-S------L----Q--Y--Q-----------GKNL--------------LNWLT-EDS-A--EAVQAL---S---D-----NA-----EQAREWRQAFIGI--T---AV--KG---
APASHSLAKQLYFPL-------PG-------------------SG---YHLLAPLFPTSLVHHVHAL-LR----E--A---R--F-G----------------DA-AKAAR-E-----AR------------S--RQESW--PH-G-FSE-----YPN-LAIQKFGG-T----KPQ---------NISQLN--SERY-------------GENWLLPSL-P----PHWQ--R-Q-DQ-RAPIRH----SSVF-----EH---DFG--R-S--PEV--SRLTRTLQRLL--------------A-----K-----------TRH-N----------
N----FT---IR--RY----RA------QLVGQICDEALQYAARVR-------E------L-E-----P--G----WS------A----T----P---G-----CQ--------LHD---AEQLWL-----D-------------PLR--AQT-------------DETFLQ--RRL--RGD-WP-AEVGNRFANWLN-----RA-----V--S------S--D-S-Q-I---------L---GSPEA--------AQWSQ-E---------LS---------------K------ELTMFKEIL-----------------EDER--------D----------
-------------------

WP_034073989 ----------------------------------------------MTSPLPTP-----------T--W---------QELR--QFIE--S-FIQ-ER-----L----QGK------------------------------------------------------------------------------------L---------D--------------K----------L----------HP------DE-----DD----------KR---QT--------------------LL----------------------------------------------------AT----------
-----HR---REAW-----LA---DA-----ARRV-------G----QLQ-L---------------VTHTLKPIHPD---A----------------R---G-S------NL----H----S----L--------P-QAP-----GQ--P--GL---------AGS--H--EL-------G-D-----K--L--V-SD-V-VG----N----AA-----A-L-----DVFKFL-S------L----Q--Y--Q-----------GKNL--------------LNWLT-EDS-A--EAVQAL---S---D-----NA-----EQAREWRQAFIGI--T---AV--KG---
APASHSLAKQLYFPL-------PG-------------------SG---YHLLAPLFPTSLVHHVHAL-LR----E--A---R--F-G----------------DA-AKAAR-E-----AR------------S--RQESW--PH-G-FSE-----YPN-LAIQKFGG-T----KPQ---------NISQLN--SERY-------------GENWLLPSL-P----PHWQ--R-Q-DQ-RAPIRH----SSVF-----EH---DFG--R-S--PEV--SRLTRTLQRLL--------------A-----K-----------TRH-N----------
N----FT---IR--RY----RA------QLVGQICDEALQYAARLR-------E------L-E-----P--G----WS------A----T----P---G-----CQ--------LHD---AEQLWL-----D-------------PLR--AQT-------------DETFLQ--RRL--RGD-WP-AEVGNRFANWLN-----RA-----V--S------S--D-S-Q-I---------L---GSPEA--------AQWSQ-E---------LS---------------K------ELTMFKEIL-----------------EDER--------D----------
-------------------

WP_121238331 ----------------------------------------------MTSPLPTP-----------T--W---------QELR--QFIE--S-FIQ-ER-----L----QGK------------------------------------------------------------------------------------L---------D--------------K----------L----------HP------DE-----DD----------KR---QT--------------------LL----------------------------------------------------AT----------
-----HR---REAW-----LA---DA-----ARRV-------G----QLQ-L---------------VTHTLKPIHPD---A----------------R---G-S------NL----H----S----L--------P-QAP-----GQ--P--GL---------AGS--H--EL-------G-D-----K--L--V-SD-V-VG----N----AA-----A-L-----DVFKFL-S------L----Q--Y--Q-----------GKNL--------------LNWLT-EDS-A--EAVQAL---S---D-----NA-----EQAREWRQAFIGI--T---AV--KG---
APASHSLAKQLYFPL-------PG-------------------SG---YHLLAPLFPTSLVHHVHAL-LR----E--A---R--F-G----------------DA-AKAAR-E-----AR------------S--RQESW--PH-G-FSE-----YPN-LAIQKFGG-T----KPQ---------NISQLN--SERY-------------GENWLLPSL-P----PHWQ--R-Q-DQ-RAPIRH----SSVF-----EH---DFG--R-S--PEV--SRLTRTLQRLL--------------A-----K-----------TRH-N----------
N----FT---IR--RY----RA------QLVGQICDEALQYAARLR-------E------L-E-----P--G----WS------A----T----P---G-----CQ--------LHD---AEQLWL-----D-------------PLR--AQT-------------DETFLQ--RRL--RGD-WP-AEVGNRFANWLN-----RA-----V--S------S--D-S-Q-I---------L---GSPEA--------AQWSQ-E---------LS---------------K------ELTMFKEIL-----------------EDER--------D----------
-------------------

WP_109413267 ----------------------------------------------MTSPLPTP-----------T--W---------QELR--QFIE--S-FIQ-ER-----L----QGK------------------------------------------------------------------------------------L---------D--------------K----------L----------HP------DE-----DD----------KR---QT--------------------LL----------------------------------------------------AT----------
-----HR---REAW-----LA---DA-----ARRV-------G----QLQ-L---------------VTHTLKPIHPD---A----------------R---G-S------NL----H----S----L--------P-QAP-----GQ--P--GL---------AGS--H--EL-------G-D-----K--L--V-SD-V-VG----N----AA-----A-L-----DVFKFL-S------L----Q--Y--Q-----------GKNL--------------LNWLT-EDS-A--EAVQAL---S---D-----NA-----EQAREWRQAFIGI--T---AV--KG---
APASHSLAKQLYFPL-------PG-------------------SG---YHLLAPLFPTSLVHHVHAL-LR----E--A---R--F-G----------------DA-AKAAR-E-----AR------------S--RQESW--PH-G-FSE-----YPN-LAIQKFGG-T----KPQ---------NISQLN--SERY-------------GENWLLPSL-P----PHWQ--R-Q-DQ-RAPIRH----SSVF-----EH---DFG--R-S--PEV--SRLTRTLQRLL--------------A-----K-----------TRH-N----------
N----FT---IR--RY----RA------QLVGQICDEALQYAARLR-------E------L-E-----P--G----WS------A----T----P---G-----CQ--------LHD---AEQLWL-----D-------------PLR--AQT-------------DETFLQ--RRL--RGD-WP-AEVGNRFANWLN-----RA-----V--S------S--D-S-Q-I---------L---GSPEA--------AQWSQ-E---------LS---------------K------ELTMFKEIL-----------------EDER--------D----------
-------------------

```
WP_034071840 -----------------------------------------------MTSPLPTP----------T--W---------QELR--QLIE--S-FIQ-ER-----L----QGK------------------------------------------------------------------------------------L---------D----------------K---------L--------HP-----DE----DD---------KR----QT---------------------LL------------------------------------------AT-----------------------
-----HR---REAW-----LA---DA-----ARRV-------G----QLQ-L-----------------VTHTLKPIHPD---A-----------------R---G-S------NL----H----S----L--------P-QAP----GQ--P--GL---------AGS--H--EL------G-D-----K--L--V-SD-V-VG----N----AA----A--L-----DVFKFL-S-----L----Q--Y--Q---------GKNL------------LNWLT-EDS-A---EAVQAL---S---D----NA----EQAREWRQAFIGI--T---AV--KG---
APASHSLAKQLYFPL-------PG-------------------SG---YHLLAPLFPTSLVHHVHGL-LR----E--A---R--F-G------------------DA-AKAAR-E-----AR------------S--RQESW--PH-G-FSE-----YPN-LAIQKFGG-T----KPQ---------NISQLN--SERY-----------GENWLLPSL-P----PHWQ--R-Q-DQ-RAPIRH----SSVF----EH---DFG--R-S--PEV--SRLTRTLQRLL-------------------A-----K-------------TRH-N-------
N----FT---IR--RY----RA------QLVGQICDEALQYAARLR--E-------L-E-----P----G----WS------A-----T---P----G-----CQ--------LHD---AEQLWL-----D-----------------PLR--AQT-------------DETFLQ--RRL--RGD-WP-AEVGNRFANWLN-----RA----V---S------S-----D--S-Q-I-------------L---GSPEA------AQWSQ-E-------LS---------------------K-----ELTMFKEIL------------------EDER------D------------------------
---------------
WP_049233520 ------------------------------------------------------------------------------------------------------------------------------------------------------------------------------------------------------------------------------AW-----LA---DA-----
ARRV-------G----QLQ-L-----------------VTHTLKPIHPD---A-----------------R---G-S------NL----H----S----L--------P-QAP----GQ--P--GL---------AGS--H--EL------G-D-----K--L--V-SD-V-VG----N----AA----A--L-----DVFKFL-S-----L----Q--Y--Q---------GKNL------------LNWLT-EDS-A---EAVQAL---S---D----NA----EQAREWRQAFIGI--T---AV--KG---APASHSLAKQLYFPL-------PG-------------------
SG---YHLLAPLFPTSLVHHVHGL-LR----E--A---R--F-G------------------DA-AKAAR-E-----AR------------S--RQESW--PH-G-FSE-----YPN-LAIQKFGG-T----KPQ---------NISQLN--SERY-----------GENWLLPSL-P----PHWQ-RQ---DQ-RAPIRH----SSVF----EH---DFG--R-S--PEV--SRLTRTLQRLL------A--KT----R-H-----N----------------------------N----FT---IR--RY----RA------
QLVGQICDEALQYAARLR--E-------L-E-----P----G----WS------A-----T---P----G-----CQ--------LHD---AEQLWL-----D-----------------PLR--AQT-------------DETFLQ--RRL--RGD-WP-AEVGNRFANWLN-----RA----V---S------SD------S-Q-I-------------L---GSPEA------AQWSQ-E-----------LS-------------------K-----ELTMFKEIL------------------EDER------D----------------------
WP_023087478 -----------------------------------------------MSERPNG----------D--S----------PTLR--QLIE--S-FIQ-ER-----L----QGK------------------------------------------------------------------------------------L---------D----------------K---------L--------KP-----DE----DV---------KR----QS---------------------LL------------------------------------------AG-----------------------
-----HR---REAW-----LA---DA-----ARRV-------G----QLQ-L-----------------VTHTLKPIHPD---A-----------------R---G-S------NL----H----S----L--------P-QAP----GQ--P--GL---------AGS--H--EL------G-D-----K--L--V-SD-V-VG----N----AA----A--L-----DVFKFL-S-----L----Q--Y--Q---------GKNL------------LNWLT-EDS-A---EAVQAL---S---D----NA----EQAREWRQAFIGI--T---AV--KG---
APASHSLAKQLYFPL-------PG-------------------SG---YHLLAPLFPTSLVHHVHGL-LR----E--A---R--F-G------------------DA-AKAAR-E-----AR------------S--RQESW--PH-G-FSE-----YPN-LAIQKFGG-T----KPQ---------NISQLN--SERY-----------GENWLLPSL-P----PHWQ-RQ---DQ-RAPIRH----SSVF----EH---DFG--R-S--PEV--SRLTRTLQRLL------A--KT----R-H-----N-----------------------------------
N----FT---IR--RY----RA------QLVGQICDEALQYAARLR--E-------L-E-----P----G----WS------A-----T---P----G-----CQ--------LHD---AEQLWL-----D-----------------PLR--AQT-------------DETFLQ--RRL--RGD-WP-AEVGNRFANWLN-----RA----V---S------S------D-S-Q------------IL---GSPEA------AQWSQ-E-------LS---------------------K-----ELTMFKEIL------------------EDER------D------------------------
---------------
WP_116805476 -----------------------------------------------MTSPLPTP----------T--W---------QELR--QFIE--S-FIQ-ER-----L----QGK------------------------------------------------------------------------------------L---------D----------------K---------L--------KP-----DE----DV---------KR----QS---------------------LL------------------------------------------AG-----------------------
-----HR---REAW-----LA---DA-----ARRV-------G----QLQ-L-----------------VTHTLKPIHPD---A-----------------R---G-S------NL----H----S----L--------P-QAP----GQ--P--GL---------AGS--H--EL------G-D-----K--L--V-SD-V-VG----N----AA----A--L-----DVFKFL-S-----L----Q--Y--Q---------GKNL------------LNWLT-EDS-A---EAVQAL---S---D----NA----EQAREWRQAFIGI--T---AV--KG---
APASHSLAKQLYFPL-------PG-------------------SG---YHLLAPLFPTSLVHHVHGL-LR----E--A---R--F-G------------------DA-AKAAR-E-----AR------------S--RQESW--PH-G-FSE-----YPN-LAIQKFGG-T----KPQ---------NISQLN--SERY-----------GENWLLPSL-P----PHWQ-RQ---DQ-RAPIRH----SSVF----EH---DFG--R-S--PEV--SRLTRTLQRLL------A--KT----R-H-----N-----------------------------------
N----FT---IR--RY----RA------QLVGQICDEALQYAARLR--E-------L-E-----P----G----WS------A-----T---P----G-----CQ--------LHD---AEQLWL-----D-----------------PLR--AQT-------------DETFLQ--RRL--RGD-WP-AEVGNRFANWLN-----RA----V---S------S------D-S-Q------------IL---GSPEA------AQWSQ-E-------LS---------------------K-----ELTMFKEIL------------------EDER------D------------------------
---------------
WP_139270494 -------------------------------------------------------------------------------------------G-----HR--------------------------------------------------------------------------------------------------------------------------------REAW-----LA---DA-----
ARRV-------G----QLQ-L-----------------VTHTLKPIHPD---A-----------------R---G-S------NL----H----S----L--------P--QA----PGQ--P--GL---------AGS--H--EL------G--------DKLV----SD-V-VG----N----AA----A--L-----DVFKFL-S-----L----Q--Y--Q---------GKNL------------LNWLT-EDS-A---EAVQAL---S---D----NA----EQAREWRQAFIGI--T---AV--KG-A--PASHSLAKQLYFPL-------PG-------------------
SG---YHLLAPLFPTSLVHHVHGL-LR----E--A---R--F-G------------------DA-AKAAR-E-----AR------------S--RQESW--PH-G-FSE-----YPN-LAIQKFGG-T----KPQ---------NISQLN--SERY-----------GENWLLPSL-P----PHWQRQ----DQ-RAPIRH----SSVF----EH---DFG--R-S--PEV--SRLTRTLQRLL--AKTRH--N---------------------------------------N----FT---IR--RY----RA------
QLVGQICDEALQYAARLR--E-------L-E-----P----G----WS------A-----T---P----G-----CQ--------LHD---AEQLWL-----D-----------------PLR--AQT-------------DETFLQ--RRL--RGD-WP-AEVGNRFANWLN-----RA----V---S------SD---------S-Q------------IL---GSPEA------AQWSQ-E-----------LS-------------------K-----ELTMFKEIL------------------EDER------D----------------------
WP_124127457 -----------------------------------------------MTSPLPTP----------T--W---------QELR--QFIE--S-FIQ-ER-----L----QGK------------------------------------------------------------------------------------L---------D----------------K---------L--------HP-----DE----DD---------KR----QT---------------------LL------------------------------------------AT-----------------------
-----HR---REAW-----LA---DA-----ARRV-------G----QLQ-L-----------------VTHTLKPIHPD---A-----------------R---G-S------NL----H----S----L--------P-QAP----GQ--P--GL---------AGS--H--EL------G-D-----K--L--V-SD-V-VG----N----AA----A--L-----DVFKFL-S-----L----Q--Y--Q---------GKNL------------LNWLT-EDS-A---EAVQAL---S---D----NA----EQAREWRQAFIDI--T---AV--KG---
APASHSLAKQLYFPL-------PG-------------------SG---YHLLAPLFPTSLVHHVHAL-LR----E--A---R--F-G------------------DA-AKAAR-E-----AR------------N--RRESW--PH-G-FSE-----YPN-LAIQKFGG-T----KPQ---------NISQLN--SERY-----------GENWLLPSL-P----PHWQ--R-Q-DQ-RAPIRH----SSVF----EH---DFG--R-S--PEV--SRLTRTLQRLL-------------------A-----K-------------TRD-N-------
N----FT---IR--RY----RA------QLVGQICDEALQYAARLR--E-------L-E-----P----G----WS------A-----T---P----G-----CQ--------LHD---AEQLWL-----D-----------------PLR--AQT-------------DETFLQ--HRL--RGD-WP-AEVGNRFANWLN-----RA----V---S------S-----D--S-Q-I-------------L---GSPEA------AQWSQ-E-------LS---------------------K-----ELTMFKEIL------------------EDER------D------------------------
---------------
WP_121305654 -----------------------------------------------MTSPLPTP----------T--W---------QELR--QFIE--S-FIQ-ER-----L----QGK------------------------------------------------------------------------------------L---------D----------------K---------L--------HP-----DE----DD---------KR----QT---------------------LL------------------------------------------AT-----------------------
-----HR---REAW-----LA---DA-----ARRV-------A----QLQ-L-----------------VTHTLKPIHPD---A-----------------R---G-S------NL----H----S----L--------P-QAP----GQ--P--GL---------AGS--H--EL------G-D-----K--L--V-SD-V-VG----N----AA----A--L-----DVFKFL-S-----L----Q--Y--Q---------GKNL------------LNWLT-EDS-A---EAVQAL---S---D----NA----EQAREWRQAFIDI--T---AV--KG---
APASHSLAKQLYFPL-------PG-------------------SG---YHLLAPLFPTSLVHHVHAL-LR----E--A---R--F-G------------------DA-AKAAR-E-----AR------------N--RRESW--PH-G-FSE-----YPN-LAIQKFGG-T----KPQ---------NISQLN--SERY-----------GENWLLPSL-P----PHWQ-RQ---DQ-RAPIRH----SSVF----EH---DFG--R-S--PEV--SRLTRTLQRLL------A--KT----R-D-----N-----------------------------------
N----FT---IR--RY----RA------QLVGQICDEALQYAARLR--E-------L-E-----P----G----WS------A-----T---P----G-----CQ--------LHD---AEQLWL-----D-----------------PLR--AQT-------------DETFLQ--HRL--RGD-WP-AEVGNRFANWLN-----RA----V---S------SD------S-Q-I-------------L---GSPEA------AQWSQ-E-----------LS-------------------K-----ELTMFKEIL------------------EDER------D------------------------
---------------
WP_074245424 -----------------------------------------------MTSPLPTP----------T--W---------QELR--QFIE--S-FIQ-ER-----L----QGK------------------------------------------------------------------------------------L---------D----------------K---------L--------HP-----DE----DD---------KR----QT---------------------LL------------------------------------------AT-----------------------
-----HR---REAW-----LA---DA-----ARRV-------G----QLQ-L-----------------VTHTLKPIHPD---A-----------------R---G-S------NL----H----S----L--------P-QAP----GQ--P--GL---------AGS--H--EL------G-D-----K--L--V-SD-V-VG----N----AA----A--L-----DVFKFL-S-----L----Q--Y--Q---------GKNL------------LNWLT-EDS-A---EAVQAL---S---D----NA----EQAREWRQAFIDI--T---AV--KG---
APASHSLAKQLYFPL-------PG-------------------SG---YHLLAPLFPTSLVHHVHAL-LR----E--A---R--F-G------------------DA-AKAAR-E-----AR------------N--RRESW--PH-G-FSE-----YPN-LAIQKFGG-T----KPQ---------NISQLN--SERY-----------GENWLLPSL-P----PHWQ--R-Q-DQ-RAPIHH----SSVF----EH---DFG--R-S--PEV--SRLTRTLQRLL-------------------A-----K-------------TRH-N-------
N----FT---IR--RY----RA------QLVGQICDEALQYAARLR--E-------L-E-----P----G----WS------A-----T---P----G-----CQ--------LHD---AEQLWL-----D-----------------PLR--AQT-------------DETFLQ--RRL--RGD-WP-AEVGNRFANWLN-----RA----V---S------S-----D--S-Q-I-------------L---GGPEA------AQWSQ-E-------LS---------------------K-----ELTMFKEIL------------------EDER------D------------------------
---------------
WP_073670440 -----------------------------------------------MTSPLPTP----------T--W---------QELR--QFIE--S-FIQ-ER-----L----QGK------------------------------------------------------------------------------------L---------D----------------K---------L--------HP-----DE----DD---------KR----QT---------------------LL------------------------------------------AT-----------------------
-----HR---REAW-----LA---DA-----ARRV-------G----QLQ-L-----------------VTHTLKPIHPD---A-----------------R---G-S------NL----H----S----L--------P-QAP----GQ--P--GL---------AGS--H--EL------G-D-----K--L--V-SD-V-AG----N----AA----A--L-----DVFKFL-S-----L----Q--Y--Q---------GKNL------------LNWLT-EDS-A---EAVQAL---S---D----NA----EQAREWRQAFIDI--T---AV--KG---
APASHSLAKQLYFPL-------PG-------------------SG---YHLLAPLFPTSLVHHVHAL-LR----E--A---R--F-G------------------DA-AKAAR-E-----AR------------N--RRESW--PH-G-FSE-----YPN-LAIQKFGG-T----KPQ---------NISQLN--SERY-----------GENWLLPSL-P----PHWQ--R-Q-DQ-RAPIHH----SSVF----EH---DFG--R-S--PEV--SRLTRTLQRLL-------------------A-----K-------------TRH-N-------
N----FT---IR--RY----RA------QLVGQICDEALQYAARLR--E-------L-E-----P----G----WS------A-----T---P----G-----CQ--------LHD---AEQLWL-----D-----------------PLR--AQT-------------DETFLQ--RRL--RGD-WP-AEVGNRFANWLN-----RA----V---S------S-----D--S-Q-I-------------L---GSPEA------AQWSQ-E-------LS---------------------K-----ELTMFKEIL------------------EDER------D------------------------
---------------
WP_034009772 -----------------------------------------------MTSPLPTP----------T--W---------QELR--QFIE--S-FIQ-ER-----L----QGK------------------------------------------------------------------------------------L---------D----------------K---------L--------HP-----DE----DD---------KR----QT---------------------LL------------------------------------------AT-----------------------
-----HR---REAW-----LA---DA-----ARRV-------G----QLQ-L-----------------VTHTLKPIHPD---A-----------------R---G-S------NL----H----S----L--------P-QAP----GQ--P--GL---------AGS--H--EL------G-D-----K--L--V-SD-V-VG----N----AA----A--L-----DVFKFL-S-----L----Q--Y--Q---------GKNL------------LNWLT-EDS-A---EAVQAL---S---D----NA----EQAREWRQAFIDI--T---AV--KG---
APASHSLAKQLYFPL-------PG-------------------SG---YHLLAPLFPTSLVHHVHAL-LR----E--A---R--F-G------------------DA-AKAAR-E-----AR------------N--RRESW--PH-G-FSE-----YPN-LAIQKFGG-T----KPQ---------NISQLN--SERY-----------GENWLLPSL-P----PHWQ-RQ---DQ-RAPIHH----SSVF----EH---DFG--R-S--PEV--SRLTRTLQRLL------A--KT----R-H-----N-----------------------------------
N----FT---IR--RY----RA------QLVGQICDEALQYAARLR--E-------L-E-----P----G----WS------A-----T---P----G-----CQ--------LHD---AEQLWL-----D-----------------PLR--AQT-------------DETFLQ--RRL--RGD-WP-AEVGNRFANWLN-----RA----V---S------SD------S-Q-I-------------L---GSPEA------AQWSQ-E-----------LS-------------------K-----ELTMFKEIL------------------EDER------D------------------------
---------------
WP_049321720 -----------------------------------------------MTSPLPTP----------T--W---------QELR--QFIE--S-FIQ-ER-----L----QGK------------------------------------------------------------------------------------L---------D----------------K---------L--------HP-----DE----DD---------KR----QT---------------------LL------------------------------------------AT-----------------------
-----HR---REAW-----LA---DA-----ARRV-------G----QLQ-L-----------------VTHTLKPIHPD---A-----------------R---G-S------NL----H----S----L--------P-QAP----GQ--P--GL---------AGS--H--EL------G-D-----R--L--V-SD-V-VG----N----AA----A--L-----DVFKFL-S-----L----Q--Y--Q---------GKNL------------LNWLT-EDS-A---EAVQAL---S---D----NA----EQAREWRQAFIGI--T---AV--KD---
APASHSLAKQLYFPL-------PG-------------------SG---YHLLAPLFPTSLVHHVHAL-LR----E--A---R--F-G------------------DA-ARAAR-E-----AR------------S--RQESW--PH-G-FSE-----YPN-LAIQKFGG-T----KPQ---------NISQLN--SERY-----------GENWLLPSL-P----PHWQ--R-Q-DQ-RAPIRH----SSVF----EH---DFG--R-S--PEV--SRLTRTLQRLL-------------------A-----K-------------TRH-N-------
N----FT---IR--RY----RA------QLVGQICDEALQYAARLR--E-------L-E-----P----G----WS------A-----T---P----G-----CQ--------LHD---AEQLWL-----D-----------------PLR--AQT-------------DETFLQ--RRL--RGD-WP-AEVGNRFANWLN-----RA----V---S------S-----D--S-Q-I-------------L---GSPEA------AQWSQ-E-------LS---------------------K-----ELTMFKEIL------------------EDER------D------------------------
---------------
WP_134308262 -----------------------------------------------MTSPLPTP----------T--W---------QELR--QFIE--S-FIQ-ER-----L----QGK------------------------------------------------------------------------------------L---------D----------------K---------L--------HP-----DE----DD---------KR----QT---------------------LL------------------------------------------AT-----------------------
-----HR---REAW-----LA---DA-----ARRV-------G----QLQ-L-----------------VTHTLKPIHPD---A-----------------R---G-S------NL----H----S----L--------P-QAP----GQ--P--GL---------AGS--H--EL------G-D-----R--L--V-SD-V-VG----N----AA----A--L-----DVFKFL-S-----L----Q--Y--Q---------GKNL------------LNWLT-EDS-A---EAVQAL---S---D----NA----EQAREWRQAFIGI--T---AV--KG---
APASHSLAKQLYFPL-------PG-------------------SG---YHLLAPLFPTSLVHHVHAL-LR----E--A---R--F-G------------------DA-AKAAR-E-----AR------------S--RQESW--PH-G-FSE-----YPN-LAIQKFGG-T----KPQ---------NISQLN--SERY-----------GENWLLPSL-P----PHWQ--R-Q-DQ-RAPIRH----SSVF----EH---DFG--R-S--PEV--SRLTRTLQRLL-------------------A-----K-------------TRD-N-------
N----FT---IR--RY----RA------QLVGQICDEALQYAARLR--E-------L-E-----P----G----WS------A-----T---P----G-----CQ--------LHD---AEQLWL-----D-----------------PLR--AQT-------------DETFLQ--RRL--RGD-WP-AEVGNRFANWLN-----RA----V---S------S-----D--S-Q-I-------------L---GSPEA------AQWSQ-E-------LS---------------------K-----ELTMFKEIL------------------EDER------D------------------------
---------------
WP_034052100 -----------------------------------------------MTSPLPTP----------T--W---------QELR--QFIE--S-FIQ-ER-----L----QGK------------------------------------------------------------------------------------L---------D----------------K---------L--------HP-----DE----DD---------KR----QT---------------------LL------------------------------------------AT-----------------------
-----HR---REAW-----LA---DA-----ARRV-------G----QLQ-L-----------------VTHTLKPIHPD---A-----------------R---G-S------NL----H----S----L--------P-QAP----GQ--P--GL---------AGS--H--EL------G-D-----R--L--V-SD-V-VG----N----AA----A--L-----DVFKFL-S-----L----Q--Y--Q---------GKNL------------LNWLT-EDS-A---EAVQAL---S---D----NA----EQAREWRQAFIGI--T---AV--KG---
APASHSLAKQLYFPL-------PG-------------------SG---YHLLAPLFPTSLVHHVHAL-LR----E--A---R--F-G------------------DA-AKAAR-E-----AR------------S--RQESW--PH-G-FSE-----YPN-LAIQKFGG-T----KPQ---------NISQLN--SERY-----------GENWLLPSL-P----PHWQ-RQ---DQ-RAPIRH----SSVF----EH---DFG--R-S--PEV--SRLTRTLQRLL------A--KT----R-D-----N-----------------------------------
N----FT---IR--RY----RA------QLLGQICDEALQYAARLR--E-------L-E-----P----G----WS------A-----T---P----G-----CQ--------LHD---AEQLWL-----D-----------------PLR--AQT-------------DETFLQ--RRL--RGD-WP-AEVGNRFANWLN-----RA----V---S------SD------S-Q-I-------------L---GSPEA------AQWSQ-E-----------LS-------------------K-----ELTMFKEIL------------------EDER------D------------------------
---------------
WP_004349919 -----------------------------------------------MTSPLPTP----------T--W---------QELR--QFIE--S-FIQ-ER-----L----QGK------------------------------------------------------------------------------------L---------D----------------K---------L--------HP-----NE----DD---------KR----QT---------------------LL------------------------------------------AT-----------------------
-----HR---REAW-----LA---DA-----ARRV-------G----QLQ-L-----------------VTHTLKPIHPD---A-----------------R---G-S------NL----H----S----L--------P-QAP----GQ--P--GL---------AGS--H--EL------G-D-----R--L--V-SD-V-VG----N----AA----A--L-----DVFKFL-S-----L----Q--Y--Q---------GKNL------------LNWLT-EDS-A---EAVQAL---S---D----NA----EQAREWRQAFIGI--T---AV--KG---
APASHSLAKQLYFPL-------PG-------------------SG---YHLLAPLFPTSLVHHVHAL-LR----E--A---R--F-G------------------DA-AKAAR-E-----AR------------S--RQESW--PH-G-FSE-----YPN-LAIQKFGG-T----KPQ---------NISQLN--SERY-----------GENWLLPSL-P----PHWQ--R-Q-DQ-RAPIRH----SSVF----EH---DFG--R-S--PEV--SRLTRTLQRLL-------------------A-----K-------------TRD-N-------
N----FT---IR--RY----RA------QLVGQICDEALQYAARLR--E-------L-E-----P----G----WS------A-----T---P----G-----CQ--------LHD---TEQLWL-----D-----------------PLR--AQT-------------DETFLQ--RRL--RGD-WP-AEVGNRFANWLN-----RA----V---S------S-----D--S-Q-I-------------L---GSPEA------AQWSQ-E-------LS---------------------K-----ELTMFKEIL------------------EDER------D------------------------
---------------
WP_134284832 -----------------------------------------------MTSPLPTP----------T--W---------QELR--QFIE--S-FIQ-ER-----L----QGK------------------------------------------------------------------------------------L---------D----------------K---------L--------HP-----DE----DD---------KR----QT---------------------LL------------------------------------------AT-----------------------
-----HR---REAW-----LA---DA-----ARRV-------G----QLQ-L-----------------VTHTLKPIHPD---A-----------------R---G-S------NL----H----S----L--------P-QAP----GQ--P--GL---------AGS--H--EL------G-D-----R--L--V-SD-V-VG----N----AA----A--L-----DVFKFL-S-----L----Q--Y--Q---------GKNL------------LNWLT-EDS-A---EAVQAL---S---D----NA----EQAREWRQAFIGI--T---AV--KG---
APASHSLAKQLYFPL-------PG-------------------SG---YHLLAPVFPTSLVHHVHAL-LR----E--A---R--F-G------------------DA-AKAAR-E-----AR------------S--RQESW--PH-G-FSE-----YPN-LAIQKFGG-T----KPQ---------NISQLN--SERY-----------GENWLLPSL-P----PHWQ--R-Q-DQ-RAPIRH----SSVF----EH---DFG--R-S--PEV--SRLTRTLQRLL-------------------A-----K-------------TRD-N-------
N----FT---IR--RY----RA------QLVGQICDEALQYAARLR--E-------L-E-----P----G----WS------A-----T---P----G-----CQ--------LHD---TEQLWL-----D-----------------PLR--AQT-------------DETFLQ--RRL--RGD-WP-AEVGNRFANWLN-----RA----V---S------S-----D--S-Q-I-------------L---GSPEA------AQWSQ-E-------LS---------------------K-----ELTMFKEIL------------------EDER------D------------------------
---------------
WP_124205178 -----------------------------------------------MTSPLPTP----------T--W---------QELR--QFIE--S-FIQ-ER-----L----QGK------------------------------------------------------------------------------------L---------D----------------K---------L--------HP-----DE----DD---------KR----QT---------------------LL------------------------------------------AT-----------------------
-----HR---REAW-----LA---DA-----ARRV-------G----QLQ-L-----------------VTHTLKPIHPD---A-----------------R---G-S------NL----H----S----L--------P-QAP----GQ--P--GL---------AGS--H--EL------G-D-----R--L--V-SD-V-VG----N----AA----A--L-----DVFKFL-S-----L----Q--Y--Q---------GKNL------------LNWLT-EDS-A---EAVQAL---S---D----NA----EQAREWRQAFIGI--T---AV--KG---
APASHSLAKQLYFPL-------PG-------------------SG---YHLLAPLFPTSLVHHVHAL-LR----E--A---R--F-G------------------DA-AKAAR-E-----AR------------S--RQESW--PH-G-FSE-----YPN-LATQKFGG-T----KPQ---------NISQLN--SERY-----------GENWLLPSL-P----PHWQ--R-Q-DQ-RAPIRH----SSVF----EH---DFG--R-S--PEV--SRLTRTLQRLL-------------------A-----K-------------TRD-N-------
N----FT---IR--RY----RA------QLVGQICDEALQYAARLR--E-------L-E-----P----G----WS------A-----T---P----G-----CQ--------LHD---TEQLWL-----D-----------------PLR--AQT-------------DETFLQ--RRL--RGD-WP-AEVGNRFANWLN-----RA----V---S------S-----D--S-Q-I-------------L---GSPEA------AQWSQ-E-------LS---------------------K-----ELTMFKEIL------------------EDER------D------------------------
---------------
WP_058151550 -----------------------------------------------MTSPLPTP----------T--W---------QELR--QFIE--S-FIQ-ER-----L----QGK------------------------------------------------------------------------------------L---------D----------------K---------L--------HP-----DE----DD---------KR----QT---------------------LL------------------------------------------AT-----------------------
-----HR---REAW-----LA---DA-----ARRV-------G----QLQ-L-----------------VTHTLKPIHPD---A-----------------R---G-S------NL----H----S----L--------P-QAP----GQ--P--GL---------AGS--H--EL------G-D-----R--L--V-SD-V-VG----N----AA----A--L-----DVFKFL-S-----L----Q--Y--Q---------GKNL------------LNWLT-EDS-A---EAVQAL---S---D----NA----EQAREWRQAFIGI--T---AV--KG---
APASHSLAKQLYFPL-------PG-------------------SG---YHLLAPLFPTSLVHHVHAL-LR----E--A---R--F-G------------------DA-AKAAR-E-----AR------------S--RQESW--PH-G-FSE-----YPN-LAIQKFGG-T----KPQ---------NISQLN--SERY-----------GENWLLPSL-P----PHWQ--R-Q-DQ-RAPIRH----SSVF----EH---DFG--R-S--PEV--SRLTRTLQRLL-------------------A-----K-------------TRD-N-------
N----FT---IR--RY----RA------HLVGQICDEALQYAARLR--E-------L-E-----P----G----WS------A-----T---P----G-----CQ--------LHD---TEQLWL-----D-----------------PLR--AQT-------------DETFLQ--RRL--RGD-WP-AEVGNRFANWLN-----RA----V---S------S-----D--S-Q-I-------------L---GSPEA------AQWSQ-E-------LS---------------------K-----ELTMFKEIL------------------EDER------D------------------------
---------------
WP_124130359 -----------------------------------------------MTSPLPTP----------T--W---------QELR--QFIE--S-FIQ-ER-----L----QGK------------------------------------------------------------------------------------L---------D----------------K---------L--------HP-----DE----DD---------KR----QT---------------------LL------------------------------------------AT-----------------------
-----HR---REAW-----LA---DA-----ARRV-------G----QLQ-L-----------------VTHTLKPIHPD---A-----------------R---G-S------NL----H----S----L--------P-QAP----GQ--P--GL---------AGS--H--EL------G-D-----R--L--V-SD-V-VG----N----AA----A--L-----DVFKFL-S-----L----Q--Y--Q---------GKNL------------LNWLT-EDS-A---EAVQAL---S---D----NA----EQAREWRQAFIGI--T---AV--KG---
APASHSLAKQLYFPL-------PG-------------------SG---YHLLAPLFPTSLVHHVHAL-LR----E--A---R--F-G------------------DA-AKAAR-E-----AR------------S--RQESW--PH-G-FSE-----YPN-LAIQKFGG-T----KPQ---------NISQLN--SERY-----------GENWLLPXL-P----PHWQ--R-Q-DQ-RAPIRH----SSVF----EH---DFG--R-S--PEV--SRLTRTLQRLL-------------------A-----K-------------TRD-N-------
N----FT---IR--RY----RA------QLVGQICDEALQYAARLR--E-------L-E-----P----G----WS------A-----T---P----G-----CQ--------LHD---TEQLWL-----D-----------------PLR--AQT-------------DETFLQ--RRL--RGD-WP-AEVGNRFANWLN-----RA----V---S------S-----D--S-Q-I-------------L---GSPEA------AQWSQ-E-------LS---------------------K-----ELTMFKEIL------------------EDER------D------------------------
---------------
WP_021205340 -----------------------------------------------MTSPLPTP----------T--W---------QELR--QFIE--S-FIQ-ER-----L----QGK------------------------------------------------------------------------------------L---------D----------------K---------L--------HP-----DE----DD---------KR----QT---------------------LL------------------------------------------AT-----------------------
-----HR---REAW-----LA---DA-----ARRV-------G----QLQ-L-----------------VTHTLKPIHPD---A-----------------R---G-S------NL----H----S----L--------P-QAP----GQ--P--GL---------AGS--H--EL------G-D-----R--L--V-SD-V-VG----N----AA----A--L-----DVFKFL-S-----L----Q--Y--Q---------GKNL------------LNWLT-EDS-A---EAVQAL---S---D----NA----EQAREWRQAFIGI--T---AV--KG---
APASHSLAKQLYFPL-------PG-------------------SG---YHLLAPLFPTSLVHHVHAL-LR----E--A---R--F-G------------------DA-AKAAR-E-----AR------------S--RQESW--PH-G-FSE-----YPN-LAIQKFGG-T----KPQ---------NISQLN--SERY-----------GENWLLPSL-P----PHWQ-RQ---DQ-RAPIRH----SSVF----EH---DFG--R-S--PEV--SRLTRTLQRLL------A--KT----R-D-----N-----------------------------------
N----FT---IR--RY----RA------QLVGQICDEALQYAARLR--E-------L-E-----P----G----WS------A-----T---P----G-----CQ--------LHD---TEQLWL-----D-----------------PLR--AQT-------------DETFLQ--RRL--RGD-WP-AEVGNRFANWLN-----RA----V---S------SD------S-Q-I-------------L---GSPEA------AQWSQ-E-----------LS-------------------K-----ELTMFKEIL------------------EDER------D------------------------
---------------
WP_128654221 -----------------------------------------------MTSPLPTP----------T--W---------QELR--QLIE--S-FIQ-ER-----L----QGK------------------------------------------------------------------------------------L---------D----------------K---------L--------HP-----DE----DD---------KR----QT---------------------LL------------------------------------------AT-----------------------
-----HR---REAW-----LA---DA-----ARRV-------G----QLQ-L-----------------VTHTLKPIHPD---A-----------------R---G-S------NL----H----S----L--------P-QAP----GQ--P--GL---------AGS--H--EL------G-D-----R--L--V-SD-V-VG----N----AA----A--L-----DVFKFL-S-----L----Q--Y--Q---------GKNL------------LNWLT-EDS-A---EAVQAL---S---D----NA----EQAREWRQAFIGI--T---AV--KG---
APASHSLXKQLYFPL-------PG-------------------SG---YHLLAPLFPTSLVHHVHAL-LR----E--A---R--F-G------------------DA-AKAAR-E-----AR------------S--RQESW--PH-G-FSE-----YPN-LAIQKFGG-T----KPQ---------NISQLN--SERY-----------GENWLLPSL-P----PHWQ--R-Q-DQ-RAPIRH----SSVF----EH---DFG--R-S--PEV--SRLTRTLQRLL-------------------A-----K-------------TRH-N-------
N----FT---IR--RY----RA------QLVGQICDEALQYAARLR--E-------L-E-----P----G----WS------A-----T---P----G-----CQ--------LHD---AEQLWL-----D-----------------PLR--AQT-------------DETFLQ--RRL--RGD-WP-AEVGNRFANWLN-----RA----V---S------S-----D--S-Q-I-------------L---GSPEA------AQWSQ-E-------LS---------------------K-----ELTMFKEIL------------------EDER------D------------------------
---------------
WP_073653177 -----------------------------------------------MTSPLPTP----------T--W---------QELR--QLIE--S-FIQ-ER-----L----QGK------------------------------------------------------------------------------------L---------D----------------K---------L--------HP-----DE----DD---------KR----QT---------------------LL------------------------------------------AT-----------------------
-----HR---REAW-----LA---DA-----ARRV-------G----QLQ-L-----------------VTHTLKPIHPD---A-----------------R---G-S------NL----H----S----L--------P-QAP----GQ--P--GL---------AGS--H--EL------G-D-----R--L--V-SD-V-VG----N----AA----A--L-----DVFKFL-S-----L----Q--Y--Q---------GKNL------------LNWLT-EDS-A---EAVQAL---S---D----NA----EQAREWRQAFIGI--T---AV--KG---
APASHSLAKQLYFPL-------PG-------------------SG---YHLLAPLFPTSLVHHVHAL-LR----E--A---R--F-G------------------DA-AKAAR-E-----AR------------S--RQESW--PH-G-FSE-----YPN-LAIQKFGG-T----KPQ---------NISQLN--SERY-----------GENWLLPSL-P----PHWQ-RQ---DQ-RAPIRH----SSVF----EH---DFG--R-S--PEV--SRLTRTLQRLL------A--KT----R-H-----N-----------------------------------
N----FT---IR--RY----RA------QLVGQICDEALQYAARLR--E-------L-E-----P----G----WS------A-----T---P----G-----CQ--------LHD---AEQLWL-----D-----------------PLR--AQT-------------DETFLQ--RRL--RGD-WP-AEVGNRFANWLN-----RA----V---S------SD------S-Q-I-------------L---GSPEA------AQWSQ-E-----------LS-------------------K-----ELTMFKEIL------------------EDER------D------------------------
---------------
WP_033940279 -----------------------------------------------MTSPLPTP----------T--W---------QELR--QFIE--S-FIQ-ER-----L----QGK------------------------------------------------------------------------------------L---------D----------------K---------L--------HP-----DE----DD---------KR----QT---------------------LL------------------------------------------AT-----------------------
-----HR---REAW-----LA---DA-----ARRV-------G----QLQ-L-----------------VTHTLKPIHPD---A-----------------R---G-S------NL----H----S----L--------P-QAP----GQ--P--GL---------AGS--H--EL------G-D-----R--L--V-SD-V-VG----N----AA----A--L-----DVFKFL-S-----L----Q--Y--Q---------GKNL------------LNWLT-EDS-A---EAVQAL---S---D----NA----EQAREWRQAFIGI--T---AV--KG---
APASHSLAKQLYFPL-------PG-------------------SG---YHLLAPLFPTSLVHHVHAL-LR----E--A---R--F-G------------------DA-AKAAR-E-----AR------------S--RQESW--PH-G-FSE-----YPN-LAIQKFGG-T----KPQ---------NISQLN--SERY-----------GENWLLPSL-P----PHWQ--R-Q-DQ-RAPIRH----SSVF----EH---DFG--R-S--PEV--SRLTRTLQRLL-------------------A-----K-------------TRH-N-------
N----FT---IR--RY----RA------QLVGQICDEALQYAARLR--E-------L-E-----P----G----WS------A-----T---P----G-----CQ--------LHD---AEQLWL-----D-----------------PLR--AQT-------------DETFLQ--RRL--RGD-WP-AEVGNRFANWLN-----RA----V---S------S-----D--R-Q-I-------------L---GSPEA------AQWSQ-E-------LS---------------------K-----ELTMFKEIL------------------EDER------D------------------------
---------------
WP_031686740 -----------------------------------------------MTSPLPTP----------T--W---------QELR--QFIE--S-FIQ-ER-----L----QGK------------------------------------------------------------------------------------L---------D----------------K---------L--------HP-----DE----DD---------KR----QT---------------------LL------------------------------------------AT-----------------------
-----HR---REAW-----LA---DA-----ARRV-------G----QLQ-L-----------------VTHTLKPIHPD---A-----------------R---G-S------NL----H----S----L--------P-QAP----GQ--P--GL---------AGS--H--EL------G-D-----R--L--V-SD-V-VG----N----AA----A--L-----DVFKFL-S-----L----Q--Y--Q---------GKNL------------LNWLT-EDS-A---EAVQAL---S---D----NA----EQAREWRQAFIGI--T---AV--KG---
APASHSLAKQLYFPL-------PG-------------------SG---YHLLAPLFPTSLVHHVHAL-LR----E--A---R--F-G------------------DA-AKAAR-E-----AR------------S--RQESW--PH-G-FSE-----YPN-LAIQKFGG-T----KPQ---------NISQLN--SERY-----------GENWLLPSL-P----PHWQ-RQ---DQ-RAPIRH----SSVF----EH---DFG--R-S--PEV--SRLTRTLQRLL------A--KT----R-H-----N-----------------------------------
N----FT---IR--RY----RA------QLVGQICDEALQYAARLR--E-------L-E-----P----G----WS------A-----T---P----G-----CQ--------LHD---AEQLWL-----D-----------------PLR--AQT-------------DETFLQ--RRL--RGD-WP-AEVGNRFANWLN-----RA----V---S------SD------T-Q-I-------------L---GSPEA------AQWSQ-E-----------LS-------------------K-----ELTMFKEIL------------------EDER------D------------------------
---------------
WP_014833996 -----------------------------------------------MTSPLPTP----------T--W---------QELR--QFIE--S-FIQ-ER-----L----QGK------------------------------------------------------------------------------------L---------D----------------K---------L--------HP-----DE----DD---------KR----QT---------------------LL------------------------------------------AT-----------------------
-----HR---REAW-----LA---DA-----ARRV-------G----QLQ-L-----------------VTHTLKPIHPD---A-----------------R---G-S------NL----H----S----L--------P-QAP----GQ--P--GL---------AGS--H--EL------G-D-----R--L--V-SD-V-VG----N----AA----A--L-----DVFKFL-S-----L----Q--Y--Q---------GKNL------------LNWLT-EDS-A---EAVQAL---S---D----NA----EQAREWRQAFIGI--T---AV--KG---
APASHSLAKQLYFPL-------PG-------------------SG---YHLLAPLFPTSLVHHVHAL-LR----E--A---R--F-G------------------DA-AKAAR-E-----AR------------S--RQESW--PH-G-FSE-----YPN-LAIQKFGG-T----KPQ---------NISQLN--SERY-----------GENWLLPSL-P----PHWQ--R-Q-DQ-RAPIRH----SSVF----EH---DFG--R-S--PEV--SRLTRTLQRLL-------------------A-----K-------------TRH-N-------
N----FT---IR--RY----RA------QLVGQICDEALQYAARLR--E-------L-E-----P----G----WS------A-----T---P----G-----CQ--------LHD---AEQLWL-----D-----------------PLR--AQT-------------DETFLQ--RRL--RGD-WP-AEVGNRFANWLN-----RA----V---S------S-----D--S-Q-I-------------L---GSPEA------AQWSP-E-------LS---------------------K-----ELTMFKEIL------------------EDER------D------------------------
---------------
WP_124167688 -----------------------------------------------MTSPLPTP----------T--W---------QELR--QFIE--S-FIQ-ER-----L----QGK------------------------------------------------------------------------------------L---------D----------------K---------L--------HP-----DE----DD---------KR----QT---------------------LL------------------------------------------AT-----------------------
-----HR---REAW-----LA---DA-----ARRV-------G----QLQ-L-----------------VTHTLKPIHPD---A-----------------R---G-S------NL----H----S----L--------P-QAP----GQ--P--GL---------AGS--H--EL------G-D-----R--L--V-SD-V-VG----N----AA----A--L-----DVFKFL-S-----L----Q--Y--Q---------GKNL------------LNWLT-EDS-A---EAVQAL---S---D----NA----EQAREWRQAFIGI--T---AV--KG---
APASHSLAKQLYFPL-------PG-------------------SG---YHLLAPLFPTSLVHHVHAL-LR----E--A---R--F-G------------------DA-AKAAR-E-----AR------------S--RQESW--PH-G-FSE-----YPN-LAIQKFGG-T----KPQ---------NISQLN--SERY-----------GENWLLPSL-P----PHWQ-RQ---DQ-RAPIRH----SSVF----EH---DFG--R-S--PEV--SRLTRTLQRLL------A--KT----R-H-----N-----------------------------------
N----FT---IR--RY----RA------QLVGQICDEALQYAARLR--E-------L-E-----P----G----WS------A-----T---P----G-----CQ--------LHD---AEQLWL-----D-----------------PLR--AQT-------------DETFLQ--RRL--RGD-WP-AEVGNRFANWLN-----RA----V---S------SD------S-Q-I-------------L---GSPEA------TQWSP-E-----------LS-------------------K-----ELTMFKEIL------------------EDER------D------------------------
---------------
WP_126621867 ----------------------------------------------------------------------------PFIE--S-FIQ-ER-----L----QGK------------------------------------------------------------------------------------L---------D----------------K---------L--------HP-----DE----DD---------KR----QT---------------------LL------------------------------------------AT----------------------------------------------HR---
REAW-----LA---DA-----ARRV-------G----QLQ-L-----------------VTHTLKPIHPD---A-----------------R---G-S------NL----H----S----L--------P-QAP----GQ--P--GL---------AGS--H--EL------G-D-----R--L--V-SD-V-VG----N----AA----A--L-----DVFKFL-S-----L----Q--Y--Q---------GKNL------------LNWLT-EDS-A---EAVQAL---S---D----NA----EQAREWRQAFIGI--T---AV--KG---APASHSLAKQLYFPL--
-----PG-------------------SG---YHLLAPLFPTSLVHHVHAL-LR----E--A---R--F-G------------------DA-AKAAR-E-----AR------------S--RQESW--PH-G-FSE-----YPN-LAIQKFGG-T----KPQ---------NISQLN--SERY-----------GENWLLPSL-P----PHWQ--R-Q-DQ-RAPIRH----SSVF----EH---DFG--R-S--PEV--SRLTRTLQRLL-------------------A-----K-------------TRH-N-------N----FT---IR--RY----RA--
----QLVGQICDEALQYAARLR--E-------L-E-----P----G----WS------A-----T---P----G-----CQ--------LHD---AEQLWL-----D-----------------PLR--AQT-------------DETFLQ--RRL--RGD-WP-AEVGNRFANWLN-----RA----V---S------S-----D--S-Q-I-------------L---GSPEA------AQWSQ-E-------LS---------------------K-----ELTMFKEIL------------------EDER------D----------------------
WP_126671529 -----------------------------------------------MTSPLPTP----------T--W---------QELR--QFIE--S-FIQ-ER-----L----QGK------------------------------------------------------------------------------------L---------D----------------K---------L--------HP-----DE----DD---------KR----QT---------------------LL------------------------------------------AT-----------------------
-----HR---REAW-----LA---DA-----ARRV-------G----QLQ-L-----------------VTHTLKPIHPD---A-----------------R---G-S------NL----H----S----L--------P-QAP----GQ--P--GL---------AGS--H--EL------G-D-----R--L--V-SD-V-VG----N----AA----A--L-----DVFKFL-G-----L----Q--Y--Q---------GKNL------------LNWLT-EDS-A---EAVQAL---S---D----NA----EQAREWRQAFIGI--T---AV--KG---
APASHSLAKQLYFPL-------PG-------------------SG---YHLLAPLFPTSLVHHVHAL-LR----E--A---R--F-G------------------DA-AKAAR-E-----AR------------S--RQESW--PH-G-FSE-----YPN-LAIQKFGG-T----KPQ---------NISQLN--SERY-----------GENWLLPSL-P----PHWQ--R-Q-DQ-RAPIRH----SSVF----EH---DFG--R-S--PEV--SRLTRTLQRLL-------------------A-----K-------------TRH-N-------
N----FT---IR--RY----RA------QLVGQICDEALQYAARLR--E-------L-E-----P----G----WS------A-----T---P----G-----CQ--------LHD---AEQLWL-----D-----------------PLR--AQT-------------DETFLQ--RRL--RGD-WP-AEVGNRFANWLN-----RA----V---S------S-----D--S-Q-I-------------L---GSPEA------AQWSQ-E-------LS---------------------K-----ELTMFKEIL------------------EDER------D------------------------
---------------
WP_070623801 -----------------------------------------------MTSPLPTP----------T--W---------QELR--QFIE--S-FIQ-ER-----L----QGK------------------------------------------------------------------------------------L---------D----------------K---------L--------HP-----DE----DD---------KR----QT---------------------LL------------------------------------------AT-----------------------
-----HR---REAW-----LA---DA-----ARRV-------G----QLQ-L-----------------VTHTLKPIHPD---A-----------------R---G-S------NL----H----S----L--------P-QAP----GQ--P--GL---------AGS--H--EL------G-D-----R--L--V-SD-V-VG----N----AA----A--L-----DVFKFL-S-----L----Q--Y--Q---------GKNL------------LNWLT-EDS-A---EAVQAL---S---D----NA----EQAREWRQAFIGI--T---AV--KG---
APASHSLAKQLYFPL-------PG-------------------SG---YHLLAPLFPTSLVHHVHAL-LR----E--A---R--F-G------------------DA-AKAAR-E-----AR------------S--RQESW--PH-G-FSE-----YPN-LAIQKFGG-T----KPQ---------NISQLN--SERY-----------GENWLLPSL-P----PHWQ--R-Q-DQ-RAPIRH----SSVF----EH---DFG--R-S--PEV--SRLTRTLQRLL-------------------A-----K-------------TRH-N-------
N----FT---IR--RY----RA------QLVGQICDEALQYAARLR--E-------L-E-----P----G----WS------A-----T---P----G-----CQ--------LHV---AEQLWL-----D-----------------PLR--AQT-------------DETFLQ--RRL--RGD-WP-AEVGNRFANWLN-----RA----V---S------S-----D--S-Q-I-------------L---GSPEA------AQWSQ-E-------LS---------------------K-----ELTMFKEIL------------------EDER------D------------------------
---------------
WP_128446510 -----------------------------------------------MTSPLPTP----------T--W---------QELR--QFIE--S-FIQ-ER-----L----QGK------------------------------------------------------------------------------------L---------D----------------K---------L--------HP-----DE----DD---------KR----QT---------------------LL------------------------------------------AT-----------------------
-----HR---REAW-----LA---DA-----ARRV-------G----QLQ-L-----------------VTHTLKPIHPD---A-----------------R---G-S------NL----H----S----L--------P-QAP----GQ--P--GL---------AGS--H--EL------G-D-----R--L--V-SD-V-VG----N----AA----A--L-----DVFKFL-S-----L----Q--Y--Q---------GKKL------------LNWLT-EDS-A---EAVQAL---S---D----NA----EQAREWRQAFIGI--T---AV--KG---
APASHSLAKQLYFPL-------PG-------------------SG---YHLLAPLFPTSLVHHVHAL-LR----E--A---R--F-G------------------DA-AKAAR-E-----AR------------S--RQESW--PH-G-FSE-----YPN-LAIQKFGG-T----KPQ---------NISQLN--SERY-----------GENWLLPSL-P----PHWQ--R-Q-DQ-RAPIRH----SSVF----EH---DFG--R-S--PEV--SRLTRTLQRLL-------------------A-----K-------------TRH-N-------
N----FT---IR--RY----RA------QLVGQICDEALQYAARLR--E-------L-E-----P----G----WS------A-----T---P----G-----CQ--------LHD---AEQLWL-----D-----------------PLR--AQT-------------DETFLQ--RRL--RGD-WP-AEVGNRFANWLN-----RA----V---S------S-----D--S-Q-I-------------L---GSPEA------AQWSQ-E-------LS---------------------K-----ELTMFKEIL------------------EDER------D------------------------
---------------
WP_126123111 -----------------------------------------------MTSPLPTP----------T--W---------QELR--QFIE--S-FIQ-ER-----L----QGK------------------------------------------------------------------------------------L---------D----------------K---------L--------HP-----DE----DD---------KR----QT---------------------LL------------------------------------------AT-----------------------
-----HR---REAW-----LA---DA-----ARRV-------G----QLQ-L-----------------VTHTLKPIHPD---A-----------------R---G-S------NL----H----S----L--------P-QAP----GQ--P--GL---------AGS--H--EL------G-D-----R--L--V-SD-V-VG----N----AA----A--L-----DVFKFL-S-----L----Q--Y--Q---------GKNL------------LNWLT-EDS-A---EAVQAL---S---D----NA----EQAREWRQAFIGI--T---AV--KG---
APASHSLAKQLYFPL-------PG-------------------SG---YHLLAPLLPTSLVHHVHAL-LR----E--A---R--F-G------------------DA-AKAAR-E-----AR------------S--RQESW--PH-G-FSE-----YPN-LAIQKFGG-T----KPQ---------NISQLN--SERY-----------GENWLLPSL-P----PHWQ--R-Q-DQ-RAPIRH----SSVF----EH---DFG--R-S--PEV--SRLTRTLQRLL-------------------A-----K-------------TRH-N-------
N----FT---IR--RY----RA------QLVGQICDEALQYAARLR--E-------L-E-----P----G----WS------A-----T---P----G-----CQ--------LHD---AEQLWL-----D-----------------PLR--AQT-------------DETFLQ--RRL--RGD-WP-AEVGNRFANWLN-----RA----V---S------S-----D--S-Q-I-------------L---GSPEA------AQWSQ-E-------LS---------------------K-----ELTMFKEIL------------------EDER------D------------------------
---------------
WP_126676667 -----------------------------------------------MTSPLPTP----------T--W---------QELR--QFIE--S-FIQ-ER-----L----QGK------------------------------------------------------------------------------------L---------D----------------K---------L--------HP-----DE----DD---------KR----QT---------------------LL------------------------------------------AT-----------------------
-----HR---REAW-----LA---DA-----ARRV-------G----QLQ-L-----------------VTHTLKPIHPD---A-----------------R---G-S------NL----H----S----L--------P-QAP----GQ--P--GL---------AGS--H--EL------G-G-----R--L--V-SD-V-VG----N----AA----A--L-----DVFKFL-S-----L----Q--Y--Q---------GKNL------------LNWLT-EDS-A---EAVQAL---S---D----NA----EQAREWRQAFIGI--T---AV--KG---
APASHSLAKQLYFPL-------PG-------------------SG---YHLLAPLFPTSLVHHVHAL-LR----E--A---R--F-G------------------DA-AKAAR-E-----AR------------S--RQESW--PH-G-FSE-----YPN-LAIQKFGG-T----KPQ---------NISQLN--SERY-----------GENWLLPSL-P----PHWQ--R-Q-DQ-RAPIRH----SSVF----EH---DFG--R-S--PEV--SRLTRTLQRLL-------------------A-----K-------------TRH-N-------
N----FT---IR--RY----RA------QLVGQICDEALQYAARLR--E-------L-E-----P----G----WS------A-----T---P----G-----CQ--------LHD---AEQLWL-----D-----------------PLR--AQT-------------DETFLQ--RRL--RGD-WP-AEVGNRFANWLN-----RA----V---S------S-----D--S-Q-I-------------L---GSPEA------AQWSQ-E-------LS---------------------K-----ELTMFKEIL------------------EDER------D------------------------
---------------
WP_004343803 -----------------------------------------------MTSPLPTP----------T--W---------QELR--QFIE--S-FIQ-ER-----L----QGK------------------------------------------------------------------------------------L---------D----------------K---------L--------HP-----DE----DD---------KR----QT---------------------LL------------------------------------------AT-----------------------
-----HR---REAW-----LA---DA-----ARRV-------G----QLQ-L-----------------VTHTLKPIHPD---A-----------------R---G-S------NL----H----S----L--------P-QAP----GQ--P--GL---------AGS--H--EL------G-D-----R--L--V-SD-V-VG----N----AA----A--L-----DVFKFL-S-----L----Q--Y--Q---------GKNL------------LNWLT-EDS-A---EAVQAL---S---D----NA----EQAREWRQAFIGI--T---AV--KG---
APASHSLAKQLYFPL-------PG-------------------SG---YHLLAPLFPTSLVHHVHAL-LR----E--A---R--F-G------------------DA-AKAAR-E-----AR------------S--RQESW--PH-G-FSE-----YPN-LAIQKFGG-T----KPQ---------NISQLN--SERY-----------GENWLLPSL-P----PHWQ-RQ---DQ-RAPIRH----SSVF----EH---DFG--R-S--PEV--SRLTRTLQRLL------A--KT----R-H-----N-----------------------------------
N----FT---IR--RY----RA------QLVGQICDEALQYAARLR--E-------L-E-----P----G----WS------A-----T---P----G-----CQ--------LHD---AEQLWL-----D-----------------PLR--AQT-------------DETFLQ--RRL--RGD-WP-AEVGNRFANWLN-----RA----V---S------SD------S-Q-I-------------L---GSPEA------AQWSQ-E-----------LS-------------------K-----ELTMFKEIL------------------EDER------D------------------------
---------------
WP_126853648 -----------------------------------------------MTSPLPTP----------T--W---------QELR--QFIE--S-FIQ-ER-----L----QGK------------------------------------------------------------------------------------L---------D----------------K---------L--------HP-----DE----DD---------KR----QT---------------------LL------------------------------------------AT-----------------------
-----HR---REAW-----LA---DA-----ARRV-------G----QLQ-L-----------------VTHTLKPIHPD---A-----------------R---G-S------NL----H----S----L--------P-QAP----GQ--P--GL---------AGS--H--EL------G-D-----K--L--V-SD-V-VG----N----AA----A--L-----DVFKFL-S-----L----Q--Y--Q---------GKNL------------LNWLT-EDS-A---EAVQAL---S---D----NA----EQAREWRQAFIGI--T---TV--KG---
APASHSLAKQLYFPL-------PG-------------------SG---YHLLAPLFPTSLVHHVHAL-LR----E--A---R--F-G------------------DA-AKAAR-E-----AR------------S--RQESW--PH-G-FSE-----YPN-LAIQKFGG-T----KPQ---------NISQLN--SERY-----------GENWLLPSL-P----PHWQ--R-Q-DQ-RAPIHH----SSVF----EH---DFG--R-S--PEV--SRLTRTLQRLL-------------------A-----K-------------TRD-N-------
N----FT---IR--RY----RA------QLVGQICDEALQYAARLR--E-------L-E-----P----G----WS------A-----T---P----G-----CQ--------LHD---AEQLWL-----D-----------------PLR--AQT-------------DETFLQ--RRL--RGD-WP-AEVGNRFANWLN-----RA----V---S------S-----D--S-Q-I-------------L---GSPEA------AQWSQ-E-------LS---------------------K-----ELTMFKEIL------------------EDER------D------------------------
---------------
WP_108112325 -----------------------------------------------MTSPLPTP----------T--W---------QELR--QFIE--S-FIQ-ER-----L----QGK------------------------------------------------------------------------------------L---------D----------------K---------L--------HP-----DE----DD---------KR----QT---------------------LL------------------------------------------AT-----------------------
-----HR---REAW-----LA---DA-----ARRV-------G----QLQ-L-----------------VTHTLKPIHPD---A-----------------R---G-S------NL----H----S----L--------P-QAP----GQ--P--GL---------AGS--H--EL------G-D-----K--L--V-SD-V-VG----N----AA----A--L-----DVFKFL-S-----L----Q--Y--Q---------GKNL------------LNWLT-EDS-A---EAVQAL---S---D----NA----EQAREWRQAFIGI--T---TV--KG---
APASHSLAKQLYFPL-------PG-------------------SG---YHLLAPLFPTSLVHHVHAL-LR----E--A---R--F-G------------------DA-AKAAR-E-----AR------------S--RQESW--PH-G-FSE-----YPN-LAIQKFGG-T----KPQ---------NISQLN--SERY-----------GENWLLPSL-P----PHWQ-RQ---DQ-RAPIHH----SSVF----EH---DFG--R-S--PEV--SRLTRTLQRLL------A--KT----R-H-----N-----------------------------------
N----FT---IR--RY----RA------QLVGQICDEALQYAARLR--E-------L-E-----P----G----WS------A-----T---P----G-----CQ--------LHD---AEQLWL-----D-----------------PLR--AQT-------------DETFLQ--RRL--RGD-WP-AEVGNRFANWLN-----RA----V---S------SD------S-Q-I-------------L---GSPEA------AQWSQ-E-----------LS-------------------K-----ELTMFKEIL------------------EDER------D------------------------
---------------
WP_128733155 -----------------------------------------------MTSPLPTP----------T--W---------QELR--QFIE--S-FIQ-ER-----L----QGK------------------------------------------------------------------------------------L---------D----------------K---------L--------HP-----DE----DD---------KR----QT---------------------LL------------------------------------------AT-----------------------
-----HR---REAW-----LA---DA-----ARRV-------G----QLQ-L-----------------VTHTLKPIHPD---A-----------------R---G-S------NL----H----S----L--------P-QAP----GQ--P--GL---------AGS--H--EL------G-D-----R--L--A-SD-V-VG----N----AA----A--L-----DVFKFL-S-----L----Q--Y--Q---------GKNL------------LNWLT-EDS-A---EALQAL---S---D----NA----EQAREWRQAFIGI--T---AV--KG---
APASHSLAKQLYFPL-------PG-------------------SG---YHLLAPLFPTSLVHHVHAL-LR----E--A---R--F-G------------------DA-AKAAR-E-----AR------------S--RQESW--PH-G-FSE-----YPN-LAIQKFGG-T----KPQ---------NISQLN--SERY-----------GENWLLPSL-P----PHWQ--R-Q-DQ-RAPIRH----SSVF----EH---DFG--R-S--PEV--SRLTRTLQRLL-------------------A-----K-------------TRH-N-------
N----FT---IR--RY----RA------QLVGQICDEALQYAARLR--E-------L-E-----P----G----WS------A-----T---P----G-----CQ--------LHD---AEQLWL-----D-----------------PLR--AQT-------------DETFLQ--RRL--RGD-WP-AEVGNRFANWLN-----RA----V---S------S-----D--S-Q-I-------------L---GSPEA------AQWSQ-E-------LS---------------------K-----ELTMFKEIL------------------EDER------D------------------------
---------------
WP_089030754 -----------------------------------------------MTSPLPTP----------T--W---------QELR--QFIE--S-FIQ-ER-----L----QGK------------------------------------------------------------------------------------L---------D----------------K---------L--------HP-----DE----DD---------KR----QT---------------------LL------------------------------------------AT-----------------------
-----HR---REAW-----LA---DA-----ARRV-------G----QLQ-L-----------------VTHTLKPIHPD---A-----------------R---G-S------NL----H----S----L--------P-QAP----GQ--P--GL---------AGS--H--EL------G-D-----K--L--V-SD-V-VG----N----AA----A--L-----DVFKFL-S-----L----Q--Y--Q---------GKNL------------LNWLT-EDS-A---EALQAL---S---D----NA----EQAREWRQAFIGI--T---AV--KG---
APASHSLAKQLYFPL-------PG-------------------SG---YHLLAPLFPTSLVHHVHAL-LR----E--A---R--F-G------------------DA-AKAAR-E-----AR------------S--RQESW--PH-G-FSE-----YPN-LAIQKFGG-T----KPQ---------NISQLN--SERY-----------GENWLLPSL-P----PHWQ--R-Q-DQ-RAPIRH----SSVF----EH---DFG--R-S--PEV--SRLTRTLQRLL-------------------A-----K-------------TRH-N-------
N----FT---IR--RY----RA------QLVGQICDEALQYAARLR--E-------L-E-----P----G----WS------A-----T---P----G-----CQ--------LHD---AEQLWL-----D-----------------PLR--AQT-------------DETFLQ--RRL--RGD-WP-AEVGNRFANWLN-----RA----V---S------S-----D--S-Q-I-------------L---GSPEA------AQWSQ-E-------LS---------------------K-----ELTMFKEIL------------------EDER------D------------------------
---------------
WP_033946188 -----------------------------------------------MTSPLPTP----------T--W---------QELR--QFIE--S-FIQ-ER-----L----QGK------------------------------------------------------------------------------------L---------D----------------K---------L--------HP-----DE----DD---------KR----QT---------------------LL------------------------------------------AT-----------------------
-----HR---REAW-----LA---DA-----ARRV-------G----QLQ-L-----------------VTHTLKPIHPD---A-----------------R---G-S------NL----H----S----L--------P-QAP----GQ--P--GL---------AGS--H--EL------G-D-----K--L--V-SD-V-VG----N----AA----A--L-----DVFKFL-S-----L----Q--Y--Q---------GKNL------------LNWLT-EDS-A---EALQAL---S---D----NA----EQAREWRQAFIGI--T---TV--KG---
APASHSLAKQLYFPL-------PG-------------------SG---YHLLAPLFPTSLVHHVHAL-LR----E--A---R--F-G------------------DA-AKAAR-E-----AR------------S--RQESW--PH-G-FSE-----YPN-LAIQKFGG-T----KPQ---------NISQLN--SERY-----------GENWLLPSL-P----PHWQ-RQ---DQ-RAPIRH----SSVF----EH---DFG--R-S--PEV--SRLTRTLQRLL------A--KT----R-H-----N-----------------------------------
N----FT---IR--RY----RA------QLVGQICDEALQYAARLR--E-------L-E-----P----G----WS------A-----T---P----G-----CQ--------LHD---AEQLWL-----D-----------------PLR--AQT-------------DETFLQ--RRL--RGD-WP-AEVGNRFANWLN-----RA----V---S------SD------S-Q-I-------------L---GSPEA------AQWSQ-E-----------LS-------------------K-----ELTMFKEIL------------------EDER------D------------------------
---------------
WP_124184559 -----------------------------------------------MTSPLPTP----------T--W---------QELR--QFIE--S-FIQ-ER-----L----QGK------------------------------------------------------------------------------------L---------D----------------K---------L--------HP-----DE----DD---------KR----QT---------------------LL------------------------------------------AT-----------------------
-----HR---REAW-----LA---DA-----ARRV-------G----QLQ-L-----------------VTHTLKPIHPD---A-----------------R---G-S------NL----H----S----L--------P-QAP----GQ--P--GL---------AGS--H--EL------G-D-----K--L--V-SD-V-VG----N----AA----A--L-----DVFKFL-S-----L----Q--Y--Q---------GKNL------------LNWLT-EDS-A---EALQAL---S---D----NA----EQAREWRQAFIGI--T---AV--KG---
APASHSLAKQLYFPL---------------------------LGSG---YHLLAPLFPTSLVHHVHAL-LR----E--A---R--F-G------------------EA-ARAAR-E-----AR------------S--RQESW--PH-G-FSE-----YPN-LAIQKFGG-T----KPQ---------NISQLN--SERY-----------GENWLLPSL-P----PHWQ--R-Q-DQ-RAPIRH----SSVF----EH---DFG--R-S--PEV--SRLTRTLQRLL-------------------A-----K-------------TRD-N-------
N----FT---IR--RY----RA------QLVGQICDEALQYAARLR--E-------L-E-----P----G----WS------A-----T---P----G-----CQ--------LHD---AEQLWL-----D-----------------PLR--AQT-------------DETFLQ--RRL--RGD-WP-AEVGNRFANWLN-----RA----V---S------S-----D--S-Q-I-------------L---GSPEA------AQWSQ-E-------LS---------------------K-----ELTMFKEIL------------------EDER------D------------------------
---------------
WP_124183302 -----------------------------------------------MTSPLPTP----------T--W---------QELR--QFIE--S-FIQ-ER-----L----QGK------------------------------------------------------------------------------------L---------D----------------K---------L--------HP-----DE----DD---------KR----QT---------------------LL------------------------------------------AT-----------------------
-----HR---REAW-----LA---DA-----ARRV-------G----QLQ-L-----------------VTHTLKPIHPD---A-----------------R---G-S------NL----H----S----L--------P-QAP----GQ--P--GL---------AGS--Y--EL------G-D-----R--L--V-SD-V-VG----N----AA----A--L-----DVFKFL-S-----L----Q--Y--Q---------GKNL------------LNWLT-EDS-A---EAVQAL---S---D----NA----EQAREWRQAFIGI--T---AV--KG---
APASHSLAKQLYFPL-------PG-------------------SG---YHLLAPLFPTSLVHHVHGL-LR----E--A---R--F-G------------------EA-ARAAR-E-----AR------------S--RQESW--PH-G-FSE-----YPN-LAIQKFGG-T----KPQ---------NISQLN--SERY-----------GENWLLPSL-P----PHWQ--R-Q-DQ-RAPIRH----SSVF----EH---DFG--R-S--PEV--SRLTRTLQRLL-------------------A-----K-------------TRH-N-------
N----FT---IR--RY----RA------QLVGQICDEALQYAARLR--E-------L-E-----P----G----WS------A-----T---P----G-----CQ--------LHD---AEQLWL-----D-----------------PLR--AQT-------------DETFLQ--RRL--RGD-WP-AEVGNRFANWLN-----RA----V---S------S-----D--S-Q-I-------------L---GSPEA------AQWSQ-E-------LS---------------------K-----ELTMFKEIL------------------EDER------D------------------------
---------------
WP_125027141 -----------------------------------------------MTSPLPTP----------T--W---------QELR--QFIE--S-FIQ-ER-----L----QGK------------------------------------------------------------------------------------L---------D----------------K---------L--------HP-----DE----DD---------KR----QT---------------------LL------------------------------------------AT-----------------------
-----HR---REAW-----LA---DA-----ARRV-------G----QLQ-L-----------------VTHTLKPIHPD---A-----------------R---G-S------NL----H----S----L--------P-QAP----GQ--P--GL---------AGS--H--EL------G-D-----R--L--V-SD-V-VG----N----AA----A--L-----DVFKFL-S-----L----Q--Y--Q---------GKNL------------LNWLT-EDS-A---EAVQAL---S---D----NA----EQAREWRQAFIGI--T---AV--KG---
APASHSLAKQLYFPL-------PG-------------------SG---YHLLAPLFPTSLVHHVHGL-LR----E--A---R--F-G------------------EA-ARAAR-E-----AR------------S--RQESW--PH-G-FSE-----YPN-LAIQKFGG-T----KPQ---------NISQLN--SERY-----------GENWLLPSL-P----PHWQ-RQ---DQ-RAPIRH----SSVF----EH---DFG--R-S--PEV--SRLTRTLQRLL------A--KT----R-H-----N-----------------------------------
N----FT---IR--RY----RA------QLVGQICDEALQYAARLR--E-------L-E-----P----G----WS------A-----T---P----G-----CQ--------LHD---AEQLWL-----D-----------------PLR--AQT-------------DETFLQ--RRL--RGD-WP-AEVGNRFANWLN-----RA----V---S------SD------S-Q-I-------------L---GSPEA------AQWSQ-E-----------LS-------------------K-----ELTMFKEIL------------------EDER------D------------------------
---------------
WP_074207530 -----------------------------------------------MTSPLPTP----------T--W---------QELR--QFIE--S-FIQ-ER-----L----QGK------------------------------------------------------------------------------------L---------D----------------K---------L--------HP-----DE----DD---------KR----QT---------------------LL------------------------------------------AT-----------------------
-----HR---REAW-----LA---DA-----ARRV-------G----QLQ-L-----------------VTHTLKPIHPD---A-----------------R---G-S------NL----H----S----L--------P-QAP----GQ--P--GL---------AGS--H--EL------G-D-----R--L--V-SD-V-VG----N----AA----A--L-----DVFKFL-S-----L----Q--Y--Q---------GKNL------------LNWLT-EDS-A---EAVQAL---S---D----NA----EQAREWRQAFIDI--T---AV--KG---
APASHSLAKQLYFPL-------PG-------------------SG---YHLLAPLFPTSLVHHVHGL-LR----E--A---R--F-G------------------DA-ARAAR-E-----AR------------S--RQESW--PH-G-FSE-----YPN-LAIQKFGG-T----KPQ---------NISQLN--SERY-----------GENWLLPSL-P----PHWQ--R-Q-DQ-RAPIRH----SSVF----EH---DFG--R-S--PEV--SRLTRTLQRLL-------------------A-----K-------------TRH-N-------
N----FT---IR--KY----RA------QLVGQICDEALQYAARLR--E-------L-E-----P----G----WS------A-----T---P----G-----CQ--------LHD---AEQLWL-----D-----------------PLR--AQT-------------DETFLQ--RRL--RGD-WP-AEVGNRFANWLN-----RA----V---S------S-----D--S-Q-I-------------L---GSPEA------AQWSQ-E-------LS---------------------K-----ELTMFKEIL------------------EDER------D------------------------
---------------
```

WP_058170033
[illegible]

WP_124090480
[illegible]

WP_003143952
[illegible]

WP_126518820
[illegible]

WP_121350174
[illegible]

WP_083556355
[illegible]

WP_124120399
[illegible]

WP_058173511
[illegible]

WP_132711961
[illegible]

WP_042170663
[illegible]

WP_058178897
[illegible]

WP_128667840
[illegible]

WP_119518997
[illegible]

WP_121290990
[illegible]

WP_033973667
[illegible]

WP_121386297
[illegible]

WP_121301534
[illegible]

WP_034020660
[illegible]

WP_059309499
[illegible]

WP_031755914
[illegible]

WP_033992769
[illegible]

WP_029610274
[illegible]

WP_073669430
[illegible]

WP_095541116
[illegible]

WP_009876497
[illegible]

WP_043104605
[illegible]

WP_024928898
[illegible]

WP_134246048
[illegible]

WP_003139224
[illegible]

WP_128672734
[illegible]

WP_034022997
[illegible]

WP_124201734
[illegible]

WP_073666572
[illegible]

WP_014602882
[illegible]

WP_088145174
[illegible]

WP_110193878
[illegible]

WP_023091296
[illegible]

WP_121573415
[illegible]

WP_034053734
[illegible]

WP_058162908
[illegible]

WP_043103236
[illegible]

APASHSLAKQLYFPL-------PG------------------SG---YHLLAPLFPTSLVHHVHGL-LR----E--A---R--F-G----------------EA-AKAAR-E-----AS-------------K--HKQPW--PH-G-FSE-----HPN-LAIQKFGG-T----KPQ---------NISQLN--SE [illegible]
------LA---IR--QH----RA------QLVGQICDEALHYADRLR--------E-L-E-----P---G----WS------A-----S---P---D-----CR--------LHE---AEQIWL-----D-------------PLR--VHT-------------DEAFMR--RRL--WAD-WP-MEVSERFANWLN [illegible]
----------------

WP_034080631 [illegible]

WP_107114087 [illegible]

WP_124200878 [illegible]

WP_128712148 [illegible]

WP_124182314 [illegible]

WP_042931586 [illegible]

WP_124172564 [illegible]

WP_124185453 [illegible]

WP_121129310 [illegible]

WP_024917184 [illegible]

WP_043095099 [illegible]

WP_126623049 [illegible]

WP_121358668 [illegible]

WP_124133303 [illegible]

WP_070748425 [illegible]

WP_033978294 [illegible]

WP_114232850 [illegible]

WP_039028030 [illegible]

WP_022654589 [illegible]

WP_093476521 [illegible]

WP_088275471 [illegible]

WP_107693142 [illegible]

WP_121081478 [illegible]

WP_091139454 [illegible]

WP_097066985 [illegible]

WP_020680648 [illegible]

WP_114281081 [illegible]

WP_121457423 [illegible]

WP_005842408 [illegible]

WP_006790330 [illegible]

WP_007069542 [illegible]

WP_023053042 [illegible]

WP_083810435 [illegible]

WP_014454507 [illegible]

WP_068258103 [illegible]

WP_020393891 [illegible]

WP_078924260 [illegible]

WP_139458403 [illegible]

WP_071565375 [illegible]

WP_020161983 [illegible]

WP_058505656 [illegible]

WP_058495128 [illegible]

WP_058451362 [illegible]

[illegible]

WP_031431426

[illegible]

WP_054759601

[illegible]

WP_104423520

[illegible]

WP_064037258

[illegible]

WP_064008668

[illegible]

WP_066977657

[illegible]

WP_103919769

[illegible]

WP_109822600

[illegible]

WP_132321189

[illegible]

WP_006461044

[illegible]

WP_110429265

[illegible]

WP_116687096

[illegible]

WP_115300579

[illegible]

WP_133131549

[illegible]

WP_014703622

[illegible]

WP_133128796

[illegible]

WP_011216754

[illegible]

WP_011212640

[illegible]

WP_061637848

[illegible]

WP_090701172

[illegible]

WP_126339901

[illegible]

WP_069367311

[illegible]

WP_021288187

[illegible]

WP_096864718

[illegible]

WP_036979129

[illegible]

WP_112860121

[illegible]

WP_140170714

[illegible]

WP_102139609

[illegible]

WP_105881018

[illegible]

WP_131679918

[illegible]

WP_042844255

[illegible]

WP_036951427

[illegible]

WP_151164165

[illegible]

WP_137021915

[illegible]

WP_140181641

[illegible]

WP_070926072

[illegible]

WP_008911673

[illegible]

WP_014658069

[illegible]

WP_076914752

[illegible]

WP_040133010

[illegible]

WP_121874877

[illegible]

WP_004918641

[illegible]

WP_110166791

[illegible]

WP_096631394

[illegible]

WP_106593224

[illegible]

WP_078744947

[illegible]

WP_104488201

[illegible]

WP_086963320

[illegible]

WP_087038341

[illegible]

WP_106761803

[illegible]

[illegible]

WP_134031978

[illegible]

WP_036179390

[illegible]

WP_138986280

[illegible]

WP_063357422

[illegible]

WP_063369624

[illegible]

WP_028115395

[illegible]

WP_010318159

[illegible]

WP_093309351

[illegible]

WP_095468801

[illegible]

WP_055033059

[illegible]

WP_138973615

[illegible]

WP_044364441

[illegible]

WP_140229015

[illegible]

WP_061055676

[illegible]

WP_113612734

[illegible]

WP_000399211

[illegible]

WP_080497333

[illegible]

WP_052141706

[illegible]

WP_142576942

[illegible]

WP_119780661

[illegible]

WP_140276092

[illegible]

WP_029821051

[illegible]

WP_102308914

[illegible]

WP_100799474

[illegible]

WP_069523959

[illegible]

WP_064369931

[illegible]

WP_046264979

[illegible]

WP_079749204

[illegible]

WP_021452717

[illegible]

WP_140382494

[illegible]

WP_140114807

[illegible]

WP_100662601

[illegible]

WP_140178011

[illegible]

WP_080277080

[illegible]

WP_079866552

[illegible]

WP_047782208

[illegible]

WP_114987157

[illegible]

WP_126468154

[illegible]

WP_083399707

[illegible]

WP_143680067

[illegible]

WP_020328487

[illegible]

WP_129532380

[illegible]

WP_094129572

[illegible]

WP_072669466

[illegible]

WP_039462015

[illegible]

WP_080371374

[illegible]

WP_033929510

[illegible]

WP_142733556

[illegible]

WP_039552587

[illegible]

WP_032468945

[illegible]

WP_142607085 ----------------------------------------------------MEGS---------------RVLD--PAIA--A-FFA-ER-----K---EAW-------------------------------------------------------------------------------L------------K-----------KNIS-------A-AM-------SD------AE-------------------------------------------------------------------------------VL----------------------------------EKQ-QE----------------------------CE---------
QN------FL---LDNW-----LP---DA-------AKRA---------G-----QIS-V-----------------ASHPCTFSHPS----A-------------R------K-NKNGYVSSI-----I----A---K------------NK---PRI--D--GF---------LRS--G--NV------S--------------V--E-PD-A-LG----N-----AA-----A--L------DVYKFL-S-----L-----E-MS--D---------QRSL-------------LVHIE-QES-E---LARQLL---D---I----PS-----CEYQVLRDGFLKM--V---DS--DQ---
ASVSSSKIKQVYFPI----------------------ADGE----YHLLSLLTHSGHLFELRKR-LD----A--L----R--F-G-----------------DA-VKEAR-E-------CK-------------K--TNHFH--PA-G-YQE--------IFG-LTTIGFGG-T------KPQ--------NISVLN---NQNA-----------------GKAHLLASI-P-----PELT-PR---DI-RPPRTD---FFKESF-----NA-------------WQA---KEVLESLHRLF----------LT-----DYN------N------------------------------------IN--
-LR--DG----RD------YRIQQYVDLVIEKMWQVR--L---------F-LE---A-YS----G------E-----L---S-----------SV-------LPL---EQKIWL-----Y-----------PEF----------------AEQRHQ---------EDE-WL-DKIIRHIARGLI-----NH-----Y---S-RSKVITN------P-V----------------TL----ADHEL------LAIEK-------------VV-------------A---------SNKENLR----------------------------------------------------------------
WP_142735590 ----------------------------------------------------MEGS---------------RVLD--PSIA--A-FFA-ER-----K---EAW------------------------------------------------------------------------------------L------------K------------KNIS------A-AM-------SD------TE-----------------------------------------------------------------------------------VL------------------------------EKQ-QE-------------------------------CE----------
QN------FL---LDNW-----LP---DA-------AKRA---------G-----QIS-V-----------------ASHPCTFSHPS----A-------------R------K-NKNGYVSSI-----I----A---K------------NK---PRI--D--GF---------LRS--G--NV------S--------------V--E-PD-A-LG----N-----AA-----A--L------DVYKFL-S-----L-----E-MS--D---------QRSL-------------LVHIE-QES-E---LARQLL---D---I----PS-----CEYQALRDGFLKM--V---DS--DQ---
ASVSSSKIKQVYFPI----------------------ADGE----YHLLSLLTHSGHLFELRKR-LD----A--L----R--F-G-----------------DA-VKEAR-E-------CK-------------K--TNHFH--PA-G-YQE--------IFG-LTTIGFGG-T------KPQ--------NISVLN---NQNA-----------------GKAHLLASI-P-----PELT-PR---DI-RLPRTE---FFKESF-----TA--------------WQA---KEVLESLHRLF--------L--TD---Y------N------------------------------N-----IN--
IR--DG----RD------YRIQQYVDLVIEKMWQVR--L---------F-LE---A----YS-----G----E---LS--------SA-------LPL---DQKIWL-----Y-----------PEF----------------AE--QR--QQE----DE-WL-DKIIRHIARGLI-----NH-----Y----S------RS--KVITNP---V-------------TL----ADQEL---------LAI-E---------KV--VA-------------------SNKENLR--------------------------------------------------------------
WP_071178037 ----------------------------------------------------MEGS---------------RVLD--PAIA--A-FFA-ER-----K---EAW------------------------------------------------------------------------------------L------------K------------KNIS------A-AM-------SD------TE-----------------------------------------------------------------------------------VL------------------------------EKQ-QE-------------------------------CE----------
QN------FL---LDNW-----LP---DA-------AKRA---------G-----QIA-V-----------------ASHPCTFSHPS----A-------------R------K-NKNGYVSSI-----I----A---K------------NK---PRI--D--GF---------LRS--G--NV------S--------------V--E-PD-A-LG----N-----AA-----A--L------DVYKFL-S-----L-----E-MS--D---------QRSL-------------LVHIE-QES-E---LARQLL---D---I----PS-----CEYQALRDGFLKM--V---DS--DQ---
ASVSSSKIKQVYFPI----------------------ADGE----YHLLSLLTHSGHLFELRKR-LD----A--L----R--F-G-----------------DA-VKEAR-E-------CK-------------K--TNHFH--PA-G-YQE--------IFG-LTTIGFGG-T------KPQ--------NISVLN---NQNA-----------------GKAHLLASI-P-----PELT-SR---DI-RLPKTD---FFKESF-----TA---------------RQA---KEVLEALHRLF--------L--TD---Y------N------------------------------N-----IN--
IR--DG----RD------YRIQQYVDLVIEKMWQVR--L---------F-LE---A----YS-----G----E---LS--------SA-------LPL---DQKIWL-----Y-----------PEF----------------AE--QR--QQE----DE-WL-DKIIRHIARGLI-----NH-----Y----S------RS--KVITNP---V-------------TL----ADQEL---------LAI-E---------KV--VA-------------------SNKENLR--------------------------------------------------------------
WP_142614055 ----------------------------------------------------MID--PVIA--A-FFA-ER-----K---EAW--------------------------------------------------------------------------------------L--K------------KN------S----A---A------MS-----DA------------------------------------------------------------------------------------------EV------------------------------------------------CE------QN----
-FL---LDNW-----LP---DA-------AKRA---------G-----QIA-V----------------ASHPCTFSHPS----A------RKNKNG-----Y---V--S-----SI----I----A-K-----------NK---PRI---D--GF---------LRS--G--NV------S--------------V--E-PD-A-LG----N-----A---A--A-----LDVYKFL-S-----L-----E-MS--D---------QRSL-------------LVHIE-QES-E---LARQLL---D---I----PS---CEYQALRDGFLKM--V---DS--DQ-A-SVSSS-
KIKQVYFPI----------------------ADGE----YHLLSLLTHSGHLFELRKR-LD----A--L----R--F-G----------------DA-VKEAR-E-------CK-------------K--TNHFH--PA-G-YQE--------IFG-LTTIGFGG-T------KPQ--------NISVLN---NQNA-----------------GKAHLLASI-P-----PELT-SR---DI-RLPKTD---FFKESF-----TA------------RQA---KEVLEALHRLF--------L--TD---Y------N------------------------------N-----IN---IR--DG---
RD------YRIQQYVDLVIEKMWQVR--L---------F-LE---A----YS-----G----E---LS--------SA-------LPL---EQKIWL-----Y-----------PEF----------------AE--QR--QQE----DE-WL-DKIIRHIARGLI-----NH-----Y----S------RS--KVITNP---V-------------TL----ADQEL---------LAI-E---------KV--VA-------------------SNKENLR-----------------------------------------------------------
WP_114710786 ----------------------------------------------------MID--PAIA--A-FFA-ER-----K---EAW--------------------------------------------------------------------------------------L--K------------KN------S----A---A------MS-----DT------------------------------------------------------------------------------------------EV------------------------------------------------L-EKQ-QE------------CE------QN----
FL---LDNW-----LP---DA-------AKRA---------G-----QIA-V----------------ASHPCTFSHPS----A------RKNKNG-----Y---V--S-----SI----I----A-K-----------NK---PRI---D--GF---------LRS--G--NV------S--------------V--E-PD-A-LG----N-----A---A--A-----LDVYKFL-S-----L-----E-MS--D---------QRSL-------------LVHIE-QES-E---LARQLL---D---I----PS---CEYQALRDGFLKM--V---DS--DQ-A-SVSSS-
KIKQVYFPI----------------------ADGE----YHLLSLLTHSGHLFELRKR-LD----A--L----R--F-G----------------DA-VKEAR-E-------CK-------------K--TNHFH--PA-G-YQE--------IFG-LTTIGFGG-T------KPQ--------NISVLN---NQNA-----------------GKAHLLASI-P-----PELT-SR---DI-RLPKTD---FFKESF-----TA------------RQA---KEVLEALHRLF--------L--TD---Y------N------------------------------N-----IN---IR--DG---
RD------YRIQQYVDLVIEKMWQVR--L---------F-LE---A----YS-----G----E---LS--------SV-------LPL---EQKIWL-----Y-----------PEF----------------AE--QR--HQE----DE-WL-DKIIRHIARGLI-----NH-----Y----S------RS--KVITSP---V-------------TL----ADQEL---------LAI-E---------KV--VA-------------------SNKENLR-----------------------------------------------------------
WP_142735063 ----------------------------------------------------MEGS---------------RVLD--PAIA--A-FFA-ER-----K---EAW------------------------------------------------------------------------------------L------------K------------KNIS------A-AM-------SD------AE-----------------------------------------------------------------------------------VL------------------------------EKQ-QE-------------------------------CE----------
QN------FL---LDNW-----LP---DA-------AKRA---------G-----QIA-V-----------------ASHPCTFSHPS----A-------------R------K-NKNGYVSSI-----I----A---K------------NK---PRI--D--GF---------LRS--G--NV------S--------------V--E-PD-A-LG----N-----AA-----A--L------DVYKFL-S-----L-----E-MS--D---------QRSL-------------LVHIE-QES-E---LARQLL---D---I----PS-----CEYQALRDGFLKM--V---DS--DQ---
ASVSSSKIKQVYFPI----------------------ADGE----YHLLSLLTHSGHLFELRKR-LD----A--L----R--F-G-----------------DA-VKEAR-E-------CK-------------K--TNHFH--PA-G-YQE--------IFG-LTTIGFGG-T------KPQ--------NISVLN---NQNA-----------------GKAHLLASI-P-----PELT-PR---DI-RLPRTD---FFKESF-----NA-------------WQA---KEVLESLHRLF----------LT-----D-Y------N------------------------------------IN--
LR--DG----RD------YRIQQYVDLVIEKMWQVR--L---F-LE---A-YS----G------E-----L---S-----------SV-------LPL---EQKIWL-----Y-----------PEF----------------AEQRHQ---------EDE-WL-DKIIRHIARGLI-----NH-----Y---S----RSK--VITS-P-I---------------TL----ADQEL---------------------------L-----AIEKVVASN-------------KENL-----------R--------------------------------------------
WP_142628887 ----------------------------------------------------MID--PAIT--A-FFA-ER-----K---EAW--------------------------------------------------------------------------------------L------------K-----KNIS--------A-AM-------SD------TE------------------------------------------------------------------------------VL----------------------------------L-EKQ-QE----------------------CE------QN----
-FL---LDNW-----LP---DA-------AKRA---------G-----QIA-V----------------ASHPCTFSHPS----A-------------R------K-NKNGYVSSI-----I----A---K------------NK---PRI--D--GF---------LRS--G--NV------S--------------V--E-PD-A-LG----N-----AA-----A--L------DVYKFL-S-----L-----E-MS--D---------QRSL-------------LVHIE-QES-E---LARQLL---D---I----PS-----CEYQALRDGFLKM--V---DS--DQ---
ASVSSSKIKQVYFPI----------------------ADGE----YHLLSLLTHSGHLFELRKR-LD----A--L----R--F-G-----------------DA-VKEAR-E-------CK-------------K--TNHFH--PA-G-YQE--------IFG-LTTIGFGG-T------KPQ--------NISVLN---NQNA-----------------GKAHLLASI-P-----PELT-LR---DI-RLPRTD---FFKESF-----NA-------------WQA---KEVLESLHRLF----------LT-----D-Y------N------------------------------------IN--
LR--DG----RD------YRIQQYVDLVIEKMWQVR--L---F-LE---A-YS----G------E-----L---S-----------SV-------LPL---EQKIWL-----Y-----------PEF----------------AEQRHQ---------EDE-WL-DKIIRHIARGLI-----NH-----Y---S----RSK--VITS-P-I---------------TL----ADQEL---------------------------L-----AIEKVVASN-------------KENL-----------R--------------------------------------------
WP_142541138 ----------------------------------------------------MID--PAIA--A-FFA-ER-----K---EAW--------------------------------------------------------------------------------------L--K------------KN------S----A---A------MS-----DT------------------------------------------------------------------------------------------EV------------------------------------------------L-EKQ-QE------------CE------QN----
FL---LDNW-----LP---DA-------AKRA---------G-----QIA-V----------------ASHPCTFSHPS----A------RKNKNG-----Y---V--S-----SI----I----A-K-----------NK---PRI---D--GF---------LRS--G--NV------S--------------V--E-PD-A-LG----N-----A---A--A-----LDVYKFL-S-----L-----E-MS--D---------QRSL-------------LVHIE-QES-E---LARQLL---D---I----PS---CEYQALRDGFLKM--V---DS--DQ-A-SVSSS-
KIKQVYFPI----------------------ADGE----YHLLSLLTHSGHLFELRKR-LD----A--L----R--F-G----------------DA-VKEAR-E-------CK-------------K--TNHFH--PA-G-YQE--------IFG-LTTIGFGG-T------KPQ--------NISVLN---NQNA-----------------GKAHLLASI-P-----PELT-LR---DI-RLPRTD---FFKESF-----NA------------WQA---KEVLESLHRLF--------L--TD---Y------N------------------------------N-----LR--DG---
-RD------YRIQQYVDLVIEKMWQVR--L---------F-LE---A-YS----G------EL----S-----------------SA-------LPL---EQKIWL-----Y-----------PEF----------------AEQRQQ---------DE-WL-DKIIRHIARGLI-----NH-----Y----S------RSK----VITN-P-V----------------TL----ADQEL----------------------------L---AIEEVVASN-------------KENL--------R-----------------------------------------
WP_119788718 ----------------------------------------------------MEGS---------------RVLD--PAIA--A-FFA-ER-----K---EAW------------------------------------------------------------------------------------L------------K------------KNIS------A-AM-------SD------TE-----------------------------------------------------------------------------------VL------------------------------EKQ-QE-------------------------------CE----------
QN------FL---LDNW-----LP---DA-------AKRA---------G-----QIA-V-----------------ASHPCTFSHPS----A-------------R------K-NKNGYVSSI-----I----A---K------------NK---PRI--D--GF---------LRS--G--NV------S--------------V--E-PD-A-LG----N-----AA-----A--L------DVYKFL-S-----L-----E-MS--D---------QRSL-------------LVHIE-QES-E---LARQLL---D---I----PS-----CEYQALRDGFLKM--V---DS--DQ---
ASVSSSKIKQVYFPI----------------------ADGE----YHLLSLLTHSGHLFELRKR-LD----A--L----R--F-G-----------------DA-VKEAR-E-------CK-------------K--TNHFH--PE-G-YQE--------IFG-LTTIGFGG-T------KPQ--------NISVLN---NQNA-----------------GKAHLLASI-P-----PELT-PR---DI-RLPKTD---FFKESF-----TA--------------RQA---KEVLESLHRLF---------QI-----D-Y------N------------------------------------IN--
LR--DG----RD------YRIQQYIDLVIEKMWQVR--L---F-LE---A-YS----G------E-----L---S-----------SA-------LPL---EQKIWL-----Y-----------PEF----------------AEQRQQ---------EDE-WL-DKIIRHIARGLI-----NH-----Y---S----RSK--VITN-P-V---------------TL----ADQEL---------------------------L-----AIEEVVASN-------------KENL-----------R--------------------------------------------
WP_032475794 ----------------------------------------------------MID--PAIA--A-FFA-ER-----K---EAW--------------------------------------------------------------------------------------L--K------------KN------S----A---A------MS-----DT------------------------------------------------------------------------------------------EV------------------------------------------------L-EKQ-QE------------CE------QN----
QN------FL---LDNW-----LP---DA-------AKRA---------G-----QIA-V----------------ASHPCTFSHPS----A------RKNKNG-----Y---V--S-----SI----I----A-K-----------NK---PRI---D--GF---------LRS--G--NV------S--------------V--E-PD-A-LG----N-----A---A--A-----LDVYKFL-S-----L-----E-MS--D---------QRSL-------------LVHIE-QES-E---LARQLL---D---I----PS---CEYQALRDGFLKM--V---DS--DQ-A-SVSSS-
KIKQVYFPI----------------------ADGE----YHLLSLLTHSGHLFELRKR-LD----A--L----R--F-G----------------DA-VKEAR-E-------CK-------------K--TNHFH--PE-G-YQE--------IFG-LTTIGFGG-T------KPQ--------NISVLN---NQNA-----------------GKAHLLASI-P-----PELT-PR---DI-RLPKTD---FFKESF-----TA------------RQA---KEVLESLHRLF--------QI--D---Y------N------------------------------N-----IN---LR--DG---
RD------YRIQQYVDLVIEKMWQVR--L---------F-LE---A-YS----G------EL----S-----------------SA-------LPL---EQKIWL-----Y-----------PDF----------------AEQRQQ---------DE-WL-DKIIRHIARGLI-----NH-----Y----S------RSK----VITN-P-V----------------TL----ADQEL----------------------------L---AIEEVVASN-------------KENL--------R-----------------------------------------
WP_102136867 ----------------------------------------------------MEGS---------------RVLD--PAIA--A-FFA-ER-----K---EAW------------------------------------------------------------------------------------L------------K------------KNIS------A-AM-------SD------AE-----------------------------------------------------------------------------------VL------------------------------EKQ-QE-------------------------------CE----------
QN------FL---LDNW-----LP---DA-------AKRA---------G-----QIA-V-----------------ASHPCTFSHPS----A-------------R------K-NKNGYVSSI-----I----A---K------------NK---PRI--D--GF---------LHS--G--NV------S--------------V--E-PD-A-LG----N-----AA-----A--L------DVYKFL-S-----L-----E-MS--D---------QRSL-------------LVHIE-QES-E---LARQLL---D---I----PS-----CEYQALRDGFLKM--V---DS--DQ---
ASVSSSKIKQVYFPI----------------------ADGE----YHLLSLLTHSGHLFELRKR-LD----A--L----R--F-G-----------------DA-VKEAR-E-------CK-------------K--TNHFH--PA-G-YQE--------IFG-LTTIGFGG-T------KPQ--------NISVLN---NQNA-----------------GKAHLLASI-P-----PELT-PR---DI-RLPKTD---FFKESF-----NA-------------WQA---KEVLESLHRLF---------QT-----D-Y------N------------------------------------IN--
LR--DG----RD------YRIQQYVDLVIEKMWQVR--L---F-LE---A-YS----G------E-----L---S-----------SA-------LPL---EQKIWL-----Y-----------PEF----------------AEQRQQ---------EDE-WL-DKIIRHIARGLI-----NH-----Y---S----RSK--VITS-P-I---------------TL----ADQEL---------------------------L-----AIEKVVASN-------------KENL-----------R--------------------------------------------
WP_080481328 ----------------------------------------------------MEGS---------------RVLD--PAIA--A-FFA-ER-----K---EAW------------------------------------------------------------------------------------L------------K------------KNIS------A-AM-------SD------AE-----------------------------------------------------------------------------------VL------------------------------EKQ-QE-------------------------------CE----------
QN------FL---LDNW-----LP---DA-------AKRA---------G-----QIS-V-----------------ASHPCTFSHPS----A-------------R------K-NKNGYVSSI-----I----A---K------------NK---PRI--D--GF---------LRS--G--NV------S--------------V--E-PD-A-LG----N-----AA-----A--L------DVYKFL-S-----L-----E-MS--D---------QRSL-------------LVHIE-QES-E---LARQLL---D---I----PS-----CEYQALRDGFLKM--V---DS--DQ---
ASVSSSKIKQVYFPI----------------------ADGE----YHLLSLLTHSGHLFELRKR-LD----A--L----R--F-G-----------------DA-VKEAR-E-------CK-------------K--TNHFH--PA-G-YQE--------IFG-LTTIGFGG-T------KPQ--------NISVLN---NQNA-----------------GKAHLLASI-P-----PELT-PR---DI-RLPKTD---FFKESF-----NA-------------WQA---KEVLESLHRLF---------QT-----D-Y------N------------------------------------IN--
LR--DG----RD------YRIQQYVDLVIEKMWQVR--L---F-LE---A-YS----G------E-----L---S-----------SA-------LPL---EQKIWL-----Y-----------PEF----------------AEQRQQ---------EDE-WL-DKIIRHIARGLI-----NH-----Y---S----RSK--VITN-P-V---------------TL----ADQEL---------------------------L-----AIEEVVASN-------------KENL-----------R--------------------------------------------
WP_096070605 ----------------------------------------------------MID--PVIA--A-FFA-ER-----K---EAW--------------------------------------------------------------------------------------L------------K-----KNIS--------A-AM-------SD------AE------------------------------------------------------------------------------VL----------------------------------EKQ-QE----------------------CE------QN----
-FL---LDNW-----LP---DA-------AKRA---------G-----QIS-V----------------ASHPCTFSHPS----A-------------R------K-NKNGYVSSI-----I----A---K------------NK---PRI--D--GF---------LRS--G--NV------S--------------V--E-PD-A-LG----N-----AA-----A--L------DVYKFL-S-----L-----E-MS--D---------QRSL-------------LVHIE-QES-E---LARQLL---D---I----PS-----CEYQALRDGFLKM--V---DS--DQ---
ASVSSSKIKQVYFPI----------------------ADGE----YHLLSLLTHSGHLFELRKR-LD----A--L----R--F-G-----------------DA-VKEAR-E-------CK-------------K--TNHFH--PA-G-YQE--------IFG-LTTIGFGG-T------KPQ--------NISVLN---NQNA-----------------GKAHLLASI-P-----PELT-PR---DI-RLPKTD---FFKESF-----NA-------------WQA---KEVLESLHRLF---------QT-----D-Y------N------------------------------------IN--
LR--DG----RD------YRIQQYVDLVIEKMWQVR--L---F-LE---A-YS----G------E-----L---S-----------SA-------LPL---EQKIWL-----Y-----------PEF----------------AEQRQQ---------EDE-WL-DKIIRHIARGLI-----NH-----Y---S----RSK--VITS-P-I---------------TL----ADQEL---------------------------L-----AIEKVVASN-------------KENL-----------R--------------------------------------------
WP_069212542 ----------------------------------------------------MID--PVIA--A-FFA-ER-----K---EAW--------------------------------------------------------------------------------------L------------K-----KNIS--------A-AM-------SD------AE------------------------------------------------------------------------------VL----------------------------------L-EKQ-QE----------------------CE------QN----
-FL---LDNW-----LP---DA-------AKRA---------G-----QIS-V----------------ASHPCTFSHPS----A-------------R------K-NKNGYVSSI-----I----A---K------------NK---PRI--D--GF---------LRS--G--NV------S--------------V--E-PD-A-LG----N-----AA-----A--L------DVYKFL-S-----L-----E-MS--D---------QRSL-------------LVHIE-QES-E---LARQLL---D---I----PS-----CEYQALRDGFLKM--V---DS--DQ---
ASVSSSKIKQVYFPI----------------------ADGE----YHLLSLLTHSGHLFELRKR-LD----A--L----R--F-G-----------------DA-VKEAR-E-------CK-------------K--TNHFH--PA-G-YQE--------IFG-LTTIGFGG-T------KPQ--------NISVLN---NQNA-----------------GKAHLLASI-P-----PELT-SR---DI-RLPKTD---FFKESF-----NA-------------WQA---KEVLESLHRLF---------QT-----D-C------N------------------------------------IN--
LR--DG----RD------YRIQQYVDLVIEKMWQVR--L---F-LE---A-YS----G------E-----L---S-----------SA-------LPL---EQKIWL-----Y-----------PEF----------------AEQRQQ---------EDE-WL-DKIIRHIARGLI-----NH-----Y---S----RSK--VITN-P-V---------------TL----ADQEL---------------------------L-----AIEEVVASN-------------KENL-----------R--------------------------------------------
WP_032481446 ----------------------------------------------------MID--PVIA--A-FFA-ER-----K---EAW--------------------------------------------------------------------------------------L--K------------KNIS------A---A------MS-----DA------------------------------------------------------------------------------------------EV------------------------------------------------L-EKQ-QE------------CE------QN----
QN------FL---LDNW-----LP---DA-------AKRA---------G-----QIS-V----------------ASHPCTFSHPS----A-------------R------K-NKNGYVSSI-----I----A---K------------NK---PRI--D--GF---------LRS--G--NV------S--------------V--E-PD-A-LG----N-----AA-----A--L------DVYKFL-S-----L-----E-MS--D---------QRSL-------------LVHIE-QES-E---LARQLL---D---I----PS-----CEYQALRDGFLKM--V---DS--DQ-A-
SVSSSSKIKQVYFPI----------------------ADGE----YHLLSLLTHSGHLFELRKR-LD----A--L----R--F-G-----------------DA-VKEAR-E-------CK-------------K--TNHFH--PA-G-YQE--------IFG-LTTIGFGG-T------KPQ--------NISVLN---NQNA-----------------GKAHLLASI-P-----PELT-PR---DI-RLPKTD---FFKESF-----NA-------------WQA---KEVLESLHRLF---------QT-----D-Y------N------------------------------------IN--
LR--DG----RD------YRIQQYVDLVIEKMWQVR--L---F-LE---A-YS----G------E-----L---S-----------SA-------LPL---EQKIWL-----Y-----------PEF----------------AEQRQQ---------EDE-WL-DKIIRHIARGLI-----NH-----Y---S----RSK--VITN-P-V---------------TL----ADQEL---------------------------L-----AIEEVVASN-------------KENL-----------R--------------------------------------------
WP_142736467 ----------------------------------------------------MID--PVIA--A-FFA-ER-----K---EAW--------------------------------------------------------------------------------------L------------K-----KNIS--------A-AM-------SD------AE------------------------------------------------------------------------------VL----------------------------------EKQ-QE----------------------CE------QN----
-FL---LDNW-----LP---DA-------AKRA---------G-----QIS-V----------------ASHPCTFSHPS----A-------------R------K-NKNGYVSSI-----I----A---K------------NK---PRI--D--GF---------LRS--G--NV------S--------------V--E-PD-A-LG----N-----AA-----A--L------DVYKFL-S-----L-----E-MS--D---------QRSL-------------LVHIE-QES-E---LARQLL---D---I----PS-----CEYQALRDGFLKM--V---DS--DQ---
ASVSSSKIKQVYFPI----------------------ADGE----YHLLSLLTHSGHLFELRKR-LD----A--L----R--F-G-----------------DA-VKEAR-E-------CK-------------K--TNHFH--PA-G-YQE--------IFG-LTTIGFGG-T------KPQ--------NISVLN---NQNA-----------------GKAHLLASI-P-----PELT-SR---DI-RLPRTD---FFKESF-----NA-------------WQA---KEVLESLHRLF---------QT-----D-Y------N------------------------------------IN--
LR--DG----RD------YRIQQYVDLVIEKMWQVR--L---F-LE---A-YS----G------E-----L---S-----------SA-------LPL---EQKIWL-----Y-----------PEF----------------AEQRQQ---------EDE-WL-DKIIRHIARGLI-----NH-----Y---S----RSK--VITN-P-V---------------TL----ADQEL---------------------------L-----AIEEVVASN-------------KENL-----------R--------------------------------------------
WP_084981128 ----------------------------------------------------MID--PVIA--A-FFA-ER-----K---EAW--------------------------------------------------------------------------------------L--K------------KNIS------A---A------MS-----DA------------------------------------------------------------------------------------------EV------------------------------------------------L-EKQ-QE------------CE------QN----
FL---LDNW-----LP---DA-------AKRA---------G-----QIS-V----------------ASHPCTFSHPS----A-------------R------K-NKNGYVSSI-----I----A---K------------NK---PRI--D--GF---------LRS--G--NV------S--------------V--E-PD-A-LG----N-----AA-----A--L------DVYKFL-S-----L-----E-MS--D---------QRSL-------------LVHIE-QES-E---LARQLL---D---I----PS-----CEYQALRDGFLKM--V---DS--DQ-A-
SVSSSSKIKQVYFPI----------------------ADGE----YHLLSLLTHSGHLFELRKR-LD----A--L----R--F-G-----------------DA-VKEAR-E-------CK-------------K--TNHFH--PA-G-YQE--------IFG-LTTIGFGG-T------KPQ--------NISVLN---NQNA-----------------GKAHLLASI-P-----PELT-SR---DI-RLPKTD---FFKESF-----NA-------------WQA---KEVLESLHRLF---------QT-----D-Y------N------------------------------------IN--
LR--DG----RD------YRIQQYVDLVIEKMWQVR--L---F-LE---A-YS----G------E-----L---S-----------SA-------LPL---EQKIWL-----Y-----------PEF----------------AEQRQQ---------EDE-WL-DKIIRHIARGLI-----NH-----Y---S----RSK--VITN-P-V---------------TL----ADQEL---------------------------L-----AIEEVVASN-------------KENL-----------R--------------------------------------------
WP_082052706 ----------------------------------------------------MEGF------------TVLD--PAIV--T-FFA-ER-----K---EAW------------------------------------------------------------------------------------L------------K-------------KNVS------T-SM-------QE------SE---------------------------------------------------------------------------------EV----------------------------------EKQ-RE---------------------------CE----------
QN------FL---LANW-----LP---DA-------AKRA---------G-----QIS-I-----------------ASHPCTFSHPS----A-------------R------K-NKNGYVSSI-----I----A---K------------NK---PSA--D--GF---------LRS--G--NV------S--------------V--E-PD-A-LG----N-----AA-----A--L------DVYKFL-S-----L-----P-MS--D---------QRSL-------------LVHIE-QES-E---LARQLL---E---V----PT-----CDYQTLRDGFLKM--I---DR--DQ---
ASVSSSKIKQVYFPV----------------------AEGE----YHLLSLLTHSGHLFELRKR-LD----A--M----R--F-G-----------------EL-VKEAR-E-------CK-------------K--ANRFH--PE-G-YQE--------IFG-LTTIGFGG-T------KPQ--------NISVLN---NQNA-----------------GKAHLLPSI-P-----PNLT-PR---EI-RLPKTD---FFKESF-----TA--------------WQA---KEVLEALHRLF---------QT-----DYN------N------------------------------------IN--
LR--DG----RD------YRIQQYVDLVIEKMWQVR--------LFL-E------------A----YS-----G---------E-LSSA------LPL---EQKVWL------Y----------------YPEFAE--QRQ--HEDEWL-DKITRHIARGLI------NH-----Y---S-RSKVIANP-------V------------TL----ADQEL---LAIEE------------VL----------------A-----SNKENLR---------------------------------------------------------------
WP_081737215 ----------------------------------------------------MEGF------------TVLD--PAIV--T-FFA-ER-----K---EAW------------------------------------------------------------------------------------L------------K-------------KNVS------T-SM-------QE------SE---------------------------------------------------------------------------------EV----------------------------------EKQ-RE---------------------------CE----------
QN------FL---LANW-----LP---DA-------AKRA---------G-----QIS-I-----------------ASHPCTFSHPS----A-------------R------K-NKNGYVSSI-----I----A---K------------NK---PSA--D--GF---------LRS--G--NV------S--------------V--E-PD-A-LG----N-----AA-----A--L------DVYKFL-S-----L-----P-MS--D---------QRSL-------------LVHIE-QES-E---LARQLL---E---V----PT-----CDYQTLRDGFLKM--I---DR--DQ---
ASVSSSKIKQVYFPV----------------------AEGE----YHLLSLLTHSGHLFELRKR-LD----A--M----R--F-G-----------------EL-VKEVR-E-------CK-------------K--ANRFH--PE-G-YQE--------IFG-LTTIGFGG-T------KPQ--------NISVLN---NQNA-----------------GKAHLLPSI-P-----PNLT-PR---EI-RLPKTD---FFKECF-----TA--------------WQA---KEVLEALHRLF---------KA-----DYN------N------------------------------------IN--
IR--SG----RD------YRIQQYVDLVIEKMWQVR--L---F-L-T--D-HQ---G------V-----L---P----------SE-------LPL---EQKVWL-----Y-----------PEF----------------ESQREQ---------ENE-WL-DKITRHIARGLI-----NH-----Y---S-RSKVISN------P-L----------LL----ADQEL-------------LAIET----------VV----------A-----SNRENFR------------------------------------------------------------
WP_144226268 ----------------------------------------------------MEGF------------TVLD--PAIV--T-FFA-ER-----K---EAW------------------------------------------------------------------------------------L------------K-------------KNVS------T-SM-------QE------SE---------------------------------------------------------------------------------EV----------------------------------EKQ-RE---------------------------CE----------
QN------FL---LANW-----LP---DA-------AKRA---------G-----QIS-I-----------------ASHPCTFSHPS----A-------------R------K-NKNGYVSSI-----I----A---K------------NK---PSV--D--GF---------LRS--G--NV------S--------------V--E-PD-A-LG----N-----AA-----A--L------DVYKFL-S-----L-----P-MS--D---------QRSL-------------LVHIE-QES-E---LARQLL---E---V----PT-----CDYQTLRDGFLKM--I---DR--DQ---
ASVSSSKIKQVYFPV----------------------AEGE----YHLLSLLTHSGHLFELRKR-LD----A--M----R--F-G-----------------EL-VKEAR-E-------CK-------------K--ANRFH--PE-G-YQE--------IFG-LTTIGFGG-T------KPQ--------NISVLN---NQNA-----------------GKAHLLPSI-P-----PNLT-PR---EI-RLPKTD---FFKECF-----TA--------------WQA---KEVLEALHRLF---------KA-----DYN------N------------------------------------IN--
IR--DG----RD------YRIQQYVDLVIEKMWQVR--L---F-L-T--G-HQ---G------D-----L---P----------SE-------LPL---EQKVWL-----Y-----------PEF----------------ESQREQ---------ENE-WL-DKITRQIARGLI-----NH-----Y---S-RSKVISN------P-L----------LL----ADQEL------------LVIET----------VV----------A-----SNRENFR------------------------------------------------------------
WP_107005250 ----------------------------------------------------M--EGF------TVLD--PAIV--T-FFA-ER-----K---EAW---------------------------------------------------------------------------------L--K-----KN------V-----S----T-S----MQ-----ES------------------------------------------------------------------------------------------EV---------------------------------------------L-EKQ-RE------------CK-------
QN------FL---LANW-----LP---DA-------AKRA---------G-----QIS-I----------------ASHPCTFSHPS----A------RKNENG-----Y---V--S-----SI----I----A-K-----------NK---PSA---D--GF---------LRS--G--NV------S--------------V--E-PD-A-LG----N-----A---A--A-----LDVYKFL-S-----L-----P-MS--D---------QRSL-------------LVHIE-QES-E---LARQLL---E---V----PT-----CDYQTLRDGFLKM--I---DS--DQ-A-SVSSS-
KIKQVYFPV----------------------AEGE----YHLLSLLTHSGHLFELRKR-LD----A--M----R--F-G-----------------EL-VKEAR-E-------CK-------------K--ANRFH--PE-G-YQE--------IFG-LTTIGFGG-T------KPQ--------NISVLN---NQNA-----------------GKAHLLPSI-P-----PNLT-PR---EI-RLPKTD---FFKECF-----TA--------------WQA---KEVLEALHRLF---------KT-----DYN------N------------------------------------IN---IR--DG--
--RD------YRIQQYVDLVIEKMWQVR--L---F-L-T--G-HQ---G------D-----L---P----------SE-------LPL---EQKVWL-----Y-----------PEF----------------ESQREQ---------ENE-WL-DKITRQIARGLI-----NH-----Y---S-RSKVISN------P-L----------LL----ADQEL------------LAIET----------VV----------A-----SNRENFR------------------------------------------------------------
WP_093981938 ----------------------------------------------------M--EGF------TVLD--PAIV--T-FFA-ER-----K---EAW---------------------------------------------------------------------------------L--K-----KN------V-----S----T-S----MQ-----ES------------------------------------------------------------------------------------------EV---------------------------------------------L-EKQ-RE------------CK-------
QN------FL---LANW-----LP---DA-------AKRA---------G-----QIS-I----------------ASHPCTFSHPS----A------RKNKNG-----Y---V--S-----SI----I----A-K-----------NK---PSA---D--GF---------LRS--G--NV------S--------------V--E-PD-A-LG----N-----A---A--A-----LDVYKFL-S-----L-----P-MS--D---------QRSL-------------LVHIE-QES-E---LARQLL---E---V----PT-----CDYQTLRDGFLKM--I---DS--DQ-A-SVSSS-
KIKQVYFPV----------------------AEGE----YHLLSLLTHSGHLFELRKR-LD----A--M----R--F-G-----------------EL-VKEAR-E-------CK-------------K--ANRFH--PE-G-YQE--------IFG-LTTIGFGG-T------KPQ--------NISVLN---NQNA-----------------GKAHLLPSI-P-----PNLT-PR---EI-RLPKTD---FFKECF-----TA--------------WQA---KEVLEALHRLF---------KT-----DYN------N------------------------------------IN---IR--DG--
--RD------YRIQQYVDLVIEKMWQVR--L---F-L-T--D-HQ---G------D-----L---P----------SE-------LPS---EQKVWL-----Y-----------PEF----------------ESQREQ---------ENE-WL-DKITRQIARGLI-----NH-----Y---S-RSKVIFN------P-L----------LL----ADQEL------------LAIET----------VV----------A-----SNRENFR------------------------------------------------------------
WP_083399742 ----------------------------------------------------M--EGF------TVLD--PAIV--T-FFA-ER-----K---EAW---------------------------------------------------------------------------------L--K-----K-NVS------T-SM------Q-----E--S------------------------------------------------------------------------------------------EV---------------------------------------------L-EKQ-RE------------CK-------
QN------FL---LANW-----LP---DA-------AKRA---------G-----QIS-I----------------ASHPCTFSHPS----A-------------R------K-NKNGYVSSI-----I----A---K------------NK---PSA--D--GF---------LRS--G--NV------S--------------V--E-PD-A-LG----N-----AA-----A--L------DVYKFL-S-----L-----P-MS--D---------QRSL-------------LVHIE-QES-E---LARQLL---E---V----PT-----CDYQTLRDGFLKM--I---DS--DQ-A-
SVSSSSKIKQVYFPV----------------------AEGE----YHLLSLLTHSGHLFELRKR-LD----A--M----R--F-G-----------------EL-VKEAR-E-------CK-------------K--ANRFH--PE-G-YQE--------IFG-LTTIGFGG-T------KPQ--------NISVLN---NQNA-----------------GKAHLLPSI-P-----PNLT-PR---EI-RLPKTD---FFKECF-----TA--------------WQA---KEVLEALHRLF---------KT-----D-Y-N------------------------------------IN--
IR--DG----RD------YRIQQYVDLVIEKMWQVR--L---F-L-T--D-HQ---G------D-----L---P----------SE-------LPL---EQKVWL-----Y-----------PEF----------------ESQREQ---------ENE-WL-DKITRQIARGLI-----NH-----Y---S-RSKVISN------P-L----------LL----ADQEL------------LAIET----------VV----------A-----SNRENFR------------------------------------------------------------
WP_107110711 ----------------------------------------------------M--EGF------TVLD--PAIV--T-FFA-ER-----K---EAW---------------------------------------------------------------------------------L--K-----KN------V-----S----T-S----MQ-----ES------------------------------------------------------------------------------------------EV---------------------------------------------L-EKQ-RE------------CK-------
QN------FL---LANW-----LP---DA-------AKRA---------G-----QIS-I----------------ASHPCTFSHPS----A------RKNKNG-----Y---V--S-----SI----I----A-K-----------NK---PSA---D--GF---------LRS--G--NV------S--------------V--E-PD-A-LG----N-----A---A--A-----LDVYKFL-S-----L-----P-MS--D---------QRSL-------------LVHIE-QES-E---LARQLL---E---V----PT-----CDYQTLRDGFLKM--I---DS--DQ-A-SVSSS-
KIKQVYFPV----------------------AEGE----YHLLSLLTHSGHLFELRKR-LD----A--M----R--F-G-----------------EL-VKEAR-E-------CK-------------K--ANRFH--PE-G-YQE--------IFG-LTTIGFGG-T------KPQ--------NISVLN---NQNA-----------------GKAHLLPSI-P-----PNLT-PR---EI-RLPKTD---FFKECF-----TA--------------WQA---KEVLEALHRLF---------KA-----DYN------N------------------------------------IN---IR--DG--
--RD------YRIQQYVDLVIEKMWQVR--L---F-L-T--G-HQ---G------D-----L---P----------SE-------LPL---EQKVWL-----Y-----------PEF----------------ESQREQ---------ENE-WL-DKITRQIARGLI-----NH-----Y---S-RSKVISN------P-L----------LL----ADQEL------------LAIET----------VV----------A-----SNRENFR------------------------------------------------------------
WP_101058358 ----------------------------------------------------M--EGF------TVLD--PAIV--T-FFA-ER-----K---EAW---------------------------------------------------------------------------------L--K-----KN------V-----S----T-S----MQ-----ES------------------------------------------------------------------------------------------EV---------------------------------------------L-EKQ-RE------------CK-------
QN------FL---LANW-----LP---DA-------AKRA---------G-----QIS-I----------------ASHPCTFSHPS----A------RKNKNG-----Y---V--S-----SI----I----A-K-----------NK---PSA---D--GF---------LRS--G--NV------S--------------V--E-PD-A-LG----N-----A---A--A-----LDVYKFL-S-----L-----P-MS--D---------QRSL-------------LVHIE-QES-E---LARQLL---E---V----PT-----CDYQTLRDGFLKM--I---DS--DQ-A-SVSSS-
KIKQVYFPV----------------------AEGE----YHLLSLLTHSGHLFELRKR-LD----A--M----R--F-G-----------------EL-VKEAR-E-------CK-------------K--ANRFH--PE-G-YQE--------IFG-LTTIGFGG-T------KPQ--------NISVLN---NQNA-----------------GKAHLLPSI-P-----PNLT-PR---EI-RLPKTD---FFKECF-----TA--------------WQA---KEVLEALHRLF---------K-TD-----Y-N------------------------------------IN---IR--DG--
--RD------YRIQQYVDLVIEKMWQVR--L---F-L-T--G-HQ---G------D-----L---P----------SE-------LPL---EQKVWL-----Y-----------PEF----------------ESQREQ---------ENE-WL-DKITRQIARGLI-----NH-----Y---S-RSKVISN------P-L----------LL----ADQEL------------LAIET----------VV----------A-----SNRENFR------------------------------------------------------------
WP_123117058 ----------------------------------------------------M--EGF------TVLD--PAIV--T-FFA-ER-----K---EAW---------------------------------------------------------------------------------L--K-----KN------V-----S----T-S----MQ-----ES------------------------------------------------------------------------------------------EV---------------------------------------------L-EKQ-RE------------CK-------
QN------FL---LANW-----LP---DA-------AKRA---------G-----QIS-I----------------ASHPCTFSHPS----A-------------R------K-NKNGYVSSI-----I----A---K------------NK---PSA--D--GF---------LRS--G--NV------S--------------V--E-PD-A-LG----N-----AA-----A--L------DVYKFL-S-----L-----P-MS--D---------QRSL-------------LVHIE-QES-E---LARQLL---E---V----PT-----CDYQTLRDGFLKM--I---DS--DQ-A-
SVSSSSKIKQVYFPV----------------------AEGE----YHLLSLLTHSGHLFELRKR-LD----A--M----R--F-G-----------------EL-VKEAR-E-------CK-------------K--ANRFH--PE-G-YQE--------IFG-LTTIGFGG-T------KPQ--------NISVLN---NQNA-----------------GKAHLLPSI-P-----PNLT-PR---EI-RLPKTD---FFKECF-----TA--------------WQA---KEVLEALHRLF---------K-TD-----Y-N------------------------------------IN--
IR--DG----RD------YRIQQYVDLVIEKMWQVR--L---F-L-T--G-HQ---G------D-----L---P----------SE-------LPL---EQKVWL-----Y-----------PEF----------------ESQREQ---------ENE-WL-DKITRQIARGLI-----NH-----Y---S-RSKVISN------P-L----------LL----ADQEL------------LAIET----------VV----------A-----SNRENFR------------------------------------------------------------
WP_018298695 ----------------------------------------------------MN--KIT--A-FFT-EK-----K---EAW---------------------------------------------------------------------------------L--K-----KN---------Q-------A----A-----S----E-----EM------------------------------------------------------------------------------------------A-VLK-SE-------------------------------------CE-----HT--
FS---LNVW-----LP---SA-------AKRA---------G-----QIS-I----------------ATHPCTFSHPS----A------RKNKNG-----Y---V-T-----SI----I----A-K-----------QAN--D--GF---------LKT--G--NV------S--------------V--M-SD-A-LG----N-----A---A--A-----LDVYKFL-S-----L-----V-ME--D---------QRSL-------------LSHIQ-QDS-K---LAKELL---N---L----DA---ANYEELKSGFLM---I--AS--ST-E-NITSS-
KIKQVYFPV--------------------------------GDD----YHQLSLLTNSGIVYELRNR-LD-----L----R--F-S----------------DS-VKELR-A-------CK-------------KY--NNEYS-EQ-G-YQE--------IYN-LTTIGYGG-T------KPQ--------NISVLN---NQHG-----------------GKAHLLSSV-P-----PSIE-KR---NT-YFPKRD---FFIESM---------QK------------AHY--IGTFKALHKIF---------QL-----DYN------N-------------------------------------------IK--IR--NA--
RD------YRLQELMNMIVENMWAVR-----L-V-S-----VA-Q---YH-------E------E------T----------ST------LEK--WQLIWL-------------------HENFEK----ERE--DSEQWL-DKIVKNISHWISQTYEKIM-EK-------Q-------A---I---KL----GEAEY---KHISN-----------IV----------T-----KNKEALR------------------------------------------------------------
WP_005497852 ----------------------------------------------------MLD--PAIT--S-FFA-ER-----K---EAW---------------------------------------------------------------------------------L--K-----KN--------I-------K------L-----A----S--------------------------------------------------------------------------------QT------------------------------------------------R-EAQ-QQ------------------CE----AA--
--FS---LSIW-----LP---KA-------AKRA---------G-----QIS-M-----------------ATHPCTFSHPS----A------RKNKNG-----Y---V-S-----SV----I----A-K-----------AE---RAT--D--GF---------LRS--G--NV-------S--------------V--D-TD-A-LG----N-----A---A--A-----LDVYKFL-S-----L-----E-MT--D-----------GGQL-------------LTHIQ-HDS-E---LATQLL---T---L----KD---VTYLELKAGFVAM--A---ES--GS-E-NVTSS-
KIKQVYFPV--------------------DDD----YHQLSLLTNSGMLYQLRKR-ID-----T----L----R--F-S---------------DS-VKVLR-D-------LK-----------K--KNEYS--AQ-G-FAE--------IYN-LTTIGYGG-T------KPQ--------NISVLN---NQHG-----------------GKAHLLSSL-P-----PQIE-KR---DL-HFPRSN---FFSESF---------R-P--YEY--KDVFQALHRIF---------KI-----DYK------N-------------------------------------------IK---LR--NS--
--RD------ARIHEVMDKVIAKMWAIR--------S-I-S--E--LQ-----YN--------S-------E------SQ------LSR---AQKIWL---------------HHDFTE--QRE-NTDEWL-DTLLNEISSWFA-----ST----Y---K-TAVGKQA-----------L---GEEER-------LHFLK--------------IV-------------T-----QHKEALR------------------------------------------------------------
WP_075473507 ----------------------------------------------------MLD--PAIK--S-FFD-ER-----K---EAW---------------------------------------------------------------------------------L--K-----KK--------V-------K------A-----S----MD-----ES---------------------------------------------------------------------------------------------------------------------------------N-ELN-IE------------------CD-------NT--
--FS---LAIW-----LP---DA-------AKRA---------G-----QIA-V-----------------ATHPCTFSHPS----A------RKNKNG-----Y---V-T-----SV----I----A-K-----------VE---RVN--D--GF---------LKS--G--NV-------S--------------V--D-TD-A-LG----N-----A---A--A-----LDVYKFL-T-----L-----E-MV--D---------QRSL-------------LDHIK-DDS-E---LAVNLL---N---L----KT---ESYLALKNGFLAM--V---NT--VG-E-NVTSS-
KIKQVYFPV--------------------DDD----YHQLSLLTNSGMLYQLRKR-LD-----T----L----R--F-S---------------DE-TKAIR-E-------LK-----------R--KNEFS--EK-G-FSE--------IYN-LTTIGYGG-T------KPQ--------NISVLN---NQNA-----------------GKAHLLSSL-P-----PQLD-KR---TI-HFPRSN---FFIESF---------K-T--HEY--RDVFQALHRIF---------RI-----DYK------N-------------------------------------------VS---LR--NS--
RD------ARIQELVDRIIDKMWAIR--------A-V-S--A-EQ----YH-------S------S-----D----LDA---NHKMWL-----H-----------SDDAQ--REE--YDD-WL-DIVVGDIASWMT----GA-----Y-------KK---------IIGKESVM-F-------GEEER-------LHFLK--------------------T-----KNKEALR-------------------------------------------------------------
WP_054453996 ----------------------------------------------------MLA--TEVD--N-FFE-KL-----K---ENW---------------------------------------------------------------------------------R-----KKIK-----A---A------MP-----IE-------------------------------------------------------------------------------------------------------------------------------------E-SLI-KE------------------CD---------SH--
YS---LNNW-----LS---YA-------SNGA---------I-----KRA-L-----------------SSHPSKFTHPD----VGASDTNIK------KKTYV-S---PV--I----H-----L-----------GN---FEN--N--GF----------LKT--G--NV-------S--------------IGIN-LD-S-TG-----N-----GA------E-V-GSIGEIGAFL-S-----LT-L-----A-----N---------GKRF-------------FEHLR-EDS-E---QARKIL---S-----Y---CT---EPYGDLKVGLLAM--I---K--DV-E-
AQTSSKIKQVYFPV--------------------DDD--YHQLSLLTNSGIVYQLRSR-LD-----K--L----R--F-S-----------------DE-VKERR-D-------KK-------------R--KNEFS--EQ-D-FSE--------IYG-LTTIGYGG-T------KPQ--------NISVLN---NQHG-----------------GKAHLLSSL-P-----PSIE-KR---SI-HFPKSD---FFTESF---------KE--YEY--ADAFKALHKIF---------QT-----DYN------N-------------------------------------------IK--
LR--DA----RD-----RNLQQIIDLIIEKMWSVR--------A-V-S-Q-AQ-----FH-----A-------D-------T----SS-----LSS---VQRTWL-----------------HDDFAD--ERE--KSEQWL-NSLTTEISAWLT-----RSYEKL---DK-------K------A--I----KL-----GDAER--------FHFAE---------IV----------T-----QNKEALR----------------------------------------------------------------
WP_011769775 ----------------------------------------------------MLD--PAIE--S-FFS-DR-----K---EAW---------------------------------------------------------------------------------L--K-----KKVN-----A------S-----MP-----IE-----------------------------------------------------------------------------------------------------------------------------------------N-VLQ-LE----------------------------QF--
--FS---LNEW-----LP---NA-------ASRA---------G-----QIS-L-----------------ATHPCTFSHPS----A------RKN-----KNGYV-S-----SV----I----A-K-----------VQ---RAE--D--GF---------LRS--G--NV-------S--------------V--V-D-TD-A-LG----N-----A---A--A-----LDVYKFL-T-----L-I-M---A--------Q---------NRSL-------------LTHIQ-QDS-A---LAVNLL---K---L----MT---KSYQDLKTGFLAM--I---ES--SV-E-
NITSSKIKQVYFPV----------------------------DND---YHQLSLLTNSGMLYQLRKR-ID-----R--L----R--F-S-----------------DQ-VKELR-D-------KK-------------R--KNEFS--EQ-G-YCD--------IYG-LTTIGYGG-T------KPQ--------NISVLN---NQHG-----------------GKAHLLSSL-P-----PTIE-KC---HT-HFPKSD---FFYESF---------QQ-----QEA--LSLFYFLQKII---------KN----KYN------N-------------------------------------------DN---IR-
-DA-----RN-----DCIQEIIDKIIDKMWAVR--------A-I-S--A-EQ-----YH-------Q-------E-------S-------SN------LKD---HQLIWL---------------HQNFAQ--KRE--ATDLWL-DQLTTEIARWLS-----RAYEKMI---GK-------Q-------A---I--KL-----GEQEL--------AHIEQ-------IV----------D-----QNREALR-----------------------------------------------------------
WP_130044827 ----------------------------------------------------MLD--PVIN--N-YFS-EL-----K---EKW----------------------------------------------------------------------------------L--K-----KNIK-----A----S-----MT-----DN-----------------------------------------------------------------------------------------------------------------------------------------N-ELK-GC----------------------------CE----
-FS---LNEW-----LP---KA-------SGRA---------G-----SRS-F----------------STHPSKFTHPS---TGVGDKNKK------NYTYV-T-----PV----I----C----N---------AE---RAN--D--GF---------LKT--G--NV-------S--------------I-TK-MD-S-LG-----N-----AA-----E-L-------DVDKFL-T-----LV-M-----E----N-----------GNSL-------------LEHIQ-QDT-D---LAVNLL---K---I----KA---VSYQELKSGFMAM--V---EI--SS-E-
NITSSKIKQVYFPI----------------------YNSDNSGSNDYHLLSLLTNSGMVYQLRER-LD-----T--L----R--F-S-----------------DE-VSELR-E-------KK-------------R--KNELS--EQ-G-ISE--------IYG-LTTIGYGG-T------KPQ--------NISVLN---NQHG-----------------GKAHLLSSL-P-----PSIE-KR---DI-HFPKSD---FFIESF---------KK--YDY--ADTFKALHKIF---------QA-----DYN------N-------------------------------------------IK--
-IR--NA----RD-----RYLQDIMDSLIEKMWAVR--------S-I-S--K-EQ----FH------K-------E------S-------SS------LSS---HQIIWL----------------YSEYEK--ERV--DSEQWL-DKVVDDISRWIS-----RSYEKLL---AK-------Q-------A----I-------KL-----GEEER--------LFVN--------------------------------------E-----QNREFLK----------------------------------------------------------
WP_138620165 ----------------------------------------------------MLD--PAIQ--V-FLS-ER-----K---EAW---------------------------------------------------------------------------------L--K-----KKIT-------T----S-----MP-----IE----------------------------------------------------------------------------------------------------------------------------------------E-ETE-QE-------------------------QV--
FS---LNEW-----LP---NA-------AKRA---------G-----QIS-M----------------STHPCTFSHPS----A------RKN-----KNGYV-S-----SV----L----A-K-----------IE---HTA--D--GF---------LKT--G--NV-------S--------------V--A-TD-A-LG----N-----A---A--A-----LDVYKFL-T-----LN-M-----E------N---------GNTL-------------LTHIE-LDT-E---LAKSLL---T---I----KS---ESYENLKKGFLAM--V---ES--AS-E-
SITSSKIKQVYFPV-------------------DDD----YHQLSLLTNSGMVYELRSR-LD-----K--L----R--F-S-----------------EN-VKILR-D-------KK-------------R--KNEFS--EQ-G-FSE--------IYG-LTTIGYGG-T------KPQ--------NISVLN---NQHG-----------------GKAHLLSSL-P-----PSIE-KR---NI-HFPKSD---FFIESF---------KQ--YEY--ADIFKALHKLF---------QA-----DHN------N-------------------------------------------IK--
EA----RD-----RYLQQIIDLIIEKMWAVR--------A-V-S-S-Q-TQ----FH------Q-------D------S-------SN------LSS---HQRAWL-----------------RSELTE--QRE--NDTQWL-DTLTQEISRWFS-----RRYEKVL---GK-------Q-------A-V------KL-----GEAER-------LLFAN---------SV----------E-----QNKEFLK-----------------------------------------------------------
WP_081178797 ----------------------------------------------------MLD--PAIQ--N-YFS-ER-----K---EAW---------------------------------------------------------------------------------L--K-----KNAK---A-----M-----P-----SD-----EL-----------------------------------------------------------------------------------------------------------------------------------Q-ELE-QL---------------------------QQ--
--FS---LNEW-----LP---NA-------AKRA---------G-----QIS-M-----------------STHPCTFSHPS----A------RKN-----KNGYV-S-----SV----I----A-K-----------ID---SVD--D--GY---------LKT--G--NV-------S--------------A-VA-TD-A-LG----N-----A---A--A-----LDVYKFL-T-----LT-M-----A------N---------GDNL-------------LTHIQ-QET-D---LAQTLL---T---L----KS---ANYQELRSGFLAM--V---ES--AT-E-
NITSSKIKQVYFPV--------------------DDD----YHQLSLLTNSGMVYQLRSR-LD-----T--L----R--F-S-----------------EE-VKELR-D-------KK-------------R--KNEFS--EQ-G-FSE--------IYG-LTTIGYGG-T------KPQ--------NISVLN---NQNG-----------------GKAHLLSSS-P-----PSIE-KR---NV-HFPKSD---FFTESF---------K-K--YEY--ANNFKALHKLF---------QT-----DYN------N-------------------------------------------IK--
LR--DA----RD-----RNLQQIIDLLIEKMWAVR--A-V-S-Q-T-Q-----YH-------A-------E-------T-------SS-----LSS---IQRTWL-----------------HDDFAD--ERE--KSAQ-WL-NSLITEISAWLT-----RS-----Y---E-KVLDKKA-----------L---GEAER-------LHFAE---------IV----------T-----KNKEALR--------------------------------------------------------
WP_086981311 ----------------------------------------------------MLD--PAIQ--S-YFA-ER-----K---EAW---------------------------------------------------------------------------------L--K-----KNVK-----N----A-----S-----MQ-----EH-----------------------------------------------------------------------------------------------------------------------------------------K-ELE-KL-----------------------------CE------QQ--
FS---LNEW-----LP---NA-------AKRA---------G-----QIS-I----------------STHPCTFSHPS----A------RKN-----KNGYV-S-----SV----I----A-K-----------ID---RIE--D--GF---------LKT--G--NV-------S--------------V-VT-TD-A-LG----N-----A---A--A-----LDVYKFL-T-----LS-M-----E------H---------GVDL-------------LTHIQ-QET-E---LAQTLL---T---I----QS---TNYQELRNGFLAM--I---ES--AS-E-
SITSSKIKQVYFPV------------------------DDD----YHQLSLLTNSGMVYQLRSR-LD-----T--L----R--F-S-----------------DE-VKELR-D-------KK-------------R--KNEFS--EQ-S-FSE--------VYG-LTTIGYGG-T------KPQ--------NISVLN---NQNG-----------------GKAHLLSSL-P-----PSIE-KR---NH-HFPRSD---FFVESF---------K-Q--YEY--ADIFKRLHKIF---------LA-----DYN------N-------------------------------------------IK--
IR--DA----RD-----RYLQQIIDLLIEKMWSVR--A-V-S-Q-Q-----YH-------E-------D-------S-------S-------LSS---VQRTWL-----------------HNDLAN--ERE--KSEQ-WL-DSLVTEISAWLT-----RS-----Y---E-QVLAKKA-----------L---GEAER-------LLFAE---------RV----------E-----RNRGFLK--------------------------------------------------------
WP_008169482 ----------------------------------------------------MLD--PAIQ--N-YFI-ER-----K---EAW---------------------------------------------------------------------------------L--K-----KNVK-----A----S-----MP-----EI-----------------------------------------------------------------------------------------------------------------------------------------K-ELE-LL-----------------------------CE------QQ--
FS---LNEW-----LP---SA-------AKRA---------G-----QIS-M----------------STHPCTFSHPS----A------RKN-----KNGYV-S-----SV----I----A-K-----------ID---RIE--D--GF---------LKT--G--NV-------S--------------A-VA-TD-A-LG----N-----A---A--A-----LDVYKFL-T-----LN-M-----E------N---------GCNL-------------LTHIQ-DET-E---LAQTLL---T---I----KS---TNYQELRAGFLTM--I---EG--AT-E-
SITSSKIKQVYFPV------------------------DDD----YHQLSLLTNSGMVYQLRSR-LD-----K--L----R--F-S-----------------DE-VKELR-D-------KK-------------R--KNEFS--EQ-G-FSE--------IYG-LTTIGYGG-T------KPQ--------NISVLN---NQNG-----------------GKAHLLSSL-P-----PSIE-KR---NI-HFPKSN---FFIESF---------K-K--YEY--ADTFKALHKLF---------QA-----DYN------N-------------------------------------------IK--
IR--DA----RD-----GYLQNIIDLIIEKMWAVR--------AVSQTQFHE---G---------T-------SS-------LSS---TQRTWL-----------------LANNIK--ERE--NSEQWL-DLLVTEIASWFS-----RSYEKL---DKQ-------A-----I-------KL-----GEAER-------LLFAD---------II----------N-----RNREFFK-----------------------------------------------------------
WP_118961040 ----------------------------------------------------MLD--PAIN--S-FFI-ER-----K---ETW---------------------------------------------------------------------------------L--K-----KNVK-----A----S-----MP-----EI-----------------------------------------------------------------------------------------------------------------------------------------K-ELS-QV-----------------------------CE------QQ--
-FS---LNEW-----LP---NA-------ASRA---------G-----QIS-M----------------STHPCTFSHPS----A------RKN-----KNGYV-S-----SV----I----A-K-----------VE---RIE--D--GF---------LKT--G--NV-------S--------------S-AA-TD-A-LG----N-----A---A--A-----LDVYKFL-T-----LT-M-----M----A---------GKSL-------------LEHIQ-QDT-D---LAKALL---T---I----KS---VSYQELKAGFMAM--A---ES--SA-E-
NITSSKIKQVYFPV-----------GD----------NDIANSETNNYHQLSLLTNSGMVYQLRSR-LD-----K--L----R--F-S-----------------DE-IKELR-D-------KK-------------R--KNEFS--EQ-G-FSE--------IYG-LTTIGYGG-T------KPQ--------NISVLN---NQNG-----------------GKAHLLSSL-P-----PSIE-KR---DV-YFPTSD---FFVESF---------KQ--------K-K-YDY--ADTFKALHKVF---------QT-----DYN------N-------------------------------------------IK--
IE---TR--DK----RD-----RYLQDIIDLLIEKMWSVR--A---ISQ-A-Q-----YH-------Q------D----------T--------SK-------IAS---HQLTWL------------------HSDYEK--ERE--SSDQ-WL-DKLVIEISRWIS-----RS-----Y---E-KVLARKA-----------L---TL-GEEER------LLFAN------------IV----------E-----HNREFLK--------------------------------------------------------
WP_033079010 ----------------------------------------------------MLD--PAIQ--S-YFS-ER-----K---EAW---------------------------------------------------------------------------------L--K-----KNVK-----A----S-----MP-----EL-----------------------------------------------------------------------------------------------------------------------------------------K-ELE-QT-----------------------------CD------QQ--
--FS---LNEW-----LP---NA-------AKRA---------G-----QIS-M----------------STHPCTFSHPS----A------RKN-----KNGYV-S-----SV----L----V-E-----------VD---RIE--D--GF---------LKT--G--NV-------S--------------S-VA-TD-A-LG----N-----A---A--A-----LDVYKFL-T-----LT-M-----E------D---------GTTL-------------LSHIQ-QDS-E---LATKLL---S---I----KS---ESYENQLKVGFLAM--V---ES--SA-E-
NITSSKIKQIYFPV-----------------------ENE----YHQLSLLTNSGMVYQLRSR-LD-----K--L----R--F-S-----------------DE-VKELR-D-------KK-------------R--KNEFS--EQ-G-YSE--------IYG-LTTIGYGG-T------KPQ--------NISVLN---NQNG-----------------GKAHLLSSL-P-----PSIE-KR---NI-HFPKSD---FFVESF---------KK----------Y-D-Y--TDTFKALHLIF---------QA-----DHN------N-------------------------------------------IK--
DA----RD-----RRLQEIIDLLIEKMWSVR--------A-V-S-Q-Q-----FH------Q-------E------A-------SS------LPS---YQRTWL-------------------HDDFSV--QRE--ESEQWL-DKLVNEIASWIT-----RSYEKTL---AK-------Q--------A----L----KL-----GEQEL-------LLFTD---------VV----------E-----QNRESLK------------------------------------------------------------
WP_042525438 ----------------------------------------------------MLN--SSID--E-FFK-KR-----K---EAW---------------------------------------------------------------------------------L--K-----KA---------K------A-----A------M-----D-----EI--------------------------------------------------------------------------------------------------------EV-----------------------------------------S-EKE-LE------------------------CE-----QV--
FA---LEHW-----LP---SA-------AKRA---------G-----QIS-I----------------STHPCTFSHPS----S-----RKNKNG-----Y---A-S-----SI----I----A-K-----------SK---RSE--D--GY---------LKS--G--NV-------S--------------V--K-AD-A-LG----N-----A---A--A-----LDVYKFL-S-----L-----E-MA--D---------GQTL-------------LSHIQ-QDS-D---LAKSLL---S---I----KS---ESYNTLKDGFLAM--I---NR--NS-D-CVTSS-
KIKQVYFPL-----------------DNG---YHQLSILSASGIMFELRKR-ID-----T--I----R--F-S-----------------DE-AKAAR-A-------CE-------------K--NNEIY--DG-D-YKQ--------IVN-ITTIGYGG-T------KPQ--------NISVLN---NQNG-----------------GKVHLISSE-P-----PKLI-KR---DAHFPTVD---FF--SQ-----SLR-----Y--RDQYSKLFSHLEF-------LNHKN-----D-------------------------------------WN--IR--HE--
RD------EYYQAIIDLIIEKMWLVR--------S-V-S-----Q-VQ-----Y-------N-------P-------D-------S-------S-----T-------LNE---TQKIWL--------------CEVNKE--NRE--TENDWL-DDLCXAIARFIF-----S-----TGYEKLL---G-----KK-------A-------F-------S---DDEF-------KHIKK-----------------------------Q-----VVKNKEALR--------------------------------------------------------
WP_005769537 ----------------------------------------------------MLN--SSID--E-FLK-KR-----K---EAW---------------------------------------------------------------------------------L--K-----KA---------K------A-----A------M-----D-----EI--------------------------------------------------------------------------------------------------------EV-----------------------------------------S-EKE-LE------------------------CE-----QV--
FA---LEHW-----LP---SA-------AKRA---------G-----QIS-I----------------STHPCTFSHPS----S-----RKNKNG-----Y---A-S-----SI----I----A-K-----------SK---RSE--D--GY---------LKS--G--NV-------S--------------V--K-AD-A-LG----N-----A---A--A-----LDVYKFL-S-----L-----E-MA--D---------GQTL-------------LSHIQ-QDS-D---LAKSLL---S---I----KS---ESYNTLKDGFLAM--I---NR--NS-D-CVTSS-
KIKQVYFPL-----------------DNG---YHQLSILSASGIMFELRKR-ID-----T--I----R--F-S-----------------DE-AKAAR-A-------CE-------------K--NNEIY--DG-D-YKQ--------IVN-ITTIGYGG-T------KPQ--------NISVLN---NQNG-----------------GKVHLISSE-P-----PKLI-KR---DAHFPTVD---FF--SQ-----SLR-----Y--RDQYSKLFSHLEF-------LNHKN-----D-------------------------------------WN--IR--HE--
RD------EYYQAIIDLIIEKMWLVR--------S-V-S-----Q-VQ-----Y-------N-------P-------D-------S-------S-----T-------LNE---TQKIWL--------------CEVNKE--NRE--TENDWL-DDLCQAIARFIF-----S-----TGYEKLL---G-----KK-------A-------F-------S---DDEF-------KHIKK-----------------------------Q-----VVKNKEALR--------------------------------------------------------
WP_005772797 ----------------------------------------------------MLN--SSID--E-FFK-KR-----K---EAW---------------------------------------------------------------------------------L--K-----KA---------K------A-----A------M-----D-----EI--------------------------------------------------------------------------------------------------------EV-----------------------------------------S-EKE-LE------------------------CE-----QV--
FA---LEHW-----LP---SA-------AKRA---------G-----QIS-I----------------STHPCTFSHPS----S-----RKNKNG-----Y---A-S-----SI----I----A-K-----------SK---RSE--D--GY---------LKS--G--NV-------S--------------V--K-AD-A-LG----N-----A---A--A-----LDVYKFL-S-----L-----E-MA--D---------GQTL-------------LSHIQ-QDS-D---LAKSLL---S---I----KS---ESYNTLKDGFLAM--I---NR--NS-D-CVTSS-
KIKQVYFPL-----------------DNG---YHQLSILSASGIMFELRKR-ID-----T--I----R--F-S-----------------DE-AKAAR-A-------CE-------------K--NNEIY--DG-D-YKQ--------IVN-ITTIGYGG-T------KPQ--------NISVLN---NQNG-----------------GKVHLISSE-P-----PKLI-KR---DAHFPTVD---FF--SQ-----SLR-----Y--RDQYSKLFSHLEF-------LNHKN-----D-------------------------------------WN--IR--HE--
RD------EYYQAIIDLIIEKMWLVR--S-V-S-----Q-V-Q-----Y-------N-------------------------------C-------EVN-------KE-NR-E-T-END-WL-DDLCQAIARFIF-----TG-----Y---E-KLLGKKA-----------F---SDDEF-------KHIKQ-----------VV----------K-----NKEALR-----------------------------------------------------------------
WP_043880971 ----------------------------------------------------MLN--SSID--E-FFK-KR-----K---EAW---------------------------------------------------------------------------------L--K-----KA---------K------A-----A------M-----D-----EI--------------------------------------------------------------------------------------------------------EV-----------------------------------------S-EKE-LE------------------------CE-----QV--
FA---LEHW-----LP---SA-------AKRA---------G-----QIS-I----------------STHPCTFSHPS----S-----RKNKNG-----Y---A-S-----SI----I----A-K-----------SK---RSE--D--GY---------LKS--G--NV-------S--------------V--K-AD-A-LG----N-----A---A--A-----LDVYKFL-S-----L-----E-MA--D---------GQTL-------------LSHIQ-QDS-D---LAKSLL---S---I----KS---ESYNTLKDGFLAM--I---NR--NS-D-CVTSS-
KIKQVYFPL-----------------DNG---YHQLSILSASGIMFELRKR-ID-----T--I----R--F-S-----------------DE-AKAAR-A-------CE-------------K--NNEIY--DG-D-YKQ--------IVN-ITTIGYGG-T------KPQ--------NISVLN---NQNG-----------------GKVHLISSE-P-----PKLI-KR---DAHFPTVD---FF--SQ-----SLR-----Y--RDQYSKLFSHLEF-------LNHKN-----D-------------------------------------WN--IR--HE--
RD------EYYQAIIDLIIEKMWLVR--S-V-S-----Q-V-Q-----Y-------N-------------------------------C-------EVN-------KE-NR-E-T-END-WL-DDLCQAIARFIF-----TG-----Y---E-KLLGKKA-----------F---SDDEF-------KHIKQ-----------VV----------K-----NKEALR-----------------------------------------------------------------
WP_094308916 ----------------------------------------------------MLN--SSID--E-FFK-KR-----K---EAW---------------------------------------------------------------------------------L--K-----KA---------K------A-----A------M-----D-----EI--------------------------------------------------------------------------------------------------------EV-----------------------------------------S-EKE-LE------------------------CE-----QV--
FA---LEHW-----LP---SA-------AKRA---------G-----QIS-I----------------STHPCTFSHPS----S-----RKNKNG-----Y---A-S-----SI----I----A-K-----------SK---RSE--D--GY---------LKS--G--NV-------S--------------V--K-AD-A-LG----N-----A---A--A-----LDVYKFL-S-----L-----E-MA--D---------GQTL-------------LSHIQ-QDS-D---LAKSLL---S---I----KS---ESYNTLKDGFLAM--I---NR--NS-D-CVTSS-
KIKQVYFPL-----------------DNG---YHQLSILSASGIMFELRKR-ID-----T--I----R--F-S-----------------DE-AKAAR-A-------CE-------------K--NNEIY--DG-D-YKQ--------IVN-ITTIGYGG-T------KPQ--------NISVLN---NQNG-----------------GKVHLISSE-P-----PKLI-KR---DAHFPTVD---FF--SQ-----SLR-----Y--RDQYSKLFSHLEF-------LNHKN-----D-------------------------------------WN--IR--HE--
RD------EYYQTIIDLIIEKMWLVR--S-V-S-----Q-V-Q-----Y-------N-------------------------------C-------EVN-------KE-NR-E-T-END-WL-DDLCQAIARFIF-----TG-----Y---E-KLLGKKA-----------F---SDDEF-------KHIKQ-----------VV----------K-----NKEALR-----------------------------------------------------------------
WP_135513942 ----------------------------------------------------M------IMLE--PVIT--G-FFR-ER-----K---EAW---------------------------------------------------------------------------------L--K-----KA---------S-----S-----------------MD-----EN------------------------------------------------------------------------------------------EI--------------------------------------------H-DKK-AE------------------------CE-------QL--
-FS---LEEW-----LP---NA-------AKRA---------G-----QIS-I----------------STHPCTFSHPS----S------RRNKNG-----D---T-T-----SV----I----A-R-----------SQ---NQR--D--GF---------LKS--G--NV------E--------------V--Q-ND-A-LG----N-----A---A--A-----LDLYKFL-S-----L-----E-TD--D---------SRTL-------------LAHLE-DES-D---LAKALL---T---I----KNA---SYTELRNGFLAM--T---KS--AT-D-SITSS-
KIKQVYFPL-----------------KDD----YHLLSILTPSGIVFDLRKR-LD-----A--M----R--F-G-----------------DE-IKLAR-D-------EK-------------R--NNAYY--EQ-G-YRE--------IYG-LTTVGYGG-T------KPQ--------NISVLN---NQNG-----------------GKAHLLSSL-P-----PELK-KR---TD-HIPKHD---FFTQTV----------R-YQHC--RDIFQALHQLD-----------SK----DLN------N-------------------------------------------MH--VR--AA-
RD-------EFYLSIMDYIVERLWQLRSVA-----N-E-Q-----YN-------S-----Q-----------------HQ-------LPK---AQQIWL-----------------HEGKEE--TRN--STDDWL-DDILGSVVNFIV-----TY--------GYEKRIL---G----K--------E-------A-----------F-----------SDTEK-------KYMTK-------------------VV-------G-----QYKECLR--------------------------------------------
WP_111568398 ----------------------------------------------------MID--DAIN--S-FFA-DR-----K---SAW---------------------------------------------------------------------------------L--K-----KN-TK-----A----S-----M-----D-----A-E-----------------------------------------------------------------------------------------VR------------------------------------------------------------------------EFECE-AQ-----------
-FS---LANW-----LP---HA-------ASRA---------G-----QIA-I----------------STHPCTFSHPS----A------------------R------K-NKNGYVSSI-----I--A-T-S-----------KAN--D--GF---------LRS--G--NV-------S--------------V--A-AD-A-LG----N-----A---A--A-----LDVYKFL-T-----L-----TLAD---------GHTL-------------IQHLE-GDS-E---QAVNIL---S---A---A---ADKDTYPDYQALKQAFLAM---TTPSSE---
NITSSRIKQVYFPL-------------------NDTDLEETD--YHQLSLLTASGIVFELRQR-LD-----V--L----R--F-G-----------------EA-IKEAR-E-------LK-------------R--KNEFS--EQ-G-YSE--------IYG-LTTIGYGG-T------KPQ--------NISVLN---NQNG-----------------GKAHLLSSL-P-----PSIK-KR---DI-HFPKQD---FFSQTL----------R-YSQC--RRQFQALHTLY-----------RD-----NKK------N--------------------------------------------------
MH---IR--AL----RD-----DYYQSVIDHIIEKMWQVR--S-V-A-Q---A--EQ-----YR-------S-------S-----E-------LPA---AQQTWL---------------EQSIAL--RET--TDD-WL-DDCSSVTSFIF------LPYEKTL---G-----KK-------A-------S-----QEEV-------RHMKA--------------------------------L-----SNKEAIR-----------------------------------
WP_025008607 ----------------------------------------------------MMD--SAIS--E-FFA-DR-----K---SAW-----------------------------------------------------------------------------------L-------------K-----KNIK-------A--------SM-----SD-----VEIKAL--------------------------------------------------------------------------------------------------------------------------ECE-TQ--------------------------
FS---LANW-----LP---NA-------ASRA---------G-----QIA-I----------------STHPCTFSHPS----A------------------R------K-NKNGYVTSI-----I--A-T-S-----------HSE--N--GF---------LKT--G--NV-------S--------------V--A-AD-A-LG----N-----A---A--A-----LDVYKFL-T-----L-----VLAD---------GHTL-------------IQHLE-RDS-E---QAVNLL---S---A---A---ANKEIYPDYQTLKQAFLAM--T---AT-SS---
ESITSSKIKQVYFPLNRGSNVNGH-------------LASIE----YHQLSLLTASGIVFELRQR-LD-----V--L----R--F-G-----------------ET-IKEAR-D-------KK-------------R--TNQFH--QG-Y-KE--------IYN-LTTIGYGG-T------KPQ--------NISVLN---NQNG-----------------GKAHLLSSL-P-----PNLK-NR---DV-HFPHSD---FFSQTL----------R-YTQC--KNQFQALHQLY-----------RG-----DDN------N--------------------------------------------------
---MH---IR--VQ----RD-----DYYQSVIDHVIEKMWQVR--S-V-A-Q-A-Q-----YR-------R-------E-------D-----L-------LPT---VQQTWL----------------CEH-------TKILRE-----TTDDWL-DVICNAITGFLF------HGYEKTL---G-----KK-------A-------KF-------GDAEY-------RQMQK--------------------------------L-----SNKEALR-----------------------------------
WP_011789484 ----------------------------------------------------MIDG-TIS--E-FFA-DR-----K---SAW-----------------------------------------------------------------------------------L-------------K-----KNIKASMSD----------EIK-------A-L---------------------------------------------------------------------------------------------------------------------------------ECE-TQ--------------------------
FS---LANW-----LP---NA-------ASRA---------G-----QIA-I----------------STHPCTFSHPS----A------------------R------K-NKNGYVTSIIAHSE-----KAN--D--GF---------LKT--G--NV-------S--------------V--A-AD-A-LG----N-----A---A--A-----LDVYKFL-T-----L-----VMAD---------GHTL-------------IQHLE-RDS-E---QAVNLL---S---A---A---ASKETYPDYQTLKQAFLAM--TATSSE---
SITSSKIKQVYFPL-------------------NRGSNVNGHLASIE---YHQLSLLTASGIVFELRQR-LD-----A--M----R--F-G-----------------ET-IKEAR-D-------KK-------------R--TNQFH--QG-Y-KE--------IYN-LTTIGYGG-T------KPQ--------NISVLN---NQNG-----------------GKAHLLSSL-P-----PNLK-NR---DV-HFPHSD---FFSQTL----------R-YTQC--KNQFQALHQLY-----------RG-----DD-N------N--------------------------------------------------
---MH---IR--AQ----RD-----DYYQSVIDHVIEKMWQVR--S-V-A-Q-A-Q-----YR-------R-------E-------D-----L-------LPT---VQQTWL----------------CEH-------TKILRE-----TTDDWL-DVICNAITVFLF------HG----Y-------E-----KTL---G-----KK-------A-------KF-------GDAEY-------RQMQK--------------------------------L-----SNKEALR-----------------------------------
WP_070993783 ----------------------------------------------------MID--KAVD--A-FFA-ER-----K---EAW---------------------------------------------------------------------------------L--K-----KN---------K-----A-----S--------M-----L-------------------------------------------------------------------------------------------------EI-------------------------------------------------------Q-QTH-ME------------------------CE-------SV--
-FS---LQNW-----LP---NA-------AKRA---------G-----QIA-I----------------STHPCTFSHPS----A------RKNKNG-----Y---V-S-----SI----I----A-K-----------SE---ASN--D--GF---------LKS--G--NV-------S--------------V--V-AD-A-LG----N-----A---A--A-----LDVYKFL-T-----L-----V-LE--D---------GQTL-------------LQHLE-QDT-D---LAQRL-F---R---T----TPMHEASDYDALKEGFLAM--T---AP-SK-E-VVTSS-
KIKQVYFPL--------------TAKNSDDVS--YHQLSLLTASGIALFALRKR-LD-----T--M----R--F-G-----------------DE-IKLAR-E-------KK-------------A--NLAH--PT-HCE------ISN-LTTIGYGG-T------KPQ--------NISVLN---NQNG-----------------GKAHLLMSM-P-----PKLQ-RR---DL-HFPRSD---FFAQVV----------R-YRHC--QAQFKALDVLY-------LLPNE------YH--------N------------------------------------------------MH--
TR---AK-----RD-----ELYQSVIDHVIEKMYLIR--------E-V-A-V-QQ-----YV-------A-------A-------S------------G----LAQ---AQRVWL-------------CEH---------DQ-------ERE--GTDDWL-DEICNAIVRFLF------HGYEKTL---G-----K--------K-------A-------V-------N-----IDDEF-------KHIRK--------------------------------L-----SNKEALR-----------------------------------
WP_119860827 ----------------------------------------------------MID--EAID--T-FFQ-ER-----K---AAW---------------------------------------------------------------------------------L--K-----KN---------I-----A-----K-----S------S----Y-------------------------------------------------------------------------------------------------AER-------------------------------------------------------K-DKH-ME-------------------------CE-------TI--
FS---LRVW-----LP---NA-------AKRA---------G-----QIS-I----------------STHPCTFSHPS----A------RKNKNG-----Y---V-S-----SI----V----A-K-----------SE---RCN--D--GF---------LKS--G--NV-------S--------------V--Q-AD-A-LG----N-----A---A--A-----LDVYKFL-T-----L-----V-TE--N---------GQTL-------------LQHLE-QDS-N---LAKNLF---R---S---TAMNEPSDYEVLKEGFLAM--I---AP-SE-E-VVTSS-
KIKQVYFPI-----------------TDESKHTQP---YHQLSLLTASGIVFELRKR-LD-----A--M----R--F-G-----------------DE-VKQAR-E-------QR-------------K--ANQVH--QS-HQE-----IID-LTTIGYGG-T------KPQ--------NISVLN---NQNG-----------------GKAHLLMSM-P-----PKLK-SR---DL-HFPTSD---FFSQTV----------R-YSQC--TKQFQALHKLY-------LLPNE------YH--------N------------------------------------------------MH--
TR---SE-----RD-----ELYQSVIDHVIEQMFTIR--------S-V-S-I-E----Q-----FN-------K-------L-------T-------S-------LPA---AQRIWL--------------CHDKAT--ERL--TTDNWL-EAICNSIVSFLF------YGYERTL-------G-------K-------K-A-------I-------K-----GDGEY-------LKIQE--------------------------------L-----SNKEILR-----------------------------------
WP_010364313 ----------------------------------------------------MID--EAID--T-FFE-ER-----K---EAW---------------------------------------------------------------------------------L--K-----KN---------I-----A-----K-----S------S----Y-------------------------------------------------------------------------------------------------AER-------------------------------------------------------K-DKH-ME-------------------------CE-------TI--
FS---LRVW-----LP---NA-------AKRA---------G-----QIS-I----------------STHPCTFSHPS----A------RKNKNG-----Y---V-S-----SI----V----A-K-----------SE---RCN--D--GF---------LKS--G--NV-------S--------------V--Q-AD-A-LG----N-----A---A--A-----LDVYKFL-T-----L-----V-TE--N---------GQTL-------------LQHLE-QDS-N---LAKNLF---R---S---TAMNEPSDYEVLKEGFLAM--I---AP-SE-E-VVTSS-
KIKQVYFPI-----------------TDESKHTQP---YHQLSLLTASGIVFELRKR-LD-----A--M----R--F-G-----------------DE-VKQAR-E-------QR-------------K--ANQVH--QS-HQE-----IID-LTTIGYGG-T------KPQ--------NISVLN---NQNG-----------------GKAHLLMSM-P-----PKLK-SR---DL-HFPTSD---FFSQTV----------R-YSQC--IKQFQALHKLY-------LLPNE------YH--------N------------------------------------------------MH--
TR---SE-----RD-----ELYQEVIDHVIEQMFTIR--------S-V-S-I-E----Q-----FN-------K-------L-------T-------S-------LPA---AQRIWL--------------CHDKAT--ERL--TTDNWL-EAICNSIVSFLF------YGYERTL-------G-------K-------K-A-------I-------K-----GDGEY-------LKIQE--------------------------------L-----SNKEILR-----------------------------------
WP_007620986 ----------------------------------------------------MVD--PAIT--T-FFD-ER-----K---EAW---------------------------------------------------------------------------------L--K-----KN---------L-----T-----A-----S------M-----S---------------------------------------------------------------------------------------------QA-------------------------------------------------------REKEQE-------------------------CESV--
--FT---LKNW-----LP---DA-------AKRA---------KSRAM----------TSHPSKFTHPS---T----------G-VGPK------NIKNLTY-----V---T-----PIICN-----AVL---ENDGF----------LRT--G--NV--------R-----------------LVDP-ID-S-IG-----N-----AG-----E-L-------DVHEFL-T-----L-----VMVD---------GRNL-------------IQHLE-QNT-E---LALSLF---T---LPNIDET---QNYASLKQGLLAM--T---SA-GS---

[illegible]

WP_101198325
[illegible]

WP_067341808
[illegible]

WP_075179156
[illegible]

WP_133504568
[illegible]

WP_013662443
[illegible]

WP_127040988
[illegible]

WP_094489615
[illegible]

WP_081222277
[illegible]

WP_063503399
[illegible]

WP_063455796
[illegible]

WP_094491019
[illegible]

WP_094493348
[illegible]

WP_094537589
[illegible]

WP_094492037
[illegible]

WP_094488664
[illegible]

WP_066569288
[illegible]

WP_136143866
[illegible]

WP_065992419
[illegible]

WP_104895765
[illegible]

WP_104511153
[illegible]

WP_130073672
[illegible]

WP_130145891
[illegible]

WP_120364305
[illegible]

WP_013818080
[illegible]

WP_107789488
[illegible]

WP_090688312
[illegible]

WP_054762952
[illegible]

WP_096876028
[illegible]

WP_064024402
[illegible]

WP_064021411
[illegible]

WP_064041139
[illegible]

WP_088621132
[illegible]

WP_127026736
[illegible]

WP_024297106
[illegible]

WP_019959484
[illegible]

WP_120375540
[illegible]

WP_126132534
[illegible]

WP_004907049
[illegible]

WP_136143749
[illegible]

WP_004758712
[illegible]

WP_126624323
[illegible]

WP_018676856
[illegible]

WP_097078011
[illegible]

WP_087618841
[illegible]

WP_046760300
[illegible]

WP_004652094
[illegible]

WP_004963263
[illegible]

WP_125297836
[illegible]

WP_075696700
[illegible]

WP_005217654
[illegible]

-------------REAQ---YHLLSILTPSGLITRLKQS-ID---A--I---R--F-S--------------ED-TKQAK-E-----SR-------------K--KNEHH--EI-G-YSD-----IFD-LTVTGYGG-T----QPQ--------NVSVLN--SQNA----------GRAYLLSSC-P----PKFE-NR---NI-KLPSQD--FFQQCL-------------Y-W--KRF--GGEFEILQKYV---------NG------I------N------------------------N-----QH---TR--KK---IK------
KILRQMVGDILFIAFCIR--------K-N-E------K---G----WS-----KKE------H----Y----------QS--------LAL---KQKIWL-----------------------------------DDAYLD--VRE--TENEWR-DEIARRTAKWIL-----DSFGDFY---G-------SK----------L-F------------------SSPEI------IELKG----------IV--FE---------------------------------ALDEDKEFF-------
WP_119064436 --------------------------------------------------------------------------------MA--ESIY--T-FLN-ER------K----ELW--------------------------------------------------------------------------------K-------E-RI-----K------K--A---------EN-----ET-------------------------------------------VI--------------------------------------S-ELQ-KQ------------------AN--------DK------
FS---LNEW-----LP---DA-----AKRV-------T----QLS-M----------VSHPSKFSHPS---A-------------K----T--S-----SV-----I-----A---K----------VE---QSN--D--GY-------LRS--G--NV------D----------Y--S-LD-V-FG-----N---AA---A---M----DVFKFL-S------L----P-LN--E-------KFTV----------------LDGFE-QQD-Q---ALKTLI---T----H---AK-----LNFDTLSTEFLKI--K---VT--DH-S-VKTDH-LVKQVYFPV-
-------------REAQ---YHLLSILTPSGLITRLKQS-ID---A--I---R--F-S--------------ED-TKQAK-E-----SR-------------K--KNEHH--EI-G-YSD-----IFD-LTVTGYGG-T----QPQ--------NVSVLN--SQNA----------GRAYLLSTC-P----PKFE-NR---NI-KLPNQD--FFQQCL-------------Y-W--KRF--GSEFESLQKYV--------T---GI-------N------------------------N-----QH---TR--KK---IK------
KILRQMVGDILFIAFCIR--------K-N-E------K---G----WS-----KKE------H----Y----------QS--------LAL---KQKIWL-----------------------------------DDAYLD--VRE--TESEWR-DEIARRTAKWIL-----DSFEDFY---G-------SK----------L-F------------------SSPEI------IELKG----------IV--FE---------------------------------ALDEDKEFF-------
WP_110974473 --------------------------------------------------------------------------------MA--ESIY--T-FLN-ER------K----ELW--------------------------------------------------------------------------------K-------E-RIK------K------K--A---------EN-----ET-------------------------------------------VI--------------------------------------S-ELN-KQ------------------AN--------DK------
FS---LNEW-----LP---DA-----AKRV-------T----QLS-M----------VSHPSKFSHPS---A--------------KTS-------SV-----I-----A---K----------VE---QSN--D--GY-------LRS--G--NV------D----------YSLD-V-FG-----N---AAA---M----DVFKFL-S------L----P-LN--E-------KFTV----------------LDGFE-QQD-Q---ALKTLI---T----H---AK-----LNFDTLSTEFLKI--KV---TD--HS----VKTDHLVKQVYFPV-
-------------REAQ---YHLLSILTPSGLITRLKQS-ID---A--I---R--F-S--------------ED-TKQAK-E-----SR-------------K--KNEHH--EI-G-YSD-----IFD-LTVTGYGG-T----QPQ--------NVSVLN--SQNA----------GRAYLLSTC-P----PKFE-NR---NI-KLPSQD--FFQQCL-------------Y-W--KRF--GSEFESLQKYV--------T---GI-------N------------------------N-----QH---TR--KK---IK------
KILRQMVGDILFIAFCIR--------K-N-E------K---G----WS-----KKE------H----Y----------QS--------LAL---KQKIWL-----------------------------------DDAYLDVRETESEWR-DEIARRTAKWIL----------------------------------------DSFED---------
WP_116762640 --------------------------------------------------------------------------------MT--ASIE--A-FLN-ER------K----ELW--------------------------------------------------------------------------------L--K-------DR------L-------K------K--A---------EN-----ET-------------------------------------------VI--------------------------------------A-ELQ-QQ------------------AN--------ER------
FS---LQEW-----LP---DA-----AKRV-------S----QLS-M----------VSHPSKFSHPS---A-------------K----T--S-----SV-----I-----V---Q----------AK---HAV--D--GY-------LRS--G--NV------D----------Y--P--LD-V-FG-----N---A---A---M----DVFKFL-S------V---N-LD--N-------GQTV----------------LDAFE-KQD-E---DLKQFI---Q----R---SS-----LFDVLRLAFLVI---K---DN--DT-S-VKTDH-
LVKQVYFPI---------GG-----------------EN-----YHLLSLLTPSGLITKLKQN-ID---A--I---R--F-S--------------DA-TKLAK-D-----RR-------------K--KNELD--DV-G-YAD-----IYD-LTVTAYGG-T----QPQ--------NVSVLN--SQNA----------GRAYLLSSA-P----PQFE-KR---KV-NLPTKD--FFVQCL-------YWKKFNHE-FS-ILQKYV--RSDLKETEQVV-------------------------S-------N-------------------------------RH---
IR---KK---IR---------QTIQQIIYEILFIAFCIR-----------R-N-Q------S---G----WS-----------------------NKEY----YQYLPL---DQRIWL--------------------------------------DEVHLE--QRE--HQEEWR-DEISRRTAKWIL-----NS----Y---E-KFMSHSN---------M-F----------------SDVEI-------KVVKI-----------VV---LE------------------------------ILEQDKEFF-------
WP_092687408 --------------------------------------------------------------------------------MT--ASIE--A-FLN-ER------K----ELW--------------------------------------------------------------------------------L--K-------DRLK-------K---V--------EN-----ET-------------------------------------------VI--------------------------------------A-ELQ-QQ------------------AN--------ER------
FS---VQEW-----LS---DA-----AKRV-------S----QLS-M----------VSHPSKFSHPS---A-------------K----T--S-----SV-----I-----V---Q----------AD---YSV--D--GY-------LRS--G--NV------D----------Y--P--LD-V-FG-----N---AA---A---M----DVFKFL-S------VN-LE--N-------GQTI----------------LDAFE-QQN-E---DLKQFI---Q----H---SS-----LFDALRLAFLVI-------KE--ND-T-
SVKTDHLVKQVYFPV----------GG-----------------EN-----YHLLSLLTPSGLITKLKQN-ID---A--I---R--F-S--------------DA-TKLAK-E-----QR-------------K--KNELD--DV-G-YAD-----VYD-LTVTAYGG-T----QPQ--------NVSVLN--SQNA----------GRAYLLSSA-P----PQFE-KR---KV-KLPTKD--FFAQCL-------YWKKFNHE-FS-ILQKYV--RSDLKETEQVV--------------------------S-------N---
---RH---IR---KK----IR---------QTIQQIIYEILFIAFCIR-----------R-N-Q------S---G----WS-----------------------NKEY----YQHLPL---DQRIWL--------------------------------------DEVHWE--QRE--HQEEWR-DEISRRTAKWIL-----DS----Y---E-KFMSHSK---------M-F----------------SDLEI-------KEVKI-----------VV---LE------------------------------ILEQDKEFF-------
WP_086183622 --------------------------------------------------------------------------------MT--ASIE--A-FLN-ER------K----ELW--------------------------------------------------------------------------------L--K-------DR------L-------K------K--A---------EN-----ET-------------------------------------------VI--------------------------------------A-ELQ-QQ------------------AN--------DK------
FS---VQEW-----LS---DA-----AKRV-------S----QLS-M----------VSHPSKFSHPS---A-------------K----T--S-----SV-----I-----V---Q----------AD---YSV--D--GY-------LRS--G--NV------D----------Y--P--LD-V-FG-----N---A---A---M----DVFKFL-S------V---N-LE--N-------GQTI----------------LDAFE-QQN-E---DLKQFI---Q----H---SS-----LFDALRLAFLVI---K---DN--DT-S-VKTDH-
LVKQVYFPV---------GG-----------------EN-----YHLLSLLTPSGLITKLKQN-ID---A--I---R--F-S--------------DA-TKLAK-E-----QR-------------K--KNELD--DV-G-YAD-----IYD-LTVTAYGG-T----QPQ--------NVSVLN--SQNA----------GRAYLLSSA-P----PQFE-KR---AI-RLPKTD--FFAQCL-------------Y-R--KNF--QDSFLQLHKLM----------QL-----DVN-----N--------------------------ID---IR---TA--
---IR-------NIIQFVIDQVMFQAFKIR-----------Q--S-YA--EG-----WS-------N-----E---DYY--------VS------------LPK---LQRIWL---------------------------------------DDFYEN--ERE--QDDEWR-DELSHEIARWIL-----RG-----Y---EK---------SIEG-S-F-------------ML----GMGEL--------VNVKQ-R------------VD-------------------K------SLQHTKEFF-------
WP_004651370 --------------------------------------------------------------------------------MT--ASIE--A-FLN-ER------K----ELW--------------------------------------------------------------------------------L--K-------DR------L-------K------K--A---------EN-----ET-------------------------------------------VI--------------------------------------A-ELQ-QQ------------------AN--------DK------
FS---LQEW-----LP---DA-----AKRV-------S----QLS-M----------VSHPSKFSHPS---A-------------K----T--S-----SV-----I-----V---Q----------AK---HSV--D--GY-------LRS--G--NV------D----------Y--P--LD-V-FG-----N---A---A---M----DVFKFL-S------V---N-LD--N-------GQTV----------------LDAFE-KQD-E---DLKQFI---Q----R---SS-----LFDVLRLAFLVI---K---DN--DT-S-VKTDH-
LVKQVYFPI---------GE----------------KN-----YHLLSLLTPSGLITKLKQS-ID---A--I---R--F-S--------------DA-TKLAK-D-----RR-------------K--KNELD--KV-G-YAD-----IYD-LTVTAYGG-T----QPQ--------NVSVLN--SQNA----------GRAYLLSSA-P----PQFE-KR---TI-RLPKTD--FFAQSLY----------R-------KNF--QDSFLQLHKLM----------QL-----DVN-----N--------------------------ID---IR---TA--
-IR-------NIIQFVIDQVMFQAFKIR-----------Q--S-YA--EG-----WS-------N-----E---DYY--------AS------------LPK---LQRVWL---------------------------------------DDFYEN--ERE--QDDEWR-DELSREIARWIL-----RS-----Y---EK---------SIEG-S-F-------------TL----GTGEL--------VELKQ-R------------VE-------------------K------SLQQTKEFF-------
WP_104440802 --------------------------------------------------------------------------------MT--ASIE--A-FLN-ER------K----ELW--------------------------------------------------------------------------------L--K-------D-RLK--KAEN-------E-------T-------------------------------------------VI--------------------------------------A-ELQ-QQ------------------AN--------ER------
FS---LQEW-----LP---DA-----AKRV-------S----QLS-M----------VSHPSKFSHPS---A-------------K----T--S-----GV-----I-----V---Q----------AK---HAV--D--GY-------LRS--G--NV------D----------Y--P-LD-V-FG-----N---A---A---M----DVFKFL-S------V---N-LD--N-------QQTV----------------LDAFE-KQD-E---DLKQLI---Q----R---SS-----LFDVLRLAFLVI-------KD--ND-T-
SVKTDHLVKQVYFPI----------------------DEKK---YHLLSLLTPSGLITKLKQS-ID---A--I---R--F-S--------------DA-TKLAK-D-----RR-------------K--KNELD--DV-G-YAD-----IYD-LTVTAYGG-T----QPQ--------NVSVLN--SQNA----------GRAYLLSSS-P----PFFE-KR---AI-RLPKTD--FFAQCL-------------Y-R--KNF--QESFIQLHNFM----------Q-LN----L-N-----N------------------------------------IE---
IR--TA---IR-------NIIQFVIDQVIFQAFKIR-----------G--S-YAE--G-----WS-------N-----E---DYY--------YYAS------------LPK---LQRVWL---------------------------------------DGFYEN--ERE--QDDEWR-DELSREIARWIL-----RS-----Y---E-K---------SIEG-S-F-------------TL----GAGEL--------VELKQ-R------------VE-------------------K------SLQQTKEFF-------
WP_117007238 --------------------------------------------------------------------------------MT--ASIH--T-FLN-ER------K----ELW--------------------------------------------------------------------------------L--K-------DR------L-------K------T--------TN-----DI-------------------------------------------EI--------------------------------------A-ELQ-QQ------------------AN--------DK------
FH---LKEW-----LL---DA-----AKRV-------T----QLS-M----------VSHPSKFSHPS---A-------------K----T--S-----SI-----I-----A---P----------VS---ANK--D--GY-------LRT--G--NV------D----------Y--P---LD-V-FG-----N---A---M----DVYKFL-C------L---S-LS--N-------ELSV----------------LDGFE-QDD-Q---NLKNLI---Q----Q---AS-----LDFETLKTNFLMI---K---QN--DT-T-SKTDH-LIKQVYFPI--
-------------SQDE---YHLLSILTPSGLITKLKQR-ID---H--I---R--F-S--------------EQ-TKFAK-E-----KR-------------K--KNKHD--TE-G-YAD-----IFD-LTVTAYGG-T----QPQ--------NVSVLN--SQNA----------GRAYLLSSA-P----PQFE-KR---AI-RLPKTD--FFAQCL-------------Y-R--KNY--QESFIQLHKFM----------QL-----DIN-----N--------------------------ID---IR---TA---IH-----
NIIQFVIDQVLLQAFKIR-----------E---T-YP--ET-----WS-------N-----------DQDY----YLGLPK---LQRIWL---------------------------------------DQQYQN--ERE--NDAEWR-DEISQNIARWIL-----RS-----Y---E-K---------AIND-S-Y--------------TL----GTGEL--------LDVQQ-R-----------VE-------------------K------SLQHAKEFF-------
WP_087512541 --------------------------------------------------------------------------------MT--ESIQ--T-FLN-ER------K----ELW--------------------------------------------------------------------------------L--K-------DR------V-------K-----------ED-----DT-------------------------------------------EV--------------------------------------A-ELQ-QQ------------------AD--------DK------
-FS---LQEW-----LP---DA-----AKRV-------S----QLSMM----------VSHPSKFSHPS---A-------------K----T--S-----SV-----I-----A---Q----------VK---SAQ--E--GY-------LRS--G--NV------H----------Y--P-LD-V-FG-----N---A---A---M----DVYKFL-S-----L---S-LT--E-------DSTV----------------LMGFE-KDD-Q---DLKSLI---E----K---SS-----LDFASLKQAFLVI---K---ND--DA-S-SKTDH-
LVKQVYFPL----------------------ENNE---YHLLSILTPSGLITKLKQS-ID---Q--L---R--F-S--------------DA-TKEAK-E-----KR-------------K--KNEHD--AE-G-YAD-----VYD-LTVTAYGG-T----QPQ--------NVSVLN--SQNA----------GRAYLLSSC-P----PSFE-ER---KI-RLPTRD--FFEQCL-------------Y-S--KGY--KQSFKELQLVI------------RKDLD-----GET-----G-------------------------------KRKNTYL---
IR---KD---ID-------KTLRKIIEQILFKAFCIR-----------K-Y-K------K---E----WS-----IND-----S----Y----------KT--------LPL---YQRIWL---------------------------------------DDFYEE--QRK--DDDEWF-EEISLRISKWII-----NV----Y---E-DKEIIEN-----------S-Y--------------ML----GSGEI--------LAVKEIVK--------------------------E------LLDEDKEFF-------
WP_104492288 --------------------------------------------------------------------------------MT--ESIH--T-FLN-ER------K----ELW--------------------------------------------------------------------------------L--K-------DR------V-------K------K--A---------EN-----ET-------------------------------------------EI--------------------------------------A-ELQ-QQ------------------AD--------DR------
FS---LKEW-----LP---DA-----AKRV-------S----QLS-M----------VSHPSKFSHPS---A-------------K----T--S-----SV-----I-----A---Q----------VK---SAQ--D--GY-------LRS--G--NV------H----------Y--P-LD-V-FG-----N---A---A---M----DVYKFL-S------N-LA--E-------DYTV----------------LTGFE-HDH-E---DLKSLI---E----K---SS-----LNFTSLKQAFLVI---K---ND--DT-S-VKTDH-LVKQVYFPI--
-------------ADTE---YHLLSILTPSGLIARLKLA-ID---Q--L---R--F-S--------------EA-TKEAK-E-----KR-------------K--KNEHD--AI-G-YAD-----IFD-LTVTAYGG-T----QPQ--------NVSVLN--SQNA----------GRAYLLSSC-P----PILE-KR---TV-RLPRTD--FFVQCL-------------Y-R--KNY--QDSFVQLHKFM----------QL-----DLN-----N--------------------------ID---IR---TA---IH-----
NIIQFVIDQVLLQGLKIR-----------E---Y-YS--EG-----WS-------Q-----Q---EYY----------AN------------LPK---LQRIWL---------------------------------------DEMYAE--QRE--NDSEWR-NELSEDIARWIL-----RS-----Y---E-KVISESE-----------M-L--------------GTGEI--------NDVKQ-R-----------VE-------------------E------SLRQAKEFF-------
WP_087552551 --------------------------------------------------------------------------------MT--ENIQ--T-FLN-ER------K----ELW--------------------------------------------------------------------------------L--K-------DRVK-----K--A----------QN-----DT-------------------------------------------EV--------------------------------------T-ELY-QQ------------------AD--------DK------
-FS---LREW-----LP---DA-----AKRG-------S----QLS-M----------VSHPSKFSHPS---A-------------K----T--S-----SV-----I-----A---Q----------VK---SAQ--D--GY-------LRS--G--NV------H----------Y--P-LD-V-FG-----N---A---A---M----DVYKFL-S------L---S-LT--E-------DSTV----------------LMGFE-KDD-Q---DLKHLI---E----K---SS-----LDFASLKQAFLVI-------KN--DD-A-
SSKTDHLVKQVYFPL----------------------ENHE---YHLLSILTPSGLITKLKQS-ID---Q--L---R--F-S--------------DA-TKEAK-E-----KR-------------K--KNEHD--AE-G-YAD-----IFD-LTVTAYGG-T----QPQ--------NVSVLN--SQNA----------GRAYLLSSC-P----PVLE-KR---TV-RLPKTD--FFAQCL-------------Y-R--KNF--QDSFIQLHKFM----------QL-DL-----N-----N---------------------------------ID---
IR---TA---IR-------NIIQFVIDQVLLQGFKIR-----------E---Y-YP--EG-----WS-------Q-----Q---DYYSG----------LPK---LQRVWL---------------------------------------DQIYAM--QRE--DDSEWR-NELSDDIARWIL-----RS-----Y---E-K---------VIND-A-F--------------KL----GTGEI--------NEVKQ-R-----------VE-------------------K------SLQQAKEFF-------
WP_111891324 --------------------------------------------------------------------------------MI--ENID--A-FLN-ER------K----ELW--------------------------------------------------------------------------------L--K-------DR------L-------K------K--A---------EN-----ET-------------------------------------------AI--------------------------------------A-ELQ-QQ------------------AN--------DK------
FD---LSEW-----LP---DA-----AKRV-------T----QLS-M----------VSHPSKFSHPS---A-------------R----T--S-----SV-----I-----A---Q----------AK---QAN--D--GY-------LRS--G--NV------D----------Y--P-LD-V-FG-----N---A---A---M----DVFKFL-S------L---S-LT--E-------DQTV----------------LDGFE-QDH-Q---DLKQFI---E----Q---AS-----LDFKTLKQAFLVI---K---HG--EN-V-NKTDH-LIKQVYFPI--
-------------KQAE---YHLLSILTPSGLMTQLKQK-ID---A--I---R--F-S--------------EQ-TKQAK-E-----KR-------------K--KNEHD--AE-G-YAD-----IYD-LTVTAYGG-T----QPQ--------NVSVLN--SQNA----------GRAYLLSSC-P----PKFE-KR---MI-RLPKTD--FFAQCL-------------Y-R--KNY--QESFMQLHKFM----------QL-----DLN-----N--------------------------ID---IR---TA---IY------
NIIQFVTDQILLQALRTR-EY-------A-T-E--------G-----WS-----NQD------Y----Y-----------AN-----LPK---VQRIWL---------------------------------------DKIHQK--ERE--ADNDWC-DELSKEIARWIL-----RS-----Y---E-K---------IISD-A-Y--------------TL----GTSEL--------LDVKQ-H----------VE-------------------K------SLQQAKEFF-------
WP_004998567 --------------------------------------------------------------------------------MT--ENIH--A-FLN-ER------K----ELW--------------------------------------------------------------------------------L--K-------D-------K-------A---------ED-----ET-------------------------------------------VI--------------------------------------L-ELN-QQ------------------AD--------DK------
FS---LNEW-----LP---DA-----AKRV-------S----QLS-M----------VSHPSKFSHPS---A-------------K----T--S-----SN-----KKS-D---GY-------LRS--G--NV------D----------Y--S-LD-V-FG-----N---A---A---M----DVFKFL-S------L---P-LT--D-------QSTV----------------LDGFE-QQN-E---DLKKLL---V----Q---NK-----LDFEQLSFEFLKI--K---TT--DS-S-VKTDH-LVKQVYFPI--
-------------AEGE---YHLLSILTPSGLITKLKQT-ID---Q--I---R--F-S--------------EE-TKQAK-E-----NR-------------K--KNELN--EI-G-YAD-----IYD-LTVTAYGG-T----QPQ--------NVSVLN--SQNA----------GRAYLLSSC-P----PVIE-KR---AI-RLPKTD--FFAQCL-------------Y-R--RNY--QESFVQLHKFM----------QL-----DL-----N--------------------------ID---IR---TA---IR------
NIIQFVIDQVLLQALKTR---Q-------Y-A-ES--G-----WS-------N-----Q---DY--Y-----VG------LPK---LQRIWL---------------------------------------QIY--------KE-ER--E-E--NTE-WR-DELSREIARWIL-----RS-----Y---E-K---------KVISDA-Y-I--------------GTGEL--------LDVKQRIE-----------KS-L---------------------Q------QIKEFF-------
WP_005003488 --------------------------------------------------------------------------------MT--ESIH--T-FLN-ER------K----ELW--------------------------------------------------------------------------------L--K-------DH-------K------A---------ET-----EA-------------------------------------------VI--------------------------------------S-ELN-QQ------------------AD--------DK------
FS---LNEW-----LP---DA-----AKRV-------S----QLS-M----------VSHPSKFSHPS---A-------------K----T--S-----SN-----KKS-D---GY-------LRS--G--NV------D----------Y--S-LD-V-FG-----N---A---A---M----DVFKFL-S------L---P-LT--D-------QSTV----------------LDGFE-QQN-E---DLKKLL---V----Q---NK-----LDFEQLSFEFLKI--K---TT--DF-S-VKTDH-LVKQVYFPI--
-------------AEGE---YHLLSILTPSGLITKLKQT-ID---Q--I---R--F-S--------------EE-TKQAK-E-----NR-------------K--KNELN--EI-G-YAD-----IYD-LTVTAYGG-T----QPQ--------NVSVLN--SQNA----------GRAYLLSSC-P----PVIE-KR---AI-RLPKTD--FFAQCL-------------Y-R--RNY--QESFVQLHKFM----------Q-L---DL-----N--------------------------ID---IR---TA---IR------
NIIQFVIDQVLLQALKTR---Y-A-ES--G-----WS-------N-----Q---DY--Y-----VG------LPK---LQRIWL---------------------------------------KE-ER--E-E--NSE-WR-NELSDEIARWIL-----RS-----Y---E-K---------KVISDA-Y-I--------------GTGEL--------LDVKQRIE-----------KS-L---------------------Q------QIKEFF-------
WP_044435845 --------------------------------------------------------------------------------MT--ESIH--T-FLN-ER------K----ELW--------------------------------------------------------------------------------L--K-------DRLKKAETETV------I-SE-------LN----------------------------------------------------------------------------------------------QQ------------------ADDK------
FS---LNEW-----LP---DA-----AKRV-------S----QLS-M----------VSHPSKFSHPS---A-------------K----T--S-----SN-----KKN-D---GY-------LRS--G--NV------D----------Y--S-LD-V-FG-----N---A---A---M----DVFKFL-S------L---P-LT--D-------QSTV----------------LDGFE-QKN-E---DLKKLL---V----Q---NK-----LDFEQLSFEFLKIKTT--DS-S-----VKTDHLVKQVYFPI--
-------------AEGE---YHLLSILTPSGLITKLKQT-ID---Q--I---R--F-S--------------EE-TKQAK-E-----NR-------------K--KNELN--EI-G-YAD-----IYD-LTVTAYGG-T----QPQ--------NVSVLN--SQNA----------GRAYLLSSC-P----PVIE-KR---AI-RLPKTD--FFAQCL-------------Y-R--RNY--QESFVQLHKFM----------Q-LD-----L-----N--------------------------ID---IR---TA---IR------
NIIQFVIDQVLLQALKTR-E-------Y-A-ES--G-----WS-------N-----Q---DYYG----------LPK---LQRIWL---------------------------------------QIY--------KE-ER--E-E--NAE-WR-DELSREIARWIL-----SS----Y---E-K---------KVISDA-Y-I--------------GTGELLDVKQRIE-----------KSL-----------------------Q------QIKEFF-------
WP_049175165 --------------------------------------------------------------------------------MT--ESIH--T-FLN-ER------K----EFW--------------------------------------------------------------------------------L--K-------KDRLKKAET-------EA-------------------------------------------VI--------------------------------------SELNQQAD--------------------------DK------
FS---LNEW-----LP---DA-----AKRV-------S----QLS-M----------VSHPSKFSHPS---A-------------K----T--S-----SN-----KRN-D---GY-------LRS--G--NV------D----------Y--S-LD-V-FG-----N---A---A---M----DVFKFL-S------L---P-LT--D-------QSTV----------------LDGFE-QKN-E---DLKKLL---V----Q---NK-----LDFEQLSFEFLKI--K---TT--DS-S-SVKTDHLVKQVYFPI--
-------------AEGE---YHLLSILTPSGLITKLKQA-ID---Q--I---R--F-S--------------EE-TKQAK-E-----NR-------------K--KNELN--EI-G-YAD-----IYD-LTVTAYGG-T----QPQ--------NVSVLN--SQNA----------GRAYLLSSC-P----PVLE-KR---AI-RLPKTD--FFAQCL-------------Y-Q--RNY--QESFIQLHKFM----------QL-----D-L-----N--------------------------ID---IR---TA---IR------
NIIQFVIDQVLLQALKTR-E-----Y-A-E-----S--G-----WS-------N-----Q---DY--Y-----VG------LPK---LQRIWL---------------------------------------DQIYKE--ERE--ENSEWR-DELSREIARWIL-----RS-----Y---E-K---------KVISD-A-Y------------IL----GTGEL--------LDVKQ------------------------Q------QIKEFF-------
WP_023271486 --------------------------------------------------------------------------------MA--ESIH--A-FLN-ER------K----ELW--------------------------------------------------------------------------------L--K-------D-------K--------ES-----DV-------------------------------------------ET--------------------------------------S-TLQ-HQ------------------AN--------DK------
-FS---LNEW-----LP---DA-----AKRV-------T----QLA-M----------VSHPSKFSHPS---A-------------K----T--S-----SS-----I-----A---K----------AK---HCN--D--GY-------LRS--G--NV------D----------Y--S-LD-V-FG-----N---A---A---M----DVFKFL-S------L---P-LT--E-------KMTV----------------LDGFE-QDN-Q---ELRALI---T----H---AD-----LNFEALSSEFLKI--K---AN--DH-S-VKTDH-LVKQVYFPI--
-------------QQSE---YHLLSILTPSGLITKLKQA-ID---A--M---R--F-S--------------EE-TKKSK-E-----LR-------------K--KNEHD--EI-G-YTD-----IFD-LTITAYGG-T----QPQ--------NVSVLN--SQNA----------GRAYLLSSR-P----PVLE-K---RTIRLPKTD-FFMQCLY----------------------L-----KNY--HDSFIQLHKFI----------QLDL-----N--------------------------N----TD---IR---NA---VK------
NIIQFVIDQVILQALKTR-------Q-YAIE--G-----WT-------N-----Q---DY--------YY-AN------LPK---LQRIWL---------------------------------------DKAYQN--ERE--ENGDWR-DELSREIARWIL-----RS-----Y---E-K---------VISD-------------------ML----GTVEL--------AEVKQ----------------------------------SLQQAKEFF-------
WP_004801339 --------------------------------------------------------------------------------MT--ESIH--T-FLN-ER------K----ELW--------------------------------------------------------------------------------L--K-------D-RLK--KAES-------ED-----------------------------------------------------------------------------------------S-TLQ-HQ------------------AN--------DK------
FS---LNEW-----LP---DA-----AKRV-------T----QLA-M----------VSHPSKFSHPS---A-------------K----T--S-----SL-----I-----A---K----------AK---HCN--D--GY-------LRS--G--NV------E----------Y--S-LD-V-FG-----N---AA---A---M----DVFKFL-S------L---P-LT--E-------KMTV----------------LDGFE-QDN-Q---ELRALI---T----H---AD-----LNFEALSSEFLKI--KA---AD--HS----VKTDHLVKQVYFPI--
-------------NQSE---YHLLSILTPSGLITKLKQV-ID---A--M---R--F-S--------------EE-TKKSK-E-----LR-------------K--KNEHD--EI-G-YTD-----IFD-LTITAYGG-T----QPQ--------NVSVLN--SQNA----------GRAYLLSSR-P----PVLE-K---RAIRLPKTD-FFMQCL-----------------------------Y-Q--KNY--HDSFIQLHKFI----------Q---LDL-----N--------------------------TD---IR---NA---IT------
NIIQFVIDQVILQALKTR-------Q-YAIE--G-----WS-------N-----Q---DY--------YYAN------LPK---LQRIWL---------------------------------------DKTYQI--ERE--ENSDWR-DELSREIARWIL-----RS-----Y---E-K---------VISD-------------------ML----GTREL--------VEVKQ----------------------------------SLQQAKEFF-------
WP_004798546 --------------------------------------------------------------------------------MT--ESIQ--N-FLN-ER------K----ELW--------------------------------------------------------------------------------L--K-------D-------K-------A---------ED-----ET-------------------------------------------VI--------------------------------------S-ELQ-QQ------------------AD--------DK------
FS---LKEW-----LP---DA-----AKRV-------T----QLS-M----------VSHPSKFSHPS---A-------------K----T--S-----SV-----I-----A---K----------AK---QAN--D--GY-------LRS--G--NV------D----------Y--P-LD-V-FG-----N---A---A---M----DVFKFL-S------L---P-LT--E-------KLTV----------------LDGFE-RQE-E---DLKSLL---V----G---AD-----LDFETLGTEFLKI--K---AT--DT-S-VKTDH-LVKQVYFPI--
-------------EQAE---YHLLSVLTPSGLITKLKQ-ID---T--I---R--F-S--------------QE-TKQAK-E-----LR-------------K--KNEHH--E-G-YAD-----IFD-LTITAYGG-T----QPQ--------NVSVLN--SQNA----------GRAYLLSSC-P----PMLE-K---RAIRLPKTD-FFAQCL-----------------------------Y-R--KNY--QESFIQLHKFM----------QLDL-----N--------------------------N----DD---IR---TA---IR------
NIIQFVIDQILVLASKTR-----------EYST-EG-----WS-------N-----Q---D-------YY-AS------LPK---LQRIWL---------------------------------------DKIYQT--ERD--EDSDWR-DELSREIARWIL-----RS-----Y---E-K---------VISD-------------------KL----GTGEL--------LDVKQ-R----------VE-------------------K------SLQKAKEFF-------
WP_004681365 --------------------------------------------------------------------------------MT--ESIQ--N-FLN-ER------K----ELW--------------------------------------------------------------------------------L--K-------DR------L-------K------K--A---------EN-----DS-------------------------------------------VI--------------------------------------S-ELQ-QQ------------------AD--------DK------
FS---LKEW-----LP---DA-----AKRV-------T----QLS-M----------VSHPSKFSHPS---A-------------K----T--S-----SV-----I-----A---K----------AK---QIN--D--GY-------LRS--G--NV------D----------Y--P-LD-V-FG-----N---A---A---M----DVFKFL-S------L---P-LT--E-------KLTV----------------LDGFE-RQE-E---DLKSLL---V----G---AD-----LDFETLGTEFLKI--K---AT--DT-S-VKTDH-LVKQVYFPI--
-------------EQAE---YHLLSVLTPSGLITKLKQS-ID---T--I---R--F-S--------------TE-TKQAK-E-----LR-------------R--KNEHH--EI-G-YAD-----IFD-LTITAYGG-T----QPQ--------NVSVLN--SQNA----------GRAYLLSSC-P----PMLE-K---RAIRLPKTD-FFAQCL-----------------------------Y-R--KNY--QESFIQLHKFM----------Q---LDL-----N--------------------------ND---IR---TA---IR------
NIIQFVIDQILVLASKTR-----------E-YSVE--G-----WS-------N-----Q---D-------YYASLPK---LQRIWL---------------------------------------DKIYQT--ERD--EDSDWR-NDLSKEIARWIL-----RS-----Y---E-K---------VISD-------------------KL----GTGEL--------LDVKQ-R----------VE-------------------K------SLQQAKEFF-------
WP_050040614 --------------------------------------------------------------------------------MA--ESIY--A-FLN-ER------K----ELW--------------------------------------------------------------------------------L--K-------DR------L-------K------K--A---------EN-----ET-------------------------------------------VI--------------------------------------A-ELQ-QQ------------------AN--------DK------
FS---LNEW-----LP---DA-----AKRV-------T----QLS-M----------VSHPSKFSHPS---A-------------K----T--S-----SV-----I-----A---K----------VE---QSN--D--GY-------LRS--G--NV------D----------Y--S-LD-V-FG-----N---AA---A---M----DVFKFL-S------L---P-LN--E-------KFTV----------------LDGFE-QQD-H---ALKTLI---T----H---AK-----LNFDTLSTEFLKI--K---VT--DH-S-VKTDH-LVKQVYFPV--
-------------REAQ---YHLLSILTPSGLITRLKQS-ID---T--I---R--F-S--------------ED-TKQAK-E-----SR-------------K--KNEHH--EM-G-YSD-----IFD-LTVTGYGG-T----QPQ--------NVSVLN--SQNA----------GRAYLLPSC-P----PVLE-K---RAIRLPKTD-FFLQCL-----------------------------Y-R--KNY--QESFIQLHKFM----------QLDLN-----N--------------------------ID---IR---NA---IR------
NIIQFVIDQILLQALKTR-----------EYAAE--G-----WS-------NQDY----Y----AS------LPK---IQRIWL---------------------------------------DKVYQT--KRE--ENDDWR-DELSREIARWIL-----L-NYEKVINDA-----------Y--------------TL----GTGEL--------LDVKQ-R----------VE-------------------N------SLQQAKEFF-------
WP_046739738 --------------------------------------------------------------------------------MA--ESIH--A-FLN-ER------K----ELW--------------------------------------------------------------------------------L--K-------DRIK-----K--A---------EN-----EA-------------------------------------------VI--------------------------------------A-ELQ-QQ------------------AN--------DK------
FS---LNEW-----LP---DA-----AKRV-------T----QLS-M----------VSHPSKFSHPS---A-------------K----T--S-----SV-----I-----A---K----------VE---QSN--D--GY-------LRS--G--NV------D----------Y--S-LD-V-FG-----N---AA---A---M----DVFKFL-S------L---P-LN--E-------KFTV----------------LDGFE-QQD-Q---ALKTLI---T----H---AK-----LNFDTLSTEFLKI--KV---TD--HS----VKTDHLVKQVYFPV--
-------------REAQ---YHLLSILTPSGLITRLKQS-ID---T--I---R--F-S--------------ED-TKQAK-E-----SR-------------K--KNEHH--EI-G-YSD-----IFG-LTVTGYGG-T----QPQ--------NVSVLN--SQNA----------GRAYLLPSC-P----PVLE-K---RAIRLPKTD-FFLQCL-----------------------------R--KNY--QESFIQLHKFM----------Q---LD-----L--------------------------N----ID---IR---NV---IR------
NIIQFVIDQILLQALKTR-----------EYAA-E--G-----WS-------NQDY----Y----VS------LPK---IQRIWL---------------------------------------DKVYQT--KRE--ENDDWR-DELSREIARWIL-----L-NYEKVIND-A-Y--------------TL----GTGEL--------LDVKQ-R----------VE-------------------N------SLQQAKEFF-------
WP_114259034 --------------------------------------------------------------------------------MA--ESIH--T-FLN-ER------K----ELW--------------------------------------------------------------------------------L--K-------D-------K-------A---------EN-----ES-------------------------------------------AI--------------------------------------A-ELQ-QQ------------------AN--------DK------
FS---LNKW-----LP---DA-----AKRV-------T----QLS-M----------VSHPSKFSHPS---A-------------K----T--S-----SV-----I-----A---K----------VE---YCN--D--GY-------LRS--G--NV------D----------Y--S-LD-V-FG-----N---AA---A---M----DVFKFL-S------L---A-LT--E-------NLTV----------------LDGFE-KQD-Q---ELKKLI---A----H---AE-----LDFDNLSAEFLKI--K---AT--DG-S-IKTDH-LVKQVYFPV--
---GE-------------AQ---YHLLSILTPSGLITRLKQS-VD---V--M---R--F-S--------------EE-TKQAK-E-----SR-------------K--KNEHH--EV-G-YSD-----IFD-LTVTGYGG-T----QPQ--------NVSVLN--SQNA----------GRAYLLPSC-P----PVLE-K---RTIRLPKAD-FFAQCLY-----------------------------R--KNY--QDSFIQLHKFM----------QLDLN-----N--------------------------ID---IR---NA---IK------
NIIQFVIDQILLQALKTR-----------EYAVE--------G-----WS-----NQDYY--------AS------LPK---IQRIWL---------------------------------------DKVHQK--ERE--ENSDWR-DELSREVARWIL-----RSYEKVI---S---------DAY--------------TL----GTGEL--------LDVKQ-R----------VE-------------------N------SLQKAKEFF-------
WP_032009099 --------------------------------------------------------------------------------MA--ESIH--T-FLN-ER------K----ELW--------------------------------------------------------------------------------L--K-------DR------L-------K------K--A---------EN-----ES-------------------------------------------AI--------------------------------------A-ELQ-QQ------------------AN--------DK------
FS---LNEW-----LP---EA-----AKRV-------T----QLS-M----------VSHPSKFSHPS---A-------------K----T--S-----SV-----I-----A---K----------VE---YCN--D--GY-------LRS--G--NV------D----------Y--S-LD-V-FG-----N---AA---A---M----DVFKFL-S------L---A-LT--E-------NLTV----------------LDGFE-KQD-Q---ELKKLI---A----H---AE-----LDFDNLSAEFLKI--K---AT--DG-S-IKTDH-LVKQVYFPV--
---GE-------------AQ---YHLLSILTPSGLITRLKQS-VD---V--M---R--F-S--------------EE-TKQAK-E-----SR-------------K--KNEHH--EV-G-YSD-----IFD-LTVTGYGG-T----QPQ--------NVSVLN--SQNA----------GRAYLLPSC-P----PVLE-K---RTIRLPKTD-FFAQCLY-----------------------------R--KNY--QDSFIQLHKFM----------QLDLN-----N--------------------------ID---IR---NA---IK------
NIIQFVIDQILLQALKTR-----------EYAVE--------WS-----NQDYY--------AS------LPK---IQRIWL---------------------------------------DKVHQK--ERE--ENSDWR-DELSREVARWIL-----RSYEKVI---S---------DAY--------------TL----GTGEL--------LDVKQ-R----------VE-------------------N------SLQKAKEFF-------
WP_031380756 --------------------------------------------------------------------------------MA--ESIH--T-FLN-ER------K----ELW--------------------------------------------------------------------------------L--K-------DR------L-------K------K--A---------EN-----ES-------------------------------------------AI--------------------------------------A-ELQ-QQ------------------AN--------YK------
FS---LNEW-----LP---DA-----AKRV-------T----QLS-M----------VSHPSKFSHPS---A-------------K----T--S-----SV-----I-----A---K----------VE---YCN--D--GY-------LRS--G--NV------D----------Y--S-LD-V-FG-----N---AA---A---M----DVFKFL-S------L---A-LT--E-------NLTV----------------LDGFE-KQD-Q---ELKKLI---A----H---AE-----LDFDNLSAEFLKI--K---AT--DD-S-IKTDH-LVKQVYFPV--
---GE-------------AQ---YHLLSILTPSGLITRLKQS-VD---V--M---R--F-S--------------EE-TKQAK-E-----SH-------------K--KNEHH--EV-G-YSD-----IFD-LTVTGYGG-T----QPQ--------NVSVLN--SQNA----------GRAYLLPSC-P----PVLE-K---RTIRLPKTD-FFAQCLY-----------------------------R--KNY--QDSFIQLHKFM----------QLDLN-----N--------------------------ID---IR---NA---IK------
NIIQFVIDQILLQALKTR-----------EYAVE--------G-----WS-----NQDYY--------AS------LPK---IQRIWL---------------------------------------DKVHQK--ERE--ENSDWR-DELSREVARWIL-----RSYEKVI---S---------DAY--------------TL----GTGEL--------LDVKQ-R----------VE-------------------N------SLQKAKEFF-------
WP_059212431 --------------------------------------------------------------------------------MA--ESIH--T-FLN-ER------K----ELW--------------------------------------------------------------------------------L--K-------DR------L-------K------K--A---------EN-----ES-------------------------------------------AI--------------------------------------A-ELQ-QQ------------------AN--------YI------
FS---LNEW-----LP---DA-----AKRV-------T----QLS-M----------VSHPSKFSHPS---A-------------K----T--S-----SV-----I-----A---K----------VE---YCN--D--GY-------LRS--G--NV------D----------Y--S-LD-V-FG-----N---AA---A---M----DVFKFL-S------L---A-LT--E-------NLTV----------------LDGFE-KQD-Q---ELKKLI---A----H---AE-----LDFDNLSAEFLKI--K---AT--DD-S-IKTDH-LVKQVYFPV--
---GE-------------AQ---YHLLSILTPSGLITRLKQS-VD---V--M---R--F-S--------------EE-TKQAK-E-----SR-------------K--KNEHH--EV-G-YSD-----IFD-LTVTGYGG-T----QPQ--------NVSVLN--SQNA----------GRAYLLPSC-P----PVLE-K---RTIRLPKTD-FFAQCLY-----------------------------R--KNY--QDSFIQLHKFM----------QLDLN-----N--------------------------ID---IR---NA---IK------
NIIQFVIDQILLQALKTR-----------EYAVE--------G-----WS-----NQDYY--------AS------LPK---IQRIWL---------------------------------------DKVHQK--ERE--ENSDWR-DELSREVARWIL-----RSYEKVI---S---------DAY--------------TL----GTGEL--------LDVKQ-R----------VE-------------------N------SLQKAKEFF-------
WP_106078368 --------------------------------------------------------------------------------MA--ESIH--T-FLN-ER------K----ELW--------------------------------------------------------------------------------L--K-------DR------L-------K------K--A---------EN-----ES-------------------------------------------AI--------------------------------------A-ELQ-QQ------------------AN--------YK------
FS---LNEW-----LP---DA-----AKRV-------T----QLS-M----------VSHPSKFSHPS---A-------------K----T--S-----SV-----I-----A---K----------VE---YCN--D--GY-------LRS--G--NV------D----------Y--S-LD-V-FG-----N---AA---A---M----DVFKFL-S------L---A-LT--E-------NLTV----------------LDGFE-KQD-Q---ELKKLI---A----H---AE-----LDFDNLSAEFLKI--K---AT--DD-S-IKTDH-LVKQVYFPV--
---GE-------------AQ---YHLLSILTPSGLITRLKQS-VD---V--M---R--F-S--------------EE-NKQAK-E-----SR-------------K--KNEHH--EV-G-YSD-----IFD-LTVTGYGG-T----QPQ--------NVSVLN--SQNA----------GRAYLLPSC-P----PVLE-K---RTIRLPKTD-FFAQCLY-----------------------------R--KNY--QDSFIQLHKFM----------QLDLN-----N--------------------------ID---IR---NA---IK------
NIIQFVIDQILLQALKTR-----------EYAVE--------G-----WS-----NQDYY--------AS------LPK---IQRIWL---------------------------------------DKVHQK--ERE--ENSDWR-DELSREVARWIL-----RSYEKVI---S---------DAY--------------TL----GTGEL--------LDVKQ-R----------VE-------------------N------SLQKAKEFF-------
WP_000841022 --------------------------------------------------------------------------------MA--ESIH--T-FLN-ER------K----ELW--------------------------------------------------------------------------------L--K-------D-RI-----K------K--A---------EN-----ES-------------------------------------------AI--------------------------------------A-ELQ-QQ------------------AN--------DK------
FS---LNEW-----LP---DA-----AKRV-------T----QLS-M----------VSHPSKFSHPS---A-------------K----T--S-----SV-----I-----A---K----------VE---YCN--D--GY-------LRS--G--NV------D----------Y--S-LD-V-FG-----N---AA---A---M----DVFKFL-S------L---A-LT--E-------NLTV----------------LDGFE-KQD-Q---ELKKLI---A----H---AE-----LDFDNLSAEFLKI--K---AT--DD-S-IKTDH-LVKQVYFPV--
---GE-------------AQ---YHLLSILTPSGLITRLKQS-VD---V--M---R--F-S--------------EE-TKQAK-E-----SR-------------K--KNEHH--EV-G-YSD-----IFD-LTVTGYGG-T----QPQ--------NVSVLN--SQNA----------GRAYLLPSC-P----PVLE-K---RTIRLPKTD-FFAQCLY-----------------------------R--KNY--QDSFIQLHKFM----------Q---LDL-----N--------------------------ID---IR---NA---IK------
NIIQFVIDQILLQALKTR-----------E-YAVE--------G-----WS-----NQDYY--------YYAS------LPK---IQRIWL---------------------------------------DKVHQK--ERE--ENSDWR-DELSREVARWIL-----RSYEKVI---S---------DAY--------------TL----GTGEL--------LDVKQ-R----------VE-------------------N------SLQKAKEFF-------
WP_057246220 --------------------------------------------------------------------------------MA--ESIH--T-FLN-ER------K----ELW--------------------------------------------------------------------------------K-DRIK-----K--A---------EN-----ES-------------------------------------------AI--------------------------------------A-ELQ-QQ------------------AN--------YK------
FS---LNEW-----LP---DA-----AKRV-------T----QLS-M----------VSHPSKFSHPS---A--------------KTS-------SV-----I-----A---K----------VE---YCN--D--GY-------LRS--G--NV------D----------YSLD-V-FG-----N---AAA---M----DVFKFL-S------L---A-LT--E-------NLTV----------------LDGFE-KQD-Q---ELKKLI---A----H---AE-----LDFDNLSAEFLKI--KA---TD--DS----IKTDHLVKQVYFPV--
---GE-------------AQ---YHLLSILTPSGLITRLKQS-VD---V--M---R--F-S--------------EE-TKQAK-E-----SR-------------K--KNEHH--EV-G-YSD-----IFD-LTVTGYGG-T----QPQ--------NVSVLN--SQNA----------GRAYLLPSC-P----PVLE-K---RTIRLPKAD-FFAQCL-----------------------------FF--AQC----------------------------------------------------LY---------RK-----NYQDSFIQ---
---------------LHK-----------------------------------------------------------FMQ----------------------------L-------------------------DL---NNIDI-----RN--------------------A----------------------------------------------------------------------------------------------------------------------------------------------------------------------------------------
WP_000841020 --------------------------------------------------------------------------------MA--ESIH--T-FLN-ER------K----ELW--------------------------------------------------------------------------------L--K-------DR------L-------K------K--A---------EN-----ES-------------------------------------------AI--------------------------------------A-ELQ-QQ------------------AN--------DK------
FS---LNEW-----LP---DA-----AKRV-------T----QLS-M----------VSHPSKFSHPS---A-------------K----T--S-----SV-----I-----A---K----------VE---YCN--D--GY-------LRS--G--NV------D----------Y--S-LD-V-FG-----N---AA---A---M----DVFKFL-S------L---A-LT--E-------NLTV----------------LDGFE-KQN-Q---ELKKLI---A----H---AE-----LDFDNLSAEFLKI--K---AT--DD-S-IKTDH-LVKQVYFPV--
---GE-------------AQ---YHLLSILTPSGLITRLKQS-VD---A--M---R--F-S--------------EE-TKQAK-E-----SR-------------K--KNEHH--EV-G-YSD-----IFD-LTVTGYGG-T----QPQ--------NVSVLN--SQNA----------GRAYLLPSC-P----PVLE-K---RTIRLPKAD-FFAQCLY-----------------------------R--KNY--HDSFIQLHKFM----------Q---LDL-----N--------------------------ID---IR---NA---IK------
NIIQFVIDQILLQALKTR-----------EYAVE--------WS-----NQDYY--------AS------LPK---IQRIWL---------------------------------------DKVHQK--ERE--ENSDWR-DELSREVARWIL-----RSYEKVI---S---------DAY--------------TL----GTGEL--------LDVKQWVE----------------------------N------SLQKAKEFF-------
WP_000841019 --------------------------------------------------------------------------------MA--ESIH--T-FLN-ER------K----ELW--------------------------------------------------------------------------------L--K-------D-RI-----K------K--A---------EN-----ES-------------------------------------------AI--------------------------------------A-ELQ-QQ------------------AN--------DK------
FS---LNEW-----LP---DA-----AKRV-------T----QLS-M----------VSHPSKFSHPS---A-------------K----T--S-----SV-----I-----A---K----------VE---YCN--D--GY-------LRS--G--NV------D----------Y--S-LD-V-FG-----N---AA---A---M----DVFKFL-S------L---A-LT--E-------NLTV----------------LDGFE-KQN-Q---ELKKLI---A----H---AE-----LDFDNLSAEFLKI--K---AT--DD-S-IKTDH-LVKQVYFPV--
---GE-------------AQ---YHLLSILTPSGLITRLKQS-VD---V--M---R--F-S--------------EE-TKQAK-E-----SR-------------K--KNEHH--EV-G-YSD-----IFD-LTVTGYGG-T----QPQ--------NVSVLN--SQNA----------GRAYLLPSC-P----PVLE-K---RTIRLPKAD-FFAQCLY-----------------------------R--KNY--HDSFIQLHKFM----------Q---LDL-----N--------------------------ID---IR---NA---IK------
NIIQFVIDQILLQALKTR-----------EYAVE--------WS-----NQDYY--------AS------LPK---IQRIWL---------------------------------------DKVHQK--ERE--ENSDWR-DELSREVARWIL-----RSYEKVI---S---------DAY--------------TL----GTGEL--------LDVKQWVE----------------------------N------SLQKAKEFF-------
WP_031965191 --------------------------------------------------------------------------------MA--ESIH--T-FLN-ER------K----ELW--------------------------------------------------------------------------------L--K-------D-RI-----K------K--A---------EN-----ES-------------------------------------------AI--------------------------------------A-ELQ-QQ------------------AN--------DK------
FS---LNEW-----LP---DA-----AKRV-------T----QLS-M----------VSHPSKFSHPS---A-------------K----T--S-----SV-----I-----A---K----------VE---YCN--D--GY-------LRS--G--NV------D----------Y--S-LD-V-FG-----N---AA---A---M----DIFKFL-S------L---A-LT--E-------NLTV----------------LDGFE-KQN-Q---ELKKLI---A----H---AE-----LDFDNLSAEFLKI--K---AT--DD-S-IKTDH-LVKQVYFPV--
---GE-------------AQ---YHLLSILTPSGLITRLKQS-VD---A--M---R--F-S--------------EE-TKQAK-E-----SR-------------K--KNEHH--EV-G-YSD-----IFD-LTVTGYGG-T----QPQ--------NVSVLN--SQNA----------GRAYLLPSC-P----PVLE-K---RTIRLPKAD-FFAQCLY-----------------------------R--KNY--HDSFIQLHKFM----------Q---LDL-----N--------------------------ID---IR---NA---IK------
NIIQFVIDQILLQALKTR-----------EYAVE--------WS-----NQDYY--------AS------LPK---IQRIWL---------------------------------------DKVHQK--ERE--ENSDWR-DELSREVARWIL-----RSYEKVI---S---------DAY--------------TL----GTGEL--------LDVKQWVE----------------------------N------SLQKAKEFF-------
WP_004739558 --------------------------------------------------------------------------------MA--ESIH--T-FLN-ER------K----ELW--------------------------------------------------------------------------------K-DRIK-----K--A---------EN-----ES-------------------------------------------AI--------------------------------------A-ELQ-QQ------------------AN--------DK------
FS---LNEW-----LP---DA-----AKRV-------T----QLS-M----------VSHPSKFSHPS---A--------------KTS-------SV-----I-----A---K----------VE---YCN--D--GY-------LRS--G--NV------D----------YSLD-V-FG-----N---AAA---M----DVFKFL-S------L---A-LT--E-------NLTV----------------LDGFE-KQD-Q---ELKKLI---A----H---AE-----LDFDNLSAEFLKI--KA---TD--DS----IKTDHLVKQVYFPV--
---GE-------------AQ---YHLLSILTPSGLITRLKQS-VD---V--M---R--F-S--------------EE-TKQAK-E-----SR-------------K--KNEHH--EV-G-YSD-----IFD-LTVTGYGG-T----QPQ--------NVSVLN--SQNA----------GRAYLLPSC-P----PVLE-K---RTIRLPKAD-FFAQCLY-----------------------------R--KNY--HDSFIQLHKFM----------Q---LDL-----N--------------------------ID---IR---NA---IK------
NIIQFVIDQILLQALKTR-----------EYAVE--------G-----WS-----NQDYY--------AS------LPK---IQRIWL---------------------------------------DKVHQK--ERE--ENSDWR-DELSREVARWIL-----RSYEKVI---S---------DAY--------------TL----GTGEL--------LDVKQWVE----------------------------N------SLQKAKEFF-------
WP_057054378 --------------------------------------------------------------------------------MA--ESIH--T-FLN-ER------K----ELW--------------------------------------------------------------------------------L--K-------DR------L-------K------K--A---------EN-----ES-------------------------------------------AI--------------------------------------A-ELQ-QQ------------------AN--------DK------
FS---LNEW-----LP---DA-----AKRV-------T----QLS-M----------VSHPSKFSHPS---A-------------K----T--S-----SV-----I-----A---K----------VE---YCN--D--GY-------LRS--G--NV------D----------Y--S-LD-V-FG-----N---AA---A---M----DVFKFL-S------L---A-LT--E-------NLTV----------------LDGFE-KQD-Q---ELKKLI---A----H---AE-----LDFDNLSAEFLKI--K---AT--DD-S-IKTDH-LVKQVYFPV--
---GE-------------AQ---YHLLSILTPSGLITRLKQS-VD---V--M---R--F-S--------------EE-TKQAK-E-----SR-------------K--KNEHH--EV-G-YSD-----IFD-LTVTGYGG-T----QPQ--------NVSVLN--SQNA----------GRAYLLPSC-P----PVLE-K---RTIRLPKAD-FFAQCLY-----------------------------R--KNY--QDSFIQLHKFM----------QLDLN-----N--------------------------ID---IR---NA---IK------
NIIQFVIDQILLQALKTR-----------EYAVE--------WS-----NQDYY--------AS------LPK---IQRIWL---------------------------------------DKVHQK--ERE--ENSDWR-DELIREVARWIL-----RSYEKVI---S---------DAY--------------TL----GTGEL--------LDVKQ-R----------VE-------------------N------SLQKAKEFF-------
WP_000841018 --------------------------------------------------------------------------------MA--ESIH--T-FLN-ER------K----ELW--------------------------------------------------------------------------------L--K-------D-RI-----K------K--A---------EN-----ES-------------------------------------------AI--------------------------------------A-ELQ-QQ------------------AN--------DK------
FS---LNEW-----LP---DA-----AKRV-------T----QLS-M----------VSHPSKFSHPS---A-------------K----T--S-----SV-----I-----A---K----------VE---YCN--D--GY-------LRS--G--NV------D----------Y--S-LD-V-FG-----N---AA---A---M----DVFKFL-S------L---A-LT--E-------NLTV----------------LDGFE-KQN-Q---ELKKLI---A----H---AE-----LDFDNLSAEFLKI--K---AT--DD-S-IKTDH-LVKQVYFPV--
---GE-------------AQ---YHLLSILTPSGLITRLKQS-VD---A--M---R--F-S--------------EE-TKQAK-E-----SR-------------K--KNEHH--EV-G-YSD-----IFD-LTVTGYGG-T----QPQ--------NVSVLN--SQNA----------GRAYLLPSC-P----PVLE-K---RTIRLPKAD-FFAQCLY-----------------------------R--KNY--HDSFIQLHKFMQLDLN-----N--------------------------ID---IR---NA---IK------
NIIQFVIDQILLQALKTR-----------EYAVE--------NQDYYAS------LPK---IQRIWL---------------------------------------DKVHQK--ERE--ENSDWR-DELSREVARWIL-----RSYEKVI---S-----DA-Y--------------TL----GTGEL--------LDVKQWVE----------------------------N------SLQKAKEFF-------
WP_119957144 --------------------------------------------------------------------------------L-WLK-DR-------------------------------------------------------------------K-------K--A---------EN-----ES-------------------------------------------AI--------------------------------------A-ELQ-QQ------------------AN--------DK------FS---
LNEW-----LP---DA-----AKRV-------T----QLS-M----------VSHPSKFSHPS---A-------------K----T--S-----SV-----I-----A---K----------VE---YCN--D--GY-------LRS--G--NV------D----------Y--S-LD-V-FG-----N---AA---A---M----DVFKFL-S------L---A-LT--E-------NLTV----------------LDGFE-KQD-Q---ELKKLI---A----H---AE-----LDFDNLSSEFLKI-KA---TD--DS----IKTDHLVKQVYFPV--
---GE-------------AQ---YHLLSILTPSGLITRLKQS-VD---A--M---R--F-S--------------EE-TKQAK-E-----SR-------------K--KNEHH--EV-G-YSD-----IFD-LTVTGYGG-T----QPQ--------NVSVLN--SQNA----------GRAYLLPSC-P----PVLE-K---RTIRLPKAD-FFAQCLY-----------------------------R--KNY--QDSFIQLHKFM----------QLDL-----N--------------------------ID---IR---NA---IK------
NIIQFVIDQILLQALKTR-----------EYAVE--------G-----WS-----N-----Q---D-----Y----AS------LPK---IQRIWL---------------------------------------DKVHQK--ERE--ENSDWR-DELSREVARWIL-----RS-----Y---E-KVISDA-----------Y--------------TL----GTGEL--------LDVKQ-R----------VE-------------------N------SLQKAKEFF-------
WP_000841017 --------------------------------------------------------------------------------MA--ESIH--T-FIN-ER------K----ELW--------------------------------------------------------------------------------L--K-------D-RI-----K------K--A---------EN-----ES-------------------------------------------AI--------------------------------------A-ELQ-QQ------------------AN--------DK------
FS---LNEW-----LP---DA-----AKRV-------T----QLS-M----------VSHPSKFSHPS---A-------------K----T--S-----SV-----I-----A---K----------VE---YCN--D--GY-------LRS--G--NV------D----------Y--S-LD-V-FG-----N---AA---A---M----DVFKFL-S------L---A-LT--E-------NLTV----------------LDGFE-KQN-Q---ELKKLI---A----H---AE-----LDFDNLSAEFLKI--K---AT--DD-S-IKTDH-LVKQVYFPV--
---GE-------------AQ---YHLLSILTPSGLITRLKQS-VD---V--M---R--F-S--------------EE-TKQAK-E-----SR-------------K--KNEHH--EV-G-YSD-----IFD-LTVTGYGG-T----QPQ--------NVSVLN--SQNA----------GRAYLLPSC-P----PVLE-K---RTIRLPKAD-FFAQCLY-----------------------------R--KNY--HDSFIQLHKFM----------Q---LDL-----N--------------------------ID---IR---NA---IK------
NIIQFVIDQILLQALNTR-----------EYAVE--------WS-----NQDYY--------AS------LPK---IQRIWL---------------------------------------DKVHQK--ERE--ENSDWR-DELSREVARWIL-----RSYEKVI---S---------DAY--------------TL----GTGEL--------LDVKQWVE----------------------------N------SLQKAKEFF-------
WP_032060690 --------------------------------------------------------------------------------MA--ESIH--T-FLN-ER------K----ELW--------------------------------------------------------------------------------L--K-------D-RI-----K------K--A---------EN-----ES-------------------------------------------AI--------------------------------------A-ELQ-QQ------------------AN--------DK------
FS---LNEW-----LP---DA-----AKRV-------T----QLS-M----------VSHPSKFSHPS---A-------------K----T--S-----SV-----I-----A---K----------VE---YCN--D--GY-------LRS--G--NV------D----------Y--S-LD-V-FG-----N---AA---A---M----DVFKFL-S------L---A-LT--E-------NLTV----------------LDGFE-KQD-Q---ELKKLI---A----H---AE-----LDFDNLSAEFLKI--K---AT--DD-S-IKTDH-LVKQVYFPV--
---GE-------------AQ---YHLLSILTPSGLITRLKQS-VD---V--M---R--F-S--------------EE-TKQAK-E-----SR-------------K--KNEHH--EV-G-YSD-----IFD-LTVTGYGG-T----QPQ--------NVSVLN--SQNA----------GRAYLLPSC-P----PVLE-K---RTIRLPKAD-FFAQCLY-----------------------------R--KNY--HDSFIQLHKFM----------Q---LDL-----N--------------------------ID---IR---NA---IK------
NIIQFVIDQILLQALNTR-----------EYAVE--------WS-----NQDYY--------AS------LPK---IQRIWL---------------------------------------DKVHQK--ERE--ENSDWR-DELSREVARWIL-----RSYEKVI---S---------DAY--------------TL----GTGEL--------LDVKQWVE----------------------------N------SLQKAKEFF-------
WP_005109633 --------------------------------------------------------------------------------MA--ESIH--T-FLN-ER------K----ELW--------------------------------------------------------------------------------K-DRIK-----K--A---------EN-----ES-------------------------------------------AI--------------------------------------A-ELQ-QQ------------------AN--------DK------
FS---LNEW-----LP---DA-----AKRV-------T----QLS-M----------VSHPSKFSHPS---A--------------KTS-------SV-----I-----A---K----------VE---YCN--D--GY-------LRS--G--NV------D----------YSLD-V-FG-----N---AAA---M----DVFKFL-S------L---A-LT--E-------NLTV----------------LDGFE-KQD-Q---ELKKLI---A----H---AE-----LDFDNLSAEFLKI--KA---TD--DS----IKTDHLVKQVYFPV--
---GE-------------AQ---YHLLSILTPSGLITRLKQS-VD---V--M---R--F-S--------------EE-TKQAK-E-----SR-------------K--KNEHH--EV-G-YSD-----IFD-LTVTGYGG-T----QPQ--------NVSVLN--SQNA----------GRAYLLPSC-P----PVLE-K---RTIRLPKAD-FFAQCLY-----------------------------R--KNY--QDSFIQLHKFM----------QLDLN-----N--------------------------ID---IR---NA---IK------
NIIQFVIDQILLQALNTR-----------E---QD-YY--------AS------LPK---IQRIWL---------------------------------------DKVHQK--ERE--ENSDWR-DELSREVARWIL-----RSYEKVI---S---------DAY--------------TL----GTGEL--------LDVKQWVE----------------------------N------SLQKAKEFF-------
WP_121929531 --------------------------------------------------------------------------------MA--ESIH--T-FLN-ER------K----ELW--------------------------------------------------------------------------------L--K-------DR------L-------K------K--A---------EN-----ES-------------------------------------------VI--------------------------------------A-ELN-QQ------------------AN--------DK------
FS---LNEW-----LP---DA-----AKRV-------T----QLS-M----------VSHPSKFSHPS---A-------------K----T--S-----SV-----I-----A---K----------VE---HRN--D--GY-------LRS--G--NV------D----------Y--S-LD-V-FG-----N---AA---A---M----DVFKFL-S------L---P-LT--D-------QSTV----------------LDGFE-EKD-Q---RLKSLI---A----E---KE-----LDFSSLAEEFLKI--K---EA--DN-S-IRTDH-LVKQVYFPV--
-------------SEAE---YHLLSILTPSGLITKLKQS-ID---A--I---R--F-S--------------EE-TKQAK-E-----SR-------------K--KNEHH--EV-G-YSD-----IFD-LTVTAYGG-T----QPQ--------NVSVLN--SQNA----------GRAYLLPSC-P----PVLE-K---RTIRLPKTD-FFVQCLY-----------------------------R--KNY--QDSFIQLHKFM----------Q---LDL-----N--------------------------ID---IR---NA---IR------
NIIQFVIDQILLQAFRTR-----------EYAVE--------G-----WS-----N-----QNRY----ES------LPK---IQRIWL---------------------------------------DKIHQK--ERE--VNTDWC-DELSREIACWIV-----RS----Y---E-K---------II---S---------DAY--------------TL----GTGEL--------FDVKN-R----------VE-------------------N------SLQKAKGYF-------
WP_017395753 --------------------------------------------------------------------------------MA--ESIH--T-FLN-ER------K----ELW--------------------------------------------------------------------------------K-DRIK-----K--A---------EN-----ES-------------------------------------------VI--------------------------------------A-ELQ-QQ------------------AK--------DK------
FS---LNEW-----LP---DA-----AKRV-------T----QLS-M----------VSHPSKFSHPS---A-------------K----T--S-----SV-----I-----A---K----------VE---YCN--D--GY-------LRS--G--NV------D----------Y--S-LD-V-FG-----N---AA---A---M----DVFKFL-S------L---A-LT--E-------ELTV----------------LDGFE-END-Q---ELRKLI---A----D---AK-----LNFTLSAEFLKI--K---TD--NS-S-IRTDH-LVKQVYFPV--
-------------SEVN---YHLLSILTPSGLITKLKQS-ID---V--M---R--F-S--------------EE-TKQAK-E-----SR-------------K--KNEHH--EV-G-YSD-----IFD-LTVTGYGG-T----QPQ--------NVSVLN--SQNA----------GRAYLLSSS-P----PVLE-K---RTIRLPKTD-FFVQCLY-----------------------------R--KNY--QDSFIQLHKFM----------Q-LDL-----N--------------------------ID---IR---NA---IR------
NIIQFVIDQILLQALKTR-----------EYAVE--------G-----WS-----NQDYY--------SS------LPK---IQRIWL---------------------------------------DKVHQM--TRD--EDNDWR-DELSREIARWIL-----RSYEKVI---S---------D-A-Y--------------TL----GTGEL--------LDVKQ-R----------VE-------------------K------SLQRAKEFF-------
;
end;

#### Cas8u2

#NEXUS
begin characters;
dimensions nchar=528;
format datatype=protein missing=? gap=-;
matrix
WP_018870931 ----------------------------------------MMAERFPVDPTNPGEILACAAIT------RLAEEET------GPNDCGF--------------------------------------------------------------IQSDARWYFEAPLPHATLTR-----------------------LLETEPREEDGAVW-----------LDDQRLDW--FQH------PY--------GLNTGFKFWAGQ-----------------------------------------QSARSVLTN--
LVE---AARTGDPADWL-----THEAPT-------TGRLGVD---PQGTWN--SLGLGWSV-----NEHNR-L---KYLCRPYVELL-------AFLG------LQHFPVQGDR---------GEGFRYHL----WQ----------PAPALIARFAIAGASRYS------LQGWRTE-IAGAGSN--------RF---LKPAHPLREEP--------
TCO79637__2_--------------------------------------------MQRVIDLRNPGEVLACGGLA------WLASLAD----------PLAETGFEPMAE-----------PGVMRFACP---------DTALQQFDAVRVVPVVDDDDALQLG--------------------------------------------------ALRLDW--WNPWG--------------LNPHMKLWSGKQ------KAESIVGN--
LFR---ASAGAASPEWL-----EFEHTT--------TGRLGVD---PRGTWN--ALGIGWSI-----NQHTH-I---EMLCRPLLELL-------AFIG------LQALPLAGS---------RRDGHVRL--HPWR----------EASFSLARLAIAGASRYA------LSGWRAP-FGKAGSN-------TI---LLTASPD-------------
RMH34963 ----------------------------------------MLLLRVRLYWKNQRCWRKWEMIKIKVDVLNPGQFFACCGLL------ELSHRLW-------PGEGHFSKCGS--------GYFFVIEPPPRGESNDVDLRCLLNKLCNCKVSGLTEEEQKERSE------------LEKKRSELEKKRKELSELEENRRKELGEKAREGTLHLGE------PFKLTLDW--WQS----------
EDEKATLKTWAGRQEI---HKIARAAQ--------------DALKALLCNGKIDLCELLNHGCILRTPA-------EYAQKDRDQA---KPVEPFC---FDTRRYVHALKVGFSL------DAQNT-----DILAYPAVELL-------ALIG------LQRFPLLPSGQ--------KWSFEYVI------WLR-----PLGVPVAAAVASGVIPFVS------SERYLFR-FVFRDDQK--------RYKAFGRATKISN--------
WP_083437694 ----------MRLTKSCADMRVCNSASGDGDLLGWSRWFAAPTLLPRGKQHNRRTGEMSNQLGFAVNVDILNPGQFFACCGLL------ELAHRLW-------PGIEGHFEARG--------SQASFIVQAPTSDGL----KCLLDHLCRCDISGLTEEEQQERRQ--------------LEERKRQNRLSDTEEERRKELQRAREGKLKLYLKD------PFHLTLDW--
WQT---------ESEDVSPKTWAGRQEI---HKIARAAQ--------------EALCEALCN-NLNLSNLLDYGCVLRTPS-------EYAQETRGTA---KPVEPFC---FDARRYVHALKVGFSL------DLQDA------EIVAYPAVELL-------ALIG------LQRFRPVPSSE---------RWSLEYTA------WFQ--------PLSAVVAAAVVSGAIPLAQ-----SKKYRFR-LLFRDDQK--------RYKAFGRASKIYSIQ-------
CDM65513 ------------------------------------------MSNQLGFAVNVDILNPGQFFACCGLL------ELAHRLW-------PGIEGHFEAR--------SQASFIVQAPTSDG--------KCLLDHLCRCDISGLTEEEQQERRQ--------------LEERKRQNRLSDTEEERRKELQRAREGKLKLYLKD------PFHLTLDW--WQT----------ESEDVSPKTWAGRQEI---
HKIARAAQ--------------EALCEALCN-NLNLSNLLDYGCVLRTPS-------EYAQETRGTA---KPVEPFC---FDARRYVHALKVGFSL------DLQDA------EIVAYPAVELL-------ALIG------LQRFRPVPSSE---------RWSLEYTA------WFQ--------PLSAVVAAAVVSGAIPLAQ-----SKKYRFR-LLFRDDQK--------RYKAFGRASKIYSIQ-------
WP_105311990__2_--------------------------MSNSANITIPVDTNPGEFFACCGLL------EVADRLW-------PGAEGWFE------------PG-------PNRSRYTV-----WTK-------PLPVNAIAFPVVCGFTPVHG-------SLAFAFH-NFFRTDQKH--------KC--FSRATQRKE--------
---EPSESDPLN--APKTVFDPPTASKQGKR-----------NEAPAK---------REPFYFD----CRRGCNAHPLDSGFSP-----DTQNL------KSECFPAVEAM-----------CFIG------LQRARPAPTDE---------PNRSRYTV-----WTK------PLPVNAIAFPVVCGFTPVHG--------SLAFAFH-NFFRTDQKH--------KC--FSRATQRKE--------
WP_114366905__2_--------------------------------MTHPKPTITVDVDVTNPGQFFACCGLL------EIADRLW-------PGAEGWF-------------EDSAFCI--------------------DAEGTLSELWCAIVD-------------AELDALDPGDETASPMRLGA----------PIDLTLDW--WKDEH-----------AGGRMLKVWAGSMRG---VRIAQSMK-------
AAIANVAKQ--VSP---FDYSAVVYDP---------DNTKKK---------VEPFYYD----ARRGWNAQPIDGFSP------DSLKM------ISAAYPAVEFM-----------CLVG------LQRFRPRQT----KQ--------RRVFEYRT-----WRT---------PLSPSLAAAVASGVMPLAA-----DVSYRFE-NAFRTDQKKH---------KA--FLPATPLGASQ-------
NP_951114__3_--------------------------------MSSNSKSAISIRVDPTNPGQFFACCGLL------ELADRLW-------PGAEGWFDK---------DGFLFSLRPYQEAKAQKCAPATLLAEITRCRLTNTMSDSELKRRD---------QLMAVPKKIESDPSLEAEKKVLDKLWREAPLVLHS-------PFNIRVDW--FLD----------
EYCGGTMFKTWAGQQSVFE-IGRGMKAA--------------LDSGDWSHT--SP---DDWLRRRMLND-----------SLPFNFD----SELGGVGSDLDVGFSF-----DPLKT-I----VQRAPLLEFL--------AFVG------LQRFRPMKINA---------ENRFQYFL-----WFG---------PLVSEVAAPAACGLFGPVR-----LKAFEFR-LLYRTK------------YLKSFLPATPIQRS--------
RJO66009 --------------------------------MNQALEPTIRVHVDPTNPGQFFACCGLL------EIADRIW--------PGAEGWF-----------EGEGFAFR-------SCIADSLADAKDMRLTNVIPKLCFE---------------SLDPGDEYSPPLALRA-------------SIEIRLDW--WKDDQ-----------AGGRMLKTWAGGQKV---INIAEGMK-------
-----REAEAMLS--RPS----WEWLQHRDDSS-------------ELPFNFD---SDLGGMGSDRDLGFSM-------DPLKM-----KATTRPMVELM-------AFIG-------LQRFRPAKLEN---------ENRYRFST------WSD----------PLVPEIASIAACGMLEGPPR---LQTYEFR-LLYRTK------------YLKSFLPANPIGGDQ-------
RLS31758 -------------------------------------MSLREPSITVNVDITNPGQFFACCGLL------ELADRMW-------PGAEGWF---------------ENGRFHITCDGTL-------RQMVDALACSDITSSVGEEGLKRLG-------------TLLSATKATLTADDERDKERLRALWQGESLLLGN------PFNLALNW--WRNET-----------NERTGLKTW---------
AAKQLILD--------------IVRPLRSAVIAN---APLAPTEDILQ---------CAVAFD-------GSSLYFD----SHAQSQSRAIDTGFSA------YDLRA-VIKEGPSTRPGLEFF-------AFIG-------MQRFRIPPRHS---------DGTFVYCA----WRT-------PLPALVAALAATGFIRSPN-----DTCYQFR-LLDRTK------------YMKAFLPATTMENHR-------
WP_010038639__2_--------------------------------MSFSVAVDPTNPGQFFACCGLL------EIADRLW-------PGAEGWFEDGTFHVST-------SCIATLTDIDQ-------LVCLRFVSSLGDDGLKERLGLLSLLSAD----------KSTLTELDKQDKERLRKAWQVERVQIPA------PFDFWLDW--WNDK-----------SGVTLLKTW-------
AAKQFVLS--------------MANPMLSALKSLTWDSSSRENCLQ------RTAKLS-------GLPFYFD----AANNTQNTPRDNGFAP------GAVKS-----APNDRPLLELL-------AFVG-------LQRFRPHRPDK----------AERILYAT-----WSI-----------PLPPSVASVAAAGMLNVTG----SRAFEFR-MLYRTE----------YMKAFLPAQAVRGVK---------

PSR23716 [illegible]
TCT03823 [illegible]
WP_058867775 [illegible]
SDE62562__2_ [illegible]
WP_076837953__2_ [illegible]
WP_100918689__2_ [illegible]
WP_083924498 [illegible]
WP_019625852 [illegible]
PKM10505 [illegible]
PPD51365__4_ [illegible]
WP_101052171__2_ [illegible]
WP_081608002 [illegible]
RMG17092 [illegible]
[illegible]

;
end;
